# Functional characterization of all *CDKN2A* missense variants and comparison to in silico models of pathogenicity

**DOI:** 10.1101/2023.12.28.573507

**Authors:** Hirokazu Kimura, Kamel Lahouel, Cristian Tomasetti, Nicholas J. Roberts

## Abstract

Interpretation of variants identified during genetic testing is a significant clinical challenge. In this study, we developed a high-throughput CDKN2A functional assay and characterized all possible *CDKN2A* missense variants. We found that 17.7% of all missense variants were functionally deleterious. We also used our functional classifications to assess the performance of in silico models that predict the effect of variants, including recently reported models based on machine learning. Notably, we found that all in silico models performed similarly when compared to our functional classifications with accuracies of 39.5-85.4%. Furthermore, while we found that functionally deleterious variants were enriched within ankyrin repeats, we did not identify any residues where all missense variants were functionally deleterious. Our functional classifications are a resource to aid the interpretation of *CDKN2A* variants and have important implications for the application of variant interpretation guidelines, particularly the use of in silico models for clinical variant interpretation.

## Introduction

Genetic testing of patients with cancer to identify variants associated with an increased cancer risk and sensitivity to targeted therapies is becoming more common as broad testing criteria are integrated into clinical care guidelines (Goggins et al., 2020; Stoffel et al., 2019). The American College of Medical Genetics (ACMG) provides a framework to integrate multiple types of evidence, including variant characteristics, disease epidemiology, clinical information, and functional classifications, to interpret variants in any gene (Richards et al., 2015). In silico variant effect predictors are also integrated into ACMG variant interpretation guidelines as supporting evidence to aid classification of variants. While numerous models have been developed, varied accuracy, poor agreement between models, and inflated performance on publicly available data have been reported (Cubuk et al., 2021; Jaffe et al., 2011; Wilcox et al., 2022). Recently developed variant effect predictors aim to overcome these limitations by incorporating deep-learning based protein structure predictions and by not training on human annotated datasets (Brandes et al., 2023; Cheng et al., 2023; Gao et al., 2023). However, post-development assessment of machine learning based variant effect predictors, to determine accuracy on novel experimental datasets and suitability for clinical use, are limited.

Variants that cannot be classified as either pathogenic or benign are categorized as variants of uncertain significance (VUSs). However, while pathogenic and benign variants identified during genetic testing are clinically actionable, VUSs are the cause of deep uncertainty for patients and their health care providers as an unknown fraction are functionally deleterious and therefore, likely pathogenic. For example, individuals with germline VUSs in a pancreatic cancer susceptibility gene are not be eligible for clinical surveillance programs that are associated with improved patient outcomes, unless they otherwise meet family history criteria (Goggins et al., 2020; Stoffel et al., 2019). Similarly, patients with breast or pancreatic cancer and a germline *BRCA2* VUS would not be eligible for treatment with olaparib, a poly (ADP-ribose) polymerase inhibitor (Golan et al., 2019; Tutt et al., 2021). Reclassification of VUSs into pathogenic or benign strata has real-world, life-or-death consequences that necessitate a high degree of accuracy.

Germline VUSs in hereditary cancer genes are a common finding in patients with cancer and frequently can be reclassified as pathogenic on the basis of in vitro functional evidence (Kimura et al., 2022). In patients with pancreatic ductal adenocarcinoma (PDAC), germline *CDKN2A* VUSs affecting p16^INK4a^, most often rare missense variants, are found in up to 4.3% of patients (Chaffee et al., 2018; Kimura et al., 2021; McWilliams et al., 2018; Roberts et al., 2016; Shindo et al., 2017; Zhen et al., 2015). As functional data from well-validated in-vitro assays are incorporated into ACMG variant interpretation guidelines, we recently determined the functional consequence of 29 *CDKN2A* VUSs identified in patients with PDAC using an in vitro cell proliferation assay (Kimura et al., 2022; Richards et al., 2015). We found that over 40% of VUSs assayed were functionally deleterious and could reclassified as likely pathogenic.

Functional characterization, however, is time-consuming, expensive, and requires technical and scientific expertise. These limitations hinder assessment of in silico variant effect predictors and patient access to functional data that may allow reclassification of VUSs into clinically actionable strata. As *CDKN2A* VUSs will continue to be identified in patients with cancer undergoing genetic testing, we developed a high-throughput functional assay to provide a broad interpretation framework for *CDKN2A* variants. We characterized all possible *CDKN2A* missense variants and compared our functional classifications to recently developed in silico models based on machine learning to determine the accuracy of variant effect predictions.

## Results

### Functional characterization of *CDKN2A* missense variants

We utilized a codon optimized *CDKN2A* sequence for our multiplexed functional assay. Expression of codon optimized CDKN2A or the synonymous CDKN2A variants, p.L32L, p.G101G, and p.V126V, in PANC-1, a PDAC cell line with a homozygous deletion of *CDKN2A*, resulted in significant reduction in cell proliferation (P value < 0.0001; **Figure 1-figure supplement 1A**). There was no significant difference between codon optimized CDKN2A and the three synonymous variants assayed. Conversely, expression of three pathogenic variants, p.L32P, p.G101W, and p.V126D, in PANC-1 cells did not result in any significant changes in cell proliferation. To determine if there were unappreciated selective effects during in vitro culture, we generated a CellTag library based on the pLJM1 plasmid that contained twenty non-functional 9 base pair barcodes of equal representation. We then transduced PANC-1 cells that stably expressed codon optimized CDKN2A with the CellTag library (Day 0) and determined representation of each barcode in the cell pool on Day 9 and at confluency (Day 45). We found no statistically significant changes in barcode representation, indicating that representation of a pool of functionally neutral variants is stable over a period of in vitro culture representing our assay time course (**Figure 1-figure supplement 1B, Appendix 1-table 1**).

We next determined whether we could identify functionally deleterious *CDKN2A* variants at a single residue when all amino acid variants were assayed simultaneously. We generated lentiviral expression plasmid libraries for all 156 CDKN2A amino acid residues, where each library contained all possible amino acids at a single residue. Twenty-seven variants (27 of 3,120, 0.87%) were represented in the plasmid libraries at ≤ 1%. Expression plasmids for each of these 27 variants were individually generated by site directed mutagenesis and added to the corresponding plasmid library to a calculated representation of 5% (**Figure 1-figure supplement 2A and 2B, Appendix 1-table 2).** Plasmid libraries were then individually amplified, and lentivirus produced. To confirm that the representation of each variant was maintained after transduction, we transduced three lentiviral libraries (amino acid residues p.R24, p.H66, and p.A127) individually into PANC-1 cells and determined the proportion of each variant in the amplified plasmid library and in the cell pool at Day 9 post-transduction. The proportion of each variant in the amplified plasmid library and in the cell pool at Day 9 were highly correlated (**Figure 1-figure supplement 2C and 2D, Appendix 1-table 3**).

For two CDKN2A amino acid residues that include pathogenic and benign variants, p.V126 and p.R144, we determined the representation of each variant in the transduced cell pool at Day 9 and at confluency after a period of in vitro culture, Day 23 and Day 31 post-transfection, respectively (**Figure 1A and 1B**, **Appendix 1-table 4**, **Appendix 1-table 5**). Two synonymous variants, p.V126V and p.R144R, as well as a previously reported benign variant, p.R144C, either decreased or maintained their representation in the cell pool during in vitro culture as determined by the number of sequence reads supporting the variant. Representation of a previously reported pathogenic variant, p.V126D, increased in the cell pool. Notably, several other variants including p.V126R, p.V126W, p.V126K, and p.V126Y, also increased in representation in the cell pool, suggesting that additional amino acid changes at this residue are functionally deleterious (**Figure 1A**).

**Figure 1.**
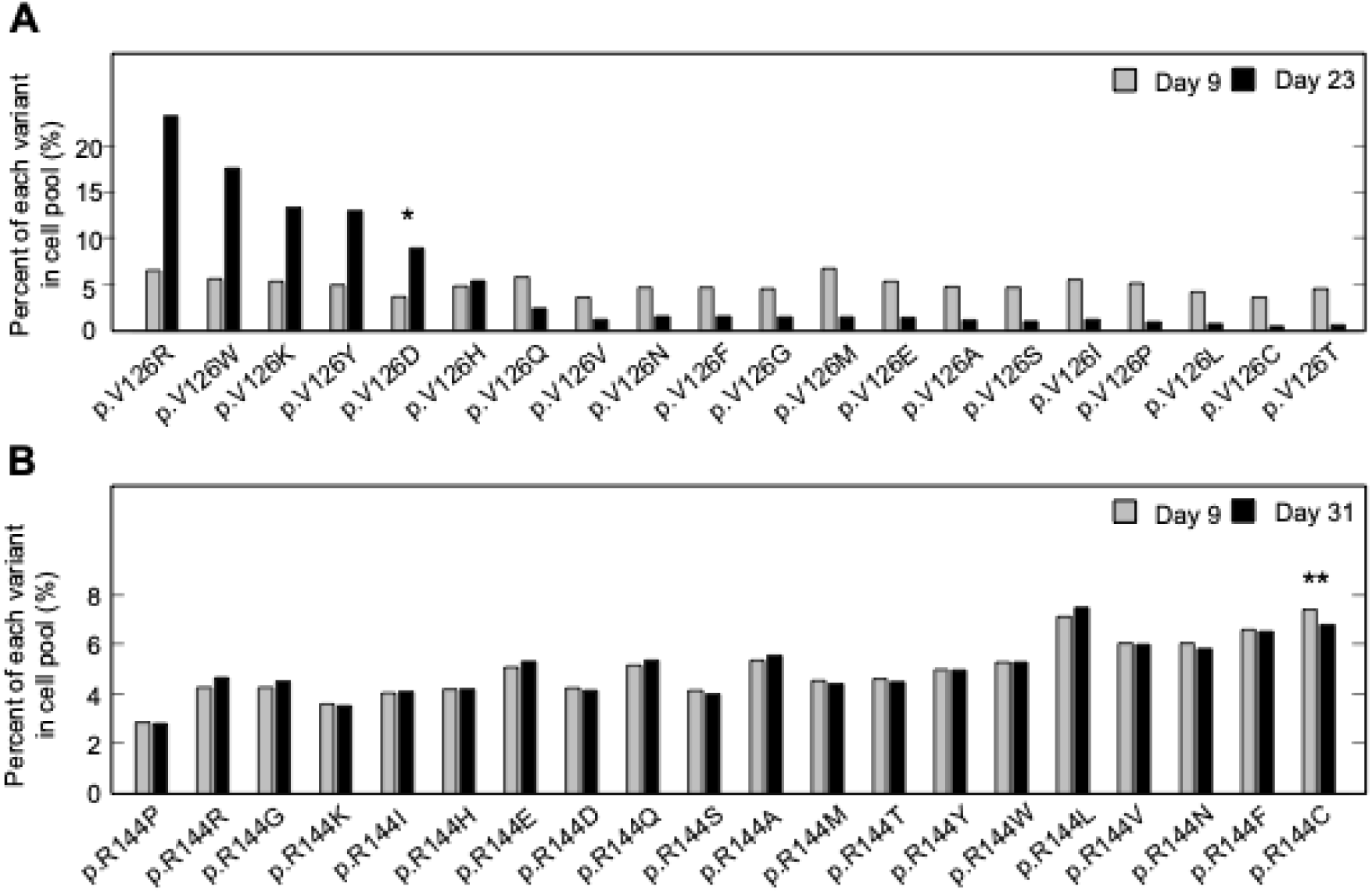
Pooled analysis of *CDKN2A* variants at two residues with previously reported pathogenic and benign variants. PANC-1 cell stably expressing one of 20 *CDKN2A* variants, 19 missense variants and 1 synonymous variant, at reside p.V126 or p.R144 were cultured. Variant representation, as the percent of reads supporting the variant sequence, before and after a period in vitro cell proliferation determined by next generation sequencing for the two residues, p.V126 (**A**) or p.R144 (**B**). *CDKN2A* variant p.V126D (*) was previously reported as pathogenic and increased representation during in vitro proliferation. CDKN2A variant p.R144C (**) was previously reported as benign variant and maintained representation during in vitro proliferation.

To functionally characterize 2,964 *CDKN2A* missense variants, PANC-1 cells were transduced with each of the 156 lentiviral expression libraries individually and representation of each *CDKN2A* variant in the resulting cell pool determined at Day 9 after transduction and at confluency (Day 16 – 40) (**Appendix 1-table 5**). Variant read counts were then analyzed using a gamma generalized linear model (GLM), that does not rely on annotation of pathogenic and benign variants to set classification thresholds, and variants with statistically significant P values were classified as functionally deleterious (log_2_ P values ≤ -53.2). Variants with P values that did not reach statistical significance were classified as either of indeterminate function (log_2_ P values > -53.2 and < -5.8) or functionally neutral (log_2_ P values ≥ -5.8).

We found that 525 of 2,964 missense variants (17.7%) were functionally deleterious in our assay (**Figure 2A, Figure 2-figure supplement 1A, Appendix 1-table 4**). In addition, 1,784 variants (60.2%) were classified as functionally neutral, with the remaining 655 variants (22.1%) classified as indeterminate function (**Figure 2A**, **Appendix 1-table 4**). In general, our results were consistent with previously reported classifications. Of variants identified in patients with cancer and previously reported to be functionally deleterious in published literature and/or reported in ClinVar as pathogenic or likely pathogenic (benchmark pathogenic variants), 27 of 32 (84.4%) were functionally deleterious in our assay (**Figure 2B, Figure 2-figure supplement 1B and 1C**, **Appendix 1-table 4**) (Chaffee et al., 2018; Chang et al., 2016; Horn et al., 2021; Hu et al., 2018; Kimura et al., 2022; McWilliams et al., 2018; Roberts et al., 2016; Zhen et al., 2015). Five benchmark pathogenic variants were characterized as indeterminate function, with log_2_ P values from -19.3 to -33.2. Of 156 synonymous variants and six missense variants previously reported to be functionally neutral in published literature and/or reported in ClinVar as benign or likely benign (benchmark benign variants), all were characterized as functionally neutral in our assay (**Figure 2B, Figure 2-figure supplement 1B and 1C, Appendix 1-table 4**) (Kimura et al., 2022; McWilliams et al., 2018; Roberts et al., 2016). Of 31 VUSs previously reported to be functionally deleterious, 28 (90.3%) were functionally deleterious and 3 (9.7%) were of indeterminate function in our assay. Similarly, of 18 VUSs previously reported to be functionally neutral, 16 (88.9%) were functionally neutral and 2 (11.1%) were of indeterminate function in our assay, (**Figure 2B, Figure 2-figure supplement 1B and 1C, Appendix 1-table 4**).

**Figure 2.**
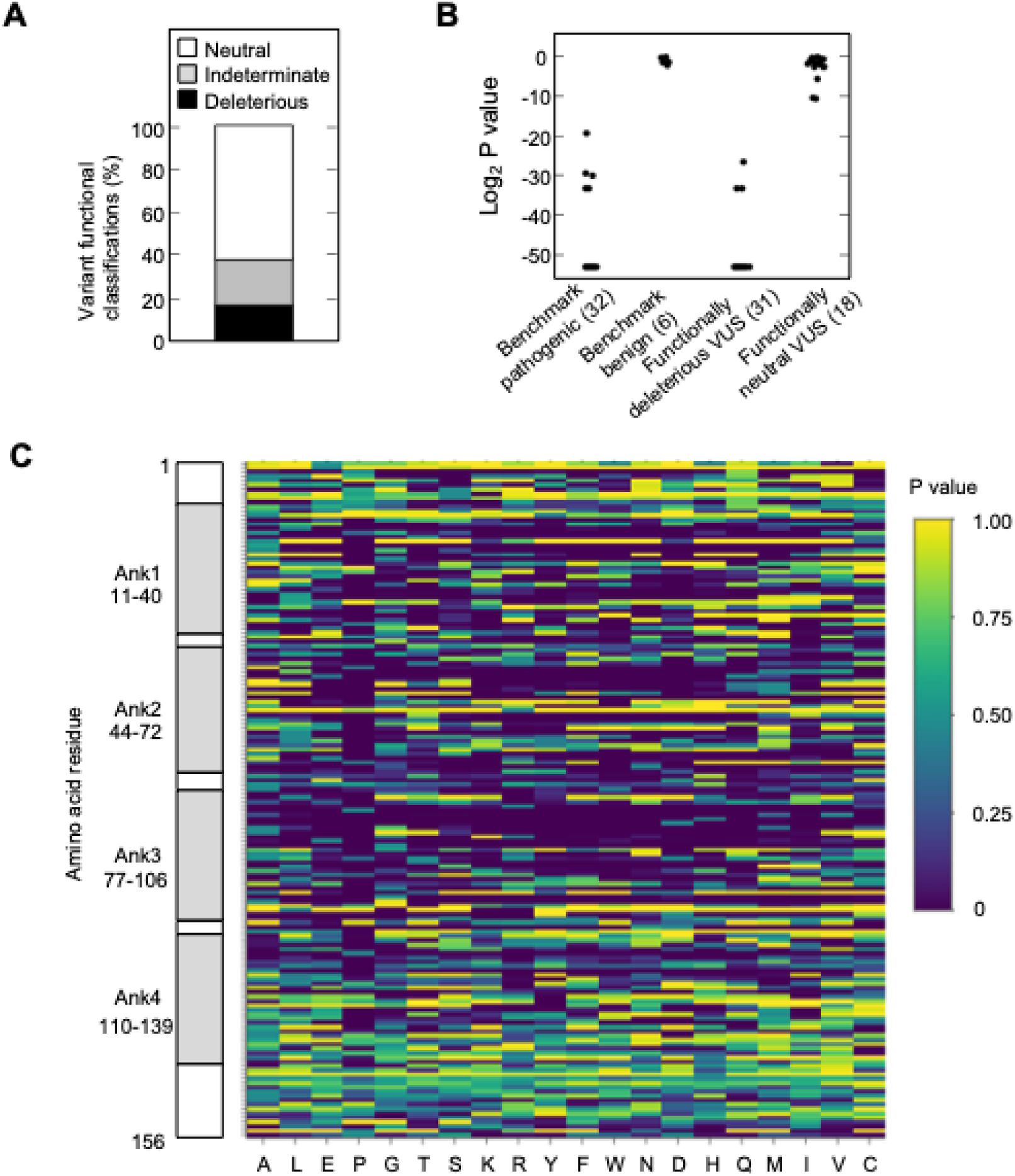
Functional characterization of all possible *CDKN2A* missense variants. (**A**) Functional classifications for 3,120 *CDKN2A* variants, including 2,964 missense variants and 156 synonymous variants. Variants were classified as functionally deleterious, indeterminate function, or neutral based on P value using gamma GLM. 525 (17.7%) variants were classified as functionally deleterious. (**B**) Log_2_ P values for 32 benchmark pathogenic variants, 6 benign variants, 31 VUSs previously reported to have functionally deleterious effects, and 18 VUSs previously reported to have functionally neutral effects. (**C**) Heat map with P values for all 3,120 *CDKN2A* variants assayed.

We next compared variant classifications using the gramma GLM to variant classifications using a normalized fold change method (Brenan et al., 2016; Giacomelli et al., 2018). Classification of missense variants using normalized fold change also differentiated benchmark pathogenic and benchmark benign variants (**Figure 2-figure supplement 2A and 2B, Appendix 1-table 6**). Using benchmark pathogenic variants and benchmark benign variants to set thresholds for classification, we classified all variants as either functionally deleterious (log_2_ normalized fold change ≤ 0.24), indeterminate function (log_2_ normalized fold change > 0.24 and < 1.09), or functionally neutral (log_2_ normalized fold change ≥1.09). Using these thresholds, 12 of 18 VUSs (66.7%) previously reported to be functionally neutral were classified as functionally neutral, while 6 (33.3%) were of indeterminate function. Similarly, of 31 VUSs previously reported to be functionally deleterious, 30 (96.8%) were functionally deleterious and 1 (3.2%) was of indeterminate function (**Figure 2-figure supplement 2A and 2B, Appendix 1-table 6**). Overall, 632 of 2,964 missense variants were functionally deleterious (21.3%), 674 variants were indeterminate function (22.7%), and 1658 variants were functionally neutral (55.9%) using log_2_ normalized fold change to classify variants (**Figure 2-figure supplement 2C, Appendix 1-table 6**). Notably, 517 of 525 variants (98.5%) classified as functionally deleterious and 1,586 of 1,784 variants (88.9%) classified as functionally neutral using the gamma GLM were similarly classified using log_2_ normalized fold change (**Figure 2-figure supplement 2D**).

To confirm the reproducibility of our variant classifications, 28 amino acid residues were assayed in duplicate, and variants classified using the gamma GLM. The majority of missense variants, 452 of 560 (80.7%), had the same functional classification in each of the two replicates (**Figure 2-figure supplement 3A and 3B, Appendix 1-table 4**). Of variants with discordant classifications, 6 (1.1%) were functionally deleterious in one replicate and of indeterminate function in another. While 102 variants (18.2%) were functionally neutral in one replicate and of indeterminate function in another. Importantly, no variant that was functionally deleterious in one replicate and functionally neutral in another (**Appendix 1 - table 4**). Furthermore, the correlation coefficient between duplicate assay results was similar using the gamma GLM and log_2_ normalized fold change (**Figure 2-figure supplement 3A and 3C**). We also determined whether underrepresentation in the cell pool at Day 9 affected variant functional classifications. Fifty-three of 2,964 missense variants (1.8%) were present in the cell pool at Day 9 of the first assay replicate (experiment 1) at < 2%, as determined by the number of sequence reads supporting the variant (**Figure 2-figure supplement 4A, Appendix 1-table 4**). There was no statistically significant difference in the proportion of variants classified as functionally deleterious for variants present in less than 2% of the cell pool at Day 9 (12 of 53 variants; 22.6%), and variants present in more than 2% of the cell pool (496 of 2,911 variants; 17.0%) (P value = 0.28) (**Figure 2-figure supplement 4B**). We also found no significant differences in the proportion of variants classified as functionally deleterious for variants present in more than 2% of the cell pool at Day 9 when variants were binned in 1% intervals (**Figure 2-figure supplement 4B**).

### Comparison to in silico prediction algorithms

As in silico predictions of variant effect are integrated into ACMG variant interpretation guidelines as supporting evidence, we compared the ability of different algorithms, including recently described algorithms that incorporate deep-learning models of protein structure, to predict the functional consequence of *CDKN2A* missense variants. We compared our functional classifications to predictions from Combined Annotation Dependent Depletion (CADD), Polymorphism Phenotyping v2 (PolyPhen-2), Sorting Intolerant From Tolerant (SIFT), Variant Effect Scoring Tool score (VEST), AlphaMissense, ESM1b, and PrimateAI-3D. In silico predictions for all missense variants were available for PolyPhen-2, SIFT, VEST, AlphaMissense, and ESM1b. For CADD and PrimateAI-3D, 910 (152 functionally deleterious, 196 indeterminate, and 562 functionally neutral) and 904 (152 functionally deleterious, 196 indeterminate, and 556 functionally neutral) missense variants had in silico predictions available respectively (**Appendix 1-table 7**). In silico variant effect predictors performed similarly across a broad range of performance characteristics (**Appendix 1-table 8**). Accuracy of in silico model predictions were 39.5 – 85.4% (CADD – 45.1%; PolyPhen-2 – 39.5%; SIFT – 60.9%; VEST – 71.9%; AlphaMissense – 71.6%; ESM1b – 59.2%; and PrimateAI-3D; 85.4%) (**Figure 3**). We also assessed sensitivity, specificity, positive predictive value, and negative predictive value for each model. We found that sensitivity was 0.25 – 0.98 (CADD – 0.97; PolyPhen-2 – 0.98; SIFT – 0.79; VEST – 0.91; AlphaMissense – 0.94; ESM1b – 0.95; and PrimateAI-3D – 0.25), specificity was 0.27 – 0.98 (CADD – 0.35; PolyPhen-2 – 0.27; SIFT – 0.57; VEST – 0.68; AlphaMissense – 0.67; ESM1b – 0.51; and PrimateAI-3D – 0.98), positive predictive value was 0.22 – 0.68 (CADD – 0.23; PolyPhen-2 – 0.22; SIFT – 0.28; VEST – 0.38; AlphaMissense – 0.38; ESM1b – 0.3; and PrimateAI-3D – 0.68), and negative predictive value was 0.87 – 0.98 (CADD – 0.98; PolyPhen-2 – 0.98; SIFT – 0.93; VEST – 0.97; AlphaMissense – 0.98; ESM1b – 0.98; and PrimateAI-3D – 0.87).

**Figure 3.**
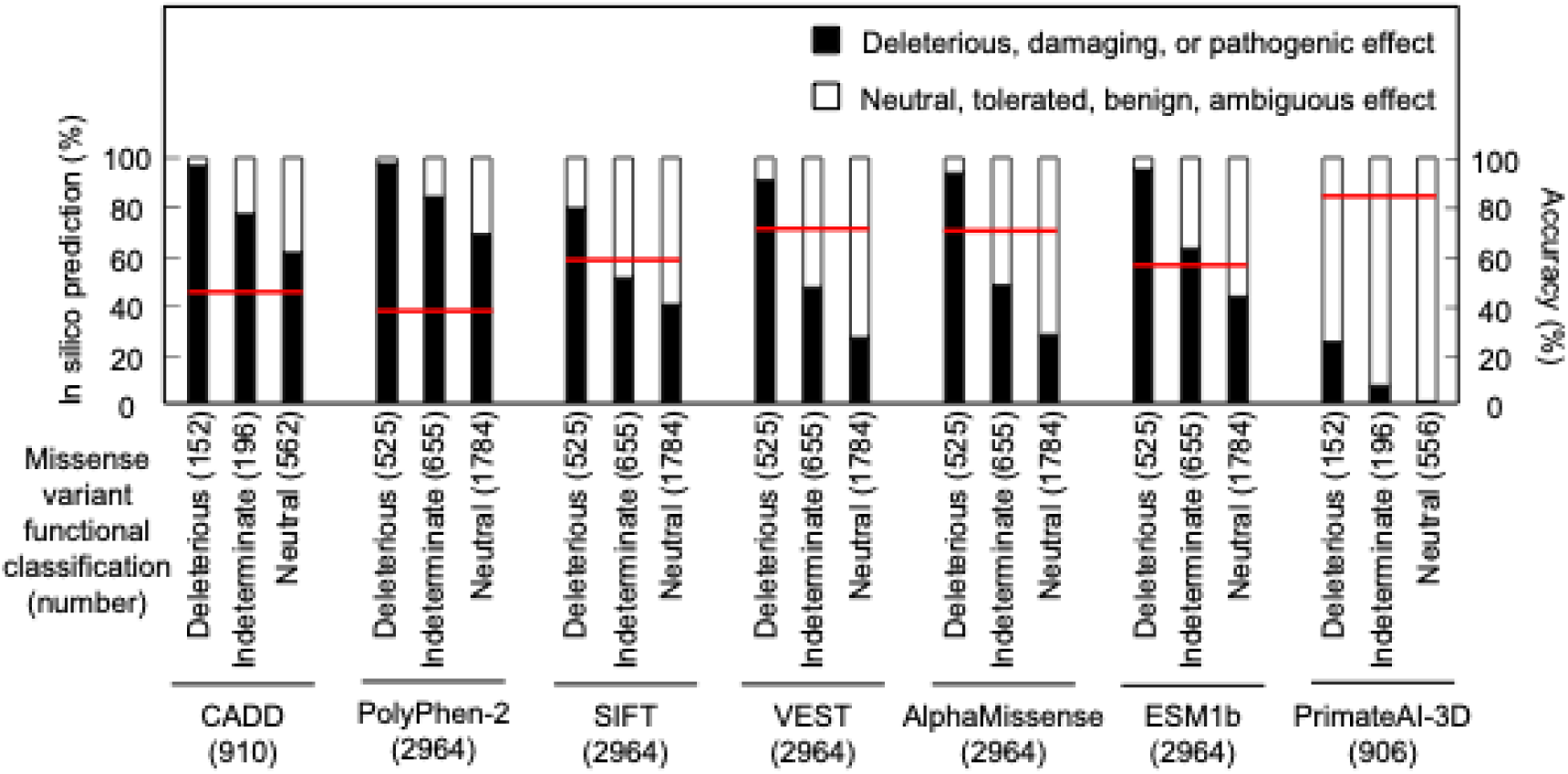
Comparison of functional classifications and in silico variant effect predictions for all possible CDKN2A missense variants. Variant effect predictions for *CDKN2A* missense variants using CADD, PolyPhen-2, SIFT, VEST, AlphaMissense, ESM1b, and PrimateAI-3D. Predicted deleterious, damaging, or pathogenic effects (black box) and predicted neutral, tolerated, benign, or ambiguous effects (white box) presented as percent of missense variants with an available prediction. Number of missense variants with an available prediction for each in silico model given in parentheses. Accuracy shown as a red line. CADD; Combined Annotation Dependent Depletion, PolyPhen-2; Polymorphism Phenotyping v2, SIFT; Sorting Intolerant From Tolerant, VEST; Variant Effect Scoring Tool score.

We also tested the effect of combining multiple in silico predictors. 904 missense variants had in silico predictions from all 7 algorithms. The remaining 2,060 missense variants had in silico predictions from 5 algorithms. Of variants with in silico predictions from all 7 algorithms, 378 (41.8%) had predictions of deleterious or pathogenic effect from a majority of algorithms (≥ 4), and of these, 137 (36.2%) were functionally deleterious in our assay. Similarly, of 2,060 missense variants that had in silico predictions from 5 algorithms, 1107 (53.7%) had predictions of deleterious or pathogenic effect from a majority of algorithms (≥ 3), of which, 361 (32.6%) were functionally deleterious in our assay (**Appendix 1-table 7**).

### Distribution of functionally deleterious variants

Analysis of functionally deleterious variants may highlight critical and non-critical resides for CDKN2A function. We found that functionally deleterious missense variants were not distributed evenly across CDKN2A. CDKN2A contains four ankyrin repeats that mediate protein-protein interactions, ankyrin repeat 1 at codon 11-40, ankyrin repeat 2 at codon 44-72, ankyrin repeat 3 at codon 77-106, and ankyrin repeat 4 at codon 110-139 (Goldstein, 2004; Ruas and Peters, 1998; Sun et al., 2010) (**Figure 2-figure supplement 5A**).

Functionally deleterious variants were enriched in ankyrin repeat 1 (21.0%, adjusted P value = 0.01), ankyrin repeat 2 (26.2%, adjusted P value = 1.0 x 10^-10^), and ankyrin repeat 3 (26.3%, adjusted P value = 2.6 x 10^-11^), while depleted in ankyrin repeat 4 (6.5%, adjusted P value = 3.2 x 10^-13^) and non-ankyrin repeat regions (6.8%, adjusted P value = 0) (**Figure 2-figure supplement 5B**). Moreover, functionally deleterious variants were further enriched within 10 residue subregions of ankyrin repeats 1-3, with 37.0% of variants in residues 16-25 of ankyrin repeat 1, 40.0% of variants in residues 46-55 of ankyrin repeat 2, and 48.0% of variants in residues 80-89 of ankyrin repeat 3 being classified as functionally deleterious (**Figure 2C**, **Appendix 1-table 4**).

Across all single residues, the mean percent of functionally deleterious missense variants was 17.7% (95% confidence interval: 12.7% - 20.9%) (**Figure 2-figure supplement 5C, Appendix 1-table 4**). At five amino acid residues, p.G23, p.G55, p.H83, p.D84, and p.G89, 17 of 19 (89.5%) possible missense variants were functionally deleterious. Notably, these residues are conserved between human and murine p16 (Byeon et al., 1998). And p.H83 has been reported to stabilize peptide loops connecting the helix-turn-helix structure of the four ankyrin repeats (Byeon et al., 1998), whereas p.D84 and p.G89 are located in a 20-residue region reported to interact with CDK4 and CDK6 (Fåhraeus et al., 1996). Conversely, 18 residues were tolerant of amino acid substitutions, with no missense variant characterized as functionally deleterious in our assay (**Figure 2-figure supplement 5C, Appendix 1-table 4**). We also determined whether the location of variants in protein domains correlated with in silico predictions for the 904 missense variants with predictions from all 7 algorithms (**Figure 3-figure supplement 1A – 1H**) and the 2,060 missense variants with predictions from 5 algorithms (**Figure 3-figure supplement 2A – 2H**). Notably, Ank2 and Ank3 domains had more variants predicted to have deleterious or pathogenic effect by the majority of algorithms compared to Ank1, Ank4, and non-Ank domains (**Figure 3-figure supplement 1C, Figure 3-figure supplement 2C**). We also found increasing agreement between in silico predictions of deleterious or pathogenic effect and functionally deleterious classification in our assay as the number of algorithms predicting deleterious or pathogenic effects increased. (**Figure 3-figure supplement 1B, Figure 3-figure supplement 2B).** This was true for all CDKN2A protein domains assessed (**Figure 3-figure supplement 1D – 1H, Figure 3-figure supplement 2D – 2H**).

### Functional effect of *CDKN2A* somatic mutations

Somatic alterations in *CDKN2A* are a frequent finding in many types of cancer. However, not all somatic alterations are unequivocally deleterious to protein function. Missense somatic mutations are particularly challenging to functionally interpret and the presence of a functionally neutral somatic mutation may impact patient care (Tung et al., 2020). To understand the functional effect of missense somatic mutations in *CDKN2A*, we functionally classified mutations reported in the Catalogue Of Somatic Mutations In Cancer (COSMIC) (Forbes et al., 2009), The Cancer Genome Atlas (TCGA) (Muddabhaktuni and Koyyala, 2021), patients with cancer undergoing sequencing at The Johns Hopkins University School of Medicine (JHU), and the Memorial Sloan Kettering-Integrated Mutation Profiling of Actionable Cancer Targets Clinical Sequencing Cohort (MSK-IMPACT) (Cheng et al., 2015). Overall, 355 unique missense somatic mutations were reported, of which 119 (33.5%) were functionally deleterious in our assay (**Appendix 1-table 9**). The percent of missense somatic mutations that were classified as functionally deleterious was greater than the percent of all possible *CDKN2A* missense variants classified as functionally deleterious, suggesting enrichment of functionally deleterious missense changes among somatic mutations (**Figure 2A**, **Appendix 1-table 4**, **Appendix 1-table 9**). The proportion of missense somatic mutations that were functionally deleterious was similar in COSMIC, TCGA, JHU, and MSK-IMPACT. We found that 34.2% - 53.4% of unique missense somatic mutations classified as functionally deleterious, with 61.4% - 67.6% of patients having a functionally deleterious somatic mutation (**Figure 4A**, **Appendix 1-table 9**). As with functionally deleterious variants, functionally deleterious missense somatic mutations were also not distributed evenly across *CDKN2A*, being enriched within the ankyrin repeat 3 (**Figure 4B**, **Appendix 1-table 9**). We found that 32.4% - 50.0% of all functionally deleterious missense somatic mutations occurred within ankyrin repeat 3, with 48.0% - 58.0% of patients in each cohort having a functionally deleterious missense somatic mutation in this domain. Notably, 65.7% - 76.0% of functionally deleterious missense somatic mutations in this domain were in residues 80-89 (**Appendix 1-table 9**).

**Figure 4.**
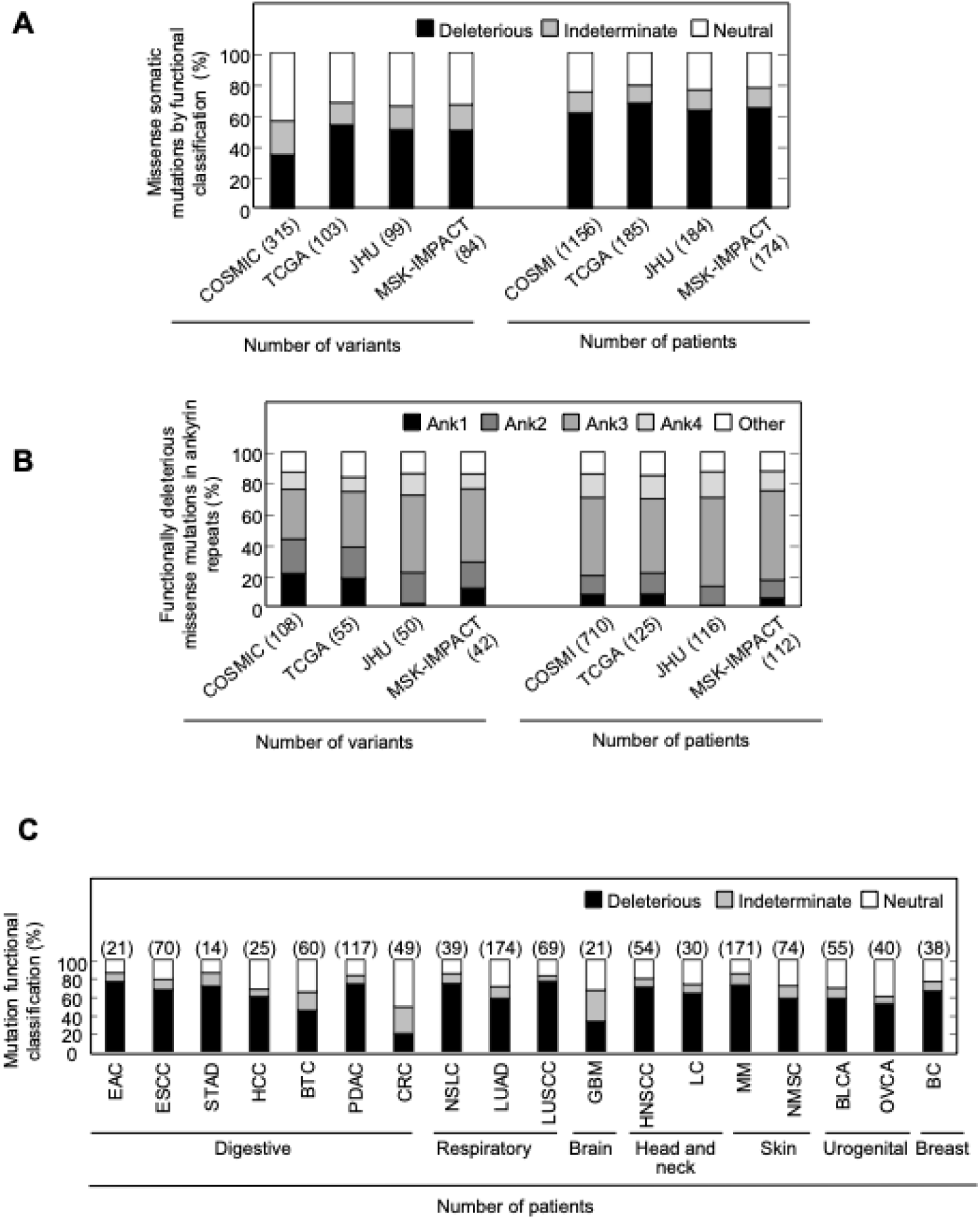
Functional classification of missense somatic mutations in *CDKN2A*. (**A**) Somatic missense variants in *CDKN2A* reported in COSMIC, TCGA, JHU, or MSK-IMPACT, by functional classification (deleterious – black box; indeterminate – gray box; neutral – white box). (**B**) Distribution of functionally deleterious missense somatic mutations *CDKN2A* reported in COSMIC, TCGA, JHU, or MSK-IMPACT by ankyrin (ANK) repeat. (**C**) Percent of missense somatic mutations in *CDKN2A* that were classified as functionally deleterious (black box), indeterminate function (gray box), or functionally neutral (white box) group by tumor type. Missense somatic mutations reported in COSMIC, TCGA, JHU, and MSK-IMPACT were combined. The number of missense somatic mutations for each tumor type given in parentheses. COSMIC; the Catalogue Of Somatic Mutations In Cancer, TCGA; The Cancer Genome Atlas, JHU; The Johns Hopkins University School of Medicine, MSK-IMPACT; Memorial Sloan Kettering-Integrated Mutation Profiling of Actionable Cancer Targets.

When considering unique missense somatic mutations, 26 of 355 (7.3%) would be classified as pathogenic or likely pathogenic by ACMG classification guidelines and these were found in 263 of 1176 (22.4%) patients in COSMIC, 45 of 185 (24.3%) patients in TCGA, 40 of 184 (21.7%) patients in JHU, and 46 of 174 (26.4%) patients in MSK-IMPACT (**Figure 4-figure supplement 1A and 1B**). In each cohort, the most prevalent of these somatic mutations were p.His83Tyr and p.Asp84Asn, with more than half of the patients with a somatic mutation that could be classified as pathogenic or likely pathogenic having either the p.His83Tyr or p.Asp84Asn alteration (**Figure 4-figure supplement 1C**). In our functional assays, these somatic mutations were both classified as functionally deleterious.

We were also able to determine the functional classification of *CDKN2A* missense somatic mutations in COSMIC, TCGA, JHU, and MSK-IMAPCT by cancer type. We found that 22.2% - 100% of *CDKN2A* missense somatic mutations were functionally deleterious depending on cancer type (**Figure 4-figure supplement 2A-2D**). When considering missense somatic mutation reported in any database, there was a statistically significant depletion of functionally deleterious mutations in colorectal adenocarcinoma (20.4%; adjusted P value = 5.4 x 10^-9^) (**Figure 4C**). As the proportion of missense somatic mutations that were functionally deleterious was less in colorectal carcinoma compared to other types of cancer, we assessed whether somatic mutations in mismatch repair genes (*MLH1*, *MLH3*, *MSH2*, *MSH6*, *PMS1*, and *PMS2*) were associated with the functional status of *CDKN2A* missense somatic mutations. Thirty-five patients in COSMIC had a *CDKN2A* missense somatic mutation, of which 12 (34.3%) had a somatic mutation in a mismatch repair gene. We found that no patients with a somatic mutation in a mismatch repair gene had a functionally deleterious *CDKN2A* missense somatic mutation compared to 6 of 23 samples (26.1%) without a somatic mutation in a mismatch repair gene (P value = 0.062).

### *CDKN2A* variants in variant databases

The Genome Aggregation Database (gnomAD) v4.1.0 reports 287 missense variants in *CDKN2A*, including the 13 pathogenic, 4 likely pathogenic, 3 likely benign, 3 benign, and 264 VUSs classified using ACMG variant interpretation guidelines (**Figure 5A and 5B**, **Appendix 1-table 10**). Of the 264 missense VUSs, 177 were functionally neutral (67.0%), 56 (21.2%) were indeterminate function, and 31 (11.7%) were functionally deleterious in our assay using the gamma GLM for classification (**Figure 5C**). Similarly, ClinVar reports 395 *CDKN2A* missense VUSs, of which 256 (64.8%) were functionally neutral, 94 (23.8%) were indeterminate function, and 45 (11.4%) were functionally deleterious in our assay (**Figure 5D**, **Appendix 1-table 11**).

**Figure 5.**
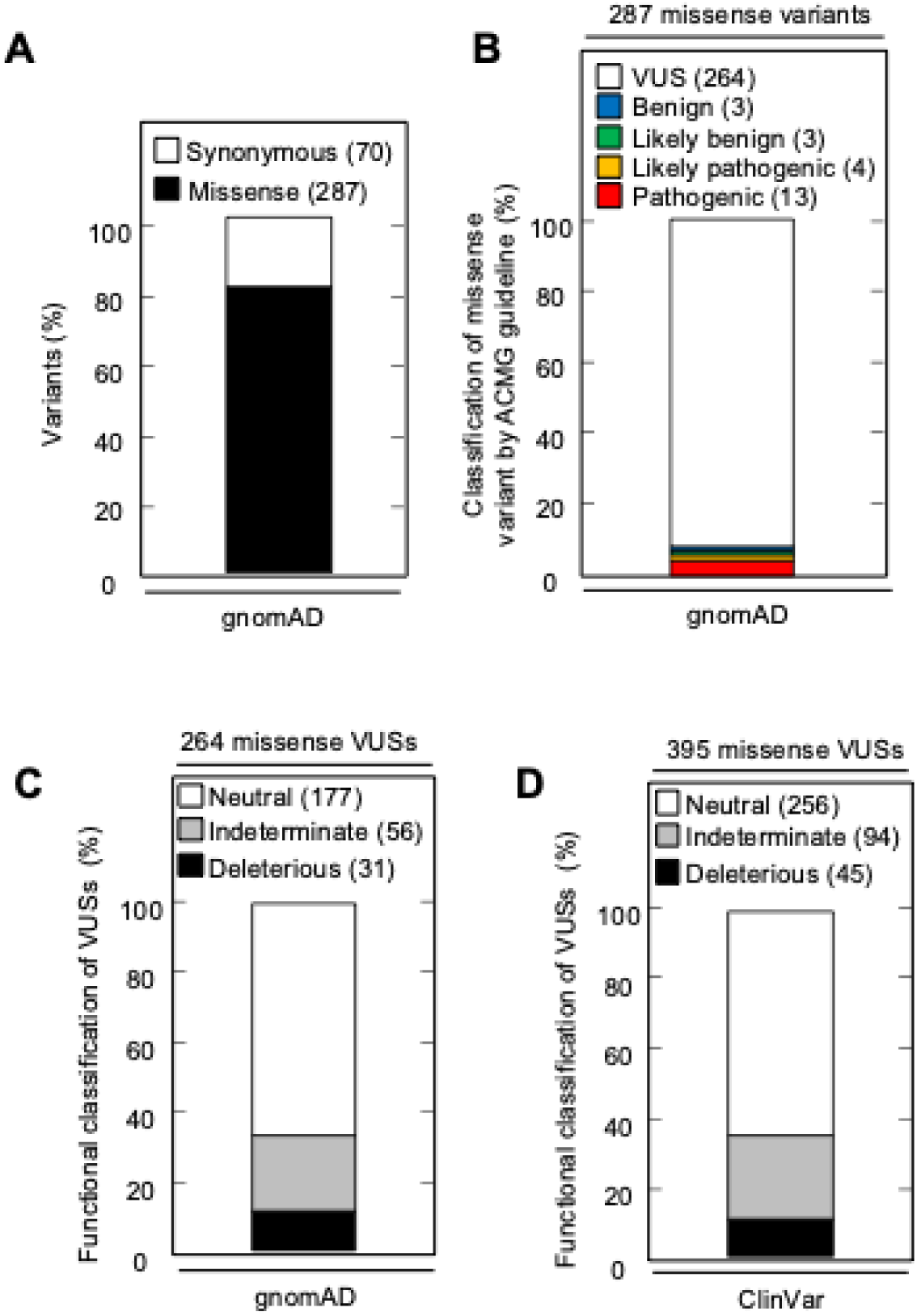
*CDKN2A* synonymous and missense variants reported in gnomAD and Clinvar. (**A**) Synonymous and missense variants in *CDKN2A* reported in gnomAD. (**B**) 287 *CDKN2A* missense variants reported in gnomAD, by ACMG guideline classification. (**C**) 264 missense variants in *CDKN2A* reported in gnomAD, by functional classification (deleterious – black box; indeterminate – gray box; neutral – white box). (**D**) 395 missense variants in *CDKN2A* reported in Clinvar, by functional classification (deleterious – black box; indeterminate – gray box; neutral – white box).

## Discussion

VUSs in hereditary cancer susceptibility genes, predominantly rare missense variants, are a frequent finding in patients undergoing genetic testing and the cause of significant uncertainty. ACMG variant interpretation guidelines incorporate functional data, as well as other evidence such as in silico predictions of variant effect, to aid classification of variants as either pathogenic or benign. *CDKN2A* VUSs are a frequent finding in patients with PDAC. We previously found that over 40% of *CDKN2A* VUSs identified in patients with PDAC were functionally deleterious and therefore could be reclassified as likely pathogenic. In this study, we developed, a high-throughput, in vitro assay and functionally characterized 2,964 *CDKN2A* missense variants, representing all possible single amino acid variants. We found that 525 missense variants (17.7%) were functionally deleterious. These pre-defined functional characterizations are resource for the scientific community and can be integrated into variant interpretation schema to aid classification of *CDKN2A* germline variants and somatic mutations.

We classified *CDKN2A* missense variants using a gamma GLM as either functionally deleterious, indeterminate functional or functionally neutral. However, we did not classify variants that may have gain-of-function effects, resulting in decreased representation in the cell pool. Future studies are necessary to determine the prevalence and significance of *CDKN2A* gain-of-function variants.

Importantly, variant classifications using a gamma GLM were not biased by assay outputs for previously reported – benchmark – pathogenic or begin variants to determine thresholds. Classification thresholds were determined using the change in representation of 20 non-functional barcodes in a pool of PANC-1 cells stably expressing CDKN2A after a period of in vitro proliferation. Even so, *CDKN2A* missense variant classifications were remarkably similar using a gamma GLM or normalized fold change with thresholds determined using benchmark pathogenic and begin variants. Of missense variants classified as functionally deleterious using a gamma GLM, 98.5% were similarly classified using normalized fold change.

We repeated our functional assay twice for 28 CDKN2A residues. For the remaining 128 residues of CDKN2A, the functional assay was completed once. While we found general agreement between functional classifications from each replicate for the 28 residues assayed in duplicate, additional repeats for each residue are necessary to determine variability in variant functional classifications.

Our characterization of all possible *CDKN2A* missense variants allowed us to assess the ability of in silico algorithms – including recently published predictors based on machine learning AlphaMissense, ESM1b, and PrimateAI-3D – to predict the pathogenicity or functional effect of *CDKN2A* missense variants. We found that all in silico variant effect predictors assessed performed similarly. Highest accuracy was observed with PrimateAI-3D at 85.4%, followed by VEST at 71.9% and AlphaMissense at 71.6%. Importantly, even in silico predictors performing best in one metric may perform poorly in others. For example, PrimateAI-3D had the highest specificity (0.98) and positive predictive values (0.68), but the lowest sensitivity (0.25) and negative predictive value (0.87). Given that reclassification of VUSs in hereditary cancer genes into inappropriate strata has significant implications for patients, use of in silico models for clinical variant interpretation, including those utilizing machine learning, may be premature. Ultimately, our data support current ACMG guidelines that include in silico predictions of variant effect as supporting evidence of pathogenicity or benign impact.

Our study also provides other insights for the implementation of variant interpretation guidelines. ACMG guidelines include presence of a missense variant at a residue with a previously reported pathogenic variant as moderate evidence of pathogenicity. We found that functionally deleterious missense variants were not evenly distributed across *CDKN2A*. We found enrichment of functionally deleterious missense variants in Ankyrin repeats 1-3 and depletion in ankyrin repeat 4. Notably, no CDKN2A residue was completely intolerant of amino acid changes. Suggesting, at least for *CDKN2A*, that the presence of a pathogenic missense variant at a residue should be used with caution when classifying other missense variants at the same residue.

We characterized variants based upon a broad cellular phenotype, cell proliferation, in a single PDAC cell line. It is possible that *CDKN2A* variant functional classifications are cell-specific and assay-specific. Our assay may not encompass all cellular functions of CDKN2A and an alternative assay of a specific CDKN2A function, such as CDK4 binding, may result in different variant functional classifications. Furthermore, *CDKN2A* variants may have different effects if alternative cell lines are used for the functional assay. However, cell-specific effects appear to be limited. In our previous study, we characterized 29 *CDKN2A* VUSs in three PDAC cell lines, using cell proliferation and cell cycle assays, and found agreement between all functional classifications (Kimura et al., 2022).

This study supports the utility of our in vitro functional assay. In general, we found that benchmark pathogenic variants, benchmark benign variants, and VUSs previously reported to be functionally deleterious had congruent functional classifications in our assay. Moreover, we found that functionally deleterious effects were enriched among somatic missense mutations, and depleted in missense VUSs in gnomAD, compared to all *CDKN2A* missense variants. Importantly, our functionally assay provides evidence to reclassify 301 of 395 (76.2%) missense VUSs reported in ClinVar and 208 of 264 (78.8%) missense VUSs reported in gnomAD. These include 45 (11.4%) VUSs in ClinVar and 31 missense VUSs in gnomAD that could be reclassified as likely pathogenic variants.

In this study, we determined functional classifications for all possible *CDKN2A* missense variants. Comparison of our functional classifications to in silico variant effect predictors, including recently described algorithms based on machine learning, provides performance benchmarks and supports current recommendations integrating data computational data into variant interpretation guidelines.

## Methods

### Cell lines

PANC-1 (American Type Culture Collection, Manassas, VA; catalog no. CRL-1469), a human PDAC cell line with a homozygous deletion of *CDKN2A* (Caldasl et al., 1994) and 293T (American Type Culture Collection; catalog no. CRL-3216), a human embryonic kidney cell line, were maintained in Dulbecco’s modified Eagle’s medium (Thermo Fisher Scientific Inc., Waltham, MA; catalog no.11995-065) supplemented with 10% fetal bovine serum (Thermo Fisher Scientific Inc.; catalog no. 26140-079). Cell line authentication and mycoplasma testing were performed using the GenePrint 10 System (Promega Corporation, Madison, WI; catalog no. B9510) and the PCR-based MycoDtect kit (Greiner Bio-One, Monroe, NC; catalog no. 463 060) (Genetics Resource Core Facility, The Johns Hopkins University, Baltimore, MD).

### *CDKN2A* somatic mutation data

*CDKN2A* (p16^INK4^; NP_000068.1) missense somatic mutation data was obtained from the Catalogue Of Somatic Mutations In Cancer (Forbes et al., 2009), The Cancer Genome Atlas (Muddabhaktuni and Koyyala, 2021), patients with cancer undergoing sequencing at The Johns Hopkins University School of Medicine (Baltimore, MD), Memorial Sloan Kettering-Integrated Mutation Profiling of Actionable Cancer Targets Clinical Sequencing Cohort (Cheng et al., 2015). *CDKN2A* variant data was obtained from gnomAD v.4.1.0. and ClinVar (Landrum et al., 2014).

### Plasmids

pHAGE-CDKN2A (Addgene, Watertown, MA; plasmid no. 116726) was created by Gordon Mills & Kenneth Scott (Ng et al., 2018). pLJM1 (Addgene; plasmid no. 91980) was created by Joshua Mendell (Golden et al., 2017). pLentiV_Blast (Addgene, plasmid no. 111887) was created by Christopher Vakoc (Tarumoto et al., 2020). psPAX2 (Addgene, plasmid no. 12260) was created by Didier Trono), and pCMV-VSV-G (Addgene, plasmid no. 8454) was created by Bob Weinberg (Stewart et al., 2003).

### CDKN2A expression plasmid libraries

Codon-optimized *CDKN2A* cDNA using p16^INK4A^ amino acid sequence (NP_000068.1), was designed (**Appendix 1-table 12**) and pLJM1 containing codon optimized CDKN2A (pLJM1-CDKN2A) generated by Twist Bioscience (South San Francisco, CA). 156 plasmid libraries were then synthesized by using pLJM1-CDKN2A, such that each library contained all possible 20 amino acids variants (19 missense and 1 synonymous) at a given position, generating 500 ng of each plasmid library (Twist Bioscience, South San Francisco, CA). The proportion of variant in each library was shown in **Appendix 1-table 2**. Variants with a representation of less than 1% in a plasmid library were individually generated using the Q5 Site-Directed Mutagenesis kit (New England Biolabs, Ipswich, MA; catalog no. E0552), and added to each library to a calculated proportion of 5%. Primers used for site-directed mutagenesis are given in **Appendix 1-table 13**. Each library was then amplified to generate at least 5 ug of plasmid DNA using QIAGEN Plasmid Midi Kit (QIAGEN, Germantown, MD; catalog no. 12143).

### Single variant CDKN2A expression plasmids

Individual pLJM1-CDKN2A expression constructs for *CDKN2A* missense variants, p.L32L, p.L32P, p.G101G, p.G101W, p.V126D, and p.V126V were generated using the Q5 Site-Directed Mutagenesis kit (New England Biolabs, Ipswich, MA; catalog no. E0552). Primers used for site-directed mutagenesis are given in **Appendix 1-table 13**. Integration of each *CDKN2A* variant was confirmed using Sanger sequencing (Genewiz, Plainsfield, NJ) using the CMV Forward sequencing primer (CGCAAATGGGCGGTAGGCGTG). The manufacturer’s protocol was followed unless otherwise specified.

### CellTag plasmid library

Twenty nonfunctional 9 base pair barcodes “CellTags” were subcloned into pLentiV_Blast using the Q5 Site-Directed Mutagenesis kit (New England Biolabs, Ipswich, MA; catalog no. E0552) (Biddy et al., 2018). Primers used to generate each CellTag plasmid are given in **Appendix 1-table 13**. Integration of each CellTag was confirmed using Sanger sequencing (Genewiz) (sequencing primer: AACTGGGAAAGTGATGTCGTG). The manufacturer’s protocol was followed unless otherwise specified. CellTag plasmids were then pooled to form a CellTag plasmid library with equal representation of each CellTag plasmid.

### Lentivirus production

Lentivirus production was performed as previously described with the following modifications (Kimura et al., 2022). pLJM1 lentiviral expression vectors (plasmid libraries and single variant expression plasmids) and lentiviral packaging vectors (psPAX2 and pCMV-VSV-G) were transfected into 293T cells using Lipofectamine 3000 Transfection Reagent (Thermo Fisher Scientific, Waltham, MA; catalog no. L3000008). Media was collected at 24 hours and 48 hours, pooled, and lentiviral particles concentrated using Lenti-X Concentrator (Clontech, Mountain View, CA; catalog no. 631231) using the manufacturer’s protocol.

### Lentiviral transduction

PANC-1 cells were used for CDKN2A plasmid library and single variant CDKN2A expression plasmid transductions. PANC-1 cells previously transduced with pLJM1-CDKN2A (PANC-1^CDKN2A^) and selected with puromycin were used for CellTag library transductions. Briefly, 1 x 10^5^ cells were cultured in media supplemented with 10 ug/ml polybrene and transduced with x 10^7^ transducing units per mL of lentivirus particles. Cells were then centrifuged at 1,200 x g for 1 hour. After 48 hours of culture at 37°C and 5% CO_2_, transduced cells were selected using 3 µg/ml puromycin (CDKN2A plasmid libraries and single variant CDKN2A expression plasmids) or 5 µg/ml blasticidin (CellTag plasmid library) for 7 days. Expected MOI was one. After selection, cells were trypsinized and 5 x 10^5^ cells were seeded into T150 flasks. DNA was collected from remaining cells and this sample was named as (Day 9). T150 flasks were cultured until confluent and then DNA was collected. The time for cells to become confluent varied for each amino acid residue (Day 16 – 40, **Appendix 1-table 5**). DNA was extracted from PANC-1 cells using the PureLink Genomic DNA Mini Kit (Invitrogen, Carlsbad, CA; catalog no. K1820-01). The assay for CellTag library was repeated in triplicate. We repeated our CDKN2A assay in duplicate for 28 residues. For the remaining 128 CDKN2A residues the assay was completed once.

### Generation of sequence libraries

Library preparation and sequencing was performed as previously described with the following modifications (Kinde et al., 2011). For the 1^st^ stage PCR, 3 target specific primers were designed to amplify *CDKN2A* amino acid positions 1 to 53, 54 to 110, and 111 to 156 (**Appendix 1-table 13**). Forward and reverse 1^st^ stage primers contained 5’ M13F (GTAAAACGACGGCCAGC) and M13R (CAGGAAACAGCTATGAC) sequence, respectively, to enable amplification and ligation of Illumina adapter sequences in a 2^nd^ stage PCR (**Appendix 1-table 13**). DNA was amplified with Q5 Hot Start High-Fidelity 2X Master Mix (New England Biolabs; catalog no. M0494S). For the 1^st^ stage PCR, each DNA sample was amplified in three reactions each containing 66 ng of DNA for 18 cycles. 1^st^ stage PCR products for each sample were then pooled and purified using the Agencourt AMPure XP system (Beckman Coulter, Inc, Brea, CA; catalog no. A63881), eluting into 50 µL of elution buffer. Purified PCR product was amplified in a 2^nd^ stage PCR to add Illumina adaptor sequences and indexes (**Appendix 1-table 13**). 2^nd^ stage PCR Amplification was performed with KAPA HiFi HotStart PCR Kit (Kapa Biosystems, Wilmington, MA; catalog no. KK2501) in 25 µL reactions containing 5X KAPA HiFi Buffer - 5 µL, 10 mM KAPA dNTP Mix - 0.75 µL, 10 μM forward primer - 0.75 µL, 10 μM reverse primer - 0.75 µL. For the 1^st^ stage PCR, 66 ng of template DNA and 12.5 µL, Q5 Hot Start High-Fidelity 2X Master Mix was used with the following cycling conditions: 98 °C for 30 seconds; 18cycles of 98 °C for 10 seconds, 72 °C for 30 seconds, 72 °C for 25 seconds; 72 °C for 2 minutes. For the 2^nd^ stage PCR, 0.25 µL of 1^st^ stage PCR product and 0.5 µL of 1 U/μL KAPA HiFi HotStart DNA Polymerase was used with the following cycling conditions: 95 °C for 3 minutes; 25 cycles of 98 °C for 20 seconds, 62 °C for 15 seconds, 72 °C for 1 minute. 2^nd^ stage PCR products were purified with the Agencourt AMPure XP system (Beckman Coulter, Inc.; catalog no. A63881) into 30 µL of elution buffer. Samples were quantified by Qubit using dsDNA HS assay kit (Invitrogen; catalog no. Q33230).

### Sequencing and analysis

Sequence libraries were pooled in equimolar amounts into groups of 16 samples and sequenced on the Illumina MiSeq System (Illumina, San Diego, CA) with the MiSeq Reagent Kit v2 (300 cycles) (Illumina catalog no. MS-102-2002) to generate 150 base pair paired-end reads. Samples were demultiplexed and FASTQ sequence read files were generated with MiSeq control software 2.5.0.5 (Illumina). Paired sequence reads were then combined into a single contiguous sequence using Paired-End Read Merger (Zhang et al., 2014). Reads supporting each variant at a given amino acid position were counted using perl.

### Functional characterization of *CDKN2A* variants using a gamma generalized linear model

We determined if a variant has a fitness advantage by assessing the significance of the observed ratio r_v,cf_ at confluence between the number of cells with a missense variant *v* and the number of cells with a synonymous variant at a given amino acid position. Using the missense variant as a benchmark variant, we assumed that the distribution of r_v,cf_ can be explained by two key covariates: *r_v,init_*, which represent the missense variant-to-synonymous variant ratio at Day 9, and *p_v,init_*, the proportion of the missense variant cells among other variants, including the synonymous variant, at the studied position. More specifically, given the variables *r_v,init_* and *p_v,init_*, the ratio at confluence follows a distribution:

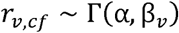

where the mean *u_v_* of the Gamma distribution is such that:

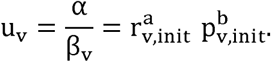

Here, the parameters of the null model to estimate are *α, a, and b*, where *α*, is the shape parameter of the Gamma distribution and is assumed to be the same for all variants. This model is a gamma Generalized Linear Model (GLM) over the response variable *r_v,cf_* with a log-link function and covariates *log*(*r_v,init_*) and *log*(*p_v,init_*). Estimating the parameters will provide a null distribution of *r_v,cf_*, generating a p-value for every observed *r_v,cf_* for any variant at a given position.

To estimate the parameters *α, a*, and *b*, we utilized three control experiments where the CellTag plasmid library was transduced into PANC-1^CDKN2Aco^ cells so that each CellTag represented a neutral variant. For a single experiment, every variant can be considered as wild-type, and we test the other 19 variants against it, knowing that they are neutral and therefore follow the null distribution. This provides us with 19 x 20 triplets (*r_v,cf_*, 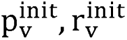), for every experiment, yielding 1140 datapoints when considering all three experiments together. To estimate the parameters using these 1140 data points, we fit the GLM corresponding GLM model using the sklearn.linear_model module.

After the estimation of parameters *α, a*, and *b*, every observation for a tested variant *v* at a given position of the triplet (*r_v,cf_*, 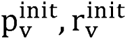) yields a p-value, defined as the probability of observing a ratio at confluence that is at least *r_v,cf_* given *p_v,init_, r_v,init_* under the null Gamma model. As some variants were tested in repeated experiments, we combined their associated p-values into a single p-value using Fisher’s method. Finally, to determine if a variant presents a fitness advantage, we apply a Benjamini-Hochberg estimator on all the tested variants p-values, fixing the False Discovery Rate at a level of 0.05.

### Functional characterization of CDKN2A variants using log_2_ normalized fold change

Fold change for each variant was calculated using the proportion of total reads representing a variant at confluency (Day 16-40, **Appendix 1-table 5**) to the proportion of total reads representing a variant on Day 9 after transfection. Fold change was then normalized to the synonymous variant at each residue and then log_2_ normalized fold change values calculated (**Appendix 1-table 4**, **Appendix 1-table 6**). Variants with log_2_ normalized fold change values greater than or equal to the minimum value of benchmark pathogenic variants were characterized as functionally deleterious, while variants with values smaller than or equal the maximum value of benchmark benign variants were characterized as functionally neutral (**Appendix 1-table 6**). Log_2_ normalized fold change values between these defined thresholds were classified as indeterminate. Mean values were used for replicated variants.

### Data visualization

Heat map of individual variant p-values by amino position was generated using R with the heatmaply package (Galili et al., 2018).

### Cell proliferation assay

Cell proliferation assay were performed as previously described with the following modifications (Kimura et al., 2022). 1 × 10^5^ cells were seeded into in vitro culture (Day 0). Cells were counted on Day 14 using a TC20 Automated Cell Counter (Bio-Rad Laboratories, Herclues, CA; catalog no. 1450102). Relative cell proliferation value was calculated as cell number normalized to empty vector control. Assays were repeated in triplicate. Mean cell proliferation value and standard deviation (s.d.) were calculated.

### Variant effect predictions

Publicly available algorithms were used to predict the consequence of *CDKN2A* missense variants. Prediction algorithms used included: CADD (Kircher et al., 2014), PolyPhen-2 (Adzhubei et al., 2010), SIFT (Kumar et al., 2009), VEST (Carter et al., 2013), AlphaMissense (Cheng et al., 2023), ESM1b (Brandes et al., 2023), and PrimateAI-3D (Gao et al., 2023) (**Appendix 1-table 7**). PolyPhen-2, SIFT, VEST, AlphaMissense, and ESM1b prediction were available for all missense variants. CADD scores were available for 910 missense variants and where multiple CADD scores were possible, mean values were used. PrimateAI-3D prediction scores were available for 904 assayed missense variants.

### Statistical analyses

Statistical analyses were performed using JMP v.11 (SAS, Cary, NC) and Python statsmodel package (version 0.14.0). Student’s t-tests was used to compare mean cell proliferation values. A chi-square test was used to compare the proportion of functionally deleterious variants for variants present in < 2% and ≥ 2% of the cell pool at Day 9. A Fisher’s exact test was used to compare prevalence of functionally deleterious *CDKN2A* variants in colorectal cancer cases from COSMIC with and without somatic mutations in mismatch repair genes. Z-tests with multiple test correction performed with the Bonferroni method was used in the following comparisons: 1) proportion of functionally deleterious variants present in < 2% of the cell pool and ≥ 2% of the cell pool at Day 9 binned in 1% intervals, 2) proportion of variants in each domain predicted to have deleterious or pathogenic effect by the majority of algorithms, 3) proportion of functionally deleterious variants in each domain, and 4) proportion of functionally deleterious missense variants and somatic mutations.

## Funding

National Institutes of Health grant P50CA62924 (NJR)

Susan Wojcicki and Dennis Troper (NR)

The Sol Goldman Pancreatic Cancer Research Center (NJR)

The Rolfe Pancreatic Cancer Foundation (NJR)

The Japanese Society of Gastroenterology Support for Young Gastroenterologists Studying in the United States (HK)

The Japan Society for the Promotion of Science Overseas Research Fellowships (HK)

## Author contributions

Conceptualization: HK, KL, CT, NJR

Resources: CT, NJR

Data curation: HK, KL, CT, NJR

Formal analysis: HK, KL, CT, NJR

Investigation: HK, KL, CT, NJR

Visualization: HK

Methodology: HK, KL, CT, NJR

Writing-original draft: HK, NJR

Project administration: NJR

Writing-review and editing: HK, KL, CT, NJR

## Data and materials availability

All data are available in the main text or the supplementary materials.

## Supplementary Information

## Figures

**Figure 1-figure supplement 1**. **Development and validation of high-throughput CDKN2A functional assay.** (**A**) Cell proliferation of PANC-1 cells stably expressing empty expression vector, codon optimized CDKN2A, one of three synonymous variants (p.L32L, p.G101G, p.V126V), or one of three pathogenic variants (p.L32P, p.G101W, p.V126D) over 14 days in culture. Cell proliferation values are given as mean of three repeats ± standard deviation normalized to PANC-1 cells that stably express empty vector. Statistically significant inhibition of cell proliferation inhibition in PANC-1 cells that stably express synonymous variants (*; P value < 0.001). (**B**) PANC-1 cells stably expressing codon optimized CDKN2A transduced with a CellTag lentiviral library of 20 nonfunctional barcodes were cultured and representation (percent of reads supporting each barcode) before (Day 9) and after a period of in vitro cell proliferation (Day 45) was determined using next generation sequencing. Percent values are given as the mean of three repeats ± standard deviation.

**Figure 1-figure supplement 2. Data for CDKN2A plasmid library.** (**A**) Dot plot showing proportion of each variant per residue in the plasmid libraries. (**B**) Variant proportion in plasmid libraries grouped in 0.5% increments. (**C**) Dot plot showing variant proportion in the amplified plasmid library compared to the Day 9 cell pool. (**D**) Normalized fold change of variant proportion between Day 9 cell pool and the amplified plasmid library based on ACMG classification.

**Figure 2-figure supplement 1. P values for all possible *CDKN2A* missense variants.** (**A**) Distribution of log_2_ P values for all possible *CDKN2A* missense variants. (**B**) Distribution of log_2_ P values for benchmark pathogenic variants (red box), benchmark benign variants (blue box), VUSs previously reported to have functionally deleterious effects (orange box), and VUSs previously reported to have functionally neutral effects (green box). (**C**) Dot plot showing log_2_ P value of all possible *CDKN2A* missense variants pre residue.

**Figure 2-figure supplement 2. Normalized fold change for all possible *CDKN2A* missense variants.** (**A**) Dot plot showing log_2_ normalized fold change of all possible *CDKN2A* missense variants by residue (**B**) Log_2_ normalized fold change for 32 benchmark pathogenic variants, 6 benign variants, 31 VUSs previously reported to have functionally deleterious effects, and 18 VUSs previously reported to have functionally neutral effects. (**C**) Functional classifications for 3,120 *CDKN2A* variants, including 2,964 missense variants and 156 synonymous variants. Variants were classified as functionally deleterious, indeterminate function, or neutral based on log_2_ normalized fold change. (**D**) Comparison of functional classification of all possible *CDKN2A* missense variants by log_2_ P value (gamma GLM) and log normalized fold change.

**Figure 2-figure supplement 3. Reproducibility of CDKN2A assay.** (**A**) Dot plot showing log_2_ P value for 560 *CDKN2A* missense variants assayed in duplicate. (**B**) Comparison of functional classifications for 560 *CDKN2A* missense variants assayed in duplicate. (**C**) Dot plot showing log_2_ normalized fold change for 560 *CDKN2A* missense variants assayed in duplicate.

**Figure 2-figure supplement 4. Proportion of variants Day 9.** (**A**) Proportion of all possible 2,964 *CDKN2A* missense variants in the Day 9 cell pool (replicate 1 if duplicated). (**B**) Percent of functionally deleterious variants (black box), variants of indeterminate function, and functionally neutral variants (white box) by variant proportion in the Day 9 cell pool (replicate 1 if duplicated). Left graph variants grouped as < 2% and ≥ 2% in Day 9 Cell Pool. Right graph, variants grouped as < 2%, 1% intervals from 2% to 8%, ≥ 8% in the Day 9 cell pool.

**Figure 2**-**figure supplement 5. Functional characterization of all possible *CDKN2A* missense variants by ankyrin domain and residue.** (**A**) Schematic representation of CDKN2A with ankyrin repeats 1-4 represented. (**B**) Percent of functionally deleterious (black box), indeterminate function (gray box), and functionally neutral variants (white box) within ankyrin repeats and non-ankyrin repeat regions of CDKN2A. Ank; Ankyrin repeat. (**C**) Dot plot showing distribution of percent functionally deleterious missense variants per residue.

**Figure 3-figure supplement 1. Combinational prediction for 7 algorithms.** (**A**) Number of algorithms predicting deleterious effect for 904 *CDKN2A* missense variants with predictions from 7 algorithms. (**B**) Percent of functionally deleterious (black box) and indeterminate function or functionally neutral (white box) variants grouped by the number of algorithms predicting deleterious effect. (**C**) Number of algorithms predicting deleterious effect for 904 *CDKN2A* missense variants grouped by ankyrin repeats and non-ankyrin repeat regions. (**D** - **H**) Percent of functionally deleterious (black box) and indeterminate function or functionally neutral (white box) variants grouped by the number of algorithms predicting deleterious effect in Ank1 (**D**), Ank2 (**E**), Ank3 (**F**), Ank4 (**G**), and non-ankyrins repeat regions (**H**) of CDKN2A.

**Figure 3-figure supplement 2. Combinational prediction for 5 algorithms.** (**A**) Number of algorithms predicting deleterious effect for 2,060 *CDKN2A* missense variants with predictions from 5 algorithms. (**B**) Percent of functionally deleterious (black box) and indeterminate function or functionally neutral (white box) variants grouped by the number of algorithms predicting deleterious effect. (**C**) Number of algorithms predicting deleterious effect for 2,060 *CDKN2A* missense variants grouped by ankyrin repeats and non-ankyrin repeat regions. (**D** - **H**) Percent of functionally deleterious (black box) and indeterminate function or functionally neutral (white box) variants grouped by the number of algorithms predicting deleterious effect in Ank1 (**D**), Ank2 (**E**), Ank3 (**F**), Ank4 (**G**), and non-ankyrins repeat regions (**H**) of CDKN2A.

**Figure 4-figure supplement 1. Missense somatic mutations in *CDKN2A*.** (**A**) Percent of missense somatic mutations in *CDKN2A* reported in either COSMIC, TCGA, JHU, or MSK-IMPACT that were classified as pathogenic or likely pathogenic (black box), VUS (gray box), or benign or likely benign (white box) using ACMG interpretation guidelines. (**B**) Percent of missense somatic mutations in *CDKN2A* that were classified as pathogenic or likely pathogenic (black box), VUS (gray box), or benign or likely benign (white box) using ACMG interpretation guidelines grouped by mutation database. (**C**) Number of patients with a pathogenic or likely pathogenic missense somatic mutation grouped by mutation database. Patients with p.His83Tyr mutation (black box), patients with p.Asp84Asn mutations (grep box), and patients with other mutations highlighted. COSMIC; the Catalogue Of Somatic Mutations In Cancer, TCGA; The Cancer Genome Atlas, JHU; The Johns Hopkins University School of Medicine, MSK-IMPACT; Memorial Sloan Kettering-Integrated Mutation Profiling of Actionable Cancer Targets.

**Figure 4-figure supplement 2. Functional classification of missense somatic mutations in *CDKN2A*.** Percent of missense somatic mutations in *CDKN2A* reported in either COSMIC (A), TCGA (B), JHU (C), or MSK-IMPACT (D) that were classified as functionally deleterious (black box), indeterminate (gray box), or functionally neutral (white box) in our CDKN2A functional assay grouped by tumor type. The number of missense somatic mutations for each tumor type given in parentheses. COSMIC; the Catalogue Of Somatic Mutations In Cancer, TCGA; The Cancer Genome Atlas, JHU; The Johns Hopkins University School of Medicine, MSK-IMPACT; Memorial Sloan Kettering-Integrated Mutation Profiling of Actionable Cancer Targets.

## Tables

Appendix 1-table 1. Assay outputs for CellTag experiments.

Appendix 1-table 2. Proportion of each variant in the initial plasmid library.

Appendix 1-table 3. Proportion of each variant in residues R24, H66, and A127.

Appendix 1-table 4. Assay outputs and functional classifications for all possible *CDKN2A* missense and synonymous variants.

Appendix 1-table 5. Day of confluency by experiment and residue.

Appendix 1-table 6. Normalized fold change for all possible *CDKN2A* missense and synonymous variants.

Appendix 1-table 7. In silico variant effect predictions for *CDKN2A* missense variants.

Appendix 1-table 8. Assessment of in silico variant effect prediction models.

Appendix 1-table 9. Missense somatic mutations in *CDKN2A* reported in COSMIC, TCGA, JHU, MSK-IMPACT.

Appendix 1-table 10. *CDKN2A* missense and synonymous variants reported in gnomAD.

Appendix 1-table 11. *CDKN2A* missense VUSs reported in ClinVar.

Appendix 1-table 12. Codon optimized *CDKN2A* sequence.

Appendix 1-table 13. Sequences of primers used in study.

## Source data

Figure 1-source data

Raw data in Figure 1

Figure 1-figure supplement 1-source data 1

Raw data in Figure 1-figure supplement 1A

Figure 1-figure supplement 1-source data 2

Raw data in Figure 1-figure supplement 1B

Figure 2-source data 1

Raw data in Figure 2B

Figure 2-figure supplement 1-source data 1

Raw data in Figure 2-figure supplement 1A and 1B

Figure 2-figure supplement 2-source data 1

Raw data in Figure 2-figure supplement 2D

Figure 2-figure supplement 3-source data 1

Raw data in Figure 2-figure supplement 3A

Figure 2-figure supplement 3-source data 2

Raw data in Figure 2-figure supplement 3B

Figure 2-figure supplement 4-source data 1

Raw data in Figure 2-figure supplement 4

Figure 2-figure supplement 5-source data 1

Raw data in Figure 2-figure supplement 5B

Figure 2-figure supplement 5-source data 2

Raw data in Figure 2-figure supplement 5C

Figure 3-source data 1

Raw data in Figure 3

Figure 3-figure supplement 1-source data 1

Raw data in Figure 3-figure supplement 1A and 1B

Figure 3-figure supplement 1-source data 2

Raw data in Figure 3-figure supplement 1C

Figure 3-figure supplement 1-source data 3

Raw data in Figure 3-figure supplement 1D - 1H

Figure 3-figure supplement 2-source data 1

Raw data in Figure 3-figure supplement 2A and 2B

Figure 3-figure supplement 2-source data 2

Raw data in Figure 3-figure supplement 2C

Figure 3-figure supplement 2-source data 3

Raw data in Figure 3-figure supplement 2D - 2H

Figure 4-source data 1

Raw data in Figure 4A and 4B

Figure 4-source data 2

Raw data in Figure 4C

Figure 4-figure supplement 1-source data 1

Raw data in Figure 4-figure supplement 1A and 1B

Figure 4-figure supplement 1-source data 2

Raw data in Figure 4-figure supplement 1C

Figure 4-figure supplement 2-source data 1

Raw data in Figure 4-figure supplement 2A - 2D

---

**Appendix 1-table 8.**
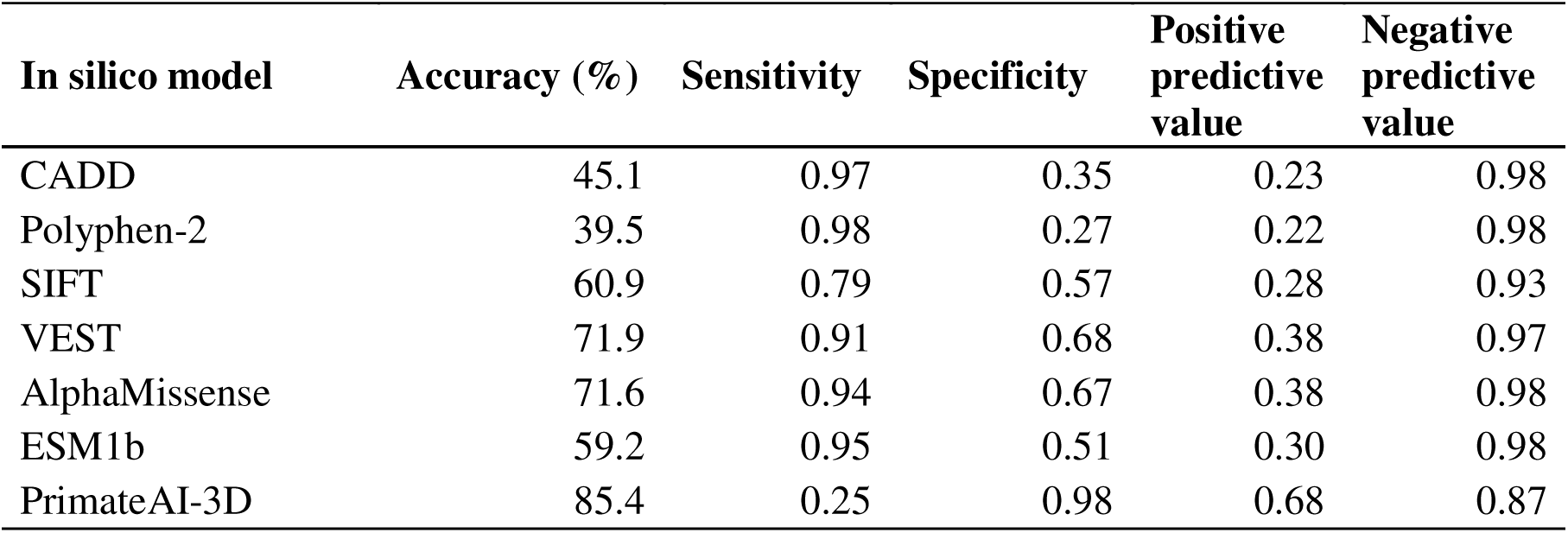
Assessment of in silico variant effect prediction models.

**Appendix 1-table 12.**
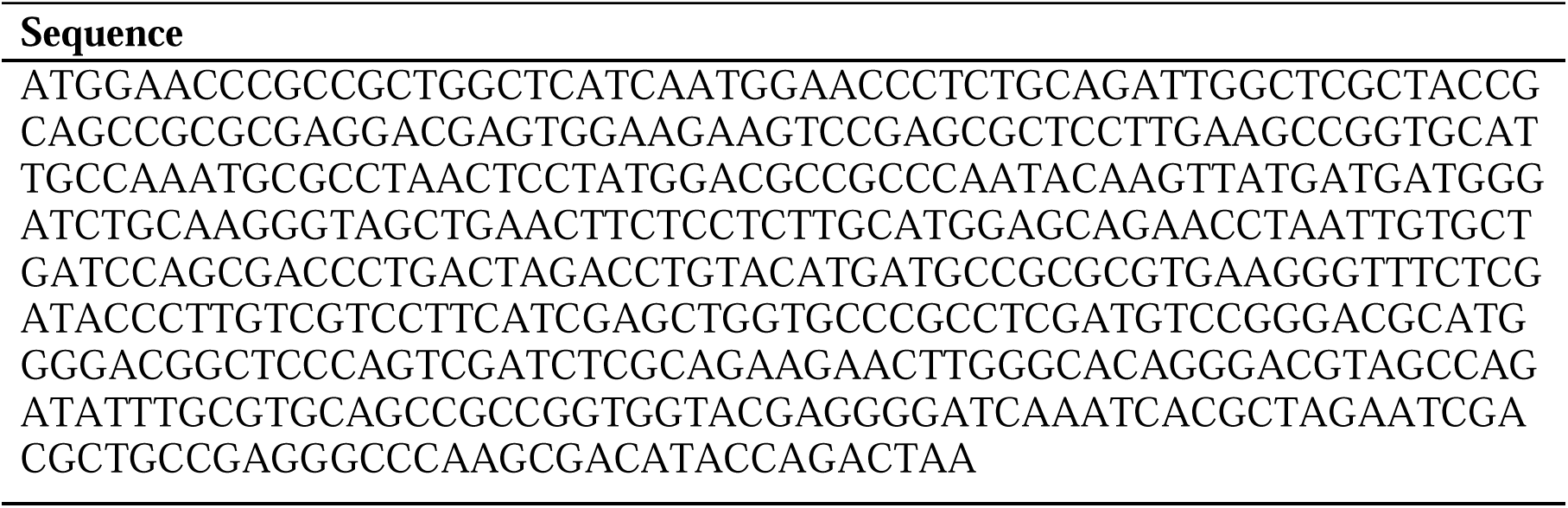
Codon optimized *CDKN2A* sequence.

