## Appendix 1-table 1 for "Functional characterization of all *CDKN2A* missense variants and comparison to in silico models of pathogenicity"

**Appendix 1-table 1. Assay outputs for CellTag experiments.**

| CellTag number | Experiment 1 | | | | Experiment 2 | | | | Experiment 3 | | | | Proprotion | | | |
|---|---|---|---|---|---|---|---|---|---|---|---|---|---|---|---|---|
| | Read count | | Proportion | | Read count | | Proportion | | Read count | | Proportion | | | | | |
| | Day 9 | Day 45 | Day 9 | Day 45 | Day 9 | Day 45 | Day 9 | Day 45 | Day 9 | Day 45 | Day 9 | Day 45 | Mean Day 9 | Mean Day 45 | Standard deviation Day 9 | Standard deviation Day 45 |
| CellTag1 | 18886 | 7358 | 4.68 | 4.40 | 9135 | 7518 | 4.49 | 4.24 | 9860 | 6608 | 4.38 | 4.42 | 4.52 | 4.35 | 0.15 | 0.10 |
| CellTag2 | 22571 | 9930 | 5.60 | 5.93 | 11498 | 10239 | 5.65 | 5.78 | 12373 | 8001 | 5.50 | 5.35 | 5.58 | 5.69 | 0.08 | 0.30 |
| CellTag3 | 18731 | 8318 | 4.65 | 4.97 | 9439 | 9057 | 4.64 | 5.11 | 11058 | 7363 | 4.91 | 4.93 | 4.73 | 5.00 | 0.16 | 0.10 |
| CellTag4 | 25031 | 10273 | 6.21 | 6.14 | 13022 | 10865 | 6.40 | 6.13 | 13515 | 9318 | 6.00 | 6.24 | 6.20 | 6.17 | 0.20 | 0.06 |
| CellTag5 | 18034 | 8229 | 4.47 | 4.92 | 9654 | 8455 | 4.74 | 4.77 | 10665 | 6944 | 4.74 | 4.65 | 4.65 | 4.78 | 0.15 | 0.13 |
| CellTag6 | 21130 | 8923 | 5.24 | 5.33 | 10702 | 9912 | 5.26 | 5.60 | 12738 | 8179 | 5.66 | 5.47 | 5.39 | 5.47 | 0.24 | 0.13 |
| CellTag7 | 17421 | 8023 | 4.32 | 4.79 | 9086 | 8475 | 4.46 | 4.79 | 9867 | 7113 | 4.38 | 4.76 | 4.39 | 4.78 | 0.07 | 0.02 |
| CellTag8 | 19780 | 8156 | 4.91 | 4.87 | 9708 | 8955 | 4.77 | 5.06 | 11174 | 7404 | 4.96 | 4.95 | 4.88 | 4.96 | 0.10 | 0.09 |
| CellTag9 | 20872 | 8931 | 5.18 | 5.33 | 10628 | 9194 | 5.22 | 5.19 | 11859 | 7849 | 5.27 | 5.25 | 5.22 | 5.26 | 0.05 | 0.07 |
| CellTag10 | 19441 | 7610 | 4.82 | 4.55 | 10308 | 7461 | 5.07 | 4.21 | 10458 | 7297 | 4.65 | 4.88 | 4.84 | 4.55 | 0.21 | 0.34 |
| CellTag11 | 24278 | 10397 | 6.02 | 6.21 | 12258 | 11333 | 6.02 | 6.40 | 14086 | 9108 | 6.26 | 6.09 | 6.10 | 6.23 | 0.14 | 0.15 |
| CellTag12 | 26268 | 11069 | 6.52 | 6.61 | 13209 | 11494 | 6.49 | 6.49 | 14782 | 9871 | 6.57 | 6.61 | 6.52 | 6.57 | 0.04 | 0.07 |
| CellTag13 | 18378 | 7440 | 4.56 | 4.44 | 8926 | 8197 | 4.39 | 4.63 | 9626 | 6363 | 4.28 | 4.26 | 4.41 | 4.44 | 0.14 | 0.19 |
| CellTag14 | 26288 | 9801 | 6.52 | 5.85 | 12418 | 11095 | 6.10 | 6.26 | 14185 | 8737 | 6.30 | 5.85 | 6.31 | 5.99 | 0.21 | 0.24 |
| CellTag15 | 23881 | 9917 | 5.92 | 5.92 | 12073 | 11203 | 5.93 | 6.33 | 13775 | 8986 | 6.12 | 6.01 | 5.99 | 6.09 | 0.11 | 0.21 |
| CellTag16 | 15131 | 6019 | 3.75 | 3.60 | 7421 | 6208 | 3.65 | 3.51 | 8175 | 5532 | 3.63 | 3.70 | 3.68 | 3.60 | 0.07 | 0.10 |
| CellTag17 | 5094 | 1861 | 1.26 | 1.11 | 2598 | 1787 | 1.28 | 1.01 | 2788 | 1607 | 1.24 | 1.08 | 1.26 | 1.07 | 0.02 | 0.05 |
| CellTag18 | 25447 | 10259 | 6.31 | 6.13 | 12235 | 10417 | 6.01 | 5.88 | 13537 | 9539 | 6.01 | 6.38 | 6.11 | 6.13 | 0.17 | 0.25 |
| CellTag19 | 25280 | 10523 | 6.27 | 6.29 | 13419 | 10627 | 6.59 | 6.00 | 14407 | 9221 | 6.40 | 6.17 | 6.42 | 6.15 | 0.16 | 0.14 |
| CellTag20 | 11225 | 4371 | 2.78 | 2.61 | 5774 | 4611 | 2.84 | 2.60 | 6170 | 4396 | 2.74 | 2.94 | 2.79 | 2.72 | 0.05 | 0.19 |
