## Appendix 1-table 2 for "Functional characterization of all *CDKN2A* missense variants and comparison to in silico models of pathogenicity"

**Appendix 1-table 2. Proportion of each variant in the initial plasmid library.**

| Residue | Variant | Respresentation in library (%) |
|---|---|---|
| 1 | p.Met1Asn | 5.0 |
| 1 | p.Met1Lys | 6.7 |
| 1 | p.Met1Thr | 5.2 |
| 1 | p.Met1Arg | 4.5 |
| 1 | p.Met1Ser | 4.6 |
| 1 | p.Met1Ile | 6.2 |
| 1 | p.Met1Met | 0.0 |
| 1 | p.Met1His | 4.6 |
| 1 | p.Met1Gln | 4.2 |
| 1 | p.Met1Pro | 4.5 |
| 1 | p.Met1Leu | 10.6 |
| 1 | p.Met1Asp | 4.3 |
| 1 | p.Met1Glu | 4.4 |
| 1 | p.Met1Ala | 4.6 |
| 1 | p.Met1Gly | 4.1 |
| 1 | p.Met1Val | 8.6 |
| 1 | p.Met1Tyr | 4.4 |
| 1 | p.Met1Cys | 4.5 |
| 1 | p.Met1Trp | 4.5 |
| 1 | p.Met1Phe | 4.6 |
| 2 | p.Glu2Asn | 3.3 |
| 2 | p.Glu2Lys | 3.4 |
| 2 | p.Glu2Thr | 2.6 |
| 2 | p.Glu2Arg | 2.8 |
| 2 | p.Glu2Ser | 2.0 |
| 2 | p.Glu2Ile | 3.9 |
| 2 | p.Glu2Met | 2.9 |
| 2 | p.Glu2His | 3.7 |
| 2 | p.Glu2Gln | 4.8 |
| 2 | p.Glu2Pro | 3.7 |
| 2 | p.Glu2Leu | 3.4 |
| 2 | p.Glu2Asp | 26.3 |
| 2 | p.Glu2Glu | 12.6 |
| 2 | p.Glu2Ala | 4.1 |
| 2 | p.Glu2Gly | 3.4 |
| 2 | p.Glu2Val | 3.4 |
| 2 | p.Glu2Tyr | 2.8 |
| 2 | p.Glu2Cys | 3.3 |
| 2 | p.Glu2Trp | 3.7 |
| 2 | p.Glu2Phe | 3.8 |
| 3 | p.Pro3Asn | 4.7 |

| | | |
|---|---|---|
| 3 | p.Pro3Lys | 3.5 |
| 3 | p.Pro3Thr | 7.5 |
| 3 | p.Pro3Arg | 4.2 |
| 3 | p.Pro3Ser | 4.3 |
| 3 | p.Pro3Ile | 3.9 |
| 3 | p.Pro3Met | 3.3 |
| 3 | p.Pro3His | 9.4 |
| 3 | p.Pro3Gln | 3.6 |
| 3 | p.Pro3Pro | 9.6 |
| 3 | p.Pro3Leu | 4.3 |
| 3 | p.Pro3Asp | 4.6 |
| 3 | p.Pro3Glu | 4.5 |
| 3 | p.Pro3Ala | 6.0 |
| 3 | p.Pro3Gly | 5.4 |
| 3 | p.Pro3Val | 4.7 |
| 3 | p.Pro3Tyr | 3.4 |
| 3 | p.Pro3Cys | 4.4 |
| 3 | p.Pro3Trp | 3.8 |
| 3 | p.Pro3Phe | 4.9 |
| 4 | p.Ala4Asn | 4.8 |
| 4 | p.Ala4Lys | 3.4 |
| 4 | p.Ala4Thr | 8.7 |
| 4 | p.Ala4Arg | 4.3 |
| 4 | p.Ala4Ser | 4.1 |
| 4 | p.Ala4Ile | 3.4 |
| 4 | p.Ala4Met | 3.6 |
| 4 | p.Ala4His | 5.8 |
| 4 | p.Ala4Gln | 3.9 |
| 4 | p.Ala4Pro | 4.6 |
| 4 | p.Ala4Leu | 3.9 |
| 4 | p.Ala4Asp | 10.1 |
| 4 | p.Ala4Glu | 3.7 |
| 4 | p.Ala4Ala | 8.5 |
| 4 | p.Ala4Gly | 6.0 |
| 4 | p.Ala4Val | 5.2 |
| 4 | p.Ala4Tyr | 3.6 |
| 4 | p.Ala4Cys | 4.2 |
| 4 | p.Ala4Trp | 4.5 |
| 4 | p.Ala4Phe | 3.7 |
| 5 | p.Ala5Asn | 6.2 |
| 5 | p.Ala5Lys | 4.0 |
| 5 | p.Ala5Thr | 4.5 |
| 5 | p.Ala5Arg | 4.5 |
| 5 | p.Ala5Ser | 4.3 |
| 5 | p.Ala5Ile | 6.0 |

| | | |
|---|---|---|
| 5 | p.Ala5Met | 4.4 |
| 5 | p.Ala5His | 5.1 |
| 5 | p.Ala5Gln | 5.7 |
| 5 | p.Ala5Pro | 5.1 |
| 5 | p.Ala5Leu | 6.4 |
| 5 | p.Ala5Asp | 5.7 |
| 5 | p.Ala5Glu | 4.5 |
| 5 | p.Ala5Ala | 7.9 |
| 5 | p.Ala5Gly | 3.6 |
| 5 | p.Ala5Val | 6.2 |
| 5 | p.Ala5Tyr | 2.8 |
| 5 | p.Ala5Cys | 3.6 |
| 5 | p.Ala5Trp | 5.9 |
| 5 | p.Ala5Phe | 3.4 |
| 6 | p.Gly6Asn | 4.3 |
| 6 | p.Gly6Lys | 4.5 |
| 6 | p.Gly6Thr | 4.0 |
| 6 | p.Gly6Arg | 4.0 |
| 6 | p.Gly6Ser | 6.7 |
| 6 | p.Gly6Ile | 2.7 |
| 6 | p.Gly6Met | 4.3 |
| 6 | p.Gly6His | 2.7 |
| 6 | p.Gly6Gln | 3.5 |
| 6 | p.Gly6Pro | 3.6 |
| 6 | p.Gly6Leu | 4.3 |
| 6 | p.Gly6Asp | 8.4 |
| 6 | p.Gly6Glu | 3.8 |
| 6 | p.Gly6Ala | 6.6 |
| 6 | p.Gly6Gly | 10.0 |
| 6 | p.Gly6Val | 5.0 |
| 6 | p.Gly6Tyr | 3.3 |
| 6 | p.Gly6Cys | 11.0 |
| 6 | p.Gly6Trp | 4.3 |
| 6 | p.Gly6Phe | 3.1 |
| 7 | p.Ser7Asn | 5.6 |
| 7 | p.Ser7Lys | 5.9 |
| 7 | p.Ser7Thr | 7.3 |
| 7 | p.Ser7Arg | 6.7 |
| 7 | p.Ser7Ser | 3.4 |
| 7 | p.Ser7Ile | 5.6 |
| 7 | p.Ser7Met | 7.3 |
| 7 | p.Ser7His | 6.7 |
| 7 | p.Ser7Gln | 6.1 |
| 7 | p.Ser7Pro | 8.9 |
| 7 | p.Ser7Leu | 6.6 |

| | | |
|---|---|---|
| 7 | p.Ser7Asp | 3.7 |
| 7 | p.Ser7Glu | 1.5 |
| 7 | p.Ser7Ala | 4.3 |
| 7 | p.Ser7Gly | 0.9 |
| 7 | p.Ser7Val | 2.6 |
| 7 | p.Ser7Tyr | 4.1 |
| 7 | p.Ser7Cys | 4.3 |
| 7 | p.Ser7Trp | 3.6 |
| 7 | p.Ser7Phe | 4.8 |
| 8 | p.Ser8Asn | 5.5 |
| 8 | p.Ser8Lys | 4.3 |
| 8 | p.Ser8Thr | 6.0 |
| 8 | p.Ser8Arg | 5.9 |
| 8 | p.Ser8Ser | 6.1 |
| 8 | p.Ser8Ile | 6.2 |
| 8 | p.Ser8Met | 3.4 |
| 8 | p.Ser8His | 3.7 |
| 8 | p.Ser8Gln | 4.7 |
| 8 | p.Ser8Pro | 4.8 |
| 8 | p.Ser8Leu | 4.8 |
| 8 | p.Ser8Asp | 5.6 |
| 8 | p.Ser8Glu | 4.6 |
| 8 | p.Ser8Ala | 4.2 |
| 8 | p.Ser8Gly | 4.7 |
| 8 | p.Ser8Val | 5.6 |
| 8 | p.Ser8Tyr | 4.4 |
| 8 | p.Ser8Cys | 4.5 |
| 8 | p.Ser8Trp | 6.1 |
| 8 | p.Ser8Phe | 4.6 |
| 9 | p.Met9Asn | 4.6 |
| 9 | p.Met9Lys | 7.0 |
| 9 | p.Met9Thr | 5.1 |
| 9 | p.Met9Arg | 6.4 |
| 9 | p.Met9Ser | 4.0 |
| 9 | p.Met9Ile | 5.9 |
| 9 | p.Met9Met | 0.0 |
| 9 | p.Met9His | 3.1 |
| 9 | p.Met9Gln | 4.1 |
| 9 | p.Met9Pro | 3.9 |
| 9 | p.Met9Leu | 6.8 |
| 9 | p.Met9Asp | 4.9 |
| 9 | p.Met9Glu | 4.6 |
| 9 | p.Met9Ala | 4.6 |
| 9 | p.Met9Gly | 4.2 |
| 9 | p.Met9Val | 8.6 |

| | | |
|---|---|---|
| 9 | p.Met9Tyr | 6.1 |
| 9 | p.Met9Cys | 5.6 |
| 9 | p.Met9Trp | 5.2 |
| 9 | p.Met9Phe | 5.5 |
| 10 | p.Glu10Asn | 3.5 |
| 10 | p.Glu10Lys | 4.9 |
| 10 | p.Glu10Thr | 2.5 |
| 10 | p.Glu10Arg | 4.9 |
| 10 | p.Glu10Ser | 2.6 |
| 10 | p.Glu10Ile | 3.4 |
| 10 | p.Glu10Met | 5.9 |
| 10 | p.Glu10His | 3.2 |
| 10 | p.Glu10Gln | 4.8 |
| 10 | p.Glu10Pro | 4.8 |
| 10 | p.Glu10Leu | 3.2 |
| 10 | p.Glu10Asp | 13.4 |
| 10 | p.Glu10Glu | 10.8 |
| 10 | p.Glu10Ala | 2.9 |
| 10 | p.Glu10Gly | 4.4 |
| 10 | p.Glu10Val | 4.6 |
| 10 | p.Glu10Tyr | 5.7 |
| 10 | p.Glu10Cys | 5.8 |
| 10 | p.Glu10Trp | 4.0 |
| 10 | p.Glu10Phe | 4.5 |
| 11 | p.Pro11Asn | 4.8 |
| 11 | p.Pro11Lys | 3.7 |
| 11 | p.Pro11Thr | 4.9 |
| 11 | p.Pro11Arg | 4.0 |
| 11 | p.Pro11Ser | 4.9 |
| 11 | p.Pro11Ile | 5.1 |
| 11 | p.Pro11Met | 5.4 |
| 11 | p.Pro11His | 7.0 |
| 11 | p.Pro11Gln | 5.1 |
| 11 | p.Pro11Pro | 5.9 |
| 11 | p.Pro11Leu | 4.3 |
| 11 | p.Pro11Asp | 5.6 |
| 11 | p.Pro11Glu | 4.6 |
| 11 | p.Pro11Ala | 6.1 |
| 11 | p.Pro11Gly | 6.0 |
| 11 | p.Pro11Val | 5.3 |
| 11 | p.Pro11Tyr | 4.6 |
| 11 | p.Pro11Cys | 3.8 |
| 11 | p.Pro11Trp | 4.7 |
| 11 | p.Pro11Phe | 4.4 |
| 12 | p.Ser12Asn | 3.8 |

| | | |
|---|---|---|
| 12 | p.Ser12Lys | 4.8 |
| 12 | p.Ser12Thr | 4.0 |
| 12 | p.Ser12Arg | 4.9 |
| 12 | p.Ser12Ser | 4.5 |
| 12 | p.Ser12Ile | 4.1 |
| 12 | p.Ser12Met | 6.3 |
| 12 | p.Ser12His | 4.9 |
| 12 | p.Ser12Gln | 6.4 |
| 12 | p.Ser12Pro | 5.9 |
| 12 | p.Ser12Leu | 6.5 |
| 12 | p.Ser12Asp | 5.4 |
| 12 | p.Ser12Glu | 4.6 |
| 12 | p.Ser12Ala | 3.5 |
| 12 | p.Ser12Gly | 4.4 |
| 12 | p.Ser12Val | 3.5 |
| 12 | p.Ser12Tyr | 4.8 |
| 12 | p.Ser12Cys | 5.3 |
| 12 | p.Ser12Trp | 6.1 |
| 12 | p.Ser12Phe | 6.2 |
| 13 | p.Ala13Asn | 4.9 |
| 13 | p.Ala13Lys | 4.3 |
| 13 | p.Ala13Thr | 5.1 |
| 13 | p.Ala13Arg | 6.3 |
| 13 | p.Ala13Ser | 5.3 |
| 13 | p.Ala13Ile | 4.6 |
| 13 | p.Ala13Met | 5.8 |
| 13 | p.Ala13His | 3.5 |
| 13 | p.Ala13Gln | 4.9 |
| 13 | p.Ala13Pro | 5.2 |
| 13 | p.Ala13Leu | 4.6 |
| 13 | p.Ala13Asp | 5.7 |
| 13 | p.Ala13Glu | 5.5 |
| 13 | p.Ala13Ala | 8.6 |
| 13 | p.Ala13Gly | 4.6 |
| 13 | p.Ala13Val | 4.0 |
| 13 | p.Ala13Tyr | 4.5 |
| 13 | p.Ala13Cys | 3.7 |
| 13 | p.Ala13Trp | 3.6 |
| 13 | p.Ala13Phe | 5.2 |
| 14 | p.Asp14Asn | 4.3 |
| 14 | p.Asp14Lys | 4.6 |
| 14 | p.Asp14Thr | 4.4 |
| 14 | p.Asp14Arg | 4.8 |
| 14 | p.Asp14Ser | 4.6 |
| 14 | p.Asp14Ile | 4.1 |

| | | |
|---|---|---|
| 14 | p.Asp14Met | 3.6 |
| 14 | p.Asp14His | 3.9 |
| 14 | p.Asp14Gln | 5.6 |
| 14 | p.Asp14Pro | 4.3 |
| 14 | p.Asp14Leu | 5.2 |
| 14 | p.Asp14Asp | 7.1 |
| 14 | p.Asp14Glu | 11.9 |
| 14 | p.Asp14Ala | 4.3 |
| 14 | p.Asp14Gly | 5.6 |
| 14 | p.Asp14Val | 4.6 |
| 14 | p.Asp14Tyr | 4.5 |
| 14 | p.Asp14Cys | 4.4 |
| 14 | p.Asp14Trp | 5.1 |
| 14 | p.Asp14Phe | 3.3 |
| 15 | p.Trp15Asn | 5.6 |
| 15 | p.Trp15Lys | 5.3 |
| 15 | p.Trp15Thr | 3.3 |
| 15 | p.Trp15Arg | 5.8 |
| 15 | p.Trp15Ser | 6.9 |
| 15 | p.Trp15Ile | 3.4 |
| 15 | p.Trp15Met | 2.6 |
| 15 | p.Trp15His | 4.1 |
| 15 | p.Trp15Gln | 5.0 |
| 15 | p.Trp15Pro | 4.1 |
| 15 | p.Trp15Leu | 7.3 |
| 15 | p.Trp15Asp | 7.6 |
| 15 | p.Trp15Glu | 5.4 |
| 15 | p.Trp15Ala | 6.1 |
| 15 | p.Trp15Gly | 4.1 |
| 15 | p.Trp15Val | 3.0 |
| 15 | p.Trp15Tyr | 5.8 |
| 15 | p.Trp15Cys | 7.4 |
| 15 | p.Trp15Trp | 0.0 |
| 15 | p.Trp15Phe | 7.1 |
| 16 | p.Leu16Asn | 4.1 |
| 16 | p.Leu16Lys | 6.6 |
| 16 | p.Leu16Thr | 4.3 |
| 16 | p.Leu16Arg | 4.7 |
| 16 | p.Leu16Ser | 4.6 |
| 16 | p.Leu16Ile | 4.2 |
| 16 | p.Leu16Met | 4.0 |
| 16 | p.Leu16His | 7.6 |
| 16 | p.Leu16Gln | 5.6 |
| 16 | p.Leu16Pro | 10.2 |
| 16 | p.Leu16Leu | 2.8 |

| | | |
|---|---|---|
| 16 | p.Leu16Asp | 3.4 |
| 16 | p.Leu16Glu | 5.4 |
| 16 | p.Leu16Ala | 3.9 |
| 16 | p.Leu16Gly | 4.7 |
| 16 | p.Leu16Val | 5.1 |
| 16 | p.Leu16Tyr | 4.3 |
| 16 | p.Leu16Cys | 4.5 |
| 16 | p.Leu16Trp | 4.8 |
| 16 | p.Leu16Phe | 5.2 |
| 17 | p.Ala17Asn | 4.6 |
| 17 | p.Ala17Lys | 6.3 |
| 17 | p.Ala17Thr | 4.8 |
| 17 | p.Ala17Arg | 5.1 |
| 17 | p.Ala17Ser | 5.6 |
| 17 | p.Ala17Ile | 4.1 |
| 17 | p.Ala17Met | 3.3 |
| 17 | p.Ala17His | 3.7 |
| 17 | p.Ala17Gln | 5.4 |
| 17 | p.Ala17Pro | 6.2 |
| 17 | p.Ala17Leu | 3.1 |
| 17 | p.Ala17Asp | 4.7 |
| 17 | p.Ala17Glu | 4.3 |
| 17 | p.Ala17Ala | 9.9 |
| 17 | p.Ala17Gly | 7.6 |
| 17 | p.Ala17Val | 5.1 |
| 17 | p.Ala17Tyr | 3.9 |
| 17 | p.Ala17Cys | 2.9 |
| 17 | p.Ala17Trp | 4.7 |
| 17 | p.Ala17Phe | 4.9 |
| 18 | p.Thr18Asn | 7.0 |
| 18 | p.Thr18Lys | 5.3 |
| 18 | p.Thr18Thr | 5.9 |
| 18 | p.Thr18Arg | 4.2 |
| 18 | p.Thr18Ser | 4.7 |
| 18 | p.Thr18Ile | 4.8 |
| 18 | p.Thr18Met | 7.2 |
| 18 | p.Thr18His | 2.7 |
| 18 | p.Thr18Gln | 5.7 |
| 18 | p.Thr18Pro | 5.0 |
| 18 | p.Thr18Leu | 6.2 |
| 18 | p.Thr18Asp | 3.8 |
| 18 | p.Thr18Glu | 5.5 |
| 18 | p.Thr18Ala | 6.8 |
| 18 | p.Thr18Gly | 3.1 |
| 18 | p.Thr18Val | 5.9 |

| | | |
|---|---|---|
| 18 | p.Thr18Tyr | 3.4 |
| 18 | p.Thr18Cys | 4.2 |
| 18 | p.Thr18Trp | 5.8 |
| 18 | p.Thr18Phe | 2.7 |
| 19 | p.Ala19Asn | 4.3 |
| 19 | p.Ala19Lys | 4.8 |
| 19 | p.Ala19Thr | 5.3 |
| 19 | p.Ala19Arg | 3.6 |
| 19 | p.Ala19Ser | 4.3 |
| 19 | p.Ala19Ile | 5.1 |
| 19 | p.Ala19Met | 3.5 |
| 19 | p.Ala19His | 6.3 |
| 19 | p.Ala19Gln | 4.7 |
| 19 | p.Ala19Pro | 5.2 |
| 19 | p.Ala19Leu | 5.2 |
| 19 | p.Ala19Asp | 4.2 |
| 19 | p.Ala19Glu | 6.9 |
| 19 | p.Ala19Ala | 11.5 |
| 19 | p.Ala19Gly | 4.4 |
| 19 | p.Ala19Val | 3.7 |
| 19 | p.Ala19Tyr | 4.3 |
| 19 | p.Ala19Cys | 3.4 |
| 19 | p.Ala19Trp | 5.0 |
| 19 | p.Ala19Phe | 4.5 |
| 20 | p.Ala20Asn | 4.7 |
| 20 | p.Ala20Lys | 6.6 |
| 20 | p.Ala20Thr | 6.5 |
| 20 | p.Ala20Arg | 3.8 |
| 20 | p.Ala20Ser | 3.3 |
| 20 | p.Ala20Ile | 4.2 |
| 20 | p.Ala20Met | 5.4 |
| 20 | p.Ala20His | 3.1 |
| 20 | p.Ala20Gln | 4.3 |
| 20 | p.Ala20Pro | 5.4 |
| 20 | p.Ala20Leu | 4.5 |
| 20 | p.Ala20Asp | 7.0 |
| 20 | p.Ala20Glu | 6.7 |
| 20 | p.Ala20Ala | 6.8 |
| 20 | p.Ala20Gly | 4.0 |
| 20 | p.Ala20Val | 5.7 |
| 20 | p.Ala20Tyr | 3.8 |
| 20 | p.Ala20Cys | 5.5 |
| 20 | p.Ala20Trp | 4.5 |
| 20 | p.Ala20Phe | 4.3 |
| 21 | p.Ala21Asn | 2.3 |

| | | |
|---|---|---|
| 21 | p.Ala21Lys | 4.0 |
| 21 | p.Ala21Thr | 3.1 |
| 21 | p.Ala21Arg | 2.7 |
| 21 | p.Ala21Ser | 2.4 |
| 21 | p.Ala21Ile | 1.8 |
| 21 | p.Ala21Met | 4.6 |
| 21 | p.Ala21His | 4.7 |
| 21 | p.Ala21Gln | 5.8 |
| 21 | p.Ala21Pro | 4.1 |
| 21 | p.Ala21Leu | 4.9 |
| 21 | p.Ala21Asp | 4.7 |
| 21 | p.Ala21Glu | 15.6 |
| 21 | p.Ala21Ala | 7.6 |
| 21 | p.Ala21Gly | 5.8 |
| 21 | p.Ala21Val | 10.4 |
| 21 | p.Ala21Tyr | 4.1 |
| 21 | p.Ala21Cys | 4.8 |
| 21 | p.Ala21Trp | 3.7 |
| 21 | p.Ala21Phe | 2.9 |
| 22 | p.Arg22Asn | 5.1 |
| 22 | p.Arg22Lys | 5.0 |
| 22 | p.Arg22Thr | 4.7 |
| 22 | p.Arg22Arg | 39.7 |
| 22 | p.Arg22Ser | 2.8 |
| 22 | p.Arg22Ile | 4.2 |
| 22 | p.Arg22Met | 4.9 |
| 22 | p.Arg22His | 1.5 |
| 22 | p.Arg22Gln | 5.1 |
| 22 | p.Arg22Pro | 1.8 |
| 22 | p.Arg22Leu | 3.4 |
| 22 | p.Arg22Asp | 1.4 |
| 22 | p.Arg22Glu | 3.6 |
| 22 | p.Arg22Ala | 1.5 |
| 22 | p.Arg22Gly | 1.5 |
| 22 | p.Arg22Val | 1.7 |
| 22 | p.Arg22Tyr | 3.8 |
| 22 | p.Arg22Cys | 1.9 |
| 22 | p.Arg22Trp | 4.1 |
| 22 | p.Arg22Phe | 2.4 |
| 23 | p.Gly23Asn | 5.1 |
| 23 | p.Gly23Lys | 4.7 |
| 23 | p.Gly23Thr | 5.4 |
| 23 | p.Gly23Arg | 8.4 |
| 23 | p.Gly23Ser | 4.0 |
| 23 | p.Gly23Ile | 4.4 |

| | | |
|---|---|---|
| 23 | p.Gly23Met | 5.6 |
| 23 | p.Gly23His | 5.1 |
| 23 | p.Gly23Gln | 4.8 |
| 23 | p.Gly23Pro | 4.7 |
| 23 | p.Gly23Leu | 1.6 |
| 23 | p.Gly23Asp | 4.6 |
| 23 | p.Gly23Glu | 5.3 |
| 23 | p.Gly23Ala | 5.3 |
| 23 | p.Gly23Gly | 5.1 |
| 23 | p.Gly23Val | 6.2 |
| 23 | p.Gly23Tyr | 4.8 |
| 23 | p.Gly23Cys | 5.1 |
| 23 | p.Gly23Trp | 4.7 |
| 23 | p.Gly23Phe | 5.1 |
| 24 | p.Arg24Asn | 4.1 |
| 24 | p.Arg24Lys | 4.9 |
| 24 | p.Arg24Thr | 3.9 |
| 24 | p.Arg24Arg | 12.1 |
| 24 | p.Arg24Ser | 5.1 |
| 24 | p.Arg24Ile | 3.6 |
| 24 | p.Arg24Met | 4.6 |
| 24 | p.Arg24His | 4.5 |
| 24 | p.Arg24Gln | 5.4 |
| 24 | p.Arg24Pro | 3.5 |
| 24 | p.Arg24Leu | 4.2 |
| 24 | p.Arg24Asp | 6.6 |
| 24 | p.Arg24Glu | 4.2 |
| 24 | p.Arg24Ala | 4.7 |
| 24 | p.Arg24Gly | 3.3 |
| 24 | p.Arg24Val | 5.7 |
| 24 | p.Arg24Tyr | 4.3 |
| 24 | p.Arg24Cys | 4.2 |
| 24 | p.Arg24Trp | 6.2 |
| 24 | p.Arg24Phe | 4.9 |
| 25 | p.Val25Asn | 3.9 |
| 25 | p.Val25Lys | 5.6 |
| 25 | p.Val25Thr | 4.2 |
| 25 | p.Val25Arg | 7.4 |
| 25 | p.Val25Ser | 4.2 |
| 25 | p.Val25Ile | 3.9 |
| 25 | p.Val25Met | 10.1 |
| 25 | p.Val25His | 2.6 |
| 25 | p.Val25Gln | 2.2 |
| 25 | p.Val25Pro | 1.5 |
| 25 | p.Val25Leu | 5.1 |

| | | |
|---|---|---|
| 25 | p.Val25Asp | 5.9 |
| 25 | p.Val25Glu | 9.8 |
| 25 | p.Val25Ala | 3.3 |
| 25 | p.Val25Gly | 5.3 |
| 25 | p.Val25Val | 6.4 |
| 25 | p.Val25Tyr | 4.6 |
| 25 | p.Val25Cys | 4.3 |
| 25 | p.Val25Trp | 5.8 |
| 25 | p.Val25Phe | 3.6 |
| 26 | p.Glu26Asn | 2.1 |
| 26 | p.Glu26Lys | 3.4 |
| 26 | p.Glu26Thr | 1.7 |
| 26 | p.Glu26Arg | 9.3 |
| 26 | p.Glu26Ser | 2.0 |
| 26 | p.Glu26Ile | 5.3 |
| 26 | p.Glu26Met | 5.8 |
| 26 | p.Glu26His | 3.1 |
| 26 | p.Glu26Gln | 3.6 |
| 26 | p.Glu26Pro | 1.0 |
| 26 | p.Glu26Leu | 4.0 |
| 26 | p.Glu26Asp | 15.7 |
| 26 | p.Glu26Glu | 15.1 |
| 26 | p.Glu26Ala | 5.5 |
| 26 | p.Glu26Gly | 2.4 |
| 26 | p.Glu26Val | 6.2 |
| 26 | p.Glu26Tyr | 3.4 |
| 26 | p.Glu26Cys | 0.8 |
| 26 | p.Glu26Trp | 5.7 |
| 26 | p.Glu26Phe | 3.8 |
| 27 | p.Glu27Asn | 1.2 |
| 27 | p.Glu27Lys | 14.1 |
| 27 | p.Glu27Thr | 0.2 |
| 27 | p.Glu27Arg | 5.1 |
| 27 | p.Glu27Ser | 3.1 |
| 27 | p.Glu27Ile | 0.6 |
| 27 | p.Glu27Met | 7.4 |
| 27 | p.Glu27His | 0.0 |
| 27 | p.Glu27Gln | 6.4 |
| 27 | p.Glu27Pro | 0.8 |
| 27 | p.Glu27Leu | 4.4 |
| 27 | p.Glu27Asp | 6.8 |
| 27 | p.Glu27Glu | 14.1 |
| 27 | p.Glu27Ala | 0.7 |
| 27 | p.Glu27Gly | 2.3 |
| 27 | p.Glu27Val | 10.5 |

| | | |
|---|---|---|
| 27 | p.Glu27Tyr | 2.5 |
| 27 | p.Glu27Cys | 2.4 |
| 27 | p.Glu27Trp | 9.2 |
| 27 | p.Glu27Phe | 8.1 |
| 28 | p.Val28Asn | 4.1 |
| 28 | p.Val28Lys | 4.4 |
| 28 | p.Val28Thr | 3.7 |
| 28 | p.Val28Arg | 4.3 |
| 28 | p.Val28Ser | 4.7 |
| 28 | p.Val28Ile | 7.0 |
| 28 | p.Val28Met | 3.8 |
| 28 | p.Val28His | 2.9 |
| 28 | p.Val28Gln | 4.2 |
| 28 | p.Val28Pro | 3.8 |
| 28 | p.Val28Leu | 5.0 |
| 28 | p.Val28Asp | 6.5 |
| 28 | p.Val28Glu | 5.7 |
| 28 | p.Val28Ala | 7.5 |
| 28 | p.Val28Gly | 8.9 |
| 28 | p.Val28Val | 7.1 |
| 28 | p.Val28Tyr | 3.8 |
| 28 | p.Val28Cys | 3.8 |
| 28 | p.Val28Trp | 4.3 |
| 28 | p.Val28Phe | 4.6 |
| 29 | p.Arg29Asn | 4.4 |
| 29 | p.Arg29Lys | 5.7 |
| 29 | p.Arg29Thr | 3.9 |
| 29 | p.Arg29Arg | 16.6 |
| 29 | p.Arg29Ser | 9.7 |
| 29 | p.Arg29Ile | 4.0 |
| 29 | p.Arg29Met | 4.8 |
| 29 | p.Arg29His | 2.0 |
| 29 | p.Arg29Gln | 5.7 |
| 29 | p.Arg29Pro | 1.4 |
| 29 | p.Arg29Leu | 5.3 |
| 29 | p.Arg29Asp | 4.7 |
| 29 | p.Arg29Glu | 3.5 |
| 29 | p.Arg29Ala | 4.7 |
| 29 | p.Arg29Gly | 5.7 |
| 29 | p.Arg29Val | 3.2 |
| 29 | p.Arg29Tyr | 4.1 |
| 29 | p.Arg29Cys | 5.7 |
| 29 | p.Arg29Trp | 3.0 |
| 29 | p.Arg29Phe | 1.8 |
| 30 | p.Ala30Asn | 5.0 |

| | | |
|---|---|---|
| 30 | p.Ala30Lys | 4.1 |
| 30 | p.Ala30Thr | 5.0 |
| 30 | p.Ala30Arg | 5.6 |
| 30 | p.Ala30Ser | 3.6 |
| 30 | p.Ala30Ile | 3.6 |
| 30 | p.Ala30Met | 6.4 |
| 30 | p.Ala30His | 3.6 |
| 30 | p.Ala30Gln | 4.4 |
| 30 | p.Ala30Pro | 3.6 |
| 30 | p.Ala30Leu | 3.3 |
| 30 | p.Ala30Asp | 3.8 |
| 30 | p.Ala30Glu | 12.6 |
| 30 | p.Ala30Ala | 4.4 |
| 30 | p.Ala30Gly | 3.6 |
| 30 | p.Ala30Val | 11.6 |
| 30 | p.Ala30Tyr | 3.6 |
| 30 | p.Ala30Cys | 2.7 |
| 30 | p.Ala30Trp | 6.8 |
| 30 | p.Ala30Phe | 2.8 |
| 31 | p.Leu31Asn | 2.5 |
| 31 | p.Leu31Lys | 3.5 |
| 31 | p.Leu31Thr | 3.1 |
| 31 | p.Leu31Arg | 3.3 |
| 31 | p.Leu31Ser | 3.3 |
| 31 | p.Leu31Ile | 10.3 |
| 31 | p.Leu31Met | 4.7 |
| 31 | p.Leu31His | 6.8 |
| 31 | p.Leu31Gln | 4.4 |
| 31 | p.Leu31Pro | 15.8 |
| 31 | p.Leu31Leu | 11.4 |
| 31 | p.Leu31Asp | 3.8 |
| 31 | p.Leu31Glu | 1.7 |
| 31 | p.Leu31Ala | 3.0 |
| 31 | p.Leu31Gly | 4.0 |
| 31 | p.Leu31Val | 2.6 |
| 31 | p.Leu31Tyr | 4.3 |
| 31 | p.Leu31Cys | 2.5 |
| 31 | p.Leu31Trp | 2.0 |
| 31 | p.Leu31Phe | 6.9 |
| 32 | p.Leu32Asn | 4.6 |
| 32 | p.Leu32Lys | 4.9 |
| 32 | p.Leu32Thr | 4.2 |
| 32 | p.Leu32Arg | 4.6 |
| 32 | p.Leu32Ser | 4.9 |
| 32 | p.Leu32Ile | 5.8 |

| | | |
|---|---|---|
| 32 | p.Leu32Met | 6.4 |
| 32 | p.Leu32His | 4.4 |
| 32 | p.Leu32Gln | 5.7 |
| 32 | p.Leu32Pro | 6.5 |
| 32 | p.Leu32Leu | 5.5 |
| 32 | p.Leu32Asp | 2.7 |
| 32 | p.Leu32Glu | 4.3 |
| 32 | p.Leu32Ala | 4.9 |
| 32 | p.Leu32Gly | 3.9 |
| 32 | p.Leu32Val | 6.4 |
| 32 | p.Leu32Tyr | 5.6 |
| 32 | p.Leu32Cys | 4.3 |
| 32 | p.Leu32Trp | 4.5 |
| 32 | p.Leu32Phe | 5.9 |
| 33 | p.Glu33Asn | 7.1 |
| 33 | p.Glu33Lys | 4.0 |
| 33 | p.Glu33Thr | 4.6 |
| 33 | p.Glu33Arg | 3.8 |
| 33 | p.Glu33Ser | 1.8 |
| 33 | p.Glu33Ile | 5.6 |
| 33 | p.Glu33Met | 4.2 |
| 33 | p.Glu33His | 5.0 |
| 33 | p.Glu33Gln | 5.0 |
| 33 | p.Glu33Pro | 2.8 |
| 33 | p.Glu33Leu | 5.2 |
| 33 | p.Glu33Asp | 8.8 |
| 33 | p.Glu33Glu | 9.0 |
| 33 | p.Glu33Ala | 3.4 |
| 33 | p.Glu33Gly | 4.4 |
| 33 | p.Glu33Val | 5.6 |
| 33 | p.Glu33Tyr | 4.8 |
| 33 | p.Glu33Cys | 4.7 |
| 33 | p.Glu33Trp | 5.5 |
| 33 | p.Glu33Phe | 4.6 |
| 34 | p.Ala34Asn | 3.6 |
| 34 | p.Ala34Lys | 6.4 |
| 34 | p.Ala34Thr | 5.3 |
| 34 | p.Ala34Arg | 6.3 |
| 34 | p.Ala34Ser | 3.4 |
| 34 | p.Ala34Ile | 2.5 |
| 34 | p.Ala34Met | 4.8 |
| 34 | p.Ala34His | 3.4 |
| 34 | p.Ala34Gln | 4.4 |
| 34 | p.Ala34Pro | 4.6 |
| 34 | p.Ala34Leu | 4.8 |

| | | |
|---|---|---|
| 34 | p.Ala34Asp | 10.6 |
| 34 | p.Ala34Glu | 5.6 |
| 34 | p.Ala34Ala | 7.2 |
| 34 | p.Ala34Gly | 5.1 |
| 34 | p.Ala34Val | 4.1 |
| 34 | p.Ala34Tyr | 4.9 |
| 34 | p.Ala34Cys | 4.5 |
| 34 | p.Ala34Trp | 4.0 |
| 34 | p.Ala34Phe | 4.4 |
| 35 | p.Gly35Asn | 3.3 |
| 35 | p.Gly35Lys | 4.3 |
| 35 | p.Gly35Thr | 4.2 |
| 35 | p.Gly35Arg | 4.9 |
| 35 | p.Gly35Ser | 3.4 |
| 35 | p.Gly35Ile | 3.1 |
| 35 | p.Gly35Met | 5.2 |
| 35 | p.Gly35His | 5.1 |
| 35 | p.Gly35Gln | 5.4 |
| 35 | p.Gly35Pro | 4.4 |
| 35 | p.Gly35Leu | 5.2 |
| 35 | p.Gly35Asp | 4.6 |
| 35 | p.Gly35Glu | 5.2 |
| 35 | p.Gly35Ala | 3.5 |
| 35 | p.Gly35Gly | 10.4 |
| 35 | p.Gly35Val | 6.6 |
| 35 | p.Gly35Tyr | 4.7 |
| 35 | p.Gly35Cys | 4.8 |
| 35 | p.Gly35Trp | 6.3 |
| 35 | p.Gly35Phe | 5.4 |
| 36 | p.Ala36Asn | 4.9 |
| 36 | p.Ala36Lys | 5.4 |
| 36 | p.Ala36Thr | 3.4 |
| 36 | p.Ala36Arg | 4.0 |
| 36 | p.Ala36Ser | 3.4 |
| 36 | p.Ala36Ile | 5.3 |
| 36 | p.Ala36Met | 4.8 |
| 36 | p.Ala36His | 4.5 |
| 36 | p.Ala36Gln | 5.8 |
| 36 | p.Ala36Pro | 5.5 |
| 36 | p.Ala36Leu | 5.5 |
| 36 | p.Ala36Asp | 4.6 |
| 36 | p.Ala36Glu | 6.4 |
| 36 | p.Ala36Ala | 8.0 |
| 36 | p.Ala36Gly | 2.6 |
| 36 | p.Ala36Val | 5.9 |

| | | |
|---|---|---|
| 36 | p.Ala36Tyr | 4.4 |
| 36 | p.Ala36Cys | 4.2 |
| 36 | p.Ala36Trp | 6.4 |
| 36 | p.Ala36Phe | 5.1 |
| 37 | p.Leu37Asn | 5.4 |
| 37 | p.Leu37Lys | 4.9 |
| 37 | p.Leu37Thr | 3.1 |
| 37 | p.Leu37Arg | 3.3 |
| 37 | p.Leu37Ser | 3.7 |
| 37 | p.Leu37Ile | 5.0 |
| 37 | p.Leu37Met | 5.9 |
| 37 | p.Leu37His | 4.8 |
| 37 | p.Leu37Gln | 7.4 |
| 37 | p.Leu37Pro | 3.6 |
| 37 | p.Leu37Leu | 7.3 |
| 37 | p.Leu37Asp | 3.4 |
| 37 | p.Leu37Glu | 4.6 |
| 37 | p.Leu37Ala | 4.7 |
| 37 | p.Leu37Gly | 3.0 |
| 37 | p.Leu37Val | 9.3 |
| 37 | p.Leu37Tyr | 3.8 |
| 37 | p.Leu37Cys | 3.2 |
| 37 | p.Leu37Trp | 6.3 |
| 37 | p.Leu37Phe | 7.1 |
| 38 | p.Pro38Asn | 4.2 |
| 38 | p.Pro38Lys | 3.5 |
| 38 | p.Pro38Thr | 3.4 |
| 38 | p.Pro38Arg | 3.9 |
| 38 | p.Pro38Ser | 3.2 |
| 38 | p.Pro38Ile | 3.1 |
| 38 | p.Pro38Met | 5.2 |
| 38 | p.Pro38His | 3.3 |
| 38 | p.Pro38Gln | 10.8 |
| 38 | p.Pro38Pro | 9.5 |
| 38 | p.Pro38Leu | 6.3 |
| 38 | p.Pro38Asp | 4.3 |
| 38 | p.Pro38Glu | 6.9 |
| 38 | p.Pro38Ala | 3.4 |
| 38 | p.Pro38Gly | 4.6 |
| 38 | p.Pro38Val | 6.0 |
| 38 | p.Pro38Tyr | 2.7 |
| 38 | p.Pro38Cys | 5.0 |
| 38 | p.Pro38Trp | 4.4 |
| 38 | p.Pro38Phe | 6.5 |
| 39 | p.Asn39Asn | 7.6 |

| | | |
|---|---|---|
| 39 | p.Asn39Lys | 8.2 |
| 39 | p.Asn39Thr | 3.0 |
| 39 | p.Asn39Arg | 4.0 |
| 39 | p.Asn39Ser | 5.0 |
| 39 | p.Asn39Ile | 4.1 |
| 39 | p.Asn39Met | 6.3 |
| 39 | p.Asn39His | 4.2 |
| 39 | p.Asn39Gln | 2.9 |
| 39 | p.Asn39Pro | 3.8 |
| 39 | p.Asn39Leu | 6.5 |
| 39 | p.Asn39Asp | 6.4 |
| 39 | p.Asn39Glu | 6.0 |
| 39 | p.Asn39Ala | 3.9 |
| 39 | p.Asn39Gly | 6.1 |
| 39 | p.Asn39Val | 5.2 |
| 39 | p.Asn39Tyr | 3.1 |
| 39 | p.Asn39Cys | 5.2 |
| 39 | p.Asn39Trp | 3.8 |
| 39 | p.Asn39Phe | 4.6 |
| 40 | p.Ala40Asn | 5.0 |
| 40 | p.Ala40Lys | 6.2 |
| 40 | p.Ala40Thr | 4.3 |
| 40 | p.Ala40Arg | 4.5 |
| 40 | p.Ala40Ser | 2.5 |
| 40 | p.Ala40Ile | 3.1 |
| 40 | p.Ala40Met | 3.4 |
| 40 | p.Ala40His | 2.9 |
| 40 | p.Ala40Gln | 4.0 |
| 40 | p.Ala40Pro | 3.9 |
| 40 | p.Ala40Leu | 4.0 |
| 40 | p.Ala40Asp | 5.1 |
| 40 | p.Ala40Glu | 14.6 |
| 40 | p.Ala40Ala | 5.8 |
| 40 | p.Ala40Gly | 4.5 |
| 40 | p.Ala40Val | 11.0 |
| 40 | p.Ala40Tyr | 4.1 |
| 40 | p.Ala40Cys | 3.3 |
| 40 | p.Ala40Trp | 3.8 |
| 40 | p.Ala40Phe | 4.1 |
| 41 | p.Pro41Asn | 4.9 |
| 41 | p.Pro41Lys | 4.3 |
| 41 | p.Pro41Thr | 4.0 |
| 41 | p.Pro41Arg | 3.4 |
| 41 | p.Pro41Ser | 6.0 |
| 41 | p.Pro41Ile | 3.9 |

| | | |
|---|---|---|
| 41 | p.Pro41Met | 6.1 |
| 41 | p.Pro41His | 3.9 |
| 41 | p.Pro41Gln | 5.7 |
| 41 | p.Pro41Pro | 4.9 |
| 41 | p.Pro41Leu | 5.1 |
| 41 | p.Pro41Asp | 6.5 |
| 41 | p.Pro41Glu | 5.2 |
| 41 | p.Pro41Ala | 6.0 |
| 41 | p.Pro41Gly | 2.9 |
| 41 | p.Pro41Val | 6.6 |
| 41 | p.Pro41Tyr | 5.5 |
| 41 | p.Pro41Cys | 5.2 |
| 41 | p.Pro41Trp | 4.0 |
| 41 | p.Pro41Phe | 5.9 |
| 42 | p.Asn42Asn | 4.8 |
| 42 | p.Asn42Lys | 6.4 |
| 42 | p.Asn42Thr | 7.5 |
| 42 | p.Asn42Arg | 5.4 |
| 42 | p.Asn42Ser | 5.8 |
| 42 | p.Asn42Ile | 5.8 |
| 42 | p.Asn42Met | 3.6 |
| 42 | p.Asn42His | 7.8 |
| 42 | p.Asn42Gln | 4.4 |
| 42 | p.Asn42Pro | 3.0 |
| 42 | p.Asn42Leu | 3.6 |
| 42 | p.Asn42Asp | 5.5 |
| 42 | p.Asn42Glu | 4.6 |
| 42 | p.Asn42Ala | 3.0 |
| 42 | p.Asn42Gly | 4.4 |
| 42 | p.Asn42Val | 5.5 |
| 42 | p.Asn42Tyr | 5.4 |
| 42 | p.Asn42Cys | 4.3 |
| 42 | p.Asn42Trp | 4.2 |
| 42 | p.Asn42Phe | 5.1 |
| 43 | p.Ser43Asn | 2.8 |
| 43 | p.Ser43Lys | 4.6 |
| 43 | p.Ser43Thr | 7.4 |
| 43 | p.Ser43Arg | 3.8 |
| 43 | p.Ser43Ser | 3.4 |
| 43 | p.Ser43Ile | 2.3 |
| 43 | p.Ser43Met | 4.0 |
| 43 | p.Ser43His | 4.3 |
| 43 | p.Ser43Gln | 4.0 |
| 43 | p.Ser43Pro | 7.7 |
| 43 | p.Ser43Leu | 2.7 |

| | | |
|---|---|---|
| 43 | p.Ser43Asp | 4.0 |
| 43 | p.Ser43Glu | 3.5 |
| 43 | p.Ser43Ala | 13.7 |
| 43 | p.Ser43Gly | 2.0 |
| 43 | p.Ser43Val | 4.3 |
| 43 | p.Ser43Tyr | 10.9 |
| 43 | p.Ser43Cys | 4.4 |
| 43 | p.Ser43Trp | 3.7 |
| 43 | p.Ser43Phe | 6.5 |
| 44 | p.Tyr44Asn | 6.1 |
| 44 | p.Tyr44Lys | 5.3 |
| 44 | p.Tyr44Thr | 5.3 |
| 44 | p.Tyr44Arg | 3.6 |
| 44 | p.Tyr44Ser | 4.6 |
| 44 | p.Tyr44Ile | 4.8 |
| 44 | p.Tyr44Met | 4.0 |
| 44 | p.Tyr44His | 6.7 |
| 44 | p.Tyr44Gln | 2.9 |
| 44 | p.Tyr44Pro | 3.5 |
| 44 | p.Tyr44Leu | 4.0 |
| 44 | p.Tyr44Asp | 3.0 |
| 44 | p.Tyr44Glu | 6.1 |
| 44 | p.Tyr44Ala | 4.4 |
| 44 | p.Tyr44Gly | 4.7 |
| 44 | p.Tyr44Val | 6.4 |
| 44 | p.Tyr44Tyr | 10.5 |
| 44 | p.Tyr44Cys | 3.9 |
| 44 | p.Tyr44Trp | 6.1 |
| 44 | p.Tyr44Phe | 4.0 |
| 45 | p.Gly45Asn | 4.3 |
| 45 | p.Gly45Lys | 4.7 |
| 45 | p.Gly45Thr | 3.0 |
| 45 | p.Gly45Arg | 8.9 |
| 45 | p.Gly45Ser | 6.3 |
| 45 | p.Gly45Ile | 5.1 |
| 45 | p.Gly45Met | 4.3 |
| 45 | p.Gly45His | 4.1 |
| 45 | p.Gly45Gln | 4.6 |
| 45 | p.Gly45Pro | 3.2 |
| 45 | p.Gly45Leu | 5.5 |
| 45 | p.Gly45Asp | 5.4 |
| 45 | p.Gly45Glu | 4.7 |
| 45 | p.Gly45Ala | 3.6 |
| 45 | p.Gly45Gly | 9.8 |
| 45 | p.Gly45Val | 4.3 |

| | | |
|---|---|---|
| 45 | p.Gly45Tyr | 4.0 |
| 45 | p.Gly45Cys | 5.0 |
| 45 | p.Gly45Trp | 3.8 |
| 45 | p.Gly45Phe | 5.4 |
| 46 | p.Arg46Asn | 4.9 |
| 46 | p.Arg46Lys | 3.4 |
| 46 | p.Arg46Thr | 5.0 |
| 46 | p.Arg46Arg | 4.4 |
| 46 | p.Arg46Ser | 9.2 |
| 46 | p.Arg46Ile | 6.0 |
| 46 | p.Arg46Met | 2.7 |
| 46 | p.Arg46His | 9.1 |
| 46 | p.Arg46Gln | 3.9 |
| 46 | p.Arg46Pro | 7.5 |
| 46 | p.Arg46Leu | 5.7 |
| 46 | p.Arg46Asp | 3.3 |
| 46 | p.Arg46Glu | 2.9 |
| 46 | p.Arg46Ala | 3.2 |
| 46 | p.Arg46Gly | 5.1 |
| 46 | p.Arg46Val | 2.7 |
| 46 | p.Arg46Tyr | 5.7 |
| 46 | p.Arg46Cys | 7.3 |
| 46 | p.Arg46Trp | 4.0 |
| 46 | p.Arg46Phe | 4.0 |
| 47 | p.Arg47Asn | 3.6 |
| 47 | p.Arg47Lys | 3.5 |
| 47 | p.Arg47Thr | 4.0 |
| 47 | p.Arg47Arg | 4.8 |
| 47 | p.Arg47Ser | 10.4 |
| 47 | p.Arg47Ile | 3.3 |
| 47 | p.Arg47Met | 4.0 |
| 47 | p.Arg47His | 9.5 |
| 47 | p.Arg47Gln | 4.5 |
| 47 | p.Arg47Pro | 4.8 |
| 47 | p.Arg47Leu | 3.8 |
| 47 | p.Arg47Asp | 4.3 |
| 47 | p.Arg47Glu | 4.9 |
| 47 | p.Arg47Ala | 3.3 |
| 47 | p.Arg47Gly | 5.0 |
| 47 | p.Arg47Val | 3.8 |
| 47 | p.Arg47Tyr | 5.0 |
| 47 | p.Arg47Cys | 8.1 |
| 47 | p.Arg47Trp | 4.5 |
| 47 | p.Arg47Phe | 4.7 |
| 48 | p.Pro48Asn | 4.7 |

| | | |
|---|---|---|
| 48 | p.Pro48Lys | 5.5 |
| 48 | p.Pro48Thr | 5.1 |
| 48 | p.Pro48Arg | 5.6 |
| 48 | p.Pro48Ser | 4.6 |
| 48 | p.Pro48Ile | 6.7 |
| 48 | p.Pro48Met | 6.6 |
| 48 | p.Pro48His | 4.4 |
| 48 | p.Pro48Gln | 5.3 |
| 48 | p.Pro48Pro | 5.2 |
| 48 | p.Pro48Leu | 7.1 |
| 48 | p.Pro48Asp | 4.6 |
| 48 | p.Pro48Glu | 4.7 |
| 48 | p.Pro48Ala | 3.5 |
| 48 | p.Pro48Gly | 4.7 |
| 48 | p.Pro48Val | 4.8 |
| 48 | p.Pro48Tyr | 3.8 |
| 48 | p.Pro48Cys | 3.4 |
| 48 | p.Pro48Trp | 5.6 |
| 48 | p.Pro48Phe | 4.3 |
| 49 | p.Ile49Asn | 3.5 |
| 49 | p.Ile49Lys | 5.2 |
| 49 | p.Ile49Thr | 3.9 |
| 49 | p.Ile49Arg | 8.3 |
| 49 | p.Ile49Ser | 4.8 |
| 49 | p.Ile49Ile | 7.3 |
| 49 | p.Ile49Met | 7.5 |
| 49 | p.Ile49His | 4.7 |
| 49 | p.Ile49Gln | 4.8 |
| 49 | p.Ile49Pro | 4.5 |
| 49 | p.Ile49Leu | 4.6 |
| 49 | p.Ile49Asp | 5.5 |
| 49 | p.Ile49Glu | 5.4 |
| 49 | p.Ile49Ala | 3.7 |
| 49 | p.Ile49Gly | 5.2 |
| 49 | p.Ile49Val | 5.5 |
| 49 | p.Ile49Tyr | 3.6 |
| 49 | p.Ile49Cys | 3.0 |
| 49 | p.Ile49Trp | 4.9 |
| 49 | p.Ile49Phe | 4.2 |
| 50 | p.Gln50Asn | 3.8 |
| 50 | p.Gln50Lys | 4.2 |
| 50 | p.Gln50Thr | 4.3 |
| 50 | p.Gln50Arg | 3.9 |
| 50 | p.Gln50Ser | 3.9 |
| 50 | p.Gln50Ile | 4.2 |

| | | |
|---|---|---|
| 50 | p.Gln50Met | 5.4 |
| 50 | p.Gln50His | 9.2 |
| 50 | p.Gln50Gln | 7.5 |
| 50 | p.Gln50Pro | 5.7 |
| 50 | p.Gln50Leu | 5.1 |
| 50 | p.Gln50Asp | 4.7 |
| 50 | p.Gln50Glu | 6.0 |
| 50 | p.Gln50Ala | 4.4 |
| 50 | p.Gln50Gly | 4.1 |
| 50 | p.Gln50Val | 6.3 |
| 50 | p.Gln50Tyr | 4.6 |
| 50 | p.Gln50Cys | 3.7 |
| 50 | p.Gln50Trp | 3.8 |
| 50 | p.Gln50Phe | 5.1 |
| 51 | p.Val51Asn | 2.1 |
| 51 | p.Val51Lys | 3.6 |
| 51 | p.Val51Thr | 2.7 |
| 51 | p.Val51Arg | 3.2 |
| 51 | p.Val51Ser | 3.1 |
| 51 | p.Val51Ile | 7.2 |
| 51 | p.Val51Met | 4.8 |
| 51 | p.Val51His | 4.1 |
| 51 | p.Val51Gln | 6.2 |
| 51 | p.Val51Pro | 5.7 |
| 51 | p.Val51Leu | 6.3 |
| 51 | p.Val51Asp | 6.2 |
| 51 | p.Val51Glu | 4.9 |
| 51 | p.Val51Ala | 3.7 |
| 51 | p.Val51Gly | 2.5 |
| 51 | p.Val51Val | 14.3 |
| 51 | p.Val51Tyr | 4.5 |
| 51 | p.Val51Cys | 7.5 |
| 51 | p.Val51Trp | 4.0 |
| 51 | p.Val51Phe | 3.5 |
| 52 | p.Met52Asn | 4.3 |
| 52 | p.Met52Lys | 7.2 |
| 52 | p.Met52Thr | 4.5 |
| 52 | p.Met52Arg | 4.2 |
| 52 | p.Met52Ser | 5.5 |
| 52 | p.Met52Ile | 5.5 |
| 52 | p.Met52Met | 0.0 |
| 52 | p.Met52His | 4.5 |
| 52 | p.Met52Gln | 4.3 |
| 52 | p.Met52Pro | 3.9 |
| 52 | p.Met52Leu | 9.7 |

| | | |
|---|---|---|
| 52 | p.Met52Asp | 3.7 |
| 52 | p.Met52Glu | 4.8 |
| 52 | p.Met52Ala | 5.7 |
| 52 | p.Met52Gly | 7.2 |
| 52 | p.Met52Val | 7.0 |
| 52 | p.Met52Tyr | 4.7 |
| 52 | p.Met52Cys | 5.1 |
| 52 | p.Met52Trp | 3.9 |
| 52 | p.Met52Phe | 4.3 |
| 53 | p.Met53Asn | 4.9 |
| 53 | p.Met53Lys | 6.9 |
| 53 | p.Met53Thr | 5.1 |
| 53 | p.Met53Arg | 4.7 |
| 53 | p.Met53Ser | 4.1 |
| 53 | p.Met53Ile | 4.7 |
| 53 | p.Met53Met | 0.0 |
| 53 | p.Met53His | 5.9 |
| 53 | p.Met53Gln | 6.5 |
| 53 | p.Met53Pro | 4.4 |
| 53 | p.Met53Leu | 8.6 |
| 53 | p.Met53Asp | 3.9 |
| 53 | p.Met53Glu | 5.7 |
| 53 | p.Met53Ala | 5.5 |
| 53 | p.Met53Gly | 4.2 |
| 53 | p.Met53Val | 6.3 |
| 53 | p.Met53Tyr | 4.5 |
| 53 | p.Met53Cys | 3.4 |
| 53 | p.Met53Trp | 6.6 |
| 53 | p.Met53Phe | 4.2 |
| 54 | p.Met54Asn | 5.1 |
| 54 | p.Met54Lys | 4.7 |
| 54 | p.Met54Thr | 5.1 |
| 54 | p.Met54Arg | 3.5 |
| 54 | p.Met54Ser | 5.8 |
| 54 | p.Met54Ile | 7.4 |
| 54 | p.Met54Met | 0.0 |
| 54 | p.Met54His | 1.4 |
| 54 | p.Met54Gln | 4.4 |
| 54 | p.Met54Pro | 3.4 |
| 54 | p.Met54Leu | 14.3 |
| 54 | p.Met54Asp | 5.1 |
| 54 | p.Met54Glu | 4.9 |
| 54 | p.Met54Ala | 3.0 |
| 54 | p.Met54Gly | 3.2 |
| 54 | p.Met54Val | 9.6 |

| | | |
|---|---|---|
| 54 | p.Met54Tyr | 2.3 |
| 54 | p.Met54Cys | 4.4 |
| 54 | p.Met54Trp | 4.7 |
| 54 | p.Met54Phe | 7.8 |
| 55 | p.Gly55Asn | 3.3 |
| 55 | p.Gly55Lys | 4.4 |
| 55 | p.Gly55Thr | 5.4 |
| 55 | p.Gly55Arg | 11.8 |
| 55 | p.Gly55Ser | 4.6 |
| 55 | p.Gly55Ile | 5.1 |
| 55 | p.Gly55Met | 4.9 |
| 55 | p.Gly55His | 4.0 |
| 55 | p.Gly55Gln | 3.5 |
| 55 | p.Gly55Pro | 6.1 |
| 55 | p.Gly55Leu | 5.8 |
| 55 | p.Gly55Asp | 4.6 |
| 55 | p.Gly55Glu | 5.8 |
| 55 | p.Gly55Ala | 2.8 |
| 55 | p.Gly55Gly | 8.6 |
| 55 | p.Gly55Val | 3.4 |
| 55 | p.Gly55Tyr | 4.9 |
| 55 | p.Gly55Cys | 4.5 |
| 55 | p.Gly55Trp | 3.4 |
| 55 | p.Gly55Phe | 3.1 |
| 56 | p.Ser56Asn | 4.9 |
| 56 | p.Ser56Lys | 7.2 |
| 56 | p.Ser56Thr | 3.6 |
| 56 | p.Ser56Arg | 3.9 |
| 56 | p.Ser56Ser | 5.0 |
| 56 | p.Ser56Ile | 4.7 |
| 56 | p.Ser56Met | 3.0 |
| 56 | p.Ser56His | 5.3 |
| 56 | p.Ser56Gln | 4.8 |
| 56 | p.Ser56Pro | 4.8 |
| 56 | p.Ser56Leu | 6.1 |
| 56 | p.Ser56Asp | 4.7 |
| 56 | p.Ser56Glu | 6.1 |
| 56 | p.Ser56Ala | 5.3 |
| 56 | p.Ser56Gly | 5.9 |
| 56 | p.Ser56Val | 5.7 |
| 56 | p.Ser56Tyr | 2.6 |
| 56 | p.Ser56Cys | 4.5 |
| 56 | p.Ser56Trp | 6.5 |
| 56 | p.Ser56Phe | 5.6 |
| 57 | p.Ala57Asn | 5.6 |

| | | |
|---|---|---|
| 57 | p.Ala57Lys | 6.2 |
| 57 | p.Ala57Thr | 3.2 |
| 57 | p.Ala57Arg | 4.0 |
| 57 | p.Ala57Ser | 5.2 |
| 57 | p.Ala57Ile | 4.5 |
| 57 | p.Ala57Met | 3.9 |
| 57 | p.Ala57His | 5.4 |
| 57 | p.Ala57Gln | 4.2 |
| 57 | p.Ala57Pro | 5.1 |
| 57 | p.Ala57Leu | 5.0 |
| 57 | p.Ala57Asp | 5.5 |
| 57 | p.Ala57Glu | 5.7 |
| 57 | p.Ala57Ala | 6.1 |
| 57 | p.Ala57Gly | 5.4 |
| 57 | p.Ala57Val | 6.7 |
| 57 | p.Ala57Tyr | 4.3 |
| 57 | p.Ala57Cys | 5.4 |
| 57 | p.Ala57Trp | 4.0 |
| 57 | p.Ala57Phe | 4.5 |
| 58 | p.Arg58Asn | 5.1 |
| 58 | p.Arg58Lys | 7.3 |
| 58 | p.Arg58Thr | 2.3 |
| 58 | p.Arg58Arg | 5.6 |
| 58 | p.Arg58Ser | 5.4 |
| 58 | p.Arg58Ile | 5.3 |
| 58 | p.Arg58Met | 11.8 |
| 58 | p.Arg58His | 3.7 |
| 58 | p.Arg58Gln | 3.5 |
| 58 | p.Arg58Pro | 3.5 |
| 58 | p.Arg58Leu | 5.1 |
| 58 | p.Arg58Asp | 3.8 |
| 58 | p.Arg58Glu | 5.2 |
| 58 | p.Arg58Ala | 3.4 |
| 58 | p.Arg58Gly | 4.1 |
| 58 | p.Arg58Val | 5.7 |
| 58 | p.Arg58Tyr | 4.6 |
| 58 | p.Arg58Cys | 4.8 |
| 58 | p.Arg58Trp | 5.3 |
| 58 | p.Arg58Phe | 4.5 |
| 59 | p.Val59Asn | 4.8 |
| 59 | p.Val59Lys | 6.2 |
| 59 | p.Val59Thr | 5.1 |
| 59 | p.Val59Arg | 5.0 |
| 59 | p.Val59Ser | 5.0 |
| 59 | p.Val59Ile | 4.7 |

| | | |
|---|---|---|
| 59 | p.Val59Met | 5.2 |
| 59 | p.Val59His | 4.4 |
| 59 | p.Val59Gln | 5.4 |
| 59 | p.Val59Pro | 4.3 |
| 59 | p.Val59Leu | 5.1 |
| 59 | p.Val59Asp | 5.3 |
| 59 | p.Val59Glu | 6.5 |
| 59 | p.Val59Ala | 3.5 |
| 59 | p.Val59Gly | 3.6 |
| 59 | p.Val59Val | 7.8 |
| 59 | p.Val59Tyr | 4.1 |
| 59 | p.Val59Cys | 4.5 |
| 59 | p.Val59Trp | 5.8 |
| 59 | p.Val59Phe | 3.9 |
| 60 | p.Ala60Asn | 3.6 |
| 60 | p.Ala60Lys | 5.3 |
| 60 | p.Ala60Thr | 5.4 |
| 60 | p.Ala60Arg | 4.5 |
| 60 | p.Ala60Ser | 3.6 |
| 60 | p.Ala60Ile | 5.0 |
| 60 | p.Ala60Met | 6.0 |
| 60 | p.Ala60His | 4.7 |
| 60 | p.Ala60Gln | 6.1 |
| 60 | p.Ala60Pro | 4.6 |
| 60 | p.Ala60Leu | 3.7 |
| 60 | p.Ala60Asp | 4.7 |
| 60 | p.Ala60Glu | 5.5 |
| 60 | p.Ala60Ala | 10.6 |
| 60 | p.Ala60Gly | 4.2 |
| 60 | p.Ala60Val | 4.5 |
| 60 | p.Ala60Tyr | 3.5 |
| 60 | p.Ala60Cys | 4.3 |
| 60 | p.Ala60Trp | 4.9 |
| 60 | p.Ala60Phe | 5.2 |
| 61 | p.Glu61Asn | 4.5 |
| 61 | p.Glu61Lys | 4.6 |
| 61 | p.Glu61Thr | 4.5 |
| 61 | p.Glu61Arg | 4.2 |
| 61 | p.Glu61Ser | 4.4 |
| 61 | p.Glu61Ile | 5.1 |
| 61 | p.Glu61Met | 5.7 |
| 61 | p.Glu61His | 4.1 |
| 61 | p.Glu61Gln | 5.1 |
| 61 | p.Glu61Pro | 3.7 |
| 61 | p.Glu61Leu | 5.8 |

| | | |
|---|---|---|
| 61 | p.Glu61Asp | 8.7 |
| 61 | p.Glu61Glu | 8.1 |
| 61 | p.Glu61Ala | 4.2 |
| 61 | p.Glu61Gly | 4.2 |
| 61 | p.Glu61Val | 4.6 |
| 61 | p.Glu61Tyr | 4.8 |
| 61 | p.Glu61Cys | 4.7 |
| 61 | p.Glu61Trp | 5.0 |
| 61 | p.Glu61Phe | 3.7 |
| 62 | p.Leu62Asn | 3.6 |
| 62 | p.Leu62Lys | 4.9 |
| 62 | p.Leu62Thr | 4.8 |
| 62 | p.Leu62Arg | 4.9 |
| 62 | p.Leu62Ser | 4.2 |
| 62 | p.Leu62Ile | 4.5 |
| 62 | p.Leu62Met | 3.9 |
| 62 | p.Leu62His | 4.7 |
| 62 | p.Leu62Gln | 4.2 |
| 62 | p.Leu62Pro | 4.8 |
| 62 | p.Leu62Leu | 8.1 |
| 62 | p.Leu62Asp | 3.0 |
| 62 | p.Leu62Glu | 4.6 |
| 62 | p.Leu62Ala | 4.1 |
| 62 | p.Leu62Gly | 6.2 |
| 62 | p.Leu62Val | 6.4 |
| 62 | p.Leu62Tyr | 4.9 |
| 62 | p.Leu62Cys | 5.5 |
| 62 | p.Leu62Trp | 4.7 |
| 62 | p.Leu62Phe | 7.8 |
| 63 | p.Leu63Asn | 4.2 |
| 63 | p.Leu63Lys | 3.8 |
| 63 | p.Leu63Thr | 3.6 |
| 63 | p.Leu63Arg | 3.8 |
| 63 | p.Leu63Ser | 4.1 |
| 63 | p.Leu63Ile | 12.4 |
| 63 | p.Leu63Met | 4.8 |
| 63 | p.Leu63His | 7.3 |
| 63 | p.Leu63Gln | 3.0 |
| 63 | p.Leu63Pro | 10.7 |
| 63 | p.Leu63Leu | 4.9 |
| 63 | p.Leu63Asp | 3.8 |
| 63 | p.Leu63Glu | 4.8 |
| 63 | p.Leu63Ala | 2.9 |
| 63 | p.Leu63Gly | 3.2 |
| 63 | p.Leu63Val | 4.6 |

| | | |
|---|---|---|
| 63 | p.Leu63Tyr | 4.5 |
| 63 | p.Leu63Cys | 2.5 |
| 63 | p.Leu63Trp | 5.1 |
| 63 | p.Leu63Phe | 6.0 |
| 64 | p.Leu64Asn | 3.9 |
| 64 | p.Leu64Lys | 4.0 |
| 64 | p.Leu64Thr | 3.9 |
| 64 | p.Leu64Arg | 4.9 |
| 64 | p.Leu64Ser | 3.8 |
| 64 | p.Leu64Ile | 10.3 |
| 64 | p.Leu64Met | 3.2 |
| 64 | p.Leu64His | 9.1 |
| 64 | p.Leu64Gln | 4.4 |
| 64 | p.Leu64Pro | 5.1 |
| 64 | p.Leu64Leu | 5.7 |
| 64 | p.Leu64Asp | 3.4 |
| 64 | p.Leu64Glu | 4.4 |
| 64 | p.Leu64Ala | 4.8 |
| 64 | p.Leu64Gly | 4.8 |
| 64 | p.Leu64Val | 3.5 |
| 64 | p.Leu64Tyr | 3.9 |
| 64 | p.Leu64Cys | 4.4 |
| 64 | p.Leu64Trp | 5.3 |
| 64 | p.Leu64Phe | 7.2 |
| 65 | p.Leu65Asn | 3.1 |
| 65 | p.Leu65Lys | 5.2 |
| 65 | p.Leu65Thr | 4.4 |
| 65 | p.Leu65Arg | 5.0 |
| 65 | p.Leu65Ser | 5.3 |
| 65 | p.Leu65Ile | 4.5 |
| 65 | p.Leu65Met | 6.5 |
| 65 | p.Leu65His | 3.5 |
| 65 | p.Leu65Gln | 4.8 |
| 65 | p.Leu65Pro | 3.2 |
| 65 | p.Leu65Leu | 6.9 |
| 65 | p.Leu65Asp | 5.6 |
| 65 | p.Leu65Glu | 5.4 |
| 65 | p.Leu65Ala | 4.0 |
| 65 | p.Leu65Gly | 5.8 |
| 65 | p.Leu65Val | 7.4 |
| 65 | p.Leu65Tyr | 4.6 |
| 65 | p.Leu65Cys | 4.4 |
| 65 | p.Leu65Trp | 6.0 |
| 65 | p.Leu65Phe | 4.7 |
| 66 | p.His66Asn | 4.6 |

| | | |
|---|---|---|
| 66 | p.His66Lys | 5.6 |
| 66 | p.His66Thr | 3.6 |
| 66 | p.His66Arg | 4.8 |
| 66 | p.His66Ser | 4.9 |
| 66 | p.His66Ile | 4.6 |
| 66 | p.His66Met | 4.7 |
| 66 | p.His66His | 5.8 |
| 66 | p.His66Gln | 5.7 |
| 66 | p.His66Pro | 4.8 |
| 66 | p.His66Leu | 5.9 |
| 66 | p.His66Asp | 2.8 |
| 66 | p.His66Glu | 6.9 |
| 66 | p.His66Ala | 5.3 |
| 66 | p.His66Gly | 5.6 |
| 66 | p.His66Val | 5.1 |
| 66 | p.His66Tyr | 4.4 |
| 66 | p.His66Cys | 4.6 |
| 66 | p.His66Trp | 6.0 |
| 66 | p.His66Phe | 4.2 |
| 67 | p.Gly67Asn | 4.2 |
| 67 | p.Gly67Lys | 5.2 |
| 67 | p.Gly67Thr | 4.3 |
| 67 | p.Gly67Arg | 7.7 |
| 67 | p.Gly67Ser | 3.9 |
| 67 | p.Gly67Ile | 4.0 |
| 67 | p.Gly67Met | 4.2 |
| 67 | p.Gly67His | 5.6 |
| 67 | p.Gly67Gln | 4.7 |
| 67 | p.Gly67Pro | 5.3 |
| 67 | p.Gly67Leu | 4.3 |
| 67 | p.Gly67Asp | 3.8 |
| 67 | p.Gly67Glu | 4.2 |
| 67 | p.Gly67Ala | 4.7 |
| 67 | p.Gly67Gly | 9.1 |
| 67 | p.Gly67Val | 4.4 |
| 67 | p.Gly67Tyr | 5.3 |
| 67 | p.Gly67Cys | 4.3 |
| 67 | p.Gly67Trp | 6.2 |
| 67 | p.Gly67Phe | 4.7 |
| 68 | p.Ala68Asn | 3.2 |
| 68 | p.Ala68Lys | 6.0 |
| 68 | p.Ala68Thr | 5.3 |
| 68 | p.Ala68Arg | 3.3 |
| 68 | p.Ala68Ser | 4.3 |
| 68 | p.Ala68Ile | 3.5 |

| | | |
|---|---|---|
| 68 | p.Ala68Met | 6.6 |
| 68 | p.Ala68His | 4.1 |
| 68 | p.Ala68Gln | 3.7 |
| 68 | p.Ala68Pro | 5.3 |
| 68 | p.Ala68Leu | 5.5 |
| 68 | p.Ala68Asp | 5.6 |
| 68 | p.Ala68Glu | 6.1 |
| 68 | p.Ala68Ala | 7.4 |
| 68 | p.Ala68Gly | 4.8 |
| 68 | p.Ala68Val | 6.6 |
| 68 | p.Ala68Tyr | 4.1 |
| 68 | p.Ala68Cys | 5.1 |
| 68 | p.Ala68Trp | 4.9 |
| 68 | p.Ala68Phe | 4.9 |
| 69 | p.Glu69Asn | 4.7 |
| 69 | p.Glu69Lys | 5.7 |
| 69 | p.Glu69Thr | 4.2 |
| 69 | p.Glu69Arg | 5.2 |
| 69 | p.Glu69Ser | 5.9 |
| 69 | p.Glu69Ile | 5.5 |
| 69 | p.Glu69Met | 4.7 |
| 69 | p.Glu69His | 5.2 |
| 69 | p.Glu69Gln | 5.4 |
| 69 | p.Glu69Pro | 5.1 |
| 69 | p.Glu69Leu | 4.7 |
| 69 | p.Glu69Asp | 4.9 |
| 69 | p.Glu69Glu | 7.5 |
| 69 | p.Glu69Ala | 0.5 |
| 69 | p.Glu69Gly | 4.8 |
| 69 | p.Glu69Val | 4.8 |
| 69 | p.Glu69Tyr | 4.9 |
| 69 | p.Glu69Cys | 6.4 |
| 69 | p.Glu69Trp | 6.1 |
| 69 | p.Glu69Phe | 3.9 |
| 70 | p.Pro70Asn | 5.9 |
| 70 | p.Pro70Lys | 3.4 |
| 70 | p.Pro70Thr | 6.1 |
| 70 | p.Pro70Arg | 5.7 |
| 70 | p.Pro70Ser | 5.7 |
| 70 | p.Pro70Ile | 6.7 |
| 70 | p.Pro70Met | 5.9 |
| 70 | p.Pro70His | 2.3 |
| 70 | p.Pro70Gln | 2.4 |
| 70 | p.Pro70Pro | 8.4 |
| 70 | p.Pro70Leu | 7.4 |

| | | |
|---|---|---|
| 70 | p.Pro70Asp | 4.8 |
| 70 | p.Pro70Glu | 5.2 |
| 70 | p.Pro70Ala | 5.3 |
| 70 | p.Pro70Gly | 4.5 |
| 70 | p.Pro70Val | 5.0 |
| 70 | p.Pro70Tyr | 3.5 |
| 70 | p.Pro70Cys | 3.0 |
| 70 | p.Pro70Trp | 4.2 |
| 70 | p.Pro70Phe | 4.7 |
| 71 | p.Asn71Asn | 6.8 |
| 71 | p.Asn71Lys | 12.4 |
| 71 | p.Asn71Thr | 4.1 |
| 71 | p.Asn71Arg | 4.5 |
| 71 | p.Asn71Ser | 3.9 |
| 71 | p.Asn71Ile | 3.5 |
| 71 | p.Asn71Met | 2.6 |
| 71 | p.Asn71His | 4.7 |
| 71 | p.Asn71Gln | 4.3 |
| 71 | p.Asn71Pro | 5.1 |
| 71 | p.Asn71Leu | 4.1 |
| 71 | p.Asn71Asp | 4.9 |
| 71 | p.Asn71Glu | 6.6 |
| 71 | p.Asn71Ala | 5.7 |
| 71 | p.Asn71Gly | 3.6 |
| 71 | p.Asn71Val | 3.8 |
| 71 | p.Asn71Tyr | 5.6 |
| 71 | p.Asn71Cys | 4.0 |
| 71 | p.Asn71Trp | 4.0 |
| 71 | p.Asn71Phe | 5.6 |
| 72 | p.Cys72Asn | 4.4 |
| 72 | p.Cys72Lys | 4.4 |
| 72 | p.Cys72Thr | 4.9 |
| 72 | p.Cys72Arg | 4.0 |
| 72 | p.Cys72Ser | 4.4 |
| 72 | p.Cys72Ile | 3.6 |
| 72 | p.Cys72Met | 3.6 |
| 72 | p.Cys72His | 3.0 |
| 72 | p.Cys72Gln | 3.9 |
| 72 | p.Cys72Pro | 4.9 |
| 72 | p.Cys72Leu | 2.8 |
| 72 | p.Cys72Asp | 5.6 |
| 72 | p.Cys72Glu | 5.5 |
| 72 | p.Cys72Ala | 3.3 |
| 72 | p.Cys72Gly | 3.1 |
| 72 | p.Cys72Val | 4.4 |

| | | |
|---|---|---|
| 72 | p.Cys72Tyr | 5.6 |
| 72 | p.Cys72Cys | 9.9 |
| 72 | p.Cys72Trp | 13.7 |
| 72 | p.Cys72Phe | 4.8 |
| 73 | p.Ala73Asn | 4.4 |
| 73 | p.Ala73Lys | 3.5 |
| 73 | p.Ala73Thr | 8.3 |
| 73 | p.Ala73Arg | 4.4 |
| 73 | p.Ala73Ser | 5.5 |
| 73 | p.Ala73Ile | 4.3 |
| 73 | p.Ala73Met | 4.4 |
| 73 | p.Ala73His | 5.4 |
| 73 | p.Ala73Gln | 3.3 |
| 73 | p.Ala73Pro | 5.1 |
| 73 | p.Ala73Leu | 5.4 |
| 73 | p.Ala73Asp | 3.0 |
| 73 | p.Ala73Glu | 5.6 |
| 73 | p.Ala73Ala | 9.6 |
| 73 | p.Ala73Gly | 4.0 |
| 73 | p.Ala73Val | 5.2 |
| 73 | p.Ala73Tyr | 5.1 |
| 73 | p.Ala73Cys | 4.7 |
| 73 | p.Ala73Trp | 5.0 |
| 73 | p.Ala73Phe | 3.9 |
| 74 | p.Asp74Asn | 5.2 |
| 74 | p.Asp74Lys | 5.8 |
| 74 | p.Asp74Thr | 4.7 |
| 74 | p.Asp74Arg | 4.5 |
| 74 | p.Asp74Ser | 4.5 |
| 74 | p.Asp74Ile | 4.6 |
| 74 | p.Asp74Met | 5.1 |
| 74 | p.Asp74His | 4.0 |
| 74 | p.Asp74Gln | 3.5 |
| 74 | p.Asp74Pro | 3.2 |
| 74 | p.Asp74Leu | 5.7 |
| 74 | p.Asp74Asp | 8.6 |
| 74 | p.Asp74Glu | 9.2 |
| 74 | p.Asp74Ala | 3.5 |
| 74 | p.Asp74Gly | 4.2 |
| 74 | p.Asp74Val | 4.5 |
| 74 | p.Asp74Tyr | 4.9 |
| 74 | p.Asp74Cys | 4.2 |
| 74 | p.Asp74Trp | 5.8 |
| 74 | p.Asp74Phe | 4.3 |
| 75 | p.Pro75Asn | 4.3 |

| | | |
|---|---|---|
| 75 | p.Pro75Lys | 5.6 |
| 75 | p.Pro75Thr | 4.8 |
| 75 | p.Pro75Arg | 5.8 |
| 75 | p.Pro75Ser | 4.0 |
| 75 | p.Pro75Ile | 4.0 |
| 75 | p.Pro75Met | 4.3 |
| 75 | p.Pro75His | 4.5 |
| 75 | p.Pro75Gln | 5.0 |
| 75 | p.Pro75Pro | 8.8 |
| 75 | p.Pro75Leu | 4.4 |
| 75 | p.Pro75Asp | 4.3 |
| 75 | p.Pro75Glu | 5.6 |
| 75 | p.Pro75Ala | 4.3 |
| 75 | p.Pro75Gly | 4.6 |
| 75 | p.Pro75Val | 5.5 |
| 75 | p.Pro75Tyr | 5.7 |
| 75 | p.Pro75Cys | 4.0 |
| 75 | p.Pro75Trp | 5.3 |
| 75 | p.Pro75Phe | 5.2 |
| 76 | p.Ala76Asn | 3.8 |
| 76 | p.Ala76Lys | 5.3 |
| 76 | p.Ala76Thr | 4.5 |
| 76 | p.Ala76Arg | 3.6 |
| 76 | p.Ala76Ser | 4.5 |
| 76 | p.Ala76Ile | 2.5 |
| 76 | p.Ala76Met | 5.2 |
| 76 | p.Ala76His | 5.1 |
| 76 | p.Ala76Gln | 4.0 |
| 76 | p.Ala76Pro | 5.3 |
| 76 | p.Ala76Leu | 3.3 |
| 76 | p.Ala76Asp | 4.3 |
| 76 | p.Ala76Glu | 12.4 |
| 76 | p.Ala76Ala | 5.0 |
| 76 | p.Ala76Gly | 4.4 |
| 76 | p.Ala76Val | 9.4 |
| 76 | p.Ala76Tyr | 3.4 |
| 76 | p.Ala76Cys | 4.5 |
| 76 | p.Ala76Trp | 5.8 |
| 76 | p.Ala76Phe | 3.5 |
| 77 | p.Thr77Asn | 10.6 |
| 77 | p.Thr77Lys | 5.2 |
| 77 | p.Thr77Thr | 14.2 |
| 77 | p.Thr77Arg | 3.9 |
| 77 | p.Thr77Ser | 5.1 |
| 77 | p.Thr77Ile | 6.4 |

| | | |
|---|---|---|
| 77 | p.Thr77Met | 2.6 |
| 77 | p.Thr77His | 2.8 |
| 77 | p.Thr77Gln | 3.1 |
| 77 | p.Thr77Pro | 6.2 |
| 77 | p.Thr77Leu | 4.6 |
| 77 | p.Thr77Asp | 2.4 |
| 77 | p.Thr77Glu | 4.2 |
| 77 | p.Thr77Ala | 5.5 |
| 77 | p.Thr77Gly | 5.1 |
| 77 | p.Thr77Val | 5.2 |
| 77 | p.Thr77Tyr | 3.1 |
| 77 | p.Thr77Cys | 5.0 |
| 77 | p.Thr77Trp | 2.6 |
| 77 | p.Thr77Phe | 2.1 |
| 78 | p.Leu78Asn | 5.4 |
| 78 | p.Leu78Lys | 6.5 |
| 78 | p.Leu78Thr | 4.0 |
| 78 | p.Leu78Arg | 4.1 |
| 78 | p.Leu78Ser | 2.9 |
| 78 | p.Leu78Ile | 6.4 |
| 78 | p.Leu78Met | 9.5 |
| 78 | p.Leu78His | 5.5 |
| 78 | p.Leu78Gln | 7.0 |
| 78 | p.Leu78Pro | 4.4 |
| 78 | p.Leu78Leu | 6.1 |
| 78 | p.Leu78Asp | 8.3 |
| 78 | p.Leu78Glu | 4.1 |
| 78 | p.Leu78Ala | 1.8 |
| 78 | p.Leu78Gly | 1.7 |
| 78 | p.Leu78Val | 5.8 |
| 78 | p.Leu78Tyr | 4.6 |
| 78 | p.Leu78Cys | 3.8 |
| 78 | p.Leu78Trp | 3.9 |
| 78 | p.Leu78Phe | 4.2 |
| 79 | p.Thr79Asn | 6.1 |
| 79 | p.Thr79Lys | 5.0 |
| 79 | p.Thr79Thr | 8.2 |
| 79 | p.Thr79Arg | 2.9 |
| 79 | p.Thr79Ser | 3.8 |
| 79 | p.Thr79Ile | 4.1 |
| 79 | p.Thr79Met | 5.8 |
| 79 | p.Thr79His | 5.2 |
| 79 | p.Thr79Gln | 4.6 |
| 79 | p.Thr79Pro | 3.8 |
| 79 | p.Thr79Leu | 4.7 |

| | | |
|---|---|---|
| 79 | p.Thr79Asp | 3.8 |
| 79 | p.Thr79Glu | 3.7 |
| 79 | p.Thr79Ala | 5.6 |
| 79 | p.Thr79Gly | 6.7 |
| 79 | p.Thr79Val | 6.8 |
| 79 | p.Thr79Tyr | 2.9 |
| 79 | p.Thr79Cys | 5.8 |
| 79 | p.Thr79Trp | 6.1 |
| 79 | p.Thr79Phe | 4.5 |
| 80 | p.Arg80Asn | 4.5 |
| 80 | p.Arg80Lys | 5.3 |
| 80 | p.Arg80Thr | 2.6 |
| 80 | p.Arg80Arg | 12.6 |
| 80 | p.Arg80Ser | 9.9 |
| 80 | p.Arg80Ile | 3.3 |
| 80 | p.Arg80Met | 4.0 |
| 80 | p.Arg80His | 3.8 |
| 80 | p.Arg80Gln | 4.3 |
| 80 | p.Arg80Pro | 4.1 |
| 80 | p.Arg80Leu | 4.5 |
| 80 | p.Arg80Asp | 5.3 |
| 80 | p.Arg80Glu | 4.2 |
| 80 | p.Arg80Ala | 4.4 |
| 80 | p.Arg80Gly | 5.1 |
| 80 | p.Arg80Val | 4.4 |
| 80 | p.Arg80Tyr | 5.2 |
| 80 | p.Arg80Cys | 3.4 |
| 80 | p.Arg80Trp | 5.8 |
| 80 | p.Arg80Phe | 3.4 |
| 81 | p.Pro81Asn | 5.8 |
| 81 | p.Pro81Lys | 3.9 |
| 81 | p.Pro81Thr | 5.0 |
| 81 | p.Pro81Arg | 5.7 |
| 81 | p.Pro81Ser | 5.2 |
| 81 | p.Pro81Ile | 3.1 |
| 81 | p.Pro81Met | 4.5 |
| 81 | p.Pro81His | 5.3 |
| 81 | p.Pro81Gln | 5.0 |
| 81 | p.Pro81Pro | 9.4 |
| 81 | p.Pro81Leu | 3.8 |
| 81 | p.Pro81Asp | 3.8 |
| 81 | p.Pro81Glu | 4.8 |
| 81 | p.Pro81Ala | 4.7 |
| 81 | p.Pro81Gly | 3.5 |
| 81 | p.Pro81Val | 4.4 |

| | | |
|---|---|---|
| 81 | p.Pro81Tyr | 5.0 |
| 81 | p.Pro81Cys | 5.9 |
| 81 | p.Pro81Trp | 5.7 |
| 81 | p.Pro81Phe | 5.4 |
| 82 | p.Val82Asn | 3.3 |
| 82 | p.Val82Lys | 4.1 |
| 82 | p.Val82Thr | 6.5 |
| 82 | p.Val82Arg | 5.4 |
| 82 | p.Val82Ser | 4.5 |
| 82 | p.Val82Ile | 4.1 |
| 82 | p.Val82Met | 4.4 |
| 82 | p.Val82His | 5.5 |
| 82 | p.Val82Gln | 5.2 |
| 82 | p.Val82Pro | 4.7 |
| 82 | p.Val82Leu | 6.0 |
| 82 | p.Val82Asp | 3.8 |
| 82 | p.Val82Glu | 4.5 |
| 82 | p.Val82Ala | 5.4 |
| 82 | p.Val82Gly | 4.0 |
| 82 | p.Val82Val | 11.1 |
| 82 | p.Val82Tyr | 3.2 |
| 82 | p.Val82Cys | 4.6 |
| 82 | p.Val82Trp | 3.2 |
| 82 | p.Val82Phe | 6.4 |
| 83 | p.His83Asn | 5.4 |
| 83 | p.His83Lys | 5.1 |
| 83 | p.His83Thr | 3.7 |
| 83 | p.His83Arg | 4.3 |
| 83 | p.His83Ser | 4.5 |
| 83 | p.His83Ile | 4.1 |
| 83 | p.His83Met | 4.1 |
| 83 | p.His83His | 9.3 |
| 83 | p.His83Gln | 8.1 |
| 83 | p.His83Pro | 4.7 |
| 83 | p.His83Leu | 4.4 |
| 83 | p.His83Asp | 4.2 |
| 83 | p.His83Glu | 5.1 |
| 83 | p.His83Ala | 4.8 |
| 83 | p.His83Gly | 5.9 |
| 83 | p.His83Val | 3.6 |
| 83 | p.His83Tyr | 4.9 |
| 83 | p.His83Cys | 3.6 |
| 83 | p.His83Trp | 4.1 |
| 83 | p.His83Phe | 5.9 |
| 84 | p.Asp84Asn | 2.9 |

| | | |
|---|---|---|
| 84 | p.Asp84Lys | 5.2 |
| 84 | p.Asp84Thr | 3.1 |
| 84 | p.Asp84Arg | 4.6 |
| 84 | p.Asp84Ser | 4.3 |
| 84 | p.Asp84Ile | 4.8 |
| 84 | p.Asp84Met | 4.2 |
| 84 | p.Asp84His | 4.9 |
| 84 | p.Asp84Gln | 4.3 |
| 84 | p.Asp84Pro | 4.9 |
| 84 | p.Asp84Leu | 5.3 |
| 84 | p.Asp84Asp | 7.6 |
| 84 | p.Asp84Glu | 9.1 |
| 84 | p.Asp84Ala | 5.6 |
| 84 | p.Asp84Gly | 4.3 |
| 84 | p.Asp84Val | 4.4 |
| 84 | p.Asp84Tyr | 5.0 |
| 84 | p.Asp84Cys | 5.6 |
| 84 | p.Asp84Trp | 4.5 |
| 84 | p.Asp84Phe | 5.1 |
| 85 | p.Ala85Asn | 3.7 |
| 85 | p.Ala85Lys | 4.0 |
| 85 | p.Ala85Thr | 7.2 |
| 85 | p.Ala85Arg | 2.8 |
| 85 | p.Ala85Ser | 4.3 |
| 85 | p.Ala85Ile | 4.5 |
| 85 | p.Ala85Met | 2.7 |
| 85 | p.Ala85His | 3.9 |
| 85 | p.Ala85Gln | 3.0 |
| 85 | p.Ala85Pro | 6.6 |
| 85 | p.Ala85Leu | 4.6 |
| 85 | p.Ala85Asp | 10.4 |
| 85 | p.Ala85Glu | 4.3 |
| 85 | p.Ala85Ala | 10.3 |
| 85 | p.Ala85Gly | 7.5 |
| 85 | p.Ala85Val | 5.3 |
| 85 | p.Ala85Tyr | 2.7 |
| 85 | p.Ala85Cys | 4.0 |
| 85 | p.Ala85Trp | 3.3 |
| 85 | p.Ala85Phe | 4.9 |
| 86 | p.Ala86Asn | 3.9 |
| 86 | p.Ala86Lys | 3.0 |
| 86 | p.Ala86Thr | 2.2 |
| 86 | p.Ala86Arg | 4.7 |
| 86 | p.Ala86Ser | 3.1 |
| 86 | p.Ala86Ile | 4.8 |

| | | |
|---|---|---|
| 86 | p.Ala86Met | 4.9 |
| 86 | p.Ala86His | 5.7 |
| 86 | p.Ala86Gln | 6.9 |
| 86 | p.Ala86Pro | 4.2 |
| 86 | p.Ala86Leu | 6.0 |
| 86 | p.Ala86Asp | 4.5 |
| 86 | p.Ala86Glu | 13.1 |
| 86 | p.Ala86Ala | 4.4 |
| 86 | p.Ala86Gly | 3.4 |
| 86 | p.Ala86Val | 7.9 |
| 86 | p.Ala86Tyr | 4.1 |
| 86 | p.Ala86Cys | 4.1 |
| 86 | p.Ala86Trp | 3.8 |
| 86 | p.Ala86Phe | 5.1 |
| 87 | p.Arg87Asn | 6.8 |
| 87 | p.Arg87Lys | 3.4 |
| 87 | p.Arg87Thr | 3.8 |
| 87 | p.Arg87Arg | 5.1 |
| 87 | p.Arg87Ser | 4.5 |
| 87 | p.Arg87Ile | 4.3 |
| 87 | p.Arg87Met | 6.0 |
| 87 | p.Arg87His | 4.2 |
| 87 | p.Arg87Gln | 5.5 |
| 87 | p.Arg87Pro | 4.9 |
| 87 | p.Arg87Leu | 4.2 |
| 87 | p.Arg87Asp | 3.6 |
| 87 | p.Arg87Glu | 5.9 |
| 87 | p.Arg87Ala | 4.9 |
| 87 | p.Arg87Gly | 4.0 |
| 87 | p.Arg87Val | 5.1 |
| 87 | p.Arg87Tyr | 3.4 |
| 87 | p.Arg87Cys | 6.7 |
| 87 | p.Arg87Trp | 6.6 |
| 87 | p.Arg87Phe | 7.2 |
| 88 | p.Glu88Asn | 4.6 |
| 88 | p.Glu88Lys | 5.3 |
| 88 | p.Glu88Thr | 2.7 |
| 88 | p.Glu88Arg | 4.5 |
| 88 | p.Glu88Ser | 3.3 |
| 88 | p.Glu88Ile | 3.1 |
| 88 | p.Glu88Met | 5.8 |
| 88 | p.Glu88His | 4.2 |
| 88 | p.Glu88Gln | 3.8 |
| 88 | p.Glu88Pro | 3.3 |
| 88 | p.Glu88Leu | 4.3 |

| | | |
|---|---|---|
| 88 | p.Glu88Asp | 11.0 |
| 88 | p.Glu88Glu | 8.4 |
| 88 | p.Glu88Ala | 3.1 |
| 88 | p.Glu88Gly | 4.5 |
| 88 | p.Glu88Val | 7.8 |
| 88 | p.Glu88Tyr | 5.9 |
| 88 | p.Glu88Cys | 4.6 |
| 88 | p.Glu88Trp | 5.6 |
| 88 | p.Glu88Phe | 4.3 |
| 89 | p.Gly89Asn | 4.0 |
| 89 | p.Gly89Lys | 5.3 |
| 89 | p.Gly89Thr | 4.1 |
| 89 | p.Gly89Arg | 4.5 |
| 89 | p.Gly89Ser | 5.0 |
| 89 | p.Gly89Ile | 4.1 |
| 89 | p.Gly89Met | 4.3 |
| 89 | p.Gly89His | 4.1 |
| 89 | p.Gly89Gln | 5.1 |
| 89 | p.Gly89Pro | 4.8 |
| 89 | p.Gly89Leu | 4.5 |
| 89 | p.Gly89Asp | 5.2 |
| 89 | p.Gly89Glu | 6.9 |
| 89 | p.Gly89Ala | 3.6 |
| 89 | p.Gly89Gly | 4.9 |
| 89 | p.Gly89Val | 7.7 |
| 89 | p.Gly89Tyr | 3.8 |
| 89 | p.Gly89Cys | 5.0 |
| 89 | p.Gly89Trp | 7.4 |
| 89 | p.Gly89Phe | 5.7 |
| 90 | p.Phe90Asn | 7.3 |
| 90 | p.Phe90Lys | 5.6 |
| 90 | p.Phe90Thr | 4.0 |
| 90 | p.Phe90Arg | 5.8 |
| 90 | p.Phe90Ser | 5.0 |
| 90 | p.Phe90Ile | 4.3 |
| 90 | p.Phe90Met | 4.1 |
| 90 | p.Phe90His | 3.0 |
| 90 | p.Phe90Gln | 3.3 |
| 90 | p.Phe90Pro | 4.8 |
| 90 | p.Phe90Leu | 4.8 |
| 90 | p.Phe90Asp | 4.3 |
| 90 | p.Phe90Glu | 5.4 |
| 90 | p.Phe90Ala | 3.0 |
| 90 | p.Phe90Gly | 4.7 |
| 90 | p.Phe90Val | 5.4 |

| | | |
|---|---|---|
| 90 | p.Phe90Tyr | 5.4 |
| 90 | p.Phe90Cys | 5.8 |
| 90 | p.Phe90Trp | 5.3 |
| 90 | p.Phe90Phe | 8.8 |
| 91 | p.Leu91Asn | 4.1 |
| 91 | p.Leu91Lys | 3.3 |
| 91 | p.Leu91Thr | 3.6 |
| 91 | p.Leu91Arg | 4.7 |
| 91 | p.Leu91Ser | 4.5 |
| 91 | p.Leu91Ile | 9.0 |
| 91 | p.Leu91Met | 4.8 |
| 91 | p.Leu91His | 5.7 |
| 91 | p.Leu91Gln | 3.2 |
| 91 | p.Leu91Pro | 11.3 |
| 91 | p.Leu91Leu | 6.3 |
| 91 | p.Leu91Asp | 3.3 |
| 91 | p.Leu91Glu | 4.1 |
| 91 | p.Leu91Ala | 4.1 |
| 91 | p.Leu91Gly | 4.0 |
| 91 | p.Leu91Val | 4.2 |
| 91 | p.Leu91Tyr | 3.2 |
| 91 | p.Leu91Cys | 3.7 |
| 91 | p.Leu91Trp | 4.4 |
| 91 | p.Leu91Phe | 8.4 |
| 92 | p.Asp92Asn | 3.7 |
| 92 | p.Asp92Lys | 3.8 |
| 92 | p.Asp92Thr | 3.8 |
| 92 | p.Asp92Arg | 3.5 |
| 92 | p.Asp92Ser | 2.7 |
| 92 | p.Asp92Ile | 5.8 |
| 92 | p.Asp92Met | 6.3 |
| 92 | p.Asp92His | 2.8 |
| 92 | p.Asp92Gln | 5.5 |
| 92 | p.Asp92Pro | 5.5 |
| 92 | p.Asp92Leu | 5.3 |
| 92 | p.Asp92Asp | 6.8 |
| 92 | p.Asp92Glu | 8.1 |
| 92 | p.Asp92Ala | 5.8 |
| 92 | p.Asp92Gly | 5.0 |
| 92 | p.Asp92Val | 7.3 |
| 92 | p.Asp92Tyr | 3.8 |
| 92 | p.Asp92Cys | 6.2 |
| 92 | p.Asp92Trp | 3.6 |
| 92 | p.Asp92Phe | 4.8 |
| 93 | p.Thr93Asn | 9.2 |

| | | |
|---|---|---|
| 93 | p.Thr93Lys | 3.1 |
| 93 | p.Thr93Thr | 13.5 |
| 93 | p.Thr93Arg | 4.0 |
| 93 | p.Thr93Ser | 3.7 |
| 93 | p.Thr93Ile | 6.2 |
| 93 | p.Thr93Met | 3.8 |
| 93 | p.Thr93His | 4.1 |
| 93 | p.Thr93Gln | 3.9 |
| 93 | p.Thr93Pro | 10.3 |
| 93 | p.Thr93Leu | 3.1 |
| 93 | p.Thr93Asp | 2.8 |
| 93 | p.Thr93Glu | 1.6 |
| 93 | p.Thr93Ala | 6.5 |
| 93 | p.Thr93Gly | 4.9 |
| 93 | p.Thr93Val | 3.3 |
| 93 | p.Thr93Tyr | 3.8 |
| 93 | p.Thr93Cys | 4.3 |
| 93 | p.Thr93Trp | 3.7 |
| 93 | p.Thr93Phe | 4.2 |
| 94 | p.Leu94Asn | 3.7 |
| 94 | p.Leu94Lys | 5.7 |
| 94 | p.Leu94Thr | 5.4 |
| 94 | p.Leu94Arg | 6.4 |
| 94 | p.Leu94Ser | 4.1 |
| 94 | p.Leu94Ile | 4.5 |
| 94 | p.Leu94Met | 2.1 |
| 94 | p.Leu94His | 5.0 |
| 94 | p.Leu94Gln | 3.8 |
| 94 | p.Leu94Pro | 3.7 |
| 94 | p.Leu94Leu | 10.6 |
| 94 | p.Leu94Asp | 4.9 |
| 94 | p.Leu94Glu | 3.8 |
| 94 | p.Leu94Ala | 3.7 |
| 94 | p.Leu94Gly | 3.2 |
| 94 | p.Leu94Val | 6.1 |
| 94 | p.Leu94Tyr | 5.0 |
| 94 | p.Leu94Cys | 5.2 |
| 94 | p.Leu94Trp | 6.6 |
| 94 | p.Leu94Phe | 6.5 |
| 95 | p.Val95Asn | 3.9 |
| 95 | p.Val95Lys | 4.7 |
| 95 | p.Val95Thr | 3.7 |
| 95 | p.Val95Arg | 5.1 |
| 95 | p.Val95Ser | 4.4 |
| 95 | p.Val95Ile | 8.2 |

| | | |
|---|---|---|
| 95 | p.Val95Met | 2.8 |
| 95 | p.Val95His | 0.8 |
| 95 | p.Val95Gln | 1.5 |
| 95 | p.Val95Pro | 3.1 |
| 95 | p.Val95Leu | 2.8 |
| 95 | p.Val95Asp | 5.7 |
| 95 | p.Val95Glu | 4.6 |
| 95 | p.Val95Ala | 5.3 |
| 95 | p.Val95Gly | 8.5 |
| 95 | p.Val95Val | 8.5 |
| 95 | p.Val95Tyr | 2.7 |
| 95 | p.Val95Cys | 3.2 |
| 95 | p.Val95Trp | 4.0 |
| 95 | p.Val95Phe | 16.5 |
| 96 | p.Val96Asn | 4.0 |
| 96 | p.Val96Lys | 4.5 |
| 96 | p.Val96Thr | 2.1 |
| 96 | p.Val96Arg | 3.5 |
| 96 | p.Val96Ser | 4.4 |
| 96 | p.Val96Ile | 7.2 |
| 96 | p.Val96Met | 3.4 |
| 96 | p.Val96His | 3.1 |
| 96 | p.Val96Gln | 3.9 |
| 96 | p.Val96Pro | 2.2 |
| 96 | p.Val96Leu | 4.6 |
| 96 | p.Val96Asp | 7.2 |
| 96 | p.Val96Glu | 3.8 |
| 96 | p.Val96Ala | 6.9 |
| 96 | p.Val96Gly | 11.5 |
| 96 | p.Val96Val | 5.3 |
| 96 | p.Val96Tyr | 3.0 |
| 96 | p.Val96Cys | 3.8 |
| 96 | p.Val96Trp | 4.2 |
| 96 | p.Val96Phe | 11.5 |
| 97 | p.Leu97Asn | 4.3 |
| 97 | p.Leu97Lys | 5.7 |
| 97 | p.Leu97Thr | 3.2 |
| 97 | p.Leu97Arg | 3.8 |
| 97 | p.Leu97Ser | 5.7 |
| 97 | p.Leu97Ile | 5.2 |
| 97 | p.Leu97Met | 5.4 |
| 97 | p.Leu97His | 3.5 |
| 97 | p.Leu97Gln | 5.5 |
| 97 | p.Leu97Pro | 3.4 |
| 97 | p.Leu97Leu | 7.2 |

| | | |
|---|---|---|
| 97 | p.Leu97Asp | 5.1 |
| 97 | p.Leu97Glu | 6.0 |
| 97 | p.Leu97Ala | 3.9 |
| 97 | p.Leu97Gly | 5.7 |
| 97 | p.Leu97Val | 5.3 |
| 97 | p.Leu97Tyr | 4.4 |
| 97 | p.Leu97Cys | 4.6 |
| 97 | p.Leu97Trp | 6.6 |
| 97 | p.Leu97Phe | 5.3 |
| 98 | p.His98Asn | 4.1 |
| 98 | p.His98Lys | 4.1 |
| 98 | p.His98Thr | 2.3 |
| 98 | p.His98Arg | 4.2 |
| 98 | p.His98Ser | 4.7 |
| 98 | p.His98Ile | 3.8 |
| 98 | p.His98Met | 4.4 |
| 98 | p.His98His | 7.3 |
| 98 | p.His98Gln | 12.0 |
| 98 | p.His98Pro | 3.1 |
| 98 | p.His98Leu | 4.1 |
| 98 | p.His98Asp | 4.7 |
| 98 | p.His98Glu | 6.5 |
| 98 | p.His98Ala | 4.0 |
| 98 | p.His98Gly | 5.0 |
| 98 | p.His98Val | 6.5 |
| 98 | p.His98Tyr | 6.0 |
| 98 | p.His98Cys | 4.0 |
| 98 | p.His98Trp | 5.3 |
| 98 | p.His98Phe | 4.1 |
| 99 | p.Arg99Asn | 4.2 |
| 99 | p.Arg99Lys | 4.3 |
| 99 | p.Arg99Thr | 4.5 |
| 99 | p.Arg99Arg | 8.9 |
| 99 | p.Arg99Ser | 5.7 |
| 99 | p.Arg99Ile | 5.4 |
| 99 | p.Arg99Met | 4.2 |
| 99 | p.Arg99His | 4.8 |
| 99 | p.Arg99Gln | 3.0 |
| 99 | p.Arg99Pro | 5.1 |
| 99 | p.Arg99Leu | 3.9 |
| 99 | p.Arg99Asp | 4.9 |
| 99 | p.Arg99Glu | 5.0 |
| 99 | p.Arg99Ala | 6.2 |
| 99 | p.Arg99Gly | 4.9 |
| 99 | p.Arg99Val | 4.0 |

| | | |
|---|---|---|
| 99 | p.Arg99Tyr | 4.2 |
| 99 | p.Arg99Cys | 5.4 |
| 99 | p.Arg99Trp | 6.9 |
| 99 | p.Arg99Phe | 4.5 |
| 100 | p.Ala100Asn | 6.8 |
| 100 | p.Ala100Lys | 8.6 |
| 100 | p.Ala100Thr | 6.2 |
| 100 | p.Ala100Arg | 8.2 |
| 100 | p.Ala100Ser | 2.7 |
| 100 | p.Ala100Ile | 5.8 |
| 100 | p.Ala100Met | 8.7 |
| 100 | p.Ala100His | 5.3 |
| 100 | p.Ala100Gln | 4.4 |
| 100 | p.Ala100Pro | 4.7 |
| 100 | p.Ala100Leu | 4.9 |
| 100 | p.Ala100Asp | 1.2 |
| 100 | p.Ala100Glu | 4.6 |
| 100 | p.Ala100Ala | 4.1 |
| 100 | p.Ala100Gly | 0.2 |
| 100 | p.Ala100Val | 5.4 |
| 100 | p.Ala100Tyr | 4.6 |
| 100 | p.Ala100Cys | 3.5 |
| 100 | p.Ala100Trp | 6.8 |
| 100 | p.Ala100Phe | 3.4 |
| 101 | p.Gly101Asn | 4.1 |
| 101 | p.Gly101Lys | 5.8 |
| 101 | p.Gly101Thr | 4.4 |
| 101 | p.Gly101Arg | 5.2 |
| 101 | p.Gly101Ser | 5.8 |
| 101 | p.Gly101Ile | 4.4 |
| 101 | p.Gly101Met | 5.9 |
| 101 | p.Gly101His | 3.6 |
| 101 | p.Gly101Gln | 5.3 |
| 101 | p.Gly101Pro | 3.7 |
| 101 | p.Gly101Leu | 4.7 |
| 101 | p.Gly101Asp | 5.5 |
| 101 | p.Gly101Glu | 4.8 |
| 101 | p.Gly101Ala | 3.4 |
| 101 | p.Gly101Gly | 5.7 |
| 101 | p.Gly101Val | 4.7 |
| 101 | p.Gly101Tyr | 5.2 |
| 101 | p.Gly101Cys | 7.0 |
| 101 | p.Gly101Trp | 5.5 |
| 101 | p.Gly101Phe | 5.4 |
| 102 | p.Ala102Asn | 2.4 |

| | | |
|---|---|---|
| 102 | p.Ala102Lys | 5.0 |
| 102 | p.Ala102Thr | 5.3 |
| 102 | p.Ala102Arg | 4.5 |
| 102 | p.Ala102Ser | 2.1 |
| 102 | p.Ala102Ile | 4.3 |
| 102 | p.Ala102Met | 4.2 |
| 102 | p.Ala102His | 3.5 |
| 102 | p.Ala102Gln | 5.5 |
| 102 | p.Ala102Pro | 3.3 |
| 102 | p.Ala102Leu | 6.0 |
| 102 | p.Ala102Asp | 11.0 |
| 102 | p.Ala102Glu | 6.8 |
| 102 | p.Ala102Ala | 9.3 |
| 102 | p.Ala102Gly | 5.1 |
| 102 | p.Ala102Val | 7.2 |
| 102 | p.Ala102Tyr | 2.3 |
| 102 | p.Ala102Cys | 2.4 |
| 102 | p.Ala102Trp | 7.6 |
| 102 | p.Ala102Phe | 2.2 |
| 103 | p.Arg103Asn | 8.0 |
| 103 | p.Arg103Lys | 0.1 |
| 103 | p.Arg103Thr | 4.6 |
| 103 | p.Arg103Arg | 2.6 |
| 103 | p.Arg103Ser | 9.1 |
| 103 | p.Arg103Ile | 4.7 |
| 103 | p.Arg103Met | 2.0 |
| 103 | p.Arg103His | 4.9 |
| 103 | p.Arg103Gln | 1.0 |
| 103 | p.Arg103Pro | 2.7 |
| 103 | p.Arg103Leu | 0.8 |
| 103 | p.Arg103Asp | 5.3 |
| 103 | p.Arg103Glu | 1.1 |
| 103 | p.Arg103Ala | 5.8 |
| 103 | p.Arg103Gly | 10.5 |
| 103 | p.Arg103Val | 2.1 |
| 103 | p.Arg103Tyr | 5.4 |
| 103 | p.Arg103Cys | 16.6 |
| 103 | p.Arg103Trp | 3.4 |
| 103 | p.Arg103Phe | 9.3 |
| 104 | p.Leu104Asn | 3.4 |
| 104 | p.Leu104Lys | 3.5 |
| 104 | p.Leu104Thr | 3.0 |
| 104 | p.Leu104Arg | 4.5 |
| 104 | p.Leu104Ser | 4.1 |
| 104 | p.Leu104Ile | 13.2 |

| | | |
|---|---|---|
| 104 | p.Leu104Met | 5.2 |
| 104 | p.Leu104His | 7.3 |
| 104 | p.Leu104Gln | 2.6 |
| 104 | p.Leu104Pro | 6.2 |
| 104 | p.Leu104Leu | 7.4 |
| 104 | p.Leu104Asp | 3.1 |
| 104 | p.Leu104Glu | 5.1 |
| 104 | p.Leu104Ala | 3.4 |
| 104 | p.Leu104Gly | 4.2 |
| 104 | p.Leu104Val | 5.6 |
| 104 | p.Leu104Tyr | 3.2 |
| 104 | p.Leu104Cys | 3.8 |
| 104 | p.Leu104Trp | 3.8 |
| 104 | p.Leu104Phe | 7.4 |
| 105 | p.Asp105Asn | 6.6 |
| 105 | p.Asp105Lys | 6.5 |
| 105 | p.Asp105Thr | 4.6 |
| 105 | p.Asp105Arg | 6.5 |
| 105 | p.Asp105Ser | 1.6 |
| 105 | p.Asp105Ile | 3.4 |
| 105 | p.Asp105Met | 2.4 |
| 105 | p.Asp105His | 4.7 |
| 105 | p.Asp105Gln | 5.3 |
| 105 | p.Asp105Pro | 2.9 |
| 105 | p.Asp105Leu | 5.0 |
| 105 | p.Asp105Asp | 6.7 |
| 105 | p.Asp105Glu | 8.3 |
| 105 | p.Asp105Ala | 4.9 |
| 105 | p.Asp105Gly | 1.9 |
| 105 | p.Asp105Val | 7.3 |
| 105 | p.Asp105Tyr | 6.5 |
| 105 | p.Asp105Cys | 2.1 |
| 105 | p.Asp105Trp | 6.7 |
| 105 | p.Asp105Phe | 6.1 |
| 106 | p.Val106Asn | 3.8 |
| 106 | p.Val106Lys | 4.2 |
| 106 | p.Val106Thr | 3.5 |
| 106 | p.Val106Arg | 2.0 |
| 106 | p.Val106Ser | 4.4 |
| 106 | p.Val106Ile | 7.2 |
| 106 | p.Val106Met | 4.4 |
| 106 | p.Val106His | 4.0 |
| 106 | p.Val106Gln | 3.8 |
| 106 | p.Val106Pro | 2.9 |
| 106 | p.Val106Leu | 2.4 |

| | | |
|---|---|---|
| 106 | p.Val106Asp | 7.5 |
| 106 | p.Val106Glu | 1.4 |
| 106 | p.Val106Ala | 8.6 |
| 106 | p.Val106Gly | 12.1 |
| 106 | p.Val106Val | 5.8 |
| 106 | p.Val106Tyr | 3.5 |
| 106 | p.Val106Cys | 5.5 |
| 106 | p.Val106Trp | 1.4 |
| 106 | p.Val106Phe | 11.7 |
| 107 | p.Arg107Asn | 3.2 |
| 107 | p.Arg107Lys | 3.2 |
| 107 | p.Arg107Thr | 4.9 |
| 107 | p.Arg107Arg | 4.6 |
| 107 | p.Arg107Ser | 3.7 |
| 107 | p.Arg107Ile | 2.4 |
| 107 | p.Arg107Met | 5.3 |
| 107 | p.Arg107His | 3.4 |
| 107 | p.Arg107Gln | 11.5 |
| 107 | p.Arg107Pro | 5.1 |
| 107 | p.Arg107Leu | 11.5 |
| 107 | p.Arg107Asp | 3.2 |
| 107 | p.Arg107Glu | 2.6 |
| 107 | p.Arg107Ala | 5.9 |
| 107 | p.Arg107Gly | 3.7 |
| 107 | p.Arg107Val | 5.4 |
| 107 | p.Arg107Tyr | 4.0 |
| 107 | p.Arg107Cys | 2.1 |
| 107 | p.Arg107Trp | 10.2 |
| 107 | p.Arg107Phe | 4.1 |
| 108 | p.Asp108Asn | 6.3 |
| 108 | p.Asp108Lys | 4.5 |
| 108 | p.Asp108Thr | 5.2 |
| 108 | p.Asp108Arg | 6.4 |
| 108 | p.Asp108Ser | 6.5 |
| 108 | p.Asp108Ile | 4.4 |
| 108 | p.Asp108Met | 4.9 |
| 108 | p.Asp108His | 3.7 |
| 108 | p.Asp108Gln | 3.1 |
| 108 | p.Asp108Pro | 4.1 |
| 108 | p.Asp108Leu | 3.0 |
| 108 | p.Asp108Asp | 5.1 |
| 108 | p.Asp108Glu | 8.1 |
| 108 | p.Asp108Ala | 3.2 |
| 108 | p.Asp108Gly | 5.2 |
| 108 | p.Asp108Val | 4.9 |

| | | |
|---|---|---|
| 108 | p.Asp108Tyr | 7.3 |
| 108 | p.Asp108Cys | 4.3 |
| 108 | p.Asp108Trp | 4.4 |
| 108 | p.Asp108Phe | 5.3 |
| 109 | p.Ala109Asn | 4.9 |
| 109 | p.Ala109Lys | 6.3 |
| 109 | p.Ala109Thr | 4.3 |
| 109 | p.Ala109Arg | 5.2 |
| 109 | p.Ala109Ser | 6.7 |
| 109 | p.Ala109Ile | 3.1 |
| 109 | p.Ala109Met | 5.1 |
| 109 | p.Ala109His | 5.4 |
| 109 | p.Ala109Gln | 5.8 |
| 109 | p.Ala109Pro | 6.2 |
| 109 | p.Ala109Leu | 4.9 |
| 109 | p.Ala109Asp | 0.8 |
| 109 | p.Ala109Glu | 5.8 |
| 109 | p.Ala109Ala | 7.3 |
| 109 | p.Ala109Gly | 3.7 |
| 109 | p.Ala109Val | 4.6 |
| 109 | p.Ala109Tyr | 3.3 |
| 109 | p.Ala109Cys | 5.2 |
| 109 | p.Ala109Trp | 6.3 |
| 109 | p.Ala109Phe | 5.0 |
| 110 | p.Trp110Asn | 4.5 |
| 110 | p.Trp110Lys | 5.8 |
| 110 | p.Trp110Thr | 5.0 |
| 110 | p.Trp110Arg | 4.5 |
| 110 | p.Trp110Ser | 7.0 |
| 110 | p.Trp110Ile | 8.0 |
| 110 | p.Trp110Met | 4.6 |
| 110 | p.Trp110His | 2.4 |
| 110 | p.Trp110Gln | 5.1 |
| 110 | p.Trp110Pro | 0.3 |
| 110 | p.Trp110Leu | 5.8 |
| 110 | p.Trp110Asp | 4.4 |
| 110 | p.Trp110Glu | 6.4 |
| 110 | p.Trp110Ala | 3.3 |
| 110 | p.Trp110Gly | 5.1 |
| 110 | p.Trp110Val | 10.8 |
| 110 | p.Trp110Tyr | 6.3 |
| 110 | p.Trp110Cys | 5.6 |
| 110 | p.Trp110Trp | 0.0 |
| 110 | p.Trp110Phe | 5.0 |
| 111 | p.Gly111Asn | 5.5 |

| | | |
|---|---|---|
| 111 | p.Gly111Lys | 5.9 |
| 111 | p.Gly111Thr | 2.4 |
| 111 | p.Gly111Arg | 8.1 |
| 111 | p.Gly111Ser | 5.6 |
| 111 | p.Gly111Ile | 3.5 |
| 111 | p.Gly111Met | 4.2 |
| 111 | p.Gly111His | 4.6 |
| 111 | p.Gly111Gln | 4.2 |
| 111 | p.Gly111Pro | 3.0 |
| 111 | p.Gly111Leu | 3.3 |
| 111 | p.Gly111Asp | 4.7 |
| 111 | p.Gly111Glu | 6.5 |
| 111 | p.Gly111Ala | 3.0 |
| 111 | p.Gly111Gly | 8.7 |
| 111 | p.Gly111Val | 5.1 |
| 111 | p.Gly111Tyr | 3.7 |
| 111 | p.Gly111Cys | 7.0 |
| 111 | p.Gly111Trp | 6.1 |
| 111 | p.Gly111Phe | 4.9 |
| 112 | p.Arg112Asn | 5.7 |
| 112 | p.Arg112Lys | 4.7 |
| 112 | p.Arg112Thr | 4.8 |
| 112 | p.Arg112Arg | 4.2 |
| 112 | p.Arg112Ser | 4.9 |
| 112 | p.Arg112Ile | 5.0 |
| 112 | p.Arg112Met | 4.7 |
| 112 | p.Arg112His | 4.3 |
| 112 | p.Arg112Gln | 8.9 |
| 112 | p.Arg112Pro | 4.1 |
| 112 | p.Arg112Leu | 6.5 |
| 112 | p.Arg112Asp | 3.9 |
| 112 | p.Arg112Glu | 4.7 |
| 112 | p.Arg112Ala | 5.2 |
| 112 | p.Arg112Gly | 4.2 |
| 112 | p.Arg112Val | 4.5 |
| 112 | p.Arg112Tyr | 4.9 |
| 112 | p.Arg112Cys | 2.7 |
| 112 | p.Arg112Trp | 7.4 |
| 112 | p.Arg112Phe | 4.6 |
| 113 | p.Leu113Asn | 2.7 |
| 113 | p.Leu113Lys | 4.3 |
| 113 | p.Leu113Thr | 4.2 |
| 113 | p.Leu113Arg | 2.2 |
| 113 | p.Leu113Ser | 3.8 |
| 113 | p.Leu113Ile | 8.8 |

| | | |
|---|---|---|
| 113 | p.Leu113Met | 2.6 |
| 113 | p.Leu113His | 8.3 |
| 113 | p.Leu113Gln | 3.1 |
| 113 | p.Leu113Pro | 22.7 |
| 113 | p.Leu113Leu | 5.3 |
| 113 | p.Leu113Asp | 3.7 |
| 113 | p.Leu113Glu | 3.4 |
| 113 | p.Leu113Ala | 2.7 |
| 113 | p.Leu113Gly | 3.6 |
| 113 | p.Leu113Val | 4.5 |
| 113 | p.Leu113Tyr | 1.8 |
| 113 | p.Leu113Cys | 3.3 |
| 113 | p.Leu113Trp | 3.2 |
| 113 | p.Leu113Phe | 5.7 |
| 114 | p.Pro114Asn | 5.1 |
| 114 | p.Pro114Lys | 3.6 |
| 114 | p.Pro114Thr | 3.7 |
| 114 | p.Pro114Arg | 5.1 |
| 114 | p.Pro114Ser | 4.7 |
| 114 | p.Pro114Ile | 3.6 |
| 114 | p.Pro114Met | 3.4 |
| 114 | p.Pro114His | 4.4 |
| 114 | p.Pro114Gln | 6.1 |
| 114 | p.Pro114Pro | 6.5 |
| 114 | p.Pro114Leu | 5.1 |
| 114 | p.Pro114Asp | 6.6 |
| 114 | p.Pro114Glu | 5.1 |
| 114 | p.Pro114Ala | 4.1 |
| 114 | p.Pro114Gly | 5.2 |
| 114 | p.Pro114Val | 5.7 |
| 114 | p.Pro114Tyr | 5.1 |
| 114 | p.Pro114Cys | 5.0 |
| 114 | p.Pro114Trp | 6.7 |
| 114 | p.Pro114Phe | 5.4 |
| 115 | p.Val115Asn | 4.1 |
| 115 | p.Val115Lys | 3.0 |
| 115 | p.Val115Thr | 3.8 |
| 115 | p.Val115Arg | 2.9 |
| 115 | p.Val115Ser | 4.0 |
| 115 | p.Val115Ile | 4.8 |
| 115 | p.Val115Met | 4.8 |
| 115 | p.Val115His | 3.7 |
| 115 | p.Val115Gln | 4.2 |
| 115 | p.Val115Pro | 5.1 |
| 115 | p.Val115Leu | 6.1 |

| | | |
|---|---|---|
| 115 | p.Val115Asp | 6.8 |
| 115 | p.Val115Glu | 3.7 |
| 115 | p.Val115Ala | 6.5 |
| 115 | p.Val115Gly | 12.5 |
| 115 | p.Val115Val | 7.0 |
| 115 | p.Val115Tyr | 2.7 |
| 115 | p.Val115Cys | 0.4 |
| 115 | p.Val115Trp | 4.0 |
| 115 | p.Val115Phe | 9.9 |
| 116 | p.Asp116Asn | 3.2 |
| 116 | p.Asp116Lys | 5.9 |
| 116 | p.Asp116Thr | 3.0 |
| 116 | p.Asp116Arg | 3.8 |
| 116 | p.Asp116Ser | 4.8 |
| 116 | p.Asp116Ile | 5.0 |
| 116 | p.Asp116Met | 3.8 |
| 116 | p.Asp116His | 5.3 |
| 116 | p.Asp116Gln | 3.7 |
| 116 | p.Asp116Pro | 2.2 |
| 116 | p.Asp116Leu | 4.5 |
| 116 | p.Asp116Asp | 9.8 |
| 116 | p.Asp116Glu | 14.4 |
| 116 | p.Asp116Ala | 3.8 |
| 116 | p.Asp116Gly | 5.1 |
| 116 | p.Asp116Val | 4.2 |
| 116 | p.Asp116Tyr | 4.1 |
| 116 | p.Asp116Cys | 5.6 |
| 116 | p.Asp116Trp | 3.4 |
| 116 | p.Asp116Phe | 4.3 |
| 117 | p.Leu117Asn | 4.0 |
| 117 | p.Leu117Lys | 5.6 |
| 117 | p.Leu117Thr | 3.9 |
| 117 | p.Leu117Arg | 4.1 |
| 117 | p.Leu117Ser | 4.4 |
| 117 | p.Leu117Ile | 10.9 |
| 117 | p.Leu117Met | 4.8 |
| 117 | p.Leu117His | 4.9 |
| 117 | p.Leu117Gln | 4.5 |
| 117 | p.Leu117Pro | 7.6 |
| 117 | p.Leu117Leu | 7.0 |
| 117 | p.Leu117Asp | 3.9 |
| 117 | p.Leu117Glu | 4.5 |
| 117 | p.Leu117Ala | 1.9 |
| 117 | p.Leu117Gly | 4.5 |
| 117 | p.Leu117Val | 4.4 |

| | | |
|---|---|---|
| 117 | p.Leu117Tyr | 4.7 |
| 117 | p.Leu117Cys | 3.8 |
| 117 | p.Leu117Trp | 3.5 |
| 117 | p.Leu117Phe | 7.0 |
| 118 | p.Ala118Asn | 5.1 |
| 118 | p.Ala118Lys | 4.1 |
| 118 | p.Ala118Thr | 4.6 |
| 118 | p.Ala118Arg | 6.0 |
| 118 | p.Ala118Ser | 4.1 |
| 118 | p.Ala118Ile | 4.4 |
| 118 | p.Ala118Met | 4.7 |
| 118 | p.Ala118His | 3.2 |
| 118 | p.Ala118Gln | 5.0 |
| 118 | p.Ala118Pro | 4.0 |
| 118 | p.Ala118Leu | 5.2 |
| 118 | p.Ala118Asp | 5.7 |
| 118 | p.Ala118Glu | 3.7 |
| 118 | p.Ala118Ala | 12.8 |
| 118 | p.Ala118Gly | 4.6 |
| 118 | p.Ala118Val | 5.7 |
| 118 | p.Ala118Tyr | 3.7 |
| 118 | p.Ala118Cys | 5.4 |
| 118 | p.Ala118Trp | 3.9 |
| 118 | p.Ala118Phe | 3.9 |
| 119 | p.Glu119Asn | 4.3 |
| 119 | p.Glu119Lys | 4.5 |
| 119 | p.Glu119Thr | 5.3 |
| 119 | p.Glu119Arg | 4.6 |
| 119 | p.Glu119Ser | 4.4 |
| 119 | p.Glu119Ile | 3.7 |
| 119 | p.Glu119Met | 5.7 |
| 119 | p.Glu119His | 5.4 |
| 119 | p.Glu119Gln | 3.5 |
| 119 | p.Glu119Pro | 4.6 |
| 119 | p.Glu119Leu | 4.6 |
| 119 | p.Glu119Asp | 9.8 |
| 119 | p.Glu119Glu | 8.5 |
| 119 | p.Glu119Ala | 4.0 |
| 119 | p.Glu119Gly | 5.3 |
| 119 | p.Glu119Val | 5.1 |
| 119 | p.Glu119Tyr | 4.2 |
| 119 | p.Glu119Cys | 5.1 |
| 119 | p.Glu119Trp | 3.9 |
| 119 | p.Glu119Phe | 3.8 |
| 120 | p.Glu120Asn | 7.3 |

| | | |
|---|---|---|
| 120 | p.Glu120Lys | 5.0 |
| 120 | p.Glu120Thr | 5.4 |
| 120 | p.Glu120Arg | 3.2 |
| 120 | p.Glu120Ser | 4.0 |
| 120 | p.Glu120Ile | 4.6 |
| 120 | p.Glu120Met | 3.7 |
| 120 | p.Glu120His | 5.2 |
| 120 | p.Glu120Gln | 5.0 |
| 120 | p.Glu120Pro | 3.7 |
| 120 | p.Glu120Leu | 3.6 |
| 120 | p.Glu120Asp | 10.2 |
| 120 | p.Glu120Glu | 8.7 |
| 120 | p.Glu120Ala | 3.5 |
| 120 | p.Glu120Gly | 2.9 |
| 120 | p.Glu120Val | 5.1 |
| 120 | p.Glu120Tyr | 2.7 |
| 120 | p.Glu120Cys | 4.7 |
| 120 | p.Glu120Trp | 5.0 |
| 120 | p.Glu120Phe | 6.4 |
| 121 | p.Leu121Asn | 3.6 |
| 121 | p.Leu121Lys | 4.2 |
| 121 | p.Leu121Thr | 4.8 |
| 121 | p.Leu121Arg | 4.2 |
| 121 | p.Leu121Ser | 3.9 |
| 121 | p.Leu121Ile | 4.5 |
| 121 | p.Leu121Met | 5.4 |
| 121 | p.Leu121His | 4.0 |
| 121 | p.Leu121Gln | 5.1 |
| 121 | p.Leu121Pro | 4.4 |
| 121 | p.Leu121Leu | 10.1 |
| 121 | p.Leu121Asp | 5.7 |
| 121 | p.Leu121Glu | 5.6 |
| 121 | p.Leu121Ala | 3.1 |
| 121 | p.Leu121Gly | 6.1 |
| 121 | p.Leu121Val | 6.2 |
| 121 | p.Leu121Tyr | 2.9 |
| 121 | p.Leu121Cys | 4.6 |
| 121 | p.Leu121Trp | 4.8 |
| 121 | p.Leu121Phe | 6.7 |
| 122 | p.Gly122Asn | 4.3 |
| 122 | p.Gly122Lys | 5.0 |
| 122 | p.Gly122Thr | 5.6 |
| 122 | p.Gly122Arg | 5.3 |
| 122 | p.Gly122Ser | 5.4 |
| 122 | p.Gly122Ile | 4.3 |

| | | |
|---|---|---|
| 122 | p.Gly122Met | 4.8 |
| 122 | p.Gly122His | 4.6 |
| 122 | p.Gly122Gln | 3.9 |
| 122 | p.Gly122Pro | 3.7 |
| 122 | p.Gly122Leu | 3.9 |
| 122 | p.Gly122Asp | 5.0 |
| 122 | p.Gly122Glu | 7.0 |
| 122 | p.Gly122Ala | 5.4 |
| 122 | p.Gly122Gly | 4.9 |
| 122 | p.Gly122Val | 6.2 |
| 122 | p.Gly122Tyr | 5.5 |
| 122 | p.Gly122Cys | 3.8 |
| 122 | p.Gly122Trp | 7.1 |
| 122 | p.Gly122Phe | 4.4 |
| 123 | p.His123Asn | 11.7 |
| 123 | p.His123Lys | 5.8 |
| 123 | p.His123Thr | 4.1 |
| 123 | p.His123Arg | 3.5 |
| 123 | p.His123Ser | 4.0 |
| 123 | p.His123Ile | 4.4 |
| 123 | p.His123Met | 5.1 |
| 123 | p.His123His | 3.8 |
| 123 | p.His123Gln | 5.8 |
| 123 | p.His123Pro | 5.0 |
| 123 | p.His123Leu | 4.1 |
| 123 | p.His123Asp | 5.1 |
| 123 | p.His123Glu | 5.3 |
| 123 | p.His123Ala | 3.7 |
| 123 | p.His123Gly | 3.1 |
| 123 | p.His123Val | 4.5 |
| 123 | p.His123Tyr | 6.1 |
| 123 | p.His123Cys | 5.4 |
| 123 | p.His123Trp | 5.3 |
| 123 | p.His123Phe | 4.0 |
| 124 | p.Arg124Asn | 3.9 |
| 124 | p.Arg124Lys | 6.7 |
| 124 | p.Arg124Thr | 3.7 |
| 124 | p.Arg124Arg | 8.4 |
| 124 | p.Arg124Ser | 5.4 |
| 124 | p.Arg124Ile | 4.7 |
| 124 | p.Arg124Met | 6.9 |
| 124 | p.Arg124His | 4.2 |
| 124 | p.Arg124Gln | 4.4 |
| 124 | p.Arg124Pro | 4.2 |
| 124 | p.Arg124Leu | 5.5 |

| | | |
|---|---|---|
| 124 | p.Arg124Asp | 3.7 |
| 124 | p.Arg124Glu | 5.3 |
| 124 | p.Arg124Ala | 4.4 |
| 124 | p.Arg124Gly | 3.4 |
| 124 | p.Arg124Val | 5.0 |
| 124 | p.Arg124Tyr | 4.2 |
| 124 | p.Arg124Cys | 4.2 |
| 124 | p.Arg124Trp | 6.7 |
| 124 | p.Arg124Phe | 5.3 |
| 125 | p.Asp125Asn | 6.5 |
| 125 | p.Asp125Lys | 5.1 |
| 125 | p.Asp125Thr | 5.4 |
| 125 | p.Asp125Arg | 4.7 |
| 125 | p.Asp125Ser | 4.0 |
| 125 | p.Asp125Ile | 4.5 |
| 125 | p.Asp125Met | 4.4 |
| 125 | p.Asp125His | 4.8 |
| 125 | p.Asp125Gln | 4.2 |
| 125 | p.Asp125Pro | 2.1 |
| 125 | p.Asp125Leu | 3.4 |
| 125 | p.Asp125Asp | 5.0 |
| 125 | p.Asp125Glu | 6.8 |
| 125 | p.Asp125Ala | 6.6 |
| 125 | p.Asp125Gly | 7.1 |
| 125 | p.Asp125Val | 4.7 |
| 125 | p.Asp125Tyr | 6.7 |
| 125 | p.Asp125Cys | 4.6 |
| 125 | p.Asp125Trp | 4.7 |
| 125 | p.Asp125Phe | 4.7 |
| 126 | p.Val126Asn | 4.2 |
| 126 | p.Val126Lys | 4.8 |
| 126 | p.Val126Thr | 3.8 |
| 126 | p.Val126Arg | 6.4 |
| 126 | p.Val126Ser | 4.6 |
| 126 | p.Val126Ile | 6.3 |
| 126 | p.Val126Met | 7.2 |
| 126 | p.Val126His | 4.9 |
| 126 | p.Val126Gln | 6.0 |
| 126 | p.Val126Pro | 5.2 |
| 126 | p.Val126Leu | 4.1 |
| 126 | p.Val126Asp | 3.5 |
| 126 | p.Val126Glu | 5.1 |
| 126 | p.Val126Ala | 4.9 |
| 126 | p.Val126Gly | 4.0 |
| 126 | p.Val126Val | 6.3 |

| | | |
|---|---|---|
| 126 | p.Val126Tyr | 4.7 |
| 126 | p.Val126Cys | 3.5 |
| 126 | p.Val126Trp | 5.4 |
| 126 | p.Val126Phe | 5.1 |
| 127 | p.Ala127Asn | 4.3 |
| 127 | p.Ala127Lys | 4.1 |
| 127 | p.Ala127Thr | 5.5 |
| 127 | p.Ala127Arg | 5.0 |
| 127 | p.Ala127Ser | 4.6 |
| 127 | p.Ala127Ile | 4.5 |
| 127 | p.Ala127Met | 5.0 |
| 127 | p.Ala127His | 4.0 |
| 127 | p.Ala127Gln | 4.5 |
| 127 | p.Ala127Pro | 2.9 |
| 127 | p.Ala127Leu | 4.4 |
| 127 | p.Ala127Asp | 7.6 |
| 127 | p.Ala127Glu | 5.2 |
| 127 | p.Ala127Ala | 6.8 |
| 127 | p.Ala127Gly | 5.2 |
| 127 | p.Ala127Val | 6.7 |
| 127 | p.Ala127Tyr | 3.9 |
| 127 | p.Ala127Cys | 4.6 |
| 127 | p.Ala127Trp | 5.6 |
| 127 | p.Ala127Phe | 5.5 |
| 128 | p.Arg128Asn | 4.1 |
| 128 | p.Arg128Lys | 4.8 |
| 128 | p.Arg128Thr | 4.7 |
| 128 | p.Arg128Arg | 5.2 |
| 128 | p.Arg128Ser | 8.5 |
| 128 | p.Arg128Ile | 4.7 |
| 128 | p.Arg128Met | 4.1 |
| 128 | p.Arg128His | 3.8 |
| 128 | p.Arg128Gln | 4.7 |
| 128 | p.Arg128Pro | 4.3 |
| 128 | p.Arg128Leu | 3.0 |
| 128 | p.Arg128Asp | 5.2 |
| 128 | p.Arg128Glu | 7.4 |
| 128 | p.Arg128Ala | 4.5 |
| 128 | p.Arg128Gly | 5.8 |
| 128 | p.Arg128Val | 5.4 |
| 128 | p.Arg128Tyr | 4.6 |
| 128 | p.Arg128Cys | 4.5 |
| 128 | p.Arg128Trp | 5.8 |
| 128 | p.Arg128Phe | 4.8 |
| 129 | p.Tyr129Asn | 3.3 |

| | | |
|---|---|---|
| 129 | p.Tyr129Lys | 4.5 |
| 129 | p.Tyr129Thr | 4.0 |
| 129 | p.Tyr129Arg | 4.7 |
| 129 | p.Tyr129Ser | 3.6 |
| 129 | p.Tyr129Ile | 4.4 |
| 129 | p.Tyr129Met | 4.6 |
| 129 | p.Tyr129His | 4.8 |
| 129 | p.Tyr129Gln | 6.6 |
| 129 | p.Tyr129Pro | 4.3 |
| 129 | p.Tyr129Leu | 3.8 |
| 129 | p.Tyr129Asp | 6.2 |
| 129 | p.Tyr129Glu | 4.7 |
| 129 | p.Tyr129Ala | 6.6 |
| 129 | p.Tyr129Gly | 5.0 |
| 129 | p.Tyr129Val | 6.8 |
| 129 | p.Tyr129Tyr | 6.5 |
| 129 | p.Tyr129Cys | 4.3 |
| 129 | p.Tyr129Trp | 5.8 |
| 129 | p.Tyr129Phe | 5.1 |
| 130 | p.Leu130Asn | 2.8 |
| 130 | p.Leu130Lys | 3.3 |
| 130 | p.Leu130Thr | 3.4 |
| 130 | p.Leu130Arg | 3.1 |
| 130 | p.Leu130Ser | 4.8 |
| 130 | p.Leu130Ile | 5.7 |
| 130 | p.Leu130Met | 6.1 |
| 130 | p.Leu130His | 2.7 |
| 130 | p.Leu130Gln | 3.0 |
| 130 | p.Leu130Pro | 2.1 |
| 130 | p.Leu130Leu | 7.5 |
| 130 | p.Leu130Asp | 3.5 |
| 130 | p.Leu130Glu | 4.4 |
| 130 | p.Leu130Ala | 3.2 |
| 130 | p.Leu130Gly | 3.7 |
| 130 | p.Leu130Val | 12.3 |
| 130 | p.Leu130Tyr | 4.1 |
| 130 | p.Leu130Cys | 4.4 |
| 130 | p.Leu130Trp | 11.1 |
| 130 | p.Leu130Phe | 8.8 |
| 131 | p.Arg131Asn | 5.1 |
| 131 | p.Arg131Lys | 5.5 |
| 131 | p.Arg131Thr | 5.3 |
| 131 | p.Arg131Arg | 4.5 |
| 131 | p.Arg131Ser | 4.6 |
| 131 | p.Arg131Ile | 4.8 |

| | | |
|---|---|---|
| 131 | p.Arg131Met | 5.0 |
| 131 | p.Arg131His | 3.9 |
| 131 | p.Arg131Gln | 5.7 |
| 131 | p.Arg131Pro | 4.1 |
| 131 | p.Arg131Leu | 4.6 |
| 131 | p.Arg131Asp | 5.9 |
| 131 | p.Arg131Glu | 5.1 |
| 131 | p.Arg131Ala | 5.9 |
| 131 | p.Arg131Gly | 4.6 |
| 131 | p.Arg131Val | 3.6 |
| 131 | p.Arg131Tyr | 4.6 |
| 131 | p.Arg131Cys | 5.3 |
| 131 | p.Arg131Trp | 6.0 |
| 131 | p.Arg131Phe | 5.9 |
| 132 | p.Ala132Asn | 3.4 |
| 132 | p.Ala132Lys | 6.1 |
| 132 | p.Ala132Thr | 1.1 |
| 132 | p.Ala132Arg | 5.9 |
| 132 | p.Ala132Ser | 3.7 |
| 132 | p.Ala132Ile | 3.4 |
| 132 | p.Ala132Met | 4.6 |
| 132 | p.Ala132His | 2.8 |
| 132 | p.Ala132Gln | 8.9 |
| 132 | p.Ala132Pro | 4.9 |
| 132 | p.Ala132Leu | 5.0 |
| 132 | p.Ala132Asp | 4.0 |
| 132 | p.Ala132Glu | 5.0 |
| 132 | p.Ala132Ala | 10.1 |
| 132 | p.Ala132Gly | 3.9 |
| 132 | p.Ala132Val | 7.3 |
| 132 | p.Ala132Tyr | 4.3 |
| 132 | p.Ala132Cys | 5.8 |
| 132 | p.Ala132Trp | 4.4 |
| 132 | p.Ala132Phe | 5.5 |
| 133 | p.Ala133Asn | 4.0 |
| 133 | p.Ala133Lys | 3.7 |
| 133 | p.Ala133Thr | 8.5 |
| 133 | p.Ala133Arg | 4.7 |
| 133 | p.Ala133Ser | 6.6 |
| 133 | p.Ala133Ile | 5.2 |
| 133 | p.Ala133Met | 3.9 |
| 133 | p.Ala133His | 2.1 |
| 133 | p.Ala133Gln | 3.8 |
| 133 | p.Ala133Pro | 8.1 |
| 133 | p.Ala133Leu | 4.4 |

| | | |
|---|---|---|
| 133 | p.Ala133Asp | 8.7 |
| 133 | p.Ala133Glu | 4.6 |
| 133 | p.Ala133Ala | 7.7 |
| 133 | p.Ala133Gly | 5.4 |
| 133 | p.Ala133Val | 4.4 |
| 133 | p.Ala133Tyr | 5.4 |
| 133 | p.Ala133Cys | 2.6 |
| 133 | p.Ala133Trp | 2.9 |
| 133 | p.Ala133Phe | 3.3 |
| 134 | p.Ala134Asn | 4.8 |
| 134 | p.Ala134Lys | 2.6 |
| 134 | p.Ala134Thr | 6.5 |
| 134 | p.Ala134Arg | 2.2 |
| 134 | p.Ala134Ser | 4.7 |
| 134 | p.Ala134Ile | 5.1 |
| 134 | p.Ala134Met | 3.9 |
| 134 | p.Ala134His | 4.7 |
| 134 | p.Ala134Gln | 3.6 |
| 134 | p.Ala134Pro | 4.3 |
| 134 | p.Ala134Leu | 4.1 |
| 134 | p.Ala134Asp | 7.9 |
| 134 | p.Ala134Glu | 6.6 |
| 134 | p.Ala134Ala | 6.0 |
| 134 | p.Ala134Gly | 7.5 |
| 134 | p.Ala134Val | 5.9 |
| 134 | p.Ala134Tyr | 4.8 |
| 134 | p.Ala134Cys | 5.7 |
| 134 | p.Ala134Trp | 4.6 |
| 134 | p.Ala134Phe | 4.5 |
| 135 | p.Gly135Asn | 3.9 |
| 135 | p.Gly135Lys | 4.1 |
| 135 | p.Gly135Thr | 3.6 |
| 135 | p.Gly135Arg | 4.9 |
| 135 | p.Gly135Ser | 2.7 |
| 135 | p.Gly135Ile | 3.8 |
| 135 | p.Gly135Met | 6.3 |
| 135 | p.Gly135His | 6.5 |
| 135 | p.Gly135Gln | 4.3 |
| 135 | p.Gly135Pro | 5.1 |
| 135 | p.Gly135Leu | 4.0 |
| 135 | p.Gly135Asp | 3.8 |
| 135 | p.Gly135Glu | 3.6 |
| 135 | p.Gly135Ala | 4.9 |
| 135 | p.Gly135Gly | 10.5 |
| 135 | p.Gly135Val | 5.4 |

| | | |
|---|---|---|
| 135 | p.Gly135Tyr | 4.3 |
| 135 | p.Gly135Cys | 5.3 |
| 135 | p.Gly135Trp | 7.6 |
| 135 | p.Gly135Phe | 5.4 |
| 136 | p.Gly136Asn | 6.1 |
| 136 | p.Gly136Lys | 4.0 |
| 136 | p.Gly136Thr | 5.2 |
| 136 | p.Gly136Arg | 5.6 |
| 136 | p.Gly136Ser | 5.2 |
| 136 | p.Gly136Ile | 5.9 |
| 136 | p.Gly136Met | 4.6 |
| 136 | p.Gly136His | 4.1 |
| 136 | p.Gly136Gln | 5.6 |
| 136 | p.Gly136Pro | 5.1 |
| 136 | p.Gly136Leu | 5.2 |
| 136 | p.Gly136Asp | 4.9 |
| 136 | p.Gly136Glu | 6.3 |
| 136 | p.Gly136Ala | 4.6 |
| 136 | p.Gly136Gly | 9.7 |
| 136 | p.Gly136Val | 6.3 |
| 136 | p.Gly136Tyr | 3.0 |
| 136 | p.Gly136Cys | 0.6 |
| 136 | p.Gly136Trp | 3.7 |
| 136 | p.Gly136Phe | 4.5 |
| 137 | p.Thr137Asn | 3.1 |
| 137 | p.Thr137Lys | 22.3 |
| 137 | p.Thr137Thr | 3.7 |
| 137 | p.Thr137Arg | 4.6 |
| 137 | p.Thr137Ser | 4.7 |
| 137 | p.Thr137Ile | 4.2 |
| 137 | p.Thr137Met | 9.2 |
| 137 | p.Thr137His | 3.0 |
| 137 | p.Thr137Gln | 3.1 |
| 137 | p.Thr137Pro | 4.8 |
| 137 | p.Thr137Leu | 4.0 |
| 137 | p.Thr137Asp | 3.5 |
| 137 | p.Thr137Glu | 3.7 |
| 137 | p.Thr137Ala | 3.7 |
| 137 | p.Thr137Gly | 3.2 |
| 137 | p.Thr137Val | 4.7 |
| 137 | p.Thr137Tyr | 3.0 |
| 137 | p.Thr137Cys | 2.6 |
| 137 | p.Thr137Trp | 4.3 |
| 137 | p.Thr137Phe | 4.5 |
| 138 | p.Arg138Asn | 4.3 |

| | | |
|---|---|---|
| 138 | p.Arg138Lys | 7.1 |
| 138 | p.Arg138Thr | 4.2 |
| 138 | p.Arg138Arg | 7.9 |
| 138 | p.Arg138Ser | 5.2 |
| 138 | p.Arg138Ile | 4.4 |
| 138 | p.Arg138Met | 14.4 |
| 138 | p.Arg138His | 3.8 |
| 138 | p.Arg138Gln | 4.2 |
| 138 | p.Arg138Pro | 3.2 |
| 138 | p.Arg138Leu | 4.9 |
| 138 | p.Arg138Asp | 2.7 |
| 138 | p.Arg138Glu | 3.6 |
| 138 | p.Arg138Ala | 1.7 |
| 138 | p.Arg138Gly | 2.4 |
| 138 | p.Arg138Val | 3.7 |
| 138 | p.Arg138Tyr | 3.3 |
| 138 | p.Arg138Cys | 3.6 |
| 138 | p.Arg138Trp | 11.1 |
| 138 | p.Arg138Phe | 4.1 |
| 139 | p.Gly139Asn | 4.0 |
| 139 | p.Gly139Lys | 5.1 |
| 139 | p.Gly139Thr | 3.7 |
| 139 | p.Gly139Arg | 7.9 |
| 139 | p.Gly139Ser | 5.5 |
| 139 | p.Gly139Ile | 4.7 |
| 139 | p.Gly139Met | 4.7 |
| 139 | p.Gly139His | 3.8 |
| 139 | p.Gly139Gln | 4.8 |
| 139 | p.Gly139Pro | 4.8 |
| 139 | p.Gly139Leu | 4.9 |
| 139 | p.Gly139Asp | 4.5 |
| 139 | p.Gly139Glu | 5.4 |
| 139 | p.Gly139Ala | 3.9 |
| 139 | p.Gly139Gly | 7.1 |
| 139 | p.Gly139Val | 5.3 |
| 139 | p.Gly139Tyr | 3.7 |
| 139 | p.Gly139Cys | 5.9 |
| 139 | p.Gly139Trp | 5.7 |
| 139 | p.Gly139Phe | 4.4 |
| 140 | p.Ser140Asn | 5.8 |
| 140 | p.Ser140Lys | 4.9 |
| 140 | p.Ser140Thr | 6.5 |
| 140 | p.Ser140Arg | 4.8 |
| 140 | p.Ser140Ser | 3.3 |
| 140 | p.Ser140Ile | 5.3 |

| | | |
|---|---|---|
| 140 | p.Ser140Met | 4.5 |
| 140 | p.Ser140His | 5.3 |
| 140 | p.Ser140Gln | 5.1 |
| 140 | p.Ser140Pro | 4.0 |
| 140 | p.Ser140Leu | 4.5 |
| 140 | p.Ser140Asp | 5.7 |
| 140 | p.Ser140Glu | 4.9 |
| 140 | p.Ser140Ala | 3.8 |
| 140 | p.Ser140Gly | 2.9 |
| 140 | p.Ser140Val | 6.5 |
| 140 | p.Ser140Tyr | 5.7 |
| 140 | p.Ser140Cys | 6.0 |
| 140 | p.Ser140Trp | 5.5 |
| 140 | p.Ser140Phe | 5.1 |
| 141 | p.Asn141Asn | 21.3 |
| 141 | p.Asn141Lys | 19.1 |
| 141 | p.Asn141Thr | 2.9 |
| 141 | p.Asn141Arg | 3.6 |
| 141 | p.Asn141Ser | 3.0 |
| 141 | p.Asn141Ile | 3.5 |
| 141 | p.Asn141Met | 4.3 |
| 141 | p.Asn141His | 2.0 |
| 141 | p.Asn141Gln | 3.3 |
| 141 | p.Asn141Pro | 2.1 |
| 141 | p.Asn141Leu | 4.8 |
| 141 | p.Asn141Asp | 2.7 |
| 141 | p.Asn141Glu | 3.6 |
| 141 | p.Asn141Ala | 3.6 |
| 141 | p.Asn141Gly | 3.4 |
| 141 | p.Asn141Val | 4.0 |
| 141 | p.Asn141Tyr | 2.7 |
| 141 | p.Asn141Cys | 2.1 |
| 141 | p.Asn141Trp | 4.1 |
| 141 | p.Asn141Phe | 4.0 |
| 142 | p.His142Asn | 19.9 |
| 142 | p.His142Lys | 2.6 |
| 142 | p.His142Thr | 2.8 |
| 142 | p.His142Arg | 2.4 |
| 142 | p.His142Ser | 1.9 |
| 142 | p.His142Ile | 2.6 |
| 142 | p.His142Met | 1.6 |
| 142 | p.His142His | 5.0 |
| 142 | p.His142Gln | 3.8 |
| 142 | p.His142Pro | 29.0 |
| 142 | p.His142Leu | 3.0 |

| | | |
|---|---|---|
| 142 | p.His142Asp | 3.8 |
| 142 | p.His142Glu | 2.8 |
| 142 | p.His142Ala | 2.4 |
| 142 | p.His142Gly | 2.3 |
| 142 | p.His142Val | 2.8 |
| 142 | p.His142Tyr | 4.7 |
| 142 | p.His142Cys | 2.3 |
| 142 | p.His142Trp | 2.4 |
| 142 | p.His142Phe | 1.7 |
| 143 | p.Ala143Asn | 5.2 |
| 143 | p.Ala143Lys | 6.3 |
| 143 | p.Ala143Thr | 5.9 |
| 143 | p.Ala143Arg | 4.3 |
| 143 | p.Ala143Ser | 4.2 |
| 143 | p.Ala143Ile | 4.3 |
| 143 | p.Ala143Met | 4.8 |
| 143 | p.Ala143His | 3.5 |
| 143 | p.Ala143Gln | 5.9 |
| 143 | p.Ala143Pro | 3.4 |
| 143 | p.Ala143Leu | 3.9 |
| 143 | p.Ala143Asp | 4.8 |
| 143 | p.Ala143Glu | 5.4 |
| 143 | p.Ala143Ala | 8.9 |
| 143 | p.Ala143Gly | 4.3 |
| 143 | p.Ala143Val | 4.6 |
| 143 | p.Ala143Tyr | 5.8 |
| 143 | p.Ala143Cys | 4.2 |
| 143 | p.Ala143Trp | 5.0 |
| 143 | p.Ala143Phe | 5.3 |
| 144 | p.Arg144Asn | 6.1 |
| 144 | p.Arg144Lys | 4.1 |
| 144 | p.Arg144Thr | 5.3 |
| 144 | p.Arg144Arg | 5.0 |
| 144 | p.Arg144Ser | 5.4 |
| 144 | p.Arg144Ile | 3.8 |
| 144 | p.Arg144Met | 4.5 |
| 144 | p.Arg144His | 4.0 |
| 144 | p.Arg144Gln | 5.1 |
| 144 | p.Arg144Pro | 3.8 |
| 144 | p.Arg144Leu | 6.4 |
| 144 | p.Arg144Asp | 4.1 |
| 144 | p.Arg144Glu | 5.7 |
| 144 | p.Arg144Ala | 4.7 |
| 144 | p.Arg144Gly | 5.7 |
| 144 | p.Arg144Val | 5.1 |

| | | |
|---|---|---|
| 144 | p.Arg144Tyr | 4.9 |
| 144 | p.Arg144Cys | 6.5 |
| 144 | p.Arg144Trp | 4.9 |
| 144 | p.Arg144Phe | 4.9 |
| 145 | p.Ile145Asn | 7.0 |
| 145 | p.Ile145Lys | 2.4 |
| 145 | p.Ile145Thr | 10.7 |
| 145 | p.Ile145Arg | 2.7 |
| 145 | p.Ile145Ser | 21.9 |
| 145 | p.Ile145Ile | 18.9 |
| 145 | p.Ile145Met | 6.5 |
| 145 | p.Ile145His | 1.5 |
| 145 | p.Ile145Gln | 2.2 |
| 145 | p.Ile145Pro | 1.9 |
| 145 | p.Ile145Leu | 2.2 |
| 145 | p.Ile145Asp | 1.8 |
| 145 | p.Ile145Glu | 2.3 |
| 145 | p.Ile145Ala | 1.4 |
| 145 | p.Ile145Gly | 2.0 |
| 145 | p.Ile145Val | 2.0 |
| 145 | p.Ile145Tyr | 1.9 |
| 145 | p.Ile145Cys | 2.0 |
| 145 | p.Ile145Trp | 1.7 |
| 145 | p.Ile145Phe | 7.1 |
| 146 | p.Asp146Asn | 8.5 |
| 146 | p.Asp146Lys | 2.3 |
| 146 | p.Asp146Thr | 2.0 |
| 146 | p.Asp146Arg | 2.7 |
| 146 | p.Asp146Ser | 2.6 |
| 146 | p.Asp146Ile | 1.7 |
| 146 | p.Asp146Met | 2.6 |
| 146 | p.Asp146His | 6.3 |
| 146 | p.Asp146Gln | 1.9 |
| 146 | p.Asp146Pro | 2.4 |
| 146 | p.Asp146Leu | 2.1 |
| 146 | p.Asp146Asp | 8.3 |
| 146 | p.Asp146Glu | 8.2 |
| 146 | p.Asp146Ala | 21.1 |
| 146 | p.Asp146Gly | 8.2 |
| 146 | p.Asp146Val | 2.5 |
| 146 | p.Asp146Tyr | 10.6 |
| 146 | p.Asp146Cys | 1.4 |
| 146 | p.Asp146Trp | 2.2 |
| 146 | p.Asp146Phe | 2.5 |
| 147 | p.Ala147Asn | 3.8 |

| | | |
|---|---|---|
| 147 | p.Ala147Lys | 2.6 |
| 147 | p.Ala147Thr | 4.5 |
| 147 | p.Ala147Arg | 4.1 |
| 147 | p.Ala147Ser | 2.2 |
| 147 | p.Ala147Ile | 5.0 |
| 147 | p.Ala147Met | 4.8 |
| 147 | p.Ala147His | 4.5 |
| 147 | p.Ala147Gln | 5.0 |
| 147 | p.Ala147Pro | 3.7 |
| 147 | p.Ala147Leu | 4.4 |
| 147 | p.Ala147Asp | 3.5 |
| 147 | p.Ala147Glu | 7.2 |
| 147 | p.Ala147Ala | 20.2 |
| 147 | p.Ala147Gly | 2.9 |
| 147 | p.Ala147Val | 5.3 |
| 147 | p.Ala147Tyr | 4.2 |
| 147 | p.Ala147Cys | 3.7 |
| 147 | p.Ala147Trp | 4.8 |
| 147 | p.Ala147Phe | 3.6 |
| 148 | p.Ala148Asn | 4.3 |
| 148 | p.Ala148Lys | 4.1 |
| 148 | p.Ala148Thr | 6.0 |
| 148 | p.Ala148Arg | 4.5 |
| 148 | p.Ala148Ser | 4.9 |
| 148 | p.Ala148Ile | 4.9 |
| 148 | p.Ala148Met | 4.2 |
| 148 | p.Ala148His | 3.8 |
| 148 | p.Ala148Gln | 3.7 |
| 148 | p.Ala148Pro | 5.8 |
| 148 | p.Ala148Leu | 4.7 |
| 148 | p.Ala148Asp | 9.7 |
| 148 | p.Ala148Glu | 5.4 |
| 148 | p.Ala148Ala | 8.4 |
| 148 | p.Ala148Gly | 6.1 |
| 148 | p.Ala148Val | 4.1 |
| 148 | p.Ala148Tyr | 4.8 |
| 148 | p.Ala148Cys | 4.3 |
| 148 | p.Ala148Trp | 1.1 |
| 148 | p.Ala148Phe | 5.2 |
| 149 | p.Glu149Asn | 3.2 |
| 149 | p.Glu149Lys | 12.7 |
| 149 | p.Glu149Thr | 3.6 |
| 149 | p.Glu149Arg | 3.5 |
| 149 | p.Glu149Ser | 2.3 |
| 149 | p.Glu149Ile | 4.3 |

| | | |
|---|---|---|
| 149 | p.Glu149Met | 3.1 |
| 149 | p.Glu149His | 3.7 |
| 149 | p.Glu149Gln | 7.3 |
| 149 | p.Glu149Pro | 4.4 |
| 149 | p.Glu149Leu | 4.5 |
| 149 | p.Glu149Asp | 8.0 |
| 149 | p.Glu149Glu | 7.6 |
| 149 | p.Glu149Ala | 3.7 |
| 149 | p.Glu149Gly | 3.9 |
| 149 | p.Glu149Val | 10.0 |
| 149 | p.Glu149Tyr | 4.3 |
| 149 | p.Glu149Cys | 2.4 |
| 149 | p.Glu149Trp | 4.6 |
| 149 | p.Glu149Phe | 2.8 |
| 150 | p.Gly150Asn | 3.3 |
| 150 | p.Gly150Lys | 2.3 |
| 150 | p.Gly150Thr | 2.6 |
| 150 | p.Gly150Arg | 4.2 |
| 150 | p.Gly150Ser | 5.6 |
| 150 | p.Gly150Ile | 4.2 |
| 150 | p.Gly150Met | 3.6 |
| 150 | p.Gly150His | 2.4 |
| 150 | p.Gly150Gln | 1.3 |
| 150 | p.Gly150Pro | 3.5 |
| 150 | p.Gly150Leu | 2.7 |
| 150 | p.Gly150Asp | 8.0 |
| 150 | p.Gly150Glu | 3.2 |
| 150 | p.Gly150Ala | 16.1 |
| 150 | p.Gly150Gly | 13.1 |
| 150 | p.Gly150Val | 2.8 |
| 150 | p.Gly150Tyr | 4.1 |
| 150 | p.Gly150Cys | 11.0 |
| 150 | p.Gly150Trp | 2.5 |
| 150 | p.Gly150Phe | 3.5 |
| 151 | p.Pro151Asn | 5.7 |
| 151 | p.Pro151Lys | 6.9 |
| 151 | p.Pro151Thr | 4.4 |
| 151 | p.Pro151Arg | 4.2 |
| 151 | p.Pro151Ser | 1.2 |
| 151 | p.Pro151Ile | 4.7 |
| 151 | p.Pro151Met | 5.9 |
| 151 | p.Pro151His | 6.1 |
| 151 | p.Pro151Gln | 5.1 |
| 151 | p.Pro151Pro | 10.3 |
| 151 | p.Pro151Leu | 5.4 |

| | | |
|---|---|---|
| 151 | p.Pro151Asp | 2.6 |
| 151 | p.Pro151Glu | 5.5 |
| 151 | p.Pro151Ala | 2.9 |
| 151 | p.Pro151Gly | 0.3 |
| 151 | p.Pro151Val | 6.3 |
| 151 | p.Pro151Tyr | 6.6 |
| 151 | p.Pro151Cys | 4.1 |
| 151 | p.Pro151Trp | 6.6 |
| 151 | p.Pro151Phe | 5.3 |
| 152 | p.Ser152Asn | 5.1 |
| 152 | p.Ser152Lys | 2.3 |
| 152 | p.Ser152Thr | 6.5 |
| 152 | p.Ser152Arg | 21.3 |
| 152 | p.Ser152Ser | 9.3 |
| 152 | p.Ser152Ile | 2.5 |
| 152 | p.Ser152Met | 2.3 |
| 152 | p.Ser152His | 2.6 |
| 152 | p.Ser152Gln | 2.4 |
| 152 | p.Ser152Pro | 1.5 |
| 152 | p.Ser152Leu | 2.6 |
| 152 | p.Ser152Asp | 2.3 |
| 152 | p.Ser152Glu | 1.8 |
| 152 | p.Ser152Ala | 18.5 |
| 152 | p.Ser152Gly | 2.8 |
| 152 | p.Ser152Val | 2.7 |
| 152 | p.Ser152Tyr | 3.2 |
| 152 | p.Ser152Cys | 5.1 |
| 152 | p.Ser152Trp | 2.6 |
| 152 | p.Ser152Phe | 2.7 |
| 153 | p.Asp153Asn | 13.3 |
| 153 | p.Asp153Lys | 5.2 |
| 153 | p.Asp153Thr | 4.0 |
| 153 | p.Asp153Arg | 4.2 |
| 153 | p.Asp153Ser | 4.6 |
| 153 | p.Asp153Ile | 4.0 |
| 153 | p.Asp153Met | 4.1 |
| 153 | p.Asp153His | 4.9 |
| 153 | p.Asp153Gln | 3.9 |
| 153 | p.Asp153Pro | 2.5 |
| 153 | p.Asp153Leu | 4.3 |
| 153 | p.Asp153Asp | 4.9 |
| 153 | p.Asp153Glu | 5.4 |
| 153 | p.Asp153Ala | 5.3 |
| 153 | p.Asp153Gly | 6.2 |
| 153 | p.Asp153Val | 3.3 |

| | | |
|---|---|---|
| 153 | p.Asp153Tyr | 6.0 |
| 153 | p.Asp153Cys | 4.8 |
| 153 | p.Asp153Trp | 4.3 |
| 153 | p.Asp153Phe | 4.6 |
| 154 | p.Ile154Asn | 3.8 |
| 154 | p.Ile154Lys | 3.0 |
| 154 | p.Ile154Thr | 3.2 |
| 154 | p.Ile154Arg | 13.0 |
| 154 | p.Ile154Ser | 3.0 |
| 154 | p.Ile154Ile | 34.2 |
| 154 | p.Ile154Met | 7.7 |
| 154 | p.Ile154His | 2.3 |
| 154 | p.Ile154Gln | 2.5 |
| 154 | p.Ile154Pro | 1.6 |
| 154 | p.Ile154Leu | 2.7 |
| 154 | p.Ile154Asp | 2.9 |
| 154 | p.Ile154Glu | 3.0 |
| 154 | p.Ile154Ala | 2.2 |
| 154 | p.Ile154Gly | 0.5 |
| 154 | p.Ile154Val | 3.2 |
| 154 | p.Ile154Tyr | 2.6 |
| 154 | p.Ile154Cys | 1.9 |
| 154 | p.Ile154Trp | 2.8 |
| 154 | p.Ile154Phe | 4.0 |
| 155 | p.Pro155Asn | 4.6 |
| 155 | p.Pro155Lys | 3.4 |
| 155 | p.Pro155Thr | 5.4 |
| 155 | p.Pro155Arg | 5.6 |
| 155 | p.Pro155Ser | 4.3 |
| 155 | p.Pro155Ile | 3.9 |
| 155 | p.Pro155Met | 4.3 |
| 155 | p.Pro155His | 5.4 |
| 155 | p.Pro155Gln | 4.8 |
| 155 | p.Pro155Pro | 20.7 |
| 155 | p.Pro155Leu | 5.4 |
| 155 | p.Pro155Asp | 4.7 |
| 155 | p.Pro155Glu | 4.3 |
| 155 | p.Pro155Ala | 2.5 |
| 155 | p.Pro155Gly | 3.9 |
| 155 | p.Pro155Val | 3.8 |
| 155 | p.Pro155Tyr | 4.0 |
| 155 | p.Pro155Cys | 3.3 |
| 155 | p.Pro155Trp | 2.8 |
| 155 | p.Pro155Phe | 3.3 |
| 156 | p.Asp156Asn | 13.4 |

| | | |
|---|---|---|
| 156 | p.Asp156Lys | 2.7 |
| 156 | p.Asp156Thr | 2.5 |
| 156 | p.Asp156Arg | 2.0 |
| 156 | p.Asp156Ser | 2.6 |
| 156 | p.Asp156Ile | 1.7 |
| 156 | p.Asp156Met | 2.2 |
| 156 | p.Asp156His | 5.6 |
| 156 | p.Asp156Gln | 1.9 |
| 156 | p.Asp156Pro | 2.1 |
| 156 | p.Asp156Leu | 2.1 |
| 156 | p.Asp156Asp | 5.1 |
| 156 | p.Asp156Glu | 8.0 |
| 156 | p.Asp156Ala | 17.7 |
| 156 | p.Asp156Gly | 6.7 |
| 156 | p.Asp156Val | 1.9 |
| 156 | p.Asp156Tyr | 14.6 |
| 156 | p.Asp156Cys | 2.2 |
| 156 | p.Asp156Trp | 1.9 |
| 156 | p.Asp156Phe | 3.1 |
