## Appendix 1-table 3 for "Functional characterization of all *CDKN2A* missense variants and comparison to in silico models of pathogenicity"

**Appendix 1-table 3. Proportion of each variant in residues R24, H66, and A127.**

| Residue | Variant | ACMG Guidline Classification | Percent (%) | | Read count | |
|---|---|---|---|---|---|---|
| | | | Amplified plasmid library | Day 9 cells | Amplified plasmid library | Day 9 cells |
| 24 | p.Arg24Asn | VUS | 4.56 | 4.34 | 8516 | 4567 |
| 24 | p.Arg24Lys | VUS | 5.15 | 4.81 | 9629 | 5056 |
| 24 | p.Arg24Thr | VUS | 4.42 | 4.18 | 8250 | 4398 |
| 24 | p.Arg24Arg | Synonymous | 4.92 | 6.32 | 9196 | 6651 |
| 24 | p.Arg24Ser | VUS | 5.09 | 5.00 | 9504 | 5259 |
| 24 | p.Arg24Ile | VUS | 4.02 | 4.13 | 7503 | 4338 |
| 24 | p.Arg24Met | VUS | 4.84 | 5.22 | 9048 | 5493 |
| 24 | p.Arg24His | VUS | 5.24 | 5.07 | 9788 | 5328 |
| 24 | p.Arg24Gln | VUS | 5.64 | 5.29 | 10542 | 5563 |
| 24 | p.Arg24Pro | Pathogenic | 3.64 | 3.79 | 6803 | 3981 |
| 24 | p.Arg24Leu | VUS | 4.45 | 4.32 | 8319 | 4538 |
| 24 | p.Arg24Asp | VUS | 7.13 | 6.91 | 13323 | 7271 |
| 24 | p.Arg24Glu | VUS | 5.19 | 5.09 | 9692 | 5349 |
| 24 | p.Arg24Ala | VUS | 4.52 | 4.28 | 8447 | 4498 |
| 24 | p.Arg24Gly | VUS | 4.06 | 4.01 | 7595 | 4217 |
| 24 | p.Arg24Val | VUS | 6.48 | 6.42 | 12100 | 6754 |
| 24 | p.Arg24Tyr | VUS | 4.58 | 4.56 | 8553 | 4797 |
| 24 | p.Arg24Cys | VUS | 4.48 | 4.40 | 8366 | 4627 |
| 24 | p.Arg24Trp | VUS | 6.23 | 6.65 | 11641 | 6995 |
| 24 | p.Arg24Phe | VUS | 5.38 | 5.21 | 10044 | 5482 |
| 66 | p.His66Asn | VUS | 4.71 | 3.52 | 5574 | 1413 |
| 66 | p.His66Lys | VUS | 5.85 | 4.48 | 6935 | 1796 |
| 66 | p.His66Thr | VUS | 4.70 | 3.60 | 5573 | 1442 |
| 66 | p.His66Arg | VUS | 5.25 | 4.85 | 6216 | 1944 |
| 66 | p.His66Ser | VUS | 5.10 | 4.57 | 6038 | 1831 |
| 66 | p.His66Ile | VUS | 4.83 | 5.04 | 5726 | 2019 |
| 66 | p.His66Met | VUS | 5.30 | 3.81 | 6278 | 1527 |
| 66 | p.His66His | Synonymous | 3.88 | 4.94 | 4591 | 1982 |
| 66 | p.His66Gln | VUS | 5.28 | 8.89 | 6250 | 3565 |
| 66 | p.His66Pro | VUS | 6.13 | 6.07 | 7260 | 2432 |
| 66 | p.His66Leu | VUS | 5.55 | 5.83 | 6575 | 2338 |
| 66 | p.His66Asp | VUS | 2.77 | 2.92 | 3286 | 1169 |
| 66 | p.His66Glu | VUS | 6.10 | 6.04 | 7229 | 2420 |
| 66 | p.His66Ala | VUS | 5.41 | 5.95 | 6414 | 2387 |
| 66 | p.His66Gly | VUS | 5.56 | 5.24 | 6587 | 2100 |
| 66 | p.His66Val | VUS | 4.58 | 4.47 | 5430 | 1793 |
| 66 | p.His66Tyr | VUS | 4.55 | 4.66 | 5384 | 1869 |
| 66 | p.His66Cys | VUS | 4.53 | 4.63 | 5367 | 1856 |
| 66 | p.His66Trp | VUS | 5.40 | 5.55 | 6397 | 2225 |
| 66 | p.His66Phe | VUS | 4.51 | 4.94 | 5346 | 1981 |
| 127 | p.Ala127Asn | Benign | 4.62 | 4.37 | 3844 | 6274 |
| 127 | p.Ala127Lys | VUS | 4.37 | 4.36 | 3631 | 6261 |
| 127 | p.Ala127Thr | VUS | 4.44 | 3.99 | 3693 | 5737 |
| 127 | p.Ala127Arg | VUS | 5.85 | 5.79 | 4862 | 8321 |
| 127 | p.Ala127Ser | VUS | 4.95 | 5.38 | 4117 | 7732 |
| 127 | p.Ala127Ile | VUS | 4.81 | 4.92 | 4000 | 7070 |
| 127 | p.Ala127Met | VUS | 5.79 | 6.01 | 4818 | 8638 |
| 127 | p.Ala127His | VUS | 4.24 | 4.31 | 3529 | 6189 |
| 127 | p.Ala127Gln | VUS | 5.09 | 5.50 | 4233 | 7900 |
| 127 | p.Ala127Pro | VUS | 2.84 | 2.63 | 2364 | 3778 |
| 127 | p.Ala127Leu | VUS | 5.06 | 4.74 | 4209 | 6805 |
| 127 | p.Ala127Asp | VUS | 6.04 | 6.53 | 5023 | 9386 |
| 127 | p.Ala127Glu | VUS | 6.07 | 5.60 | 5052 | 8045 |
| 127 | p.Ala127Ala | Synonymous | 4.27 | 3.68 | 3548 | 5292 |
| 127 | p.Ala127Gly | VUS | 4.75 | 4.74 | 3949 | 6809 |
| 127 | p.Ala127Val | VUS | 6.33 | 6.83 | 5268 | 9821 |
| 127 | p.Ala127Tyr | VUS | 4.33 | 3.86 | 3599 | 5544 |
| 127 | p.Ala127Cys | VUS | 4.44 | 4.93 | 3696 | 7087 |
| 127 | p.Ala127Trp | VUS | 5.97 | 5.92 | 4962 | 8513 |
| 127 | p.Ala127Phe | VUS | 5.73 | 5.92 | 4767 | 8512 |
