## Appendix 1-table 4 for "Functional characterization of all *CDKN2A* missense variants and comparison to in silico models of pathogenicity"

**Appendix 1-table 4. Assay outputs and functional classifications for all possible *CDKN2A* missense and synonymous variants.**

| Residue | Variant | Benchmark | Functionally reported VUS | Experiment_1 Read count | | Experiment_1 Proportion | | Experiment_2 Read count | | Experiment_2 Proportion | | Experiment_1 | | Experiment_2 | | Merged | |
|---|---|---|---|---|---|---|---|---|---|---|---|---|---|---|---|---|---|
| | | | | Day 9_1 | Confluent_1 | Day 9_1 | Confluent_1 | Day 9_2 | Confluent_2 | Day 9_2 | Confluent_2 | Log P value_1 | Fucntional characterization_1 | Log P value_2 | Fucntional characterization_2 | Log P value_merged | Fucntional characterization _merged |
| 1 | p.Met1Asn | | | 7917 | 3240 | 4.45 | 4.27 | | | | | -1.53E-01 | Neutral | | | -1.53E-01 | Neutral |
| 1 | p.Met1Lys | | | 10656 | 3866 | 5.99 | 5.10 | | | | | -8.17E-03 | Neutral | | | -8.17E-03 | Neutral |
| 1 | p.Met1Thr | | | 8682 | 3625 | 4.88 | 4.78 | | | | | -1.36E-01 | Neutral | | | -1.36E-01 | Neutral |
| 1 | p.Met1Arg | | | 4562 | 1607 | 2.56 | 2.12 | | | | | -2.18E-01 | Neutral | | | -2.18E-01 | Neutral |
| 1 | p.Met1Ser | | | 14850 | 6066 | 8.35 | 8.00 | | | | | -9.32E-03 | Neutral | | | -9.32E-03 | Neutral |
| 1 | p.Met1Ile | | | 5660 | 2304 | 3.18 | 3.04 | | | | | -4.38E-01 | Neutral | | | -4.38E-01 | Neutral |
| 1 | p.Met1Met | Synonymous | | 5258 | 2332 | 2.96 | 3.08 | | | | | -1.06E+00 | Neutral | | | -1.06E+00 | Neutral |
| 1 | p.Met1His | | | 5078 | 2285 | 2.85 | 3.01 | | | | | -1.27E+00 | Neutral | | | -1.27E+00 | Neutral |
| 1 | p.Met1Gln | | | 9585 | 4102 | 5.39 | 5.41 | | | | | -1.24E-01 | Neutral | | | -1.24E-01 | Neutral |
| 1 | p.Met1Pro | | | 5141 | 1985 | 2.89 | 2.62 | | | | | -3.62E-01 | Neutral | | | -3.62E-01 | Neutral |
| 1 | p.Met1Leu | | | 14664 | 6252 | 8.24 | 8.25 | | | | | -1.90E-02 | Neutral | | | -1.90E-02 | Neutral |
| 1 | p.Met1Asp | | | 14894 | 5681 | 8.37 | 7.49 | | | | | -2.98E-03 | Neutral | | | -2.98E-03 | Neutral |
| 1 | p.Met1Glu | | | 5078 | 2195 | 2.85 | 2.89 | | | | | -9.57E-01 | Neutral | | | -9.57E-01 | Neutral |
| 1 | p.Met1Ala | | | 8804 | 3502 | 4.95 | 4.62 | | | | | -7.30E-02 | Neutral | | | -7.30E-02 | Neutral |
| 1 | p.Met1Gly | | | 6523 | 2720 | 3.67 | 3.59 | | | | | -3.55E-01 | Neutral | | | -3.55E-01 | Neutral |
| 1 | p.Met1Val | | | 11052 | 7911 | 6.21 | 10.43 | | | | | -4.16E+00 | Neutral | | | -4.16E+00 | Neutral |
| 1 | p.Met1Tyr | | | 12992 | 5282 | 7.30 | 6.97 | | | | | -1.70E-02 | Neutral | | | -1.70E-02 | Neutral |
| 1 | p.Met1Cys | | | 14675 | 5858 | 8.25 | 7.73 | | | | | -6.89E-03 | Neutral | | | -6.89E-03 | Neutral |
| 1 | p.Met1Trp | | | 3926 | 1599 | 2.21 | 2.11 | | | | | -1.15E+00 | Neutral | | | -1.15E+00 | Neutral |
| 1 | p.Met1Phe | | | 7910 | 3411 | 4.45 | 4.50 | | | | | -2.65E-01 | Neutral | | | -2.65E-01 | Neutral |
| 2 | p.Glu2Asn | | | 6727 | 2813 | 4.99 | 2.50 | | | | | -2.75E-14 | Neutral | | | 0.00E+00 | Neutral |
| 2 | p.Glu2Lys | | | 6744 | 5413 | 5.01 | 4.81 | | | | | -2.54E-06 | Neutral | | | -2.54E-06 | Neutral |
| 2 | p.Glu2Thr | | | 5212 | 4425 | 3.87 | 3.93 | | | | | -6.21E-05 | Neutral | | | -6.21E-05 | Neutral |
| 2 | p.Glu2Arg | | | 5222 | 2299 | 3.88 | 2.04 | | | | | -1.88E-12 | Neutral | | | 0.00E+00 | Neutral |
| 2 | p.Glu2Ser | | | 4630 | 6041 | 3.44 | 5.36 | | | | | -1.34E-01 | Neutral | | | -1.34E-01 | Neutral |
| 2 | p.Glu2Ile | | | 7541 | 3779 | 5.60 | 3.36 | | | | | -2.35E-12 | Neutral | | | 0.00E+00 | Neutral |
| 2 | p.Glu2Met | | | 6300 | 7369 | 4.68 | 6.54 | | | | | -7.63E-03 | Neutral | | | -7.63E-03 | Neutral |
| 2 | p.Glu2His | | | 7815 | 6349 | 5.80 | 5.64 | | | | | -1.05E-06 | Neutral | | | -1.05E-06 | Neutral |
| 2 | p.Glu2Gln | | | 9277 | 5087 | 6.89 | 4.52 | | | | | -4.48E-12 | Neutral | | | 0.00E+00 | Neutral |
| 2 | p.Glu2Pro | | | 8059 | 5092 | 5.98 | 4.52 | | | | | -1.09E-09 | Neutral | | | -9.43E-10 | Neutral |
| 2 | p.Glu2Leu | | | 7335 | 3699 | 5.44 | 3.28 | | | | | -3.77E-12 | Neutral | | | 0.00E+00 | Neutral |
| 2 | p.Glu2Asp | | | 4954 | 1515 | 3.68 | 1.35 | | | | | 0.00E+00 | Neutral | | | 0.00E+00 | Neutral |
| 2 | p.Glu2Glu | Synonymous | | 9092 | 18647 | 6.75 | 16.56 | | | | | -1.06E+00 | Neutral | | | -1.06E+00 | Neutral |
| 2 | p.Glu2Ala | | | 6504 | 8164 | 4.83 | 7.25 | | | | | -1.91E-02 | Neutral | | | -1.91E-02 | Neutral |
| 2 | p.Glu2Gly | | | 5629 | 4118 | 4.18 | 3.66 | | | | | -1.16E-06 | Neutral | | | -1.16E-06 | Neutral |
| 2 | p.Glu2Val | | | 6747 | 3223 | 5.01 | 2.86 | | | | | -1.69E-12 | Neutral | | | 0.00E+00 | Neutral |
| 2 | p.Glu2Tyr | | | 4650 | 2961 | 3.45 | 2.63 | | | | | -1.78E-07 | Neutral | | | -1.78E-07 | Neutral |
| 2 | p.Glu2Cys | | | 6088 | 7985 | 4.52 | 7.09 | | | | | -4.74E-02 | Neutral | | | -4.74E-02 | Neutral |
| 2 | p.Glu2Trp | | | 7642 | 6902 | 5.67 | 6.13 | | | | | -1.49E-05 | Neutral | | | -1.49E-05 | Neutral |
| 2 | p.Glu2Phe | | | 8566 | 6750 | 6.36 | 5.99 | | | | | -2.31E-07 | Neutral | | | -2.31E-07 | Neutral |
| 3 | p.Pro3Asn | | | 10479 | 4032 | 5.98 | 5.43 | | | | | -7.64E+00 | Indeterminate | | | -7.64E+00 | Indeterminate |
| 3 | p.Pro3Lys | | | 6030 | 2500 | 3.44 | 3.37 | | | | | -1.84E+01 | Indeterminate | | | -1.84E+01 | Indeterminate |
| 3 | p.Pro3Thr | | | 8211 | 3197 | 4.69 | 4.31 | | | | | -1.08E+01 | Indeterminate | | | -1.08E+01 | Indeterminate |
| 3 | p.Pro3Arg | | | 8154 | 5051 | 4.66 | 6.81 | | | | | -4.02E+01 | Indeterminate | | | -3.32E+01 | Indeterminate |
| 3 | p.Pro3Ser | | | 7465 | 3568 | 4.26 | 4.81 | | | | | -2.25E+01 | Indeterminate | | | -2.25E+01 | Indeterminate |
| 3 | p.Pro3Ile | | | 6870 | 2552 | 3.92 | 3.44 | | | | | -1.13E+01 | Indeterminate | | | -1.13E+01 | Indeterminate |
| 3 | p.Pro3Met | | | 6862 | 3240 | 3.92 | 4.37 | | | | | -2.35E+01 | Indeterminate | | | -2.35E+01 | Indeterminate |
| 3 | p.Pro3His | | | 6669 | 2007 | 3.81 | 2.71 | | | | | -5.11E+00 | Neutral | | | -5.11E+00 | Neutral |
| 3 | p.Pro3Gln | | | 7887 | 1624 | 4.50 | 2.19 | | | | | -2.77E-01 | Neutral | | | -2.77E-01 | Neutral |
| 3 | p.Pro3Pro | Synonymous | | 12225 | 3414 | 6.98 | 4.60 | | | | | -1.06E+00 | Neutral | | | -1.06E+00 | Neutral |
| 3 | p.Pro3Leu | | | 7911 | 3359 | 4.52 | 4.53 | | | | | -1.50E+01 | Indeterminate | | | -1.50E+01 | Indeterminate |
| 3 | p.Pro3Asp | | | 11034 | 5168 | 6.30 | 6.97 | | | | | -1.44E+01 | Indeterminate | | | -1.44E+01 | Indeterminate |
| 3 | p.Pro3Glu | | | 8543 | 2741 | 4.88 | 3.69 | | | | | -4.70E+00 | Neutral | | | -4.70E+00 | Neutral |
| 3 | p.Pro3Ala | | | 7836 | 2325 | 4.47 | 3.13 | | | | | -3.70E+00 | Neutral | | | -3.70E+00 | Neutral |
| 3 | p.Pro3Gly | | | 9287 | 7121 | 5.30 | 9.60 | | | | | -5.32E+01 | Deleterious | | | -5.32E+01 | Deleterious |
| 3 | p.Pro3Val | | | 11755 | 5610 | 6.71 | 7.56 | | | | | -1.43E+01 | Indeterminate | | | -1.43E+01 | Indeterminate |
| 3 | p.Pro3Tyr | | | 8286 | 2722 | 4.73 | 3.67 | | | | | -5.47E+00 | Neutral | | | -5.47E+00 | Neutral |
| 3 | p.Pro3Cys | | | 9700 | 4045 | 5.54 | 5.45 | | | | | -1.13E+01 | Indeterminate | | | -1.13E+01 | Indeterminate |
| 3 | p.Pro3Trp | | | 7181 | 3898 | 4.10 | 5.25 | | | | | -3.25E+01 | Indeterminate | | | -3.18E+01 | Indeterminate |
| 3 | p.Pro3Phe | | | 12773 | 6021 | 7.29 | 8.12 | | | | | -1.26E+01 | Indeterminate | | | -1.26E+01 | Indeterminate |
| 4 | p.Ala4Asn | | | 6496 | 5119 | 5.55 | 5.97 | | | | | -5.71E+00 | Neutral | | | -5.71E+00 | Neutral |
| 4 | p.Ala4Lys | | | 5599 | 3201 | 4.79 | 3.73 | | | | | -1.31E+00 | Neutral | | | -1.31E+00 | Neutral |
| 4 | p.Ala4Thr | | | 6049 | 9770 | 5.17 | 11.39 | | | | | -5.14E+01 | Indeterminate | | | -3.32E+01 | Indeterminate |
| 4 | p.Ala4Arg | | | 6946 | 4316 | 5.94 | 5.03 | | | | | -1.41E+00 | Neutral | | | -1.41E+00 | Neutral |
| 4 | p.Ala4Ser | | | 5204 | 8721 | 4.45 | 10.16 | | | | | -5.32E+01 | Deleterious | | | -5.32E+01 | Deleterious |
| 4 | p.Ala4Ile | | | 5102 | 2244 | 4.36 | 2.62 | | | | | -1.79E-01 | Neutral | | | -1.79E-01 | Neutral |
| 4 | p.Ala4Met | | | 6306 | 5309 | 5.39 | 6.19 | | | | | -7.81E+00 | Indeterminate | | | -7.81E+00 | Indeterminate |
| 4 | p.Ala4His | | | 9180 | 6186 | 7.85 | 7.21 | | | | | -1.31E+00 | Neutral | | | -1.31E+00 | Neutral |
| 4 | p.Ala4Gln | | | 4555 | 2044 | 3.89 | 2.38 | | | | | -3.21E-01 | Neutral | | | -3.21E-01 | Neutral |
| 4 | p.Ala4Pro | | | 4683 | 2290 | 4.00 | 2.67 | | | | | -6.33E-01 | Neutral | | | -6.33E-01 | Neutral |
| 4 | p.Ala4Leu | | | 6006 | 3343 | 5.13 | 3.90 | | | | | -9.14E-01 | Neutral | | | -9.14E-01 | Neutral |
| 4 | p.Ala4Asp | | | 4923 | 3133 | 4.21 | 3.65 | | | | | -3.20E+00 | Neutral | | | -3.20E+00 | Neutral |
| 4 | p.Ala4Glu | | | 5399 | 2024 | 4.62 | 2.36 | | | | | -1.89E-02 | Neutral | | | -1.89E-02 | Neutral |
| 4 | p.Ala4Ala | Synonymous | | 6423 | 3731 | 5.49 | 4.35 | | | | | -1.06E+00 | Neutral | | | -1.06E+00 | Neutral |
| 4 | p.Ala4Gly | | | 5628 | 4372 | 4.81 | 5.10 | | | | | -6.58E+00 | Indeterminate | | | -6.58E+00 | Indeterminate |
| 4 | p.Ala4Val | | | 6269 | 2830 | 5.36 | 3.30 | | | | | -1.12E-01 | Neutral | | | -1.12E-01 | Neutral |
| 4 | p.Ala4Tyr | | | 5640 | 2955 | 4.82 | 3.44 | | | | | -6.72E-01 | Neutral | | | -6.72E-01 | Neutral |
| 4 | p.Ala4Cys | | | 4485 | 5419 | 3.83 | 6.32 | | | | | -3.33E+01 | Indeterminate | | | -3.23E+01 | Indeterminate |
| 4 | p.Ala4Trp | | | 6700 | 4351 | 5.73 | 5.07 | | | | | -2.02E+00 | Neutral | | | -2.02E+00 | Neutral |
| 4 | p.Ala4Phe | | | 5374 | 4447 | 4.59 | 5.18 | | | | | -8.96E+00 | Indeterminate | | | -8.96E+00 | Indeterminate |
| 5 | p.Ala5Asn | | | 10398 | 3217 | 7.08 | 4.47 | | | | | -1.02E-02 | Neutral | | | -1.02E-02 | Neutral |
| 5 | p.Ala5Lys | | | 6616 | 943 | 4.50 | 1.31 | | | | | -1.32E-08 | Neutral | | | -1.31E-08 | Neutral |
| 5 | p.Ala5Thr | | | 7235 | 2897 | 4.92 | 4.02 | | | | | -7.68E-01 | Neutral | | | -7.68E-01 | Neutral |
| 5 | p.Ala5Arg | | | 6763 | 1495 | 4.60 | 2.08 | | | | | -3.35E-04 | Neutral | | | -3.35E-04 | Neutral |
| 5 | p.Ala5Ser | | | 6901 | 7643 | 4.70 | 10.62 | | | | | -4.59E+01 | Indeterminate | | | -3.32E+01 | Indeterminate |
| 5 | p.Ala5Ile | | | 9416 | 1133 | 6.41 | 1.57 | | | | | -3.62E-12 | Neutral | | | 0.00E+00 | Neutral |
| 5 | p.Ala5Met | | | 6059 | 2031 | 4.12 | 2.82 | | | | | -2.70E-01 | Neutral | | | -2.70E-01 | Neutral |
| 5 | p.Ala5His | | | 9217 | 3925 | 6.27 | 5.45 | | | | | -6.61E-01 | Neutral | | | -6.61E-01 | Neutral |
| 5 | p.Ala5Gln | | | 8603 | 3685 | 5.86 | 5.12 | | | | | -8.29E-01 | Neutral | | | -8.29E-01 | Neutral |
| 5 | p.Ala5Pro | | | 8547 | 2852 | 5.82 | 3.96 | | | | | -7.23E-02 | Neutral | | | -7.23E-02 | Neutral |
| 5 | p.Ala5Leu | | | 10779 | 7833 | 7.34 | 10.88 | | | | | -9.21E+00 | Indeterminate | | | -9.21E+00 | Indeterminate |
| 5 | p.Ala5Asp | | | 7000 | 3082 | 4.76 | 4.28 | | | | | -1.63E+00 | Neutral | | | -1.63E+00 | Neutral |
| 5 | p.Ala5Glu | | | 5477 | 2907 | 3.73 | 4.04 | | | | | -6.48E+00 | Indeterminate | | | -6.48E+00 | Indeterminate |
| 5 | p.Ala5Ala | Synonymous | | 5728 | 2216 | 3.90 | 3.08 | | | | | -1.06E+00 | Neutral | | | -1.06E+00 | Neutral |
| 5 | p.Ala5Gly | | | 4494 | 1339 | 3.06 | 1.86 | | | | | -2.24E-01 | Neutral | | | -2.24E-01 | Neutral |
| 5 | p.Ala5Val | | | 8831 | 3044 | 6.01 | 4.23 | | | | | -9.39E-02 | Neutral | | | -9.39E-02 | Neutral |
| 5 | p.Ala5Tyr | | | 4625 | 8628 | 3.15 | 11.99 | | | | | -5.32E+01 | Deleterious | | | -5.32E+01 | Deleterious |
| 5 | p.Ala5Cys | | | 5753 | 2808 | 3.92 | 3.90 | | | | | -4.14E+00 | Neutral | | | -4.14E+00 | Neutral |
| 5 | p.Ala5Trp | | | 8851 | 8353 | 6.02 | 11.60 | | | | | -2.55E+01 | Indeterminate | | | -2.55E+01 | Indeterminate |
| 5 | p.Ala5Phe | | | 5632 | 1954 | 3.83 | 2.71 | | | | | -4.66E-01 | Neutral | | | -4.66E-01 | Neutral |
| 6 | p.Gly6Asn | | | 5526 | 2196 | 5.53 | 2.96 | | | | | -9.29E-02 | Neutral | | | -9.29E-02 | Neutral |
| 6 | p.Gly6Lys | | | 5673 | 3312 | 5.68 | 4.46 | | | | | -2.25E+00 | Neutral | | | -2.25E+00 | Neutral |
| 6 | p.Gly6Thr | | | 4626 | 2843 | 4.63 | 3.83 | | | | | -4.22E+00 | Neutral | | | -4.22E+00 | Neutral |
| 6 | p.Gly6Arg | | | 4400 | 8247 | 4.40 | 11.12 | | | | | -5.32E+01 | Deleterious | | | -5.32E+01 | Deleterious |
| 6 | p.Gly6Ser | | | 5199 | 7507 | 5.20 | 10.12 | | | | | -5.20E+01 | Indeterminate | | | -3.32E+01 | Indeterminate |
| 6 | p.Gly6Ile | | | 4539 | 7126 | 4.54 | 9.61 | | | | | -5.32E+01 | Deleterious | | | -5.32E+01 | Deleterious |
| 6 | p.Gly6Met | | | 7887 | 11562 | 7.89 | 15.59 | | | | | -3.98E+01 | Indeterminate | | | -3.32E+01 | Indeterminate |
| 6 | p.Gly6His | | | 4522 | 1360 | 4.53 | 1.83 | | | | | -4.50E-03 | Neutral | | | -4.50E-03 | Neutral |
| 6 | p.Gly6Gln | | | 4363 | 1586 | 4.37 | 2.14 | | | | | -8.17E-02 | Neutral | | | -8.17E-02 | Neutral |
| 6 | p.Gly6Pro | | | 3564 | 3083 | 3.57 | 4.16 | | | | | -2.03E+01 | Indeterminate | | | -2.03E+01 | Indeterminate |
| 6 | p.Gly6Leu | | | 6157 | 3203 | 6.16 | 4.32 | | | | | -8.72E-01 | Neutral | | | -8.72E-01 | Neutral |
| 6 | p.Gly6Asp | | | 4366 | 1905 | 4.37 | 2.57 | | | | | -5.31E-01 | Neutral | | | -5.31E-01 | Neutral |
| 6 | p.Gly6Glu | | | 5386 | 2616 | 5.39 | 3.53 | | | | | -7.22E-01 | Neutral | | | -7.22E-01 | Neutral |
| 6 | p.Gly6Ala | | | 6037 | 2648 | 6.04 | 3.57 | | | | | -2.01E-01 | Neutral | | | -2.01E-01 | Neutral |
| 6 | p.Gly6Gly | Synonymous | | 2942 | 1230 | 2.94 | 1.66 | | | | | -1.06E+00 | Neutral | | | -1.06E+00 | Neutral |
| 6 | p.Gly6Val | | | 7551 | 3158 | 7.56 | 4.26 | | | | | -4.82E-02 | Neutral | | | -4.82E-02 | Neutral |
| 6 | p.Gly6Tyr | | | 3436 | 1449 | 3.44 | 1.95 | | | | | -7.69E-01 | Neutral | | | -7.69E-01 | Neutral |
| 6 | p.Gly6Cys | | | 3736 | 2856 | 3.74 | 3.85 | | | | | -1.32E+01 | Indeterminate | | | -1.32E+01 | Indeterminate |
| 6 | p.Gly6Trp | | | 5083 | 3058 | 5.09 | 4.12 | | | | | -3.24E+00 | Neutral | | | -3.24E+00 | Neutral |
| 6 | p.Gly6Phe | | | 4916 | 3241 | 4.92 | 4.37 | | | | | -5.31E+00 | Neutral | | | -5.31E+00 | Neutral |
| 7 | p.Ser7Asn | | | 21503 | 2770 | 5.64 | 3.85 | | | | | -7.01E-02 | Neutral | | | -7.01E-02 | Neutral |
| 7 | p.Ser7Lys | | | 17957 | 2366 | 4.71 | 3.28 | | | | | -1.81E-01 | Neutral | | | -1.81E-01 | Neutral |
| 7 | p.Ser7Thr | | | 26503 | 7810 | 6.95 | 10.84 | | | | | -1.12E+01 | Indeterminate | | | -1.12E+01 | Indeterminate |
| 7 | p.Ser7Arg | | | 20121 | 2384 | 5.28 | 3.31 | | | | | -3.13E-02 | Neutral | | | -3.13E-02 | Neutral |
| 7 | p.Ser7Ser | Synonymous | | 13032 | 1887 | 3.42 | 2.62 | | | | | -1.06E+00 | Neutral | | | -1.06E+00 | Neutral |
| 7 | p.Ser7Ile | | | 21968 | 3908 | 5.76 | 5.43 | | | | | -1.33E+00 | Neutral | | | -1.33E+00 | Neutral |
| 7 | p.Ser7Met | | | 25433 | 4362 | 6.67 | 6.06 | | | | | -7.12E-01 | Neutral | | | -7.12E-01 | Neutral |
| 7 | p.Ser7His | | | 23901 | 5103 | 6.27 | 7.08 | | | | | -3.32E+00 | Neutral | | | -3.32E+00 | Neutral |
| 7 | p.Ser7Gln | | | 25626 | 4097 | 6.72 | 5.69 | | | | | -3.82E-01 | Neutral | | | -3.82E-01 | Neutral |
| 7 | p.Ser7Pro | | | 30865 | 5648 | 8.09 | 7.84 | | | | | -7.15E-01 | Neutral | | | -7.15E-01 | Neutral |
| 7 | p.Ser7Leu | | | 22752 | 5920 | 5.97 | 8.22 | | | | | -8.57E+00 | Indeterminate | | | -8.57E+00 | Indeterminate |
| 7 | p.Ser7Asp | | | 13045 | 1864 | 3.42 | 2.59 | | | | | -9.61E-01 | Neutral | | | -9.61E-01 | Neutral |

| | | | | | | | | | | | |
|---|---|---|---|---|---|---|---|---|---|---|---|
| 7 | p.Ser7Glu | | | 5274 | 1263 | 1.38 | 1.75 | -2.95E+01 | Indeterminate | -2.94E+01 | Indeterminate |
| 7 | p.Ser7Ala | | | 13797 | 3900 | 3.62 | 5.41 | -1.95E+01 | Indeterminate | -1.95E+01 | Indeterminate |
| 7 | p.Ser7Gly | | | 34546 | 4664 | 9.06 | 6.48 | -1.55E-02 | Neutral | -1.55E-02 | Neutral |
| 7 | p.Ser7Val | | | 6451 | 937 | 1.69 | 1.30 | -4.25E+00 | Neutral | -4.25E+00 | Neutral |
| 7 | p.Ser7Tyr | | | 14033 | 5213 | 3.68 | 7.24 | -3.89E+01 | Indeterminate | -3.32E+01 | Indeterminate |
| 7 | p.Ser7Cys | | | 15237 | 2771 | 4.00 | 3.85 | -3.14E+00 | Neutral | -3.14E+00 | Neutral |
| 7 | p.Ser7Trp | | | 10664 | 2020 | 2.80 | 2.80 | -6.60E+00 | Indeterminate | -6.60E+00 | Indeterminate |
| 7 | p.Ser7Phe | | | 18617 | 3141 | 4.88 | 4.36 | -1.35E+00 | Neutral | -1.35E+00 | Neutral |
| 8 | p.Ser8Asn | | | 5102 | 2754 | 5.51 | 4.26 | -1.03E-03 | Neutral | -1.03E-03 | Neutral |
| 8 | p.Ser8Lys | | | 3518 | 5765 | 3.80 | 8.91 | -2.11E+01 | Indeterminate | -2.11E+01 | Indeterminate |
| 8 | p.Ser8Thr | | | 5291 | 4626 | 5.72 | 7.15 | -5.41E-01 | Neutral | -5.41E-01 | Neutral |
| 8 | p.Ser8Arg | | | 4648 | 3049 | 5.02 | 4.71 | -3.81E-02 | Neutral | -3.81E-02 | Neutral |
| 8 | p.Ser8Ser | Synonymous | | 5221 | 4953 | 5.64 | 7.66 | -1.06E+00 | Neutral | -1.06E+00 | Neutral |
| 8 | p.Ser8Ile | | | 6586 | 4481 | 7.11 | 6.93 | -1.22E-02 | Neutral | -1.22E-02 | Neutral |
| 8 | p.Ser8Met | | | 3576 | 1885 | 3.86 | 2.91 | -5.33E-03 | Neutral | -5.33E-03 | Neutral |
| 8 | p.Ser8His | | | 4006 | 2293 | 4.33 | 3.54 | -1.09E-02 | Neutral | -1.09E-02 | Neutral |
| 8 | p.Ser8Gln | | | 5026 | 3177 | 5.43 | 4.91 | -1.56E-02 | Neutral | -1.56E-02 | Neutral |
| 8 | p.Ser8Pro | | | 3368 | 2661 | 3.64 | 4.11 | -8.03E-01 | Neutral | -8.03E-01 | Neutral |
| 8 | p.Ser8Leu | | | 4268 | 3385 | 4.61 | 5.23 | -4.31E-01 | Neutral | -4.31E-01 | Neutral |
| 8 | p.Ser8Asp | | | 5645 | 3906 | 6.10 | 6.04 | -3.28E-02 | Neutral | -3.28E-02 | Neutral |
| 8 | p.Ser8Glu | | | 5235 | 2231 | 5.65 | 3.45 | -6.17E-06 | Neutral | -6.17E-06 | Neutral |
| 8 | p.Ser8Ala | | | 3572 | 2303 | 3.86 | 3.56 | -9.28E-02 | Neutral | -9.28E-02 | Neutral |
| 8 | p.Ser8Gly | | | 4071 | 3170 | 4.40 | 4.90 | -4.21E-01 | Neutral | -4.21E-01 | Neutral |
| 8 | p.Ser8Val | | | 5367 | 3622 | 5.80 | 5.60 | -2.92E-02 | Neutral | -2.92E-02 | Neutral |
| 8 | p.Ser8Tyr | | | 4234 | 2444 | 4.57 | 3.78 | -9.30E-03 | Neutral | -9.30E-03 | Neutral |
| 8 | p.Ser8Cys | | | 4875 | 2414 | 5.27 | 3.73 | -2.62E-04 | Neutral | -2.62E-04 | Neutral |
| 8 | p.Ser8Trp | | | 5137 | 2823 | 5.55 | 4.36 | -1.37E-03 | Neutral | -1.37E-03 | Neutral |
| 8 | p.Ser8Phe | | | 3833 | 2741 | 4.14 | 4.24 | -2.25E-01 | Neutral | -2.25E-01 | Neutral |
| 9 | p.Met9Asn | | | 7469 | 3933 | 4.84 | 4.81 | -2.11E-01 | Neutral | -2.11E-01 | Neutral |
| 9 | p.Met9Lys | | | 9609 | 5540 | 6.23 | 6.77 | -2.27E-01 | Neutral | -2.27E-01 | Neutral |
| 9 | p.Met9Thr | | Neutral | 8791 | 5010 | 5.70 | 6.12 | -2.72E-01 | Neutral | -2.72E-01 | Neutral |
| 9 | p.Met9Arg | | | 10762 | 5831 | 6.97 | 7.12 | -7.40E-02 | Neutral | -7.40E-02 | Neutral |
| 9 | p.Met9Ser | | | 5046 | 2796 | 3.27 | 3.42 | -1.02E+00 | Neutral | -1.02E+00 | Neutral |
| 9 | p.Met9Ile | | | 8383 | 4731 | 5.43 | 5.78 | -2.88E-01 | Neutral | -2.88E-01 | Neutral |
| 9 | p.Met9Met | Synonymous | | 3169 | 1512 | 2.05 | 1.85 | -1.06E+00 | Neutral | -1.06E+00 | Neutral |
| 9 | p.Met9His | | | 4318 | 2163 | 2.80 | 2.64 | -7.05E-01 | Neutral | -7.05E-01 | Neutral |
| 9 | p.Met9Gln | | | 5190 | 2565 | 3.36 | 3.13 | -3.71E-01 | Neutral | -3.71E-01 | Neutral |
| 9 | p.Met9Pro | | | 6382 | 3588 | 4.14 | 4.38 | -6.27E-01 | Neutral | -6.27E-01 | Neutral |
| 9 | p.Met9Leu | | | 8726 | 3851 | 5.66 | 4.71 | -1.14E-02 | Neutral | -1.14E-02 | Neutral |
| 9 | p.Met9Asp | | | 9850 | 5446 | 6.38 | 6.65 | -1.33E-01 | Neutral | -1.33E-01 | Neutral |
| 9 | p.Met9Glu | | | 8636 | 4204 | 5.60 | 5.14 | -4.81E-02 | Neutral | -4.81E-02 | Neutral |
| 9 | p.Met9Ala | | | 8064 | 4131 | 5.23 | 5.05 | -1.19E-01 | Neutral | -1.19E-01 | Neutral |
| 9 | p.Met9Gly | | | 6394 | 3368 | 4.14 | 4.12 | -3.52E-01 | Neutral | -3.52E-01 | Neutral |
| 9 | p.Met9Val | | | 8528 | 5087 | 5.53 | 6.22 | -4.58E-01 | Neutral | -4.58E-01 | Neutral |
| 9 | p.Met9Tyr | | | 9867 | 4712 | 6.39 | 5.76 | -2.01E-02 | Neutral | -2.01E-02 | Neutral |
| 9 | p.Met9Cys | | | 8011 | 4120 | 5.19 | 5.03 | -1.27E-01 | Neutral | -1.27E-01 | Neutral |
| 9 | p.Met9Trp | | | 9040 | 4855 | 5.86 | 5.93 | -1.32E-01 | Neutral | -1.32E-01 | Neutral |
| 9 | p.Met9Phe | | | 8063 | 4400 | 5.23 | 5.38 | -2.35E-01 | Neutral | -2.35E-01 | Neutral |
| 10 | p.Glu10Asn | | | 6098 | 2242 | 4.21 | 3.54 | -2.65E+00 | Neutral | -2.65E+00 | Neutral |
| 10 | p.Glu10Lys | | | 5462 | 4301 | 3.77 | 6.80 | -4.13E+01 | Indeterminate | -3.32E+01 | Indeterminate |
| 10 | p.Glu10Thr | | | 5428 | 2183 | 3.75 | 3.45 | -5.04E+00 | Neutral | -5.04E+00 | Neutral |
| 10 | p.Glu10Arg | | | 6859 | 2202 | 4.74 | 3.48 | -8.52E-01 | Neutral | -8.52E-01 | Neutral |
| 10 | p.Glu10Ser | | | 4238 | 2388 | 2.93 | 3.77 | -2.20E+01 | Indeterminate | -2.20E+01 | Indeterminate |
| 10 | p.Glu10Ile | | | 5736 | 3004 | 3.96 | 4.75 | -1.30E+01 | Indeterminate | -1.30E+01 | Indeterminate |
| 10 | p.Glu10Met | | | 10083 | 5755 | 6.96 | 9.09 | -9.12E+00 | Indeterminate | -9.12E+00 | Indeterminate |
| 10 | p.Glu10His | | | 7085 | 5545 | 4.89 | 8.76 | -3.32E+01 | Indeterminate | -3.22E+01 | Indeterminate |
| 10 | p.Glu10Gln | | | 8938 | 2836 | 6.17 | 4.48 | -3.68E-01 | Neutral | -3.68E-01 | Neutral |
| 10 | p.Glu10Pro | | | 9004 | 3151 | 6.22 | 4.98 | -8.25E-01 | Neutral | -8.25E-01 | Neutral |
| 10 | p.Glu10Leu | | | 4693 | 1833 | 3.24 | 2.90 | -5.49E+00 | Neutral | -5.49E+00 | Neutral |
| 10 | p.Glu10Asp | | | 7860 | 3023 | 5.43 | 4.78 | -2.12E+00 | Neutral | -2.12E+00 | Neutral |
| 10 | p.Glu10Glu | Synonymous | | 7526 | 2567 | 5.20 | 4.06 | -1.06E+00 | Neutral | -1.06E+00 | Neutral |
| 10 | p.Glu10Ala | | | 3045 | 1836 | 2.10 | 2.90 | -3.48E+01 | Indeterminate | -3.28E+01 | Indeterminate |
| 10 | p.Glu10Gly | | | 7598 | 2700 | 5.25 | 4.27 | -1.38E+00 | Neutral | -1.38E+00 | Neutral |
| 10 | p.Glu10Val | | | 8257 | 2931 | 5.70 | 4.63 | -1.13E+00 | Neutral | -1.13E+00 | Neutral |
| 10 | p.Glu10Tyr | | | 11783 | 5357 | 8.14 | 8.47 | -2.56E+00 | Neutral | -2.56E+00 | Neutral |
| 10 | p.Glu10Cys | | | 11223 | 4410 | 7.75 | 6.97 | -1.13E+00 | Neutral | -1.13E+00 | Neutral |
| 10 | p.Glu10Trp | | | 7320 | 2533 | 5.05 | 4.00 | -1.25E+00 | Neutral | -1.25E+00 | Neutral |
| 10 | p.Glu10Phe | | | 6589 | 2486 | 4.55 | 3.93 | -2.65E+00 | Neutral | -2.65E+00 | Neutral |
| 11 | p.Pro11Asn | | | 8506 | 1300 | 5.16 | 5.15 | -1.32E+00 | Neutral | -1.32E+00 | Neutral |
| 11 | p.Pro11Lys | | | 6439 | 1122 | 3.91 | 4.44 | -4.63E+00 | Neutral | -4.63E+00 | Neutral |
| 11 | p.Pro11Thr | | | 5725 | 1120 | 3.47 | 4.43 | -8.84E+00 | Indeterminate | -8.84E+00 | Indeterminate |
| 11 | p.Pro11Arg | | | 6357 | 933 | 3.86 | 3.69 | -1.90E+00 | Neutral | -1.90E+00 | Neutral |
| 11 | p.Pro11Ser | | | 8111 | 1330 | 4.92 | 5.27 | -2.27E+00 | Neutral | -2.27E+00 | Neutral |
| 11 | p.Pro11Ile | | | 8262 | 1268 | 5.01 | 5.02 | -1.45E+00 | Neutral | -1.45E+00 | Neutral |
| 11 | p.Pro11Met | | | 12247 | 1773 | 7.43 | 7.02 | -3.21E-01 | Neutral | -3.21E-01 | Neutral |
| 11 | p.Pro11His | | | 7706 | 1174 | 4.68 | 4.65 | -1.60E+00 | Neutral | -1.60E+00 | Neutral |
| 11 | p.Pro11Gln | | | 8805 | 1147 | 5.34 | 4.54 | -3.33E-01 | Neutral | -3.33E-01 | Neutral |
| 11 | p.Pro11Pro | Synonymous | | 6518 | 884 | 3.95 | 3.50 | -1.06E+00 | Neutral | -1.06E+00 | Neutral |
| 11 | p.Pro11Leu | | Neutral | 7222 | 1095 | 4.38 | 4.34 | -1.78E+00 | Neutral | -1.78E+00 | Neutral |
| 11 | p.Pro11Asp | | | 10405 | 1506 | 6.31 | 5.96 | -5.22E-01 | Neutral | -5.22E-01 | Neutral |
| 11 | p.Pro11Glu | | | 7910 | 1079 | 4.80 | 4.27 | -6.80E-01 | Neutral | -6.80E-01 | Neutral |
| 11 | p.Pro11Ala | | | 12124 | 1807 | 7.36 | 7.15 | -4.33E-01 | Neutral | -4.33E-01 | Neutral |
| 11 | p.Pro11Gly | | | 8624 | 1400 | 5.23 | 5.54 | -1.89E+00 | Neutral | -1.89E+00 | Neutral |
| 11 | p.Pro11Val | | | 8786 | 1298 | 5.33 | 5.14 | -9.61E-01 | Neutral | -9.61E-01 | Neutral |
| 11 | p.Pro11Tyr | | | 7352 | 1214 | 4.46 | 4.81 | -2.83E+00 | Neutral | -2.83E+00 | Neutral |
| 11 | p.Pro11Cys | | | 7776 | 1069 | 4.72 | 4.23 | -7.57E-01 | Neutral | -7.57E-01 | Neutral |
| 11 | p.Pro11Trp | | | 7900 | 1402 | 4.79 | 5.55 | -3.64E+00 | Neutral | -3.64E+00 | Neutral |
| 11 | p.Pro11Phe | | | 8057 | 1336 | 4.89 | 5.29 | -2.45E+00 | Neutral | -2.45E+00 | Neutral |
| 12 | p.Ser12Asn | | | 4278 | 1302 | 3.95 | 3.63 | -1.24E+00 | Neutral | -1.24E+00 | Neutral |
| 12 | p.Ser12Lys | | | 4907 | 1159 | 4.53 | 3.23 | -7.52E-02 | Neutral | -7.52E-02 | Neutral |
| 12 | p.Ser12Thr | | | 5668 | 1434 | 5.24 | 4.00 | -9.67E-02 | Neutral | -9.67E-02 | Neutral |
| 12 | p.Ser12Arg | | | 4891 | 1365 | 4.52 | 3.81 | -4.41E-01 | Neutral | -4.41E-01 | Neutral |
| 12 | p.Ser12Ser | Synonymous | | 5874 | 1944 | 5.43 | 5.42 | -1.06E+00 | Neutral | -1.06E+00 | Neutral |
| 12 | p.Ser12Ile | | | 4931 | 1318 | 4.56 | 3.67 | -2.87E-01 | Neutral | -2.87E-01 | Neutral |
| 12 | p.Ser12Met | | | 5639 | 1332 | 5.21 | 3.71 | -4.17E-02 | Neutral | -4.17E-02 | Neutral |
| 12 | p.Ser12His | | | 5177 | 1913 | 4.78 | 5.33 | -2.76E+00 | Neutral | -2.76E+00 | Neutral |
| 12 | p.Ser12Gln | | | 6165 | 2351 | 5.70 | 6.55 | -2.38E+00 | Neutral | -2.38E+00 | Neutral |
| 12 | p.Ser12Pro | | | 6926 | 1787 | 6.40 | 4.98 | -5.42E-02 | Neutral | -5.42E-02 | Neutral |
| 12 | p.Ser12Leu | | | 7110 | 5519 | 6.57 | 15.38 | -2.85E+01 | Indeterminate | -2.84E+01 | Indeterminate |
| 12 | p.Ser12Asp | | | 6223 | 1364 | 5.75 | 3.80 | -8.73E-03 | Neutral | -8.73E-03 | Neutral |
| 12 | p.Ser12Glu | | | 3816 | 1115 | 3.53 | 3.11 | -1.21E+00 | Neutral | -1.21E+00 | Neutral |
| 12 | p.Ser12Ala | | | 4304 | 1215 | 3.98 | 3.39 | -6.95E-01 | Neutral | -6.95E-01 | Neutral |
| 12 | p.Ser12Gly | | | 6165 | 1672 | 5.70 | 4.66 | -1.55E-01 | Neutral | -1.55E-01 | Neutral |
| 12 | p.Ser12Val | | | 3830 | 1433 | 3.54 | 3.99 | -4.85E+00 | Neutral | -4.85E+00 | Neutral |
| 12 | p.Ser12Tyr | | | 5082 | 1268 | 4.70 | 3.53 | -1.25E-01 | Neutral | -1.25E-01 | Neutral |
| 12 | p.Ser12Cys | | | 5540 | 2071 | 5.12 | 5.77 | -2.60E+00 | Neutral | -2.60E+00 | Neutral |
| 12 | p.Ser12Trp | | | 4602 | 1720 | 4.25 | 4.79 | -3.60E+00 | Neutral | -3.60E+00 | Neutral |
| 12 | p.Ser12Phe | | | 7080 | 2591 | 6.54 | 7.22 | -1.39E+00 | Neutral | -1.39E+00 | Neutral |
| 13 | p.Ala13Asn | | | 4218 | 1562 | 5.37 | 4.39 | -1.90E-03 | Neutral | -1.90E-03 | Neutral |
| 13 | p.Ala13Lys | | | 3029 | 1313 | 3.85 | 3.69 | -9.89E-02 | Neutral | -9.89E-02 | Neutral |
| 13 | p.Ala13Thr | | | 3969 | 1691 | 5.05 | 4.75 | -2.47E-02 | Neutral | -2.47E-02 | Neutral |
| 13 | p.Ala13Arg | | | 4846 | 1789 | 6.17 | 5.03 | -7.73E-04 | Neutral | -7.73E-04 | Neutral |
| 13 | p.Ala13Ser | | | 6020 | 2315 | 7.66 | 6.51 | -4.26E-04 | Neutral | -4.26E-04 | Neutral |
| 13 | p.Ala13Ile | | | 3905 | 1553 | 4.97 | 4.37 | -9.58E-03 | Neutral | -9.58E-03 | Neutral |
| 13 | p.Ala13Met | | | 5301 | 3446 | 6.75 | 9.69 | -8.15E-01 | Neutral | -8.15E-01 | Neutral |
| 13 | p.Ala13His | | | 3118 | 1431 | 3.97 | 4.02 | -1.69E-01 | Neutral | -1.69E-01 | Neutral |
| 13 | p.Ala13Gln | | | 3692 | 1490 | 4.70 | 4.19 | -1.59E-02 | Neutral | -1.59E-02 | Neutral |
| 13 | p.Ala13Pro | | | 4393 | 1429 | 5.59 | 4.02 | -1.21E-04 | Neutral | -1.21E-04 | Neutral |
| 13 | p.Ala13Leu | | | 2886 | 1301 | 3.67 | 3.66 | -1.84E-01 | Neutral | -1.84E-01 | Neutral |
| 13 | p.Ala13Asp | | | 4736 | 1965 | 6.03 | 5.52 | -6.79E-03 | Neutral | -6.79E-03 | Neutral |
| 13 | p.Ala13Glu | | | 3561 | 1393 | 4.53 | 3.92 | -1.19E-02 | Neutral | -1.19E-02 | Neutral |
| 13 | p.Ala13Ala | Synonymous | | 3906 | 2377 | 4.97 | 6.68 | -1.06E+00 | Neutral | -1.06E+00 | Neutral |
| 13 | p.Ala13Gly | | | 3294 | 1146 | 4.19 | 3.22 | -2.68E-03 | Neutral | -2.68E-03 | Neutral |
| 13 | p.Ala13Val | | | 3458 | 1473 | 4.40 | 4.14 | -4.60E-02 | Neutral | -4.60E-02 | Neutral |
| 13 | p.Ala13Tyr | | | 3988 | 2503 | 5.08 | 7.04 | -1.26E+00 | Neutral | -1.26E+00 | Neutral |
| 13 | p.Ala13Cys | | | 2683 | 1335 | 3.41 | 3.75 | -5.86E-01 | Neutral | -5.86E-01 | Neutral |
| 13 | p.Ala13Trp | | | 2538 | 1184 | 3.23 | 3.33 | -3.92E-01 | Neutral | -3.92E-01 | Neutral |
| 13 | p.Ala13Phe | | | 5038 | 2876 | 6.41 | 8.09 | -3.08E-01 | Neutral | -3.08E-01 | Neutral |
| 14 | p.Asp14Asn | | | 3574 | 1772 | 3.96 | 3.01 | -3.70E+00 | Neutral | -3.70E+00 | Neutral |
| 14 | p.Asp14Lys | | | 4011 | 1920 | 4.44 | 3.26 | -2.50E+00 | Neutral | -2.50E+00 | Neutral |
| 14 | p.Asp14Thr | | | 5279 | 2173 | 5.85 | 3.69 | -4.49E-01 | Neutral | -4.49E-01 | Neutral |
| 14 | p.Asp14Arg | | | 3601 | 2556 | 3.99 | 4.34 | -1.51E+01 | Indeterminate | -1.51E+01 | Indeterminate |
| 14 | p.Asp14Ser | | | 4983 | 1986 | 5.52 | 3.37 | -3.99E-01 | Neutral | -3.99E-01 | Neutral |
| 14 | p.Asp14Ile | | | 4429 | 4433 | 4.91 | 7.53 | -3.23E+01 | Indeterminate | -3.17E+01 | Indeterminate |
| 14 | p.Asp14Met | | | 3172 | 2437 | 3.51 | 4.14 | -2.16E+01 | Indeterminate | -2.16E+01 | Indeterminate |
| 14 | p.Asp14His | | | 4685 | 1942 | 5.19 | 3.30 | -6.67E-01 | Neutral | -6.67E-01 | Neutral |
| 14 | p.Asp14Gln | | | 5284 | 2584 | 5.85 | 4.39 | -1.64E+00 | Neutral | -1.64E+00 | Neutral |
| 14 | p.Asp14Pro | | | 4429 | 4941 | 4.91 | 8.39 | -4.17E+01 | Indeterminate | -3.32E+01 | Indeterminate |
| 14 | p.Asp14Leu | | | 6098 | 4767 | 6.76 | 8.09 | -1.16E+01 | Indeterminate | -1.16E+01 | Indeterminate |

| | | | | | | | | | | | | | | | | | |
|---|---|---|---|---|---|---|---|---|---|---|---|---|---|---|---|---|---|
| 14 | p.Asp14Asp | Synonymous | | 4519 | 1967 | 5.01 | 3.34 | | | | | -1.06E+00 | Neutral | | | -1.06E+00 | Neutral |
| 14 | p.Asp14Glu | | | 4387 | 2080 | 4.86 | 3.53 | | | | | -1.99E+00 | Neutral | | | -1.99E+00 | Neutral |
| 14 | p.Asp14Ala | | | 3103 | 1857 | 3.44 | 3.15 | | | | | -1.00E+01 | Indeterminate | | | -1.00E+01 | Indeterminate |
| 14 | p.Asp14Gly | | | 5701 | 2590 | 6.32 | 4.40 | | | | | -8.22E-01 | Neutral | | | -8.22E-01 | Neutral |
| 14 | p.Asp14Val | | | 5632 | 6061 | 6.24 | 10.29 | | | | | -3.17E+01 | Indeterminate | | | -3.13E+01 | Indeterminate |
| 14 | p.Asp14Tyr | | | 4781 | 4014 | 5.30 | 6.81 | | | | | -1.88E+01 | Indeterminate | | | -1.88E+01 | Indeterminate |
| 14 | p.Asp14Cys | | | 4284 | 1950 | 4.75 | 3.31 | | | | | -1.62E+00 | Neutral | | | -1.62E+00 | Neutral |
| 14 | p.Asp14Trp | | | 4208 | 3422 | 4.66 | 5.81 | | | | | -1.94E+01 | Indeterminate | | | -1.94E+01 | Indeterminate |
| 14 | p.Asp14Phe | | | 4096 | 3448 | 4.54 | 5.85 | | | | | -2.20E+01 | Indeterminate | | | -2.20E+01 | Indeterminate |
| 15 | p.Trp15Asn | | | 6806 | 1782 | 4.70 | 5.32 | 11345 | 3522 | 4.53 | 3.40 | -7.59E+00 | Indeterminate | -2.14E-02 | Neutral | -4.97E+00 | Neutral |
| 15 | p.Trp15Lys | | | 7236 | 1725 | 5.00 | 5.15 | 13768 | 4227 | 5.49 | 4.09 | -4.66E+00 | Neutral | -6.78E-03 | Neutral | -2.58E+00 | Neutral |
| 15 | p.Trp15Thr | | | 5550 | 1595 | 3.84 | 4.76 | 9633 | 4223 | 3.84 | 4.08 | -1.35E+01 | Indeterminate | -1.29E+00 | Neutral | -1.13E+01 | Indeterminate |
| 15 | p.Trp15Arg | | | 5767 | 1195 | 3.99 | 3.57 | 9685 | 2686 | 3.86 | 2.60 | -3.39E+00 | Neutral | -8.64E-03 | Neutral | -1.65E+00 | Neutral |
| 15 | p.Trp15Ser | | | 7958 | 1900 | 5.50 | 5.67 | 13848 | 4083 | 5.52 | 3.95 | -4.04E+00 | Neutral | -3.36E-03 | Neutral | -2.11E+00 | Neutral |
| 15 | p.Trp15Ile | | | 5763 | 1034 | 3.98 | 3.09 | 9127 | 3846 | 3.64 | 3.72 | -1.47E+00 | Neutral | -1.11E+00 | Neutral | -1.10E+00 | Neutral |
| 15 | p.Trp15Met | | | 5274 | 1033 | 3.64 | 3.08 | 9117 | 3643 | 3.64 | 3.52 | -2.95E+00 | Neutral | -7.47E-01 | Neutral | -1.86E+00 | Neutral |
| 15 | p.Trp15His | | | 4145 | 1068 | 2.86 | 3.19 | 7396 | 5106 | 2.95 | 4.93 | -1.30E+01 | Indeterminate | -1.53E+01 | Indeterminate | -2.39E+01 | Indeterminate |
| 15 | p.Trp15Gln | | | 7425 | 1883 | 5.13 | 5.62 | 12925 | 6246 | 5.16 | 6.04 | -5.91E+00 | Indeterminate | -1.29E+00 | Neutral | -4.61E+00 | Neutral |
| 15 | p.Trp15Pro | | | 4346 | 1381 | 3.00 | 4.12 | 8773 | 5495 | 3.50 | 5.31 | -2.33E+01 | Indeterminate | -9.05E+00 | Indeterminate | -2.78E+01 | Indeterminate |
| 15 | p.Trp15Leu | | | 12335 | 2792 | 8.52 | 8.34 | 22595 | 11054 | 9.01 | 10.68 | -1.28E+00 | Neutral | -3.16E-01 | Neutral | -5.23E-01 | Neutral |
| 15 | p.Trp15Asp | | | 11922 | 2648 | 8.24 | 7.91 | 18631 | 8445 | 7.43 | 8.16 | -1.22E+00 | Neutral | -2.81E-01 | Neutral | -4.71E-01 | Neutral |
| 15 | p.Trp15Glu | | | 10331 | 1960 | 7.14 | 5.85 | 16533 | 7828 | 6.60 | 7.57 | -5.20E-01 | Neutral | -5.96E-01 | Neutral | -2.89E-01 | Neutral |
| 15 | p.Trp15Ala | | | 8846 | 2089 | 6.11 | 6.24 | 15101 | 4990 | 6.02 | 4.82 | -3.20E+00 | Neutral | -1.32E-02 | Neutral | -1.52E+00 | Neutral |
| 15 | p.Trp15Gly | | | 5208 | 1067 | 3.60 | 3.19 | 9097 | 5665 | 3.63 | 5.48 | -3.80E+00 | Neutral | -8.46E+00 | Indeterminate | -9.01E+00 | Indeterminate |
| 15 | p.Trp15Val | | | 4607 | 847 | 3.18 | 2.53 | 7933 | 2119 | 3.16 | 2.05 | -2.67E+00 | Neutral | -1.35E-02 | Neutral | -1.17E+00 | Neutral |
| 15 | p.Trp15Tyr | | | 8482 | 2468 | 5.86 | 7.37 | 14113 | 5497 | 5.63 | 5.31 | -8.61E+00 | Indeterminate | -1.51E-01 | Neutral | -5.94E+00 | Indeterminate |
| 15 | p.Trp15Cys | | | 6277 | 1646 | 4.34 | 4.91 | 12412 | 4374 | 4.95 | 4.23 | -8.48E+00 | Indeterminate | -7.78E-02 | Neutral | -5.76E+00 | Neutral |
| 15 | p.Trp15Trp | Synonymous | | 6045 | 1051 | 4.18 | 3.14 | 9713 | 4153 | 3.87 | 4.01 | -1.06E+00 | Neutral | -1.06E+00 | Neutral | -8.18E-01 | Neutral |
| 15 | p.Trp15Phe | | | 10391 | 2331 | 7.18 | 6.96 | 18937 | 6268 | 7.55 | 6.06 | -1.76E+00 | Neutral | -4.10E-03 | Neutral | -6.10E-01 | Neutral |
| 16 | p.Leu16Asn | | | 3055 | 6743 | 4.59 | 5.25 | | | | | -5.32E+01 | Deleterious | | | -5.32E+01 | Deleterious |
| 16 | p.Leu16Lys | | | 5427 | 16967 | 8.15 | 13.21 | | | | | -5.32E+01 | Deleterious | | | -5.32E+01 | Deleterious |
| 16 | p.Leu16Thr | | | 2209 | 660 | 3.32 | 0.51 | | | | | -2.76E+01 | Indeterminate | | | -2.76E+01 | Indeterminate |
| 16 | p.Leu16Arg | Pathogenic | | 3583 | 13291 | 5.38 | 10.35 | | | | | -5.32E+01 | Deleterious | | | -5.32E+01 | Deleterious |
| 16 | p.Leu16Ser | | | 3934 | 6544 | 5.91 | 5.09 | | | | | -5.32E+01 | Deleterious | | | -5.32E+01 | Deleterious |
| 16 | p.Leu16Ile | | | 1553 | 269 | 2.33 | 0.21 | | | | | -6.96E+00 | Indeterminate | | | -6.96E+00 | Indeterminate |
| 16 | p.Leu16Met | | | 1995 | 306 | 3.00 | 0.24 | | | | | -2.63E+00 | Neutral | | | -2.63E+00 | Neutral |
| 16 | p.Leu16His | | | 4913 | 10696 | 7.38 | 8.33 | | | | | -5.32E+01 | Deleterious | | | -5.32E+01 | Deleterious |
| 16 | p.Leu16Gln | | | 4828 | 10448 | 7.25 | 8.13 | | | | | -5.32E+01 | Deleterious | | | -5.32E+01 | Deleterious |
| 16 | p.Leu16Pro | Pathogenic | | 2946 | 6033 | 4.43 | 4.70 | | | | | -5.32E+01 | Deleterious | | | -5.32E+01 | Deleterious |
| 16 | p.Leu16Leu | Synonymous | | 1846 | 239 | 2.77 | 0.19 | | | | | -1.06E+00 | Neutral | | | -1.06E+00 | Neutral |
| 16 | p.Leu16Asp | | | 3226 | 10025 | 4.85 | 7.80 | | | | | -5.32E+01 | Deleterious | | | -5.32E+01 | Deleterious |
| 16 | p.Leu16Glu | | | 4055 | 11396 | 6.09 | 8.87 | | | | | -5.32E+01 | Deleterious | | | -5.32E+01 | Deleterious |
| 16 | p.Leu16Ala | | | 3236 | 2061 | 4.86 | 1.60 | | | | | -5.32E+01 | Deleterious | | | -5.32E+01 | Deleterious |
| 16 | p.Leu16Gly | | | 4238 | 10949 | 6.37 | 8.52 | | | | | -5.32E+01 | Deleterious | | | -5.32E+01 | Deleterious |
| 16 | p.Leu16Val | | | 3046 | 804 | 4.58 | 0.63 | | | | | -1.42E+01 | Indeterminate | | | -1.42E+01 | Indeterminate |
| 16 | p.Leu16Tyr | | | 3070 | 8610 | 4.61 | 6.70 | | | | | -5.32E+01 | Deleterious | | | -5.32E+01 | Deleterious |
| 16 | p.Leu16Cys | | | 2604 | 687 | 3.91 | 0.53 | | | | | -1.67E+01 | Indeterminate | | | -1.67E+01 | Indeterminate |
| 16 | p.Leu16Trp | | | 4013 | 11211 | 6.03 | 8.73 | | | | | -5.32E+01 | Deleterious | | | -5.32E+01 | Deleterious |
| 16 | p.Leu16Phe | | | 2774 | 530 | 4.17 | 0.41 | | | | | -4.61E+00 | Neutral | | | -4.61E+00 | Neutral |
| 17 | p.Ala17Asn | | | 3109 | 1043 | 5.30 | 1.20 | | | | | -9.84E+00 | Indeterminate | | | -9.84E+00 | Indeterminate |
| 17 | p.Ala17Lys | | | 3328 | 25579 | 5.67 | 29.52 | | | | | -5.32E+01 | Deleterious | | | -5.32E+01 | Deleterious |
| 17 | p.Ala17Thr | | | 2004 | 476 | 3.41 | 0.55 | | | | | -4.30E+00 | Neutral | | | -4.30E+00 | Neutral |
| 17 | p.Ala17Arg | | | 3397 | 27882 | 5.79 | 32.18 | | | | | -5.32E+01 | Deleterious | | | -5.32E+01 | Deleterious |
| 17 | p.Ala17Ser | | | 3222 | 774 | 5.49 | 0.89 | | | | | -1.96E+00 | Neutral | | | -1.96E+00 | Neutral |
| 17 | p.Ala17Ile | | | 3809 | 1359 | 6.49 | 1.57 | | | | | -9.63E+00 | Indeterminate | | | -9.63E+00 | Indeterminate |
| 17 | p.Ala17Met | | | 1845 | 1285 | 3.14 | 1.48 | | | | | -5.32E+01 | Deleterious | | | -5.32E+01 | Deleterious |
| 17 | p.Ala17His | | | 2589 | 622 | 4.41 | 0.72 | | | | | -2.96E+00 | Neutral | | | -2.96E+00 | Neutral |
| 17 | p.Ala17Gln | | | 3620 | 2691 | 6.17 | 3.11 | | | | | -5.32E+01 | Deleterious | | | -5.32E+01 | Deleterious |
| 17 | p.Ala17Pro | | | 3101 | 714 | 5.28 | 0.82 | | | | | -1.63E+00 | Neutral | | | -1.63E+00 | Neutral |
| 17 | p.Ala17Leu | | | 2565 | 3353 | 4.37 | 3.87 | | | | | -5.32E+01 | Deleterious | | | -5.32E+01 | Deleterious |
| 17 | p.Ala17Asp | | | 3473 | 2181 | 5.92 | 2.52 | | | | | -4.98E+01 | Indeterminate | | | -3.32E+01 | Indeterminate |
| 17 | p.Ala17Glu | | | 2642 | 8434 | 4.50 | 9.73 | | | | | -5.32E+01 | Deleterious | | | -5.32E+01 | Deleterious |
| 17 | p.Ala17Ala | Synonymous | | 2835 | 595 | 4.83 | 0.69 | | | | | -1.06E+00 | Neutral | | | -1.06E+00 | Neutral |
| 17 | p.Ala17Gly | | | 3409 | 720 | 5.81 | 0.83 | | | | | -7.00E-01 | Neutral | | | -7.00E-01 | Neutral |
| 17 | p.Ala17Val | | | 3266 | 716 | 5.56 | 0.83 | | | | | -1.04E+00 | Neutral | | | -1.04E+00 | Neutral |
| 17 | p.Ala17Tyr | | | 2603 | 2585 | 4.43 | 2.98 | | | | | -5.32E+01 | Deleterious | | | -5.32E+01 | Deleterious |
| 17 | p.Ala17Cys | | | 1754 | 279 | 2.99 | 0.32 | | | | | -4.10E-01 | Neutral | | | -4.10E-01 | Neutral |
| 17 | p.Ala17Trp | | | 2913 | 3306 | 4.96 | 3.82 | | | | | -5.32E+01 | Deleterious | | | -5.32E+01 | Deleterious |
| 17 | p.Ala17Phe | | | 3210 | 2051 | 5.47 | 2.37 | | | | | -5.32E+01 | Deleterious | | | -5.32E+01 | Deleterious |
| 18 | p.Thr18Asn | | | 1065 | 1200 | 4.51 | 5.39 | | | | | -4.09E+01 | Indeterminate | | | -3.32E+01 | Indeterminate |
| 18 | p.Thr18Lys | | | 1322 | 1076 | 5.60 | 4.83 | | | | | -1.40E+01 | Indeterminate | | | -1.40E+01 | Indeterminate |
| 18 | p.Thr18Thr | Synonymous | | 1088 | 483 | 4.61 | 2.17 | | | | | -1.06E+00 | Neutral | | | -1.06E+00 | Neutral |
| 18 | p.Thr18Arg | | | 1372 | 1100 | 5.81 | 4.94 | | | | | -1.28E+01 | Indeterminate | | | -1.28E+01 | Indeterminate |
| 18 | p.Thr18Ser | | | 1365 | 878 | 5.78 | 3.94 | | | | | -5.57E+00 | Neutral | | | -5.57E+00 | Neutral |
| 18 | p.Thr18Ile | | | 921 | 852 | 3.90 | 3.82 | | | | | -2.85E+01 | Indeterminate | | | -2.85E+01 | Indeterminate |
| 18 | p.Thr18Met | | | 1963 | 2078 | 8.31 | 9.33 | | | | | -2.08E+01 | Indeterminate | | | -2.08E+01 | Indeterminate |
| 18 | p.Thr18His | | | 697 | 420 | 2.95 | 1.89 | | | | | -1.04E+01 | Indeterminate | | | -1.04E+01 | Indeterminate |
| 18 | p.Thr18Gln | | | 1320 | 973 | 5.59 | 4.37 | | | | | -1.00E+01 | Indeterminate | | | -1.00E+01 | Indeterminate |
| 18 | p.Thr18Pro | | Deleterious | 762 | 3836 | 3.23 | 17.22 | | | | | -5.32E+01 | Deleterious | | | -5.32E+01 | Deleterious |
| 18 | p.Thr18Leu | | | 1397 | 1503 | 5.92 | 6.75 | | | | | -2.94E+01 | Indeterminate | | | -2.93E+01 | Indeterminate |
| 18 | p.Thr18Asp | | | 1125 | 829 | 4.76 | 3.72 | | | | | -1.20E+01 | Indeterminate | | | -1.20E+01 | Indeterminate |
| 18 | p.Thr18Glu | | | 1271 | 825 | 5.38 | 3.70 | | | | | -6.41E+00 | Indeterminate | | | -6.41E+00 | Indeterminate |
| 18 | p.Thr18Ala | | | 1230 | 1342 | 5.21 | 6.02 | | | | | -3.39E+01 | Indeterminate | | | -3.25E+01 | Indeterminate |
| 18 | p.Thr18Gly | | | 842 | 544 | 3.57 | 2.44 | | | | | -1.06E+01 | Indeterminate | | | -1.06E+01 | Indeterminate |
| 18 | p.Thr18Val | | | 1693 | 1130 | 7.17 | 5.07 | | | | | -4.77E+00 | Neutral | | | -4.77E+00 | Neutral |
| 18 | p.Thr18Tyr | | | 785 | 884 | 3.32 | 3.97 | | | | | -5.10E+01 | Indeterminate | | | -3.32E+01 | Indeterminate |
| 18 | p.Thr18Cys | | | 1182 | 574 | 5.01 | 2.58 | | | | | -1.62E+00 | Neutral | | | -1.62E+00 | Neutral |
| 18 | p.Thr18Trp | | | 1679 | 1410 | 7.11 | 6.33 | | | | | -1.19E+01 | Indeterminate | | | -1.19E+01 | Indeterminate |
| 18 | p.Thr18Phe | | | 533 | 340 | 2.26 | 1.53 | | | | | -1.68E+01 | Indeterminate | | | -1.68E+01 | Indeterminate |
| 19 | p.Ala19Asn | | | 2103 | 1690 | 6.11 | 2.62 | | | | | -7.89E-08 | Neutral | | | -7.87E-08 | Neutral |
| 19 | p.Ala19Lys | | | 1123 | 2217 | 3.26 | 3.44 | | | | | -2.42E+00 | Neutral | | | -2.42E+00 | Neutral |
| 19 | p.Ala19Thr | | | 1850 | 2296 | 5.38 | 3.56 | | | | | -2.73E-03 | Neutral | | | -2.73E-03 | Neutral |
| 19 | p.Ala19Arg | | | 1712 | 3057 | 4.98 | 4.75 | | | | | -4.32E-01 | Neutral | | | -4.32E-01 | Neutral |
| 19 | p.Ala19Ser | | | 1700 | 1327 | 4.94 | 2.06 | | | | | -2.29E-07 | Neutral | | | -2.29E-07 | Neutral |
| 19 | p.Ala19Ile | | | 2099 | 2243 | 6.10 | 3.48 | | | | | -6.78E-05 | Neutral | | | -6.78E-05 | Neutral |
| 19 | p.Ala19Met | | | 1357 | 1021 | 3.94 | 1.59 | | | | | -5.92E-07 | Neutral | | | -5.92E-07 | Neutral |
| 19 | p.Ala19His | | | 2393 | 3205 | 6.96 | 4.98 | | | | | -2.33E-03 | Neutral | | | -2.33E-03 | Neutral |
| 19 | p.Ala19Gln | | | 1314 | 1013 | 3.82 | 1.57 | | | | | -1.39E-06 | Neutral | | | -1.39E-06 | Neutral |
| 19 | p.Ala19Pro | | | 2107 | 27833 | 6.12 | 43.22 | | | | | -5.32E+01 | Deleterious | | | -5.32E+01 | Deleterious |
| 19 | p.Ala19Leu | | | 1886 | 1318 | 5.48 | 2.05 | | | | | -5.00E-09 | Neutral | | | -4.86E-09 | Neutral |
| 19 | p.Ala19Asp | | | 1341 | 2749 | 3.90 | 4.27 | | | | | -2.14E+00 | Neutral | | | -2.14E+00 | Neutral |
| 19 | p.Ala19Glu | | | 2092 | 1938 | 6.08 | 3.01 | | | | | -2.81E-06 | Neutral | | | -2.81E-06 | Neutral |
| 19 | p.Ala19Ala | Synonymous | | 1342 | 2472 | 3.90 | 3.84 | | | | | -1.06E+00 | Neutral | | | -1.06E+00 | Neutral |
| 19 | p.Ala19Gly | | | 1446 | 1223 | 4.20 | 1.90 | | | | | -5.89E-06 | Neutral | | | -5.89E-06 | Neutral |
| 19 | p.Ala19Val | | | 1157 | 1160 | 3.36 | 1.80 | | | | | -9.74E-04 | Neutral | | | -9.74E-04 | Neutral |
| 19 | p.Ala19Tyr | | | 1572 | 1657 | 4.57 | 2.57 | | | | | -3.63E-04 | Neutral | | | -3.63E-04 | Neutral |
| 19 | p.Ala19Cys | | | 1675 | 3514 | 4.87 | 5.46 | | | | | -1.56E+00 | Neutral | | | -1.56E+00 | Neutral |
| 19 | p.Ala19Trp | | | 1976 | 1448 | 5.74 | 2.25 | | | | | -1.19E-08 | Neutral | | | -1.18E-08 | Neutral |
| 19 | p.Ala19Phe | | | 2158 | 1024 | 6.27 | 1.59 | | | | | -1.28E-14 | Neutral | | | 0.00E+00 | Neutral |
| 20 | p.Ala20Asn | | | 9582 | 6012 | 5.43 | 6.63 | 5392 | 5033 | 5.32 | 6.92 | -5.32E+01 | Deleterious | -5.32E+01 | Deleterious | -5.32E+01 | Deleterious |
| 20 | p.Ala20Lys | | | 13641 | 10495 | 7.72 | 11.57 | 7118 | 8147 | 7.02 | 11.21 | -5.32E+01 | Deleterious | -5.32E+01 | Deleterious | -5.32E+01 | Deleterious |
| 20 | p.Ala20Thr | | | 6922 | 432 | 3.92 | 0.48 | 4653 | 296 | 4.59 | 0.41 | -1.61E+01 | Indeterminate | -1.01E+01 | Indeterminate | -2.19E+01 | Indeterminate |
| 20 | p.Ala20Arg | | | 8181 | 5927 | 4.63 | 6.54 | 4101 | 4835 | 4.05 | 6.65 | -5.32E+01 | Deleterious | -5.32E+01 | Deleterious | -5.32E+01 | Deleterious |
| 20 | p.Ala20Ser | | | 4887 | 170 | 2.77 | 0.19 | 3069 | 235 | 3.03 | 0.32 | -2.25E+00 | Neutral | -2.71E+01 | Indeterminate | -2.49E+01 | Indeterminate |
| 20 | p.Ala20Ile | | | 7212 | 2564 | 4.08 | 2.83 | 4277 | 1841 | 4.22 | 2.53 | -5.32E+01 | Deleterious | -5.32E+01 | Deleterious | -5.32E+01 | Deleterious |
| 20 | p.Ala20Met | | | 8089 | 2111 | 4.58 | 2.33 | 4693 | 1696 | 4.63 | 2.33 | -5.32E+01 | Deleterious | -5.32E+01 | Deleterious | -5.32E+01 | Deleterious |
| 20 | p.Ala20His | | | 6923 | 4925 | 3.92 | 5.43 | 3697 | 4261 | 3.65 | 5.86 | -5.32E+01 | Deleterious | -5.32E+01 | Deleterious | -5.32E+01 | Deleterious |
| 20 | p.Ala20Gln | | | 10148 | 7891 | 5.75 | 8.70 | 5584 | 6254 | 5.51 | 8.60 | -5.32E+01 | Deleterious | -5.32E+01 | Deleterious | -5.32E+01 | Deleterious |
| 20 | p.Ala20Pro | | Deleterious | 9331 | 7379 | 5.28 | 8.14 | 5105 | 5579 | 5.04 | 7.67 | -5.32E+01 | Deleterious | -5.32E+01 | Deleterious | -5.32E+01 | Deleterious |
| 20 | p.Ala20Leu | | | 9121 | 2392 | 5.16 | 2.64 | 5598 | 1953 | 5.52 | 2.69 | -5.32E+01 | Deleterious | -5.32E+01 | Deleterious | -5.32E+01 | Deleterious |
| 20 | p.Ala20Asp | | | 10217 | 7096 | 5.78 | 7.82 | 5645 | 6060 | 5.57 | 8.33 | -5.32E+01 | Deleterious | -5.32E+01 | Deleterious | -5.32E+01 | Deleterious |
| 20 | p.Ala20Glu | | | 12616 | 9358 | 7.14 | 10.32 | 6694 | 7998 | 6.60 | 11.00 | -5.32E+01 | Deleterious | -5.32E+01 | Deleterious | -5.32E+01 | Deleterious |
| 20 | p.Ala20Ala | Synonymous | | 8085 | 296 | 4.58 | 0.33 | 5293 | 225 | 5.22 | 0.31 | -1.06E+00 | Neutral | -1.06E+00 | Neutral | -8.18E-01 | Neutral |
| 20 | p.Ala20Gly | | Deleterious | 7764 | 1542 | 4.40 | 1.70 | 5051 | 413 | 4.98 | 0.57 | -5.32E+01 | Deleterious | -2.06E+01 | Indeterminate | -5.32E+01 | Deleterious |
| 20 | p.Ala20Val | | | 7744 | 325 | 4.38 | 0.36 | 5039 | 329 | 4.97 | 0.45 | -2.74E+00 | Neutral | -1.01E+01 | Indeterminate | -9.53E+00 | Indeterminate |
| 20 | p.Ala20Tyr | | | 8424 | 6722 | 4.77 | 7.41 | 4712 | 5356 | 4.65 | 7.37 | -5.32E+01 | Deleterious | -5.32E+01 | Deleterious | -5.32E+01 | Deleterious |
| 20 | p.Ala20Cys | | | 8337 | 361 | 4.72 | 0.40 | 5338 | 304 | 5.27 | 0.42 | -2.84E+00 | Neutral | -5.43E+00 | Neutral | -5.52E+00 | Neutral |
| 20 | p.Ala20Trp | | | 9412 | 7408 | 5.33 | 8.17 | 4993 | 5865 | 4.93 | 8.07 | -5.32E+01 | Deleterious | -5.32E+01 | Deleterious | -5.32E+01 | Deleterious |
| 20 | p.Ala20Phe | | | 9980 | 7279 | 5.65 | 8.03 | 5306 | 6026 | 5.23 | 8.29 | -5.32E+01 | Deleterious | -5.32E+01 | Deleterious | -5.32E+01 | Deleterious |
| 21 | p.Ala21Asn | | | 1371 | 1131 | 2.99 | 1.73 | | | | | -5.32E+01 | Deleterious | | | -5.32E+01 | Deleterious |
| 21 | p.Ala21Lys | | | 2163 | 10988 | 4.72 | 16.83 | | | | | -5.32E+01 | Deleterious | | | -5.32E+01 | Deleterious |
| 21 | p.Ala21Thr | | | 1828 | 453 | 3.99 | 0.69 | | | | | -9.18E+00 | Indeterminate | | | -9.18E+00 | Indeterminate |
| 21 | p.Ala21Arg | | | 1481 | 2986 | 3.23 | 4.57 | | | | | -5.32E+01 | Deleterious | | | -5.32E+01 | Deleterious |
| 21 | p.Ala21Ser | | | 1688 | 297 | 3.68 | 0.45 | | | | | -2.09E+00 | Neutral | | | -2.09E+00 | Neutral |
| 21 | p.Ala21Ile | | | 1041 | 426 | 2.27 | 0.65 | | | | | -5.32E+01 | Deleterious | | | -5.32E+01 | Deleterious |
| 21 | p.Ala21Met | | | 2168 | 1207 | 4.73 | 1.85 | | | | | -5.32E+01 | Deleterious | | | -5.32E+01 | Deleterious |
| 21 | p.Ala21His | | | 3495 | 5459 | 7.62 | 8.36 | | | | | -5.32E+01 | Deleterious | | | -5.32E+01 | Deleterious |
| 21 | p.Ala21Gln | | | 3485 | 3708 | 7.60 | 5.68 | | | | | -5.32E+01 | Deleterious | | | -5.32E+01 | Deleterious |
| 21 | p.Ala21Pro | | | 2404 | 15276 | 5.24 | 23.40 | | | | | -5.32E+01 | Deleterious | | | -5.32E+01 | Deleterious |

| | | | | | | | | | | | | | | | | |
|---|---|---|---|---|---|---|---|---|---|---|---|---|---|---|---|---|
| 21 | p.Ala21Leu | | 3758 | 1034 | 8.19 | 1.58 | | | | | -5.31E+00 | Neutral | | | -5.31E+00 | Neutral |
| 21 | p.Ala21Asp | | 2436 | 9530 | 5.31 | 14.60 | | | | | -5.32E+01 | Deleterious | | | -5.32E+01 | Deleterious |
| 21 | p.Ala21Glu | | 1636 | 2674 | 3.57 | 4.10 | | | | | -5.32E+01 | Deleterious | | | -5.32E+01 | Deleterious |
| 21 | p.Ala21Ala | Synonymous | 1979 | 331 | 4.31 | 0.51 | | | | | -1.06E+00 | Neutral | | | -1.06E+00 | Neutral |
| 21 | p.Ala21Gly | | 2395 | 390 | 5.22 | 0.60 | | | | | -5.23E-01 | Neutral | | | -5.23E-01 | Neutral |
| 21 | p.Ala21Val | | 2784 | 547 | 6.07 | 0.84 | | | | | -1.47E+00 | Neutral | | | -1.47E+00 | Neutral |
| 21 | p.Ala21Tyr | | 2718 | 3295 | 5.93 | 5.05 | | | | | -5.32E+01 | Deleterious | | | -5.32E+01 | Deleterious |
| 21 | p.Ala21Cys | | 2659 | 468 | 5.80 | 0.72 | | | | | -7.45E-01 | Neutral | | | -7.45E-01 | Neutral |
| 21 | p.Ala21Trp | | 2399 | 3082 | 5.23 | 4.72 | | | | | -5.32E+01 | Deleterious | | | -5.32E+01 | Deleterious |
| 21 | p.Ala21Phe | | 1984 | 1999 | 4.33 | 3.06 | | | | | -5.32E+01 | Deleterious | | | -5.32E+01 | Deleterious |
| 22 | p.Arg22Asn | | 1177 | 1020 | 8.01 | 2.54 | | | | | -1.24E-07 | Neutral | | | -1.24E-07 | Neutral |
| 22 | p.Arg22Lys | | 1049 | 2391 | 7.14 | 5.96 | | | | | -1.44E+00 | Neutral | | | -1.44E+00 | Neutral |
| 22 | p.Arg22Thr | | 1153 | 5650 | 7.85 | 14.09 | | | | | -2.77E+01 | Indeterminate | | | -2.77E+01 | Indeterminate |
| 22 | p.Arg22Arg | Synonymous | 1233 | 2837 | 8.39 | 7.08 | | | | | -1.06E+00 | Neutral | | | -1.06E+00 | Neutral |
| 22 | p.Arg22Ser | | 473 | 1630 | 3.22 | 4.07 | | | | | -2.39E+01 | Indeterminate | | | -2.39E+01 | Indeterminate |
| 22 | p.Arg22Ile | | 819 | 1627 | 5.58 | 4.06 | | | | | -9.80E-01 | Neutral | | | -9.80E-01 | Neutral |
| 22 | p.Arg22Met | | 1029 | 3315 | 7.01 | 8.27 | | | | | -8.42E+00 | Indeterminate | | | -8.42E+00 | Indeterminate |
| 22 | p.Arg22His | | 338 | 167 | 2.30 | 0.42 | | | | | -2.42E-09 | Neutral | | | -2.28E-09 | Neutral |
| 22 | p.Arg22Gln | | 1275 | 1139 | 8.68 | 2.84 | | | | | -1.37E-07 | Neutral | | | -1.37E-07 | Neutral |
| 22 | p.Arg22Pro | | 424 | 6555 | 2.89 | 16.35 | | | | | -5.32E+01 | Deleterious | | | -5.32E+01 | Deleterious |
| 22 | p.Arg22Leu | | 617 | 764 | 4.20 | 1.91 | | | | | -1.65E-02 | Neutral | | | -1.65E-02 | Neutral |
| 22 | p.Arg22Asp | | 324 | 671 | 2.21 | 1.67 | | | | | -6.69E+00 | Indeterminate | | | -6.69E+00 | Indeterminate |
| 22 | p.Arg22Glu | | 737 | 522 | 5.02 | 1.30 | | | | | -3.73E-08 | Neutral | | | -3.72E-08 | Neutral |
| 22 | p.Arg22Ala | | 310 | 287 | 2.11 | 0.72 | | | | | -6.27E-03 | Neutral | | | -6.27E-03 | Neutral |
| 22 | p.Arg22Gly | | 323 | 2465 | 2.20 | 6.15 | | | | | -5.32E+01 | Deleterious | | | -5.32E+01 | Deleterious |
| 22 | p.Arg22Val | | 312 | 367 | 2.12 | 0.92 | | | | | -1.56E-01 | Neutral | | | -1.56E-01 | Neutral |
| 22 | p.Arg22Tyr | | 596 | 1333 | 4.06 | 3.32 | | | | | -3.79E+00 | Neutral | | | -3.79E+00 | Neutral |
| 22 | p.Arg22Cys | | 639 | 324 | 4.35 | 0.81 | | | | | -1.16E-11 | Neutral | | | 0.00E+00 | Neutral |
| 22 | p.Arg22Trp | | 1319 | 4784 | 8.98 | 11.93 | | | | | -9.64E+00 | Indeterminate | | | -9.64E+00 | Indeterminate |
| 22 | p.Arg22Phe | | 542 | 2250 | 3.69 | 5.61 | | | | | -3.42E+01 | Indeterminate | | | -3.26E+01 | Indeterminate |
| 23 | p.Gly23Asn | | 1971 | 217 | 4.60 | 0.67 | | | | | -2.16E+01 | Indeterminate | | | -2.16E+01 | Indeterminate |
| 23 | p.Gly23Lys | | 2291 | 2054 | 5.34 | 6.32 | | | | | -5.32E+01 | Deleterious | | | -5.32E+01 | Deleterious |
| 23 | p.Gly23Thr | | 2549 | 2313 | 5.94 | 7.12 | | | | | -5.32E+01 | Deleterious | | | -5.32E+01 | Deleterious |
| 23 | p.Gly23Arg | Likely pathogenic | 2176 | 1816 | 5.07 | 5.59 | | | | | -5.32E+01 | Deleterious | | | -5.32E+01 | Deleterious |
| 23 | p.Gly23Ser | Likely pathogenic | 1609 | 285 | 3.75 | 0.88 | | | | | -5.32E+01 | Deleterious | | | -5.32E+01 | Deleterious |
| 23 | p.Gly23Ile | | 2654 | 2752 | 6.19 | 8.47 | | | | | -5.32E+01 | Deleterious | | | -5.32E+01 | Deleterious |
| 23 | p.Gly23Met | | 2276 | 2025 | 5.31 | 6.23 | | | | | -5.32E+01 | Deleterious | | | -5.32E+01 | Deleterious |
| 23 | p.Gly23His | | 2587 | 2185 | 6.03 | 6.72 | | | | | -5.32E+01 | Deleterious | | | -5.32E+01 | Deleterious |
| 23 | p.Gly23Gln | | 2634 | 2152 | 6.14 | 6.62 | | | | | -5.32E+01 | Deleterious | | | -5.32E+01 | Deleterious |
| 23 | p.Gly23Pro | | 2042 | 1771 | 4.76 | 5.45 | | | | | -5.32E+01 | Deleterious | | | -5.32E+01 | Deleterious |
| 23 | p.Gly23Leu | | 724 | 730 | 1.69 | 2.25 | | | | | -5.32E+01 | Deleterious | | | -5.32E+01 | Deleterious |
| 23 | p.Gly23Asp | Pathogenic | 1669 | 1066 | 3.89 | 3.28 | | | | | -5.32E+01 | Deleterious | | | -5.32E+01 | Deleterious |
| 23 | p.Gly23Glu | | 2812 | 2666 | 6.56 | 8.20 | | | | | -5.32E+01 | Deleterious | | | -5.32E+01 | Deleterious |
| 23 | p.Gly23Ala | | 1966 | 112 | 4.58 | 0.34 | | | | | -1.29E+00 | Neutral | | | -1.29E+00 | Neutral |
| 23 | p.Gly23Gly | Synonymous | 1029 | 46 | 2.40 | 0.14 | | | | | -1.06E+00 | Neutral | | | -1.06E+00 | Neutral |
| 23 | p.Gly23Val | | 2948 | 2588 | 6.87 | 7.96 | | | | | -5.32E+01 | Deleterious | | | -5.32E+01 | Deleterious |
| 23 | p.Gly23Tyr | | 1763 | 1578 | 4.11 | 4.85 | | | | | -5.32E+01 | Deleterious | | | -5.32E+01 | Deleterious |
| 23 | p.Gly23Cys | | 2210 | 1286 | 5.15 | 3.96 | | | | | -5.32E+01 | Deleterious | | | -5.32E+01 | Deleterious |
| 23 | p.Gly23Trp | | 2196 | 2273 | 5.12 | 6.99 | | | | | -5.32E+01 | Deleterious | | | -5.32E+01 | Deleterious |
| 23 | p.Gly23Phe | | 2787 | 2593 | 6.50 | 7.98 | | | | | -5.32E+01 | Deleterious | | | -5.32E+01 | Deleterious |
| 24 | p.Arg24Asn | | 2933 | 1156 | 4.19 | 2.75 | 7521 | 6903 | 4.31 | 3.38 | -9.14E-02 | Neutral | -1.70E+00 | Neutral | -6.25E-01 | Neutral |
| 24 | p.Arg24Lys | | 3506 | 1119 | 5.01 | 2.66 | 8038 | 8566 | 4.61 | 4.19 | -1.60E-03 | Neutral | -3.49E+00 | Neutral | -1.72E+00 | Neutral |
| 24 | p.Arg24Thr | | 3180 | 1226 | 4.54 | 2.92 | 6972 | 8076 | 4.00 | 3.95 | -4.97E-02 | Neutral | -6.38E+00 | Neutral | -3.98E+00 | Neutral |
| 24 | p.Arg24Arg | Synonymous | 3351 | 1793 | 4.79 | 4.27 | 8843 | 7990 | 5.07 | 3.91 | -1.06E+00 | Neutral | -1.06E+00 | Neutral | -8.18E-01 | Neutral |
| 24 | p.Arg24Ser | | 3650 | 1132 | 5.21 | 2.69 | 8814 | 7806 | 5.06 | 3.82 | -7.40E-04 | Neutral | -9.24E-01 | Neutral | -2.10E-01 | Neutral |
| 24 | p.Arg24Ile | | 2629 | 1067 | 3.76 | 2.54 | 6448 | 5232 | 3.70 | 2.56 | -1.91E-01 | Neutral | -1.04E+00 | Neutral | -3.42E-01 | Neutral |
| 24 | p.Arg24Met | | 3751 | 1815 | 5.36 | 4.32 | 9977 | 9768 | 5.72 | 4.78 | -3.40E-01 | Neutral | -1.40E+00 | Neutral | -6.00E-01 | Neutral |
| 24 | p.Arg24His | | 3570 | 1006 | 5.10 | 2.39 | 9360 | 7018 | 5.37 | 3.44 | -1.29E-04 | Neutral | -1.72E-01 | Neutral | -9.56E-03 | Neutral |
| 24 | p.Arg24Gln | | 3595 | 2952 | 5.14 | 7.02 | 8545 | 12341 | 4.90 | 6.04 | -8.69E+00 | Indeterminate | -1.14E+01 | Indeterminate | -1.62E+01 | Indeterminate |
| 24 | p.Arg24Pro | Pathogenic | 2814 | 13992 | 4.02 | 33.29 | 7157 | 46361 | 4.11 | 22.70 | -5.32E+01 | Deleterious | -5.32E+01 | Deleterious | -5.32E+01 | Deleterious |
| 24 | p.Arg24Leu | | 2863 | 1249 | 4.09 | 2.97 | 7372 | 7029 | 4.23 | 3.44 | -2.98E-01 | Neutral | -2.23E+00 | Neutral | -1.07E+00 | Neutral |
| 24 | p.Arg24Asp | | 4737 | 1883 | 6.77 | 4.48 | 12409 | 9485 | 7.12 | 4.64 | -1.18E-02 | Neutral | -7.32E-02 | Neutral | -2.41E-03 | Neutral |
| 24 | p.Arg24Glu | | 3650 | 1301 | 5.21 | 3.09 | 8804 | 9043 | 5.05 | 4.43 | -8.28E-03 | Neutral | -2.43E+00 | Neutral | -1.01E+00 | Neutral |
| 24 | p.Arg24Ala | | 3165 | 902 | 4.52 | 2.15 | 8126 | 6212 | 4.66 | 3.04 | -3.62E-04 | Neutral | -3.36E-01 | Neutral | -3.41E-02 | Neutral |
| 24 | p.Arg24Gly | | 2844 | 1050 | 4.06 | 2.50 | 7234 | 8057 | 4.15 | 3.94 | -4.58E-02 | Neutral | -5.10E+00 | Neutral | -2.96E+00 | Neutral |
| 24 | p.Arg24Val | | 4782 | 1725 | 6.83 | 4.10 | 11195 | 10279 | 6.42 | 5.03 | -2.26E-03 | Neutral | -6.60E-01 | Neutral | -1.17E-01 | Neutral |
| 24 | p.Arg24Tyr | | 3352 | 1179 | 4.79 | 2.80 | 8524 | 8763 | 4.89 | 4.29 | -1.05E-02 | Neutral | -2.60E+00 | Neutral | -1.12E+00 | Neutral |
| 24 | p.Arg24Cys | | 2826 | 1270 | 4.04 | 3.02 | 6942 | 4683 | 3.98 | 2.29 | -4.09E-01 | Neutral | -1.61E-01 | Neutral | -8.97E-02 | Neutral |
| 24 | p.Arg24Trp | | 4945 | 2525 | 7.06 | 6.01 | 11886 | 12036 | 6.82 | 5.89 | -2.34E-01 | Neutral | -1.19E+00 | Neutral | -4.32E-01 | Neutral |
| 24 | p.Arg24Phe | | 3862 | 1694 | 5.52 | 4.03 | 10145 | 8607 | 5.82 | 4.21 | -1.10E-01 | Neutral | -4.45E-01 | Neutral | -8.53E-02 | Neutral |
| 25 | p.Val25Asn | | 8478 | 1522 | 4.84 | 3.94 | | | | | -4.24E+00 | Neutral | | | -4.24E+00 | Neutral |
| 25 | p.Val25Lys | | 9210 | 2493 | 5.25 | 6.45 | | | | | -1.78E+01 | Indeterminate | | | -1.78E+01 | Indeterminate |
| 25 | p.Val25Thr | | 8108 | 1182 | 4.62 | 3.06 | | | | | -1.42E+00 | Neutral | | | -1.42E+00 | Neutral |
| 25 | p.Val25Arg | | 13506 | 2689 | 7.70 | 6.96 | | | | | -3.28E+00 | Neutral | | | -3.28E+00 | Neutral |
| 25 | p.Val25Ser | | 8701 | 1111 | 4.96 | 2.87 | | | | | -4.26E-01 | Neutral | | | -4.26E-01 | Neutral |
| 25 | p.Val25Ile | | 6987 | 845 | 3.98 | 2.19 | | | | | -5.01E-01 | Neutral | | | -5.01E-01 | Neutral |
| 25 | p.Val25Met | | 10879 | 3296 | 6.20 | 8.53 | | | | | -2.10E+01 | Indeterminate | | | -2.10E+01 | Indeterminate |
| 25 | p.Val25His | | 7159 | 812 | 4.08 | 2.10 | | | | | -2.56E-01 | Neutral | | | -2.56E-01 | Neutral |
| 25 | p.Val25Gln | | 5328 | 592 | 3.04 | 1.53 | | | | | -5.25E-01 | Neutral | | | -5.25E-01 | Neutral |
| 25 | p.Val25Pro | | 4054 | 267 | 2.31 | 0.69 | | | | | -1.91E-03 | Neutral | | | -1.91E-03 | Neutral |
| 25 | p.Val25Leu | | 4540 | 391 | 2.59 | 1.01 | | | | | -6.34E-02 | Neutral | | | -6.34E-02 | Neutral |
| 25 | p.Val25Asp | | 11884 | 3484 | 6.78 | 9.01 | | | | | -1.76E+01 | Indeterminate | | | -1.76E+01 | Indeterminate |
| 25 | p.Val25Glu | | 11601 | 1495 | 6.62 | 3.87 | | | | | -1.83E-01 | Neutral | | | -1.83E-01 | Neutral |
| 25 | p.Val25Ala | | 6216 | 5757 | 3.55 | 14.90 | | | | | -5.32E+01 | Deleterious | | | -5.32E+01 | Deleterious |
| 25 | p.Val25Gly | | 10784 | 3197 | 6.15 | 8.27 | | | | | -1.99E+01 | Indeterminate | | | -1.99E+01 | Indeterminate |
| 25 | p.Val25Val | Synonymous | 9182 | 1337 | 5.24 | 3.46 | | | | | -1.06E+00 | Neutral | | | -1.06E+00 | Neutral |
| 25 | p.Val25Tyr | | 9422 | 3948 | 5.37 | 10.21 | | | | | -5.20E+01 | Indeterminate | | | -3.32E+01 | Indeterminate |
| 25 | p.Val25Cys | | 9741 | 871 | 5.56 | 2.25 | | | | | -2.47E-03 | Neutral | | | -2.47E-03 | Neutral |
| 25 | p.Val25Trp | | 11563 | 2037 | 6.59 | 5.27 | | | | | -2.22E+00 | Neutral | | | -2.22E+00 | Neutral |
| 25 | p.Val25Phe | | 7991 | 1324 | 4.56 | 3.43 | | | | | -3.13E+00 | Neutral | | | -3.13E+00 | Neutral |
| 26 | p.Glu26Asn | | 2618 | 1398 | 2.56 | 2.14 | | | | | -1.41E+00 | Neutral | | | -1.41E+00 | Neutral |
| 26 | p.Glu26Lys | | 4162 | 2229 | 4.06 | 3.42 | | | | | -4.39E-01 | Neutral | | | -4.39E-01 | Neutral |
| 26 | p.Glu26Thr | | 1175 | 706 | 1.15 | 1.08 | | | | | -9.37E+00 | Indeterminate | | | -9.37E+00 | Indeterminate |
| 26 | p.Glu26Arg | | 8484 | 4992 | 8.28 | 7.65 | | | | | -1.02E-01 | Neutral | | | -1.02E-01 | Neutral |
| 26 | p.Glu26Ser | | 903 | 705 | 0.88 | 1.08 | | | | | -2.74E+01 | Indeterminate | | | -2.74E+01 | Indeterminate |
| 26 | p.Glu26Ile | | 6233 | 3893 | 6.09 | 5.97 | | | | | -5.21E-01 | Neutral | | | -5.21E-01 | Neutral |
| 26 | p.Glu26Met | | 5772 | 3272 | 5.64 | 5.02 | | | | | -2.70E-01 | Neutral | | | -2.70E-01 | Neutral |
| 26 | p.Glu26His | | 3306 | 2308 | 3.23 | 3.54 | | | | | -4.15E+00 | Neutral | | | -4.15E+00 | Neutral |
| 26 | p.Glu26Gln | | 4189 | 2887 | 4.09 | 4.43 | | | | | -2.58E+00 | Neutral | | | -2.58E+00 | Neutral |
| 26 | p.Glu26Pro | | 9408 | 6512 | 9.19 | 9.98 | | | | | -3.86E-01 | Neutral | | | -3.86E-01 | Neutral |
| 26 | p.Glu26Leu | | 4257 | 2348 | 4.16 | 3.60 | | | | | -5.32E-01 | Neutral | | | -5.32E-01 | Neutral |
| 26 | p.Glu26Asp | | 3303 | 2162 | 3.23 | 3.31 | | | | | -3.00E+00 | Neutral | | | -3.00E+00 | Neutral |
| 26 | p.Glu26Glu | Synonymous | 5487 | 3596 | 5.36 | 5.51 | | | | | -1.06E+00 | Neutral | | | -1.06E+00 | Neutral |
| 26 | p.Glu26Ala | | 5080 | 3561 | 4.96 | 5.46 | | | | | -1.96E+00 | Neutral | | | -1.96E+00 | Neutral |
| 26 | p.Glu26Gly | | 2672 | 1759 | 2.61 | 2.70 | | | | | -4.39E+00 | Neutral | | | -4.39E+00 | Neutral |
| 26 | p.Glu26Val | | 9172 | 5895 | 8.96 | 9.04 | | | | | -2.03E-01 | Neutral | | | -2.03E-01 | Neutral |
| 26 | p.Glu26Tyr | | 3224 | 2365 | 3.15 | 3.63 | | | | | -5.42E+00 | Neutral | | | -5.42E+00 | Neutral |
| 26 | p.Glu26Cys | | 14018 | 8717 | 13.69 | 13.36 | | | | | -2.39E-02 | Neutral | | | -2.39E-02 | Neutral |
| 26 | p.Glu26Trp | | 5619 | 3660 | 5.49 | 5.61 | | | | | -9.58E-01 | Neutral | | | -9.58E-01 | Neutral |
| 26 | p.Glu26Phe | | 3324 | 2258 | 3.25 | 3.46 | | | | | -3.59E+00 | Neutral | | | -3.59E+00 | Neutral |
| 27 | p.Glu27Asn | | 3049 | 116 | 1.35 | 0.54 | 2068 | 545 | 1.19 | 1.12 | -1.15E-02 | Neutral | -1.91E+01 | Indeterminate | -1.53E+01 | Indeterminate |
| 27 | p.Glu27Lys | | 23016 | 2287 | 10.17 | 10.63 | 16797 | 4397 | 9.63 | 9.04 | -3.85E-01 | Neutral | -6.79E-01 | Neutral | -2.67E-01 | Neutral |
| 27 | p.Glu27Thr | | 9777 | 1746 | 4.32 | 8.12 | 7364 | 2199 | 4.22 | 4.52 | -2.54E+01 | Indeterminate | -6.92E+00 | Indeterminate | -2.78E+01 | Indeterminate |
| 27 | p.Glu27Arg | | 9627 | 961 | 4.26 | 4.47 | 7549 | 2188 | 4.33 | 4.50 | -3.03E+00 | Neutral | -5.90E+00 | Neutral | -6.08E+00 | Indeterminate |
| 27 | p.Glu27Ser | | 7381 | 589 | 3.26 | 2.74 | 6110 | 1601 | 3.50 | 3.29 | -1.34E+00 | Neutral | -5.15E+00 | Neutral | -4.04E+00 | Neutral |
| 27 | p.Glu27Ile | | 12256 | 855 | 5.42 | 3.97 | 9316 | 2786 | 5.34 | 5.73 | -8.34E-02 | Neutral | -4.97E+00 | Neutral | -2.89E+00 | Neutral |
| 27 | p.Glu27Met | | 13701 | 1347 | 6.06 | 6.26 | 10283 | 2849 | 5.89 | 5.85 | -1.37E+00 | Neutral | -2.90E+00 | Neutral | -2.28E+00 | Neutral |
| 27 | p.Glu27His | | 11903 | 1119 | 5.26 | 5.20 | 8714 | 2582 | 5.00 | 5.31 | -1.39E+00 | Neutral | -5.27E+00 | Neutral | -4.17E+00 | Neutral |
| 27 | p.Glu27Gln | | 12459 | 1195 | 5.51 | 5.56 | 9245 | 2630 | 5.30 | 5.40 | -1.44E+00 | Neutral | -3.98E+00 | Neutral | -3.17E+00 | Neutral |
| 27 | p.Glu27Pro | | 14286 | 1751 | 6.31 | 8.14 | 11214 | 3736 | 6.43 | 7.68 | -4.40E+00 | Neutral | -6.10E+00 | Neutral | -7.45E+00 | Indeterminate |
| 27 | p.Glu27Leu | | 11139 | 661 | 4.92 | 3.07 | 8014 | 2183 | 4.59 | 4.49 | -1.43E-02 | Neutral | -4.06E+00 | Neutral | -2.14E+00 | Neutral |
| 27 | p.Glu27Asp | | 4309 | 353 | 1.90 | 1.64 | 3024 | 1006 | 1.73 | 2.07 | -4.30E+00 | Neutral | -2.57E+01 | Indeterminate | -2.56E+01 | Indeterminate |
| 27 | p.Glu27Glu | Synonymous | 18925 | 1996 | 8.37 | 9.28 | 14627 | 3878 | 8.38 | 7.97 | -1.06E+00 | Neutral | -1.06E+00 | Neutral | -8.18E-01 | Neutral |
| 27 | p.Glu27Ala | | 10932 | 963 | 4.83 | 4.48 | 8227 | 2282 | 4.72 | 4.69 | -1.08E+00 | Neutral | -4.24E+00 | Neutral | -3.09E+00 | Neutral |
| 27 | p.Glu27Gly | | 3311 | 311 | 1.46 | 1.45 | 3177 | 813 | 1.82 | 1.67 | -1.07E+01 | Indeterminate | -1.11E+01 | Indeterminate | -1.78E+01 | Indeterminate |
| 27 | p.Glu27Val | | 18682 | 1607 | 8.26 | 7.47 | 14601 | 4170 | 8.37 | 8.57 | -1.84E-01 | Neutral | -1.76E+00 | Neutral | -7.10E-01 | Neutral |
| 27 | p.Glu27Tyr | | 4149 | 804 | 1.83 | 3.74 | 3759 | 949 | 2.15 | 1.95 | -5.32E+01 | Deleterious | -8.60E+00 | Indeterminate | -5.32E+01 | Deleterious |
| 27 | p.Glu27Cys | | 4188 | 379 | 1.85 | 1.76 | 3656 | 1088 | 2.10 | 2.24 | -6.94E+00 | Indeterminate | -1.56E+01 | Indeterminate | -1.85E+01 | Indeterminate |
| 27 | p.Glu27Trp | | 18527 | 1531 | 8.19 | 7.12 | 13839 | 3589 | 7.93 | 7.38 | -1.22E-01 | Neutral | -1.03E+00 | Neutral | -3.07E-01 | Neutral |
| 27 | p.Glu27Phe | | 14611 | 941 | 6.46 | 4.37 | 12869 | 3189 | 7.38 | 6.55 | -1.26E-02 | Neutral | -8.83E-01 | Neutral | -1.99E-01 | Neutral |
| 28 | p.Val28Asn | | 9727 | 11536 | 4.96 | 7.12 | | | | | -5.32E+01 | Deleterious | | | -5.32E+01 | Deleterious |
| 28 | p.Val28Lys | | 9389 | 15308 | 4.79 | 9.45 | | | | | -5.32E+01 | Deleterious | | | -5.32E+01 | Deleterious |
| 28 | p.Val28Thr | | 9288 | 1619 | 4.74 | 1.00 | | | | | -1.58E-01 | Neutral | | | -1.58E-01 | Neutral |
| 28 | p.Val28Arg | | 8484 | 12431 | 4.33 | 7.67 | | | | | -5.32E+01 | Deleterious | | | -5.32E+01 | Deleterious |
| 28 | p.Val28Ser | | 11984 | 2938 | 6.12 | 1.81 | | | | | -1.43E+00 | Neutral | | | -1.43E+00 | Neutral |
| 28 | p.Val28Ile | | 10439 | 1906 | 5.33 | 1.18 | | | | | -1.71E-01 | Neutral | | | -1.71E-01 | Neutral |
| 28 | p.Val28Met | | 7038 | 1186 | 3.59 | 0.73 | | | | | -2.88E-01 | Neutral | | | -2.88E-01 | Neutral |
| 28 | p.Val28His | | 7337 | 11305 | 3.74 | 6.98 | | | | | -5.32E+01 | Deleterious | | | -5.32E+01 | Deleterious |
| 28 | p.Val28Gln | | 9411 | 9712 | 4.80 | 5.99 | | | | | -5.32E+01 | Deleterious | | | -5.32E+01 | Deleterious |

| | | | | | | | | | | | | | | | | | |
|---|---|---|---|---|---|---|---|---|---|---|---|---|---|---|---|---|---|
| 28 | p.Val28Pro | | | 8623 | 12625 | 4.40 | 7.79 | | | | | -5.32E+01 | Deleterious | | | -5.32E+01 | Deleterious |
| 28 | p.Val28Leu | | | 11339 | 1580 | 5.79 | 0.98 | | | | | -2.63E-03 | Neutral | | | -2.63E-03 | Neutral |
| 28 | p.Val28Asp | | | 10547 | 15278 | 5.38 | 9.43 | | | | | -5.32E+01 | Deleterious | | | -5.32E+01 | Deleterious |
| 28 | p.Val28Glu | | | 12987 | 12241 | 6.63 | 7.56 | | | | | -5.32E+01 | Deleterious | | | -5.32E+01 | Deleterious |
| 28 | p.Val28Ala | | | 11044 | 1828 | 5.64 | 1.13 | | | | | -4.21E-02 | Neutral | | | -4.21E-02 | Neutral |
| 28 | p.Val28Gly | | | 8378 | 3399 | 4.28 | 2.10 | | | | | -2.04E+01 | Indeterminate | | | -2.04E+01 | Indeterminate |
| 28 | p.Val28Val | Synonymous | | 11421 | 2640 | 5.83 | 1.63 | | | | | -1.06E+00 | Neutral | | | -1.06E+00 | Neutral |
| 28 | p.Val28Tyr | | | 8913 | 13513 | 4.55 | 8.34 | | | | | -5.32E+01 | Deleterious | | | -5.32E+01 | Deleterious |
| 28 | p.Val28Cys | | | 8264 | 1344 | 4.22 | 0.83 | | | | | -1.12E-01 | Neutral | | | -1.12E-01 | Neutral |
| 28 | p.Val28Trp | | | 11572 | 19076 | 5.91 | 11.77 | | | | | -5.32E+01 | Deleterious | | | -5.32E+01 | Deleterious |
| 28 | p.Val28Phe | | | 9745 | 10542 | 4.97 | 6.51 | | | | | -5.32E+01 | Deleterious | | | -5.32E+01 | Deleterious |
| 29 | p.Arg29Asn | | | 20602 | 4626 | 6.41 | 6.71 | | | | | -4.64E+00 | Neutral | | | -4.64E+00 | Neutral |
| 29 | p.Arg29Lys | | | 20053 | 2816 | 6.24 | 4.08 | | | | | -1.73E-01 | Neutral | | | -1.73E-01 | Neutral |
| 29 | p.Arg29Thr | | | 12052 | 2884 | 3.75 | 4.18 | | | | | -1.21E+01 | Indeterminate | | | -1.21E+01 | Indeterminate |
| 29 | p.Arg29Arg | Synonymous | | 17229 | 2824 | 5.36 | 4.09 | | | | | -1.06E+00 | Neutral | | | -1.06E+00 | Neutral |
| 29 | p.Arg29Ser | | | 32631 | 6359 | 10.16 | 9.22 | | | | | -7.93E-01 | Neutral | | | -7.93E-01 | Neutral |
| 29 | p.Arg29Ile | | | 15556 | 4861 | 4.84 | 7.05 | | | | | -2.12E+01 | Indeterminate | | | -2.12E+01 | Indeterminate |
| 29 | p.Arg29Met | | | 16707 | 2888 | 5.20 | 4.19 | | | | | -1.63E+00 | Neutral | | | -1.63E+00 | Neutral |
| 29 | p.Arg29His | | | 7538 | 1143 | 2.35 | 1.66 | | | | | -3.49E+00 | Neutral | | | -3.49E+00 | Neutral |
| 29 | p.Arg29Gln | | | 19103 | 2763 | 5.95 | 4.01 | | | | | -2.77E-01 | Neutral | | | -2.77E-01 | Neutral |
| 29 | p.Arg29Pro | | | 4160 | 10588 | 1.30 | 15.35 | | | | | -5.32E+01 | Deleterious | | | -5.32E+01 | Deleterious |
| 29 | p.Arg29Leu | | | 19150 | 3168 | 5.96 | 4.59 | | | | | -8.77E-01 | Neutral | | | -8.77E-01 | Neutral |
| 29 | p.Arg29Asp | | | 17057 | 5258 | 5.31 | 7.62 | | | | | -1.87E+01 | Indeterminate | | | -1.87E+01 | Indeterminate |
| 29 | p.Arg29Glu | | | 13773 | 2739 | 4.29 | 3.97 | | | | | -4.92E+00 | Neutral | | | -4.92E+00 | Neutral |
| 29 | p.Arg29Ala | | | 17777 | 1898 | 5.53 | 2.75 | | | | | -7.71E-03 | Neutral | | | -7.71E-03 | Neutral |
| 29 | p.Arg29Gly | | | 19312 | 3544 | 6.01 | 5.14 | | | | | -1.76E+00 | Neutral | | | -1.76E+00 | Neutral |
| 29 | p.Arg29Val | | | 10545 | 1606 | 3.28 | 2.33 | | | | | -1.92E+00 | Neutral | | | -1.92E+00 | Neutral |
| 29 | p.Arg29Tyr | | | 18295 | 2604 | 5.70 | 3.77 | | | | | -2.72E-01 | Neutral | | | -2.72E-01 | Neutral |
| 29 | p.Arg29Cys | | | 22937 | 4095 | 7.14 | 5.94 | | | | | -9.90E-01 | Neutral | | | -9.90E-01 | Neutral |
| 29 | p.Arg29Trp | | | 9347 | 1224 | 2.91 | 1.77 | | | | | -9.07E-01 | Neutral | | | -9.07E-01 | Neutral |
| 29 | p.Arg29Phe | | | 7409 | 1098 | 2.31 | 1.59 | | | | | -3.19E+00 | Neutral | | | -3.19E+00 | Neutral |
| 30 | p.Ala30Asn | | | 21267 | 13488 | 5.27 | 11.35 | | | | | -5.32E+01 | Deleterious | | | -5.32E+01 | Deleterious |
| 30 | p.Ala30Lys | | | 17443 | 2724 | 4.32 | 2.29 | | | | | -1.25E+00 | Neutral | | | -1.25E+00 | Neutral |
| 30 | p.Ala30Thr | | | 24071 | 8424 | 5.96 | 7.09 | | | | | -2.40E+01 | Indeterminate | | | -2.40E+01 | Indeterminate |
| 30 | p.Ala30Arg | | | 26854 | 5725 | 6.65 | 4.82 | | | | | -3.40E+00 | Neutral | | | -3.40E+00 | Neutral |
| 30 | p.Ala30Ser | | | 20268 | 5206 | 5.02 | 4.38 | | | | | -1.11E+01 | Indeterminate | | | -1.11E+01 | Indeterminate |
| 30 | p.Ala30Ile | | | 18034 | 4333 | 4.47 | 3.64 | | | | | -1.00E+01 | Indeterminate | | | -1.00E+01 | Indeterminate |
| 30 | p.Ala30Met | | | 27133 | 6428 | 6.72 | 5.41 | | | | | -5.52E+00 | Neutral | | | -5.52E+00 | Neutral |
| 30 | p.Ala30His | | | 18096 | 3956 | 4.48 | 3.33 | | | | | -6.94E+00 | Indeterminate | | | -6.94E+00 | Indeterminate |
| 30 | p.Ala30Gln | | | 18462 | 4653 | 4.57 | 3.91 | | | | | -1.15E+01 | Indeterminate | | | -1.15E+01 | Indeterminate |
| 30 | p.Ala30Pro | | | 18318 | 9216 | 4.54 | 7.75 | | | | | -5.32E+01 | Deleterious | | | -5.32E+01 | Deleterious |
| 30 | p.Ala30Leu | | | 14439 | 3198 | 3.58 | 2.69 | | | | | -9.74E+00 | Indeterminate | | | -9.74E+00 | Indeterminate |
| 30 | p.Ala30Asp | | | 11706 | 4156 | 2.90 | 3.50 | | | | | -4.46E+01 | Indeterminate | | | -3.32E+01 | Indeterminate |
| 30 | p.Ala30Glu | | | 21369 | 3913 | 5.30 | 3.29 | | | | | -2.24E+00 | Neutral | | | -2.24E+00 | Neutral |
| 30 | p.Ala30Ala | Synonymous | | 25468 | 4405 | 6.31 | 3.71 | | | | | -1.06E+00 | Neutral | | | -1.06E+00 | Neutral |
| 30 | p.Ala30Gly | | | 17570 | 10117 | 4.35 | 8.51 | | | | | -5.32E+01 | Deleterious | | | -5.32E+01 | Deleterious |
| 30 | p.Ala30Val | | | 33236 | 8979 | 8.24 | 7.55 | | | | | -7.22E+00 | Indeterminate | | | -7.22E+00 | Indeterminate |
| 30 | p.Ala30Tyr | | | 16045 | 5032 | 3.98 | 4.23 | | | | | -2.56E+01 | Indeterminate | | | -2.56E+01 | Indeterminate |
| 30 | p.Ala30Cys | | | 15100 | 4355 | 3.74 | 3.66 | | | | | -2.15E+01 | Indeterminate | | | -2.15E+01 | Indeterminate |
| 30 | p.Ala30Trp | | | 26402 | 7996 | 6.54 | 6.73 | | | | | -1.43E+01 | Indeterminate | | | -1.43E+01 | Indeterminate |
| 30 | p.Ala30Phe | | | 12276 | 2580 | 3.04 | 2.17 | | | | | -9.80E+00 | Indeterminate | | | -9.80E+00 | Indeterminate |
| 31 | p.Leu31Asn | | | 17658 | 3677 | 4.70 | 3.62 | | | | | -1.58E+01 | Indeterminate | | | -1.58E+01 | Indeterminate |
| 31 | p.Leu31Lys | | | 22252 | 3043 | 5.93 | 3.00 | | | | | -1.84E+00 | Neutral | | | -1.84E+00 | Neutral |
| 31 | p.Leu31Thr | | | 16130 | 2611 | 4.30 | 2.57 | | | | | -7.23E+00 | Indeterminate | | | -7.23E+00 | Indeterminate |
| 31 | p.Leu31Arg | | | 19399 | 2812 | 5.17 | 2.77 | | | | | -3.31E+00 | Neutral | | | -3.31E+00 | Neutral |
| 31 | p.Leu31Ser | | | 18159 | 3537 | 4.84 | 3.49 | | | | | -1.24E+01 | Indeterminate | | | -1.24E+01 | Indeterminate |
| 31 | p.Leu31Ile | | | 26461 | 3657 | 7.05 | 3.61 | | | | | -1.36E+00 | Neutral | | | -1.36E+00 | Neutral |
| 31 | p.Leu31Met | | | 25568 | 4775 | 6.81 | 4.71 | | | | | -6.95E+00 | Indeterminate | | | -6.95E+00 | Indeterminate |
| 31 | p.Leu31His | | | 14102 | 2056 | 3.76 | 2.03 | | | | | -5.62E+00 | Neutral | | | -5.62E+00 | Neutral |
| 31 | p.Leu31Gln | | | 20265 | 4695 | 5.40 | 4.63 | | | | | -1.90E+01 | Indeterminate | | | -1.90E+01 | Indeterminate |
| 31 | p.Leu31Pro | | | 18066 | 30967 | 4.81 | 30.53 | | | | | -5.32E+01 | Deleterious | | | -5.32E+01 | Deleterious |
| 31 | p.Leu31Leu | Synonymous | | 29114 | 4009 | 7.76 | 3.95 | | | | | -1.06E+00 | Neutral | | | -1.06E+00 | Neutral |
| 31 | p.Leu31Asp | | | 17841 | 8877 | 4.75 | 8.75 | | | | | -5.32E+01 | Deleterious | | | -5.32E+01 | Deleterious |
| 31 | p.Leu31Glu | | | 12731 | 2060 | 3.39 | 2.03 | | | | | -9.76E+00 | Indeterminate | | | -9.76E+00 | Indeterminate |
| 31 | p.Leu31Ala | | | 17448 | 3125 | 4.65 | 3.08 | | | | | -9.65E+00 | Indeterminate | | | -9.65E+00 | Indeterminate |
| 31 | p.Leu31Gly | | | 16782 | 2693 | 4.47 | 2.65 | | | | | -6.62E+00 | Indeterminate | | | -6.62E+00 | Indeterminate |
| 31 | p.Leu31Val | | | 16042 | 3588 | 4.27 | 3.54 | | | | | -2.14E+01 | Indeterminate | | | -2.14E+01 | Indeterminate |
| 31 | p.Leu31Tyr | | | 29006 | 7869 | 7.73 | 7.76 | | | | | -2.12E+01 | Indeterminate | | | -2.12E+01 | Indeterminate |
| 31 | p.Leu31Cys | | | 10884 | 2718 | 2.90 | 2.68 | | | | | -3.93E+01 | Indeterminate | | | -3.32E+01 | Indeterminate |
| 31 | p.Leu31Trp | | | 13277 | 2652 | 3.54 | 2.61 | | | | | -1.85E+01 | Indeterminate | | | -1.85E+01 | Indeterminate |
| 31 | p.Leu31Phe | | | 14121 | 2017 | 3.76 | 1.99 | | | | | -5.13E+00 | Neutral | | | -5.13E+00 | Neutral |
| 32 | p.Leu32Asn | | | 2757 | 2086 | 5.77 | 6.42 | 1361 | 1177 | 5.80 | 7.89 | -2.49E+01 | Indeterminate | -2.98E+01 | Indeterminate | -5.32E+01 | Deleterious |
| 32 | p.Leu32Lys | | | 4132 | 3792 | 8.65 | 11.66 | 1940 | 1823 | 8.26 | 12.23 | -2.90E+01 | Indeterminate | -2.73E+01 | Indeterminate | -5.32E+01 | Deleterious |
| 32 | p.Leu32Thr | | | 1677 | 990 | 3.51 | 3.05 | 942 | 231 | 4.01 | 1.55 | -1.98E+01 | Indeterminate | -3.76E-02 | Neutral | -1.59E+01 | Indeterminate |
| 32 | p.Leu32Arg | | | 1964 | 1636 | 4.11 | 5.03 | 944 | 914 | 4.02 | 6.13 | -4.17E+01 | Indeterminate | -5.14E+01 | Indeterminate | -5.32E+01 | Deleterious |
| 32 | p.Leu32Ser | | | 2088 | 1649 | 4.37 | 5.07 | 1437 | 721 | 6.12 | 4.84 | -3.51E+01 | Indeterminate | -4.41E+00 | Neutral | -3.25E+01 | Indeterminate |
| 32 | p.Leu32Ile | | | 2006 | 586 | 4.20 | 1.80 | 1122 | 207 | 4.78 | 1.39 | -4.47E-01 | Neutral | -1.01E-04 | Neutral | -5.77E-02 | Neutral |
| 32 | p.Leu32Met | | | 3442 | 1252 | 7.21 | 3.85 | 1437 | 260 | 6.12 | 1.74 | -6.29E-01 | Neutral | -1.11E-05 | Neutral | -1.07E-01 | Neutral |
| 32 | p.Leu32His | | | 1717 | 1516 | 3.60 | 4.66 | 795 | 651 | 3.39 | 4.37 | -5.20E+01 | Indeterminate | -4.02E+01 | Indeterminate | -5.32E+01 | Deleterious |
| 32 | p.Leu32Gln | | | 2563 | 1371 | 5.37 | 4.22 | 1457 | 756 | 6.21 | 5.07 | -9.05E+00 | Indeterminate | -5.05E+00 | Neutral | -1.07E+01 | Indeterminate |
| 32 | p.Leu32Pro | Pathogenic | | 4912 | 4768 | 10.29 | 14.67 | 1841 | 2065 | 7.84 | 13.85 | -2.89E+01 | Indeterminate | -4.36E+01 | Indeterminate | -5.32E+01 | Deleterious |
| 32 | p.Leu32Leu | Synonymous | | 1981 | 642 | 4.15 | 1.97 | 851 | 281 | 3.63 | 1.88 | -1.06E+00 | Neutral | -1.06E+00 | Neutral | -8.18E-01 | Neutral |
| 32 | p.Leu32Asp | | | 2011 | 1682 | 4.21 | 5.17 | 723 | 788 | 3.08 | 5.29 | -4.13E+01 | Indeterminate | -5.32E+01 | Deleterious | -5.32E+01 | Deleterious |
| 32 | p.Leu32Glu | | | 2518 | 2363 | 5.27 | 7.27 | 1148 | 1069 | 4.89 | 7.17 | -4.52E+01 | Indeterminate | -4.08E+01 | Indeterminate | -5.32E+01 | Deleterious |
| 32 | p.Leu32Ala | | | 876 | 322 | 1.83 | 0.99 | 256 | 98 | 1.09 | 0.66 | -8.43E+00 | Indeterminate | -1.42E+01 | Indeterminate | -1.86E+01 | Indeterminate |
| 32 | p.Leu32Gly | | | 1595 | 1406 | 3.34 | 4.32 | 878 | 716 | 3.74 | 4.80 | -5.32E+01 | Deleterious | -3.68E+01 | Indeterminate | -5.32E+01 | Deleterious |
| 32 | p.Leu32Val | | | 3564 | 1243 | 7.46 | 3.82 | 1582 | 300 | 6.74 | 2.01 | -3.96E-01 | Neutral | -1.56E-05 | Neutral | -4.60E-02 | Neutral |
| 32 | p.Leu32Tyr | | | 1947 | 1414 | 4.08 | 4.35 | 1105 | 911 | 4.71 | 6.11 | -3.02E+01 | Indeterminate | -3.14E+01 | Indeterminate | -5.32E+01 | Deleterious |
| 32 | p.Leu32Cys | | | 1419 | 550 | 2.97 | 1.69 | 793 | 133 | 3.38 | 0.89 | -5.43E+00 | Neutral | -1.53E-04 | Neutral | -3.18E+00 | Neutral |
| 32 | p.Leu32Trp | | | 2952 | 2525 | 6.18 | 7.77 | 1653 | 1521 | 7.04 | 10.20 | -3.22E+01 | Indeterminate | -2.96E+01 | Indeterminate | -5.32E+01 | Deleterious |
| 32 | p.Leu32Phe | | | 1622 | 716 | 3.40 | 2.20 | 1208 | 288 | 5.15 | 1.93 | -7.76E+00 | Indeterminate | -7.36E-03 | Neutral | -5.09E+00 | Neutral |
| 33 | p.Glu33Asn | | | 2405 | 1987 | 6.29 | 6.03 | | | | | -1.19E-01 | Neutral | | | -1.19E-01 | Neutral |
| 33 | p.Glu33Lys | | | 2310 | 2125 | 6.04 | 6.45 | | | | | -4.10E-01 | Neutral | | | -4.10E-01 | Neutral |
| 33 | p.Glu33Thr | | | 2551 | 2244 | 6.67 | 6.81 | | | | | -1.92E-01 | Neutral | | | -1.92E-01 | Neutral |
| 33 | p.Glu33Arg | | | 323 | 265 | 0.84 | 0.80 | | | | | -9.24E+00 | Indeterminate | | | -9.24E+00 | Indeterminate |
| 33 | p.Glu33Ser | | | 894 | 826 | 2.34 | 2.51 | | | | | -3.59E+00 | Neutral | | | -3.59E+00 | Neutral |
| 33 | p.Glu33Ile | | | 3960 | 3269 | 10.35 | 9.92 | | | | | -1.31E-02 | Neutral | | | -1.31E-02 | Neutral |
| 33 | p.Glu33Met | | | 4009 | 3279 | 10.48 | 9.95 | | | | | -1.07E-02 | Neutral | | | -1.07E-02 | Neutral |
| 33 | p.Glu33His | | | 2694 | 2397 | 7.04 | 7.28 | | | | | -1.79E-01 | Neutral | | | -1.79E-01 | Neutral |
| 33 | p.Glu33Gln | | | 3450 | 3143 | 9.02 | 9.54 | | | | | -9.23E-02 | Neutral | | | -9.23E-02 | Neutral |
| 33 | p.Glu33Pro | | | 2489 | 1886 | 6.51 | 5.73 | | | | | -3.51E-02 | Neutral | | | -3.51E-02 | Neutral |
| 33 | p.Glu33Leu | | | 2006 | 1914 | 5.25 | 5.81 | | | | | -8.22E-01 | Neutral | | | -8.22E-01 | Neutral |
| 33 | p.Glu33Asp | | | 1895 | 1477 | 4.96 | 4.48 | | | | | -1.51E-01 | Neutral | | | -1.51E-01 | Neutral |
| 33 | p.Glu33Glu | Synonymous | | 1646 | 1522 | 4.30 | 4.62 | | | | | -1.06E+00 | Neutral | | | -1.06E+00 | Neutral |
| 33 | p.Glu33Ala | | | 425 | 405 | 1.11 | 1.23 | | | | | -1.14E+01 | Indeterminate | | | -1.14E+01 | Indeterminate |
| 33 | p.Glu33Gly | | | 2167 | 1901 | 5.67 | 5.77 | | | | | -3.21E-01 | Neutral | | | -3.21E-01 | Neutral |
| 33 | p.Glu33Val | | | 1148 | 984 | 3.00 | 2.99 | | | | | -1.44E+00 | Neutral | | | -1.44E+00 | Neutral |
| 33 | p.Glu33Tyr | | | 374 | 322 | 0.98 | 0.98 | | | | | -9.22E+00 | Indeterminate | | | -9.22E+00 | Indeterminate |
| 33 | p.Glu33Cys | | | 784 | 708 | 2.05 | 2.15 | | | | | -3.97E+00 | Neutral | | | -3.97E+00 | Neutral |
| 33 | p.Glu33Trp | | | 1942 | 1609 | 5.08 | 4.88 | | | | | -2.63E-01 | Neutral | | | -2.63E-01 | Neutral |
| 33 | p.Glu33Phe | | | 771 | 678 | 2.02 | 2.06 | | | | | -3.58E+00 | Neutral | | | -3.58E+00 | Neutral |
| 34 | p.Ala34Asn | | | 1048 | 380 | 2.48 | 2.47 | 974 | 864 | 2.39 | 3.01 | -5.06E+00 | Neutral | -9.96E+00 | Indeterminate | -1.15E+01 | Indeterminate |
| 34 | p.Ala34Lys | | | 1978 | 717 | 4.68 | 4.66 | 1977 | 1399 | 4.84 | 4.88 | -1.63E+00 | Neutral | -9.22E-01 | Neutral | -1.08E+00 | Neutral |
| 34 | p.Ala34Thr | | | 1206 | 360 | 2.86 | 2.34 | 1040 | 862 | 2.55 | 3.01 | -1.34E+00 | Neutral | -7.07E+00 | Indeterminate | -5.63E+00 | Neutral |
| 34 | p.Ala34Arg | | | 4462 | 1497 | 10.56 | 9.73 | 4243 | 2972 | 10.39 | 10.37 | -6.89E-02 | Neutral | -7.03E-02 | Neutral | -6.31E-03 | Neutral |
| 34 | p.Ala34Ser | | | 1003 | 246 | 2.37 | 1.60 | 903 | 642 | 2.21 | 2.24 | -4.70E-01 | Neutral | -4.44E+00 | Neutral | -2.77E+00 | Neutral |
| 34 | p.Ala34Ile | | | 1160 | 348 | 2.75 | 2.26 | 1259 | 760 | 3.08 | 2.65 | -1.50E+00 | Neutral | -8.61E-01 | Neutral | -9.65E-01 | Neutral |
| 34 | p.Ala34Met | | | 2790 | 1090 | 6.61 | 7.08 | 2925 | 1910 | 7.16 | 6.66 | -1.25E+00 | Neutral | -1.32E-01 | Neutral | -4.14E-01 | Neutral |
| 34 | p.Ala34His | | | 1987 | 510 | 4.70 | 3.31 | 1486 | 902 | 3.64 | 3.15 | -6.96E-02 | Neutral | -5.80E-01 | Neutral | -1.13E-01 | Neutral |
| 34 | p.Ala34Gln | | | 2756 | 1108 | 6.53 | 7.20 | 2539 | 2187 | 6.22 | 7.63 | -1.56E+00 | Neutral | -1.98E+00 | Neutral | -1.75E+00 | Neutral |
| 34 | p.Ala34Pro | | | 1736 | 907 | 4.11 | 5.89 | 1555 | 1319 | 3.81 | 4.60 | -1.11E+01 | Indeterminate | -4.36E+00 | Neutral | -1.19E+01 | Indeterminate |
| 34 | p.Ala34Leu | | | 2439 | 838 | 5.77 | 5.45 | 2468 | 1808 | 6.05 | 6.31 | -6.70E-01 | Neutral | -6.81E-01 | Neutral | -3.98E-01 | Neutral |
| 34 | p.Ala34Asp | | | 2130 | 868 | 5.04 | 5.64 | 2207 | 1477 | 5.41 | 5.15 | -2.81E+00 | Neutral | -4.33E-01 | Neutral | -1.54E+00 | Neutral |
| 34 | p.Ala34Glu | | | 2272 | 972 | 5.38 | 6.32 | 2212 | 1619 | 5.42 | 5.65 | -3.25E+00 | Neutral | -8.98E-01 | Neutral | -2.19E+00 | Neutral |
| 34 | p.Ala34Ala | Synonymous | | 2329 | 837 | 5.51 | 5.44 | 2131 | 1576 | 5.22 | 5.50 | -1.06E+00 | Neutral | -1.06E+00 | Neutral | -8.18E-01 | Neutral |
| 34 | p.Ala34Gly | | | 1829 | 780 | 4.33 | 5.07 | 1820 | 1092 | 4.46 | 3.81 | -4.56E+00 | Neutral | -2.84E-01 | Neutral | -2.72E+00 | Neutral |
| 34 | p.Ala34Val | | | 1093 | 525 | 2.59 | 3.41 | 1289 | 878 | 3.16 | 3.06 | -1.40E+01 | Indeterminate | -1.87E+00 | Neutral | -1.23E+01 | Indeterminate |
| 34 | p.Ala34Tyr | | | 2781 | 751 | 6.58 | 4.88 | 2256 | 1675 | 5.53 | 5.84 | -3.14E-02 | Neutral | -9.51E-01 | Neutral | -2.33E-01 | Neutral |
| 34 | p.Ala34Cys | | | 2203 | 772 | 5.22 | 5.02 | 2252 | 1303 | 5.52 | 4.55 | -1.01E+00 | Neutral | -8.89E-02 | Neutral | -2.82E-01 | Neutral |
| 34 | p.Ala34Trp | | | 1513 | 537 | 3.58 | 3.49 | 1712 | 1111 | 4.19 | 3.88 | -2.45E+00 | Neutral | -6.83E-01 | Neutral | -1.47E+00 | Neutral |
| 34 | p.Ala34Phe | | | 3520 | 1345 | 8.33 | 8.74 | 3578 | 2302 | 8.76 | 8.03 | -5.90E-01 | Neutral | -4.90E-02 | Neutral | -1.10E-01 | Neutral |
| 35 | p.Gly35Asn | | | 1949 | 799 | 3.50 | 1.78 | 10005 | 6273 | 4.43 | 2.61 | -8.73E+00 | Indeterminate | -1.65E+00 | Neutral | -7.34E+00 | Indeterminate |
| 35 | p.Gly35Lys | | | 2552 | 1260 | 4.58 | 2.80 | 10948 | 8539 | 4.85 | 3.55 | -1.23E+01 | Indeterminate | -4.60E+00 | Neutral | -1.32E+01 | Indeterminate |
| 35 | p.Gly35Thr | | | 2584 | 1647 | 4.63 | 3.66 | 10143 | 10610 | 4.49 | 4.41 | -2.60E+01 | Indeterminate | -1.54E+01 | Indeterminate | -5.32E+01 | Deleterious |
| 35 | p.Gly35Arg | | Neutral | 2733 | 924 | 4.90 | 2.06 | 11916 | 8419 | 5.28 | 3.50 | -2.06E+00 | Neutral | -2.39E+00 | Neutral | -2.42E+00 | Neutral |
| 35 | p.Gly35Ser | | | 2126 | 992 | 3.81 | 2.21 | 8773 | 6513 | 3.89 | 2.71 | -1.25E+01 | Indeterminate | -5.14E+00 | Neutral | -1.39E+01 | Indeterminate |
| 35 | p.Gly35Ile | | | 2090 | 4458 | 3.75 | 9.92 | 9346 | 19604 | 4.14 | 8.15 | -5.32E+01 | Deleterious | -5.32E+01 | Deleterious | -5.32E+01 | Deleterious |
| 35 | p.Gly35Met | | | 3439 | 1998 | 6.17 | 4.44 | 12679 | 12683 | 5.62 | 5.27 | -1.51E+01 | Indeterminate | -1.04E+01 | Indeterminate | -2.13E+01 | Indeterminate |
| 35 | p.Gly35His | | | 2988 | 1047 | 5.36 | 2.33 | 11525 | 8225 | 5.10 | 3.42 | -2.14E+00 | Neutral | -2.69E+00 | Neutral | -2.71E+00 | Neutral |

| | | | | | | | | | | | | | | | | | |
|---|---|---|---|---|---|---|---|---|---|---|---|---|---|---|---|---|---|
| 35 | p.Gly35Gln | | | 4241 | 1991 | 7.61 | 4.43 | 14962 | 14359 | 6.63 | 5.97 | -5.28E+00 | Neutral | -7.21E+00 | Indeterminate | -9.22E+00 | Indeterminate |
| 35 | p.Gly35Pro | | | 2624 | 9100 | 4.71 | 20.24 | 10587 | 31189 | 4.69 | 12.96 | -5.32E+01 | Deleterious | -5.32E+01 | Deleterious | -5.32E+01 | Deleterious |
| 35 | p.Gly35Leu | | | 3145 | 2964 | 5.64 | 6.59 | 12395 | 15692 | 5.49 | 6.52 | -5.32E+01 | Deleterious | -2.23E+01 | Indeterminate | -5.32E+01 | Deleterious |
| 35 | p.Gly35Asp | | | 2568 | 1419 | 4.61 | 3.16 | 10551 | 7765 | 4.67 | 3.23 | -1.75E+01 | Indeterminate | -3.69E+00 | Neutral | -1.72E+01 | Indeterminate |
| 35 | p.Gly35Glu | | Neutral | 2733 | 1146 | 4.90 | 2.55 | 10753 | 9407 | 4.76 | 3.91 | -6.09E+00 | Indeterminate | -7.72E+00 | Indeterminate | -1.04E+01 | Indeterminate |
| 35 | p.Gly35Ala | | | 2285 | 903 | 4.10 | 2.01 | 9298 | 6841 | 4.12 | 2.84 | -6.10E+00 | Indeterminate | -4.52E+00 | Neutral | -7.55E+00 | Indeterminate |
| 35 | p.Gly35Gly | Synonymous | | 1358 | 329 | 2.44 | 0.73 | 5570 | 2694 | 2.47 | 1.12 | -1.06E+00 | Neutral | -1.06E+00 | Neutral | -8.18E-01 | Neutral |
| 35 | p.Gly35Val | | Deleterious | 3854 | 5902 | 6.91 | 13.13 | 15424 | 25699 | 6.83 | 10.68 | -5.32E+01 | Deleterious | -3.73E+01 | Indeterminate | -5.32E+01 | Deleterious |
| 35 | p.Gly35Tyr | | | 2312 | 1427 | 4.15 | 3.17 | 10542 | 8067 | 4.67 | 3.35 | -2.63E+01 | Indeterminate | -4.47E+00 | Neutral | -2.63E+01 | Indeterminate |
| 35 | p.Gly35Cys | | | 3042 | 1561 | 5.46 | 3.47 | 11585 | 9259 | 5.13 | 3.85 | -1.15E+01 | Indeterminate | -4.73E+00 | Neutral | -1.26E+01 | Indeterminate |
| 35 | p.Gly35Trp | | Deleterious | 4083 | 3648 | 7.32 | 8.11 | 15410 | 18194 | 6.83 | 7.56 | -4.10E+01 | Indeterminate | -1.46E+01 | Indeterminate | -5.32E+01 | Deleterious |
| 35 | p.Gly35Phe | | | 3051 | 1446 | 5.47 | 3.22 | 13371 | 10574 | 5.92 | 4.39 | -8.60E+00 | Indeterminate | -3.57E+00 | Neutral | -8.93E+00 | Indeterminate |
| 36 | p.Ala36Asn | | | 5581 | 3345 | 5.14 | 6.08 | | | | | -2.13E+01 | Indeterminate | | | -2.13E+01 | Indeterminate |
| 36 | p.Ala36Lys | | | 8028 | 18657 | 7.40 | 33.90 | | | | | -5.32E+01 | Deleterious | | | -5.32E+01 | Deleterious |
| 36 | p.Ala36Thr | | | 2783 | 1191 | 2.56 | 2.16 | | | | | -1.56E+01 | Indeterminate | | | -1.56E+01 | Indeterminate |
| 36 | p.Ala36Arg | | | 4397 | 1199 | 4.05 | 2.18 | | | | | -8.65E-01 | Neutral | | | -8.65E-01 | Neutral |
| 36 | p.Ala36Ser | | | 4348 | 1450 | 4.01 | 2.64 | | | | | -3.16E+00 | Neutral | | | -3.16E+00 | Neutral |
| 36 | p.Ala36Ile | | | 5237 | 1240 | 4.83 | 2.25 | | | | | -1.36E-01 | Neutral | | | -1.36E-01 | Neutral |
| 36 | p.Ala36Met | | | 6481 | 232 | 5.97 | 0.42 | | | | | 0.00E+00 | Neutral | | | 0.00E+00 | Neutral |
| 36 | p.Ala36His | | | 5535 | 1160 | 5.10 | 2.11 | | | | | -2.28E-02 | Neutral | | | -2.28E-02 | Neutral |
| 36 | p.Ala36Gln | | | 5409 | 1222 | 4.98 | 2.22 | | | | | -6.88E-02 | Neutral | | | -6.88E-02 | Neutral |
| 36 | p.Ala36Pro | | | 8410 | 9367 | 7.75 | 17.02 | | | | | -5.32E+01 | Deleterious | | | -5.32E+01 | Deleterious |
| 36 | p.Ala36Leu | | | 6369 | 2754 | 5.87 | 5.00 | | | | | -5.94E+00 | Indeterminate | | | -5.94E+00 | Indeterminate |
| 36 | p.Ala36Asp | | | 3629 | 1165 | 3.34 | 2.12 | | | | | -3.54E+00 | Neutral | | | -3.54E+00 | Neutral |
| 36 | p.Ala36Glu | | | 7209 | 3345 | 6.64 | 6.08 | | | | | -6.72E+00 | Indeterminate | | | -6.72E+00 | Indeterminate |
| 36 | p.Ala36Ala | Synonymous | | 4309 | 1200 | 3.97 | 2.18 | | | | | -1.06E+00 | Neutral | | | -1.06E+00 | Neutral |
| 36 | p.Ala36Gly | | | 2425 | 903 | 2.23 | 1.64 | | | | | -1.15E+01 | Indeterminate | | | -1.15E+01 | Indeterminate |
| 36 | p.Ala36Val | | | 5770 | 1324 | 5.32 | 2.41 | | | | | -6.40E-02 | Neutral | | | -6.40E-02 | Neutral |
| 36 | p.Ala36Tyr | | | 4470 | 1637 | 4.12 | 2.97 | | | | | -4.78E+00 | Neutral | | | -4.78E+00 | Neutral |
| 36 | p.Ala36Cys | | | 4719 | 296 | 4.35 | 0.54 | | | | | -1.04E-14 | Neutral | | | 0.00E+00 | Neutral |
| 36 | p.Ala36Trp | | | 7861 | 1433 | 7.24 | 2.60 | | | | | -2.83E-04 | Neutral | | | -2.83E-04 | Neutral |
| 36 | p.Ala36Phe | | | 5551 | 1908 | 5.12 | 3.47 | | | | | -2.39E+00 | Neutral | | | -2.39E+00 | Neutral |
| 37 | p.Leu37Asn | | | 15659 | 807 | 7.21 | 4.34 | | | | | -7.48E-02 | Neutral | | | -7.48E-02 | Neutral |
| 37 | p.Leu37Lys | | | 9469 | 805 | 4.36 | 4.33 | | | | | -8.20E+00 | Indeterminate | | | -8.20E+00 | Indeterminate |
| 37 | p.Leu37Thr | | | 7000 | 646 | 3.22 | 3.48 | | | | | -1.54E+01 | Indeterminate | | | -1.54E+01 | Indeterminate |
| 37 | p.Leu37Arg | | | 10261 | 287 | 4.73 | 1.54 | | | | | -2.02E-05 | Neutral | | | -2.02E-05 | Neutral |
| 37 | p.Leu37Ser | | | 7945 | 512 | 3.66 | 2.76 | | | | | -3.05E+00 | Neutral | | | -3.05E+00 | Neutral |
| 37 | p.Leu37Ile | | | 13094 | 1494 | 6.03 | 8.04 | | | | | -1.56E+01 | Indeterminate | | | -1.56E+01 | Indeterminate |
| 37 | p.Leu37Met | | | 11651 | 487 | 5.37 | 2.62 | | | | | -1.66E-02 | Neutral | | | -1.66E-02 | Neutral |
| 37 | p.Leu37His | | | 11818 | 1084 | 5.44 | 5.83 | | | | | -8.27E+00 | Indeterminate | | | -8.27E+00 | Indeterminate |
| 37 | p.Leu37Gln | | | 17769 | 1937 | 8.18 | 10.42 | | | | | -9.51E+00 | Indeterminate | | | -9.51E+00 | Indeterminate |
| 37 | p.Leu37Pro | | | 9322 | 1163 | 4.29 | 6.26 | | | | | -2.77E+01 | Indeterminate | | | -2.77E+01 | Indeterminate |
| 37 | p.Leu37Leu | Synonymous | | 14149 | 931 | 6.52 | 5.01 | | | | | -1.06E+00 | Neutral | | | -1.06E+00 | Neutral |
| 37 | p.Leu37Asp | | | 10884 | 742 | 5.01 | 3.99 | | | | | -2.34E+00 | Neutral | | | -2.34E+00 | Neutral |
| 37 | p.Leu37Glu | | | 9551 | 634 | 4.40 | 3.41 | | | | | -2.57E+00 | Neutral | | | -2.57E+00 | Neutral |
| 37 | p.Leu37Ala | | | 11018 | 534 | 5.08 | 2.87 | | | | | -1.43E-01 | Neutral | | | -1.43E-01 | Neutral |
| 37 | p.Leu37Gly | | | 9306 | 588 | 4.29 | 3.16 | | | | | -2.03E+00 | Neutral | | | -2.03E+00 | Neutral |
| 37 | p.Leu37Val | | | 13136 | 1307 | 6.05 | 7.03 | | | | | -9.83E+00 | Indeterminate | | | -9.83E+00 | Indeterminate |
| 37 | p.Leu37Tyr | | | 8482 | 697 | 3.91 | 3.75 | | | | | -8.28E+00 | Indeterminate | | | -8.28E+00 | Indeterminate |
| 37 | p.Leu37Cys | | | 9322 | 518 | 4.29 | 2.79 | | | | | -8.47E-01 | Neutral | | | -8.47E-01 | Neutral |
| 37 | p.Leu37Trp | | | 6345 | 2856 | 2.92 | 15.37 | | | | | -5.32E+01 | Deleterious | | | -5.32E+01 | Deleterious |
| 37 | p.Leu37Phe | | | 10921 | 555 | 5.03 | 2.99 | | | | | -2.43E-01 | Neutral | | | -2.43E-01 | Neutral |
| 38 | p.Pro38Asn | | | 7663 | 2366 | 4.62 | 2.91 | | | | | -5.32E+01 | Deleterious | | | -5.32E+01 | Deleterious |
| 38 | p.Pro38Lys | | | 6677 | 2439 | 4.03 | 3.00 | | | | | -5.32E+01 | Deleterious | | | -5.32E+01 | Deleterious |
| 38 | p.Pro38Thr | | | 6647 | 881 | 4.01 | 1.08 | | | | | -6.12E+00 | Indeterminate | | | -6.12E+00 | Indeterminate |
| 38 | p.Pro38Arg | | | 6252 | 3126 | 3.77 | 3.84 | | | | | -5.32E+01 | Deleterious | | | -5.32E+01 | Deleterious |
| 38 | p.Pro38Ser | | | 6013 | 1033 | 3.63 | 1.27 | | | | | -1.73E+01 | Indeterminate | | | -1.73E+01 | Indeterminate |
| 38 | p.Pro38Ile | | | 6138 | 1077 | 3.70 | 1.32 | | | | | -1.81E+01 | Indeterminate | | | -1.81E+01 | Indeterminate |
| 38 | p.Pro38Met | | | 8518 | 1945 | 5.14 | 2.39 | | | | | -2.77E+01 | Indeterminate | | | -2.77E+01 | Indeterminate |
| 38 | p.Pro38His | | | 5342 | 2012 | 3.22 | 2.47 | | | | | -5.32E+01 | Deleterious | | | -5.32E+01 | Deleterious |
| 38 | p.Pro38Gln | | | 17977 | 7277 | 10.84 | 8.94 | | | | | -5.32E+01 | Deleterious | | | -5.32E+01 | Deleterious |
| 38 | p.Pro38Pro | Synonymous | | 7226 | 717 | 4.36 | 0.88 | | | | | -1.06E+00 | Neutral | | | -1.06E+00 | Neutral |
| 38 | p.Pro38Leu | | | 10269 | 3279 | 6.19 | 4.03 | | | | | -5.30E+01 | Indeterminate | | | -3.32E+01 | Indeterminate |
| 38 | p.Pro38Asp | | | 7287 | 14782 | 4.39 | 18.15 | | | | | -5.32E+01 | Deleterious | | | -5.32E+01 | Deleterious |
| 38 | p.Pro38Glu | | | 14248 | 3748 | 8.59 | 4.60 | | | | | -2.57E+01 | Indeterminate | | | -2.57E+01 | Indeterminate |
| 38 | p.Pro38Ala | | | 6236 | 3456 | 3.76 | 4.24 | | | | | -5.32E+01 | Deleterious | | | -5.32E+01 | Deleterious |
| 38 | p.Pro38Gly | | | 8191 | 804 | 4.94 | 0.99 | | | | | -7.14E-01 | Neutral | | | -7.14E-01 | Neutral |
| 38 | p.Pro38Val | | | 10026 | 1016 | 6.04 | 1.25 | | | | | -5.34E-01 | Neutral | | | -5.34E-01 | Neutral |
| 38 | p.Pro38Tyr | | | 4282 | 1445 | 2.58 | 1.77 | | | | | -5.32E+01 | Deleterious | | | -5.32E+01 | Deleterious |
| 38 | p.Pro38Cys | | | 8165 | 611 | 4.92 | 0.75 | | | | | -4.22E-02 | Neutral | | | -4.22E-02 | Neutral |
| 38 | p.Pro38Trp | | | 7975 | 24617 | 4.81 | 30.23 | | | | | -5.32E+01 | Deleterious | | | -5.32E+01 | Deleterious |
| 38 | p.Pro38Phe | | | 10730 | 4800 | 6.47 | 5.89 | | | | | -5.32E+01 | Deleterious | | | -5.32E+01 | Deleterious |
| 39 | p.Asn39Asn | Synonymous | | 11025 | 1321 | 4.88 | 1.95 | | | | | -1.06E+00 | Neutral | | | -1.06E+00 | Neutral |
| 39 | p.Asn39Lys | | | 12759 | 3256 | 5.65 | 4.81 | | | | | -2.29E+01 | Indeterminate | | | -2.29E+01 | Indeterminate |
| 39 | p.Asn39Thr | | | 9531 | 725 | 4.22 | 1.07 | | | | | -1.29E-02 | Neutral | | | -1.29E-02 | Neutral |
| 39 | p.Asn39Arg | | | 7919 | 1061 | 3.51 | 1.57 | | | | | -3.95E+00 | Neutral | | | -3.95E+00 | Neutral |
| 39 | p.Asn39Ser | | | 8858 | 1238 | 3.92 | 1.83 | | | | | -4.05E+00 | Neutral | | | -4.05E+00 | Neutral |
| 39 | p.Asn39Ile | | | 10639 | 1845 | 4.71 | 2.73 | | | | | -7.95E+00 | Indeterminate | | | -7.95E+00 | Indeterminate |
| 39 | p.Asn39Met | | | 12667 | 3339 | 5.61 | 4.93 | | | | | -2.52E+01 | Indeterminate | | | -2.52E+01 | Indeterminate |
| 39 | p.Asn39His | | | 11236 | 950 | 4.98 | 1.40 | | | | | -2.72E-02 | Neutral | | | -2.72E-02 | Neutral |
| 39 | p.Asn39Gln | | | 8347 | 932 | 3.70 | 1.38 | | | | | -1.24E+00 | Neutral | | | -1.24E+00 | Neutral |
| 39 | p.Asn39Pro | | | 9695 | 16800 | 4.30 | 24.82 | | | | | -5.32E+01 | Deleterious | | | -5.32E+01 | Deleterious |
| 39 | p.Asn39Leu | | | 13886 | 6932 | 6.15 | 10.24 | | | | | -5.32E+01 | Deleterious | | | -5.32E+01 | Deleterious |
| 39 | p.Asn39Asp | | | 15016 | 1873 | 6.65 | 2.77 | | | | | -6.59E-01 | Neutral | | | -6.59E-01 | Neutral |
| 39 | p.Asn39Glu | | | 15471 | 2195 | 6.85 | 3.24 | | | | | -1.56E+00 | Neutral | | | -1.56E+00 | Neutral |
| 39 | p.Asn39Ala | | | 8785 | 1365 | 3.89 | 2.02 | | | | | -6.60E+00 | Indeterminate | | | -6.60E+00 | Indeterminate |
| 39 | p.Asn39Gly | | | 15618 | 1831 | 6.92 | 2.70 | | | | | -3.42E-01 | Neutral | | | -3.42E-01 | Neutral |
| 39 | p.Asn39Val | | | 12208 | 1947 | 5.41 | 2.88 | | | | | -4.58E+00 | Neutral | | | -4.58E+00 | Neutral |
| 39 | p.Asn39Tyr | | | 8959 | 952 | 3.97 | 1.41 | | | | | -7.21E-01 | Neutral | | | -7.21E-01 | Neutral |
| 39 | p.Asn39Cys | | | 13589 | 7404 | 6.02 | 10.94 | | | | | -5.32E+01 | Deleterious | | | -5.32E+01 | Deleterious |
| 39 | p.Asn39Trp | | | 9837 | 1931 | 4.36 | 2.85 | | | | | -1.36E+01 | Indeterminate | | | -1.36E+01 | Indeterminate |
| 39 | p.Asn39Phe | | | 9654 | 9801 | 4.28 | 14.48 | | | | | -5.32E+01 | Deleterious | | | -5.32E+01 | Deleterious |
| 40 | p.Ala40Asn | | | 17799 | 1463 | 5.87 | 4.98 | | | | | -6.02E+00 | Indeterminate | | | -6.02E+00 | Indeterminate |
| 40 | p.Ala40Lys | | | 21626 | 1443 | 7.13 | 4.91 | | | | | -1.40E+00 | Neutral | | | -1.40E+00 | Neutral |
| 40 | p.Ala40Thr | | | 19138 | 8921 | 6.31 | 30.36 | | | | | -5.32E+01 | Deleterious | | | -5.32E+01 | Deleterious |
| 40 | p.Ala40Arg | | | 12717 | 763 | 4.19 | 2.60 | | | | | -2.18E+00 | Neutral | | | -2.18E+00 | Neutral |
| 40 | p.Ala40Ser | | | 7528 | 329 | 2.48 | 1.12 | | | | | -8.34E-01 | Neutral | | | -8.34E-01 | Neutral |
| 40 | p.Ala40Ile | | | 9646 | 648 | 3.18 | 2.21 | | | | | -6.06E+00 | Indeterminate | | | -6.06E+00 | Indeterminate |
| 40 | p.Ala40Met | | | 12826 | 464 | 4.23 | 1.58 | | | | | -1.44E-02 | Neutral | | | -1.44E-02 | Neutral |
| 40 | p.Ala40His | | | 10373 | 460 | 3.42 | 1.57 | | | | | -3.86E-01 | Neutral | | | -3.86E-01 | Neutral |
| 40 | p.Ala40Gln | | | 12433 | 454 | 4.10 | 1.54 | | | | | -1.93E-02 | Neutral | | | -1.93E-02 | Neutral |
| 40 | p.Ala40Pro | | | 13362 | 1203 | 4.41 | 4.09 | | | | | -1.22E+01 | Indeterminate | | | -1.22E+01 | Indeterminate |
| 40 | p.Ala40Leu | | | 14549 | 3440 | 4.80 | 11.71 | | | | | -5.32E+01 | Deleterious | | | -5.32E+01 | Deleterious |
| 40 | p.Ala40Asp | | | 12913 | 1289 | 4.26 | 4.39 | | | | | -1.76E+01 | Indeterminate | | | -1.76E+01 | Indeterminate |
| 40 | p.Ala40Glu | | | 22310 | 1055 | 7.36 | 3.59 | | | | | -4.71E-02 | Neutral | | | -4.71E-02 | Neutral |
| 40 | p.Ala40Ala | Synonymous | | 17869 | 1075 | 5.89 | 3.66 | | | | | -1.06E+00 | Neutral | | | -1.06E+00 | Neutral |
| 40 | p.Ala40Gly | | | 20764 | 1292 | 6.85 | 4.40 | | | | | -9.43E-01 | Neutral | | | -9.43E-01 | Neutral |
| 40 | p.Ala40Val | | | 17468 | 1452 | 5.76 | 4.94 | | | | | -6.48E+00 | Indeterminate | | | -6.48E+00 | Indeterminate |
| 40 | p.Ala40Tyr | | | 14320 | 633 | 4.72 | 2.15 | | | | | -1.25E-01 | Neutral | | | -1.25E-01 | Neutral |
| 40 | p.Ala40Cys | | | 15434 | 936 | 5.09 | 3.19 | | | | | -1.56E+00 | Neutral | | | -1.56E+00 | Neutral |
| 40 | p.Ala40Trp | | | 12309 | 863 | 4.06 | 2.94 | | | | | -5.12E+00 | Neutral | | | -5.12E+00 | Neutral |
| 40 | p.Ala40Phe | | | 17875 | 1204 | 5.89 | 4.10 | | | | | -2.21E+00 | Neutral | | | -2.21E+00 | Neutral |
| 41 | p.Pro41Asn | | | 3018 | 760 | 4.29 | 3.16 | | | | | -3.89E-01 | Neutral | | | -3.89E-01 | Neutral |
| 41 | p.Pro41Lys | | | 2802 | 980 | 3.98 | 4.07 | | | | | -4.11E+00 | Neutral | | | -4.11E+00 | Neutral |
| 41 | p.Pro41Thr | | | 2775 | 782 | 3.95 | 3.25 | | | | | -1.20E+00 | Neutral | | | -1.20E+00 | Neutral |
| 41 | p.Pro41Arg | | | 2832 | 1777 | 4.03 | 7.39 | | | | | -3.01E+01 | Indeterminate | | | -2.99E+01 | Indeterminate |
| 41 | p.Pro41Ser | | | 4466 | 1181 | 6.35 | 4.91 | | | | | -1.74E-01 | Neutral | | | -1.74E-01 | Neutral |
| 41 | p.Pro41Ile | | | 2598 | 1141 | 3.69 | 4.74 | | | | | -1.15E+01 | Indeterminate | | | -1.15E+01 | Indeterminate |
| 41 | p.Pro41Met | | | 4122 | 874 | 5.86 | 3.63 | | | | | -1.44E-02 | Neutral | | | -1.44E-02 | Neutral |
| 41 | p.Pro41His | | | 2358 | 1155 | 3.35 | 4.80 | | | | | -1.81E+01 | Indeterminate | | | -1.81E+01 | Indeterminate |
| 41 | p.Pro41Gln | | Neutral | 3840 | 1354 | 5.46 | 5.63 | | | | | -2.48E+00 | Neutral | | | -2.48E+00 | Neutral |
| 41 | p.Pro41Pro | Synonymous | | 2669 | 730 | 3.79 | 3.03 | | | | | -1.06E+00 | Neutral | | | -1.06E+00 | Neutral |
| 41 | p.Pro41Leu | | | 3638 | 786 | 5.17 | 3.27 | | | | | -3.41E-02 | Neutral | | | -3.41E-02 | Neutral |
| 41 | p.Pro41Asp | | | 3619 | 1263 | 5.15 | 5.25 | | | | | -2.62E+00 | Neutral | | | -2.62E+00 | Neutral |
| 41 | p.Pro41Glu | | | 3875 | 956 | 5.51 | 3.97 | | | | | -1.36E-01 | Neutral | | | -1.36E-01 | Neutral |
| 41 | p.Pro41Ala | | | 4397 | 1208 | 6.25 | 5.02 | | | | | -2.71E-01 | Neutral | | | -2.71E-01 | Neutral |
| 41 | p.Pro41Gly | | | 2512 | 730 | 3.57 | 3.03 | | | | | -1.82E+00 | Neutral | | | -1.82E+00 | Neutral |
| 41 | p.Pro41Val | | | 5452 | 1628 | 7.75 | 6.77 | | | | | -3.05E-01 | Neutral | | | -3.05E-01 | Neutral |
| 41 | p.Pro41Tyr | | | 4042 | 2317 | 5.75 | 9.63 | | | | | -1.69E+01 | Indeterminate | | | -1.69E+01 | Indeterminate |
| 41 | p.Pro41Cys | | | 3683 | 1331 | 5.24 | 5.53 | | | | | -3.07E+00 | Neutral | | | -3.07E+00 | Neutral |
| 41 | p.Pro41Trp | | | 2884 | 1129 | 4.10 | 4.69 | | | | | -6.53E+00 | Indeterminate | | | -6.53E+00 | Indeterminate |
| 41 | p.Pro41Phe | | | 4753 | 1980 | 6.76 | 8.23 | | | | | -4.10E+00 | Neutral | | | -4.10E+00 | Neutral |
| 42 | p.Asn42Asn | Synonymous | | 8645 | 341 | 4.30 | 0.55 | | | | | -1.06E+00 | Neutral | | | -1.06E+00 | Neutral |
| 42 | p.Asn42Lys | | | 9397 | 4647 | 4.67 | 7.54 | | | | | -5.32E+01 | Deleterious | | | -5.32E+01 | Deleterious |
| 42 | p.Asn42Thr | | | 10798 | 564 | 5.37 | 0.92 | | | | | -3.68E+00 | Neutral | | | -3.68E+00 | Neutral |
| 42 | p.Asn42Arg | | | 13173 | 6054 | 6.55 | 9.82 | | | | | -5.32E+01 | Deleterious | | | -5.32E+01 | Deleterious |
| 42 | p.Asn42Ser | | | 9213 | 444 | 4.58 | 0.72 | | | | | -3.19E+00 | Neutral | | | -3.19E+00 | Neutral |
| 42 | p.Asn42Ile | | | 10911 | 4981 | 5.42 | 8.08 | | | | | -5.32E+01 | Deleterious | | | -5.32E+01 | Deleterious |
| 42 | p.Asn42Met | | | 8586 | 3308 | 4.27 | 5.37 | | | | | -5.32E+01 | Deleterious | | | -5.32E+01 | Deleterious |

| | | | | | | | | | | | | | | | | |
|---|---|---|---|---|---|---|---|---|---|---|---|---|---|---|---|---|
| 42 | p.Asn42His | | 9638 | 2787 | 4.79 | 4.52 | | | | | -5.32E+01 | Deleterious | | | -5.32E+01 | Deleterious |
| 42 | p.Asn42Gln | | 10388 | 2910 | 5.16 | 4.72 | | | | | -5.32E+01 | Deleterious | | | -5.32E+01 | Deleterious |
| 42 | p.Asn42Pro | | 6267 | 2676 | 3.11 | 4.34 | | | | | -5.32E+01 | Deleterious | | | -5.32E+01 | Deleterious |
| 42 | p.Asn42Leu | | 9292 | 4188 | 4.62 | 6.79 | | | | | -5.32E+01 | Deleterious | | | -5.32E+01 | Deleterious |
| 42 | p.Asn42Asp | | 9064 | 576 | 4.50 | 0.93 | | | | | -1.07E+01 | Indeterminate | | | -1.07E+01 | Indeterminate |
| 42 | p.Asn42Glu | | 12901 | 6353 | 6.41 | 10.31 | | | | | -5.32E+01 | Deleterious | | | -5.32E+01 | Deleterious |
| 42 | p.Asn42Ala | | 7254 | 328 | 3.61 | 0.53 | | | | | -3.46E+00 | Neutral | | | -3.46E+00 | Neutral |
| 42 | p.Asn42Gly | | 12142 | 451 | 6.03 | 0.73 | | | | | -2.42E-01 | Neutral | | | -2.42E-01 | Neutral |
| 42 | p.Asn42Val | | 13261 | 5223 | 6.59 | 8.47 | | | | | -5.32E+01 | Deleterious | | | -5.32E+01 | Deleterious |
| 42 | p.Asn42Tyr | | 9832 | 4604 | 4.89 | 7.47 | | | | | -5.32E+01 | Deleterious | | | -5.32E+01 | Deleterious |
| 42 | p.Asn42Cys | | 10625 | 622 | 5.28 | 1.01 | | | | | -6.36E+00 | Indeterminate | | | -6.36E+00 | Indeterminate |
| 42 | p.Asn42Trp | | 8933 | 5163 | 4.44 | 8.38 | | | | | -5.32E+01 | Deleterious | | | -5.32E+01 | Deleterious |
| 42 | p.Asn42Phe | | 10895 | 5419 | 5.41 | 8.79 | | | | | -5.32E+01 | Deleterious | | | -5.32E+01 | Deleterious |
| 43 | p.Ser43Asn | | 8865 | 6639 | 3.69 | 3.91 | | | | | -3.02E+00 | Neutral | | | -3.02E+00 | Neutral |
| 43 | p.Ser43Lys | | 16747 | 11835 | 6.97 | 6.97 | | | | | -4.78E-01 | Neutral | | | -4.78E-01 | Neutral |
| 43 | p.Ser43Thr | | 14488 | 9553 | 6.03 | 5.62 | | | | | -3.96E-01 | Neutral | | | -3.96E-01 | Neutral |
| 43 | p.Ser43Arg | | 16871 | 11411 | 7.02 | 6.72 | | | | | -3.13E-01 | Neutral | | | -3.13E-01 | Neutral |
| 43 | p.Ser43Ser | Synonymous | 11014 | 7488 | 4.58 | 4.41 | | | | | -1.06E+00 | Neutral | | | -1.06E+00 | Neutral |
| 43 | p.Ser43Ile | | 7943 | 6763 | 3.30 | 3.98 | | | | | -6.55E+00 | Indeterminate | | | -6.55E+00 | Indeterminate |
| 43 | p.Ser43Met | | 15797 | 11730 | 6.57 | 6.90 | | | | | -8.40E-01 | Neutral | | | -8.40E-01 | Neutral |
| 43 | p.Ser43His | | 16521 | 11177 | 6.87 | 6.58 | | | | | -3.35E-01 | Neutral | | | -3.35E-01 | Neutral |
| 43 | p.Ser43Gln | | 17314 | 11767 | 7.20 | 6.93 | | | | | -3.01E-01 | Neutral | | | -3.01E-01 | Neutral |
| 43 | p.Ser43Pro | | 12527 | 8469 | 5.21 | 4.98 | | | | | -7.37E-01 | Neutral | | | -7.37E-01 | Neutral |
| 43 | p.Ser43Leu | | 10449 | 7515 | 4.35 | 4.42 | | | | | -1.74E+00 | Neutral | | | -1.74E+00 | Neutral |
| 43 | p.Ser43Asp | | 11537 | 8545 | 4.80 | 5.03 | | | | | -1.70E+00 | Neutral | | | -1.70E+00 | Neutral |
| 43 | p.Ser43Glu | | 14597 | 10128 | 6.07 | 5.96 | | | | | -6.03E-01 | Neutral | | | -6.03E-01 | Neutral |
| 43 | p.Ser43Ala | | 11830 | 7793 | 4.92 | 4.59 | | | | | -6.98E-01 | Neutral | | | -6.98E-01 | Neutral |
| 43 | p.Ser43Gly | | 4691 | 3009 | 1.95 | 1.77 | | | | | -4.02E+00 | Neutral | | | -4.02E+00 | Neutral |
| 43 | p.Ser43Val | | 11779 | 8766 | 4.90 | 5.16 | | | | | -1.68E+00 | Neutral | | | -1.68E+00 | Neutral |
| 43 | p.Ser43Tyr | | 9565 | 6955 | 3.98 | 4.09 | | | | | -2.22E+00 | Neutral | | | -2.22E+00 | Neutral |
| 43 | p.Ser43Cys | | 5091 | 3662 | 2.12 | 2.16 | | | | | -6.00E+00 | Indeterminate | | | -6.00E+00 | Indeterminate |
| 43 | p.Ser43Trp | | 10965 | 8262 | 4.56 | 4.86 | | | | | -2.10E+00 | Neutral | | | -2.10E+00 | Neutral |
| 43 | p.Ser43Phe | | 11776 | 8442 | 4.90 | 4.97 | | | | | -1.32E+00 | Neutral | | | -1.32E+00 | Neutral |
| 44 | p.Tyr44Asn | | 7443 | 4844 | 6.01 | 6.02 | | | | | -3.69E+00 | Neutral | | | -3.69E+00 | Neutral |
| 44 | p.Tyr44Lys | | 4300 | 2790 | 3.47 | 3.47 | | | | | -8.10E+00 | Indeterminate | | | -8.10E+00 | Indeterminate |
| 44 | p.Tyr44Thr | | 7466 | 3963 | 6.03 | 4.92 | | | | | -1.08E+00 | Neutral | | | -1.08E+00 | Neutral |
| 44 | p.Tyr44Arg | | 4924 | 2723 | 3.98 | 3.38 | | | | | -3.23E+00 | Neutral | | | -3.23E+00 | Neutral |
| 44 | p.Tyr44Ser | | 6073 | 3554 | 4.90 | 4.42 | | | | | -3.02E+00 | Neutral | | | -3.02E+00 | Neutral |
| 44 | p.Tyr44Ile | | 5380 | 3030 | 4.34 | 3.76 | | | | | -3.05E+00 | Neutral | | | -3.05E+00 | Neutral |
| 44 | p.Tyr44Met | | 5878 | 3444 | 4.75 | 4.28 | | | | | -3.22E+00 | Neutral | | | -3.22E+00 | Neutral |
| 44 | p.Tyr44His | | 8889 | 5272 | 7.18 | 6.55 | | | | | -1.55E+00 | Neutral | | | -1.55E+00 | Neutral |
| 44 | p.Tyr44Gln | | 2488 | 1711 | 2.01 | 2.13 | | | | | -1.82E+01 | Indeterminate | | | -1.82E+01 | Indeterminate |
| 44 | p.Tyr44Pro | | 6918 | 8564 | 5.59 | 10.64 | | | | | -3.52E+01 | Indeterminate | | | -3.29E+01 | Indeterminate |
| 44 | p.Tyr44Leu | | 6515 | 3822 | 5.26 | 4.75 | | | | | -2.69E+00 | Neutral | | | -2.69E+00 | Neutral |
| 44 | p.Tyr44Asp | | 5508 | 3671 | 4.45 | 4.56 | | | | | -6.48E+00 | Indeterminate | | | -6.48E+00 | Indeterminate |
| 44 | p.Tyr44Glu | | 4288 | 3217 | 3.46 | 4.00 | | | | | -1.36E+01 | Indeterminate | | | -1.36E+01 | Indeterminate |
| 44 | p.Tyr44Ala | | 4861 | 4765 | 3.92 | 5.92 | | | | | -2.63E+01 | Indeterminate | | | -2.63E+01 | Indeterminate |
| 44 | p.Tyr44Gly | | 4818 | 3568 | 3.89 | 4.43 | | | | | -1.14E+01 | Indeterminate | | | -1.14E+01 | Indeterminate |
| 44 | p.Tyr44Val | | 11028 | 6477 | 8.90 | 8.05 | | | | | -8.77E-01 | Neutral | | | -8.77E-01 | Neutral |
| 44 | p.Tyr44Tyr | Synonymous | 7815 | 4201 | 6.31 | 5.22 | | | | | -1.06E+00 | Neutral | | | -1.06E+00 | Neutral |
| 44 | p.Tyr44Cys | | 5174 | 2688 | 4.18 | 3.34 | | | | | -2.08E+00 | Neutral | | | -2.08E+00 | Neutral |
| 44 | p.Tyr44Trp | | 7945 | 4712 | 6.41 | 5.85 | | | | | -1.96E+00 | Neutral | | | -1.96E+00 | Neutral |
| 44 | p.Tyr44Phe | | 6154 | 3465 | 4.97 | 4.31 | | | | | -2.38E+00 | Neutral | | | -2.38E+00 | Neutral |
| 45 | p.Gly45Asn | | 9066 | 190 | 4.25 | 1.00 | 4002 | 2167 | 4.05 | 3.80 | -3.62E-02 | Neutral | -3.51E+00 | Neutral | -1.75E+00 | Neutral |
| 45 | p.Gly45Lys | | 10852 | 426 | 5.08 | 2.25 | 4963 | 2650 | 5.02 | 4.65 | -4.19E+00 | Neutral | -2.20E+00 | Neutral | -3.95E+00 | Neutral |
| 45 | p.Gly45Thr | | 8428 | 159 | 3.95 | 0.84 | 3637 | 1917 | 3.68 | 3.36 | -1.12E-02 | Neutral | -3.59E+00 | Neutral | -1.80E+00 | Neutral |
| 45 | p.Gly45Arg | | 13226 | 375 | 6.19 | 1.98 | 5999 | 3130 | 6.07 | 5.49 | -2.82E-01 | Neutral | -1.27E+00 | Neutral | -5.00E-01 | Neutral |
| 45 | p.Gly45Ser | | 13865 | 787 | 6.49 | 4.15 | 6599 | 3360 | 6.68 | 5.89 | -1.31E+01 | Indeterminate | -8.49E-01 | Neutral | -1.05E+01 | Indeterminate |
| 45 | p.Gly45Ile | | 11823 | 669 | 5.54 | 3.53 | 6125 | 3452 | 6.20 | 6.05 | -1.54E+01 | Indeterminate | -2.00E+00 | Neutral | -1.37E+01 | Indeterminate |
| 45 | p.Gly45Met | | 11011 | 402 | 5.16 | 2.12 | 5143 | 2690 | 5.21 | 4.72 | -2.82E+00 | Neutral | -1.81E+00 | Neutral | -2.56E+00 | Neutral |
| 45 | p.Gly45His | | 9932 | 322 | 4.65 | 1.70 | 4752 | 2496 | 4.81 | 4.38 | -1.72E+00 | Neutral | -2.17E+00 | Neutral | -2.01E+00 | Neutral |
| 45 | p.Gly45Gln | | 10648 | 224 | 4.99 | 1.18 | 5023 | 2411 | 5.08 | 4.23 | -1.80E-02 | Neutral | -1.07E+00 | Neutral | -2.78E-01 | Neutral |
| 45 | p.Gly45Pro | | 8515 | 12301 | 3.99 | 64.90 | 3482 | 7178 | 3.52 | 12.59 | -5.32E+01 | Deleterious | -5.32E+01 | Deleterious | -5.32E+01 | Deleterious |
| 45 | p.Gly45Leu | | 13995 | 472 | 6.55 | 2.49 | 6319 | 3279 | 6.40 | 5.75 | -1.04E+00 | Neutral | -1.09E+00 | Neutral | -8.21E-01 | Neutral |
| 45 | p.Gly45Asp | | 11282 | 237 | 5.28 | 1.25 | 5537 | 2729 | 5.60 | 4.79 | -1.32E-02 | Neutral | -1.03E+00 | Neutral | -2.57E-01 | Neutral |
| 45 | p.Gly45Glu | | 10953 | 350 | 5.13 | 1.85 | 5234 | 2644 | 5.30 | 4.64 | -1.26E+00 | Neutral | -1.39E+00 | Neutral | -1.15E+00 | Neutral |
| 45 | p.Gly45Ala | | 8210 | 153 | 3.84 | 0.81 | 3412 | 1811 | 3.45 | 3.18 | -1.06E-02 | Neutral | -4.13E+00 | Neutral | -2.19E+00 | Neutral |
| 45 | p.Gly45Gly | Synonymous | 9184 | 270 | 4.30 | 1.42 | 4688 | 2196 | 4.74 | 3.85 | -1.06E+00 | Neutral | -1.06E+00 | Neutral | -8.18E-01 | Neutral |
| 45 | p.Gly45Val | | 8564 | 481 | 4.01 | 2.54 | 4028 | 2387 | 4.08 | 4.19 | -2.07E+01 | Indeterminate | -5.33E+00 | Neutral | -2.18E+01 | Indeterminate |
| 45 | p.Gly45Tyr | | 8720 | 186 | 4.08 | 0.98 | 3827 | 2257 | 3.87 | 3.96 | -5.42E-02 | Neutral | -5.62E+00 | Neutral | -3.37E+00 | Neutral |
| 45 | p.Gly45Cys | | 11799 | 295 | 5.53 | 1.56 | 5280 | 2701 | 5.34 | 4.74 | -1.11E-01 | Neutral | -1.48E+00 | Neutral | -5.20E-01 | Neutral |
| 45 | p.Gly45Trp | | 9020 | 275 | 4.22 | 1.45 | 3974 | 2313 | 4.02 | 4.06 | -1.42E+00 | Neutral | -5.01E+00 | Neutral | -3.99E+00 | Neutral |
| 45 | p.Gly45Phe | | 14446 | 381 | 6.77 | 2.01 | 6777 | 3246 | 6.86 | 5.69 | -9.41E-02 | Neutral | -4.79E-01 | Neutral | -9.06E-02 | Neutral |
| 46 | p.Arg46Asn | | 12380 | 531 | 5.08 | 1.57 | | | | | -7.10E-01 | Neutral | | | -7.10E-01 | Neutral |
| 46 | p.Arg46Lys | | 9730 | 361 | 3.99 | 1.07 | | | | | -4.08E-01 | Neutral | | | -4.08E-01 | Neutral |
| 46 | p.Arg46Thr | | 16955 | 2071 | 6.96 | 6.12 | | | | | -3.64E+01 | Indeterminate | | | -3.31E+01 | Indeterminate |
| 46 | p.Arg46Arg | Synonymous | 13782 | 646 | 5.65 | 1.91 | | | | | -1.06E+00 | Neutral | | | -1.06E+00 | Neutral |
| 46 | p.Arg46Ser | | 13112 | 523 | 5.38 | 1.54 | | | | | -3.19E-01 | Neutral | | | -3.19E-01 | Neutral |
| 46 | p.Arg46Ile | | 17693 | 2934 | 7.26 | 8.67 | | | | | -5.32E+01 | Deleterious | | | -5.32E+01 | Deleterious |
| 46 | p.Arg46Met | | 7746 | 476 | 3.18 | 1.41 | | | | | -1.07E+01 | Indeterminate | | | -1.07E+01 | Indeterminate |
| 46 | p.Arg46His | | 8963 | 333 | 3.68 | 0.98 | | | | | -5.25E-01 | Neutral | | | -5.25E-01 | Neutral |
| 46 | p.Arg46Gln | | 9746 | 704 | 4.00 | 2.08 | | | | | -1.43E+01 | Indeterminate | | | -1.43E+01 | Indeterminate |
| 46 | p.Arg46Pro | | 18485 | 15419 | 7.58 | 45.54 | | | | | -5.32E+01 | Deleterious | | | -5.32E+01 | Deleterious |
| 46 | p.Arg46Leu | | 16241 | 1098 | 6.66 | 3.24 | | | | | -5.97E+00 | Indeterminate | | | -5.97E+00 | Indeterminate |
| 46 | p.Arg46Asp | | 8698 | 2412 | 3.57 | 7.12 | | | | | -5.32E+01 | Deleterious | | | -5.32E+01 | Deleterious |
| 46 | p.Arg46Glu | | 7379 | 1044 | 3.03 | 3.08 | | | | | -5.32E+01 | Deleterious | | | -5.32E+01 | Deleterious |
| 46 | p.Arg46Ala | | 9496 | 620 | 3.90 | 1.83 | | | | | -1.04E+01 | Indeterminate | | | -1.04E+01 | Indeterminate |
| 46 | p.Arg46Gly | | 12625 | 802 | 5.18 | 2.37 | | | | | -6.54E+00 | Indeterminate | | | -6.54E+00 | Indeterminate |
| 46 | p.Arg46Val | | 8953 | 791 | 3.67 | 2.34 | | | | | -2.76E+01 | Indeterminate | | | -2.76E+01 | Indeterminate |
| 46 | p.Arg46Tyr | | 14312 | 1176 | 5.87 | 3.47 | | | | | -1.43E+01 | Indeterminate | | | -1.43E+01 | Indeterminate |
| 46 | p.Arg46Cys | | 14267 | 907 | 5.85 | 2.68 | | | | | -5.51E+00 | Neutral | | | -5.51E+00 | Neutral |
| 46 | p.Arg46Trp | | 11216 | 426 | 4.60 | 1.26 | | | | | -3.27E-01 | Neutral | | | -3.27E-01 | Neutral |
| 46 | p.Arg46Phe | | 11963 | 581 | 4.91 | 1.72 | | | | | -1.84E+00 | Neutral | | | -1.84E+00 | Neutral |
| 47 | p.Arg47Asn | | 4828 | 1429 | 3.10 | 2.33 | | | | | -2.20E+00 | Neutral | | | -2.20E+00 | Neutral |
| 47 | p.Arg47Lys | | 6158 | 1982 | 3.96 | 3.24 | | | | | -2.24E+00 | Neutral | | | -2.24E+00 | Neutral |
| 47 | p.Arg47Thr | | 7976 | 2557 | 5.12 | 4.18 | | | | | -1.27E+00 | Neutral | | | -1.27E+00 | Neutral |
| 47 | p.Arg47Arg | Synonymous | 8947 | 2906 | 5.75 | 4.75 | | | | | -1.06E+00 | Neutral | | | -1.06E+00 | Neutral |
| 47 | p.Arg47Ser | | 6416 | 2760 | 4.12 | 4.51 | | | | | -8.25E+00 | Indeterminate | | | -8.25E+00 | Indeterminate |
| 47 | p.Arg47Ile | | 6040 | 2038 | 3.88 | 3.33 | | | | | -3.02E+00 | Neutral | | | -3.02E+00 | Neutral |
| 47 | p.Arg47Met | | 8024 | 2754 | 5.15 | 4.50 | | | | | -1.94E+00 | Neutral | | | -1.94E+00 | Neutral |
| 47 | p.Arg47His | | 7289 | 2516 | 4.68 | 4.11 | | | | | -2.42E+00 | Neutral | | | -2.42E+00 | Neutral |
| 47 | p.Arg47Gln | | 7598 | 3704 | 4.88 | 6.05 | | | | | -1.06E+01 | Indeterminate | | | -1.06E+01 | Indeterminate |
| 47 | p.Arg47Pro | | 8480 | 4396 | 5.45 | 7.18 | | | | | -1.16E+01 | Indeterminate | | | -1.16E+01 | Indeterminate |
| 47 | p.Arg47Leu | | 8455 | 2369 | 5.43 | 3.87 | | | | | -3.70E-01 | Neutral | | | -3.70E-01 | Neutral |
| 47 | p.Arg47Asp | | 6458 | 7409 | 4.15 | 12.10 | | | | | -5.32E+01 | Deleterious | | | -5.32E+01 | Deleterious |
| 47 | p.Arg47Glu | | 9515 | 3930 | 6.11 | 6.42 | | | | | -3.92E+00 | Neutral | | | -3.92E+00 | Neutral |
| 47 | p.Arg47Ala | | 5555 | 1795 | 3.57 | 2.93 | | | | | -2.77E+00 | Neutral | | | -2.77E+00 | Neutral |
| 47 | p.Arg47Gly | | 7345 | 3399 | 4.72 | 5.55 | | | | | -9.18E+00 | Indeterminate | | | -9.18E+00 | Indeterminate |
| 47 | p.Arg47Val | | 7749 | 2449 | 4.98 | 4.00 | | | | | -1.23E+00 | Neutral | | | -1.23E+00 | Neutral |
| 47 | p.Arg47Tyr | | 9908 | 3661 | 6.36 | 5.98 | | | | | -1.99E+00 | Neutral | | | -1.99E+00 | Neutral |
| 47 | p.Arg47Cys | | 9687 | 2786 | 6.22 | 4.55 | | | | | -3.09E-01 | Neutral | | | -3.09E-01 | Neutral |
| 47 | p.Arg47Trp | | 8071 | 3147 | 5.18 | 5.14 | | | | | -3.86E+00 | Neutral | | | -3.86E+00 | Neutral |
| 47 | p.Arg47Phe | | 11181 | 3229 | 7.18 | 5.27 | | | | | -2.00E-01 | Neutral | | | -2.00E-01 | Neutral |
| 48 | p.Pro48Asn | | 2904 | 1850 | 5.11 | 6.04 | | | | | -5.32E+01 | Deleterious | | | -5.32E+01 | Deleterious |
| 48 | p.Pro48Lys | | 3193 | 2371 | 5.62 | 7.75 | | | | | -5.32E+01 | Deleterious | | | -5.32E+01 | Deleterious |
| 48 | p.Pro48Thr | Pathogenic | 2253 | 231 | 3.97 | 0.75 | | | | | -1.93E+01 | Indeterminate | | | -1.93E+01 | Indeterminate |
| 48 | p.Pro48Arg | | 2576 | 1662 | 4.53 | 5.43 | | | | | -5.32E+01 | Deleterious | | | -5.32E+01 | Deleterious |
| 48 | p.Pro48Ser | | 2673 | 193 | 4.70 | 0.63 | | | | | -4.30E+00 | Neutral | | | -4.30E+00 | Neutral |
| 48 | p.Pro48Ile | | 3400 | 1282 | 5.98 | 4.19 | | | | | -5.32E+01 | Deleterious | | | -5.32E+01 | Deleterious |
| 48 | p.Pro48Met | | 3721 | 2133 | 6.55 | 6.97 | | | | | -5.32E+01 | Deleterious | | | -5.32E+01 | Deleterious |
| 48 | p.Pro48His | | 2798 | 1554 | 4.92 | 5.08 | | | | | -5.32E+01 | Deleterious | | | -5.32E+01 | Deleterious |
| 48 | p.Pro48Gln | | 2850 | 2049 | 5.02 | 6.69 | | | | | -5.32E+01 | Deleterious | | | -5.32E+01 | Deleterious |
| 48 | p.Pro48Pro | Synonymous | 1620 | 78 | 2.85 | 0.25 | | | | | -1.06E+00 | Neutral | | | -1.06E+00 | Neutral |
| 48 | p.Pro48Leu | | 4208 | 3546 | 7.41 | 11.59 | | | | | -5.32E+01 | Deleterious | | | -5.32E+01 | Deleterious |
| 48 | p.Pro48Asp | | 3106 | 2678 | 5.47 | 8.75 | | | | | -5.32E+01 | Deleterious | | | -5.32E+01 | Deleterious |
| 48 | p.Pro48Glu | | 2942 | 2356 | 5.18 | 7.70 | | | | | -5.32E+01 | Deleterious | | | -5.32E+01 | Deleterious |
| 48 | p.Pro48Ala | | 1831 | 66 | 3.22 | 0.22 | | | | | -3.82E-02 | Neutral | | | -3.82E-02 | Neutral |
| 48 | p.Pro48Gly | | 2682 | 268 | 4.72 | 0.88 | | | | | -1.50E+01 | Indeterminate | | | -1.50E+01 | Indeterminate |
| 48 | p.Pro48Val | | 2679 | 279 | 4.72 | 0.91 | | | | | -1.70E+01 | Indeterminate | | | -1.70E+01 | Indeterminate |
| 48 | p.Pro48Tyr | | 2687 | 2346 | 4.73 | 7.66 | | | | | -5.32E+01 | Deleterious | | | -5.32E+01 | Deleterious |
| 48 | p.Pro48Cys | | 1917 | 219 | 3.37 | 0.72 | | | | | -2.99E+01 | Indeterminate | | | -2.98E+01 | Indeterminate |
| 48 | p.Pro48Trp | | 3810 | 3295 | 6.71 | 10.77 | | | | | -5.32E+01 | Deleterious | | | -5.32E+01 | Deleterious |
| 48 | p.Pro48Phe | | 2967 | 2151 | 5.22 | 7.03 | | | | | -5.32E+01 | Deleterious | | | -5.32E+01 | Deleterious |
| 49 | p.Ile49Asn | | 2237 | 1425 | 4.04 | 5.04 | | | | | -5.32E+01 | Deleterious | | | -5.32E+01 | Deleterious |
| 49 | p.Ile49Lys | | 2695 | 2221 | 4.87 | 7.85 | | | | | -5.32E+01 | Deleterious | | | -5.32E+01 | Deleterious |
| 49 | p.Ile49Thr | Likely pathogenic | 2164 | 487 | 3.91 | 1.72 | | | | | -3.01E+01 | Indeterminate | | | -3.00E+01 | Indeterminate |
| 49 | p.Ile49Arg | | 3841 | 2810 | 6.94 | 9.94 | | | | | -5.32E+01 | Deleterious | | | -5.32E+01 | Deleterious |
| 49 | p.Ile49Ser | Likely pathogenic | 3236 | 1418 | 5.85 | 5.01 | | | | | -5.32E+01 | Deleterious | | | -5.32E+01 | Deleterious |
| 49 | p.Ile49Ile | Synonymous | 1978 | 192 | 3.57 | 0.68 | | | | | -1.06E+00 | Neutral | | | -1.06E+00 | Neutral |

| | | | | | | | | | | | |
|---|---|---|---|---|---|---|---|---|---|---|---|
| 49 | p.Ile49Met | | | 2492 | 263 | 4.50 | 0.93 | -1.12E+00 | Neutral | -1.12E+00 | Neutral |
| 49 | p.Ile49His | | | 2474 | 1184 | 4.47 | 4.19 | -5.32E+01 | Deleterious | -5.32E+01 | Deleterious |
| 49 | p.Ile49Gln | | | 3367 | 1300 | 6.08 | 4.60 | -5.32E+01 | Deleterious | -5.32E+01 | Deleterious |
| 49 | p.Ile49Pro | | | 3123 | 2176 | 5.64 | 7.69 | -5.32E+01 | Deleterious | -5.32E+01 | Deleterious |
| 49 | p.Ile49Leu | | | 2625 | 246 | 4.74 | 0.87 | -3.69E-01 | Neutral | -3.69E-01 | Neutral |
| 49 | p.Ile49Asp | | | 3604 | 3212 | 6.51 | 11.36 | -5.32E+01 | Deleterious | -5.32E+01 | Deleterious |
| 49 | p.Ile49Glu | | | 3855 | 3143 | 6.96 | 11.11 | -5.32E+01 | Deleterious | -5.32E+01 | Deleterious |
| 49 | p.Ile49Ala | | | 2079 | 459 | 3.76 | 1.62 | -2.97E+01 | Indeterminate | -2.96E+01 | Indeterminate |
| 49 | p.Ile49Gly | | | 3080 | 2656 | 5.56 | 9.39 | -5.32E+01 | Deleterious | -5.32E+01 | Deleterious |
| 49 | p.Ile49Val | | | 2891 | 306 | 5.22 | 1.08 | -7.90E-01 | Neutral | -7.90E-01 | Neutral |
| 49 | p.Ile49Tyr | | | 1968 | 1294 | 3.56 | 4.58 | -5.32E+01 | Deleterious | -5.32E+01 | Deleterious |
| 49 | p.Ile49Cys | | | 1784 | 321 | 3.22 | 1.14 | -1.96E+01 | Indeterminate | -1.96E+01 | Indeterminate |
| 49 | p.Ile49Trp | | | 3459 | 2624 | 6.25 | 9.28 | -5.32E+01 | Deleterious | -5.32E+01 | Deleterious |
| 49 | p.Ile49Phe | | | 2399 | 543 | 4.33 | 1.92 | -2.80E+01 | Indeterminate | -2.80E+01 | Indeterminate |
| 50 | p.Gln50Asn | | | 6179 | 763 | 3.25 | 0.67 | -1.25E+01 | Indeterminate | -1.25E+01 | Indeterminate |
| 50 | p.Gln50Lys | | | 7027 | 8327 | 3.70 | 7.36 | -5.32E+01 | Deleterious | -5.32E+01 | Deleterious |
| 50 | p.Gln50Thr | | | 5937 | 695 | 3.12 | 0.61 | -1.09E+01 | Indeterminate | -1.09E+01 | Indeterminate |
| 50 | p.Gln50Arg | | Deleterious | 7334 | 11605 | 3.86 | 10.26 | -5.32E+01 | Deleterious | -5.32E+01 | Deleterious |
| 50 | p.Gln50Ser | | | 8421 | 926 | 4.43 | 0.82 | -5.39E+00 | Neutral | -5.39E+00 | Neutral |
| 50 | p.Gln50Ile | | | 8078 | 3493 | 4.25 | 3.09 | -5.32E+01 | Deleterious | -5.32E+01 | Deleterious |
| 50 | p.Gln50Met | | | 9962 | 960 | 5.24 | 0.85 | -2.05E+00 | Neutral | -2.05E+00 | Neutral |
| 50 | p.Gln50His | | | 10223 | 4095 | 5.38 | 3.62 | -5.32E+01 | Deleterious | -5.32E+01 | Deleterious |
| 50 | p.Gln50Gln | Synonymous | | 12106 | 1126 | 6.37 | 1.00 | -1.06E+00 | Neutral | -1.06E+00 | Neutral |
| 50 | p.Gln50Pro | | | 9549 | 16066 | 5.03 | 14.20 | -5.32E+01 | Deleterious | -5.32E+01 | Deleterious |
| 50 | p.Gln50Leu | | | 11007 | 1915 | 5.79 | 1.69 | -2.00E+01 | Indeterminate | -2.00E+01 | Indeterminate |
| 50 | p.Gln50Asp | | | 10090 | 6468 | 5.31 | 5.72 | -5.32E+01 | Deleterious | -5.32E+01 | Deleterious |
| 50 | p.Gln50Glu | | | 12681 | 1243 | 6.67 | 1.10 | -1.37E+00 | Neutral | -1.37E+00 | Neutral |
| 50 | p.Gln50Ala | | | 7885 | 1001 | 4.15 | 0.88 | -1.04E+01 | Indeterminate | -1.04E+01 | Indeterminate |
| 50 | p.Gln50Gly | | | 7736 | 1057 | 4.07 | 0.93 | -1.36E+01 | Indeterminate | -1.36E+01 | Indeterminate |
| 50 | p.Gln50Val | | | 14010 | 5143 | 7.37 | 4.55 | -5.32E+01 | Deleterious | -5.32E+01 | Deleterious |
| 50 | p.Gln50Tyr | | | 13564 | 21314 | 7.14 | 18.84 | -5.32E+01 | Deleterious | -5.32E+01 | Deleterious |
| 50 | p.Gln50Cys | | | 9439 | 1625 | 4.97 | 1.44 | -2.24E+01 | Indeterminate | -2.24E+01 | Indeterminate |
| 50 | p.Gln50Trp | | | 10854 | 15054 | 5.71 | 13.30 | -5.32E+01 | Deleterious | -5.32E+01 | Deleterious |
| 50 | p.Gln50Phe | | | 7924 | 10276 | 4.17 | 9.08 | -5.32E+01 | Deleterious | -5.32E+01 | Deleterious |
| 51 | p.Val51Asn | | | 2467 | 455 | 1.70 | 0.40 | -5.32E+01 | Deleterious | -5.32E+01 | Deleterious |
| 51 | p.Val51Lys | | | 3393 | 9087 | 2.34 | 8.00 | -5.32E+01 | Deleterious | -5.32E+01 | Deleterious |
| 51 | p.Val51Thr | | | 3961 | 298 | 2.73 | 0.26 | -4.90E+00 | Neutral | -4.90E+00 | Neutral |
| 51 | p.Val51Arg | | | 3536 | 10023 | 2.44 | 8.83 | -5.32E+01 | Deleterious | -5.32E+01 | Deleterious |
| 51 | p.Val51Ser | | | 7469 | 443 | 5.14 | 0.39 | -2.22E-01 | Neutral | -2.22E-01 | Neutral |
| 51 | p.Val51Ile | | | 8945 | 908 | 6.16 | 0.80 | -5.58E+00 | Neutral | -5.58E+00 | Neutral |
| 51 | p.Val51Met | | | 6116 | 1205 | 4.21 | 1.06 | -5.32E+01 | Deleterious | -5.32E+01 | Deleterious |
| 51 | p.Val51His | | | 6561 | 10183 | 4.52 | 8.97 | -5.32E+01 | Deleterious | -5.32E+01 | Deleterious |
| 51 | p.Val51Gln | | | 10283 | 2904 | 7.08 | 2.56 | -5.32E+01 | Deleterious | -5.32E+01 | Deleterious |
| 51 | p.Val51Pro | | | 12358 | 22046 | 8.51 | 19.41 | -5.32E+01 | Deleterious | -5.32E+01 | Deleterious |
| 51 | p.Val51Leu | | | 9977 | 848 | 6.87 | 0.75 | -1.84E+00 | Neutral | -1.84E+00 | Neutral |
| 51 | p.Val51Asp | | | 11158 | 18353 | 7.68 | 16.16 | -5.32E+01 | Deleterious | -5.32E+01 | Deleterious |
| 51 | p.Val51Glu | | | 7184 | 13435 | 4.95 | 11.83 | -5.32E+01 | Deleterious | -5.32E+01 | Deleterious |
| 51 | p.Val51Ala | | | 6834 | 308 | 4.71 | 0.27 | -9.40E-03 | Neutral | -9.40E-03 | Neutral |
| 51 | p.Val51Gly | | | 5966 | 473 | 4.11 | 0.42 | -3.26E+00 | Neutral | -3.26E+00 | Neutral |
| 51 | p.Val51Val | Synonymous | | 10471 | 833 | 7.21 | 0.73 | -1.06E+00 | Neutral | -1.06E+00 | Neutral |
| 51 | p.Val51Tyr | | | 4583 | 4152 | 3.16 | 3.66 | -5.32E+01 | Deleterious | -5.32E+01 | Deleterious |
| 51 | p.Val51Cys | | | 13104 | 988 | 9.02 | 0.87 | -3.73E-01 | Neutral | -3.73E-01 | Neutral |
| 51 | p.Val51Trp | | | 7433 | 14300 | 5.12 | 12.59 | -5.32E+01 | Deleterious | -5.32E+01 | Deleterious |
| 51 | p.Val51Phe | | | 3400 | 2313 | 2.34 | 2.04 | -5.32E+01 | Deleterious | -5.32E+01 | Deleterious |
| 52 | p.Met52Asn | | | 4478 | 6062 | 2.15 | 2.76 | -1.64E+01 | Indeterminate | -1.64E+01 | Indeterminate |
| 52 | p.Met52Lys | | | 3389 | 6716 | 1.63 | 3.06 | -5.32E+01 | Deleterious | -5.32E+01 | Deleterious |
| 52 | p.Met52Thr | | | 7325 | 6547 | 3.52 | 2.98 | -1.17E+00 | Neutral | -1.17E+00 | Neutral |
| 52 | p.Met52Arg | | | 3644 | 7476 | 1.75 | 3.41 | -5.32E+01 | Deleterious | -5.32E+01 | Deleterious |
| 52 | p.Met52Ser | | | 12692 | 10076 | 6.09 | 4.59 | -6.65E-02 | Neutral | -6.65E-02 | Neutral |
| 52 | p.Met52Ile | | | 4566 | 5019 | 2.19 | 2.29 | -7.82E+00 | Indeterminate | -7.82E+00 | Indeterminate |
| 52 | p.Met52Met | Synonymous | | 4922 | 3803 | 2.36 | 1.73 | -1.06E+00 | Neutral | -1.06E+00 | Neutral |
| 52 | p.Met52His | | | 8903 | 10578 | 4.27 | 4.82 | -4.19E+00 | Neutral | -4.19E+00 | Neutral |
| 52 | p.Met52Gln | | | 8262 | 7816 | 3.97 | 3.56 | -1.31E+00 | Neutral | -1.31E+00 | Neutral |
| 52 | p.Met52Pro | | | 11304 | 19362 | 5.43 | 8.83 | -1.31E+01 | Indeterminate | -1.31E+01 | Indeterminate |
| 52 | p.Met52Leu | | | 19024 | 15490 | 9.13 | 7.06 | -1.49E-02 | Neutral | -1.49E-02 | Neutral |
| 52 | p.Met52Asp | | | 7647 | 15009 | 3.67 | 6.84 | -2.82E+01 | Indeterminate | -2.82E+01 | Indeterminate |
| 52 | p.Met52Glu | | | 10223 | 16152 | 4.91 | 7.36 | -1.11E+01 | Indeterminate | -1.11E+01 | Indeterminate |
| 52 | p.Met52Ala | | | 20519 | 16257 | 9.85 | 7.41 | -6.58E-03 | Neutral | -6.58E-03 | Neutral |
| 52 | p.Met52Gly | | | 28403 | 21442 | 13.64 | 9.77 | -4.04E-04 | Neutral | -4.04E-04 | Neutral |
| 52 | p.Met52Val | | | 16148 | 12224 | 7.75 | 5.57 | -1.12E-02 | Neutral | -1.12E-02 | Neutral |
| 52 | p.Met52Tyr | | | 5210 | 6051 | 2.50 | 2.76 | -8.17E+00 | Indeterminate | -8.17E+00 | Indeterminate |
| 52 | p.Met52Cys | | | 15883 | 11778 | 7.63 | 5.37 | -8.88E-03 | Neutral | -8.88E-03 | Neutral |
| 52 | p.Met52Trp | | | 10918 | 16716 | 5.24 | 7.62 | -9.17E+00 | Indeterminate | -9.17E+00 | Indeterminate |
| 52 | p.Met52Phe | | | 4809 | 4797 | 2.31 | 2.19 | -4.81E+00 | Neutral | -4.81E+00 | Neutral |
| 53 | p.Met53Asn | | | 5148 | 1876 | 4.45 | 2.67 | -3.58E+01 | Indeterminate | -3.30E+01 | Indeterminate |
| 53 | p.Met53Lys | | | 6247 | 1516 | 5.40 | 2.16 | -8.88E+00 | Indeterminate | -8.88E+00 | Indeterminate |
| 53 | p.Met53Thr | | | 5939 | 2515 | 5.13 | 3.58 | -4.53E+01 | Indeterminate | -3.32E+01 | Indeterminate |
| 53 | p.Met53Arg | | | 5012 | 915 | 4.33 | 1.30 | -3.54E+00 | Neutral | -3.54E+00 | Neutral |
| 53 | p.Met53Ser | | | 5763 | 1238 | 4.98 | 1.76 | -6.07E+00 | Indeterminate | -6.07E+00 | Indeterminate |
| 53 | p.Met53Ile | Pathogenic | | 5291 | 4631 | 4.57 | 6.59 | -5.32E+01 | Deleterious | -5.32E+01 | Deleterious |
| 53 | p.Met53Met | Synonymous | | 3139 | 402 | 2.71 | 0.57 | -1.06E+00 | Neutral | -1.06E+00 | Neutral |
| 53 | p.Met53His | | | 7214 | 3027 | 6.23 | 4.31 | -3.83E+01 | Indeterminate | -3.32E+01 | Indeterminate |
| 53 | p.Met53Gln | | | 8241 | 1527 | 7.12 | 2.17 | -1.49E+00 | Neutral | -1.49E+00 | Neutral |
| 53 | p.Met53Pro | | | 5318 | 6610 | 4.59 | 9.40 | -5.32E+01 | Deleterious | -5.32E+01 | Deleterious |
| 53 | p.Met53Leu | | | 6554 | 1519 | 5.66 | 2.16 | -6.97E+00 | Indeterminate | -6.97E+00 | Indeterminate |
| 53 | p.Met53Asp | | | 5058 | 9025 | 4.37 | 12.84 | -5.32E+01 | Deleterious | -5.32E+01 | Deleterious |
| 53 | p.Met53Glu | | | 7104 | 3670 | 6.14 | 5.22 | -5.32E+01 | Deleterious | -5.32E+01 | Deleterious |
| 53 | p.Met53Ala | | | 6055 | 1223 | 5.23 | 1.74 | -4.27E+00 | Neutral | -4.27E+00 | Neutral |
| 53 | p.Met53Gly | | | 5478 | 4287 | 4.73 | 6.10 | -5.32E+01 | Deleterious | -5.32E+01 | Deleterious |
| 53 | p.Met53Val | Likely pathogenic | | 5837 | 5714 | 5.04 | 8.13 | -5.32E+01 | Deleterious | -5.32E+01 | Deleterious |
| 53 | p.Met53Tyr | | | 5550 | 4771 | 4.80 | 6.79 | -5.32E+01 | Deleterious | -5.32E+01 | Deleterious |
| 53 | p.Met53Cys | | | 4558 | 1062 | 3.94 | 1.51 | -1.12E+01 | Indeterminate | -1.12E+01 | Indeterminate |
| 53 | p.Met53Trp | | | 7931 | 11517 | 6.85 | 16.38 | -5.32E+01 | Deleterious | -5.32E+01 | Deleterious |
| 53 | p.Met53Phe | | | 4299 | 3259 | 3.71 | 4.64 | -5.32E+01 | Deleterious | -5.32E+01 | Deleterious |
| 54 | p.Met54Asn | | | 4729 | 6229 | 4.82 | 4.68 | -1.00E+00 | Neutral | -1.00E+00 | Neutral |
| 54 | p.Met54Lys | | | 1457 | 2678 | 1.48 | 2.01 | -2.35E+01 | Indeterminate | -2.35E+01 | Indeterminate |
| 54 | p.Met54Thr | | | 9921 | 9389 | 10.11 | 7.06 | -6.02E-04 | Neutral | -6.02E-04 | Neutral |
| 54 | p.Met54Arg | | | 2692 | 6197 | 2.74 | 4.66 | -2.49E+01 | Indeterminate | -2.49E+01 | Indeterminate |
| 54 | p.Met54Ser | | | 7785 | 8580 | 7.93 | 6.45 | -2.84E-02 | Neutral | -2.84E-02 | Neutral |
| 54 | p.Met54Ile | | | 4949 | 5748 | 5.04 | 4.32 | -3.06E-01 | Neutral | -3.06E-01 | Neutral |
| 54 | p.Met54Met | Synonymous | | 1901 | 1865 | 1.94 | 1.40 | -1.06E+00 | Neutral | -1.06E+00 | Neutral |
| 54 | p.Met54His | | | 1990 | 2453 | 2.03 | 1.84 | -3.79E+00 | Neutral | -3.79E+00 | Neutral |
| 54 | p.Met54Gln | | | 3851 | 6289 | 3.92 | 4.73 | -5.09E+00 | Neutral | -5.09E+00 | Neutral |
| 54 | p.Met54Pro | | | 7720 | 20805 | 7.86 | 15.63 | -1.42E+01 | Indeterminate | -1.42E+01 | Indeterminate |
| 54 | p.Met54Leu | | | 5644 | 6063 | 5.75 | 4.56 | -8.18E-02 | Neutral | -8.18E-02 | Neutral |
| 54 | p.Met54Asp | | | 5244 | 12085 | 5.34 | 9.08 | -1.29E+01 | Indeterminate | -1.29E+01 | Indeterminate |
| 54 | p.Met54Glu | | | 3404 | 6990 | 3.47 | 5.25 | -1.42E+01 | Indeterminate | -1.42E+01 | Indeterminate |
| 54 | p.Met54Ala | | | 4236 | 4218 | 4.32 | 3.17 | -1.03E-01 | Neutral | -1.03E-01 | Neutral |
| 54 | p.Met54Gly | | | 6165 | 6242 | 6.28 | 4.69 | -2.56E-02 | Neutral | -2.56E-02 | Neutral |
| 54 | p.Met54Val | | | 5273 | 5438 | 5.37 | 4.09 | -6.54E-02 | Neutral | -6.54E-02 | Neutral |
| 54 | p.Met54Tyr | | | 2374 | 2422 | 2.42 | 1.82 | -8.20E-01 | Neutral | -8.20E-01 | Neutral |
| 54 | p.Met54Cys | | | 7763 | 6885 | 7.91 | 5.17 | -8.22E-04 | Neutral | -8.22E-04 | Neutral |
| 54 | p.Met54Trp | | | 4101 | 4916 | 4.18 | 3.69 | -7.00E-01 | Neutral | -7.00E-01 | Neutral |
| 54 | p.Met54Phe | | | 6961 | 7588 | 7.09 | 5.70 | -4.06E-02 | Neutral | -4.06E-02 | Neutral |
| 55 | p.Gly55Asn | | | 7992 | 2142 | 3.29 | 3.35 | -5.32E+01 | Deleterious | -5.32E+01 | Deleterious |
| 55 | p.Gly55Lys | | | 10951 | 2798 | 4.51 | 4.37 | -5.32E+01 | Deleterious | -5.32E+01 | Deleterious |
| 55 | p.Gly55Thr | | | 17033 | 3602 | 7.01 | 5.63 | -5.32E+01 | Deleterious | -5.32E+01 | Deleterious |
| 55 | p.Gly55Arg | | | 11124 | 2826 | 4.58 | 4.42 | -5.32E+01 | Deleterious | -5.32E+01 | Deleterious |
| 55 | p.Gly55Ser | | | 12413 | 706 | 5.11 | 1.10 | -3.27E+01 | Indeterminate | -3.19E+01 | Indeterminate |
| 55 | p.Gly55Ile | | | 11813 | 2953 | 4.86 | 4.62 | -5.32E+01 | Deleterious | -5.32E+01 | Deleterious |
| 55 | p.Gly55Met | | | 10258 | 3064 | 4.22 | 4.79 | -5.32E+01 | Deleterious | -5.32E+01 | Deleterious |
| 55 | p.Gly55His | | | 14055 | 4676 | 5.78 | 7.31 | -5.32E+01 | Deleterious | -5.32E+01 | Deleterious |
| 55 | p.Gly55Gln | | | 14040 | 3922 | 5.78 | 6.13 | -5.32E+01 | Deleterious | -5.32E+01 | Deleterious |
| 55 | p.Gly55Pro | | | 24068 | 10469 | 9.90 | 16.36 | -5.32E+01 | Deleterious | -5.32E+01 | Deleterious |
| 55 | p.Gly55Leu | | | 21350 | 7759 | 8.79 | 12.13 | -5.32E+01 | Deleterious | -5.32E+01 | Deleterious |
| 55 | p.Gly55Asp | | | 12218 | 4275 | 5.03 | 6.68 | -5.32E+01 | Deleterious | -5.32E+01 | Deleterious |
| 55 | p.Gly55Glu | | | 16079 | 4459 | 6.62 | 6.97 | -5.32E+01 | Deleterious | -5.32E+01 | Deleterious |
| 55 | p.Gly55Ala | | | 8380 | 288 | 3.45 | 0.45 | -1.10E+01 | Indeterminate | -1.10E+01 | Indeterminate |
| 55 | p.Gly55Gly | Synonymous | | 8104 | 172 | 3.34 | 0.27 | -1.06E+00 | Neutral | -1.06E+00 | Neutral |
| 55 | p.Gly55Val | | | 7187 | 2149 | 2.96 | 3.36 | -5.32E+01 | Deleterious | -5.32E+01 | Deleterious |
| 55 | p.Gly55Tyr | | | 10835 | 2494 | 4.46 | 3.90 | -5.32E+01 | Deleterious | -5.32E+01 | Deleterious |
| 55 | p.Gly55Cys | | | 9250 | 737 | 3.81 | 1.15 | -5.32E+01 | Deleterious | -5.32E+01 | Deleterious |
| 55 | p.Gly55Trp | | | 8904 | 2798 | 3.66 | 4.37 | -5.32E+01 | Deleterious | -5.32E+01 | Deleterious |
| 55 | p.Gly55Phe | | | 6940 | 1690 | 2.86 | 2.64 | -5.32E+01 | Deleterious | -5.32E+01 | Deleterious |
| 56 | p.Ser56Asn | | | 15782 | 229 | 5.04 | 0.81 | -8.15E-10 | Neutral | -6.71E-10 | Neutral |
| 56 | p.Ser56Lys | | | 21348 | 835 | 6.82 | 2.94 | -9.71E-02 | Neutral | -9.71E-02 | Neutral |
| 56 | p.Ser56Thr | | | 12135 | 459 | 3.88 | 1.62 | -4.71E-01 | Neutral | -4.71E-01 | Neutral |
| 56 | p.Ser56Arg | | | 12834 | 231 | 4.10 | 0.81 | -1.41E-06 | Neutral | -1.41E-06 | Neutral |
| 56 | p.Ser56Ser | Synonymous | | 18053 | 863 | 5.77 | 3.04 | -1.06E+00 | Neutral | -1.06E+00 | Neutral |

| | | | | | | | | | | | | | | | | | |
|---|---|---|---|---|---|---|---|---|---|---|---|---|---|---|---|---|---|
| 56 | p.Ser56Ile | Pathogenic | | 15771 | 4860 | 5.04 | 17.13 | | | | | -5.32E+01 | Deleterious | | | -5.32E+01 | Deleterious |
| 56 | p.Ser56Met | | | 9103 | 181 | 2.91 | 0.64 | | | | | -1.66E-04 | Neutral | | | -1.66E-04 | Neutral |
| 56 | p.Ser56His | | | 16781 | 342 | 5.36 | 1.21 | | | | | -3.27E-06 | Neutral | | | -3.27E-06 | Neutral |
| 56 | p.Ser56Gln | | | 13828 | 378 | 4.42 | 1.33 | | | | | -4.63E-03 | Neutral | | | -4.63E-03 | Neutral |
| 56 | p.Ser56Pro | | | 15799 | 9165 | 5.05 | 32.30 | | | | | -5.32E+01 | Deleterious | | | -5.32E+01 | Deleterious |
| 56 | p.Ser56Leu | | | 18803 | 1683 | 6.01 | 5.93 | | | | | -1.75E+01 | Indeterminate | | | -1.75E+01 | Indeterminate |
| 56 | p.Ser56Asp | | | 15719 | 594 | 5.02 | 2.09 | | | | | -2.05E-01 | Neutral | | | -2.05E-01 | Neutral |
| 56 | p.Ser56Glu | | | 16419 | 941 | 5.25 | 3.32 | | | | | -3.79E+00 | Neutral | | | -3.79E+00 | Neutral |
| 56 | p.Ser56Ala | | | 17828 | 655 | 5.70 | 2.31 | | | | | -9.40E-02 | Neutral | | | -9.40E-02 | Neutral |
| 56 | p.Ser56Gly | | | 20474 | 1154 | 6.54 | 4.07 | | | | | -2.34E+00 | Neutral | | | -2.34E+00 | Neutral |
| 56 | p.Ser56Val | | | 17532 | 973 | 5.60 | 3.43 | | | | | -2.86E+00 | Neutral | | | -2.86E+00 | Neutral |
| 56 | p.Ser56Tyr | | | 7771 | 187 | 2.48 | 0.66 | | | | | -1.28E-02 | Neutral | | | -1.28E-02 | Neutral |
| 56 | p.Ser56Cys | | | 14573 | 299 | 4.66 | 1.05 | | | | | -1.13E-05 | Neutral | | | -1.13E-05 | Neutral |
| 56 | p.Ser56Trp | | | 17186 | 3817 | 5.49 | 13.45 | | | | | -5.32E+01 | Deleterious | | | -5.32E+01 | Deleterious |
| 56 | p.Ser56Phe | | | 15288 | 528 | 4.88 | 1.86 | | | | | -8.26E-02 | Neutral | | | -8.26E-02 | Neutral |
| 57 | p.Ala57Asn | | | 4228 | 707 | 5.51 | 4.84 | 12028 | 2228 | 5.44 | 4.68 | -8.42E-07 | Neutral | -1.69E+00 | Neutral | -5.69E-01 | Neutral |
| 57 | p.Ala57Lys | | | 3617 | 548 | 4.71 | 3.75 | 10968 | 2317 | 4.96 | 4.86 | -2.63E-07 | Neutral | -4.13E+00 | Neutral | -2.18E+00 | Neutral |
| 57 | p.Ala57Thr | | | 2864 | 570 | 3.73 | 3.90 | 8869 | 1879 | 4.01 | 3.95 | -6.42E-04 | Neutral | -5.78E+00 | Neutral | -3.45E+00 | Neutral |
| 57 | p.Ala57Arg | | | 3189 | 550 | 4.16 | 3.77 | 11295 | 1899 | 5.10 | 3.99 | -1.57E-05 | Neutral | -1.01E+00 | Neutral | -2.42E-01 | Neutral |
| 57 | p.Ala57Ser | | | 4843 | 991 | 6.31 | 6.79 | 14163 | 3613 | 6.40 | 7.59 | -3.20E-05 | Neutral | -6.57E+00 | Neutral | -4.10E+00 | Neutral |
| 57 | p.Ala57Ile | | | 2093 | 525 | 2.73 | 3.60 | 10521 | 1993 | 4.75 | 4.18 | -1.10E-01 | Neutral | -2.51E+00 | Neutral | -1.13E+00 | Neutral |
| 57 | p.Ala57Met | | | 3187 | 502 | 4.15 | 3.44 | 8084 | 1645 | 3.65 | 3.45 | -1.92E-06 | Neutral | -5.54E+00 | Neutral | -3.27E+00 | Neutral |
| 57 | p.Ala57His | | | 4842 | 956 | 6.31 | 6.55 | 13594 | 1983 | 6.14 | 4.16 | -1.45E-05 | Neutral | -1.62E-01 | Neutral | -8.49E-03 | Neutral |
| 57 | p.Ala57Gln | | | 4779 | 911 | 6.23 | 6.24 | 11318 | 2326 | 5.11 | 4.88 | -7.21E-06 | Neutral | -3.42E+00 | Neutral | -1.67E+00 | Neutral |
| 57 | p.Ala57Pro | | | 4088 | 887 | 5.33 | 6.08 | 10929 | 2644 | 4.94 | 5.55 | -3.47E-04 | Neutral | -7.51E+00 | Indeterminate | -4.88E+00 | Neutral |
| 57 | p.Ala57Leu | | | 3855 | 603 | 5.02 | 4.13 | 9126 | 2396 | 4.12 | 5.03 | -3.45E-07 | Neutral | -1.26E+01 | Indeterminate | -9.30E+00 | Indeterminate |
| 57 | p.Ala57Asp | | | 4487 | 1216 | 5.85 | 8.33 | 13571 | 2300 | 6.13 | 4.83 | -1.01E-02 | Neutral | -6.69E-01 | Neutral | -1.23E-01 | Neutral |
| 57 | p.Ala57Glu | | | 3993 | 794 | 5.20 | 5.44 | 13025 | 3886 | 5.89 | 8.16 | -6.86E-05 | Neutral | -1.30E+01 | Indeterminate | -9.67E+00 | Indeterminate |
| 57 | p.Ala57Ala | Synonymous | | 2787 | 983 | 3.63 | 6.73 | 9862 | 1597 | 4.46 | 3.35 | -1.06E+00 | Neutral | -1.06E+00 | Neutral | -8.18E-01 | Neutral |
| 57 | p.Ala57Gly | | | 5677 | 999 | 7.40 | 6.84 | 13161 | 2709 | 5.95 | 5.69 | -2.63E-07 | Neutral | -2.64E+00 | Neutral | -1.14E+00 | Neutral |
| 57 | p.Ala57Val | Likely benign | | 4184 | 783 | 5.45 | 5.36 | 14947 | 2673 | 6.75 | 5.61 | -1.29E-05 | Neutral | -7.94E-01 | Neutral | -1.61E-01 | Neutral |
| 57 | p.Ala57Tyr | | | 3130 | 578 | 4.08 | 3.96 | 8794 | 2642 | 3.97 | 5.55 | -7.93E-05 | Neutral | -1.99E+01 | Indeterminate | -1.60E+01 | Indeterminate |
| 57 | p.Ala57Cys | | | 4967 | 664 | 6.47 | 4.55 | 11591 | 2870 | 5.24 | 6.03 | -5.42E-10 | Neutral | -7.62E+00 | Indeterminate | -4.97E+00 | Neutral |
| 57 | p.Ala57Trp | | | 3220 | 520 | 4.20 | 3.56 | 8448 | 2110 | 3.82 | 4.43 | -3.21E-06 | Neutral | -1.16E+01 | Indeterminate | -8.41E+00 | Indeterminate |
| 57 | p.Ala57Phe | | | 2695 | 310 | 3.51 | 2.12 | 7000 | 1919 | 3.16 | 4.03 | -2.37E-09 | Neutral | -1.90E+01 | Indeterminate | -1.51E+01 | Indeterminate |
| 58 | p.Arg58Asn | | | 11388 | 6167 | 5.68 | 5.29 | | | | | -3.71E-04 | Neutral | | | -3.71E-04 | Neutral |
| 58 | p.Arg58Lys | | | 12667 | 6509 | 6.32 | 5.59 | | | | | -6.17E-05 | Neutral | | | -6.17E-05 | Neutral |
| 58 | p.Arg58Thr | | | 4945 | 2887 | 2.47 | 2.48 | | | | | -9.58E-02 | Neutral | | | -9.58E-02 | Neutral |
| 58 | p.Arg58Arg | Synonymous | | 4309 | 3109 | 2.15 | 2.67 | | | | | -1.06E+00 | Neutral | | | -1.06E+00 | Neutral |
| 58 | p.Arg58Ser | | | 8923 | 4891 | 4.45 | 4.20 | | | | | -2.10E-03 | Neutral | | | -2.10E-03 | Neutral |
| 58 | p.Arg58Ile | | | 14106 | 8230 | 7.04 | 7.06 | | | | | -3.97E-04 | Neutral | | | -3.97E-04 | Neutral |
| 58 | p.Arg58Met | | | 8462 | 4305 | 4.22 | 3.70 | | | | | -7.43E-04 | Neutral | | | -7.43E-04 | Neutral |
| 58 | p.Arg58His | | | 10078 | 4647 | 5.03 | 3.99 | | | | | -3.07E-05 | Neutral | | | -3.07E-05 | Neutral |
| 58 | p.Arg58Gln | | | 8727 | 5863 | 4.35 | 5.03 | | | | | -5.28E-02 | Neutral | | | -5.28E-02 | Neutral |
| 58 | p.Arg58Pro | | | 9411 | 7328 | 4.70 | 6.29 | | | | | -2.18E-01 | Neutral | | | -2.18E-01 | Neutral |
| 58 | p.Arg58Leu | | | 10691 | 6368 | 5.33 | 5.47 | | | | | -3.14E-03 | Neutral | | | -3.14E-03 | Neutral |
| 58 | p.Arg58Asp | | | 10734 | 5563 | 5.36 | 4.78 | | | | | -2.29E-04 | Neutral | | | -2.29E-04 | Neutral |
| 58 | p.Arg58Glu | | | 12402 | 7460 | 6.19 | 6.40 | | | | | -1.57E-03 | Neutral | | | -1.57E-03 | Neutral |
| 58 | p.Arg58Ala | | | 8563 | 5464 | 4.27 | 4.69 | | | | | -2.88E-02 | Neutral | | | -2.88E-02 | Neutral |
| 58 | p.Arg58Gly | | | 9407 | 6284 | 4.69 | 5.39 | | | | | -3.52E-02 | Neutral | | | -3.52E-02 | Neutral |
| 58 | p.Arg58Val | | | 13045 | 8449 | 6.51 | 7.25 | | | | | -4.23E-03 | Neutral | | | -4.23E-03 | Neutral |
| 58 | p.Arg58Tyr | | | 11421 | 5102 | 5.70 | 4.38 | | | | | -5.90E-06 | Neutral | | | -5.90E-06 | Neutral |
| 58 | p.Arg58Cys | | | 10605 | 5418 | 5.29 | 4.65 | | | | | -1.86E-04 | Neutral | | | -1.86E-04 | Neutral |
| 58 | p.Arg58Trp | | | 8533 | 4672 | 4.26 | 4.01 | | | | | -2.67E-03 | Neutral | | | -2.67E-03 | Neutral |
| 58 | p.Arg58Phe | | | 12024 | 7781 | 6.00 | 6.68 | | | | | -6.51E-03 | Neutral | | | -6.51E-03 | Neutral |
| 59 | p.Val59Asn | | | 2415 | 585 | 4.15 | 1.45 | | | | | -5.32E+01 | Deleterious | | | -5.32E+01 | Deleterious |
| 59 | p.Val59Lys | | | 3708 | 3626 | 6.38 | 8.99 | | | | | -5.32E+01 | Deleterious | | | -5.32E+01 | Deleterious |
| 59 | p.Val59Thr | | | 3352 | 218 | 5.77 | 0.54 | | | | | -2.12E-01 | Neutral | | | -2.12E-01 | Neutral |
| 59 | p.Val59Arg | | | 2890 | 4202 | 4.97 | 10.42 | | | | | -5.32E+01 | Deleterious | | | -5.32E+01 | Deleterious |
| 59 | p.Val59Ser | | | 2742 | 260 | 4.72 | 0.64 | | | | | -4.70E+00 | Neutral | | | -4.70E+00 | Neutral |
| 59 | p.Val59Ile | | | 2316 | 130 | 3.98 | 0.32 | | | | | -1.64E-01 | Neutral | | | -1.64E-01 | Neutral |
| 59 | p.Val59Met | | | 2652 | 203 | 4.56 | 0.50 | | | | | -1.55E+00 | Neutral | | | -1.55E+00 | Neutral |
| 59 | p.Val59His | | | 2849 | 3587 | 4.90 | 8.90 | | | | | -5.32E+01 | Deleterious | | | -5.32E+01 | Deleterious |
| 59 | p.Val59Gln | | | 2872 | 790 | 4.94 | 1.96 | | | | | -5.32E+01 | Deleterious | | | -5.32E+01 | Deleterious |
| 59 | p.Val59Pro | | | 3188 | 3072 | 5.48 | 7.62 | | | | | -5.32E+01 | Deleterious | | | -5.32E+01 | Deleterious |
| 59 | p.Val59Leu | | | 3295 | 327 | 5.67 | 0.81 | | | | | -4.37E+00 | Neutral | | | -4.37E+00 | Neutral |
| 59 | p.Val59Asp | | | 2993 | 4745 | 5.15 | 11.77 | | | | | -5.32E+01 | Deleterious | | | -5.32E+01 | Deleterious |
| 59 | p.Val59Glu | | | 3323 | 3022 | 5.72 | 7.50 | | | | | -5.32E+01 | Deleterious | | | -5.32E+01 | Deleterious |
| 59 | p.Val59Ala | | | 2057 | 183 | 3.54 | 0.45 | | | | | -5.43E+00 | Neutral | | | -5.43E+00 | Neutral |
| 59 | p.Val59Gly | Pathogenic | | 1783 | 295 | 3.07 | 0.73 | | | | | -4.27E+01 | Indeterminate | | | -3.32E+01 | Indeterminate |
| 59 | p.Val59Val | Synonymous | | 3055 | 232 | 5.26 | 0.58 | | | | | -1.06E+00 | Neutral | | | -1.06E+00 | Neutral |
| 59 | p.Val59Tyr | | | 3471 | 4948 | 5.97 | 12.27 | | | | | -5.32E+01 | Deleterious | | | -5.32E+01 | Deleterious |
| 59 | p.Val59Cys | | | 2693 | 193 | 4.63 | 0.48 | | | | | -9.44E-01 | Neutral | | | -9.44E-01 | Neutral |
| 59 | p.Val59Trp | | | 3706 | 6000 | 6.38 | 14.88 | | | | | -5.32E+01 | Deleterious | | | -5.32E+01 | Deleterious |
| 59 | p.Val59Phe | | | 2767 | 3698 | 4.76 | 9.17 | | | | | -5.32E+01 | Deleterious | | | -5.32E+01 | Deleterious |
| 60 | p.Ala60Asn | | | 8575 | 4388 | 3.78 | 4.49 | | | | | -5.32E+01 | Deleterious | | | -5.32E+01 | Deleterious |
| 60 | p.Ala60Lys | | | 13108 | 8103 | 5.78 | 8.29 | | | | | -5.32E+01 | Deleterious | | | -5.32E+01 | Deleterious |
| 60 | p.Ala60Thr | | | 14496 | 2990 | 6.39 | 3.06 | | | | | -1.18E+00 | Neutral | | | -1.18E+00 | Neutral |
| 60 | p.Ala60Arg | | Deleterious | 11246 | 10734 | 4.96 | 10.98 | | | | | -5.32E+01 | Deleterious | | | -5.32E+01 | Deleterious |
| 60 | p.Ala60Ser | | | 13309 | 4554 | 5.86 | 4.66 | | | | | -1.44E+01 | Indeterminate | | | -1.44E+01 | Indeterminate |
| 60 | p.Ala60Ile | | | 11680 | 4451 | 5.15 | 4.55 | | | | | -2.25E+01 | Indeterminate | | | -2.25E+01 | Indeterminate |
| 60 | p.Ala60Met | | | 13714 | 3222 | 6.04 | 3.30 | | | | | -2.94E+00 | Neutral | | | -2.94E+00 | Neutral |
| 60 | p.Ala60His | | | 14244 | 7153 | 6.28 | 7.32 | | | | | -3.83E+01 | Indeterminate | | | -3.32E+01 | Indeterminate |
| 60 | p.Ala60Gln | | | 13494 | 8204 | 5.95 | 8.40 | | | | | -5.32E+01 | Deleterious | | | -5.32E+01 | Deleterious |
| 60 | p.Ala60Pro | | | 8977 | 2945 | 3.96 | 3.01 | | | | | -1.89E+01 | Indeterminate | | | -1.89E+01 | Indeterminate |
| 60 | p.Ala60Leu | | | 9673 | 1966 | 4.26 | 2.01 | | | | | -2.51E+00 | Neutral | | | -2.51E+00 | Neutral |
| 60 | p.Ala60Asp | | | 10802 | 5546 | 4.76 | 5.68 | | | | | -4.95E+01 | Indeterminate | | | -3.32E+01 | Indeterminate |
| 60 | p.Ala60Glu | Likely pathogenic | | 12346 | 6697 | 5.44 | 6.85 | | | | | -5.07E+01 | Indeterminate | | | -3.32E+01 | Indeterminate |
| 60 | p.Ala60Ala | Synonymous | | 12155 | 2329 | 5.36 | 2.38 | | | | | -1.06E+00 | Neutral | | | -1.06E+00 | Neutral |
| 60 | p.Ala60Gly | | | 10466 | 2098 | 4.61 | 2.15 | | | | | -1.98E+00 | Neutral | | | -1.98E+00 | Neutral |
| 60 | p.Ala60Val | | | 9945 | 2357 | 4.38 | 2.41 | | | | | -5.18E+00 | Neutral | | | -5.18E+00 | Neutral |
| 60 | p.Ala60Tyr | | | 9740 | 5258 | 4.29 | 5.38 | | | | | -5.32E+01 | Deleterious | | | -5.32E+01 | Deleterious |
| 60 | p.Ala60Cys | | | 9566 | 2423 | 4.21 | 2.48 | | | | | -7.26E+00 | Indeterminate | | | -7.26E+00 | Indeterminate |
| 60 | p.Ala60Trp | | | 8828 | 5434 | 3.89 | 5.56 | | | | | -5.32E+01 | Deleterious | | | -5.32E+01 | Deleterious |
| 60 | p.Ala60Phe | | | 10592 | 6867 | 4.67 | 7.03 | | | | | -5.32E+01 | Deleterious | | | -5.32E+01 | Deleterious |
| 61 | p.Glu61Asn | | | 14015 | 2650 | 5.27 | 4.43 | | | | | -1.02E+00 | Neutral | | | -1.02E+00 | Neutral |
| 61 | p.Glu61Lys | | | 12182 | 1826 | 4.58 | 3.05 | | | | | -2.01E-01 | Neutral | | | -2.01E-01 | Neutral |
| 61 | p.Glu61Thr | | | 15061 | 2530 | 5.66 | 4.23 | | | | | -3.11E-01 | Neutral | | | -3.11E-01 | Neutral |
| 61 | p.Glu61Arg | | | 12012 | 2820 | 4.52 | 4.71 | | | | | -4.80E+00 | Neutral | | | -4.80E+00 | Neutral |
| 61 | p.Glu61Ser | | | 14189 | 2881 | 5.34 | 4.82 | | | | | -1.62E+00 | Neutral | | | -1.62E+00 | Neutral |
| 61 | p.Glu61Ile | | | 12732 | 2674 | 4.79 | 4.47 | | | | | -2.46E+00 | Neutral | | | -2.46E+00 | Neutral |
| 61 | p.Glu61Met | | | 14829 | 3250 | 5.58 | 5.43 | | | | | -2.35E+00 | Neutral | | | -2.35E+00 | Neutral |
| 61 | p.Glu61His | | | 13568 | 2686 | 5.10 | 4.49 | | | | | -1.51E+00 | Neutral | | | -1.51E+00 | Neutral |
| 61 | p.Glu61Gln | | | 13805 | 2206 | 5.19 | 3.69 | | | | | -2.53E-01 | Neutral | | | -2.53E-01 | Neutral |
| 61 | p.Glu61Pro | | | 10268 | 11949 | 3.86 | 19.97 | | | | | -5.32E+01 | Deleterious | | | -5.32E+01 | Deleterious |
| 61 | p.Glu61Leu | | | 17939 | 3142 | 6.75 | 5.25 | | | | | -2.65E-01 | Neutral | | | -2.65E-01 | Neutral |
| 61 | p.Glu61Asp | | | 12883 | 2304 | 4.85 | 3.85 | | | | | -8.32E-01 | Neutral | | | -8.32E-01 | Neutral |
| 61 | p.Glu61Glu | Synonymous | | 14218 | 2715 | 5.35 | 4.54 | | | | | -1.06E+00 | Neutral | | | -1.06E+00 | Neutral |
| 61 | p.Glu61Ala | | | 13232 | 2302 | 4.98 | 3.85 | | | | | -6.23E-01 | Neutral | | | -6.23E-01 | Neutral |
| 61 | p.Glu61Gly | | | 11310 | 1790 | 4.25 | 2.99 | | | | | -4.32E-01 | Neutral | | | -4.32E-01 | Neutral |
| 61 | p.Glu61Val | | | 14302 | 2388 | 5.38 | 3.99 | | | | | -3.46E-01 | Neutral | | | -3.46E-01 | Neutral |
| 61 | p.Glu61Tyr | | | 11817 | 2739 | 4.44 | 4.58 | | | | | -4.64E+00 | Neutral | | | -4.64E+00 | Neutral |
| 61 | p.Glu61Cys | | | 13774 | 2388 | 5.18 | 3.99 | | | | | -5.41E-01 | Neutral | | | -5.41E-01 | Neutral |
| 61 | p.Glu61Trp | | | 14023 | 2781 | 5.27 | 4.65 | | | | | -1.42E+00 | Neutral | | | -1.42E+00 | Neutral |
| 61 | p.Glu61Phe | | | 9711 | 1799 | 3.65 | 3.01 | | | | | -2.00E+00 | Neutral | | | -2.00E+00 | Neutral |
| 62 | p.Leu62Asn | | | 5284 | 1310 | 4.68 | 2.44 | 3577 | 4531 | 4.52 | 4.40 | -2.92E-03 | Neutral | -2.05E-02 | Neutral | -1.88E-04 | Neutral |
| 62 | p.Leu62Lys | | | 5818 | 2856 | 5.15 | 5.32 | 4271 | 5506 | 5.40 | 5.34 | -2.78E+00 | Neutral | -1.10E-02 | Neutral | -1.24E+00 | Neutral |
| 62 | p.Leu62Thr | | | 4915 | 1901 | 4.35 | 3.54 | 4822 | 6010 | 6.10 | 5.83 | -8.57E-01 | Neutral | -3.32E-03 | Neutral | -1.86E-01 | Neutral |
| 62 | p.Leu62Arg | | | 3035 | 2548 | 2.69 | 4.74 | 2222 | 1857 | 2.81 | 1.80 | -3.79E+01 | Indeterminate | -2.36E-04 | Neutral | -2.86E+01 | Indeterminate |
| 62 | p.Leu62Ser | | | 3653 | 1159 | 3.23 | 2.16 | 2675 | 3190 | 3.38 | 3.10 | -3.71E-01 | Neutral | -3.40E-02 | Neutral | -4.81E-02 | Neutral |
| 62 | p.Leu62Ile | | | 1275 | 513 | 1.13 | 0.96 | 1037 | 743 | 1.31 | 0.72 | -1.06E+01 | Indeterminate | -1.55E-03 | Neutral | -7.52E+00 | Indeterminate |
| 62 | p.Leu62Met | | | 7821 | 2419 | 6.92 | 4.50 | 5702 | 4735 | 7.21 | 4.59 | -1.28E-02 | Neutral | -1.57E-07 | Neutral | -5.61E-05 | Neutral |
| 62 | p.Leu62His | | | 14346 | 3742 | 12.70 | 6.97 | 10997 | 15496 | 13.91 | 15.04 | -1.03E-05 | Neutral | -1.98E-04 | Neutral | -1.50E-08 | Neutral |
| 62 | p.Leu62Gln | | | 10306 | 14558 | 9.12 | 27.11 | 6533 | 14265 | 8.26 | 13.84 | -4.91E+01 | Indeterminate | -8.48E-01 | Neutral | -2.94E+01 | Indeterminate |
| 62 | p.Leu62Pro | | | 56 | 9 | 0.05 | 0.02 | 31 | 12 | 0.04 | 0.01 | -1.58E+01 | Indeterminate | -1.19E+00 | Neutral | -1.33E+01 | Indeterminate |
| 62 | p.Leu62Leu | Synonymous | | 3953 | 1464 | 3.50 | 2.73 | 2244 | 3542 | 2.84 | 3.44 | -1.06E+00 | Neutral | -1.06E+00 | Neutral | -8.18E-01 | Neutral |
| 62 | p.Leu62Asp | | | 8215 | 3191 | 7.27 | 5.94 | 5245 | 9043 | 6.63 | 8.78 | -1.92E-01 | Neutral | -1.95E-01 | Neutral | -4.43E-02 | Neutral |
| 62 | p.Leu62Glu | | | 8076 | 3464 | 7.15 | 6.45 | 5397 | 3735 | 6.83 | 3.62 | -5.21E-01 | Neutral | -1.98E-09 | Neutral | -7.63E-02 | Neutral |
| 62 | p.Leu62Ala | | | 5822 | 2021 | 5.15 | 3.76 | 3711 | 2780 | 4.69 | 2.70 | -1.96E-01 | Neutral | -4.26E-07 | Neutral | -1.22E-02 | Neutral |
| 62 | p.Leu62Gly | | | 5101 | 2717 | 4.52 | 5.06 | 3194 | 5893 | 4.04 | 5.72 | -5.17E+00 | Neutral | -1.39E+00 | Neutral | -4.09E+00 | Neutral |
| 62 | p.Leu62Val | | | 5797 | 2364 | 5.13 | 4.40 | 4147 | 3770 | 5.25 | 3.66 | -8.45E-01 | Neutral | -1.73E-05 | Neutral | -1.80E-01 | Neutral |
| 62 | p.Leu62Tyr | | | 5132 | 3349 | 4.54 | 6.24 | 3645 | 5500 | 4.61 | 5.34 | -1.14E+01 | Indeterminate | -1.71E-01 | Neutral | -8.39E+00 | Indeterminate |
| 62 | p.Leu62Cys | | | 1257 | 423 | 1.11 | 0.79 | 849 | 1112 | 1.07 | 1.08 | -5.31E+00 | Neutral | -2.53E+00 | Neutral | -5.15E+00 | Neutral |
| 62 | p.Leu62Trp | | | 7748 | 2419 | 6.86 | 4.50 | 5120 | 7229 | 6.48 | 7.01 | -1.54E-02 | Neutral | -1.75E-02 | Neutral | -3.70E-04 | Neutral |
| 62 | p.Leu62Phe | | | 5365 | 1278 | 4.75 | 2.38 | 3646 | 4104 | 4.61 | 3.98 | -1.31E-03 | Neutral | -2.84E-03 | Neutral | -5.97E-06 | Neutral |
| 63 | p.Leu63Asn | | | 13544 | 5590 | 7.01 | 6.89 | | | | | -5.32E+01 | Deleterious | | | -5.32E+01 | Deleterious |
| 63 | p.Leu63Lys | | | 6835 | 4935 | 3.54 | 6.09 | | | | | -5.32E+01 | Deleterious | | | -5.32E+01 | Deleterious |
| 63 | p.Leu63Thr | | | 10869 | 4259 | 5.63 | 5.25 | | | | | -5.32E+01 | Deleterious | | | -5.32E+01 | Deleterious |
| 63 | p.Leu63Arg | | | 9152 | 4924 | 4.74 | 6.07 | | | | | -5.32E+01 | Deleterious | | | -5.32E+01 | Deleterious |

| | | | | | | | | | | | |
|---|---|---|---|---|---|---|---|---|---|---|---|
| 63 | p.Leu63Ser | | | 8714 | 4308 | 4.51 | 5.31 | -5.32E+01 | Deleterious | -5.32E+01 | Deleterious |
| 63 | p.Leu63Ile | | | 10567 | 462 | 5.47 | 0.57 | -5.21E+00 | Neutral | -5.21E+00 | Neutral |
| 63 | p.Leu63Met | | | 8538 | 226 | 4.42 | 0.28 | -3.02E-01 | Neutral | -3.02E-01 | Neutral |
| 63 | p.Leu63His | | | 11608 | 6109 | 6.01 | 7.53 | -5.32E+01 | Deleterious | -5.32E+01 | Deleterious |
| 63 | p.Leu63Gln | | | 7276 | 4368 | 3.77 | 5.39 | -5.32E+01 | Deleterious | -5.32E+01 | Deleterious |
| 63 | p.Leu63Pro | | | 10214 | 6547 | 5.29 | 8.07 | -5.32E+01 | Deleterious | -5.32E+01 | Deleterious |
| 63 | p.Leu63Leu | Synonymous | | 8789 | 273 | 4.55 | 0.34 | -1.06E+00 | Neutral | -1.06E+00 | Neutral |
| 63 | p.Leu63Asp | | | 6798 | 6741 | 3.52 | 8.31 | -5.32E+01 | Deleterious | -5.32E+01 | Deleterious |
| 63 | p.Leu63Glu | | | 13660 | 6285 | 7.07 | 7.75 | -5.32E+01 | Deleterious | -5.32E+01 | Deleterious |
| 63 | p.Leu63Ala | | | 9691 | 4744 | 5.02 | 5.85 | -5.32E+01 | Deleterious | -5.32E+01 | Deleterious |
| 63 | p.Leu63Gly | | | 8561 | 5277 | 4.43 | 6.51 | -5.32E+01 | Deleterious | -5.32E+01 | Deleterious |
| 63 | p.Leu63Val | | | 6705 | 689 | 3.47 | 0.85 | -5.32E+01 | Deleterious | -5.32E+01 | Deleterious |
| 63 | p.Leu63Tyr | | | 15623 | 7883 | 8.09 | 9.72 | -5.32E+01 | Deleterious | -5.32E+01 | Deleterious |
| 63 | p.Leu63Cys | | | 7726 | 791 | 4.00 | 0.98 | -5.32E+01 | Deleterious | -5.32E+01 | Deleterious |
| 63 | p.Leu63Trp | | | 11790 | 6222 | 6.11 | 7.67 | -5.32E+01 | Deleterious | -5.32E+01 | Deleterious |
| 63 | p.Leu63Phe | | | 6454 | 448 | 3.34 | 0.55 | -3.80E+01 | Indeterminate | -3.32E+01 | Indeterminate |
| 64 | p.Leu64Asn | | | 12833 | 6821 | 4.39 | 5.64 | -1.11E+01 | Indeterminate | -1.11E+01 | Indeterminate |
| 64 | p.Leu64Lys | | | 16339 | 6603 | 5.59 | 5.46 | -2.27E+00 | Neutral | -2.27E+00 | Neutral |
| 64 | p.Leu64Thr | | | 14425 | 4621 | 4.94 | 3.82 | -6.00E-01 | Neutral | -6.00E-01 | Neutral |
| 64 | p.Leu64Arg | | | 20825 | 9191 | 7.13 | 7.60 | -2.37E+00 | Neutral | -2.37E+00 | Neutral |
| 64 | p.Leu64Ser | | | 13437 | 4243 | 4.60 | 3.51 | -6.47E-01 | Neutral | -6.47E-01 | Neutral |
| 64 | p.Leu64Ile | | | 6398 | 2634 | 2.19 | 2.18 | -1.02E+01 | Indeterminate | -1.02E+01 | Indeterminate |
| 64 | p.Leu64Met | | | 11781 | 4017 | 4.03 | 3.32 | -1.56E+00 | Neutral | -1.56E+00 | Neutral |
| 64 | p.Leu64His | | | 21371 | 9431 | 7.31 | 7.80 | -2.25E+00 | Neutral | -2.25E+00 | Neutral |
| 64 | p.Leu64Gln | | | 15398 | 7296 | 5.27 | 6.03 | -5.58E+00 | Neutral | -5.58E+00 | Neutral |
| 64 | p.Leu64Pro | | | 1303 | 1822 | 0.45 | 1.51 | -5.32E+01 | Deleterious | -5.32E+01 | Deleterious |
| 64 | p.Leu64Leu | Synonymous | | 15457 | 5454 | 5.29 | 4.51 | -1.06E+00 | Neutral | -1.06E+00 | Neutral |
| 64 | p.Leu64Asp | | | 14256 | 9066 | 4.88 | 7.50 | -1.76E+01 | Indeterminate | -1.76E+01 | Indeterminate |
| 64 | p.Leu64Glu | | | 14674 | 7300 | 5.02 | 6.04 | -7.32E+00 | Indeterminate | -7.32E+00 | Indeterminate |
| 64 | p.Leu64Ala | | | 18044 | 7247 | 6.17 | 5.99 | -1.79E+00 | Neutral | -1.79E+00 | Neutral |
| 64 | p.Leu64Gly | | | 19651 | 9506 | 6.72 | 7.86 | -4.23E+00 | Neutral | -4.23E+00 | Neutral |
| 64 | p.Leu64Val | | | 10172 | 2912 | 3.48 | 2.41 | -6.18E-01 | Neutral | -6.18E-01 | Neutral |
| 64 | p.Leu64Tyr | | | 14135 | 5042 | 4.84 | 4.17 | -1.41E+00 | Neutral | -1.41E+00 | Neutral |
| 64 | p.Leu64Cys | | | 15527 | 5308 | 5.31 | 4.39 | -8.30E-01 | Neutral | -8.30E-01 | Neutral |
| 64 | p.Leu64Trp | | | 20389 | 7820 | 6.98 | 6.47 | -9.98E-01 | Neutral | -9.98E-01 | Neutral |
| 64 | p.Leu64Phe | | | 15813 | 4622 | 5.41 | 3.82 | -1.91E-01 | Neutral | -1.91E-01 | Neutral |
| 65 | p.Leu65Asn | | | 1677 | 543 | 3.60 | 3.38 | -1.20E+01 | Indeterminate | -1.20E+01 | Indeterminate |
| 65 | p.Leu65Lys | | | 2335 | 440 | 5.01 | 2.74 | -2.88E-01 | Neutral | -2.88E-01 | Neutral |
| 65 | p.Leu65Thr | | | 2434 | 656 | 5.22 | 4.09 | -3.35E+00 | Neutral | -3.35E+00 | Neutral |
| 65 | p.Leu65Arg | | | 2730 | 595 | 5.85 | 3.71 | -6.63E-01 | Neutral | -6.63E-01 | Neutral |
| 65 | p.Leu65Ser | | | 2983 | 833 | 6.40 | 5.19 | -2.83E+00 | Neutral | -2.83E+00 | Neutral |
| 65 | p.Leu65Ile | | | 1967 | 438 | 4.22 | 2.73 | -1.69E+00 | Neutral | -1.69E+00 | Neutral |
| 65 | p.Leu65Met | | | 2487 | 463 | 5.33 | 2.89 | -2.06E-01 | Neutral | -2.06E-01 | Neutral |
| 65 | p.Leu65His | | | 1410 | 419 | 3.02 | 2.61 | -1.09E+01 | Indeterminate | -1.09E+01 | Indeterminate |
| 65 | p.Leu65Gln | | | 2753 | 794 | 5.90 | 4.95 | -3.84E+00 | Neutral | -3.84E+00 | Neutral |
| 65 | p.Leu65Pro | Likely pathogenic | | 1740 | 4497 | 3.73 | 28.03 | -5.32E+01 | Deleterious | -5.32E+01 | Deleterious |
| 65 | p.Leu65Leu | Synonymous | | 2636 | 604 | 5.65 | 3.76 | -1.06E+00 | Neutral | -1.06E+00 | Neutral |
| 65 | p.Leu65Asp | | | 2598 | 572 | 5.57 | 3.57 | -8.17E-01 | Neutral | -8.17E-01 | Neutral |
| 65 | p.Leu65Glu | | | 2759 | 572 | 5.92 | 3.57 | -4.19E-01 | Neutral | -4.19E-01 | Neutral |
| 65 | p.Leu65Ala | | | 2039 | 549 | 4.37 | 3.42 | -4.46E+00 | Neutral | -4.46E+00 | Neutral |
| 65 | p.Leu65Gly | | | 2677 | 911 | 5.74 | 5.68 | -8.22E+00 | Indeterminate | -8.22E+00 | Indeterminate |
| 65 | p.Leu65Val | | | 2667 | 680 | 5.72 | 4.24 | -2.08E+00 | Neutral | -2.08E+00 | Neutral |
| 65 | p.Leu65Tyr | | | 2428 | 553 | 5.21 | 3.45 | -1.23E+00 | Neutral | -1.23E+00 | Neutral |
| 65 | p.Leu65Cys | | | 2119 | 761 | 4.54 | 4.74 | -1.31E+01 | Indeterminate | -1.31E+01 | Indeterminate |
| 65 | p.Leu65Trp | | | 2348 | 753 | 5.03 | 4.69 | -7.73E+00 | Indeterminate | -7.73E+00 | Indeterminate |
| 65 | p.Leu65Phe | | | 1851 | 410 | 3.97 | 2.56 | -1.85E+00 | Neutral | -1.85E+00 | Neutral |
| 66 | p.His66Asn | | | 11379 | 793 | 4.83 | 1.96 | -1.65E-01 | Neutral | -1.65E-01 | Neutral |
| 66 | p.His66Lys | | | 12099 | 869 | 5.14 | 2.15 | -1.83E-01 | Neutral | -1.83E-01 | Neutral |
| 66 | p.His66Thr | | | 9820 | 910 | 4.17 | 2.25 | -2.36E+00 | Neutral | -2.36E+00 | Neutral |
| 66 | p.His66Arg | | Neutral | 12631 | 1230 | 5.36 | 3.05 | -1.93E+00 | Neutral | -1.93E+00 | Neutral |
| 66 | p.His66Ser | | | 10876 | 904 | 4.62 | 2.24 | -9.28E-01 | Neutral | -9.28E-01 | Neutral |
| 66 | p.His66Ile | | | 11804 | 1490 | 5.01 | 3.69 | -7.66E+00 | Indeterminate | -7.66E+00 | Indeterminate |
| 66 | p.His66Met | | | 11778 | 860 | 5.00 | 2.13 | -2.38E-01 | Neutral | -2.38E-01 | Neutral |
| 66 | p.His66His | Synonymous | | 9315 | 749 | 3.96 | 1.86 | -1.06E+00 | Neutral | -1.06E+00 | Neutral |
| 66 | p.His66Gln | | | 13561 | 1242 | 5.76 | 3.08 | -1.10E+00 | Neutral | -1.10E+00 | Neutral |
| 66 | p.His66Pro | | Deleterious | 14437 | 20924 | 6.13 | 51.84 | -5.32E+01 | Deleterious | -5.32E+01 | Deleterious |
| 66 | p.His66Leu | | | 13015 | 1236 | 5.53 | 3.06 | -1.55E+00 | Neutral | -1.55E+00 | Neutral |
| 66 | p.His66Asp | | | 6613 | 509 | 2.81 | 1.26 | -1.70E+00 | Neutral | -1.70E+00 | Neutral |
| 66 | p.His66Glu | | | 13336 | 1439 | 5.66 | 3.56 | -3.12E+00 | Neutral | -3.12E+00 | Neutral |
| 66 | p.His66Ala | | | 12769 | 1270 | 5.42 | 3.15 | -2.15E+00 | Neutral | -2.15E+00 | Neutral |
| 66 | p.His66Gly | | | 13669 | 1191 | 5.81 | 2.95 | -7.45E-01 | Neutral | -7.45E-01 | Neutral |
| 66 | p.His66Val | | | 10655 | 1203 | 4.53 | 2.98 | -5.55E+00 | Neutral | -5.55E+00 | Neutral |
| 66 | p.His66Tyr | | | 10205 | 865 | 4.33 | 2.14 | -1.24E+00 | Neutral | -1.24E+00 | Neutral |
| 66 | p.His66Cys | | | 12241 | 683 | 5.20 | 1.69 | -5.86E-03 | Neutral | -5.86E-03 | Neutral |
| 66 | p.His66Trp | | | 14222 | 1111 | 6.04 | 2.75 | -2.50E-01 | Neutral | -2.50E-01 | Neutral |
| 66 | p.His66Phe | | | 11036 | 888 | 4.69 | 2.20 | -6.96E-01 | Neutral | -6.96E-01 | Neutral |
| 67 | p.Gly67Asn | | | 1613 | 199 | 4.19 | 1.41 | -1.34E-01 | Neutral | -1.34E-01 | Neutral |
| 67 | p.Gly67Lys | | | 1931 | 310 | 5.01 | 2.20 | -8.83E-01 | Neutral | -8.83E-01 | Neutral |
| 67 | p.Gly67Thr | | | 1987 | 510 | 5.16 | 3.62 | -9.71E+00 | Indeterminate | -9.71E+00 | Indeterminate |
| 67 | p.Gly67Arg | | Neutral | 2225 | 231 | 5.77 | 1.64 | -2.44E-03 | Neutral | -2.44E-03 | Neutral |
| 67 | p.Gly67Ser | | | 1578 | 182 | 4.09 | 1.29 | -6.56E-02 | Neutral | -6.56E-02 | Neutral |
| 67 | p.Gly67Ile | | | 1553 | 1876 | 4.03 | 13.31 | -5.32E+01 | Deleterious | -5.32E+01 | Deleterious |
| 67 | p.Gly67Met | | | 1703 | 310 | 4.42 | 2.20 | -2.62E+00 | Neutral | -2.62E+00 | Neutral |
| 67 | p.Gly67His | | | 2343 | 275 | 6.08 | 1.95 | -1.36E-02 | Neutral | -1.36E-02 | Neutral |
| 67 | p.Gly67Gln | | | 1984 | 187 | 5.15 | 1.33 | -8.48E-04 | Neutral | -8.48E-04 | Neutral |
| 67 | p.Gly67Pro | | | 2448 | 5867 | 6.35 | 41.61 | -5.32E+01 | Deleterious | -5.32E+01 | Deleterious |
| 67 | p.Gly67Leu | | | 2000 | 298 | 5.19 | 2.11 | -4.35E-01 | Neutral | -4.35E-01 | Neutral |
| 67 | p.Gly67Asp | | | 1535 | 195 | 3.98 | 1.38 | -2.19E-01 | Neutral | -2.19E-01 | Neutral |
| 67 | p.Gly67Glu | | | 1601 | 201 | 4.15 | 1.43 | -1.67E-01 | Neutral | -1.67E-01 | Neutral |
| 67 | p.Gly67Ala | | | 2068 | 206 | 5.37 | 1.46 | -1.81E-03 | Neutral | -1.81E-03 | Neutral |
| 67 | p.Gly67Gly | Synonymous | | 1598 | 247 | 4.15 | 1.75 | -1.06E+00 | Neutral | -1.06E+00 | Neutral |
| 67 | p.Gly67Val | | | 2037 | 1509 | 5.29 | 10.70 | -5.32E+01 | Deleterious | -5.32E+01 | Deleterious |
| 67 | p.Gly67Tyr | | | 2243 | 315 | 5.82 | 2.23 | -1.67E-01 | Neutral | -1.67E-01 | Neutral |
| 67 | p.Gly67Cys | | | 1798 | 317 | 4.67 | 2.25 | -1.96E+00 | Neutral | -1.96E+00 | Neutral |
| 67 | p.Gly67Trp | | | 1957 | 441 | 5.08 | 3.13 | -5.92E+00 | Indeterminate | -5.92E+00 | Indeterminate |
| 67 | p.Gly67Phe | | | 2337 | 423 | 6.06 | 3.00 | -1.32E+00 | Neutral | -1.32E+00 | Neutral |
| 68 | p.Ala68Asn | | | 10578 | 6466 | 3.32 | 4.05 | -5.32E+01 | Deleterious | -5.32E+01 | Deleterious |
| 68 | p.Ala68Lys | | | 16843 | 13015 | 5.29 | 8.16 | -5.32E+01 | Deleterious | -5.32E+01 | Deleterious |
| 68 | p.Ala68Thr | | | 20629 | 5342 | 6.48 | 3.35 | -2.88E+00 | Neutral | -2.88E+00 | Neutral |
| 68 | p.Ala68Arg | | | 9508 | 7401 | 2.99 | 4.64 | -5.32E+01 | Deleterious | -5.32E+01 | Deleterious |
| 68 | p.Ala68Ser | | | 15363 | 2849 | 4.83 | 1.79 | -5.90E-01 | Neutral | -5.90E-01 | Neutral |
| 68 | p.Ala68Ile | | | 11369 | 7162 | 3.57 | 4.49 | -5.32E+01 | Deleterious | -5.32E+01 | Deleterious |
| 68 | p.Ala68Met | | | 20851 | 11459 | 6.55 | 7.18 | -3.82E+01 | Indeterminate | -3.32E+01 | Indeterminate |
| 68 | p.Ala68His | | | 14597 | 6003 | 4.59 | 3.76 | -2.49E+01 | Indeterminate | -2.49E+01 | Indeterminate |
| 68 | p.Ala68Gln | | | 14112 | 8817 | 4.43 | 5.53 | -5.32E+01 | Deleterious | -5.32E+01 | Deleterious |
| 68 | p.Ala68Pro | | | 19019 | 12912 | 5.98 | 8.09 | -5.32E+01 | Deleterious | -5.32E+01 | Deleterious |
| 68 | p.Ala68Leu | Likely pathogenic | | 17116 | 12054 | 5.38 | 7.56 | -5.32E+01 | Deleterious | -5.32E+01 | Deleterious |
| 68 | p.Ala68Asp | | | 14679 | 6960 | 4.61 | 4.36 | -3.55E+01 | Indeterminate | -3.29E+01 | Indeterminate |
| 68 | p.Ala68Glu | | | 19547 | 9872 | 6.14 | 6.19 | -3.28E+01 | Indeterminate | -3.20E+01 | Indeterminate |
| 68 | p.Ala68Ala | Synonymous | | 13663 | 2629 | 4.29 | 1.65 | -1.06E+00 | Neutral | -1.06E+00 | Neutral |
| 68 | p.Ala68Gly | | | 15153 | 3134 | 4.76 | 1.96 | -1.38E+00 | Neutral | -1.38E+00 | Neutral |
| 68 | p.Ala68Val | | | 20388 | 6977 | 6.41 | 4.37 | -1.00E+01 | Indeterminate | -1.00E+01 | Indeterminate |
| 68 | p.Ala68Tyr | | | 10108 | 7330 | 3.18 | 4.60 | -5.32E+01 | Deleterious | -5.32E+01 | Deleterious |
| 68 | p.Ala68Cys | | | 21550 | 4671 | 6.77 | 2.93 | -8.36E-01 | Neutral | -8.36E-01 | Neutral |
| 68 | p.Ala68Trp | | | 16659 | 12867 | 5.23 | 8.07 | -5.32E+01 | Deleterious | -5.32E+01 | Deleterious |
| 68 | p.Ala68Phe | | | 16504 | 11587 | 5.19 | 7.26 | -5.32E+01 | Deleterious | -5.32E+01 | Deleterious |
| 69 | p.Glu69Asn | | | 1335 | 244 | 3.53 | 5.26 | -5.32E+01 | Deleterious | -5.32E+01 | Deleterious |
| 69 | p.Glu69Lys | | | 1741 | 274 | 4.60 | 5.91 | -3.58E+01 | Indeterminate | -3.30E+01 | Indeterminate |
| 69 | p.Glu69Thr | | | 1856 | 192 | 4.91 | 4.14 | -9.93E+00 | Indeterminate | -9.93E+00 | Indeterminate |
| 69 | p.Glu69Arg | | | 1897 | 137 | 5.02 | 2.95 | -1.75E+00 | Neutral | -1.75E+00 | Neutral |
| 69 | p.Glu69Ser | | | 2201 | 266 | 5.82 | 5.73 | -1.39E+01 | Indeterminate | -1.39E+01 | Indeterminate |
| 69 | p.Glu69Ile | | | 2528 | 228 | 6.68 | 4.91 | -3.57E+00 | Neutral | -3.57E+00 | Neutral |
| 69 | p.Glu69Met | | | 2022 | 315 | 5.35 | 6.79 | -3.09E+01 | Indeterminate | -3.07E+01 | Indeterminate |
| 69 | p.Glu69His | | | 1853 | 178 | 4.90 | 3.84 | -7.51E+00 | Indeterminate | -7.51E+00 | Indeterminate |
| 69 | p.Glu69Gln | | | 1715 | 155 | 4.53 | 3.34 | -6.51E+00 | Indeterminate | -6.51E+00 | Indeterminate |
| 69 | p.Glu69Pro | | | 1958 | 639 | 5.18 | 13.77 | -5.32E+01 | Deleterious | -5.32E+01 | Deleterious |
| 69 | p.Glu69Leu | | | 1850 | 329 | 4.89 | 7.09 | -4.53E+01 | Indeterminate | -3.32E+01 | Indeterminate |
| 69 | p.Glu69Asp | | | 1695 | 145 | 4.48 | 3.13 | -5.22E+00 | Neutral | -5.22E+00 | Neutral |
| 69 | p.Glu69Glu | Synonymous | | 1906 | 128 | 5.04 | 2.76 | -1.06E+00 | Neutral | -1.06E+00 | Neutral |
| 69 | p.Glu69Ala | | | 1875 | 102 | 4.96 | 2.20 | -1.73E-01 | Neutral | -1.73E-01 | Neutral |
| 69 | p.Glu69Gly | | | 1878 | 211 | 4.96 | 4.55 | -1.30E+01 | Indeterminate | -1.30E+01 | Indeterminate |
| 69 | p.Glu69Val | | | 2070 | 301 | 5.47 | 6.49 | -2.54E+01 | Indeterminate | -2.54E+01 | Indeterminate |
| 69 | p.Glu69Tyr | | | 1544 | 146 | 4.08 | 3.15 | -8.93E+00 | Indeterminate | -8.93E+00 | Indeterminate |
| 69 | p.Glu69Cys | | | 2103 | 199 | 5.56 | 4.29 | -5.94E+00 | Indeterminate | -5.94E+00 | Indeterminate |
| 69 | p.Glu69Trp | | | 1943 | 251 | 5.14 | 5.41 | -1.94E+01 | Indeterminate | -1.94E+01 | Indeterminate |
| 69 | p.Glu69Phe | | | 1855 | 199 | 4.90 | 4.29 | -1.13E+01 | Indeterminate | -1.13E+01 | Indeterminate |
| 70 | p.Pro70Asn | | | 1067 | 1056 | 7.80 | 6.39 | -3.10E+00 | Neutral | -3.10E+00 | Neutral |
| 70 | p.Pro70Lys | | | 292 | 407 | 2.13 | 2.46 | -4.16E+01 | Indeterminate | -3.32E+01 | Indeterminate |
| 70 | p.Pro70Thr | | | 534 | 451 | 3.90 | 2.73 | -4.42E+00 | Neutral | -4.42E+00 | Neutral |

| | | | | | | | | | | | | | | | | | |
|---|---|---|---|---|---|---|---|---|---|---|---|---|---|---|---|---|---|
| 70 | p.Pro70Arg | | | 1225 | 962 | 8.95 | 5.82 | | | | | -4.65E-01 | Neutral | | | -4.65E-01 | Neutral |
| 70 | p.Pro70Ser | | | 611 | 1001 | 4.46 | 6.06 | | | | | -3.42E+01 | Indeterminate | | | -3.26E+01 | Indeterminate |
| 70 | p.Pro70Ile | | | 868 | 595 | 6.34 | 3.60 | | | | | -3.83E-01 | Neutral | | | -3.83E-01 | Neutral |
| 70 | p.Pro70Met | | | 1191 | 437 | 8.70 | 2.65 | | | | | -1.73E-06 | Neutral | | | -1.73E-06 | Neutral |
| 70 | p.Pro70His | | | 344 | 116 | 2.51 | 0.70 | | | | | -1.74E-03 | Neutral | | | -1.74E-03 | Neutral |
| 70 | p.Pro70Gln | | | 453 | 139 | 3.31 | 0.84 | | | | | -4.43E-05 | Neutral | | | -4.43E-05 | Neutral |
| 70 | p.Pro70Pro | Synonymous | | 683 | 490 | 4.99 | 2.97 | | | | | -1.06E+00 | Neutral | | | -1.06E+00 | Neutral |
| 70 | p.Pro70Leu | | | 725 | 642 | 5.30 | 3.89 | | | | | -3.39E+00 | Neutral | | | -3.39E+00 | Neutral |
| 70 | p.Pro70Asp | | | 669 | 5207 | 4.89 | 31.52 | | | | | -5.32E+01 | Deleterious | | | -5.32E+01 | Deleterious |
| 70 | p.Pro70Glu | | | 405 | 1516 | 2.96 | 9.18 | | | | | -5.32E+01 | Deleterious | | | -5.32E+01 | Deleterious |
| 70 | p.Pro70Ala | | | 506 | 362 | 3.70 | 2.19 | | | | | -2.01E+00 | Neutral | | | -2.01E+00 | Neutral |
| 70 | p.Pro70Gly | | | 500 | 371 | 3.65 | 2.25 | | | | | -2.54E+00 | Neutral | | | -2.54E+00 | Neutral |
| 70 | p.Pro70Val | | | 703 | 347 | 5.14 | 2.10 | | | | | -1.80E-02 | Neutral | | | -1.80E-02 | Neutral |
| 70 | p.Pro70Tyr | | | 458 | 314 | 3.35 | 1.90 | | | | | -1.90E+00 | Neutral | | | -1.90E+00 | Neutral |
| 70 | p.Pro70Cys | | | 1010 | 291 | 7.38 | 1.76 | | | | | -1.48E-08 | Neutral | | | -1.46E-08 | Neutral |
| 70 | p.Pro70Trp | | | 557 | 1275 | 4.07 | 7.72 | | | | | -5.32E+01 | Deleterious | | | -5.32E+01 | Deleterious |
| 70 | p.Pro70Phe | | | 887 | 542 | 6.48 | 3.28 | | | | | -1.08E-01 | Neutral | | | -1.08E-01 | Neutral |
| 71 | p.Asn71Asn | Synonymous | | 7181 | 171 | 4.62 | 0.30 | | | | | -1.06E+00 | Neutral | | | -1.06E+00 | Neutral |
| 71 | p.Asn71Lys | | Deleterious | 8578 | 3658 | 5.51 | 6.33 | | | | | -5.32E+01 | Deleterious | | | -5.32E+01 | Deleterious |
| 71 | p.Asn71Thr | | | 7155 | 580 | 4.60 | 1.00 | | | | | -5.32E+01 | Deleterious | | | -5.32E+01 | Deleterious |
| 71 | p.Asn71Arg | | | 7444 | 2468 | 4.78 | 4.27 | | | | | -5.32E+01 | Deleterious | | | -5.32E+01 | Deleterious |
| 71 | p.Asn71Ser | Likely pathogenic | | 5704 | 4086 | 3.67 | 7.07 | | | | | -5.32E+01 | Deleterious | | | -5.32E+01 | Deleterious |
| 71 | p.Asn71Ile | | Deleterious | 7003 | 6534 | 4.50 | 11.31 | | | | | -5.32E+01 | Deleterious | | | -5.32E+01 | Deleterious |
| 71 | p.Asn71Met | | | 4089 | 487 | 2.63 | 0.84 | | | | | -5.32E+01 | Deleterious | | | -5.32E+01 | Deleterious |
| 71 | p.Asn71His | | | 7484 | 353 | 4.81 | 0.61 | | | | | -2.06E+01 | Indeterminate | | | -2.06E+01 | Indeterminate |
| 71 | p.Asn71Gln | | | 6723 | 266 | 4.32 | 0.46 | | | | | -1.35E+01 | Indeterminate | | | -1.35E+01 | Indeterminate |
| 71 | p.Asn71Pro | | | 8614 | 17352 | 5.54 | 30.02 | | | | | -5.32E+01 | Deleterious | | | -5.32E+01 | Deleterious |
| 71 | p.Asn71Leu | | | 7663 | 5928 | 4.93 | 10.26 | | | | | -5.32E+01 | Deleterious | | | -5.32E+01 | Deleterious |
| 71 | p.Asn71Asp | | | 9069 | 498 | 5.83 | 0.86 | | | | | -2.63E+01 | Indeterminate | | | -2.63E+01 | Indeterminate |
| 71 | p.Asn71Glu | | | 10749 | 485 | 6.91 | 0.84 | | | | | -1.24E+01 | Indeterminate | | | -1.24E+01 | Indeterminate |
| 71 | p.Asn71Ala | | | 11275 | 608 | 7.25 | 1.05 | | | | | -2.05E+01 | Indeterminate | | | -2.05E+01 | Indeterminate |
| 71 | p.Asn71Gly | | | 6444 | 162 | 4.14 | 0.28 | | | | | -1.93E+00 | Neutral | | | -1.93E+00 | Neutral |
| 71 | p.Asn71Val | | | 7148 | 4897 | 4.59 | 8.47 | | | | | -5.32E+01 | Deleterious | | | -5.32E+01 | Deleterious |
| 71 | p.Asn71Tyr | | | 8986 | 3770 | 5.78 | 6.52 | | | | | -5.32E+01 | Deleterious | | | -5.32E+01 | Deleterious |
| 71 | p.Asn71Cys | | | 6712 | 283 | 4.31 | 0.49 | | | | | -1.65E+01 | Indeterminate | | | -1.65E+01 | Indeterminate |
| 71 | p.Asn71Trp | | | 8041 | 1238 | 5.17 | 2.14 | | | | | -5.32E+01 | Deleterious | | | -5.32E+01 | Deleterious |
| 71 | p.Asn71Phe | | | 9509 | 3973 | 6.11 | 6.87 | | | | | -5.32E+01 | Deleterious | | | -5.32E+01 | Deleterious |
| 72 | p.Cys72Asn | | | 8802 | 378 | 5.55 | 6.25 | 3988 | 1635 | 5.17 | 4.66 | -7.98E+00 | Indeterminate | -3.88E-01 | Neutral | -5.60E+00 | Neutral |
| 72 | p.Cys72Lys | | | 6740 | 268 | 4.25 | 4.43 | 3506 | 1606 | 4.54 | 4.57 | -8.34E+00 | Indeterminate | -1.31E+00 | Neutral | -6.71E+00 | Indeterminate |
| 72 | p.Cys72Thr | | | 10096 | 417 | 6.36 | 6.90 | 4817 | 1804 | 6.24 | 5.14 | -5.63E+00 | Neutral | -7.58E-02 | Neutral | -3.40E+00 | Neutral |
| 72 | p.Cys72Arg | | | 5486 | 148 | 3.46 | 2.45 | 4246 | 1315 | 5.50 | 3.74 | -1.75E+00 | Neutral | -9.63E-03 | Neutral | -6.11E-01 | Neutral |
| 72 | p.Cys72Ser | | | 7807 | 304 | 4.92 | 5.03 | 2617 | 1397 | 3.39 | 3.98 | -6.31E+00 | Indeterminate | -5.20E+00 | Neutral | -8.34E+00 | Indeterminate |
| 72 | p.Cys72Ile | | | 7381 | 305 | 4.65 | 5.05 | 3965 | 1960 | 5.14 | 5.58 | -8.61E+00 | Indeterminate | -1.65E+00 | Neutral | -7.24E+00 | Indeterminate |
| 72 | p.Cys72Met | | | 6176 | 187 | 3.89 | 3.09 | 2324 | 1050 | 3.01 | 2.99 | -2.74E+00 | Neutral | -2.77E+00 | Neutral | -3.24E+00 | Neutral |
| 72 | p.Cys72His | | | 4117 | 134 | 2.59 | 2.22 | 1768 | 865 | 2.29 | 2.46 | -7.23E+00 | Indeterminate | -6.27E+00 | Neutral | -1.01E+01 | Indeterminate |
| 72 | p.Cys72Gln | | | 6432 | 304 | 4.05 | 5.03 | 3140 | 1134 | 4.07 | 3.23 | -1.59E+01 | Indeterminate | -2.46E-01 | Neutral | -1.26E+01 | Indeterminate |
| 72 | p.Cys72Pro | | | 9516 | 455 | 6.00 | 7.53 | 6995 | 2515 | 9.07 | 7.16 | -1.08E+01 | Indeterminate | -7.42E-03 | Neutral | -7.71E+00 | Indeterminate |
| 72 | p.Cys72Leu | | | 4547 | 198 | 2.87 | 3.28 | 1856 | 930 | 2.41 | 2.65 | -1.76E+01 | Indeterminate | -6.48E+00 | Neutral | -2.00E+01 | Indeterminate |
| 72 | p.Cys72Asp | | | 9589 | 296 | 6.04 | 4.90 | 4756 | 2676 | 6.16 | 7.62 | -1.26E+00 | Neutral | -2.50E+00 | Neutral | -1.91E+00 | Neutral |
| 72 | p.Cys72Glu | | | 10351 | 375 | 6.52 | 6.20 | 4290 | 2323 | 5.56 | 6.61 | -2.86E+00 | Neutral | -2.44E+00 | Neutral | -3.08E+00 | Neutral |
| 72 | p.Cys72Ala | | | 8098 | 266 | 5.10 | 4.40 | 2928 | 1595 | 3.79 | 4.54 | -2.61E+00 | Neutral | -4.81E+00 | Neutral | -4.81E+00 | Neutral |
| 72 | p.Cys72Gly | | | 4850 | 154 | 3.06 | 2.55 | 3719 | 984 | 4.82 | 2.80 | -5.16E+00 | Neutral | -1.42E-03 | Neutral | -2.97E+00 | Neutral |
| 72 | p.Cys72Val | | | 9374 | 377 | 5.91 | 6.24 | 5039 | 2858 | 6.53 | 8.14 | -5.58E+00 | Neutral | -2.35E+00 | Neutral | -5.23E+00 | Neutral |
| 72 | p.Cys72Tyr | | | 11042 | 374 | 6.96 | 6.19 | 5452 | 2304 | 7.07 | 6.56 | -1.70E+00 | Neutral | -1.88E-01 | Neutral | -6.82E-01 | Neutral |
| 72 | p.Cys72Cys | Synonymous | | 9589 | 289 | 6.04 | 4.78 | 3417 | 1506 | 4.43 | 4.29 | -1.06E+00 | Neutral | -1.06E+00 | Neutral | -8.18E-01 | Neutral |
| 72 | p.Cys72Trp | | | 10835 | 582 | 6.83 | 9.63 | 4896 | 2290 | 6.35 | 6.52 | -1.38E+01 | Indeterminate | -6.72E-01 | Neutral | -1.10E+01 | Indeterminate |
| 72 | p.Cys72Phe | | | 7835 | 234 | 4.94 | 3.87 | 3442 | 2376 | 4.46 | 6.76 | -1.58E+00 | Neutral | -1.01E+01 | Indeterminate | -8.51E+00 | Indeterminate |
| 73 | p.Ala73Asn | | | 6226 | 776 | 4.30 | 5.20 | 11983 | 6680 | 4.44 | 4.78 | -7.10E+00 | Indeterminate | -3.83E+00 | Neutral | -7.83E+00 | Indeterminate |
| 73 | p.Ala73Lys | | | 5464 | 431 | 3.77 | 2.89 | 9537 | 5074 | 3.54 | 3.63 | -6.66E-01 | Neutral | -4.41E+00 | Neutral | -2.90E+00 | Neutral |
| 73 | p.Ala73Thr | | | 11273 | 1232 | 7.78 | 8.26 | 21395 | 10364 | 7.93 | 7.42 | -1.26E+00 | Neutral | -4.10E-01 | Neutral | -5.62E-01 | Neutral |
| 73 | p.Ala73Arg | | | 6861 | 819 | 4.74 | 5.49 | 13885 | 6784 | 5.15 | 4.86 | -5.15E+00 | Neutral | -1.35E+00 | Neutral | -4.04E+00 | Neutral |
| 73 | p.Ala73Ser | | | 7157 | 807 | 4.94 | 5.41 | 13597 | 7093 | 5.04 | 5.08 | -3.67E+00 | Neutral | -2.13E+00 | Neutral | -3.48E+00 | Neutral |
| 73 | p.Ala73Ile | | | 6816 | 562 | 4.71 | 3.77 | 11736 | 5986 | 4.35 | 4.29 | -5.23E-01 | Neutral | -2.49E+00 | Neutral | -1.38E+00 | Neutral |
| 73 | p.Ala73Met | | | 4904 | 408 | 3.39 | 2.73 | 9903 | 5188 | 3.67 | 3.71 | -1.29E+00 | Neutral | -3.86E+00 | Neutral | -2.96E+00 | Neutral |
| 73 | p.Ala73His | | | 12140 | 1111 | 8.38 | 7.45 | 19805 | 9884 | 7.34 | 7.08 | -2.31E-01 | Neutral | -6.60E-01 | Neutral | -1.97E-01 | Neutral |
| 73 | p.Ala73Gln | | | 4919 | 518 | 3.40 | 3.47 | 8952 | 4454 | 3.32 | 3.19 | -4.81E+00 | Neutral | -3.52E+00 | Neutral | -5.57E+00 | Neutral |
| 73 | p.Ala73Pro | | | 8448 | 860 | 5.83 | 5.76 | 14367 | 8124 | 5.33 | 5.82 | -1.49E+00 | Neutral | -3.03E+00 | Neutral | -2.47E+00 | Neutral |
| 73 | p.Ala73Leu | | | 6766 | 587 | 4.67 | 3.93 | 12546 | 6520 | 4.65 | 4.67 | -8.07E-01 | Neutral | -2.44E+00 | Neutral | -1.54E+00 | Neutral |
| 73 | p.Ala73Asp | | | 4742 | 488 | 3.27 | 3.27 | 7482 | 3511 | 2.77 | 2.51 | -4.57E+00 | Neutral | -3.54E+00 | Neutral | -5.39E+00 | Neutral |
| 73 | p.Ala73Glu | | | 7289 | 687 | 5.03 | 4.60 | 13173 | 7029 | 4.88 | 5.03 | -1.23E+00 | Neutral | -2.58E+00 | Neutral | -1.95E+00 | Neutral |
| 73 | p.Ala73Ala | Synonymous | | 9811 | 999 | 6.77 | 6.70 | 18989 | 9945 | 7.04 | 7.12 | -1.06E+00 | Neutral | -1.06E+00 | Neutral | -8.18E-01 | Neutral |
| 73 | p.Ala73Gly | | | 7551 | 854 | 5.21 | 5.72 | 15359 | 8099 | 5.70 | 5.80 | -3.41E+00 | Neutral | -1.78E+00 | Neutral | -2.99E+00 | Neutral |
| 73 | p.Ala73Val | | | 7585 | 867 | 5.24 | 5.81 | 16194 | 7473 | 6.01 | 5.35 | -3.57E+00 | Neutral | -6.00E-01 | Neutral | -2.21E+00 | Neutral |
| 73 | p.Ala73Tyr | | | 8714 | 958 | 6.02 | 6.42 | 14375 | 8705 | 5.33 | 6.23 | -2.25E+00 | Neutral | -4.27E+00 | Neutral | -4.06E+00 | Neutral |
| 73 | p.Ala73Cys | | | 7123 | 790 | 4.92 | 5.30 | 13784 | 7017 | 5.11 | 5.02 | -3.40E+00 | Neutral | -1.79E+00 | Neutral | -2.99E+00 | Neutral |
| 73 | p.Ala73Trp | | | 5446 | 663 | 3.76 | 4.44 | 11835 | 6004 | 4.39 | 4.30 | -7.72E+00 | Indeterminate | -2.37E+00 | Neutral | -7.09E+00 | Indeterminate |
| 73 | p.Ala73Phe | | | 5632 | 502 | 3.89 | 3.36 | 10777 | 5759 | 4.00 | 4.12 | -1.51E+00 | Neutral | -3.70E+00 | Neutral | -3.00E+00 | Neutral |
| 74 | p.Asp74Asn | | | 1709 | 639 | 4.77 | 1.32 | 1637 | 440 | 4.33 | 2.22 | -1.61E+01 | Indeterminate | -2.63E+01 | Indeterminate | -5.32E+01 | Deleterious |
| 74 | p.Asp74Lys | | | 2080 | 5773 | 5.81 | 11.95 | 2112 | 2212 | 5.59 | 11.17 | -5.32E+01 | Deleterious | -5.32E+01 | Deleterious | -5.32E+01 | Deleterious |
| 74 | p.Asp74Thr | | | 1807 | 2120 | 5.05 | 4.39 | 2075 | 1153 | 5.49 | 5.82 | -5.32E+01 | Deleterious | -5.32E+01 | Deleterious | -5.32E+01 | Deleterious |
| 74 | p.Asp74Arg | | | 1651 | 4359 | 4.61 | 9.02 | 1825 | 1757 | 4.83 | 8.87 | -5.32E+01 | Deleterious | -5.32E+01 | Deleterious | -5.32E+01 | Deleterious |
| 74 | p.Asp74Ser | | | 1832 | 475 | 5.12 | 0.98 | 1970 | 439 | 5.21 | 2.22 | -3.50E+00 | Neutral | -1.26E+01 | Indeterminate | -1.25E+01 | Indeterminate |
| 74 | p.Asp74Ile | | | 1665 | 5131 | 4.65 | 10.62 | 1904 | 1506 | 5.04 | 7.60 | -5.32E+01 | Deleterious | -5.32E+01 | Deleterious | -5.32E+01 | Deleterious |
| 74 | p.Asp74Met | | | 2121 | 2922 | 5.92 | 6.05 | 2291 | 1390 | 6.06 | 7.02 | -5.32E+01 | Deleterious | -5.32E+01 | Deleterious | -5.32E+01 | Deleterious |
| 74 | p.Asp74His | | Neutral | 1436 | 376 | 4.01 | 0.78 | 1554 | 225 | 4.11 | 1.14 | -5.38E+00 | Neutral | -2.84E+00 | Neutral | -5.47E+00 | Neutral |
| 74 | p.Asp74Gln | | | 1572 | 2339 | 4.39 | 4.84 | 1722 | 962 | 4.56 | 4.86 | -5.32E+01 | Deleterious | -5.32E+01 | Deleterious | -5.32E+01 | Deleterious |
| 74 | p.Asp74Pro | | | 1306 | 3867 | 3.65 | 8.00 | 1439 | 1350 | 3.81 | 6.82 | -5.32E+01 | Deleterious | -5.32E+01 | Deleterious | -5.32E+01 | Deleterious |
| 74 | p.Asp74Leu | | | 2179 | 3931 | 6.09 | 8.14 | 2204 | 1598 | 5.83 | 8.07 | -5.32E+01 | Deleterious | -5.32E+01 | Deleterious | -5.32E+01 | Deleterious |
| 74 | p.Asp74Asp | Synonymous | | 2293 | 526 | 6.40 | 1.09 | 2368 | 337 | 6.26 | 1.70 | -1.06E+00 | Neutral | -1.06E+00 | Neutral | -8.18E-01 | Neutral |
| 74 | p.Asp74Glu | | | 1910 | 219 | 5.33 | 0.45 | 1855 | 108 | 4.91 | 0.55 | -2.35E-04 | Neutral | -4.72E-06 | Neutral | -1.99E-08 | Neutral |
| 74 | p.Asp74Ala | | Deleterious | 1340 | 933 | 3.74 | 1.93 | 1438 | 505 | 3.80 | 2.55 | -5.32E+01 | Deleterious | -5.32E+01 | Deleterious | -5.32E+01 | Deleterious |
| 74 | p.Asp74Gly | | | 1450 | 722 | 4.05 | 1.49 | 1509 | 400 | 3.99 | 2.02 | -3.98E+01 | Indeterminate | -2.72E+01 | Indeterminate | -5.32E+01 | Deleterious |
| 74 | p.Asp74Val | | | 2034 | 4350 | 5.68 | 9.00 | 2153 | 1594 | 5.70 | 8.05 | -5.32E+01 | Deleterious | -5.32E+01 | Deleterious | -5.32E+01 | Deleterious |
| 74 | p.Asp74Tyr | | Deleterious | 1872 | 2840 | 5.23 | 5.88 | 1987 | 1081 | 5.26 | 5.46 | -5.32E+01 | Deleterious | -5.32E+01 | Deleterious | -5.32E+01 | Deleterious |
| 74 | p.Asp74Cys | | | 1766 | 266 | 4.93 | 0.55 | 1743 | 175 | 4.61 | 0.88 | -3.63E-02 | Neutral | -1.08E-01 | Neutral | -6.79E-03 | Neutral |
| 74 | p.Asp74Trp | | | 2243 | 3647 | 6.26 | 7.55 | 2314 | 1334 | 6.12 | 6.73 | -5.32E+01 | Deleterious | -5.32E+01 | Deleterious | -5.32E+01 | Deleterious |
| 74 | p.Asp74Phe | | | 1538 | 2881 | 4.30 | 5.96 | 1700 | 1243 | 4.50 | 6.27 | -5.32E+01 | Deleterious | -5.32E+01 | Deleterious | -5.32E+01 | Deleterious |
| 75 | p.Pro75Asn | | | 9670 | 524 | 4.87 | 4.68 | 11723 | 3674 | 4.98 | 5.59 | -1.84E+00 | Neutral | -2.33E+00 | Neutral | -2.21E+00 | Neutral |
| 75 | p.Pro75Lys | | | 9337 | 609 | 4.70 | 5.43 | 11391 | 3490 | 4.84 | 5.31 | -5.18E+00 | Neutral | -2.16E+00 | Neutral | -4.73E+00 | Neutral |
| 75 | p.Pro75Thr | | | 11192 | 685 | 5.64 | 6.11 | 12660 | 3467 | 5.38 | 5.27 | -2.81E+00 | Neutral | -7.98E-01 | Neutral | -1.80E+00 | Neutral |
| 75 | p.Pro75Arg | | | 9880 | 459 | 4.98 | 4.10 | 10942 | 3084 | 4.65 | 4.69 | -5.75E-01 | Neutral | -1.39E+00 | Neutral | -7.26E-01 | Neutral |
| 75 | p.Pro75Ser | | | 10276 | 621 | 5.18 | 5.54 | 11437 | 2762 | 4.86 | 4.20 | -3.06E+00 | Neutral | -3.66E-01 | Neutral | -1.67E+00 | Neutral |
| 75 | p.Pro75Ile | | | 9905 | 491 | 4.99 | 4.38 | 11194 | 3667 | 4.76 | 5.57 | -9.52E-01 | Neutral | -3.23E+00 | Neutral | -2.22E+00 | Neutral |
| 75 | p.Pro75Met | | | 9184 | 602 | 4.63 | 5.37 | 11731 | 3461 | 4.99 | 5.26 | -5.43E+00 | Neutral | -1.61E+00 | Neutral | -4.48E+00 | Neutral |
| 75 | p.Pro75His | | | 9528 | 472 | 4.80 | 4.21 | 10826 | 2310 | 4.60 | 3.51 | -1.04E+00 | Neutral | -1.23E-01 | Neutral | -3.11E-01 | Neutral |
| 75 | p.Pro75Gln | | | 8492 | 532 | 4.28 | 4.75 | 11101 | 3043 | 4.72 | 4.63 | -4.98E+00 | Neutral | -1.11E+00 | Neutral | -3.70E+00 | Neutral |
| 75 | p.Pro75Pro | Synonymous | | 11272 | 592 | 5.68 | 5.28 | 11825 | 3289 | 5.03 | 5.00 | -1.06E+00 | Neutral | -1.06E+00 | Neutral | -8.18E-01 | Neutral |
| 75 | p.Pro75Leu | | | 8173 | 440 | 4.12 | 3.93 | 10218 | 2734 | 4.34 | 4.16 | -2.46E+00 | Neutral | -1.14E+00 | Neutral | -1.79E+00 | Neutral |
| 75 | p.Pro75Asp | | | 7922 | 428 | 3.99 | 3.82 | 9264 | 2396 | 3.94 | 3.64 | -2.66E+00 | Neutral | -1.13E+00 | Neutral | -1.93E+00 | Neutral |
| 75 | p.Pro75Glu | | | 10198 | 576 | 5.14 | 5.14 | 12698 | 3162 | 5.40 | 4.81 | -2.12E+00 | Neutral | -3.52E-01 | Neutral | -1.03E+00 | Neutral |
| 75 | p.Pro75Ala | | | 10907 | 615 | 5.49 | 5.49 | 13939 | 3795 | 5.93 | 5.77 | -1.83E+00 | Neutral | -5.89E-01 | Neutral | -1.00E+00 | Neutral |
| 75 | p.Pro75Gly | | | 8363 | 444 | 4.21 | 3.96 | 10388 | 2646 | 4.42 | 4.02 | -2.17E+00 | Neutral | -7.57E-01 | Neutral | -1.33E+00 | Neutral |
| 75 | p.Pro75Val | | | 11370 | 636 | 5.73 | 5.67 | 14557 | 4466 | 6.19 | 6.79 | -1.60E+00 | Neutral | -1.30E+00 | Neutral | -1.31E+00 | Neutral |
| 75 | p.Pro75Tyr | | | 11247 | 621 | 5.66 | 5.54 | 12810 | 3790 | 5.45 | 5.76 | -1.50E+00 | Neutral | -1.36E+00 | Neutral | -1.28E+00 | Neutral |
| 75 | p.Pro75Cys | | | 9823 | 570 | 4.95 | 5.09 | 11358 | 2913 | 4.83 | 4.43 | -2.66E+00 | Neutral | -6.31E-01 | Neutral | -1.58E+00 | Neutral |
| 75 | p.Pro75Trp | | | 11015 | 691 | 5.55 | 6.17 | 13000 | 4054 | 5.53 | 6.16 | -3.29E+00 | Neutral | -1.85E+00 | Neutral | -2.95E+00 | Neutral |
| 75 | p.Pro75Phe | | | 10783 | 600 | 5.43 | 5.35 | 12116 | 3578 | 5.15 | 5.44 | -1.73E+00 | Neutral | -1.51E+00 | Neutral | -1.54E+00 | Neutral |
| 76 | p.Ala76Asn | | | 1971 | 8478 | 4.41 | 5.28 | | | | | -1.34E+01 | Indeterminate | | | -1.34E+01 | Indeterminate |
| 76 | p.Ala76Lys | | | 1935 | 7710 | 4.33 | 4.80 | | | | | -1.05E+01 | Indeterminate | | | -1.05E+01 | Indeterminate |
| 76 | p.Ala76Thr | | | 2338 | 7340 | 5.23 | 4.57 | | | | | -2.83E+00 | Neutral | | | -2.83E+00 | Neutral |
| 76 | p.Ala76Arg | | | 1828 | 7490 | 4.09 | 4.66 | | | | | -1.24E+01 | Indeterminate | | | -1.24E+01 | Indeterminate |
| 76 | p.Ala76Ser | | | 2564 | 10119 | 5.73 | 6.30 | | | | | -7.14E+00 | Indeterminate | | | -7.14E+00 | Indeterminate |
| 76 | p.Ala76Ile | | | 1198 | 4992 | 2.68 | 3.11 | | | | | -2.02E+01 | Indeterminate | | | -2.02E+01 | Indeterminate |
| 76 | p.Ala76Met | | | 2897 | 9059 | 6.48 | 5.64 | | | | | -1.83E+00 | Neutral | | | -1.83E+00 | Neutral |
| 76 | p.Ala76His | | | 2955 | 11104 | 6.61 | 6.91 | | | | | -4.71E+00 | Neutral | | | -4.71E+00 | Neutral |
| 76 | p.Ala76Gln | | | 1360 | 5492 | 3.04 | 3.42 | | | | | -1.62E+01 | Indeterminate | | | -1.62E+01 | Indeterminate |
| 76 | p.Ala76Pro | | | 2850 | 9900 | 6.37 | 6.16 | | | | | -3.41E+00 | Neutral | | | -3.41E+00 | Neutral |
| 76 | p.Ala76Leu | | | 1821 | 6818 | 4.07 | 4.24 | | | | | -9.05E+00 | Indeterminate | | | -9.05E+00 | Indeterminate |
| 76 | p.Ala76Asp | | | 2380 | 8354 | 5.32 | 5.20 | | | | | -4.79E+00 | Neutral | | | -4.79E+00 | Neutral |
| 76 | p.Ala76Glu | | | 2284 | 9100 | 5.11 | 5.66 | | | | | -8.61E+00 | Indeterminate | | | -8.61E+00 | Indeterminate |
| 76 | p.Ala76Ala | Synonymous | | 1395 | 3155 | 3.12 | 1.96 | | | | | -1.06E+00 | Neutral | | | -1.06E+00 | Neutral |
| 76 | p.Ala76Gly | | | 2810 | 9192 | 6.28 | 5.72 | | | | | -2.53E+00 | Neutral | | | -2.53E+00 | Neutral |
| 76 | p.Ala76Val | | | 1949 | 6091 | 4.36 | 3.79 | | | | | -3.78E+00 | Neutral | | | -3.78E+00 | Neutral |
| 76 | p.Ala76Tyr | | | 2465 | 8009 | 5.51 | 4.99 | | | | | -3.09E+00 | Neutral | | | -3.09E+00 | Neutral |
| 76 | p.Ala76Cys | | | 2123 | 9589 | 4.75 | 5.97 | | | | | -1.45E+01 | Indeterminate | | | -1.45E+01 | Indeterminate |
| 76 | p.Ala76Trp | | | 3462 | 10823 | 7.74 | 6.74 | | | | | -1.24E+00 | Neutral | | | -1.24E+00 | Neutral |
| 76 | p.Ala76Phe | | | 2136 | 7846 | 4.78 | 4.88 | | | | | -6.82E+00 | Indeterminate | | | -6.82E+00 | Indeterminate |
| 77 | p.Thr77Asn | | | 10090 | 1639 | 5.69 | 5.55 | | | | | -4.52E+00 | Neutral | | | -4.52E+00 | Neutral |
| 77 | p.Thr77Lys | | | 12148 | 2087 | 6.85 | 7.06 | | | | | -4.40E+00 | Neutral | | | -4.40E+00 | Neutral |
| 77 | p.Thr77Thr | Synonymous | | 6814 | 757 | 3.84 | 2.56 | | | | | -1.06E+00 | Neutral | | | -1.06E+00 | Neutral |

| | | | | | | | | | | | |
|---|---|---|---|---|---|---|---|---|---|---|---|
| 77 | p.Thr77Arg | | | 7484 | 1416 | 4.22 | 4.79 | -1.23E+01 | Indeterminate | -1.23E+01 | Indeterminate |
| 77 | p.Thr77Ser | | | 8715 | 1463 | 4.91 | 4.95 | -6.48E+00 | Indeterminate | -6.48E+00 | Indeterminate |
| 77 | p.Thr77Ile | | | 9404 | 1326 | 5.30 | 4.49 | -2.46E+00 | Neutral | -2.46E+00 | Neutral |
| 77 | p.Thr77Met | | | 5988 | 879 | 3.37 | 2.97 | -6.23E+00 | Indeterminate | -6.23E+00 | Indeterminate |
| 77 | p.Thr77His | | | 7330 | 1092 | 4.13 | 3.69 | -4.95E+00 | Neutral | -4.95E+00 | Neutral |
| 77 | p.Thr77Gln | | | 8278 | 1295 | 4.66 | 4.38 | -5.14E+00 | Neutral | -5.14E+00 | Neutral |
| 77 | p.Thr77Pro | | Deleterious | 7110 | 2118 | 4.01 | 7.17 | -4.41E+01 | Indeterminate | -3.32E+01 | Indeterminate |
| 77 | p.Thr77Leu | | | 11158 | 1918 | 6.29 | 6.49 | -5.03E+00 | Neutral | -5.03E+00 | Neutral |
| 77 | p.Thr77Asp | | | 4915 | 701 | 2.77 | 2.37 | -7.25E+00 | Indeterminate | -7.25E+00 | Indeterminate |
| 77 | p.Thr77Glu | | | 9748 | 2211 | 5.49 | 7.48 | -1.66E+01 | Indeterminate | -1.66E+01 | Indeterminate |
| 77 | p.Thr77Ala | | | 9298 | 1602 | 5.24 | 5.42 | -6.61E+00 | Indeterminate | -6.61E+00 | Indeterminate |
| 77 | p.Thr77Gly | | | 12368 | 1745 | 6.97 | 5.90 | -1.41E+00 | Neutral | -1.41E+00 | Neutral |
| 77 | p.Thr77Val | | | 11478 | 2244 | 6.47 | 7.59 | -8.27E+00 | Indeterminate | -8.27E+00 | Indeterminate |
| 77 | p.Thr77Tyr | | | 9931 | 1465 | 5.60 | 4.96 | -2.86E+00 | Neutral | -2.86E+00 | Neutral |
| 77 | p.Thr77Cys | | | 13304 | 2046 | 7.50 | 6.92 | -2.08E+00 | Neutral | -2.08E+00 | Neutral |
| 77 | p.Thr77Trp | | | 6374 | 853 | 3.59 | 2.89 | -3.70E+00 | Neutral | -3.70E+00 | Neutral |
| 77 | p.Thr77Phe | | | 5533 | 701 | 3.12 | 2.37 | -3.56E+00 | Neutral | -3.56E+00 | Neutral |
| 78 | p.Leu78Asn | | | 2675 | 1771 | 7.71 | 6.34 | -4.47E-03 | Neutral | -4.47E-03 | Neutral |
| 78 | p.Leu78Lys | | | 1995 | 1493 | 5.75 | 5.34 | -1.01E-01 | Neutral | -1.01E-01 | Neutral |
| 78 | p.Leu78Thr | | | 1668 | 1264 | 4.81 | 4.52 | -2.23E-01 | Neutral | -2.23E-01 | Neutral |
| 78 | p.Leu78Arg | | | 943 | 1294 | 2.72 | 4.63 | -1.69E+01 | Indeterminate | -1.69E+01 | Indeterminate |
| 78 | p.Leu78Ser | | | 1338 | 660 | 3.86 | 2.36 | -1.46E-03 | Neutral | -1.46E-03 | Neutral |
| 78 | p.Leu78Ile | | | 2099 | 1782 | 6.05 | 6.38 | -3.23E-01 | Neutral | -3.23E-01 | Neutral |
| 78 | p.Leu78Met | | | 1020 | 1289 | 2.94 | 4.61 | -1.20E+01 | Indeterminate | -1.20E+01 | Indeterminate |
| 78 | p.Leu78His | | | 1588 | 1438 | 4.58 | 5.15 | -1.17E+00 | Neutral | -1.17E+00 | Neutral |
| 78 | p.Leu78Gln | | | 1438 | 1676 | 4.14 | 6.00 | -5.66E+00 | Neutral | -5.66E+00 | Neutral |
| 78 | p.Leu78Pro | | | 1788 | 2655 | 5.15 | 9.50 | -1.10E+01 | Indeterminate | -1.10E+01 | Indeterminate |
| 78 | p.Leu78Leu | Synonymous | | 1779 | 1650 | 5.13 | 5.90 | -1.06E+00 | Neutral | -1.06E+00 | Neutral |
| 78 | p.Leu78Asp | | | 5455 | 2299 | 15.72 | 8.23 | -5.61E-10 | Neutral | -4.16E-10 | Neutral |
| 78 | p.Leu78Glu | | | 1427 | 1642 | 4.11 | 5.88 | -5.41E+00 | Neutral | -5.41E+00 | Neutral |
| 78 | p.Leu78Ala | | | 637 | 727 | 1.84 | 2.60 | -1.43E+01 | Indeterminate | -1.43E+01 | Indeterminate |
| 78 | p.Leu78Gly | | | 1290 | 219 | 3.72 | 0.78 | -1.12E-15 | Neutral | 0.00E+00 | Neutral |
| 78 | p.Leu78Val | | | 2096 | 1560 | 6.04 | 5.58 | -7.80E-02 | Neutral | -7.80E-02 | Neutral |
| 78 | p.Leu78Tyr | | | 1478 | 1553 | 4.26 | 5.56 | -3.31E+00 | Neutral | -3.31E+00 | Neutral |
| 78 | p.Leu78Cys | | | 1343 | 1282 | 3.87 | 4.59 | -2.33E+00 | Neutral | -2.33E+00 | Neutral |
| 78 | p.Leu78Trp | | | 1194 | 791 | 3.44 | 2.83 | -1.75E-01 | Neutral | -1.75E-01 | Neutral |
| 78 | p.Leu78Phe | | | 1456 | 900 | 4.20 | 3.22 | -3.38E-02 | Neutral | -3.38E-02 | Neutral |
| 79 | p.Thr79Asn | | | 6357 | 2367 | 6.19 | 4.94 | -3.23E-01 | Neutral | -3.23E-01 | Neutral |
| 79 | p.Thr79Lys | | | 4790 | 2080 | 4.67 | 4.34 | -2.10E+00 | Neutral | -2.10E+00 | Neutral |
| 79 | p.Thr79Thr | Synonymous | | 7615 | 3475 | 7.42 | 7.25 | -1.06E+00 | Neutral | -1.06E+00 | Neutral |
| 79 | p.Thr79Arg | | | 3106 | 1753 | 3.03 | 3.66 | -1.26E+01 | Indeterminate | -1.26E+01 | Indeterminate |
| 79 | p.Thr79Ser | | | 5296 | 2013 | 5.16 | 4.20 | -6.60E-01 | Neutral | -6.60E-01 | Neutral |
| 79 | p.Thr79Ile | | | 5523 | 1971 | 5.38 | 4.11 | -3.36E-01 | Neutral | -3.36E-01 | Neutral |
| 79 | p.Thr79Met | | | 6304 | 2761 | 6.14 | 5.76 | -1.24E+00 | Neutral | -1.24E+00 | Neutral |
| 79 | p.Thr79His | | | 5309 | 2284 | 5.17 | 4.77 | -1.61E+00 | Neutral | -1.61E+00 | Neutral |
| 79 | p.Thr79Gln | | | 4412 | 1628 | 4.30 | 3.40 | -8.42E-01 | Neutral | -8.42E-01 | Neutral |
| 79 | p.Thr79Pro | | | 3460 | 4139 | 3.37 | 8.64 | -5.32E+01 | Deleterious | -5.32E+01 | Deleterious |
| 79 | p.Thr79Leu | | | 4103 | 1982 | 4.00 | 4.14 | -4.80E+00 | Neutral | -4.80E+00 | Neutral |
| 79 | p.Thr79Asp | | | 3999 | 1769 | 3.90 | 3.69 | -3.25E+00 | Neutral | -3.25E+00 | Neutral |
| 79 | p.Thr79Glu | | | 3681 | 1420 | 3.59 | 2.96 | -1.74E+00 | Neutral | -1.74E+00 | Neutral |
| 79 | p.Thr79Ala | | | 5581 | 2518 | 5.44 | 5.25 | -1.95E+00 | Neutral | -1.95E+00 | Neutral |
| 79 | p.Thr79Gly | | | 5518 | 2596 | 5.38 | 5.42 | -2.54E+00 | Neutral | -2.54E+00 | Neutral |
| 79 | p.Thr79Val | | | 8644 | 4107 | 8.42 | 8.57 | -1.05E+00 | Neutral | -1.05E+00 | Neutral |
| 79 | p.Thr79Tyr | | | 3452 | 1556 | 3.36 | 3.25 | -4.54E+00 | Neutral | -4.54E+00 | Neutral |
| 79 | p.Thr79Cys | | | 5814 | 2910 | 5.66 | 6.07 | -3.24E+00 | Neutral | -3.24E+00 | Neutral |
| 79 | p.Thr79Trp | | | 4719 | 2447 | 4.60 | 5.11 | -5.36E+00 | Neutral | -5.36E+00 | Neutral |
| 79 | p.Thr79Phe | | | 4968 | 2150 | 4.84 | 4.49 | -1.91E+00 | Neutral | -1.91E+00 | Neutral |
| 80 | p.Arg80Asn | | | 9628 | 1551 | 4.97 | 5.34 | -1.78E+01 | Indeterminate | -1.78E+01 | Indeterminate |
| 80 | p.Arg80Lys | | | 11473 | 1355 | 5.92 | 4.67 | -4.55E+00 | Neutral | -4.55E+00 | Neutral |
| 80 | p.Arg80Thr | | | 5402 | 616 | 2.79 | 2.12 | -1.09E+01 | Indeterminate | -1.09E+01 | Indeterminate |
| 80 | p.Arg80Arg | Synonymous | | 13295 | 1283 | 6.86 | 4.42 | -1.06E+00 | Neutral | -1.06E+00 | Neutral |
| 80 | p.Arg80Ser | | | 12287 | 1392 | 6.34 | 4.80 | -3.31E+00 | Neutral | -3.31E+00 | Neutral |
| 80 | p.Arg80Ile | | | 7889 | 1862 | 4.07 | 6.41 | -5.32E+01 | Deleterious | -5.32E+01 | Deleterious |
| 80 | p.Arg80Met | | | 8839 | 1240 | 4.56 | 4.27 | -1.26E+01 | Indeterminate | -1.26E+01 | Indeterminate |
| 80 | p.Arg80His | | | 9521 | 1203 | 4.92 | 4.14 | -7.91E+00 | Indeterminate | -7.91E+00 | Indeterminate |
| 80 | p.Arg80Gln | | | 7688 | 1142 | 3.97 | 3.93 | -1.75E+01 | Indeterminate | -1.75E+01 | Indeterminate |
| 80 | p.Arg80Pro | | | 8976 | 4035 | 4.63 | 13.90 | -5.32E+01 | Deleterious | -5.32E+01 | Deleterious |
| 80 | p.Arg80Leu | | | 10680 | 1517 | 5.51 | 5.23 | -1.06E+01 | Indeterminate | -1.06E+01 | Indeterminate |
| 80 | p.Arg80Asp | | | 10178 | 1582 | 5.26 | 5.45 | -1.51E+01 | Indeterminate | -1.51E+01 | Indeterminate |
| 80 | p.Arg80Glu | | | 9465 | 1058 | 4.89 | 3.64 | -4.74E+00 | Neutral | -4.74E+00 | Neutral |
| 80 | p.Arg80Ala | | | 11000 | 1400 | 5.68 | 4.82 | -6.72E+00 | Indeterminate | -6.72E+00 | Indeterminate |
| 80 | p.Arg80Gly | | | 10632 | 1370 | 5.49 | 4.72 | -7.39E+00 | Indeterminate | -7.39E+00 | Indeterminate |
| 80 | p.Arg80Val | | | 8051 | 1272 | 4.16 | 4.38 | -2.01E+01 | Indeterminate | -2.01E+01 | Indeterminate |
| 80 | p.Arg80Tyr | | | 11274 | 1590 | 5.82 | 5.48 | -9.69E+00 | Indeterminate | -9.69E+00 | Indeterminate |
| 80 | p.Arg80Cys | | | 6897 | 589 | 3.56 | 2.03 | -2.00E+00 | Neutral | -2.00E+00 | Neutral |
| 80 | p.Arg80Trp | | | 11399 | 1615 | 5.89 | 5.56 | -9.72E+00 | Indeterminate | -9.72E+00 | Indeterminate |
| 80 | p.Arg80Phe | | | 9095 | 1358 | 4.70 | 4.68 | -1.50E+01 | Indeterminate | -1.50E+01 | Indeterminate |
| 81 | p.Pro81Asn | | | 3270 | 2138 | 4.49 | 5.03 | -5.32E+01 | Deleterious | -5.32E+01 | Deleterious |
| 81 | p.Pro81Lys | | | 4056 | 3564 | 5.57 | 8.39 | -5.32E+01 | Deleterious | -5.32E+01 | Deleterious |
| 81 | p.Pro81Thr | | Deleterious | 3644 | 1370 | 5.01 | 3.23 | -4.95E+01 | Indeterminate | -3.32E+01 | Indeterminate |
| 81 | p.Pro81Arg | | | 4210 | 2742 | 5.79 | 6.46 | -5.32E+01 | Deleterious | -5.32E+01 | Deleterious |
| 81 | p.Pro81Ser | | | 2453 | 1122 | 3.37 | 2.64 | -5.32E+01 | Deleterious | -5.32E+01 | Deleterious |
| 81 | p.Pro81Ile | | | 2461 | 1198 | 3.38 | 2.82 | -5.32E+01 | Deleterious | -5.32E+01 | Deleterious |
| 81 | p.Pro81Met | | | 4378 | 2646 | 6.02 | 6.23 | -5.32E+01 | Deleterious | -5.32E+01 | Deleterious |
| 81 | p.Pro81His | | | 3889 | 2773 | 5.35 | 6.53 | -5.32E+01 | Deleterious | -5.32E+01 | Deleterious |
| 81 | p.Pro81Gln | | | 4314 | 4646 | 5.93 | 10.94 | -5.32E+01 | Deleterious | -5.32E+01 | Deleterious |
| 81 | p.Pro81Pro | Synonymous | | 3937 | 545 | 5.41 | 1.28 | -1.06E+00 | Neutral | -1.06E+00 | Neutral |
| 81 | p.Pro81Leu | | Deleterious | 2688 | 1382 | 3.69 | 3.25 | -5.32E+01 | Deleterious | -5.32E+01 | Deleterious |
| 81 | p.Pro81Asp | | | 3627 | 2528 | 4.99 | 5.95 | -5.32E+01 | Deleterious | -5.32E+01 | Deleterious |
| 81 | p.Pro81Glu | | | 3955 | 3393 | 5.44 | 7.99 | -5.32E+01 | Deleterious | -5.32E+01 | Deleterious |
| 81 | p.Pro81Ala | | | 2546 | 590 | 3.50 | 1.39 | -2.01E+01 | Indeterminate | -2.01E+01 | Indeterminate |
| 81 | p.Pro81Gly | | | 3018 | 1629 | 4.15 | 3.84 | -5.32E+01 | Deleterious | -5.32E+01 | Deleterious |
| 81 | p.Pro81Val | | | 3803 | 1377 | 5.23 | 3.24 | -4.42E+01 | Indeterminate | -3.32E+01 | Indeterminate |
| 81 | p.Pro81Tyr | | | 3466 | 2061 | 4.76 | 4.85 | -5.32E+01 | Deleterious | -5.32E+01 | Deleterious |
| 81 | p.Pro81Cys | | | 5033 | 1452 | 6.92 | 3.42 | -1.97E+01 | Indeterminate | -1.97E+01 | Indeterminate |
| 81 | p.Pro81Trp | | | 4048 | 2847 | 5.56 | 6.70 | -5.32E+01 | Deleterious | -5.32E+01 | Deleterious |
| 81 | p.Pro81Phe | | | 3961 | 2474 | 5.44 | 5.82 | -5.32E+01 | Deleterious | -5.32E+01 | Deleterious |
| 82 | p.Val82Asn | | | 5118 | 927 | 3.84 | 4.94 | -5.32E+01 | Deleterious | -5.32E+01 | Deleterious |
| 82 | p.Val82Lys | | | 4194 | 977 | 3.15 | 5.20 | -5.32E+01 | Deleterious | -5.32E+01 | Deleterious |
| 82 | p.Val82Thr | | | 11017 | 1270 | 8.28 | 6.76 | -8.29E+00 | Indeterminate | -8.29E+00 | Indeterminate |
| 82 | p.Val82Arg | | | 8316 | 1978 | 6.25 | 10.53 | -5.32E+01 | Deleterious | -5.32E+01 | Deleterious |
| 82 | p.Val82Ser | | | 8583 | 897 | 6.45 | 4.78 | -7.82E+00 | Indeterminate | -7.82E+00 | Indeterminate |
| 82 | p.Val82Ile | | | 7271 | 717 | 5.46 | 3.82 | -7.73E+00 | Indeterminate | -7.73E+00 | Indeterminate |
| 82 | p.Val82Met | | | 5075 | 324 | 3.81 | 1.73 | -1.59E+00 | Neutral | -1.59E+00 | Neutral |
| 82 | p.Val82His | | | 8957 | 1689 | 6.73 | 9.00 | -4.23E+01 | Indeterminate | -3.32E+01 | Indeterminate |
| 82 | p.Val82Gln | | | 6300 | 686 | 4.73 | 3.65 | -1.31E+01 | Indeterminate | -1.31E+01 | Indeterminate |
| 82 | p.Val82Pro | | | 7701 | 1207 | 5.78 | 6.43 | -3.07E+01 | Indeterminate | -3.05E+01 | Indeterminate |
| 82 | p.Val82Leu | | | 6686 | 586 | 5.02 | 3.12 | -5.30E+00 | Neutral | -5.30E+00 | Neutral |
| 82 | p.Val82Asp | | | 3241 | 694 | 2.43 | 3.70 | -5.32E+01 | Deleterious | -5.32E+01 | Deleterious |
| 82 | p.Val82Glu | | | 6491 | 1072 | 4.88 | 5.71 | -3.99E+01 | Indeterminate | -3.32E+01 | Indeterminate |
| 82 | p.Val82Ala | | | 8058 | 625 | 6.05 | 3.33 | -2.05E+00 | Neutral | -2.05E+00 | Neutral |
| 82 | p.Val82Gly | | | 4572 | 717 | 3.43 | 3.82 | -4.61E+01 | Indeterminate | -3.32E+01 | Indeterminate |
| 82 | p.Val82Val | Synonymous | | 10662 | 821 | 8.01 | 4.37 | -1.06E+00 | Neutral | -1.06E+00 | Neutral |
| 82 | p.Val82Tyr | | | 3733 | 830 | 2.80 | 4.42 | -5.32E+01 | Deleterious | -5.32E+01 | Deleterious |
| 82 | p.Val82Cys | | | 4968 | 509 | 3.73 | 2.71 | -1.39E+01 | Indeterminate | -1.39E+01 | Indeterminate |
| 82 | p.Val82Trp | | | 4187 | 1143 | 3.14 | 6.09 | -5.32E+01 | Deleterious | -5.32E+01 | Deleterious |
| 82 | p.Val82Phe | | | 8005 | 1107 | 6.01 | 5.90 | -2.13E+01 | Indeterminate | -2.13E+01 | Indeterminate |
| 83 | p.His83Asn | | | 2209 | 1716 | 5.20 | 5.30 | -5.32E+01 | Deleterious | -5.32E+01 | Deleterious |
| 83 | p.His83Lys | | | 2282 | 2477 | 5.37 | 7.65 | -5.32E+01 | Deleterious | -5.32E+01 | Deleterious |
| 83 | p.His83Thr | | | 1704 | 1097 | 4.01 | 3.39 | -5.32E+01 | Deleterious | -5.32E+01 | Deleterious |
| 83 | p.His83Arg | Pathogenic | | 2229 | 2555 | 5.25 | 7.89 | -5.32E+01 | Deleterious | -5.32E+01 | Deleterious |
| 83 | p.His83Ser | | | 2121 | 1323 | 4.99 | 4.09 | -5.32E+01 | Deleterious | -5.32E+01 | Deleterious |
| 83 | p.His83Ile | | | 2239 | 1501 | 5.27 | 4.64 | -5.32E+01 | Deleterious | -5.32E+01 | Deleterious |
| 83 | p.His83Met | | | 1978 | 448 | 4.66 | 1.38 | -3.07E+01 | Indeterminate | -3.05E+01 | Indeterminate |
| 83 | p.His83His | Synonymous | | 2145 | 220 | 5.05 | 0.68 | -1.06E+00 | Neutral | -1.06E+00 | Neutral |
| 83 | p.His83Gln | | | 2092 | 858 | 4.92 | 2.65 | -5.32E+01 | Deleterious | -5.32E+01 | Deleterious |
| 83 | p.His83Pro | | | 2039 | 2411 | 4.80 | 7.45 | -5.32E+01 | Deleterious | -5.32E+01 | Deleterious |
| 83 | p.His83Leu | | | 2061 | 1244 | 4.85 | 3.84 | -5.32E+01 | Deleterious | -5.32E+01 | Deleterious |
| 83 | p.His83Asp | Likely pathogenic | | 1956 | 2148 | 4.60 | 6.63 | -5.32E+01 | Deleterious | -5.32E+01 | Deleterious |
| 83 | p.His83Glu | | | 2438 | 2516 | 5.74 | 7.77 | -5.32E+01 | Deleterious | -5.32E+01 | Deleterious |
| 83 | p.His83Ala | | | 2190 | 589 | 5.15 | 1.82 | -4.26E+01 | Indeterminate | -3.32E+01 | Indeterminate |
| 83 | p.His83Gly | | | 2738 | 2060 | 6.44 | 6.36 | -5.32E+01 | Deleterious | -5.32E+01 | Deleterious |
| 83 | p.His83Val | | | 1653 | 1644 | 3.89 | 5.08 | -5.32E+01 | Deleterious | -5.32E+01 | Deleterious |
| 83 | p.His83Tyr | Likely pathogenic | | 1726 | 1905 | 4.06 | 5.88 | -5.32E+01 | Deleterious | -5.32E+01 | Deleterious |
| 83 | p.His83Cys | | | 1753 | 1232 | 4.13 | 3.80 | -5.32E+01 | Deleterious | -5.32E+01 | Deleterious |
| 83 | p.His83Trp | | | 2279 | 2636 | 5.36 | 8.14 | -5.32E+01 | Deleterious | -5.32E+01 | Deleterious |
| 83 | p.His83Phe | | | 2658 | 1802 | 6.26 | 5.56 | -5.32E+01 | Deleterious | -5.32E+01 | Deleterious |
| 84 | p.Asp84Asn | Likely pathogenic | | 2322 | 945 | 3.70 | 3.51 | -5.32E+01 | Deleterious | -5.32E+01 | Deleterious |
| 84 | p.Asp84Lys | | | 3482 | 2031 | 5.55 | 7.54 | -5.32E+01 | Deleterious | -5.32E+01 | Deleterious |

| | | | | | | | | | | | |
|---|---|---|---|---|---|---|---|---|---|---|---|
| 84 | p.Asp84Thr | | | 2464 | 1033 | 3.92 | 3.83 | -5.32E+01 | Deleterious | -5.32E+01 | Deleterious |
| 84 | p.Asp84Arg | | | 3099 | 1815 | 4.94 | 6.73 | -5.32E+01 | Deleterious | -5.32E+01 | Deleterious |
| 84 | p.Asp84Ser | | | 2805 | 443 | 4.47 | 1.64 | -5.32E+01 | Deleterious | -5.32E+01 | Deleterious |
| 84 | p.Asp84Ile | | | 3566 | 2122 | 5.68 | 7.87 | -5.32E+01 | Deleterious | -5.32E+01 | Deleterious |
| 84 | p.Asp84Met | | | 2901 | 1475 | 4.62 | 5.47 | -5.32E+01 | Deleterious | -5.32E+01 | Deleterious |
| 84 | p.Asp84His | | Deleterious | 3544 | 821 | 5.64 | 3.05 | -5.32E+01 | Deleterious | -5.32E+01 | Deleterious |
| 84 | p.Asp84Gln | | | 2535 | 744 | 4.04 | 2.76 | -5.32E+01 | Deleterious | -5.32E+01 | Deleterious |
| 84 | p.Asp84Pro | | | 3263 | 1893 | 5.20 | 7.02 | -5.32E+01 | Deleterious | -5.32E+01 | Deleterious |
| 84 | p.Asp84Leu | | | 3617 | 2184 | 5.76 | 8.10 | -5.32E+01 | Deleterious | -5.32E+01 | Deleterious |
| 84 | p.Asp84Asp | Synonymous | | 3069 | 79 | 4.89 | 0.29 | -1.06E+00 | Neutral | -1.06E+00 | Neutral |
| 84 | p.Asp84Glu | | | 2431 | 94 | 3.87 | 0.35 | -1.16E+01 | Indeterminate | -1.16E+01 | Indeterminate |
| 84 | p.Asp84Ala | | Deleterious | 4167 | 1843 | 6.64 | 6.84 | -5.32E+01 | Deleterious | -5.32E+01 | Deleterious |
| 84 | p.Asp84Gly | | Deleterious | 2603 | 1289 | 4.15 | 4.78 | -5.32E+01 | Deleterious | -5.32E+01 | Deleterious |
| 84 | p.Asp84Val | | Deleterious | 2863 | 1540 | 4.56 | 5.71 | -5.32E+01 | Deleterious | -5.32E+01 | Deleterious |
| 84 | p.Asp84Tyr | | Deleterious | 3406 | 1751 | 5.42 | 6.50 | -5.32E+01 | Deleterious | -5.32E+01 | Deleterious |
| 84 | p.Asp84Cys | | | 3373 | 512 | 5.37 | 1.90 | -5.32E+01 | Deleterious | -5.32E+01 | Deleterious |
| 84 | p.Asp84Trp | | | 3522 | 2243 | 5.61 | 8.32 | -5.32E+01 | Deleterious | -5.32E+01 | Deleterious |
| 84 | p.Asp84Phe | | | 3758 | 2096 | 5.99 | 7.78 | -5.32E+01 | Deleterious | -5.32E+01 | Deleterious |
| 85 | p.Ala85Asn | | | 5212 | 1526 | 2.82 | 2.47 | -1.83E+01 | Indeterminate | -1.83E+01 | Indeterminate |
| 85 | p.Ala85Lys | | | 9383 | 4778 | 5.07 | 7.73 | -4.54E+01 | Indeterminate | -3.32E+01 | Indeterminate |
| 85 | p.Ala85Thr | | | 7427 | 2426 | 4.01 | 3.92 | -1.78E+01 | Indeterminate | -1.78E+01 | Indeterminate |
| 85 | p.Ala85Arg | | | 8838 | 4487 | 4.78 | 7.26 | -4.72E+01 | Indeterminate | -3.32E+01 | Indeterminate |
| 85 | p.Ala85Ser | | | 10574 | 2582 | 5.72 | 4.18 | -3.75E+00 | Neutral | -3.75E+00 | Neutral |
| 85 | p.Ala85Ile | | | 11678 | 4130 | 6.31 | 6.68 | -1.43E+01 | Indeterminate | -1.43E+01 | Indeterminate |
| 85 | p.Ala85Met | | | 6476 | 1133 | 3.50 | 1.83 | -1.40E+00 | Neutral | -1.40E+00 | Neutral |
| 85 | p.Ala85His | | | 7202 | 2718 | 3.89 | 4.40 | -2.74E+01 | Indeterminate | -2.74E+01 | Indeterminate |
| 85 | p.Ala85Gln | | | 8493 | 2892 | 4.59 | 4.68 | -1.77E+01 | Indeterminate | -1.77E+01 | Indeterminate |
| 85 | p.Ala85Pro | | | 10997 | 3768 | 5.94 | 6.09 | -1.38E+01 | Indeterminate | -1.38E+01 | Indeterminate |
| 85 | p.Ala85Leu | | | 10857 | 2524 | 5.87 | 4.08 | -2.77E+00 | Neutral | -2.77E+00 | Neutral |
| 85 | p.Ala85Asp | | | 7167 | 3294 | 3.87 | 5.33 | -4.41E+01 | Indeterminate | -3.32E+01 | Indeterminate |
| 85 | p.Ala85Glu | | | 7645 | 3405 | 4.13 | 5.51 | -3.91E+01 | Indeterminate | -3.32E+01 | Indeterminate |
| 85 | p.Ala85Ala | Synonymous | | 15969 | 3621 | 8.63 | 5.86 | -1.06E+00 | Neutral | -1.06E+00 | Neutral |
| 85 | p.Ala85Gly | | | 13936 | 2718 | 7.53 | 4.40 | -4.57E-01 | Neutral | -4.57E-01 | Neutral |
| 85 | p.Ala85Val | | | 10776 | 3252 | 5.82 | 5.26 | -9.08E+00 | Indeterminate | -9.08E+00 | Indeterminate |
| 85 | p.Ala85Tyr | | | 6635 | 2743 | 3.59 | 4.44 | -3.67E+01 | Indeterminate | -3.31E+01 | Indeterminate |
| 85 | p.Ala85Cys | | | 9406 | 1933 | 5.08 | 3.13 | -1.78E+00 | Neutral | -1.78E+00 | Neutral |
| 85 | p.Ala85Trp | | | 6098 | 3377 | 3.30 | 5.46 | -5.32E+01 | Deleterious | -5.32E+01 | Deleterious |
| 85 | p.Ala85Phe | | | 10242 | 4529 | 5.54 | 7.32 | -3.04E+01 | Indeterminate | -3.02E+01 | Indeterminate |
| 86 | p.Ala86Asn | | | 2738 | 7263 | 5.42 | 6.88 | -5.32E+01 | Deleterious | -5.32E+01 | Deleterious |
| 86 | p.Ala86Lys | | | 1712 | 5339 | 3.39 | 5.06 | -5.32E+01 | Deleterious | -5.32E+01 | Deleterious |
| 86 | p.Ala86Thr | | | 1173 | 286 | 2.32 | 0.27 | -6.25E-02 | Neutral | -6.25E-02 | Neutral |
| 86 | p.Ala86Arg | | | 2701 | 8049 | 5.35 | 7.62 | -5.32E+01 | Deleterious | -5.32E+01 | Deleterious |
| 86 | p.Ala86Ser | | | 1742 | 753 | 3.45 | 0.71 | -2.71E+00 | Neutral | -2.71E+00 | Neutral |
| 86 | p.Ala86Ile | | | 2571 | 1478 | 5.09 | 1.40 | -5.80E+00 | Neutral | -5.80E+00 | Indeterminate |
| 86 | p.Ala86Met | | | 2759 | 6416 | 5.46 | 6.08 | -5.32E+01 | Deleterious | -5.32E+01 | Deleterious |
| 86 | p.Ala86His | | | 3907 | 10251 | 7.74 | 9.71 | -5.32E+01 | Deleterious | -5.32E+01 | Deleterious |
| 86 | p.Ala86Gln | | | 3641 | 10410 | 7.21 | 9.86 | -5.32E+01 | Deleterious | -5.32E+01 | Deleterious |
| 86 | p.Ala86Pro | | | 3199 | 8498 | 6.34 | 8.05 | -5.32E+01 | Deleterious | -5.32E+01 | Deleterious |
| 86 | p.Ala86Leu | | | 3102 | 7242 | 6.14 | 6.86 | -5.32E+01 | Deleterious | -5.32E+01 | Deleterious |
| 86 | p.Ala86Asp | | | 2882 | 7986 | 5.71 | 7.56 | -5.32E+01 | Deleterious | -5.32E+01 | Deleterious |
| 86 | p.Ala86Glu | | | 3145 | 8838 | 6.23 | 8.37 | -5.32E+01 | Deleterious | -5.32E+01 | Deleterious |
| 86 | p.Ala86Ala | Synonymous | | 986 | 304 | 1.95 | 0.29 | -1.06E+00 | Neutral | -1.06E+00 | Neutral |
| 86 | p.Ala86Gly | | | 1964 | 522 | 3.89 | 0.49 | -1.98E-02 | Neutral | -1.98E-02 | Neutral |
| 86 | p.Ala86Val | | | 2113 | 649 | 4.18 | 0.61 | -9.64E-02 | Neutral | -9.64E-02 | Neutral |
| 86 | p.Ala86Tyr | | | 2032 | 5898 | 4.02 | 5.59 | -5.32E+01 | Deleterious | -5.32E+01 | Deleterious |
| 86 | p.Ala86Cys | | | 2481 | 507 | 4.91 | 0.48 | -4.00E-05 | Neutral | -4.00E-05 | Neutral |
| 86 | p.Ala86Trp | | | 2661 | 6920 | 5.27 | 6.55 | -5.32E+01 | Deleterious | -5.32E+01 | Deleterious |
| 86 | p.Ala86Phe | | | 2986 | 7970 | 5.91 | 7.55 | -5.32E+01 | Deleterious | -5.32E+01 | Deleterious |
| 87 | p.Arg87Asn | | | 7612 | 236 | 5.72 | 1.26 | -3.51E+00 | Neutral | -3.51E+00 | Neutral |
| 87 | p.Arg87Lys | | | 3377 | 37 | 2.54 | 0.20 | -7.53E-04 | Neutral | -7.53E-04 | Neutral |
| 87 | p.Arg87Thr | | | 4254 | 127 | 3.20 | 0.68 | -7.09E+00 | Indeterminate | -7.09E+00 | Indeterminate |
| 87 | p.Arg87Arg | Synonymous | | 5869 | 137 | 5.16 | 1.03 | -1.06E+00 | Neutral | -1.06E+00 | Neutral |
| 87 | p.Arg87Ser | | | 3450 | 208 | 2.59 | 1.11 | -5.32E+01 | Deleterious | -5.32E+01 | Deleterious |
| 87 | p.Arg87Ile | | | 4046 | 129 | 3.04 | 0.69 | -9.73E+00 | Indeterminate | -9.73E+00 | Indeterminate |
| 87 | p.Arg87Met | | | 7168 | 345 | 5.38 | 1.84 | -1.99E+01 | Indeterminate | -1.99E+01 | Indeterminate |
| 87 | p.Arg87His | | | 4551 | 247 | 3.42 | 1.32 | -3.99E+01 | Indeterminate | -3.32E+01 | Indeterminate |
| 87 | p.Arg87Gln | | | 7646 | 452 | 5.74 | 2.41 | -3.25E+01 | Indeterminate | -3.18E+01 | Indeterminate |
| 87 | p.Arg87Pro | Pathogenic | | 3416 | 3365 | 2.57 | 17.92 | -5.32E+01 | Deleterious | -5.32E+01 | Deleterious |
| 87 | p.Arg87Leu | | | 4276 | 160 | 3.21 | 0.85 | -1.57E+01 | Indeterminate | -1.57E+01 | Indeterminate |
| 87 | p.Arg87Asp | | | 4569 | 777 | 3.43 | 4.14 | -5.32E+01 | Deleterious | -5.32E+01 | Deleterious |
| 87 | p.Arg87Glu | | | 6786 | 626 | 5.10 | 3.33 | -5.32E+01 | Deleterious | -5.32E+01 | Deleterious |
| 87 | p.Arg87Ala | | | 7639 | 562 | 5.74 | 2.99 | -5.30E+01 | Indeterminate | -3.32E+01 | Indeterminate |
| 87 | p.Arg87Gly | | | 5541 | 127 | 4.16 | 0.68 | -1.07E+00 | Neutral | -1.07E+00 | Neutral |
| 87 | p.Arg87Val | | | 6100 | 198 | 4.58 | 1.05 | -6.11E+00 | Indeterminate | -6.11E+00 | Indeterminate |
| 87 | p.Arg87Tyr | | | 2406 | 123 | 1.81 | 0.66 | -5.32E+01 | Deleterious | -5.32E+01 | Deleterious |
| 87 | p.Arg87Cys | | | 8971 | 188 | 6.74 | 1.00 | -1.06E-01 | Neutral | -1.06E-01 | Neutral |
| 87 | p.Arg87Trp | Likely pathogenic | | 8616 | 4952 | 6.47 | 26.37 | -5.32E+01 | Deleterious | -5.32E+01 | Deleterious |
| 87 | p.Arg87Phe | | | 7490 | 267 | 5.63 | 1.42 | -6.79E+00 | Indeterminate | -6.79E+00 | Indeterminate |
| 88 | p.Glu88Asn | | | 7680 | 5284 | 4.42 | 4.15 | -8.96E+00 | Indeterminate | -8.96E+00 | Indeterminate |
| 88 | p.Glu88Lys | | Deleterious | 9106 | 9400 | 5.25 | 7.39 | -2.66E+01 | Indeterminate | -2.65E+01 | Indeterminate |
| 88 | p.Glu88Thr | | | 5283 | 3622 | 3.04 | 2.85 | -1.37E+01 | Indeterminate | -1.37E+01 | Indeterminate |
| 88 | p.Glu88Arg | | | 9435 | 9436 | 5.44 | 7.42 | -2.37E+01 | Indeterminate | -2.37E+01 | Indeterminate |
| 88 | p.Glu88Ser | | | 6793 | 4359 | 3.91 | 3.43 | -8.02E+00 | Indeterminate | -8.02E+00 | Indeterminate |
| 88 | p.Glu88Ile | | | 7738 | 4822 | 4.46 | 3.79 | -5.97E+00 | Indeterminate | -5.97E+00 | Indeterminate |
| 88 | p.Glu88Met | | | 9862 | 6110 | 5.68 | 4.80 | -4.02E+00 | Neutral | -4.02E+00 | Neutral |
| 88 | p.Glu88His | | | 6938 | 5151 | 4.00 | 4.05 | -1.32E+01 | Indeterminate | -1.32E+01 | Indeterminate |
| 88 | p.Glu88Gln | | | 7089 | 4779 | 4.08 | 3.76 | -9.17E+00 | Indeterminate | -9.17E+00 | Indeterminate |
| 88 | p.Glu88Pro | | | 6230 | 10321 | 3.59 | 8.11 | -5.32E+01 | Deleterious | -5.32E+01 | Deleterious |
| 88 | p.Glu88Leu | | | 7368 | 5329 | 4.24 | 4.19 | -1.13E+01 | Indeterminate | -1.13E+01 | Indeterminate |
| 88 | p.Glu88Asp | | | 8787 | 6308 | 5.06 | 4.96 | -8.89E+00 | Indeterminate | -8.89E+00 | Indeterminate |
| 88 | p.Glu88Glu | Synonymous | | 12109 | 6412 | 6.98 | 5.04 | -1.06E+00 | Neutral | -1.06E+00 | Neutral |
| 88 | p.Glu88Ala | | | 6884 | 3920 | 3.97 | 3.08 | -4.75E+00 | Neutral | -4.75E+00 | Neutral |
| 88 | p.Glu88Gly | | | 8821 | 5602 | 5.08 | 4.40 | -5.36E+00 | Neutral | -5.36E+00 | Neutral |
| 88 | p.Glu88Val | | | 16505 | 11466 | 9.51 | 9.01 | -3.02E+00 | Neutral | -3.02E+00 | Neutral |
| 88 | p.Glu88Tyr | | | 9528 | 6744 | 5.49 | 5.30 | -7.59E+00 | Indeterminate | -7.59E+00 | Indeterminate |
| 88 | p.Glu88Cys | | | 8363 | 5109 | 4.82 | 4.02 | -4.88E+00 | Neutral | -4.88E+00 | Neutral |
| 88 | p.Glu88Trp | | | 10256 | 6510 | 5.91 | 5.12 | -4.25E+00 | Neutral | -4.25E+00 | Neutral |
| 88 | p.Glu88Phe | | | 8799 | 6563 | 5.07 | 5.16 | -1.02E+01 | Indeterminate | -1.02E+01 | Indeterminate |
| 89 | p.Gly89Asn | | | 2627 | 478 | 3.60 | 0.67 | -1.70E+01 | Indeterminate | -1.70E+01 | Indeterminate |
| 89 | p.Gly89Lys | | | 4247 | 3480 | 5.82 | 4.87 | -5.32E+01 | Deleterious | -5.32E+01 | Deleterious |
| 89 | p.Gly89Thr | | | 3807 | 5129 | 5.22 | 7.18 | -5.32E+01 | Deleterious | -5.32E+01 | Deleterious |
| 89 | p.Gly89Arg | | | 3355 | 3087 | 4.60 | 4.32 | -5.32E+01 | Deleterious | -5.32E+01 | Deleterious |
| 89 | p.Gly89Ser | | | 3824 | 1590 | 5.24 | 2.23 | -5.32E+01 | Deleterious | -5.32E+01 | Deleterious |
| 89 | p.Gly89Ile | | | 3356 | 5608 | 4.60 | 7.85 | -5.32E+01 | Deleterious | -5.32E+01 | Deleterious |
| 89 | p.Gly89Met | | | 3081 | 3692 | 4.22 | 5.17 | -5.32E+01 | Deleterious | -5.32E+01 | Deleterious |
| 89 | p.Gly89His | | | 3103 | 3096 | 4.25 | 4.33 | -5.32E+01 | Deleterious | -5.32E+01 | Deleterious |
| 89 | p.Gly89Gln | | | 3330 | 2815 | 4.56 | 3.94 | -5.32E+01 | Deleterious | -5.32E+01 | Deleterious |
| 89 | p.Gly89Pro | | | 4288 | 4970 | 5.87 | 6.96 | -5.32E+01 | Deleterious | -5.32E+01 | Deleterious |
| 89 | p.Gly89Leu | | | 3791 | 5275 | 5.19 | 7.38 | -5.32E+01 | Deleterious | -5.32E+01 | Deleterious |
| 89 | p.Gly89Asp | | Deleterious | 3504 | 1266 | 4.80 | 1.77 | -5.32E+01 | Deleterious | -5.32E+01 | Deleterious |
| 89 | p.Gly89Glu | | | 4701 | 4959 | 6.44 | 6.94 | -5.32E+01 | Deleterious | -5.32E+01 | Deleterious |
| 89 | p.Gly89Ala | | | 2968 | 631 | 4.07 | 0.88 | -2.36E+01 | Indeterminate | -2.36E+01 | Indeterminate |
| 89 | p.Gly89Gly | Synonymous | | 2868 | 294 | 3.93 | 0.41 | -1.06E+00 | Neutral | -1.06E+00 | Neutral |
| 89 | p.Gly89Val | | | 3667 | 5622 | 5.02 | 7.87 | -5.32E+01 | Deleterious | -5.32E+01 | Deleterious |
| 89 | p.Gly89Tyr | | | 3624 | 5208 | 4.96 | 7.29 | -5.32E+01 | Deleterious | -5.32E+01 | Deleterious |
| 89 | p.Gly89Cys | | | 4087 | 1699 | 5.60 | 2.38 | -5.32E+01 | Deleterious | -5.32E+01 | Deleterious |
| 89 | p.Gly89Trp | | | 4212 | 6145 | 5.77 | 8.60 | -5.32E+01 | Deleterious | -5.32E+01 | Deleterious |
| 89 | p.Gly89Phe | | | 4558 | 6406 | 6.24 | 8.97 | -5.32E+01 | Deleterious | -5.32E+01 | Deleterious |
| 90 | p.Phe90Asn | | | 17215 | 3468 | 7.19 | 6.33 | -7.65E-02 | Neutral | -7.65E-02 | Neutral |
| 90 | p.Phe90Lys | | | 12298 | 3370 | 5.14 | 6.15 | -2.63E+00 | Neutral | -2.63E+00 | Neutral |
| 90 | p.Phe90Thr | | | 8757 | 2286 | 3.66 | 4.17 | -3.69E+00 | Neutral | -3.69E+00 | Neutral |
| 90 | p.Phe90Arg | | | 14675 | 3186 | 6.13 | 5.81 | -3.08E-01 | Neutral | -3.08E-01 | Neutral |
| 90 | p.Phe90Ser | | | 12195 | 2607 | 5.09 | 4.76 | -4.70E-01 | Neutral | -4.70E-01 | Neutral |
| 90 | p.Phe90Ile | | | 9858 | 1941 | 4.12 | 3.54 | -4.24E-01 | Neutral | -4.24E-01 | Neutral |
| 90 | p.Phe90Met | | | 9884 | 1676 | 4.13 | 3.06 | -8.85E-02 | Neutral | -8.85E-02 | Neutral |
| 90 | p.Phe90His | | | 9400 | 1661 | 3.93 | 3.03 | -1.71E-01 | Neutral | -1.71E-01 | Neutral |
| 90 | p.Phe90Gln | | | 11418 | 1955 | 4.77 | 3.57 | -5.51E-02 | Neutral | -5.51E-02 | Neutral |
| 90 | p.Phe90Pro | | | 13543 | 4974 | 5.66 | 9.08 | -8.66E+00 | Indeterminate | -8.66E+00 | Indeterminate |
| 90 | p.Phe90Leu | | | 12336 | 2179 | 5.15 | 3.98 | -5.91E-02 | Neutral | -5.91E-02 | Neutral |
| 90 | p.Phe90Asp | | | 8808 | 3457 | 3.68 | 6.31 | -1.75E+01 | Indeterminate | -1.75E+01 | Indeterminate |
| 90 | p.Phe90Glu | | | 12532 | 3273 | 5.23 | 5.97 | -1.91E+00 | Neutral | -1.91E+00 | Neutral |
| 90 | p.Phe90Ala | | | 9217 | 1762 | 3.85 | 3.22 | -3.98E-01 | Neutral | -3.98E-01 | Neutral |
| 90 | p.Phe90Gly | | | 12639 | 2896 | 5.28 | 5.28 | -7.58E-01 | Neutral | -7.58E-01 | Neutral |
| 90 | p.Phe90Val | | | 12878 | 3230 | 5.38 | 5.89 | -1.40E+00 | Neutral | -1.40E+00 | Neutral |
| 90 | p.Phe90Tyr | | | 13813 | 2502 | 5.77 | 4.57 | -5.03E-02 | Neutral | -5.03E-02 | Neutral |
| 90 | p.Phe90Cys | | | 14267 | 3205 | 5.96 | 5.85 | -4.60E-01 | Neutral | -4.60E-01 | Neutral |
| 90 | p.Phe90Trp | | | 13085 | 2766 | 5.46 | 5.05 | -3.43E-01 | Neutral | -3.43E-01 | Neutral |
| 90 | p.Phe90Phe | Synonymous | | 10647 | 2410 | 4.45 | 4.40 | -1.06E+00 | Neutral | -1.06E+00 | Neutral |
| 91 | p.Leu91Asn | | | 8432 | 493 | 5.01 | 2.36 | -2.02E-02 | Neutral | -2.02E-02 | Neutral |

| | | | | | | | | | | | | | | | | | |
|---|---|---|---|---|---|---|---|---|---|---|---|---|---|---|---|---|---|
| 91 | p.Leu91Lys | | | 7857 | 938 | 4.67 | 4.49 | | | | | -7.34E+00 | Indeterminate | | | -7.34E+00 | Indeterminate |
| 91 | p.Leu91Thr | | | 7096 | 862 | 4.22 | 4.13 | | | | | -8.95E+00 | Indeterminate | | | -8.95E+00 | Indeterminate |
| 91 | p.Leu91Arg | | | 10604 | 1024 | 6.30 | 4.91 | | | | | -1.49E+00 | Neutral | | | -1.49E+00 | Neutral |
| 91 | p.Leu91Ser | | | 8848 | 1433 | 5.26 | 6.86 | | | | | -1.82E+01 | Indeterminate | | | -1.82E+01 | Indeterminate |
| 91 | p.Leu91Ile | | | 5518 | 782 | 3.28 | 3.75 | | | | | -1.95E+01 | Indeterminate | | | -1.95E+01 | Indeterminate |
| 91 | p.Leu91Met | | | 9789 | 2126 | 5.82 | 10.18 | | | | | -3.62E+01 | Indeterminate | | | -3.31E+01 | Indeterminate |
| 91 | p.Leu91His | | | 6249 | 874 | 3.71 | 4.19 | | | | | -1.66E+01 | Indeterminate | | | -1.66E+01 | Indeterminate |
| 91 | p.Leu91Gln | | | 7540 | 500 | 4.48 | 2.40 | | | | | -1.59E-01 | Neutral | | | -1.59E-01 | Neutral |
| 91 | p.Leu91Pro | | | 9411 | 1199 | 5.59 | 5.74 | | | | | -7.49E+00 | Indeterminate | | | -7.49E+00 | Indeterminate |
| 91 | p.Leu91Leu | Synonymous | | 7934 | 662 | 4.72 | 3.17 | | | | | -1.06E+00 | Neutral | | | -1.06E+00 | Neutral |
| 91 | p.Leu91Asp | | | 6698 | 730 | 3.98 | 3.50 | | | | | -6.28E+00 | Indeterminate | | | -6.28E+00 | Indeterminate |
| 91 | p.Leu91Glu | | | 8804 | 987 | 5.23 | 4.73 | | | | | -4.76E+00 | Neutral | | | -4.76E+00 | Neutral |
| 91 | p.Leu91Ala | | | 8686 | 744 | 5.16 | 3.56 | | | | | -1.03E+00 | Neutral | | | -1.03E+00 | Neutral |
| 91 | p.Leu91Gly | | | 9719 | 1125 | 5.78 | 5.39 | | | | | -4.74E+00 | Neutral | | | -4.74E+00 | Neutral |
| 91 | p.Leu91Val | | | 11061 | 1191 | 6.58 | 5.71 | | | | | -2.65E+00 | Neutral | | | -2.65E+00 | Neutral |
| 91 | p.Leu91Tyr | | | 5704 | 1472 | 3.39 | 7.05 | | | | | -5.32E+01 | Deleterious | | | -5.32E+01 | Deleterious |
| 91 | p.Leu91Cys | | | 8443 | 679 | 5.02 | 3.25 | | | | | -6.85E-01 | Neutral | | | -6.85E-01 | Neutral |
| 91 | p.Leu91Trp | | | 10058 | 1333 | 5.98 | 6.39 | | | | | -8.02E+00 | Indeterminate | | | -8.02E+00 | Indeterminate |
| 91 | p.Leu91Phe | | | 9772 | 1720 | 5.81 | 8.24 | | | | | -2.11E+01 | Indeterminate | | | -2.11E+01 | Indeterminate |
| 92 | p.Asp92Asn | | | 3059 | 1748 | 3.61 | 3.58 | | | | | -3.34E+00 | Neutral | | | -3.34E+00 | Neutral |
| 92 | p.Asp92Lys | | | 2980 | 1988 | 3.51 | 4.07 | | | | | -7.05E+00 | Indeterminate | | | -7.05E+00 | Indeterminate |
| 92 | p.Asp92Thr | | | 3372 | 1636 | 3.97 | 3.35 | | | | | -1.00E+00 | Neutral | | | -1.00E+00 | Neutral |
| 92 | p.Asp92Arg | | | 5018 | 2780 | 5.92 | 5.69 | | | | | -1.01E+00 | Neutral | | | -1.01E+00 | Neutral |
| 92 | p.Asp92Ser | | | 2101 | 1098 | 2.48 | 2.25 | | | | | -3.99E+00 | Neutral | | | -3.99E+00 | Neutral |
| 92 | p.Asp92Ile | | | 5343 | 3437 | 6.30 | 7.04 | | | | | -2.32E+00 | Neutral | | | -2.32E+00 | Neutral |
| 92 | p.Asp92Met | | | 5318 | 2816 | 6.27 | 5.77 | | | | | -6.09E-01 | Neutral | | | -6.09E-01 | Neutral |
| 92 | p.Asp92His | | | 2930 | 1735 | 3.45 | 3.55 | | | | | -4.28E+00 | Neutral | | | -4.28E+00 | Neutral |
| 92 | p.Asp92Gln | | | 5536 | 3118 | 6.53 | 6.39 | | | | | -8.92E-01 | Neutral | | | -8.92E-01 | Neutral |
| 92 | p.Asp92Pro | | | 5268 | 3172 | 6.21 | 6.50 | | | | | -1.59E+00 | Neutral | | | -1.59E+00 | Neutral |
| 92 | p.Asp92Leu | | | 6269 | 3445 | 7.39 | 7.06 | | | | | -5.25E-01 | Neutral | | | -5.25E-01 | Neutral |
| 92 | p.Asp92Asp | Synonymous | | 4573 | 2475 | 5.39 | 5.07 | | | | | -1.06E+00 | Neutral | | | -1.06E+00 | Neutral |
| 92 | p.Asp92Glu | | | 3137 | 1867 | 3.70 | 3.82 | | | | | -3.93E+00 | Neutral | | | -3.93E+00 | Neutral |
| 92 | p.Asp92Ala | | | 6689 | 4069 | 7.88 | 8.33 | | | | | -9.91E-01 | Neutral | | | -9.91E-01 | Neutral |
| 92 | p.Asp92Gly | | | 3137 | 1980 | 3.70 | 4.05 | | | | | -5.17E+00 | Neutral | | | -5.17E+00 | Neutral |
| 92 | p.Asp92Val | | | 5507 | 3063 | 6.49 | 6.27 | | | | | -8.22E-01 | Neutral | | | -8.22E-01 | Neutral |
| 92 | p.Asp92Tyr | | | 2568 | 1440 | 3.03 | 2.95 | | | | | -4.06E+00 | Neutral | | | -4.06E+00 | Neutral |
| 92 | p.Asp92Cys | | | 5479 | 2909 | 6.46 | 5.96 | | | | | -5.73E-01 | Neutral | | | -5.73E-01 | Neutral |
| 92 | p.Asp92Trp | | | 2854 | 1628 | 3.36 | 3.33 | | | | | -3.72E+00 | Neutral | | | -3.72E+00 | Neutral |
| 92 | p.Asp92Phe | | | 3697 | 2426 | 4.36 | 4.97 | | | | | -4.82E+00 | Neutral | | | -4.82E+00 | Neutral |
| 93 | p.Thr93Asn | | | 12872 | 1136 | 4.39 | 3.41 | | | | | -1.65E+01 | Indeterminate | | | -1.65E+01 | Indeterminate |
| 93 | p.Thr93Lys | | | 10656 | 1950 | 3.64 | 5.86 | | | | | -5.32E+01 | Deleterious | | | -5.32E+01 | Deleterious |
| 93 | p.Thr93Thr | Synonymous | | 12435 | 599 | 4.24 | 1.80 | | | | | -1.06E+00 | Neutral | | | -1.06E+00 | Neutral |
| 93 | p.Thr93Arg | | | 12522 | 1486 | 4.27 | 4.47 | | | | | -3.73E+01 | Indeterminate | | | -3.31E+01 | Indeterminate |
| 93 | p.Thr93Ser | | | 11324 | 798 | 3.86 | 2.40 | | | | | -8.91E+00 | Indeterminate | | | -8.91E+00 | Indeterminate |
| 93 | p.Thr93Ile | | | 13017 | 614 | 4.44 | 1.85 | | | | | -8.11E-01 | Neutral | | | -8.11E-01 | Neutral |
| 93 | p.Thr93Met | | | 14173 | 1206 | 4.84 | 3.62 | | | | | -1.33E+01 | Indeterminate | | | -1.33E+01 | Indeterminate |
| 93 | p.Thr93His | | | 19146 | 2259 | 6.53 | 6.79 | | | | | -2.59E+01 | Indeterminate | | | -2.59E+01 | Indeterminate |
| 93 | p.Thr93Gln | | | 17574 | 1861 | 6.00 | 5.59 | | | | | -2.08E+01 | Indeterminate | | | -2.08E+01 | Indeterminate |
| 93 | p.Thr93Pro | | | 18332 | 5525 | 6.26 | 16.61 | | | | | -5.32E+01 | Deleterious | | | -5.32E+01 | Deleterious |
| 93 | p.Thr93Leu | | | 15110 | 1179 | 5.16 | 3.54 | | | | | -9.12E+00 | Indeterminate | | | -9.12E+00 | Indeterminate |
| 93 | p.Thr93Asp | | | 11110 | 1576 | 3.79 | 4.74 | | | | | -5.32E+01 | Deleterious | | | -5.32E+01 | Deleterious |
| 93 | p.Thr93Glu | | | 5928 | 693 | 2.02 | 2.08 | | | | | -5.32E+01 | Deleterious | | | -5.32E+01 | Deleterious |
| 93 | p.Thr93Ala | | | 18371 | 2050 | 6.27 | 6.16 | | | | | -2.31E+01 | Indeterminate | | | -2.31E+01 | Indeterminate |
| 93 | p.Thr93Gly | | | 21093 | 2490 | 7.20 | 7.48 | | | | | -2.38E+01 | Indeterminate | | | -2.38E+01 | Indeterminate |
| 93 | p.Thr93Val | | | 14806 | 1033 | 5.05 | 3.10 | | | | | -5.99E+00 | Indeterminate | | | -5.99E+00 | Indeterminate |
| 93 | p.Thr93Tyr | | | 15176 | 2095 | 5.18 | 6.30 | | | | | -4.55E+01 | Indeterminate | | | -3.32E+01 | Indeterminate |
| 93 | p.Thr93Cys | | | 14861 | 1164 | 5.07 | 3.50 | | | | | -9.44E+00 | Indeterminate | | | -9.44E+00 | Indeterminate |
| 93 | p.Thr93Trp | | | 16976 | 1890 | 5.79 | 5.68 | | | | | -2.47E+01 | Indeterminate | | | -2.47E+01 | Indeterminate |
| 93 | p.Thr93Phe | | | 17510 | 1668 | 5.98 | 5.01 | | | | | -1.52E+01 | Indeterminate | | | -1.52E+01 | Indeterminate |
| 94 | p.Leu94Asn | | | 1461 | 780 | 4.07 | 2.20 | | | | | -3.44E+01 | Indeterminate | | | -3.27E+01 | Indeterminate |
| 94 | p.Leu94Lys | | | 2375 | 3419 | 6.61 | 9.64 | | | | | -5.32E+01 | Deleterious | | | -5.32E+01 | Deleterious |
| 94 | p.Leu94Thr | | | 1880 | 1076 | 5.23 | 3.03 | | | | | -3.32E+01 | Indeterminate | | | -3.22E+01 | Indeterminate |
| 94 | p.Leu94Arg | | | 2092 | 3934 | 5.82 | 11.09 | | | | | -5.32E+01 | Deleterious | | | -5.32E+01 | Deleterious |
| 94 | p.Leu94Ser | | | 1901 | 421 | 5.29 | 1.19 | | | | | -4.70E-01 | Neutral | | | -4.70E-01 | Neutral |
| 94 | p.Leu94Ile | | | 1867 | 295 | 5.20 | 0.83 | | | | | -8.36E-03 | Neutral | | | -8.36E-03 | Neutral |
| 94 | p.Leu94Met | | | 432 | 42 | 1.20 | 0.12 | | | | | -8.41E-03 | Neutral | | | -8.41E-03 | Neutral |
| 94 | p.Leu94His | | | 1805 | 3659 | 5.02 | 10.31 | | | | | -5.32E+01 | Deleterious | | | -5.32E+01 | Deleterious |
| 94 | p.Leu94Gln | Likely pathogenic | | 1288 | 1435 | 3.58 | 4.04 | | | | | -5.32E+01 | Deleterious | | | -5.32E+01 | Deleterious |
| 94 | p.Leu94Pro | Likely pathogenic | | 2142 | 2900 | 5.96 | 8.17 | | | | | -5.32E+01 | Deleterious | | | -5.32E+01 | Deleterious |
| 94 | p.Leu94Leu | Synonymous | | 1575 | 363 | 4.38 | 1.02 | | | | | -1.06E+00 | Neutral | | | -1.06E+00 | Neutral |
| 94 | p.Leu94Asp | | | 2051 | 3654 | 5.71 | 10.30 | | | | | -5.32E+01 | Deleterious | | | -5.32E+01 | Deleterious |
| 94 | p.Leu94Glu | | | 2084 | 2145 | 5.80 | 6.04 | | | | | -5.32E+01 | Deleterious | | | -5.32E+01 | Deleterious |
| 94 | p.Leu94Ala | | | 946 | 501 | 2.63 | 1.41 | | | | | -4.70E+01 | Indeterminate | | | -3.32E+01 | Indeterminate |
| 94 | p.Leu94Gly | | | 1187 | 1220 | 3.30 | 3.44 | | | | | -5.32E+01 | Deleterious | | | -5.32E+01 | Deleterious |
| 94 | p.Leu94Val | | | 1555 | 436 | 4.33 | 1.23 | | | | | -3.56E+00 | Neutral | | | -3.56E+00 | Neutral |
| 94 | p.Leu94Tyr | | | 1270 | 2783 | 3.53 | 7.84 | | | | | -5.32E+01 | Deleterious | | | -5.32E+01 | Deleterious |
| 94 | p.Leu94Cys | | | 2850 | 1202 | 7.93 | 3.39 | | | | | -8.77E+00 | Indeterminate | | | -8.77E+00 | Indeterminate |
| 94 | p.Leu94Trp | | | 2406 | 4027 | 6.70 | 11.35 | | | | | -5.32E+01 | Deleterious | | | -5.32E+01 | Deleterious |
| 94 | p.Leu94Phe | | | 2763 | 1193 | 7.69 | 3.36 | | | | | -9.94E+00 | Indeterminate | | | -9.94E+00 | Indeterminate |
| 95 | p.Val95Asn | | | 7232 | 2182 | 3.02 | 3.98 | | | | | -4.64E+00 | Neutral | | | -4.64E+00 | Neutral |
| 95 | p.Val95Lys | | | 10963 | 2844 | 4.58 | 5.19 | | | | | -8.25E-01 | Neutral | | | -8.25E-01 | Neutral |
| 95 | p.Val95Thr | | | 5808 | 1609 | 2.43 | 2.94 | | | | | -4.38E+00 | Neutral | | | -4.38E+00 | Neutral |
| 95 | p.Val95Arg | | | 9203 | 2809 | 3.84 | 5.13 | | | | | -3.32E+00 | Neutral | | | -3.32E+00 | Neutral |
| 95 | p.Val95Ser | | | 6861 | 2066 | 2.87 | 3.77 | | | | | -4.99E+00 | Neutral | | | -4.99E+00 | Neutral |
| 95 | p.Val95Ile | | | 8681 | 2394 | 3.63 | 4.37 | | | | | -2.11E+00 | Neutral | | | -2.11E+00 | Neutral |
| 95 | p.Val95Met | | | 4894 | 1367 | 2.04 | 2.49 | | | | | -5.88E+00 | Indeterminate | | | -5.88E+00 | Indeterminate |
| 95 | p.Val95His | | | 7829 | 1699 | 3.27 | 3.10 | | | | | -4.78E-01 | Neutral | | | -4.78E-01 | Neutral |
| 95 | p.Val95Gln | | | 2718 | 740 | 1.14 | 1.35 | | | | | -1.13E+01 | Indeterminate | | | -1.13E+01 | Indeterminate |
| 95 | p.Val95Pro | | | 5472 | 7300 | 2.29 | 13.32 | | | | | -5.32E+01 | Deleterious | | | -5.32E+01 | Deleterious |
| 95 | p.Val95Leu | | | 5183 | 1187 | 2.16 | 2.17 | | | | | -1.93E+00 | Neutral | | | -1.93E+00 | Neutral |
| 95 | p.Val95Asp | | | 7731 | 2073 | 3.23 | 3.78 | | | | | -2.24E+00 | Neutral | | | -2.24E+00 | Neutral |
| 95 | p.Val95Glu | | | 7407 | 2177 | 3.09 | 3.97 | | | | | -3.95E+00 | Neutral | | | -3.95E+00 | Neutral |
| 95 | p.Val95Ala | | | 6406 | 1317 | 2.68 | 2.40 | | | | | -5.32E-01 | Neutral | | | -5.32E-01 | Neutral |
| 95 | p.Val95Gly | | | 9626 | 2766 | 4.02 | 5.05 | | | | | -2.19E+00 | Neutral | | | -2.19E+00 | Neutral |
| 95 | p.Val95Val | Synonymous | | 12313 | 3434 | 5.14 | 6.27 | | | | | -1.06E+00 | Neutral | | | -1.06E+00 | Neutral |
| 95 | p.Val95Tyr | | | 5221 | 1479 | 2.18 | 2.70 | | | | | -5.69E+00 | Neutral | | | -5.69E+00 | Neutral |
| 95 | p.Val95Cys | | | 6989 | 1603 | 2.92 | 2.92 | | | | | -1.01E+00 | Neutral | | | -1.01E+00 | Neutral |
| 95 | p.Val95Trp | | | 6758 | 2279 | 2.82 | 4.16 | | | | | -8.18E+00 | Indeterminate | | | -8.18E+00 | Indeterminate |
| 95 | p.Val95Phe | | | 5986 | 1846 | 2.50 | 3.37 | | | | | -6.73E+00 | Indeterminate | | | -6.73E+00 | Indeterminate |
| 96 | p.Val96Asn | | | 3048 | 665 | 5.44 | 4.93 | 5216 | 4257 | 5.13 | 6.03 | -2.60E-01 | Neutral | -7.47E+00 | Indeterminate | -5.06E+00 | Neutral |
| 96 | p.Val96Lys | | | 4337 | 1119 | 7.74 | 8.30 | 7262 | 4566 | 7.14 | 6.47 | -4.20E-01 | Neutral | -1.03E+00 | Neutral | -4.44E-01 | Neutral |
| 96 | p.Val96Thr | | | 3524 | 706 | 6.29 | 5.23 | 5839 | 3601 | 5.74 | 5.10 | -5.74E-02 | Neutral | -1.48E+00 | Neutral | -4.91E-01 | Neutral |
| 96 | p.Val96Arg | | | 4400 | 1080 | 7.85 | 8.01 | 8595 | 5833 | 8.45 | 8.26 | -2.49E-01 | Neutral | -1.19E+00 | Neutral | -4.39E-01 | Neutral |
| 96 | p.Val96Ser | | | 3519 | 765 | 6.28 | 5.67 | 6849 | 3584 | 6.73 | 5.08 | -1.52E-01 | Neutral | -2.52E-01 | Neutral | -4.78E-02 | Neutral |
| 96 | p.Val96Ile | | | 1854 | 453 | 3.31 | 3.36 | 3334 | 2258 | 3.28 | 3.20 | -2.21E+00 | Neutral | -6.40E+00 | Neutral | -5.81E+00 | Indeterminate |
| 96 | p.Val96Met | | | 2021 | 408 | 3.61 | 3.02 | 4039 | 2200 | 3.97 | 3.12 | -4.52E-01 | Neutral | -1.46E+00 | Neutral | -6.96E-01 | Neutral |
| 96 | p.Val96His | | | 1796 | 486 | 3.21 | 3.60 | 3759 | 2299 | 3.70 | 3.26 | -4.03E+00 | Neutral | -3.31E+00 | Neutral | -4.73E+00 | Neutral |
| 96 | p.Val96Gln | | | 2870 | 711 | 5.12 | 5.27 | 5857 | 4319 | 5.76 | 6.12 | -9.26E-01 | Neutral | -4.04E+00 | Neutral | -2.82E+00 | Neutral |
| 96 | p.Val96Pro | | | 2688 | 727 | 4.80 | 5.39 | 6159 | 3568 | 6.06 | 5.05 | -1.94E+00 | Neutral | -8.38E-01 | Neutral | -1.23E+00 | Neutral |
| 96 | p.Val96Leu | | | 1684 | 426 | 3.01 | 3.16 | 3163 | 2563 | 3.11 | 3.63 | -3.19E+00 | Neutral | -1.33E+01 | Indeterminate | -1.28E+01 | Indeterminate |
| 96 | p.Val96Asp | | | 2748 | 651 | 4.90 | 4.83 | 3473 | 2913 | 3.41 | 4.13 | -7.34E-01 | Neutral | -1.34E+01 | Indeterminate | -1.07E+01 | Indeterminate |
| 96 | p.Val96Glu | | | 3956 | 740 | 7.06 | 5.49 | 6847 | 4820 | 6.73 | 6.83 | -1.29E-02 | Neutral | -2.41E+00 | Neutral | -1.00E+00 | Neutral |
| 96 | p.Val96Ala | | | 1027 | 328 | 1.83 | 2.43 | 1739 | 1778 | 1.71 | 2.52 | -1.57E+01 | Indeterminate | -4.25E+01 | Indeterminate | -5.32E+01 | Deleterious |
| 96 | p.Val96Gly | | | 2331 | 764 | 4.16 | 5.66 | 4461 | 2549 | 4.39 | 3.61 | -6.46E+00 | Indeterminate | -1.61E+00 | Neutral | -5.35E+00 | Neutral |
| 96 | p.Val96Val | Synonymous | | 4346 | 1259 | 7.76 | 9.33 | 8569 | 5719 | 8.43 | 8.10 | -1.06E+00 | Neutral | -1.06E+00 | Neutral | -8.18E-01 | Neutral |
| 96 | p.Val96Tyr | | | 1651 | 394 | 2.95 | 2.92 | 3353 | 2410 | 3.30 | 3.41 | -2.41E+00 | Neutral | -8.07E+00 | Indeterminate | -7.43E+00 | Indeterminate |
| 96 | p.Val96Cys | | | 3437 | 820 | 6.13 | 6.08 | 5233 | 4773 | 5.15 | 6.76 | -4.16E-01 | Neutral | -1.13E+01 | Indeterminate | -8.48E+00 | Indeterminate |
| 96 | p.Val96Trp | | | 2216 | 450 | 3.96 | 3.34 | 3357 | 3261 | 3.30 | 4.62 | -3.61E-01 | Neutral | -2.16E+01 | Indeterminate | -1.80E+01 | Indeterminate |
| 96 | p.Val96Phe | | | 2574 | 536 | 4.59 | 3.97 | 4599 | 3318 | 4.52 | 4.70 | -2.85E-01 | Neutral | -5.30E+00 | Neutral | -3.30E+00 | Neutral |
| 97 | p.Leu97Asn | | | 2906 | 1928 | 3.97 | 5.53 | | | | | -5.32E+01 | Deleterious | | | -5.32E+01 | Deleterious |
| 97 | p.Leu97Lys | | | 5234 | 3867 | 7.14 | 11.09 | | | | | -5.32E+01 | Deleterious | | | -5.32E+01 | Deleterious |
| 97 | p.Leu97Thr | | | 2517 | 1324 | 3.43 | 3.80 | | | | | -5.32E+01 | Deleterious | | | -5.32E+01 | Deleterious |
| 97 | p.Leu97Arg | | Deleterious | 3861 | 2940 | 5.27 | 8.43 | | | | | -5.32E+01 | Deleterious | | | -5.32E+01 | Deleterious |
| 97 | p.Leu97Ser | | | 5025 | 3430 | 6.86 | 9.84 | | | | | -5.32E+01 | Deleterious | | | -5.32E+01 | Deleterious |
| 97 | p.Leu97Ile | | | 3028 | 229 | 4.13 | 0.66 | | | | | -2.64E-01 | Neutral | | | -2.64E-01 | Neutral |
| 97 | p.Leu97Met | | | 2875 | 178 | 3.92 | 0.51 | | | | | -3.03E-02 | Neutral | | | -3.03E-02 | Neutral |
| 97 | p.Leu97His | | | 2448 | 1356 | 3.34 | 3.89 | | | | | -5.32E+01 | Deleterious | | | -5.32E+01 | Deleterious |
| 97 | p.Leu97Gln | | | 3798 | 1421 | 5.18 | 4.08 | | | | | -5.32E+01 | Deleterious | | | -5.32E+01 | Deleterious |
| 97 | p.Leu97Pro | | | 2652 | 2066 | 3.62 | 5.93 | | | | | -5.32E+01 | Deleterious | | | -5.32E+01 | Deleterious |
| 97 | p.Leu97Leu | Synonymous | | 3046 | 272 | 4.16 | 0.78 | | | | | -1.06E+00 | Neutral | | | -1.06E+00 | Neutral |
| 97 | p.Leu97Asp | | | 4997 | 3900 | 6.82 | 11.19 | | | | | -5.32E+01 | Deleterious | | | -5.32E+01 | Deleterious |
| 97 | p.Leu97Glu | | | 4583 | 2785 | 6.25 | 7.99 | | | | | -5.32E+01 | Deleterious | | | -5.32E+01 | Deleterious |
| 97 | p.Leu97Ala | | | 3439 | 1472 | 4.69 | 4.22 | | | | | -5.32E+01 | Deleterious | | | -5.32E+01 | Deleterious |
| 97 | p.Leu97Gly | | | 4508 | 3527 | 6.15 | 10.12 | | | | | -5.32E+01 | Deleterious | | | -5.32E+01 | Deleterious |
| 97 | p.Leu97Val | | | 3310 | 497 | 4.52 | 1.43 | | | | | -1.19E+01 | Indeterminate | | | -1.19E+01 | Indeterminate |
| 97 | p.Leu97Tyr | | | 3469 | 313 | 4.73 | 0.90 | | | | | -8.32E-01 | Neutral | | | -8.32E-01 | Neutral |
| 97 | p.Leu97Cys | | | 3690 | 339 | 5.04 | 0.97 | | | | | -8.16E-01 | Neutral | | | -8.16E-01 | Neutral |
| 97 | p.Leu97Trp | | | 4586 | 2750 | 6.26 | 7.89 | | | | | -5.32E+01 | Deleterious | | | -5.32E+01 | Deleterious |
| 97 | p.Leu97Phe | | | 3313 | 273 | 4.52 | 0.78 | | | | | -4.46E-01 | Neutral | | | -4.46E-01 | Neutral |

| | | | | | | | | | | | |
|---|---|---|---|---|---|---|---|---|---|---|---|
| 98 | p.His98Asn | | | 6962 | 3204 | 5.23 | 4.40 | -6.69E-01 | Neutral | -6.69E-01 | Neutral |
| 98 | p.His98Lys | | | 3696 | 1535 | 2.77 | 2.11 | -1.47E+00 | Neutral | -1.47E+00 | Neutral |
| 98 | p.His98Thr | | | 4272 | 2069 | 3.21 | 2.84 | -2.77E+00 | Neutral | -2.77E+00 | Neutral |
| 98 | p.His98Arg | | | 7635 | 3456 | 5.73 | 4.75 | -4.48E-01 | Neutral | -4.48E-01 | Neutral |
| 98 | p.His98Ser | | | 6500 | 3794 | 4.88 | 5.21 | -3.55E+00 | Neutral | -3.55E+00 | Neutral |
| 98 | p.His98Ile | | | 6867 | 3037 | 5.15 | 4.17 | -4.98E-01 | Neutral | -4.98E-01 | Neutral |
| 98 | p.His98Met | | | 5455 | 2395 | 4.09 | 3.29 | -8.72E-01 | Neutral | -8.72E-01 | Neutral |
| 98 | p.His98His | Synonymous | | 5527 | 2503 | 4.15 | 3.44 | -1.06E+00 | Neutral | -1.06E+00 | Neutral |
| 98 | p.His98Gln | | | 4315 | 2331 | 3.24 | 3.20 | -4.71E+00 | Neutral | -4.71E+00 | Neutral |
| 98 | p.His98Pro | | Deleterious | 4682 | 11420 | 3.51 | 15.68 | -5.32E+01 | Deleterious | -5.32E+01 | Deleterious |
| 98 | p.His98Leu | | | 5788 | 3026 | 4.34 | 4.16 | -2.42E+00 | Neutral | -2.42E+00 | Neutral |
| 98 | p.His98Asp | | | 5699 | 3256 | 4.28 | 4.47 | -3.97E+00 | Neutral | -3.97E+00 | Neutral |
| 98 | p.His98Glu | | | 8665 | 4094 | 6.50 | 5.62 | -4.51E-01 | Neutral | -4.51E-01 | Neutral |
| 98 | p.His98Ala | | | 8015 | 3389 | 6.02 | 4.65 | -1.99E-01 | Neutral | -1.99E-01 | Neutral |
| 98 | p.His98Gly | | | 5917 | 3101 | 4.44 | 4.26 | -2.35E+00 | Neutral | -2.35E+00 | Neutral |
| 98 | p.His98Val | | | 12966 | 6294 | 9.73 | 8.64 | -1.58E-01 | Neutral | -1.58E-01 | Neutral |
| 98 | p.His98Tyr | | | 8668 | 4200 | 6.51 | 5.77 | -5.61E-01 | Neutral | -5.61E-01 | Neutral |
| 98 | p.His98Cys | | | 7543 | 3723 | 5.66 | 5.11 | -9.36E-01 | Neutral | -9.36E-01 | Neutral |
| 98 | p.His98Trp | | | 7222 | 3059 | 5.42 | 4.20 | -2.87E-01 | Neutral | -2.87E-01 | Neutral |
| 98 | p.His98Phe | | | 6849 | 2927 | 5.14 | 4.02 | -3.69E-01 | Neutral | -3.69E-01 | Neutral |
| 99 | p.Arg99Asn | | | 2603 | 447 | 5.58 | 2.32 | -8.76E+00 | Indeterminate | -8.76E+00 | Indeterminate |
| 99 | p.Arg99Lys | | | 2018 | 299 | 4.32 | 1.55 | -6.84E+00 | Indeterminate | -6.84E+00 | Indeterminate |
| 99 | p.Arg99Thr | | | 1591 | 288 | 3.41 | 1.49 | -1.80E+01 | Indeterminate | -1.80E+01 | Indeterminate |
| 99 | p.Arg99Arg | Synonymous | | 3293 | 407 | 7.05 | 2.11 | -1.06E+00 | Neutral | -1.06E+00 | Neutral |
| 99 | p.Arg99Ser | | | 2572 | 463 | 5.51 | 2.40 | -1.06E+01 | Indeterminate | -1.06E+01 | Indeterminate |
| 99 | p.Arg99Ile | | | 2237 | 347 | 4.79 | 1.80 | -7.17E+00 | Indeterminate | -7.17E+00 | Indeterminate |
| 99 | p.Arg99Met | | | 2430 | 480 | 5.20 | 2.49 | -1.53E+01 | Indeterminate | -1.53E+01 | Indeterminate |
| 99 | p.Arg99His | | | 2305 | 267 | 4.94 | 1.38 | -1.53E+00 | Neutral | -1.53E+00 | Neutral |
| 99 | p.Arg99Gln | | | 1726 | 197 | 3.70 | 1.02 | -2.51E+00 | Neutral | -2.51E+00 | Neutral |
| 99 | p.Arg99Pro | Likely pathogenic | | 2397 | 11387 | 5.13 | 59.05 | -5.32E+01 | Deleterious | -5.32E+01 | Deleterious |
| 99 | p.Arg99Leu | | | 1736 | 369 | 3.72 | 1.91 | -2.60E+01 | Indeterminate | -2.60E+01 | Indeterminate |
| 99 | p.Arg99Asp | | | 1995 | 474 | 4.27 | 2.46 | -3.08E+01 | Indeterminate | -3.05E+01 | Indeterminate |
| 99 | p.Arg99Glu | | | 2272 | 392 | 4.87 | 2.03 | -1.05E+01 | Indeterminate | -1.05E+01 | Indeterminate |
| 99 | p.Arg99Ala | | | 2818 | 806 | 6.04 | 4.18 | -3.66E+01 | Indeterminate | -3.31E+01 | Indeterminate |
| 99 | p.Arg99Gly | | Neutral | 3322 | 420 | 7.12 | 2.18 | -1.22E+00 | Neutral | -1.22E+00 | Neutral |
| 99 | p.Arg99Val | | | 1847 | 518 | 3.96 | 2.69 | -4.81E+01 | Indeterminate | -3.32E+01 | Indeterminate |
| 99 | p.Arg99Tyr | | | 2348 | 308 | 5.03 | 1.60 | -3.07E+00 | Neutral | -3.07E+00 | Neutral |
| 99 | p.Arg99Cys | | | 2310 | 505 | 4.95 | 2.62 | -2.17E+01 | Indeterminate | -2.17E+01 | Indeterminate |
| 99 | p.Arg99Trp | | | 2612 | 591 | 5.59 | 3.06 | -2.13E+01 | Indeterminate | -2.13E+01 | Indeterminate |
| 99 | p.Arg99Phe | | | 2255 | 318 | 4.83 | 1.65 | -4.69E+00 | Neutral | -4.69E+00 | Neutral |
| 100 | p.Ala100Asn | | | 3418 | 332 | 2.33 | 1.28 | -1.08E+00 | Neutral | -1.08E+00 | Neutral |
| 100 | p.Ala100Lys | | | 13759 | 1148 | 9.36 | 4.42 | -3.54E-04 | Neutral | -3.54E-04 | Neutral |
| 100 | p.Ala100Thr | | | 5473 | 648 | 3.72 | 2.49 | -1.44E+00 | Neutral | -1.44E+00 | Neutral |
| 100 | p.Ala100Arg | | | 14131 | 1152 | 9.61 | 4.43 | -1.87E-04 | Neutral | -1.87E-04 | Neutral |
| 100 | p.Ala100Ser | Likely benign | | 3813 | 249 | 2.59 | 0.96 | -8.77E-03 | Neutral | -8.77E-03 | Neutral |
| 100 | p.Ala100Ile | | | 6693 | 620 | 4.55 | 2.38 | -8.43E-02 | Neutral | -8.43E-02 | Neutral |
| 100 | p.Ala100Met | | | 14631 | 1686 | 9.95 | 6.49 | -5.02E-02 | Neutral | -5.02E-02 | Neutral |
| 100 | p.Ala100His | | | 7412 | 516 | 5.04 | 1.98 | -5.68E-04 | Neutral | -5.68E-04 | Neutral |
| 100 | p.Ala100Gln | | | 6694 | 604 | 4.55 | 2.32 | -6.09E-02 | Neutral | -6.09E-02 | Neutral |
| 100 | p.Ala100Pro | | Deleterious | 10531 | 12616 | 7.16 | 48.53 | -5.32E+01 | Deleterious | -5.32E+01 | Deleterious |
| 100 | p.Ala100Leu | | | 6791 | 525 | 4.62 | 2.02 | -6.08E-03 | Neutral | -6.08E-03 | Neutral |
| 100 | p.Ala100Asp | | | 1017 | 66 | 0.69 | 0.25 | -1.07E+00 | Neutral | -1.07E+00 | Neutral |
| 100 | p.Ala100Glu | | | 7528 | 1165 | 5.12 | 4.48 | -3.66E+00 | Neutral | -3.66E+00 | Neutral |
| 100 | p.Ala100Ala | Synonymous | | 2608 | 231 | 1.77 | 0.89 | -1.06E+00 | Neutral | -1.06E+00 | Neutral |
| 100 | p.Ala100Gly | | | 5224 | 1793 | 3.55 | 6.90 | -5.32E+01 | Deleterious | -5.32E+01 | Deleterious |
| 100 | p.Ala100Val | | | 9737 | 705 | 6.62 | 2.71 | -2.07E-04 | Neutral | -2.07E-04 | Neutral |
| 100 | p.Ala100Tyr | | | 3636 | 375 | 2.47 | 1.44 | -1.42E+00 | Neutral | -1.42E+00 | Neutral |
| 100 | p.Ala100Cys | | | 4099 | 233 | 2.79 | 0.90 | -5.06E-04 | Neutral | -5.06E-04 | Neutral |
| 100 | p.Ala100Trp | | | 15058 | 1024 | 10.24 | 3.94 | -2.12E-06 | Neutral | -2.12E-06 | Neutral |
| 100 | p.Ala100Phe | | | 4733 | 309 | 3.22 | 1.19 | -2.66E-03 | Neutral | -2.66E-03 | Neutral |
| 101 | p.Gly101Asn | | | 16195 | 7975 | 4.69 | 2.29 | -6.43E+00 | Indeterminate | -6.43E+00 | Indeterminate |
| 101 | p.Gly101Lys | | | 18728 | 11869 | 5.42 | 3.41 | -1.36E+01 | Indeterminate | -1.36E+01 | Indeterminate |
| 101 | p.Gly101Thr | | | 14607 | 10765 | 4.23 | 3.09 | -2.69E+01 | Indeterminate | -2.68E+01 | Indeterminate |
| 101 | p.Gly101Arg | | Neutral | 17888 | 10294 | 5.18 | 2.95 | -1.03E+01 | Indeterminate | -1.03E+01 | Indeterminate |
| 101 | p.Gly101Ser | | | 20113 | 10885 | 5.82 | 3.12 | -7.05E+00 | Indeterminate | -7.05E+00 | Indeterminate |
| 101 | p.Gly101Ile | | | 17140 | 56282 | 4.96 | 16.15 | -5.32E+01 | Deleterious | -5.32E+01 | Deleterious |
| 101 | p.Gly101Met | | | 24756 | 17567 | 7.16 | 5.04 | -1.45E+01 | Indeterminate | -1.45E+01 | Indeterminate |
| 101 | p.Gly101His | | | 12394 | 6745 | 3.59 | 1.94 | -1.30E+01 | Indeterminate | -1.30E+01 | Indeterminate |
| 101 | p.Gly101Gln | | | 19989 | 10844 | 5.78 | 3.11 | -7.18E+00 | Indeterminate | -7.18E+00 | Indeterminate |
| 101 | p.Gly101Pro | | | 13085 | 38084 | 3.79 | 10.93 | -5.32E+01 | Deleterious | -5.32E+01 | Deleterious |
| 101 | p.Gly101Leu | | | 17138 | 20603 | 4.96 | 5.91 | -5.32E+01 | Deleterious | -5.32E+01 | Deleterious |
| 101 | p.Gly101Asp | | | 16905 | 7563 | 4.89 | 2.17 | -3.90E+00 | Neutral | -3.90E+00 | Neutral |
| 101 | p.Gly101Glu | | | 16990 | 10748 | 4.92 | 3.08 | -1.50E+01 | Indeterminate | -1.50E+01 | Indeterminate |
| 101 | p.Gly101Ala | | | 14073 | 7741 | 4.07 | 2.22 | -1.17E+01 | Indeterminate | -1.17E+01 | Indeterminate |
| 101 | p.Gly101Gly | Synonymous | | 14598 | 4998 | 4.22 | 1.43 | -1.06E+00 | Neutral | -1.06E+00 | Neutral |
| 101 | p.Gly101Val | | | 14907 | 44442 | 4.31 | 12.75 | -5.32E+01 | Deleterious | -5.32E+01 | Deleterious |
| 101 | p.Gly101Tyr | | | 15482 | 13387 | 4.48 | 3.84 | -3.80E+01 | Indeterminate | -3.32E+01 | Indeterminate |
| 101 | p.Gly101Cys | | | 26741 | 15085 | 7.74 | 4.33 | -5.64E+00 | Neutral | -5.64E+00 | Neutral |
| 101 | p.Gly101Trp | Pathogenic | | 21842 | 33554 | 6.32 | 9.63 | -5.32E+01 | Deleterious | -5.32E+01 | Deleterious |
| 101 | p.Gly101Phe | | | 12038 | 9008 | 3.48 | 2.59 | -3.28E+01 | Indeterminate | -3.20E+01 | Indeterminate |
| 102 | p.Ala102Asn | | | 1416 | 161 | 0.51 | 0.23 | -5.32E+01 | Deleterious | -5.32E+01 | Deleterious |
| 102 | p.Ala102Lys | | | 25422 | 7632 | 9.21 | 10.79 | -5.32E+01 | Deleterious | -5.32E+01 | Deleterious |
| 102 | p.Ala102Thr | | | 90 | 11 | 0.03 | 0.02 | -5.32E+01 | Deleterious | -5.32E+01 | Deleterious |
| 102 | p.Ala102Arg | | | 16482 | 8234 | 5.97 | 11.65 | -5.32E+01 | Deleterious | -5.32E+01 | Deleterious |
| 102 | p.Ala102Ser | | | 6001 | 154 | 2.17 | 0.22 | -5.32E+01 | Deleterious | -5.32E+01 | Deleterious |
| 102 | p.Ala102Ile | | | 14717 | 2928 | 5.33 | 4.14 | -5.32E+01 | Deleterious | -5.32E+01 | Deleterious |
| 102 | p.Ala102Met | | | 12041 | 3575 | 4.36 | 5.06 | -5.32E+01 | Deleterious | -5.32E+01 | Deleterious |
| 102 | p.Ala102His | | | 10705 | 2037 | 3.88 | 2.88 | -5.32E+01 | Deleterious | -5.32E+01 | Deleterious |
| 102 | p.Ala102Gln | | | 20898 | 5855 | 7.57 | 8.28 | -5.32E+01 | Deleterious | -5.32E+01 | Deleterious |
| 102 | p.Ala102Pro | | | 1015 | 335 | 0.37 | 0.47 | -5.32E+01 | Deleterious | -5.32E+01 | Deleterious |
| 102 | p.Ala102Leu | | | 19464 | 8187 | 7.05 | 11.58 | -5.32E+01 | Deleterious | -5.32E+01 | Deleterious |
| 102 | p.Ala102Asp | | | 15694 | 4695 | 5.69 | 6.64 | -5.32E+01 | Deleterious | -5.32E+01 | Deleterious |
| 102 | p.Ala102Glu | | | 22596 | 8973 | 8.19 | 12.69 | -5.32E+01 | Deleterious | -5.32E+01 | Deleterious |
| 102 | p.Ala102Ala | Synonymous | | 17069 | 184 | 6.18 | 0.26 | -1.06E+00 | Neutral | -1.06E+00 | Neutral |
| 102 | p.Ala102Gly | | | 8592 | 52 | 3.11 | 0.07 | -2.94E-02 | Neutral | -2.94E-02 | Neutral |
| 102 | p.Ala102Val | | | 31839 | 787 | 11.54 | 1.11 | -1.80E+01 | Indeterminate | -1.80E+01 | Indeterminate |
| 102 | p.Ala102Tyr | | | 7303 | 3519 | 2.65 | 4.98 | -5.32E+01 | Deleterious | -5.32E+01 | Deleterious |
| 102 | p.Ala102Cys | | | 9054 | 131 | 3.28 | 0.19 | -1.24E+01 | Indeterminate | -1.24E+01 | Indeterminate |
| 102 | p.Ala102Trp | | | 27880 | 10063 | 10.10 | 14.23 | -5.32E+01 | Deleterious | -5.32E+01 | Deleterious |
| 102 | p.Ala102Phe | | | 7718 | 3189 | 2.80 | 4.51 | -5.32E+01 | Deleterious | -5.32E+01 | Deleterious |
| 103 | p.Arg103Asn | | | 6943 | 7127 | 8.93 | 8.95 | -1.36E-02 | Neutral | -1.36E-02 | Neutral |
| 103 | p.Arg103Lys | | | 3330 | 3626 | 4.28 | 4.55 | -4.82E-01 | Neutral | -4.82E-01 | Neutral |
| 103 | p.Arg103Thr | | | 3053 | 4907 | 3.93 | 6.16 | -6.16E+00 | Indeterminate | -6.16E+00 | Indeterminate |
| 103 | p.Arg103Arg | Synonymous | | 2349 | 2516 | 3.02 | 3.16 | -1.06E+00 | Neutral | -1.06E+00 | Neutral |
| 103 | p.Arg103Ser | | | 2005 | 1352 | 2.58 | 1.70 | -1.21E-02 | Neutral | -1.21E-02 | Neutral |
| 103 | p.Arg103Ile | | | 3142 | 4421 | 4.04 | 5.55 | -3.14E+00 | Neutral | -3.14E+00 | Neutral |
| 103 | p.Arg103Met | | | 1311 | 1440 | 1.69 | 1.81 | -3.90E+00 | Neutral | -3.90E+00 | Neutral |
| 103 | p.Arg103His | | | 1356 | 1198 | 1.74 | 1.50 | -9.86E-01 | Neutral | -9.86E-01 | Neutral |
| 103 | p.Arg103Gln | | | 2359 | 2500 | 3.03 | 3.14 | -9.72E-01 | Neutral | -9.72E-01 | Neutral |
| 103 | p.Arg103Pro | | | 1548 | 1348 | 1.99 | 1.69 | -6.26E-01 | Neutral | -6.26E-01 | Neutral |
| 103 | p.Arg103Leu | | | 2506 | 1742 | 3.22 | 2.19 | -6.08E-03 | Neutral | -6.08E-03 | Neutral |
| 103 | p.Arg103Asp | | | 5113 | 4011 | 6.58 | 5.03 | -7.98E-04 | Neutral | -7.98E-04 | Neutral |
| 103 | p.Arg103Glu | | | 753 | 642 | 0.97 | 0.81 | -2.72E+00 | Neutral | -2.72E+00 | Neutral |
| 103 | p.Arg103Ala | | | 6092 | 5272 | 7.83 | 6.62 | -1.66E-03 | Neutral | -1.66E-03 | Neutral |
| 103 | p.Arg103Gly | | | 7131 | 7369 | 9.17 | 9.25 | -1.32E-02 | Neutral | -1.32E-02 | Neutral |
| 103 | p.Arg103Val | | | 2730 | 2488 | 3.51 | 3.12 | -1.62E-01 | Neutral | -1.62E-01 | Neutral |
| 103 | p.Arg103Tyr | | | 3731 | 4354 | 4.80 | 5.47 | -6.30E-01 | Neutral | -6.30E-01 | Neutral |
| 103 | p.Arg103Cys | | | 10170 | 11475 | 13.08 | 14.40 | -8.28E-03 | Neutral | -8.28E-03 | Neutral |
| 103 | p.Arg103Trp | | | 2854 | 2738 | 3.67 | 3.44 | -2.37E-01 | Neutral | -2.37E-01 | Neutral |
| 103 | p.Arg103Phe | | | 9286 | 9137 | 11.94 | 11.47 | -1.35E-03 | Neutral | -1.35E-03 | Neutral |
| 104 | p.Leu104Asn | | | 5163 | 1602 | 5.50 | 3.07 | -6.56E-09 | Neutral | -6.41E-09 | Neutral |
| 104 | p.Leu104Lys | | | 4859 | 6969 | 5.17 | 13.35 | -1.14E+01 | Indeterminate | -1.14E+01 | Indeterminate |
| 104 | p.Leu104Thr | | | 2937 | 952 | 3.13 | 1.82 | -2.40E-06 | Neutral | -2.40E-06 | Neutral |
| 104 | p.Leu104Arg | | | 5550 | 15703 | 5.91 | 30.09 | -5.32E+01 | Deleterious | -5.32E+01 | Deleterious |
| 104 | p.Leu104Ser | | | 2630 | 681 | 2.80 | 1.31 | -2.09E-08 | Neutral | -2.08E-08 | Neutral |
| 104 | p.Leu104Ile | | | 4061 | 634 | 4.32 | 1.21 | 0.00E+00 | Neutral | 0.00E+00 | Neutral |
| 104 | p.Leu104Met | | | 6322 | 1029 | 6.73 | 1.97 | 0.00E+00 | Neutral | 0.00E+00 | Neutral |
| 104 | p.Leu104His | | | 3101 | 650 | 3.30 | 1.25 | -1.12E-11 | Neutral | 0.00E+00 | Neutral |
| 104 | p.Leu104Gln | | | 4063 | 2097 | 4.32 | 4.02 | -3.69E-03 | Neutral | -3.69E-03 | Neutral |
| 104 | p.Leu104Pro | | | 1101 | 388 | 1.17 | 0.74 | -8.64E-03 | Neutral | -8.64E-03 | Neutral |
| 104 | p.Leu104Leu | Synonymous | | 2345 | 1636 | 2.50 | 3.14 | -1.06E+00 | Neutral | -1.06E+00 | Neutral |
| 104 | p.Leu104Asp | | | 5290 | 1684 | 5.63 | 3.23 | -1.05E-08 | Neutral | -1.04E-08 | Neutral |
| 104 | p.Leu104Glu | | | 6303 | 3154 | 6.71 | 6.04 | -1.33E-04 | Neutral | -1.33E-04 | Neutral |
| 104 | p.Leu104Ala | | | 3493 | 486 | 3.72 | 0.93 | 0.00E+00 | Neutral | 0.00E+00 | Neutral |
| 104 | p.Leu104Gly | | | 3859 | 11170 | 4.11 | 21.41 | -5.32E+01 | Deleterious | -5.32E+01 | Deleterious |
| 104 | p.Leu104Val | | | 10072 | 959 | 10.72 | 1.84 | 0.00E+00 | Neutral | 0.00E+00 | Neutral |
| 104 | p.Leu104Tyr | | | 5704 | 811 | 6.07 | 1.55 | 0.00E+00 | Neutral | 0.00E+00 | Neutral |
| 104 | p.Leu104Cys | | | 3887 | 660 | 4.14 | 1.26 | -1.44E-15 | Neutral | 0.00E+00 | Neutral |
| 104 | p.Leu104Trp | | | 7298 | 511 | 7.77 | 0.98 | 0.00E+00 | Neutral | 0.00E+00 | Neutral |

| | | | | | | | | | | | | | | | | | |
|---|---|---|---|---|---|---|---|---|---|---|---|---|---|---|---|---|---|
| 104 | p.Leu104Phe | | | 5907 | 407 | 6.29 | 0.78 | | | | | 0.00E+00 | Neutral | | | 0.00E+00 | Neutral |
| 105 | p.Asp105Asn | | | 2615 | 618 | 3.97 | 2.03 | | | | | -2.86E+00 | Neutral | | | -2.86E+00 | Neutral |
| 105 | p.Asp105Lys | | | 4759 | 2093 | 7.22 | 6.86 | | | | | -1.59E+01 | Indeterminate | | | -1.59E+01 | Indeterminate |
| 105 | p.Asp105Thr | | | 2587 | 857 | 3.92 | 2.81 | | | | | -1.22E+01 | Indeterminate | | | -1.22E+01 | Indeterminate |
| 105 | p.Asp105Arg | | | 4066 | 2281 | 6.17 | 7.48 | | | | | -3.54E+01 | Indeterminate | | | -3.29E+01 | Indeterminate |
| 105 | p.Asp105Ser | | | 985 | 469 | 1.49 | 1.54 | | | | | -5.32E+01 | Deleterious | | | -5.32E+01 | Deleterious |
| 105 | p.Asp105Ile | | | 2485 | 1504 | 3.77 | 4.93 | | | | | -5.32E+01 | Deleterious | | | -5.32E+01 | Deleterious |
| 105 | p.Asp105Met | | | 2222 | 756 | 3.37 | 2.48 | | | | | -1.57E+01 | Indeterminate | | | -1.57E+01 | Indeterminate |
| 105 | p.Asp105His | | | 3979 | 1502 | 6.04 | 4.92 | | | | | -1.17E+01 | Indeterminate | | | -1.17E+01 | Indeterminate |
| 105 | p.Asp105Gln | | | 3702 | 1586 | 5.62 | 5.20 | | | | | -1.89E+01 | Indeterminate | | | -1.89E+01 | Indeterminate |
| 105 | p.Asp105Pro | | | 2531 | 5497 | 3.84 | 18.01 | | | | | -5.32E+01 | Deleterious | | | -5.32E+01 | Deleterious |
| 105 | p.Asp105Leu | | | 3705 | 1568 | 5.62 | 5.14 | | | | | -1.82E+01 | Indeterminate | | | -1.82E+01 | Indeterminate |
| 105 | p.Asp105Asp | Synonymous | | 3838 | 879 | 5.82 | 2.88 | | | | | -1.06E+00 | Neutral | | | -1.06E+00 | Neutral |
| 105 | p.Asp105Glu | | | 5734 | 2764 | 8.70 | 9.06 | | | | | -1.74E+01 | Indeterminate | | | -1.74E+01 | Indeterminate |
| 105 | p.Asp105Ala | | | 4005 | 1276 | 6.08 | 4.18 | | | | | -6.08E+00 | Indeterminate | | | -6.08E+00 | Indeterminate |
| 105 | p.Asp105Gly | | | 1562 | 963 | 2.37 | 3.16 | | | | | -5.32E+01 | Deleterious | | | -5.32E+01 | Deleterious |
| 105 | p.Asp105Val | | | 4690 | 2554 | 7.11 | 8.37 | | | | | -2.92E+01 | Indeterminate | | | -2.92E+01 | Indeterminate |
| 105 | p.Asp105Tyr | | | 3757 | 461 | 5.70 | 1.51 | | | | | -3.22E-04 | Neutral | | | -3.22E-04 | Neutral |
| 105 | p.Asp105Cys | | | 1256 | 166 | 1.91 | 0.54 | | | | | -2.28E-01 | Neutral | | | -2.28E-01 | Neutral |
| 105 | p.Asp105Trp | | | 3585 | 1128 | 5.44 | 3.70 | | | | | -6.73E+00 | Indeterminate | | | -6.73E+00 | Indeterminate |
| 105 | p.Asp105Phe | | | 3861 | 1593 | 5.86 | 5.22 | | | | | -1.61E+01 | Indeterminate | | | -1.61E+01 | Indeterminate |
| 106 | p.Val106Asn | | | 1125 | 3424 | 4.79 | 5.65 | 5445 | 1988 | 4.36 | 4.81 | -7.07E+00 | Indeterminate | -4.25E+00 | Neutral | -8.17E+00 | Indeterminate |
| 106 | p.Val106Lys | | | 1023 | 4025 | 4.36 | 6.64 | 7514 | 2575 | 6.02 | 6.23 | -1.90E+01 | Indeterminate | -1.66E+00 | Neutral | -1.67E+01 | Indeterminate |
| 106 | p.Val106Thr | | | 979 | 2138 | 4.17 | 3.53 | 5546 | 1736 | 4.44 | 4.20 | -1.66E+00 | Neutral | -1.77E+00 | Neutral | -1.68E+00 | Neutral |
| 106 | p.Val106Arg | | | 641 | 2444 | 2.73 | 4.03 | 3991 | 1125 | 3.20 | 2.72 | -2.67E+01 | Indeterminate | -1.82E+00 | Neutral | -2.41E+01 | Indeterminate |
| 106 | p.Val106Ser | | | 1510 | 3703 | 6.44 | 6.11 | 7597 | 2262 | 6.08 | 5.47 | -1.39E+00 | Neutral | -5.78E-01 | Neutral | -7.25E-01 | Neutral |
| 106 | p.Val106Ile | | | 1265 | 1935 | 5.39 | 3.19 | 6656 | 2352 | 5.33 | 5.69 | -2.05E-02 | Neutral | -2.54E+00 | Neutral | -1.09E+00 | Neutral |
| 106 | p.Val106Met | | | 1360 | 3085 | 5.80 | 5.09 | 7476 | 2399 | 5.99 | 5.80 | -1.03E+00 | Neutral | -1.08E+00 | Neutral | -8.05E-01 | Neutral |
| 106 | p.Val106His | | | 1415 | 3084 | 6.03 | 5.09 | 7083 | 2284 | 5.67 | 5.52 | -6.83E-01 | Neutral | -1.26E+00 | Neutral | -7.14E-01 | Neutral |
| 106 | p.Val106Gln | | | 1752 | 2941 | 7.47 | 4.85 | 6941 | 2628 | 5.56 | 6.35 | -1.64E-02 | Neutral | -3.41E+00 | Neutral | -1.67E+00 | Neutral |
| 106 | p.Val106Pro | | | 827 | 3185 | 3.52 | 5.26 | 4716 | 1641 | 3.78 | 3.97 | -2.18E+01 | Indeterminate | -4.24E+00 | Neutral | -2.18E+01 | Indeterminate |
| 106 | p.Val106Leu | | | 633 | 1732 | 2.70 | 2.86 | 4193 | 1337 | 3.36 | 3.23 | -9.78E+00 | Indeterminate | -3.33E+00 | Neutral | -9.78E+00 | Indeterminate |
| 106 | p.Val106Asp | | | 1686 | 3754 | 7.19 | 6.19 | 7863 | 2900 | 6.30 | 7.01 | -5.03E-01 | Neutral | -2.36E+00 | Neutral | -1.29E+00 | Neutral |
| 106 | p.Val106Glu | | | 517 | 1854 | 2.20 | 3.06 | 2857 | 932 | 2.29 | 2.25 | -2.74E+01 | Indeterminate | -6.67E+00 | Indeterminate | -2.94E+01 | Indeterminate |
| 106 | p.Val106Ala | | | 1537 | 5106 | 6.55 | 8.43 | 7541 | 2613 | 6.04 | 6.32 | -6.62E+00 | Indeterminate | -1.77E+00 | Neutral | -5.62E+00 | Neutral |
| 106 | p.Val106Gly | | | 1091 | 3828 | 4.65 | 6.32 | 7234 | 2084 | 5.79 | 5.04 | -1.24E+01 | Indeterminate | -5.01E-01 | Neutral | -9.59E+00 | Indeterminate |
| 106 | p.Val106Val | Synonymous | | 1338 | 3033 | 5.70 | 5.00 | 6396 | 1945 | 5.12 | 4.70 | -1.06E+00 | Neutral | -1.06E+00 | Neutral | -8.18E-01 | Neutral |
| 106 | p.Val106Tyr | | | 1292 | 2558 | 5.51 | 4.22 | 5942 | 2069 | 4.76 | 5.00 | -3.88E-01 | Neutral | -2.88E+00 | Neutral | -1.56E+00 | Neutral |
| 106 | p.Val106Cys | | | 1632 | 4660 | 6.96 | 7.69 | 9313 | 2868 | 7.46 | 6.93 | -2.96E+00 | Neutral | -4.31E-01 | Neutral | -1.64E+00 | Neutral |
| 106 | p.Val106Trp | | | 338 | 818 | 1.44 | 1.35 | 2610 | 734 | 2.09 | 1.77 | -1.32E+01 | Indeterminate | -3.91E+00 | Neutral | -1.34E+01 | Indeterminate |
| 106 | p.Val106Phe | | | 1504 | 3293 | 6.41 | 5.43 | 7973 | 2886 | 6.38 | 6.98 | -6.01E-01 | Neutral | -2.06E+00 | Neutral | -1.15E+00 | Neutral |
| 107 | p.Arg107Asn | | | 4635 | 525 | 4.04 | 4.09 | 1667 | 4330 | 4.10 | 4.93 | -1.50E+00 | Neutral | -3.32E+00 | Neutral | -2.70E+00 | Neutral |
| 107 | p.Arg107Lys | | | 4356 | 469 | 3.80 | 3.65 | 1642 | 3493 | 4.04 | 3.98 | -1.23E+00 | Neutral | -1.00E+00 | Neutral | -8.82E-01 | Neutral |
| 107 | p.Arg107Thr | | | 7105 | 785 | 6.19 | 6.11 | 2387 | 4172 | 5.87 | 4.75 | -4.16E-01 | Neutral | -3.90E-02 | Neutral | -5.96E-02 | Neutral |
| 107 | p.Arg107Arg | Synonymous | | 6674 | 811 | 5.82 | 6.31 | 2006 | 4597 | 4.94 | 5.24 | -1.06E+00 | Neutral | -1.06E+00 | Neutral | -8.18E-01 | Neutral |
| 107 | p.Arg107Ser | | | 5046 | 684 | 4.40 | 5.32 | 1875 | 4722 | 4.61 | 5.38 | -3.59E+00 | Neutral | -2.26E+00 | Neutral | -3.51E+00 | Neutral |
| 107 | p.Arg107Ile | | | 3528 | 390 | 3.08 | 3.04 | 1126 | 1863 | 2.77 | 2.12 | -2.26E+00 | Neutral | -3.53E-01 | Neutral | -1.12E+00 | Neutral |
| 107 | p.Arg107Met | | | 7376 | 804 | 6.43 | 6.26 | 2724 | 5759 | 6.70 | 6.56 | -3.28E-01 | Neutral | -2.20E-01 | Neutral | -8.34E-02 | Neutral |
| 107 | p.Arg107His | | | 4098 | 438 | 3.57 | 3.41 | 1418 | 3207 | 3.49 | 3.65 | -1.34E+00 | Neutral | -2.07E+00 | Neutral | -1.66E+00 | Neutral |
| 107 | p.Arg107Gln | | | 10107 | 1081 | 8.81 | 8.41 | 4069 | 7524 | 10.01 | 8.57 | -8.71E-02 | Neutral | -6.53E-03 | Neutral | -2.91E-03 | Neutral |
| 107 | p.Arg107Pro | | | 6819 | 747 | 5.94 | 5.81 | 2662 | 5187 | 6.55 | 5.91 | -4.36E-01 | Neutral | -9.73E-02 | Neutral | -7.95E-02 | Neutral |
| 107 | p.Arg107Leu | | | 6193 | 712 | 5.40 | 5.54 | 2041 | 5611 | 5.02 | 6.39 | -8.46E-01 | Neutral | -3.14E+00 | Neutral | -2.08E+00 | Neutral |
| 107 | p.Arg107Asp | | | 3847 | 431 | 3.35 | 3.35 | 1546 | 3341 | 3.80 | 3.81 | -2.07E+00 | Neutral | -1.29E+00 | Neutral | -1.62E+00 | Neutral |
| 107 | p.Arg107Glu | | | 3892 | 481 | 3.39 | 3.74 | 1098 | 2698 | 2.70 | 3.07 | -3.48E+00 | Neutral | -4.97E+00 | Neutral | -5.67E+00 | Neutral |
| 107 | p.Arg107Ala | | | 8763 | 943 | 7.64 | 7.34 | 3149 | 6188 | 7.75 | 7.05 | -1.61E-01 | Neutral | -5.43E-02 | Neutral | -1.46E-02 | Neutral |
| 107 | p.Arg107Gly | | | 5027 | 563 | 4.38 | 4.38 | 1622 | 3862 | 3.99 | 4.40 | -1.16E+00 | Neutral | -2.16E+00 | Neutral | -1.60E+00 | Neutral |
| 107 | p.Arg107Val | | | 7473 | 764 | 6.51 | 5.95 | 2752 | 6282 | 6.77 | 7.16 | -1.64E-01 | Neutral | -4.44E-01 | Neutral | -1.00E-01 | Neutral |
| 107 | p.Arg107Tyr | | | 5383 | 660 | 4.69 | 5.14 | 1758 | 3648 | 4.33 | 4.16 | -1.82E+00 | Neutral | -6.97E-01 | Neutral | -1.06E+00 | Neutral |
| 107 | p.Arg107Cys | | | 2751 | 264 | 2.40 | 2.05 | 1066 | 3094 | 2.62 | 3.52 | -1.58E+00 | Neutral | -1.01E+01 | Indeterminate | -8.51E+00 | Indeterminate |
| 107 | p.Arg107Trp | | | 6378 | 683 | 5.56 | 5.32 | 2287 | 4275 | 5.63 | 4.87 | -4.34E-01 | Neutral | -1.08E-01 | Neutral | -8.19E-02 | Neutral |
| 107 | p.Arg107Phe | | | 5273 | 613 | 4.60 | 4.77 | 1737 | 3940 | 4.27 | 4.49 | -1.35E+00 | Neutral | -1.38E+00 | Neutral | -1.19E+00 | Neutral |
| 108 | p.Asp108Asn | | Neutral | 2885 | 264 | 5.26 | 1.09 | | | | | -7.77E-01 | Neutral | | | -7.77E-01 | Neutral |
| 108 | p.Asp108Lys | | | 2742 | 1549 | 5.00 | 6.37 | | | | | -5.32E+01 | Deleterious | | | -5.32E+01 | Deleterious |
| 108 | p.Asp108Thr | | | 3646 | 860 | 6.65 | 3.54 | | | | | -3.14E+01 | Indeterminate | | | -3.10E+01 | Indeterminate |
| 108 | p.Asp108Arg | | | 3490 | 2166 | 6.36 | 8.91 | | | | | -5.32E+01 | Deleterious | | | -5.32E+01 | Deleterious |
| 108 | p.Asp108Ser | | | 4074 | 1584 | 7.43 | 6.52 | | | | | -5.32E+01 | Deleterious | | | -5.32E+01 | Deleterious |
| 108 | p.Asp108Ile | | | 3055 | 1607 | 5.57 | 6.61 | | | | | -5.32E+01 | Deleterious | | | -5.32E+01 | Deleterious |
| 108 | p.Asp108Met | | | 3341 | 1731 | 6.09 | 7.12 | | | | | -5.32E+01 | Deleterious | | | -5.32E+01 | Deleterious |
| 108 | p.Asp108His | | Deleterious | 1882 | 1059 | 3.43 | 4.36 | | | | | -5.32E+01 | Deleterious | | | -5.32E+01 | Deleterious |
| 108 | p.Asp108Gln | | | 2051 | 1059 | 3.74 | 4.36 | | | | | -5.32E+01 | Deleterious | | | -5.32E+01 | Deleterious |
| 108 | p.Asp108Pro | | | 2292 | 1346 | 4.18 | 5.54 | | | | | -5.32E+01 | Deleterious | | | -5.32E+01 | Deleterious |
| 108 | p.Asp108Leu | | | 1845 | 1018 | 3.36 | 4.19 | | | | | -5.32E+01 | Deleterious | | | -5.32E+01 | Deleterious |
| 108 | p.Asp108Asp | Synonymous | | 2828 | 268 | 5.16 | 1.10 | | | | | -1.06E+00 | Neutral | | | -1.06E+00 | Neutral |
| 108 | p.Asp108Glu | | | 1895 | 256 | 3.46 | 1.05 | | | | | -1.17E+01 | Indeterminate | | | -1.17E+01 | Indeterminate |
| 108 | p.Asp108Ala | | | 1228 | 713 | 2.24 | 2.93 | | | | | -5.32E+01 | Deleterious | | | -5.32E+01 | Deleterious |
| 108 | p.Asp108Gly | | | 2077 | 1202 | 3.79 | 4.95 | | | | | -5.32E+01 | Deleterious | | | -5.32E+01 | Deleterious |
| 108 | p.Asp108Val | | | 2618 | 1408 | 4.77 | 5.79 | | | | | -5.32E+01 | Deleterious | | | -5.32E+01 | Deleterious |
| 108 | p.Asp108Tyr | | Deleterious | 3107 | 1750 | 5.67 | 7.20 | | | | | -5.32E+01 | Deleterious | | | -5.32E+01 | Deleterious |
| 108 | p.Asp108Cys | | | 3596 | 1073 | 6.56 | 4.42 | | | | | -5.32E+01 | Deleterious | | | -5.32E+01 | Deleterious |
| 108 | p.Asp108Trp | | | 2630 | 1506 | 4.80 | 6.20 | | | | | -5.32E+01 | Deleterious | | | -5.32E+01 | Deleterious |
| 108 | p.Asp108Phe | | | 3562 | 1880 | 6.49 | 7.74 | | | | | -5.32E+01 | Deleterious | | | -5.32E+01 | Deleterious |
| 109 | p.Ala109Asn | | | 5087 | 2168 | 5.92 | 3.29 | 2356 | 497 | 5.70 | 3.39 | -5.43E-02 | Neutral | -6.55E-05 | Neutral | -1.00E-03 | Neutral |
| 109 | p.Ala109Lys | | | 4243 | 2332 | 4.94 | 3.54 | 2242 | 926 | 5.42 | 6.32 | -1.15E+00 | Neutral | -1.01E+00 | Neutral | -8.41E-01 | Neutral |
| 109 | p.Ala109Thr | | | 3577 | 1765 | 4.16 | 2.68 | 1732 | 348 | 4.19 | 2.38 | -7.87E-01 | Neutral | -1.98E-04 | Neutral | -1.59E-01 | Neutral |
| 109 | p.Ala109Arg | | | 4277 | 1773 | 4.98 | 2.69 | 2178 | 929 | 5.27 | 6.35 | -7.90E-02 | Neutral | -1.36E+00 | Neutral | -4.40E-01 | Neutral |
| 109 | p.Ala109Ser | | | 5242 | 1956 | 6.10 | 2.97 | 2496 | 754 | 6.03 | 5.15 | -6.86E-03 | Neutral | -2.73E-02 | Neutral | -3.97E-04 | Neutral |
| 109 | p.Ala109Ile | | | 2094 | 593 | 2.44 | 0.90 | 1056 | 155 | 2.55 | 1.06 | -1.08E-02 | Neutral | -7.81E-06 | Neutral | -4.05E-05 | Neutral |
| 109 | p.Ala109Met | | | 4264 | 1398 | 4.96 | 2.12 | 2308 | 561 | 5.58 | 3.83 | -2.48E-03 | Neutral | -1.24E-03 | Neutral | -4.77E-06 | Neutral |
| 109 | p.Ala109His | | | 4833 | 1775 | 5.62 | 2.70 | 2328 | 605 | 5.63 | 4.13 | -8.20E-03 | Neutral | -3.81E-03 | Neutral | -4.97E-05 | Neutral |
| 109 | p.Ala109Gln | | | 3927 | 1804 | 4.57 | 2.74 | 1861 | 497 | 4.50 | 3.39 | -3.26E-01 | Neutral | -1.89E-02 | Neutral | -3.56E-02 | Neutral |
| 109 | p.Ala109Pro | | Deleterious | 4529 | 30146 | 5.27 | 45.77 | 2031 | 3032 | 4.91 | 20.71 | -5.32E+01 | Deleterious | -5.32E+01 | Deleterious | -5.32E+01 | Deleterious |
| 109 | p.Ala109Leu | | | 4138 | 1033 | 4.81 | 1.57 | 2106 | 955 | 5.09 | 6.52 | -1.26E-05 | Neutral | -2.15E+00 | Neutral | -8.31E-01 | Neutral |
| 109 | p.Ala109Asp | | | 5728 | 2678 | 6.66 | 4.07 | 2458 | 789 | 5.94 | 5.39 | -1.04E-01 | Neutral | -6.53E-02 | Neutral | -9.24E-03 | Neutral |
| 109 | p.Ala109Glu | | | 5017 | 3315 | 5.84 | 5.03 | 2333 | 784 | 5.64 | 5.35 | -2.58E+00 | Neutral | -1.38E-01 | Neutral | -1.19E+00 | Neutral |
| 109 | p.Ala109Ala | Synonymous | | 2870 | 1370 | 3.34 | 2.08 | 1272 | 438 | 3.08 | 2.99 | -1.06E+00 | Neutral | -1.06E+00 | Neutral | -8.18E-01 | Neutral |
| 109 | p.Ala109Gly | | | 3264 | 1198 | 3.80 | 1.82 | 1630 | 426 | 3.94 | 2.91 | -5.29E-02 | Neutral | -2.60E-02 | Neutral | -2.08E-03 | Neutral |
| 109 | p.Ala109Val | | | 3862 | 1619 | 4.49 | 2.46 | 1980 | 602 | 4.79 | 4.11 | -1.34E-01 | Neutral | -8.11E-02 | Neutral | -1.46E-02 | Neutral |
| 109 | p.Ala109Tyr | | | 3063 | 1160 | 3.56 | 1.76 | 1340 | 362 | 3.24 | 2.47 | -1.01E-01 | Neutral | -9.30E-02 | Neutral | -1.20E-02 | Neutral |
| 109 | p.Ala109Cys | | | 5733 | 2308 | 6.67 | 3.50 | 2540 | 570 | 6.14 | 3.89 | -1.41E-02 | Neutral | -1.41E-04 | Neutral | -6.98E-05 | Neutral |
| 109 | p.Ala109Trp | | | 5488 | 2705 | 6.38 | 4.11 | 2690 | 766 | 6.50 | 5.23 | -2.18E-01 | Neutral | -7.76E-03 | Neutral | -1.60E-02 | Neutral |
| 109 | p.Ala109Phe | | | 4731 | 2761 | 5.50 | 4.19 | 2428 | 645 | 5.87 | 4.41 | -1.35E+00 | Neutral | -4.35E-03 | Neutral | -4.01E-01 | Neutral |
| 110 | p.Trp110Asn | | | 4679 | 4750 | 5.07 | 5.08 | | | | | -4.11E-01 | Neutral | | | -4.11E-01 | Neutral |
| 110 | p.Trp110Lys | | | 4116 | 4922 | 4.46 | 5.27 | | | | | -1.92E+00 | Neutral | | | -1.92E+00 | Neutral |
| 110 | p.Trp110Thr | | | 4345 | 3844 | 4.71 | 4.11 | | | | | -1.31E-01 | Neutral | | | -1.31E-01 | Neutral |
| 110 | p.Trp110Arg | | | 5145 | 4792 | 5.58 | 5.13 | | | | | -1.24E-01 | Neutral | | | -1.24E-01 | Neutral |
| 110 | p.Trp110Ser | | | 4655 | 5660 | 5.05 | 6.06 | | | | | -1.65E+00 | Neutral | | | -1.65E+00 | Neutral |
| 110 | p.Trp110Ile | | | 6799 | 7011 | 7.37 | 7.50 | | | | | -1.38E-01 | Neutral | | | -1.38E-01 | Neutral |
| 110 | p.Trp110Met | | | 3650 | 3895 | 3.96 | 4.17 | | | | | -1.18E+00 | Neutral | | | -1.18E+00 | Neutral |
| 110 | p.Trp110His | | | 2242 | 2520 | 2.43 | 2.70 | | | | | -4.10E+00 | Neutral | | | -4.10E+00 | Neutral |
| 110 | p.Trp110Gln | | | 4280 | 4927 | 4.64 | 5.27 | | | | | -1.38E+00 | Neutral | | | -1.38E+00 | Neutral |
| 110 | p.Trp110Pro | | | 2564 | 3394 | 2.78 | 3.63 | | | | | -6.95E+00 | Indeterminate | | | -6.95E+00 | Indeterminate |
| 110 | p.Trp110Leu | | | 5206 | 4558 | 5.64 | 4.88 | | | | | -5.62E-02 | Neutral | | | -5.62E-02 | Neutral |
| 110 | p.Trp110Asp | | | 5506 | 4245 | 5.97 | 4.54 | | | | | -6.85E-03 | Neutral | | | -6.85E-03 | Neutral |
| 110 | p.Trp110Glu | | | 4669 | 5741 | 5.06 | 6.14 | | | | | -1.76E+00 | Neutral | | | -1.76E+00 | Neutral |
| 110 | p.Trp110Ala | | | 2347 | 3010 | 2.54 | 3.22 | | | | | -6.88E+00 | Indeterminate | | | -6.88E+00 | Indeterminate |
| 110 | p.Trp110Gly | | | 4650 | 4291 | 5.04 | 4.59 | | | | | -1.63E-01 | Neutral | | | -1.63E-01 | Neutral |
| 110 | p.Trp110Val | | | 8901 | 8507 | 9.65 | 9.10 | | | | | -1.61E-02 | Neutral | | | -1.61E-02 | Neutral |
| 110 | p.Trp110Tyr | | | 5931 | 5740 | 6.43 | 6.14 | | | | | -1.12E-01 | Neutral | | | -1.12E-01 | Neutral |
| 110 | p.Trp110Cys | | | 4011 | 3729 | 4.35 | 3.99 | | | | | -2.91E-01 | Neutral | | | -2.91E-01 | Neutral |
| 110 | p.Trp110Trp | Synonymous | | 3742 | 3966 | 4.06 | 4.24 | | | | | -1.06E+00 | Neutral | | | -1.06E+00 | Neutral |
| 110 | p.Trp110Phe | | | 4803 | 3940 | 5.21 | 4.22 | | | | | -3.39E-02 | Neutral | | | -3.39E-02 | Neutral |
| 111 | p.Gly111Asn | | | 6372 | 1964 | 5.15 | 2.01 | | | | | -1.98E-01 | Neutral | | | -1.98E-01 | Neutral |
| 111 | p.Gly111Lys | | | 6688 | 3205 | 5.41 | 3.28 | | | | | -4.17E+00 | Neutral | | | -4.17E+00 | Neutral |
| 111 | p.Gly111Thr | | | 3788 | 3224 | 3.06 | 3.30 | | | | | -4.65E+01 | Indeterminate | | | -3.32E+01 | Indeterminate |
| 111 | p.Gly111Arg | | | 6851 | 2649 | 5.54 | 2.71 | | | | | -1.14E+00 | Neutral | | | -1.14E+00 | Neutral |
| 111 | p.Gly111Ser | | Neutral | 6085 | 1922 | 4.92 | 1.97 | | | | | -2.95E-01 | Neutral | | | -2.95E-01 | Neutral |
| 111 | p.Gly111Ile | | | 6288 | 9894 | 5.08 | 10.12 | | | | | -5.32E+01 | Deleterious | | | -5.32E+01 | Deleterious |
| 111 | p.Gly111Met | | | 5548 | 1895 | 4.49 | 1.94 | | | | | -7.64E-01 | Neutral | | | -7.64E-01 | Neutral |
| 111 | p.Gly111His | | | 7031 | 3824 | 5.68 | 3.91 | | | | | -6.80E+00 | Indeterminate | | | -6.80E+00 | Indeterminate |
| 111 | p.Gly111Gln | | | 6371 | 1696 | 5.15 | 1.73 | | | | | -3.39E-02 | Neutral | | | -3.39E-02 | Neutral |
| 111 | p.Gly111Pro | | | 3949 | 35782 | 3.19 | 36.59 | | | | | -5.32E+01 | Deleterious | | | -5.32E+01 | Deleterious |
| 111 | p.Gly111Leu | | | 4980 | 1649 | 4.03 | 1.69 | | | | | -7.94E-01 | Neutral | | | -7.94E-01 | Neutral |
| 111 | p.Gly111Asp | | | 5458 | 1653 | 4.41 | 1.69 | | | | | -2.79E-01 | Neutral | | | -2.79E-01 | Neutral |
| 111 | p.Gly111Glu | | | 8722 | 3773 | 7.05 | 3.86 | | | | | -1.42E+00 | Neutral | | | -1.42E+00 | Neutral |
| 111 | p.Gly111Ala | | | 4049 | 1655 | 3.27 | 1.69 | | | | | -4.33E+00 | Neutral | | | -4.33E+00 | Neutral |
| 111 | p.Gly111Gly | Synonymous | | 5356 | 1889 | 4.33 | 1.93 | | | | | -1.06E+00 | Neutral | | | -1.06E+00 | Neutral |
| 111 | p.Gly111Val | | | 8116 | 10825 | 6.56 | 11.07 | | | | | -5.32E+01 | Deleterious | | | -5.32E+01 | Deleterious |
| 111 | p.Gly111Tyr | | | 5450 | 1959 | 4.41 | 2.00 | | | | | -1.17E+00 | Neutral | | | -1.17E+00 | Neutral |
| 111 | p.Gly111Cys | | | 8487 | 3356 | 6.86 | 3.43 | | | | | -7.96E-01 | Neutral | | | -7.96E-01 | Neutral |

| | | | | | | | | | | | | | | | | | |
|---|---|---|---|---|---|---|---|---|---|---|---|---|---|---|---|---|---|
| 111 | p.Gly111Trp | | | 7849 | 2880 | 6.35 | 2.95 | | | | | -5.35E-01 | Neutral | | | -5.35E-01 | Neutral |
| 111 | p.Gly111Phe | | | 6259 | 2098 | 5.06 | 2.15 | | | | | -4.69E-01 | Neutral | | | -4.69E-01 | Neutral |
| 112 | p.Arg112Asn | | | 1991 | 616 | 5.55 | 1.56 | | | | | -2.08E-01 | Neutral | | | -2.08E-01 | Neutral |
| 112 | p.Arg112Lys | | | 1846 | 746 | 5.14 | 1.89 | | | | | -2.10E+00 | Neutral | | | -2.10E+00 | Neutral |
| 112 | p.Arg112Thr | | | 1776 | 536 | 4.95 | 1.36 | | | | | -2.38E-01 | Neutral | | | -2.38E-01 | Neutral |
| 112 | p.Arg112Arg | Synonymous | | 1452 | 488 | 4.05 | 1.24 | | | | | -1.06E+00 | Neutral | | | -1.06E+00 | Neutral |
| 112 | p.Arg112Ser | | | 1978 | 644 | 5.51 | 1.63 | | | | | -3.51E-01 | Neutral | | | -3.51E-01 | Neutral |
| 112 | p.Arg112Ile | | | 2053 | 798 | 5.72 | 2.02 | | | | | -1.31E+00 | Neutral | | | -1.31E+00 | Neutral |
| 112 | p.Arg112Met | | | 1638 | 361 | 4.56 | 0.91 | | | | | -5.32E-03 | Neutral | | | -5.32E-03 | Neutral |
| 112 | p.Arg112His | | | 2167 | 850 | 6.04 | 2.15 | | | | | -1.24E+00 | Neutral | | | -1.24E+00 | Neutral |
| 112 | p.Arg112Gln | | | 2010 | 873 | 5.60 | 2.21 | | | | | -2.70E+00 | Neutral | | | -2.70E+00 | Neutral |
| 112 | p.Arg112Pro | | | 1820 | 24747 | 5.07 | 62.68 | | | | | -5.32E+01 | Deleterious | | | -5.32E+01 | Deleterious |
| 112 | p.Arg112Leu | | | 1796 | 1126 | 5.00 | 2.85 | | | | | -1.45E+01 | Indeterminate | | | -1.45E+01 | Indeterminate |
| 112 | p.Arg112Asp | | | 1639 | 814 | 4.57 | 2.06 | | | | | -7.02E+00 | Indeterminate | | | -7.02E+00 | Indeterminate |
| 112 | p.Arg112Glu | | | 1519 | 919 | 4.23 | 2.33 | | | | | -1.55E+01 | Indeterminate | | | -1.55E+01 | Indeterminate |
| 112 | p.Arg112Ala | | | 1691 | 765 | 4.71 | 1.94 | | | | | -4.46E+00 | Neutral | | | -4.46E+00 | Neutral |
| 112 | p.Arg112Gly | Pathogenic | | 1652 | 1294 | 4.60 | 3.28 | | | | | -2.95E+01 | Indeterminate | | | -2.94E+01 | Indeterminate |
| 112 | p.Arg112Val | | | 1610 | 761 | 4.49 | 1.93 | | | | | -5.85E+00 | Indeterminate | | | -5.85E+00 | Indeterminate |
| 112 | p.Arg112Tyr | | | 2411 | 944 | 6.72 | 2.39 | | | | | -9.47E-01 | Neutral | | | -9.47E-01 | Neutral |
| 112 | p.Arg112Cys | | | 1023 | 549 | 2.85 | 1.39 | | | | | -1.60E+01 | Indeterminate | | | -1.60E+01 | Indeterminate |
| 112 | p.Arg112Trp | | | 1893 | 729 | 5.27 | 1.85 | | | | | -1.48E+00 | Neutral | | | -1.48E+00 | Neutral |
| 112 | p.Arg112Phe | | | 1931 | 923 | 5.38 | 2.34 | | | | | -4.69E+00 | Neutral | | | -4.69E+00 | Neutral |
| 113 | p.Leu113Asn | | | 5379 | 2633 | 3.76 | 3.88 | 4836 | 1048 | 3.79 | 4.54 | -5.05E+00 | Neutral | -6.01E+00 | Neutral | -7.95E+00 | Indeterminate |
| 113 | p.Leu113Lys | | | 10008 | 4687 | 7.00 | 6.90 | 9167 | 1337 | 7.19 | 5.79 | -1.25E+00 | Neutral | -6.04E-02 | Neutral | -3.79E-01 | Neutral |
| 113 | p.Leu113Thr | | | 7489 | 4108 | 5.24 | 6.05 | 7289 | 1294 | 5.72 | 5.61 | -5.15E+00 | Neutral | -9.00E-01 | Neutral | -3.67E+00 | Neutral |
| 113 | p.Leu113Arg | | | 5302 | 2864 | 3.71 | 4.22 | 5345 | 800 | 4.19 | 3.47 | -7.81E+00 | Indeterminate | -5.23E-01 | Neutral | -5.57E+00 | Neutral |
| 113 | p.Leu113Ser | | | 9675 | 4603 | 6.77 | 6.78 | 9003 | 1272 | 7.06 | 5.51 | -1.50E+00 | Neutral | -4.32E-02 | Neutral | -4.94E-01 | Neutral |
| 113 | p.Leu113Ile | | | 7432 | 3761 | 5.20 | 5.54 | 6762 | 832 | 5.31 | 3.61 | -3.54E+00 | Neutral | -2.39E-02 | Neutral | -1.77E+00 | Neutral |
| 113 | p.Leu113Met | | | 7071 | 2518 | 4.95 | 3.71 | 5559 | 1368 | 4.36 | 5.93 | -3.49E-01 | Neutral | -8.34E+00 | Indeterminate | -5.88E+00 | Indeterminate |
| 113 | p.Leu113His | | | 5034 | 2915 | 3.52 | 4.29 | 4779 | 1071 | 3.75 | 4.64 | -1.08E+01 | Indeterminate | -7.03E+00 | Indeterminate | -1.41E+01 | Indeterminate |
| 113 | p.Leu113Gln | | | 6683 | 3104 | 4.67 | 4.57 | 6294 | 360 | 4.94 | 1.56 | -2.72E+00 | Neutral | -2.58E-09 | Neutral | -1.19E+00 | Neutral |
| 113 | p.Leu113Pro | | | 2367 | 1619 | 1.66 | 2.38 | 2507 | 642 | 1.97 | 2.78 | -3.57E+01 | Indeterminate | -2.24E+01 | Indeterminate | -5.32E+01 | Deleterious |
| 113 | p.Leu113Leu | Synonymous | | 8768 | 3839 | 6.13 | 5.65 | 7778 | 1443 | 6.10 | 6.25 | -1.06E+00 | Neutral | -1.06E+00 | Neutral | -8.18E-01 | Neutral |
| 113 | p.Leu113Asp | | | 8948 | 4600 | 6.26 | 6.78 | 8096 | 1453 | 6.35 | 6.30 | -2.78E+00 | Neutral | -7.48E-01 | Neutral | -1.75E+00 | Neutral |
| 113 | p.Leu113Glu | | | 9912 | 4090 | 6.93 | 6.02 | 7870 | 1492 | 6.18 | 6.46 | -4.80E-01 | Neutral | -1.20E+00 | Neutral | -5.68E-01 | Neutral |
| 113 | p.Leu113Ala | | | 4998 | 2415 | 3.50 | 3.56 | 4479 | 1363 | 3.51 | 5.91 | -5.32E+00 | Neutral | -2.11E+01 | Indeterminate | -2.22E+01 | Indeterminate |
| 113 | p.Leu113Gly | | | 8660 | 4353 | 6.06 | 6.41 | 7215 | 1802 | 5.66 | 7.81 | -2.61E+00 | Neutral | -6.25E+00 | Neutral | -6.02E+00 | Indeterminate |
| 113 | p.Leu113Val | | | 10162 | 4313 | 7.11 | 6.35 | 9204 | 1432 | 7.22 | 6.20 | -5.69E-01 | Neutral | -1.29E-01 | Neutral | -1.29E-01 | Neutral |
| 113 | p.Leu113Tyr | | | 4520 | 2034 | 3.16 | 3.00 | 3746 | 470 | 2.94 | 2.04 | -4.48E+00 | Neutral | -3.07E-01 | Neutral | -2.67E+00 | Neutral |
| 113 | p.Leu113Cys | | | 7692 | 3300 | 5.38 | 4.86 | 6668 | 1154 | 5.23 | 5.00 | -1.25E+00 | Neutral | -9.28E-01 | Neutral | -8.50E-01 | Neutral |
| 113 | p.Leu113Trp | | | 8114 | 3571 | 5.67 | 5.26 | 6388 | 1407 | 5.01 | 6.10 | -1.32E+00 | Neutral | -4.26E+00 | Neutral | -3.29E+00 | Neutral |
| 113 | p.Leu113Phe | | | 4770 | 2565 | 3.34 | 3.78 | 4461 | 1039 | 3.50 | 4.50 | -8.78E+00 | Indeterminate | -8.93E+00 | Indeterminate | -1.40E+01 | Indeterminate |
| 114 | p.Pro114Asn | | | 1817 | 3514 | 6.12 | 6.50 | | | | | -5.32E+01 | Deleterious | | | -5.32E+01 | Deleterious |
| 114 | p.Pro114Lys | | | 995 | 2390 | 3.35 | 4.42 | | | | | -5.32E+01 | Deleterious | | | -5.32E+01 | Deleterious |
| 114 | p.Pro114Thr | Likely pathogenic | | 936 | 1298 | 3.15 | 2.40 | | | | | -5.32E+01 | Deleterious | | | -5.32E+01 | Deleterious |
| 114 | p.Pro114Arg | | | 1780 | 3499 | 6.00 | 6.47 | | | | | -5.32E+01 | Deleterious | | | -5.32E+01 | Deleterious |
| 114 | p.Pro114Ser | | | 1360 | 987 | 4.58 | 1.83 | | | | | -9.07E+00 | Indeterminate | | | -9.07E+00 | Indeterminate |
| 114 | p.Pro114Ile | | | 1197 | 2618 | 4.03 | 4.84 | | | | | -5.32E+01 | Deleterious | | | -5.32E+01 | Deleterious |
| 114 | p.Pro114Met | | | 1129 | 2500 | 3.81 | 4.62 | | | | | -5.32E+01 | Deleterious | | | -5.32E+01 | Deleterious |
| 114 | p.Pro114His | Likely pathogenic | | 1368 | 3436 | 4.61 | 6.35 | | | | | -5.32E+01 | Deleterious | | | -5.32E+01 | Deleterious |
| 114 | p.Pro114Gln | | | 1806 | 4048 | 6.09 | 7.49 | | | | | -5.32E+01 | Deleterious | | | -5.32E+01 | Deleterious |
| 114 | p.Pro114Pro | Synonymous | | 970 | 415 | 3.27 | 0.77 | | | | | -1.06E+00 | Neutral | | | -1.06E+00 | Neutral |
| 114 | p.Pro114Leu | | Deleterious | 1586 | 3408 | 5.35 | 6.30 | | | | | -5.32E+01 | Deleterious | | | -5.32E+01 | Deleterious |
| 114 | p.Pro114Asp | | | 1846 | 3955 | 6.22 | 7.31 | | | | | -5.32E+01 | Deleterious | | | -5.32E+01 | Deleterious |
| 114 | p.Pro114Glu | | | 1828 | 3725 | 6.16 | 6.89 | | | | | -5.32E+01 | Deleterious | | | -5.32E+01 | Deleterious |
| 114 | p.Pro114Ala | | | 394 | 187 | 1.33 | 0.35 | | | | | -8.63E+00 | Indeterminate | | | -8.63E+00 | Indeterminate |
| 114 | p.Pro114Gly | | | 1358 | 1887 | 4.58 | 3.49 | | | | | -5.32E+01 | Deleterious | | | -5.32E+01 | Deleterious |
| 114 | p.Pro114Val | | | 1936 | 2488 | 6.53 | 4.60 | | | | | -3.48E+01 | Indeterminate | | | -3.28E+01 | Indeterminate |
| 114 | p.Pro114Tyr | | | 1770 | 4188 | 5.97 | 7.74 | | | | | -5.32E+01 | Deleterious | | | -5.32E+01 | Deleterious |
| 114 | p.Pro114Cys | | | 1321 | 903 | 4.45 | 1.67 | | | | | -7.47E+00 | Indeterminate | | | -7.47E+00 | Indeterminate |
| 114 | p.Pro114Trp | | | 2579 | 5476 | 8.69 | 10.13 | | | | | -5.32E+01 | Deleterious | | | -5.32E+01 | Deleterious |
| 114 | p.Pro114Phe | | | 1693 | 3153 | 5.71 | 5.83 | | | | | -5.32E+01 | Deleterious | | | -5.32E+01 | Deleterious |
| 115 | p.Val115Asn | | | 6328 | 3214 | 5.11 | 4.25 | | | | | -2.41E+00 | Neutral | | | -2.41E+00 | Neutral |
| 115 | p.Val115Lys | | | 5786 | 2973 | 4.67 | 3.93 | | | | | -3.03E+00 | Neutral | | | -3.03E+00 | Neutral |
| 115 | p.Val115Thr | | | 6751 | 3265 | 5.45 | 4.31 | | | | | -1.57E+00 | Neutral | | | -1.57E+00 | Neutral |
| 115 | p.Val115Arg | | | 3621 | 2684 | 2.92 | 3.55 | | | | | -2.19E+01 | Indeterminate | | | -2.19E+01 | Indeterminate |
| 115 | p.Val115Ser | | | 5887 | 3318 | 4.75 | 4.38 | | | | | -4.66E+00 | Neutral | | | -4.66E+00 | Neutral |
| 115 | p.Val115Ile | | | 4855 | 2279 | 3.92 | 3.01 | | | | | -2.54E+00 | Neutral | | | -2.54E+00 | Neutral |
| 115 | p.Val115Met | | | 7340 | 3600 | 5.92 | 4.76 | | | | | -1.44E+00 | Neutral | | | -1.44E+00 | Neutral |
| 115 | p.Val115His | | | 4929 | 2890 | 3.98 | 3.82 | | | | | -7.13E+00 | Indeterminate | | | -7.13E+00 | Indeterminate |
| 115 | p.Val115Gln | | | 4345 | 3257 | 3.51 | 4.30 | | | | | -1.91E+01 | Indeterminate | | | -1.91E+01 | Indeterminate |
| 115 | p.Val115Pro | | | 9037 | 11154 | 7.29 | 14.74 | | | | | -3.72E+01 | Indeterminate | | | -3.31E+01 | Indeterminate |
| 115 | p.Val115Leu | | | 9713 | 5878 | 7.84 | 7.77 | | | | | -2.89E+00 | Neutral | | | -2.89E+00 | Neutral |
| 115 | p.Val115Asp | | | 5570 | 3650 | 4.50 | 4.82 | | | | | -9.37E+00 | Indeterminate | | | -9.37E+00 | Indeterminate |
| 115 | p.Val115Glu | | | 4451 | 2694 | 3.59 | 3.56 | | | | | -9.20E+00 | Indeterminate | | | -9.20E+00 | Indeterminate |
| 115 | p.Val115Ala | | | 3262 | 2371 | 2.63 | 3.13 | | | | | -2.29E+01 | Indeterminate | | | -2.29E+01 | Indeterminate |
| 115 | p.Val115Gly | | | 9239 | 5617 | 7.46 | 7.42 | | | | | -3.24E+00 | Neutral | | | -3.24E+00 | Neutral |
| 115 | p.Val115Val | Synonymous | | 8999 | 4521 | 7.26 | 5.97 | | | | | -1.06E+00 | Neutral | | | -1.06E+00 | Neutral |
| 115 | p.Val115Tyr | | | 5619 | 2709 | 4.54 | 3.58 | | | | | -2.24E+00 | Neutral | | | -2.24E+00 | Neutral |
| 115 | p.Val115Cys | | | 4727 | 2228 | 3.82 | 2.94 | | | | | -2.73E+00 | Neutral | | | -2.73E+00 | Neutral |
| 115 | p.Val115Trp | | | 6028 | 3240 | 4.87 | 4.28 | | | | | -3.56E+00 | Neutral | | | -3.56E+00 | Neutral |
| 115 | p.Val115Phe | | | 7402 | 4130 | 5.97 | 5.46 | | | | | -3.04E+00 | Neutral | | | -3.04E+00 | Neutral |
| 116 | p.Asp116Asn | | | 4943 | 1914 | 2.89 | 2.39 | | | | | -3.49E+00 | Neutral | | | -3.49E+00 | Neutral |
| 116 | p.Asp116Lys | | | 12435 | 4890 | 7.27 | 6.11 | | | | | -5.04E-01 | Neutral | | | -5.04E-01 | Neutral |
| 116 | p.Asp116Thr | | | 6247 | 2920 | 3.65 | 3.65 | | | | | -5.85E+00 | Indeterminate | | | -5.85E+00 | Indeterminate |
| 116 | p.Asp116Arg | | | 7079 | 3300 | 4.14 | 4.12 | | | | | -4.81E+00 | Neutral | | | -4.81E+00 | Neutral |
| 116 | p.Asp116Ser | | | 10328 | 4273 | 6.04 | 5.34 | | | | | -1.21E+00 | Neutral | | | -1.21E+00 | Neutral |
| 116 | p.Asp116Ile | | | 8211 | 3857 | 4.80 | 4.82 | | | | | -3.94E+00 | Neutral | | | -3.94E+00 | Neutral |
| 116 | p.Asp116Met | | | 7955 | 2576 | 4.65 | 3.22 | | | | | -3.32E-01 | Neutral | | | -3.32E-01 | Neutral |
| 116 | p.Asp116His | | | 10717 | 4466 | 6.26 | 5.58 | | | | | -1.16E+00 | Neutral | | | -1.16E+00 | Neutral |
| 116 | p.Asp116Gln | | | 8325 | 3445 | 4.87 | 4.30 | | | | | -1.92E+00 | Neutral | | | -1.92E+00 | Neutral |
| 116 | p.Asp116Pro | | | 4065 | 7636 | 2.38 | 9.54 | | | | | -5.32E+01 | Deleterious | | | -5.32E+01 | Deleterious |
| 116 | p.Asp116Leu | | | 7761 | 3355 | 4.54 | 4.19 | | | | | -2.83E+00 | Neutral | | | -2.83E+00 | Neutral |
| 116 | p.Asp116Asp | Synonymous | | 9567 | 3790 | 5.59 | 4.73 | | | | | -1.06E+00 | Neutral | | | -1.06E+00 | Neutral |
| 116 | p.Asp116Glu | | | 10329 | 4515 | 6.04 | 5.64 | | | | | -1.73E+00 | Neutral | | | -1.73E+00 | Neutral |
| 116 | p.Asp116Ala | | | 6842 | 2732 | 4.00 | 3.41 | | | | | -2.29E+00 | Neutral | | | -2.29E+00 | Neutral |
| 116 | p.Asp116Gly | | | 10318 | 6029 | 6.03 | 7.53 | | | | | -7.40E+00 | Indeterminate | | | -7.40E+00 | Indeterminate |
| 116 | p.Asp116Val | | | 8773 | 3898 | 5.13 | 4.87 | | | | | -2.63E+00 | Neutral | | | -2.63E+00 | Neutral |
| 116 | p.Asp116Tyr | | | 7958 | 2869 | 4.65 | 3.58 | | | | | -8.30E-01 | Neutral | | | -8.30E-01 | Neutral |
| 116 | p.Asp116Cys | | | 12620 | 5342 | 7.38 | 6.67 | | | | | -8.78E-01 | Neutral | | | -8.78E-01 | Neutral |
| 116 | p.Asp116Trp | | | 7328 | 3954 | 4.28 | 4.94 | | | | | -8.45E+00 | Indeterminate | | | -8.45E+00 | Indeterminate |
| 116 | p.Asp116Phe | | | 9303 | 4319 | 5.44 | 5.39 | | | | | -3.01E+00 | Neutral | | | -3.01E+00 | Neutral |
| 117 | p.Leu117Asn | | | 7070 | 2516 | 4.56 | 4.40 | | | | | -1.87E+00 | Neutral | | | -1.87E+00 | Neutral |
| 117 | p.Leu117Lys | | | 9877 | 3587 | 6.37 | 6.28 | | | | | -1.02E+00 | Neutral | | | -1.02E+00 | Neutral |
| 117 | p.Leu117Thr | | | 7864 | 2594 | 5.07 | 4.54 | | | | | -8.74E-01 | Neutral | | | -8.74E-01 | Neutral |
| 117 | p.Leu117Arg | | | 8872 | 3342 | 5.72 | 5.85 | | | | | -1.66E+00 | Neutral | | | -1.66E+00 | Neutral |
| 117 | p.Leu117Ser | | | 8065 | 2946 | 5.20 | 5.15 | | | | | -1.67E+00 | Neutral | | | -1.67E+00 | Neutral |
| 117 | p.Leu117Ile | | | 8208 | 2847 | 5.30 | 4.98 | | | | | -1.14E+00 | Neutral | | | -1.14E+00 | Neutral |
| 117 | p.Leu117Met | | | 6340 | 2230 | 4.09 | 3.90 | | | | | -2.16E+00 | Neutral | | | -2.16E+00 | Neutral |
| 117 | p.Leu117His | | | 6546 | 1898 | 4.22 | 3.32 | | | | | -5.04E-01 | Neutral | | | -5.04E-01 | Neutral |
| 117 | p.Leu117Gln | | | 7223 | 2954 | 4.66 | 5.17 | | | | | -3.86E+00 | Neutral | | | -3.86E+00 | Neutral |
| 117 | p.Leu117Pro | | | 5349 | 4175 | 3.45 | 7.30 | | | | | -4.32E+01 | Indeterminate | | | -3.32E+01 | Indeterminate |
| 117 | p.Leu117Leu | Synonymous | | 8344 | 2882 | 5.38 | 5.04 | | | | | -1.06E+00 | Neutral | | | -1.06E+00 | Neutral |
| 117 | p.Leu117Asp | | | 8914 | 3001 | 5.75 | 5.25 | | | | | -7.42E-01 | Neutral | | | -7.42E-01 | Neutral |
| 117 | p.Leu117Glu | | | 9669 | 3670 | 6.24 | 6.42 | | | | | -1.45E+00 | Neutral | | | -1.45E+00 | Neutral |
| 117 | p.Leu117Ala | | | 4021 | 1315 | 2.59 | 2.30 | | | | | -3.34E+00 | Neutral | | | -3.34E+00 | Neutral |
| 117 | p.Leu117Gly | | | 8262 | 3050 | 5.33 | 5.34 | | | | | -1.70E+00 | Neutral | | | -1.70E+00 | Neutral |
| 117 | p.Leu117Val | | | 9084 | 3239 | 5.86 | 5.67 | | | | | -1.09E+00 | Neutral | | | -1.09E+00 | Neutral |
| 117 | p.Leu117Tyr | | | 9820 | 3189 | 6.34 | 5.58 | | | | | -4.17E-01 | Neutral | | | -4.17E-01 | Neutral |
| 117 | p.Leu117Cys | | | 8025 | 3045 | 5.18 | 5.33 | | | | | -2.13E+00 | Neutral | | | -2.13E+00 | Neutral |
| 117 | p.Leu117Trp | | | 5931 | 1964 | 3.83 | 3.44 | | | | | -1.72E+00 | Neutral | | | -1.72E+00 | Neutral |
| 117 | p.Leu117Phe | | | 7491 | 2716 | 4.83 | 4.75 | | | | | -1.86E+00 | Neutral | | | -1.86E+00 | Neutral |
| 118 | p.Ala118Asn | | | 5316 | 3171 | 4.50 | 5.33 | | | | | -3.28E+01 | Indeterminate | | | -3.20E+01 | Indeterminate |
| 118 | p.Ala118Lys | | | 5355 | 3669 | 4.53 | 6.16 | | | | | -4.51E+01 | Indeterminate | | | -3.32E+01 | Indeterminate |
| 118 | p.Ala118Thr | | | 7459 | 2789 | 6.31 | 4.69 | | | | | -4.89E+00 | Neutral | | | -4.89E+00 | Neutral |
| 118 | p.Ala118Arg | | | 8551 | 5560 | 7.24 | 9.34 | | | | | -2.74E+01 | Indeterminate | | | -2.74E+01 | Indeterminate |
| 118 | p.Ala118Ser | | | 4903 | 1694 | 4.15 | 2.85 | | | | | -6.37E+00 | Indeterminate | | | -6.37E+00 | Indeterminate |
| 118 | p.Ala118Ile | | | 7141 | 3330 | 6.05 | 5.59 | | | | | -1.23E+01 | Indeterminate | | | -1.23E+01 | Indeterminate |
| 118 | p.Ala118Met | | | 4319 | 2202 | 3.66 | 3.70 | | | | | -2.63E+01 | Indeterminate | | | -2.63E+01 | Indeterminate |
| 118 | p.Ala118His | | | 4487 | 2865 | 3.80 | 4.81 | | | | | -4.39E+01 | Indeterminate | | | -3.32E+01 | Indeterminate |
| 118 | p.Ala118Gln | | | 5718 | 3355 | 4.84 | 5.64 | | | | | -2.97E+01 | Indeterminate | | | -2.96E+01 | Indeterminate |
| 118 | p.Ala118Pro | | | 2401 | 1145 | 2.03 | 1.92 | | | | | -3.62E+01 | Indeterminate | | | -3.30E+01 | Indeterminate |
| 118 | p.Ala118Leu | | | 6468 | 2765 | 5.48 | 4.64 | | | | | -1.02E+01 | Indeterminate | | | -1.02E+01 | Indeterminate |
| 118 | p.Ala118Asp | | | 7364 | 4617 | 6.23 | 7.76 | | | | | -2.84E+01 | Indeterminate | | | -2.83E+01 | Indeterminate |
| 118 | p.Ala118Glu | | | 6132 | 3782 | 5.19 | 6.35 | | | | | -3.17E+01 | Indeterminate | | | -3.13E+01 | Indeterminate |
| 118 | p.Ala118Ala | Synonymous | | 7922 | 2315 | 6.71 | 3.89 | | | | | -1.06E+00 | Neutral | | | -1.06E+00 | Neutral |
| 118 | p.Ala118Gly | | | 4629 | 1330 | 3.92 | 2.23 | | | | | -2.90E+00 | Neutral | | | -2.90E+00 | Neutral |
| 118 | p.Ala118Val | | | 6165 | 2323 | 5.22 | 3.90 | | | | | -6.66E+00 | Indeterminate | | | -6.66E+00 | Indeterminate |
| 118 | p.Ala118Tyr | | | 4780 | 2825 | 4.05 | 4.75 | | | | | -3.50E+01 | Indeterminate | | | -3.28E+01 | Indeterminate |

| | | | | | | | | | | | | | | | | | |
|---|---|---|---|---|---|---|---|---|---|---|---|---|---|---|---|---|---|
| 118 | p.Ala118Cys | | | 7840 | 2535 | 6.64 | 4.26 | | | | | -2.10E+00 | Neutral | | | -2.10E+00 | Neutral |
| 118 | p.Ala118Trp | | | 5510 | 3866 | 4.66 | 6.49 | | | | | -4.65E+01 | Indeterminate | | | -3.32E+01 | Indeterminate |
| 118 | p.Ala118Phe | | | 5657 | 3391 | 4.79 | 5.70 | | | | | -3.16E+01 | Indeterminate | | | -3.12E+01 | Indeterminate |
| 119 | p.Glu119Asn | | | 4365 | 494 | 4.22 | 3.26 | | | | | -1.46E+00 | Neutral | | | -1.46E+00 | Neutral |
| 119 | p.Glu119Lys | | | 4474 | 419 | 4.33 | 2.77 | | | | | -3.00E-01 | Neutral | | | -3.00E-01 | Neutral |
| 119 | p.Glu119Thr | | | 6227 | 566 | 6.02 | 3.74 | | | | | -6.40E-02 | Neutral | | | -6.40E-02 | Neutral |
| 119 | p.Glu119Arg | | | 3977 | 649 | 3.85 | 4.29 | | | | | -1.00E+01 | Indeterminate | | | -1.00E+01 | Indeterminate |
| 119 | p.Glu119Ser | | | 5648 | 290 | 5.46 | 1.92 | | | | | -3.20E-06 | Neutral | | | -3.20E-06 | Neutral |
| 119 | p.Glu119Ile | | | 4576 | 367 | 4.42 | 2.42 | | | | | -4.77E-02 | Neutral | | | -4.77E-02 | Neutral |
| 119 | p.Glu119Met | | | 6932 | 1140 | 6.70 | 7.53 | | | | | -4.89E+00 | Neutral | | | -4.89E+00 | Neutral |
| 119 | p.Glu119His | | | 5341 | 1407 | 5.16 | 9.29 | | | | | -3.11E+01 | Indeterminate | | | -3.08E+01 | Indeterminate |
| 119 | p.Glu119Gln | | | 4684 | 1065 | 4.53 | 7.03 | | | | | -2.38E+01 | Indeterminate | | | -2.38E+01 | Indeterminate |
| 119 | p.Glu119Pro | | | 4894 | 1402 | 4.73 | 9.26 | | | | | -4.08E+01 | Indeterminate | | | -3.32E+01 | Indeterminate |
| 119 | p.Glu119Leu | | | 3932 | 414 | 3.80 | 2.73 | | | | | -1.13E+00 | Neutral | | | -1.13E+00 | Neutral |
| 119 | p.Glu119Asp | | | 4939 | 865 | 4.78 | 5.71 | | | | | -9.98E+00 | Indeterminate | | | -9.98E+00 | Indeterminate |
| 119 | p.Glu119Glu | Synonymous | | 7139 | 908 | 6.90 | 6.00 | | | | | -1.06E+00 | Neutral | | | -1.06E+00 | Neutral |
| 119 | p.Glu119Ala | | | 3939 | 268 | 3.81 | 1.77 | | | | | -9.16E-03 | Neutral | | | -9.16E-03 | Neutral |
| 119 | p.Glu119Gly | | | 6152 | 699 | 5.95 | 4.62 | | | | | -6.53E-01 | Neutral | | | -6.53E-01 | Neutral |
| 119 | p.Glu119Val | | | 5805 | 939 | 5.61 | 6.20 | | | | | -5.91E+00 | Indeterminate | | | -5.91E+00 | Indeterminate |
| 119 | p.Glu119Tyr | | | 4931 | 262 | 4.77 | 1.73 | | | | | -1.97E-05 | Neutral | | | -1.97E-05 | Neutral |
| 119 | p.Glu119Cys | | | 5729 | 354 | 5.54 | 2.34 | | | | | -1.70E-04 | Neutral | | | -1.70E-04 | Neutral |
| 119 | p.Glu119Trp | | | 5340 | 1418 | 5.16 | 9.37 | | | | | -3.18E+01 | Indeterminate | | | -3.13E+01 | Indeterminate |
| 119 | p.Glu119Phe | | | 4406 | 1213 | 4.26 | 8.01 | | | | | -4.03E+01 | Indeterminate | | | -3.32E+01 | Indeterminate |
| 120 | p.Glu120Asn | | | 6668 | 1715 | 6.69 | 7.09 | 5879 | 924 | 6.09 | 6.77 | -2.64E+00 | Neutral | -9.94E-01 | Neutral | -1.82E+00 | Neutral |
| 120 | p.Glu120Lys | | | 5140 | 1178 | 5.16 | 4.87 | 4764 | 741 | 4.94 | 5.43 | -2.24E+00 | Neutral | -1.50E+00 | Neutral | -1.90E+00 | Neutral |
| 120 | p.Glu120Thr | | | 4944 | 1177 | 4.96 | 4.87 | 5718 | 665 | 5.93 | 4.87 | -2.98E+00 | Neutral | -5.61E-02 | Neutral | -1.40E+00 | Neutral |
| 120 | p.Glu120Arg | | | 3381 | 806 | 3.39 | 3.33 | 3318 | 513 | 3.44 | 3.76 | -5.53E+00 | Neutral | -2.95E+00 | Neutral | -5.70E+00 | Neutral |
| 120 | p.Glu120Ser | | | 4204 | 949 | 4.22 | 3.93 | 3955 | 492 | 4.10 | 3.60 | -2.99E+00 | Neutral | -4.35E-01 | Neutral | -1.67E+00 | Neutral |
| 120 | p.Glu120Ile | | | 5170 | 1508 | 5.19 | 6.24 | 5138 | 717 | 5.32 | 5.25 | -7.14E+00 | Indeterminate | -5.55E-01 | Neutral | -5.03E+00 | Neutral |
| 120 | p.Glu120Met | | | 3518 | 1081 | 3.53 | 4.47 | 3493 | 547 | 3.62 | 4.01 | -1.37E+01 | Indeterminate | -2.88E+00 | Neutral | -1.29E+01 | Indeterminate |
| 120 | p.Glu120His | | | 4893 | 1157 | 4.91 | 4.79 | 4467 | 725 | 4.63 | 5.31 | -2.93E+00 | Neutral | -2.23E+00 | Neutral | -2.96E+00 | Neutral |
| 120 | p.Glu120Gln | | | 5102 | 1362 | 5.12 | 5.63 | 4390 | 675 | 4.55 | 4.95 | -4.98E+00 | Neutral | -1.66E+00 | Neutral | -4.15E+00 | Neutral |
| 120 | p.Glu120Pro | | | 3878 | 1004 | 3.89 | 4.15 | 4187 | 558 | 4.34 | 4.09 | -6.48E+00 | Indeterminate | -6.65E-01 | Neutral | -4.57E+00 | Neutral |
| 120 | p.Glu120Leu | | | 4277 | 850 | 4.29 | 3.52 | 3887 | 565 | 4.03 | 4.14 | -1.35E+00 | Neutral | -1.50E+00 | Neutral | -1.27E+00 | Neutral |
| 120 | p.Glu120Asp | | | 5594 | 1357 | 5.61 | 5.61 | 5098 | 821 | 5.28 | 6.02 | -2.64E+00 | Neutral | -1.62E+00 | Neutral | -2.28E+00 | Neutral |
| 120 | p.Glu120Glu | Synonymous | | 5994 | 1288 | 6.01 | 5.33 | 5329 | 818 | 5.52 | 5.99 | -1.06E+00 | Neutral | -1.06E+00 | Neutral | -8.18E-01 | Neutral |
| 120 | p.Glu120Ala | | | 4804 | 1061 | 4.82 | 4.39 | 5170 | 515 | 5.36 | 3.77 | -2.06E+00 | Neutral | -1.00E-02 | Neutral | -7.85E-01 | Neutral |
| 120 | p.Glu120Gly | | | 4006 | 926 | 4.02 | 3.83 | 4433 | 514 | 4.59 | 3.77 | -3.67E+00 | Neutral | -1.48E-01 | Neutral | -1.95E+00 | Neutral |
| 120 | p.Glu120Val | | | 5499 | 1316 | 5.52 | 5.44 | 5200 | 764 | 5.39 | 5.60 | -2.53E+00 | Neutral | -8.14E-01 | Neutral | -1.61E+00 | Neutral |
| 120 | p.Glu120Tyr | | | 2339 | 529 | 2.35 | 2.19 | 2119 | 350 | 2.20 | 2.56 | -7.40E+00 | Indeterminate | -7.88E+00 | Indeterminate | -1.18E+01 | Indeterminate |
| 120 | p.Glu120Cys | | | 6757 | 1679 | 6.78 | 6.94 | 6708 | 1017 | 6.95 | 7.45 | -2.12E+00 | Neutral | -5.29E-01 | Neutral | -1.14E+00 | Neutral |
| 120 | p.Glu120Trp | | | 5516 | 1474 | 5.53 | 6.10 | 5475 | 719 | 5.67 | 5.27 | -4.44E+00 | Neutral | -2.62E-01 | Neutral | -2.61E+00 | Neutral |
| 120 | p.Glu120Phe | | | 7976 | 1759 | 8.00 | 7.28 | 7777 | 1009 | 8.06 | 7.39 | -6.25E-01 | Neutral | -6.15E-02 | Neutral | -1.25E-01 | Neutral |
| 121 | p.Leu121Asn | | | 5557 | 2222 | 3.85 | 4.08 | | | | | -4.57E+00 | Neutral | | | -4.57E+00 | Neutral |
| 121 | p.Leu121Lys | | | 5913 | 2024 | 4.10 | 3.72 | | | | | -1.77E+00 | Neutral | | | -1.77E+00 | Neutral |
| 121 | p.Leu121Thr | | | 7287 | 2639 | 5.05 | 4.85 | | | | | -1.63E+00 | Neutral | | | -1.63E+00 | Neutral |
| 121 | p.Leu121Arg | | | 6565 | 3538 | 4.55 | 6.50 | | | | | -1.20E+01 | Indeterminate | | | -1.20E+01 | Indeterminate |
| 121 | p.Leu121Ser | | | 4991 | 1747 | 3.46 | 3.21 | | | | | -2.80E+00 | Neutral | | | -2.80E+00 | Neutral |
| 121 | p.Leu121Ile | | | 6708 | 2326 | 4.65 | 4.27 | | | | | -1.47E+00 | Neutral | | | -1.47E+00 | Neutral |
| 121 | p.Leu121Met | | | 7054 | 9643 | 4.89 | 17.72 | | | | | -5.32E+01 | Deleterious | | | -5.32E+01 | Deleterious |
| 121 | p.Leu121His | | | 6679 | 1883 | 4.63 | 3.46 | | | | | -2.79E-01 | Neutral | | | -2.79E-01 | Neutral |
| 121 | p.Leu121Gln | | | 6870 | 2290 | 4.76 | 4.21 | | | | | -1.06E+00 | Neutral | | | -1.06E+00 | Neutral |
| 121 | p.Leu121Pro | | | 6118 | 1809 | 4.24 | 3.32 | | | | | -5.61E-01 | Neutral | | | -5.61E-01 | Neutral |
| 121 | p.Leu121Leu | Synonymous | | 5345 | 1640 | 3.71 | 3.01 | | | | | -1.06E+00 | Neutral | | | -1.06E+00 | Neutral |
| 121 | p.Leu121Asp | | | 20642 | 4114 | 14.32 | 7.56 | | | | | -9.42E-07 | Neutral | | | -9.42E-07 | Neutral |
| 121 | p.Leu121Glu | | | 8089 | 3520 | 5.61 | 6.47 | | | | | -3.77E+00 | Neutral | | | -3.77E+00 | Neutral |
| 121 | p.Leu121Ala | | | 5113 | 1790 | 3.55 | 3.29 | | | | | -2.68E+00 | Neutral | | | -2.68E+00 | Neutral |
| 121 | p.Leu121Gly | | | 7464 | 3126 | 5.18 | 5.74 | | | | | -3.55E+00 | Neutral | | | -3.55E+00 | Neutral |
| 121 | p.Leu121Val | | | 8431 | 2171 | 5.85 | 3.99 | | | | | -4.01E-02 | Neutral | | | -4.01E-02 | Neutral |
| 121 | p.Leu121Tyr | | | 4665 | 1066 | 3.24 | 1.96 | | | | | -1.05E-01 | Neutral | | | -1.05E-01 | Neutral |
| 121 | p.Leu121Cys | | | 6774 | 1898 | 4.70 | 3.49 | | | | | -2.50E-01 | Neutral | | | -2.50E-01 | Neutral |
| 121 | p.Leu121Trp | | | 7344 | 2981 | 5.09 | 5.48 | | | | | -3.11E+00 | Neutral | | | -3.11E+00 | Neutral |
| 121 | p.Leu121Phe | | | 6583 | 1996 | 4.57 | 3.67 | | | | | -5.65E-01 | Neutral | | | -5.65E-01 | Neutral |
| 122 | p.Gly122Asn | | | 7569 | 3399 | 4.56 | 3.94 | 11922 | 382 | 4.65 | 2.73 | -4.61E-01 | Neutral | -1.76E-05 | Neutral | -6.09E-02 | Neutral |
| 122 | p.Gly122Lys | | | 8834 | 4858 | 5.33 | 5.63 | 14027 | 527 | 5.47 | 3.77 | -1.46E+00 | Neutral | -1.66E-04 | Neutral | -4.53E-01 | Neutral |
| 122 | p.Gly122Thr | | | 9750 | 4942 | 5.88 | 5.73 | 14935 | 744 | 5.83 | 5.32 | -6.31E-01 | Neutral | -1.67E-02 | Neutral | -1.13E-01 | Neutral |
| 122 | p.Gly122Arg | | | 9626 | 6094 | 5.80 | 7.07 | 13461 | 1005 | 5.25 | 7.18 | -2.86E+00 | Neutral | -1.50E+00 | Neutral | -2.36E+00 | Neutral |
| 122 | p.Gly122Ser | | | 10109 | 4589 | 6.09 | 5.32 | 13433 | 666 | 5.24 | 4.76 | -2.03E-01 | Neutral | -2.60E-02 | Neutral | -1.65E-02 | Neutral |
| 122 | p.Gly122Ile | | | 5859 | 3683 | 3.53 | 4.27 | 11299 | 488 | 4.41 | 3.49 | -6.10E+00 | Indeterminate | -7.76E-03 | Neutral | -3.72E+00 | Neutral |
| 122 | p.Gly122Met | | | 8522 | 3997 | 5.14 | 4.63 | 12277 | 772 | 4.79 | 5.52 | -4.78E-01 | Neutral | -5.21E-01 | Neutral | -2.40E-01 | Neutral |
| 122 | p.Gly122His | | | 8831 | 4052 | 5.32 | 4.70 | 13991 | 782 | 5.46 | 5.59 | -3.52E-01 | Neutral | -1.02E-01 | Neutral | -5.93E-02 | Neutral |
| 122 | p.Gly122Gln | | | 9295 | 4626 | 5.60 | 5.36 | 13146 | 527 | 5.13 | 3.77 | -6.19E-01 | Neutral | -8.97E-04 | Neutral | -1.04E-01 | Neutral |
| 122 | p.Gly122Pro | | | 7279 | 3497 | 4.39 | 4.05 | 11943 | 942 | 4.66 | 6.73 | -8.88E-01 | Neutral | -2.66E+00 | Neutral | -1.75E+00 | Neutral |
| 122 | p.Gly122Leu | | | 6446 | 3447 | 3.89 | 4.00 | 10134 | 422 | 3.95 | 3.02 | -2.35E+00 | Neutral | -7.71E-03 | Neutral | -9.59E-01 | Neutral |
| 122 | p.Gly122Asp | | | 8712 | 4580 | 5.25 | 5.31 | 14156 | 564 | 5.52 | 4.03 | -1.11E+00 | Neutral | -5.01E-04 | Neutral | -2.87E-01 | Neutral |
| 122 | p.Gly122Glu | | | 9974 | 5662 | 6.01 | 6.56 | 15722 | 654 | 6.13 | 4.67 | -1.38E+00 | Neutral | -5.87E-04 | Neutral | -4.14E-01 | Neutral |
| 122 | p.Gly122Ala | | | 8715 | 4648 | 5.25 | 5.39 | 14349 | 673 | 5.60 | 4.81 | -1.23E+00 | Neutral | -8.15E-03 | Neutral | -3.43E-01 | Neutral |
| 122 | p.Gly122Gly | Synonymous | | 6971 | 3382 | 4.20 | 3.92 | 9670 | 615 | 3.77 | 4.40 | -1.06E+00 | Neutral | -1.06E+00 | Neutral | -8.18E-01 | Neutral |
| 122 | p.Gly122Val | | Neutral | 7691 | 3268 | 4.64 | 3.79 | 12215 | 711 | 4.77 | 5.08 | -2.61E-01 | Neutral | -2.59E-01 | Neutral | -7.59E-02 | Neutral |
| 122 | p.Gly122Tyr | | | 9213 | 4716 | 5.55 | 5.47 | 14745 | 1063 | 5.75 | 7.60 | -7.93E-01 | Neutral | -9.53E-01 | Neutral | -6.02E-01 | Neutral |
| 122 | p.Gly122Cys | | | 6227 | 4023 | 3.75 | 4.66 | 9379 | 574 | 3.66 | 4.10 | -6.29E+00 | Indeterminate | -8.61E-01 | Neutral | -4.58E+00 | Neutral |
| 122 | p.Gly122Trp | | | 6552 | 2705 | 3.95 | 3.14 | 8809 | 789 | 3.44 | 5.64 | -3.31E-01 | Neutral | -7.69E+00 | Indeterminate | -5.31E+00 | Neutral |
| 122 | p.Gly122Phe | | | 9687 | 6084 | 5.84 | 7.05 | 16655 | 1092 | 6.50 | 7.80 | -2.71E+00 | Neutral | -3.03E-01 | Neutral | -1.39E+00 | Neutral |
| 123 | p.His123Asn | | | 7749 | 1318 | 5.54 | 5.67 | 4136 | 391 | 5.18 | 4.24 | -6.16E+00 | Indeterminate | -2.67E+00 | Neutral | -6.00E+00 | Indeterminate |
| 123 | p.His123Lys | | | 8216 | 869 | 5.87 | 3.74 | 4134 | 343 | 5.17 | 3.72 | -2.36E-01 | Neutral | -1.19E+00 | Neutral | -4.32E-01 | Neutral |
| 123 | p.His123Thr | | | 5633 | 1158 | 4.03 | 4.98 | 3132 | 397 | 3.92 | 4.31 | -1.76E+01 | Indeterminate | -1.36E+01 | Indeterminate | -2.66E+01 | Indeterminate |
| 123 | p.His123Arg | | | 7158 | 1021 | 5.12 | 4.39 | 3689 | 331 | 4.62 | 3.59 | -3.03E+00 | Neutral | -2.46E+00 | Neutral | -3.23E+00 | Neutral |
| 123 | p.His123Ser | | | 6242 | 951 | 4.46 | 4.09 | 3377 | 270 | 4.23 | 2.93 | -5.21E+00 | Neutral | -1.45E+00 | Neutral | -4.18E+00 | Neutral |
| 123 | p.His123Ile | | | 6667 | 1082 | 4.76 | 4.65 | 3468 | 304 | 4.34 | 3.30 | -6.24E+00 | Indeterminate | -2.42E+00 | Neutral | -5.85E+00 | Indeterminate |
| 123 | p.His123Met | | | 7540 | 1467 | 5.39 | 6.31 | 4059 | 452 | 5.08 | 4.91 | -1.07E+01 | Indeterminate | -6.08E+00 | Neutral | -1.31E+01 | Indeterminate |
| 123 | p.His123His | Synonymous | | 8307 | 1051 | 5.94 | 4.52 | 4588 | 388 | 5.74 | 4.21 | -1.06E+00 | Neutral | -1.06E+00 | Neutral | -8.18E-01 | Neutral |
| 123 | p.His123Gln | Likely benign | | 7177 | 959 | 5.13 | 4.12 | 4451 | 353 | 5.57 | 3.83 | -2.09E+00 | Neutral | -7.04E-01 | Neutral | -1.24E+00 | Neutral |
| 123 | p.His123Pro | | | 4158 | 1844 | 2.97 | 7.93 | 2773 | 1007 | 3.47 | 10.93 | -5.32E+01 | Deleterious | -5.32E+01 | Deleterious | -5.32E+01 | Deleterious |
| 123 | p.His123Leu | | | 6904 | 1120 | 4.93 | 4.82 | 4153 | 458 | 5.20 | 4.97 | -5.93E+00 | Indeterminate | -5.64E+00 | Neutral | -8.40E+00 | Indeterminate |
| 123 | p.His123Asp | | | 7108 | 1408 | 5.08 | 6.05 | 4396 | 588 | 5.50 | 6.38 | -1.22E+01 | Indeterminate | -1.11E+01 | Indeterminate | -1.92E+01 | Indeterminate |
| 123 | p.His123Glu | | | 8263 | 1312 | 5.91 | 5.64 | 4672 | 718 | 5.85 | 7.79 | -4.10E+00 | Neutral | -1.64E+01 | Indeterminate | -1.65E+01 | Indeterminate |
| 123 | p.His123Ala | | | 5540 | 763 | 3.96 | 3.28 | 3137 | 305 | 3.93 | 3.31 | -3.91E+00 | Neutral | -4.87E+00 | Neutral | -5.96E+00 | Indeterminate |
| 123 | p.His123Gly | | | 4257 | 992 | 3.04 | 4.27 | 2165 | 292 | 2.71 | 3.17 | -3.18E+01 | Indeterminate | -2.35E+01 | Indeterminate | -5.32E+01 | Deleterious |
| 123 | p.His123Val | | | 7367 | 1424 | 5.26 | 6.12 | 4585 | 769 | 5.74 | 8.35 | -1.08E+01 | Indeterminate | -2.15E+01 | Indeterminate | -2.78E+01 | Indeterminate |
| 123 | p.His123Tyr | | | 6410 | 817 | 4.58 | 3.51 | 3853 | 265 | 4.82 | 2.88 | -1.97E+00 | Neutral | -3.07E-01 | Neutral | -9.09E-01 | Neutral |
| 123 | p.His123Cys | | | 10430 | 1689 | 7.45 | 7.26 | 5920 | 478 | 7.41 | 5.19 | -3.06E+00 | Neutral | -3.63E-01 | Neutral | -1.67E+00 | Neutral |
| 123 | p.His123Trp | | | 9243 | 1217 | 6.61 | 5.23 | 5457 | 491 | 6.83 | 5.33 | -1.10E+00 | Neutral | -1.10E+00 | Neutral | -8.58E-01 | Neutral |
| 123 | p.His123Phe | | | 5563 | 793 | 3.98 | 3.41 | 3757 | 612 | 4.70 | 6.64 | -4.58E+00 | Neutral | -2.39E+01 | Indeterminate | -2.41E+01 | Indeterminate |
| 124 | p.Arg124Asn | | | 13117 | 2444 | 5.17 | 4.54 | 3663 | 474 | 4.87 | 3.58 | -3.29E-01 | Neutral | -2.58E-02 | Neutral | -3.76E-02 | Neutral |
| 124 | p.Arg124Lys | | | 12813 | 2748 | 5.05 | 5.10 | 3694 | 668 | 4.91 | 5.04 | -1.12E+00 | Neutral | -8.71E-01 | Neutral | -7.38E-01 | Neutral |
| 124 | p.Arg124Thr | | | 9192 | 1893 | 3.62 | 3.52 | 2863 | 759 | 3.80 | 5.73 | -1.78E+00 | Neutral | -9.91E+00 | Indeterminate | -8.50E+00 | Indeterminate |
| 124 | p.Arg124Arg | Synonymous | | 17487 | 4129 | 6.90 | 7.67 | 4768 | 964 | 6.34 | 7.27 | -1.06E+00 | Neutral | -1.06E+00 | Neutral | -8.18E-01 | Neutral |
| 124 | p.Arg124Ser | | | 13488 | 2606 | 5.32 | 4.84 | 4071 | 483 | 5.41 | 3.65 | -4.23E-01 | Neutral | -3.91E-03 | Neutral | -5.28E-02 | Neutral |
| 124 | p.Arg124Ile | | | 12169 | 3269 | 4.80 | 6.07 | 3406 | 1712 | 4.53 | 12.92 | -4.52E+00 | Neutral | -4.93E+01 | Indeterminate | -3.30E+01 | Indeterminate |
| 124 | p.Arg124Met | | | 12928 | 2709 | 5.10 | 5.03 | 3850 | 244 | 5.12 | 1.84 | -9.23E-01 | Neutral | -4.15E-09 | Neutral | -2.09E-01 | Neutral |
| 124 | p.Arg124His | | | 11854 | 2494 | 4.67 | 4.63 | 3574 | 303 | 4.75 | 2.29 | -1.17E+00 | Neutral | -1.16E-05 | Neutral | -3.13E-01 | Neutral |
| 124 | p.Arg124Gln | | | 11493 | 2278 | 4.53 | 4.23 | 3369 | 459 | 4.48 | 3.46 | -8.14E-01 | Neutral | -7.35E-02 | Neutral | -1.96E-01 | Neutral |
| 124 | p.Arg124Pro | | | 11653 | 2437 | 4.60 | 4.53 | 4001 | 473 | 5.32 | 3.57 | -1.17E+00 | Neutral | -4.05E-03 | Neutral | -3.14E-01 | Neutral |
| 124 | p.Arg124Leu | | | 12505 | 2677 | 4.93 | 4.97 | 4088 | 679 | 5.43 | 5.12 | -1.17E+00 | Neutral | -3.17E-01 | Neutral | -4.63E-01 | Neutral |
| 124 | p.Arg124Asp | | | 12156 | 2226 | 4.79 | 4.13 | 3222 | 321 | 4.28 | 2.42 | -3.55E-01 | Neutral | -6.88E-04 | Neutral | -3.77E-02 | Neutral |
| 124 | p.Arg124Glu | | | 14476 | 2761 | 5.71 | 5.13 | 4456 | 841 | 5.92 | 6.35 | -3.01E-01 | Neutral | -7.49E-01 | Neutral | -2.61E-01 | Neutral |
| 124 | p.Arg124Ala | | | 11408 | 2306 | 4.50 | 4.28 | 3663 | 419 | 4.87 | 3.16 | -9.62E-01 | Neutral | -3.81E-03 | Neutral | -2.26E-01 | Neutral |
| 124 | p.Arg124Gly | | | 9347 | 2760 | 3.69 | 5.13 | 2629 | 570 | 3.49 | 4.30 | -9.60E+00 | Indeterminate | -4.91E+00 | Neutral | -1.10E+01 | Indeterminate |
| 124 | p.Arg124Val | | | 14183 | 3045 | 5.59 | 5.66 | 4315 | 946 | 5.73 | 7.14 | -8.79E-01 | Neutral | -2.23E+00 | Neutral | -1.45E+00 | Neutral |
| 124 | p.Arg124Tyr | | | 14546 | 2873 | 5.74 | 5.34 | 3803 | 1355 | 5.05 | 10.23 | -4.11E-01 | Neutral | -1.90E+01 | Indeterminate | -1.56E+01 | Indeterminate |
| 124 | p.Arg124Cys | | Neutral | 10657 | 2523 | 4.20 | 4.69 | 3124 | 287 | 4.15 | 2.17 | -2.99E+00 | Neutral | -1.71E-04 | Neutral | -1.37E+00 | Neutral |
| 124 | p.Arg124Trp | | | 14282 | 3094 | 5.63 | 5.75 | 4499 | 564 | 5.98 | 4.26 | -9.24E-01 | Neutral | -5.62E-03 | Neutral | -2.12E-01 | Neutral |
| 124 | p.Arg124Phe | | | 13826 | 2570 | 5.45 | 4.77 | 4199 | 730 | 5.58 | 5.51 | -2.72E-01 | Neutral | -4.44E-01 | Neutral | -1.35E-01 | Neutral |
| 125 | p.Asp125Asn | | | 2848 | 2460 | 5.20 | 6.10 | 3408 | 1349 | 5.14 | 4.32 | -1.78E+00 | Neutral | -1.18E-01 | Neutral | -6.88E-01 | Neutral |
| 125 | p.Asp125Lys | | | 3101 | 2194 | 5.66 | 5.44 | 4037 | 1492 | 6.09 | 4.78 | -3.02E-01 | Neutral | -2.37E-02 | Neutral | -3.20E-02 | Neutral |
| 125 | p.Asp125Thr | | | 3289 | 1858 | 6.00 | 4.61 | 3748 | 1509 | 5.65 | 4.84 | -1.52E-02 | Neutral | -9.95E-02 | Neutral | -4.33E-03 | Neutral |
| 125 | p.Asp125Arg | | | 3171 | 2491 | 5.79 | 6.18 | 3969 | 1800 | 5.99 | 5.77 | -7.10E-01 | Neutral | -2.91E-01 | Neutral | -2.40E-01 | Neutral |
| 125 | p.Asp125Ser | | | 2543 | 1437 | 4.64 | 3.57 | 2832 | 1341 | 4.27 | 4.30 | -5.04E-02 | Neutral | -1.07E+00 | Neutral | -2.91E-01 | Neutral |
| 125 | p.Asp125Ile | | | 2818 | 1560 | 5.14 | 3.87 | 3472 | 1190 | 5.24 | 3.81 | -2.39E-02 | Neutral | -1.64E-02 | Neutral | -5.54E-04 | Neutral |
| 125 | p.Asp125Met | | | 2373 | 1268 | 4.33 | 3.15 | 2705 | 1017 | 4.08 | 3.26 | -3.24E-02 | Neutral | -1.56E-01 | Neutral | -1.14E-02 | Neutral |
| 125 | p.Asp125His | | Neutral | 2521 | 1745 | 4.60 | 4.33 | 3237 | 1195 | 4.88 | 3.83 | -4.65E-01 | Neutral | -6.25E-02 | Neutral | -7.80E-02 | Neutral |
| 125 | p.Asp125Gln | | | 2478 | 1355 | 4.52 | 3.36 | 3307 | 1133 | 4.99 | 3.63 | -3.65E-02 | Neutral | -2.07E-02 | Neutral | -1.10E-03 | Neutral |
| 125 | p.Asp125Pro | | | 1616 | 908 | 2.95 | 2.25 | 1907 | 835 | 2.88 | 2.68 | -2.63E-01 | Neutral | -1.54E+00 | Neutral | -6.31E-01 | Neutral |
| 125 | p.Asp125Leu | | | 2325 | 1463 | 4.24 | 3.63 | 2786 | 1357 | 4.20 | 4.35 | -2.44E-01 | Neutral | -1.35E+00 | Neutral | -5.21E-01 | Neutral |
| 125 | p.Asp125Asp | Synonymous | | 2689 | 2109 | 4.91 | 5.23 | 3254 | 1612 | 4.91 | 5.17 | -1.06E+00 | Neutral | -1.06E+00 | Neutral | -8.18E-01 | Neutral |
| 125 | p.Asp125Glu | | | 2333 | 1284 | 4.26 | 3.19 | 2599 | 1204 | 3.92 | 3.86 | -5.18E-02 | Neutral | -1.12E+00 | Neutral | -3.14E-01 | Neutral |
| 125 | p.Asp125Ala | | | 2163 | 1221 | 3.95 | 3.03 | 2290 | 1343 | 3.46 | 4.31 | -9.68E-02 | Neutral | -5.33E+00 | Neutral | -3.17E+00 | Neutral |
| 125 | p.Asp125Gly | | | 2889 | 2643 | 5.27 | 6.56 | 3529 | 1120 | 5.32 | 3.59 | -2.44E+00 | Neutral | -4.49E-03 | Neutral | -1.02E+00 | Neutral |
| 125 | p.Asp125Val | | | 2562 | 1031 | 4.68 | 2.56 | 2832 | 1017 | 4.27 | 3.26 | -1.62E-04 | Neutral | -7.70E-02 | Neutral | -1.99E-03 | Neutral |

| | | | | | | | | | | | | | | | | | |
|---|---|---|---|---|---|---|---|---|---|---|---|---|---|---|---|---|---|
| 125 | p.Asp125Tyr | | | 3302 | 6200 | 6.03 | 15.38 | 4205 | 5560 | 6.34 | 17.82 | -2.96E+01 | Indeterminate | -3.71E+01 | Indeterminate | -5.32E+01 | Deleterious |
| 125 | p.Asp125Cys | | | 3684 | 3165 | 6.72 | 7.85 | 4837 | 1450 | 7.30 | 4.65 | -9.62E-01 | Neutral | -2.29E-04 | Neutral | -2.25E-01 | Neutral |
| 125 | p.Asp125Trp | | | 3184 | 2210 | 5.81 | 5.48 | 3864 | 1751 | 5.83 | 5.61 | -2.29E-01 | Neutral | -3.15E-01 | Neutral | -8.23E-02 | Neutral |
| 125 | p.Asp125Phe | | | 2896 | 1702 | 5.29 | 4.22 | 3460 | 1919 | 5.22 | 6.15 | -4.79E-02 | Neutral | -1.96E+00 | Neutral | -7.48E-01 | Neutral |
| 126 | p.Val126Asn | | | 13592 | 1396 | 4.72 | 1.65 | | | | | -4.72E-01 | Neutral | | | -4.72E-01 | Neutral |
| 126 | p.Val126Lys | | | 15516 | 11295 | 5.38 | 13.32 | | | | | -5.32E+01 | Deleterious | | | -5.32E+01 | Deleterious |
| 126 | p.Val126Thr | | | 13332 | 577 | 4.63 | 0.68 | | | | | -1.29E-07 | Neutral | | | -1.28E-07 | Neutral |
| 126 | p.Val126Arg | | | 18926 | 19643 | 6.57 | 23.16 | | | | | -5.32E+01 | Deleterious | | | -5.32E+01 | Deleterious |
| 126 | p.Val126Ser | | | 13725 | 918 | 4.76 | 1.08 | | | | | -1.54E-03 | Neutral | | | -1.54E-03 | Neutral |
| 126 | p.Val126Ile | | | 16166 | 1121 | 5.61 | 1.32 | | | | | -1.10E-03 | Neutral | | | -1.10E-03 | Neutral |
| 126 | p.Val126Met | | | 19562 | 1317 | 6.79 | 1.55 | | | | | -1.83E-04 | Neutral | | | -1.83E-04 | Neutral |
| 126 | p.Val126His | | | 14043 | 4631 | 4.87 | 5.46 | | | | | -5.30E+01 | Indeterminate | | | -3.32E+01 | Indeterminate |
| 126 | p.Val126Gln | | | 16828 | 2136 | 5.84 | 2.52 | | | | | -1.39E+00 | Neutral | | | -1.39E+00 | Neutral |
| 126 | p.Val126Pro | | | 14969 | 885 | 5.19 | 1.04 | | | | | -7.83E-05 | Neutral | | | -7.83E-05 | Neutral |
| 126 | p.Val126Leu | | | 12287 | 717 | 4.26 | 0.85 | | | | | -2.30E-04 | Neutral | | | -2.30E-04 | Neutral |
| 126 | p.Val126Asp | Pathogenic | | 10860 | 7618 | 3.77 | 8.98 | | | | | -5.32E+01 | Deleterious | | | -5.32E+01 | Deleterious |
| 126 | p.Val126Glu | | | 15495 | 1254 | 5.38 | 1.48 | | | | | -1.80E-02 | Neutral | | | -1.80E-02 | Neutral |
| 126 | p.Val126Ala | | | 13910 | 1058 | 4.83 | 1.25 | | | | | -1.21E-02 | Neutral | | | -1.21E-02 | Neutral |
| 126 | p.Val126Gly | | | 13329 | 1339 | 4.62 | 1.58 | | | | | -4.10E-01 | Neutral | | | -4.10E-01 | Neutral |
| 126 | p.Val126Val | Synonymous | | 10679 | 1121 | 3.70 | 1.32 | | | | | -1.06E+00 | Neutral | | | -1.06E+00 | Neutral |
| 126 | p.Val126Tyr | | | 14358 | 11033 | 4.98 | 13.01 | | | | | -5.32E+01 | Deleterious | | | -5.32E+01 | Deleterious |
| 126 | p.Val126Cys | | | 10597 | 503 | 3.68 | 0.59 | | | | | -7.77E-06 | Neutral | | | -7.77E-06 | Neutral |
| 126 | p.Val126Trp | | | 16355 | 14867 | 5.67 | 17.53 | | | | | -5.32E+01 | Deleterious | | | -5.32E+01 | Deleterious |
| 126 | p.Val126Phe | | | 13728 | 1397 | 4.76 | 1.65 | | | | | -4.22E-01 | Neutral | | | -4.22E-01 | Neutral |
| 127 | p.Ala127Asn | | | 7524 | 1490 | 4.20 | 2.73 | | | | | -1.01E+00 | Neutral | | | -1.01E+00 | Neutral |
| 127 | p.Ala127Lys | | | 7364 | 2041 | 4.11 | 3.74 | | | | | -6.63E+00 | Indeterminate | | | -6.63E+00 | Indeterminate |
| 127 | p.Ala127Thr | | | 7875 | 3331 | 4.40 | 6.11 | | | | | -2.48E+01 | Indeterminate | | | -2.48E+01 | Indeterminate |
| 127 | p.Ala127Arg | | | 10173 | 4327 | 5.68 | 7.93 | | | | | -1.99E+01 | Indeterminate | | | -1.99E+01 | Indeterminate |
| 127 | p.Ala127Ser | Benign | | 9439 | 2253 | 5.27 | 4.13 | | | | | -2.10E+00 | Neutral | | | -2.10E+00 | Neutral |
| 127 | p.Ala127Ile | | | 8459 | 3367 | 4.72 | 6.17 | | | | | -1.96E+01 | Indeterminate | | | -1.96E+01 | Indeterminate |
| 127 | p.Ala127Met | | | 10187 | 2188 | 5.69 | 4.01 | | | | | -8.74E-01 | Neutral | | | -8.74E-01 | Neutral |
| 127 | p.Ala127His | | | 7847 | 3375 | 4.38 | 6.19 | | | | | -2.60E+01 | Indeterminate | | | -2.60E+01 | Indeterminate |
| 127 | p.Ala127Gln | | | 9425 | 2342 | 5.26 | 4.29 | | | | | -2.66E+00 | Neutral | | | -2.66E+00 | Neutral |
| 127 | p.Ala127Pro | | Deleterious | 5169 | 5329 | 2.89 | 9.77 | | | | | -5.32E+01 | Deleterious | | | -5.32E+01 | Deleterious |
| 127 | p.Ala127Leu | | | 9185 | 2356 | 5.13 | 4.32 | | | | | -3.30E+00 | Neutral | | | -3.30E+00 | Neutral |
| 127 | p.Ala127Asp | | | 10308 | 3893 | 5.76 | 7.14 | | | | | -1.37E+01 | Indeterminate | | | -1.37E+01 | Indeterminate |
| 127 | p.Ala127Glu | | | 11153 | 2432 | 6.23 | 4.46 | | | | | -7.79E-01 | Neutral | | | -7.79E-01 | Neutral |
| 127 | p.Ala127Ala | Synonymous | | 7852 | 1588 | 4.38 | 2.91 | | | | | -1.06E+00 | Neutral | | | -1.06E+00 | Neutral |
| 127 | p.Ala127Gly | | | 8262 | 2466 | 4.61 | 4.52 | | | | | -7.66E+00 | Indeterminate | | | -7.66E+00 | Indeterminate |
| 127 | p.Ala127Val | | | 11122 | 2140 | 6.21 | 3.92 | | | | | -2.61E-01 | Neutral | | | -2.61E-01 | Neutral |
| 127 | p.Ala127Tyr | | | 7626 | 2133 | 4.26 | 3.91 | | | | | -6.56E+00 | Indeterminate | | | -6.56E+00 | Indeterminate |
| 127 | p.Ala127Cys | | | 8750 | 1407 | 4.89 | 2.58 | | | | | -8.73E-02 | Neutral | | | -8.73E-02 | Neutral |
| 127 | p.Ala127Trp | | | 10780 | 3177 | 6.02 | 5.82 | | | | | -5.00E+00 | Neutral | | | -5.00E+00 | Neutral |
| 127 | p.Ala127Phe | | | 10582 | 2912 | 5.91 | 5.34 | | | | | -3.71E+00 | Neutral | | | -3.71E+00 | Neutral |
| 128 | p.Arg128Asn | | | 2284 | 409 | 3.90 | 1.59 | | | | | -5.62E-05 | Neutral | | | -5.62E-05 | Neutral |
| 128 | p.Arg128Lys | | | 3039 | 1401 | 5.19 | 5.46 | | | | | -2.71E+00 | Neutral | | | -2.71E+00 | Neutral |
| 128 | p.Arg128Thr | | | 3022 | 1036 | 5.16 | 4.04 | | | | | -3.17E-01 | Neutral | | | -3.17E-01 | Neutral |
| 128 | p.Arg128Arg | Synonymous | | 5105 | 2409 | 8.72 | 9.39 | | | | | -1.06E+00 | Neutral | | | -1.06E+00 | Neutral |
| 128 | p.Arg128Ser | | | 2145 | 1444 | 3.66 | 5.63 | | | | | -1.91E+01 | Indeterminate | | | -1.91E+01 | Indeterminate |
| 128 | p.Arg128Ile | | | 3252 | 1959 | 5.55 | 7.63 | | | | | -8.33E+00 | Indeterminate | | | -8.33E+00 | Indeterminate |
| 128 | p.Arg128Met | | | 2925 | 1150 | 4.99 | 4.48 | | | | | -1.08E+00 | Neutral | | | -1.08E+00 | Neutral |
| 128 | p.Arg128His | | | 1803 | 728 | 3.08 | 2.84 | | | | | -3.37E+00 | Neutral | | | -3.37E+00 | Neutral |
| 128 | p.Arg128Gln | | | 2303 | 651 | 3.93 | 2.54 | | | | | -1.07E-01 | Neutral | | | -1.07E-01 | Neutral |
| 128 | p.Arg128Pro | | Deleterious | 2278 | 2859 | 3.89 | 11.14 | | | | | -5.32E+01 | Deleterious | | | -5.32E+01 | Deleterious |
| 128 | p.Arg128Leu | | | 1945 | 682 | 3.32 | 2.66 | | | | | -1.25E+00 | Neutral | | | -1.25E+00 | Neutral |
| 128 | p.Arg128Asp | | | 2937 | 906 | 5.02 | 3.53 | | | | | -1.15E-01 | Neutral | | | -1.15E-01 | Neutral |
| 128 | p.Arg128Glu | | | 4092 | 1701 | 6.99 | 6.63 | | | | | -7.08E-01 | Neutral | | | -7.08E-01 | Neutral |
| 128 | p.Arg128Ala | | | 2727 | 946 | 4.66 | 3.69 | | | | | -4.82E-01 | Neutral | | | -4.82E-01 | Neutral |
| 128 | p.Arg128Gly | | | 3260 | 1838 | 5.57 | 7.16 | | | | | -6.37E+00 | Indeterminate | | | -6.37E+00 | Indeterminate |
| 128 | p.Arg128Val | | | 3216 | 980 | 5.49 | 3.82 | | | | | -6.95E-02 | Neutral | | | -6.95E-02 | Neutral |
| 128 | p.Arg128Tyr | | | 2778 | 1025 | 4.74 | 3.99 | | | | | -7.63E-01 | Neutral | | | -7.63E-01 | Neutral |
| 128 | p.Arg128Cys | | | 2996 | 723 | 5.12 | 2.82 | | | | | -3.13E-03 | Neutral | | | -3.13E-03 | Neutral |
| 128 | p.Arg128Trp | | | 3307 | 1044 | 5.65 | 4.07 | | | | | -9.54E-02 | Neutral | | | -9.54E-02 | Neutral |
| 128 | p.Arg128Phe | | | 3149 | 1776 | 5.38 | 6.92 | | | | | -6.69E+00 | Indeterminate | | | -6.69E+00 | Indeterminate |
| 129 | p.Tyr129Asn | | | 3172 | 1902 | 2.84 | 4.42 | | | | | -4.63E+01 | Indeterminate | | | -3.32E+01 | Indeterminate |
| 129 | p.Tyr129Lys | | | 5186 | 1692 | 4.64 | 3.93 | | | | | -3.97E+00 | Neutral | | | -3.97E+00 | Neutral |
| 129 | p.Tyr129Thr | | | 3503 | 1741 | 3.14 | 4.05 | | | | | -2.74E+01 | Indeterminate | | | -2.74E+01 | Indeterminate |
| 129 | p.Tyr129Arg | | | 6578 | 1764 | 5.89 | 4.10 | | | | | -7.14E-01 | Neutral | | | -7.14E-01 | Neutral |
| 129 | p.Tyr129Ser | | | 3563 | 958 | 3.19 | 2.23 | | | | | -2.79E+00 | Neutral | | | -2.79E+00 | Neutral |
| 129 | p.Tyr129Ile | | | 5889 | 2057 | 5.27 | 4.78 | | | | | -4.50E+00 | Neutral | | | -4.50E+00 | Neutral |
| 129 | p.Tyr129Met | | | 4837 | 1185 | 4.33 | 2.76 | | | | | -7.83E-01 | Neutral | | | -7.83E-01 | Neutral |
| 129 | p.Tyr129His | | | 5494 | 1466 | 4.92 | 3.41 | | | | | -1.08E+00 | Neutral | | | -1.08E+00 | Neutral |
| 129 | p.Tyr129Gln | | | 6782 | 1841 | 6.07 | 4.28 | | | | | -7.26E-01 | Neutral | | | -7.26E-01 | Neutral |
| 129 | p.Tyr129Pro | | | 4492 | 9559 | 4.02 | 22.23 | | | | | -5.32E+01 | Deleterious | | | -5.32E+01 | Deleterious |
| 129 | p.Tyr129Leu | | | 4705 | 1043 | 4.21 | 2.43 | | | | | -3.61E-01 | Neutral | | | -3.61E-01 | Neutral |
| 129 | p.Tyr129Asp | | | 7326 | 3885 | 6.56 | 9.03 | | | | | -1.65E+01 | Indeterminate | | | -1.65E+01 | Indeterminate |
| 129 | p.Tyr129Glu | | | 5802 | 1453 | 5.19 | 3.38 | | | | | -5.74E-01 | Neutral | | | -5.74E-01 | Neutral |
| 129 | p.Tyr129Ala | | | 8100 | 2262 | 7.25 | 5.26 | | | | | -5.65E-01 | Neutral | | | -5.65E-01 | Neutral |
| 129 | p.Tyr129Gly | | | 6216 | 2149 | 5.56 | 5.00 | | | | | -3.93E+00 | Neutral | | | -3.93E+00 | Neutral |
| 129 | p.Tyr129Val | | | 7809 | 2285 | 6.99 | 5.31 | | | | | -9.02E-01 | Neutral | | | -9.02E-01 | Neutral |
| 129 | p.Tyr129Tyr | Synonymous | | 4524 | 1129 | 4.05 | 2.63 | | | | | -1.06E+00 | Neutral | | | -1.06E+00 | Neutral |
| 129 | p.Tyr129Cys | | | 5819 | 1556 | 5.21 | 3.62 | | | | | -9.55E-01 | Neutral | | | -9.55E-01 | Neutral |
| 129 | p.Tyr129Trp | | | 6000 | 1338 | 5.37 | 3.11 | | | | | -1.71E-01 | Neutral | | | -1.71E-01 | Neutral |
| 129 | p.Tyr129Phe | | | 5935 | 1736 | 5.31 | 4.04 | | | | | -1.69E+00 | Neutral | | | -1.69E+00 | Neutral |
| 130 | p.Leu130Asn | | | 6622 | 5730 | 3.98 | 4.80 | | | | | -2.91E+01 | Indeterminate | | | -2.90E+01 | Indeterminate |
| 130 | p.Leu130Lys | | | 6586 | 10703 | 3.96 | 8.96 | | | | | -5.32E+01 | Deleterious | | | -5.32E+01 | Deleterious |
| 130 | p.Leu130Thr | | | 8414 | 4005 | 5.06 | 3.35 | | | | | -2.31E+00 | Neutral | | | -2.31E+00 | Neutral |
| 130 | p.Leu130Arg | | | 6954 | 11309 | 4.18 | 9.47 | | | | | -5.32E+01 | Deleterious | | | -5.32E+01 | Deleterious |
| 130 | p.Leu130Ser | | | 10649 | 5793 | 6.40 | 4.85 | | | | | -3.13E+00 | Neutral | | | -3.13E+00 | Neutral |
| 130 | p.Leu130Ile | | | 11747 | 4756 | 7.06 | 3.98 | | | | | -2.97E-01 | Neutral | | | -2.97E-01 | Neutral |
| 130 | p.Leu130Met | | | 8403 | 3454 | 5.05 | 2.89 | | | | | -8.77E-01 | Neutral | | | -8.77E-01 | Neutral |
| 130 | p.Leu130His | | | 5760 | 4572 | 3.46 | 3.83 | | | | | -2.62E+01 | Indeterminate | | | -2.62E+01 | Indeterminate |
| 130 | p.Leu130Gln | | | 6416 | 4936 | 3.86 | 4.13 | | | | | -2.19E+01 | Indeterminate | | | -2.19E+01 | Indeterminate |
| 130 | p.Leu130Pro | | | 5787 | 9545 | 3.48 | 7.99 | | | | | -5.32E+01 | Deleterious | | | -5.32E+01 | Deleterious |
| 130 | p.Leu130Leu | Synonymous | | 9828 | 4367 | 5.91 | 3.66 | | | | | -1.06E+00 | Neutral | | | -1.06E+00 | Neutral |
| 130 | p.Leu130Asp | | | 7930 | 10361 | 4.77 | 8.67 | | | | | -5.32E+01 | Deleterious | | | -5.32E+01 | Deleterious |
| 130 | p.Leu130Glu | | | 9503 | 11579 | 5.71 | 9.69 | | | | | -4.90E+01 | Indeterminate | | | -3.32E+01 | Indeterminate |
| 130 | p.Leu130Ala | | | 6719 | 3124 | 4.04 | 2.62 | | | | | -3.07E+00 | Neutral | | | -3.07E+00 | Neutral |
| 130 | p.Leu130Gly | | | 9249 | 4455 | 5.56 | 3.73 | | | | | -2.06E+00 | Neutral | | | -2.06E+00 | Neutral |
| 130 | p.Leu130Val | | | 9958 | 3841 | 5.99 | 3.22 | | | | | -3.15E-01 | Neutral | | | -3.15E-01 | Neutral |
| 130 | p.Leu130Tyr | | | 7425 | 3577 | 4.46 | 2.99 | | | | | -3.10E+00 | Neutral | | | -3.10E+00 | Neutral |
| 130 | p.Leu130Cys | | | 10316 | 4878 | 6.20 | 4.08 | | | | | -1.46E+00 | Neutral | | | -1.46E+00 | Neutral |
| 130 | p.Leu130Trp | | | 4919 | 2621 | 2.96 | 2.19 | | | | | -8.91E+00 | Indeterminate | | | -8.91E+00 | Indeterminate |
| 130 | p.Leu130Phe | | | 13178 | 5836 | 7.92 | 4.89 | | | | | -4.77E-01 | Neutral | | | -4.77E-01 | Neutral |
| 131 | p.Arg131Asn | | | 5042 | 2882 | 3.58 | 2.81 | | | | | -6.79E-01 | Neutral | | | -6.79E-01 | Neutral |
| 131 | p.Arg131Lys | | | 9318 | 4561 | 6.62 | 4.44 | | | | | -1.08E-02 | Neutral | | | -1.08E-02 | Neutral |
| 131 | p.Arg131Thr | | | 7380 | 7597 | 5.25 | 7.40 | | | | | -8.63E+00 | Indeterminate | | | -8.63E+00 | Indeterminate |
| 131 | p.Arg131Arg | Synonymous | | 4654 | 2747 | 3.31 | 2.68 | | | | | -1.06E+00 | Neutral | | | -1.06E+00 | Neutral |
| 131 | p.Arg131Ser | | | 6240 | 4107 | 4.44 | 4.00 | | | | | -1.15E+00 | Neutral | | | -1.15E+00 | Neutral |
| 131 | p.Arg131Ile | | | 7882 | 4014 | 5.60 | 3.91 | | | | | -4.23E-02 | Neutral | | | -4.23E-02 | Neutral |
| 131 | p.Arg131Met | | | 6830 | 3996 | 4.86 | 3.89 | | | | | -3.46E-01 | Neutral | | | -3.46E-01 | Neutral |
| 131 | p.Arg131His | | | 8168 | 5443 | 5.81 | 5.30 | | | | | -6.42E-01 | Neutral | | | -6.42E-01 | Neutral |
| 131 | p.Arg131Gln | | | 8309 | 11946 | 5.91 | 11.63 | | | | | -2.23E+01 | Indeterminate | | | -2.23E+01 | Indeterminate |
| 131 | p.Arg131Pro | | | 6340 | 7930 | 4.51 | 7.72 | | | | | -1.94E+01 | Indeterminate | | | -1.94E+01 | Indeterminate |
| 131 | p.Arg131Leu | | | 6498 | 2904 | 4.62 | 2.83 | | | | | -1.66E-02 | Neutral | | | -1.66E-02 | Neutral |
| 131 | p.Arg131Asp | | | 9630 | 4408 | 6.85 | 4.29 | | | | | -3.02E-03 | Neutral | | | -3.02E-03 | Neutral |
| 131 | p.Arg131Glu | | | 8105 | 4164 | 5.76 | 4.06 | | | | | -4.21E-02 | Neutral | | | -4.21E-02 | Neutral |
| 131 | p.Arg131Ala | | | 8275 | 5635 | 5.88 | 5.49 | | | | | -7.38E-01 | Neutral | | | -7.38E-01 | Neutral |
| 131 | p.Arg131Gly | | | 4870 | 2924 | 3.46 | 2.85 | | | | | -1.08E+00 | Neutral | | | -1.08E+00 | Neutral |
| 131 | p.Arg131Val | | | 4142 | 2191 | 2.94 | 2.13 | | | | | -6.14E-01 | Neutral | | | -6.14E-01 | Neutral |
| 131 | p.Arg131Tyr | | | 4020 | 2220 | 2.86 | 2.16 | | | | | -9.31E-01 | Neutral | | | -9.31E-01 | Neutral |
| 131 | p.Arg131Cys | | | 5494 | 12172 | 3.91 | 11.85 | | | | | -5.32E+01 | Deleterious | | | -5.32E+01 | Deleterious |
| 131 | p.Arg131Trp | | | 9383 | 6720 | 6.67 | 6.54 | | | | | -7.90E-01 | Neutral | | | -7.90E-01 | Neutral |
| 131 | p.Arg131Phe | | | 10081 | 4121 | 7.17 | 4.01 | | | | | -2.80E-04 | Neutral | | | -2.80E-04 | Neutral |
| 132 | p.Ala132Asn | | | 3857 | 2920 | 3.55 | 4.75 | | | | | -3.99E+01 | Indeterminate | | | -3.32E+01 | Indeterminate |
| 132 | p.Ala132Lys | | | 6605 | 2921 | 6.08 | 4.75 | | | | | -3.78E+00 | Neutral | | | -3.78E+00 | Neutral |
| 132 | p.Ala132Thr | | | 1388 | 575 | 1.28 | 0.94 | | | | | -2.04E+01 | Indeterminate | | | -2.04E+01 | Indeterminate |
| 132 | p.Ala132Arg | | | 8297 | 3403 | 7.63 | 5.54 | | | | | -1.61E+00 | Neutral | | | -1.61E+00 | Neutral |
| 132 | p.Ala132Ser | | | 4216 | 3329 | 3.88 | 5.42 | | | | | -4.11E+01 | Indeterminate | | | -3.32E+01 | Indeterminate |
| 132 | p.Ala132Ile | | | 4164 | 2141 | 3.83 | 3.48 | | | | | -1.29E+01 | Indeterminate | | | -1.29E+01 | Indeterminate |
| 132 | p.Ala132Met | | | 4378 | 2043 | 4.03 | 3.33 | | | | | -8.62E+00 | Indeterminate | | | -8.62E+00 | Indeterminate |
| 132 | p.Ala132His | | | 3831 | 2206 | 3.52 | 3.59 | | | | | -1.99E+01 | Indeterminate | | | -1.99E+01 | Indeterminate |
| 132 | p.Ala132Gln | | | 8852 | 3713 | 8.14 | 6.04 | | | | | -1.62E+00 | Neutral | | | -1.62E+00 | Neutral |
| 132 | p.Ala132Pro | | | 5579 | 2302 | 5.13 | 3.75 | | | | | -3.53E+00 | Neutral | | | -3.53E+00 | Neutral |
| 132 | p.Ala132Leu | | | 5922 | 3129 | 5.45 | 5.09 | | | | | -9.42E+00 | Indeterminate | | | -9.42E+00 | Indeterminate |
| 132 | p.Ala132Asp | | | 4119 | 2805 | 3.79 | 4.57 | | | | | -2.93E+01 | Indeterminate | | | -2.92E+01 | Indeterminate |
| 132 | p.Ala132Glu | | | 5719 | 3008 | 5.26 | 4.90 | | | | | -9.67E+00 | Indeterminate | | | -9.67E+00 | Indeterminate |
| 132 | p.Ala132Ala | Synonymous | | 6066 | 2109 | 5.58 | 3.43 | | | | | -1.06E+00 | Neutral | | | -1.06E+00 | Neutral |
| 132 | p.Ala132Gly | | | 4391 | 8761 | 4.04 | 14.26 | | | | | -5.32E+01 | Deleterious | | | -5.32E+01 | Deleterious |

| | | | | | | | | | | | | | | | | | |
|---|---|---|---|---|---|---|---|---|---|---|---|---|---|---|---|---|---|
| 132 | p.Ala132Val | | | 7251 | 3409 | 6.67 | 5.55 | | | | | -4.37E+00 | Neutral | | | -4.37E+00 | Neutral |
| 132 | p.Ala132Tyr | | | 5165 | 2556 | 4.75 | 4.16 | | | | | -8.74E+00 | Indeterminate | | | -8.74E+00 | Indeterminate |
| 132 | p.Ala132Cys | | | 8345 | 5592 | 7.68 | 9.10 | | | | | -1.44E+01 | Indeterminate | | | -1.44E+01 | Indeterminate |
| 132 | p.Ala132Trp | | | 5564 | 2457 | 5.12 | 4.00 | | | | | -4.92E+00 | Neutral | | | -4.92E+00 | Neutral |
| 132 | p.Ala132Phe | | | 4977 | 2057 | 4.58 | 3.35 | | | | | -4.29E+00 | Neutral | | | -4.29E+00 | Neutral |
| 133 | p.Ala133Asn | | | 4372 | 938 | 4.80 | 3.57 | | | | | -2.23E-02 | Neutral | | | -2.23E-02 | Neutral |
| 133 | p.Ala133Lys | | | 4380 | 1041 | 4.81 | 3.96 | | | | | -8.53E-02 | Neutral | | | -8.53E-02 | Neutral |
| 133 | p.Ala133Thr | | | 4331 | 1060 | 4.76 | 4.03 | | | | | -1.26E-01 | Neutral | | | -1.26E-01 | Neutral |
| 133 | p.Ala133Arg | | | 3954 | 1335 | 4.34 | 5.08 | | | | | -2.27E+00 | Neutral | | | -2.27E+00 | Neutral |
| 133 | p.Ala133Ser | | | 6658 | 1862 | 7.31 | 7.08 | | | | | -1.13E-01 | Neutral | | | -1.13E-01 | Neutral |
| 133 | p.Ala133Ile | | | 4627 | 1388 | 5.08 | 5.28 | | | | | -7.13E-01 | Neutral | | | -7.13E-01 | Neutral |
| 133 | p.Ala133Met | | | 6154 | 1438 | 6.76 | 5.47 | | | | | -1.48E-02 | Neutral | | | -1.48E-02 | Neutral |
| 133 | p.Ala133His | | | 2367 | 508 | 2.60 | 1.93 | | | | | -2.62E-01 | Neutral | | | -2.62E-01 | Neutral |
| 133 | p.Ala133Gln | | | 4448 | 1056 | 4.89 | 4.02 | | | | | -7.91E-02 | Neutral | | | -7.91E-02 | Neutral |
| 133 | p.Ala133Pro | | | 6362 | 1967 | 6.99 | 7.48 | | | | | -3.73E-01 | Neutral | | | -3.73E-01 | Neutral |
| 133 | p.Ala133Leu | | | 5900 | 1653 | 6.48 | 6.29 | | | | | -1.80E-01 | Neutral | | | -1.80E-01 | Neutral |
| 133 | p.Ala133Asp | | | 2399 | 878 | 2.63 | 3.34 | | | | | -7.43E+00 | Indeterminate | | | -7.43E+00 | Indeterminate |
| 133 | p.Ala133Glu | | | 4841 | 1739 | 5.32 | 6.62 | | | | | -2.21E+00 | Neutral | | | -2.21E+00 | Neutral |
| 133 | p.Ala133Ala | Synonymous | | 4332 | 1340 | 4.76 | 5.10 | | | | | -1.06E+00 | Neutral | | | -1.06E+00 | Neutral |
| 133 | p.Ala133Gly | | | 4928 | 1597 | 5.41 | 6.08 | | | | | -1.09E+00 | Neutral | | | -1.09E+00 | Neutral |
| 133 | p.Ala133Val | | | 5234 | 1964 | 5.75 | 7.47 | | | | | -2.44E+00 | Neutral | | | -2.44E+00 | Neutral |
| 133 | p.Ala133Tyr | | | 5290 | 1737 | 5.81 | 6.61 | | | | | -1.01E+00 | Neutral | | | -1.01E+00 | Neutral |
| 133 | p.Ala133Cys | | | 2580 | 572 | 2.83 | 2.18 | | | | | -2.72E-01 | Neutral | | | -2.72E-01 | Neutral |
| 133 | p.Ala133Trp | | | 4232 | 1160 | 4.65 | 4.41 | | | | | -4.27E-01 | Neutral | | | -4.27E-01 | Neutral |
| 133 | p.Ala133Phe | | | 3655 | 1054 | 4.01 | 4.01 | | | | | -9.60E-01 | Neutral | | | -9.60E-01 | Neutral |
| 134 | p.Ala134Asn | | | 8503 | 16401 | 4.84 | 4.71 | 3843 | 656 | 4.51 | 4.15 | -5.88E-01 | Neutral | -3.23E-02 | Neutral | -1.04E-01 | Neutral |
| 134 | p.Ala134Lys | | | 6884 | 14985 | 3.92 | 4.30 | 3773 | 681 | 4.43 | 4.31 | -2.24E+00 | Neutral | -7.23E-02 | Neutral | -9.30E-01 | Neutral |
| 134 | p.Ala134Thr | | Neutral | 7442 | 13995 | 4.24 | 4.02 | 2573 | 670 | 3.02 | 4.24 | -6.87E-01 | Neutral | -3.79E+00 | Neutral | -2.44E+00 | Neutral |
| 134 | p.Ala134Arg | | | 5325 | 12020 | 3.03 | 3.45 | 6217 | 601 | 7.30 | 3.80 | -4.25E+00 | Neutral | -6.81E-09 | Neutral | -2.27E+00 | Neutral |
| 134 | p.Ala134Ser | | | 9157 | 17350 | 5.21 | 4.98 | 3396 | 901 | 3.99 | 5.70 | -4.07E-01 | Neutral | -2.57E+00 | Neutral | -1.36E+00 | Neutral |
| 134 | p.Ala134Ile | | | 8079 | 14846 | 4.60 | 4.26 | 3017 | 641 | 3.54 | 4.06 | -4.49E-01 | Neutral | -7.67E-01 | Neutral | -3.34E-01 | Neutral |
| 134 | p.Ala134Met | | | 7883 | 15439 | 4.49 | 4.43 | 3536 | 666 | 4.15 | 4.22 | -8.14E-01 | Neutral | -1.53E-01 | Neutral | -2.27E-01 | Neutral |
| 134 | p.Ala134His | | | 9304 | 18154 | 5.29 | 5.21 | 4549 | 925 | 5.34 | 5.86 | -5.03E-01 | Neutral | -1.41E-01 | Neutral | -1.12E-01 | Neutral |
| 134 | p.Ala134Gln | | | 8265 | 16273 | 4.70 | 4.67 | 3771 | 773 | 4.43 | 4.89 | -7.50E-01 | Neutral | -2.94E-01 | Neutral | -2.58E-01 | Neutral |
| 134 | p.Ala134Pro | | | 5493 | 10521 | 3.13 | 3.02 | 3057 | 490 | 3.59 | 3.10 | -1.62E+00 | Neutral | -3.85E-02 | Neutral | -5.53E-01 | Neutral |
| 134 | p.Ala134Leu | | | 9637 | 18558 | 5.48 | 5.33 | 4933 | 845 | 5.79 | 5.35 | -4.04E-01 | Neutral | -1.01E-02 | Neutral | -5.00E-02 | Neutral |
| 134 | p.Ala134Asp | | | 10375 | 19782 | 5.90 | 5.68 | 4062 | 849 | 4.77 | 5.37 | -2.92E-01 | Neutral | -2.79E-01 | Neutral | -9.00E-02 | Neutral |
| 134 | p.Ala134Glu | | | 11179 | 23452 | 6.36 | 6.74 | 7244 | 1068 | 8.51 | 6.76 | -5.55E-01 | Neutral | -5.89E-05 | Neutral | -8.56E-02 | Neutral |
| 134 | p.Ala134Ala | Synonymous | | 6807 | 13159 | 3.87 | 3.78 | 2772 | 598 | 3.26 | 3.79 | -1.06E+00 | Neutral | -1.06E+00 | Neutral | -8.18E-01 | Neutral |
| 134 | p.Ala134Gly | | | 7599 | 15473 | 4.32 | 4.44 | 3125 | 574 | 3.67 | 3.63 | -1.18E+00 | Neutral | -1.81E-01 | Neutral | -4.05E-01 | Neutral |
| 134 | p.Ala134Val | | | 11867 | 23859 | 6.75 | 6.85 | 5268 | 1204 | 6.19 | 7.62 | -3.17E-01 | Neutral | -2.87E-01 | Neutral | -9.96E-02 | Neutral |
| 134 | p.Ala134Tyr | | | 11715 | 22207 | 6.67 | 6.38 | 4204 | 1130 | 4.94 | 7.15 | -1.83E-01 | Neutral | -1.83E+00 | Neutral | -7.53E-01 | Neutral |
| 134 | p.Ala134Cys | | | 11760 | 21834 | 6.69 | 6.27 | 3466 | 958 | 4.07 | 6.06 | -1.44E-01 | Neutral | -3.10E+00 | Neutral | -1.54E+00 | Neutral |
| 134 | p.Ala134Trp | | | 9783 | 23774 | 5.57 | 6.83 | 8758 | 932 | 10.28 | 5.90 | -2.15E+00 | Neutral | -4.19E-09 | Neutral | -8.35E-01 | Neutral |
| 134 | p.Ala134Phe | | | 8664 | 16053 | 4.93 | 4.61 | 3592 | 634 | 4.22 | 4.01 | -3.92E-01 | Neutral | -6.71E-02 | Neutral | -6.06E-02 | Neutral |
| 135 | p.Gly135Asn | | | 7098 | 1823 | 4.57 | 3.10 | | | | | -1.29E-02 | Neutral | | | -1.29E-02 | Neutral |
| 135 | p.Gly135Lys | | | 5037 | 2084 | 3.24 | 3.55 | | | | | -3.29E+00 | Neutral | | | -3.29E+00 | Neutral |
| 135 | p.Gly135Thr | | | 5864 | 2062 | 3.77 | 3.51 | | | | | -8.61E-01 | Neutral | | | -8.61E-01 | Neutral |
| 135 | p.Gly135Arg | | | 13028 | 3421 | 8.38 | 5.82 | | | | | -5.94E-04 | Neutral | | | -5.94E-04 | Neutral |
| 135 | p.Gly135Ser | | | 4979 | 1773 | 3.20 | 3.02 | | | | | -1.39E+00 | Neutral | | | -1.39E+00 | Neutral |
| 135 | p.Gly135Ile | | | 8051 | 3609 | 5.18 | 6.14 | | | | | -2.16E+00 | Neutral | | | -2.16E+00 | Neutral |
| 135 | p.Gly135Met | | | 12233 | 4835 | 7.87 | 8.23 | | | | | -2.87E-01 | Neutral | | | -2.87E-01 | Neutral |
| 135 | p.Gly135His | | | 11507 | 4705 | 7.40 | 8.01 | | | | | -4.74E-01 | Neutral | | | -4.74E-01 | Neutral |
| 135 | p.Gly135Gln | | | 6056 | 2112 | 3.90 | 3.59 | | | | | -7.44E-01 | Neutral | | | -7.44E-01 | Neutral |
| 135 | p.Gly135Pro | | | 8775 | 3358 | 5.65 | 5.71 | | | | | -5.81E-01 | Neutral | | | -5.81E-01 | Neutral |
| 135 | p.Gly135Leu | | | 8259 | 3502 | 5.31 | 5.96 | | | | | -1.45E+00 | Neutral | | | -1.45E+00 | Neutral |
| 135 | p.Gly135Asp | | | 5506 | 1975 | 3.54 | 3.36 | | | | | -1.16E+00 | Neutral | | | -1.16E+00 | Neutral |
| 135 | p.Gly135Glu | | | 4893 | 3812 | 3.15 | 6.49 | | | | | -3.14E+01 | Indeterminate | | | -3.10E+01 | Indeterminate |
| 135 | p.Gly135Ala | | | 6693 | 2173 | 4.31 | 3.70 | | | | | -2.97E-01 | Neutral | | | -2.97E-01 | Neutral |
| 135 | p.Gly135Gly | Synonymous | | 8277 | 3357 | 5.33 | 5.71 | | | | | -1.06E+00 | Neutral | | | -1.06E+00 | Neutral |
| 135 | p.Gly135Val | | | 5528 | 2052 | 3.56 | 3.49 | | | | | -1.45E+00 | Neutral | | | -1.45E+00 | Neutral |
| 135 | p.Gly135Tyr | | | 7744 | 2852 | 4.98 | 4.85 | | | | | -5.97E-01 | Neutral | | | -5.97E-01 | Neutral |
| 135 | p.Gly135Cys | | | 8671 | 2609 | 5.58 | 4.44 | | | | | -4.73E-02 | Neutral | | | -4.73E-02 | Neutral |
| 135 | p.Gly135Trp | | | 9281 | 3513 | 5.97 | 5.98 | | | | | -4.51E-01 | Neutral | | | -4.51E-01 | Neutral |
| 135 | p.Gly135Phe | | | 7947 | 3144 | 5.11 | 5.35 | | | | | -9.77E-01 | Neutral | | | -9.77E-01 | Neutral |
| 136 | p.Gly136Asn | | | 10420 | 5992 | 5.71 | 5.62 | 12915 | 6828 | 5.88 | 9.49 | -5.84E-01 | Neutral | -9.84E+00 | Indeterminate | -7.38E+00 | Indeterminate |
| 136 | p.Gly136Lys | | | 9133 | 4870 | 5.00 | 4.57 | 10618 | 2631 | 4.83 | 3.66 | -4.44E-01 | Neutral | -9.93E-02 | Neutral | -8.23E-02 | Neutral |
| 136 | p.Gly136Thr | | | 11568 | 6636 | 6.33 | 6.22 | 13564 | 1561 | 6.18 | 2.17 | -4.23E-01 | Neutral | -2.85E-09 | Neutral | -5.21E-02 | Neutral |
| 136 | p.Gly136Arg | | | 13087 | 7645 | 7.17 | 7.17 | 15369 | 3820 | 7.00 | 5.31 | -3.43E-01 | Neutral | -2.07E-02 | Neutral | -3.94E-02 | Neutral |
| 136 | p.Gly136Ser | | | 9479 | 5468 | 5.19 | 5.13 | 11411 | 5318 | 5.20 | 7.39 | -7.72E-01 | Neutral | -7.10E+00 | Indeterminate | -5.18E+00 | Neutral |
| 136 | p.Gly136Ile | | | 7498 | 4411 | 4.11 | 4.14 | 9506 | 1638 | 4.33 | 2.28 | -1.55E+00 | Neutral | -6.45E-04 | Neutral | -4.96E-01 | Neutral |
| 136 | p.Gly136Met | | | 10463 | 6084 | 5.73 | 5.70 | 13373 | 2252 | 6.09 | 3.13 | -6.33E-01 | Neutral | -4.03E-05 | Neutral | -1.08E-01 | Neutral |
| 136 | p.Gly136His | | | 10513 | 5586 | 5.76 | 5.24 | 11604 | 6173 | 5.28 | 8.58 | -2.78E-01 | Neutral | -1.14E+01 | Indeterminate | -8.50E+00 | Indeterminate |
| 136 | p.Gly136Gln | | | 8073 | 4302 | 4.42 | 4.03 | 9656 | 3063 | 4.40 | 4.26 | -6.27E-01 | Neutral | -1.26E+00 | Neutral | -6.79E-01 | Neutral |
| 136 | p.Gly136Pro | | | 6566 | 3880 | 3.59 | 3.64 | 8140 | 4546 | 3.71 | 6.32 | -2.09E+00 | Neutral | -1.93E+01 | Indeterminate | -1.74E+01 | Indeterminate |
| 136 | p.Gly136Leu | | | 9190 | 5242 | 5.03 | 4.92 | 11122 | 5391 | 5.06 | 7.49 | -7.66E-01 | Neutral | -8.57E+00 | Indeterminate | -6.43E+00 | Indeterminate |
| 136 | p.Gly136Asp | | | 9243 | 5278 | 5.06 | 4.95 | 10964 | 7479 | 4.99 | 10.40 | -7.61E-01 | Neutral | -2.56E+01 | Indeterminate | -2.20E+01 | Indeterminate |
| 136 | p.Gly136Glu | | | 6896 | 3746 | 3.78 | 3.51 | 8983 | 1778 | 4.09 | 2.47 | -1.09E+00 | Neutral | -9.84E-03 | Neutral | -2.80E-01 | Neutral |
| 136 | p.Gly136Ala | | | 8356 | 5198 | 4.58 | 4.87 | 9671 | 1125 | 4.40 | 1.56 | -1.76E+00 | Neutral | -7.70E-08 | Neutral | -6.07E-01 | Neutral |
| 136 | p.Gly136Gly | Synonymous | | 11614 | 7478 | 6.36 | 7.01 | 14031 | 4919 | 6.39 | 6.84 | -1.06E+00 | Neutral | -1.06E+00 | Neutral | -8.18E-01 | Neutral |
| 136 | p.Gly136Val | | | 12033 | 7432 | 6.59 | 6.97 | 14744 | 2105 | 6.71 | 2.93 | -7.08E-01 | Neutral | -4.00E-07 | Neutral | -1.32E-01 | Neutral |
| 136 | p.Gly136Tyr | | | 7640 | 5300 | 4.18 | 4.97 | 9352 | 2412 | 4.26 | 3.35 | -3.80E+00 | Neutral | -2.43E-01 | Neutral | -2.12E+00 | Neutral |
| 136 | p.Gly136Cys | | | 7914 | 4580 | 4.33 | 4.29 | 9143 | 4462 | 4.16 | 6.20 | -1.23E+00 | Neutral | -1.11E+01 | Indeterminate | -9.10E+00 | Indeterminate |
| 136 | p.Gly136Trp | | | 4677 | 2955 | 2.56 | 2.77 | 5657 | 474 | 2.58 | 0.66 | -5.27E+00 | Neutral | -1.35E-09 | Neutral | -3.05E+00 | Neutral |
| 136 | p.Gly136Phe | | | 8281 | 4565 | 4.53 | 4.28 | 9815 | 3962 | 4.47 | 5.51 | -7.69E-01 | Neutral | -4.74E+00 | Neutral | -3.24E+00 | Neutral |
| 137 | p.Thr137Asn | | | 12637 | 7950 | 6.42 | 6.37 | | | | | -5.48E-01 | Neutral | | | -5.48E-01 | Neutral |
| 137 | p.Thr137Lys | | | 10056 | 5827 | 5.11 | 4.67 | | | | | -5.19E-01 | Neutral | | | -5.19E-01 | Neutral |
| 137 | p.Thr137Thr | Synonymous | | 9153 | 5630 | 4.65 | 4.51 | | | | | -1.06E+00 | Neutral | | | -1.06E+00 | Neutral |
| 137 | p.Thr137Arg | | | 10802 | 7256 | 5.48 | 5.81 | | | | | -1.34E+00 | Neutral | | | -1.34E+00 | Neutral |
| 137 | p.Thr137Ser | | | 15695 | 10181 | 7.97 | 8.15 | | | | | -3.80E-01 | Neutral | | | -3.80E-01 | Neutral |
| 137 | p.Thr137Ile | | | 5926 | 3761 | 3.01 | 3.01 | | | | | -3.13E+00 | Neutral | | | -3.13E+00 | Neutral |
| 137 | p.Thr137Met | | | 15098 | 10222 | 7.67 | 8.18 | | | | | -6.18E-01 | Neutral | | | -6.18E-01 | Neutral |
| 137 | p.Thr137His | | | 7921 | 5006 | 4.02 | 4.01 | | | | | -1.76E+00 | Neutral | | | -1.76E+00 | Neutral |
| 137 | p.Thr137Gln | | | 2342 | 1107 | 1.19 | 0.89 | | | | | -3.36E+00 | Neutral | | | -3.36E+00 | Neutral |
| 137 | p.Thr137Pro | | | 6861 | 3500 | 3.48 | 2.80 | | | | | -5.17E-01 | Neutral | | | -5.17E-01 | Neutral |
| 137 | p.Thr137Leu | | | 11186 | 7011 | 5.68 | 5.61 | | | | | -7.40E-01 | Neutral | | | -7.40E-01 | Neutral |
| 137 | p.Thr137Asp | | | 6374 | 3772 | 3.24 | 3.02 | | | | | -1.83E+00 | Neutral | | | -1.83E+00 | Neutral |
| 137 | p.Thr137Glu | | | 9691 | 6160 | 4.92 | 4.93 | | | | | -1.17E+00 | Neutral | | | -1.17E+00 | Neutral |
| 137 | p.Thr137Ala | | | 8690 | 6226 | 4.41 | 4.98 | | | | | -3.04E+00 | Neutral | | | -3.04E+00 | Neutral |
| 137 | p.Thr137Gly | | | 14994 | 10033 | 7.61 | 8.03 | | | | | -5.71E-01 | Neutral | | | -5.71E-01 | Neutral |
| 137 | p.Thr137Val | | | 17827 | 12301 | 9.05 | 9.85 | | | | | -4.53E-01 | Neutral | | | -4.53E-01 | Neutral |
| 137 | p.Thr137Tyr | | | 7543 | 4012 | 3.83 | 3.21 | | | | | -5.64E-01 | Neutral | | | -5.64E-01 | Neutral |
| 137 | p.Thr137Cys | | | 4535 | 3046 | 2.30 | 2.44 | | | | | -6.22E+00 | Indeterminate | | | -6.22E+00 | Indeterminate |
| 137 | p.Thr137Trp | | | 9317 | 5548 | 4.73 | 4.44 | | | | | -7.97E-01 | Neutral | | | -7.97E-01 | Neutral |
| 137 | p.Thr137Phe | | | 10305 | 6346 | 5.23 | 5.08 | | | | | -7.98E-01 | Neutral | | | -7.98E-01 | Neutral |
| 138 | p.Arg138Asn | | | 9124 | 4360 | 3.23 | 3.99 | | | | | -1.57E+00 | Neutral | | | -1.57E+00 | Neutral |
| 138 | p.Arg138Lys | | | 16369 | 6363 | 5.80 | 5.82 | | | | | -3.22E-02 | Neutral | | | -3.22E-02 | Neutral |
| 138 | p.Arg138Thr | | | 14620 | 6499 | 5.18 | 5.94 | | | | | -2.46E-01 | Neutral | | | -2.46E-01 | Neutral |
| 138 | p.Arg138Arg | Synonymous | | 7495 | 3167 | 2.65 | 2.90 | | | | | -1.06E+00 | Neutral | | | -1.06E+00 | Neutral |
| 138 | p.Arg138Ser | | | 15989 | 5361 | 5.66 | 4.90 | | | | | -3.76E-03 | Neutral | | | -3.76E-03 | Neutral |
| 138 | p.Arg138Ile | | | 21820 | 8258 | 7.73 | 7.55 | | | | | -5.06E-03 | Neutral | | | -5.06E-03 | Neutral |
| 138 | p.Arg138Met | | | 27553 | 10505 | 9.76 | 9.61 | | | | | -1.51E-03 | Neutral | | | -1.51E-03 | Neutral |
| 138 | p.Arg138His | | | 7027 | 2951 | 2.49 | 2.70 | | | | | -1.18E+00 | Neutral | | | -1.18E+00 | Neutral |
| 138 | p.Arg138Gln | | | 15327 | 6033 | 5.43 | 5.52 | | | | | -5.09E-02 | Neutral | | | -5.09E-02 | Neutral |
| 138 | p.Arg138Pro | | | 6749 | 3535 | 2.39 | 3.23 | | | | | -4.52E+00 | Neutral | | | -4.52E+00 | Neutral |
| 138 | p.Arg138Leu | | | 33430 | 13311 | 11.84 | 12.17 | | | | | -1.03E-03 | Neutral | | | -1.03E-03 | Neutral |
| 138 | p.Arg138Asp | | | 16062 | 6554 | 5.69 | 5.99 | | | | | -6.60E-02 | Neutral | | | -6.60E-02 | Neutral |
| 138 | p.Arg138Glu | | | 6604 | 2634 | 2.34 | 2.41 | | | | | -9.47E-01 | Neutral | | | -9.47E-01 | Neutral |
| 138 | p.Arg138Ala | | | 1275 | 310 | 0.45 | 0.28 | | | | | -1.36E+00 | Neutral | | | -1.36E+00 | Neutral |
| 138 | p.Arg138Gly | | | 9750 | 3959 | 3.45 | 3.62 | | | | | -3.76E-01 | Neutral | | | -3.76E-01 | Neutral |
| 138 | p.Arg138Val | | | 10559 | 4030 | 3.74 | 3.69 | | | | | -1.54E-01 | Neutral | | | -1.54E-01 | Neutral |
| 138 | p.Arg138Tyr | | | 15818 | 4829 | 5.60 | 4.42 | | | | | -7.49E-04 | Neutral | | | -7.49E-04 | Neutral |
| 138 | p.Arg138Cys | | | 18784 | 6819 | 6.65 | 6.24 | | | | | -5.78E-03 | Neutral | | | -5.78E-03 | Neutral |
| 138 | p.Arg138Trp | | | 17148 | 5835 | 6.07 | 5.34 | | | | | -3.25E-03 | Neutral | | | -3.25E-03 | Neutral |
| 138 | p.Arg138Phe | | | 10816 | 4048 | 3.83 | 3.70 | | | | | -1.13E-01 | Neutral | | | -1.13E-01 | Neutral |
| 139 | p.Gly139Asn | | | 8496 | 2550 | 3.53 | 3.45 | 3712 | 1812 | 3.95 | 3.41 | -2.29E+00 | Neutral | -2.33E+00 | Neutral | -2.55E+00 | Neutral |
| 139 | p.Gly139Lys | | | 12403 | 4023 | 5.15 | 5.44 | 4555 | 2814 | 4.84 | 5.29 | -1.71E+00 | Neutral | -5.43E+00 | Neutral | -4.57E+00 | Neutral |
| 139 | p.Gly139Thr | | | 8144 | 2197 | 3.38 | 2.97 | 2926 | 1791 | 3.11 | 3.37 | -1.28E+00 | Neutral | -9.45E+00 | Indeterminate | -7.65E+00 | Indeterminate |
| 139 | p.Gly139Arg | | Neutral | 12860 | 3446 | 5.34 | 4.66 | 4701 | 2594 | 5.00 | 4.88 | -3.60E-01 | Neutral | -2.98E+00 | Neutral | -1.61E+00 | Neutral |
| 139 | p.Gly139Ser | | | 13019 | 4819 | 5.41 | 6.51 | 5676 | 3432 | 6.04 | 6.45 | -3.32E+00 | Neutral | -3.46E+00 | Neutral | -4.27E+00 | Neutral |
| 139 | p.Gly139Ile | | | 15538 | 4725 | 6.46 | 6.39 | 5905 | 3339 | 6.28 | 6.28 | -6.28E-01 | Neutral | -2.24E+00 | Neutral | -1.29E+00 | Neutral |
| 139 | p.Gly139Met | | | 11545 | 4056 | 4.80 | 5.48 | 4565 | 2731 | 4.85 | 5.14 | -3.11E+00 | Neutral | -4.68E+00 | Neutral | -5.11E+00 | Neutral |
| 139 | p.Gly139His | | | 11470 | 3114 | 4.77 | 4.21 | 4453 | 2398 | 4.74 | 4.51 | -5.65E-01 | Neutral | -2.88E+00 | Neutral | -1.68E+00 | Neutral |
| 139 | p.Gly139Gln | | | 10561 | 3014 | 4.39 | 4.07 | 4842 | 2382 | 5.15 | 4.48 | -1.03E+00 | Neutral | -1.42E+00 | Neutral | -1.02E+00 | Neutral |
| 139 | p.Gly139Pro | | | 8278 | 3312 | 3.44 | 4.48 | 4736 | 2102 | 5.04 | 3.95 | -9.03E+00 | Indeterminate | -7.05E-01 | Neutral | -6.78E+00 | Indeterminate |
| 139 | p.Gly139Leu | | | 13458 | 3515 | 5.59 | 4.75 | 4746 | 3057 | 5.05 | 5.75 | -2.42E-01 | Neutral | -6.13E+00 | Neutral | -3.93E+00 | Neutral |
| 139 | p.Gly139Asp | | | 12264 | 3368 | 5.10 | 4.55 | 3626 | 2391 | 3.86 | 4.50 | -5.16E-01 | Neutral | -9.57E+00 | Indeterminate | -7.08E+00 | Indeterminate |
| 139 | p.Gly139Glu | | | 14048 | 4837 | 5.84 | 6.54 | 5228 | 3110 | 5.56 | 5.85 | -1.92E+00 | Neutral | -3.66E+00 | Neutral | -3.30E+00 | Neutral |
| 139 | p.Gly139Ala | | | 10712 | 2633 | 4.45 | 3.56 | 3613 | 2393 | 3.84 | 4.50 | -2.81E-01 | Neutral | -9.76E+00 | Indeterminate | -7.05E+00 | Indeterminate |
| 139 | p.Gly139Gly | Synonymous | | 13109 | 4034 | 5.45 | 5.45 | 5129 | 2467 | 5.45 | 4.64 | -1.06E+00 | Neutral | -1.06E+00 | Neutral | -8.18E-01 | Neutral |

| | | | | | | | | | | | | | | | | | |
|---|---|---|---|---|---|---|---|---|---|---|---|---|---|---|---|---|---|
| 139 | p.Gly139Val | | | 15407 | 4203 | 6.40 | 5.68 | 6369 | 3174 | 6.77 | 5.97 | -2.39E-01 | Neutral | -8.09E-01 | Neutral | -2.60E-01 | Neutral |
| 139 | p.Gly139Tyr | | | 10886 | 4620 | 4.52 | 6.24 | 4241 | 2748 | 4.51 | 5.17 | -8.00E+00 | Indeterminate | -7.33E+00 | Indeterminate | -1.18E+01 | Indeterminate |
| 139 | p.Gly139Cys | | | 12796 | 3369 | 5.32 | 4.55 | 5017 | 2479 | 5.34 | 4.66 | -3.09E-01 | Neutral | -1.35E+00 | Neutral | -5.54E-01 | Neutral |
| 139 | p.Gly139Trp | | | 12302 | 3446 | 5.11 | 4.66 | 4799 | 2716 | 5.10 | 5.11 | -6.04E-01 | Neutral | -3.28E+00 | Neutral | -2.00E+00 | Neutral |
| 139 | p.Gly139Phe | | | 13311 | 4715 | 5.53 | 6.37 | 5200 | 3243 | 5.53 | 6.10 | -2.52E+00 | Neutral | -4.64E+00 | Neutral | -4.58E+00 | Neutral |
| 140 | p.Ser140Asn | | | 10552 | 7922 | 5.17 | 5.24 | | | | | -7.53E-01 | Neutral | | | -7.53E-01 | Neutral |
| 140 | p.Ser140Lys | | | 11164 | 7762 | 5.47 | 5.14 | | | | | -3.30E-01 | Neutral | | | -3.30E-01 | Neutral |
| 140 | p.Ser140Thr | | | 12855 | 9304 | 6.30 | 6.16 | | | | | -3.09E-01 | Neutral | | | -3.09E-01 | Neutral |
| 140 | p.Ser140Arg | | | 7945 | 6865 | 3.89 | 4.54 | | | | | -3.36E+00 | Neutral | | | -3.36E+00 | Neutral |
| 140 | p.Ser140Ser | Synonymous | | 8620 | 6334 | 4.23 | 4.19 | | | | | -1.06E+00 | Neutral | | | -1.06E+00 | Neutral |
| 140 | p.Ser140Ile | | | 15602 | 10874 | 7.65 | 7.20 | | | | | -1.05E-01 | Neutral | | | -1.05E-01 | Neutral |
| 140 | p.Ser140Met | | | 7484 | 5193 | 3.67 | 3.44 | | | | | -9.84E-01 | Neutral | | | -9.84E-01 | Neutral |
| 140 | p.Ser140His | | | 9812 | 7506 | 4.81 | 4.97 | | | | | -1.04E+00 | Neutral | | | -1.04E+00 | Neutral |
| 140 | p.Ser140Gln | | | 11521 | 8353 | 5.65 | 5.53 | | | | | -4.40E-01 | Neutral | | | -4.40E-01 | Neutral |
| 140 | p.Ser140Pro | | | 5355 | 4736 | 2.62 | 3.13 | | | | | -6.82E+00 | Indeterminate | | | -6.82E+00 | Indeterminate |
| 140 | p.Ser140Leu | | | 9344 | 6183 | 4.58 | 4.09 | | | | | -3.61E-01 | Neutral | | | -3.61E-01 | Neutral |
| 140 | p.Ser140Asp | | | 15150 | 10921 | 7.43 | 7.23 | | | | | -1.71E-01 | Neutral | | | -1.71E-01 | Neutral |
| 140 | p.Ser140Glu | | | 9632 | 6353 | 4.72 | 4.20 | | | | | -3.18E-01 | Neutral | | | -3.18E-01 | Neutral |
| 140 | p.Ser140Ala | | | 4610 | 3616 | 2.26 | 2.39 | | | | | -5.04E+00 | Neutral | | | -5.04E+00 | Neutral |
| 140 | p.Ser140Gly | | | 6200 | 5842 | 3.04 | 3.87 | | | | | -7.24E+00 | Indeterminate | | | -7.24E+00 | Indeterminate |
| 140 | p.Ser140Val | | | 17043 | 12142 | 8.35 | 8.04 | | | | | -9.60E-02 | Neutral | | | -9.60E-02 | Neutral |
| 140 | p.Ser140Tyr | | | 15829 | 12798 | 7.76 | 8.47 | | | | | -4.54E-01 | Neutral | | | -4.54E-01 | Neutral |
| 140 | p.Ser140Cys | | | 11151 | 8478 | 5.47 | 5.61 | | | | | -7.20E-01 | Neutral | | | -7.20E-01 | Neutral |
| 140 | p.Ser140Trp | | | 7713 | 5562 | 3.78 | 3.68 | | | | | -1.21E+00 | Neutral | | | -1.21E+00 | Neutral |
| 140 | p.Ser140Phe | | | 6438 | 4352 | 3.16 | 2.88 | | | | | -1.17E+00 | Neutral | | | -1.17E+00 | Neutral |
| 141 | p.Asn141Asn | Synonymous | | 9534 | 7176 | 4.94 | 5.44 | | | | | -1.06E+00 | Neutral | | | -1.06E+00 | Neutral |
| 141 | p.Asn141Lys | | | 6445 | 3767 | 3.34 | 2.86 | | | | | -3.91E-01 | Neutral | | | -3.91E-01 | Neutral |
| 141 | p.Asn141Thr | | | 5628 | 4142 | 2.91 | 3.14 | | | | | -2.77E+00 | Neutral | | | -2.77E+00 | Neutral |
| 141 | p.Asn141Arg | | | 9129 | 6690 | 4.73 | 5.07 | | | | | -9.71E-01 | Neutral | | | -9.71E-01 | Neutral |
| 141 | p.Asn141Ser | | | 9765 | 6185 | 5.06 | 4.69 | | | | | -2.24E-01 | Neutral | | | -2.24E-01 | Neutral |
| 141 | p.Asn141Ile | | | 11867 | 8445 | 6.14 | 6.41 | | | | | -3.71E-01 | Neutral | | | -3.71E-01 | Neutral |
| 141 | p.Asn141Met | | | 11856 | 8168 | 6.14 | 6.20 | | | | | -2.73E-01 | Neutral | | | -2.73E-01 | Neutral |
| 141 | p.Asn141His | | | 4415 | 2868 | 2.29 | 2.18 | | | | | -2.15E+00 | Neutral | | | -2.15E+00 | Neutral |
| 141 | p.Asn141Gln | | | 10967 | 8321 | 5.68 | 6.31 | | | | | -7.96E-01 | Neutral | | | -7.96E-01 | Neutral |
| 141 | p.Asn141Pro | | | 6338 | 4134 | 3.28 | 3.14 | | | | | -1.00E+00 | Neutral | | | -1.00E+00 | Neutral |
| 141 | p.Asn141Leu | | | 11622 | 7486 | 6.02 | 5.68 | | | | | -1.45E-01 | Neutral | | | -1.45E-01 | Neutral |
| 141 | p.Asn141Asp | | | 13047 | 9029 | 6.76 | 6.85 | | | | | -2.07E-01 | Neutral | | | -2.07E-01 | Neutral |
| 141 | p.Asn141Glu | | | 7938 | 5777 | 4.11 | 4.38 | | | | | -1.28E+00 | Neutral | | | -1.28E+00 | Neutral |
| 141 | p.Asn141Ala | | | 12198 | 8306 | 6.32 | 6.30 | | | | | -2.20E-01 | Neutral | | | -2.20E-01 | Neutral |
| 141 | p.Asn141Gly | | | 9905 | 6558 | 5.13 | 4.97 | | | | | -3.31E-01 | Neutral | | | -3.31E-01 | Neutral |
| 141 | p.Asn141Val | | | 15454 | 9860 | 8.00 | 7.48 | | | | | -4.06E-02 | Neutral | | | -4.06E-02 | Neutral |
| 141 | p.Asn141Tyr | | | 6746 | 4426 | 3.49 | 3.36 | | | | | -9.00E-01 | Neutral | | | -9.00E-01 | Neutral |
| 141 | p.Asn141Cys | | | 4124 | 2702 | 2.14 | 2.05 | | | | | -2.57E+00 | Neutral | | | -2.57E+00 | Neutral |
| 141 | p.Asn141Trp | | | 15020 | 9827 | 7.78 | 7.45 | | | | | -6.36E-02 | Neutral | | | -6.36E-02 | Neutral |
| 141 | p.Asn141Phe | | | 11136 | 7974 | 5.77 | 6.05 | | | | | -4.74E-01 | Neutral | | | -4.74E-01 | Neutral |
| 142 | p.His142Asn | | | 8383 | 1168 | 4.70 | 4.66 | | | | | -1.78E-04 | Neutral | | | -1.78E-04 | Neutral |
| 142 | p.His142Lys | | | 10971 | 1665 | 6.15 | 6.65 | | | | | -1.65E-04 | Neutral | | | -1.65E-04 | Neutral |
| 142 | p.His142Thr | | | 8686 | 1345 | 4.87 | 5.37 | | | | | -1.10E-03 | Neutral | | | -1.10E-03 | Neutral |
| 142 | p.His142Arg | | | 10575 | 1237 | 5.93 | 4.94 | | | | | -6.01E-07 | Neutral | | | -6.01E-07 | Neutral |
| 142 | p.His142Ser | | | 7833 | 1082 | 4.39 | 4.32 | | | | | -2.36E-04 | Neutral | | | -2.36E-04 | Neutral |
| 142 | p.His142Ile | | | 11025 | 1519 | 6.18 | 6.06 | | | | | -2.03E-05 | Neutral | | | -2.03E-05 | Neutral |
| 142 | p.His142Met | | | 7777 | 1104 | 4.36 | 4.41 | | | | | -4.24E-04 | Neutral | | | -4.24E-04 | Neutral |
| 142 | p.His142His | Synonymous | | 1320 | 190 | 0.74 | 0.76 | | | | | -1.06E+00 | Neutral | | | -1.06E+00 | Neutral |
| 142 | p.His142Gln | | | 6699 | 1050 | 3.76 | 4.19 | | | | | -6.05E-03 | Neutral | | | -6.05E-03 | Neutral |
| 142 | p.His142Pro | | | 5548 | 762 | 3.11 | 3.04 | | | | | -1.85E-03 | Neutral | | | -1.85E-03 | Neutral |
| 142 | p.His142Leu | | | 15858 | 2076 | 8.89 | 8.29 | | | | | -3.48E-07 | Neutral | | | -3.48E-07 | Neutral |
| 142 | p.His142Asp | | | 8253 | 1295 | 4.63 | 5.17 | | | | | -1.89E-03 | Neutral | | | -1.89E-03 | Neutral |
| 142 | p.His142Glu | | | 10412 | 1458 | 5.84 | 5.82 | | | | | -4.39E-05 | Neutral | | | -4.39E-05 | Neutral |
| 142 | p.His142Ala | | | 12203 | 1693 | 6.84 | 6.76 | | | | | -1.11E-05 | Neutral | | | -1.11E-05 | Neutral |
| 142 | p.His142Gly | | | 7160 | 966 | 4.01 | 3.86 | | | | | -2.68E-04 | Neutral | | | -2.68E-04 | Neutral |
| 142 | p.His142Val | | | 11560 | 1684 | 6.48 | 6.72 | | | | | -4.87E-05 | Neutral | | | -4.87E-05 | Neutral |
| 142 | p.His142Tyr | | | 8206 | 1063 | 4.60 | 4.24 | | | | | -4.50E-05 | Neutral | | | -4.50E-05 | Neutral |
| 142 | p.His142Cys | | | 9000 | 1249 | 5.05 | 4.99 | | | | | -1.01E-04 | Neutral | | | -1.01E-04 | Neutral |
| 142 | p.His142Trp | | | 9332 | 1224 | 5.23 | 4.89 | | | | | -2.33E-05 | Neutral | | | -2.33E-05 | Neutral |
| 142 | p.His142Phe | | | 7571 | 1225 | 4.24 | 4.89 | | | | | -5.23E-03 | Neutral | | | -5.23E-03 | Neutral |
| 143 | p.Ala143Asn | | | 10595 | 21434 | 4.99 | 5.07 | | | | | -1.04E+00 | Neutral | | | -1.04E+00 | Neutral |
| 143 | p.Ala143Lys | | | 11917 | 23232 | 5.61 | 5.49 | | | | | -5.77E-01 | Neutral | | | -5.77E-01 | Neutral |
| 143 | p.Ala143Thr | | Neutral | 12320 | 23747 | 5.80 | 5.61 | | | | | -4.77E-01 | Neutral | | | -4.77E-01 | Neutral |
| 143 | p.Ala143Arg | | | 10547 | 20563 | 4.96 | 4.86 | | | | | -8.01E-01 | Neutral | | | -8.01E-01 | Neutral |
| 143 | p.Ala143Ser | | | 9145 | 17169 | 4.30 | 4.06 | | | | | -8.62E-01 | Neutral | | | -8.62E-01 | Neutral |
| 143 | p.Ala143Ile | | | 9506 | 17634 | 4.47 | 4.17 | | | | | -7.11E-01 | Neutral | | | -7.11E-01 | Neutral |
| 143 | p.Ala143Met | | | 10605 | 21003 | 4.99 | 4.96 | | | | | -8.91E-01 | Neutral | | | -8.91E-01 | Neutral |
| 143 | p.Ala143His | | | 9550 | 18973 | 4.49 | 4.48 | | | | | -1.17E+00 | Neutral | | | -1.17E+00 | Neutral |
| 143 | p.Ala143Gln | | | 11085 | 22354 | 5.22 | 5.28 | | | | | -9.13E-01 | Neutral | | | -9.13E-01 | Neutral |
| 143 | p.Ala143Pro | | | 8704 | 17930 | 4.10 | 4.24 | | | | | -1.82E+00 | Neutral | | | -1.82E+00 | Neutral |
| 143 | p.Ala143Leu | | | 9306 | 17759 | 4.38 | 4.20 | | | | | -9.33E-01 | Neutral | | | -9.33E-01 | Neutral |
| 143 | p.Ala143Asp | | | 8557 | 15731 | 4.03 | 3.72 | | | | | -8.69E-01 | Neutral | | | -8.69E-01 | Neutral |
| 143 | p.Ala143Glu | | | 11690 | 23121 | 5.50 | 5.47 | | | | | -6.85E-01 | Neutral | | | -6.85E-01 | Neutral |
| 143 | p.Ala143Ala | Synonymous | | 10390 | 20931 | 4.89 | 4.95 | | | | | -1.06E+00 | Neutral | | | -1.06E+00 | Neutral |
| 143 | p.Ala143Gly | | | 12401 | 25612 | 5.84 | 6.05 | | | | | -8.25E-01 | Neutral | | | -8.25E-01 | Neutral |
| 143 | p.Ala143Val | | | 11134 | 21640 | 5.24 | 5.12 | | | | | -6.78E-01 | Neutral | | | -6.78E-01 | Neutral |
| 143 | p.Ala143Tyr | | | 10660 | 21796 | 5.02 | 5.15 | | | | | -1.11E+00 | Neutral | | | -1.11E+00 | Neutral |
| 143 | p.Ala143Cys | | | 10965 | 22961 | 5.16 | 5.43 | | | | | -1.23E+00 | Neutral | | | -1.23E+00 | Neutral |
| 143 | p.Ala143Trp | | | 11086 | 24426 | 5.22 | 5.77 | | | | | -1.68E+00 | Neutral | | | -1.68E+00 | Neutral |
| 143 | p.Ala143Phe | | | 12322 | 25053 | 5.80 | 5.92 | | | | | -7.43E-01 | Neutral | | | -7.43E-01 | Neutral |
| 144 | p.Arg144Asn | | | 19759 | 3661 | 6.06 | 5.83 | | | | | -1.14E-01 | Neutral | | | -1.14E-01 | Neutral |
| 144 | p.Arg144Lys | | | 11748 | 2225 | 3.60 | 3.54 | | | | | -7.41E-01 | Neutral | | | -7.41E-01 | Neutral |
| 144 | p.Arg144Thr | | | 15043 | 2830 | 4.61 | 4.50 | | | | | -3.45E-01 | Neutral | | | -3.45E-01 | Neutral |
| 144 | p.Arg144Arg | Synonymous | | 13943 | 2932 | 4.28 | 4.67 | | | | | -1.06E+00 | Neutral | | | -1.06E+00 | Neutral |
| 144 | p.Arg144Ser | | | 13559 | 2514 | 4.16 | 4.00 | | | | | -4.14E-01 | Neutral | | | -4.14E-01 | Neutral |
| 144 | p.Arg144Ile | | | 13225 | 2572 | 4.06 | 4.09 | | | | | -6.69E-01 | Neutral | | | -6.69E-01 | Neutral |
| 144 | p.Arg144Met | | | 14812 | 2776 | 4.54 | 4.42 | | | | | -3.49E-01 | Neutral | | | -3.49E-01 | Neutral |
| 144 | p.Arg144His | | | 13721 | 2644 | 4.21 | 4.21 | | | | | -5.60E-01 | Neutral | | | -5.60E-01 | Neutral |
| 144 | p.Arg144Gln | | | 16857 | 3371 | 5.17 | 5.37 | | | | | -4.27E-01 | Neutral | | | -4.27E-01 | Neutral |
| 144 | p.Arg144Pro | | | 9345 | 1766 | 2.87 | 2.81 | | | | | -1.27E+00 | Neutral | | | -1.27E+00 | Neutral |
| 144 | p.Arg144Leu | | | 23232 | 4702 | 7.13 | 7.48 | | | | | -1.68E-01 | Neutral | | | -1.68E-01 | Neutral |
| 144 | p.Arg144Asp | | | 13837 | 2604 | 4.24 | 4.15 | | | | | -4.45E-01 | Neutral | | | -4.45E-01 | Neutral |
| 144 | p.Arg144Glu | | | 16608 | 3337 | 5.09 | 5.31 | | | | | -4.65E-01 | Neutral | | | -4.65E-01 | Neutral |
| 144 | p.Arg144Ala | | | 17489 | 3491 | 5.36 | 5.56 | | | | | -3.75E-01 | Neutral | | | -3.75E-01 | Neutral |
| 144 | p.Arg144Gly | | | 13921 | 2836 | 4.27 | 4.51 | | | | | -8.42E-01 | Neutral | | | -8.42E-01 | Neutral |
| 144 | p.Arg144Val | | | 19701 | 3764 | 6.04 | 5.99 | | | | | -1.63E-01 | Neutral | | | -1.63E-01 | Neutral |
| 144 | p.Arg144Tyr | | | 16291 | 3119 | 5.00 | 4.96 | | | | | -3.17E-01 | Neutral | | | -3.17E-01 | Neutral |
| 144 | p.Arg144Cys | Benign | | 24189 | 4270 | 7.42 | 6.80 | | | | | -2.61E-02 | Neutral | | | -2.61E-02 | Neutral |
| 144 | p.Arg144Trp | | | 17236 | 3316 | 5.29 | 5.28 | | | | | -2.77E-01 | Neutral | | | -2.77E-01 | Neutral |
| 144 | p.Arg144Phe | | | 21521 | 4091 | 6.60 | 6.51 | | | | | -1.11E-01 | Neutral | | | -1.11E-01 | Neutral |
| 145 | p.Ile145Asn | | | 7612 | 1838 | 4.67 | 4.38 | | | | | -6.79E-01 | Neutral | | | -6.79E-01 | Neutral |
| 145 | p.Ile145Lys | | | 11244 | 3123 | 6.90 | 7.45 | | | | | -7.38E-01 | Neutral | | | -7.38E-01 | Neutral |
| 145 | p.Ile145Thr | | | 7751 | 1989 | 4.76 | 4.74 | | | | | -1.03E+00 | Neutral | | | -1.03E+00 | Neutral |
| 145 | p.Ile145Arg | | | 8952 | 2589 | 5.50 | 6.17 | | | | | -1.68E+00 | Neutral | | | -1.68E+00 | Neutral |
| 145 | p.Ile145Ser | | | 10994 | 2980 | 6.75 | 7.11 | | | | | -6.44E-01 | Neutral | | | -6.44E-01 | Neutral |
| 145 | p.Ile145Ile | Synonymous | | 6650 | 1628 | 4.08 | 3.88 | | | | | -1.06E+00 | Neutral | | | -1.06E+00 | Neutral |
| 145 | p.Ile145Met | | | 7048 | 2095 | 4.33 | 5.00 | | | | | -3.12E+00 | Neutral | | | -3.12E+00 | Neutral |
| 145 | p.Ile145His | | | 7200 | 1665 | 4.42 | 3.97 | | | | | -5.54E-01 | Neutral | | | -5.54E-01 | Neutral |
| 145 | p.Ile145Gln | | | 8916 | 2404 | 5.47 | 5.73 | | | | | -1.06E+00 | Neutral | | | -1.06E+00 | Neutral |
| 145 | p.Ile145Pro | | | 8974 | 2142 | 5.51 | 5.11 | | | | | -3.84E-01 | Neutral | | | -3.84E-01 | Neutral |
| 145 | p.Ile145Leu | | | 8437 | 2140 | 5.18 | 5.10 | | | | | -7.65E-01 | Neutral | | | -7.65E-01 | Neutral |
| 145 | p.Ile145Asp | | | 7325 | 1859 | 4.50 | 4.43 | | | | | -1.09E+00 | Neutral | | | -1.09E+00 | Neutral |
| 145 | p.Ile145Glu | | | 9464 | 2422 | 5.81 | 5.78 | | | | | -6.05E-01 | Neutral | | | -6.05E-01 | Neutral |
| 145 | p.Ile145Ala | | | 5278 | 1486 | 3.24 | 3.54 | | | | | -3.86E+00 | Neutral | | | -3.86E+00 | Neutral |
| 145 | p.Ile145Gly | | | 9634 | 2314 | 5.92 | 5.52 | | | | | -3.27E-01 | Neutral | | | -3.27E-01 | Neutral |
| 145 | p.Ile145Val | | | 6709 | 1848 | 4.12 | 4.41 | | | | | -2.24E+00 | Neutral | | | -2.24E+00 | Neutral |
| 145 | p.Ile145Tyr | | | 7802 | 1946 | 4.79 | 4.64 | | | | | -8.22E-01 | Neutral | | | -8.22E-01 | Neutral |
| 145 | p.Ile145Cys | | | 8714 | 1957 | 5.35 | 4.67 | | | | | -2.34E-01 | Neutral | | | -2.34E-01 | Neutral |
| 145 | p.Ile145Trp | | | 7610 | 1935 | 4.67 | 4.61 | | | | | -1.01E+00 | Neutral | | | -1.01E+00 | Neutral |
| 145 | p.Ile145Phe | | | 6546 | 1571 | 4.02 | 3.75 | | | | | -9.54E-01 | Neutral | | | -9.54E-01 | Neutral |
| 146 | p.Asp146Asn | | | 10106 | 2780 | 4.71 | 5.30 | | | | | -3.08E+00 | Neutral | | | -3.08E+00 | Neutral |
| 146 | p.Asp146Lys | | | 9313 | 1900 | 4.34 | 3.62 | | | | | -4.83E-01 | Neutral | | | -4.83E-01 | Neutral |
| 146 | p.Asp146Thr | | | 8674 | 2156 | 4.04 | 4.11 | | | | | -2.33E+00 | Neutral | | | -2.33E+00 | Neutral |
| 146 | p.Asp146Arg | | | 15511 | 3860 | 7.23 | 7.35 | | | | | -6.19E-01 | Neutral | | | -6.19E-01 | Neutral |
| 146 | p.Asp146Ser | | | 11188 | 2585 | 5.21 | 4.93 | | | | | -8.14E-01 | Neutral | | | -8.14E-01 | Neutral |
| 146 | p.Asp146Ile | | | 9107 | 1893 | 4.24 | 3.61 | | | | | -6.03E-01 | Neutral | | | -6.03E-01 | Neutral |
| 146 | p.Asp146Met | | | 12934 | 3264 | 6.03 | 6.22 | | | | | -1.10E+00 | Neutral | | | -1.10E+00 | Neutral |
| 146 | p.Asp146His | | | 10636 | 2801 | 4.96 | 5.34 | | | | | -2.20E+00 | Neutral | | | -2.20E+00 | Neutral |
| 146 | p.Asp146Gln | | | 10024 | 2577 | 4.67 | 4.91 | | | | | -2.14E+00 | Neutral | | | -2.14E+00 | Neutral |
| 146 | p.Asp146Pro | | | 11243 | 2945 | 5.24 | 5.61 | | | | | -1.91E+00 | Neutral | | | -1.91E+00 | Neutral |
| 146 | p.Asp146Leu | | | 8863 | 2148 | 4.13 | 4.09 | | | | | -1.92E+00 | Neutral | | | -1.92E+00 | Neutral |
| 146 | p.Asp146Asp | Synonymous | | 14629 | 3829 | 6.82 | 7.30 | | | | | -1.06E+00 | Neutral | | | -1.06E+00 | Neutral |
| 146 | p.Asp146Glu | | | 11304 | 2581 | 5.27 | 4.92 | | | | | -7.22E-01 | Neutral | | | -7.22E-01 | Neutral |
| 146 | p.Asp146Ala | | | 9378 | 2111 | 4.37 | 4.02 | | | | | -1.04E+00 | Neutral | | | -1.04E+00 | Neutral |

| | | | | | | | | | | | | | | | | |
|---|---|---|---|---|---|---|---|---|---|---|---|---|---|---|---|---|
| 146 | p.Asp146Gly | | 10422 | 2344 | 4.86 | 4.47 | | | | | -7.93E-01 | Neutral | | | -7.93E-01 | Neutral |
| 146 | p.Asp146Val | | 10670 | 2408 | 4.97 | 4.59 | | | | | -7.66E-01 | Neutral | | | -7.66E-01 | Neutral |
| 146 | p.Asp146Tyr | | 8216 | 2149 | 3.83 | 4.09 | | | | | -3.39E+00 | Neutral | | | -3.39E+00 | Neutral |
| 146 | p.Asp146Cys | | 8017 | 2125 | 3.74 | 4.05 | | | | | -3.78E+00 | Neutral | | | -3.78E+00 | Neutral |
| 146 | p.Asp146Trp | | 11676 | 2611 | 5.44 | 4.97 | | | | | -5.58E-01 | Neutral | | | -5.58E-01 | Neutral |
| 146 | p.Asp146Phe | | 12657 | 3420 | 5.90 | 6.52 | | | | | -1.81E+00 | Neutral | | | -1.81E+00 | Neutral |
| 147 | p.Ala147Asn | | 9204 | 2354 | 5.80 | 6.05 | | | | | -9.33E-01 | Neutral | | | -9.33E-01 | Neutral |
| 147 | p.Ala147Lys | | 5763 | 1279 | 3.63 | 3.29 | | | | | -1.03E+00 | Neutral | | | -1.03E+00 | Neutral |
| 147 | p.Ala147Thr | | 8653 | 2165 | 5.45 | 5.56 | | | | | -9.22E-01 | Neutral | | | -9.22E-01 | Neutral |
| 147 | p.Ala147Arg | | 9200 | 2032 | 5.79 | 5.22 | | | | | -2.64E-01 | Neutral | | | -2.64E-01 | Neutral |
| 147 | p.Ala147Ser | | 4918 | 1083 | 3.10 | 2.78 | | | | | -1.41E+00 | Neutral | | | -1.41E+00 | Neutral |
| 147 | p.Ala147Ile | | 12599 | 3435 | 7.93 | 8.82 | | | | | -6.82E-01 | Neutral | | | -6.82E-01 | Neutral |
| 147 | p.Ala147Met | | 8368 | 2192 | 5.27 | 5.63 | | | | | -1.38E+00 | Neutral | | | -1.38E+00 | Neutral |
| 147 | p.Ala147His | | 8991 | 1948 | 5.66 | 5.00 | | | | | -2.35E-01 | Neutral | | | -2.35E-01 | Neutral |
| 147 | p.Ala147Gln | | 8751 | 3013 | 5.51 | 7.74 | | | | | -5.59E+00 | Neutral | | | -5.59E+00 | Neutral |
| 147 | p.Ala147Pro | | 4007 | 861 | 2.52 | 2.21 | | | | | -1.86E+00 | Neutral | | | -1.86E+00 | Neutral |
| 147 | p.Ala147Leu | | 7838 | 2032 | 4.94 | 5.22 | | | | | -1.49E+00 | Neutral | | | -1.49E+00 | Neutral |
| 147 | p.Ala147Asp | | 7218 | 1972 | 4.55 | 5.07 | | | | | -2.43E+00 | Neutral | | | -2.43E+00 | Neutral |
| 147 | p.Ala147Glu | | 10401 | 2371 | 6.55 | 6.09 | | | | | -2.41E-01 | Neutral | | | -2.41E-01 | Neutral |
| 147 | p.Ala147Ala | Synonymous | 7509 | 1827 | 4.73 | 4.69 | | | | | -1.06E+00 | Neutral | | | -1.06E+00 | Neutral |
| 147 | p.Ala147Gly | | 7263 | 1725 | 4.57 | 4.43 | | | | | -9.65E-01 | Neutral | | | -9.65E-01 | Neutral |
| 147 | p.Ala147Val | | 6913 | 1751 | 4.35 | 4.50 | | | | | -1.68E+00 | Neutral | | | -1.68E+00 | Neutral |
| 147 | p.Ala147Tyr | | 7642 | 1659 | 4.81 | 4.26 | | | | | -4.02E-01 | Neutral | | | -4.02E-01 | Neutral |
| 147 | p.Ala147Cys | | 6487 | 1451 | 4.08 | 3.73 | | | | | -8.15E-01 | Neutral | | | -8.15E-01 | Neutral |
| 147 | p.Ala147Trp | | 7819 | 1773 | 4.92 | 4.55 | | | | | -5.51E-01 | Neutral | | | -5.51E-01 | Neutral |
| 147 | p.Ala147Phe | | 9259 | 2010 | 5.83 | 5.16 | | | | | -2.17E-01 | Neutral | | | -2.17E-01 | Neutral |
| 148 | p.Ala148Asn | | 5500 | 1127 | 5.24 | 5.45 | | | | | -6.43E+00 | Indeterminate | | | -6.43E+00 | Indeterminate |
| 148 | p.Ala148Lys | | 5191 | 1606 | 4.94 | 7.77 | | | | | -2.64E+01 | Indeterminate | | | -2.64E+01 | Indeterminate |
| 148 | p.Ala148Thr | Benign | 4478 | 636 | 4.27 | 3.08 | | | | | -1.32E+00 | Neutral | | | -1.32E+00 | Neutral |
| 148 | p.Ala148Arg | | 5505 | 1227 | 5.24 | 5.94 | | | | | -8.95E+00 | Indeterminate | | | -8.95E+00 | Indeterminate |
| 148 | p.Ala148Ser | | 5604 | 937 | 5.34 | 4.53 | | | | | -2.30E+00 | Neutral | | | -2.30E+00 | Neutral |
| 148 | p.Ala148Ile | | 5331 | 951 | 5.08 | 4.60 | | | | | -3.57E+00 | Neutral | | | -3.57E+00 | Neutral |
| 148 | p.Ala148Met | | 5054 | 1202 | 4.81 | 5.81 | | | | | -1.25E+01 | Indeterminate | | | -1.25E+01 | Indeterminate |
| 148 | p.Ala148His | | 4976 | 975 | 4.74 | 4.72 | | | | | -6.12E+00 | Indeterminate | | | -6.12E+00 | Indeterminate |
| 148 | p.Ala148Gln | | 4007 | 815 | 3.82 | 3.94 | | | | | -9.43E+00 | Indeterminate | | | -9.43E+00 | Indeterminate |
| 148 | p.Ala148Pro | | 5297 | 975 | 5.05 | 4.72 | | | | | -4.20E+00 | Neutral | | | -4.20E+00 | Neutral |
| 148 | p.Ala148Leu | | 5329 | 998 | 5.08 | 4.83 | | | | | -4.52E+00 | Neutral | | | -4.52E+00 | Neutral |
| 148 | p.Ala148Asp | | 7534 | 1486 | 7.18 | 7.19 | | | | | -3.31E+00 | Neutral | | | -3.31E+00 | Neutral |
| 148 | p.Ala148Glu | | 5527 | 1188 | 5.26 | 5.75 | | | | | -7.75E+00 | Indeterminate | | | -7.75E+00 | Indeterminate |
| 148 | p.Ala148Ala | Synonymous | 6364 | 985 | 6.06 | 4.76 | | | | | -1.06E+00 | Neutral | | | -1.06E+00 | Neutral |
| 148 | p.Ala148Gly | | 5784 | 901 | 5.51 | 4.36 | | | | | -1.38E+00 | Neutral | | | -1.38E+00 | Neutral |
| 148 | p.Ala148Val | | 5833 | 1397 | 5.56 | 6.76 | | | | | -1.08E+01 | Indeterminate | | | -1.08E+01 | Indeterminate |
| 148 | p.Ala148Tyr | | 5919 | 862 | 5.64 | 4.17 | | | | | -8.07E-01 | Neutral | | | -8.07E-01 | Neutral |
| 148 | p.Ala148Cys | | 4878 | 497 | 4.65 | 2.40 | | | | | -3.84E-02 | Neutral | | | -3.84E-02 | Neutral |
| 148 | p.Ala148Trp | | 1553 | 307 | 1.48 | 1.49 | | | | | -2.33E+01 | Indeterminate | | | -2.33E+01 | Indeterminate |
| 148 | p.Ala148Phe | | 5316 | 1600 | 5.06 | 7.74 | | | | | -2.40E+01 | Indeterminate | | | -2.40E+01 | Indeterminate |
| 149 | p.Glu149Asn | | 11526 | 2539 | 5.21 | 4.39 | | | | | -2.13E-01 | Neutral | | | -2.13E-01 | Neutral |
| 149 | p.Glu149Lys | | 12918 | 3636 | 5.84 | 6.28 | | | | | -1.24E+00 | Neutral | | | -1.24E+00 | Neutral |
| 149 | p.Glu149Thr | | 8983 | 2667 | 4.06 | 4.61 | | | | | -3.47E+00 | Neutral | | | -3.47E+00 | Neutral |
| 149 | p.Glu149Arg | | 11953 | 2775 | 5.40 | 4.79 | | | | | -3.18E-01 | Neutral | | | -3.18E-01 | Neutral |
| 149 | p.Glu149Ser | | 7632 | 2377 | 3.45 | 4.11 | | | | | -5.62E+00 | Neutral | | | -5.62E+00 | Neutral |
| 149 | p.Glu149Ile | | 9830 | 2957 | 4.44 | 5.11 | | | | | -3.18E+00 | Neutral | | | -3.18E+00 | Neutral |
| 149 | p.Glu149Met | | 8397 | 1815 | 3.79 | 3.14 | | | | | -4.83E-01 | Neutral | | | -4.83E-01 | Neutral |
| 149 | p.Glu149His | | 9745 | 2478 | 4.40 | 4.28 | | | | | -1.17E+00 | Neutral | | | -1.17E+00 | Neutral |
| 149 | p.Glu149Gln | | 14693 | 3524 | 6.64 | 6.09 | | | | | -2.22E-01 | Neutral | | | -2.22E-01 | Neutral |
| 149 | p.Glu149Pro | | 18087 | 4495 | 8.17 | 7.77 | | | | | -1.56E-01 | Neutral | | | -1.56E-01 | Neutral |
| 149 | p.Glu149Leu | | 19825 | 4976 | 8.96 | 8.60 | | | | | -1.24E-01 | Neutral | | | -1.24E-01 | Neutral |
| 149 | p.Glu149Asp | | 6538 | 1538 | 2.95 | 2.66 | | | | | -1.68E+00 | Neutral | | | -1.68E+00 | Neutral |
| 149 | p.Glu149Glu | Synonymous | 12702 | 3479 | 5.74 | 6.01 | | | | | -1.06E+00 | Neutral | | | -1.06E+00 | Neutral |
| 149 | p.Glu149Ala | | 9398 | 2578 | 4.25 | 4.45 | | | | | -2.07E+00 | Neutral | | | -2.07E+00 | Neutral |
| 149 | p.Glu149Gly | | 10131 | 2730 | 4.58 | 4.72 | | | | | -1.58E+00 | Neutral | | | -1.58E+00 | Neutral |
| 149 | p.Glu149Val | | 11244 | 3057 | 5.08 | 5.28 | | | | | -1.34E+00 | Neutral | | | -1.34E+00 | Neutral |
| 149 | p.Glu149Tyr | | 13264 | 3705 | 5.99 | 6.40 | | | | | -1.10E+00 | Neutral | | | -1.10E+00 | Neutral |
| 149 | p.Glu149Cys | | 7128 | 1678 | 3.22 | 2.90 | | | | | -1.40E+00 | Neutral | | | -1.40E+00 | Neutral |
| 149 | p.Glu149Trp | | 10289 | 2823 | 4.65 | 4.88 | | | | | -1.72E+00 | Neutral | | | -1.72E+00 | Neutral |
| 149 | p.Glu149Phe | | 7043 | 2050 | 3.18 | 3.54 | | | | | -4.67E+00 | Neutral | | | -4.67E+00 | Neutral |
| 150 | p.Gly150Asn | | 7889 | 2318 | 5.48 | 5.15 | | | | | -2.17E-01 | Neutral | | | -2.17E-01 | Neutral |
| 150 | p.Gly150Lys | | 5103 | 1727 | 3.55 | 3.84 | | | | | -2.08E+00 | Neutral | | | -2.08E+00 | Neutral |
| 150 | p.Gly150Thr | | 5460 | 1724 | 3.79 | 3.83 | | | | | -1.15E+00 | Neutral | | | -1.15E+00 | Neutral |
| 150 | p.Gly150Arg | | 10512 | 3556 | 7.31 | 7.91 | | | | | -3.40E-01 | Neutral | | | -3.40E-01 | Neutral |
| 150 | p.Gly150Ser | | 5701 | 1862 | 3.96 | 4.14 | | | | | -1.31E+00 | Neutral | | | -1.31E+00 | Neutral |
| 150 | p.Gly150Ile | | 9376 | 2842 | 6.52 | 6.32 | | | | | -1.65E-01 | Neutral | | | -1.65E-01 | Neutral |
| 150 | p.Gly150Met | | 8873 | 3128 | 6.17 | 6.95 | | | | | -7.85E-01 | Neutral | | | -7.85E-01 | Neutral |
| 150 | p.Gly150His | | 6463 | 2210 | 4.49 | 4.91 | | | | | -1.35E+00 | Neutral | | | -1.35E+00 | Neutral |
| 150 | p.Gly150Gln | | 4578 | 1412 | 3.18 | 3.14 | | | | | -1.46E+00 | Neutral | | | -1.46E+00 | Neutral |
| 150 | p.Gly150Pro | | 7208 | 2368 | 5.01 | 5.26 | | | | | -7.75E-01 | Neutral | | | -7.75E-01 | Neutral |
| 150 | p.Gly150Leu | | 6458 | 1932 | 4.49 | 4.30 | | | | | -4.84E-01 | Neutral | | | -4.84E-01 | Neutral |
| 150 | p.Gly150Asp | | 4629 | 1476 | 3.22 | 3.28 | | | | | -1.76E+00 | Neutral | | | -1.76E+00 | Neutral |
| 150 | p.Gly150Glu | | 7971 | 2314 | 5.54 | 5.14 | | | | | -1.85E-01 | Neutral | | | -1.85E-01 | Neutral |
| 150 | p.Gly150Ala | | 8197 | 2338 | 5.70 | 5.20 | | | | | -1.38E-01 | Neutral | | | -1.38E-01 | Neutral |
| 150 | p.Gly150Gly | Synonymous | 6435 | 2123 | 4.47 | 4.72 | | | | | -1.06E+00 | Neutral | | | -1.06E+00 | Neutral |
| 150 | p.Gly150Val | | 6648 | 2121 | 4.62 | 4.72 | | | | | -7.61E-01 | Neutral | | | -7.61E-01 | Neutral |
| 150 | p.Gly150Tyr | | 9875 | 2976 | 6.86 | 6.62 | | | | | -1.28E-01 | Neutral | | | -1.28E-01 | Neutral |
| 150 | p.Gly150Cys | | 9027 | 2296 | 6.27 | 5.10 | | | | | -2.12E-02 | Neutral | | | -2.12E-02 | Neutral |
| 150 | p.Gly150Trp | | 6653 | 2013 | 4.62 | 4.48 | | | | | -4.90E-01 | Neutral | | | -4.90E-01 | Neutral |
| 150 | p.Gly150Phe | | 6845 | 2245 | 4.76 | 4.99 | | | | | -8.74E-01 | Neutral | | | -8.74E-01 | Neutral |
| 151 | p.Pro151Asn | | 7141 | 5866 | 5.72 | 6.09 | | | | | -7.22E-01 | Neutral | | | -7.22E-01 | Neutral |
| 151 | p.Pro151Lys | | 10023 | 7533 | 8.03 | 7.83 | | | | | -1.02E-01 | Neutral | | | -1.02E-01 | Neutral |
| 151 | p.Pro151Thr | | 4705 | 3885 | 3.77 | 4.04 | | | | | -1.96E+00 | Neutral | | | -1.96E+00 | Neutral |
| 151 | p.Pro151Arg | | 4020 | 2684 | 3.22 | 2.79 | | | | | -6.32E-01 | Neutral | | | -6.32E-01 | Neutral |
| 151 | p.Pro151Ser | | 1092 | 898 | 0.87 | 0.93 | | | | | -1.55E+01 | Indeterminate | | | -1.55E+01 | Indeterminate |
| 151 | p.Pro151Ile | | 7374 | 5843 | 5.91 | 6.07 | | | | | -4.90E-01 | Neutral | | | -4.90E-01 | Neutral |
| 151 | p.Pro151Met | | 10441 | 7670 | 8.36 | 7.97 | | | | | -6.54E-02 | Neutral | | | -6.54E-02 | Neutral |
| 151 | p.Pro151His | | 7271 | 6167 | 5.82 | 6.41 | | | | | -8.83E-01 | Neutral | | | -8.83E-01 | Neutral |
| 151 | p.Pro151Gln | | 7412 | 5338 | 5.94 | 5.55 | | | | | -1.93E-01 | Neutral | | | -1.93E-01 | Neutral |
| 151 | p.Pro151Pro | Synonymous | 5643 | 4511 | 4.52 | 4.69 | | | | | -1.06E+00 | Neutral | | | -1.06E+00 | Neutral |
| 151 | p.Pro151Leu | | 3192 | 2581 | 2.56 | 2.68 | | | | | -3.56E+00 | Neutral | | | -3.56E+00 | Neutral |
| 151 | p.Pro151Asp | | 3871 | 3149 | 3.10 | 3.27 | | | | | -2.62E+00 | Neutral | | | -2.62E+00 | Neutral |
| 151 | p.Pro151Glu | | 7344 | 6204 | 5.88 | 6.45 | | | | | -8.35E-01 | Neutral | | | -8.35E-01 | Neutral |
| 151 | p.Pro151Ala | | 5061 | 4390 | 4.05 | 4.56 | | | | | -2.27E+00 | Neutral | | | -2.27E+00 | Neutral |
| 151 | p.Pro151Gly | | 2399 | 2158 | 1.92 | 2.24 | | | | | -8.60E+00 | Indeterminate | | | -8.60E+00 | Indeterminate |
| 151 | p.Pro151Val | | 8770 | 6632 | 7.02 | 6.89 | | | | | -1.79E-01 | Neutral | | | -1.79E-01 | Neutral |
| 151 | p.Pro151Tyr | | 10716 | 7570 | 8.58 | 7.87 | | | | | -3.50E-02 | Neutral | | | -3.50E-02 | Neutral |
| 151 | p.Pro151Cys | | 5307 | 3759 | 4.25 | 3.91 | | | | | -4.77E-01 | Neutral | | | -4.77E-01 | Neutral |
| 151 | p.Pro151Trp | | 6639 | 4694 | 5.32 | 4.88 | | | | | -2.33E-01 | Neutral | | | -2.33E-01 | Neutral |
| 151 | p.Pro151Phe | | 6424 | 4716 | 5.15 | 4.90 | | | | | -3.72E-01 | Neutral | | | -3.72E-01 | Neutral |
| 152 | p.Ser152Asn | | 997 | 2607 | 2.80 | 2.66 | | | | | -3.22E+00 | Neutral | | | -3.22E+00 | Neutral |
| 152 | p.Ser152Lys | | 1636 | 4727 | 4.59 | 4.82 | | | | | -2.25E+00 | Neutral | | | -2.25E+00 | Neutral |
| 152 | p.Ser152Thr | | 1526 | 4399 | 4.28 | 4.48 | | | | | -2.54E+00 | Neutral | | | -2.54E+00 | Neutral |
| 152 | p.Ser152Arg | | 1486 | 3962 | 4.17 | 4.04 | | | | | -1.68E+00 | Neutral | | | -1.68E+00 | Neutral |
| 152 | p.Ser152Ser | Synonymous | 2605 | 7820 | 7.31 | 7.97 | | | | | -1.06E+00 | Neutral | | | -1.06E+00 | Neutral |
| 152 | p.Ser152Ile | | 1530 | 3475 | 4.29 | 3.54 | | | | | -4.71E-01 | Neutral | | | -4.71E-01 | Neutral |
| 152 | p.Ser152Met | | 1280 | 3711 | 3.59 | 3.78 | | | | | -3.57E+00 | Neutral | | | -3.57E+00 | Neutral |
| 152 | p.Ser152His | | 2226 | 5927 | 6.25 | 6.04 | | | | | -6.31E-01 | Neutral | | | -6.31E-01 | Neutral |
| 152 | p.Ser152Gln | | 2013 | 4802 | 5.65 | 4.90 | | | | | -3.20E-01 | Neutral | | | -3.20E-01 | Neutral |
| 152 | p.Ser152Pro | | 1571 | 4606 | 4.41 | 4.70 | | | | | -2.64E+00 | Neutral | | | -2.64E+00 | Neutral |
| 152 | p.Ser152Leu | | 1380 | 3970 | 3.87 | 4.05 | | | | | -3.01E+00 | Neutral | | | -3.01E+00 | Neutral |
| 152 | p.Ser152Asp | | 2675 | 7145 | 7.51 | 7.28 | | | | | -3.81E-01 | Neutral | | | -3.81E-01 | Neutral |
| 152 | p.Ser152Glu | | 1732 | 4745 | 4.86 | 4.84 | | | | | -1.44E+00 | Neutral | | | -1.44E+00 | Neutral |
| 152 | p.Ser152Ala | | 1609 | 5022 | 4.52 | 5.12 | | | | | -3.53E+00 | Neutral | | | -3.53E+00 | Neutral |
| 152 | p.Ser152Gly | | 1064 | 3104 | 2.99 | 3.16 | | | | | -4.94E+00 | Neutral | | | -4.94E+00 | Neutral |
| 152 | p.Ser152Val | | 2808 | 8240 | 7.88 | 8.40 | | | | | -7.41E-01 | Neutral | | | -7.41E-01 | Neutral |
| 152 | p.Ser152Tyr | | 1351 | 4459 | 3.79 | 4.55 | | | | | -6.00E+00 | Indeterminate | | | -6.00E+00 | Indeterminate |
| 152 | p.Ser152Cys | | 2129 | 4765 | 5.98 | 4.86 | | | | | -1.37E-01 | Neutral | | | -1.37E-01 | Neutral |
| 152 | p.Ser152Trp | | 2066 | 5505 | 5.80 | 5.61 | | | | | -7.73E-01 | Neutral | | | -7.73E-01 | Neutral |
| 152 | p.Ser152Phe | | 1942 | 5105 | 5.45 | 5.20 | | | | | -8.17E-01 | Neutral | | | -8.17E-01 | Neutral |
| 153 | p.Asp153Asn | | 6218 | 2873 | 4.13 | 4.91 | 8856 | 2743 | 4.08 | 3.33 | -5.02E+00 | Neutral | -2.25E-01 | Neutral | -3.03E+00 | Neutral |
| 153 | p.Asp153Lys | | 8629 | 2960 | 5.73 | 5.06 | 12644 | 6300 | 5.83 | 7.65 | -3.51E-01 | Neutral | -3.16E+00 | Neutral | -1.73E+00 | Neutral |
| 153 | p.Asp153Thr | | 7168 | 3671 | 4.76 | 6.28 | 10937 | 4515 | 5.04 | 5.48 | -6.40E+00 | Indeterminate | -1.38E+00 | Neutral | -5.10E+00 | Neutral |
| 153 | p.Asp153Arg | | 7408 | 3213 | 4.92 | 5.49 | 11470 | 5904 | 5.29 | 7.17 | -2.74E+00 | Neutral | -4.37E+00 | Neutral | -4.54E+00 | Neutral |
| 153 | p.Asp153Ser | | 8767 | 2503 | 5.82 | 4.28 | 12287 | 3331 | 5.67 | 4.05 | -4.19E-02 | Neutral | -1.01E-02 | Neutral | -9.15E-04 | Neutral |
| 153 | p.Asp153Ile | | 7764 | 2451 | 5.16 | 4.19 | 11706 | 5110 | 5.40 | 6.21 | -2.19E-01 | Neutral | -1.71E+00 | Neutral | -7.04E-01 | Neutral |
| 153 | p.Asp153Met | | 8272 | 3776 | 5.49 | 6.46 | 12502 | 4304 | 5.77 | 5.23 | -2.96E+00 | Neutral | -2.06E-01 | Neutral | -1.49E+00 | Neutral |
| 153 | p.Asp153His | | 7310 | 3413 | 4.86 | 5.84 | 10786 | 3711 | 4.97 | 4.51 | -4.10E+00 | Neutral | -3.33E-01 | Neutral | -2.40E+00 | Neutral |
| 153 | p.Asp153Gln | | 8065 | 3909 | 5.36 | 6.68 | 11217 | 3993 | 5.17 | 4.85 | -4.19E+00 | Neutral | -4.04E-01 | Neutral | -2.53E+00 | Neutral |
| 153 | p.Asp153Pro | | 5129 | 1214 | 3.41 | 2.08 | 7233 | 1428 | 3.34 | 1.73 | -3.69E-02 | Neutral | -9.05E-04 | Neutral | -4.86E-04 | Neutral |
| 153 | p.Asp153Leu | | 8094 | 3783 | 5.38 | 6.47 | 11872 | 4179 | 5.48 | 5.08 | -3.48E+00 | Neutral | -3.05E-01 | Neutral | -1.93E+00 | Neutral |
| 153 | p.Asp153Asp | Synonymous | 6009 | 2091 | 3.99 | 3.58 | 9161 | 3435 | 4.23 | 4.17 | -1.06E+00 | Neutral | -1.06E+00 | Neutral | -8.18E-01 | Neutral |
| 153 | p.Asp153Glu | | 7756 | 3190 | 5.15 | 5.46 | 10546 | 3242 | 4.86 | 3.94 | -1.84E+00 | Neutral | -1.10E-01 | Neutral | -7.16E-01 | Neutral |

| | | | | | | | | | | | | | | | | |
|---|---|---|---|---|---|---|---|---|---|---|---|---|---|---|---|---|
| 153 | p.Asp153Ala | | 8429 | 1977 | 5.60 | 3.38 | 11857 | 4363 | 5.47 | 5.30 | -2.49E-03 | Neutral | -4.61E-01 | Neutral | -6.16E-02 | Neutral |
| 153 | p.Asp153Gly | | 7582 | 1910 | 5.04 | 3.27 | 11395 | 2748 | 5.26 | 3.34 | -1.41E-02 | Neutral | -2.20E-03 | Neutral | -9.11E-05 | Neutral |
| 153 | p.Asp153Val | | 6468 | 2574 | 4.30 | 4.40 | 8833 | 3958 | 4.07 | 4.81 | -2.17E+00 | Neutral | -3.38E+00 | Neutral | -3.27E+00 | Neutral |
| 153 | p.Asp153Tyr | | 6325 | 2136 | 4.20 | 3.65 | 8714 | 4339 | 4.02 | 5.27 | -7.45E-01 | Neutral | -5.68E+00 | Neutral | -3.98E+00 | Neutral |
| 153 | p.Asp153Cys | | 8817 | 4328 | 5.86 | 7.40 | 12054 | 4932 | 5.56 | 5.99 | -3.86E+00 | Neutral | -1.03E+00 | Neutral | -2.76E+00 | Neutral |
| 153 | p.Asp153Trp | | 8197 | 3569 | 5.44 | 6.10 | 11348 | 5242 | 5.23 | 6.37 | -2.32E+00 | Neutral | -2.54E+00 | Neutral | -2.73E+00 | Neutral |
| 153 | p.Asp153Phe | | 8150 | 2936 | 5.41 | 5.02 | 11400 | 4560 | 5.26 | 5.54 | -6.37E-01 | Neutral | -1.00E+00 | Neutral | -5.44E-01 | Neutral |
| 154 | p.Ile154Asn | | 2296 | 1516 | 4.81 | 5.02 | | | | | -4.12E+00 | Neutral | | | -4.12E+00 | Neutral |
| 154 | p.Ile154Lys | | 2910 | 1596 | 6.10 | 5.29 | | | | | -8.25E-01 | Neutral | | | -8.25E-01 | Neutral |
| 154 | p.Ile154Thr | | 2308 | 1518 | 4.84 | 5.03 | | | | | -4.01E+00 | Neutral | | | -4.01E+00 | Neutral |
| 154 | p.Ile154Arg | | 3084 | 1930 | 6.46 | 6.40 | | | | | -1.80E+00 | Neutral | | | -1.80E+00 | Neutral |
| 154 | p.Ile154Ser | | 2108 | 1096 | 4.42 | 3.63 | | | | | -1.22E+00 | Neutral | | | -1.22E+00 | Neutral |
| 154 | p.Ile154Ile | Synonymous | 3477 | 2093 | 7.29 | 6.94 | | | | | -1.06E+00 | Neutral | | | -1.06E+00 | Neutral |
| 154 | p.Ile154Met | | 2376 | 1476 | 4.98 | 4.89 | | | | | -2.85E+00 | Neutral | | | -2.85E+00 | Neutral |
| 154 | p.Ile154His | | 1658 | 1126 | 3.47 | 3.73 | | | | | -7.49E+00 | Indeterminate | | | -7.49E+00 | Indeterminate |
| 154 | p.Ile154Gln | | 2231 | 1345 | 4.67 | 4.46 | | | | | -2.71E+00 | Neutral | | | -2.71E+00 | Neutral |
| 154 | p.Ile154Pro | | 1218 | 831 | 2.55 | 2.75 | | | | | -1.12E+01 | Indeterminate | | | -1.12E+01 | Indeterminate |
| 154 | p.Ile154Leu | | 2537 | 1415 | 5.32 | 4.69 | | | | | -1.30E+00 | Neutral | | | -1.30E+00 | Neutral |
| 154 | p.Ile154Asp | | 2677 | 1599 | 5.61 | 5.30 | | | | | -1.80E+00 | Neutral | | | -1.80E+00 | Neutral |
| 154 | p.Ile154Glu | | 3009 | 1926 | 6.30 | 6.38 | | | | | -2.16E+00 | Neutral | | | -2.16E+00 | Neutral |
| 154 | p.Ile154Ala | | 1684 | 1283 | 3.53 | 4.25 | | | | | -1.13E+01 | Indeterminate | | | -1.13E+01 | Indeterminate |
| 154 | p.Ile154Gly | | 2477 | 1294 | 5.19 | 4.29 | | | | | -8.56E-01 | Neutral | | | -8.56E-01 | Neutral |
| 154 | p.Ile154Val | | 2522 | 1819 | 5.28 | 6.03 | | | | | -5.37E+00 | Neutral | | | -5.37E+00 | Neutral |
| 154 | p.Ile154Tyr | | 1981 | 1370 | 4.15 | 4.54 | | | | | -6.33E+00 | Indeterminate | | | -6.33E+00 | Indeterminate |
| 154 | p.Ile154Cys | | 1607 | 1097 | 3.37 | 3.64 | | | | | -7.96E+00 | Indeterminate | | | -7.96E+00 | Indeterminate |
| 154 | p.Ile154Trp | | 2865 | 2125 | 6.00 | 7.04 | | | | | -5.04E+00 | Neutral | | | -5.04E+00 | Neutral |
| 154 | p.Ile154Phe | | 2701 | 1718 | 5.66 | 5.69 | | | | | -2.57E+00 | Neutral | | | -2.57E+00 | Neutral |
| 155 | p.Pro155Asn | | 7516 | 1212 | 4.52 | 4.44 | | | | | -1.38E+00 | Neutral | | | -1.38E+00 | Neutral |
| 155 | p.Pro155Lys | | 6712 | 1306 | 4.04 | 4.79 | | | | | -4.81E+00 | Neutral | | | -4.81E+00 | Neutral |
| 155 | p.Pro155Thr | | 9334 | 283 | 5.62 | 1.04 | | | | | 0.00E+00 | Neutral | | | 0.00E+00 | Neutral |
| 155 | p.Pro155Arg | | 9747 | 1439 | 5.87 | 5.28 | | | | | -3.44E-01 | Neutral | | | -3.44E-01 | Neutral |
| 155 | p.Pro155Ser | | 8082 | 2822 | 4.86 | 10.35 | | | | | -2.83E+01 | Indeterminate | | | -2.82E+01 | Indeterminate |
| 155 | p.Pro155Ile | | 7620 | 1707 | 4.59 | 6.26 | | | | | -7.35E+00 | Indeterminate | | | -7.35E+00 | Indeterminate |
| 155 | p.Pro155Met | | 8315 | 793 | 5.00 | 2.91 | | | | | -1.94E-03 | Neutral | | | -1.94E-03 | Neutral |
| 155 | p.Pro155His | | 9630 | 2577 | 5.80 | 9.45 | | | | | -1.07E+01 | Indeterminate | | | -1.07E+01 | Indeterminate |
| 155 | p.Pro155Gln | | 7547 | 572 | 4.54 | 2.10 | | | | | -3.92E-05 | Neutral | | | -3.92E-05 | Neutral |
| 155 | p.Pro155Pro | Synonymous | 8846 | 1450 | 5.32 | 5.32 | | | | | -1.06E+00 | Neutral | | | -1.06E+00 | Neutral |
| 155 | p.Pro155Leu | | 12220 | 1003 | 7.35 | 3.68 | | | | | -6.49E-06 | Neutral | | | -6.49E-06 | Neutral |
| 155 | p.Pro155Asp | | 9525 | 3729 | 5.73 | 13.67 | | | | | -3.28E+01 | Indeterminate | | | -3.20E+01 | Indeterminate |
| 155 | p.Pro155Glu | | 8909 | 797 | 5.36 | 2.92 | | | | | -3.84E-04 | Neutral | | | -3.84E-04 | Neutral |
| 155 | p.Pro155Ala | | 4663 | 140 | 2.81 | 0.51 | | | | | -3.27E-14 | Neutral | | | 0.00E+00 | Neutral |
| 155 | p.Pro155Gly | | 8335 | 970 | 5.02 | 3.56 | | | | | -4.22E-02 | Neutral | | | -4.22E-02 | Neutral |
| 155 | p.Pro155Val | | 8856 | 2882 | 5.33 | 10.57 | | | | | -2.16E+01 | Indeterminate | | | -2.16E+01 | Indeterminate |
| 155 | p.Pro155Tyr | | 9258 | 1066 | 5.57 | 3.91 | | | | | -2.25E-02 | Neutral | | | -2.25E-02 | Neutral |
| 155 | p.Pro155Cys | | 7343 | 471 | 4.42 | 1.73 | | | | | -1.03E-06 | Neutral | | | -1.03E-06 | Neutral |
| 155 | p.Pro155Trp | | 7079 | 1840 | 4.26 | 6.75 | | | | | -1.37E+01 | Indeterminate | | | -1.37E+01 | Indeterminate |
| 155 | p.Pro155Phe | | 6611 | 211 | 3.98 | 0.77 | | | | | -5.77E-15 | Neutral | | | 0.00E+00 | Neutral |
| 156 | p.Asp156Asn | | 5066 | 1583 | 4.64 | 4.64 | | | | | -3.19E+00 | Neutral | | | -3.19E+00 | Neutral |
| 156 | p.Asp156Lys | | 2453 | 728 | 2.25 | 2.13 | | | | | -7.54E+00 | Indeterminate | | | -7.54E+00 | Indeterminate |
| 156 | p.Asp156Thr | | 7734 | 2477 | 7.09 | 7.26 | | | | | -1.63E+00 | Neutral | | | -1.63E+00 | Neutral |
| 156 | p.Asp156Arg | | 3732 | 1122 | 3.42 | 3.29 | | | | | -4.36E+00 | Neutral | | | -4.36E+00 | Neutral |
| 156 | p.Asp156Ser | | 8975 | 2716 | 8.23 | 7.96 | | | | | -7.69E-01 | Neutral | | | -7.69E-01 | Neutral |
| 156 | p.Asp156Ile | | 4671 | 1355 | 4.28 | 3.97 | | | | | -2.46E+00 | Neutral | | | -2.46E+00 | Neutral |
| 156 | p.Asp156Met | | 5697 | 1689 | 5.22 | 4.95 | | | | | -1.90E+00 | Neutral | | | -1.90E+00 | Neutral |
| 156 | p.Asp156His | | 3364 | 981 | 3.08 | 2.87 | | | | | -4.44E+00 | Neutral | | | -4.44E+00 | Neutral |
| 156 | p.Asp156Gln | | 4870 | 1727 | 4.46 | 5.06 | | | | | -6.15E+00 | Indeterminate | | | -6.15E+00 | Indeterminate |
| 156 | p.Asp156Pro | | 7585 | 2236 | 6.95 | 6.55 | | | | | -9.64E-01 | Neutral | | | -9.64E-01 | Neutral |
| 156 | p.Asp156Leu | | 5547 | 1799 | 5.08 | 5.27 | | | | | -3.31E+00 | Neutral | | | -3.31E+00 | Neutral |
| 156 | p.Asp156Asp | Synonymous | 7593 | 2269 | 6.96 | 6.65 | | | | | -1.06E+00 | Neutral | | | -1.06E+00 | Neutral |
| 156 | p.Asp156Glu | | 6739 | 2108 | 6.18 | 6.17 | | | | | -1.87E+00 | Neutral | | | -1.87E+00 | Neutral |
| 156 | p.Asp156Ala | | 4389 | 1665 | 4.02 | 4.88 | | | | | -9.22E+00 | Indeterminate | | | -9.22E+00 | Indeterminate |
| 156 | p.Asp156Gly | | 5406 | 1778 | 4.96 | 5.21 | | | | | -3.71E+00 | Neutral | | | -3.71E+00 | Neutral |
| 156 | p.Asp156Val | | 5781 | 1953 | 5.30 | 5.72 | | | | | -3.79E+00 | Neutral | | | -3.79E+00 | Neutral |
| 156 | p.Asp156Tyr | | 4106 | 1384 | 3.76 | 4.05 | | | | | -6.31E+00 | Indeterminate | | | -6.31E+00 | Indeterminate |
| 156 | p.Asp156Cys | | 3646 | 1198 | 3.34 | 3.51 | | | | | -6.69E+00 | Indeterminate | | | -6.69E+00 | Indeterminate |
| 156 | p.Asp156Trp | | 4783 | 1291 | 4.38 | 3.78 | | | | | -1.51E+00 | Neutral | | | -1.51E+00 | Neutral |
| 156 | p.Asp156Phe | | 6952 | 2080 | 6.37 | 6.09 | | | | | -1.31E+00 | Neutral | | | -1.31E+00 | Neutral |
