## Appendix 1-table 5 for "Functional characterization of all *CDKN2A* missense variants and comparison to in silico models of pathogenicity"

**Appendix 1-table 5. Day of confluency by experiment and residue.**

| Residue | Exp_1 | Exp_2 |
|---|---|---|
| 1 | 32 | |
| 2 | 31 | |
| 3 | 33 | |
| 4 | 31 | |
| 5 | 29 | |
| 6 | 29 | |
| 7 | 33 | |
| 8 | 29 | |
| 9 | 25 | |
| 10 | 29 | |
| 11 | 25 | |
| 12 | 34 | |
| 13 | 33 | |
| 14 | 32 | |
| 15 | 29 | 30 |
| 16 | 21 | |
| 17 | 23 | |
| 18 | 29 | |
| 19 | 30 | |
| 20 | 23 | 17 |
| 21 | 17 | |
| 22 | 29 | |
| 23 | 23 | |
| 24 | 27 | 25 |
| 25 | 30 | |
| 26 | 33 | |
| 27 | 31 | 30 |
| 28 | 22 | |
| 29 | 29 | |
| 30 | 27 | |
| 31 | 29 | |
| 32 | 16 | 20 |
| 33 | 33 | |
| 34 | 34 | 30 |
| 35 | 25 | 25 |
| 36 | 28 | |
| 37 | 30 | |
| 38 | 22 | |
| 39 | 24 | |
| 40 | 28 | |
| 41 | 30 | |
| 42 | 20 | |
| 43 | 30 | |
| 44 | 32 | |
| 45 | 31 | 30 |
| 46 | 27 | |
| 47 | 31 | |
| 48 | 16 | |
| 49 | 23 | |

| | | |
|---|---|---|
| 50 | 22 | |
| 51 | 23 | |
| 52 | 30 | |
| 53 | 25 | |
| 54 | 33 | |
| 55 | 21 | |
| 56 | 27 | |
| 57 | 30 | 31 |
| 58 | 27 | |
| 59 | 21 | |
| 60 | 21 | |
| 61 | 28 | |
| 62 | 33 | 33 |
| 63 | 20 | |
| 64 | 28 | |
| 65 | 23 | |
| 66 | 23 | |
| 67 | 27 | |
| 68 | 20 | |
| 69 | 31 | |
| 70 | 26 | |
| 71 | 20 | |
| 72 | 29 | 28 |
| 73 | 27 | 29 |
| 74 | 21 | 23 |
| 75 | 27 | 28 |
| 76 | 27 | |
| 77 | 28 | |
| 78 | 30 | |
| 79 | 29 | |
| 80 | 27 | |
| 81 | 20 | |
| 82 | 23 | |
| 83 | 20 | |
| 84 | 23 | |
| 85 | 30 | |
| 86 | 20 | |
| 87 | 22 | |
| 88 | 29 | |
| 89 | 19 | |
| 90 | 28 | |
| 91 | 28 | |
| 92 | 30 | |
| 93 | 25 | |
| 94 | 21 | |
| 95 | 34 | |
| 96 | 34 | 30 |
| 97 | 22 | |
| 98 | 30 | |
| 99 | 27 | |
| 100 | 23 | |
| 101 | 23 | |
| 102 | 20 | |
| 103 | 29 | |

| | | |
|---|---|---|
| 104 | 28 | |
| 105 | 27 | |
| 106 | 30 | 28 |
| 107 | 33 | 30 |
| 108 | 18 | |
| 109 | 28 | 30 |
| 110 | 38 | |
| 111 | 24 | |
| 112 | 27 | |
| 113 | 38 | 35 |
| 114 | 17 | |
| 115 | 33 | |
| 116 | 32 | |
| 117 | 30 | |
| 118 | 35 | |
| 119 | 38 | |
| 120 | 29 | 30 |
| 121 | 28 | |
| 122 | 27 | 27 |
| 123 | 29 | 27 |
| 124 | 30 | 30 |
| 125 | 32 | 30 |
| 126 | 23 | |
| 127 | 30 | |
| 128 | 29 | |
| 129 | 30 | |
| 130 | 25 | |
| 131 | 31 | |
| 132 | 30 | |
| 133 | 38 | |
| 134 | 32 | 28 |
| 135 | 38 | |
| 136 | 40 | 35 |
| 137 | 40 | |
| 138 | 40 | |
| 139 | 32 | 30 |
| 140 | 35 | |
| 141 | 33 | |
| 142 | 30 | |
| 143 | 32 | |
| 144 | 31 | |
| 145 | 30 | |
| 146 | 35 | |
| 147 | 29 | |
| 148 | 31 | |
| 149 | 30 | |
| 150 | 31 | |
| 151 | 32 | |
| 152 | 29 | |
| 153 | 31 | 28 |
| 154 | 30 | |
| 155 | 32 | |
| 156 | 30 | |
