## Appendix 1-table 6 for "Functional characterization of all *CDKN2A* missense variants and comparison to in silico models of pathogenicity"

**Appendix 1-table 6. Normalized fold change for all possible *CDKN2A* missense and synonymous variants.**

| Residue | Variant | Normalized fold change_1 | Log2 normalized fold change_1 | Classification_1 | Normalized fold change_2 | Log2 normalized fold change_2 | Classification_2 | Mean of normalized fold change | Mean of log normalized fold change | Classification_merged |
|---|---|---|---|---|---|---|---|---|---|---|
| 1 | p.Met1Asn | 0.92 | -0.12 | Neutral | | | | 0.92 | -0.12 | Neutral |
| 1 | p.Met1Lys | 0.82 | -0.29 | Neutral | | | | 0.82 | -0.29 | Neutral |
| 1 | p.Met1Thr | 0.94 | -0.09 | Neutral | | | | 0.94 | -0.09 | Neutral |
| 1 | p.Met1Arg | 0.79 | -0.33 | Neutral | | | | 0.79 | -0.33 | Neutral |
| 1 | p.Met1Ser | 0.92 | -0.12 | Neutral | | | | 0.92 | -0.12 | Neutral |
| 1 | p.Met1Ile | 0.92 | -0.12 | Neutral | | | | 0.92 | -0.12 | Neutral |
| 1 | p.Met1Met | 1.00 | 0.00 | Neutral | | | | 1.00 | 0.00 | Neutral |
| 1 | p.Met1His | 1.01 | 0.02 | Neutral | | | | 1.01 | 0.02 | Neutral |
| 1 | p.Met1Gln | 0.96 | -0.05 | Neutral | | | | 0.96 | -0.05 | Neutral |
| 1 | p.Met1Pro | 0.87 | -0.20 | Neutral | | | | 0.87 | -0.20 | Neutral |
| 1 | p.Met1Leu | 0.96 | -0.06 | Neutral | | | | 0.96 | -0.06 | Neutral |
| 1 | p.Met1Asp | 0.86 | -0.22 | Neutral | | | | 0.86 | -0.22 | Neutral |
| 1 | p.Met1Glu | 0.97 | -0.04 | Neutral | | | | 0.97 | -0.04 | Neutral |
| 1 | p.Met1Ala | 0.90 | -0.16 | Neutral | | | | 0.90 | -0.16 | Neutral |
| 1 | p.Met1Gly | 0.94 | -0.09 | Neutral | | | | 0.94 | -0.09 | Neutral |
| 1 | p.Met1Val | 1.61 | 0.69 | Indeterminate | | | | 1.61 | 0.69 | Indeterminate |
| 1 | p.Met1Tyr | 0.92 | -0.13 | Neutral | | | | 0.92 | -0.13 | Neutral |
| 1 | p.Met1Cys | 0.90 | -0.15 | Neutral | | | | 0.90 | -0.15 | Neutral |
| 1 | p.Met1Trp | 0.92 | -0.12 | Neutral | | | | 0.92 | -0.12 | Neutral |
| 1 | p.Met1Phe | 0.97 | -0.04 | Neutral | | | | 0.97 | -0.04 | Neutral |
| 2 | p.Glu2Asn | 0.20 | -2.29 | Neutral | | | | 0.20 | -2.29 | Neutral |
| 2 | p.Glu2Lys | 0.39 | -1.35 | Neutral | | | | 0.39 | -1.35 | Neutral |
| 2 | p.Glu2Thr | 0.41 | -1.27 | Neutral | | | | 0.41 | -1.27 | Neutral |
| 2 | p.Glu2Arg | 0.21 | -2.22 | Neutral | | | | 0.21 | -2.22 | Neutral |
| 2 | p.Glu2Ser | 0.64 | -0.65 | Neutral | | | | 0.64 | -0.65 | Neutral |
| 2 | p.Glu2Ile | 0.24 | -2.03 | Neutral | | | | 0.24 | -2.03 | Neutral |
| 2 | p.Glu2Met | 0.57 | -0.81 | Neutral | | | | 0.57 | -0.81 | Neutral |
| 2 | p.Glu2His | 0.40 | -1.34 | Neutral | | | | 0.40 | -1.34 | Neutral |
| 2 | p.Glu2Gln | 0.27 | -1.90 | Neutral | | | | 0.27 | -1.90 | Neutral |
| 2 | p.Glu2Pro | 0.31 | -1.70 | Neutral | | | | 0.31 | -1.70 | Neutral |
| 2 | p.Glu2Leu | 0.25 | -2.02 | Neutral | | | | 0.25 | -2.02 | Neutral |
| 2 | p.Glu2Asp | 0.15 | -2.75 | Neutral | | | | 0.15 | -2.75 | Neutral |
| 2 | p.Glu2Glu | 1.00 | 0.00 | Neutral | | | | 1.00 | 0.00 | Neutral |
| 2 | p.Glu2Ala | 0.61 | -0.71 | Neutral | | | | 0.61 | -0.71 | Neutral |
| 2 | p.Glu2Gly | 0.36 | -1.49 | Neutral | | | | 0.36 | -1.49 | Neutral |
| 2 | p.Glu2Val | 0.23 | -2.10 | Neutral | | | | 0.23 | -2.10 | Neutral |
| 2 | p.Glu2Tyr | 0.31 | -1.69 | Neutral | | | | 0.31 | -1.69 | Neutral |
| 2 | p.Glu2Cys | 0.64 | -0.64 | Neutral | | | | 0.64 | -0.64 | Neutral |
| 2 | p.Glu2Trp | 0.44 | -1.18 | Neutral | | | | 0.44 | -1.18 | Neutral |
| 2 | p.Glu2Phe | 0.38 | -1.38 | Neutral | | | | 0.38 | -1.38 | Neutral |
| 3 | p.Pro3Asn | 1.38 | 0.46 | Indeterminate | | | | 1.38 | 0.46 | Indeterminate |
| 3 | p.Pro3Lys | 1.48 | 0.57 | Indeterminate | | | | 1.48 | 0.57 | Indeterminate |
| 3 | p.Pro3Thr | 1.39 | 0.48 | Indeterminate | | | | 1.39 | 0.48 | Indeterminate |
| 3 | p.Pro3Arg | 2.22 | 1.15 | Deleterious | | | | 2.22 | 1.15 | Deleterious |
| 3 | p.Pro3Ser | 1.71 | 0.78 | Indeterminate | | | | 1.71 | 0.78 | Indeterminate |
| 3 | p.Pro3Ile | 1.33 | 0.41 | Indeterminate | | | | 1.33 | 0.41 | Indeterminate |
| 3 | p.Pro3Met | 1.69 | 0.76 | Indeterminate | | | | 1.69 | 0.76 | Indeterminate |
| 3 | p.Pro3His | 1.08 | 0.11 | Neutral | | | | 1.08 | 0.11 | Neutral |
| 3 | p.Pro3Gln | 0.74 | -0.44 | Neutral | | | | 0.74 | -0.44 | Neutral |
| 3 | p.Pro3Pro | 1.00 | 0.00 | Neutral | | | | 1.00 | 0.00 | Neutral |
| 3 | p.Pro3Leu | 1.52 | 0.60 | Indeterminate | | | | 1.52 | 0.60 | Indeterminate |
| 3 | p.Pro3Asp | 1.68 | 0.75 | Indeterminate | | | | 1.68 | 0.75 | Indeterminate |
| 3 | p.Pro3Glu | 1.15 | 0.20 | Neutral | | | | 1.15 | 0.20 | Neutral |
| 3 | p.Pro3Ala | 1.06 | 0.09 | Neutral | | | | 1.06 | 0.09 | Neutral |
| 3 | p.Pro3Gly | 2.75 | 1.46 | Deleterious | | | | 2.75 | 1.46 | Deleterious |
| 3 | p.Pro3Val | 1.71 | 0.77 | Indeterminate | | | | 1.71 | 0.77 | Indeterminate |
| 3 | p.Pro3Tyr | 1.18 | 0.23 | Neutral | | | | 1.18 | 0.23 | Neutral |
| 3 | p.Pro3Cys | 1.49 | 0.58 | Indeterminate | | | | 1.49 | 0.58 | Indeterminate |
| 3 | p.Pro3Trp | 1.94 | 0.96 | Indeterminate | | | | 1.94 | 0.96 | Indeterminate |
| 3 | p.Pro3Phe | 1.69 | 0.76 | Indeterminate | | | | 1.69 | 0.76 | Indeterminate |
| 4 | p.Ala4Asn | 1.36 | 0.44 | Indeterminate | | | | 1.36 | 0.44 | Indeterminate |
| 4 | p.Ala4Lys | 0.98 | -0.02 | Neutral | | | | 0.98 | -0.02 | Neutral |
| 4 | p.Ala4Thr | 2.78 | 1.48 | Deleterious | | | | 2.78 | 1.48 | Deleterious |
| 4 | p.Ala4Arg | 1.07 | 0.10 | Neutral | | | | 1.07 | 0.10 | Neutral |
| 4 | p.Ala4Ser | 2.88 | 1.53 | Deleterious | | | | 2.88 | 1.53 | Deleterious |
| 4 | p.Ala4Ile | 0.76 | -0.40 | Neutral | | | | 0.76 | -0.40 | Neutral |
| 4 | p.Ala4Met | 1.45 | 0.54 | Indeterminate | | | | 1.45 | 0.54 | Indeterminate |
| 4 | p.Ala4His | 1.16 | 0.21 | Neutral | | | | 1.16 | 0.21 | Neutral |
| 4 | p.Ala4Gln | 0.77 | -0.37 | Neutral | | | | 0.77 | -0.37 | Neutral |
| 4 | p.Ala4Pro | 0.84 | -0.25 | Neutral | | | | 0.84 | -0.25 | Neutral |
| 4 | p.Ala4Leu | 0.96 | -0.06 | Neutral | | | | 0.96 | -0.06 | Neutral |
| 4 | p.Ala4Asp | 1.10 | 0.13 | Neutral | | | | 1.10 | 0.13 | Neutral |
| 4 | p.Ala4Glu | 0.65 | -0.63 | Neutral | | | | 0.65 | -0.63 | Neutral |
| 4 | p.Ala4Ala | 1.00 | 0.00 | Neutral | | | | 1.00 | 0.00 | Neutral |
| 4 | p.Ala4Gly | 1.34 | 0.42 | Indeterminate | | | | 1.34 | 0.42 | Indeterminate |
| 4 | p.Ala4Val | 0.78 | -0.36 | Neutral | | | | 0.78 | -0.36 | Neutral |
| 4 | p.Ala4Tyr | 0.90 | -0.15 | Neutral | | | | 0.90 | -0.15 | Neutral |
| 4 | p.Ala4Cys | 2.08 | 1.06 | Indeterminate | | | | 2.08 | 1.06 | Indeterminate |
| 4 | p.Ala4Trp | 1.12 | 0.16 | Neutral | | | | 1.12 | 0.16 | Neutral |
| 4 | p.Ala4Phe | 1.42 | 0.51 | Indeterminate | | | | 1.42 | 0.51 | Indeterminate |
| 5 | p.Ala5Asn | 0.80 | -0.32 | Neutral | | | | 0.80 | -0.32 | Neutral |
| 5 | p.Ala5Lys | 0.37 | -1.44 | Neutral | | | | 0.37 | -1.44 | Neutral |
| 5 | p.Ala5Thr | 1.04 | 0.05 | Neutral | | | | 1.04 | 0.05 | Neutral |
| 5 | p.Ala5Arg | 0.57 | -0.81 | Neutral | | | | 0.57 | -0.81 | Neutral |
| 5 | p.Ala5Ser | 2.86 | 1.52 | Deleterious | | | | 2.86 | 1.52 | Deleterious |
| 5 | p.Ala5Ile | 0.31 | -1.68 | Neutral | | | | 0.31 | -1.68 | Neutral |
| 5 | p.Ala5Met | 0.87 | -0.21 | Neutral | | | | 0.87 | -0.21 | Neutral |
| 5 | p.Ala5His | 1.10 | 0.14 | Neutral | | | | 1.10 | 0.14 | Neutral |
| 5 | p.Ala5Gln | 1.11 | 0.15 | Neutral | | | | 1.11 | 0.15 | Neutral |
| 5 | p.Ala5Pro | 0.86 | -0.21 | Neutral | | | | 0.86 | -0.21 | Neutral |
| 5 | p.Ala5Leu | 1.88 | 0.91 | Indeterminate | | | | 1.88 | 0.91 | Indeterminate |
| 5 | p.Ala5Asp | 1.14 | 0.19 | Neutral | | | | 1.14 | 0.19 | Neutral |
| 5 | p.Ala5Glu | 1.37 | 0.46 | Indeterminate | | | | 1.37 | 0.46 | Indeterminate |
| 5 | p.Ala5Ala | 1.00 | 0.00 | Neutral | | | | 1.00 | 0.00 | Neutral |
| 5 | p.Ala5Gly | 0.77 | -0.38 | Neutral | | | | 0.77 | -0.38 | Neutral |
| 5 | p.Ala5Val | 0.89 | -0.17 | Neutral | | | | 0.89 | -0.17 | Neutral |

| | | | | | | | |
|---|---|---|---|---|---|---|---|
| 5 | p.Ala5Tyr | 4.82 | 2.27 | Deleterious | 4.82 | 2.27 | Deleterious |
| 5 | p.Ala5Cys | 1.26 | 0.34 | Indeterminate | 1.26 | 0.34 | Indeterminate |
| 5 | p.Ala5Trp | 2.44 | 1.29 | Deleterious | 2.44 | 1.29 | Deleterious |
| 5 | p.Ala5Phe | 0.90 | -0.16 | Neutral | 0.90 | -0.16 | Neutral |
| 6 | p.Gly6Asn | 0.95 | -0.07 | Neutral | 0.95 | -0.07 | Neutral |
| 6 | p.Gly6Lys | 1.40 | 0.48 | Indeterminate | 1.40 | 0.48 | Indeterminate |
| 6 | p.Gly6Thr | 1.47 | 0.56 | Indeterminate | 1.47 | 0.56 | Indeterminate |
| 6 | p.Gly6Arg | 4.48 | 2.16 | Deleterious | 4.48 | 2.16 | Deleterious |
| 6 | p.Gly6Ser | 3.45 | 1.79 | Deleterious | 3.45 | 1.79 | Deleterious |
| 6 | p.Gly6Ile | 3.76 | 1.91 | Deleterious | 3.76 | 1.91 | Deleterious |
| 6 | p.Gly6Met | 3.51 | 1.81 | Deleterious | 3.51 | 1.81 | Deleterious |
| 6 | p.Gly6His | 0.72 | -0.48 | Neutral | 0.72 | -0.48 | Neutral |
| 6 | p.Gly6Gln | 0.87 | -0.20 | Neutral | 0.87 | -0.20 | Neutral |
| 6 | p.Gly6Pro | 2.07 | 1.05 | Indeterminate | 2.07 | 1.05 | Indeterminate |
| 6 | p.Gly6Leu | 1.24 | 0.32 | Indeterminate | 1.24 | 0.32 | Indeterminate |
| 6 | p.Gly6Asp | 1.04 | 0.06 | Neutral | 1.04 | 0.06 | Neutral |
| 6 | p.Gly6Glu | 1.16 | 0.22 | Neutral | 1.16 | 0.22 | Neutral |
| 6 | p.Gly6Ala | 1.05 | 0.07 | Neutral | 1.05 | 0.07 | Neutral |
| 6 | p.Gly6Gly | 1.00 | 0.00 | Neutral | 1.00 | 0.00 | Neutral |
| 6 | p.Gly6Val | 1.00 | 0.00 | Neutral | 1.00 | 0.00 | Neutral |
| 6 | p.Gly6Tyr | 1.01 | 0.01 | Neutral | 1.01 | 0.01 | Neutral |
| 6 | p.Gly6Cys | 1.83 | 0.87 | Indeterminate | 1.83 | 0.87 | Indeterminate |
| 6 | p.Gly6Trp | 1.44 | 0.53 | Indeterminate | 1.44 | 0.53 | Indeterminate |
| 6 | p.Gly6Phe | 1.58 | 0.66 | Indeterminate | 1.58 | 0.66 | Indeterminate |
| 7 | p.Ser7Asn | 0.89 | -0.17 | Neutral | 0.89 | -0.17 | Neutral |
| 7 | p.Ser7Lys | 0.91 | -0.14 | Neutral | 0.91 | -0.14 | Neutral |
| 7 | p.Ser7Thr | 2.04 | 1.03 | Indeterminate | 2.04 | 1.03 | Indeterminate |
| 7 | p.Ser7Arg | 0.82 | -0.29 | Neutral | 0.82 | -0.29 | Neutral |
| 7 | p.Ser7Ser | 1.00 | 0.00 | Neutral | 1.00 | 0.00 | Neutral |
| 7 | p.Ser7Ile | 1.23 | 0.30 | Indeterminate | 1.23 | 0.30 | Indeterminate |
| 7 | p.Ser7Met | 1.18 | 0.24 | Indeterminate | 1.18 | 0.24 | Indeterminate |
| 7 | p.Ser7His | 1.47 | 0.56 | Indeterminate | 1.47 | 0.56 | Indeterminate |
| 7 | p.Ser7Gln | 1.10 | 0.14 | Neutral | 1.10 | 0.14 | Neutral |
| 7 | p.Ser7Pro | 1.26 | 0.34 | Indeterminate | 1.26 | 0.34 | Indeterminate |
| 7 | p.Ser7Leu | 1.80 | 0.85 | Indeterminate | 1.80 | 0.85 | Indeterminate |
| 7 | p.Ser7Asp | 0.99 | -0.02 | Neutral | 0.99 | -0.02 | Neutral |
| 7 | p.Ser7Glu | 1.65 | 0.73 | Indeterminate | 1.65 | 0.73 | Indeterminate |
| 7 | p.Ser7Ala | 1.95 | 0.97 | Indeterminate | 1.95 | 0.97 | Indeterminate |
| 7 | p.Ser7Gly | 0.93 | -0.10 | Neutral | 0.93 | -0.10 | Neutral |
| 7 | p.Ser7Val | 1.00 | 0.00 | Neutral | 1.00 | 0.00 | Neutral |
| 7 | p.Ser7Tyr | 2.57 | 1.36 | Deleterious | 2.57 | 1.36 | Deleterious |
| 7 | p.Ser7Cys | 1.26 | 0.33 | Indeterminate | 1.26 | 0.33 | Indeterminate |
| 7 | p.Ser7Trp | 1.31 | 0.39 | Indeterminate | 1.31 | 0.39 | Indeterminate |
| 7 | p.Ser7Phe | 1.17 | 0.22 | Neutral | 1.17 | 0.22 | Neutral |
| 8 | p.Ser8Asn | 0.57 | -0.81 | Neutral | 0.57 | -0.81 | Neutral |
| 8 | p.Ser8Lys | 1.73 | 0.79 | Indeterminate | 1.73 | 0.79 | Indeterminate |
| 8 | p.Ser8Thr | 0.92 | -0.12 | Neutral | 0.92 | -0.12 | Neutral |
| 8 | p.Ser8Arg | 0.69 | -0.53 | Neutral | 0.69 | -0.53 | Neutral |
| 8 | p.Ser8Ser | 1.00 | 0.00 | Neutral | 1.00 | 0.00 | Neutral |
| 8 | p.Ser8Ile | 0.72 | -0.48 | Neutral | 0.72 | -0.48 | Neutral |
| 8 | p.Ser8Met | 0.56 | -0.85 | Neutral | 0.56 | -0.85 | Neutral |
| 8 | p.Ser8His | 0.60 | -0.73 | Neutral | 0.60 | -0.73 | Neutral |
| 8 | p.Ser8Gln | 0.67 | -0.59 | Neutral | 0.67 | -0.59 | Neutral |
| 8 | p.Ser8Pro | 0.83 | -0.26 | Neutral | 0.83 | -0.26 | Neutral |
| 8 | p.Ser8Leu | 0.84 | -0.26 | Neutral | 0.84 | -0.26 | Neutral |
| 8 | p.Ser8Asp | 0.73 | -0.46 | Neutral | 0.73 | -0.46 | Neutral |
| 8 | p.Ser8Glu | 0.45 | -1.15 | Neutral | 0.45 | -1.15 | Neutral |
| 8 | p.Ser8Ala | 0.68 | -0.56 | Neutral | 0.68 | -0.56 | Neutral |
| 8 | p.Ser8Gly | 0.82 | -0.28 | Neutral | 0.82 | -0.28 | Neutral |
| 8 | p.Ser8Val | 0.71 | -0.49 | Neutral | 0.71 | -0.49 | Neutral |
| 8 | p.Ser8Tyr | 0.61 | -0.72 | Neutral | 0.61 | -0.72 | Neutral |
| 8 | p.Ser8Cys | 0.52 | -0.94 | Neutral | 0.52 | -0.94 | Neutral |
| 8 | p.Ser8Trp | 0.58 | -0.79 | Neutral | 0.58 | -0.79 | Neutral |
| 8 | p.Ser8Phe | 0.75 | -0.41 | Neutral | 0.75 | -0.41 | Neutral |
| 9 | p.Met9Asn | 1.10 | 0.14 | Neutral | 1.10 | 0.14 | Neutral |
| 9 | p.Met9Lys | 1.21 | 0.27 | Indeterminate | 1.21 | 0.27 | Indeterminate |
| 9 | p.Met9Thr | 1.19 | 0.26 | Indeterminate | 1.19 | 0.26 | Indeterminate |
| 9 | p.Met9Arg | 1.14 | 0.18 | Neutral | 1.14 | 0.18 | Neutral |
| 9 | p.Met9Ser | 1.16 | 0.22 | Neutral | 1.16 | 0.22 | Neutral |
| 9 | p.Met9Ile | 1.18 | 0.24 | Indeterminate | 1.18 | 0.24 | Indeterminate |
| 9 | p.Met9Met | 1.00 | 0.00 | Neutral | 1.00 | 0.00 | Neutral |
| 9 | p.Met9His | 1.05 | 0.07 | Neutral | 1.05 | 0.07 | Neutral |
| 9 | p.Met9Gln | 1.04 | 0.05 | Neutral | 1.04 | 0.05 | Neutral |
| 9 | p.Met9Pro | 1.18 | 0.24 | Neutral | 1.18 | 0.24 | Neutral |
| 9 | p.Met9Leu | 0.92 | -0.11 | Neutral | 0.92 | -0.11 | Neutral |
| 9 | p.Met9Asp | 1.16 | 0.21 | Neutral | 1.16 | 0.21 | Neutral |
| 9 | p.Met9Glu | 1.02 | 0.03 | Neutral | 1.02 | 0.03 | Neutral |
| 9 | p.Met9Ala | 1.07 | 0.10 | Neutral | 1.07 | 0.10 | Neutral |
| 9 | p.Met9Gly | 1.10 | 0.14 | Neutral | 1.10 | 0.14 | Neutral |
| 9 | p.Met9Val | 1.25 | 0.32 | Indeterminate | 1.25 | 0.32 | Indeterminate |
| 9 | p.Met9Tyr | 1.00 | 0.00 | Neutral | 1.00 | 0.00 | Neutral |
| 9 | p.Met9Cys | 1.08 | 0.11 | Neutral | 1.08 | 0.11 | Neutral |
| 9 | p.Met9Trp | 1.13 | 0.17 | Neutral | 1.13 | 0.17 | Neutral |
| 9 | p.Met9Phe | 1.14 | 0.19 | Neutral | 1.14 | 0.19 | Neutral |
| 10 | p.Glu10Asn | 1.08 | 0.11 | Neutral | 1.08 | 0.11 | Neutral |
| 10 | p.Glu10Lys | 2.31 | 1.21 | Deleterious | 2.31 | 1.21 | Deleterious |
| 10 | p.Glu10Thr | 1.18 | 0.24 | Neutral | 1.18 | 0.24 | Neutral |
| 10 | p.Glu10Arg | 0.94 | -0.09 | Neutral | 0.94 | -0.09 | Neutral |
| 10 | p.Glu10Ser | 1.65 | 0.72 | Indeterminate | 1.65 | 0.72 | Indeterminate |
| 10 | p.Glu10Ile | 1.54 | 0.62 | Indeterminate | 1.54 | 0.62 | Indeterminate |
| 10 | p.Glu10Met | 1.67 | 0.74 | Indeterminate | 1.67 | 0.74 | Indeterminate |
| 10 | p.Glu10His | 2.29 | 1.20 | Deleterious | 2.29 | 1.20 | Deleterious |
| 10 | p.Glu10Gln | 0.93 | -0.10 | Neutral | 0.93 | -0.10 | Neutral |
| 10 | p.Glu10Pro | 1.03 | 0.04 | Neutral | 1.03 | 0.04 | Neutral |
| 10 | p.Glu10Leu | 1.15 | 0.20 | Neutral | 1.15 | 0.20 | Neutral |
| 10 | p.Glu10Asp | 1.13 | 0.17 | Neutral | 1.13 | 0.17 | Neutral |
| 10 | p.Glu10Glu | 1.00 | 0.00 | Neutral | 1.00 | 0.00 | Neutral |
| 10 | p.Glu10Ala | 1.77 | 0.82 | Indeterminate | 1.77 | 0.82 | Indeterminate |
| 10 | p.Glu10Gly | 1.04 | 0.06 | Neutral | 1.04 | 0.06 | Neutral |
| 10 | p.Glu10Val | 1.04 | 0.06 | Neutral | 1.04 | 0.06 | Neutral |
| 10 | p.Glu10Tyr | 1.33 | 0.41 | Indeterminate | 1.33 | 0.41 | Indeterminate |

| | | | | | | | | | | |
|---|---|---|---|---|---|---|---|---|---|---|
| 10 | p.Glu10Cys | 1.15 | 0.20 | Neutral | | | | 1.15 | 0.20 | Neutral |
| 10 | p.Glu10Trp | 1.01 | 0.02 | Neutral | | | | 1.01 | 0.02 | Neutral |
| 10 | p.Glu10Phe | 1.11 | 0.15 | Neutral | | | | 1.11 | 0.15 | Neutral |
| 11 | p.Pro11Asn | 1.13 | 0.17 | Neutral | | | | 1.13 | 0.17 | Neutral |
| 11 | p.Pro11Lys | 1.28 | 0.36 | Indeterminate | | | | 1.28 | 0.36 | Indeterminate |
| 11 | p.Pro11Thr | 1.44 | 0.53 | Indeterminate | | | | 1.44 | 0.53 | Indeterminate |
| 11 | p.Pro11Arg | 1.08 | 0.11 | Neutral | | | | 1.08 | 0.11 | Neutral |
| 11 | p.Pro11Ser | 1.21 | 0.27 | Indeterminate | | | | 1.21 | 0.27 | Indeterminate |
| 11 | p.Pro11Ile | 1.13 | 0.18 | Neutral | | | | 1.13 | 0.18 | Neutral |
| 11 | p.Pro11Met | 1.07 | 0.09 | Neutral | | | | 1.07 | 0.09 | Neutral |
| 11 | p.Pro11His | 1.12 | 0.17 | Neutral | | | | 1.12 | 0.17 | Neutral |
| 11 | p.Pro11Gln | 0.96 | -0.06 | Neutral | | | | 0.96 | -0.06 | Neutral |
| 11 | p.Pro11Pro | 1.00 | 0.00 | Neutral | | | | 1.00 | 0.00 | Neutral |
| 11 | p.Pro11Leu | 1.12 | 0.16 | Neutral | | | | 1.12 | 0.16 | Neutral |
| 11 | p.Pro11Asp | 1.07 | 0.09 | Neutral | | | | 1.07 | 0.09 | Neutral |
| 11 | p.Pro11Glu | 1.01 | 0.01 | Neutral | | | | 1.01 | 0.01 | Neutral |
| 11 | p.Pro11Ala | 1.10 | 0.14 | Neutral | | | | 1.10 | 0.14 | Neutral |
| 11 | p.Pro11Gly | 1.20 | 0.26 | Indeterminate | | | | 1.20 | 0.26 | Indeterminate |
| 11 | p.Pro11Val | 1.09 | 0.12 | Neutral | | | | 1.09 | 0.12 | Neutral |
| 11 | p.Pro11Tyr | 1.22 | 0.28 | Indeterminate | | | | 1.22 | 0.28 | Indeterminate |
| 11 | p.Pro11Cys | 1.01 | 0.02 | Neutral | | | | 1.01 | 0.02 | Neutral |
| 11 | p.Pro11Trp | 1.31 | 0.39 | Indeterminate | | | | 1.31 | 0.39 | Indeterminate |
| 11 | p.Pro11Phe | 1.22 | 0.29 | Indeterminate | | | | 1.22 | 0.29 | Indeterminate |
| 12 | p.Ser12Asn | 0.92 | -0.12 | Neutral | | | | 0.92 | -0.12 | Neutral |
| 12 | p.Ser12Lys | 0.71 | -0.49 | Neutral | | | | 0.71 | -0.49 | Neutral |
| 12 | p.Ser12Thr | 0.76 | -0.39 | Neutral | | | | 0.76 | -0.39 | Neutral |
| 12 | p.Ser12Arg | 0.84 | -0.25 | Neutral | | | | 0.84 | -0.25 | Neutral |
| 12 | p.Ser12Ser | 1.00 | 0.00 | Neutral | | | | 1.00 | 0.00 | Neutral |
| 12 | p.Ser12Ile | 0.81 | -0.31 | Neutral | | | | 0.81 | -0.31 | Neutral |
| 12 | p.Ser12Met | 0.71 | -0.49 | Neutral | | | | 0.71 | -0.49 | Neutral |
| 12 | p.Ser12His | 1.12 | 0.16 | Neutral | | | | 1.12 | 0.16 | Neutral |
| 12 | p.Ser12Gln | 1.15 | 0.20 | Neutral | | | | 1.15 | 0.20 | Neutral |
| 12 | p.Ser12Pro | 0.78 | -0.36 | Neutral | | | | 0.78 | -0.36 | Neutral |
| 12 | p.Ser12Leu | 2.35 | 1.23 | Deleterious | | | | 2.35 | 1.23 | Deleterious |
| 12 | p.Ser12Asp | 0.66 | -0.59 | Neutral | | | | 0.66 | -0.59 | Neutral |
| 12 | p.Ser12Glu | 0.88 | -0.18 | Neutral | | | | 0.88 | -0.18 | Neutral |
| 12 | p.Ser12Ala | 0.85 | -0.23 | Neutral | | | | 0.85 | -0.23 | Neutral |
| 12 | p.Ser12Gly | 0.82 | -0.29 | Neutral | | | | 0.82 | -0.29 | Neutral |
| 12 | p.Ser12Val | 1.13 | 0.18 | Neutral | | | | 1.13 | 0.18 | Neutral |
| 12 | p.Ser12Tyr | 0.75 | -0.41 | Neutral | | | | 0.75 | -0.41 | Neutral |
| 12 | p.Ser12Cys | 1.13 | 0.18 | Neutral | | | | 1.13 | 0.18 | Neutral |
| 12 | p.Ser12Trp | 1.13 | 0.18 | Neutral | | | | 1.13 | 0.18 | Neutral |
| 12 | p.Ser12Phe | 1.11 | 0.15 | Neutral | | | | 1.11 | 0.15 | Neutral |
| 13 | p.Ala13Asn | 0.61 | -0.72 | Neutral | | | | 0.61 | -0.72 | Neutral |
| 13 | p.Ala13Lys | 0.71 | -0.49 | Neutral | | | | 0.71 | -0.49 | Neutral |
| 13 | p.Ala13Thr | 0.70 | -0.51 | Neutral | | | | 0.70 | -0.51 | Neutral |
| 13 | p.Ala13Arg | 0.61 | -0.72 | Neutral | | | | 0.61 | -0.72 | Neutral |
| 13 | p.Ala13Ser | 0.63 | -0.66 | Neutral | | | | 0.63 | -0.66 | Neutral |
| 13 | p.Ala13Ile | 0.65 | -0.61 | Neutral | | | | 0.65 | -0.61 | Neutral |
| 13 | p.Ala13Met | 1.07 | 0.10 | Neutral | | | | 1.07 | 0.10 | Neutral |
| 13 | p.Ala13His | 0.75 | -0.41 | Neutral | | | | 0.75 | -0.41 | Neutral |
| 13 | p.Ala13Gln | 0.66 | -0.59 | Neutral | | | | 0.66 | -0.59 | Neutral |
| 13 | p.Ala13Pro | 0.53 | -0.90 | Neutral | | | | 0.53 | -0.90 | Neutral |
| 13 | p.Ala13Leu | 0.74 | -0.43 | Neutral | | | | 0.74 | -0.43 | Neutral |
| 13 | p.Ala13Asp | 0.68 | -0.55 | Neutral | | | | 0.68 | -0.55 | Neutral |
| 13 | p.Ala13Glu | 0.64 | -0.64 | Neutral | | | | 0.64 | -0.64 | Neutral |
| 13 | p.Ala13Ala | 1.00 | 0.00 | Neutral | | | | 1.00 | 0.00 | Neutral |
| 13 | p.Ala13Gly | 0.57 | -0.81 | Neutral | | | | 0.57 | -0.81 | Neutral |
| 13 | p.Ala13Val | 0.70 | -0.51 | Neutral | | | | 0.70 | -0.51 | Neutral |
| 13 | p.Ala13Tyr | 1.03 | 0.04 | Neutral | | | | 1.03 | 0.04 | Neutral |
| 13 | p.Ala13Cys | 0.82 | -0.29 | Neutral | | | | 0.82 | -0.29 | Neutral |
| 13 | p.Ala13Trp | 0.77 | -0.38 | Neutral | | | | 0.77 | -0.38 | Neutral |
| 13 | p.Ala13Phe | 0.94 | -0.09 | Neutral | | | | 0.94 | -0.09 | Neutral |
| 14 | p.Asp14Asn | 1.14 | 0.19 | Neutral | | | | 1.14 | 0.19 | Neutral |
| 14 | p.Asp14Lys | 1.10 | 0.14 | Neutral | | | | 1.10 | 0.14 | Neutral |
| 14 | p.Asp14Thr | 0.95 | -0.08 | Neutral | | | | 0.95 | -0.08 | Neutral |
| 14 | p.Asp14Arg | 1.63 | 0.71 | Indeterminate | | | | 1.63 | 0.71 | Indeterminate |
| 14 | p.Asp14Ser | 0.92 | -0.13 | Neutral | | | | 0.92 | -0.13 | Neutral |
| 14 | p.Asp14Ile | 2.30 | 1.20 | Deleterious | | | | 2.30 | 1.20 | Deleterious |
| 14 | p.Asp14Met | 1.77 | 0.82 | Indeterminate | | | | 1.77 | 0.82 | Indeterminate |
| 14 | p.Asp14His | 0.95 | -0.07 | Neutral | | | | 0.95 | -0.07 | Neutral |
| 14 | p.Asp14Gln | 1.12 | 0.17 | Neutral | | | | 1.12 | 0.17 | Neutral |
| 14 | p.Asp14Pro | 2.56 | 1.36 | Deleterious | | | | 2.56 | 1.36 | Deleterious |
| 14 | p.Asp14Leu | 1.80 | 0.84 | Indeterminate | | | | 1.80 | 0.84 | Indeterminate |
| 14 | p.Asp14Asp | 1.00 | 0.00 | Neutral | | | | 1.00 | 0.00 | Neutral |
| 14 | p.Asp14Glu | 1.09 | 0.12 | Neutral | | | | 1.09 | 0.12 | Neutral |
| 14 | p.Asp14Ala | 1.37 | 0.46 | Indeterminate | | | | 1.37 | 0.46 | Indeterminate |
| 14 | p.Asp14Gly | 1.04 | 0.06 | Neutral | | | | 1.04 | 0.06 | Neutral |
| 14 | p.Asp14Val | 2.47 | 1.31 | Deleterious | | | | 2.47 | 1.31 | Deleterious |
| 14 | p.Asp14Tyr | 1.93 | 0.95 | Indeterminate | | | | 1.93 | 0.95 | Indeterminate |
| 14 | p.Asp14Cys | 1.05 | 0.06 | Neutral | | | | 1.05 | 0.06 | Neutral |
| 14 | p.Asp14Trp | 1.87 | 0.90 | Indeterminate | | | | 1.87 | 0.90 | Indeterminate |
| 14 | p.Asp14Phe | 1.93 | 0.95 | Indeterminate | | | | 1.93 | 0.95 | Indeterminate |
| 15 | p.Trp15Asn | 1.51 | 0.59 | Indeterminate | 0.73 | -0.46 | Neutral | 1.12 | 0.16 | Neutral |
| 15 | p.Trp15Lys | 1.37 | 0.46 | Indeterminate | 0.72 | -0.48 | Neutral | 1.04 | 0.06 | Neutral |
| 15 | p.Trp15Thr | 1.65 | 0.73 | Indeterminate | 1.03 | 0.04 | Neutral | 1.34 | 0.42 | Indeterminate |
| 15 | p.Trp15Arg | 1.19 | 0.25 | Indeterminate | 0.65 | -0.62 | Neutral | 0.92 | -0.12 | Neutral |
| 15 | p.Trp15Ser | 1.37 | 0.46 | Indeterminate | 0.69 | -0.54 | Neutral | 1.03 | 0.04 | Neutral |
| 15 | p.Trp15Ile | 1.03 | 0.05 | Neutral | 0.99 | -0.02 | Neutral | 1.01 | 0.01 | Neutral |
| 15 | p.Trp15Met | 1.13 | 0.17 | Neutral | 0.93 | -0.10 | Neutral | 1.03 | 0.04 | Neutral |
| 15 | p.Trp15His | 1.48 | 0.57 | Indeterminate | 1.61 | 0.69 | Indeterminate | 1.55 | 0.63 | Indeterminate |
| 15 | p.Trp15Gln | 1.46 | 0.54 | Indeterminate | 1.13 | 0.18 | Indeterminate | 1.29 | 0.37 | Indeterminate |
| 15 | p.Trp15Pro | 1.83 | 0.87 | Indeterminate | 1.46 | 0.55 | Indeterminate | 1.65 | 0.72 | Indeterminate |
| 15 | p.Trp15Leu | 1.30 | 0.38 | Indeterminate | 1.14 | 0.19 | Indeterminate | 1.22 | 0.29 | Indeterminate |
| 15 | p.Trp15Asp | 1.28 | 0.35 | Indeterminate | 1.06 | 0.08 | Neutral | 1.17 | 0.23 | Neutral |
| 15 | p.Trp15Glu | 1.09 | 0.13 | Neutral | 1.11 | 0.15 | Indeterminate | 1.10 | 0.14 | Neutral |
| 15 | p.Trp15Ala | 1.36 | 0.44 | Indeterminate | 0.77 | -0.37 | Neutral | 1.07 | 0.09 | Neutral |
| 15 | p.Trp15Gly | 1.18 | 0.24 | Neutral | 1.46 | 0.54 | Indeterminate | 1.32 | 0.40 | Indeterminate |
| 15 | p.Trp15Val | 1.06 | 0.08 | Neutral | 0.62 | -0.68 | Neutral | 0.84 | -0.25 | Neutral |
| 15 | p.Trp15Tyr | 1.67 | 0.74 | Indeterminate | 0.91 | -0.13 | Neutral | 1.29 | 0.37 | Indeterminate |
| 15 | p.Trp15Cys | 1.51 | 0.59 | Indeterminate | 0.82 | -0.28 | Neutral | 1.17 | 0.22 | Neutral |

| | | | | | | | | | | |
|---|---|---|---|---|---|---|---|---|---|---|
| 15 | p.Trp15Trp | 1.00 | 0.00 | Neutral | 1.00 | 0.00 | Neutral | 1.00 | 0.00 | Neutral |
| 15 | p.Trp15Phe | 1.29 | 0.37 | Indeterminate | 0.77 | -0.37 | Neutral | 1.03 | 0.05 | Neutral |
| 16 | p.Leu16Asn | 17.05 | 4.09 | Deleterious | | | | 17.05 | 4.09 | Deleterious |
| 16 | p.Leu16Lys | 24.15 | 4.59 | Deleterious | | | | 24.15 | 4.59 | Deleterious |
| 16 | p.Leu16Thr | 2.31 | 1.21 | Deleterious | | | | 2.31 | 1.21 | Deleterious |
| 16 | p.Leu16Arg | 28.65 | 4.84 | Deleterious | | | | 28.65 | 4.84 | Deleterious |
| 16 | p.Leu16Ser | 12.85 | 3.68 | Deleterious | | | | 12.85 | 3.68 | Deleterious |
| 16 | p.Leu16Ile | 1.34 | 0.42 | Indeterminate | | | | 1.34 | 0.42 | Indeterminate |
| 16 | p.Leu16Met | 1.18 | 0.24 | Indeterminate | | | | 1.18 | 0.24 | Indeterminate |
| 16 | p.Leu16His | 16.82 | 4.07 | Deleterious | | | | 16.82 | 4.07 | Deleterious |
| 16 | p.Leu16Gln | 16.71 | 4.06 | Deleterious | | | | 16.71 | 4.06 | Deleterious |
| 16 | p.Leu16Pro | 15.82 | 3.98 | Deleterious | | | | 15.82 | 3.98 | Deleterious |
| 16 | p.Leu16Leu | 1.00 | 0.00 | Neutral | | | | 1.00 | 0.00 | Neutral |
| 16 | p.Leu16Asp | 24.00 | 4.59 | Deleterious | | | | 24.00 | 4.59 | Deleterious |
| 16 | p.Leu16Glu | 21.71 | 4.44 | Deleterious | | | | 21.71 | 4.44 | Deleterious |
| 16 | p.Leu16Ala | 4.92 | 2.30 | Deleterious | | | | 4.92 | 2.30 | Deleterious |
| 16 | p.Leu16Gly | 19.95 | 4.32 | Deleterious | | | | 19.95 | 4.32 | Deleterious |
| 16 | p.Leu16Val | 2.04 | 1.03 | Indeterminate | | | | 2.04 | 1.03 | Indeterminate |
| 16 | p.Leu16Tyr | 21.66 | 4.44 | Deleterious | | | | 21.66 | 4.44 | Deleterious |
| 16 | p.Leu16Cys | 2.04 | 1.03 | Indeterminate | | | | 2.04 | 1.03 | Indeterminate |
| 16 | p.Leu16Trp | 21.58 | 4.43 | Deleterious | | | | 21.58 | 4.43 | Deleterious |
| 16 | p.Leu16Phe | 1.48 | 0.56 | Indeterminate | | | | 1.48 | 0.56 | Indeterminate |
| 17 | p.Ala17Asn | 1.60 | 0.68 | Indeterminate | | | | 1.60 | 0.68 | Indeterminate |
| 17 | p.Ala17Lys | 36.62 | 5.19 | Deleterious | | | | 36.62 | 5.19 | Deleterious |
| 17 | p.Ala17Thr | 1.13 | 0.18 | Neutral | | | | 1.13 | 0.18 | Neutral |
| 17 | p.Ala17Arg | 39.11 | 5.29 | Deleterious | | | | 39.11 | 5.29 | Deleterious |
| 17 | p.Ala17Ser | 1.14 | 0.19 | Neutral | | | | 1.14 | 0.19 | Neutral |
| 17 | p.Ala17Ile | 1.70 | 0.77 | Indeterminate | | | | 1.70 | 0.77 | Indeterminate |
| 17 | p.Ala17Met | 3.32 | 1.73 | Deleterious | | | | 3.32 | 1.73 | Deleterious |
| 17 | p.Ala17His | 1.14 | 0.19 | Neutral | | | | 1.14 | 0.19 | Neutral |
| 17 | p.Ala17Gln | 3.54 | 1.82 | Deleterious | | | | 3.54 | 1.82 | Deleterious |
| 17 | p.Ala17Pro | 1.10 | 0.13 | Neutral | | | | 1.10 | 0.13 | Neutral |
| 17 | p.Ala17Leu | 6.23 | 2.64 | Deleterious | | | | 6.23 | 2.64 | Deleterious |
| 17 | p.Ala17Asp | 2.99 | 1.58 | Deleterious | | | | 2.99 | 1.58 | Deleterious |
| 17 | p.Ala17Glu | 15.21 | 3.93 | Deleterious | | | | 15.21 | 3.93 | Deleterious |
| 17 | p.Ala17Ala | 1.00 | 0.00 | Neutral | | | | 1.00 | 0.00 | Neutral |
| 17 | p.Ala17Gly | 1.01 | 0.01 | Neutral | | | | 1.01 | 0.01 | Neutral |
| 17 | p.Ala17Val | 1.04 | 0.06 | Neutral | | | | 1.04 | 0.06 | Neutral |
| 17 | p.Ala17Tyr | 4.73 | 2.24 | Deleterious | | | | 4.73 | 2.24 | Deleterious |
| 17 | p.Ala17Cys | 0.76 | -0.40 | Neutral | | | | 0.76 | -0.40 | Neutral |
| 17 | p.Ala17Trp | 5.41 | 2.43 | Deleterious | | | | 5.41 | 2.43 | Deleterious |
| 17 | p.Ala17Phe | 3.04 | 1.61 | Deleterious | | | | 3.04 | 1.61 | Deleterious |
| 18 | p.Thr18Asn | 2.54 | 1.34 | Deleterious | | | | 2.54 | 1.34 | Deleterious |
| 18 | p.Thr18Lys | 1.83 | 0.87 | Indeterminate | | | | 1.83 | 0.87 | Indeterminate |
| 18 | p.Thr18Thr | 1.00 | 0.00 | Neutral | | | | 1.00 | 0.00 | Neutral |
| 18 | p.Thr18Arg | 1.81 | 0.85 | Indeterminate | | | | 1.81 | 0.85 | Indeterminate |
| 18 | p.Thr18Ser | 1.45 | 0.53 | Indeterminate | | | | 1.45 | 0.53 | Indeterminate |
| 18 | p.Thr18Ile | 2.08 | 1.06 | Indeterminate | | | | 2.08 | 1.06 | Indeterminate |
| 18 | p.Thr18Met | 2.38 | 1.25 | Deleterious | | | | 2.38 | 1.25 | Deleterious |
| 18 | p.Thr18His | 1.36 | 0.44 | Indeterminate | | | | 1.36 | 0.44 | Indeterminate |
| 18 | p.Thr18Gln | 1.66 | 0.73 | Indeterminate | | | | 1.66 | 0.73 | Indeterminate |
| 18 | p.Thr18Pro | 11.34 | 3.50 | Deleterious | | | | 11.34 | 3.50 | Deleterious |
| 18 | p.Thr18Leu | 2.42 | 1.28 | Deleterious | | | | 2.42 | 1.28 | Deleterious |
| 18 | p.Thr18Asp | 1.66 | 0.73 | Indeterminate | | | | 1.66 | 0.73 | Indeterminate |
| 18 | p.Thr18Glu | 1.46 | 0.55 | Indeterminate | | | | 1.46 | 0.55 | Indeterminate |
| 18 | p.Thr18Ala | 2.46 | 1.30 | Deleterious | | | | 2.46 | 1.30 | Deleterious |
| 18 | p.Thr18Gly | 1.46 | 0.54 | Indeterminate | | | | 1.46 | 0.54 | Indeterminate |
| 18 | p.Thr18Val | 1.50 | 0.59 | Indeterminate | | | | 1.50 | 0.59 | Indeterminate |
| 18 | p.Thr18Tyr | 2.54 | 1.34 | Deleterious | | | | 2.54 | 1.34 | Deleterious |
| 18 | p.Thr18Cys | 1.09 | 0.13 | Neutral | | | | 1.09 | 0.13 | Neutral |
| 18 | p.Thr18Trp | 1.89 | 0.92 | Indeterminate | | | | 1.89 | 0.92 | Indeterminate |
| 18 | p.Thr18Phe | 1.44 | 0.52 | Indeterminate | | | | 1.44 | 0.52 | Indeterminate |
| 19 | p.Ala19Asn | 0.44 | -1.20 | Neutral | | | | 0.44 | -1.20 | Neutral |
| 19 | p.Ala19Lys | 1.07 | 0.10 | Neutral | | | | 1.07 | 0.10 | Neutral |
| 19 | p.Ala19Thr | 0.67 | -0.57 | Neutral | | | | 0.67 | -0.57 | Neutral |
| 19 | p.Ala19Arg | 0.97 | -0.04 | Neutral | | | | 0.97 | -0.04 | Neutral |
| 19 | p.Ala19Ser | 0.42 | -1.24 | Neutral | | | | 0.42 | -1.24 | Neutral |
| 19 | p.Ala19Ile | 0.58 | -0.79 | Neutral | | | | 0.58 | -0.79 | Neutral |
| 19 | p.Ala19Met | 0.41 | -1.29 | Neutral | | | | 0.41 | -1.29 | Neutral |
| 19 | p.Ala19His | 0.73 | -0.46 | Neutral | | | | 0.73 | -0.46 | Neutral |
| 19 | p.Ala19Gln | 0.42 | -1.26 | Neutral | | | | 0.42 | -1.26 | Neutral |
| 19 | p.Ala19Pro | 7.17 | 2.84 | Deleterious | | | | 7.17 | 2.84 | Deleterious |
| 19 | p.Ala19Leu | 0.38 | -1.40 | Neutral | | | | 0.38 | -1.40 | Neutral |
| 19 | p.Ala19Asp | 1.11 | 0.15 | Neutral | | | | 1.11 | 0.15 | Neutral |
| 19 | p.Ala19Glu | 0.50 | -0.99 | Neutral | | | | 0.50 | -0.99 | Neutral |
| 19 | p.Ala19Ala | 1.00 | 0.00 | Neutral | | | | 1.00 | 0.00 | Neutral |
| 19 | p.Ala19Gly | 0.46 | -1.12 | Neutral | | | | 0.46 | -1.12 | Neutral |
| 19 | p.Ala19Val | 0.54 | -0.88 | Neutral | | | | 0.54 | -0.88 | Neutral |
| 19 | p.Ala19Tyr | 0.57 | -0.81 | Neutral | | | | 0.57 | -0.81 | Neutral |
| 19 | p.Ala19Cys | 1.14 | 0.19 | Neutral | | | | 1.14 | 0.19 | Neutral |
| 19 | p.Ala19Trp | 0.40 | -1.33 | Neutral | | | | 0.40 | -1.33 | Neutral |
| 19 | p.Ala19Phe | 0.26 | -1.96 | Neutral | | | | 0.26 | -1.96 | Neutral |
| 20 | p.Ala20Asn | 17.14 | 4.10 | Deleterious | 21.96 | 4.46 | Deleterious | 19.55 | 4.29 | Deleterious |
| 20 | p.Ala20Lys | 21.01 | 4.39 | Deleterious | 26.93 | 4.75 | Deleterious | 23.97 | 4.58 | Deleterious |
| 20 | p.Ala20Thr | 1.70 | 0.77 | Indeterminate | 1.50 | 0.58 | Indeterminate | 1.60 | 0.68 | Indeterminate |
| 20 | p.Ala20Arg | 19.79 | 4.31 | Deleterious | 27.73 | 4.79 | Deleterious | 23.76 | 4.57 | Deleterious |
| 20 | p.Ala20Ser | 0.95 | -0.07 | Neutral | 1.80 | 0.85 | Indeterminate | 1.38 | 0.46 | Indeterminate |
| 20 | p.Ala20Ile | 9.71 | 3.28 | Deleterious | 10.13 | 3.34 | Deleterious | 9.92 | 3.31 | Deleterious |
| 20 | p.Ala20Met | 7.13 | 2.83 | Deleterious | 8.50 | 3.09 | Deleterious | 7.81 | 2.97 | Deleterious |
| 20 | p.Ala20His | 19.43 | 4.28 | Deleterious | 27.11 | 4.76 | Deleterious | 23.27 | 4.54 | Deleterious |
| 20 | p.Ala20Gln | 21.24 | 4.41 | Deleterious | 26.35 | 4.72 | Deleterious | 23.79 | 4.57 | Deleterious |
| 20 | p.Ala20Pro | 21.60 | 4.43 | Deleterious | 25.71 | 4.68 | Deleterious | 23.65 | 4.56 | Deleterious |
| 20 | p.Ala20Leu | 7.16 | 2.84 | Deleterious | 8.21 | 3.04 | Deleterious | 7.69 | 2.94 | Deleterious |
| 20 | p.Ala20Asp | 18.97 | 4.25 | Deleterious | 25.25 | 4.66 | Deleterious | 22.11 | 4.47 | Deleterious |
| 20 | p.Ala20Glu | 20.26 | 4.34 | Deleterious | 28.11 | 4.81 | Deleterious | 24.18 | 4.60 | Deleterious |
| 20 | p.Ala20Ala | 1.00 | 0.00 | Neutral | 1.00 | 0.00 | Neutral | 1.00 | 0.00 | Neutral |
| 20 | p.Ala20Gly | 5.42 | 2.44 | Deleterious | 1.92 | 0.94 | Indeterminate | 3.67 | 1.88 | Deleterious |
| 20 | p.Ala20Val | 1.15 | 0.20 | Neutral | 1.54 | 0.62 | Indeterminate | 1.34 | 0.42 | Indeterminate |
| 20 | p.Ala20Tyr | 21.80 | 4.45 | Deleterious | 26.74 | 4.74 | Deleterious | 24.27 | 4.60 | Deleterious |
| 20 | p.Ala20Cys | 1.18 | 0.24 | Indeterminate | 1.34 | 0.42 | Indeterminate | 1.26 | 0.33 | Indeterminate |
| 20 | p.Ala20Trp | 21.50 | 4.43 | Deleterious | 27.63 | 4.79 | Deleterious | 24.57 | 4.62 | Deleterious |

| | | | | | | | | | | |
|---|---|---|---|---|---|---|---|---|---|---|
| 20 | p.Ala20Phe | 19.92 | 4.32 | Deleterious | 26.72 | 4.74 | Deleterious | 23.32 | 4.54 | Deleterious |
| 21 | p.Ala21Asn | 4.93 | 2.30 | Deleterious | | | | 4.93 | 2.30 | Deleterious |
| 21 | p.Ala21Lys | 30.37 | 4.92 | Deleterious | | | | 30.37 | 4.92 | Deleterious |
| 21 | p.Ala21Thr | 1.48 | 0.57 | Indeterminate | | | | 1.48 | 0.57 | Indeterminate |
| 21 | p.Ala21Arg | 12.05 | 3.59 | Deleterious | | | | 12.05 | 3.59 | Deleterious |
| 21 | p.Ala21Ser | 1.05 | 0.07 | Neutral | | | | 1.05 | 0.07 | Neutral |
| 21 | p.Ala21Ile | 2.45 | 1.29 | Deleterious | | | | 2.45 | 1.29 | Deleterious |
| 21 | p.Ala21Met | 3.33 | 1.73 | Deleterious | | | | 3.33 | 1.73 | Deleterious |
| 21 | p.Ala21His | 9.34 | 3.22 | Deleterious | | | | 9.34 | 3.22 | Deleterious |
| 21 | p.Ala21Gln | 6.36 | 2.67 | Deleterious | | | | 6.36 | 2.67 | Deleterious |
| 21 | p.Ala21Pro | 37.99 | 5.25 | Deleterious | | | | 37.99 | 5.25 | Deleterious |
| 21 | p.Ala21Leu | 1.65 | 0.72 | Indeterminate | | | | 1.65 | 0.72 | Indeterminate |
| 21 | p.Ala21Asp | 23.39 | 4.55 | Deleterious | | | | 23.39 | 4.55 | Deleterious |
| 21 | p.Ala21Glu | 9.77 | 3.29 | Deleterious | | | | 9.77 | 3.29 | Deleterious |
| 21 | p.Ala21Ala | 1.00 | 0.00 | Neutral | | | | 1.00 | 0.00 | Neutral |
| 21 | p.Ala21Gly | 0.97 | -0.04 | Neutral | | | | 0.97 | -0.04 | Neutral |
| 21 | p.Ala21Val | 1.17 | 0.23 | Neutral | | | | 1.17 | 0.23 | Neutral |
| 21 | p.Ala21Tyr | 7.25 | 2.86 | Deleterious | | | | 7.25 | 2.86 | Deleterious |
| 21 | p.Ala21Cys | 1.05 | 0.07 | Neutral | | | | 1.05 | 0.07 | Neutral |
| 21 | p.Ala21Trp | 7.68 | 2.94 | Deleterious | | | | 7.68 | 2.94 | Deleterious |
| 21 | p.Ala21Phe | 6.02 | 2.59 | Deleterious | | | | 6.02 | 2.59 | Deleterious |
| 22 | p.Arg22Asn | 0.38 | -1.41 | Neutral | | | | 0.38 | -1.41 | Neutral |
| 22 | p.Arg22Lys | 0.99 | -0.01 | Neutral | | | | 0.99 | -0.01 | Neutral |
| 22 | p.Arg22Thr | 2.13 | 1.09 | Deleterious | | | | 2.13 | 1.09 | Deleterious |
| 22 | p.Arg22Arg | 1.00 | 0.00 | Neutral | | | | 1.00 | 0.00 | Neutral |
| 22 | p.Arg22Ser | 1.50 | 0.58 | Indeterminate | | | | 1.50 | 0.58 | Indeterminate |
| 22 | p.Arg22Ile | 0.86 | -0.21 | Neutral | | | | 0.86 | -0.21 | Neutral |
| 22 | p.Arg22Met | 1.40 | 0.49 | Indeterminate | | | | 1.40 | 0.49 | Indeterminate |
| 22 | p.Arg22His | 0.21 | -2.22 | Neutral | | | | 0.21 | -2.22 | Neutral |
| 22 | p.Arg22Gln | 0.39 | -1.36 | Neutral | | | | 0.39 | -1.36 | Neutral |
| 22 | p.Arg22Pro | 6.72 | 2.75 | Deleterious | | | | 6.72 | 2.75 | Deleterious |
| 22 | p.Arg22Leu | 0.54 | -0.89 | Neutral | | | | 0.54 | -0.89 | Neutral |
| 22 | p.Arg22Asp | 0.90 | -0.15 | Neutral | | | | 0.90 | -0.15 | Neutral |
| 22 | p.Arg22Glu | 0.31 | -1.70 | Neutral | | | | 0.31 | -1.70 | Neutral |
| 22 | p.Arg22Ala | 0.40 | -1.31 | Neutral | | | | 0.40 | -1.31 | Neutral |
| 22 | p.Arg22Gly | 3.32 | 1.73 | Deleterious | | | | 3.32 | 1.73 | Deleterious |
| 22 | p.Arg22Val | 0.51 | -0.97 | Neutral | | | | 0.51 | -0.97 | Neutral |
| 22 | p.Arg22Tyr | 0.97 | -0.04 | Neutral | | | | 0.97 | -0.04 | Neutral |
| 22 | p.Arg22Cys | 0.22 | -2.18 | Neutral | | | | 0.22 | -2.18 | Neutral |
| 22 | p.Arg22Trp | 1.58 | 0.66 | Indeterminate | | | | 1.58 | 0.66 | Indeterminate |
| 22 | p.Arg22Phe | 1.80 | 0.85 | Indeterminate | | | | 1.80 | 0.85 | Indeterminate |
| 23 | p.Gly23Asn | 2.46 | 1.30 | Deleterious | | | | 2.46 | 1.30 | Deleterious |
| 23 | p.Gly23Lys | 20.06 | 4.33 | Deleterious | | | | 20.06 | 4.33 | Deleterious |
| 23 | p.Gly23Thr | 20.30 | 4.34 | Deleterious | | | | 20.30 | 4.34 | Deleterious |
| 23 | p.Gly23Arg | 18.67 | 4.22 | Deleterious | | | | 18.67 | 4.22 | Deleterious |
| 23 | p.Gly23Ser | 3.96 | 1.99 | Deleterious | | | | 3.96 | 1.99 | Deleterious |
| 23 | p.Gly23Ile | 23.20 | 4.54 | Deleterious | | | | 23.20 | 4.54 | Deleterious |
| 23 | p.Gly23Met | 19.90 | 4.31 | Deleterious | | | | 19.90 | 4.31 | Deleterious |
| 23 | p.Gly23His | 18.89 | 4.24 | Deleterious | | | | 18.89 | 4.24 | Deleterious |
| 23 | p.Gly23Gln | 18.28 | 4.19 | Deleterious | | | | 18.28 | 4.19 | Deleterious |
| 23 | p.Gly23Pro | 19.40 | 4.28 | Deleterious | | | | 19.40 | 4.28 | Deleterious |
| 23 | p.Gly23Leu | 22.55 | 4.50 | Deleterious | | | | 22.55 | 4.50 | Deleterious |
| 23 | p.Gly23Asp | 14.29 | 3.84 | Deleterious | | | | 14.29 | 3.84 | Deleterious |
| 23 | p.Gly23Glu | 21.21 | 4.41 | Deleterious | | | | 21.21 | 4.41 | Deleterious |
| 23 | p.Gly23Ala | 1.27 | 0.35 | Indeterminate | | | | 1.27 | 0.35 | Indeterminate |
| 23 | p.Gly23Gly | 1.00 | 0.00 | Neutral | | | | 1.00 | 0.00 | Neutral |
| 23 | p.Gly23Val | 19.64 | 4.30 | Deleterious | | | | 19.64 | 4.30 | Deleterious |
| 23 | p.Gly23Tyr | 20.02 | 4.32 | Deleterious | | | | 20.02 | 4.32 | Deleterious |
| 23 | p.Gly23Cys | 13.02 | 3.70 | Deleterious | | | | 13.02 | 3.70 | Deleterious |
| 23 | p.Gly23Trp | 23.15 | 4.53 | Deleterious | | | | 23.15 | 4.53 | Deleterious |
| 23 | p.Gly23Phe | 20.81 | 4.38 | Deleterious | | | | 20.81 | 4.38 | Deleterious |
| 24 | p.Arg24Asn | 0.74 | -0.44 | Neutral | 1.02 | 0.02 | Neutral | 0.88 | -0.19 | Neutral |
| 24 | p.Arg24Lys | 0.60 | -0.75 | Neutral | 1.18 | 0.24 | Indeterminate | 0.89 | -0.17 | Neutral |
| 24 | p.Arg24Thr | 0.72 | -0.47 | Neutral | 1.28 | 0.36 | Indeterminate | 1.00 | 0.00 | Neutral |
| 24 | p.Arg24Arg | 1.00 | 0.00 | Neutral | 1.00 | 0.00 | Neutral | 1.00 | 0.00 | Neutral |
| 24 | p.Arg24Ser | 0.58 | -0.79 | Neutral | 0.98 | -0.03 | Neutral | 0.78 | -0.36 | Neutral |
| 24 | p.Arg24Ile | 0.76 | -0.40 | Neutral | 0.90 | -0.16 | Neutral | 0.83 | -0.27 | Neutral |
| 24 | p.Arg24Met | 0.90 | -0.15 | Neutral | 1.08 | 0.12 | Neutral | 0.99 | -0.01 | Neutral |
| 24 | p.Arg24His | 0.53 | -0.93 | Neutral | 0.83 | -0.27 | Neutral | 0.68 | -0.56 | Neutral |
| 24 | p.Arg24Gln | 1.53 | 0.62 | Indeterminate | 1.60 | 0.68 | Indeterminate | 1.57 | 0.65 | Indeterminate |
| 24 | p.Arg24Pro | 9.29 | 3.22 | Deleterious | 7.17 | 2.84 | Deleterious | 8.23 | 3.04 | Deleterious |
| 24 | p.Arg24Leu | 0.82 | -0.29 | Neutral | 1.06 | 0.08 | Neutral | 0.94 | -0.10 | Neutral |
| 24 | p.Arg24Asp | 0.74 | -0.43 | Neutral | 0.85 | -0.24 | Neutral | 0.79 | -0.33 | Neutral |
| 24 | p.Arg24Glu | 0.67 | -0.59 | Neutral | 1.14 | 0.18 | Indeterminate | 0.90 | -0.15 | Neutral |
| 24 | p.Arg24Ala | 0.53 | -0.91 | Neutral | 0.85 | -0.24 | Neutral | 0.69 | -0.54 | Neutral |
| 24 | p.Arg24Gly | 0.69 | -0.54 | Neutral | 1.23 | 0.30 | Indeterminate | 0.96 | -0.06 | Neutral |
| 24 | p.Arg24Val | 0.67 | -0.57 | Neutral | 1.02 | 0.02 | Neutral | 0.85 | -0.24 | Neutral |
| 24 | p.Arg24Tyr | 0.66 | -0.61 | Neutral | 1.14 | 0.19 | Indeterminate | 0.90 | -0.16 | Neutral |
| 24 | p.Arg24Cys | 0.84 | -0.25 | Neutral | 0.75 | -0.42 | Neutral | 0.79 | -0.33 | Neutral |
| 24 | p.Arg24Trp | 0.95 | -0.07 | Neutral | 1.12 | 0.16 | Indeterminate | 1.04 | 0.05 | Neutral |
| 24 | p.Arg24Phe | 0.82 | -0.29 | Neutral | 0.94 | -0.09 | Neutral | 0.88 | -0.19 | Neutral |
| 25 | p.Val25Asn | 1.23 | 0.30 | Indeterminate | | | | 1.23 | 0.30 | Indeterminate |
| 25 | p.Val25Lys | 1.86 | 0.89 | Indeterminate | | | | 1.86 | 0.89 | Indeterminate |
| 25 | p.Val25Thr | 1.00 | 0.00 | Neutral | | | | 1.00 | 0.00 | Neutral |
| 25 | p.Val25Arg | 1.37 | 0.45 | Indeterminate | | | | 1.37 | 0.45 | Indeterminate |
| 25 | p.Val25Ser | 0.88 | -0.19 | Neutral | | | | 0.88 | -0.19 | Neutral |
| 25 | p.Val25Ile | 0.83 | -0.27 | Neutral | | | | 0.83 | -0.27 | Neutral |
| 25 | p.Val25Met | 2.08 | 1.06 | Indeterminate | | | | 2.08 | 1.06 | Indeterminate |
| 25 | p.Val25His | 0.78 | -0.36 | Neutral | | | | 0.78 | -0.36 | Neutral |
| 25 | p.Val25Gln | 0.76 | -0.39 | Neutral | | | | 0.76 | -0.39 | Neutral |
| 25 | p.Val25Pro | 0.45 | -1.14 | Neutral | | | | 0.45 | -1.14 | Neutral |
| 25 | p.Val25Leu | 0.59 | -0.76 | Neutral | | | | 0.59 | -0.76 | Neutral |
| 25 | p.Val25Asp | 2.01 | 1.01 | Indeterminate | | | | 2.01 | 1.01 | Indeterminate |
| 25 | p.Val25Glu | 0.89 | -0.18 | Neutral | | | | 0.89 | -0.18 | Neutral |
| 25 | p.Val25Ala | 6.36 | 2.67 | Deleterious | | | | 6.36 | 2.67 | Deleterious |
| 25 | p.Val25Gly | 2.04 | 1.03 | Indeterminate | | | | 2.04 | 1.03 | Indeterminate |
| 25 | p.Val25Val | 1.00 | 0.00 | Neutral | | | | 1.00 | 0.00 | Neutral |
| 25 | p.Val25Tyr | 2.88 | 1.52 | Deleterious | | | | 2.88 | 1.52 | Deleterious |
| 25 | p.Val25Cys | 0.61 | -0.70 | Neutral | | | | 0.61 | -0.70 | Neutral |
| 25 | p.Val25Trp | 1.21 | 0.27 | Indeterminate | | | | 1.21 | 0.27 | Indeterminate |
| 25 | p.Val25Phe | 1.14 | 0.19 | Neutral | | | | 1.14 | 0.19 | Neutral |

| | | | | | | | | | | |
|---|---|---|---|---|---|---|---|---|---|---|
| 26 | p.Glu26Asn | 0.81 | -0.30 | Neutral | | | | 0.81 | -0.30 | Neutral |
| 26 | p.Glu26Lys | 0.82 | -0.29 | Neutral | | | | 0.82 | -0.29 | Neutral |
| 26 | p.Glu26Thr | 0.92 | -0.13 | Neutral | | | | 0.92 | -0.13 | Neutral |
| 26 | p.Glu26Arg | 0.90 | -0.16 | Neutral | | | | 0.90 | -0.16 | Neutral |
| 26 | p.Glu26Ser | 1.19 | 0.25 | Indeterminate | | | | 1.19 | 0.25 | Indeterminate |
| 26 | p.Glu26Ile | 0.95 | -0.07 | Neutral | | | | 0.95 | -0.07 | Neutral |
| 26 | p.Glu26Met | 0.86 | -0.21 | Neutral | | | | 0.86 | -0.21 | Neutral |
| 26 | p.Glu26His | 1.07 | 0.09 | Neutral | | | | 1.07 | 0.09 | Neutral |
| 26 | p.Glu26Gln | 1.05 | 0.07 | Neutral | | | | 1.05 | 0.07 | Neutral |
| 26 | p.Glu26Pro | 1.06 | 0.08 | Neutral | | | | 1.06 | 0.08 | Neutral |
| 26 | p.Glu26Leu | 0.84 | -0.25 | Neutral | | | | 0.84 | -0.25 | Neutral |
| 26 | p.Glu26Asp | 1.00 | 0.00 | Neutral | | | | 1.00 | 0.00 | Neutral |
| 26 | p.Glu26Glu | 1.00 | 0.00 | Neutral | | | | 1.00 | 0.00 | Neutral |
| 26 | p.Glu26Ala | 1.07 | 0.10 | Neutral | | | | 1.07 | 0.10 | Neutral |
| 26 | p.Glu26Gly | 1.00 | 0.01 | Neutral | | | | 1.00 | 0.01 | Neutral |
| 26 | p.Glu26Val | 0.98 | -0.03 | Neutral | | | | 0.98 | -0.03 | Neutral |
| 26 | p.Glu26Tyr | 1.12 | 0.16 | Neutral | | | | 1.12 | 0.16 | Neutral |
| 26 | p.Glu26Cys | 0.95 | -0.08 | Neutral | | | | 0.95 | -0.08 | Neutral |
| 26 | p.Glu26Trp | 0.99 | -0.01 | Neutral | | | | 0.99 | -0.01 | Neutral |
| 26 | p.Glu26Phe | 1.04 | 0.05 | Neutral | | | | 1.04 | 0.05 | Neutral |
| 27 | p.Glu27Asn | 0.36 | -1.47 | Neutral | 0.99 | -0.01 | Neutral | 0.68 | -0.56 | Neutral |
| 27 | p.Glu27Lys | 0.94 | -0.09 | Neutral | 0.99 | -0.02 | Neutral | 0.96 | -0.05 | Neutral |
| 27 | p.Glu27Thr | 1.69 | 0.76 | Indeterminate | 1.13 | 0.17 | Indeterminate | 1.41 | 0.50 | Indeterminate |
| 27 | p.Glu27Arg | 0.95 | -0.08 | Neutral | 1.09 | 0.13 | Neutral | 1.02 | 0.03 | Neutral |
| 27 | p.Glu27Ser | 0.76 | -0.40 | Neutral | 0.99 | -0.02 | Neutral | 0.87 | -0.20 | Neutral |
| 27 | p.Glu27Ile | 0.66 | -0.60 | Neutral | 1.13 | 0.17 | Indeterminate | 0.89 | -0.16 | Neutral |
| 27 | p.Glu27Met | 0.93 | -0.10 | Neutral | 1.05 | 0.06 | Neutral | 0.99 | -0.02 | Neutral |
| 27 | p.Glu27His | 0.89 | -0.17 | Neutral | 1.12 | 0.16 | Indeterminate | 1.00 | 0.01 | Neutral |
| 27 | p.Glu27Gln | 0.91 | -0.14 | Neutral | 1.07 | 0.10 | Neutral | 0.99 | -0.01 | Neutral |
| 27 | p.Glu27Pro | 1.16 | 0.22 | Neutral | 1.26 | 0.33 | Indeterminate | 1.21 | 0.27 | Indeterminate |
| 27 | p.Glu27Leu | 0.56 | -0.83 | Neutral | 1.03 | 0.04 | Neutral | 0.80 | -0.33 | Neutral |
| 27 | p.Glu27Asp | 0.78 | -0.36 | Neutral | 1.25 | 0.33 | Indeterminate | 1.02 | 0.02 | Neutral |
| 27 | p.Glu27Glu | 1.00 | 0.00 | Neutral | 1.00 | 0.00 | Neutral | 1.00 | 0.00 | Neutral |
| 27 | p.Glu27Ala | 0.84 | -0.26 | Neutral | 1.05 | 0.07 | Neutral | 0.94 | -0.09 | Neutral |
| 27 | p.Glu27Gly | 0.89 | -0.17 | Neutral | 0.97 | -0.05 | Neutral | 0.93 | -0.11 | Neutral |
| 27 | p.Glu27Val | 0.82 | -0.29 | Neutral | 1.08 | 0.11 | Neutral | 0.95 | -0.08 | Neutral |
| 27 | p.Glu27Tyr | 1.84 | 0.88 | Indeterminate | 0.95 | -0.07 | Neutral | 1.39 | 0.48 | Indeterminate |
| 27 | p.Glu27Cys | 0.86 | -0.22 | Neutral | 1.12 | 0.17 | Indeterminate | 0.99 | -0.01 | Neutral |
| 27 | p.Glu27Trp | 0.78 | -0.35 | Neutral | 0.98 | -0.03 | Neutral | 0.88 | -0.18 | Neutral |
| 27 | p.Glu27Phe | 0.61 | -0.71 | Neutral | 0.93 | -0.10 | Neutral | 0.77 | -0.37 | Neutral |
| 28 | p.Val28Asn | 5.13 | 2.36 | Deleterious | | | | 5.13 | 2.36 | Deleterious |
| 28 | p.Val28Lys | 7.05 | 2.82 | Deleterious | | | | 7.05 | 2.82 | Deleterious |
| 28 | p.Val28Thr | 0.75 | -0.41 | Neutral | | | | 0.75 | -0.41 | Neutral |
| 28 | p.Val28Arg | 6.34 | 2.66 | Deleterious | | | | 6.34 | 2.66 | Deleterious |
| 28 | p.Val28Ser | 1.06 | 0.08 | Neutral | | | | 1.06 | 0.08 | Neutral |
| 28 | p.Val28Ile | 0.79 | -0.34 | Neutral | | | | 0.79 | -0.34 | Neutral |
| 28 | p.Val28Met | 0.73 | -0.46 | Neutral | | | | 0.73 | -0.46 | Neutral |
| 28 | p.Val28His | 6.67 | 2.74 | Deleterious | | | | 6.67 | 2.74 | Deleterious |
| 28 | p.Val28Gln | 4.46 | 2.16 | Deleterious | | | | 4.46 | 2.16 | Deleterious |
| 28 | p.Val28Pro | 6.33 | 2.66 | Deleterious | | | | 6.33 | 2.66 | Deleterious |
| 28 | p.Val28Leu | 0.60 | -0.73 | Neutral | | | | 0.60 | -0.73 | Neutral |
| 28 | p.Val28Asp | 6.27 | 2.65 | Deleterious | | | | 6.27 | 2.65 | Deleterious |
| 28 | p.Val28Glu | 4.08 | 2.03 | Deleterious | | | | 4.08 | 2.03 | Deleterious |
| 28 | p.Val28Ala | 0.72 | -0.48 | Neutral | | | | 0.72 | -0.48 | Neutral |
| 28 | p.Val28Gly | 1.76 | 0.81 | Indeterminate | | | | 1.76 | 0.81 | Indeterminate |
| 28 | p.Val28Val | 1.00 | 0.00 | Neutral | | | | 1.00 | 0.00 | Neutral |
| 28 | p.Val28Tyr | 6.56 | 2.71 | Deleterious | | | | 6.56 | 2.71 | Deleterious |
| 28 | p.Val28Cys | 0.70 | -0.51 | Neutral | | | | 0.70 | -0.51 | Neutral |
| 28 | p.Val28Trp | 7.13 | 2.83 | Deleterious | | | | 7.13 | 2.83 | Deleterious |
| 28 | p.Val28Phe | 4.68 | 2.23 | Deleterious | | | | 4.68 | 2.23 | Deleterious |
| 29 | p.Arg29Asn | 1.37 | 0.45 | Indeterminate | | | | 1.37 | 0.45 | Indeterminate |
| 29 | p.Arg29Lys | 0.86 | -0.22 | Neutral | | | | 0.86 | -0.22 | Neutral |
| 29 | p.Arg29Thr | 1.46 | 0.55 | Indeterminate | | | | 1.46 | 0.55 | Indeterminate |
| 29 | p.Arg29Arg | 1.00 | 0.00 | Neutral | | | | 1.00 | 0.00 | Neutral |
| 29 | p.Arg29Ser | 1.19 | 0.25 | Indeterminate | | | | 1.19 | 0.25 | Indeterminate |
| 29 | p.Arg29Ile | 1.91 | 0.93 | Indeterminate | | | | 1.91 | 0.93 | Indeterminate |
| 29 | p.Arg29Met | 1.05 | 0.08 | Neutral | | | | 1.05 | 0.08 | Neutral |
| 29 | p.Arg29His | 0.93 | -0.11 | Neutral | | | | 0.93 | -0.11 | Neutral |
| 29 | p.Arg29Gln | 0.88 | -0.18 | Neutral | | | | 0.88 | -0.18 | Neutral |
| 29 | p.Arg29Pro | 15.53 | 3.96 | Deleterious | | | | 15.53 | 3.96 | Deleterious |
| 29 | p.Arg29Leu | 1.01 | 0.01 | Neutral | | | | 1.01 | 0.01 | Neutral |
| 29 | p.Arg29Asp | 1.88 | 0.91 | Indeterminate | | | | 1.88 | 0.91 | Indeterminate |
| 29 | p.Arg29Glu | 1.21 | 0.28 | Indeterminate | | | | 1.21 | 0.28 | Indeterminate |
| 29 | p.Arg29Ala | 0.65 | -0.62 | Neutral | | | | 0.65 | -0.62 | Neutral |
| 29 | p.Arg29Gly | 1.12 | 0.16 | Neutral | | | | 1.12 | 0.16 | Neutral |
| 29 | p.Arg29Val | 0.93 | -0.11 | Neutral | | | | 0.93 | -0.11 | Neutral |
| 29 | p.Arg29Tyr | 0.87 | -0.20 | Neutral | | | | 0.87 | -0.20 | Neutral |
| 29 | p.Arg29Cys | 1.09 | 0.12 | Neutral | | | | 1.09 | 0.12 | Neutral |
| 29 | p.Arg29Trp | 0.80 | -0.32 | Neutral | | | | 0.80 | -0.32 | Neutral |
| 29 | p.Arg29Phe | 0.87 | -0.20 | Neutral | | | | 0.87 | -0.20 | Neutral |
| 30 | p.Ala30Asn | 3.67 | 1.87 | Deleterious | | | | 3.67 | 1.87 | Deleterious |
| 30 | p.Ala30Lys | 0.90 | -0.15 | Neutral | | | | 0.90 | -0.15 | Neutral |
| 30 | p.Ala30Thr | 2.02 | 1.02 | Indeterminate | | | | 2.02 | 1.02 | Indeterminate |
| 30 | p.Ala30Arg | 1.23 | 0.30 | Indeterminate | | | | 1.23 | 0.30 | Indeterminate |
| 30 | p.Ala30Ser | 1.49 | 0.57 | Indeterminate | | | | 1.49 | 0.57 | Indeterminate |
| 30 | p.Ala30Ile | 1.39 | 0.47 | Indeterminate | | | | 1.39 | 0.47 | Indeterminate |
| 30 | p.Ala30Met | 1.37 | 0.45 | Indeterminate | | | | 1.37 | 0.45 | Indeterminate |
| 30 | p.Ala30His | 1.26 | 0.34 | Indeterminate | | | | 1.26 | 0.34 | Indeterminate |
| 30 | p.Ala30Gln | 1.46 | 0.54 | Indeterminate | | | | 1.46 | 0.54 | Indeterminate |
| 30 | p.Ala30Pro | 2.91 | 1.54 | Deleterious | | | | 2.91 | 1.54 | Deleterious |
| 30 | p.Ala30Leu | 1.28 | 0.36 | Indeterminate | | | | 1.28 | 0.36 | Indeterminate |
| 30 | p.Ala30Asp | 2.05 | 1.04 | Indeterminate | | | | 2.05 | 1.04 | Indeterminate |
| 30 | p.Ala30Glu | 1.06 | 0.08 | Neutral | | | | 1.06 | 0.08 | Neutral |
| 30 | p.Ala30Ala | 1.00 | 0.00 | Neutral | | | | 1.00 | 0.00 | Neutral |
| 30 | p.Ala30Gly | 3.33 | 1.74 | Deleterious | | | | 3.33 | 1.74 | Deleterious |
| 30 | p.Ala30Val | 1.56 | 0.64 | Indeterminate | | | | 1.56 | 0.64 | Indeterminate |
| 30 | p.Ala30Tyr | 1.81 | 0.86 | Indeterminate | | | | 1.81 | 0.86 | Indeterminate |
| 30 | p.Ala30Cys | 1.67 | 0.74 | Indeterminate | | | | 1.67 | 0.74 | Indeterminate |
| 30 | p.Ala30Trp | 1.75 | 0.81 | Indeterminate | | | | 1.75 | 0.81 | Indeterminate |
| 30 | p.Ala30Phe | 1.22 | 0.28 | Indeterminate | | | | 1.22 | 0.28 | Indeterminate |
| 31 | p.Leu31Asn | 1.51 | 0.60 | Indeterminate | | | | 1.51 | 0.60 | Indeterminate |

| | | | | | | | | | | |
|---|---|---|---|---|---|---|---|---|---|---|
| 31 | p.Leu31Lys | 0.99 | -0.01 | Neutral | | | | 0.99 | -0.01 | Neutral |
| 31 | p.Leu31Thr | 1.18 | 0.23 | Neutral | | | | 1.18 | 0.23 | Neutral |
| 31 | p.Leu31Arg | 1.05 | 0.07 | Neutral | | | | 1.05 | 0.07 | Neutral |
| 31 | p.Leu31Ser | 1.41 | 0.50 | Indeterminate | | | | 1.41 | 0.50 | Indeterminate |
| 31 | p.Leu31Ile | 1.00 | 0.01 | Neutral | | | | 1.00 | 0.01 | Neutral |
| 31 | p.Leu31Met | 1.36 | 0.44 | Indeterminate | | | | 1.36 | 0.44 | Indeterminate |
| 31 | p.Leu31His | 1.06 | 0.08 | Neutral | | | | 1.06 | 0.08 | Neutral |
| 31 | p.Leu31Gln | 1.68 | 0.75 | Indeterminate | | | | 1.68 | 0.75 | Indeterminate |
| 31 | p.Leu31Pro | 12.45 | 3.64 | Deleterious | | | | 12.45 | 3.64 | Deleterious |
| 31 | p.Leu31Leu | 1.00 | 0.00 | Neutral | | | | 1.00 | 0.00 | Neutral |
| 31 | p.Leu31Asp | 3.61 | 1.85 | Deleterious | | | | 3.61 | 1.85 | Deleterious |
| 31 | p.Leu31Glu | 1.18 | 0.23 | Neutral | | | | 1.18 | 0.23 | Neutral |
| 31 | p.Leu31Ala | 1.30 | 0.38 | Indeterminate | | | | 1.30 | 0.38 | Indeterminate |
| 31 | p.Leu31Gly | 1.17 | 0.22 | Neutral | | | | 1.17 | 0.22 | Neutral |
| 31 | p.Leu31Val | 1.62 | 0.70 | Indeterminate | | | | 1.62 | 0.70 | Indeterminate |
| 31 | p.Leu31Tyr | 1.97 | 0.98 | Indeterminate | | | | 1.97 | 0.98 | Indeterminate |
| 31 | p.Leu31Cys | 1.81 | 0.86 | Indeterminate | | | | 1.81 | 0.86 | Indeterminate |
| 31 | p.Leu31Trp | 1.45 | 0.54 | Indeterminate | | | | 1.45 | 0.54 | Indeterminate |
| 31 | p.Leu31Phe | 1.04 | 0.05 | Neutral | | | | 1.04 | 0.05 | Neutral |
| 32 | p.Leu32Asn | 2.33 | 1.22 | Deleterious | 2.62 | 1.39 | Indeterminate | 2.48 | 1.31 | Deleterious |
| 32 | p.Leu32Lys | 2.83 | 1.50 | Deleterious | 2.85 | 1.51 | Indeterminate | 2.84 | 1.51 | Deleterious |
| 32 | p.Leu32Thr | 1.82 | 0.87 | Indeterminate | 0.74 | -0.43 | Neutral | 1.28 | 0.36 | Indeterminate |
| 32 | p.Leu32Arg | 2.57 | 1.36 | Deleterious | 2.93 | 1.55 | Indeterminate | 2.75 | 1.46 | Deleterious |
| 32 | p.Leu32Ser | 2.44 | 1.29 | Deleterious | 1.52 | 0.60 | Indeterminate | 1.98 | 0.98 | Indeterminate |
| 32 | p.Leu32Ile | 0.90 | -0.15 | Neutral | 0.56 | -0.84 | Neutral | 0.73 | -0.45 | Neutral |
| 32 | p.Leu32Met | 1.12 | 0.17 | Neutral | 0.55 | -0.87 | Neutral | 0.84 | -0.26 | Neutral |
| 32 | p.Leu32His | 2.72 | 1.45 | Deleterious | 2.48 | 1.31 | Indeterminate | 2.60 | 1.38 | Deleterious |
| 32 | p.Leu32Gln | 1.65 | 0.72 | Indeterminate | 1.57 | 0.65 | Indeterminate | 1.61 | 0.69 | Indeterminate |
| 32 | p.Leu32Pro | 3.00 | 1.58 | Deleterious | 3.40 | 1.76 | Deleterious | 3.20 | 1.68 | Deleterious |
| 32 | p.Leu32Leu | 1.00 | 0.00 | Neutral | 1.00 | 0.00 | Neutral | 1.00 | 0.00 | Neutral |
| 32 | p.Leu32Asp | 2.58 | 1.37 | Deleterious | 3.30 | 1.72 | Indeterminate | 2.94 | 1.56 | Deleterious |
| 32 | p.Leu32Glu | 2.90 | 1.53 | Deleterious | 2.82 | 1.50 | Indeterminate | 2.86 | 1.51 | Deleterious |
| 32 | p.Leu32Ala | 1.13 | 0.18 | Neutral | 1.16 | 0.21 | Indeterminate | 1.15 | 0.20 | Neutral |
| 32 | p.Leu32Gly | 2.72 | 1.44 | Deleterious | 2.47 | 1.30 | Indeterminate | 2.59 | 1.38 | Deleterious |
| 32 | p.Leu32Val | 1.08 | 0.11 | Neutral | 0.57 | -0.80 | Neutral | 0.83 | -0.28 | Neutral |
| 32 | p.Leu32Tyr | 2.24 | 1.16 | Deleterious | 2.50 | 1.32 | Indeterminate | 2.37 | 1.24 | Deleterious |
| 32 | p.Leu32Cys | 1.20 | 0.26 | Indeterminate | 0.51 | -0.98 | Neutral | 0.85 | -0.23 | Neutral |
| 32 | p.Leu32Trp | 2.64 | 1.40 | Deleterious | 2.79 | 1.48 | Indeterminate | 2.71 | 1.44 | Deleterious |
| 32 | p.Leu32Phe | 1.36 | 0.45 | Indeterminate | 0.72 | -0.47 | Neutral | 1.04 | 0.06 | Neutral |
| 33 | p.Glu33Asn | 0.89 | -0.16 | Neutral | | | | 0.89 | -0.16 | Neutral |
| 33 | p.Glu33Lys | 0.99 | -0.01 | Neutral | | | | 0.99 | -0.01 | Neutral |
| 33 | p.Glu33Thr | 0.95 | -0.07 | Neutral | | | | 0.95 | -0.07 | Neutral |
| 33 | p.Glu33Arg | 0.89 | -0.17 | Neutral | | | | 0.89 | -0.17 | Neutral |
| 33 | p.Glu33Ser | 1.00 | 0.00 | Neutral | | | | 1.00 | 0.00 | Neutral |
| 33 | p.Glu33Ile | 0.89 | -0.16 | Neutral | | | | 0.89 | -0.16 | Neutral |
| 33 | p.Glu33Met | 0.88 | -0.18 | Neutral | | | | 0.88 | -0.18 | Neutral |
| 33 | p.Glu33His | 0.96 | -0.06 | Neutral | | | | 0.96 | -0.06 | Neutral |
| 33 | p.Glu33Gln | 0.99 | -0.02 | Neutral | | | | 0.99 | -0.02 | Neutral |
| 33 | p.Glu33Pro | 0.82 | -0.29 | Neutral | | | | 0.82 | -0.29 | Neutral |
| 33 | p.Glu33Leu | 1.03 | 0.05 | Neutral | | | | 1.03 | 0.05 | Neutral |
| 33 | p.Glu33Asp | 0.84 | -0.25 | Neutral | | | | 0.84 | -0.25 | Neutral |
| 33 | p.Glu33Glu | 1.00 | 0.00 | Neutral | | | | 1.00 | 0.00 | Neutral |
| 33 | p.Glu33Ala | 1.03 | 0.04 | Neutral | | | | 1.03 | 0.04 | Neutral |
| 33 | p.Glu33Gly | 0.95 | -0.08 | Neutral | | | | 0.95 | -0.08 | Neutral |
| 33 | p.Glu33Val | 0.93 | -0.11 | Neutral | | | | 0.93 | -0.11 | Neutral |
| 33 | p.Glu33Tyr | 0.93 | -0.10 | Neutral | | | | 0.93 | -0.10 | Neutral |
| 33 | p.Glu33Cys | 0.98 | -0.03 | Neutral | | | | 0.98 | -0.03 | Neutral |
| 33 | p.Glu33Trp | 0.90 | -0.16 | Neutral | | | | 0.90 | -0.16 | Neutral |
| 33 | p.Glu33Phe | 0.95 | -0.07 | Neutral | | | | 0.95 | -0.07 | Neutral |
| 34 | p.Ala34Asn | 1.01 | 0.01 | Neutral | 1.20 | 0.26 | Indeterminate | 1.10 | 0.14 | Neutral |
| 34 | p.Ala34Lys | 1.01 | 0.01 | Neutral | 0.96 | -0.06 | Neutral | 0.98 | -0.03 | Neutral |
| 34 | p.Ala34Thr | 0.83 | -0.27 | Neutral | 1.12 | 0.16 | Indeterminate | 0.98 | -0.04 | Neutral |
| 34 | p.Ala34Arg | 0.93 | -0.10 | Neutral | 0.95 | -0.08 | Neutral | 0.94 | -0.09 | Neutral |
| 34 | p.Ala34Ser | 0.68 | -0.55 | Neutral | 0.96 | -0.06 | Neutral | 0.82 | -0.28 | Neutral |
| 34 | p.Ala34Ile | 0.83 | -0.26 | Neutral | 0.82 | -0.29 | Neutral | 0.83 | -0.28 | Neutral |
| 34 | p.Ala34Met | 1.09 | 0.12 | Neutral | 0.88 | -0.18 | Neutral | 0.99 | -0.02 | Neutral |
| 34 | p.Ala34His | 0.71 | -0.49 | Neutral | 0.82 | -0.28 | Neutral | 0.77 | -0.38 | Neutral |
| 34 | p.Ala34Gln | 1.12 | 0.16 | Neutral | 1.16 | 0.22 | Indeterminate | 1.14 | 0.19 | Neutral |
| 34 | p.Ala34Pro | 1.45 | 0.54 | Indeterminate | 1.15 | 0.20 | Indeterminate | 1.30 | 0.38 | Indeterminate |
| 34 | p.Ala34Leu | 0.96 | -0.06 | Neutral | 0.99 | -0.01 | Neutral | 0.97 | -0.04 | Neutral |
| 34 | p.Ala34Asp | 1.13 | 0.18 | Neutral | 0.90 | -0.14 | Neutral | 1.02 | 0.03 | Neutral |
| 34 | p.Ala34Glu | 1.19 | 0.25 | Indeterminate | 0.99 | -0.01 | Neutral | 1.09 | 0.12 | Neutral |
| 34 | p.Ala34Ala | 1.00 | 0.00 | Neutral | 1.00 | 0.00 | Neutral | 1.00 | 0.00 | Neutral |
| 34 | p.Ala34Gly | 1.19 | 0.25 | Indeterminate | 0.81 | -0.30 | Neutral | 1.00 | 0.00 | Neutral |
| 34 | p.Ala34Val | 1.34 | 0.42 | Indeterminate | 0.92 | -0.12 | Neutral | 1.13 | 0.17 | Neutral |
| 34 | p.Ala34Tyr | 0.75 | -0.41 | Neutral | 1.00 | 0.01 | Neutral | 0.88 | -0.19 | Neutral |
| 34 | p.Ala34Cys | 0.98 | -0.04 | Neutral | 0.78 | -0.35 | Neutral | 0.88 | -0.19 | Neutral |
| 34 | p.Ala34Trp | 0.99 | -0.02 | Neutral | 0.88 | -0.19 | Neutral | 0.93 | -0.10 | Neutral |
| 34 | p.Ala34Phe | 1.06 | 0.09 | Neutral | 0.87 | -0.20 | Neutral | 0.97 | -0.05 | Neutral |
| 35 | p.Gly35Asn | 1.69 | 0.76 | Indeterminate | 1.30 | 0.37 | Indeterminate | 1.49 | 0.58 | Indeterminate |
| 35 | p.Gly35Lys | 2.04 | 1.03 | Indeterminate | 1.61 | 0.69 | Indeterminate | 1.83 | 0.87 | Indeterminate |
| 35 | p.Gly35Thr | 2.63 | 1.40 | Deleterious | 2.16 | 1.11 | Indeterminate | 2.40 | 1.26 | Deleterious |
| 35 | p.Gly35Arg | 1.40 | 0.48 | Indeterminate | 1.46 | 0.55 | Indeterminate | 1.43 | 0.51 | Indeterminate |
| 35 | p.Gly35Ser | 1.93 | 0.95 | Indeterminate | 1.53 | 0.62 | Indeterminate | 1.73 | 0.79 | Indeterminate |
| 35 | p.Gly35Ile | 8.80 | 3.14 | Deleterious | 4.34 | 2.12 | Deleterious | 6.57 | 2.72 | Deleterious |
| 35 | p.Gly35Met | 2.40 | 1.26 | Deleterious | 2.07 | 1.05 | Indeterminate | 2.23 | 1.16 | Deleterious |
| 35 | p.Gly35His | 1.45 | 0.53 | Indeterminate | 1.48 | 0.56 | Indeterminate | 1.46 | 0.55 | Indeterminate |
| 35 | p.Gly35Gln | 1.94 | 0.95 | Indeterminate | 1.98 | 0.99 | Indeterminate | 1.96 | 0.97 | Indeterminate |
| 35 | p.Gly35Pro | 14.31 | 3.84 | Deleterious | 6.09 | 2.61 | Deleterious | 10.20 | 3.35 | Deleterious |
| 35 | p.Gly35Leu | 3.89 | 1.96 | Deleterious | 2.62 | 1.39 | Indeterminate | 3.25 | 1.70 | Deleterious |
| 35 | p.Gly35Asp | 2.28 | 1.19 | Deleterious | 1.52 | 0.61 | Indeterminate | 1.90 | 0.93 | Indeterminate |
| 35 | p.Gly35Glu | 1.73 | 0.79 | Indeterminate | 1.81 | 0.85 | Indeterminate | 1.77 | 0.82 | Indeterminate |
| 35 | p.Gly35Ala | 1.63 | 0.71 | Indeterminate | 1.52 | 0.61 | Indeterminate | 1.58 | 0.66 | Indeterminate |
| 35 | p.Gly35Gly | 1.00 | 0.00 | Neutral | 1.00 | 0.00 | Neutral | 1.00 | 0.00 | Neutral |
| 35 | p.Gly35Val | 6.32 | 2.66 | Deleterious | 3.44 | 1.78 | Deleterious | 4.88 | 2.29 | Deleterious |
| 35 | p.Gly35Tyr | 2.55 | 1.35 | Deleterious | 1.58 | 0.66 | Indeterminate | 2.06 | 1.05 | Indeterminate |
| 35 | p.Gly35Cys | 2.12 | 1.08 | Indeterminate | 1.65 | 0.72 | Indeterminate | 1.89 | 0.91 | Indeterminate |
| 35 | p.Gly35Trp | 3.69 | 1.88 | Deleterious | 2.44 | 1.29 | Indeterminate | 3.06 | 1.62 | Deleterious |
| 35 | p.Gly35Phe | 1.96 | 0.97 | Indeterminate | 1.64 | 0.71 | Indeterminate | 1.80 | 0.84 | Indeterminate |
| 36 | p.Ala36Asn | 2.15 | 1.11 | Deleterious | | | | 2.15 | 1.11 | Deleterious |
| 36 | p.Ala36Lys | 8.35 | 3.06 | Deleterious | | | | 8.35 | 3.06 | Deleterious |

| | | | | | | | |
|---|---|---|---|---|---|---|---|
| 36 | p.Ala36Thr | 1.54 | 0.62 | Indeterminate | 1.54 | 0.62 | Indeterminate |
| 36 | p.Ala36Arg | 0.98 | -0.03 | Neutral | 0.98 | -0.03 | Neutral |
| 36 | p.Ala36Ser | 1.20 | 0.26 | Indeterminate | 1.20 | 0.26 | Indeterminate |
| 36 | p.Ala36Ile | 0.85 | -0.23 | Neutral | 0.85 | -0.23 | Neutral |
| 36 | p.Ala36Met | 0.13 | -2.96 | Neutral | 0.13 | -2.96 | Neutral |
| 36 | p.Ala36His | 0.75 | -0.41 | Neutral | 0.75 | -0.41 | Neutral |
| 36 | p.Ala36Gln | 0.81 | -0.30 | Neutral | 0.81 | -0.30 | Neutral |
| 36 | p.Ala36Pro | 4.00 | 2.00 | Deleterious | 4.00 | 2.00 | Deleterious |
| 36 | p.Ala36Leu | 1.55 | 0.63 | Indeterminate | 1.55 | 0.63 | Indeterminate |
| 36 | p.Ala36Asp | 1.15 | 0.21 | Neutral | 1.15 | 0.21 | Neutral |
| 36 | p.Ala36Glu | 1.67 | 0.74 | Indeterminate | 1.67 | 0.74 | Indeterminate |
| 36 | p.Ala36Ala | 1.00 | 0.00 | Neutral | 1.00 | 0.00 | Neutral |
| 36 | p.Ala36Gly | 1.34 | 0.42 | Indeterminate | 1.34 | 0.42 | Indeterminate |
| 36 | p.Ala36Val | 0.82 | -0.28 | Neutral | 0.82 | -0.28 | Neutral |
| 36 | p.Ala36Tyr | 1.32 | 0.40 | Indeterminate | 1.32 | 0.40 | Indeterminate |
| 36 | p.Ala36Cys | 0.23 | -2.15 | Neutral | 0.23 | -2.15 | Neutral |
| 36 | p.Ala36Trp | 0.65 | -0.61 | Neutral | 0.65 | -0.61 | Neutral |
| 36 | p.Ala36Phe | 1.23 | 0.30 | Indeterminate | 1.23 | 0.30 | Indeterminate |
| 37 | p.Leu37Asn | 0.78 | -0.35 | Neutral | 0.78 | -0.35 | Neutral |
| 37 | p.Leu37Lys | 1.29 | 0.37 | Indeterminate | 1.29 | 0.37 | Indeterminate |
| 37 | p.Leu37Thr | 1.40 | 0.49 | Indeterminate | 1.40 | 0.49 | Indeterminate |
| 37 | p.Leu37Arg | 0.43 | -1.23 | Neutral | 0.43 | -1.23 | Neutral |
| 37 | p.Leu37Ser | 0.98 | -0.03 | Neutral | 0.98 | -0.03 | Neutral |
| 37 | p.Leu37Ile | 1.73 | 0.79 | Indeterminate | 1.73 | 0.79 | Indeterminate |
| 37 | p.Leu37Met | 0.64 | -0.65 | Neutral | 0.64 | -0.65 | Neutral |
| 37 | p.Leu37His | 1.39 | 0.48 | Indeterminate | 1.39 | 0.48 | Indeterminate |
| 37 | p.Leu37Gln | 1.66 | 0.73 | Indeterminate | 1.66 | 0.73 | Indeterminate |
| 37 | p.Leu37Pro | 1.90 | 0.92 | Indeterminate | 1.90 | 0.92 | Indeterminate |
| 37 | p.Leu37Leu | 1.00 | 0.00 | Neutral | 1.00 | 0.00 | Neutral |
| 37 | p.Leu37Asp | 1.04 | 0.05 | Neutral | 1.04 | 0.05 | Neutral |
| 37 | p.Leu37Glu | 1.01 | 0.01 | Neutral | 1.01 | 0.01 | Neutral |
| 37 | p.Leu37Ala | 0.74 | -0.44 | Neutral | 0.74 | -0.44 | Neutral |
| 37 | p.Leu37Gly | 0.96 | -0.06 | Neutral | 0.96 | -0.06 | Neutral |
| 37 | p.Leu37Val | 1.51 | 0.60 | Indeterminate | 1.51 | 0.60 | Indeterminate |
| 37 | p.Leu37Tyr | 1.25 | 0.32 | Indeterminate | 1.25 | 0.32 | Indeterminate |
| 37 | p.Leu37Cys | 0.84 | -0.24 | Neutral | 0.84 | -0.24 | Neutral |
| 37 | p.Leu37Trp | 6.84 | 2.77 | Deleterious | 6.84 | 2.77 | Deleterious |
| 37 | p.Leu37Phe | 0.77 | -0.37 | Neutral | 0.77 | -0.37 | Neutral |
| 38 | p.Pro38Asn | 3.11 | 1.64 | Deleterious | 3.11 | 1.64 | Deleterious |
| 38 | p.Pro38Lys | 3.68 | 1.88 | Deleterious | 3.68 | 1.88 | Deleterious |
| 38 | p.Pro38Thr | 1.34 | 0.42 | Indeterminate | 1.34 | 0.42 | Indeterminate |
| 38 | p.Pro38Arg | 5.04 | 2.33 | Deleterious | 5.04 | 2.33 | Deleterious |
| 38 | p.Pro38Ser | 1.73 | 0.79 | Indeterminate | 1.73 | 0.79 | Indeterminate |
| 38 | p.Pro38Ile | 1.77 | 0.82 | Indeterminate | 1.77 | 0.82 | Indeterminate |
| 38 | p.Pro38Met | 2.30 | 1.20 | Deleterious | 2.30 | 1.20 | Deleterious |
| 38 | p.Pro38His | 3.80 | 1.92 | Deleterious | 3.80 | 1.92 | Deleterious |
| 38 | p.Pro38Gln | 4.08 | 2.03 | Deleterious | 4.08 | 2.03 | Deleterious |
| 38 | p.Pro38Pro | 1.00 | 0.00 | Neutral | 1.00 | 0.00 | Neutral |
| 38 | p.Pro38Leu | 3.22 | 1.69 | Deleterious | 3.22 | 1.69 | Deleterious |
| 38 | p.Pro38Asp | 20.44 | 4.35 | Deleterious | 20.44 | 4.35 | Deleterious |
| 38 | p.Pro38Glu | 2.65 | 1.41 | Deleterious | 2.65 | 1.41 | Deleterious |
| 38 | p.Pro38Ala | 5.59 | 2.48 | Deleterious | 5.59 | 2.48 | Deleterious |
| 38 | p.Pro38Gly | 0.99 | -0.02 | Neutral | 0.99 | -0.02 | Neutral |
| 38 | p.Pro38Val | 1.02 | 0.03 | Neutral | 1.02 | 0.03 | Neutral |
| 38 | p.Pro38Tyr | 3.40 | 1.77 | Deleterious | 3.40 | 1.77 | Deleterious |
| 38 | p.Pro38Cys | 0.75 | -0.41 | Neutral | 0.75 | -0.41 | Neutral |
| 38 | p.Pro38Trp | 31.11 | 4.96 | Deleterious | 31.11 | 4.96 | Deleterious |
| 38 | p.Pro38Phe | 4.51 | 2.17 | Deleterious | 4.51 | 2.17 | Deleterious |
| 39 | p.Asn39Asn | 1.00 | 0.00 | Neutral | 1.00 | 0.00 | Neutral |
| 39 | p.Asn39Lys | 2.13 | 1.09 | Deleterious | 2.13 | 1.09 | Deleterious |
| 39 | p.Asn39Thr | 0.63 | -0.66 | Neutral | 0.63 | -0.66 | Neutral |
| 39 | p.Asn39Arg | 1.12 | 0.16 | Neutral | 1.12 | 0.16 | Neutral |
| 39 | p.Asn39Ser | 1.17 | 0.22 | Neutral | 1.17 | 0.22 | Neutral |
| 39 | p.Asn39Ile | 1.45 | 0.53 | Indeterminate | 1.45 | 0.53 | Indeterminate |
| 39 | p.Asn39Met | 2.20 | 1.14 | Deleterious | 2.20 | 1.14 | Deleterious |
| 39 | p.Asn39His | 0.71 | -0.50 | Neutral | 0.71 | -0.50 | Neutral |
| 39 | p.Asn39Gln | 0.93 | -0.10 | Neutral | 0.93 | -0.10 | Neutral |
| 39 | p.Asn39Pro | 14.46 | 3.85 | Deleterious | 14.46 | 3.85 | Deleterious |
| 39 | p.Asn39Leu | 4.17 | 2.06 | Deleterious | 4.17 | 2.06 | Deleterious |
| 39 | p.Asn39Asp | 1.04 | 0.06 | Neutral | 1.04 | 0.06 | Neutral |
| 39 | p.Asn39Glu | 1.18 | 0.24 | Indeterminate | 1.18 | 0.24 | Indeterminate |
| 39 | p.Asn39Ala | 1.30 | 0.37 | Indeterminate | 1.30 | 0.37 | Indeterminate |
| 39 | p.Asn39Gly | 0.98 | -0.03 | Neutral | 0.98 | -0.03 | Neutral |
| 39 | p.Asn39Val | 1.33 | 0.41 | Indeterminate | 1.33 | 0.41 | Indeterminate |
| 39 | p.Asn39Tyr | 0.89 | -0.17 | Neutral | 0.89 | -0.17 | Neutral |
| 39 | p.Asn39Cys | 4.55 | 2.19 | Deleterious | 4.55 | 2.19 | Deleterious |
| 39 | p.Asn39Trp | 1.64 | 0.71 | Indeterminate | 1.64 | 0.71 | Indeterminate |
| 39 | p.Asn39Phe | 8.47 | 3.08 | Deleterious | 8.47 | 3.08 | Deleterious |
| 40 | p.Ala40Asn | 1.37 | 0.45 | Indeterminate | 1.37 | 0.45 | Indeterminate |
| 40 | p.Ala40Lys | 1.11 | 0.15 | Neutral | 1.11 | 0.15 | Neutral |
| 40 | p.Ala40Thr | 7.75 | 2.95 | Deleterious | 7.75 | 2.95 | Deleterious |
| 40 | p.Ala40Arg | 1.00 | 0.00 | Neutral | 1.00 | 0.00 | Neutral |
| 40 | p.Ala40Ser | 0.73 | -0.46 | Neutral | 0.73 | -0.46 | Neutral |
| 40 | p.Ala40Ile | 1.12 | 0.16 | Neutral | 1.12 | 0.16 | Neutral |
| 40 | p.Ala40Met | 0.60 | -0.73 | Neutral | 0.60 | -0.73 | Neutral |
| 40 | p.Ala40His | 0.74 | -0.44 | Neutral | 0.74 | -0.44 | Neutral |
| 40 | p.Ala40Gln | 0.61 | -0.72 | Neutral | 0.61 | -0.72 | Neutral |
| 40 | p.Ala40Pro | 1.50 | 0.58 | Indeterminate | 1.50 | 0.58 | Indeterminate |
| 40 | p.Ala40Leu | 3.93 | 1.97 | Deleterious | 3.93 | 1.97 | Deleterious |
| 40 | p.Ala40Asp | 1.66 | 0.73 | Indeterminate | 1.66 | 0.73 | Indeterminate |
| 40 | p.Ala40Glu | 0.79 | -0.35 | Neutral | 0.79 | -0.35 | Neutral |
| 40 | p.Ala40Ala | 1.00 | 0.00 | Neutral | 1.00 | 0.00 | Neutral |
| 40 | p.Ala40Gly | 1.03 | 0.05 | Neutral | 1.03 | 0.05 | Neutral |
| 40 | p.Ala40Val | 1.38 | 0.47 | Indeterminate | 1.38 | 0.47 | Indeterminate |
| 40 | p.Ala40Tyr | 0.73 | -0.44 | Neutral | 0.73 | -0.44 | Neutral |
| 40 | p.Ala40Cys | 1.01 | 0.01 | Neutral | 1.01 | 0.01 | Neutral |
| 40 | p.Ala40Trp | 1.17 | 0.22 | Neutral | 1.17 | 0.22 | Neutral |
| 40 | p.Ala40Phe | 1.12 | 0.16 | Neutral | 1.12 | 0.16 | Neutral |
| 41 | p.Pro41Asn | 0.92 | -0.12 | Neutral | 0.92 | -0.12 | Neutral |
| 41 | p.Pro41Lys | 1.28 | 0.35 | Indeterminate | 1.28 | 0.35 | Indeterminate |
| 41 | p.Pro41Thr | 1.03 | 0.04 | Neutral | 1.03 | 0.04 | Neutral |

| | | | | | | | | | | |
|---|---|---|---|---|---|---|---|---|---|---|
| 41 | p.Pro41Arg | 2.29 | 1.20 | Deleterious | | | | 2.29 | 1.20 | Deleterious |
| 41 | p.Pro41Ser | 0.97 | -0.05 | Neutral | | | | 0.97 | -0.05 | Neutral |
| 41 | p.Pro41Ile | 1.61 | 0.68 | Indeterminate | | | | 1.61 | 0.68 | Indeterminate |
| 41 | p.Pro41Met | 0.78 | -0.37 | Neutral | | | | 0.78 | -0.37 | Neutral |
| 41 | p.Pro41His | 1.79 | 0.84 | Indeterminate | | | | 1.79 | 0.84 | Indeterminate |
| 41 | p.Pro41Gln | 1.29 | 0.37 | Indeterminate | | | | 1.29 | 0.37 | Indeterminate |
| 41 | p.Pro41Pro | 1.00 | 0.00 | Neutral | | | | 1.00 | 0.00 | Neutral |
| 41 | p.Pro41Leu | 0.79 | -0.34 | Neutral | | | | 0.79 | -0.34 | Neutral |
| 41 | p.Pro41Asp | 1.28 | 0.35 | Indeterminate | | | | 1.28 | 0.35 | Indeterminate |
| 41 | p.Pro41Glu | 0.90 | -0.15 | Neutral | | | | 0.90 | -0.15 | Neutral |
| 41 | p.Pro41Ala | 1.00 | 0.01 | Neutral | | | | 1.00 | 0.01 | Neutral |
| 41 | p.Pro41Gly | 1.06 | 0.09 | Neutral | | | | 1.06 | 0.09 | Neutral |
| 41 | p.Pro41Val | 1.09 | 0.13 | Neutral | | | | 1.09 | 0.13 | Neutral |
| 41 | p.Pro41Tyr | 2.10 | 1.07 | Indeterminate | | | | 2.10 | 1.07 | Indeterminate |
| 41 | p.Pro41Cys | 1.32 | 0.40 | Indeterminate | | | | 1.32 | 0.40 | Indeterminate |
| 41 | p.Pro41Trp | 1.43 | 0.52 | Indeterminate | | | | 1.43 | 0.52 | Indeterminate |
| 41 | p.Pro41Phe | 1.52 | 0.61 | Indeterminate | | | | 1.52 | 0.61 | Indeterminate |
| 42 | p.Asn42Asn | 1.00 | 0.00 | Neutral | | | | 1.00 | 0.00 | Neutral |
| 42 | p.Asn42Lys | 12.54 | 3.65 | Deleterious | | | | 12.54 | 3.65 | Deleterious |
| 42 | p.Asn42Thr | 1.32 | 0.41 | Indeterminate | | | | 1.32 | 0.41 | Indeterminate |
| 42 | p.Asn42Arg | 11.65 | 3.54 | Deleterious | | | | 11.65 | 3.54 | Deleterious |
| 42 | p.Asn42Ser | 1.22 | 0.29 | Indeterminate | | | | 1.22 | 0.29 | Indeterminate |
| 42 | p.Asn42Ile | 11.57 | 3.53 | Deleterious | | | | 11.57 | 3.53 | Deleterious |
| 42 | p.Asn42Met | 9.77 | 3.29 | Deleterious | | | | 9.77 | 3.29 | Deleterious |
| 42 | p.Asn42His | 7.33 | 2.87 | Deleterious | | | | 7.33 | 2.87 | Deleterious |
| 42 | p.Asn42Gln | 7.10 | 2.83 | Deleterious | | | | 7.10 | 2.83 | Deleterious |
| 42 | p.Asn42Pro | 10.83 | 3.44 | Deleterious | | | | 10.83 | 3.44 | Deleterious |
| 42 | p.Asn42Leu | 11.43 | 3.51 | Deleterious | | | | 11.43 | 3.51 | Deleterious |
| 42 | p.Asn42Asp | 1.61 | 0.69 | Indeterminate | | | | 1.61 | 0.69 | Indeterminate |
| 42 | p.Asn42Glu | 12.48 | 3.64 | Deleterious | | | | 12.48 | 3.64 | Deleterious |
| 42 | p.Asn42Ala | 1.15 | 0.20 | Neutral | | | | 1.15 | 0.20 | Neutral |
| 42 | p.Asn42Gly | 0.94 | -0.09 | Neutral | | | | 0.94 | -0.09 | Neutral |
| 42 | p.Asn42Val | 9.99 | 3.32 | Deleterious | | | | 9.99 | 3.32 | Deleterious |
| 42 | p.Asn42Tyr | 11.87 | 3.57 | Deleterious | | | | 11.87 | 3.57 | Deleterious |
| 42 | p.Asn42Cys | 1.48 | 0.57 | Indeterminate | | | | 1.48 | 0.57 | Indeterminate |
| 42 | p.Asn42Trp | 14.65 | 3.87 | Deleterious | | | | 14.65 | 3.87 | Deleterious |
| 42 | p.Asn42Phe | 12.61 | 3.66 | Deleterious | | | | 12.61 | 3.66 | Deleterious |
| 43 | p.Ser43Asn | 1.10 | 0.14 | Neutral | | | | 1.10 | 0.14 | Neutral |
| 43 | p.Ser43Lys | 1.04 | 0.06 | Neutral | | | | 1.04 | 0.06 | Neutral |
| 43 | p.Ser43Thr | 0.97 | -0.04 | Neutral | | | | 0.97 | -0.04 | Neutral |
| 43 | p.Ser43Arg | 0.99 | -0.01 | Neutral | | | | 0.99 | -0.01 | Neutral |
| 43 | p.Ser43Ser | 1.00 | 0.00 | Neutral | | | | 1.00 | 0.00 | Neutral |
| 43 | p.Ser43Ile | 1.25 | 0.32 | Indeterminate | | | | 1.25 | 0.32 | Indeterminate |
| 43 | p.Ser43Met | 1.09 | 0.13 | Neutral | | | | 1.09 | 0.13 | Neutral |
| 43 | p.Ser43His | 1.00 | -0.01 | Neutral | | | | 1.00 | -0.01 | Neutral |
| 43 | p.Ser43Gln | 1.00 | 0.00 | Neutral | | | | 1.00 | 0.00 | Neutral |
| 43 | p.Ser43Pro | 0.99 | -0.01 | Neutral | | | | 0.99 | -0.01 | Neutral |
| 43 | p.Ser43Leu | 1.06 | 0.08 | Neutral | | | | 1.06 | 0.08 | Neutral |
| 43 | p.Ser43Asp | 1.09 | 0.12 | Neutral | | | | 1.09 | 0.12 | Neutral |
| 43 | p.Ser43Glu | 1.02 | 0.03 | Neutral | | | | 1.02 | 0.03 | Neutral |
| 43 | p.Ser43Ala | 0.97 | -0.05 | Neutral | | | | 0.97 | -0.05 | Neutral |
| 43 | p.Ser43Gly | 0.94 | -0.08 | Neutral | | | | 0.94 | -0.08 | Neutral |
| 43 | p.Ser43Val | 1.09 | 0.13 | Neutral | | | | 1.09 | 0.13 | Neutral |
| 43 | p.Ser43Tyr | 1.07 | 0.10 | Neutral | | | | 1.07 | 0.10 | Neutral |
| 43 | p.Ser43Cys | 1.06 | 0.08 | Neutral | | | | 1.06 | 0.08 | Neutral |
| 43 | p.Ser43Trp | 1.11 | 0.15 | Neutral | | | | 1.11 | 0.15 | Neutral |
| 43 | p.Ser43Phe | 1.05 | 0.08 | Neutral | | | | 1.05 | 0.08 | Neutral |
| 44 | p.Tyr44Asn | 1.21 | 0.28 | Indeterminate | | | | 1.21 | 0.28 | Indeterminate |
| 44 | p.Tyr44Lys | 1.21 | 0.27 | Indeterminate | | | | 1.21 | 0.27 | Indeterminate |
| 44 | p.Tyr44Thr | 0.99 | -0.02 | Neutral | | | | 0.99 | -0.02 | Neutral |
| 44 | p.Tyr44Arg | 1.03 | 0.04 | Neutral | | | | 1.03 | 0.04 | Neutral |
| 44 | p.Tyr44Ser | 1.09 | 0.12 | Neutral | | | | 1.09 | 0.12 | Neutral |
| 44 | p.Tyr44Ile | 1.05 | 0.07 | Neutral | | | | 1.05 | 0.07 | Neutral |
| 44 | p.Tyr44Met | 1.09 | 0.12 | Neutral | | | | 1.09 | 0.12 | Neutral |
| 44 | p.Tyr44His | 1.10 | 0.14 | Neutral | | | | 1.10 | 0.14 | Neutral |
| 44 | p.Tyr44Gln | 1.28 | 0.36 | Indeterminate | | | | 1.28 | 0.36 | Indeterminate |
| 44 | p.Tyr44Pro | 2.30 | 1.20 | Deleterious | | | | 2.30 | 1.20 | Deleterious |
| 44 | p.Tyr44Leu | 1.09 | 0.13 | Neutral | | | | 1.09 | 0.13 | Neutral |
| 44 | p.Tyr44Asp | 1.24 | 0.31 | Indeterminate | | | | 1.24 | 0.31 | Indeterminate |
| 44 | p.Tyr44Glu | 1.40 | 0.48 | Indeterminate | | | | 1.40 | 0.48 | Indeterminate |
| 44 | p.Tyr44Ala | 1.82 | 0.87 | Indeterminate | | | | 1.82 | 0.87 | Indeterminate |
| 44 | p.Tyr44Gly | 1.38 | 0.46 | Indeterminate | | | | 1.38 | 0.46 | Indeterminate |
| 44 | p.Tyr44Val | 1.09 | 0.13 | Neutral | | | | 1.09 | 0.13 | Neutral |
| 44 | p.Tyr44Tyr | 1.00 | 0.00 | Neutral | | | | 1.00 | 0.00 | Neutral |
| 44 | p.Tyr44Cys | 0.97 | -0.05 | Neutral | | | | 0.97 | -0.05 | Neutral |
| 44 | p.Tyr44Trp | 1.10 | 0.14 | Neutral | | | | 1.10 | 0.14 | Neutral |
| 44 | p.Tyr44Phe | 1.05 | 0.07 | Neutral | | | | 1.05 | 0.07 | Neutral |
| 45 | p.Gly45Asn | 0.71 | -0.49 | Neutral | 1.16 | 0.21 | Indeterminate | 0.93 | -0.10 | Neutral |
| 45 | p.Gly45Lys | 1.34 | 0.42 | Indeterminate | 1.14 | 0.19 | Indeterminate | 1.24 | 0.31 | Indeterminate |
| 45 | p.Gly45Thr | 0.64 | -0.64 | Neutral | 1.13 | 0.17 | Indeterminate | 0.88 | -0.18 | Neutral |
| 45 | p.Gly45Arg | 0.96 | -0.05 | Neutral | 1.11 | 0.16 | Indeterminate | 1.04 | 0.06 | Neutral |
| 45 | p.Gly45Ser | 1.93 | 0.95 | Indeterminate | 1.09 | 0.12 | Neutral | 1.51 | 0.59 | Indeterminate |
| 45 | p.Gly45Ile | 1.92 | 0.94 | Indeterminate | 1.20 | 0.27 | Indeterminate | 1.56 | 0.65 | Indeterminate |
| 45 | p.Gly45Met | 1.24 | 0.31 | Indeterminate | 1.12 | 0.16 | Indeterminate | 1.18 | 0.24 | Neutral |
| 45 | p.Gly45His | 1.10 | 0.14 | Neutral | 1.12 | 0.17 | Indeterminate | 1.11 | 0.15 | Neutral |
| 45 | p.Gly45Gln | 0.72 | -0.48 | Neutral | 1.02 | 0.04 | Neutral | 0.87 | -0.20 | Neutral |
| 45 | p.Gly45Pro | 49.14 | 5.62 | Deleterious | 4.40 | 2.14 | Deleterious | 26.77 | 4.74 | Deleterious |
| 45 | p.Gly45Leu | 1.15 | 0.20 | Neutral | 1.11 | 0.15 | Indeterminate | 1.13 | 0.17 | Neutral |
| 45 | p.Gly45Asp | 0.71 | -0.48 | Neutral | 1.05 | 0.07 | Neutral | 0.88 | -0.18 | Neutral |
| 45 | p.Gly45Glu | 1.09 | 0.12 | Neutral | 1.08 | 0.11 | Neutral | 1.08 | 0.11 | Neutral |
| 45 | p.Gly45Ala | 0.63 | -0.66 | Neutral | 1.13 | 0.18 | Indeterminate | 0.88 | -0.18 | Neutral |
| 45 | p.Gly45Gly | 1.00 | 0.00 | Neutral | 1.00 | 0.00 | Neutral | 1.00 | 0.00 | Neutral |
| 45 | p.Gly45Val | 1.91 | 0.93 | Indeterminate | 1.27 | 0.34 | Indeterminate | 1.59 | 0.67 | Indeterminate |
| 45 | p.Gly45Tyr | 0.73 | -0.46 | Neutral | 1.26 | 0.33 | Indeterminate | 0.99 | -0.01 | Neutral |
| 45 | p.Gly45Cys | 0.85 | -0.23 | Neutral | 1.09 | 0.13 | Neutral | 0.97 | -0.04 | Neutral |
| 45 | p.Gly45Trp | 1.04 | 0.05 | Neutral | 1.24 | 0.31 | Indeterminate | 1.14 | 0.19 | Neutral |
| 45 | p.Gly45Phe | 0.90 | -0.16 | Neutral | 1.02 | 0.03 | Neutral | 0.96 | -0.06 | Neutral |
| 46 | p.Arg46Asn | 0.92 | -0.13 | Neutral | | | | 0.92 | -0.13 | Neutral |
| 46 | p.Arg46Lys | 0.79 | -0.34 | Neutral | | | | 0.79 | -0.34 | Neutral |
| 46 | p.Arg46Thr | 2.61 | 1.38 | Deleterious | | | | 2.61 | 1.38 | Deleterious |
| 46 | p.Arg46Arg | 1.00 | 0.00 | Neutral | | | | 1.00 | 0.00 | Neutral |

| | | | | | | | |
|---|---|---|---|---|---|---|---|
| 46 | p.Arg46Ser | 0.85 | -0.23 | Neutral | 0.85 | -0.23 | Neutral |
| 46 | p.Arg46Ile | 3.54 | 1.82 | Deleterious | 3.54 | 1.82 | Deleterious |
| 46 | p.Arg46Met | 1.31 | 0.39 | Indeterminate | 1.31 | 0.39 | Indeterminate |
| 46 | p.Arg46His | 0.79 | -0.34 | Neutral | 0.79 | -0.34 | Neutral |
| 46 | p.Arg46Gln | 1.54 | 0.62 | Indeterminate | 1.54 | 0.62 | Indeterminate |
| 46 | p.Arg46Pro | 17.80 | 4.15 | Deleterious | 17.80 | 4.15 | Deleterious |
| 46 | p.Arg46Leu | 1.44 | 0.53 | Indeterminate | 1.44 | 0.53 | Indeterminate |
| 46 | p.Arg46Asp | 5.92 | 2.56 | Deleterious | 5.92 | 2.56 | Deleterious |
| 46 | p.Arg46Glu | 3.02 | 1.59 | Deleterious | 3.02 | 1.59 | Deleterious |
| 46 | p.Arg46Ala | 1.39 | 0.48 | Indeterminate | 1.39 | 0.48 | Indeterminate |
| 46 | p.Arg46Gly | 1.36 | 0.44 | Indeterminate | 1.36 | 0.44 | Indeterminate |
| 46 | p.Arg46Val | 1.88 | 0.91 | Indeterminate | 1.88 | 0.91 | Indeterminate |
| 46 | p.Arg46Tyr | 1.75 | 0.81 | Indeterminate | 1.75 | 0.81 | Indeterminate |
| 46 | p.Arg46Cys | 1.36 | 0.44 | Indeterminate | 1.36 | 0.44 | Indeterminate |
| 46 | p.Arg46Trp | 0.81 | -0.30 | Neutral | 0.81 | -0.30 | Neutral |
| 46 | p.Arg46Phe | 1.04 | 0.05 | Neutral | 1.04 | 0.05 | Neutral |
| 47 | p.Arg47Asn | 0.91 | -0.13 | Neutral | 0.91 | -0.13 | Neutral |
| 47 | p.Arg47Lys | 0.99 | -0.01 | Neutral | 0.99 | -0.01 | Neutral |
| 47 | p.Arg47Thr | 0.99 | -0.02 | Neutral | 0.99 | -0.02 | Neutral |
| 47 | p.Arg47Arg | 1.00 | 0.00 | Neutral | 1.00 | 0.00 | Neutral |
| 47 | p.Arg47Ser | 1.32 | 0.41 | Indeterminate | 1.32 | 0.41 | Indeterminate |
| 47 | p.Arg47Ile | 1.04 | 0.05 | Neutral | 1.04 | 0.05 | Neutral |
| 47 | p.Arg47Met | 1.06 | 0.08 | Neutral | 1.06 | 0.08 | Neutral |
| 47 | p.Arg47His | 1.06 | 0.09 | Neutral | 1.06 | 0.09 | Neutral |
| 47 | p.Arg47Gln | 1.50 | 0.59 | Indeterminate | 1.50 | 0.59 | Indeterminate |
| 47 | p.Arg47Pro | 1.60 | 0.67 | Indeterminate | 1.60 | 0.67 | Indeterminate |
| 47 | p.Arg47Leu | 0.86 | -0.21 | Neutral | 0.86 | -0.21 | Neutral |
| 47 | p.Arg47Asp | 3.53 | 1.82 | Deleterious | 3.53 | 1.82 | Deleterious |
| 47 | p.Arg47Glu | 1.27 | 0.35 | Indeterminate | 1.27 | 0.35 | Indeterminate |
| 47 | p.Arg47Ala | 0.99 | -0.01 | Neutral | 0.99 | -0.01 | Neutral |
| 47 | p.Arg47Gly | 1.42 | 0.51 | Indeterminate | 1.42 | 0.51 | Indeterminate |
| 47 | p.Arg47Val | 0.97 | -0.04 | Neutral | 0.97 | -0.04 | Neutral |
| 47 | p.Arg47Tyr | 1.14 | 0.19 | Neutral | 1.14 | 0.19 | Neutral |
| 47 | p.Arg47Cys | 0.89 | -0.18 | Neutral | 0.89 | -0.18 | Neutral |
| 47 | p.Arg47Trp | 1.20 | 0.26 | Indeterminate | 1.20 | 0.26 | Indeterminate |
| 47 | p.Arg47Phe | 0.89 | -0.17 | Neutral | 0.89 | -0.17 | Neutral |
| 48 | p.Pro48Asn | 13.23 | 3.73 | Deleterious | 13.23 | 3.73 | Deleterious |
| 48 | p.Pro48Lys | 15.42 | 3.95 | Deleterious | 15.42 | 3.95 | Deleterious |
| 48 | p.Pro48Thr | 2.13 | 1.09 | Deleterious | 2.13 | 1.09 | Deleterious |
| 48 | p.Pro48Arg | 13.40 | 3.74 | Deleterious | 13.40 | 3.74 | Deleterious |
| 48 | p.Pro48Ser | 1.50 | 0.58 | Indeterminate | 1.50 | 0.58 | Indeterminate |
| 48 | p.Pro48Ile | 7.83 | 2.97 | Deleterious | 7.83 | 2.97 | Deleterious |
| 48 | p.Pro48Met | 11.91 | 3.57 | Deleterious | 11.91 | 3.57 | Deleterious |
| 48 | p.Pro48His | 11.54 | 3.53 | Deleterious | 11.54 | 3.53 | Deleterious |
| 48 | p.Pro48Gln | 14.93 | 3.90 | Deleterious | 14.93 | 3.90 | Deleterious |
| 48 | p.Pro48Pro | 1.00 | 0.00 | Neutral | 1.00 | 0.00 | Neutral |
| 48 | p.Pro48Leu | 17.50 | 4.13 | Deleterious | 17.50 | 4.13 | Deleterious |
| 48 | p.Pro48Asp | 17.91 | 4.16 | Deleterious | 17.91 | 4.16 | Deleterious |
| 48 | p.Pro48Glu | 16.63 | 4.06 | Deleterious | 16.63 | 4.06 | Deleterious |
| 48 | p.Pro48Ala | 0.75 | -0.42 | Neutral | 0.75 | -0.42 | Neutral |
| 48 | p.Pro48Gly | 2.08 | 1.05 | Indeterminate | 2.08 | 1.05 | Indeterminate |
| 48 | p.Pro48Val | 2.16 | 1.11 | Deleterious | 2.16 | 1.11 | Deleterious |
| 48 | p.Pro48Tyr | 18.13 | 4.18 | Deleterious | 18.13 | 4.18 | Deleterious |
| 48 | p.Pro48Cys | 2.37 | 1.25 | Deleterious | 2.37 | 1.25 | Deleterious |
| 48 | p.Pro48Trp | 17.96 | 4.17 | Deleterious | 17.96 | 4.17 | Deleterious |
| 48 | p.Pro48Phe | 15.06 | 3.91 | Deleterious | 15.06 | 3.91 | Deleterious |
| 49 | p.Ile49Asn | 6.56 | 2.71 | Deleterious | 6.56 | 2.71 | Deleterious |
| 49 | p.Ile49Lys | 8.49 | 3.09 | Deleterious | 8.49 | 3.09 | Deleterious |
| 49 | p.Ile49Thr | 2.32 | 1.21 | Deleterious | 2.32 | 1.21 | Deleterious |
| 49 | p.Ile49Arg | 7.54 | 2.91 | Deleterious | 7.54 | 2.91 | Deleterious |
| 49 | p.Ile49Ser | 4.51 | 2.17 | Deleterious | 4.51 | 2.17 | Deleterious |
| 49 | p.Ile49Ile | 1.00 | 0.00 | Neutral | 1.00 | 0.00 | Neutral |
| 49 | p.Ile49Met | 1.09 | 0.12 | Neutral | 1.09 | 0.12 | Neutral |
| 49 | p.Ile49His | 4.93 | 2.30 | Deleterious | 4.93 | 2.30 | Deleterious |
| 49 | p.Ile49Gln | 3.98 | 1.99 | Deleterious | 3.98 | 1.99 | Deleterious |
| 49 | p.Ile49Pro | 7.18 | 2.84 | Deleterious | 7.18 | 2.84 | Deleterious |
| 49 | p.Ile49Leu | 0.97 | -0.05 | Neutral | 0.97 | -0.05 | Neutral |
| 49 | p.Ile49Asp | 9.18 | 3.20 | Deleterious | 9.18 | 3.20 | Deleterious |
| 49 | p.Ile49Glu | 8.40 | 3.07 | Deleterious | 8.40 | 3.07 | Deleterious |
| 49 | p.Ile49Ala | 2.27 | 1.19 | Deleterious | 2.27 | 1.19 | Deleterious |
| 49 | p.Ile49Gly | 8.88 | 3.15 | Deleterious | 8.88 | 3.15 | Deleterious |
| 49 | p.Ile49Val | 1.09 | 0.12 | Neutral | 1.09 | 0.12 | Neutral |
| 49 | p.Ile49Tyr | 6.77 | 2.76 | Deleterious | 6.77 | 2.76 | Deleterious |
| 49 | p.Ile49Cys | 1.85 | 0.89 | Indeterminate | 1.85 | 0.89 | Indeterminate |
| 49 | p.Ile49Trp | 7.82 | 2.97 | Deleterious | 7.82 | 2.97 | Deleterious |
| 49 | p.Ile49Phe | 2.33 | 1.22 | Deleterious | 2.33 | 1.22 | Deleterious |
| 50 | p.Gln50Asn | 1.33 | 0.41 | Indeterminate | 1.33 | 0.41 | Indeterminate |
| 50 | p.Gln50Lys | 12.74 | 3.67 | Deleterious | 12.74 | 3.67 | Deleterious |
| 50 | p.Gln50Thr | 1.26 | 0.33 | Indeterminate | 1.26 | 0.33 | Indeterminate |
| 50 | p.Gln50Arg | 17.01 | 4.09 | Deleterious | 17.01 | 4.09 | Deleterious |
| 50 | p.Gln50Ser | 1.18 | 0.24 | Indeterminate | 1.18 | 0.24 | Indeterminate |
| 50 | p.Gln50Ile | 4.65 | 2.22 | Deleterious | 4.65 | 2.22 | Deleterious |
| 50 | p.Gln50Met | 1.04 | 0.05 | Neutral | 1.04 | 0.05 | Neutral |
| 50 | p.Gln50His | 4.31 | 2.11 | Deleterious | 4.31 | 2.11 | Deleterious |
| 50 | p.Gln50Gln | 1.00 | 0.00 | Neutral | 1.00 | 0.00 | Neutral |
| 50 | p.Gln50Pro | 18.09 | 4.18 | Deleterious | 18.09 | 4.18 | Deleterious |
| 50 | p.Gln50Leu | 1.87 | 0.90 | Indeterminate | 1.87 | 0.90 | Indeterminate |
| 50 | p.Gln50Asp | 6.89 | 2.78 | Deleterious | 6.89 | 2.78 | Deleterious |
| 50 | p.Gln50Glu | 1.05 | 0.08 | Neutral | 1.05 | 0.08 | Neutral |
| 50 | p.Gln50Ala | 1.36 | 0.45 | Indeterminate | 1.36 | 0.45 | Indeterminate |
| 50 | p.Gln50Gly | 1.47 | 0.55 | Indeterminate | 1.47 | 0.55 | Indeterminate |
| 50 | p.Gln50Val | 3.95 | 1.98 | Deleterious | 3.95 | 1.98 | Deleterious |
| 50 | p.Gln50Tyr | 16.89 | 4.08 | Deleterious | 16.89 | 4.08 | Deleterious |
| 50 | p.Gln50Cys | 1.85 | 0.89 | Indeterminate | 1.85 | 0.89 | Indeterminate |
| 50 | p.Gln50Trp | 14.91 | 3.90 | Deleterious | 14.91 | 3.90 | Deleterious |
| 50 | p.Gln50Phe | 13.94 | 3.80 | Deleterious | 13.94 | 3.80 | Deleterious |
| 51 | p.Val51Asn | 2.32 | 1.21 | Deleterious | 2.32 | 1.21 | Deleterious |
| 51 | p.Val51Lys | 33.67 | 5.07 | Deleterious | 33.67 | 5.07 | Deleterious |
| 51 | p.Val51Thr | 0.95 | -0.08 | Neutral | 0.95 | -0.08 | Neutral |
| 51 | p.Val51Arg | 35.63 | 5.16 | Deleterious | 35.63 | 5.16 | Deleterious |
| 51 | p.Val51Ser | 0.75 | -0.42 | Neutral | 0.75 | -0.42 | Neutral |

| | | | | | | | |
|---|---|---|---|---|---|---|---|
| 51 | p.Val51Ile | 1.28 | 0.35 | Indeterminate | 1.28 | 0.35 | Indeterminate |
| 51 | p.Val51Met | 2.48 | 1.31 | Deleterious | 2.48 | 1.31 | Deleterious |
| 51 | p.Val51His | 19.51 | 4.29 | Deleterious | 19.51 | 4.29 | Deleterious |
| 51 | p.Val51Gln | 3.55 | 1.83 | Deleterious | 3.55 | 1.83 | Deleterious |
| 51 | p.Val51Pro | 22.42 | 4.49 | Deleterious | 22.42 | 4.49 | Deleterious |
| 51 | p.Val51Leu | 1.07 | 0.10 | Neutral | 1.07 | 0.10 | Neutral |
| 51 | p.Val51Asp | 20.68 | 4.37 | Deleterious | 20.68 | 4.37 | Deleterious |
| 51 | p.Val51Glu | 23.51 | 4.56 | Deleterious | 23.51 | 4.56 | Deleterious |
| 51 | p.Val51Ala | 0.57 | -0.82 | Neutral | 0.57 | -0.82 | Neutral |
| 51 | p.Val51Gly | 1.00 | 0.00 | Neutral | 1.00 | 0.00 | Neutral |
| 51 | p.Val51Val | 1.00 | 0.00 | Neutral | 1.00 | 0.00 | Neutral |
| 51 | p.Val51Tyr | 11.39 | 3.51 | Deleterious | 11.39 | 3.51 | Deleterious |
| 51 | p.Val51Cys | 0.95 | -0.08 | Neutral | 0.95 | -0.08 | Neutral |
| 51 | p.Val51Trp | 24.18 | 4.60 | Deleterious | 24.18 | 4.60 | Deleterious |
| 51 | p.Val51Phe | 8.55 | 3.10 | Deleterious | 8.55 | 3.10 | Deleterious |
| 52 | p.Met52Asn | 1.75 | 0.81 | Indeterminate | 1.75 | 0.81 | Indeterminate |
| 52 | p.Met52Lys | 2.56 | 1.36 | Deleterious | 2.56 | 1.36 | Deleterious |
| 52 | p.Met52Thr | 1.16 | 0.21 | Neutral | 1.16 | 0.21 | Neutral |
| 52 | p.Met52Arg | 2.66 | 1.41 | Deleterious | 2.66 | 1.41 | Deleterious |
| 52 | p.Met52Ser | 1.03 | 0.04 | Neutral | 1.03 | 0.04 | Neutral |
| 52 | p.Met52Ile | 1.42 | 0.51 | Indeterminate | 1.42 | 0.51 | Indeterminate |
| 52 | p.Met52Met | 1.00 | 0.00 | Neutral | 1.00 | 0.00 | Neutral |
| 52 | p.Met52His | 1.54 | 0.62 | Indeterminate | 1.54 | 0.62 | Indeterminate |
| 52 | p.Met52Gln | 1.22 | 0.29 | Indeterminate | 1.22 | 0.29 | Indeterminate |
| 52 | p.Met52Pro | 2.22 | 1.15 | Deleterious | 2.22 | 1.15 | Deleterious |
| 52 | p.Met52Leu | 1.05 | 0.08 | Neutral | 1.05 | 0.08 | Neutral |
| 52 | p.Met52Asp | 2.54 | 1.34 | Deleterious | 2.54 | 1.34 | Deleterious |
| 52 | p.Met52Glu | 2.04 | 1.03 | Indeterminate | 2.04 | 1.03 | Indeterminate |
| 52 | p.Met52Ala | 1.03 | 0.04 | Neutral | 1.03 | 0.04 | Neutral |
| 52 | p.Met52Gly | 0.98 | -0.03 | Neutral | 0.98 | -0.03 | Neutral |
| 52 | p.Met52Val | 0.98 | -0.03 | Neutral | 0.98 | -0.03 | Neutral |
| 52 | p.Met52Tyr | 1.50 | 0.59 | Indeterminate | 1.50 | 0.59 | Indeterminate |
| 52 | p.Met52Cys | 0.96 | -0.06 | Neutral | 0.96 | -0.06 | Neutral |
| 52 | p.Met52Trp | 1.98 | 0.99 | Indeterminate | 1.98 | 0.99 | Indeterminate |
| 52 | p.Met52Phe | 1.29 | 0.37 | Indeterminate | 1.29 | 0.37 | Indeterminate |
| 53 | p.Met53Asn | 2.85 | 1.51 | Deleterious | 2.85 | 1.51 | Deleterious |
| 53 | p.Met53Lys | 1.89 | 0.92 | Indeterminate | 1.89 | 0.92 | Indeterminate |
| 53 | p.Met53Thr | 3.31 | 1.73 | Deleterious | 3.31 | 1.73 | Deleterious |
| 53 | p.Met53Arg | 1.43 | 0.51 | Indeterminate | 1.43 | 0.51 | Indeterminate |
| 53 | p.Met53Ser | 1.68 | 0.75 | Indeterminate | 1.68 | 0.75 | Indeterminate |
| 53 | p.Met53Ile | 6.83 | 2.77 | Deleterious | 6.83 | 2.77 | Deleterious |
| 53 | p.Met53Met | 1.00 | 0.00 | Neutral | 1.00 | 0.00 | Neutral |
| 53 | p.Met53His | 3.28 | 1.71 | Deleterious | 3.28 | 1.71 | Deleterious |
| 53 | p.Met53Gln | 1.45 | 0.53 | Indeterminate | 1.45 | 0.53 | Indeterminate |
| 53 | p.Met53Pro | 9.71 | 3.28 | Deleterious | 9.71 | 3.28 | Deleterious |
| 53 | p.Met53Leu | 1.81 | 0.86 | Indeterminate | 1.81 | 0.86 | Indeterminate |
| 53 | p.Met53Asp | 13.93 | 3.80 | Deleterious | 13.93 | 3.80 | Deleterious |
| 53 | p.Met53Glu | 4.03 | 2.01 | Deleterious | 4.03 | 2.01 | Deleterious |
| 53 | p.Met53Ala | 1.58 | 0.66 | Indeterminate | 1.58 | 0.66 | Indeterminate |
| 53 | p.Met53Gly | 6.11 | 2.61 | Deleterious | 6.11 | 2.61 | Deleterious |
| 53 | p.Met53Val | 7.64 | 2.93 | Deleterious | 7.64 | 2.93 | Deleterious |
| 53 | p.Met53Tyr | 6.71 | 2.75 | Deleterious | 6.71 | 2.75 | Deleterious |
| 53 | p.Met53Cys | 1.82 | 0.86 | Indeterminate | 1.82 | 0.86 | Indeterminate |
| 53 | p.Met53Trp | 11.34 | 3.50 | Deleterious | 11.34 | 3.50 | Deleterious |
| 53 | p.Met53Phe | 5.92 | 2.57 | Deleterious | 5.92 | 2.57 | Deleterious |
| 54 | p.Met54Asn | 1.34 | 0.43 | Indeterminate | 1.34 | 0.43 | Indeterminate |
| 54 | p.Met54Lys | 1.87 | 0.91 | Indeterminate | 1.87 | 0.91 | Indeterminate |
| 54 | p.Met54Thr | 0.96 | -0.05 | Neutral | 0.96 | -0.05 | Neutral |
| 54 | p.Met54Arg | 2.35 | 1.23 | Deleterious | 2.35 | 1.23 | Deleterious |
| 54 | p.Met54Ser | 1.12 | 0.17 | Neutral | 1.12 | 0.17 | Neutral |
| 54 | p.Met54Ile | 1.18 | 0.24 | Indeterminate | 1.18 | 0.24 | Indeterminate |
| 54 | p.Met54Met | 1.00 | 0.00 | Neutral | 1.00 | 0.00 | Neutral |
| 54 | p.Met54His | 1.26 | 0.33 | Indeterminate | 1.26 | 0.33 | Indeterminate |
| 54 | p.Met54Gln | 1.66 | 0.74 | Indeterminate | 1.66 | 0.74 | Indeterminate |
| 54 | p.Met54Pro | 2.75 | 1.46 | Deleterious | 2.75 | 1.46 | Deleterious |
| 54 | p.Met54Leu | 1.09 | 0.13 | Neutral | 1.09 | 0.13 | Neutral |
| 54 | p.Met54Asp | 2.35 | 1.23 | Deleterious | 2.35 | 1.23 | Deleterious |
| 54 | p.Met54Glu | 2.09 | 1.07 | Indeterminate | 2.09 | 1.07 | Indeterminate |
| 54 | p.Met54Ala | 1.01 | 0.02 | Neutral | 1.01 | 0.02 | Neutral |
| 54 | p.Met54Gly | 1.03 | 0.05 | Neutral | 1.03 | 0.05 | Neutral |
| 54 | p.Met54Val | 1.05 | 0.07 | Neutral | 1.05 | 0.07 | Neutral |
| 54 | p.Met54Tyr | 1.04 | 0.06 | Neutral | 1.04 | 0.06 | Neutral |
| 54 | p.Met54Cys | 0.90 | -0.15 | Neutral | 0.90 | -0.15 | Neutral |
| 54 | p.Met54Trp | 1.22 | 0.29 | Indeterminate | 1.22 | 0.29 | Indeterminate |
| 54 | p.Met54Phe | 1.11 | 0.15 | Neutral | 1.11 | 0.15 | Neutral |
| 55 | p.Gly55Asn | 12.63 | 3.66 | Deleterious | 12.63 | 3.66 | Deleterious |
| 55 | p.Gly55Lys | 12.04 | 3.59 | Deleterious | 12.04 | 3.59 | Deleterious |
| 55 | p.Gly55Thr | 9.96 | 3.32 | Deleterious | 9.96 | 3.32 | Deleterious |
| 55 | p.Gly55Arg | 11.97 | 3.58 | Deleterious | 11.97 | 3.58 | Deleterious |
| 55 | p.Gly55Ser | 2.68 | 1.42 | Deleterious | 2.68 | 1.42 | Deleterious |
| 55 | p.Gly55Ile | 11.78 | 3.56 | Deleterious | 11.78 | 3.56 | Deleterious |
| 55 | p.Gly55Met | 14.07 | 3.81 | Deleterious | 14.07 | 3.81 | Deleterious |
| 55 | p.Gly55His | 15.68 | 3.97 | Deleterious | 15.68 | 3.97 | Deleterious |
| 55 | p.Gly55Gln | 13.16 | 3.72 | Deleterious | 13.16 | 3.72 | Deleterious |
| 55 | p.Gly55Pro | 20.49 | 4.36 | Deleterious | 20.49 | 4.36 | Deleterious |
| 55 | p.Gly55Leu | 17.12 | 4.10 | Deleterious | 17.12 | 4.10 | Deleterious |
| 55 | p.Gly55Asp | 16.49 | 4.04 | Deleterious | 16.49 | 4.04 | Deleterious |
| 55 | p.Gly55Glu | 13.07 | 3.71 | Deleterious | 13.07 | 3.71 | Deleterious |
| 55 | p.Gly55Ala | 1.62 | 0.70 | Indeterminate | 1.62 | 0.70 | Indeterminate |
| 55 | p.Gly55Gly | 1.00 | 0.00 | Neutral | 1.00 | 0.00 | Neutral |
| 55 | p.Gly55Val | 14.09 | 3.82 | Deleterious | 14.09 | 3.82 | Deleterious |
| 55 | p.Gly55Tyr | 10.85 | 3.44 | Deleterious | 10.85 | 3.44 | Deleterious |
| 55 | p.Gly55Cys | 3.75 | 1.91 | Deleterious | 3.75 | 1.91 | Deleterious |
| 55 | p.Gly55Trp | 14.81 | 3.89 | Deleterious | 14.81 | 3.89 | Deleterious |
| 55 | p.Gly55Phe | 11.47 | 3.52 | Deleterious | 11.47 | 3.52 | Deleterious |
| 56 | p.Ser56Asn | 0.30 | -1.72 | Neutral | 0.30 | -1.72 | Neutral |
| 56 | p.Ser56Lys | 0.82 | -0.29 | Neutral | 0.82 | -0.29 | Neutral |
| 56 | p.Ser56Thr | 0.79 | -0.34 | Neutral | 0.79 | -0.34 | Neutral |
| 56 | p.Ser56Arg | 0.38 | -1.41 | Neutral | 0.38 | -1.41 | Neutral |
| 56 | p.Ser56Ser | 1.00 | 0.00 | Neutral | 1.00 | 0.00 | Neutral |
| 56 | p.Ser56Ile | 6.45 | 2.69 | Deleterious | 6.45 | 2.69 | Deleterious |

| | | | | | | | | | | |
|---|---|---|---|---|---|---|---|---|---|---|
| 56 | p.Ser56Met | 0.42 | -1.27 | Neutral | | | | 0.42 | -1.27 | Neutral |
| 56 | p.Ser56His | 0.43 | -1.23 | Neutral | | | | 0.43 | -1.23 | Neutral |
| 56 | p.Ser56Gln | 0.57 | -0.81 | Neutral | | | | 0.57 | -0.81 | Neutral |
| 56 | p.Ser56Pro | 12.14 | 3.60 | Deleterious | | | | 12.14 | 3.60 | Deleterious |
| 56 | p.Ser56Leu | 1.87 | 0.90 | Indeterminate | | | | 1.87 | 0.90 | Indeterminate |
| 56 | p.Ser56Asp | 0.79 | -0.34 | Neutral | | | | 0.79 | -0.34 | Neutral |
| 56 | p.Ser56Glu | 1.20 | 0.26 | Indeterminate | | | | 1.20 | 0.26 | Indeterminate |
| 56 | p.Ser56Ala | 0.77 | -0.38 | Neutral | | | | 0.77 | -0.38 | Neutral |
| 56 | p.Ser56Gly | 1.18 | 0.24 | Neutral | | | | 1.18 | 0.24 | Neutral |
| 56 | p.Ser56Val | 1.16 | 0.22 | Neutral | | | | 1.16 | 0.22 | Neutral |
| 56 | p.Ser56Tyr | 0.50 | -0.99 | Neutral | | | | 0.50 | -0.99 | Neutral |
| 56 | p.Ser56Cys | 0.43 | -1.22 | Neutral | | | | 0.43 | -1.22 | Neutral |
| 56 | p.Ser56Trp | 4.65 | 2.22 | Deleterious | | | | 4.65 | 2.22 | Deleterious |
| 56 | p.Ser56Phe | 0.72 | -0.47 | Neutral | | | | 0.72 | -0.47 | Neutral |
| 57 | p.Ala57Asn | 0.47 | -1.08 | Neutral | 1.14 | 0.19 | Indeterminate | 0.81 | -0.31 | Neutral |
| 57 | p.Ala57Lys | 0.43 | -1.22 | Neutral | 1.30 | 0.38 | Indeterminate | 0.87 | -0.21 | Neutral |
| 57 | p.Ala57Thr | 0.56 | -0.83 | Neutral | 1.31 | 0.39 | Indeterminate | 0.94 | -0.09 | Neutral |
| 57 | p.Ala57Arg | 0.49 | -1.03 | Neutral | 1.04 | 0.05 | Neutral | 0.76 | -0.39 | Neutral |
| 57 | p.Ala57Ser | 0.58 | -0.79 | Neutral | 1.58 | 0.66 | Indeterminate | 1.08 | 0.11 | Neutral |
| 57 | p.Ala57Ile | 0.71 | -0.49 | Neutral | 1.17 | 0.23 | Indeterminate | 0.94 | -0.09 | Neutral |
| 57 | p.Ala57Met | 0.45 | -1.16 | Neutral | 1.26 | 0.33 | Indeterminate | 0.85 | -0.23 | Neutral |
| 57 | p.Ala57His | 0.56 | -0.84 | Neutral | 0.90 | -0.15 | Neutral | 0.73 | -0.45 | Neutral |
| 57 | p.Ala57Gln | 0.54 | -0.89 | Neutral | 1.27 | 0.34 | Indeterminate | 0.90 | -0.14 | Neutral |
| 57 | p.Ala57Pro | 0.62 | -0.70 | Neutral | 1.49 | 0.58 | Indeterminate | 1.05 | 0.08 | Neutral |
| 57 | p.Ala57Leu | 0.44 | -1.17 | Neutral | 1.62 | 0.70 | Indeterminate | 1.03 | 0.05 | Neutral |
| 57 | p.Ala57Asp | 0.77 | -0.38 | Neutral | 1.05 | 0.07 | Neutral | 0.91 | -0.14 | Neutral |
| 57 | p.Ala57Glu | 0.56 | -0.83 | Neutral | 1.84 | 0.88 | Indeterminate | 1.20 | 0.27 | Indeterminate |
| 57 | p.Ala57Ala | 1.00 | 0.00 | Neutral | 1.00 | 0.00 | Neutral | 1.00 | 0.00 | Neutral |
| 57 | p.Ala57Gly | 0.50 | -1.00 | Neutral | 1.27 | 0.35 | Indeterminate | 0.89 | -0.18 | Neutral |
| 57 | p.Ala57Val | 0.53 | -0.91 | Neutral | 1.10 | 0.14 | Neutral | 0.82 | -0.29 | Neutral |
| 57 | p.Ala57Tyr | 0.52 | -0.93 | Neutral | 1.86 | 0.89 | Indeterminate | 1.19 | 0.25 | Indeterminate |
| 57 | p.Ala57Cys | 0.38 | -1.40 | Neutral | 1.53 | 0.61 | Indeterminate | 0.95 | -0.07 | Neutral |
| 57 | p.Ala57Trp | 0.46 | -1.13 | Neutral | 1.54 | 0.63 | Indeterminate | 1.00 | 0.00 | Neutral |
| 57 | p.Ala57Phe | 0.33 | -1.62 | Neutral | 1.69 | 0.76 | Indeterminate | 1.01 | 0.01 | Neutral |
| 58 | p.Arg58Asn | 0.75 | -0.41 | Neutral | | | | 0.75 | -0.41 | Neutral |
| 58 | p.Arg58Lys | 0.71 | -0.49 | Neutral | | | | 0.71 | -0.49 | Neutral |
| 58 | p.Arg58Thr | 0.81 | -0.31 | Neutral | | | | 0.81 | -0.31 | Neutral |
| 58 | p.Arg58Arg | 1.00 | 0.00 | Neutral | | | | 1.00 | 0.00 | Neutral |
| 58 | p.Arg58Ser | 0.76 | -0.40 | Neutral | | | | 0.76 | -0.40 | Neutral |
| 58 | p.Arg58Ile | 0.81 | -0.31 | Neutral | | | | 0.81 | -0.31 | Neutral |
| 58 | p.Arg58Met | 0.71 | -0.50 | Neutral | | | | 0.71 | -0.50 | Neutral |
| 58 | p.Arg58His | 0.64 | -0.65 | Neutral | | | | 0.64 | -0.65 | Neutral |
| 58 | p.Arg58Gln | 0.93 | -0.10 | Neutral | | | | 0.93 | -0.10 | Neutral |
| 58 | p.Arg58Pro | 1.08 | 0.11 | Neutral | | | | 1.08 | 0.11 | Neutral |
| 58 | p.Arg58Leu | 0.83 | -0.28 | Neutral | | | | 0.83 | -0.28 | Neutral |
| 58 | p.Arg58Asp | 0.72 | -0.48 | Neutral | | | | 0.72 | -0.48 | Neutral |
| 58 | p.Arg58Glu | 0.83 | -0.26 | Neutral | | | | 0.83 | -0.26 | Neutral |
| 58 | p.Arg58Ala | 0.88 | -0.18 | Neutral | | | | 0.88 | -0.18 | Neutral |
| 58 | p.Arg58Gly | 0.93 | -0.11 | Neutral | | | | 0.93 | -0.11 | Neutral |
| 58 | p.Arg58Val | 0.90 | -0.16 | Neutral | | | | 0.90 | -0.16 | Neutral |
| 58 | p.Arg58Tyr | 0.62 | -0.69 | Neutral | | | | 0.62 | -0.69 | Neutral |
| 58 | p.Arg58Cys | 0.71 | -0.50 | Neutral | | | | 0.71 | -0.50 | Neutral |
| 58 | p.Arg58Trp | 0.76 | -0.40 | Neutral | | | | 0.76 | -0.40 | Neutral |
| 58 | p.Arg58Phe | 0.90 | -0.16 | Neutral | | | | 0.90 | -0.16 | Neutral |
| 59 | p.Val59Asn | 3.19 | 1.67 | Deleterious | | | | 3.19 | 1.67 | Deleterious |
| 59 | p.Val59Lys | 12.88 | 3.69 | Deleterious | | | | 12.88 | 3.69 | Deleterious |
| 59 | p.Val59Thr | 0.86 | -0.22 | Neutral | | | | 0.86 | -0.22 | Neutral |
| 59 | p.Val59Arg | 19.15 | 4.26 | Deleterious | | | | 19.15 | 4.26 | Deleterious |
| 59 | p.Val59Ser | 1.25 | 0.32 | Indeterminate | | | | 1.25 | 0.32 | Indeterminate |
| 59 | p.Val59Ile | 0.74 | -0.44 | Neutral | | | | 0.74 | -0.44 | Neutral |
| 59 | p.Val59Met | 1.01 | 0.01 | Neutral | | | | 1.01 | 0.01 | Neutral |
| 59 | p.Val59His | 16.58 | 4.05 | Deleterious | | | | 16.58 | 4.05 | Deleterious |
| 59 | p.Val59Gln | 3.62 | 1.86 | Deleterious | | | | 3.62 | 1.86 | Deleterious |
| 59 | p.Val59Pro | 12.69 | 3.67 | Deleterious | | | | 12.69 | 3.67 | Deleterious |
| 59 | p.Val59Leu | 1.31 | 0.39 | Indeterminate | | | | 1.31 | 0.39 | Indeterminate |
| 59 | p.Val59Asp | 20.88 | 4.38 | Deleterious | | | | 20.88 | 4.38 | Deleterious |
| 59 | p.Val59Glu | 11.98 | 3.58 | Deleterious | | | | 11.98 | 3.58 | Deleterious |
| 59 | p.Val59Ala | 1.17 | 0.23 | Neutral | | | | 1.17 | 0.23 | Neutral |
| 59 | p.Val59Gly | 2.18 | 1.12 | Deleterious | | | | 2.18 | 1.12 | Deleterious |
| 59 | p.Val59Val | 1.00 | 0.00 | Neutral | | | | 1.00 | 0.00 | Neutral |
| 59 | p.Val59Tyr | 18.77 | 4.23 | Deleterious | | | | 18.77 | 4.23 | Deleterious |
| 59 | p.Val59Cys | 0.94 | -0.08 | Neutral | | | | 0.94 | -0.08 | Neutral |
| 59 | p.Val59Trp | 21.32 | 4.41 | Deleterious | | | | 21.32 | 4.41 | Deleterious |
| 59 | p.Val59Phe | 17.60 | 4.14 | Deleterious | | | | 17.60 | 4.14 | Deleterious |
| 60 | p.Ala60Asn | 2.67 | 1.42 | Deleterious | | | | 2.67 | 1.42 | Deleterious |
| 60 | p.Ala60Lys | 3.23 | 1.69 | Deleterious | | | | 3.23 | 1.69 | Deleterious |
| 60 | p.Ala60Thr | 1.08 | 0.11 | Neutral | | | | 1.08 | 0.11 | Neutral |
| 60 | p.Ala60Arg | 4.98 | 2.32 | Deleterious | | | | 4.98 | 2.32 | Deleterious |
| 60 | p.Ala60Ser | 1.79 | 0.84 | Indeterminate | | | | 1.79 | 0.84 | Indeterminate |
| 60 | p.Ala60Ile | 1.99 | 0.99 | Indeterminate | | | | 1.99 | 0.99 | Indeterminate |
| 60 | p.Ala60Met | 1.23 | 0.29 | Indeterminate | | | | 1.23 | 0.29 | Indeterminate |
| 60 | p.Ala60His | 2.62 | 1.39 | Deleterious | | | | 2.62 | 1.39 | Deleterious |
| 60 | p.Ala60Gln | 3.17 | 1.67 | Deleterious | | | | 3.17 | 1.67 | Deleterious |
| 60 | p.Ala60Pro | 1.71 | 0.78 | Indeterminate | | | | 1.71 | 0.78 | Indeterminate |
| 60 | p.Ala60Leu | 1.06 | 0.09 | Neutral | | | | 1.06 | 0.09 | Neutral |
| 60 | p.Ala60Asp | 2.68 | 1.42 | Deleterious | | | | 2.68 | 1.42 | Deleterious |
| 60 | p.Ala60Glu | 2.83 | 1.50 | Deleterious | | | | 2.83 | 1.50 | Deleterious |
| 60 | p.Ala60Ala | 1.00 | 0.00 | Neutral | | | | 1.00 | 0.00 | Neutral |
| 60 | p.Ala60Gly | 1.05 | 0.07 | Neutral | | | | 1.05 | 0.07 | Neutral |
| 60 | p.Ala60Val | 1.24 | 0.31 | Indeterminate | | | | 1.24 | 0.31 | Indeterminate |
| 60 | p.Ala60Tyr | 2.82 | 1.49 | Deleterious | | | | 2.82 | 1.49 | Deleterious |
| 60 | p.Ala60Cys | 1.32 | 0.40 | Indeterminate | | | | 1.32 | 0.40 | Indeterminate |
| 60 | p.Ala60Trp | 3.21 | 1.68 | Deleterious | | | | 3.21 | 1.68 | Deleterious |
| 60 | p.Ala60Phe | 3.38 | 1.76 | Deleterious | | | | 3.38 | 1.76 | Deleterious |
| 61 | p.Glu61Asn | 0.99 | -0.01 | Neutral | | | | 0.99 | -0.01 | Neutral |
| 61 | p.Glu61Lys | 0.78 | -0.35 | Neutral | | | | 0.78 | -0.35 | Neutral |
| 61 | p.Glu61Thr | 0.88 | -0.18 | Neutral | | | | 0.88 | -0.18 | Neutral |
| 61 | p.Glu61Arg | 1.23 | 0.30 | Indeterminate | | | | 1.23 | 0.30 | Indeterminate |
| 61 | p.Glu61Ser | 1.06 | 0.09 | Neutral | | | | 1.06 | 0.09 | Neutral |
| 61 | p.Glu61Ile | 1.10 | 0.14 | Neutral | | | | 1.10 | 0.14 | Neutral |
| 61 | p.Glu61Met | 1.15 | 0.20 | Neutral | | | | 1.15 | 0.20 | Neutral |

| | | | | | | | | | | |
|---|---|---|---|---|---|---|---|---|---|---|
| 61 | p.Glu61His | 1.04 | 0.05 | Neutral | | | | 1.04 | 0.05 | Neutral |
| 61 | p.Glu61Gln | 0.84 | -0.26 | Neutral | | | | 0.84 | -0.26 | Neutral |
| 61 | p.Glu61Pro | 6.09 | 2.61 | Deleterious | | | | 6.09 | 2.61 | Deleterious |
| 61 | p.Glu61Leu | 0.92 | -0.12 | Neutral | | | | 0.92 | -0.12 | Neutral |
| 61 | p.Glu61Asp | 0.94 | -0.09 | Neutral | | | | 0.94 | -0.09 | Neutral |
| 61 | p.Glu61Glu | 1.00 | 0.00 | Neutral | | | | 1.00 | 0.00 | Neutral |
| 61 | p.Glu61Ala | 0.91 | -0.13 | Neutral | | | | 0.91 | -0.13 | Neutral |
| 61 | p.Glu61Gly | 0.83 | -0.27 | Neutral | | | | 0.83 | -0.27 | Neutral |
| 61 | p.Glu61Val | 0.87 | -0.19 | Neutral | | | | 0.87 | -0.19 | Neutral |
| 61 | p.Glu61Tyr | 1.21 | 0.28 | Indeterminate | | | | 1.21 | 0.28 | Indeterminate |
| 61 | p.Glu61Cys | 0.91 | -0.14 | Neutral | | | | 0.91 | -0.14 | Neutral |
| 61 | p.Glu61Trp | 1.04 | 0.05 | Neutral | | | | 1.04 | 0.05 | Neutral |
| 61 | p.Glu61Phe | 0.97 | -0.04 | Neutral | | | | 0.97 | -0.04 | Neutral |
| 62 | p.Leu62Asn | 0.67 | -0.58 | Neutral | 0.80 | -0.32 | Neutral | 0.74 | -0.44 | Neutral |
| 62 | p.Leu62Lys | 1.33 | 0.41 | Indeterminate | 0.82 | -0.29 | Neutral | 1.07 | 0.10 | Neutral |
| 62 | p.Leu62Thr | 1.04 | 0.06 | Neutral | 0.79 | -0.34 | Neutral | 0.92 | -0.13 | Neutral |
| 62 | p.Leu62Arg | 2.27 | 1.18 | Deleterious | 0.53 | -0.92 | Neutral | 1.40 | 0.48 | Indeterminate |
| 62 | p.Leu62Ser | 0.86 | -0.22 | Neutral | 0.76 | -0.40 | Neutral | 0.81 | -0.31 | Neutral |
| 62 | p.Leu62Ile | 1.09 | 0.12 | Neutral | 0.45 | -1.14 | Neutral | 0.77 | -0.38 | Neutral |
| 62 | p.Leu62Met | 0.84 | -0.26 | Neutral | 0.53 | -0.93 | Neutral | 0.68 | -0.56 | Neutral |
| 62 | p.Leu62His | 0.70 | -0.51 | Neutral | 0.89 | -0.16 | Neutral | 0.80 | -0.32 | Neutral |
| 62 | p.Leu62Gln | 3.81 | 1.93 | Deleterious | 1.38 | 0.47 | Indeterminate | 2.60 | 1.38 | Deleterious |
| 62 | p.Leu62Pro | 0.43 | -1.20 | Neutral | 0.25 | -2.03 | Neutral | 0.34 | -1.56 | Neutral |
| 62 | p.Leu62Leu | 1.00 | 0.00 | Neutral | 1.00 | 0.00 | Neutral | 1.00 | 0.00 | Neutral |
| 62 | p.Leu62Asp | 1.05 | 0.07 | Neutral | 1.09 | 0.13 | Neutral | 1.07 | 0.10 | Neutral |
| 62 | p.Leu62Glu | 1.16 | 0.21 | Neutral | 0.44 | -1.19 | Neutral | 0.80 | -0.32 | Neutral |
| 62 | p.Leu62Ala | 0.94 | -0.09 | Neutral | 0.47 | -1.08 | Neutral | 0.71 | -0.50 | Neutral |
| 62 | p.Leu62Gly | 1.44 | 0.52 | Indeterminate | 1.17 | 0.23 | Indeterminate | 1.30 | 0.38 | Indeterminate |
| 62 | p.Leu62Val | 1.10 | 0.14 | Neutral | 0.58 | -0.80 | Neutral | 0.84 | -0.25 | Neutral |
| 62 | p.Leu62Tyr | 1.76 | 0.82 | Indeterminate | 0.96 | -0.06 | Neutral | 1.36 | 0.44 | Indeterminate |
| 62 | p.Leu62Cys | 0.91 | -0.14 | Neutral | 0.83 | -0.27 | Neutral | 0.87 | -0.20 | Neutral |
| 62 | p.Leu62Trp | 0.84 | -0.25 | Neutral | 0.89 | -0.16 | Neutral | 0.87 | -0.20 | Neutral |
| 62 | p.Leu62Phe | 0.64 | -0.64 | Neutral | 0.71 | -0.49 | Neutral | 0.68 | -0.56 | Neutral |
| 63 | p.Leu63Asn | 13.29 | 3.73 | Deleterious | | | | 13.29 | 3.73 | Deleterious |
| 63 | p.Leu63Lys | 23.24 | 4.54 | Deleterious | | | | 23.24 | 4.54 | Deleterious |
| 63 | p.Leu63Thr | 12.62 | 3.66 | Deleterious | | | | 12.62 | 3.66 | Deleterious |
| 63 | p.Leu63Arg | 17.32 | 4.11 | Deleterious | | | | 17.32 | 4.11 | Deleterious |
| 63 | p.Leu63Ser | 15.92 | 3.99 | Deleterious | | | | 15.92 | 3.99 | Deleterious |
| 63 | p.Leu63Ile | 1.41 | 0.49 | Indeterminate | | | | 1.41 | 0.49 | Indeterminate |
| 63 | p.Leu63Met | 0.85 | -0.23 | Neutral | | | | 0.85 | -0.23 | Neutral |
| 63 | p.Leu63His | 16.94 | 4.08 | Deleterious | | | | 16.94 | 4.08 | Deleterious |
| 63 | p.Leu63Gln | 19.33 | 4.27 | Deleterious | | | | 19.33 | 4.27 | Deleterious |
| 63 | p.Leu63Pro | 20.64 | 4.37 | Deleterious | | | | 20.64 | 4.37 | Deleterious |
| 63 | p.Leu63Leu | 1.00 | 0.00 | Neutral | | | | 1.00 | 0.00 | Neutral |
| 63 | p.Leu63Asp | 31.92 | 5.00 | Deleterious | | | | 31.92 | 5.00 | Deleterious |
| 63 | p.Leu63Glu | 14.81 | 3.89 | Deleterious | | | | 14.81 | 3.89 | Deleterious |
| 63 | p.Leu63Ala | 15.76 | 3.98 | Deleterious | | | | 15.76 | 3.98 | Deleterious |
| 63 | p.Leu63Gly | 19.84 | 4.31 | Deleterious | | | | 19.84 | 4.31 | Deleterious |
| 63 | p.Leu63Val | 3.31 | 1.73 | Deleterious | | | | 3.31 | 1.73 | Deleterious |
| 63 | p.Leu63Tyr | 16.24 | 4.02 | Deleterious | | | | 16.24 | 4.02 | Deleterious |
| 63 | p.Leu63Cys | 3.30 | 1.72 | Deleterious | | | | 3.30 | 1.72 | Deleterious |
| 63 | p.Leu63Trp | 16.99 | 4.09 | Deleterious | | | | 16.99 | 4.09 | Deleterious |
| 63 | p.Leu63Phe | 2.23 | 1.16 | Deleterious | | | | 2.23 | 1.16 | Deleterious |
| 64 | p.Leu64Asn | 1.51 | 0.59 | Indeterminate | | | | 1.51 | 0.59 | Indeterminate |
| 64 | p.Leu64Lys | 1.15 | 0.20 | Neutral | | | | 1.15 | 0.20 | Neutral |
| 64 | p.Leu64Thr | 0.91 | -0.14 | Neutral | | | | 0.91 | -0.14 | Neutral |
| 64 | p.Leu64Arg | 1.25 | 0.32 | Indeterminate | | | | 1.25 | 0.32 | Indeterminate |
| 64 | p.Leu64Ser | 0.89 | -0.16 | Neutral | | | | 0.89 | -0.16 | Neutral |
| 64 | p.Leu64Ile | 1.17 | 0.22 | Neutral | | | | 1.17 | 0.22 | Neutral |
| 64 | p.Leu64Met | 0.97 | -0.05 | Neutral | | | | 0.97 | -0.05 | Neutral |
| 64 | p.Leu64His | 1.25 | 0.32 | Indeterminate | | | | 1.25 | 0.32 | Indeterminate |
| 64 | p.Leu64Gln | 1.34 | 0.43 | Indeterminate | | | | 1.34 | 0.43 | Indeterminate |
| 64 | p.Leu64Pro | 3.96 | 1.99 | Deleterious | | | | 3.96 | 1.99 | Deleterious |
| 64 | p.Leu64Leu | 1.00 | 0.00 | Neutral | | | | 1.00 | 0.00 | Neutral |
| 64 | p.Leu64Asp | 1.80 | 0.85 | Indeterminate | | | | 1.80 | 0.85 | Indeterminate |
| 64 | p.Leu64Glu | 1.41 | 0.50 | Indeterminate | | | | 1.41 | 0.50 | Indeterminate |
| 64 | p.Leu64Ala | 1.14 | 0.19 | Neutral | | | | 1.14 | 0.19 | Neutral |
| 64 | p.Leu64Gly | 1.37 | 0.46 | Indeterminate | | | | 1.37 | 0.46 | Indeterminate |
| 64 | p.Leu64Val | 0.81 | -0.30 | Neutral | | | | 0.81 | -0.30 | Neutral |
| 64 | p.Leu64Tyr | 1.01 | 0.02 | Neutral | | | | 1.01 | 0.02 | Neutral |
| 64 | p.Leu64Cys | 0.97 | -0.05 | Neutral | | | | 0.97 | -0.05 | Neutral |
| 64 | p.Leu64Trp | 1.09 | 0.12 | Neutral | | | | 1.09 | 0.12 | Neutral |
| 64 | p.Leu64Phe | 0.83 | -0.27 | Neutral | | | | 0.83 | -0.27 | Neutral |
| 65 | p.Leu65Asn | 1.41 | 0.50 | Indeterminate | | | | 1.41 | 0.50 | Indeterminate |
| 65 | p.Leu65Lys | 0.82 | -0.28 | Neutral | | | | 0.82 | -0.28 | Neutral |
| 65 | p.Leu65Thr | 1.18 | 0.23 | Neutral | | | | 1.18 | 0.23 | Neutral |
| 65 | p.Leu65Arg | 0.95 | -0.07 | Neutral | | | | 0.95 | -0.07 | Neutral |
| 65 | p.Leu65Ser | 1.22 | 0.29 | Indeterminate | | | | 1.22 | 0.29 | Indeterminate |
| 65 | p.Leu65Ile | 0.97 | -0.04 | Neutral | | | | 0.97 | -0.04 | Neutral |
| 65 | p.Leu65Met | 0.81 | -0.30 | Neutral | | | | 0.81 | -0.30 | Neutral |
| 65 | p.Leu65His | 1.30 | 0.38 | Indeterminate | | | | 1.30 | 0.38 | Indeterminate |
| 65 | p.Leu65Gln | 1.26 | 0.33 | Indeterminate | | | | 1.26 | 0.33 | Indeterminate |
| 65 | p.Leu65Pro | 11.28 | 3.50 | Deleterious | | | | 11.28 | 3.50 | Deleterious |
| 65 | p.Leu65Leu | 1.00 | 0.00 | Neutral | | | | 1.00 | 0.00 | Neutral |
| 65 | p.Leu65Asp | 0.96 | -0.06 | Neutral | | | | 0.96 | -0.06 | Neutral |
| 65 | p.Leu65Glu | 0.90 | -0.14 | Neutral | | | | 0.90 | -0.14 | Neutral |
| 65 | p.Leu65Ala | 1.18 | 0.23 | Neutral | | | | 1.18 | 0.23 | Neutral |
| 65 | p.Leu65Gly | 1.49 | 0.57 | Indeterminate | | | | 1.49 | 0.57 | Indeterminate |
| 65 | p.Leu65Val | 1.11 | 0.15 | Neutral | | | | 1.11 | 0.15 | Neutral |
| 65 | p.Leu65Tyr | 0.99 | -0.01 | Neutral | | | | 0.99 | -0.01 | Neutral |
| 65 | p.Leu65Cys | 1.57 | 0.65 | Indeterminate | | | | 1.57 | 0.65 | Indeterminate |
| 65 | p.Leu65Trp | 1.40 | 0.49 | Indeterminate | | | | 1.40 | 0.49 | Indeterminate |
| 65 | p.Leu65Phe | 0.97 | -0.05 | Neutral | | | | 0.97 | -0.05 | Neutral |
| 66 | p.His66Asn | 0.87 | -0.21 | Neutral | | | | 0.87 | -0.21 | Neutral |
| 66 | p.His66Lys | 0.89 | -0.16 | Neutral | | | | 0.89 | -0.16 | Neutral |
| 66 | p.His66Thr | 1.15 | 0.20 | Neutral | | | | 1.15 | 0.20 | Neutral |
| 66 | p.His66Arg | 1.21 | 0.28 | Indeterminate | | | | 1.21 | 0.28 | Indeterminate |
| 66 | p.His66Ser | 1.03 | 0.05 | Neutral | | | | 1.03 | 0.05 | Neutral |
| 66 | p.His66Ile | 1.57 | 0.65 | Indeterminate | | | | 1.57 | 0.65 | Indeterminate |
| 66 | p.His66Met | 0.91 | -0.14 | Neutral | | | | 0.91 | -0.14 | Neutral |
| 66 | p.His66His | 1.00 | 0.00 | Neutral | | | | 1.00 | 0.00 | Neutral |

| | | | | | | | |
|---|---|---|---|---|---|---|---|
| 66 | p.His66Gln | 1.14 | 0.19 | Neutral | 1.14 | 0.19 | Neutral |
| 66 | p.His66Pro | 18.02 | 4.17 | Deleterious | 18.02 | 4.17 | Deleterious |
| 66 | p.His66Leu | 1.18 | 0.24 | Indeterminate | 1.18 | 0.24 | Indeterminate |
| 66 | p.His66Asp | 0.96 | -0.06 | Neutral | 0.96 | -0.06 | Neutral |
| 66 | p.His66Glu | 1.34 | 0.42 | Indeterminate | 1.34 | 0.42 | Indeterminate |
| 66 | p.His66Ala | 1.24 | 0.31 | Indeterminate | 1.24 | 0.31 | Indeterminate |
| 66 | p.His66Gly | 1.08 | 0.12 | Neutral | 1.08 | 0.12 | Neutral |
| 66 | p.His66Val | 1.40 | 0.49 | Indeterminate | 1.40 | 0.49 | Indeterminate |
| 66 | p.His66Tyr | 1.05 | 0.08 | Neutral | 1.05 | 0.08 | Neutral |
| 66 | p.His66Cys | 0.69 | -0.53 | Neutral | 0.69 | -0.53 | Neutral |
| 66 | p.His66Trp | 0.97 | -0.04 | Neutral | 0.97 | -0.04 | Neutral |
| 66 | p.His66Phe | 1.00 | 0.00 | Neutral | 1.00 | 0.00 | Neutral |
| 67 | p.Gly67Asn | 0.80 | -0.33 | Neutral | 0.80 | -0.33 | Neutral |
| 67 | p.Gly67Lys | 1.04 | 0.05 | Neutral | 1.04 | 0.05 | Neutral |
| 67 | p.Gly67Thr | 1.66 | 0.73 | Indeterminate | 1.66 | 0.73 | Indeterminate |
| 67 | p.Gly67Arg | 0.67 | -0.57 | Neutral | 0.67 | -0.57 | Neutral |
| 67 | p.Gly67Ser | 0.75 | -0.42 | Neutral | 0.75 | -0.42 | Neutral |
| 67 | p.Gly67Ile | 7.82 | 2.97 | Deleterious | 7.82 | 2.97 | Deleterious |
| 67 | p.Gly67Met | 1.18 | 0.24 | Neutral | 1.18 | 0.24 | Neutral |
| 67 | p.Gly67His | 0.76 | -0.40 | Neutral | 0.76 | -0.40 | Neutral |
| 67 | p.Gly67Gln | 0.61 | -0.71 | Neutral | 0.61 | -0.71 | Neutral |
| 67 | p.Gly67Pro | 15.51 | 3.95 | Deleterious | 15.51 | 3.95 | Deleterious |
| 67 | p.Gly67Leu | 0.96 | -0.05 | Neutral | 0.96 | -0.05 | Neutral |
| 67 | p.Gly67Asp | 0.82 | -0.28 | Neutral | 0.82 | -0.28 | Neutral |
| 67 | p.Gly67Glu | 0.81 | -0.30 | Neutral | 0.81 | -0.30 | Neutral |
| 67 | p.Gly67Ala | 0.64 | -0.63 | Neutral | 0.64 | -0.63 | Neutral |
| 67 | p.Gly67Gly | 1.00 | 0.00 | Neutral | 1.00 | 0.00 | Neutral |
| 67 | p.Gly67Val | 4.79 | 2.26 | Deleterious | 4.79 | 2.26 | Deleterious |
| 67 | p.Gly67Tyr | 0.91 | -0.14 | Neutral | 0.91 | -0.14 | Neutral |
| 67 | p.Gly67Cys | 1.14 | 0.19 | Neutral | 1.14 | 0.19 | Neutral |
| 67 | p.Gly67Trp | 1.46 | 0.54 | Indeterminate | 1.46 | 0.54 | Indeterminate |
| 67 | p.Gly67Phe | 1.17 | 0.23 | Neutral | 1.17 | 0.23 | Neutral |
| 68 | p.Ala68Asn | 3.18 | 1.67 | Deleterious | 3.18 | 1.67 | Deleterious |
| 68 | p.Ala68Lys | 4.02 | 2.01 | Deleterious | 4.02 | 2.01 | Deleterious |
| 68 | p.Ala68Thr | 1.35 | 0.43 | Indeterminate | 1.35 | 0.43 | Indeterminate |
| 68 | p.Ala68Arg | 4.05 | 2.02 | Deleterious | 4.05 | 2.02 | Deleterious |
| 68 | p.Ala68Ser | 0.96 | -0.05 | Neutral | 0.96 | -0.05 | Neutral |
| 68 | p.Ala68Ile | 3.27 | 1.71 | Deleterious | 3.27 | 1.71 | Deleterious |
| 68 | p.Ala68Met | 2.86 | 1.51 | Deleterious | 2.86 | 1.51 | Deleterious |
| 68 | p.Ala68His | 2.14 | 1.10 | Deleterious | 2.14 | 1.10 | Deleterious |
| 68 | p.Ala68Gln | 3.25 | 1.70 | Deleterious | 3.25 | 1.70 | Deleterious |
| 68 | p.Ala68Pro | 3.53 | 1.82 | Deleterious | 3.53 | 1.82 | Deleterious |
| 68 | p.Ala68Leu | 3.66 | 1.87 | Deleterious | 3.66 | 1.87 | Deleterious |
| 68 | p.Ala68Asp | 2.46 | 1.30 | Deleterious | 2.46 | 1.30 | Deleterious |
| 68 | p.Ala68Glu | 2.62 | 1.39 | Deleterious | 2.62 | 1.39 | Deleterious |
| 68 | p.Ala68Ala | 1.00 | 0.00 | Neutral | 1.00 | 0.00 | Neutral |
| 68 | p.Ala68Gly | 1.07 | 0.10 | Neutral | 1.07 | 0.10 | Neutral |
| 68 | p.Ala68Val | 1.78 | 0.83 | Indeterminate | 1.78 | 0.83 | Indeterminate |
| 68 | p.Ala68Tyr | 3.77 | 1.91 | Deleterious | 3.77 | 1.91 | Deleterious |
| 68 | p.Ala68Cys | 1.13 | 0.17 | Neutral | 1.13 | 0.17 | Neutral |
| 68 | p.Ala68Trp | 4.01 | 2.01 | Deleterious | 4.01 | 2.01 | Deleterious |
| 68 | p.Ala68Phe | 3.65 | 1.87 | Deleterious | 3.65 | 1.87 | Deleterious |
| 69 | p.Glu69Asn | 2.72 | 1.44 | Deleterious | 2.72 | 1.44 | Deleterious |
| 69 | p.Glu69Lys | 2.34 | 1.23 | Deleterious | 2.34 | 1.23 | Deleterious |
| 69 | p.Glu69Thr | 1.54 | 0.62 | Indeterminate | 1.54 | 0.62 | Indeterminate |
| 69 | p.Glu69Arg | 1.08 | 0.10 | Neutral | 1.08 | 0.10 | Neutral |
| 69 | p.Glu69Ser | 1.80 | 0.85 | Indeterminate | 1.80 | 0.85 | Indeterminate |
| 69 | p.Glu69Ile | 1.34 | 0.43 | Indeterminate | 1.34 | 0.43 | Indeterminate |
| 69 | p.Glu69Met | 2.32 | 1.21 | Deleterious | 2.32 | 1.21 | Deleterious |
| 69 | p.Glu69His | 1.43 | 0.52 | Indeterminate | 1.43 | 0.52 | Indeterminate |
| 69 | p.Glu69Gln | 1.35 | 0.43 | Indeterminate | 1.35 | 0.43 | Indeterminate |
| 69 | p.Glu69Pro | 4.86 | 2.28 | Deleterious | 4.86 | 2.28 | Deleterious |
| 69 | p.Glu69Leu | 2.65 | 1.40 | Deleterious | 2.65 | 1.40 | Deleterious |
| 69 | p.Glu69Asp | 1.27 | 0.35 | Indeterminate | 1.27 | 0.35 | Indeterminate |
| 69 | p.Glu69Glu | 1.00 | 0.00 | Neutral | 1.00 | 0.00 | Neutral |
| 69 | p.Glu69Ala | 0.81 | -0.30 | Neutral | 0.81 | -0.30 | Neutral |
| 69 | p.Glu69Gly | 1.67 | 0.74 | Indeterminate | 1.67 | 0.74 | Indeterminate |
| 69 | p.Glu69Val | 2.17 | 1.11 | Deleterious | 2.17 | 1.11 | Deleterious |
| 69 | p.Glu69Tyr | 1.41 | 0.49 | Indeterminate | 1.41 | 0.49 | Indeterminate |
| 69 | p.Glu69Cys | 1.41 | 0.49 | Indeterminate | 1.41 | 0.49 | Indeterminate |
| 69 | p.Glu69Trp | 1.92 | 0.94 | Indeterminate | 1.92 | 0.94 | Indeterminate |
| 69 | p.Glu69Phe | 1.60 | 0.68 | Indeterminate | 1.60 | 0.68 | Indeterminate |
| 70 | p.Pro70Asn | 1.38 | 0.46 | Indeterminate | 1.38 | 0.46 | Indeterminate |
| 70 | p.Pro70Lys | 1.94 | 0.96 | Indeterminate | 1.94 | 0.96 | Indeterminate |
| 70 | p.Pro70Thr | 1.18 | 0.24 | Neutral | 1.18 | 0.24 | Neutral |
| 70 | p.Pro70Arg | 1.09 | 0.13 | Neutral | 1.09 | 0.13 | Neutral |
| 70 | p.Pro70Ser | 2.28 | 1.19 | Deleterious | 2.28 | 1.19 | Deleterious |
| 70 | p.Pro70Ile | 0.96 | -0.07 | Neutral | 0.96 | -0.07 | Neutral |
| 70 | p.Pro70Met | 0.51 | -0.97 | Neutral | 0.51 | -0.97 | Neutral |
| 70 | p.Pro70His | 0.47 | -1.09 | Neutral | 0.47 | -1.09 | Neutral |
| 70 | p.Pro70Gln | 0.43 | -1.23 | Neutral | 0.43 | -1.23 | Neutral |
| 70 | p.Pro70Pro | 1.00 | 0.00 | Neutral | 1.00 | 0.00 | Neutral |
| 70 | p.Pro70Leu | 1.23 | 0.30 | Indeterminate | 1.23 | 0.30 | Indeterminate |
| 70 | p.Pro70Asp | 10.85 | 3.44 | Deleterious | 10.85 | 3.44 | Deleterious |
| 70 | p.Pro70Glu | 5.22 | 2.38 | Deleterious | 5.22 | 2.38 | Deleterious |
| 70 | p.Pro70Ala | 1.00 | 0.00 | Neutral | 1.00 | 0.00 | Neutral |
| 70 | p.Pro70Gly | 1.03 | 0.05 | Neutral | 1.03 | 0.05 | Neutral |
| 70 | p.Pro70Val | 0.69 | -0.54 | Neutral | 0.69 | -0.54 | Neutral |
| 70 | p.Pro70Tyr | 0.96 | -0.07 | Neutral | 0.96 | -0.07 | Neutral |
| 70 | p.Pro70Cys | 0.40 | -1.32 | Neutral | 0.40 | -1.32 | Neutral |
| 70 | p.Pro70Trp | 3.19 | 1.67 | Deleterious | 3.19 | 1.67 | Deleterious |
| 70 | p.Pro70Phe | 0.85 | -0.23 | Neutral | 0.85 | -0.23 | Neutral |
| 71 | p.Asn71Asn | 1.00 | 0.00 | Neutral | 1.00 | 0.00 | Neutral |
| 71 | p.Asn71Lys | 17.91 | 4.16 | Deleterious | 17.91 | 4.16 | Deleterious |
| 71 | p.Asn71Thr | 3.40 | 1.77 | Deleterious | 3.40 | 1.77 | Deleterious |
| 71 | p.Asn71Arg | 13.92 | 3.80 | Deleterious | 13.92 | 3.80 | Deleterious |
| 71 | p.Asn71Ser | 30.08 | 4.91 | Deleterious | 30.08 | 4.91 | Deleterious |
| 71 | p.Asn71Ile | 39.18 | 5.29 | Deleterious | 39.18 | 5.29 | Deleterious |
| 71 | p.Asn71Met | 5.00 | 2.32 | Deleterious | 5.00 | 2.32 | Deleterious |
| 71 | p.Asn71His | 1.98 | 0.99 | Indeterminate | 1.98 | 0.99 | Indeterminate |
| 71 | p.Asn71Gln | 1.66 | 0.73 | Indeterminate | 1.66 | 0.73 | Indeterminate |

| | | | | | | | | | | |
|---|---|---|---|---|---|---|---|---|---|---|
| 71 | p.Asn71Pro | 84.59 | 6.40 | Deleterious | | | | 84.59 | 6.40 | Deleterious |
| 71 | p.Asn71Leu | 32.49 | 5.02 | Deleterious | | | | 32.49 | 5.02 | Deleterious |
| 71 | p.Asn71Asp | 2.31 | 1.21 | Deleterious | | | | 2.31 | 1.21 | Deleterious |
| 71 | p.Asn71Glu | 1.89 | 0.92 | Indeterminate | | | | 1.89 | 0.92 | Indeterminate |
| 71 | p.Asn71Ala | 2.26 | 1.18 | Deleterious | | | | 2.26 | 1.18 | Deleterious |
| 71 | p.Asn71Gly | 1.06 | 0.08 | Neutral | | | | 1.06 | 0.08 | Neutral |
| 71 | p.Asn71Val | 28.77 | 4.85 | Deleterious | | | | 28.77 | 4.85 | Deleterious |
| 71 | p.Asn71Tyr | 17.62 | 4.14 | Deleterious | | | | 17.62 | 4.14 | Deleterious |
| 71 | p.Asn71Cys | 1.77 | 0.82 | Indeterminate | | | | 1.77 | 0.82 | Indeterminate |
| 71 | p.Asn71Trp | 6.47 | 2.69 | Deleterious | | | | 6.47 | 2.69 | Deleterious |
| 71 | p.Asn71Phe | 17.55 | 4.13 | Deleterious | | | | 17.55 | 4.13 | Deleterious |
| 72 | p.Cys72Asn | 1.42 | 0.51 | Indeterminate | 0.93 | -0.10 | Neutral | 1.18 | 0.24 | Neutral |
| 72 | p.Cys72Lys | 1.32 | 0.40 | Indeterminate | 1.04 | 0.06 | Neutral | 1.18 | 0.24 | Neutral |
| 72 | p.Cys72Thr | 1.37 | 0.45 | Indeterminate | 0.85 | -0.23 | Neutral | 1.11 | 0.15 | Neutral |
| 72 | p.Cys72Arg | 0.90 | -0.16 | Neutral | 0.70 | -0.51 | Neutral | 0.80 | -0.32 | Neutral |
| 72 | p.Cys72Ser | 1.29 | 0.37 | Indeterminate | 1.21 | 0.28 | Indeterminate | 1.25 | 0.32 | Indeterminate |
| 72 | p.Cys72Ile | 1.37 | 0.46 | Indeterminate | 1.12 | 0.17 | Indeterminate | 1.25 | 0.32 | Indeterminate |
| 72 | p.Cys72Met | 1.00 | 0.01 | Neutral | 1.03 | 0.04 | Neutral | 1.01 | 0.02 | Neutral |
| 72 | p.Cys72His | 1.08 | 0.11 | Neutral | 1.11 | 0.15 | Indeterminate | 1.10 | 0.13 | Neutral |
| 72 | p.Cys72Gln | 1.57 | 0.65 | Indeterminate | 0.82 | -0.29 | Neutral | 1.19 | 0.26 | Indeterminate |
| 72 | p.Cys72Pro | 1.59 | 0.67 | Indeterminate | 0.82 | -0.29 | Neutral | 1.20 | 0.26 | Indeterminate |
| 72 | p.Cys72Leu | 1.44 | 0.53 | Indeterminate | 1.14 | 0.19 | Indeterminate | 1.29 | 0.37 | Indeterminate |
| 72 | p.Cys72Asp | 1.02 | 0.03 | Neutral | 1.28 | 0.35 | Indeterminate | 1.15 | 0.20 | Neutral |
| 72 | p.Cys72Glu | 1.20 | 0.27 | Indeterminate | 1.23 | 0.30 | Indeterminate | 1.22 | 0.28 | Indeterminate |
| 72 | p.Cys72Ala | 1.09 | 0.12 | Neutral | 1.24 | 0.31 | Indeterminate | 1.16 | 0.22 | Neutral |
| 72 | p.Cys72Gly | 1.05 | 0.08 | Neutral | 0.60 | -0.74 | Neutral | 0.83 | -0.27 | Neutral |
| 72 | p.Cys72Val | 1.33 | 0.42 | Indeterminate | 1.29 | 0.36 | Indeterminate | 1.31 | 0.39 | Indeterminate |
| 72 | p.Cys72Tyr | 1.12 | 0.17 | Neutral | 0.96 | -0.06 | Neutral | 1.04 | 0.06 | Neutral |
| 72 | p.Cys72Cys | 1.00 | 0.00 | Neutral | 1.00 | 0.00 | Neutral | 1.00 | 0.00 | Neutral |
| 72 | p.Cys72Trp | 1.78 | 0.83 | Indeterminate | 1.06 | 0.09 | Neutral | 1.42 | 0.51 | Indeterminate |
| 72 | p.Cys72Phe | 0.99 | -0.01 | Neutral | 1.57 | 0.65 | Indeterminate | 1.28 | 0.35 | Indeterminate |
| 73 | p.Ala73Asn | 1.22 | 0.29 | Indeterminate | 1.06 | 0.09 | Neutral | 1.14 | 0.19 | Neutral |
| 73 | p.Ala73Lys | 0.77 | -0.37 | Neutral | 1.02 | 0.02 | Neutral | 0.90 | -0.16 | Neutral |
| 73 | p.Ala73Thr | 1.07 | 0.10 | Neutral | 0.92 | -0.11 | Neutral | 1.00 | 0.00 | Neutral |
| 73 | p.Ala73Arg | 1.17 | 0.23 | Neutral | 0.93 | -0.10 | Neutral | 1.05 | 0.07 | Neutral |
| 73 | p.Ala73Ser | 1.11 | 0.15 | Neutral | 1.00 | -0.01 | Neutral | 1.05 | 0.07 | Neutral |
| 73 | p.Ala73Ile | 0.81 | -0.30 | Neutral | 0.97 | -0.04 | Neutral | 0.89 | -0.17 | Neutral |
| 73 | p.Ala73Met | 0.82 | -0.29 | Neutral | 1.00 | 0.00 | Neutral | 0.91 | -0.14 | Neutral |
| 73 | p.Ala73His | 0.90 | -0.15 | Neutral | 0.95 | -0.07 | Neutral | 0.93 | -0.11 | Neutral |
| 73 | p.Ala73Gln | 1.03 | 0.05 | Neutral | 0.95 | -0.07 | Neutral | 0.99 | -0.01 | Neutral |
| 73 | p.Ala73Pro | 1.00 | 0.00 | Neutral | 1.08 | 0.11 | Neutral | 1.04 | 0.06 | Neutral |
| 73 | p.Ala73Leu | 0.85 | -0.23 | Neutral | 0.99 | -0.01 | Neutral | 0.92 | -0.12 | Neutral |
| 73 | p.Ala73Asp | 1.01 | 0.02 | Neutral | 0.90 | -0.16 | Neutral | 0.95 | -0.07 | Neutral |
| 73 | p.Ala73Glu | 0.93 | -0.11 | Neutral | 1.02 | 0.03 | Neutral | 0.97 | -0.04 | Neutral |
| 73 | p.Ala73Ala | 1.00 | 0.00 | Neutral | 1.00 | 0.00 | Neutral | 1.00 | 0.00 | Neutral |
| 73 | p.Ala73Gly | 1.11 | 0.15 | Neutral | 1.01 | 0.01 | Neutral | 1.06 | 0.08 | Neutral |
| 73 | p.Ala73Val | 1.12 | 0.17 | Neutral | 0.88 | -0.18 | Neutral | 1.00 | 0.00 | Neutral |
| 73 | p.Ala73Tyr | 1.08 | 0.11 | Neutral | 1.16 | 0.21 | Indeterminate | 1.12 | 0.16 | Neutral |
| 73 | p.Ala73Cys | 1.09 | 0.12 | Neutral | 0.97 | -0.04 | Neutral | 1.03 | 0.04 | Neutral |
| 73 | p.Ala73Trp | 1.20 | 0.26 | Indeterminate | 0.97 | -0.05 | Neutral | 1.08 | 0.11 | Neutral |
| 73 | p.Ala73Phe | 0.88 | -0.19 | Neutral | 1.02 | 0.03 | Neutral | 0.95 | -0.08 | Neutral |
| 74 | p.Asp74Asn | 1.63 | 0.70 | Indeterminate | 1.89 | 0.92 | Indeterminate | 1.76 | 0.82 | Indeterminate |
| 74 | p.Asp74Lys | 12.10 | 3.60 | Deleterious | 7.36 | 2.88 | Deleterious | 9.73 | 3.28 | Deleterious |
| 74 | p.Asp74Thr | 5.11 | 2.35 | Deleterious | 3.90 | 1.97 | Deleterious | 4.51 | 2.17 | Deleterious |
| 74 | p.Asp74Arg | 11.51 | 3.52 | Deleterious | 6.76 | 2.76 | Deleterious | 9.14 | 3.19 | Deleterious |
| 74 | p.Asp74Ser | 1.13 | 0.18 | Neutral | 1.57 | 0.65 | Indeterminate | 1.35 | 0.43 | Indeterminate |
| 74 | p.Asp74Ile | 13.43 | 3.75 | Deleterious | 5.56 | 2.47 | Deleterious | 9.50 | 3.25 | Deleterious |
| 74 | p.Asp74Met | 6.01 | 2.59 | Deleterious | 4.26 | 2.09 | Deleterious | 5.13 | 2.36 | Deleterious |
| 74 | p.Asp74His | 1.14 | 0.19 | Neutral | 1.02 | 0.02 | Neutral | 1.08 | 0.11 | Neutral |
| 74 | p.Asp74Gln | 6.49 | 2.70 | Deleterious | 3.93 | 1.97 | Deleterious | 5.21 | 2.38 | Deleterious |
| 74 | p.Asp74Pro | 12.91 | 3.69 | Deleterious | 6.59 | 2.72 | Deleterious | 9.75 | 3.29 | Deleterious |
| 74 | p.Asp74Leu | 7.86 | 2.98 | Deleterious | 5.09 | 2.35 | Deleterious | 6.48 | 2.70 | Deleterious |
| 74 | p.Asp74Asp | 1.00 | 0.00 | Neutral | 1.00 | 0.00 | Neutral | 1.00 | 0.00 | Neutral |
| 74 | p.Asp74Glu | 0.50 | -1.00 | Neutral | 0.41 | -1.29 | Neutral | 0.45 | -1.14 | Neutral |
| 74 | p.Asp74Ala | 3.04 | 1.60 | Deleterious | 2.47 | 1.30 | Indeterminate | 2.75 | 1.46 | Deleterious |
| 74 | p.Asp74Gly | 2.17 | 1.12 | Deleterious | 1.86 | 0.90 | Indeterminate | 2.02 | 1.01 | Indeterminate |
| 74 | p.Asp74Val | 9.32 | 3.22 | Deleterious | 5.20 | 2.38 | Deleterious | 7.26 | 2.86 | Deleterious |
| 74 | p.Asp74Tyr | 6.61 | 2.73 | Deleterious | 3.82 | 1.93 | Deleterious | 5.22 | 2.38 | Deleterious |
| 74 | p.Asp74Cys | 0.66 | -0.61 | Neutral | 0.71 | -0.50 | Neutral | 0.68 | -0.55 | Neutral |
| 74 | p.Asp74Trp | 7.09 | 2.83 | Deleterious | 4.05 | 2.02 | Deleterious | 5.57 | 2.48 | Deleterious |
| 74 | p.Asp74Phe | 8.17 | 3.03 | Deleterious | 5.14 | 2.36 | Deleterious | 6.65 | 2.73 | Deleterious |
| 75 | p.Pro75Asn | 1.03 | 0.05 | Neutral | 1.13 | 0.17 | Indeterminate | 1.08 | 0.11 | Neutral |
| 75 | p.Pro75Lys | 1.24 | 0.31 | Indeterminate | 1.10 | 0.14 | Neutral | 1.17 | 0.23 | Neutral |
| 75 | p.Pro75Thr | 1.17 | 0.22 | Neutral | 0.98 | -0.02 | Neutral | 1.07 | 0.10 | Neutral |
| 75 | p.Pro75Arg | 0.88 | -0.18 | Neutral | 1.01 | 0.02 | Neutral | 0.95 | -0.08 | Neutral |
| 75 | p.Pro75Ser | 1.15 | 0.20 | Neutral | 0.87 | -0.20 | Neutral | 1.01 | 0.01 | Neutral |
| 75 | p.Pro75Ile | 0.94 | -0.08 | Neutral | 1.18 | 0.24 | Indeterminate | 1.06 | 0.09 | Neutral |
| 75 | p.Pro75Met | 1.25 | 0.32 | Indeterminate | 1.06 | 0.09 | Neutral | 1.15 | 0.21 | Neutral |
| 75 | p.Pro75His | 0.94 | -0.08 | Neutral | 0.77 | -0.38 | Neutral | 0.86 | -0.23 | Neutral |
| 75 | p.Pro75Gln | 1.19 | 0.25 | Indeterminate | 0.99 | -0.02 | Neutral | 1.09 | 0.12 | Neutral |
| 75 | p.Pro75Pro | 1.00 | 0.00 | Neutral | 1.00 | 0.00 | Neutral | 1.00 | 0.00 | Neutral |
| 75 | p.Pro75Leu | 1.03 | 0.04 | Neutral | 0.96 | -0.06 | Neutral | 0.99 | -0.01 | Neutral |
| 75 | p.Pro75Asp | 1.03 | 0.04 | Neutral | 0.93 | -0.10 | Neutral | 0.98 | -0.03 | Neutral |
| 75 | p.Pro75Glu | 1.08 | 0.10 | Neutral | 0.90 | -0.16 | Neutral | 0.99 | -0.02 | Neutral |
| 75 | p.Pro75Ala | 1.07 | 0.10 | Neutral | 0.98 | -0.03 | Neutral | 1.03 | 0.04 | Neutral |
| 75 | p.Pro75Gly | 1.01 | 0.02 | Neutral | 0.92 | -0.13 | Neutral | 0.96 | -0.05 | Neutral |
| 75 | p.Pro75Val | 1.07 | 0.09 | Neutral | 1.10 | 0.14 | Neutral | 1.08 | 0.12 | Neutral |
| 75 | p.Pro75Tyr | 1.05 | 0.07 | Neutral | 1.06 | 0.09 | Neutral | 1.06 | 0.08 | Neutral |
| 75 | p.Pro75Cys | 1.10 | 0.14 | Neutral | 0.92 | -0.12 | Neutral | 1.01 | 0.02 | Neutral |
| 75 | p.Pro75Trp | 1.19 | 0.26 | Indeterminate | 1.12 | 0.17 | Indeterminate | 1.16 | 0.21 | Neutral |
| 75 | p.Pro75Phe | 1.06 | 0.08 | Neutral | 1.06 | 0.09 | Neutral | 1.06 | 0.08 | Neutral |
| 76 | p.Ala76Asn | 1.90 | 0.93 | Indeterminate | | | | 1.90 | 0.93 | Indeterminate |
| 76 | p.Ala76Lys | 1.76 | 0.82 | Indeterminate | | | | 1.76 | 0.82 | Indeterminate |
| 76 | p.Ala76Thr | 1.39 | 0.47 | Indeterminate | | | | 1.39 | 0.47 | Indeterminate |
| 76 | p.Ala76Arg | 1.81 | 0.86 | Indeterminate | | | | 1.81 | 0.86 | Indeterminate |
| 76 | p.Ala76Ser | 1.74 | 0.80 | Indeterminate | | | | 1.74 | 0.80 | Indeterminate |
| 76 | p.Ala76Ile | 1.84 | 0.88 | Indeterminate | | | | 1.84 | 0.88 | Indeterminate |
| 76 | p.Ala76Met | 1.38 | 0.47 | Indeterminate | | | | 1.38 | 0.47 | Indeterminate |
| 76 | p.Ala76His | 1.66 | 0.73 | Indeterminate | | | | 1.66 | 0.73 | Indeterminate |
| 76 | p.Ala76Gln | 1.79 | 0.84 | Indeterminate | | | | 1.79 | 0.84 | Indeterminate |
| 76 | p.Ala76Pro | 1.54 | 0.62 | Indeterminate | | | | 1.54 | 0.62 | Indeterminate |

| | | | | | | | |
|---|---|---|---|---|---|---|---|
| 76 | p.Ala76Leu | 1.66 | 0.73 | Indeterminate | 1.66 | 0.73 | Indeterminate |
| 76 | p.Ala76Asp | 1.55 | 0.63 | Indeterminate | 1.55 | 0.63 | Indeterminate |
| 76 | p.Ala76Glu | 1.76 | 0.82 | Indeterminate | 1.76 | 0.82 | Indeterminate |
| 76 | p.Ala76Ala | 1.00 | 0.00 | Neutral | 1.00 | 0.00 | Neutral |
| 76 | p.Ala76Gly | 1.45 | 0.53 | Indeterminate | 1.45 | 0.53 | Indeterminate |
| 76 | p.Ala76Val | 1.38 | 0.47 | Indeterminate | 1.38 | 0.47 | Indeterminate |
| 76 | p.Ala76Tyr | 1.44 | 0.52 | Indeterminate | 1.44 | 0.52 | Indeterminate |
| 76 | p.Ala76Cys | 2.00 | 1.00 | Indeterminate | 2.00 | 1.00 | Indeterminate |
| 76 | p.Ala76Trp | 1.38 | 0.47 | Indeterminate | 1.38 | 0.47 | Indeterminate |
| 76 | p.Ala76Phe | 1.62 | 0.70 | Indeterminate | 1.62 | 0.70 | Indeterminate |
| 77 | p.Thr77Asn | 1.46 | 0.55 | Indeterminate | 1.46 | 0.55 | Indeterminate |
| 77 | p.Thr77Lys | 1.55 | 0.63 | Indeterminate | 1.55 | 0.63 | Indeterminate |
| 77 | p.Thr77Thr | 1.00 | 0.00 | Neutral | 1.00 | 0.00 | Neutral |
| 77 | p.Thr77Arg | 1.70 | 0.77 | Indeterminate | 1.70 | 0.77 | Indeterminate |
| 77 | p.Thr77Ser | 1.51 | 0.60 | Indeterminate | 1.51 | 0.60 | Indeterminate |
| 77 | p.Thr77Ile | 1.27 | 0.34 | Indeterminate | 1.27 | 0.34 | Indeterminate |
| 77 | p.Thr77Met | 1.32 | 0.40 | Indeterminate | 1.32 | 0.40 | Indeterminate |
| 77 | p.Thr77His | 1.34 | 0.42 | Indeterminate | 1.34 | 0.42 | Indeterminate |
| 77 | p.Thr77Gln | 1.41 | 0.49 | Indeterminate | 1.41 | 0.49 | Indeterminate |
| 77 | p.Thr77Pro | 2.68 | 1.42 | Deleterious | 2.68 | 1.42 | Deleterious |
| 77 | p.Thr77Leu | 1.55 | 0.63 | Indeterminate | 1.55 | 0.63 | Indeterminate |
| 77 | p.Thr77Asp | 1.28 | 0.36 | Indeterminate | 1.28 | 0.36 | Indeterminate |
| 77 | p.Thr77Glu | 2.04 | 1.03 | Indeterminate | 2.04 | 1.03 | Indeterminate |
| 77 | p.Thr77Ala | 1.55 | 0.63 | Indeterminate | 1.55 | 0.63 | Indeterminate |
| 77 | p.Thr77Gly | 1.27 | 0.34 | Indeterminate | 1.27 | 0.34 | Indeterminate |
| 77 | p.Thr77Val | 1.76 | 0.82 | Indeterminate | 1.76 | 0.82 | Indeterminate |
| 77 | p.Thr77Tyr | 1.33 | 0.41 | Indeterminate | 1.33 | 0.41 | Indeterminate |
| 77 | p.Thr77Cys | 1.38 | 0.47 | Indeterminate | 1.38 | 0.47 | Indeterminate |
| 77 | p.Thr77Trp | 1.20 | 0.27 | Indeterminate | 1.20 | 0.27 | Indeterminate |
| 77 | p.Thr77Phe | 1.14 | 0.19 | Neutral | 1.14 | 0.19 | Neutral |
| 78 | p.Leu78Asn | 0.71 | -0.49 | Neutral | 0.71 | -0.49 | Neutral |
| 78 | p.Leu78Lys | 0.81 | -0.31 | Neutral | 0.81 | -0.31 | Neutral |
| 78 | p.Leu78Thr | 0.82 | -0.29 | Neutral | 0.82 | -0.29 | Neutral |
| 78 | p.Leu78Arg | 1.48 | 0.57 | Indeterminate | 1.48 | 0.57 | Indeterminate |
| 78 | p.Leu78Ser | 0.53 | -0.91 | Neutral | 0.53 | -0.91 | Neutral |
| 78 | p.Leu78Ile | 0.92 | -0.13 | Neutral | 0.92 | -0.13 | Neutral |
| 78 | p.Leu78Met | 1.36 | 0.45 | Indeterminate | 1.36 | 0.45 | Indeterminate |
| 78 | p.Leu78His | 0.98 | -0.03 | Neutral | 0.98 | -0.03 | Neutral |
| 78 | p.Leu78Gln | 1.26 | 0.33 | Indeterminate | 1.26 | 0.33 | Indeterminate |
| 78 | p.Leu78Pro | 1.60 | 0.68 | Indeterminate | 1.60 | 0.68 | Indeterminate |
| 78 | p.Leu78Leu | 1.00 | 0.00 | Neutral | 1.00 | 0.00 | Neutral |
| 78 | p.Leu78Asp | 0.45 | -1.14 | Neutral | 0.45 | -1.14 | Neutral |
| 78 | p.Leu78Glu | 1.24 | 0.31 | Indeterminate | 1.24 | 0.31 | Indeterminate |
| 78 | p.Leu78Ala | 1.23 | 0.30 | Indeterminate | 1.23 | 0.30 | Indeterminate |
| 78 | p.Leu78Gly | 0.18 | -2.45 | Neutral | 0.18 | -2.45 | Neutral |
| 78 | p.Leu78Val | 0.80 | -0.32 | Neutral | 0.80 | -0.32 | Neutral |
| 78 | p.Leu78Tyr | 1.13 | 0.18 | Neutral | 1.13 | 0.18 | Neutral |
| 78 | p.Leu78Cys | 1.03 | 0.04 | Neutral | 1.03 | 0.04 | Neutral |
| 78 | p.Leu78Trp | 0.71 | -0.49 | Neutral | 0.71 | -0.49 | Neutral |
| 78 | p.Leu78Phe | 0.67 | -0.59 | Neutral | 0.67 | -0.59 | Neutral |
| 79 | p.Thr79Asn | 0.82 | -0.29 | Neutral | 0.82 | -0.29 | Neutral |
| 79 | p.Thr79Lys | 0.95 | -0.07 | Neutral | 0.95 | -0.07 | Neutral |
| 79 | p.Thr79Thr | 1.00 | 0.00 | Neutral | 1.00 | 0.00 | Neutral |
| 79 | p.Thr79Arg | 1.24 | 0.31 | Indeterminate | 1.24 | 0.31 | Indeterminate |
| 79 | p.Thr79Ser | 0.83 | -0.26 | Neutral | 0.83 | -0.26 | Neutral |
| 79 | p.Thr79Ile | 0.78 | -0.35 | Neutral | 0.78 | -0.35 | Neutral |
| 79 | p.Thr79Met | 0.96 | -0.06 | Neutral | 0.96 | -0.06 | Neutral |
| 79 | p.Thr79His | 0.94 | -0.09 | Neutral | 0.94 | -0.09 | Neutral |
| 79 | p.Thr79Gln | 0.81 | -0.31 | Neutral | 0.81 | -0.31 | Neutral |
| 79 | p.Thr79Pro | 2.62 | 1.39 | Deleterious | 2.62 | 1.39 | Deleterious |
| 79 | p.Thr79Leu | 1.06 | 0.08 | Neutral | 1.06 | 0.08 | Neutral |
| 79 | p.Thr79Asp | 0.97 | -0.04 | Neutral | 0.97 | -0.04 | Neutral |
| 79 | p.Thr79Glu | 0.85 | -0.24 | Neutral | 0.85 | -0.24 | Neutral |
| 79 | p.Thr79Ala | 0.99 | -0.02 | Neutral | 0.99 | -0.02 | Neutral |
| 79 | p.Thr79Gly | 1.03 | 0.04 | Neutral | 1.03 | 0.04 | Neutral |
| 79 | p.Thr79Val | 1.04 | 0.06 | Neutral | 1.04 | 0.06 | Neutral |
| 79 | p.Thr79Tyr | 0.99 | -0.02 | Neutral | 0.99 | -0.02 | Neutral |
| 79 | p.Thr79Cys | 1.10 | 0.13 | Neutral | 1.10 | 0.13 | Neutral |
| 79 | p.Thr79Trp | 1.14 | 0.18 | Neutral | 1.14 | 0.18 | Neutral |
| 79 | p.Thr79Phe | 0.95 | -0.08 | Neutral | 0.95 | -0.08 | Neutral |
| 80 | p.Arg80Asn | 1.67 | 0.74 | Indeterminate | 1.67 | 0.74 | Indeterminate |
| 80 | p.Arg80Lys | 1.22 | 0.29 | Indeterminate | 1.22 | 0.29 | Indeterminate |
| 80 | p.Arg80Thr | 1.18 | 0.24 | Indeterminate | 1.18 | 0.24 | Indeterminate |
| 80 | p.Arg80Arg | 1.00 | 0.00 | Neutral | 1.00 | 0.00 | Neutral |
| 80 | p.Arg80Ser | 1.17 | 0.23 | Neutral | 1.17 | 0.23 | Neutral |
| 80 | p.Arg80Ile | 2.45 | 1.29 | Deleterious | 2.45 | 1.29 | Deleterious |
| 80 | p.Arg80Met | 1.45 | 0.54 | Indeterminate | 1.45 | 0.54 | Indeterminate |
| 80 | p.Arg80His | 1.31 | 0.39 | Indeterminate | 1.31 | 0.39 | Indeterminate |
| 80 | p.Arg80Gln | 1.54 | 0.62 | Indeterminate | 1.54 | 0.62 | Indeterminate |
| 80 | p.Arg80Pro | 4.66 | 2.22 | Deleterious | 4.66 | 2.22 | Deleterious |
| 80 | p.Arg80Leu | 1.47 | 0.56 | Indeterminate | 1.47 | 0.56 | Indeterminate |
| 80 | p.Arg80Asp | 1.61 | 0.69 | Indeterminate | 1.61 | 0.69 | Indeterminate |
| 80 | p.Arg80Glu | 1.16 | 0.21 | Neutral | 1.16 | 0.21 | Neutral |
| 80 | p.Arg80Ala | 1.32 | 0.40 | Indeterminate | 1.32 | 0.40 | Indeterminate |
| 80 | p.Arg80Gly | 1.34 | 0.42 | Indeterminate | 1.34 | 0.42 | Indeterminate |
| 80 | p.Arg80Val | 1.64 | 0.71 | Indeterminate | 1.64 | 0.71 | Indeterminate |
| 80 | p.Arg80Tyr | 1.46 | 0.55 | Indeterminate | 1.46 | 0.55 | Indeterminate |
| 80 | p.Arg80Cys | 0.88 | -0.18 | Neutral | 0.88 | -0.18 | Neutral |
| 80 | p.Arg80Trp | 1.47 | 0.55 | Indeterminate | 1.47 | 0.55 | Indeterminate |
| 80 | p.Arg80Phe | 1.55 | 0.63 | Indeterminate | 1.55 | 0.63 | Indeterminate |
| 81 | p.Pro81Asn | 4.72 | 2.24 | Deleterious | 4.72 | 2.24 | Deleterious |
| 81 | p.Pro81Lys | 6.35 | 2.67 | Deleterious | 6.35 | 2.67 | Deleterious |
| 81 | p.Pro81Thr | 2.72 | 1.44 | Deleterious | 2.72 | 1.44 | Deleterious |
| 81 | p.Pro81Arg | 4.70 | 2.23 | Deleterious | 4.70 | 2.23 | Deleterious |
| 81 | p.Pro81Ser | 3.30 | 1.72 | Deleterious | 3.30 | 1.72 | Deleterious |
| 81 | p.Pro81Ile | 3.52 | 1.81 | Deleterious | 3.52 | 1.81 | Deleterious |
| 81 | p.Pro81Met | 4.37 | 2.13 | Deleterious | 4.37 | 2.13 | Deleterious |
| 81 | p.Pro81His | 5.15 | 2.36 | Deleterious | 5.15 | 2.36 | Deleterious |
| 81 | p.Pro81Gln | 7.78 | 2.96 | Deleterious | 7.78 | 2.96 | Deleterious |
| 81 | p.Pro81Pro | 1.00 | 0.00 | Neutral | 1.00 | 0.00 | Neutral |
| 81 | p.Pro81Leu | 3.71 | 1.89 | Deleterious | 3.71 | 1.89 | Deleterious |

| | | | | | | | |
|---|---|---|---|---|---|---|---|
| 81 | p.Pro81Asp | 5.03 | 2.33 | Deleterious | 5.03 | 2.33 | Deleterious |
| 81 | p.Pro81Glu | 6.20 | 2.63 | Deleterious | 6.20 | 2.63 | Deleterious |
| 81 | p.Pro81Ala | 1.67 | 0.74 | Indeterminate | 1.67 | 0.74 | Indeterminate |
| 81 | p.Pro81Gly | 3.90 | 1.96 | Deleterious | 3.90 | 1.96 | Deleterious |
| 81 | p.Pro81Val | 2.62 | 1.39 | Deleterious | 2.62 | 1.39 | Deleterious |
| 81 | p.Pro81Tyr | 4.30 | 2.10 | Deleterious | 4.30 | 2.10 | Deleterious |
| 81 | p.Pro81Cys | 2.08 | 1.06 | Indeterminate | 2.08 | 1.06 | Indeterminate |
| 81 | p.Pro81Trp | 5.08 | 2.35 | Deleterious | 5.08 | 2.35 | Deleterious |
| 81 | p.Pro81Phe | 4.51 | 2.17 | Deleterious | 4.51 | 2.17 | Deleterious |
| 82 | p.Val82Asn | 2.35 | 1.23 | Deleterious | 2.35 | 1.23 | Deleterious |
| 82 | p.Val82Lys | 3.03 | 1.60 | Deleterious | 3.03 | 1.60 | Deleterious |
| 82 | p.Val82Thr | 1.50 | 0.58 | Indeterminate | 1.50 | 0.58 | Indeterminate |
| 82 | p.Val82Arg | 3.09 | 1.63 | Deleterious | 3.09 | 1.63 | Deleterious |
| 82 | p.Val82Ser | 1.36 | 0.44 | Indeterminate | 1.36 | 0.44 | Indeterminate |
| 82 | p.Val82Ile | 1.28 | 0.36 | Indeterminate | 1.28 | 0.36 | Indeterminate |
| 82 | p.Val82Met | 0.83 | -0.27 | Neutral | 0.83 | -0.27 | Neutral |
| 82 | p.Val82His | 2.45 | 1.29 | Deleterious | 2.45 | 1.29 | Deleterious |
| 82 | p.Val82Gln | 1.41 | 0.50 | Indeterminate | 1.41 | 0.50 | Indeterminate |
| 82 | p.Val82Pro | 2.04 | 1.03 | Indeterminate | 2.04 | 1.03 | Indeterminate |
| 82 | p.Val82Leu | 1.14 | 0.19 | Neutral | 1.14 | 0.19 | Neutral |
| 82 | p.Val82Asp | 2.78 | 1.48 | Deleterious | 2.78 | 1.48 | Deleterious |
| 82 | p.Val82Glu | 2.14 | 1.10 | Deleterious | 2.14 | 1.10 | Deleterious |
| 82 | p.Val82Ala | 1.01 | 0.01 | Neutral | 1.01 | 0.01 | Neutral |
| 82 | p.Val82Gly | 2.04 | 1.03 | Indeterminate | 2.04 | 1.03 | Indeterminate |
| 82 | p.Val82Val | 1.00 | 0.00 | Neutral | 1.00 | 0.00 | Neutral |
| 82 | p.Val82Tyr | 2.89 | 1.53 | Deleterious | 2.89 | 1.53 | Deleterious |
| 82 | p.Val82Cys | 1.33 | 0.41 | Indeterminate | 1.33 | 0.41 | Indeterminate |
| 82 | p.Val82Trp | 3.55 | 1.83 | Deleterious | 3.55 | 1.83 | Deleterious |
| 82 | p.Val82Phe | 1.80 | 0.84 | Indeterminate | 1.80 | 0.84 | Indeterminate |
| 83 | p.His83Asn | 7.57 | 2.92 | Deleterious | 7.57 | 2.92 | Deleterious |
| 83 | p.His83Lys | 10.58 | 3.40 | Deleterious | 10.58 | 3.40 | Deleterious |
| 83 | p.His83Thr | 6.28 | 2.65 | Deleterious | 6.28 | 2.65 | Deleterious |
| 83 | p.His83Arg | 11.18 | 3.48 | Deleterious | 11.18 | 3.48 | Deleterious |
| 83 | p.His83Ser | 6.08 | 2.60 | Deleterious | 6.08 | 2.60 | Deleterious |
| 83 | p.His83Ile | 6.54 | 2.71 | Deleterious | 6.54 | 2.71 | Deleterious |
| 83 | p.His83Met | 2.21 | 1.14 | Deleterious | 2.21 | 1.14 | Deleterious |
| 83 | p.His83His | 1.00 | 0.00 | Neutral | 1.00 | 0.00 | Neutral |
| 83 | p.His83Gln | 4.00 | 2.00 | Deleterious | 4.00 | 2.00 | Deleterious |
| 83 | p.His83Pro | 11.53 | 3.53 | Deleterious | 11.53 | 3.53 | Deleterious |
| 83 | p.His83Leu | 5.89 | 2.56 | Deleterious | 5.89 | 2.56 | Deleterious |
| 83 | p.His83Asp | 10.71 | 3.42 | Deleterious | 10.71 | 3.42 | Deleterious |
| 83 | p.His83Glu | 10.06 | 3.33 | Deleterious | 10.06 | 3.33 | Deleterious |
| 83 | p.His83Ala | 2.62 | 1.39 | Deleterious | 2.62 | 1.39 | Deleterious |
| 83 | p.His83Gly | 7.34 | 2.87 | Deleterious | 7.34 | 2.87 | Deleterious |
| 83 | p.His83Val | 9.70 | 3.28 | Deleterious | 9.70 | 3.28 | Deleterious |
| 83 | p.His83Tyr | 10.76 | 3.43 | Deleterious | 10.76 | 3.43 | Deleterious |
| 83 | p.His83Cys | 6.85 | 2.78 | Deleterious | 6.85 | 2.78 | Deleterious |
| 83 | p.His83Trp | 11.28 | 3.50 | Deleterious | 11.28 | 3.50 | Deleterious |
| 83 | p.His83Phe | 6.61 | 2.72 | Deleterious | 6.61 | 2.72 | Deleterious |
| 84 | p.Asp84Asn | 15.81 | 3.98 | Deleterious | 15.81 | 3.98 | Deleterious |
| 84 | p.Asp84Lys | 22.66 | 4.50 | Deleterious | 22.66 | 4.50 | Deleterious |
| 84 | p.Asp84Thr | 16.29 | 4.03 | Deleterious | 16.29 | 4.03 | Deleterious |
| 84 | p.Asp84Arg | 22.75 | 4.51 | Deleterious | 22.75 | 4.51 | Deleterious |
| 84 | p.Asp84Ser | 6.14 | 2.62 | Deleterious | 6.14 | 2.62 | Deleterious |
| 84 | p.Asp84Ile | 23.12 | 4.53 | Deleterious | 23.12 | 4.53 | Deleterious |
| 84 | p.Asp84Met | 19.75 | 4.30 | Deleterious | 19.75 | 4.30 | Deleterious |
| 84 | p.Asp84His | 9.00 | 3.17 | Deleterious | 9.00 | 3.17 | Deleterious |
| 84 | p.Asp84Gln | 11.40 | 3.51 | Deleterious | 11.40 | 3.51 | Deleterious |
| 84 | p.Asp84Pro | 22.54 | 4.49 | Deleterious | 22.54 | 4.49 | Deleterious |
| 84 | p.Asp84Leu | 23.46 | 4.55 | Deleterious | 23.46 | 4.55 | Deleterious |
| 84 | p.Asp84Asp | 1.00 | 0.00 | Neutral | 1.00 | 0.00 | Neutral |
| 84 | p.Asp84Glu | 1.50 | 0.59 | Indeterminate | 1.50 | 0.59 | Indeterminate |
| 84 | p.Asp84Ala | 17.18 | 4.10 | Deleterious | 17.18 | 4.10 | Deleterious |
| 84 | p.Asp84Gly | 19.24 | 4.27 | Deleterious | 19.24 | 4.27 | Deleterious |
| 84 | p.Asp84Val | 20.90 | 4.39 | Deleterious | 20.90 | 4.39 | Deleterious |
| 84 | p.Asp84Tyr | 19.97 | 4.32 | Deleterious | 19.97 | 4.32 | Deleterious |
| 84 | p.Asp84Cys | 5.90 | 2.56 | Deleterious | 5.90 | 2.56 | Deleterious |
| 84 | p.Asp84Trp | 24.74 | 4.63 | Deleterious | 24.74 | 4.63 | Deleterious |
| 84 | p.Asp84Phe | 21.67 | 4.44 | Deleterious | 21.67 | 4.44 | Deleterious |
| 85 | p.Ala85Asn | 1.29 | 0.37 | Indeterminate | 1.29 | 0.37 | Indeterminate |
| 85 | p.Ala85Lys | 2.25 | 1.17 | Deleterious | 2.25 | 1.17 | Deleterious |
| 85 | p.Ala85Thr | 1.44 | 0.53 | Indeterminate | 1.44 | 0.53 | Indeterminate |
| 85 | p.Ala85Arg | 2.24 | 1.16 | Deleterious | 2.24 | 1.16 | Deleterious |
| 85 | p.Ala85Ser | 1.08 | 0.11 | Neutral | 1.08 | 0.11 | Neutral |
| 85 | p.Ala85Ile | 1.56 | 0.64 | Indeterminate | 1.56 | 0.64 | Indeterminate |
| 85 | p.Ala85Met | 0.77 | -0.37 | Neutral | 0.77 | -0.37 | Neutral |
| 85 | p.Ala85His | 1.66 | 0.73 | Indeterminate | 1.66 | 0.73 | Indeterminate |
| 85 | p.Ala85Gln | 1.50 | 0.59 | Indeterminate | 1.50 | 0.59 | Indeterminate |
| 85 | p.Ala85Pro | 1.51 | 0.60 | Indeterminate | 1.51 | 0.60 | Indeterminate |
| 85 | p.Ala85Leu | 1.03 | 0.04 | Neutral | 1.03 | 0.04 | Neutral |
| 85 | p.Ala85Asp | 2.03 | 1.02 | Indeterminate | 2.03 | 1.02 | Indeterminate |
| 85 | p.Ala85Glu | 1.96 | 0.97 | Indeterminate | 1.96 | 0.97 | Indeterminate |
| 85 | p.Ala85Ala | 1.00 | 0.00 | Neutral | 1.00 | 0.00 | Neutral |
| 85 | p.Ala85Gly | 0.86 | -0.22 | Neutral | 0.86 | -0.22 | Neutral |
| 85 | p.Ala85Val | 1.33 | 0.41 | Indeterminate | 1.33 | 0.41 | Indeterminate |
| 85 | p.Ala85Tyr | 1.82 | 0.87 | Indeterminate | 1.82 | 0.87 | Indeterminate |
| 85 | p.Ala85Cys | 0.91 | -0.14 | Neutral | 0.91 | -0.14 | Neutral |
| 85 | p.Ala85Trp | 2.44 | 1.29 | Deleterious | 2.44 | 1.29 | Deleterious |
| 85 | p.Ala85Phe | 1.95 | 0.96 | Indeterminate | 1.95 | 0.96 | Indeterminate |
| 86 | p.Ala86Asn | 8.60 | 3.10 | Deleterious | 8.60 | 3.10 | Deleterious |
| 86 | p.Ala86Lys | 10.11 | 3.34 | Deleterious | 10.11 | 3.34 | Deleterious |
| 86 | p.Ala86Thr | 0.79 | -0.34 | Neutral | 0.79 | -0.34 | Neutral |
| 86 | p.Ala86Arg | 9.67 | 3.27 | Deleterious | 9.67 | 3.27 | Deleterious |
| 86 | p.Ala86Ser | 1.40 | 0.49 | Indeterminate | 1.40 | 0.49 | Indeterminate |
| 86 | p.Ala86Ile | 1.86 | 0.90 | Indeterminate | 1.86 | 0.90 | Indeterminate |
| 86 | p.Ala86Met | 7.54 | 2.92 | Deleterious | 7.54 | 2.92 | Deleterious |
| 86 | p.Ala86His | 8.51 | 3.09 | Deleterious | 8.51 | 3.09 | Deleterious |
| 86 | p.Ala86Gln | 9.27 | 3.21 | Deleterious | 9.27 | 3.21 | Deleterious |
| 86 | p.Ala86Pro | 8.62 | 3.11 | Deleterious | 8.62 | 3.11 | Deleterious |
| 86 | p.Ala86Leu | 7.57 | 2.92 | Deleterious | 7.57 | 2.92 | Deleterious |
| 86 | p.Ala86Asp | 8.99 | 3.17 | Deleterious | 8.99 | 3.17 | Deleterious |

| | | | | | | | |
|---|---|---|---|---|---|---|---|
| 86 | p.Ala86Glu | 9.11 | 3.19 | Deleterious | 9.11 | 3.19 | Deleterious |
| 86 | p.Ala86Ala | 1.00 | 0.00 | Neutral | 1.00 | 0.00 | Neutral |
| 86 | p.Ala86Gly | 0.86 | -0.21 | Neutral | 0.86 | -0.21 | Neutral |
| 86 | p.Ala86Val | 1.00 | -0.01 | Neutral | 1.00 | -0.01 | Neutral |
| 86 | p.Ala86Tyr | 9.41 | 3.23 | Deleterious | 9.41 | 3.23 | Deleterious |
| 86 | p.Ala86Cys | 0.66 | -0.59 | Neutral | 0.66 | -0.59 | Neutral |
| 86 | p.Ala86Trp | 8.43 | 3.08 | Deleterious | 8.43 | 3.08 | Deleterious |
| 86 | p.Ala86Phe | 8.66 | 3.11 | Deleterious | 8.66 | 3.11 | Deleterious |
| 87 | p.Arg87Asn | 1.10 | 0.13 | Neutral | 1.10 | 0.13 | Neutral |
| 87 | p.Arg87Lys | 0.39 | -1.37 | Neutral | 0.39 | -1.37 | Neutral |
| 87 | p.Arg87Thr | 1.06 | 0.08 | Neutral | 1.06 | 0.08 | Neutral |
| 87 | p.Arg87Arg | 1.00 | 0.00 | Neutral | 1.00 | 0.00 | Neutral |
| 87 | p.Arg87Ser | 2.13 | 1.09 | Deleterious | 2.13 | 1.09 | Deleterious |
| 87 | p.Arg87Ile | 1.13 | 0.17 | Neutral | 1.13 | 0.17 | Neutral |
| 87 | p.Arg87Met | 1.70 | 0.77 | Indeterminate | 1.70 | 0.77 | Indeterminate |
| 87 | p.Arg87His | 1.92 | 0.94 | Indeterminate | 1.92 | 0.94 | Indeterminate |
| 87 | p.Arg87Gln | 2.09 | 1.07 | Indeterminate | 2.09 | 1.07 | Indeterminate |
| 87 | p.Arg87Pro | 34.88 | 5.12 | Deleterious | 34.88 | 5.12 | Deleterious |
| 87 | p.Arg87Leu | 1.32 | 0.41 | Indeterminate | 1.32 | 0.41 | Indeterminate |
| 87 | p.Arg87Asp | 6.02 | 2.59 | Deleterious | 6.02 | 2.59 | Deleterious |
| 87 | p.Arg87Glu | 3.27 | 1.71 | Deleterious | 3.27 | 1.71 | Deleterious |
| 87 | p.Arg87Ala | 2.60 | 1.38 | Deleterious | 2.60 | 1.38 | Deleterious |
| 87 | p.Arg87Gly | 0.81 | -0.30 | Neutral | 0.81 | -0.30 | Neutral |
| 87 | p.Arg87Val | 1.15 | 0.20 | Neutral | 1.15 | 0.20 | Neutral |
| 87 | p.Arg87Tyr | 1.81 | 0.86 | Indeterminate | 1.81 | 0.86 | Indeterminate |
| 87 | p.Arg87Cys | 0.74 | -0.43 | Neutral | 0.74 | -0.43 | Neutral |
| 87 | p.Arg87Trp | 20.35 | 4.35 | Deleterious | 20.35 | 4.35 | Deleterious |
| 87 | p.Arg87Phe | 1.26 | 0.34 | Indeterminate | 1.26 | 0.34 | Indeterminate |
| 88 | p.Glu88Asn | 1.30 | 0.38 | Indeterminate | 1.30 | 0.38 | Indeterminate |
| 88 | p.Glu88Lys | 1.95 | 0.96 | Indeterminate | 1.95 | 0.96 | Indeterminate |
| 88 | p.Glu88Thr | 1.29 | 0.37 | Indeterminate | 1.29 | 0.37 | Indeterminate |
| 88 | p.Glu88Arg | 1.89 | 0.92 | Indeterminate | 1.89 | 0.92 | Indeterminate |
| 88 | p.Glu88Ser | 1.21 | 0.28 | Indeterminate | 1.21 | 0.28 | Indeterminate |
| 88 | p.Glu88Ile | 1.18 | 0.23 | Neutral | 1.18 | 0.23 | Neutral |
| 88 | p.Glu88Met | 1.17 | 0.23 | Neutral | 1.17 | 0.23 | Neutral |
| 88 | p.Glu88His | 1.40 | 0.49 | Indeterminate | 1.40 | 0.49 | Indeterminate |
| 88 | p.Glu88Gln | 1.27 | 0.35 | Indeterminate | 1.27 | 0.35 | Indeterminate |
| 88 | p.Glu88Pro | 3.13 | 1.65 | Deleterious | 3.13 | 1.65 | Deleterious |
| 88 | p.Glu88Leu | 1.37 | 0.45 | Indeterminate | 1.37 | 0.45 | Indeterminate |
| 88 | p.Glu88Asp | 1.36 | 0.44 | Indeterminate | 1.36 | 0.44 | Indeterminate |
| 88 | p.Glu88Glu | 1.00 | 0.00 | Neutral | 1.00 | 0.00 | Neutral |
| 88 | p.Glu88Ala | 1.08 | 0.10 | Neutral | 1.08 | 0.10 | Neutral |
| 88 | p.Glu88Gly | 1.20 | 0.26 | Indeterminate | 1.20 | 0.26 | Indeterminate |
| 88 | p.Glu88Val | 1.31 | 0.39 | Indeterminate | 1.31 | 0.39 | Indeterminate |
| 88 | p.Glu88Tyr | 1.34 | 0.42 | Indeterminate | 1.34 | 0.42 | Indeterminate |
| 88 | p.Glu88Cys | 1.15 | 0.21 | Neutral | 1.15 | 0.21 | Neutral |
| 88 | p.Glu88Trp | 1.20 | 0.26 | Indeterminate | 1.20 | 0.26 | Indeterminate |
| 88 | p.Glu88Phe | 1.41 | 0.49 | Indeterminate | 1.41 | 0.49 | Indeterminate |
| 89 | p.Gly89Asn | 1.78 | 0.83 | Indeterminate | 1.78 | 0.83 | Indeterminate |
| 89 | p.Gly89Lys | 7.99 | 3.00 | Deleterious | 7.99 | 3.00 | Deleterious |
| 89 | p.Gly89Thr | 13.14 | 3.72 | Deleterious | 13.14 | 3.72 | Deleterious |
| 89 | p.Gly89Arg | 8.98 | 3.17 | Deleterious | 8.98 | 3.17 | Deleterious |
| 89 | p.Gly89Ser | 4.06 | 2.02 | Deleterious | 4.06 | 2.02 | Deleterious |
| 89 | p.Gly89Ile | 16.30 | 4.03 | Deleterious | 16.30 | 4.03 | Deleterious |
| 89 | p.Gly89Met | 11.69 | 3.55 | Deleterious | 11.69 | 3.55 | Deleterious |
| 89 | p.Gly89His | 9.73 | 3.28 | Deleterious | 9.73 | 3.28 | Deleterious |
| 89 | p.Gly89Gln | 8.25 | 3.04 | Deleterious | 8.25 | 3.04 | Deleterious |
| 89 | p.Gly89Pro | 11.31 | 3.50 | Deleterious | 11.31 | 3.50 | Deleterious |
| 89 | p.Gly89Leu | 13.57 | 3.76 | Deleterious | 13.57 | 3.76 | Deleterious |
| 89 | p.Gly89Asp | 3.52 | 1.82 | Deleterious | 3.52 | 1.82 | Deleterious |
| 89 | p.Gly89Glu | 10.29 | 3.36 | Deleterious | 10.29 | 3.36 | Deleterious |
| 89 | p.Gly89Ala | 2.07 | 1.05 | Indeterminate | 2.07 | 1.05 | Indeterminate |
| 89 | p.Gly89Gly | 1.00 | 0.00 | Neutral | 1.00 | 0.00 | Neutral |
| 89 | p.Gly89Val | 14.96 | 3.90 | Deleterious | 14.96 | 3.90 | Deleterious |
| 89 | p.Gly89Tyr | 14.02 | 3.81 | Deleterious | 14.02 | 3.81 | Deleterious |
| 89 | p.Gly89Cys | 4.06 | 2.02 | Deleterious | 4.06 | 2.02 | Deleterious |
| 89 | p.Gly89Trp | 14.23 | 3.83 | Deleterious | 14.23 | 3.83 | Deleterious |
| 89 | p.Gly89Phe | 13.71 | 3.78 | Deleterious | 13.71 | 3.78 | Deleterious |
| 90 | p.Phe90Asn | 0.89 | -0.17 | Neutral | 0.89 | -0.17 | Neutral |
| 90 | p.Phe90Lys | 1.21 | 0.28 | Indeterminate | 1.21 | 0.28 | Indeterminate |
| 90 | p.Phe90Thr | 1.15 | 0.21 | Neutral | 1.15 | 0.21 | Neutral |
| 90 | p.Phe90Arg | 0.96 | -0.06 | Neutral | 0.96 | -0.06 | Neutral |
| 90 | p.Phe90Ser | 0.94 | -0.08 | Neutral | 0.94 | -0.08 | Neutral |
| 90 | p.Phe90Ile | 0.87 | -0.20 | Neutral | 0.87 | -0.20 | Neutral |
| 90 | p.Phe90Met | 0.75 | -0.42 | Neutral | 0.75 | -0.42 | Neutral |
| 90 | p.Phe90His | 0.78 | -0.36 | Neutral | 0.78 | -0.36 | Neutral |
| 90 | p.Phe90Gln | 0.76 | -0.40 | Neutral | 0.76 | -0.40 | Neutral |
| 90 | p.Phe90Pro | 1.62 | 0.70 | Indeterminate | 1.62 | 0.70 | Indeterminate |
| 90 | p.Phe90Leu | 0.78 | -0.36 | Neutral | 0.78 | -0.36 | Neutral |
| 90 | p.Phe90Asp | 1.73 | 0.79 | Indeterminate | 1.73 | 0.79 | Indeterminate |
| 90 | p.Phe90Glu | 1.15 | 0.21 | Neutral | 1.15 | 0.21 | Neutral |
| 90 | p.Phe90Ala | 0.84 | -0.24 | Neutral | 0.84 | -0.24 | Neutral |
| 90 | p.Phe90Gly | 1.01 | 0.02 | Neutral | 1.01 | 0.02 | Neutral |
| 90 | p.Phe90Val | 1.11 | 0.15 | Neutral | 1.11 | 0.15 | Neutral |
| 90 | p.Phe90Tyr | 0.80 | -0.32 | Neutral | 0.80 | -0.32 | Neutral |
| 90 | p.Phe90Cys | 0.99 | -0.01 | Neutral | 0.99 | -0.01 | Neutral |
| 90 | p.Phe90Trp | 0.93 | -0.10 | Neutral | 0.93 | -0.10 | Neutral |
| 90 | p.Phe90Phe | 1.00 | 0.00 | Neutral | 1.00 | 0.00 | Neutral |
| 91 | p.Leu91Asn | 0.70 | -0.51 | Neutral | 0.70 | -0.51 | Neutral |
| 91 | p.Leu91Lys | 1.43 | 0.52 | Indeterminate | 1.43 | 0.52 | Indeterminate |
| 91 | p.Leu91Thr | 1.46 | 0.54 | Indeterminate | 1.46 | 0.54 | Indeterminate |
| 91 | p.Leu91Arg | 1.16 | 0.21 | Neutral | 1.16 | 0.21 | Neutral |
| 91 | p.Leu91Ser | 1.94 | 0.96 | Indeterminate | 1.94 | 0.96 | Indeterminate |
| 91 | p.Leu91Ile | 1.70 | 0.76 | Indeterminate | 1.70 | 0.76 | Indeterminate |
| 91 | p.Leu91Met | 2.60 | 1.38 | Deleterious | 2.60 | 1.38 | Deleterious |
| 91 | p.Leu91His | 1.68 | 0.75 | Indeterminate | 1.68 | 0.75 | Indeterminate |
| 91 | p.Leu91Gln | 0.79 | -0.33 | Neutral | 0.79 | -0.33 | Neutral |
| 91 | p.Leu91Pro | 1.53 | 0.61 | Indeterminate | 1.53 | 0.61 | Indeterminate |
| 91 | p.Leu91Leu | 1.00 | 0.00 | Neutral | 1.00 | 0.00 | Neutral |
| 91 | p.Leu91Asp | 1.31 | 0.39 | Indeterminate | 1.31 | 0.39 | Indeterminate |
| 91 | p.Leu91Glu | 1.34 | 0.43 | Indeterminate | 1.34 | 0.43 | Indeterminate |

| | | | | | | | | | | |
|---|---|---|---|---|---|---|---|---|---|---|
| 91 | p.Leu91Ala | 1.03 | 0.04 | Neutral | | | | 1.03 | 0.04 | Neutral |
| 91 | p.Leu91Gly | 1.39 | 0.47 | Indeterminate | | | | 1.39 | 0.47 | Indeterminate |
| 91 | p.Leu91Val | 1.29 | 0.37 | Indeterminate | | | | 1.29 | 0.37 | Indeterminate |
| 91 | p.Leu91Tyr | 3.09 | 1.63 | Deleterious | | | | 3.09 | 1.63 | Deleterious |
| 91 | p.Leu91Cys | 0.96 | -0.05 | Neutral | | | | 0.96 | -0.05 | Neutral |
| 91 | p.Leu91Trp | 1.59 | 0.67 | Indeterminate | | | | 1.59 | 0.67 | Indeterminate |
| 91 | p.Leu91Phe | 2.11 | 1.08 | Indeterminate | | | | 2.11 | 1.08 | Indeterminate |
| 92 | p.Asp92Asn | 1.06 | 0.08 | Neutral | | | | 1.06 | 0.08 | Neutral |
| 92 | p.Asp92Lys | 1.23 | 0.30 | Indeterminate | | | | 1.23 | 0.30 | Indeterminate |
| 92 | p.Asp92Thr | 0.90 | -0.16 | Neutral | | | | 0.90 | -0.16 | Neutral |
| 92 | p.Asp92Arg | 1.02 | 0.03 | Neutral | | | | 1.02 | 0.03 | Neutral |
| 92 | p.Asp92Ser | 0.97 | -0.05 | Neutral | | | | 0.97 | -0.05 | Neutral |
| 92 | p.Asp92Ile | 1.19 | 0.25 | Indeterminate | | | | 1.19 | 0.25 | Indeterminate |
| 92 | p.Asp92Met | 0.98 | -0.03 | Neutral | | | | 0.98 | -0.03 | Neutral |
| 92 | p.Asp92His | 1.09 | 0.13 | Neutral | | | | 1.09 | 0.13 | Neutral |
| 92 | p.Asp92Gln | 1.04 | 0.06 | Neutral | | | | 1.04 | 0.06 | Neutral |
| 92 | p.Asp92Pro | 1.11 | 0.15 | Neutral | | | | 1.11 | 0.15 | Neutral |
| 92 | p.Asp92Leu | 1.02 | 0.02 | Neutral | | | | 1.02 | 0.02 | Neutral |
| 92 | p.Asp92Asp | 1.00 | 0.00 | Neutral | | | | 1.00 | 0.00 | Neutral |
| 92 | p.Asp92Glu | 1.10 | 0.14 | Neutral | | | | 1.10 | 0.14 | Neutral |
| 92 | p.Asp92Ala | 1.12 | 0.17 | Neutral | | | | 1.12 | 0.17 | Neutral |
| 92 | p.Asp92Gly | 1.17 | 0.22 | Neutral | | | | 1.17 | 0.22 | Neutral |
| 92 | p.Asp92Val | 1.03 | 0.04 | Neutral | | | | 1.03 | 0.04 | Neutral |
| 92 | p.Asp92Tyr | 1.04 | 0.05 | Neutral | | | | 1.04 | 0.05 | Neutral |
| 92 | p.Asp92Cys | 0.98 | -0.03 | Neutral | | | | 0.98 | -0.03 | Neutral |
| 92 | p.Asp92Trp | 1.05 | 0.08 | Neutral | | | | 1.05 | 0.08 | Neutral |
| 92 | p.Asp92Phe | 1.21 | 0.28 | Indeterminate | | | | 1.21 | 0.28 | Indeterminate |
| 93 | p.Thr93Asn | 1.83 | 0.87 | Indeterminate | | | | 1.83 | 0.87 | Indeterminate |
| 93 | p.Thr93Lys | 3.80 | 1.93 | Deleterious | | | | 3.80 | 1.93 | Deleterious |
| 93 | p.Thr93Thr | 1.00 | 0.00 | Neutral | | | | 1.00 | 0.00 | Neutral |
| 93 | p.Thr93Arg | 2.46 | 1.30 | Deleterious | | | | 2.46 | 1.30 | Deleterious |
| 93 | p.Thr93Ser | 1.46 | 0.55 | Indeterminate | | | | 1.46 | 0.55 | Indeterminate |
| 93 | p.Thr93Ile | 0.98 | -0.03 | Neutral | | | | 0.98 | -0.03 | Neutral |
| 93 | p.Thr93Met | 1.77 | 0.82 | Indeterminate | | | | 1.77 | 0.82 | Indeterminate |
| 93 | p.Thr93His | 2.45 | 1.29 | Deleterious | | | | 2.45 | 1.29 | Deleterious |
| 93 | p.Thr93Gln | 2.20 | 1.14 | Deleterious | | | | 2.20 | 1.14 | Deleterious |
| 93 | p.Thr93Pro | 6.26 | 2.65 | Deleterious | | | | 6.26 | 2.65 | Deleterious |
| 93 | p.Thr93Leu | 1.62 | 0.70 | Indeterminate | | | | 1.62 | 0.70 | Indeterminate |
| 93 | p.Thr93Asp | 2.94 | 1.56 | Deleterious | | | | 2.94 | 1.56 | Deleterious |
| 93 | p.Thr93Glu | 2.43 | 1.28 | Deleterious | | | | 2.43 | 1.28 | Deleterious |
| 93 | p.Thr93Ala | 2.32 | 1.21 | Deleterious | | | | 2.32 | 1.21 | Deleterious |
| 93 | p.Thr93Gly | 2.45 | 1.29 | Deleterious | | | | 2.45 | 1.29 | Deleterious |
| 93 | p.Thr93Val | 1.45 | 0.53 | Indeterminate | | | | 1.45 | 0.53 | Indeterminate |
| 93 | p.Thr93Tyr | 2.87 | 1.52 | Deleterious | | | | 2.87 | 1.52 | Deleterious |
| 93 | p.Thr93Cys | 1.63 | 0.70 | Indeterminate | | | | 1.63 | 0.70 | Indeterminate |
| 93 | p.Thr93Trp | 2.31 | 1.21 | Deleterious | | | | 2.31 | 1.21 | Deleterious |
| 93 | p.Thr93Phe | 1.98 | 0.98 | Indeterminate | | | | 1.98 | 0.98 | Indeterminate |
| 94 | p.Leu94Asn | 2.32 | 1.21 | Deleterious | | | | 2.32 | 1.21 | Deleterious |
| 94 | p.Leu94Lys | 6.25 | 2.64 | Deleterious | | | | 6.25 | 2.64 | Deleterious |
| 94 | p.Leu94Thr | 2.48 | 1.31 | Deleterious | | | | 2.48 | 1.31 | Deleterious |
| 94 | p.Leu94Arg | 8.16 | 3.03 | Deleterious | | | | 8.16 | 3.03 | Deleterious |
| 94 | p.Leu94Ser | 0.96 | -0.06 | Neutral | | | | 0.96 | -0.06 | Neutral |
| 94 | p.Leu94Ile | 0.69 | -0.54 | Neutral | | | | 0.69 | -0.54 | Neutral |
| 94 | p.Leu94Met | 0.42 | -1.25 | Neutral | | | | 0.42 | -1.25 | Neutral |
| 94 | p.Leu94His | 8.80 | 3.14 | Deleterious | | | | 8.80 | 3.14 | Deleterious |
| 94 | p.Leu94Gln | 4.83 | 2.27 | Deleterious | | | | 4.83 | 2.27 | Deleterious |
| 94 | p.Leu94Pro | 5.87 | 2.55 | Deleterious | | | | 5.87 | 2.55 | Deleterious |
| 94 | p.Leu94Leu | 1.00 | 0.00 | Neutral | | | | 1.00 | 0.00 | Neutral |
| 94 | p.Leu94Asp | 7.73 | 2.95 | Deleterious | | | | 7.73 | 2.95 | Deleterious |
| 94 | p.Leu94Glu | 4.47 | 2.16 | Deleterious | | | | 4.47 | 2.16 | Deleterious |
| 94 | p.Leu94Ala | 2.30 | 1.20 | Deleterious | | | | 2.30 | 1.20 | Deleterious |
| 94 | p.Leu94Gly | 4.46 | 2.16 | Deleterious | | | | 4.46 | 2.16 | Deleterious |
| 94 | p.Leu94Val | 1.22 | 0.28 | Indeterminate | | | | 1.22 | 0.28 | Indeterminate |
| 94 | p.Leu94Tyr | 9.51 | 3.25 | Deleterious | | | | 9.51 | 3.25 | Deleterious |
| 94 | p.Leu94Cys | 1.83 | 0.87 | Indeterminate | | | | 1.83 | 0.87 | Indeterminate |
| 94 | p.Leu94Trp | 7.26 | 2.86 | Deleterious | | | | 7.26 | 2.86 | Deleterious |
| 94 | p.Leu94Phe | 1.87 | 0.91 | Indeterminate | | | | 1.87 | 0.91 | Indeterminate |
| 95 | p.Val95Asn | 1.08 | 0.11 | Neutral | | | | 1.08 | 0.11 | Neutral |
| 95 | p.Val95Lys | 0.93 | -0.10 | Neutral | | | | 0.93 | -0.10 | Neutral |
| 95 | p.Val95Thr | 0.99 | -0.01 | Neutral | | | | 0.99 | -0.01 | Neutral |
| 95 | p.Val95Arg | 1.09 | 0.13 | Neutral | | | | 1.09 | 0.13 | Neutral |
| 95 | p.Val95Ser | 1.08 | 0.11 | Neutral | | | | 1.08 | 0.11 | Neutral |
| 95 | p.Val95Ile | 0.99 | -0.02 | Neutral | | | | 0.99 | -0.02 | Neutral |
| 95 | p.Val95Met | 1.00 | 0.00 | Neutral | | | | 1.00 | 0.00 | Neutral |
| 95 | p.Val95His | 0.78 | -0.36 | Neutral | | | | 0.78 | -0.36 | Neutral |
| 95 | p.Val95Gln | 0.98 | -0.03 | Neutral | | | | 0.98 | -0.03 | Neutral |
| 95 | p.Val95Pro | 4.78 | 2.26 | Deleterious | | | | 4.78 | 2.26 | Deleterious |
| 95 | p.Val95Leu | 0.82 | -0.28 | Neutral | | | | 0.82 | -0.28 | Neutral |
| 95 | p.Val95Asp | 0.96 | -0.06 | Neutral | | | | 0.96 | -0.06 | Neutral |
| 95 | p.Val95Glu | 1.05 | 0.08 | Neutral | | | | 1.05 | 0.08 | Neutral |
| 95 | p.Val95Ala | 0.74 | -0.44 | Neutral | | | | 0.74 | -0.44 | Neutral |
| 95 | p.Val95Gly | 1.03 | 0.04 | Neutral | | | | 1.03 | 0.04 | Neutral |
| 95 | p.Val95Val | 1.00 | 0.00 | Neutral | | | | 1.00 | 0.00 | Neutral |
| 95 | p.Val95Tyr | 1.02 | 0.02 | Neutral | | | | 1.02 | 0.02 | Neutral |
| 95 | p.Val95Cys | 0.82 | -0.28 | Neutral | | | | 0.82 | -0.28 | Neutral |
| 95 | p.Val95Trp | 1.21 | 0.27 | Indeterminate | | | | 1.21 | 0.27 | Indeterminate |
| 95 | p.Val95Phe | 1.11 | 0.15 | Neutral | | | | 1.11 | 0.15 | Neutral |
| 96 | p.Val96Asn | 0.75 | -0.41 | Neutral | 1.22 | 0.29 | Indeterminate | 0.99 | -0.02 | Neutral |
| 96 | p.Val96Lys | 0.89 | -0.17 | Neutral | 0.94 | -0.09 | Neutral | 0.92 | -0.13 | Neutral |
| 96 | p.Val96Thr | 0.69 | -0.53 | Neutral | 0.92 | -0.11 | Neutral | 0.81 | -0.31 | Neutral |
| 96 | p.Val96Arg | 0.85 | -0.24 | Neutral | 1.02 | 0.02 | Neutral | 0.93 | -0.10 | Neutral |
| 96 | p.Val96Ser | 0.75 | -0.41 | Neutral | 0.78 | -0.35 | Neutral | 0.77 | -0.38 | Neutral |
| 96 | p.Val96Ile | 0.84 | -0.25 | Neutral | 1.01 | 0.02 | Neutral | 0.93 | -0.11 | Neutral |
| 96 | p.Val96Met | 0.70 | -0.52 | Neutral | 0.82 | -0.29 | Neutral | 0.76 | -0.40 | Neutral |
| 96 | p.Val96His | 0.93 | -0.10 | Neutral | 0.92 | -0.13 | Neutral | 0.93 | -0.11 | Neutral |
| 96 | p.Val96Gln | 0.86 | -0.23 | Neutral | 1.10 | 0.14 | Indeterminate | 0.98 | -0.03 | Neutral |
| 96 | p.Val96Pro | 0.93 | -0.10 | Neutral | 0.87 | -0.20 | Neutral | 0.90 | -0.15 | Neutral |
| 96 | p.Val96Leu | 0.87 | -0.20 | Neutral | 1.21 | 0.28 | Indeterminate | 1.04 | 0.06 | Neutral |
| 96 | p.Val96Asp | 0.82 | -0.29 | Neutral | 1.26 | 0.33 | Indeterminate | 1.04 | 0.05 | Neutral |
| 96 | p.Val96Glu | 0.65 | -0.63 | Neutral | 1.05 | 0.08 | Neutral | 0.85 | -0.23 | Neutral |
| 96 | p.Val96Ala | 1.10 | 0.14 | Neutral | 1.53 | 0.62 | Indeterminate | 1.32 | 0.40 | Indeterminate |

| | | | | | | | | | | |
|---|---|---|---|---|---|---|---|---|---|---|
| 96 | p.Val96Gly | 1.13 | 0.18 | Neutral | 0.86 | -0.22 | Neutral | 0.99 | -0.01 | Neutral |
| 96 | p.Val96Val | 1.00 | 0.00 | Neutral | 1.00 | 0.00 | Neutral | 1.00 | 0.00 | Neutral |
| 96 | p.Val96Tyr | 0.82 | -0.28 | Neutral | 1.08 | 0.11 | Neutral | 0.95 | -0.07 | Neutral |
| 96 | p.Val96Cys | 0.82 | -0.28 | Neutral | 1.37 | 0.45 | Indeterminate | 1.10 | 0.13 | Neutral |
| 96 | p.Val96Trp | 0.70 | -0.51 | Neutral | 1.46 | 0.54 | Indeterminate | 1.08 | 0.11 | Neutral |
| 96 | p.Val96Phe | 0.72 | -0.48 | Neutral | 1.08 | 0.11 | Neutral | 0.90 | -0.15 | Neutral |
| 97 | p.Leu97Asn | 7.43 | 2.89 | Deleterious | | | | 7.43 | 2.89 | Deleterious |
| 97 | p.Leu97Lys | 8.27 | 3.05 | Deleterious | | | | 8.27 | 3.05 | Deleterious |
| 97 | p.Leu97Thr | 5.89 | 2.56 | Deleterious | | | | 5.89 | 2.56 | Deleterious |
| 97 | p.Leu97Arg | 8.53 | 3.09 | Deleterious | | | | 8.53 | 3.09 | Deleterious |
| 97 | p.Leu97Ser | 7.64 | 2.93 | Deleterious | | | | 7.64 | 2.93 | Deleterious |
| 97 | p.Leu97Ile | 0.85 | -0.24 | Neutral | | | | 0.85 | -0.24 | Neutral |
| 97 | p.Leu97Met | 0.69 | -0.53 | Neutral | | | | 0.69 | -0.53 | Neutral |
| 97 | p.Leu97His | 6.20 | 2.63 | Deleterious | | | | 6.20 | 2.63 | Deleterious |
| 97 | p.Leu97Gln | 4.19 | 2.07 | Deleterious | | | | 4.19 | 2.07 | Deleterious |
| 97 | p.Leu97Pro | 8.72 | 3.12 | Deleterious | | | | 8.72 | 3.12 | Deleterious |
| 97 | p.Leu97Leu | 1.00 | 0.00 | Neutral | | | | 1.00 | 0.00 | Neutral |
| 97 | p.Leu97Asp | 8.74 | 3.13 | Deleterious | | | | 8.74 | 3.13 | Deleterious |
| 97 | p.Leu97Glu | 6.81 | 2.77 | Deleterious | | | | 6.81 | 2.77 | Deleterious |
| 97 | p.Leu97Ala | 4.79 | 2.26 | Deleterious | | | | 4.79 | 2.26 | Deleterious |
| 97 | p.Leu97Gly | 8.76 | 3.13 | Deleterious | | | | 8.76 | 3.13 | Deleterious |
| 97 | p.Leu97Val | 1.68 | 0.75 | Indeterminate | | | | 1.68 | 0.75 | Indeterminate |
| 97 | p.Leu97Tyr | 1.01 | 0.01 | Neutral | | | | 1.01 | 0.01 | Neutral |
| 97 | p.Leu97Cys | 1.03 | 0.04 | Neutral | | | | 1.03 | 0.04 | Neutral |
| 97 | p.Leu97Trp | 6.72 | 2.75 | Deleterious | | | | 6.72 | 2.75 | Deleterious |
| 97 | p.Leu97Phe | 0.92 | -0.12 | Neutral | | | | 0.92 | -0.12 | Neutral |
| 98 | p.His98Asn | 1.02 | 0.02 | Neutral | | | | 1.02 | 0.02 | Neutral |
| 98 | p.His98Lys | 0.92 | -0.12 | Neutral | | | | 0.92 | -0.12 | Neutral |
| 98 | p.His98Thr | 1.07 | 0.10 | Neutral | | | | 1.07 | 0.10 | Neutral |
| 98 | p.His98Arg | 1.00 | 0.00 | Neutral | | | | 1.00 | 0.00 | Neutral |
| 98 | p.His98Ser | 1.29 | 0.37 | Indeterminate | | | | 1.29 | 0.37 | Indeterminate |
| 98 | p.His98Ile | 0.98 | -0.03 | Neutral | | | | 0.98 | -0.03 | Neutral |
| 98 | p.His98Met | 0.97 | -0.04 | Neutral | | | | 0.97 | -0.04 | Neutral |
| 98 | p.His98His | 1.00 | 0.00 | Neutral | | | | 1.00 | 0.00 | Neutral |
| 98 | p.His98Gln | 1.19 | 0.25 | Indeterminate | | | | 1.19 | 0.25 | Indeterminate |
| 98 | p.His98Pro | 5.39 | 2.43 | Deleterious | | | | 5.39 | 2.43 | Deleterious |
| 98 | p.His98Leu | 1.15 | 0.21 | Neutral | | | | 1.15 | 0.21 | Neutral |
| 98 | p.His98Asp | 1.26 | 0.34 | Indeterminate | | | | 1.26 | 0.34 | Indeterminate |
| 98 | p.His98Glu | 1.04 | 0.06 | Neutral | | | | 1.04 | 0.06 | Neutral |
| 98 | p.His98Ala | 0.93 | -0.10 | Neutral | | | | 0.93 | -0.10 | Neutral |
| 98 | p.His98Gly | 1.16 | 0.21 | Neutral | | | | 1.16 | 0.21 | Neutral |
| 98 | p.His98Val | 1.07 | 0.10 | Neutral | | | | 1.07 | 0.10 | Neutral |
| 98 | p.His98Tyr | 1.07 | 0.10 | Neutral | | | | 1.07 | 0.10 | Neutral |
| 98 | p.His98Cys | 1.09 | 0.12 | Neutral | | | | 1.09 | 0.12 | Neutral |
| 98 | p.His98Trp | 0.94 | -0.10 | Neutral | | | | 0.94 | -0.10 | Neutral |
| 98 | p.His98Phe | 0.94 | -0.08 | Neutral | | | | 0.94 | -0.08 | Neutral |
| 99 | p.Arg99Asn | 1.39 | 0.47 | Indeterminate | | | | 1.39 | 0.47 | Indeterminate |
| 99 | p.Arg99Lys | 1.20 | 0.26 | Indeterminate | | | | 1.20 | 0.26 | Indeterminate |
| 99 | p.Arg99Thr | 1.46 | 0.55 | Indeterminate | | | | 1.46 | 0.55 | Indeterminate |
| 99 | p.Arg99Arg | 1.00 | 0.00 | Neutral | | | | 1.00 | 0.00 | Neutral |
| 99 | p.Arg99Ser | 1.46 | 0.54 | Indeterminate | | | | 1.46 | 0.54 | Indeterminate |
| 99 | p.Arg99Ile | 1.26 | 0.33 | Indeterminate | | | | 1.26 | 0.33 | Indeterminate |
| 99 | p.Arg99Met | 1.60 | 0.68 | Indeterminate | | | | 1.60 | 0.68 | Indeterminate |
| 99 | p.Arg99His | 0.94 | -0.09 | Neutral | | | | 0.94 | -0.09 | Neutral |
| 99 | p.Arg99Gln | 0.92 | -0.11 | Neutral | | | | 0.92 | -0.11 | Neutral |
| 99 | p.Arg99Pro | 38.44 | 5.26 | Deleterious | | | | 38.44 | 5.26 | Deleterious |
| 99 | p.Arg99Leu | 1.72 | 0.78 | Indeterminate | | | | 1.72 | 0.78 | Indeterminate |
| 99 | p.Arg99Asp | 1.92 | 0.94 | Indeterminate | | | | 1.92 | 0.94 | Indeterminate |
| 99 | p.Arg99Glu | 1.40 | 0.48 | Indeterminate | | | | 1.40 | 0.48 | Indeterminate |
| 99 | p.Arg99Ala | 2.31 | 1.21 | Deleterious | | | | 2.31 | 1.21 | Deleterious |
| 99 | p.Arg99Gly | 1.02 | 0.03 | Neutral | | | | 1.02 | 0.03 | Neutral |
| 99 | p.Arg99Val | 2.27 | 1.18 | Deleterious | | | | 2.27 | 1.18 | Deleterious |
| 99 | p.Arg99Tyr | 1.06 | 0.09 | Neutral | | | | 1.06 | 0.09 | Neutral |
| 99 | p.Arg99Cys | 1.77 | 0.82 | Indeterminate | | | | 1.77 | 0.82 | Indeterminate |
| 99 | p.Arg99Trp | 1.83 | 0.87 | Indeterminate | | | | 1.83 | 0.87 | Indeterminate |
| 99 | p.Arg99Phe | 1.14 | 0.19 | Neutral | | | | 1.14 | 0.19 | Neutral |
| 100 | p.Ala100Asn | 1.10 | 0.13 | Neutral | | | | 1.10 | 0.13 | Neutral |
| 100 | p.Ala100Lys | 0.94 | -0.09 | Neutral | | | | 0.94 | -0.09 | Neutral |
| 100 | p.Ala100Thr | 1.34 | 0.42 | Indeterminate | | | | 1.34 | 0.42 | Indeterminate |
| 100 | p.Ala100Arg | 0.92 | -0.12 | Neutral | | | | 0.92 | -0.12 | Neutral |
| 100 | p.Ala100Ser | 0.74 | -0.44 | Neutral | | | | 0.74 | -0.44 | Neutral |
| 100 | p.Ala100Ile | 1.05 | 0.06 | Neutral | | | | 1.05 | 0.06 | Neutral |
| 100 | p.Ala100Met | 1.30 | 0.38 | Indeterminate | | | | 1.30 | 0.38 | Indeterminate |
| 100 | p.Ala100His | 0.79 | -0.35 | Neutral | | | | 0.79 | -0.35 | Neutral |
| 100 | p.Ala100Gln | 1.02 | 0.03 | Neutral | | | | 1.02 | 0.03 | Neutral |
| 100 | p.Ala100Pro | 13.53 | 3.76 | Deleterious | | | | 13.53 | 3.76 | Deleterious |
| 100 | p.Ala100Leu | 0.87 | -0.20 | Neutral | | | | 0.87 | -0.20 | Neutral |
| 100 | p.Ala100Asp | 0.73 | -0.45 | Neutral | | | | 0.73 | -0.45 | Neutral |
| 100 | p.Ala100Glu | 1.75 | 0.81 | Indeterminate | | | | 1.75 | 0.81 | Indeterminate |
| 100 | p.Ala100Ala | 1.00 | 0.00 | Neutral | | | | 1.00 | 0.00 | Neutral |
| 100 | p.Ala100Gly | 3.88 | 1.95 | Deleterious | | | | 3.88 | 1.95 | Deleterious |
| 100 | p.Ala100Val | 0.82 | -0.29 | Neutral | | | | 0.82 | -0.29 | Neutral |
| 100 | p.Ala100Tyr | 1.16 | 0.22 | Neutral | | | | 1.16 | 0.22 | Neutral |
| 100 | p.Ala100Cys | 0.64 | -0.64 | Neutral | | | | 0.64 | -0.64 | Neutral |
| 100 | p.Ala100Trp | 0.77 | -0.38 | Neutral | | | | 0.77 | -0.38 | Neutral |
| 100 | p.Ala100Phe | 0.74 | -0.44 | Neutral | | | | 0.74 | -0.44 | Neutral |
| 101 | p.Gly101Asn | 1.44 | 0.52 | Indeterminate | | | | 1.44 | 0.52 | Indeterminate |
| 101 | p.Gly101Lys | 1.85 | 0.89 | Indeterminate | | | | 1.85 | 0.89 | Indeterminate |
| 101 | p.Gly101Thr | 2.15 | 1.11 | Deleterious | | | | 2.15 | 1.11 | Deleterious |
| 101 | p.Gly101Arg | 1.68 | 0.75 | Indeterminate | | | | 1.68 | 0.75 | Indeterminate |
| 101 | p.Gly101Ser | 1.58 | 0.66 | Indeterminate | | | | 1.58 | 0.66 | Indeterminate |
| 101 | p.Gly101Ile | 9.59 | 3.26 | Deleterious | | | | 9.59 | 3.26 | Deleterious |
| 101 | p.Gly101Met | 2.07 | 1.05 | Indeterminate | | | | 2.07 | 1.05 | Indeterminate |
| 101 | p.Gly101His | 1.59 | 0.67 | Indeterminate | | | | 1.59 | 0.67 | Indeterminate |
| 101 | p.Gly101Gln | 1.58 | 0.66 | Indeterminate | | | | 1.58 | 0.66 | Indeterminate |
| 101 | p.Gly101Pro | 8.50 | 3.09 | Deleterious | | | | 8.50 | 3.09 | Deleterious |
| 101 | p.Gly101Leu | 3.51 | 1.81 | Deleterious | | | | 3.51 | 1.81 | Deleterious |
| 101 | p.Gly101Asp | 1.31 | 0.39 | Indeterminate | | | | 1.31 | 0.39 | Indeterminate |
| 101 | p.Gly101Glu | 1.85 | 0.89 | Indeterminate | | | | 1.85 | 0.89 | Indeterminate |
| 101 | p.Gly101Ala | 1.61 | 0.68 | Indeterminate | | | | 1.61 | 0.68 | Indeterminate |
| 101 | p.Gly101Gly | 1.00 | 0.00 | Neutral | | | | 1.00 | 0.00 | Neutral |

| | | | | | | | | | | |
|---|---|---|---|---|---|---|---|---|---|---|
| 101 | p.Gly101Val | 8.71 | 3.12 | Deleterious | | | | 8.71 | 3.12 | Deleterious |
| 101 | p.Gly101Tyr | 2.53 | 1.34 | Deleterious | | | | 2.53 | 1.34 | Deleterious |
| 101 | p.Gly101Cys | 1.65 | 0.72 | Indeterminate | | | | 1.65 | 0.72 | Indeterminate |
| 101 | p.Gly101Trp | 4.49 | 2.17 | Deleterious | | | | 4.49 | 2.17 | Deleterious |
| 101 | p.Gly101Phe | 2.19 | 1.13 | Deleterious | | | | 2.19 | 1.13 | Deleterious |
| 102 | p.Ala102Asn | 10.55 | 3.40 | Deleterious | | | | 10.55 | 3.40 | Deleterious |
| 102 | p.Ala102Lys | 27.85 | 4.80 | Deleterious | | | | 27.85 | 4.80 | Deleterious |
| 102 | p.Ala102Thr | 11.34 | 3.50 | Deleterious | | | | 11.34 | 3.50 | Deleterious |
| 102 | p.Ala102Arg | 46.34 | 5.53 | Deleterious | | | | 46.34 | 5.53 | Deleterious |
| 102 | p.Ala102Ser | 2.38 | 1.25 | Deleterious | | | | 2.38 | 1.25 | Deleterious |
| 102 | p.Ala102Ile | 18.46 | 4.21 | Deleterious | | | | 18.46 | 4.21 | Deleterious |
| 102 | p.Ala102Met | 27.54 | 4.78 | Deleterious | | | | 27.54 | 4.78 | Deleterious |
| 102 | p.Ala102His | 17.65 | 4.14 | Deleterious | | | | 17.65 | 4.14 | Deleterious |
| 102 | p.Ala102Gln | 25.99 | 4.70 | Deleterious | | | | 25.99 | 4.70 | Deleterious |
| 102 | p.Ala102Pro | 30.62 | 4.94 | Deleterious | | | | 30.62 | 4.94 | Deleterious |
| 102 | p.Ala102Leu | 39.02 | 5.29 | Deleterious | | | | 39.02 | 5.29 | Deleterious |
| 102 | p.Ala102Asp | 27.75 | 4.79 | Deleterious | | | | 27.75 | 4.79 | Deleterious |
| 102 | p.Ala102Glu | 36.84 | 5.20 | Deleterious | | | | 36.84 | 5.20 | Deleterious |
| 102 | p.Ala102Ala | 1.00 | 0.00 | Neutral | | | | 1.00 | 0.00 | Neutral |
| 102 | p.Ala102Gly | 0.56 | -0.83 | Neutral | | | | 0.56 | -0.83 | Neutral |
| 102 | p.Ala102Val | 2.29 | 1.20 | Deleterious | | | | 2.29 | 1.20 | Deleterious |
| 102 | p.Ala102Tyr | 44.70 | 5.48 | Deleterious | | | | 44.70 | 5.48 | Deleterious |
| 102 | p.Ala102Cys | 1.34 | 0.42 | Indeterminate | | | | 1.34 | 0.42 | Indeterminate |
| 102 | p.Ala102Trp | 33.48 | 5.07 | Deleterious | | | | 33.48 | 5.07 | Deleterious |
| 102 | p.Ala102Phe | 38.33 | 5.26 | Deleterious | | | | 38.33 | 5.26 | Deleterious |
| 103 | p.Arg103Asn | 0.96 | -0.06 | Neutral | | | | 0.96 | -0.06 | Neutral |
| 103 | p.Arg103Lys | 1.02 | 0.02 | Neutral | | | | 1.02 | 0.02 | Neutral |
| 103 | p.Arg103Thr | 1.50 | 0.59 | Indeterminate | | | | 1.50 | 0.59 | Indeterminate |
| 103 | p.Arg103Arg | 1.00 | 0.00 | Neutral | | | | 1.00 | 0.00 | Neutral |
| 103 | p.Arg103Ser | 0.63 | -0.67 | Neutral | | | | 0.63 | -0.67 | Neutral |
| 103 | p.Arg103Ile | 1.31 | 0.39 | Indeterminate | | | | 1.31 | 0.39 | Indeterminate |
| 103 | p.Arg103Met | 1.03 | 0.04 | Neutral | | | | 1.03 | 0.04 | Neutral |
| 103 | p.Arg103His | 0.82 | -0.28 | Neutral | | | | 0.82 | -0.28 | Neutral |
| 103 | p.Arg103Gln | 0.99 | -0.02 | Neutral | | | | 0.99 | -0.02 | Neutral |
| 103 | p.Arg103Pro | 0.81 | -0.30 | Neutral | | | | 0.81 | -0.30 | Neutral |
| 103 | p.Arg103Leu | 0.65 | -0.62 | Neutral | | | | 0.65 | -0.62 | Neutral |
| 103 | p.Arg103Asp | 0.73 | -0.45 | Neutral | | | | 0.73 | -0.45 | Neutral |
| 103 | p.Arg103Glu | 0.80 | -0.33 | Neutral | | | | 0.80 | -0.33 | Neutral |
| 103 | p.Arg103Ala | 0.81 | -0.31 | Neutral | | | | 0.81 | -0.31 | Neutral |
| 103 | p.Arg103Gly | 0.96 | -0.05 | Neutral | | | | 0.96 | -0.05 | Neutral |
| 103 | p.Arg103Val | 0.85 | -0.23 | Neutral | | | | 0.85 | -0.23 | Neutral |
| 103 | p.Arg103Tyr | 1.09 | 0.12 | Neutral | | | | 1.09 | 0.12 | Neutral |
| 103 | p.Arg103Cys | 1.05 | 0.08 | Neutral | | | | 1.05 | 0.08 | Neutral |
| 103 | p.Arg103Trp | 0.90 | -0.16 | Neutral | | | | 0.90 | -0.16 | Neutral |
| 103 | p.Arg103Phe | 0.92 | -0.12 | Neutral | | | | 0.92 | -0.12 | Neutral |
| 104 | p.Leu104Asn | 0.44 | -1.17 | Neutral | | | | 0.44 | -1.17 | Neutral |
| 104 | p.Leu104Lys | 2.06 | 1.04 | Indeterminate | | | | 2.06 | 1.04 | Indeterminate |
| 104 | p.Leu104Thr | 0.46 | -1.11 | Neutral | | | | 0.46 | -1.11 | Neutral |
| 104 | p.Leu104Arg | 4.06 | 2.02 | Deleterious | | | | 4.06 | 2.02 | Deleterious |
| 104 | p.Leu104Ser | 0.37 | -1.43 | Neutral | | | | 0.37 | -1.43 | Neutral |
| 104 | p.Leu104Ile | 0.22 | -2.16 | Neutral | | | | 0.22 | -2.16 | Neutral |
| 104 | p.Leu104Met | 0.23 | -2.10 | Neutral | | | | 0.23 | -2.10 | Neutral |
| 104 | p.Leu104His | 0.30 | -1.73 | Neutral | | | | 0.30 | -1.73 | Neutral |
| 104 | p.Leu104Gln | 0.74 | -0.43 | Neutral | | | | 0.74 | -0.43 | Neutral |
| 104 | p.Leu104Pro | 0.51 | -0.99 | Neutral | | | | 0.51 | -0.99 | Neutral |
| 104 | p.Leu104Leu | 1.00 | 0.00 | Neutral | | | | 1.00 | 0.00 | Neutral |
| 104 | p.Leu104Asp | 0.46 | -1.13 | Neutral | | | | 0.46 | -1.13 | Neutral |
| 104 | p.Leu104Glu | 0.72 | -0.48 | Neutral | | | | 0.72 | -0.48 | Neutral |
| 104 | p.Leu104Ala | 0.20 | -2.33 | Neutral | | | | 0.20 | -2.33 | Neutral |
| 104 | p.Leu104Gly | 4.15 | 2.05 | Deleterious | | | | 4.15 | 2.05 | Deleterious |
| 104 | p.Leu104Val | 0.14 | -2.87 | Neutral | | | | 0.14 | -2.87 | Neutral |
| 104 | p.Leu104Tyr | 0.20 | -2.29 | Neutral | | | | 0.20 | -2.29 | Neutral |
| 104 | p.Leu104Cys | 0.24 | -2.04 | Neutral | | | | 0.24 | -2.04 | Neutral |
| 104 | p.Leu104Trp | 0.10 | -3.32 | Neutral | | | | 0.10 | -3.32 | Neutral |
| 104 | p.Leu104Phe | 0.10 | -3.34 | Neutral | | | | 0.10 | -3.34 | Neutral |
| 105 | p.Asp105Asn | 1.03 | 0.05 | Neutral | | | | 1.03 | 0.05 | Neutral |
| 105 | p.Asp105Lys | 1.92 | 0.94 | Indeterminate | | | | 1.92 | 0.94 | Indeterminate |
| 105 | p.Asp105Thr | 1.45 | 0.53 | Indeterminate | | | | 1.45 | 0.53 | Indeterminate |
| 105 | p.Asp105Arg | 2.45 | 1.29 | Deleterious | | | | 2.45 | 1.29 | Deleterious |
| 105 | p.Asp105Ser | 2.08 | 1.06 | Indeterminate | | | | 2.08 | 1.06 | Indeterminate |
| 105 | p.Asp105Ile | 2.64 | 1.40 | Deleterious | | | | 2.64 | 1.40 | Deleterious |
| 105 | p.Asp105Met | 1.49 | 0.57 | Indeterminate | | | | 1.49 | 0.57 | Indeterminate |
| 105 | p.Asp105His | 1.65 | 0.72 | Indeterminate | | | | 1.65 | 0.72 | Indeterminate |
| 105 | p.Asp105Gln | 1.87 | 0.90 | Indeterminate | | | | 1.87 | 0.90 | Indeterminate |
| 105 | p.Asp105Pro | 9.48 | 3.25 | Deleterious | | | | 9.48 | 3.25 | Deleterious |
| 105 | p.Asp105Leu | 1.85 | 0.89 | Indeterminate | | | | 1.85 | 0.89 | Indeterminate |
| 105 | p.Asp105Asp | 1.00 | 0.00 | Neutral | | | | 1.00 | 0.00 | Neutral |
| 105 | p.Asp105Glu | 2.10 | 1.07 | Indeterminate | | | | 2.10 | 1.07 | Indeterminate |
| 105 | p.Asp105Ala | 1.39 | 0.48 | Indeterminate | | | | 1.39 | 0.48 | Indeterminate |
| 105 | p.Asp105Gly | 2.69 | 1.43 | Deleterious | | | | 2.69 | 1.43 | Deleterious |
| 105 | p.Asp105Val | 2.38 | 1.25 | Deleterious | | | | 2.38 | 1.25 | Deleterious |
| 105 | p.Asp105Tyr | 0.54 | -0.90 | Neutral | | | | 0.54 | -0.90 | Neutral |
| 105 | p.Asp105Cys | 0.58 | -0.79 | Neutral | | | | 0.58 | -0.79 | Neutral |
| 105 | p.Asp105Trp | 1.37 | 0.46 | Indeterminate | | | | 1.37 | 0.46 | Indeterminate |
| 105 | p.Asp105Phe | 1.80 | 0.85 | Indeterminate | | | | 1.80 | 0.85 | Indeterminate |
| 106 | p.Val106Asn | 1.34 | 0.43 | Indeterminate | 1.20 | 0.26 | Indeterminate | 1.27 | 0.35 | Indeterminate |
| 106 | p.Val106Lys | 1.74 | 0.80 | Indeterminate | 1.13 | 0.17 | Indeterminate | 1.43 | 0.52 | Indeterminate |
| 106 | p.Val106Thr | 0.96 | -0.05 | Neutral | 1.03 | 0.04 | Neutral | 1.00 | -0.01 | Neutral |
| 106 | p.Val106Arg | 1.68 | 0.75 | Indeterminate | 0.93 | -0.11 | Neutral | 1.30 | 0.38 | Indeterminate |
| 106 | p.Val106Ser | 1.08 | 0.11 | Neutral | 0.98 | -0.03 | Neutral | 1.03 | 0.04 | Neutral |
| 106 | p.Val106Ile | 0.67 | -0.57 | Neutral | 1.16 | 0.22 | Indeterminate | 0.92 | -0.12 | Neutral |
| 106 | p.Val106Met | 1.00 | 0.00 | Neutral | 1.06 | 0.08 | Neutral | 1.03 | 0.04 | Neutral |
| 106 | p.Val106His | 0.96 | -0.06 | Neutral | 1.06 | 0.08 | Neutral | 1.01 | 0.02 | Neutral |
| 106 | p.Val106Gln | 0.74 | -0.43 | Neutral | 1.25 | 0.32 | Indeterminate | 0.99 | -0.01 | Neutral |
| 106 | p.Val106Pro | 1.70 | 0.76 | Indeterminate | 1.14 | 0.19 | Indeterminate | 1.42 | 0.51 | Indeterminate |
| 106 | p.Val106Leu | 1.21 | 0.27 | Indeterminate | 1.05 | 0.07 | Neutral | 1.13 | 0.17 | Neutral |
| 106 | p.Val106Asp | 0.98 | -0.03 | Neutral | 1.21 | 0.28 | Indeterminate | 1.10 | 0.13 | Neutral |
| 106 | p.Val106Glu | 1.58 | 0.66 | Indeterminate | 1.07 | 0.10 | Neutral | 1.33 | 0.41 | Indeterminate |
| 106 | p.Val106Ala | 1.47 | 0.55 | Indeterminate | 1.14 | 0.19 | Indeterminate | 1.30 | 0.38 | Indeterminate |
| 106 | p.Val106Gly | 1.55 | 0.63 | Indeterminate | 0.95 | -0.08 | Neutral | 1.25 | 0.32 | Indeterminate |
| 106 | p.Val106Val | 1.00 | 0.00 | Neutral | 1.00 | 0.00 | Neutral | 1.00 | 0.00 | Neutral |

| | | | | | | | | | | |
|---|---|---|---|---|---|---|---|---|---|---|
| 106 | p.Val106Tyr | 0.87 | -0.20 | Neutral | 1.15 | 0.20 | Indeterminate | 1.01 | 0.01 | Neutral |
| 106 | p.Val106Cys | 1.26 | 0.33 | Indeterminate | 1.01 | 0.02 | Neutral | 1.14 | 0.18 | Neutral |
| 106 | p.Val106Trp | 1.07 | 0.09 | Neutral | 0.92 | -0.11 | Neutral | 1.00 | -0.01 | Neutral |
| 106 | p.Val106Phe | 0.97 | -0.05 | Neutral | 1.19 | 0.25 | Indeterminate | 1.08 | 0.11 | Neutral |
| 107 | p.Arg107Asn | 0.93 | -0.10 | Neutral | 1.13 | 0.18 | Indeterminate | 1.03 | 0.05 | Neutral |
| 107 | p.Arg107Lys | 0.89 | -0.17 | Neutral | 0.93 | -0.11 | Neutral | 0.91 | -0.14 | Neutral |
| 107 | p.Arg107Thr | 0.91 | -0.14 | Neutral | 0.76 | -0.39 | Neutral | 0.84 | -0.26 | Neutral |
| 107 | p.Arg107Arg | 1.00 | 0.00 | Neutral | 1.00 | 0.00 | Neutral | 1.00 | 0.00 | Neutral |
| 107 | p.Arg107Ser | 1.12 | 0.16 | Neutral | 1.10 | 0.14 | Neutral | 1.11 | 0.15 | Neutral |
| 107 | p.Arg107Ile | 0.91 | -0.14 | Neutral | 0.72 | -0.47 | Neutral | 0.82 | -0.29 | Neutral |
| 107 | p.Arg107Met | 0.90 | -0.16 | Neutral | 0.92 | -0.12 | Neutral | 0.91 | -0.14 | Neutral |
| 107 | p.Arg107His | 0.88 | -0.19 | Neutral | 0.99 | -0.02 | Neutral | 0.93 | -0.10 | Neutral |
| 107 | p.Arg107Gln | 0.88 | -0.18 | Neutral | 0.81 | -0.31 | Neutral | 0.84 | -0.25 | Neutral |
| 107 | p.Arg107Pro | 0.90 | -0.15 | Neutral | 0.85 | -0.23 | Neutral | 0.88 | -0.19 | Neutral |
| 107 | p.Arg107Leu | 0.95 | -0.08 | Neutral | 1.20 | 0.26 | Indeterminate | 1.07 | 0.10 | Neutral |
| 107 | p.Arg107Asp | 0.92 | -0.12 | Neutral | 0.94 | -0.08 | Neutral | 0.93 | -0.10 | Neutral |
| 107 | p.Arg107Glu | 1.02 | 0.02 | Neutral | 1.07 | 0.10 | Neutral | 1.04 | 0.06 | Neutral |
| 107 | p.Arg107Ala | 0.89 | -0.18 | Neutral | 0.86 | -0.22 | Neutral | 0.87 | -0.20 | Neutral |
| 107 | p.Arg107Gly | 0.92 | -0.12 | Neutral | 1.04 | 0.06 | Neutral | 0.98 | -0.03 | Neutral |
| 107 | p.Arg107Val | 0.84 | -0.25 | Neutral | 1.00 | -0.01 | Neutral | 0.92 | -0.12 | Neutral |
| 107 | p.Arg107Tyr | 1.01 | 0.01 | Neutral | 0.91 | -0.14 | Neutral | 0.96 | -0.06 | Neutral |
| 107 | p.Arg107Cys | 0.79 | -0.34 | Neutral | 1.27 | 0.34 | Indeterminate | 1.03 | 0.04 | Neutral |
| 107 | p.Arg107Trp | 0.88 | -0.18 | Neutral | 0.82 | -0.29 | Neutral | 0.85 | -0.24 | Neutral |
| 107 | p.Arg107Phe | 0.96 | -0.06 | Neutral | 0.99 | -0.01 | Neutral | 0.97 | -0.04 | Neutral |
| 108 | p.Asp108Asn | 0.97 | -0.05 | Neutral | | | | 0.97 | -0.05 | Neutral |
| 108 | p.Asp108Lys | 5.96 | 2.58 | Deleterious | | | | 5.96 | 2.58 | Deleterious |
| 108 | p.Asp108Thr | 2.49 | 1.32 | Deleterious | | | | 2.49 | 1.32 | Deleterious |
| 108 | p.Asp108Arg | 6.55 | 2.71 | Deleterious | | | | 6.55 | 2.71 | Deleterious |
| 108 | p.Asp108Ser | 4.10 | 2.04 | Deleterious | | | | 4.10 | 2.04 | Deleterious |
| 108 | p.Asp108Ile | 5.55 | 2.47 | Deleterious | | | | 5.55 | 2.47 | Deleterious |
| 108 | p.Asp108Met | 5.47 | 2.45 | Deleterious | | | | 5.47 | 2.45 | Deleterious |
| 108 | p.Asp108His | 5.94 | 2.57 | Deleterious | | | | 5.94 | 2.57 | Deleterious |
| 108 | p.Asp108Gln | 5.45 | 2.45 | Deleterious | | | | 5.45 | 2.45 | Deleterious |
| 108 | p.Asp108Pro | 6.20 | 2.63 | Deleterious | | | | 6.20 | 2.63 | Deleterious |
| 108 | p.Asp108Leu | 5.82 | 2.54 | Deleterious | | | | 5.82 | 2.54 | Deleterious |
| 108 | p.Asp108Asp | 1.00 | 0.00 | Neutral | | | | 1.00 | 0.00 | Neutral |
| 108 | p.Asp108Glu | 1.43 | 0.51 | Indeterminate | | | | 1.43 | 0.51 | Indeterminate |
| 108 | p.Asp108Ala | 6.13 | 2.62 | Deleterious | | | | 6.13 | 2.62 | Deleterious |
| 108 | p.Asp108Gly | 6.11 | 2.61 | Deleterious | | | | 6.11 | 2.61 | Deleterious |
| 108 | p.Asp108Val | 5.68 | 2.50 | Deleterious | | | | 5.68 | 2.50 | Deleterious |
| 108 | p.Asp108Tyr | 5.94 | 2.57 | Deleterious | | | | 5.94 | 2.57 | Deleterious |
| 108 | p.Asp108Cys | 3.15 | 1.65 | Deleterious | | | | 3.15 | 1.65 | Deleterious |
| 108 | p.Asp108Trp | 6.04 | 2.60 | Deleterious | | | | 6.04 | 2.60 | Deleterious |
| 108 | p.Asp108Phe | 5.57 | 2.48 | Deleterious | | | | 5.57 | 2.48 | Deleterious |
| 109 | p.Ala109Asn | 0.89 | -0.16 | Neutral | 0.61 | -0.71 | Neutral | 0.75 | -0.41 | Neutral |
| 109 | p.Ala109Lys | 1.15 | 0.20 | Neutral | 1.20 | 0.26 | Indeterminate | 1.18 | 0.23 | Neutral |
| 109 | p.Ala109Thr | 1.03 | 0.05 | Neutral | 0.58 | -0.78 | Neutral | 0.81 | -0.31 | Neutral |
| 109 | p.Ala109Arg | 0.87 | -0.20 | Neutral | 1.24 | 0.31 | Indeterminate | 1.05 | 0.08 | Neutral |
| 109 | p.Ala109Ser | 0.78 | -0.36 | Neutral | 0.88 | -0.19 | Neutral | 0.83 | -0.27 | Neutral |
| 109 | p.Ala109Ile | 0.59 | -0.75 | Neutral | 0.43 | -1.23 | Neutral | 0.51 | -0.97 | Neutral |
| 109 | p.Ala109Met | 0.69 | -0.54 | Neutral | 0.71 | -0.50 | Neutral | 0.70 | -0.52 | Neutral |
| 109 | p.Ala109His | 0.77 | -0.38 | Neutral | 0.75 | -0.41 | Neutral | 0.76 | -0.39 | Neutral |
| 109 | p.Ala109Gln | 0.96 | -0.06 | Neutral | 0.78 | -0.37 | Neutral | 0.87 | -0.20 | Neutral |
| 109 | p.Ala109Pro | 13.94 | 3.80 | Deleterious | 4.34 | 2.12 | Deleterious | 9.14 | 3.19 | Deleterious |
| 109 | p.Ala109Leu | 0.52 | -0.94 | Neutral | 1.32 | 0.40 | Indeterminate | 0.92 | -0.12 | Neutral |
| 109 | p.Ala109Asp | 0.98 | -0.03 | Neutral | 0.93 | -0.10 | Neutral | 0.96 | -0.07 | Neutral |
| 109 | p.Ala109Glu | 1.38 | 0.47 | Indeterminate | 0.98 | -0.04 | Neutral | 1.18 | 0.24 | Neutral |
| 109 | p.Ala109Ala | 1.00 | 0.00 | Neutral | 1.00 | 0.00 | Neutral | 1.00 | 0.00 | Neutral |
| 109 | p.Ala109Gly | 0.77 | -0.38 | Neutral | 0.76 | -0.40 | Neutral | 0.76 | -0.39 | Neutral |
| 109 | p.Ala109Val | 0.88 | -0.19 | Neutral | 0.88 | -0.18 | Neutral | 0.88 | -0.18 | Neutral |
| 109 | p.Ala109Tyr | 0.79 | -0.33 | Neutral | 0.78 | -0.35 | Neutral | 0.79 | -0.34 | Neutral |
| 109 | p.Ala109Cys | 0.84 | -0.25 | Neutral | 0.65 | -0.62 | Neutral | 0.75 | -0.42 | Neutral |
| 109 | p.Ala109Trp | 1.03 | 0.05 | Neutral | 0.83 | -0.27 | Neutral | 0.93 | -0.11 | Neutral |
| 109 | p.Ala109Phe | 1.22 | 0.29 | Indeterminate | 0.77 | -0.37 | Neutral | 1.00 | 0.00 | Neutral |
| 110 | p.Trp110Asn | 0.96 | -0.06 | Neutral | | | | 0.96 | -0.06 | Neutral |
| 110 | p.Trp110Lys | 1.13 | 0.17 | Neutral | | | | 1.13 | 0.17 | Neutral |
| 110 | p.Trp110Thr | 0.83 | -0.26 | Neutral | | | | 0.83 | -0.26 | Neutral |
| 110 | p.Trp110Arg | 0.88 | -0.19 | Neutral | | | | 0.88 | -0.19 | Neutral |
| 110 | p.Trp110Ser | 1.15 | 0.20 | Neutral | | | | 1.15 | 0.20 | Neutral |
| 110 | p.Trp110Ile | 0.97 | -0.04 | Neutral | | | | 0.97 | -0.04 | Neutral |
| 110 | p.Trp110Met | 1.01 | 0.01 | Neutral | | | | 1.01 | 0.01 | Neutral |
| 110 | p.Trp110His | 1.06 | 0.08 | Neutral | | | | 1.06 | 0.08 | Neutral |
| 110 | p.Trp110Gln | 1.09 | 0.12 | Neutral | | | | 1.09 | 0.12 | Neutral |
| 110 | p.Trp110Pro | 1.25 | 0.32 | Indeterminate | | | | 1.25 | 0.32 | Indeterminate |
| 110 | p.Trp110Leu | 0.83 | -0.28 | Neutral | | | | 0.83 | -0.28 | Neutral |
| 110 | p.Trp110Asp | 0.73 | -0.46 | Neutral | | | | 0.73 | -0.46 | Neutral |
| 110 | p.Trp110Glu | 1.16 | 0.21 | Neutral | | | | 1.16 | 0.21 | Neutral |
| 110 | p.Trp110Ala | 1.21 | 0.28 | Indeterminate | | | | 1.21 | 0.28 | Indeterminate |
| 110 | p.Trp110Gly | 0.87 | -0.20 | Neutral | | | | 0.87 | -0.20 | Neutral |
| 110 | p.Trp110Val | 0.90 | -0.15 | Neutral | | | | 0.90 | -0.15 | Neutral |
| 110 | p.Trp110Tyr | 0.91 | -0.13 | Neutral | | | | 0.91 | -0.13 | Neutral |
| 110 | p.Trp110Cys | 0.88 | -0.19 | Neutral | | | | 0.88 | -0.19 | Neutral |
| 110 | p.Trp110Trp | 1.00 | 0.00 | Neutral | | | | 1.00 | 0.00 | Neutral |
| 110 | p.Trp110Phe | 0.77 | -0.37 | Neutral | | | | 0.77 | -0.37 | Neutral |
| 111 | p.Gly111Asn | 0.87 | -0.19 | Neutral | | | | 0.87 | -0.19 | Neutral |
| 111 | p.Gly111Lys | 1.36 | 0.44 | Indeterminate | | | | 1.36 | 0.44 | Indeterminate |
| 111 | p.Gly111Thr | 2.41 | 1.27 | Deleterious | | | | 2.41 | 1.27 | Deleterious |
| 111 | p.Gly111Arg | 1.10 | 0.13 | Neutral | | | | 1.10 | 0.13 | Neutral |
| 111 | p.Gly111Ser | 0.90 | -0.16 | Neutral | | | | 0.90 | -0.16 | Neutral |
| 111 | p.Gly111Ile | 4.46 | 2.16 | Deleterious | | | | 4.46 | 2.16 | Deleterious |
| 111 | p.Gly111Met | 0.97 | -0.05 | Neutral | | | | 0.97 | -0.05 | Neutral |
| 111 | p.Gly111His | 1.54 | 0.62 | Indeterminate | | | | 1.54 | 0.62 | Indeterminate |
| 111 | p.Gly111Gln | 0.75 | -0.41 | Neutral | | | | 0.75 | -0.41 | Neutral |
| 111 | p.Gly111Pro | 25.69 | 4.68 | Deleterious | | | | 25.69 | 4.68 | Deleterious |
| 111 | p.Gly111Leu | 0.94 | -0.09 | Neutral | | | | 0.94 | -0.09 | Neutral |
| 111 | p.Gly111Asp | 0.86 | -0.22 | Neutral | | | | 0.86 | -0.22 | Neutral |
| 111 | p.Gly111Glu | 1.23 | 0.29 | Indeterminate | | | | 1.23 | 0.29 | Indeterminate |
| 111 | p.Gly111Ala | 1.16 | 0.21 | Neutral | | | | 1.16 | 0.21 | Neutral |
| 111 | p.Gly111Gly | 1.00 | 0.00 | Neutral | | | | 1.00 | 0.00 | Neutral |
| 111 | p.Gly111Val | 3.78 | 1.92 | Deleterious | | | | 3.78 | 1.92 | Deleterious |
| 111 | p.Gly111Tyr | 1.02 | 0.03 | Neutral | | | | 1.02 | 0.03 | Neutral |

| | | | | | | | | | | |
|---|---|---|---|---|---|---|---|---|---|---|
| 111 | p.Gly111Cys | 1.12 | 0.17 | Neutral | | | | 1.12 | 0.17 | Neutral |
| 111 | p.Gly111Trp | 1.04 | 0.06 | Neutral | | | | 1.04 | 0.06 | Neutral |
| 111 | p.Gly111Phe | 0.95 | -0.07 | Neutral | | | | 0.95 | -0.07 | Neutral |
| 112 | p.Arg112Asn | 0.92 | -0.12 | Neutral | | | | 0.92 | -0.12 | Neutral |
| 112 | p.Arg112Lys | 1.20 | 0.27 | Indeterminate | | | | 1.20 | 0.27 | Indeterminate |
| 112 | p.Arg112Thr | 0.90 | -0.16 | Neutral | | | | 0.90 | -0.16 | Neutral |
| 112 | p.Arg112Arg | 1.00 | 0.00 | Neutral | | | | 1.00 | 0.00 | Neutral |
| 112 | p.Arg112Ser | 0.97 | -0.05 | Neutral | | | | 0.97 | -0.05 | Neutral |
| 112 | p.Arg112Ile | 1.16 | 0.21 | Neutral | | | | 1.16 | 0.21 | Neutral |
| 112 | p.Arg112Met | 0.66 | -0.61 | Neutral | | | | 0.66 | -0.61 | Neutral |
| 112 | p.Arg112His | 1.17 | 0.22 | Neutral | | | | 1.17 | 0.22 | Neutral |
| 112 | p.Arg112Gln | 1.29 | 0.37 | Indeterminate | | | | 1.29 | 0.37 | Indeterminate |
| 112 | p.Arg112Pro | 40.46 | 5.34 | Deleterious | | | | 40.46 | 5.34 | Deleterious |
| 112 | p.Arg112Leu | 1.87 | 0.90 | Indeterminate | | | | 1.87 | 0.90 | Indeterminate |
| 112 | p.Arg112Asp | 1.48 | 0.56 | Indeterminate | | | | 1.48 | 0.56 | Indeterminate |
| 112 | p.Arg112Glu | 1.80 | 0.85 | Indeterminate | | | | 1.80 | 0.85 | Indeterminate |
| 112 | p.Arg112Ala | 1.35 | 0.43 | Indeterminate | | | | 1.35 | 0.43 | Indeterminate |
| 112 | p.Arg112Gly | 2.33 | 1.22 | Deleterious | | | | 2.33 | 1.22 | Deleterious |
| 112 | p.Arg112Val | 1.41 | 0.49 | Indeterminate | | | | 1.41 | 0.49 | Indeterminate |
| 112 | p.Arg112Tyr | 1.16 | 0.22 | Neutral | | | | 1.16 | 0.22 | Neutral |
| 112 | p.Arg112Cys | 1.60 | 0.68 | Indeterminate | | | | 1.60 | 0.68 | Indeterminate |
| 112 | p.Arg112Trp | 1.15 | 0.20 | Neutral | | | | 1.15 | 0.20 | Neutral |
| 112 | p.Arg112Phe | 1.42 | 0.51 | Indeterminate | | | | 1.42 | 0.51 | Indeterminate |
| 113 | p.Leu113Asn | 1.12 | 0.16 | Neutral | 1.17 | 0.22 | Indeterminate | 1.14 | 0.19 | Neutral |
| 113 | p.Leu113Lys | 1.07 | 0.10 | Neutral | 0.79 | -0.35 | Neutral | 0.93 | -0.11 | Neutral |
| 113 | p.Leu113Thr | 1.25 | 0.33 | Indeterminate | 0.96 | -0.06 | Neutral | 1.10 | 0.14 | Neutral |
| 113 | p.Leu113Arg | 1.23 | 0.30 | Indeterminate | 0.81 | -0.31 | Neutral | 1.02 | 0.03 | Neutral |
| 113 | p.Leu113Ser | 1.09 | 0.12 | Neutral | 0.76 | -0.39 | Neutral | 0.92 | -0.11 | Neutral |
| 113 | p.Leu113Ile | 1.16 | 0.21 | Neutral | 0.66 | -0.59 | Neutral | 0.91 | -0.14 | Neutral |
| 113 | p.Leu113Met | 0.81 | -0.30 | Neutral | 1.33 | 0.41 | Indeterminate | 1.07 | 0.10 | Neutral |
| 113 | p.Leu113His | 1.32 | 0.40 | Indeterminate | 1.21 | 0.27 | Indeterminate | 1.27 | 0.34 | Indeterminate |
| 113 | p.Leu113Gln | 1.06 | 0.09 | Neutral | 0.31 | -1.70 | Neutral | 0.68 | -0.55 | Neutral |
| 113 | p.Leu113Pro | 1.56 | 0.64 | Indeterminate | 1.38 | 0.47 | Indeterminate | 1.47 | 0.56 | Indeterminate |
| 113 | p.Leu113Leu | 1.00 | 0.00 | Neutral | 1.00 | 0.00 | Neutral | 1.00 | 0.00 | Neutral |
| 113 | p.Leu113Asp | 1.17 | 0.23 | Neutral | 0.97 | -0.05 | Neutral | 1.07 | 0.10 | Neutral |
| 113 | p.Leu113Glu | 0.94 | -0.09 | Neutral | 1.02 | 0.03 | Neutral | 0.98 | -0.03 | Neutral |
| 113 | p.Leu113Ala | 1.10 | 0.14 | Neutral | 1.64 | 0.71 | Indeterminate | 1.37 | 0.46 | Indeterminate |
| 113 | p.Leu113Gly | 1.15 | 0.20 | Neutral | 1.35 | 0.43 | Indeterminate | 1.25 | 0.32 | Indeterminate |
| 113 | p.Leu113Val | 0.97 | -0.04 | Neutral | 0.84 | -0.25 | Neutral | 0.90 | -0.15 | Neutral |
| 113 | p.Leu113Tyr | 1.03 | 0.04 | Neutral | 0.68 | -0.56 | Neutral | 0.85 | -0.23 | Neutral |
| 113 | p.Leu113Cys | 0.98 | -0.03 | Neutral | 0.93 | -0.10 | Neutral | 0.96 | -0.06 | Neutral |
| 113 | p.Leu113Trp | 1.01 | 0.01 | Neutral | 1.19 | 0.25 | Indeterminate | 1.10 | 0.13 | Neutral |
| 113 | p.Leu113Phe | 1.23 | 0.30 | Indeterminate | 1.26 | 0.33 | Indeterminate | 1.24 | 0.31 | Indeterminate |
| 114 | p.Pro114Asn | 4.52 | 2.18 | Deleterious | | | | 4.52 | 2.18 | Deleterious |
| 114 | p.Pro114Lys | 5.61 | 2.49 | Deleterious | | | | 5.61 | 2.49 | Deleterious |
| 114 | p.Pro114Thr | 3.24 | 1.70 | Deleterious | | | | 3.24 | 1.70 | Deleterious |
| 114 | p.Pro114Arg | 4.59 | 2.20 | Deleterious | | | | 4.59 | 2.20 | Deleterious |
| 114 | p.Pro114Ser | 1.70 | 0.76 | Indeterminate | | | | 1.70 | 0.76 | Indeterminate |
| 114 | p.Pro114Ile | 5.11 | 2.35 | Deleterious | | | | 5.11 | 2.35 | Deleterious |
| 114 | p.Pro114Met | 5.18 | 2.37 | Deleterious | | | | 5.18 | 2.37 | Deleterious |
| 114 | p.Pro114His | 5.87 | 2.55 | Deleterious | | | | 5.87 | 2.55 | Deleterious |
| 114 | p.Pro114Gln | 5.24 | 2.39 | Deleterious | | | | 5.24 | 2.39 | Deleterious |
| 114 | p.Pro114Pro | 1.00 | 0.00 | Neutral | | | | 1.00 | 0.00 | Neutral |
| 114 | p.Pro114Leu | 5.02 | 2.33 | Deleterious | | | | 5.02 | 2.33 | Deleterious |
| 114 | p.Pro114Asp | 5.01 | 2.32 | Deleterious | | | | 5.01 | 2.32 | Deleterious |
| 114 | p.Pro114Glu | 4.76 | 2.25 | Deleterious | | | | 4.76 | 2.25 | Deleterious |
| 114 | p.Pro114Ala | 1.11 | 0.15 | Neutral | | | | 1.11 | 0.15 | Neutral |
| 114 | p.Pro114Gly | 3.25 | 1.70 | Deleterious | | | | 3.25 | 1.70 | Deleterious |
| 114 | p.Pro114Val | 3.00 | 1.59 | Deleterious | | | | 3.00 | 1.59 | Deleterious |
| 114 | p.Pro114Tyr | 5.53 | 2.47 | Deleterious | | | | 5.53 | 2.47 | Deleterious |
| 114 | p.Pro114Cys | 1.60 | 0.68 | Indeterminate | | | | 1.60 | 0.68 | Indeterminate |
| 114 | p.Pro114Trp | 4.96 | 2.31 | Deleterious | | | | 4.96 | 2.31 | Deleterious |
| 114 | p.Pro114Phe | 4.35 | 2.12 | Deleterious | | | | 4.35 | 2.12 | Deleterious |
| 115 | p.Val115Asn | 1.01 | 0.02 | Neutral | | | | 1.01 | 0.02 | Neutral |
| 115 | p.Val115Lys | 1.02 | 0.03 | Neutral | | | | 1.02 | 0.03 | Neutral |
| 115 | p.Val115Thr | 0.96 | -0.05 | Neutral | | | | 0.96 | -0.05 | Neutral |
| 115 | p.Val115Arg | 1.48 | 0.56 | Indeterminate | | | | 1.48 | 0.56 | Indeterminate |
| 115 | p.Val115Ser | 1.12 | 0.17 | Neutral | | | | 1.12 | 0.17 | Neutral |
| 115 | p.Val115Ile | 0.93 | -0.10 | Neutral | | | | 0.93 | -0.10 | Neutral |
| 115 | p.Val115Met | 0.98 | -0.03 | Neutral | | | | 0.98 | -0.03 | Neutral |
| 115 | p.Val115His | 1.17 | 0.22 | Neutral | | | | 1.17 | 0.22 | Neutral |
| 115 | p.Val115Gln | 1.49 | 0.58 | Indeterminate | | | | 1.49 | 0.58 | Indeterminate |
| 115 | p.Val115Pro | 2.46 | 1.30 | Deleterious | | | | 2.46 | 1.30 | Deleterious |
| 115 | p.Val115Leu | 1.20 | 0.27 | Indeterminate | | | | 1.20 | 0.27 | Indeterminate |
| 115 | p.Val115Asp | 1.30 | 0.38 | Indeterminate | | | | 1.30 | 0.38 | Indeterminate |
| 115 | p.Val115Glu | 1.20 | 0.27 | Indeterminate | | | | 1.20 | 0.27 | Indeterminate |
| 115 | p.Val115Ala | 1.45 | 0.53 | Indeterminate | | | | 1.45 | 0.53 | Indeterminate |
| 115 | p.Val115Gly | 1.21 | 0.28 | Indeterminate | | | | 1.21 | 0.28 | Indeterminate |
| 115 | p.Val115Val | 1.00 | 0.00 | Neutral | | | | 1.00 | 0.00 | Neutral |
| 115 | p.Val115Tyr | 0.96 | -0.06 | Neutral | | | | 0.96 | -0.06 | Neutral |
| 115 | p.Val115Cys | 0.94 | -0.09 | Neutral | | | | 0.94 | -0.09 | Neutral |
| 115 | p.Val115Trp | 1.07 | 0.10 | Neutral | | | | 1.07 | 0.10 | Neutral |
| 115 | p.Val115Phe | 1.11 | 0.15 | Neutral | | | | 1.11 | 0.15 | Neutral |
| 116 | p.Asp116Asn | 0.98 | -0.03 | Neutral | | | | 0.98 | -0.03 | Neutral |
| 116 | p.Asp116Lys | 0.99 | -0.01 | Neutral | | | | 0.99 | -0.01 | Neutral |
| 116 | p.Asp116Thr | 1.18 | 0.24 | Neutral | | | | 1.18 | 0.24 | Neutral |
| 116 | p.Asp116Arg | 1.18 | 0.23 | Neutral | | | | 1.18 | 0.23 | Neutral |
| 116 | p.Asp116Ser | 1.04 | 0.06 | Neutral | | | | 1.04 | 0.06 | Neutral |
| 116 | p.Asp116Ile | 1.19 | 0.25 | Indeterminate | | | | 1.19 | 0.25 | Indeterminate |
| 116 | p.Asp116Met | 0.82 | -0.29 | Neutral | | | | 0.82 | -0.29 | Neutral |
| 116 | p.Asp116His | 1.05 | 0.07 | Neutral | | | | 1.05 | 0.07 | Neutral |
| 116 | p.Asp116Gln | 1.04 | 0.06 | Neutral | | | | 1.04 | 0.06 | Neutral |
| 116 | p.Asp116Pro | 4.74 | 2.25 | Deleterious | | | | 4.74 | 2.25 | Deleterious |
| 116 | p.Asp116Leu | 1.09 | 0.13 | Neutral | | | | 1.09 | 0.13 | Neutral |
| 116 | p.Asp116Asp | 1.00 | 0.00 | Neutral | | | | 1.00 | 0.00 | Neutral |
| 116 | p.Asp116Glu | 1.10 | 0.14 | Neutral | | | | 1.10 | 0.14 | Neutral |
| 116 | p.Asp116Ala | 1.01 | 0.01 | Neutral | | | | 1.01 | 0.01 | Neutral |
| 116 | p.Asp116Gly | 1.47 | 0.56 | Indeterminate | | | | 1.47 | 0.56 | Indeterminate |
| 116 | p.Asp116Val | 1.12 | 0.17 | Neutral | | | | 1.12 | 0.17 | Neutral |
| 116 | p.Asp116Tyr | 0.91 | -0.14 | Neutral | | | | 0.91 | -0.14 | Neutral |
| 116 | p.Asp116Cys | 1.07 | 0.10 | Neutral | | | | 1.07 | 0.10 | Neutral |

| | | | | | | | | | | |
|---|---|---|---|---|---|---|---|---|---|---|
| 116 | p.Asp116Trp | 1.36 | 0.45 | Indeterminate | | | | 1.36 | 0.45 | Indeterminate |
| 116 | p.Asp116Phe | 1.17 | 0.23 | Neutral | | | | 1.17 | 0.23 | Neutral |
| 117 | p.Leu117Asn | 1.03 | 0.04 | Neutral | | | | 1.03 | 0.04 | Neutral |
| 117 | p.Leu117Lys | 1.05 | 0.07 | Neutral | | | | 1.05 | 0.07 | Neutral |
| 117 | p.Leu117Thr | 0.96 | -0.07 | Neutral | | | | 0.96 | -0.07 | Neutral |
| 117 | p.Leu117Arg | 1.09 | 0.13 | Neutral | | | | 1.09 | 0.13 | Neutral |
| 117 | p.Leu117Ser | 1.06 | 0.08 | Neutral | | | | 1.06 | 0.08 | Neutral |
| 117 | p.Leu117Ile | 1.00 | 0.01 | Neutral | | | | 1.00 | 0.01 | Neutral |
| 117 | p.Leu117Met | 1.02 | 0.03 | Neutral | | | | 1.02 | 0.03 | Neutral |
| 117 | p.Leu117His | 0.84 | -0.25 | Neutral | | | | 0.84 | -0.25 | Neutral |
| 117 | p.Leu117Gln | 1.18 | 0.24 | Indeterminate | | | | 1.18 | 0.24 | Indeterminate |
| 117 | p.Leu117Pro | 2.26 | 1.18 | Deleterious | | | | 2.26 | 1.18 | Deleterious |
| 117 | p.Leu117Leu | 1.00 | 0.00 | Neutral | | | | 1.00 | 0.00 | Neutral |
| 117 | p.Leu117Asp | 0.97 | -0.04 | Neutral | | | | 0.97 | -0.04 | Neutral |
| 117 | p.Leu117Glu | 1.10 | 0.14 | Neutral | | | | 1.10 | 0.14 | Neutral |
| 117 | p.Leu117Ala | 0.95 | -0.08 | Neutral | | | | 0.95 | -0.08 | Neutral |
| 117 | p.Leu117Gly | 1.07 | 0.10 | Neutral | | | | 1.07 | 0.10 | Neutral |
| 117 | p.Leu117Val | 1.03 | 0.05 | Neutral | | | | 1.03 | 0.05 | Neutral |
| 117 | p.Leu117Tyr | 0.94 | -0.09 | Neutral | | | | 0.94 | -0.09 | Neutral |
| 117 | p.Leu117Cys | 1.10 | 0.14 | Neutral | | | | 1.10 | 0.14 | Neutral |
| 117 | p.Leu117Trp | 0.96 | -0.06 | Neutral | | | | 0.96 | -0.06 | Neutral |
| 117 | p.Leu117Phe | 1.05 | 0.07 | Neutral | | | | 1.05 | 0.07 | Neutral |
| 118 | p.Ala118Asn | 2.04 | 1.03 | Indeterminate | | | | 2.04 | 1.03 | Indeterminate |
| 118 | p.Ala118Lys | 2.34 | 1.23 | Deleterious | | | | 2.34 | 1.23 | Deleterious |
| 118 | p.Ala118Thr | 1.28 | 0.36 | Indeterminate | | | | 1.28 | 0.36 | Indeterminate |
| 118 | p.Ala118Arg | 2.23 | 1.15 | Deleterious | | | | 2.23 | 1.15 | Deleterious |
| 118 | p.Ala118Ser | 1.18 | 0.24 | Indeterminate | | | | 1.18 | 0.24 | Indeterminate |
| 118 | p.Ala118Ile | 1.60 | 0.67 | Indeterminate | | | | 1.60 | 0.67 | Indeterminate |
| 118 | p.Ala118Met | 1.74 | 0.80 | Indeterminate | | | | 1.74 | 0.80 | Indeterminate |
| 118 | p.Ala118His | 2.19 | 1.13 | Deleterious | | | | 2.19 | 1.13 | Deleterious |
| 118 | p.Ala118Gln | 2.01 | 1.01 | Indeterminate | | | | 2.01 | 1.01 | Indeterminate |
| 118 | p.Ala118Pro | 1.63 | 0.71 | Indeterminate | | | | 1.63 | 0.71 | Indeterminate |
| 118 | p.Ala118Leu | 1.46 | 0.55 | Indeterminate | | | | 1.46 | 0.55 | Indeterminate |
| 118 | p.Ala118Asp | 2.15 | 1.10 | Deleterious | | | | 2.15 | 1.10 | Deleterious |
| 118 | p.Ala118Glu | 2.11 | 1.08 | Indeterminate | | | | 2.11 | 1.08 | Indeterminate |
| 118 | p.Ala118Ala | 1.00 | 0.00 | Neutral | | | | 1.00 | 0.00 | Neutral |
| 118 | p.Ala118Gly | 0.98 | -0.02 | Neutral | | | | 0.98 | -0.02 | Neutral |
| 118 | p.Ala118Val | 1.29 | 0.37 | Indeterminate | | | | 1.29 | 0.37 | Indeterminate |
| 118 | p.Ala118Tyr | 2.02 | 1.02 | Indeterminate | | | | 2.02 | 1.02 | Indeterminate |
| 118 | p.Ala118Cys | 1.11 | 0.15 | Neutral | | | | 1.11 | 0.15 | Neutral |
| 118 | p.Ala118Trp | 2.40 | 1.26 | Deleterious | | | | 2.40 | 1.26 | Deleterious |
| 118 | p.Ala118Phe | 2.05 | 1.04 | Indeterminate | | | | 2.05 | 1.04 | Indeterminate |
| 119 | p.Glu119Asn | 0.89 | -0.17 | Neutral | | | | 0.89 | -0.17 | Neutral |
| 119 | p.Glu119Lys | 0.74 | -0.44 | Neutral | | | | 0.74 | -0.44 | Neutral |
| 119 | p.Glu119Thr | 0.71 | -0.48 | Neutral | | | | 0.71 | -0.48 | Neutral |
| 119 | p.Glu119Arg | 1.28 | 0.36 | Indeterminate | | | | 1.28 | 0.36 | Indeterminate |
| 119 | p.Glu119Ser | 0.40 | -1.31 | Neutral | | | | 0.40 | -1.31 | Neutral |
| 119 | p.Glu119Ile | 0.63 | -0.67 | Neutral | | | | 0.63 | -0.67 | Neutral |
| 119 | p.Glu119Met | 1.29 | 0.37 | Indeterminate | | | | 1.29 | 0.37 | Indeterminate |
| 119 | p.Glu119His | 2.07 | 1.05 | Indeterminate | | | | 2.07 | 1.05 | Indeterminate |
| 119 | p.Glu119Gln | 1.79 | 0.84 | Indeterminate | | | | 1.79 | 0.84 | Indeterminate |
| 119 | p.Glu119Pro | 2.25 | 1.17 | Deleterious | | | | 2.25 | 1.17 | Deleterious |
| 119 | p.Glu119Leu | 0.83 | -0.27 | Neutral | | | | 0.83 | -0.27 | Neutral |
| 119 | p.Glu119Asp | 1.38 | 0.46 | Indeterminate | | | | 1.38 | 0.46 | Indeterminate |
| 119 | p.Glu119Glu | 1.00 | 0.00 | Neutral | | | | 1.00 | 0.00 | Neutral |
| 119 | p.Glu119Ala | 0.53 | -0.90 | Neutral | | | | 0.53 | -0.90 | Neutral |
| 119 | p.Glu119Gly | 0.89 | -0.16 | Neutral | | | | 0.89 | -0.16 | Neutral |
| 119 | p.Glu119Val | 1.27 | 0.35 | Indeterminate | | | | 1.27 | 0.35 | Indeterminate |
| 119 | p.Glu119Tyr | 0.42 | -1.26 | Neutral | | | | 0.42 | -1.26 | Neutral |
| 119 | p.Glu119Cys | 0.49 | -1.04 | Neutral | | | | 0.49 | -1.04 | Neutral |
| 119 | p.Glu119Trp | 2.09 | 1.06 | Indeterminate | | | | 2.09 | 1.06 | Indeterminate |
| 119 | p.Glu119Phe | 2.16 | 1.11 | Deleterious | | | | 2.16 | 1.11 | Deleterious |
| 120 | p.Glu120Asn | 1.20 | 0.26 | Indeterminate | 1.02 | 0.03 | Neutral | 1.11 | 0.15 | Neutral |
| 120 | p.Glu120Lys | 1.07 | 0.09 | Neutral | 1.01 | 0.02 | Neutral | 1.04 | 0.06 | Neutral |
| 120 | p.Glu120Thr | 1.11 | 0.15 | Neutral | 0.76 | -0.40 | Neutral | 0.93 | -0.10 | Neutral |
| 120 | p.Glu120Arg | 1.11 | 0.15 | Neutral | 1.01 | 0.01 | Neutral | 1.06 | 0.08 | Neutral |
| 120 | p.Glu120Ser | 1.05 | 0.07 | Neutral | 0.81 | -0.30 | Neutral | 0.93 | -0.10 | Neutral |
| 120 | p.Glu120Ile | 1.36 | 0.44 | Indeterminate | 0.91 | -0.14 | Neutral | 1.13 | 0.18 | Neutral |
| 120 | p.Glu120Met | 1.43 | 0.52 | Indeterminate | 1.02 | 0.03 | Neutral | 1.23 | 0.29 | Indeterminate |
| 120 | p.Glu120His | 1.10 | 0.14 | Neutral | 1.06 | 0.08 | Neutral | 1.08 | 0.11 | Neutral |
| 120 | p.Glu120Gln | 1.24 | 0.31 | Indeterminate | 1.00 | 0.00 | Neutral | 1.12 | 0.17 | Neutral |
| 120 | p.Glu120Pro | 1.20 | 0.27 | Indeterminate | 0.87 | -0.20 | Neutral | 1.04 | 0.05 | Neutral |
| 120 | p.Glu120Leu | 0.92 | -0.11 | Neutral | 0.95 | -0.08 | Neutral | 0.94 | -0.10 | Neutral |
| 120 | p.Glu120Asp | 1.13 | 0.17 | Neutral | 1.05 | 0.07 | Neutral | 1.09 | 0.12 | Neutral |
| 120 | p.Glu120Glu | 1.00 | 0.00 | Neutral | 1.00 | 0.00 | Neutral | 1.00 | 0.00 | Neutral |
| 120 | p.Glu120Ala | 1.03 | 0.04 | Neutral | 0.65 | -0.62 | Neutral | 0.84 | -0.25 | Neutral |
| 120 | p.Glu120Gly | 1.08 | 0.11 | Neutral | 0.76 | -0.40 | Neutral | 0.92 | -0.13 | Neutral |
| 120 | p.Glu120Val | 1.11 | 0.16 | Neutral | 0.96 | -0.06 | Neutral | 1.04 | 0.05 | Neutral |
| 120 | p.Glu120Tyr | 1.05 | 0.07 | Neutral | 1.08 | 0.11 | Neutral | 1.06 | 0.09 | Neutral |
| 120 | p.Glu120Cys | 1.16 | 0.21 | Neutral | 0.99 | -0.02 | Neutral | 1.07 | 0.10 | Neutral |
| 120 | p.Glu120Trp | 1.24 | 0.31 | Indeterminate | 0.86 | -0.23 | Neutral | 1.05 | 0.07 | Neutral |
| 120 | p.Glu120Phe | 1.03 | 0.04 | Neutral | 0.85 | -0.24 | Neutral | 0.94 | -0.10 | Neutral |
| 121 | p.Leu121Asn | 1.30 | 0.38 | Indeterminate | | | | 1.30 | 0.38 | Indeterminate |
| 121 | p.Leu121Lys | 1.12 | 0.16 | Neutral | | | | 1.12 | 0.16 | Neutral |
| 121 | p.Leu121Thr | 1.18 | 0.24 | Indeterminate | | | | 1.18 | 0.24 | Indeterminate |
| 121 | p.Leu121Arg | 1.76 | 0.81 | Indeterminate | | | | 1.76 | 0.81 | Indeterminate |
| 121 | p.Leu121Ser | 1.14 | 0.19 | Neutral | | | | 1.14 | 0.19 | Neutral |
| 121 | p.Leu121Ile | 1.13 | 0.18 | Neutral | | | | 1.13 | 0.18 | Neutral |
| 121 | p.Leu121Met | 4.46 | 2.16 | Deleterious | | | | 4.46 | 2.16 | Deleterious |
| 121 | p.Leu121His | 0.92 | -0.12 | Neutral | | | | 0.92 | -0.12 | Neutral |
| 121 | p.Leu121Gln | 1.09 | 0.12 | Neutral | | | | 1.09 | 0.12 | Neutral |
| 121 | p.Leu121Pro | 0.96 | -0.05 | Neutral | | | | 0.96 | -0.05 | Neutral |
| 121 | p.Leu121Leu | 1.00 | 0.00 | Neutral | | | | 1.00 | 0.00 | Neutral |
| 121 | p.Leu121Asp | 0.65 | -0.62 | Neutral | | | | 0.65 | -0.62 | Neutral |
| 121 | p.Leu121Glu | 1.42 | 0.50 | Indeterminate | | | | 1.42 | 0.50 | Indeterminate |
| 121 | p.Leu121Ala | 1.14 | 0.19 | Neutral | | | | 1.14 | 0.19 | Neutral |
| 121 | p.Leu121Gly | 1.36 | 0.45 | Indeterminate | | | | 1.36 | 0.45 | Indeterminate |
| 121 | p.Leu121Val | 0.84 | -0.25 | Neutral | | | | 0.84 | -0.25 | Neutral |
| 121 | p.Leu121Tyr | 0.74 | -0.43 | Neutral | | | | 0.74 | -0.43 | Neutral |
| 121 | p.Leu121Cys | 0.91 | -0.13 | Neutral | | | | 0.91 | -0.13 | Neutral |
| 121 | p.Leu121Trp | 1.32 | 0.40 | Indeterminate | | | | 1.32 | 0.40 | Indeterminate |

| | | | | | | | | | | |
|---|---|---|---|---|---|---|---|---|---|---|
| 121 | p.Leu121Phe | 0.99 | -0.02 | Neutral | | | | 0.99 | -0.02 | Neutral |
| 122 | p.Gly122Asn | 0.93 | -0.11 | Neutral | 0.50 | -0.99 | Neutral | 0.71 | -0.48 | Neutral |
| 122 | p.Gly122Lys | 1.13 | 0.18 | Neutral | 0.59 | -0.76 | Neutral | 0.86 | -0.21 | Neutral |
| 122 | p.Gly122Thr | 1.04 | 0.06 | Neutral | 0.78 | -0.35 | Neutral | 0.91 | -0.13 | Neutral |
| 122 | p.Gly122Arg | 1.30 | 0.38 | Indeterminate | 1.17 | 0.23 | Indeterminate | 1.24 | 0.31 | Indeterminate |
| 122 | p.Gly122Ser | 0.94 | -0.10 | Neutral | 0.78 | -0.36 | Neutral | 0.86 | -0.22 | Neutral |
| 122 | p.Gly122Ile | 1.30 | 0.37 | Indeterminate | 0.68 | -0.56 | Neutral | 0.99 | -0.02 | Neutral |
| 122 | p.Gly122Met | 0.97 | -0.05 | Neutral | 0.99 | -0.02 | Neutral | 0.98 | -0.03 | Neutral |
| 122 | p.Gly122His | 0.95 | -0.08 | Neutral | 0.88 | -0.19 | Neutral | 0.91 | -0.13 | Neutral |
| 122 | p.Gly122Gln | 1.03 | 0.04 | Neutral | 0.63 | -0.67 | Neutral | 0.83 | -0.27 | Neutral |
| 122 | p.Gly122Pro | 0.99 | -0.01 | Neutral | 1.24 | 0.31 | Indeterminate | 1.12 | 0.16 | Neutral |
| 122 | p.Gly122Leu | 1.10 | 0.14 | Neutral | 0.65 | -0.61 | Neutral | 0.88 | -0.19 | Neutral |
| 122 | p.Gly122Asp | 1.08 | 0.12 | Neutral | 0.63 | -0.67 | Neutral | 0.86 | -0.23 | Neutral |
| 122 | p.Gly122Glu | 1.17 | 0.23 | Neutral | 0.65 | -0.61 | Neutral | 0.91 | -0.13 | Neutral |
| 122 | p.Gly122Ala | 1.10 | 0.14 | Neutral | 0.74 | -0.44 | Neutral | 0.92 | -0.12 | Neutral |
| 122 | p.Gly122Gly | 1.00 | 0.00 | Neutral | 1.00 | 0.00 | Neutral | 1.00 | 0.00 | Neutral |
| 122 | p.Gly122Val | 0.88 | -0.19 | Neutral | 0.92 | -0.13 | Neutral | 0.90 | -0.16 | Neutral |
| 122 | p.Gly122Tyr | 1.06 | 0.08 | Neutral | 1.13 | 0.18 | Indeterminate | 1.09 | 0.13 | Neutral |
| 122 | p.Gly122Cys | 1.33 | 0.41 | Indeterminate | 0.96 | -0.06 | Neutral | 1.15 | 0.20 | Neutral |
| 122 | p.Gly122Trp | 0.85 | -0.23 | Neutral | 1.41 | 0.49 | Indeterminate | 1.13 | 0.18 | Neutral |
| 122 | p.Gly122Phe | 1.29 | 0.37 | Indeterminate | 1.03 | 0.04 | Neutral | 1.16 | 0.22 | Neutral |
| 123 | p.His123Asn | 1.34 | 0.43 | Indeterminate | 1.12 | 0.16 | Indeterminate | 1.23 | 0.30 | Indeterminate |
| 123 | p.His123Lys | 0.84 | -0.26 | Neutral | 0.98 | -0.03 | Neutral | 0.91 | -0.14 | Neutral |
| 123 | p.His123Thr | 1.62 | 0.70 | Indeterminate | 1.50 | 0.58 | Indeterminate | 1.56 | 0.64 | Indeterminate |
| 123 | p.His123Arg | 1.13 | 0.17 | Neutral | 1.06 | 0.09 | Neutral | 1.09 | 0.13 | Neutral |
| 123 | p.His123Ser | 1.20 | 0.27 | Indeterminate | 0.95 | -0.08 | Neutral | 1.07 | 0.10 | Neutral |
| 123 | p.His123Ile | 1.28 | 0.36 | Indeterminate | 1.04 | 0.05 | Neutral | 1.16 | 0.21 | Neutral |
| 123 | p.His123Met | 1.54 | 0.62 | Indeterminate | 1.32 | 0.40 | Indeterminate | 1.43 | 0.51 | Indeterminate |
| 123 | p.His123His | 1.00 | 0.00 | Neutral | 1.00 | 0.00 | Neutral | 1.00 | 0.00 | Neutral |
| 123 | p.His123Gln | 1.06 | 0.08 | Neutral | 0.94 | -0.09 | Neutral | 1.00 | 0.00 | Neutral |
| 123 | p.His123Pro | 3.51 | 1.81 | Deleterious | 4.29 | 2.10 | Deleterious | 3.90 | 1.96 | Deleterious |
| 123 | p.His123Leu | 1.28 | 0.36 | Indeterminate | 1.30 | 0.38 | Indeterminate | 1.29 | 0.37 | Indeterminate |
| 123 | p.His123Asp | 1.57 | 0.65 | Indeterminate | 1.58 | 0.66 | Indeterminate | 1.57 | 0.65 | Indeterminate |
| 123 | p.His123Glu | 1.25 | 0.33 | Indeterminate | 1.82 | 0.86 | Indeterminate | 1.54 | 0.62 | Indeterminate |
| 123 | p.His123Ala | 1.09 | 0.12 | Neutral | 1.15 | 0.20 | Indeterminate | 1.12 | 0.16 | Neutral |
| 123 | p.His123Gly | 1.84 | 0.88 | Indeterminate | 1.59 | 0.67 | Indeterminate | 1.72 | 0.78 | Indeterminate |
| 123 | p.His123Val | 1.53 | 0.61 | Indeterminate | 1.98 | 0.99 | Indeterminate | 1.76 | 0.81 | Indeterminate |
| 123 | p.His123Tyr | 1.01 | 0.01 | Neutral | 0.81 | -0.30 | Neutral | 0.91 | -0.14 | Neutral |
| 123 | p.His123Cys | 1.28 | 0.36 | Indeterminate | 0.95 | -0.07 | Neutral | 1.12 | 0.16 | Neutral |
| 123 | p.His123Trp | 1.04 | 0.06 | Neutral | 1.06 | 0.09 | Neutral | 1.05 | 0.07 | Neutral |
| 123 | p.His123Phe | 1.13 | 0.17 | Neutral | 1.93 | 0.95 | Indeterminate | 1.53 | 0.61 | Indeterminate |
| 124 | p.Arg124Asn | 0.79 | -0.34 | Neutral | 0.64 | -0.64 | Neutral | 0.71 | -0.48 | Neutral |
| 124 | p.Arg124Lys | 0.91 | -0.14 | Neutral | 0.89 | -0.16 | Neutral | 0.90 | -0.15 | Neutral |
| 124 | p.Arg124Thr | 0.87 | -0.20 | Neutral | 1.31 | 0.39 | Indeterminate | 1.09 | 0.13 | Neutral |
| 124 | p.Arg124Arg | 1.00 | 0.00 | Neutral | 1.00 | 0.00 | Neutral | 1.00 | 0.00 | Neutral |
| 124 | p.Arg124Ser | 0.82 | -0.29 | Neutral | 0.59 | -0.77 | Neutral | 0.70 | -0.51 | Neutral |
| 124 | p.Arg124Ile | 1.14 | 0.19 | Neutral | 2.49 | 1.31 | Indeterminate | 1.81 | 0.86 | Indeterminate |
| 124 | p.Arg124Met | 0.89 | -0.17 | Neutral | 0.31 | -1.67 | Neutral | 0.60 | -0.74 | Neutral |
| 124 | p.Arg124His | 0.89 | -0.17 | Neutral | 0.42 | -1.25 | Neutral | 0.66 | -0.61 | Neutral |
| 124 | p.Arg124Gln | 0.84 | -0.25 | Neutral | 0.67 | -0.57 | Neutral | 0.76 | -0.40 | Neutral |
| 124 | p.Arg124Pro | 0.89 | -0.18 | Neutral | 0.58 | -0.77 | Neutral | 0.74 | -0.44 | Neutral |
| 124 | p.Arg124Leu | 0.91 | -0.14 | Neutral | 0.82 | -0.28 | Neutral | 0.86 | -0.21 | Neutral |
| 124 | p.Arg124Asp | 0.78 | -0.37 | Neutral | 0.49 | -1.02 | Neutral | 0.63 | -0.66 | Neutral |
| 124 | p.Arg124Glu | 0.81 | -0.31 | Neutral | 0.93 | -0.10 | Neutral | 0.87 | -0.20 | Neutral |
| 124 | p.Arg124Ala | 0.86 | -0.22 | Neutral | 0.57 | -0.82 | Neutral | 0.71 | -0.49 | Neutral |
| 124 | p.Arg124Gly | 1.25 | 0.32 | Indeterminate | 1.07 | 0.10 | Neutral | 1.16 | 0.22 | Neutral |
| 124 | p.Arg124Val | 0.91 | -0.14 | Neutral | 1.08 | 0.12 | Neutral | 1.00 | 0.00 | Neutral |
| 124 | p.Arg124Tyr | 0.84 | -0.26 | Neutral | 1.76 | 0.82 | Indeterminate | 1.30 | 0.38 | Indeterminate |
| 124 | p.Arg124Cys | 1.00 | 0.00 | Neutral | 0.45 | -1.14 | Neutral | 0.73 | -0.46 | Neutral |
| 124 | p.Arg124Trp | 0.92 | -0.12 | Neutral | 0.62 | -0.69 | Neutral | 0.77 | -0.38 | Neutral |
| 124 | p.Arg124Phe | 0.79 | -0.35 | Neutral | 0.86 | -0.22 | Neutral | 0.82 | -0.28 | Neutral |
| 125 | p.Asp125Asn | 1.10 | 0.14 | Neutral | 0.80 | -0.32 | Neutral | 0.95 | -0.07 | Neutral |
| 125 | p.Asp125Lys | 0.90 | -0.15 | Neutral | 0.75 | -0.42 | Neutral | 0.82 | -0.28 | Neutral |
| 125 | p.Asp125Thr | 0.72 | -0.47 | Neutral | 0.81 | -0.30 | Neutral | 0.77 | -0.38 | Neutral |
| 125 | p.Asp125Arg | 1.00 | 0.00 | Neutral | 0.92 | -0.13 | Neutral | 0.96 | -0.06 | Neutral |
| 125 | p.Asp125Ser | 0.72 | -0.47 | Neutral | 0.96 | -0.07 | Neutral | 0.84 | -0.25 | Neutral |
| 125 | p.Asp125Ile | 0.71 | -0.50 | Neutral | 0.69 | -0.53 | Neutral | 0.70 | -0.52 | Neutral |
| 125 | p.Asp125Met | 0.68 | -0.55 | Neutral | 0.76 | -0.40 | Neutral | 0.72 | -0.47 | Neutral |
| 125 | p.Asp125His | 0.88 | -0.18 | Neutral | 0.75 | -0.42 | Neutral | 0.81 | -0.30 | Neutral |
| 125 | p.Asp125Gln | 0.70 | -0.52 | Neutral | 0.69 | -0.53 | Neutral | 0.69 | -0.53 | Neutral |
| 125 | p.Asp125Pro | 0.72 | -0.48 | Neutral | 0.88 | -0.18 | Neutral | 0.80 | -0.32 | Neutral |
| 125 | p.Asp125Leu | 0.80 | -0.32 | Neutral | 0.98 | -0.02 | Neutral | 0.89 | -0.16 | Neutral |
| 125 | p.Asp125Asp | 1.00 | 0.00 | Neutral | 1.00 | 0.00 | Neutral | 1.00 | 0.00 | Neutral |
| 125 | p.Asp125Glu | 0.70 | -0.51 | Neutral | 0.94 | -0.10 | Neutral | 0.82 | -0.29 | Neutral |
| 125 | p.Asp125Ala | 0.72 | -0.47 | Neutral | 1.18 | 0.24 | Indeterminate | 0.95 | -0.07 | Neutral |
| 125 | p.Asp125Gly | 1.17 | 0.22 | Neutral | 0.64 | -0.64 | Neutral | 0.90 | -0.15 | Neutral |
| 125 | p.Asp125Val | 0.51 | -0.96 | Neutral | 0.72 | -0.46 | Neutral | 0.62 | -0.69 | Neutral |
| 125 | p.Asp125Tyr | 2.39 | 1.26 | Deleterious | 2.67 | 1.42 | Indeterminate | 2.53 | 1.34 | Deleterious |
| 125 | p.Asp125Cys | 1.10 | 0.13 | Neutral | 0.61 | -0.72 | Neutral | 0.85 | -0.23 | Neutral |
| 125 | p.Asp125Trp | 0.88 | -0.18 | Neutral | 0.91 | -0.13 | Neutral | 0.90 | -0.15 | Neutral |
| 125 | p.Asp125Phe | 0.75 | -0.42 | Neutral | 1.12 | 0.16 | Indeterminate | 0.93 | -0.10 | Neutral |
| 126 | p.Val126Asn | 0.98 | -0.03 | Neutral | | | | 0.98 | -0.03 | Neutral |
| 126 | p.Val126Lys | 6.93 | 2.79 | Deleterious | | | | 6.93 | 2.79 | Deleterious |
| 126 | p.Val126Thr | 0.41 | -1.28 | Neutral | | | | 0.41 | -1.28 | Neutral |
| 126 | p.Val126Arg | 9.89 | 3.31 | Deleterious | | | | 9.89 | 3.31 | Deleterious |
| 126 | p.Val126Ser | 0.64 | -0.65 | Neutral | | | | 0.64 | -0.65 | Neutral |
| 126 | p.Val126Ile | 0.66 | -0.60 | Neutral | | | | 0.66 | -0.60 | Neutral |
| 126 | p.Val126Met | 0.64 | -0.64 | Neutral | | | | 0.64 | -0.64 | Neutral |
| 126 | p.Val126His | 3.14 | 1.65 | Deleterious | | | | 3.14 | 1.65 | Deleterious |
| 126 | p.Val126Gln | 1.21 | 0.27 | Indeterminate | | | | 1.21 | 0.27 | Indeterminate |
| 126 | p.Val126Pro | 0.56 | -0.83 | Neutral | | | | 0.56 | -0.83 | Neutral |
| 126 | p.Val126Leu | 0.56 | -0.85 | Neutral | | | | 0.56 | -0.85 | Neutral |
| 126 | p.Val126Asp | 6.68 | 2.74 | Deleterious | | | | 6.68 | 2.74 | Deleterious |
| 126 | p.Val126Glu | 0.77 | -0.38 | Neutral | | | | 0.77 | -0.38 | Neutral |
| 126 | p.Val126Ala | 0.72 | -0.46 | Neutral | | | | 0.72 | -0.46 | Neutral |
| 126 | p.Val126Gly | 0.96 | -0.06 | Neutral | | | | 0.96 | -0.06 | Neutral |
| 126 | p.Val126Val | 1.00 | 0.00 | Neutral | | | | 1.00 | 0.00 | Neutral |
| 126 | p.Val126Tyr | 7.32 | 2.87 | Deleterious | | | | 7.32 | 2.87 | Deleterious |
| 126 | p.Val126Cys | 0.45 | -1.15 | Neutral | | | | 0.45 | -1.15 | Neutral |
| 126 | p.Val126Trp | 8.66 | 3.11 | Deleterious | | | | 8.66 | 3.11 | Deleterious |
| 126 | p.Val126Phe | 0.97 | -0.04 | Neutral | | | | 0.97 | -0.04 | Neutral |

| | | | | | | | |
|---|---|---|---|---|---|---|---|
| 127 | p.Ala127Asn | 0.98 | -0.03 | Neutral | 0.98 | -0.03 | Neutral |
| 127 | p.Ala127Lys | 1.37 | 0.45 | Indeterminate | 1.37 | 0.45 | Indeterminate |
| 127 | p.Ala127Thr | 2.09 | 1.06 | Indeterminate | 2.09 | 1.06 | Indeterminate |
| 127 | p.Ala127Arg | 2.10 | 1.07 | Indeterminate | 2.10 | 1.07 | Indeterminate |
| 127 | p.Ala127Ser | 1.18 | 0.24 | Neutral | 1.18 | 0.24 | Neutral |
| 127 | p.Ala127Ile | 1.97 | 0.98 | Indeterminate | 1.97 | 0.98 | Indeterminate |
| 127 | p.Ala127Met | 1.06 | 0.09 | Neutral | 1.06 | 0.09 | Neutral |
| 127 | p.Ala127His | 2.13 | 1.09 | Indeterminate | 2.13 | 1.09 | Indeterminate |
| 127 | p.Ala127Gln | 1.23 | 0.30 | Indeterminate | 1.23 | 0.30 | Indeterminate |
| 127 | p.Ala127Pro | 5.10 | 2.35 | Deleterious | 5.10 | 2.35 | Deleterious |
| 127 | p.Ala127Leu | 1.27 | 0.34 | Indeterminate | 1.27 | 0.34 | Indeterminate |
| 127 | p.Ala127Asp | 1.87 | 0.90 | Indeterminate | 1.87 | 0.90 | Indeterminate |
| 127 | p.Ala127Glu | 1.08 | 0.11 | Neutral | 1.08 | 0.11 | Neutral |
| 127 | p.Ala127Ala | 1.00 | 0.00 | Neutral | 1.00 | 0.00 | Neutral |
| 127 | p.Ala127Gly | 1.48 | 0.56 | Indeterminate | 1.48 | 0.56 | Indeterminate |
| 127 | p.Ala127Val | 0.95 | -0.07 | Neutral | 0.95 | -0.07 | Neutral |
| 127 | p.Ala127Tyr | 1.38 | 0.47 | Indeterminate | 1.38 | 0.47 | Indeterminate |
| 127 | p.Ala127Cys | 0.80 | -0.33 | Neutral | 0.80 | -0.33 | Neutral |
| 127 | p.Ala127Trp | 1.46 | 0.54 | Indeterminate | 1.46 | 0.54 | Indeterminate |
| 127 | p.Ala127Phe | 1.36 | 0.44 | Indeterminate | 1.36 | 0.44 | Indeterminate |
| 128 | p.Arg128Asn | 0.38 | -1.40 | Neutral | 0.38 | -1.40 | Neutral |
| 128 | p.Arg128Lys | 0.98 | -0.03 | Neutral | 0.98 | -0.03 | Neutral |
| 128 | p.Arg128Thr | 0.73 | -0.46 | Neutral | 0.73 | -0.46 | Neutral |
| 128 | p.Arg128Arg | 1.00 | 0.00 | Neutral | 1.00 | 0.00 | Neutral |
| 128 | p.Arg128Ser | 1.43 | 0.51 | Indeterminate | 1.43 | 0.51 | Indeterminate |
| 128 | p.Arg128Ile | 1.28 | 0.35 | Indeterminate | 1.28 | 0.35 | Indeterminate |
| 128 | p.Arg128Met | 0.83 | -0.26 | Neutral | 0.83 | -0.26 | Neutral |
| 128 | p.Arg128His | 0.86 | -0.22 | Neutral | 0.86 | -0.22 | Neutral |
| 128 | p.Arg128Gln | 0.60 | -0.74 | Neutral | 0.60 | -0.74 | Neutral |
| 128 | p.Arg128Pro | 2.66 | 1.41 | Deleterious | 2.66 | 1.41 | Deleterious |
| 128 | p.Arg128Leu | 0.74 | -0.43 | Neutral | 0.74 | -0.43 | Neutral |
| 128 | p.Arg128Asp | 0.65 | -0.61 | Neutral | 0.65 | -0.61 | Neutral |
| 128 | p.Arg128Glu | 0.88 | -0.18 | Neutral | 0.88 | -0.18 | Neutral |
| 128 | p.Arg128Ala | 0.74 | -0.44 | Neutral | 0.74 | -0.44 | Neutral |
| 128 | p.Arg128Gly | 1.19 | 0.26 | Indeterminate | 1.19 | 0.26 | Indeterminate |
| 128 | p.Arg128Val | 0.65 | -0.63 | Neutral | 0.65 | -0.63 | Neutral |
| 128 | p.Arg128Tyr | 0.78 | -0.35 | Neutral | 0.78 | -0.35 | Neutral |
| 128 | p.Arg128Cys | 0.51 | -0.97 | Neutral | 0.51 | -0.97 | Neutral |
| 128 | p.Arg128Trp | 0.67 | -0.58 | Neutral | 0.67 | -0.58 | Neutral |
| 128 | p.Arg128Phe | 1.20 | 0.26 | Indeterminate | 1.20 | 0.26 | Indeterminate |
| 129 | p.Tyr129Asn | 2.40 | 1.26 | Deleterious | 2.40 | 1.26 | Deleterious |
| 129 | p.Tyr129Lys | 1.31 | 0.39 | Indeterminate | 1.31 | 0.39 | Indeterminate |
| 129 | p.Tyr129Thr | 1.99 | 0.99 | Indeterminate | 1.99 | 0.99 | Indeterminate |
| 129 | p.Tyr129Arg | 1.07 | 0.10 | Neutral | 1.07 | 0.10 | Neutral |
| 129 | p.Tyr129Ser | 1.08 | 0.11 | Neutral | 1.08 | 0.11 | Neutral |
| 129 | p.Tyr129Ile | 1.40 | 0.49 | Indeterminate | 1.40 | 0.49 | Indeterminate |
| 129 | p.Tyr129Met | 0.98 | -0.03 | Neutral | 0.98 | -0.03 | Neutral |
| 129 | p.Tyr129His | 1.07 | 0.10 | Neutral | 1.07 | 0.10 | Neutral |
| 129 | p.Tyr129Gln | 1.09 | 0.12 | Neutral | 1.09 | 0.12 | Neutral |
| 129 | p.Tyr129Pro | 8.53 | 3.09 | Deleterious | 8.53 | 3.09 | Deleterious |
| 129 | p.Tyr129Leu | 0.89 | -0.17 | Neutral | 0.89 | -0.17 | Neutral |
| 129 | p.Tyr129Asp | 2.12 | 1.09 | Indeterminate | 2.12 | 1.09 | Indeterminate |
| 129 | p.Tyr129Glu | 1.00 | 0.01 | Neutral | 1.00 | 0.01 | Neutral |
| 129 | p.Tyr129Ala | 1.12 | 0.16 | Neutral | 1.12 | 0.16 | Neutral |
| 129 | p.Tyr129Gly | 1.39 | 0.47 | Indeterminate | 1.39 | 0.47 | Indeterminate |
| 129 | p.Tyr129Val | 1.17 | 0.23 | Neutral | 1.17 | 0.23 | Neutral |
| 129 | p.Tyr129Tyr | 1.00 | 0.00 | Neutral | 1.00 | 0.00 | Neutral |
| 129 | p.Tyr129Cys | 1.07 | 0.10 | Neutral | 1.07 | 0.10 | Neutral |
| 129 | p.Tyr129Trp | 0.89 | -0.16 | Neutral | 0.89 | -0.16 | Neutral |
| 129 | p.Tyr129Phe | 1.17 | 0.23 | Neutral | 1.17 | 0.23 | Neutral |
| 130 | p.Leu130Asn | 1.95 | 0.96 | Indeterminate | 1.95 | 0.96 | Indeterminate |
| 130 | p.Leu130Lys | 3.66 | 1.87 | Deleterious | 3.66 | 1.87 | Deleterious |
| 130 | p.Leu130Thr | 1.07 | 0.10 | Neutral | 1.07 | 0.10 | Neutral |
| 130 | p.Leu130Arg | 3.66 | 1.87 | Deleterious | 3.66 | 1.87 | Deleterious |
| 130 | p.Leu130Ser | 1.22 | 0.29 | Indeterminate | 1.22 | 0.29 | Indeterminate |
| 130 | p.Leu130Ile | 0.91 | -0.13 | Neutral | 0.91 | -0.13 | Neutral |
| 130 | p.Leu130Met | 0.93 | -0.11 | Neutral | 0.93 | -0.11 | Neutral |
| 130 | p.Leu130His | 1.79 | 0.84 | Indeterminate | 1.79 | 0.84 | Indeterminate |
| 130 | p.Leu130Gln | 1.73 | 0.79 | Indeterminate | 1.73 | 0.79 | Indeterminate |
| 130 | p.Leu130Pro | 3.71 | 1.89 | Deleterious | 3.71 | 1.89 | Deleterious |
| 130 | p.Leu130Leu | 1.00 | 0.00 | Neutral | 1.00 | 0.00 | Neutral |
| 130 | p.Leu130Asp | 2.94 | 1.56 | Deleterious | 2.94 | 1.56 | Deleterious |
| 130 | p.Leu130Glu | 2.74 | 1.46 | Deleterious | 2.74 | 1.46 | Deleterious |
| 130 | p.Leu130Ala | 1.05 | 0.07 | Neutral | 1.05 | 0.07 | Neutral |
| 130 | p.Leu130Gly | 1.08 | 0.12 | Neutral | 1.08 | 0.12 | Neutral |
| 130 | p.Leu130Val | 0.87 | -0.20 | Neutral | 0.87 | -0.20 | Neutral |
| 130 | p.Leu130Tyr | 1.08 | 0.12 | Neutral | 1.08 | 0.12 | Neutral |
| 130 | p.Leu130Cys | 1.06 | 0.09 | Neutral | 1.06 | 0.09 | Neutral |
| 130 | p.Leu130Trp | 1.20 | 0.26 | Indeterminate | 1.20 | 0.26 | Indeterminate |
| 130 | p.Leu130Phe | 1.00 | 0.00 | Neutral | 1.00 | 0.00 | Neutral |
| 131 | p.Arg131Asn | 0.97 | -0.05 | Neutral | 0.97 | -0.05 | Neutral |
| 131 | p.Arg131Lys | 0.83 | -0.27 | Neutral | 0.83 | -0.27 | Neutral |
| 131 | p.Arg131Thr | 1.74 | 0.80 | Indeterminate | 1.74 | 0.80 | Indeterminate |
| 131 | p.Arg131Arg | 1.00 | 0.00 | Neutral | 1.00 | 0.00 | Neutral |
| 131 | p.Arg131Ser | 1.12 | 0.16 | Neutral | 1.12 | 0.16 | Neutral |
| 131 | p.Arg131Ile | 0.86 | -0.21 | Neutral | 0.86 | -0.21 | Neutral |
| 131 | p.Arg131Met | 0.99 | -0.01 | Neutral | 0.99 | -0.01 | Neutral |
| 131 | p.Arg131His | 1.13 | 0.18 | Neutral | 1.13 | 0.18 | Neutral |
| 131 | p.Arg131Gln | 2.44 | 1.28 | Deleterious | 2.44 | 1.28 | Deleterious |
| 131 | p.Arg131Pro | 2.12 | 1.08 | Indeterminate | 2.12 | 1.08 | Indeterminate |
| 131 | p.Arg131Leu | 0.76 | -0.40 | Neutral | 0.76 | -0.40 | Neutral |
| 131 | p.Arg131Asp | 0.78 | -0.37 | Neutral | 0.78 | -0.37 | Neutral |
| 131 | p.Arg131Glu | 0.87 | -0.20 | Neutral | 0.87 | -0.20 | Neutral |
| 131 | p.Arg131Ala | 1.15 | 0.21 | Neutral | 1.15 | 0.21 | Neutral |
| 131 | p.Arg131Gly | 1.02 | 0.02 | Neutral | 1.02 | 0.02 | Neutral |
| 131 | p.Arg131Val | 0.90 | -0.16 | Neutral | 0.90 | -0.16 | Neutral |
| 131 | p.Arg131Tyr | 0.94 | -0.10 | Neutral | 0.94 | -0.10 | Neutral |
| 131 | p.Arg131Cys | 3.75 | 1.91 | Deleterious | 3.75 | 1.91 | Deleterious |
| 131 | p.Arg131Trp | 1.21 | 0.28 | Indeterminate | 1.21 | 0.28 | Indeterminate |
| 131 | p.Arg131Phe | 0.69 | -0.53 | Neutral | 0.69 | -0.53 | Neutral |
| 132 | p.Ala132Asn | 2.18 | 1.12 | Deleterious | 2.18 | 1.12 | Deleterious |

| | | | | | | | | | | |
|---|---|---|---|---|---|---|---|---|---|---|
| 132 | p.Ala132Lys | 1.27 | 0.35 | Indeterminate | | | | 1.27 | 0.35 | Indeterminate |
| 132 | p.Ala132Thr | 1.19 | 0.25 | Indeterminate | | | | 1.19 | 0.25 | Indeterminate |
| 132 | p.Ala132Arg | 1.18 | 0.24 | Neutral | | | | 1.18 | 0.24 | Neutral |
| 132 | p.Ala132Ser | 2.27 | 1.18 | Deleterious | | | | 2.27 | 1.18 | Deleterious |
| 132 | p.Ala132Ile | 1.48 | 0.56 | Indeterminate | | | | 1.48 | 0.56 | Indeterminate |
| 132 | p.Ala132Met | 1.34 | 0.42 | Indeterminate | | | | 1.34 | 0.42 | Indeterminate |
| 132 | p.Ala132His | 1.66 | 0.73 | Indeterminate | | | | 1.66 | 0.73 | Indeterminate |
| 132 | p.Ala132Gln | 1.21 | 0.27 | Indeterminate | | | | 1.21 | 0.27 | Indeterminate |
| 132 | p.Ala132Pro | 1.19 | 0.25 | Indeterminate | | | | 1.19 | 0.25 | Indeterminate |
| 132 | p.Ala132Leu | 1.52 | 0.60 | Indeterminate | | | | 1.52 | 0.60 | Indeterminate |
| 132 | p.Ala132Asp | 1.96 | 0.97 | Indeterminate | | | | 1.96 | 0.97 | Indeterminate |
| 132 | p.Ala132Glu | 1.51 | 0.60 | Indeterminate | | | | 1.51 | 0.60 | Indeterminate |
| 132 | p.Ala132Ala | 1.00 | 0.00 | Neutral | | | | 1.00 | 0.00 | Neutral |
| 132 | p.Ala132Gly | 5.74 | 2.52 | Deleterious | | | | 5.74 | 2.52 | Deleterious |
| 132 | p.Ala132Val | 1.35 | 0.44 | Indeterminate | | | | 1.35 | 0.44 | Indeterminate |
| 132 | p.Ala132Tyr | 1.42 | 0.51 | Indeterminate | | | | 1.42 | 0.51 | Indeterminate |
| 132 | p.Ala132Cys | 1.93 | 0.95 | Indeterminate | | | | 1.93 | 0.95 | Indeterminate |
| 132 | p.Ala132Trp | 1.27 | 0.34 | Indeterminate | | | | 1.27 | 0.34 | Indeterminate |
| 132 | p.Ala132Phe | 1.19 | 0.25 | Indeterminate | | | | 1.19 | 0.25 | Indeterminate |
| 133 | p.Ala133Asn | 0.69 | -0.53 | Neutral | | | | 0.69 | -0.53 | Neutral |
| 133 | p.Ala133Lys | 0.77 | -0.38 | Neutral | | | | 0.77 | -0.38 | Neutral |
| 133 | p.Ala133Thr | 0.79 | -0.34 | Neutral | | | | 0.79 | -0.34 | Neutral |
| 133 | p.Ala133Arg | 1.09 | 0.13 | Neutral | | | | 1.09 | 0.13 | Neutral |
| 133 | p.Ala133Ser | 0.90 | -0.15 | Neutral | | | | 0.90 | -0.15 | Neutral |
| 133 | p.Ala133Ile | 0.97 | -0.04 | Neutral | | | | 0.97 | -0.04 | Neutral |
| 133 | p.Ala133Met | 0.76 | -0.40 | Neutral | | | | 0.76 | -0.40 | Neutral |
| 133 | p.Ala133His | 0.69 | -0.53 | Neutral | | | | 0.69 | -0.53 | Neutral |
| 133 | p.Ala133Gln | 0.77 | -0.38 | Neutral | | | | 0.77 | -0.38 | Neutral |
| 133 | p.Ala133Pro | 1.00 | 0.00 | Neutral | | | | 1.00 | 0.00 | Neutral |
| 133 | p.Ala133Leu | 0.91 | -0.14 | Neutral | | | | 0.91 | -0.14 | Neutral |
| 133 | p.Ala133Asp | 1.18 | 0.24 | Indeterminate | | | | 1.18 | 0.24 | Indeterminate |
| 133 | p.Ala133Glu | 1.16 | 0.22 | Neutral | | | | 1.16 | 0.22 | Neutral |
| 133 | p.Ala133Ala | 1.00 | 0.00 | Neutral | | | | 1.00 | 0.00 | Neutral |
| 133 | p.Ala133Gly | 1.05 | 0.07 | Neutral | | | | 1.05 | 0.07 | Neutral |
| 133 | p.Ala133Val | 1.21 | 0.28 | Indeterminate | | | | 1.21 | 0.28 | Indeterminate |
| 133 | p.Ala133Tyr | 1.06 | 0.09 | Neutral | | | | 1.06 | 0.09 | Neutral |
| 133 | p.Ala133Cys | 0.72 | -0.48 | Neutral | | | | 0.72 | -0.48 | Neutral |
| 133 | p.Ala133Trp | 0.89 | -0.17 | Neutral | | | | 0.89 | -0.17 | Neutral |
| 133 | p.Ala133Phe | 0.93 | -0.10 | Neutral | | | | 0.93 | -0.10 | Neutral |
| 134 | p.Ala134Asn | 1.00 | 0.00 | Neutral | 0.79 | -0.34 | Neutral | 0.89 | -0.16 | Neutral |
| 134 | p.Ala134Lys | 1.13 | 0.17 | Neutral | 0.84 | -0.26 | Neutral | 0.98 | -0.03 | Neutral |
| 134 | p.Ala134Thr | 0.97 | -0.04 | Neutral | 1.21 | 0.27 | Indeterminate | 1.09 | 0.12 | Neutral |
| 134 | p.Ala134Arg | 1.17 | 0.22 | Neutral | 0.45 | -1.16 | Neutral | 0.81 | -0.31 | Neutral |
| 134 | p.Ala134Ser | 0.98 | -0.03 | Neutral | 1.23 | 0.30 | Indeterminate | 1.10 | 0.14 | Neutral |
| 134 | p.Ala134Ile | 0.95 | -0.07 | Neutral | 0.98 | -0.02 | Neutral | 0.97 | -0.05 | Neutral |
| 134 | p.Ala134Met | 1.01 | 0.02 | Neutral | 0.87 | -0.20 | Neutral | 0.94 | -0.08 | Neutral |
| 134 | p.Ala134His | 1.01 | 0.01 | Neutral | 0.94 | -0.09 | Neutral | 0.98 | -0.04 | Neutral |
| 134 | p.Ala134Gln | 1.02 | 0.03 | Neutral | 0.95 | -0.07 | Neutral | 0.98 | -0.02 | Neutral |
| 134 | p.Ala134Pro | 0.99 | -0.01 | Neutral | 0.74 | -0.43 | Neutral | 0.87 | -0.21 | Neutral |
| 134 | p.Ala134Leu | 1.00 | -0.01 | Neutral | 0.79 | -0.33 | Neutral | 0.90 | -0.16 | Neutral |
| 134 | p.Ala134Asp | 0.99 | -0.02 | Neutral | 0.97 | -0.05 | Neutral | 0.98 | -0.03 | Neutral |
| 134 | p.Ala134Glu | 1.09 | 0.12 | Neutral | 0.68 | -0.55 | Neutral | 0.88 | -0.18 | Neutral |
| 134 | p.Ala134Ala | 1.00 | 0.00 | Neutral | 1.00 | 0.00 | Neutral | 1.00 | 0.00 | Neutral |
| 134 | p.Ala134Gly | 1.05 | 0.07 | Neutral | 0.85 | -0.23 | Neutral | 0.95 | -0.07 | Neutral |
| 134 | p.Ala134Val | 1.04 | 0.06 | Neutral | 1.06 | 0.08 | Neutral | 1.05 | 0.07 | Neutral |
| 134 | p.Ala134Tyr | 0.98 | -0.03 | Neutral | 1.25 | 0.32 | Indeterminate | 1.11 | 0.15 | Neutral |
| 134 | p.Ala134Cys | 0.96 | -0.06 | Neutral | 1.28 | 0.36 | Indeterminate | 1.12 | 0.16 | Neutral |
| 134 | p.Ala134Trp | 1.26 | 0.33 | Indeterminate | 0.49 | -1.02 | Neutral | 0.88 | -0.19 | Neutral |
| 134 | p.Ala134Phe | 0.96 | -0.06 | Neutral | 0.82 | -0.29 | Neutral | 0.89 | -0.17 | Neutral |
| 135 | p.Gly135Asn | 0.63 | -0.66 | Neutral | | | | 0.63 | -0.66 | Neutral |
| 135 | p.Gly135Lys | 1.02 | 0.03 | Neutral | | | | 1.02 | 0.03 | Neutral |
| 135 | p.Gly135Thr | 0.87 | -0.21 | Neutral | | | | 0.87 | -0.21 | Neutral |
| 135 | p.Gly135Arg | 0.65 | -0.63 | Neutral | | | | 0.65 | -0.63 | Neutral |
| 135 | p.Gly135Ser | 0.88 | -0.19 | Neutral | | | | 0.88 | -0.19 | Neutral |
| 135 | p.Gly135Ile | 1.11 | 0.14 | Neutral | | | | 1.11 | 0.14 | Neutral |
| 135 | p.Gly135Met | 0.97 | -0.04 | Neutral | | | | 0.97 | -0.04 | Neutral |
| 135 | p.Gly135His | 1.01 | 0.01 | Neutral | | | | 1.01 | 0.01 | Neutral |
| 135 | p.Gly135Gln | 0.86 | -0.22 | Neutral | | | | 0.86 | -0.22 | Neutral |
| 135 | p.Gly135Pro | 0.94 | -0.08 | Neutral | | | | 0.94 | -0.08 | Neutral |
| 135 | p.Gly135Leu | 1.05 | 0.06 | Neutral | | | | 1.05 | 0.06 | Neutral |
| 135 | p.Gly135Asp | 0.88 | -0.18 | Neutral | | | | 0.88 | -0.18 | Neutral |
| 135 | p.Gly135Glu | 1.92 | 0.94 | Indeterminate | | | | 1.92 | 0.94 | Indeterminate |
| 135 | p.Gly135Ala | 0.80 | -0.32 | Neutral | | | | 0.80 | -0.32 | Neutral |
| 135 | p.Gly135Gly | 1.00 | 0.00 | Neutral | | | | 1.00 | 0.00 | Neutral |
| 135 | p.Gly135Val | 0.92 | -0.13 | Neutral | | | | 0.92 | -0.13 | Neutral |
| 135 | p.Gly135Tyr | 0.91 | -0.14 | Neutral | | | | 0.91 | -0.14 | Neutral |
| 135 | p.Gly135Cys | 0.74 | -0.43 | Neutral | | | | 0.74 | -0.43 | Neutral |
| 135 | p.Gly135Trp | 0.93 | -0.10 | Neutral | | | | 0.93 | -0.10 | Neutral |
| 135 | p.Gly135Phe | 0.98 | -0.04 | Neutral | | | | 0.98 | -0.04 | Neutral |
| 136 | p.Gly136Asn | 0.89 | -0.16 | Neutral | 1.51 | 0.59 | Indeterminate | 1.20 | 0.26 | Indeterminate |
| 136 | p.Gly136Lys | 0.83 | -0.27 | Neutral | 0.71 | -0.50 | Neutral | 0.77 | -0.38 | Neutral |
| 136 | p.Gly136Thr | 0.89 | -0.17 | Neutral | 0.33 | -1.61 | Neutral | 0.61 | -0.71 | Neutral |
| 136 | p.Gly136Arg | 0.91 | -0.14 | Neutral | 0.71 | -0.50 | Neutral | 0.81 | -0.31 | Neutral |
| 136 | p.Gly136Ser | 0.90 | -0.16 | Neutral | 1.33 | 0.41 | Indeterminate | 1.11 | 0.15 | Neutral |
| 136 | p.Gly136Ile | 0.91 | -0.13 | Neutral | 0.49 | -1.02 | Neutral | 0.70 | -0.51 | Neutral |
| 136 | p.Gly136Met | 0.90 | -0.15 | Neutral | 0.48 | -1.06 | Neutral | 0.69 | -0.53 | Neutral |
| 136 | p.Gly136His | 0.83 | -0.28 | Neutral | 1.52 | 0.60 | Indeterminate | 1.17 | 0.23 | Neutral |
| 136 | p.Gly136Gln | 0.83 | -0.27 | Neutral | 0.90 | -0.14 | Neutral | 0.87 | -0.21 | Neutral |
| 136 | p.Gly136Pro | 0.92 | -0.12 | Neutral | 1.59 | 0.67 | Indeterminate | 1.26 | 0.33 | Indeterminate |
| 136 | p.Gly136Leu | 0.89 | -0.17 | Neutral | 1.38 | 0.47 | Indeterminate | 1.13 | 0.18 | Neutral |
| 136 | p.Gly136Asp | 0.89 | -0.17 | Neutral | 1.95 | 0.96 | Indeterminate | 1.42 | 0.50 | Indeterminate |
| 136 | p.Gly136Glu | 0.84 | -0.25 | Neutral | 0.56 | -0.82 | Neutral | 0.70 | -0.51 | Neutral |
| 136 | p.Gly136Ala | 0.97 | -0.05 | Neutral | 0.33 | -1.59 | Neutral | 0.65 | -0.62 | Neutral |
| 136 | p.Gly136Gly | 1.00 | 0.00 | Neutral | 1.00 | 0.00 | Neutral | 1.00 | 0.00 | Neutral |
| 136 | p.Gly136Val | 0.96 | -0.06 | Neutral | 0.41 | -1.30 | Neutral | 0.68 | -0.55 | Neutral |
| 136 | p.Gly136Tyr | 1.08 | 0.11 | Neutral | 0.74 | -0.44 | Neutral | 0.91 | -0.14 | Neutral |
| 136 | p.Gly136Cys | 0.90 | -0.15 | Neutral | 1.39 | 0.48 | Indeterminate | 1.15 | 0.20 | Neutral |
| 136 | p.Gly136Trp | 0.98 | -0.03 | Neutral | 0.24 | -2.06 | Neutral | 0.61 | -0.71 | Neutral |
| 136 | p.Gly136Phe | 0.86 | -0.22 | Neutral | 1.15 | 0.20 | Indeterminate | 1.00 | 0.01 | Neutral |
| 137 | p.Thr137Asn | 1.02 | 0.03 | Neutral | | | | 1.02 | 0.03 | Neutral |
| 137 | p.Thr137Lys | 0.94 | -0.09 | Neutral | | | | 0.94 | -0.09 | Neutral |

| | | | | | | | | | | |
|---|---|---|---|---|---|---|---|---|---|---|
| 137 | p.Thr137Thr | 1.00 | 0.00 | Neutral | | | | 1.00 | 0.00 | Neutral |
| 137 | p.Thr137Arg | 1.09 | 0.13 | Neutral | | | | 1.09 | 0.13 | Neutral |
| 137 | p.Thr137Ser | 1.05 | 0.08 | Neutral | | | | 1.05 | 0.08 | Neutral |
| 137 | p.Thr137Ile | 1.03 | 0.05 | Neutral | | | | 1.03 | 0.05 | Neutral |
| 137 | p.Thr137Met | 1.10 | 0.14 | Neutral | | | | 1.10 | 0.14 | Neutral |
| 137 | p.Thr137His | 1.03 | 0.04 | Neutral | | | | 1.03 | 0.04 | Neutral |
| 137 | p.Thr137Gln | 0.77 | -0.38 | Neutral | | | | 0.77 | -0.38 | Neutral |
| 137 | p.Thr137Pro | 0.83 | -0.27 | Neutral | | | | 0.83 | -0.27 | Neutral |
| 137 | p.Thr137Leu | 1.02 | 0.03 | Neutral | | | | 1.02 | 0.03 | Neutral |
| 137 | p.Thr137Asp | 0.96 | -0.06 | Neutral | | | | 0.96 | -0.06 | Neutral |
| 137 | p.Thr137Glu | 1.03 | 0.05 | Neutral | | | | 1.03 | 0.05 | Neutral |
| 137 | p.Thr137Ala | 1.16 | 0.22 | Neutral | | | | 1.16 | 0.22 | Neutral |
| 137 | p.Thr137Gly | 1.09 | 0.12 | Neutral | | | | 1.09 | 0.12 | Neutral |
| 137 | p.Thr137Val | 1.12 | 0.17 | Neutral | | | | 1.12 | 0.17 | Neutral |
| 137 | p.Thr137Tyr | 0.86 | -0.21 | Neutral | | | | 0.86 | -0.21 | Neutral |
| 137 | p.Thr137Cys | 1.09 | 0.13 | Neutral | | | | 1.09 | 0.13 | Neutral |
| 137 | p.Thr137Trp | 0.97 | -0.05 | Neutral | | | | 0.97 | -0.05 | Neutral |
| 137 | p.Thr137Phe | 1.00 | 0.00 | Neutral | | | | 1.00 | 0.00 | Neutral |
| 138 | p.Arg138Asn | 1.13 | 0.18 | Neutral | | | | 1.13 | 0.18 | Neutral |
| 138 | p.Arg138Lys | 0.92 | -0.12 | Neutral | | | | 0.92 | -0.12 | Neutral |
| 138 | p.Arg138Thr | 1.05 | 0.07 | Neutral | | | | 1.05 | 0.07 | Neutral |
| 138 | p.Arg138Arg | 1.00 | 0.00 | Neutral | | | | 1.00 | 0.00 | Neutral |
| 138 | p.Arg138Ser | 0.79 | -0.33 | Neutral | | | | 0.79 | -0.33 | Neutral |
| 138 | p.Arg138Ile | 0.90 | -0.16 | Neutral | | | | 0.90 | -0.16 | Neutral |
| 138 | p.Arg138Met | 0.90 | -0.15 | Neutral | | | | 0.90 | -0.15 | Neutral |
| 138 | p.Arg138His | 0.99 | -0.01 | Neutral | | | | 0.99 | -0.01 | Neutral |
| 138 | p.Arg138Gln | 0.93 | -0.10 | Neutral | | | | 0.93 | -0.10 | Neutral |
| 138 | p.Arg138Pro | 1.24 | 0.31 | Indeterminate | | | | 1.24 | 0.31 | Indeterminate |
| 138 | p.Arg138Leu | 0.94 | -0.09 | Neutral | | | | 0.94 | -0.09 | Neutral |
| 138 | p.Arg138Asp | 0.97 | -0.05 | Neutral | | | | 0.97 | -0.05 | Neutral |
| 138 | p.Arg138Glu | 0.94 | -0.08 | Neutral | | | | 0.94 | -0.08 | Neutral |
| 138 | p.Arg138Ala | 0.58 | -0.80 | Neutral | | | | 0.58 | -0.80 | Neutral |
| 138 | p.Arg138Gly | 0.96 | -0.06 | Neutral | | | | 0.96 | -0.06 | Neutral |
| 138 | p.Arg138Val | 0.90 | -0.15 | Neutral | | | | 0.90 | -0.15 | Neutral |
| 138 | p.Arg138Tyr | 0.72 | -0.47 | Neutral | | | | 0.72 | -0.47 | Neutral |
| 138 | p.Arg138Cys | 0.86 | -0.22 | Neutral | | | | 0.86 | -0.22 | Neutral |
| 138 | p.Arg138Trp | 0.81 | -0.31 | Neutral | | | | 0.81 | -0.31 | Neutral |
| 138 | p.Arg138Phe | 0.89 | -0.18 | Neutral | | | | 0.89 | -0.18 | Neutral |
| 139 | p.Gly139Asn | 0.98 | -0.04 | Neutral | 1.01 | 0.02 | Neutral | 1.00 | -0.01 | Neutral |
| 139 | p.Gly139Lys | 1.05 | 0.08 | Neutral | 1.28 | 0.36 | Indeterminate | 1.17 | 0.23 | Neutral |
| 139 | p.Gly139Thr | 0.88 | -0.19 | Neutral | 1.27 | 0.35 | Indeterminate | 1.07 | 0.10 | Neutral |
| 139 | p.Gly139Arg | 0.87 | -0.20 | Neutral | 1.15 | 0.20 | Indeterminate | 1.01 | 0.01 | Neutral |
| 139 | p.Gly139Ser | 1.20 | 0.27 | Indeterminate | 1.26 | 0.33 | Indeterminate | 1.23 | 0.30 | Indeterminate |
| 139 | p.Gly139Ile | 0.99 | -0.02 | Neutral | 1.18 | 0.23 | Indeterminate | 1.08 | 0.11 | Neutral |
| 139 | p.Gly139Met | 1.14 | 0.19 | Neutral | 1.24 | 0.31 | Indeterminate | 1.19 | 0.25 | Indeterminate |
| 139 | p.Gly139His | 0.88 | -0.18 | Neutral | 1.12 | 0.16 | Indeterminate | 1.00 | 0.00 | Neutral |
| 139 | p.Gly139Gln | 0.93 | -0.11 | Neutral | 1.02 | 0.03 | Neutral | 0.98 | -0.04 | Neutral |
| 139 | p.Gly139Pro | 1.30 | 0.38 | Indeterminate | 0.92 | -0.12 | Neutral | 1.11 | 0.15 | Neutral |
| 139 | p.Gly139Leu | 0.85 | -0.24 | Neutral | 1.34 | 0.42 | Indeterminate | 1.09 | 0.13 | Neutral |
| 139 | p.Gly139Asp | 0.89 | -0.16 | Neutral | 1.37 | 0.46 | Indeterminate | 1.13 | 0.18 | Neutral |
| 139 | p.Gly139Glu | 1.12 | 0.16 | Neutral | 1.24 | 0.31 | Indeterminate | 1.18 | 0.24 | Neutral |
| 139 | p.Gly139Ala | 0.80 | -0.32 | Neutral | 1.38 | 0.46 | Indeterminate | 1.09 | 0.12 | Neutral |
| 139 | p.Gly139Gly | 1.00 | 0.00 | Neutral | 1.00 | 0.00 | Neutral | 1.00 | 0.00 | Neutral |
| 139 | p.Gly139Val | 0.89 | -0.17 | Neutral | 1.04 | 0.05 | Neutral | 0.96 | -0.06 | Neutral |
| 139 | p.Gly139Tyr | 1.38 | 0.46 | Indeterminate | 1.35 | 0.43 | Indeterminate | 1.36 | 0.45 | Indeterminate |
| 139 | p.Gly139Cys | 0.86 | -0.23 | Neutral | 1.03 | 0.04 | Neutral | 0.94 | -0.09 | Neutral |
| 139 | p.Gly139Trp | 0.91 | -0.14 | Neutral | 1.18 | 0.23 | Indeterminate | 1.04 | 0.06 | Neutral |
| 139 | p.Gly139Phe | 1.15 | 0.20 | Neutral | 1.30 | 0.37 | Indeterminate | 1.22 | 0.29 | Indeterminate |
| 140 | p.Ser140Asn | 1.02 | 0.03 | Neutral | | | | 1.02 | 0.03 | Neutral |
| 140 | p.Ser140Lys | 0.95 | -0.08 | Neutral | | | | 0.95 | -0.08 | Neutral |
| 140 | p.Ser140Thr | 0.98 | -0.02 | Neutral | | | | 0.98 | -0.02 | Neutral |
| 140 | p.Ser140Arg | 1.18 | 0.23 | Neutral | | | | 1.18 | 0.23 | Neutral |
| 140 | p.Ser140Ser | 1.00 | 0.00 | Neutral | | | | 1.00 | 0.00 | Neutral |
| 140 | p.Ser140Ile | 0.95 | -0.08 | Neutral | | | | 0.95 | -0.08 | Neutral |
| 140 | p.Ser140Met | 0.94 | -0.08 | Neutral | | | | 0.94 | -0.08 | Neutral |
| 140 | p.Ser140His | 1.04 | 0.06 | Neutral | | | | 1.04 | 0.06 | Neutral |
| 140 | p.Ser140Gln | 0.99 | -0.02 | Neutral | | | | 0.99 | -0.02 | Neutral |
| 140 | p.Ser140Pro | 1.20 | 0.27 | Indeterminate | | | | 1.20 | 0.27 | Indeterminate |
| 140 | p.Ser140Leu | 0.90 | -0.15 | Neutral | | | | 0.90 | -0.15 | Neutral |
| 140 | p.Ser140Asp | 0.98 | -0.03 | Neutral | | | | 0.98 | -0.03 | Neutral |
| 140 | p.Ser140Glu | 0.90 | -0.16 | Neutral | | | | 0.90 | -0.16 | Neutral |
| 140 | p.Ser140Ala | 1.07 | 0.09 | Neutral | | | | 1.07 | 0.09 | Neutral |
| 140 | p.Ser140Gly | 1.28 | 0.36 | Indeterminate | | | | 1.28 | 0.36 | Indeterminate |
| 140 | p.Ser140Val | 0.97 | -0.04 | Neutral | | | | 0.97 | -0.04 | Neutral |
| 140 | p.Ser140Tyr | 1.10 | 0.14 | Neutral | | | | 1.10 | 0.14 | Neutral |
| 140 | p.Ser140Cys | 1.03 | 0.05 | Neutral | | | | 1.03 | 0.05 | Neutral |
| 140 | p.Ser140Trp | 0.98 | -0.03 | Neutral | | | | 0.98 | -0.03 | Neutral |
| 140 | p.Ser140Phe | 0.92 | -0.12 | Neutral | | | | 0.92 | -0.12 | Neutral |
| 141 | p.Asn141Asn | 1.00 | 0.00 | Neutral | | | | 1.00 | 0.00 | Neutral |
| 141 | p.Asn141Lys | 0.78 | -0.36 | Neutral | | | | 0.78 | -0.36 | Neutral |
| 141 | p.Asn141Thr | 0.98 | -0.03 | Neutral | | | | 0.98 | -0.03 | Neutral |
| 141 | p.Asn141Arg | 0.97 | -0.04 | Neutral | | | | 0.97 | -0.04 | Neutral |
| 141 | p.Asn141Ser | 0.84 | -0.25 | Neutral | | | | 0.84 | -0.25 | Neutral |
| 141 | p.Asn141Ile | 0.95 | -0.08 | Neutral | | | | 0.95 | -0.08 | Neutral |
| 141 | p.Asn141Met | 0.92 | -0.13 | Neutral | | | | 0.92 | -0.13 | Neutral |
| 141 | p.Asn141His | 0.86 | -0.21 | Neutral | | | | 0.86 | -0.21 | Neutral |
| 141 | p.Asn141Gln | 1.01 | 0.01 | Neutral | | | | 1.01 | 0.01 | Neutral |
| 141 | p.Asn141Pro | 0.87 | -0.21 | Neutral | | | | 0.87 | -0.21 | Neutral |
| 141 | p.Asn141Leu | 0.86 | -0.22 | Neutral | | | | 0.86 | -0.22 | Neutral |
| 141 | p.Asn141Asp | 0.92 | -0.12 | Neutral | | | | 0.92 | -0.12 | Neutral |
| 141 | p.Asn141Glu | 0.97 | -0.05 | Neutral | | | | 0.97 | -0.05 | Neutral |
| 141 | p.Asn141Ala | 0.90 | -0.14 | Neutral | | | | 0.90 | -0.14 | Neutral |
| 141 | p.Asn141Gly | 0.88 | -0.18 | Neutral | | | | 0.88 | -0.18 | Neutral |
| 141 | p.Asn141Val | 0.85 | -0.24 | Neutral | | | | 0.85 | -0.24 | Neutral |
| 141 | p.Asn141Tyr | 0.87 | -0.20 | Neutral | | | | 0.87 | -0.20 | Neutral |
| 141 | p.Asn141Cys | 0.87 | -0.20 | Neutral | | | | 0.87 | -0.20 | Neutral |
| 141 | p.Asn141Trp | 0.87 | -0.20 | Neutral | | | | 0.87 | -0.20 | Neutral |
| 141 | p.Asn141Phe | 0.95 | -0.07 | Neutral | | | | 0.95 | -0.07 | Neutral |
| 142 | p.His142Asn | 0.97 | -0.05 | Neutral | | | | 0.97 | -0.05 | Neutral |
| 142 | p.His142Lys | 1.05 | 0.08 | Neutral | | | | 1.05 | 0.08 | Neutral |
| 142 | p.His142Thr | 1.08 | 0.11 | Neutral | | | | 1.08 | 0.11 | Neutral |

| | | | | | | | |
|---|---|---|---|---|---|---|---|
| 142 | p.His142Arg | 0.81 | -0.30 | Neutral | 0.81 | -0.30 | Neutral |
| 142 | p.His142Ser | 0.96 | -0.06 | Neutral | 0.96 | -0.06 | Neutral |
| 142 | p.His142Ile | 0.96 | -0.06 | Neutral | 0.96 | -0.06 | Neutral |
| 142 | p.His142Met | 0.99 | -0.02 | Neutral | 0.99 | -0.02 | Neutral |
| 142 | p.His142His | 1.00 | 0.00 | Neutral | 1.00 | 0.00 | Neutral |
| 142 | p.His142Gln | 1.09 | 0.12 | Neutral | 1.09 | 0.12 | Neutral |
| 142 | p.His142Pro | 0.95 | -0.07 | Neutral | 0.95 | -0.07 | Neutral |
| 142 | p.His142Leu | 0.91 | -0.14 | Neutral | 0.91 | -0.14 | Neutral |
| 142 | p.His142Asp | 1.09 | 0.12 | Neutral | 1.09 | 0.12 | Neutral |
| 142 | p.His142Glu | 0.97 | -0.04 | Neutral | 0.97 | -0.04 | Neutral |
| 142 | p.His142Ala | 0.96 | -0.05 | Neutral | 0.96 | -0.05 | Neutral |
| 142 | p.His142Gly | 0.94 | -0.09 | Neutral | 0.94 | -0.09 | Neutral |
| 142 | p.His142Val | 1.01 | 0.02 | Neutral | 1.01 | 0.02 | Neutral |
| 142 | p.His142Tyr | 0.90 | -0.15 | Neutral | 0.90 | -0.15 | Neutral |
| 142 | p.His142Cys | 0.96 | -0.05 | Neutral | 0.96 | -0.05 | Neutral |
| 142 | p.His142Trp | 0.91 | -0.13 | Neutral | 0.91 | -0.13 | Neutral |
| 142 | p.His142Phe | 1.12 | 0.17 | Neutral | 1.12 | 0.17 | Neutral |
| 143 | p.Ala143Asn | 1.00 | 0.01 | Neutral | 1.00 | 0.01 | Neutral |
| 143 | p.Ala143Lys | 0.97 | -0.05 | Neutral | 0.97 | -0.05 | Neutral |
| 143 | p.Ala143Thr | 0.96 | -0.06 | Neutral | 0.96 | -0.06 | Neutral |
| 143 | p.Ala143Arg | 0.97 | -0.05 | Neutral | 0.97 | -0.05 | Neutral |
| 143 | p.Ala143Ser | 0.93 | -0.10 | Neutral | 0.93 | -0.10 | Neutral |
| 143 | p.Ala143Ile | 0.92 | -0.12 | Neutral | 0.92 | -0.12 | Neutral |
| 143 | p.Ala143Met | 0.98 | -0.02 | Neutral | 0.98 | -0.02 | Neutral |
| 143 | p.Ala143His | 0.99 | -0.02 | Neutral | 0.99 | -0.02 | Neutral |
| 143 | p.Ala143Gln | 1.00 | 0.00 | Neutral | 1.00 | 0.00 | Neutral |
| 143 | p.Ala143Pro | 1.02 | 0.03 | Neutral | 1.02 | 0.03 | Neutral |
| 143 | p.Ala143Leu | 0.95 | -0.08 | Neutral | 0.95 | -0.08 | Neutral |
| 143 | p.Ala143Asp | 0.91 | -0.13 | Neutral | 0.91 | -0.13 | Neutral |
| 143 | p.Ala143Glu | 0.98 | -0.03 | Neutral | 0.98 | -0.03 | Neutral |
| 143 | p.Ala143Ala | 1.00 | 0.00 | Neutral | 1.00 | 0.00 | Neutral |
| 143 | p.Ala143Gly | 1.03 | 0.04 | Neutral | 1.03 | 0.04 | Neutral |
| 143 | p.Ala143Val | 0.96 | -0.05 | Neutral | 0.96 | -0.05 | Neutral |
| 143 | p.Ala143Tyr | 1.01 | 0.02 | Neutral | 1.01 | 0.02 | Neutral |
| 143 | p.Ala143Cys | 1.04 | 0.06 | Neutral | 1.04 | 0.06 | Neutral |
| 143 | p.Ala143Trp | 1.09 | 0.13 | Neutral | 1.09 | 0.13 | Neutral |
| 143 | p.Ala143Phe | 1.01 | 0.01 | Neutral | 1.01 | 0.01 | Neutral |
| 144 | p.Arg144Asn | 0.88 | -0.18 | Neutral | 0.88 | -0.18 | Neutral |
| 144 | p.Arg144Lys | 0.90 | -0.15 | Neutral | 0.90 | -0.15 | Neutral |
| 144 | p.Arg144Thr | 0.89 | -0.16 | Neutral | 0.89 | -0.16 | Neutral |
| 144 | p.Arg144Arg | 1.00 | 0.00 | Neutral | 1.00 | 0.00 | Neutral |
| 144 | p.Arg144Ser | 0.88 | -0.18 | Neutral | 0.88 | -0.18 | Neutral |
| 144 | p.Arg144Ile | 0.92 | -0.11 | Neutral | 0.92 | -0.11 | Neutral |
| 144 | p.Arg144Met | 0.89 | -0.17 | Neutral | 0.89 | -0.17 | Neutral |
| 144 | p.Arg144His | 0.92 | -0.13 | Neutral | 0.92 | -0.13 | Neutral |
| 144 | p.Arg144Gln | 0.95 | -0.07 | Neutral | 0.95 | -0.07 | Neutral |
| 144 | p.Arg144Pro | 0.90 | -0.15 | Neutral | 0.90 | -0.15 | Neutral |
| 144 | p.Arg144Leu | 0.96 | -0.06 | Neutral | 0.96 | -0.06 | Neutral |
| 144 | p.Arg144Asp | 0.89 | -0.16 | Neutral | 0.89 | -0.16 | Neutral |
| 144 | p.Arg144Glu | 0.96 | -0.07 | Neutral | 0.96 | -0.07 | Neutral |
| 144 | p.Arg144Ala | 0.95 | -0.08 | Neutral | 0.95 | -0.08 | Neutral |
| 144 | p.Arg144Gly | 0.97 | -0.05 | Neutral | 0.97 | -0.05 | Neutral |
| 144 | p.Arg144Val | 0.91 | -0.14 | Neutral | 0.91 | -0.14 | Neutral |
| 144 | p.Arg144Tyr | 0.91 | -0.14 | Neutral | 0.91 | -0.14 | Neutral |
| 144 | p.Arg144Cys | 0.84 | -0.25 | Neutral | 0.84 | -0.25 | Neutral |
| 144 | p.Arg144Trp | 0.91 | -0.13 | Neutral | 0.91 | -0.13 | Neutral |
| 144 | p.Arg144Phe | 0.90 | -0.15 | Neutral | 0.90 | -0.15 | Neutral |
| 145 | p.Ile145Asn | 0.99 | -0.02 | Neutral | 0.99 | -0.02 | Neutral |
| 145 | p.Ile145Lys | 1.13 | 0.18 | Neutral | 1.13 | 0.18 | Neutral |
| 145 | p.Ile145Thr | 1.05 | 0.07 | Neutral | 1.05 | 0.07 | Neutral |
| 145 | p.Ile145Arg | 1.18 | 0.24 | Indeterminate | 1.18 | 0.24 | Indeterminate |
| 145 | p.Ile145Ser | 1.11 | 0.15 | Neutral | 1.11 | 0.15 | Neutral |
| 145 | p.Ile145Ile | 1.00 | 0.00 | Neutral | 1.00 | 0.00 | Neutral |
| 145 | p.Ile145Met | 1.21 | 0.28 | Indeterminate | 1.21 | 0.28 | Indeterminate |
| 145 | p.Ile145His | 0.94 | -0.08 | Neutral | 0.94 | -0.08 | Neutral |
| 145 | p.Ile145Gln | 1.10 | 0.14 | Neutral | 1.10 | 0.14 | Neutral |
| 145 | p.Ile145Pro | 0.97 | -0.04 | Neutral | 0.97 | -0.04 | Neutral |
| 145 | p.Ile145Leu | 1.04 | 0.05 | Neutral | 1.04 | 0.05 | Neutral |
| 145 | p.Ile145Asp | 1.04 | 0.05 | Neutral | 1.04 | 0.05 | Neutral |
| 145 | p.Ile145Glu | 1.05 | 0.06 | Neutral | 1.05 | 0.06 | Neutral |
| 145 | p.Ile145Ala | 1.15 | 0.20 | Neutral | 1.15 | 0.20 | Neutral |
| 145 | p.Ile145Gly | 0.98 | -0.03 | Neutral | 0.98 | -0.03 | Neutral |
| 145 | p.Ile145Val | 1.13 | 0.17 | Neutral | 1.13 | 0.17 | Neutral |
| 145 | p.Ile145Tyr | 1.02 | 0.03 | Neutral | 1.02 | 0.03 | Neutral |
| 145 | p.Ile145Cys | 0.92 | -0.12 | Neutral | 0.92 | -0.12 | Neutral |
| 145 | p.Ile145Trp | 1.04 | 0.05 | Neutral | 1.04 | 0.05 | Neutral |
| 145 | p.Ile145Phe | 0.98 | -0.03 | Neutral | 0.98 | -0.03 | Neutral |
| 146 | p.Asp146Asn | 1.05 | 0.07 | Neutral | 1.05 | 0.07 | Neutral |
| 146 | p.Asp146Lys | 0.78 | -0.36 | Neutral | 0.78 | -0.36 | Neutral |
| 146 | p.Asp146Thr | 0.95 | -0.07 | Neutral | 0.95 | -0.07 | Neutral |
| 146 | p.Asp146Arg | 0.95 | -0.07 | Neutral | 0.95 | -0.07 | Neutral |
| 146 | p.Asp146Ser | 0.88 | -0.18 | Neutral | 0.88 | -0.18 | Neutral |
| 146 | p.Asp146Ile | 0.79 | -0.33 | Neutral | 0.79 | -0.33 | Neutral |
| 146 | p.Asp146Met | 0.96 | -0.05 | Neutral | 0.96 | -0.05 | Neutral |
| 146 | p.Asp146His | 1.01 | 0.01 | Neutral | 1.01 | 0.01 | Neutral |
| 146 | p.Asp146Gln | 0.98 | -0.03 | Neutral | 0.98 | -0.03 | Neutral |
| 146 | p.Asp146Pro | 1.00 | 0.00 | Neutral | 1.00 | 0.00 | Neutral |
| 146 | p.Asp146Leu | 0.93 | -0.11 | Neutral | 0.93 | -0.11 | Neutral |
| 146 | p.Asp146Asp | 1.00 | 0.00 | Neutral | 1.00 | 0.00 | Neutral |
| 146 | p.Asp146Glu | 0.87 | -0.20 | Neutral | 0.87 | -0.20 | Neutral |
| 146 | p.Asp146Ala | 0.86 | -0.22 | Neutral | 0.86 | -0.22 | Neutral |
| 146 | p.Asp146Gly | 0.86 | -0.22 | Neutral | 0.86 | -0.22 | Neutral |
| 146 | p.Asp146Val | 0.86 | -0.21 | Neutral | 0.86 | -0.21 | Neutral |
| 146 | p.Asp146Tyr | 1.00 | 0.00 | Neutral | 1.00 | 0.00 | Neutral |
| 146 | p.Asp146Cys | 1.01 | 0.02 | Neutral | 1.01 | 0.02 | Neutral |
| 146 | p.Asp146Trp | 0.85 | -0.23 | Neutral | 0.85 | -0.23 | Neutral |
| 146 | p.Asp146Phe | 1.03 | 0.05 | Neutral | 1.03 | 0.05 | Neutral |
| 147 | p.Ala147Asn | 1.05 | 0.07 | Neutral | 1.05 | 0.07 | Neutral |
| 147 | p.Ala147Lys | 0.91 | -0.13 | Neutral | 0.91 | -0.13 | Neutral |
| 147 | p.Ala147Thr | 1.03 | 0.04 | Neutral | 1.03 | 0.04 | Neutral |
| 147 | p.Ala147Arg | 0.91 | -0.14 | Neutral | 0.91 | -0.14 | Neutral |

| | | | | | | | |
|---|---|---|---|---|---|---|---|
| 147 | p.Ala147Ser | 0.91 | -0.14 | Neutral | 0.91 | -0.14 | Neutral |
| 147 | p.Ala147Ile | 1.12 | 0.16 | Neutral | 1.12 | 0.16 | Neutral |
| 147 | p.Ala147Met | 1.08 | 0.11 | Neutral | 1.08 | 0.11 | Neutral |
| 147 | p.Ala147His | 0.89 | -0.17 | Neutral | 0.89 | -0.17 | Neutral |
| 147 | p.Ala147Gln | 1.42 | 0.50 | Indeterminate | 1.42 | 0.50 | Indeterminate |
| 147 | p.Ala147Pro | 0.88 | -0.18 | Neutral | 0.88 | -0.18 | Neutral |
| 147 | p.Ala147Leu | 1.07 | 0.09 | Neutral | 1.07 | 0.09 | Neutral |
| 147 | p.Ala147Asp | 1.12 | 0.17 | Neutral | 1.12 | 0.17 | Neutral |
| 147 | p.Ala147Glu | 0.94 | -0.09 | Neutral | 0.94 | -0.09 | Neutral |
| 147 | p.Ala147Ala | 1.00 | 0.00 | Neutral | 1.00 | 0.00 | Neutral |
| 147 | p.Ala147Gly | 0.98 | -0.03 | Neutral | 0.98 | -0.03 | Neutral |
| 147 | p.Ala147Val | 1.04 | 0.06 | Neutral | 1.04 | 0.06 | Neutral |
| 147 | p.Ala147Tyr | 0.89 | -0.16 | Neutral | 0.89 | -0.16 | Neutral |
| 147 | p.Ala147Cys | 0.92 | -0.12 | Neutral | 0.92 | -0.12 | Neutral |
| 147 | p.Ala147Trp | 0.93 | -0.10 | Neutral | 0.93 | -0.10 | Neutral |
| 147 | p.Ala147Phe | 0.89 | -0.16 | Neutral | 0.89 | -0.16 | Neutral |
| 148 | p.Ala148Asn | 1.32 | 0.40 | Indeterminate | 1.32 | 0.40 | Indeterminate |
| 148 | p.Ala148Lys | 2.00 | 1.00 | Indeterminate | 2.00 | 1.00 | Indeterminate |
| 148 | p.Ala148Thr | 0.92 | -0.12 | Neutral | 0.92 | -0.12 | Neutral |
| 148 | p.Ala148Arg | 1.44 | 0.53 | Indeterminate | 1.44 | 0.53 | Indeterminate |
| 148 | p.Ala148Ser | 1.08 | 0.11 | Neutral | 1.08 | 0.11 | Neutral |
| 148 | p.Ala148Ile | 1.15 | 0.20 | Neutral | 1.15 | 0.20 | Neutral |
| 148 | p.Ala148Met | 1.54 | 0.62 | Indeterminate | 1.54 | 0.62 | Indeterminate |
| 148 | p.Ala148His | 1.27 | 0.34 | Indeterminate | 1.27 | 0.34 | Indeterminate |
| 148 | p.Ala148Gln | 1.31 | 0.39 | Indeterminate | 1.31 | 0.39 | Indeterminate |
| 148 | p.Ala148Pro | 1.19 | 0.25 | Indeterminate | 1.19 | 0.25 | Indeterminate |
| 148 | p.Ala148Leu | 1.21 | 0.27 | Indeterminate | 1.21 | 0.27 | Indeterminate |
| 148 | p.Ala148Asp | 1.27 | 0.35 | Indeterminate | 1.27 | 0.35 | Indeterminate |
| 148 | p.Ala148Glu | 1.39 | 0.47 | Indeterminate | 1.39 | 0.47 | Indeterminate |
| 148 | p.Ala148Ala | 1.00 | 0.00 | Neutral | 1.00 | 0.00 | Neutral |
| 148 | p.Ala148Gly | 1.01 | 0.01 | Neutral | 1.01 | 0.01 | Neutral |
| 148 | p.Ala148Val | 1.55 | 0.63 | Indeterminate | 1.55 | 0.63 | Indeterminate |
| 148 | p.Ala148Tyr | 0.94 | -0.09 | Neutral | 0.94 | -0.09 | Neutral |
| 148 | p.Ala148Cys | 0.66 | -0.60 | Neutral | 0.66 | -0.60 | Neutral |
| 148 | p.Ala148Trp | 1.28 | 0.35 | Indeterminate | 1.28 | 0.35 | Indeterminate |
| 148 | p.Ala148Phe | 1.94 | 0.96 | Indeterminate | 1.94 | 0.96 | Indeterminate |
| 149 | p.Glu149Asn | 0.80 | -0.31 | Neutral | 0.80 | -0.31 | Neutral |
| 149 | p.Glu149Lys | 1.03 | 0.04 | Neutral | 1.03 | 0.04 | Neutral |
| 149 | p.Glu149Thr | 1.08 | 0.12 | Neutral | 1.08 | 0.12 | Neutral |
| 149 | p.Glu149Arg | 0.85 | -0.24 | Neutral | 0.85 | -0.24 | Neutral |
| 149 | p.Glu149Ser | 1.14 | 0.19 | Neutral | 1.14 | 0.19 | Neutral |
| 149 | p.Glu149Ile | 1.10 | 0.14 | Neutral | 1.10 | 0.14 | Neutral |
| 149 | p.Glu149Met | 0.79 | -0.34 | Neutral | 0.79 | -0.34 | Neutral |
| 149 | p.Glu149His | 0.93 | -0.11 | Neutral | 0.93 | -0.11 | Neutral |
| 149 | p.Glu149Gln | 0.88 | -0.19 | Neutral | 0.88 | -0.19 | Neutral |
| 149 | p.Glu149Pro | 0.91 | -0.14 | Neutral | 0.91 | -0.14 | Neutral |
| 149 | p.Glu149Leu | 0.92 | -0.13 | Neutral | 0.92 | -0.13 | Neutral |
| 149 | p.Glu149Asp | 0.86 | -0.22 | Neutral | 0.86 | -0.22 | Neutral |
| 149 | p.Glu149Glu | 1.00 | 0.00 | Neutral | 1.00 | 0.00 | Neutral |
| 149 | p.Glu149Ala | 1.00 | 0.00 | Neutral | 1.00 | 0.00 | Neutral |
| 149 | p.Glu149Gly | 0.98 | -0.02 | Neutral | 0.98 | -0.02 | Neutral |
| 149 | p.Glu149Val | 0.99 | -0.01 | Neutral | 0.99 | -0.01 | Neutral |
| 149 | p.Glu149Tyr | 1.02 | 0.03 | Neutral | 1.02 | 0.03 | Neutral |
| 149 | p.Glu149Cys | 0.86 | -0.22 | Neutral | 0.86 | -0.22 | Neutral |
| 149 | p.Glu149Trp | 1.00 | 0.00 | Neutral | 1.00 | 0.00 | Neutral |
| 149 | p.Glu149Phe | 1.06 | 0.09 | Neutral | 1.06 | 0.09 | Neutral |
| 150 | p.Gly150Asn | 0.89 | -0.17 | Neutral | 0.89 | -0.17 | Neutral |
| 150 | p.Gly150Lys | 1.03 | 0.04 | Neutral | 1.03 | 0.04 | Neutral |
| 150 | p.Gly150Thr | 0.96 | -0.06 | Neutral | 0.96 | -0.06 | Neutral |
| 150 | p.Gly150Arg | 1.03 | 0.04 | Neutral | 1.03 | 0.04 | Neutral |
| 150 | p.Gly150Ser | 0.99 | -0.01 | Neutral | 0.99 | -0.01 | Neutral |
| 150 | p.Gly150Ile | 0.92 | -0.12 | Neutral | 0.92 | -0.12 | Neutral |
| 150 | p.Gly150Met | 1.07 | 0.10 | Neutral | 1.07 | 0.10 | Neutral |
| 150 | p.Gly150His | 1.04 | 0.05 | Neutral | 1.04 | 0.05 | Neutral |
| 150 | p.Gly150Gln | 0.93 | -0.10 | Neutral | 0.93 | -0.10 | Neutral |
| 150 | p.Gly150Pro | 1.00 | -0.01 | Neutral | 1.00 | -0.01 | Neutral |
| 150 | p.Gly150Leu | 0.91 | -0.14 | Neutral | 0.91 | -0.14 | Neutral |
| 150 | p.Gly150Asp | 0.97 | -0.05 | Neutral | 0.97 | -0.05 | Neutral |
| 150 | p.Gly150Glu | 0.88 | -0.18 | Neutral | 0.88 | -0.18 | Neutral |
| 150 | p.Gly150Ala | 0.86 | -0.21 | Neutral | 0.86 | -0.21 | Neutral |
| 150 | p.Gly150Gly | 1.00 | 0.00 | Neutral | 1.00 | 0.00 | Neutral |
| 150 | p.Gly150Val | 0.97 | -0.05 | Neutral | 0.97 | -0.05 | Neutral |
| 150 | p.Gly150Tyr | 0.91 | -0.13 | Neutral | 0.91 | -0.13 | Neutral |
| 150 | p.Gly150Cys | 0.77 | -0.38 | Neutral | 0.77 | -0.38 | Neutral |
| 150 | p.Gly150Trp | 0.92 | -0.12 | Neutral | 0.92 | -0.12 | Neutral |
| 150 | p.Gly150Phe | 0.99 | -0.01 | Neutral | 0.99 | -0.01 | Neutral |
| 151 | p.Pro151Asn | 1.03 | 0.04 | Neutral | 1.03 | 0.04 | Neutral |
| 151 | p.Pro151Lys | 0.94 | -0.09 | Neutral | 0.94 | -0.09 | Neutral |
| 151 | p.Pro151Thr | 1.03 | 0.05 | Neutral | 1.03 | 0.05 | Neutral |
| 151 | p.Pro151Arg | 0.84 | -0.26 | Neutral | 0.84 | -0.26 | Neutral |
| 151 | p.Pro151Ser | 1.03 | 0.04 | Neutral | 1.03 | 0.04 | Neutral |
| 151 | p.Pro151Ile | 0.99 | -0.01 | Neutral | 0.99 | -0.01 | Neutral |
| 151 | p.Pro151Met | 0.92 | -0.12 | Neutral | 0.92 | -0.12 | Neutral |
| 151 | p.Pro151His | 1.06 | 0.09 | Neutral | 1.06 | 0.09 | Neutral |
| 151 | p.Pro151Gln | 0.90 | -0.15 | Neutral | 0.90 | -0.15 | Neutral |
| 151 | p.Pro151Pro | 1.00 | 0.00 | Neutral | 1.00 | 0.00 | Neutral |
| 151 | p.Pro151Leu | 1.01 | 0.02 | Neutral | 1.01 | 0.02 | Neutral |
| 151 | p.Pro151Asp | 1.02 | 0.03 | Neutral | 1.02 | 0.03 | Neutral |
| 151 | p.Pro151Glu | 1.06 | 0.08 | Neutral | 1.06 | 0.08 | Neutral |
| 151 | p.Pro151Ala | 1.09 | 0.12 | Neutral | 1.09 | 0.12 | Neutral |
| 151 | p.Pro151Gly | 1.13 | 0.17 | Neutral | 1.13 | 0.17 | Neutral |
| 151 | p.Pro151Val | 0.95 | -0.08 | Neutral | 0.95 | -0.08 | Neutral |
| 151 | p.Pro151Tyr | 0.88 | -0.18 | Neutral | 0.88 | -0.18 | Neutral |
| 151 | p.Pro151Cys | 0.89 | -0.17 | Neutral | 0.89 | -0.17 | Neutral |
| 151 | p.Pro151Trp | 0.88 | -0.18 | Neutral | 0.88 | -0.18 | Neutral |
| 151 | p.Pro151Phe | 0.92 | -0.12 | Neutral | 0.92 | -0.12 | Neutral |
| 152 | p.Ser152Asn | 0.87 | -0.20 | Neutral | 0.87 | -0.20 | Neutral |
| 152 | p.Ser152Lys | 0.96 | -0.06 | Neutral | 0.96 | -0.06 | Neutral |
| 152 | p.Ser152Thr | 0.96 | -0.06 | Neutral | 0.96 | -0.06 | Neutral |
| 152 | p.Ser152Arg | 0.89 | -0.17 | Neutral | 0.89 | -0.17 | Neutral |
| 152 | p.Ser152Ser | 1.00 | 0.00 | Neutral | 1.00 | 0.00 | Neutral |

| | | | | | | | | | | |
|---|---|---|---|---|---|---|---|---|---|---|
| 152 | p.Ser152Ile | 0.76 | -0.40 | Neutral | | | | 0.76 | -0.40 | Neutral |
| 152 | p.Ser152Met | 0.97 | -0.05 | Neutral | | | | 0.97 | -0.05 | Neutral |
| 152 | p.Ser152His | 0.89 | -0.17 | Neutral | | | | 0.89 | -0.17 | Neutral |
| 152 | p.Ser152Gln | 0.79 | -0.33 | Neutral | | | | 0.79 | -0.33 | Neutral |
| 152 | p.Ser152Pro | 0.98 | -0.03 | Neutral | | | | 0.98 | -0.03 | Neutral |
| 152 | p.Ser152Leu | 0.96 | -0.06 | Neutral | | | | 0.96 | -0.06 | Neutral |
| 152 | p.Ser152Asp | 0.89 | -0.17 | Neutral | | | | 0.89 | -0.17 | Neutral |
| 152 | p.Ser152Glu | 0.91 | -0.13 | Neutral | | | | 0.91 | -0.13 | Neutral |
| 152 | p.Ser152Ala | 1.04 | 0.06 | Neutral | | | | 1.04 | 0.06 | Neutral |
| 152 | p.Ser152Gly | 0.97 | -0.04 | Neutral | | | | 0.97 | -0.04 | Neutral |
| 152 | p.Ser152Val | 0.98 | -0.03 | Neutral | | | | 0.98 | -0.03 | Neutral |
| 152 | p.Ser152Tyr | 1.10 | 0.14 | Neutral | | | | 1.10 | 0.14 | Neutral |
| 152 | p.Ser152Cys | 0.75 | -0.42 | Neutral | | | | 0.75 | -0.42 | Neutral |
| 152 | p.Ser152Trp | 0.89 | -0.17 | Neutral | | | | 0.89 | -0.17 | Neutral |
| 152 | p.Ser152Phe | 0.88 | -0.19 | Neutral | | | | 0.88 | -0.19 | Neutral |
| 153 | p.Asp153Asn | 1.33 | 0.41 | Indeterminate | 0.83 | -0.28 | Neutral | 1.08 | 0.11 | Neutral |
| 153 | p.Asp153Lys | 0.99 | -0.02 | Neutral | 1.33 | 0.41 | Indeterminate | 1.16 | 0.21 | Neutral |
| 153 | p.Asp153Thr | 1.47 | 0.56 | Indeterminate | 1.10 | 0.14 | Neutral | 1.29 | 0.36 | Indeterminate |
| 153 | p.Asp153Arg | 1.25 | 0.32 | Indeterminate | 1.37 | 0.46 | Indeterminate | 1.31 | 0.39 | Indeterminate |
| 153 | p.Asp153Ser | 0.82 | -0.29 | Neutral | 0.72 | -0.47 | Neutral | 0.77 | -0.37 | Neutral |
| 153 | p.Asp153Ile | 0.91 | -0.14 | Neutral | 1.16 | 0.22 | Indeterminate | 1.04 | 0.05 | Neutral |
| 153 | p.Asp153Met | 1.31 | 0.39 | Indeterminate | 0.92 | -0.12 | Neutral | 1.11 | 0.16 | Neutral |
| 153 | p.Asp153His | 1.34 | 0.42 | Indeterminate | 0.92 | -0.12 | Neutral | 1.13 | 0.18 | Neutral |
| 153 | p.Asp153Gln | 1.39 | 0.48 | Indeterminate | 0.95 | -0.07 | Neutral | 1.17 | 0.23 | Neutral |
| 153 | p.Asp153Pro | 0.68 | -0.56 | Neutral | 0.53 | -0.93 | Neutral | 0.60 | -0.73 | Neutral |
| 153 | p.Asp153Leu | 1.34 | 0.43 | Indeterminate | 0.94 | -0.09 | Neutral | 1.14 | 0.19 | Neutral |
| 153 | p.Asp153Asp | 1.00 | 0.00 | Neutral | 1.00 | 0.00 | Neutral | 1.00 | 0.00 | Neutral |
| 153 | p.Asp153Glu | 1.18 | 0.24 | Indeterminate | 0.82 | -0.29 | Neutral | 1.00 | 0.00 | Neutral |
| 153 | p.Asp153Ala | 0.67 | -0.57 | Neutral | 0.98 | -0.03 | Neutral | 0.83 | -0.27 | Neutral |
| 153 | p.Asp153Gly | 0.72 | -0.47 | Neutral | 0.64 | -0.64 | Neutral | 0.68 | -0.55 | Neutral |
| 153 | p.Asp153Val | 1.14 | 0.19 | Neutral | 1.20 | 0.26 | Indeterminate | 1.17 | 0.23 | Neutral |
| 153 | p.Asp153Tyr | 0.97 | -0.04 | Neutral | 1.33 | 0.41 | Indeterminate | 1.15 | 0.20 | Neutral |
| 153 | p.Asp153Cys | 1.41 | 0.50 | Indeterminate | 1.09 | 0.13 | Neutral | 1.25 | 0.32 | Indeterminate |
| 153 | p.Asp153Trp | 1.25 | 0.32 | Indeterminate | 1.23 | 0.30 | Indeterminate | 1.24 | 0.31 | Indeterminate |
| 153 | p.Asp153Phe | 1.04 | 0.05 | Neutral | 1.07 | 0.09 | Neutral | 1.05 | 0.07 | Neutral |
| 154 | p.Ile154Asn | 1.10 | 0.13 | Neutral | | | | 1.10 | 0.13 | Neutral |
| 154 | p.Ile154Lys | 0.91 | -0.13 | Neutral | | | | 0.91 | -0.13 | Neutral |
| 154 | p.Ile154Thr | 1.09 | 0.13 | Neutral | | | | 1.09 | 0.13 | Neutral |
| 154 | p.Ile154Arg | 1.04 | 0.06 | Neutral | | | | 1.04 | 0.06 | Neutral |
| 154 | p.Ile154Ser | 0.86 | -0.21 | Neutral | | | | 0.86 | -0.21 | Neutral |
| 154 | p.Ile154Ile | 1.00 | 0.00 | Neutral | | | | 1.00 | 0.00 | Neutral |
| 154 | p.Ile154Met | 1.03 | 0.05 | Neutral | | | | 1.03 | 0.05 | Neutral |
| 154 | p.Ile154His | 1.13 | 0.17 | Neutral | | | | 1.13 | 0.17 | Neutral |
| 154 | p.Ile154Gln | 1.00 | 0.00 | Neutral | | | | 1.00 | 0.00 | Neutral |
| 154 | p.Ile154Pro | 1.13 | 0.18 | Neutral | | | | 1.13 | 0.18 | Neutral |
| 154 | p.Ile154Leu | 0.93 | -0.11 | Neutral | | | | 0.93 | -0.11 | Neutral |
| 154 | p.Ile154Asp | 0.99 | -0.01 | Neutral | | | | 0.99 | -0.01 | Neutral |
| 154 | p.Ile154Glu | 1.06 | 0.09 | Neutral | | | | 1.06 | 0.09 | Neutral |
| 154 | p.Ile154Ala | 1.27 | 0.34 | Indeterminate | | | | 1.27 | 0.34 | Indeterminate |
| 154 | p.Ile154Gly | 0.87 | -0.20 | Neutral | | | | 0.87 | -0.20 | Neutral |
| 154 | p.Ile154Val | 1.20 | 0.26 | Indeterminate | | | | 1.20 | 0.26 | Indeterminate |
| 154 | p.Ile154Tyr | 1.15 | 0.20 | Neutral | | | | 1.15 | 0.20 | Neutral |
| 154 | p.Ile154Cys | 1.13 | 0.18 | Neutral | | | | 1.13 | 0.18 | Neutral |
| 154 | p.Ile154Trp | 1.23 | 0.30 | Indeterminate | | | | 1.23 | 0.30 | Indeterminate |
| 154 | p.Ile154Phe | 1.06 | 0.08 | Neutral | | | | 1.06 | 0.08 | Neutral |
| 155 | p.Pro155Asn | 0.98 | -0.02 | Neutral | | | | 0.98 | -0.02 | Neutral |
| 155 | p.Pro155Lys | 1.19 | 0.25 | Indeterminate | | | | 1.19 | 0.25 | Indeterminate |
| 155 | p.Pro155Thr | 0.18 | -2.43 | Neutral | | | | 0.18 | -2.43 | Neutral |
| 155 | p.Pro155Arg | 0.90 | -0.15 | Neutral | | | | 0.90 | -0.15 | Neutral |
| 155 | p.Pro155Ser | 2.13 | 1.09 | Deleterious | | | | 2.13 | 1.09 | Deleterious |
| 155 | p.Pro155Ile | 1.37 | 0.45 | Indeterminate | | | | 1.37 | 0.45 | Indeterminate |
| 155 | p.Pro155Met | 0.58 | -0.78 | Neutral | | | | 0.58 | -0.78 | Neutral |
| 155 | p.Pro155His | 1.63 | 0.71 | Indeterminate | | | | 1.63 | 0.71 | Indeterminate |
| 155 | p.Pro155Gln | 0.46 | -1.11 | Neutral | | | | 0.46 | -1.11 | Neutral |
| 155 | p.Pro155Pro | 1.00 | 0.00 | Neutral | | | | 1.00 | 0.00 | Neutral |
| 155 | p.Pro155Leu | 0.50 | -1.00 | Neutral | | | | 0.50 | -1.00 | Neutral |
| 155 | p.Pro155Asp | 2.39 | 1.26 | Deleterious | | | | 2.39 | 1.26 | Deleterious |
| 155 | p.Pro155Glu | 0.55 | -0.87 | Neutral | | | | 0.55 | -0.87 | Neutral |
| 155 | p.Pro155Ala | 0.18 | -2.45 | Neutral | | | | 0.18 | -2.45 | Neutral |
| 155 | p.Pro155Gly | 0.71 | -0.49 | Neutral | | | | 0.71 | -0.49 | Neutral |
| 155 | p.Pro155Val | 1.99 | 0.99 | Indeterminate | | | | 1.99 | 0.99 | Indeterminate |
| 155 | p.Pro155Tyr | 0.70 | -0.51 | Neutral | | | | 0.70 | -0.51 | Neutral |
| 155 | p.Pro155Cys | 0.39 | -1.35 | Neutral | | | | 0.39 | -1.35 | Neutral |
| 155 | p.Pro155Trp | 1.59 | 0.67 | Indeterminate | | | | 1.59 | 0.67 | Indeterminate |
| 155 | p.Pro155Phe | 0.19 | -2.36 | Neutral | | | | 0.19 | -2.36 | Neutral |
| 156 | p.Asp156Asn | 1.05 | 0.06 | Neutral | | | | 1.05 | 0.06 | Neutral |
| 156 | p.Asp156Lys | 0.99 | -0.01 | Neutral | | | | 0.99 | -0.01 | Neutral |
| 156 | p.Asp156Thr | 1.07 | 0.10 | Neutral | | | | 1.07 | 0.10 | Neutral |
| 156 | p.Asp156Arg | 1.01 | 0.01 | Neutral | | | | 1.01 | 0.01 | Neutral |
| 156 | p.Asp156Ser | 1.01 | 0.02 | Neutral | | | | 1.01 | 0.02 | Neutral |
| 156 | p.Asp156Ile | 0.97 | -0.04 | Neutral | | | | 0.97 | -0.04 | Neutral |
| 156 | p.Asp156Met | 0.99 | -0.01 | Neutral | | | | 0.99 | -0.01 | Neutral |
| 156 | p.Asp156His | 0.98 | -0.04 | Neutral | | | | 0.98 | -0.04 | Neutral |
| 156 | p.Asp156Gln | 1.19 | 0.25 | Indeterminate | | | | 1.19 | 0.25 | Indeterminate |
| 156 | p.Asp156Pro | 0.99 | -0.02 | Neutral | | | | 0.99 | -0.02 | Neutral |
| 156 | p.Asp156Leu | 1.09 | 0.12 | Neutral | | | | 1.09 | 0.12 | Neutral |
| 156 | p.Asp156Asp | 1.00 | 0.00 | Neutral | | | | 1.00 | 0.00 | Neutral |
| 156 | p.Asp156Glu | 1.05 | 0.07 | Neutral | | | | 1.05 | 0.07 | Neutral |
| 156 | p.Asp156Ala | 1.27 | 0.34 | Indeterminate | | | | 1.27 | 0.34 | Indeterminate |
| 156 | p.Asp156Gly | 1.10 | 0.14 | Neutral | | | | 1.10 | 0.14 | Neutral |
| 156 | p.Asp156Val | 1.13 | 0.18 | Neutral | | | | 1.13 | 0.18 | Neutral |
| 156 | p.Asp156Tyr | 1.13 | 0.17 | Neutral | | | | 1.13 | 0.17 | Neutral |
| 156 | p.Asp156Cys | 1.10 | 0.14 | Neutral | | | | 1.10 | 0.14 | Neutral |
| 156 | p.Asp156Trp | 0.90 | -0.15 | Neutral | | | | 0.90 | -0.15 | Neutral |
| 156 | p.Asp156Phe | 1.00 | 0.00 | Neutral | | | | 1.00 | 0.00 | Neutral |
