## Appendix 1-table 7 for "Functional characterization of all *CDKN2A* missense variants and comparison to in silico models of pathogenicity"

Appendix 1-table 7. In silico variant effect predictions for *CDKN2A* missense variants.

| Protein | ACMG Guidline | This study | | AlphaMisssense | | ESM1b | | VEST | | Polyphen-2 | | SIFT | | CADD | |
|---|---|---|---|---|---|---|---|---|---|---|---|---|---|---|---|
| Consequence | Classification | Log P value | Characteritic | Score | Prediction | Score | Prediction | Score | Prediction | Score | Prediction | Score | Prediction | Score | Prediction |
| p.Met1Asn | VUS | -0.15 | Neutral | 0.34 | Benign | -11.02 | Pathogenic | 0.44 | Neutral | 0.19 | Benign | 0 | Deleterious | | |
| p.Met1Lys | VUS | -0.01 | Neutral | 0.18 | Benign | -10.15 | Pathogenic | 0.48 | Neutral | 0.04 | Benign | 0 | Deleterious | 14.26 | Neutral |
| p.Met1Thr | VUS | -0.14 | Neutral | 0.11 | Benign | -8.46 | Pathogenic | 0.43 | Neutral | 0.00 | Benign | 0 | Deleterious | 12.50 | Neutral |
| p.Met1Arg | VUS | -0.22 | Neutral | 0.10 | Benign | -8.48 | Pathogenic | 0.57 | Disease | 0.00 | Benign | 0 | Deleterious | 14.15 | Neutral |
| p.Met1Ser | VUS | -0.01 | Neutral | 0.15 | Benign | -7.86 | Pathogenic | 0.43 | Neutral | 0.05 | Benign | 0 | Deleterious | | |
| p.Met1Ile | VUS | -0.44 | Neutral | 0.22 | Benign | -10.40 | Pathogenic | 0.36 | Neutral | 0.15 | Benign | 0 | Deleterious | 18.53 | Deleterious |
| p.Met1His | VUS | -1.27 | Neutral | 0.22 | Benign | -10.63 | Pathogenic | 0.41 | Neutral | 0.62 | Possibly damaging | 0 | Deleterious | | |
| p.Met1Gln | VUS | -0.12 | Neutral | 0.07 | Benign | -9.16 | Pathogenic | 0.40 | Neutral | 0.10 | Benign | 0 | Deleterious | | |
| p.Met1Pro | VUS | -0.36 | Neutral | 0.22 | Benign | -7.53 | Pathogenic | 0.44 | Neutral | 0.32 | Benign | 0 | Deleterious | | |
| p.Met1Leu | VUS | -0.02 | Neutral | 0.11 | Benign | -8.18 | Pathogenic | 0.35 | Neutral | 0.03 | Benign | 0 | Deleterious | 16.05 | Deleterious |
| p.Met1Asp | VUS | 0.00 | Neutral | 0.33 | Benign | -9.42 | Pathogenic | 0.43 | Neutral | 0.05 | Benign | 0 | Deleterious | | |
| p.Met1Glu | VUS | -0.96 | Neutral | 0.18 | Benign | -8.42 | Pathogenic | 0.46 | Neutral | 0.00 | Benign | 0 | Deleterious | | |
| p.Met1Ala | VUS | -0.07 | Neutral | 0.08 | Benign | -7.47 | Benign | 0.39 | Neutral | 0.02 | Benign | 0 | Deleterious | | |
| p.Met1Gly | VUS | -0.36 | Neutral | 0.16 | Benign | -7.93 | Pathogenic | 0.44 | Neutral | 0.00 | Benign | 0 | Deleterious | | |
| p.Met1Val | VUS | -4.16 | Neutral | 0.07 | Benign | -8.26 | Pathogenic | 0.30 | Neutral | 0.04 | Benign | 0 | Deleterious | 15.11 | Deleterious |
| p.Met1Tyr | VUS | -0.02 | Neutral | 0.31 | Benign | -11.51 | Pathogenic | 0.37 | Neutral | 0.83 | Possibly damaging | 0 | Deleterious | | |
| p.Met1Cys | VUS | -0.01 | Neutral | 0.14 | Benign | -10.28 | Pathogenic | 0.41 | Neutral | 0.95 | Possibly damaging | 0 | Deleterious | | |
| p.Met1Trp | VUS | -1.15 | Neutral | 0.19 | Benign | -10.35 | Pathogenic | 0.37 | Neutral | 0.95 | Possibly damaging | 0 | Deleterious | | |
| p.Met1Phe | VUS | -0.27 | Neutral | 0.12 | Benign | -10.79 | Pathogenic | 0.34 | Neutral | 0.83 | Possibly damaging | 0 | Deleterious | | |
| p.Glu2Asn | VUS | 0.00 | Neutral | 0.16 | Benign | -6.38 | Benign | 0.25 | Neutral | 0.21 | Benign | 0 | Deleterious | | |
| p.Glu2Lys | VUS | 0.00 | Neutral | 0.18 | Benign | -6.28 | Benign | 0.22 | Neutral | 0.08 | Benign | 0 | Deleterious | 18.01 | Deleterious |
| p.Glu2Thr | VUS | 0.00 | Neutral | 0.14 | Benign | -5.22 | Benign | 0.26 | Neutral | 0.34 | Benign | 0 | Deleterious | | |
| p.Glu2Arg | VUS | 0.00 | Neutral | 0.14 | Benign | -5.17 | Benign | 0.29 | Neutral | 0.00 | Benign | 0 | Deleterious | | |
| p.Glu2Ser | VUS | -0.13 | Neutral | 0.11 | Benign | -4.30 | Benign | 0.30 | Neutral | 0.34 | Benign | 0 | Deleterious | | |
| p.Glu2Ile | VUS | 0.00 | Neutral | 0.25 | Benign | -7.33 | Benign | 0.33 | Neutral | 0.65 | Possibly damaging | 0 | Deleterious | | |
| p.Glu2Met | VUS | -0.01 | Neutral | 0.36 | Ambiguous | -7.24 | Benign | 0.33 | Neutral | 0.93 | Possibly damaging | 0 | Deleterious | | |
| p.Glu2His | VUS | 0.00 | Neutral | 0.18 | Benign | -6.40 | Benign | 0.28 | Neutral | 0.65 | Possibly damaging | 0 | Deleterious | | |
| p.Glu2Gln | VUS | 0.00 | Neutral | 0.09 | Benign | -5.39 | Benign | 0.17 | Neutral | 0.01 | Benign | 0 | Deleterious | 12.97 | Neutral |
| p.Glu2Pro | VUS | 0.00 | Neutral | 0.12 | Benign | -4.12 | Benign | 0.32 | Neutral | 0.79 | Possibly damaging | 0 | Deleterious | | |
| p.Glu2Leu | VUS | 0.00 | Neutral | 0.20 | Benign | -5.40 | Benign | 0.32 | Neutral | 0.21 | Benign | 0 | Deleterious | | |
| p.Glu2Asp | VUS | 0.00 | Neutral | 0.10 | Benign | -4.35 | Benign | 0.17 | Neutral | 0.00 | Benign | 0 | Deleterious | 13.41 | Neutral |
| p.Glu2Ala | VUS | -0.02 | Neutral | 0.09 | Benign | -3.30 | Benign | 0.19 | Neutral | 0.16 | Benign | 0 | Deleterious | 18.03 | Deleterious |
| p.Glu2Gly | VUS | 0.00 | Neutral | 0.10 | Benign | -4.46 | Benign | 0.24 | Neutral | 0.28 | Benign | 0 | Deleterious | 20.50 | Deleterious |
| p.Glu2Val | VUS | 0.00 | Neutral | 0.18 | Benign | -5.70 | Benign | 0.21 | Neutral | 0.58 | Possibly damaging | 0 | Deleterious | 19.22 | Deleterious |
| p.Glu2Tyr | VUS | 0.00 | Neutral | 0.29 | Benign | -7.99 | Pathogenic | 0.32 | Neutral | 0.00 | Benign | 0 | Deleterious | | |
| p.Glu2Cys | VUS | -0.05 | Neutral | 0.57 | Pathogenic | -6.92 | Benign | 0.34 | Neutral | 0.98 | Probably damaging | 0 | Deleterious | | |
| p.Glu2Trp | VUS | 0.00 | Neutral | 0.56 | Ambiguous | -7.47 | Benign | 0.35 | Neutral | 0.93 | Possibly damaging | 0 | Deleterious | | |
| p.Glu2Phe | VUS | 0.00 | Neutral | 0.33 | Benign | -7.07 | Benign | 0.34 | Neutral | 0.48 | Possibly damaging | 0 | Deleterious | | |
| p.Pro3Asn | VUS | -7.64 | Indeterminate | 0.15 | Benign | -6.32 | Benign | 0.28 | Neutral | 0.05 | Benign | 0 | Deleterious | | |
| p.Pro3Lys | VUS | -18.39 | Indeterminate | 0.14 | Benign | -6.26 | Benign | 0.24 | Neutral | 0.01 | Benign | 0 | Deleterious | | |
| p.Pro3Thr | VUS | -10.80 | Indeterminate | 0.06 | Benign | -4.43 | Benign | 0.18 | Neutral | 0.02 | Benign | 0 | Deleterious | 12.72 | Neutral |
| p.Pro3Arg | VUS | -33.21 | Indeterminate | 0.08 | Benign | -4.81 | Benign | 0.21 | Neutral | 0.00 | Benign | 0 | Deleterious | 0.00 | Neutral |
| p.Pro3Ser | VUS | -22.54 | Indeterminate | 0.07 | Benign | -3.83 | Benign | 0.16 | Neutral | 0.00 | Benign | 0 | Deleterious | 12.92 | Neutral |
| p.Pro3Ile | VUS | -11.27 | Indeterminate | 0.17 | Benign | -6.97 | Benign | 0.25 | Neutral | 0.03 | Benign | 0 | Deleterious | | |
| p.Pro3Met | VUS | -23.54 | Indeterminate | 0.21 | Benign | -6.54 | Benign | 0.24 | Neutral | 0.32 | Benign | 0 | Deleterious | | |
| p.Pro3His | VUS | -5.11 | Neutral | 0.11 | Benign | -6.20 | Benign | 0.18 | Neutral | 0.00 | Benign | 0 | Deleterious | | |
| p.Pro3Gln | VUS | -0.28 | Neutral | 0.08 | Benign | -5.60 | Benign | 0.25 | Neutral | 0.00 | Benign | 0 | Deleterious | 0.01 | Neutral |
| p.Pro3Leu | VUS | -15.02 | Indeterminate | 0.11 | Benign | -5.31 | Benign | 0.14 | Neutral | 0.00 | Benign | 0 | Deleterious | 0.02 | Neutral |
| p.Pro3Asp | VUS | -14.44 | Indeterminate | 0.12 | Benign | -4.22 | Benign | 0.26 | Neutral | 0.05 | Benign | 0 | Deleterious | | |
| p.Pro3Glu | VUS | -4.70 | Neutral | 0.08 | Benign | -3.98 | Benign | 0.28 | Neutral | 0.01 | Benign | 0 | Deleterious | | |
| p.Pro3Ala | VUS | -3.70 | Neutral | 0.04 | Benign | -2.61 | Benign | 0.21 | Neutral | 0.01 | Benign | 0 | Deleterious | 10.50 | Neutral |
| p.Pro3Gly | VUS | -53.15 | Deleterious | 0.10 | Benign | -3.74 | Benign | 0.25 | Neutral | 0.03 | Benign | 0 | Deleterious | | |
| p.Pro3Val | VUS | -14.34 | Indeterminate | 0.10 | Benign | -5.05 | Benign | 0.24 | Neutral | 0.03 | Benign | 0 | Deleterious | | |
| p.Pro3Tyr | VUS | -5.47 | Neutral | 0.25 | Benign | -7.97 | Pathogenic | 0.26 | Neutral | 0.00 | Benign | 0 | Deleterious | | |
| p.Pro3Cys | VUS | -11.32 | Indeterminate | 0.33 | Benign | -6.71 | Benign | 0.25 | Neutral | 0.75 | Possibly damaging | 0 | Deleterious | | |
| p.Pro3Trp | VUS | -31.79 | Indeterminate | 0.32 | Benign | -7.79 | Pathogenic | 0.27 | Neutral | 0.49 | Possibly damaging | 0 | Deleterious | | |
| p.Pro3Phe | VUS | -12.58 | Indeterminate | 0.25 | Benign | -7.36 | Benign | 0.26 | Neutral | 0.06 | Benign | 0 | Deleterious | | |
| p.Ala4Asn | VUS | -5.71 | Neutral | 0.20 | Benign | -5.54 | Benign | 0.24 | Neutral | 0.19 | Benign | 0.05 | Deleterious | | |
| p.Ala4Lys | VUS | -1.31 | Neutral | 0.22 | Benign | -6.17 | Benign | 0.26 | Neutral | 0.19 | Benign | 0.07 | Tolerated | | |
| p.Ala4Thr | VUS | -33.22 | Indeterminate | 0.09 | Benign | -4.69 | Benign | 0.20 | Neutral | 0.15 | Benign | 0.11 | Tolerated | 15.19 | Deleterious |
| p.Ala4Arg | VUS | -1.41 | Neutral | 0.14 | Benign | -4.55 | Benign | 0.26 | Neutral | 0.45 | Benign | 0.05 | Deleterious | | |
| p.Ala4Ser | VUS | -53.15 | Deleterious | 0.09 | Benign | -2.95 | Benign | 0.19 | Neutral | 0.07 | Benign | 0.5 | Tolerated | 10.21 | Neutral |
| p.Ala4Ile | VUS | -0.18 | Neutral | 0.22 | Benign | -6.32 | Benign | 0.24 | Neutral | 0.10 | Benign | 0.03 | Deleterious | | |
| p.Ala4Met | VUS | -7.81 | Indeterminate | 0.25 | Benign | -6.25 | Benign | 0.23 | Neutral | 0.62 | Possibly damaging | 0.02 | Deleterious | | |
| p.Ala4His | VUS | -1.31 | Neutral | 0.20 | Benign | -6.23 | Benign | 0.24 | Neutral | 0.83 | Possibly damaging | 0.02 | Deleterious | | |
| p.Ala4Gln | VUS | -0.32 | Neutral | 0.14 | Benign | -5.70 | Benign | 0.24 | Neutral | 0.45 | Benign | 0.05 | Deleterious | | |
| p.Ala4Pro | VUS | -0.63 | Neutral | 0.08 | Benign | -3.99 | Benign | 0.23 | Neutral | 0.79 | Possibly damaging | 0.07 | Tolerated | 15.45 | Deleterious |
| p.Ala4Leu | VUS | -0.91 | Neutral | 0.14 | Benign | -4.80 | Benign | 0.27 | Neutral | 0.05 | Benign | 0.04 | Deleterious | | |
| p.Ala4Asp | VUS | -3.20 | Neutral | 0.10 | Benign | -4.74 | Benign | 0.23 | Neutral | 0.10 | Benign | 0.06 | Tolerated | | |
| p.Ala4Glu | VUS | -0.02 | Neutral | 0.10 | Benign | -4.54 | Benign | 0.21 | Neutral | 0.01 | Benign | 0.08 | Tolerated | 12.91 | Neutral |
| p.Ala4Gly | VUS | -6.58 | Indeterminate | 0.08 | Benign | -3.58 | Benign | 0.21 | Neutral | 0.00 | Benign | 0.13 | Tolerated | 14.74 | Neutral |
| p.Ala4Val | VUS | -0.11 | Neutral | 0.11 | Benign | -4.86 | Benign | 0.13 | Neutral | 0.00 | Benign | 0.06 | Tolerated | 14.00 | Neutral |
| p.Ala4Tyr | VUS | -0.67 | Neutral | 0.27 | Benign | -7.04 | Benign | 0.27 | Neutral | 0.45 | Benign | 0.02 | Deleterious | | |
| p.Ala4Cys | VUS | -32.26 | Indeterminate | 0.29 | Benign | -5.69 | Benign | 0.25 | Neutral | 0.95 | Possibly damaging | 0.02 | Deleterious | | |
| p.Ala4Trp | VUS | -2.02 | Neutral | 0.33 | Benign | -7.03 | Benign | 0.23 | Neutral | 0.00 | Benign | 0.01 | Deleterious | | |
| p.Ala4Phe | VUS | -8.96 | Indeterminate | 0.20 | Benign | -6.43 | Benign | 0.25 | Neutral | 0.45 | Benign | 0.02 | Deleterious | | |
| p.Ala5Asn | VUS | -0.01 | Neutral | 0.19 | Benign | -5.78 | Benign | 0.27 | Neutral | 0.48 | Possibly damaging | 0.06 | Tolerated | | |
| p.Ala5Lys | VUS | 0.00 | Neutral | 0.21 | Benign | -5.65 | Benign | 0.27 | Neutral | 0.21 | Benign | 0.07 | Tolerated | | |
| p.Ala5Thr | VUS | -0.77 | Neutral | 0.09 | Benign | -4.08 | Benign | 0.17 | Neutral | 0.28 | Benign | 0.08 | Tolerated | 9.47 | Neutral |
| p.Ala5Arg | VUS | 0.00 | Neutral | 0.13 | Benign | -4.42 | Benign | 0.29 | Neutral | 0.65 | Possibly damaging | 0.05 | Deleterious | | |
| p.Ala5Ser | VUS | -33.22 | Indeterminate | 0.08 | Benign | -3.80 | Benign | 0.18 | Neutral | 0.02 | Benign | 0.19 | Tolerated | 6.32 | Neutral |
| p.Ala5Ile | VUS | 0.00 | Neutral | 0.22 | Benign | -6.07 | Benign | 0.28 | Neutral | 0.21 | Benign | 0.02 | Deleterious | | |
| p.Ala5Met | VUS | -0.27 | Neutral | 0.25 | Benign | -4.19 | Benign | 0.29 | Neutral | 0.65 | Possibly damaging | 0.02 | Deleterious | | |
| p.Ala5His | VUS | -0.66 | Neutral | 0.20 | Benign | -6.30 | Benign | 0.28 | Neutral | 0.93 | Possibly damaging | 0.02 | Deleterious | | |
| p.Ala5Gln | VUS | -0.83 | Neutral | 0.14 | Benign | -5.76 | Benign | 0.27 | Neutral | 0.48 | Possibly damaging | 0.05 | Deleterious | | |
| p.Ala5Pro | VUS | -0.07 | Neutral | 0.07 | Benign | -3.82 | Benign | 0.26 | Neutral | 0.74 | Possibly damaging | 0.09 | Tolerated | 9.87 | Neutral |
| p.Ala5Leu | VUS | -9.21 | Indeterminate | 0.13 | Benign | -4.31 | Benign | 0.28 | Neutral | 0.00 | Benign | 0.05 | Deleterious | | |
| p.Ala5Asp | VUS | -1.63 | Neutral | 0.11 | Benign | -4.48 | Benign | 0.23 | Neutral | 0.00 | Benign | 0.06 | Tolerated | | |
| p.Ala5Glu | VUS | -6.48 | Indeterminate | 0.10 | Benign | -4.50 | Benign | 0.23 | Neutral | 0.00 | Benign | 0.08 | Tolerated | 7.82 | Neutral |
| p.Ala5Gly | VUS | -0.22 | Neutral | 0.07 | Benign | -3.84 | Benign | 0.21 | Neutral | 0.29 | Benign | 0.48 | Tolerated | 9.20 | Neutral |
| p.Ala5Val | VUS | -0.09 | Neutral | 0.12 | Benign | -4.34 | Benign | 0.20 | Neutral | 0.08 | Benign | 0.06 | Tolerated | 9.18 | Neutral |
| p.Ala5Tyr | VUS | -53.15 | Deleterious | 0.27 | Benign | -7.06 | Benign | 0.31 | Neutral | 0.79 | Possibly damaging | 0.02 | Deleterious | | |
| p.Ala5Cys | VUS | -4.14 | Neutral | 0.29 | Benign | -5.94 | Benign | 0.26 | Neutral | 0.98 | Probably damaging | 0.03 | Deleterious | | |
| p.Ala5Trp | VUS | -25.53 | Indeterminate | 0.30 | Benign | -6.63 | Benign | 0.32 | Neutral | 0.98 | Probably damaging | 0.01 | Deleterious | | |
| p.Ala5Phe | VUS | -0.47 | Neutral | 0.19 | Benign | -6.40 | Benign | 0.27 | Neutral | 0.65 | Possibly damaging | 0.02 | Deleterious | | |
| p.Gly6Asn | VUS | -0.09 | Neutral | 0.19 | Benign | -6.09 | Benign | 0.23 | Neutral | 1.00 | Probably damaging | 0 | Deleterious | | |
| p.Gly6Lys | VUS | -2.25 | Neutral | 0.26 | Benign | -5.94 | Benign | 0.23 | Neutral | 1.00 | Probably damaging | 0 | Deleterious | | |
| p.Gly6Thr | VUS | -4.22 | Neutral | 0.08 | Benign | -4.52 | Benign | 0.25 | Neutral | 1.00 | Probably damaging | 0 | Deleterious | | |
| p.Gly6Arg | VUS | -53.15 | Deleterious | 0.16 | Benign | -4.42 | Benign | 0.14 | Neutral | 1.00 | Probably damaging | 0 | Deleterious | 14.96 | Neutral |
| p.Gly6Ser | VUS | -33.22 | Indeterminate | 0.09 | Benign | -3.21 | Benign | 0.18 | Neutral | 1.00 | Probably damaging | 0 | Deleterious | | |
| p.Gly6Ile | VUS | -53.15 | Deleterious | 0.16 | Benign | -6.38 | Benign | 0.22 | Neutral | 1.00 | Probably damaging | 0 | Deleterious | | |
| p.Gly6Met | VUS | -33.20 | Indeterminate | 0.30 | Benign | -6.24 | Benign | 0.21 | Neutral | 1.00 | Probably damaging | 0 | Deleterious | | |
| p.Gly6His | VUS | 0.00 | Neutral | 0.19 | Benign | -6.10 | Benign | 0.24 | Neutral | 1.00 | Probably damaging | 0 | Deleterious | | |
| p.Gly6Gln | VUS | -0.08 | Neutral | 0.17 | Benign | -5.44 | Benign | 0.25 | Neutral | 1.00 | Probably damaging | 0 | Deleterious | | |
| p.Gly6Pro | VUS | -20.27 | Indeterminate | 0.14 | Benign | -3.48 | Benign | 0.25 | Neutral | 1.00 | Probably damaging | 0 | Deleterious | | |
| p.Gly6Leu | VUS | -0.87 | Neutral | 0.19 | Benign | -4.70 | Benign | 0.21 | Neutral | 1.00 | Probably damaging | 0 | Deleterious | | |
| p.Gly6Asp | VUS | -0.53 | Neutral | 0.09 | Benign | -3.84 | Benign | 0.16 | Neutral | 1.00 | Probably damaging | 0 | Deleterious | | |

| | | | | | | | | | | | | | | | |
|---|---|---|---|---|---|---|---|---|---|---|---|---|---|---|---|
| p.Gly6Glu | VUS | -0.72 | Neutral | 0.09 | Benign | -3.51 | Benign | 0.13 | Neutral | 1.00 | Probably damaging | 0 | Deleterious | 6.13 | Neutral |
| p.Gly6Ala | VUS | -0.20 | Neutral | 0.07 | Benign | -3.05 | Benign | 0.11 | Neutral | 1.00 | Probably damaging | 0 | Deleterious | 0.10 | Neutral |
| p.Gly6Val | VUS | -0.05 | Neutral | 0.10 | Benign | -4.63 | Benign | 0.14 | Neutral | 1.00 | Probably damaging | 0 | Deleterious | 7.89 | Neutral |
| p.Gly6Tyr | VUS | -0.77 | Neutral | 0.24 | Benign | -7.14 | Benign | 0.23 | Neutral | 1.00 | Probably damaging | 0 | Deleterious | | |
| p.Gly6Cys | VUS | -13.23 | Indeterminate | 0.15 | Benign | -5.88 | Benign | 0.18 | Neutral | 1.00 | Probably damaging | 0 | Deleterious | | |
| p.Gly6Trp | VUS | -3.24 | Neutral | 0.18 | Benign | -6.33 | Benign | 0.21 | Neutral | 1.00 | Probably damaging | 0 | Deleterious | 20.50 | Deleterious |
| p.Gly6Phe | VUS | -5.31 | Neutral | 0.27 | Benign | -6.22 | Benign | 0.21 | Neutral | 1.00 | Probably damaging | 0 | Deleterious | | |
| p.Ser7Asn | VUS | -0.07 | Neutral | 0.16 | Benign | -6.24 | Benign | 0.16 | Neutral | 0.91 | Possibly damaging | 0.41 | Tolerated | 10.26 | Neutral |
| p.Ser7Lys | VUS | -0.18 | Neutral | 0.31 | Benign | -6.64 | Benign | 0.27 | Neutral | 0.93 | Possibly damaging | 0.31 | Tolerated | | |
| p.Ser7Thr | VUS | -11.20 | Indeterminate | 0.09 | Benign | -4.38 | Benign | 0.19 | Neutral | 0.81 | Possibly damaging | 0.38 | Tolerated | 10.06 | Neutral |
| p.Ser7Arg | VUS | -0.03 | Neutral | 0.19 | Benign | -4.15 | Benign | 0.16 | Neutral | 0.97 | Probably damaging | 0.19 | Tolerated | 11.49 | Neutral |
| p.Ser7Ile | VUS | -1.33 | Neutral | 0.16 | Benign | -6.23 | Benign | 0.25 | Neutral | 0.97 | Probably damaging | 0.09 | Tolerated | 15.55 | Deleterious |
| p.Ser7Met | VUS | -0.71 | Neutral | 0.25 | Benign | -6.72 | Benign | 0.30 | Neutral | 0.98 | Probably damaging | 0.06 | Tolerated | | |
| p.Ser7His | VUS | -3.32 | Neutral | 0.15 | Benign | -5.85 | Benign | 0.28 | Neutral | 0.98 | Probably damaging | 0.1 | Tolerated | | |
| p.Ser7Gln | VUS | -0.38 | Neutral | 0.20 | Benign | -5.54 | Benign | 0.25 | Neutral | 0.98 | Probably damaging | 0.2 | Tolerated | | |
| p.Ser7Pro | VUS | -0.71 | Neutral | 0.08 | Benign | -2.24 | Benign | 0.22 | Neutral | 0.93 | Possibly damaging | 0.25 | Tolerated | | |
| p.Ser7Leu | VUS | -8.57 | Indeterminate | 0.15 | Benign | -4.35 | Benign | 0.25 | Neutral | 0.93 | Possibly damaging | 0.15 | Tolerated | | |
| p.Ser7Asp | VUS | -0.96 | Neutral | 0.15 | Benign | -4.70 | Benign | 0.24 | Neutral | 0.93 | Possibly damaging | 0.39 | Tolerated | | |
| p.Ser7Glu | VUS | -29.43 | Indeterminate | 0.16 | Benign | -4.81 | Benign | 0.25 | Neutral | 0.93 | Possibly damaging | 0.31 | Tolerated | | |
| p.Ser7Ala | VUS | -19.51 | Indeterminate | 0.07 | Benign | -3.25 | Benign | 0.20 | Neutral | 0.71 | Possibly damaging | 0.82 | Tolerated | | |
| p.Ser7Gly | VUS | -0.02 | Neutral | 0.07 | Benign | -3.27 | Benign | 0.16 | Neutral | 0.81 | Possibly damaging | 0.72 | Tolerated | 9.07 | Neutral |
| p.Ser7Val | VUS | -4.25 | Neutral | 0.15 | Benign | -4.90 | Benign | 0.31 | Neutral | 0.93 | Possibly damaging | 0.18 | Tolerated | | |
| p.Ser7Tyr | VUS | -33.19 | Indeterminate | 0.14 | Benign | -7.30 | Benign | 0.25 | Neutral | 0.98 | Probably damaging | 0.09 | Tolerated | | |
| p.Ser7Cys | VUS | -3.14 | Neutral | 0.11 | Benign | -5.61 | Benign | 0.23 | Neutral | 0.99 | Probably damaging | 0.09 | Tolerated | 13.90 | Neutral |
| p.Ser7Trp | VUS | -6.60 | Indeterminate | 0.14 | Benign | -6.50 | Benign | 0.32 | Neutral | 0.99 | Probably damaging | 0.02 | Deleterious | | |
| p.Ser7Phe | VUS | -1.35 | Neutral | 0.15 | Benign | -6.30 | Benign | 0.27 | Neutral | 0.98 | Probably damaging | 0.07 | Tolerated | | |
| p.Ser8Asn | VUS | 0.00 | Neutral | 0.19 | Benign | -5.65 | Benign | 0.15 | Neutral | 0.91 | Possibly damaging | 0 | Deleterious | 11.20 | Neutral |
| p.Ser8Lys | VUS | -21.11 | Indeterminate | 0.35 | Ambiguous | -6.11 | Benign | 0.26 | Neutral | 0.93 | Possibly damaging | 0 | Deleterious | | |
| p.Ser8Thr | VUS | -0.54 | Neutral | 0.09 | Benign | -4.19 | Benign | 0.15 | Neutral | 0.81 | Possibly damaging | 0 | Deleterious | 9.91 | Neutral |
| p.Ser8Arg | VUS | -0.04 | Neutral | 0.20 | Benign | -4.19 | Benign | 0.21 | Neutral | 0.97 | Probably damaging | 0 | Deleterious | 13.46 | Neutral |
| p.Ser8Ile | VUS | -0.01 | Neutral | 0.20 | Benign | -5.85 | Benign | 0.21 | Neutral | 0.97 | Probably damaging | 0 | Deleterious | 19.43 | Deleterious |
| p.Ser8Met | VUS | -0.01 | Neutral | 0.29 | Benign | -6.03 | Benign | 0.28 | Neutral | 0.98 | Probably damaging | 0 | Deleterious | | |
| p.Ser8His | VUS | -0.01 | Neutral | 0.18 | Benign | -6.03 | Benign | 0.28 | Neutral | 0.98 | Probably damaging | 0 | Deleterious | | |
| p.Ser8Gln | VUS | -0.02 | Neutral | 0.22 | Benign | -5.47 | Benign | 0.26 | Neutral | 0.98 | Probably damaging | 0 | Deleterious | | |
| p.Ser8Pro | VUS | -0.80 | Neutral | 0.08 | Benign | -3.55 | Benign | 0.20 | Neutral | 0.93 | Possibly damaging | 0 | Deleterious | | |
| p.Ser8Leu | VUS | -0.43 | Neutral | 0.17 | Benign | -4.58 | Benign | 0.23 | Neutral | 0.93 | Possibly damaging | 0 | Deleterious | | |
| p.Ser8Asp | VUS | -0.03 | Neutral | 0.18 | Benign | -4.38 | Benign | 0.26 | Neutral | 0.93 | Possibly damaging | 0 | Deleterious | | |
| p.Ser8Glu | VUS | 0.00 | Neutral | 0.18 | Benign | -4.18 | Benign | 0.26 | Neutral | 0.93 | Possibly damaging | 0 | Deleterious | | |
| p.Ser8Ala | VUS | -0.09 | Neutral | 0.07 | Benign | -2.85 | Benign | 0.18 | Neutral | 0.71 | Possibly damaging | 0 | Deleterious | | |
| p.Ser8Gly | VUS | -0.42 | Neutral | 0.07 | Benign | -3.03 | Benign | 0.16 | Neutral | 0.81 | Possibly damaging | 0 | Deleterious | 11.53 | Neutral |
| p.Ser8Val | VUS | -0.03 | Neutral | 0.15 | Benign | -4.45 | Benign | 0.27 | Neutral | 0.93 | Possibly damaging | 0 | Deleterious | | |
| p.Ser8Tyr | VUS | -0.01 | Neutral | 0.18 | Benign | -6.89 | Benign | 0.25 | Neutral | 0.98 | Probably damaging | 0 | Deleterious | | |
| p.Ser8Cys | VUS | 0.00 | Neutral | 0.12 | Benign | -5.77 | Benign | 0.22 | Neutral | 0.99 | Probably damaging | 0 | Deleterious | 19.53 | Deleterious |
| p.Ser8Trp | VUS | 0.00 | Neutral | 0.17 | Benign | -5.94 | Benign | 0.32 | Neutral | 0.99 | Probably damaging | 0 | Deleterious | | |
| p.Ser8Phe | VUS | -0.23 | Neutral | 0.18 | Benign | -6.25 | Benign | 0.24 | Neutral | 0.98 | Probably damaging | 0 | Deleterious | | |
| p.Met9Asn | VUS | -0.21 | Neutral | 0.28 | Benign | -4.02 | Benign | 0.33 | Neutral | 0.13 | Benign | 0.49 | Tolerated | | |
| p.Met9Lys | VUS | -0.23 | Neutral | 0.13 | Benign | -3.57 | Benign | 0.35 | Neutral | 0.04 | Benign | 0.52 | Tolerated | 10.96 | Neutral |
| p.Met9Thr | VUS | -0.27 | Neutral | 0.11 | Benign | -1.57 | Benign | 0.17 | Neutral | 0.00 | Benign | 0.51 | Tolerated | 9.04 | Neutral |
| p.Met9Arg | VUS | -0.07 | Neutral | 0.08 | Benign | -1.77 | Benign | 0.33 | Neutral | 0.10 | Benign | 0.31 | Tolerated | 11.88 | Neutral |
| p.Met9Ser | VUS | -1.02 | Neutral | 0.14 | Benign | -1.00 | Benign | 0.33 | Neutral | 0.01 | Benign | 0.6 | Tolerated | | |
| p.Met9Ile | VUS | -0.29 | Neutral | 0.33 | Benign | -3.66 | Benign | 0.27 | Neutral | 0.04 | Benign | 0.36 | Tolerated | 14.95 | Neutral |
| p.Met9His | VUS | -0.70 | Neutral | 0.19 | Benign | -3.54 | Benign | 0.33 | Neutral | 0.76 | Possibly damaging | 0.21 | Tolerated | | |
| p.Met9Gln | VUS | -0.37 | Neutral | 0.08 | Benign | -2.68 | Benign | 0.30 | Neutral | 0.23 | Benign | 0.42 | Tolerated | | |
| p.Met9Pro | VUS | -0.63 | Neutral | 0.12 | Benign | -0.65 | Benign | 0.35 | Neutral | 0.23 | Benign | 0.28 | Tolerated | | |
| p.Met9Leu | VUS | -0.01 | Neutral | 0.12 | Benign | -1.65 | Benign | 0.21 | Neutral | 0.00 | Benign | 0.68 | Tolerated | 6.18 | Neutral |
| p.Met9Asp | VUS | -0.13 | Neutral | 0.21 | Benign | -2.24 | Benign | 0.33 | Neutral | 0.13 | Benign | 0.77 | Tolerated | | |
| p.Met9Glu | VUS | -0.05 | Neutral | 0.12 | Benign | -1.93 | Benign | 0.35 | Neutral | 0.06 | Benign | 1 | Tolerated | | |
| p.Met9Ala | VUS | -0.12 | Neutral | 0.11 | Benign | -0.64 | Benign | 0.32 | Neutral | 0.01 | Benign | 0.68 | Tolerated | | |
| p.Met9Gly | VUS | -0.35 | Neutral | 0.10 | Benign | -1.14 | Benign | 0.28 | Neutral | 0.06 | Benign | 0.44 | Tolerated | | |
| p.Met9Val | VUS | -0.46 | Neutral | 0.07 | Benign | -2.11 | Benign | 0.22 | Neutral | 0.00 | Benign | 0.43 | Tolerated | 9.64 | Neutral |
| p.Met9Tyr | VUS | -0.02 | Neutral | 0.31 | Benign | -4.70 | Benign | 0.34 | Neutral | 0.47 | Possibly damaging | 0.24 | Tolerated | | |
| p.Met9Cys | VUS | -0.13 | Neutral | 0.30 | Benign | -3.72 | Benign | 0.34 | Neutral | 0.76 | Possibly damaging | 0.09 | Tolerated | | |
| p.Met9Trp | VUS | -0.13 | Neutral | 0.25 | Benign | -3.88 | Benign | 0.37 | Neutral | 0.76 | Possibly damaging | 0.06 | Tolerated | | |
| p.Met9Phe | VUS | -0.23 | Neutral | 0.17 | Benign | -4.09 | Benign | 0.33 | Neutral | 0.23 | Benign | 0.24 | Tolerated | | |
| p.Glu10Asn | VUS | -2.65 | Neutral | 0.33 | Benign | -6.10 | Benign | 0.30 | Neutral | 0.26 | Benign | 0 | Deleterious | | |
| p.Glu10Lys | VUS | -33.21 | Indeterminate | 0.26 | Benign | -5.90 | Benign | 0.25 | Neutral | 0.35 | Benign | 0 | Deleterious | 13.32 | Neutral |
| p.Glu10Thr | VUS | -5.04 | Neutral | 0.22 | Benign | -4.77 | Benign | 0.30 | Neutral | 0.59 | Possibly damaging | 0 | Deleterious | | |
| p.Glu10Arg | VUS | -0.85 | Neutral | 0.20 | Benign | -4.48 | Benign | 0.25 | Neutral | 0.83 | Possibly damaging | 0 | Deleterious | | |
| p.Glu10Ser | VUS | -22.02 | Indeterminate | 0.21 | Benign | -4.04 | Benign | 0.31 | Neutral | 0.41 | Benign | 0 | Deleterious | | |
| p.Glu10Ile | VUS | -13.01 | Indeterminate | 0.38 | Ambiguous | -6.67 | Benign | 0.33 | Neutral | 0.94 | Possibly damaging | 0 | Deleterious | | |
| p.Glu10Met | VUS | -9.12 | Indeterminate | 0.50 | Ambiguous | -6.58 | Benign | 0.36 | Neutral | 0.98 | Probably damaging | 0 | Deleterious | | |
| p.Glu10His | VUS | -32.21 | Indeterminate | 0.31 | Benign | -5.89 | Benign | 0.31 | Neutral | 0.94 | Possibly damaging | 0 | Deleterious | | |
| p.Glu10Gln | VUS | -0.37 | Neutral | 0.12 | Benign | -5.01 | Benign | 0.22 | Neutral | 0.52 | Possibly damaging | 0 | Deleterious | 11.32 | Neutral |
| p.Glu10Pro | VUS | -0.83 | Neutral | 0.20 | Benign | -3.49 | Benign | 0.35 | Neutral | 0.94 | Possibly damaging | 0 | Deleterious | | |
| p.Glu10Leu | VUS | -5.49 | Neutral | 0.30 | Benign | -4.45 | Benign | 0.30 | Neutral | 0.83 | Possibly damaging | 0 | Deleterious | | |
| p.Glu10Asp | VUS | -2.12 | Neutral | 0.16 | Benign | -3.82 | Benign | 0.14 | Neutral | 0.00 | Benign | 0 | Deleterious | 3.76 | Neutral |
| p.Glu10Ala | VUS | -32.81 | Indeterminate | 0.12 | Benign | -3.21 | Benign | 0.23 | Neutral | 0.35 | Benign | 0 | Deleterious | 11.44 | Neutral |
| p.Glu10Gly | VUS | -1.38 | Neutral | 0.15 | Benign | -2.92 | Benign | 0.21 | Neutral | 0.35 | Benign | 0 | Deleterious | 13.55 | Neutral |
| p.Glu10Val | VUS | -1.13 | Neutral | 0.22 | Benign | -4.69 | Benign | 0.26 | Neutral | 0.79 | Possibly damaging | 0 | Deleterious | 12.78 | Neutral |
| p.Glu10Tyr | VUS | -2.56 | Neutral | 0.45 | Ambiguous | -6.99 | Benign | 0.32 | Neutral | 0.94 | Possibly damaging | 0 | Deleterious | | |
| p.Glu10Cys | VUS | -1.13 | Neutral | 0.77 | Pathogenic | -6.29 | Benign | 0.34 | Neutral | 0.98 | Probably damaging | 0 | Deleterious | | |
| p.Glu10Trp | VUS | -1.25 | Neutral | 0.76 | Pathogenic | -6.63 | Benign | 0.36 | Neutral | 0.98 | Probably damaging | 0 | Deleterious | | |
| p.Glu10Phe | VUS | -2.65 | Neutral | 0.53 | Ambiguous | -6.37 | Benign | 0.33 | Neutral | 0.94 | Possibly damaging | 0 | Deleterious | | |
| p.Pro11Asn | VUS | -1.32 | Neutral | 0.32 | Benign | -6.47 | Benign | 0.27 | Neutral | 0.02 | Benign | 0.04 | Deleterious | | |
| p.Pro11Lys | VUS | -4.63 | Neutral | 0.28 | Benign | -7.00 | Benign | 0.25 | Neutral | 0.12 | Benign | 0.03 | Deleterious | | |
| p.Pro11Thr | VUS | -8.84 | Indeterminate | 0.10 | Benign | -4.72 | Benign | 0.36 | Neutral | 0.00 | Benign | 0.08 | Tolerated | 12.11 | Neutral |
| p.Pro11Arg | VUS | -1.90 | Neutral | 0.14 | Benign | -4.49 | Benign | 0.18 | Neutral | 0.26 | Benign | 0.02 | Deleterious | 15.68 | Deleterious |
| p.Pro11Ser | VUS | -2.27 | Neutral | 0.13 | Benign | -3.34 | Benign | 0.19 | Neutral | 0.00 | Benign | 0.39 | Tolerated | 13.98 | Neutral |
| p.Pro11Ile | VUS | -1.45 | Neutral | 0.27 | Benign | -6.46 | Benign | 0.29 | Neutral | 0.06 | Benign | 0.01 | Deleterious | | |
| p.Pro11Met | VUS | -0.32 | Neutral | 0.35 | Ambiguous | -6.10 | Benign | 0.28 | Neutral | 0.15 | Benign | 0.01 | Deleterious | | |
| p.Pro11His | VUS | -1.60 | Neutral | 0.21 | Benign | -6.18 | Benign | 0.21 | Neutral | 0.45 | Benign | 0.01 | Deleterious | 16.24 | Deleterious |
| p.Pro11Gln | VUS | -0.33 | Neutral | 0.16 | Benign | -5.73 | Benign | 0.28 | Neutral | 0.26 | Benign | 0.02 | Deleterious | | |
| p.Pro11Leu | VUS | -1.78 | Neutral | 0.17 | Benign | -4.57 | Benign | 0.17 | Neutral | 0.00 | Benign | 0.02 | Deleterious | 16.66 | Deleterious |
| p.Pro11Asp | VUS | -0.52 | Neutral | 0.29 | Benign | -4.51 | Benign | 0.27 | Neutral | 0.01 | Benign | 0.03 | Deleterious | | |
| p.Pro11Glu | VUS | -0.68 | Neutral | 0.20 | Benign | -4.43 | Benign | 0.28 | Neutral | 0.00 | Benign | 0.03 | Deleterious | | |
| p.Pro11Ala | VUS | -0.43 | Neutral | 0.07 | Benign | -3.36 | Benign | 0.24 | Neutral | 0.00 | Benign | 0.12 | Tolerated | 8.49 | Neutral |
| p.Pro11Gly | VUS | -1.89 | Neutral | 0.19 | Benign | -3.65 | Benign | 0.27 | Neutral | 0.03 | Benign | 0.07 | Tolerated | | |
| p.Pro11Val | VUS | -0.96 | Neutral | 0.16 | Benign | -4.78 | Benign | 0.27 | Neutral | 0.01 | Benign | 0.03 | Deleterious | | |
| p.Pro11Tyr | VUS | -2.83 | Neutral | 0.35 | Ambiguous | -6.83 | Benign | 0.26 | Neutral | 0.49 | Possibly damaging | 0.01 | Deleterious | | |
| p.Pro11Cys | VUS | -0.76 | Neutral | 0.50 | Ambiguous | -6.12 | Benign | 0.27 | Neutral | 0.99 | Probably damaging | 0.01 | Deleterious | | |
| p.Pro11Trp | VUS | -3.64 | Neutral | 0.51 | Ambiguous | -6.73 | Benign | 0.28 | Neutral | 0.92 | Possibly damaging | 0 | Deleterious | | |
| p.Pro11Phe | VUS | -2.45 | Neutral | 0.38 | Ambiguous | -6.30 | Benign | 0.28 | Neutral | 0.32 | Benign | 0.01 | Deleterious | | |
| p.Ser12Asn | VUS | -1.24 | Neutral | 0.22 | Benign | -5.32 | Benign | 0.21 | Neutral | 0.01 | Benign | 0.19 | Tolerated | | |
| p.Ser12Lys | VUS | -0.08 | Neutral | 0.49 | Ambiguous | -5.64 | Benign | 0.28 | Neutral | 0.10 | Benign | 0.2 | Tolerated | | |
| p.Ser12Thr | VUS | -0.10 | Neutral | 0.10 | Benign | -3.60 | Benign | 0.12 | Neutral | 0.00 | Benign | 0.56 | Tolerated | 9.20 | Neutral |
| p.Ser12Arg | VUS | -0.44 | Neutral | 0.36 | Ambiguous | -4.09 | Benign | 0.24 | Neutral | 0.18 | Benign | 0.15 | Tolerated | | |
| p.Ser12Ile | VUS | -0.29 | Neutral | 0.37 | Ambiguous | -6.38 | Benign | 0.23 | Neutral | 0.05 | Benign | 0.12 | Tolerated | | |
| p.Ser12Met | VUS | -0.04 | Neutral | 0.39 | Ambiguous | -5.38 | Benign | 0.22 | Neutral | 0.28 | Benign | 0.08 | Tolerated | | |
| p.Ser12His | VUS | -2.76 | Neutral | 0.28 | Benign | -5.49 | Benign | 0.29 | Neutral | 0.70 | Possibly damaging | 0.09 | Tolerated | | |
| p.Ser12Gln | VUS | -2.38 | Neutral | 0.33 | Benign | -4.82 | Benign | 0.26 | Neutral | 0.44 | Benign | 0.14 | Tolerated | | |
| p.Ser12Pro | VUS | -0.05 | Neutral | 0.10 | Benign | -3.18 | Benign | 0.27 | Neutral | 0.00 | Benign | 0.27 | Tolerated | 13.08 | Neutral |

| | | | | | | | | | | | | | | | |
|---|---|---|---|---|---|---|---|---|---|---|---|---|---|---|---|
| p.Ser12Leu | VUS | -28.42 | Indeterminate | 0.23 | Benign | -3.77 | Benign | 0.16 | Neutral | 0.00 | Benign | 0.19 | Tolerated | 17.06 | Deleterious |
| p.Ser12Asp | VUS | -0.01 | Neutral | 0.26 | Benign | -3.26 | Benign | 0.28 | Neutral | 0.06 | Benign | 0.17 | Tolerated | | |
| p.Ser12Glu | VUS | -1.21 | Neutral | 0.30 | Benign | -3.82 | Benign | 0.28 | Neutral | 0.04 | Benign | 0.18 | Tolerated | | |
| p.Ser12Ala | VUS | -0.70 | Neutral | 0.08 | Benign | -1.94 | Benign | 0.15 | Neutral | 0.00 | Benign | 1 | Tolerated | 8.47 | Neutral |
| p.Ser12Gly | VUS | -0.16 | Neutral | 0.09 | Benign | -3.33 | Benign | 0.16 | Neutral | 0.14 | Benign | 0.53 | Tolerated | | |
| p.Ser12Val | VUS | -4.85 | Neutral | 0.25 | Benign | -4.02 | Benign | 0.21 | Neutral | 0.01 | Benign | 0.25 | Tolerated | | |
| p.Ser12Tyr | VUS | -0.13 | Neutral | 0.28 | Benign | -6.41 | Benign | 0.24 | Neutral | 0.44 | Benign | 0.15 | Tolerated | | |
| p.Ser12Cys | VUS | -2.60 | Neutral | 0.15 | Benign | -5.63 | Benign | 0.23 | Neutral | 0.39 | Benign | 0.12 | Tolerated | | |
| p.Ser12Trp | VUS | -3.60 | Neutral | 0.30 | Benign | -6.43 | Benign | 0.24 | Neutral | 0.95 | Possibly damaging | 0.05 | Deleterious | 20.80 | Deleterious |
| p.Ser12Phe | VUS | -1.39 | Neutral | 0.35 | Ambiguous | -6.44 | Benign | 0.24 | Neutral | 0.28 | Benign | 0.13 | Tolerated | | |
| p.Ala13Asn | VUS | 0.00 | Neutral | 0.44 | Ambiguous | -6.94 | Benign | 0.29 | Neutral | 0.09 | Benign | 0.02 | Deleterious | | |
| p.Ala13Lys | VUS | -0.10 | Neutral | 0.62 | Pathogenic | -7.44 | Benign | 0.49 | Neutral | 0.65 | Possibly damaging | 0.02 | Deleterious | | |
| p.Ala13Thr | VUS | -0.02 | Neutral | 0.17 | Benign | -5.53 | Benign | 0.43 | Neutral | 0.33 | Benign | 0.04 | Deleterious | 23.30 | Deleterious |
| p.Ala13Arg | VUS | 0.00 | Neutral | 0.39 | Ambiguous | -5.27 | Benign | 0.48 | Neutral | 0.93 | Possibly damaging | 0.02 | Deleterious | | |
| p.Ala13Ser | VUS | 0.00 | Neutral | 0.13 | Benign | -4.23 | Benign | 0.43 | Neutral | 0.03 | Benign | 0.11 | Tolerated | 21.70 | Deleterious |
| p.Ala13Ile | VUS | -0.01 | Neutral | 0.50 | Ambiguous | -8.08 | Pathogenic | 0.49 | Neutral | 0.93 | Possibly damaging | 0.01 | Deleterious | | |
| p.Ala13Met | VUS | -0.82 | Neutral | 0.55 | Ambiguous | -7.42 | Benign | 0.45 | Neutral | 0.99 | Probably damaging | 0.01 | Deleterious | | |
| p.Ala13His | VUS | -0.17 | Neutral | 0.51 | Ambiguous | -6.96 | Benign | 0.46 | Neutral | 0.99 | Probably damaging | 0.01 | Deleterious | | |
| p.Ala13Gln | VUS | -0.02 | Neutral | 0.39 | Ambiguous | -5.95 | Benign | 0.47 | Neutral | 0.87 | Possibly damaging | 0.02 | Deleterious | | |
| p.Ala13Pro | VUS | 0.00 | Neutral | 0.11 | Benign | -4.31 | Benign | 0.53 | Disease | 0.73 | Possibly damaging | 0.05 | Deleterious | 23.60 | Deleterious |
| p.Ala13Leu | VUS | -0.18 | Neutral | 0.35 | Ambiguous | -6.06 | Benign | 0.33 | Neutral | 0.79 | Possibly damaging | 0.02 | Deleterious | | |
| p.Ala13Asp | VUS | -0.01 | Neutral | 0.37 | Ambiguous | -4.88 | Benign | 0.33 | Neutral | 0.00 | Benign | 0.02 | Deleterious | 15.22 | Deleterious |
| p.Ala13Glu | VUS | -0.01 | Neutral | 0.32 | Benign | -4.65 | Benign | 0.46 | Neutral | 0.03 | Benign | 0.02 | Deleterious | | |
| p.Ala13Gly | VUS | 0.00 | Neutral | 0.14 | Benign | -2.03 | Benign | 0.26 | Neutral | 0.00 | Benign | 1 | Tolerated | 10.78 | Neutral |
| p.Ala13Val | VUS | -0.05 | Neutral | 0.25 | Benign | -5.39 | Benign | 0.28 | Neutral | 0.52 | Possibly damaging | 0.03 | Deleterious | 16.07 | Deleterious |
| p.Ala13Tyr | VUS | -1.26 | Neutral | 0.58 | Pathogenic | -8.64 | Pathogenic | 0.50 | Neutral | 0.98 | Probably damaging | 0.01 | Deleterious | | |
| p.Ala13Cys | VUS | -0.59 | Neutral | 0.54 | Ambiguous | -6.73 | Benign | 0.44 | Neutral | 0.98 | Probably damaging | 0.02 | Deleterious | | |
| p.Ala13Trp | VUS | -0.39 | Neutral | 0.66 | Pathogenic | -6.89 | Benign | 0.56 | Disease | 0.99 | Probably damaging | 0 | Deleterious | | |
| p.Ala13Phe | VUS | -0.31 | Neutral | 0.51 | Ambiguous | -8.31 | Pathogenic | 0.50 | Neutral | 0.98 | Probably damaging | 0.01 | Deleterious | | |
| p.Asp14Asn | VUS | -3.70 | Neutral | 0.26 | Benign | -6.07 | Benign | 0.45 | Neutral | 0.87 | Possibly damaging | 0.51 | Tolerated | 23.20 | Deleterious |
| p.Asp14Lys | VUS | -2.50 | Neutral | 0.72 | Pathogenic | -7.26 | Benign | 0.38 | Neutral | 0.97 | Probably damaging | 0.05 | Deleterious | | |
| p.Asp14Thr | VUS | -0.45 | Neutral | 0.43 | Ambiguous | -5.05 | Benign | 0.55 | Disease | 0.90 | Possibly damaging | 0.03 | Deleterious | | |
| p.Asp14Arg | VUS | -15.07 | Indeterminate | 0.53 | Ambiguous | -4.63 | Benign | 0.55 | Disease | 0.99 | Probably damaging | 0.02 | Deleterious | | |
| p.Asp14Ser | VUS | -0.40 | Neutral | 0.25 | Benign | -3.92 | Benign | 0.50 | Disease | 0.94 | Possibly damaging | 0.06 | Tolerated | | |
| p.Asp14Ile | VUS | -31.69 | Indeterminate | 0.67 | Pathogenic | -7.71 | Pathogenic | 0.55 | Disease | 0.99 | Probably damaging | 0.01 | Deleterious | | |
| p.Asp14Met | VUS | -21.59 | Indeterminate | 0.87 | Pathogenic | -7.47 | Benign | 0.58 | Disease | 1.00 | Probably damaging | 0 | Deleterious | | |
| p.Asp14His | VUS | -0.67 | Neutral | 0.44 | Ambiguous | -5.88 | Benign | 0.50 | Disease | 0.99 | Probably damaging | 0.02 | Deleterious | 24.40 | Deleterious |
| p.Asp14Gln | VUS | -1.64 | Neutral | 0.50 | Ambiguous | -5.12 | Benign | 0.45 | Neutral | 0.96 | Probably damaging | 0.04 | Deleterious | | |
| p.Asp14Pro | VUS | -33.22 | Indeterminate | 0.62 | Pathogenic | -4.81 | Benign | 0.40 | Neutral | 0.73 | Possibly damaging | 0.03 | Deleterious | | |
| p.Asp14Leu | VUS | -11.64 | Indeterminate | 0.59 | Pathogenic | -5.32 | Benign | 0.58 | Disease | 0.99 | Probably damaging | 0.01 | Deleterious | | |
| p.Asp14Glu | VUS | -1.99 | Neutral | 0.28 | Benign | -3.12 | Benign | 0.23 | Neutral | 0.00 | Benign | 0.23 | Tolerated | 15.97 | Deleterious |
| p.Asp14Ala | VUS | -10.01 | Indeterminate | 0.30 | Benign | -3.02 | Benign | 0.30 | Neutral | 0.14 | Benign | 0.04 | Deleterious | 19.29 | Deleterious |
| p.Asp14Gly | VUS | -0.82 | Neutral | 0.31 | Benign | -3.87 | Benign | 0.51 | Disease | 0.96 | Possibly damaging | 0.08 | Tolerated | 22.80 | Deleterious |
| p.Asp14Val | VUS | -31.29 | Indeterminate | 0.46 | Ambiguous | -5.24 | Benign | 0.60 | Disease | 0.82 | Possibly damaging | 0.01 | Deleterious | 22.80 | Deleterious |
| p.Asp14Tyr | VUS | -18.85 | Indeterminate | 0.41 | Ambiguous | -8.06 | Pathogenic | 0.54 | Disease | 1.00 | Probably damaging | 0.01 | Deleterious | 24.80 | Deleterious |
| p.Asp14Cys | VUS | -1.62 | Neutral | 0.78 | Pathogenic | -6.51 | Benign | 0.56 | Disease | 1.00 | Probably damaging | 0.01 | Deleterious | | |
| p.Asp14Trp | VUS | -19.43 | Indeterminate | 0.88 | Pathogenic | -7.13 | Benign | 0.59 | Disease | 1.00 | Probably damaging | 0 | Deleterious | | |
| p.Asp14Phe | VUS | -21.99 | Indeterminate | 0.76 | Pathogenic | -7.76 | Pathogenic | 0.58 | Disease | 1.00 | Probably damaging | 0.01 | Deleterious | | |
| p.Trp15Asn | VUS | -4.97 | Neutral | 0.23 | Benign | -3.04 | Benign | 0.27 | Neutral | 0.89 | Possibly damaging | 0.28 | Tolerated | | |
| p.Trp15Lys | VUS | -2.58 | Neutral | 0.23 | Benign | -2.62 | Benign | 0.26 | Neutral | 0.01 | Benign | 1 | Tolerated | | |
| p.Trp15Thr | VUS | -11.33 | Indeterminate | 0.08 | Benign | -1.18 | Benign | 0.30 | Neutral | 0.64 | Possibly damaging | 0.31 | Tolerated | | |
| p.Trp15Arg | VUS | -1.65 | Neutral | 0.11 | Benign | 0.64 | Benign | 0.12 | Neutral | 0.01 | Benign | 0.78 | Tolerated | 0.23 | Neutral |
| p.Trp15Ser | VUS | -2.11 | Neutral | 0.06 | Benign | 0.17 | Benign | 0.19 | Neutral | 0.08 | Benign | 0.34 | Tolerated | 8.05 | Neutral |
| p.Trp15Ile | VUS | -1.10 | Neutral | 0.16 | Benign | -4.07 | Benign | 0.31 | Neutral | 0.49 | Possibly damaging | 0.12 | Tolerated | | |
| p.Trp15Met | VUS | -1.86 | Neutral | 0.26 | Benign | -3.17 | Benign | 0.25 | Neutral | 0.44 | Benign | 0.08 | Tolerated | | |
| p.Trp15His | VUS | -23.94 | Indeterminate | 0.15 | Benign | -2.56 | Benign | 0.33 | Neutral | 0.99 | Probably damaging | 0.16 | Tolerated | | |
| p.Trp15Gln | VUS | -4.61 | Neutral | 0.17 | Benign | -0.15 | Benign | 0.22 | Neutral | 0.10 | Benign | 0.44 | Tolerated | | |
| p.Trp15Pro | VUS | -27.76 | Indeterminate | 0.31 | Benign | 1.37 | Benign | 0.32 | Neutral | 0.98 | Probably damaging | 0.18 | Tolerated | | |
| p.Trp15Leu | VUS | -0.52 | Neutral | 0.10 | Benign | -0.60 | Benign | 0.18 | Neutral | 0.01 | Benign | 0.23 | Tolerated | 9.65 | Neutral |
| p.Trp15Asp | VUS | -0.47 | Neutral | 0.11 | Benign | 0.24 | Benign | 0.28 | Neutral | 0.89 | Possibly damaging | 0.26 | Tolerated | | |
| p.Trp15Glu | VUS | -0.29 | Neutral | 0.09 | Benign | 0.93 | Benign | 0.26 | Neutral | 0.10 | Benign | 0.53 | Tolerated | | |
| p.Trp15Ala | VUS | -1.52 | Neutral | 0.04 | Benign | 1.49 | Benign | 0.31 | Neutral | 0.03 | Benign | 0.37 | Tolerated | | |
| p.Trp15Gly | VUS | -9.01 | Indeterminate | 0.05 | Benign | -0.71 | Benign | 0.20 | Neutral | 0.57 | Possibly damaging | 0.23 | Tolerated | 1.39 | Neutral |
| p.Trp15Val | VUS | -1.17 | Neutral | 0.09 | Benign | -1.43 | Benign | 0.32 | Neutral | 0.64 | Possibly damaging | 0.18 | Tolerated | | |
| p.Trp15Tyr | VUS | -5.94 | Indeterminate | 0.24 | Benign | -5.65 | Benign | 0.24 | Neutral | 0.74 | Possibly damaging | 0.12 | Tolerated | | |
| p.Trp15Cys | VUS | -5.76 | Neutral | 0.44 | Ambiguous | -3.13 | Benign | 0.30 | Neutral | 0.99 | Probably damaging | 0.04 | Deleterious | 22.95 | Deleterious |
| p.Trp15Phe | VUS | -0.61 | Neutral | 0.14 | Benign | -4.96 | Benign | 0.25 | Neutral | 0.86 | Possibly damaging | 0.08 | Tolerated | | |
| p.Leu16Asn | VUS | -53.15 | Deleterious | 0.97 | Pathogenic | -15.85 | Pathogenic | 0.80 | Disease | 1.00 | Probably damaging | 0 | Deleterious | | |
| p.Leu16Lys | VUS | -53.15 | Deleterious | 0.93 | Pathogenic | -15.37 | Pathogenic | 0.80 | Disease | 0.99 | Probably damaging | 0 | Deleterious | | |
| p.Leu16Thr | VUS | -27.55 | Indeterminate | 0.83 | Pathogenic | -12.66 | Pathogenic | 0.79 | Disease | 1.00 | Probably damaging | 0 | Deleterious | | |
| p.Leu16Arg | Pathogenic | -53.15 | Deleterious | 0.84 | Pathogenic | -11.98 | Pathogenic | 0.92 | Disease | 1.00 | Probably damaging | 0 | Deleterious | 22.80 | Deleterious |
| p.Leu16Ser | VUS | -53.15 | Deleterious | 0.97 | Pathogenic | -12.29 | Pathogenic | 0.89 | Disease | 1.00 | Probably damaging | 0 | Deleterious | | |
| p.Leu16Ile | VUS | -6.96 | Indeterminate | 0.23 | Benign | -9.94 | Pathogenic | 0.53 | Disease | 1.00 | Probably damaging | 0 | Deleterious | | |
| p.Leu16Met | VUS | -2.63 | Neutral | 0.60 | Pathogenic | -9.00 | Pathogenic | 0.66 | Disease | 1.00 | Probably damaging | 0 | Deleterious | 20.30 | Deleterious |
| p.Leu16His | VUS | -53.15 | Deleterious | 0.97 | Pathogenic | -12.95 | Pathogenic | 0.86 | Disease | 1.00 | Probably damaging | 0 | Deleterious | | |
| p.Leu16Gln | VUS | -53.15 | Deleterious | 0.92 | Pathogenic | -12.59 | Pathogenic | 0.74 | Disease | 1.00 | Probably damaging | 0 | Deleterious | 22.60 | Deleterious |
| p.Leu16Pro | Pathogenic | -53.15 | Deleterious | 0.87 | Pathogenic | -9.98 | Pathogenic | 0.94 | Disease | 1.00 | Probably damaging | 0 | Deleterious | 22.70 | Deleterious |
| p.Leu16Asp | VUS | -53.15 | Deleterious | 0.99 | Pathogenic | -14.19 | Pathogenic | 0.80 | Disease | 1.00 | Probably damaging | 0 | Deleterious | | |
| p.Leu16Glu | VUS | -53.15 | Deleterious | 0.94 | Pathogenic | -13.68 | Pathogenic | 0.82 | Disease | 1.00 | Probably damaging | 0 | Deleterious | | |
| p.Leu16Ala | VUS | -53.15 | Deleterious | 0.85 | Pathogenic | -11.52 | Pathogenic | 0.78 | Disease | 1.00 | Probably damaging | 0 | Deleterious | | |
| p.Leu16Gly | VUS | -53.15 | Deleterious | 0.95 | Pathogenic | -12.24 | Pathogenic | 0.81 | Disease | 1.00 | Probably damaging | 0 | Deleterious | | |
| p.Leu16Val | VUS | -14.21 | Indeterminate | 0.29 | Benign | -9.22 | Pathogenic | 0.61 | Disease | 0.98 | Probably damaging | 0 | Deleterious | 19.96 | Deleterious |
| p.Leu16Tyr | VUS | -53.15 | Deleterious | 0.97 | Pathogenic | -13.00 | Pathogenic | 0.75 | Disease | 0.92 | Possibly damaging | 0 | Deleterious | | |
| p.Leu16Cys | VUS | -16.70 | Indeterminate | 0.97 | Pathogenic | -12.75 | Pathogenic | 0.74 | Disease | 1.00 | Probably damaging | 0 | Deleterious | | |
| p.Leu16Trp | VUS | -53.15 | Deleterious | 0.95 | Pathogenic | -11.31 | Pathogenic | 0.81 | Disease | 0.78 | Possibly damaging | 0 | Deleterious | | |
| p.Leu16Phe | VUS | -4.61 | Neutral | 0.87 | Pathogenic | -9.34 | Pathogenic | 0.67 | Disease | 0.99 | Probably damaging | 0 | Deleterious | | |
| p.Ala17Asn | VUS | -9.84 | Indeterminate | 0.69 | Pathogenic | -9.33 | Pathogenic | 0.50 | Neutral | 0.74 | Possibly damaging | 0.31 | Tolerated | | |
| p.Ala17Lys | VUS | -53.15 | Deleterious | 0.93 | Pathogenic | -14.13 | Pathogenic | 0.55 | Disease | 0.99 | Probably damaging | 0.34 | Tolerated | | |
| p.Ala17Thr | VUS | -4.30 | Neutral | 0.33 | Benign | -5.99 | Benign | 0.20 | Neutral | 0.04 | Benign | 1 | Tolerated | 8.81 | Neutral |
| p.Ala17Arg | VUS | -53.15 | Deleterious | 0.77 | Pathogenic | -7.44 | Benign | 0.52 | Disease | 0.99 | Probably damaging | 0.26 | Tolerated | | |
| p.Ala17Ser | VUS | -1.96 | Neutral | 0.15 | Benign | -1.74 | Benign | 0.20 | Neutral | 0.01 | Benign | 0.86 | Tolerated | 7.00 | Neutral |
| p.Ala17Ile | VUS | -9.63 | Indeterminate | 0.73 | Pathogenic | -11.13 | Pathogenic | 0.48 | Neutral | 0.99 | Probably damaging | 0.24 | Tolerated | | |
| p.Ala17Met | VUS | -53.15 | Deleterious | 0.64 | Pathogenic | -10.49 | Pathogenic | 0.44 | Neutral | 1.00 | Probably damaging | 0.12 | Tolerated | | |
| p.Ala17His | VUS | -2.96 | Neutral | 0.64 | Pathogenic | -9.79 | Pathogenic | 0.51 | Disease | 1.00 | Probably damaging | 0.3 | Tolerated | | |
| p.Ala17Gln | VUS | -53.15 | Deleterious | 0.64 | Pathogenic | -10.72 | Pathogenic | 0.50 | Neutral | 1.00 | Probably damaging | 0.25 | Tolerated | | |
| p.Ala17Pro | VUS | -1.63 | Neutral | 0.70 | Pathogenic | -8.27 | Pathogenic | 0.59 | Disease | 0.97 | Probably damaging | 0.22 | Tolerated | 18.05 | Deleterious |
| p.Ala17Leu | VUS | -53.15 | Deleterious | 0.46 | Ambiguous | -8.16 | Pathogenic | 0.47 | Neutral | 0.93 | Possibly damaging | 0.44 | Tolerated | | |
| p.Ala17Asp | VUS | -33.22 | Indeterminate | 0.83 | Pathogenic | -11.49 | Pathogenic | 0.62 | Disease | 0.95 | Possibly damaging | 0.25 | Tolerated | 19.75 | Deleterious |
| p.Ala17Glu | VUS | -53.15 | Deleterious | 0.72 | Pathogenic | -11.65 | Pathogenic | 0.49 | Neutral | 0.97 | Probably damaging | 0.31 | Tolerated | | |
| p.Ala17Gly | VUS | -0.70 | Neutral | 0.23 | Benign | -3.66 | Benign | 0.28 | Neutral | 0.02 | Benign | 0.28 | Tolerated | 12.08 | Neutral |
| p.Ala17Val | VUS | -1.04 | Neutral | 0.40 | Ambiguous | -8.46 | Pathogenic | 0.42 | Neutral | 0.88 | Possibly damaging | 0.37 | Tolerated | 18.67 | Deleterious |
| p.Ala17Tyr | VUS | -53.15 | Deleterious | 0.71 | Pathogenic | -8.94 | Pathogenic | 0.49 | Neutral | 1.00 | Probably damaging | 0.35 | Tolerated | | |
| p.Ala17Cys | VUS | -0.41 | Neutral | 0.44 | Ambiguous | -0.72 | Benign | 0.29 | Neutral | 0.10 | Benign | 0.11 | Tolerated | | |
| p.Ala17Trp | VUS | -53.15 | Deleterious | 0.79 | Pathogenic | -8.74 | Pathogenic | 0.54 | Disease | 1.00 | Probably damaging | 0.07 | Tolerated | | |
| p.Ala17Phe | VUS | -53.15 | Deleterious | 0.56 | Ambiguous | -8.36 | Pathogenic | 0.47 | Neutral | 1.00 | Probably damaging | 0.26 | Tolerated | | |
| p.Thr18Asn | VUS | -33.21 | Indeterminate | 0.18 | Benign | -4.86 | Benign | 0.19 | Neutral | 0.12 | Benign | 0.54 | Tolerated | | |
| p.Thr18Lys | VUS | -14.00 | Indeterminate | 0.24 | Benign | -3.90 | Benign | 0.32 | Neutral | 0.76 | Possibly damaging | 0.86 | Tolerated | 11.22 | Neutral |
| p.Thr18Arg | VUS | -12.80 | Indeterminate | 0.16 | Benign | -1.73 | Benign | 0.25 | Neutral | 0.05 | Benign | 0.52 | Tolerated | 10.25 | Neutral |
| p.Thr18Ser | VUS | -5.57 | Neutral | 0.14 | Benign | -2.82 | Benign | 0.09 | Neutral | 0.01 | Benign | 0.88 | Tolerated | 0.39 | Neutral |
| p.Thr18Ile | VUS | -28.46 | Indeterminate | 0.33 | Benign | -5.32 | Benign | 0.25 | Neutral | 0.92 | Possibly damaging | 0.18 | Tolerated | | |
| p.Thr18Met | VUS | -20.84 | Indeterminate | 0.19 | Benign | -5.46 | Benign | 0.25 | Neutral | 1.00 | Probably damaging | 0.1 | Tolerated | 18.95 | Deleterious |

| | | | | | | | | | | | | | | | |
|---|---|---|---|---|---|---|---|---|---|---|---|---|---|---|---|
| p.Thr18His | VUS | -10.39 | Indeterminate | 0.27 | Benign | -4.59 | Benign | 0.33 | Neutral | 0.99 | Probably damaging | 0.13 | Tolerated | | |
| p.Thr18Gln | VUS | -10.00 | Indeterminate | 0.23 | Benign | -2.84 | Benign | 0.25 | Neutral | 0.84 | Possibly damaging | 0.55 | Tolerated | | |
| p.Thr18Pro | VUS | -53.15 | Deleterious | 0.32 | Benign | -5.58 | Benign | 0.31 | Neutral | 1.00 | Probably damaging | 0.27 | Tolerated | 16.16 | Deleterious |
| p.Thr18Leu | VUS | -29.34 | Indeterminate | 0.17 | Benign | -3.26 | Benign | 0.31 | Neutral | 0.62 | Possibly damaging | 0.28 | Tolerated | | |
| p.Thr18Asp | VUS | -12.03 | Indeterminate | 0.35 | Ambiguous | -0.56 | Benign | 0.30 | Neutral | 0.83 | Possibly damaging | 0.63 | Tolerated | | |
| p.Thr18Glu | VUS | -6.41 | Indeterminate | 0.27 | Benign | 0.00 | Benign | 0.34 | Neutral | 0.03 | Benign | 1 | Tolerated | | |
| p.Thr18Ala | VUS | -32.50 | Indeterminate | 0.09 | Benign | -1.41 | Benign | 0.11 | Neutral | 0.00 | Benign | 0.71 | Tolerated | 4.84 | Neutral |
| p.Thr18Gly | VUS | -10.65 | Indeterminate | 0.18 | Benign | -3.12 | Benign | 0.34 | Neutral | 0.99 | Probably damaging | 0.44 | Tolerated | | |
| p.Thr18Val | VUS | -4.77 | Neutral | 0.20 | Benign | -3.12 | Benign | 0.28 | Neutral | 0.54 | Possibly damaging | 0.27 | Tolerated | | |
| p.Thr18Tyr | VUS | -33.22 | Indeterminate | 0.37 | Ambiguous | -7.10 | Benign | 0.32 | Neutral | 1.00 | Probably damaging | 0.05 | Deleterious | | |
| p.Thr18Cys | VUS | -1.62 | Neutral | 0.52 | Ambiguous | -4.88 | Benign | 0.34 | Neutral | 1.00 | Probably damaging | 0.06 | Tolerated | | |
| p.Thr18Trp | VUS | -11.94 | Indeterminate | 0.56 | Ambiguous | -5.04 | Benign | 0.36 | Neutral | 1.00 | Probably damaging | 0.02 | Deleterious | | |
| p.Thr18Phe | VUS | -16.79 | Indeterminate | 0.35 | Ambiguous | -6.88 | Benign | 0.32 | Neutral | 0.98 | Probably damaging | 0.05 | Deleterious | | |
| p.Ala19Asn | VUS | 0.00 | Neutral | 0.92 | Pathogenic | -10.76 | Pathogenic | 0.66 | Disease | 1.00 | Probably damaging | 0 | Deleterious | | |
| p.Ala19Lys | VUS | -2.42 | Neutral | 0.97 | Pathogenic | -11.61 | Pathogenic | 0.68 | Disease | 1.00 | Probably damaging | 0 | Deleterious | | |
| p.Ala19Thr | VUS | 0.00 | Neutral | 0.62 | Pathogenic | -8.56 | Pathogenic | 0.40 | Neutral | 1.00 | Probably damaging | 0 | Deleterious | 24.90 | Deleterious |
| p.Ala19Arg | VUS | -0.43 | Neutral | 0.90 | Pathogenic | -9.41 | Pathogenic | 0.69 | Disease | 1.00 | Probably damaging | 0 | Deleterious | | |
| p.Ala19Ser | VUS | 0.00 | Neutral | 0.30 | Benign | -7.07 | Benign | 0.63 | Disease | 0.99 | Probably damaging | 0 | Deleterious | 24.40 | Deleterious |
| p.Ala19Ile | VUS | 0.00 | Neutral | 0.93 | Pathogenic | -10.54 | Pathogenic | 0.71 | Disease | 1.00 | Probably damaging | 0 | Deleterious | | |
| p.Ala19Met | VUS | 0.00 | Neutral | 0.86 | Pathogenic | -9.74 | Pathogenic | 0.68 | Disease | 1.00 | Probably damaging | 0 | Deleterious | | |
| p.Ala19His | VUS | 0.00 | Neutral | 0.92 | Pathogenic | -9.09 | Pathogenic | 0.70 | Disease | 1.00 | Probably damaging | 0 | Deleterious | | |
| p.Ala19Gln | VUS | 0.00 | Neutral | 0.84 | Pathogenic | -8.78 | Pathogenic | 0.70 | Disease | 1.00 | Probably damaging | 0 | Deleterious | | |
| p.Ala19Pro | VUS | -53.15 | Deleterious | 0.94 | Pathogenic | -9.95 | Pathogenic | 0.81 | Disease | 1.00 | Probably damaging | 0 | Deleterious | 25.00 | Deleterious |
| p.Ala19Leu | VUS | 0.00 | Neutral | 0.71 | Pathogenic | -6.76 | Benign | 0.70 | Disease | 1.00 | Probably damaging | 0 | Deleterious | | |
| p.Ala19Asp | VUS | -2.14 | Neutral | 0.94 | Pathogenic | -9.76 | Pathogenic | 0.71 | Disease | 0.99 | Probably damaging | 0 | Deleterious | 24.40 | Deleterious |
| p.Ala19Glu | VUS | 0.00 | Neutral | 0.84 | Pathogenic | -7.55 | Pathogenic | 0.75 | Disease | 1.00 | Probably damaging | 0 | Deleterious | | |
| p.Ala19Gly | VUS | 0.00 | Neutral | 0.43 | Ambiguous | -8.65 | Pathogenic | 0.44 | Neutral | 1.00 | Probably damaging | 0 | Deleterious | 24.30 | Deleterious |
| p.Ala19Val | VUS | 0.00 | Neutral | 0.65 | Pathogenic | -7.64 | Pathogenic | 0.42 | Neutral | 1.00 | Probably damaging | 0 | Deleterious | 24.40 | Deleterious |
| p.Ala19Tyr | VUS | 0.00 | Neutral | 0.94 | Pathogenic | -10.25 | Pathogenic | 0.70 | Disease | 1.00 | Probably damaging | 0 | Deleterious | | |
| p.Ala19Cys | VUS | -1.56 | Neutral | 0.76 | Pathogenic | -8.25 | Pathogenic | 0.65 | Disease | 1.00 | Probably damaging | 0 | Deleterious | | |
| p.Ala19Trp | VUS | 0.00 | Neutral | 0.98 | Pathogenic | -10.40 | Pathogenic | 0.69 | Disease | 1.00 | Probably damaging | 0 | Deleterious | | |
| p.Ala19Phe | VUS | 0.00 | Neutral | 0.93 | Pathogenic | -9.57 | Pathogenic | 0.71 | Disease | 1.00 | Probably damaging | 0 | Deleterious | | |
| p.Ala20Asn | VUS | -53.15 | Deleterious | 0.99 | Pathogenic | -20.00 | Pathogenic | 0.64 | Disease | 0.96 | Probably damaging | 0.02 | Deleterious | | |
| p.Ala20Lys | VUS | -53.15 | Deleterious | 0.99 | Pathogenic | -21.46 | Pathogenic | 0.68 | Disease | 1.00 | Probably damaging | 0.02 | Deleterious | | |
| p.Ala20Thr | VUS | -21.91 | Indeterminate | 0.50 | Ambiguous | -8.44 | Pathogenic | 0.62 | Disease | 1.00 | Probably damaging | 0.21 | Tolerated | 22.50 | Deleterious |
| p.Ala20Arg | VUS | -53.15 | Deleterious | 0.97 | Pathogenic | -18.12 | Pathogenic | 0.69 | Disease | 1.00 | Probably damaging | 0.06 | Tolerated | | |
| p.Ala20Ser | VUS | -24.89 | Indeterminate | 0.31 | Benign | -7.00 | Benign | 0.37 | Neutral | 0.20 | Benign | 0.26 | Tolerated | 14.86 | Neutral |
| p.Ala20Ile | VUS | -53.15 | Deleterious | 0.86 | Pathogenic | -14.21 | Pathogenic | 0.69 | Disease | 1.00 | Probably damaging | 0.01 | Deleterious | | |
| p.Ala20Met | VUS | -53.15 | Deleterious | 0.92 | Pathogenic | -17.29 | Pathogenic | 0.64 | Disease | 1.00 | Probably damaging | 0.01 | Deleterious | | |
| p.Ala20His | VUS | -53.15 | Deleterious | 0.99 | Pathogenic | -21.47 | Pathogenic | 0.68 | Disease | 1.00 | Probably damaging | 0 | Deleterious | | |
| p.Ala20Gln | VUS | -53.15 | Deleterious | 0.96 | Pathogenic | -19.57 | Pathogenic | 0.68 | Disease | 1.00 | Probably damaging | 0.01 | Deleterious | | |
| p.Ala20Pro | VUS | -53.15 | Deleterious | 0.95 | Pathogenic | -15.03 | Pathogenic | 0.77 | Disease | 1.00 | Probably damaging | 0.02 | Deleterious | 22.70 | Deleterious |
| p.Ala20Leu | VUS | -53.15 | Deleterious | 0.81 | Pathogenic | -16.05 | Pathogenic | 0.68 | Disease | 0.99 | Probably damaging | 0.01 | Deleterious | | |
| p.Ala20Asp | VUS | -53.15 | Deleterious | 0.99 | Pathogenic | -19.89 | Pathogenic | 0.73 | Disease | 1.00 | Probably damaging | 0.01 | Deleterious | | |
| p.Ala20Glu | VUS | -53.15 | Deleterious | 0.98 | Pathogenic | -18.55 | Pathogenic | 0.72 | Disease | 1.00 | Probably damaging | 0.01 | Deleterious | 22.90 | Deleterious |
| p.Ala20Gly | VUS | -53.15 | Deleterious | 0.54 | Ambiguous | -11.28 | Pathogenic | 0.55 | Disease | 0.99 | Probably damaging | 0.03 | Deleterious | 22.90 | Deleterious |
| p.Ala20Val | VUS | -9.53 | Indeterminate | 0.50 | Ambiguous | -9.61 | Pathogenic | 0.62 | Disease | 0.85 | Possibly damaging | 0.02 | Deleterious | 20.60 | Deleterious |
| p.Ala20Tyr | VUS | -53.15 | Deleterious | 0.99 | Pathogenic | -19.34 | Pathogenic | 0.68 | Disease | 1.00 | Probably damaging | 0 | Deleterious | | |
| p.Ala20Cys | VUS | -5.52 | Neutral | 0.74 | Pathogenic | -8.55 | Pathogenic | 0.59 | Disease | 1.00 | Probably damaging | 0.01 | Deleterious | | |
| p.Ala20Trp | VUS | -53.15 | Deleterious | 1.00 | Pathogenic | -18.45 | Pathogenic | 0.69 | Disease | 1.00 | Probably damaging | 0 | Deleterious | | |
| p.Ala20Phe | VUS | -53.15 | Deleterious | 0.98 | Pathogenic | -19.05 | Pathogenic | 0.69 | Disease | 1.00 | Probably damaging | 0 | Deleterious | | |
| p.Ala21Asn | VUS | -53.15 | Deleterious | 0.8048 | Pathogenic | -13.28 | Pathogenic | 0.63 | Disease | 0.99 | Probably damaging | 0.02 | Deleterious | | |
| p.Ala21Lys | VUS | -53.15 | Deleterious | 0.8769 | Pathogenic | -13.50 | Pathogenic | 0.64 | Disease | 1.00 | Probably damaging | 0.03 | Deleterious | | |
| p.Ala21Thr | VUS | -9.18 | Indeterminate | 0.4417 | Ambiguous | -9.89 | Pathogenic | 0.53 | Disease | 1.00 | Probably damaging | 0.06 | Tolerated | 23.30 | Deleterious |
| p.Ala21Arg | VUS | -53.15 | Deleterious | 0.6339 | Pathogenic | -11.05 | Pathogenic | 0.61 | Disease | 1.00 | Probably damaging | 0.02 | Deleterious | | |
| p.Ala21Ser | VUS | -2.09 | Neutral | 0.1823 | Benign | -6.29 | Benign | 0.52 | Disease | 0.52 | Possibly damaging | 0.22 | Tolerated | 21.10 | Deleterious |
| p.Ala21Ile | VUS | -53.15 | Deleterious | 0.74 | Pathogenic | -13.32 | Pathogenic | 0.64 | Disease | 1.00 | Probably damaging | 0.03 | Deleterious | | |
| p.Ala21Met | VUS | -53.15 | Deleterious | 0.6696 | Pathogenic | -11.97 | Pathogenic | 0.63 | Disease | 1.00 | Probably damaging | 0.02 | Deleterious | | |
| p.Ala21His | VUS | -53.15 | Deleterious | 0.80 | Pathogenic | -13.25 | Pathogenic | 0.63 | Disease | 1.00 | Probably damaging | 0.01 | Deleterious | | |
| p.Ala21Gln | VUS | -53.15 | Deleterious | 0.5501 | Ambiguous | -11.63 | Pathogenic | 0.62 | Disease | 1.00 | Probably damaging | 0.02 | Deleterious | | |
| p.Ala21Pro | VUS | -53.15 | Deleterious | 0.9341 | Pathogenic | -14.22 | Pathogenic | 0.67 | Disease | 1.00 | Probably damaging | 0.06 | Tolerated | 23.50 | Deleterious |
| p.Ala21Leu | VUS | -5.31 | Neutral | 0.5239 | Ambiguous | -11.90 | Pathogenic | 0.63 | Disease | 1.00 | Probably damaging | 0.18 | Tolerated | | |
| p.Ala21Asp | VUS | -53.15 | Deleterious | 0.91 | Pathogenic | -11.71 | Pathogenic | 0.74 | Disease | 1.00 | Probably damaging | 0.02 | Deleterious | 25.20 | Deleterious |
| p.Ala21Glu | VUS | -53.15 | Deleterious | 0.71 | Pathogenic | -9.30 | Pathogenic | 0.67 | Disease | 1.00 | Probably damaging | 0.03 | Deleterious | | |
| p.Ala21Gly | VUS | -0.52 | Neutral | 0.25 | Benign | -0.84 | Benign | 0.51 | Disease | 0.99 | Probably damaging | 0.13 | Tolerated | 24.60 | Deleterious |
| p.Ala21Val | VUS | -1.47 | Neutral | 0.4395 | Ambiguous | -10.01 | Pathogenic | 0.46 | Neutral | 1.00 | Probably damaging | 0.06 | Tolerated | 25.00 | Deleterious |
| p.Ala21Tyr | VUS | -53.15 | Deleterious | 0.7762 | Pathogenic | -11.79 | Pathogenic | 0.66 | Disease | 1.00 | Probably damaging | 0.01 | Deleterious | | |
| p.Ala21Cys | VUS | -0.74 | Neutral | 0.63 | Pathogenic | -8.76 | Pathogenic | 0.59 | Disease | 1.00 | Probably damaging | 0.03 | Deleterious | | |
| p.Ala21Trp | VUS | -53.15 | Deleterious | 0.8406 | Pathogenic | -11.16 | Pathogenic | 0.67 | Disease | 1.00 | Probably damaging | 0 | Deleterious | | |
| p.Ala21Phe | VUS | -53.15 | Deleterious | 0.70 | Pathogenic | -12.11 | Pathogenic | 0.67 | Disease | 1.00 | Probably damaging | 0.01 | Deleterious | | |
| p.Arg22Asn | VUS | 0.00 | Neutral | 0.5974 | Pathogenic | -6.51 | Benign | 0.44 | Neutral | 0.99 | Probably damaging | 0.07 | Tolerated | | |
| p.Arg22Lys | VUS | -1.44 | Neutral | 0.2087 | Benign | -6.50 | Benign | 0.16 | Neutral | 0.28 | Benign | 0.25 | Tolerated | | |
| p.Arg22Thr | VUS | -27.66 | Indeterminate | 0.3244 | Benign | -7.75 | Pathogenic | 0.28 | Neutral | 0.77 | Possibly damaging | 0.09 | Tolerated | | |
| p.Arg22Ser | VUS | -23.87 | Indeterminate | 0.4527 | Ambiguous | -6.49 | Benign | 0.36 | Neutral | 0.99 | Probably damaging | 0.23 | Tolerated | | |
| p.Arg22Ile | VUS | -0.98 | Neutral | 0.4942 | Ambiguous | -11.18 | Pathogenic | 0.45 | Neutral | 1.00 | Probably damaging | 0.03 | Deleterious | | |
| p.Arg22Met | VUS | -8.42 | Indeterminate | 0.4456 | Ambiguous | -9.14 | Pathogenic | 0.47 | Neutral | 0.98 | Probably damaging | 0.02 | Deleterious | | |
| p.Arg22His | VUS | 0.00 | Neutral | 0.16 | Benign | -4.40 | Benign | 0.32 | Neutral | 0.99 | Probably damaging | 0.04 | Deleterious | | |
| p.Arg22Gln | VUS | 0.00 | Neutral | 0.1157 | Benign | -5.42 | Benign | 0.16 | Neutral | 0.78 | Possibly damaging | 0.15 | Tolerated | 12.71 | Neutral |
| p.Arg22Pro | VUS | -53.15 | Deleterious | 0.8905 | Pathogenic | -11.55 | Pathogenic | 0.54 | Disease | 1.00 | Probably damaging | 0.05 | Deleterious | 20.40 | Deleterious |
| p.Arg22Leu | VUS | -0.02 | Neutral | 0.2206 | Benign | -6.54 | Benign | 0.22 | Neutral | 0.51 | Possibly damaging | 0.1 | Tolerated | 12.51 | Neutral |
| p.Arg22Asp | VUS | -6.69 | Indeterminate | 0.6151 | Pathogenic | -6.66 | Benign | 0.47 | Neutral | 1.00 | Probably damaging | 0.05 | Deleterious | | |
| p.Arg22Glu | VUS | 0.00 | Neutral | 0.2995 | Benign | -6.72 | Benign | 0.37 | Neutral | 0.84 | Possibly damaging | 0.11 | Tolerated | | |
| p.Arg22Ala | VUS | -0.01 | Neutral | 0.2965 | Benign | -5.51 | Benign | 0.41 | Neutral | 0.95 | Possibly damaging | 0.22 | Tolerated | | |
| p.Arg22Gly | VUS | -53.15 | Deleterious | 0.2016 | Benign | -6.85 | Benign | 0.25 | Neutral | 0.47 | Possibly damaging | 0.06 | Tolerated | 16.11 | Deleterious |
| p.Arg22Val | VUS | -0.16 | Neutral | 0.4546 | Ambiguous | -8.96 | Pathogenic | 0.45 | Neutral | 0.99 | Probably damaging | 0.05 | Deleterious | | |
| p.Arg22Tyr | VUS | -3.79 | Neutral | 0.4817 | Ambiguous | -6.03 | Benign | 0.41 | Neutral | 0.94 | Possibly damaging | 0.03 | Deleterious | | |
| p.Arg22Cys | VUS | 0.00 | Neutral | 0.2916 | Benign | -6.74 | Benign | 0.27 | Neutral | 1.00 | Probably damaging | 0.01 | Deleterious | | |
| p.Arg22Trp | VUS | -9.64 | Indeterminate | 0.2423 | Benign | -8.27 | Pathogenic | 0.53 | Disease | 1.00 | Probably damaging | 0.01 | Deleterious | 23.50 | Deleterious |
| p.Arg22Phe | VUS | -32.64 | Indeterminate | 0.5572 | Ambiguous | -7.96 | Pathogenic | 0.45 | Neutral | 0.89 | Possibly damaging | 0.02 | Deleterious | | |
| p.Gly23Asn | VUS | -21.58 | Indeterminate | 0.88 | Pathogenic | -9.28 | Pathogenic | 0.66 | Disease | 1.00 | Probably damaging | 0.21 | Tolerated | | |
| p.Gly23Lys | VUS | -53.15 | Deleterious | 0.9761 | Pathogenic | -13.72 | Pathogenic | 0.70 | Disease | 1.00 | Probably damaging | 0.01 | Deleterious | | |
| p.Gly23Thr | VUS | -53.15 | Deleterious | 0.9417 | Pathogenic | -11.02 | Pathogenic | 0.64 | Disease | 1.00 | Probably damaging | 0.01 | Deleterious | | |
| p.Gly23Arg | Likely pathogenic | -53.15 | Deleterious | 0.9175 | Pathogenic | -11.14 | Pathogenic | 0.49 | Neutral | 1.00 | Probably damaging | 0.01 | Deleterious | 27.20 | Deleterious |
| p.Gly23Ser | Likely pathogenic | -53.15 | Deleterious | 0.6877 | Pathogenic | -8.75 | Pathogenic | 0.67 | Disease | 0.99 | Probably damaging | 0.03 | Deleterious | 26.90 | Deleterious |
| p.Gly23Ile | VUS | -53.15 | Deleterious | 0.9805 | Pathogenic | -13.32 | Pathogenic | 0.71 | Disease | 1.00 | Probably damaging | 0 | Deleterious | | |
| p.Gly23Met | VUS | -53.15 | Deleterious | 0.9793 | Pathogenic | -13.18 | Pathogenic | 0.71 | Disease | 1.00 | Probably damaging | 0 | Deleterious | | |
| p.Gly23His | VUS | -53.15 | Deleterious | 0.9519 | Pathogenic | -12.10 | Pathogenic | 0.68 | Disease | 1.00 | Probably damaging | 0.01 | Deleterious | | |
| p.Gly23Gln | VUS | -53.15 | Deleterious | 0.9087 | Pathogenic | -11.13 | Pathogenic | 0.65 | Disease | 1.00 | Probably damaging | 0.01 | Deleterious | | |
| p.Gly23Pro | VUS | -53.15 | Deleterious | 0.9957 | Pathogenic | -12.38 | Pathogenic | 0.47 | Neutral | 1.00 | Probably damaging | 0.01 | Deleterious | | |
| p.Gly23Leu | VUS | -53.15 | Deleterious | 0.9462 | Pathogenic | -11.12 | Pathogenic | 0.70 | Disease | 1.00 | Probably damaging | 0 | Deleterious | | |
| p.Gly23Asp | Pathogenic | -53.15 | Deleterious | 0.8537 | Pathogenic | -9.16 | Pathogenic | 0.83 | Disease | 1.00 | Probably damaging | 0.04 | Deleterious | 27.70 | Deleterious |
| p.Gly23Glu | VUS | -53.15 | Deleterious | 0.9158 | Pathogenic | -9.00 | Pathogenic | 0.78 | Disease | 1.00 | Probably damaging | 0.01 | Deleterious | | |
| p.Gly23Ala | VUS | -1.29 | Neutral | 0.7359 | Pathogenic | -7.21 | Benign | 0.65 | Disease | 1.00 | Probably damaging | 0.02 | Deleterious | 26.10 | Deleterious |
| p.Gly23Val | VUS | -53.15 | Deleterious | 0.955 | Pathogenic | -10.22 | Pathogenic | 0.81 | Disease | 1.00 | Probably damaging | 0 | Deleterious | 27.30 | Deleterious |
| p.Gly23Tyr | VUS | -53.15 | Deleterious | 0.9386 | Pathogenic | -15.29 | Pathogenic | 0.71 | Disease | 1.00 | Probably damaging | 0 | Deleterious | | |
| p.Gly23Cys | VUS | -53.15 | Deleterious | 0.8139 | Pathogenic | -9.87 | Pathogenic | 0.84 | Disease | 1.00 | Probably damaging | 0 | Deleterious | 28.10 | Deleterious |
| p.Gly23Trp | VUS | -53.15 | Deleterious | 0.9369 | Pathogenic | -13.94 | Pathogenic | 0.76 | Disease | 1.00 | Probably damaging | 0 | Deleterious | | |
| p.Gly23Phe | VUS | -53.15 | Deleterious | 0.9795 | Pathogenic | -14.73 | Pathogenic | 0.73 | Disease | 1.00 | Probably damaging | 0 | Deleterious | | |
| p.Arg24Asn | VUS | -0.62 | Neutral | 0.4681 | Ambiguous | -4.43 | Benign | 0.22 | Neutral | 0.85 | Possibly damaging | 0.43 | Tolerated | | |
| p.Arg24Lys | VUS | -1.72 | Neutral | 0.2416 | Benign | -5.53 | Benign | 0.23 | Neutral | 0.62 | Possibly damaging | 1 | Tolerated | | |
| p.Arg24Thr | VUS | -3.98 | Neutral | 0.4732 | Ambiguous | -5.85 | Benign | 0.39 | Neutral | 0.99 | Probably damaging | 0.42 | Tolerated | | |

| | | | | | | | | | | | | | | | |
|---|---|---|---|---|---|---|---|---|---|---|---|---|---|---|---|
| p.Arg24Ser | VUS | -0.21 | Neutral | 0.4235 | Ambiguous | -3.88 | Benign | 0.38 | Neutral | 1.00 | Probably damaging | 0.49 | Tolerated | | |
| p.Arg24Ile | VUS | -0.34 | Neutral | 0.6604 | Pathogenic | -9.38 | Pathogenic | 0.48 | Neutral | 1.00 | Probably damaging | 0.19 | Tolerated | | |
| p.Arg24Met | VUS | -0.60 | Neutral | 0.5378 | Ambiguous | -8.07 | Pathogenic | 0.46 | Neutral | 0.68 | Possibly damaging | 0.11 | Tolerated | | |
| p.Arg24His | VUS | -0.01 | Neutral | 0.1286 | Benign | -5.36 | Benign | 0.11 | Neutral | 0.99 | Probably damaging | 0.27 | Tolerated | | |
| p.Arg24Gln | VUS | -16.21 | Indeterminate | 0.12 | Benign | -4.47 | Benign | 0.08 | Neutral | 0.73 | Possibly damaging | 0.61 | Tolerated | 18.45 | Deleterious |
| p.Arg24Pro | Pathogenic | -53.15 | Deleterious | 0.9136 | Pathogenic | -8.08 | Pathogenic | 0.63 | Disease | 1.00 | Probably damaging | 0.24 | Tolerated | 22.80 | Deleterious |
| p.Arg24Leu | VUS | -1.07 | Neutral | 0.2995 | Benign | -5.72 | Benign | 0.13 | Neutral | 1.00 | Probably damaging | 0.35 | Tolerated | 19.08 | Deleterious |
| p.Arg24Asp | VUS | 0.00 | Neutral | 0.4363 | Ambiguous | -1.34 | Benign | 0.19 | Neutral | 0.79 | Possibly damaging | 0.64 | Tolerated | | |
| p.Arg24Glu | VUS | -1.01 | Neutral | 0.3064 | Benign | -2.84 | Benign | 0.20 | Neutral | 0.80 | Possibly damaging | 0.8 | Tolerated | | |
| p.Arg24Ala | VUS | -0.03 | Neutral | 0.3784 | Ambiguous | -3.76 | Benign | 0.39 | Neutral | 0.99 | Probably damaging | 0.53 | Tolerated | | |
| p.Arg24Gly | VUS | -2.96 | Neutral | 0.2927 | Benign | -5.24 | Benign | 0.14 | Neutral | 1.00 | Probably damaging | 0.33 | Tolerated | 23.00 | Deleterious |
| p.Arg24Val | VUS | -0.12 | Neutral | 0.5914 | Pathogenic | -6.67 | Benign | 0.48 | Neutral | 1.00 | Probably damaging | 0.26 | Tolerated | | |
| p.Arg24Tyr | VUS | -1.12 | Neutral | 0.5087 | Ambiguous | -8.91 | Pathogenic | 0.38 | Neutral | 0.76 | Possibly damaging | 0.24 | Tolerated | | |
| p.Arg24Cys | VUS | -0.09 | Neutral | 0.2899 | Benign | -5.58 | Benign | 0.17 | Neutral | 1.00 | Probably damaging | 0.07 | Tolerated | | |
| p.Arg24Trp | VUS | -0.43 | Neutral | 0.2888 | Benign | -8.64 | Pathogenic | 0.19 | Neutral | 0.32 | Benign | 0.06 | Tolerated | 22.70 | Deleterious |
| p.Arg24Phe | VUS | -0.09 | Neutral | 0.6672 | Pathogenic | -9.23 | Pathogenic | 0.46 | Neutral | 0.76 | Possibly damaging | 0.17 | Tolerated | | |
| p.Val25Asn | VUS | -4.24 | Neutral | 0.4525 | Ambiguous | -10.67 | Pathogenic | 0.30 | Neutral | 0.84 | Possibly damaging | 0.02 | Deleterious | | |
| p.Val25Lys | VUS | -17.81 | Indeterminate | 0.4628 | Ambiguous | -10.72 | Pathogenic | 0.29 | Neutral | 0.12 | Benign | 0.03 | Deleterious | | |
| p.Val25Thr | VUS | -1.42 | Neutral | 0.1379 | Benign | -5.74 | Benign | 0.16 | Neutral | 0.04 | Benign | 0.32 | Tolerated | | |
| p.Val25Arg | VUS | -3.28 | Neutral | 0.3039 | Benign | -8.25 | Pathogenic | 0.18 | Neutral | 0.38 | Benign | 0.02 | Deleterious | | |
| p.Val25Ser | VUS | -0.43 | Neutral | 0.2142 | Benign | -7.24 | Benign | 0.27 | Neutral | 0.80 | Possibly damaging | 0.03 | Deleterious | | |
| p.Val25Ile | VUS | -0.50 | Neutral | 0.1369 | Benign | -7.02 | Benign | 0.13 | Neutral | 0.01 | Benign | 0.39 | Tolerated | 14.12 | Neutral |
| p.Val25Met | VUS | -21.03 | Indeterminate | 0.3185 | Benign | -7.60 | Pathogenic | 0.16 | Neutral | 0.60 | Possibly damaging | 0.05 | Deleterious | | |
| p.Val25His | VUS | -0.26 | Neutral | 0.4947 | Ambiguous | -7.06 | Benign | 0.20 | Neutral | 0.90 | Possibly damaging | 0.01 | Deleterious | | |
| p.Val25Gln | VUS | -0.52 | Neutral | 0.3048 | Benign | -8.28 | Pathogenic | 0.27 | Neutral | 0.41 | Benign | 0.03 | Deleterious | | |
| p.Val25Pro | VUS | 0.00 | Neutral | 0.2132 | Benign | -6.08 | Benign | 0.29 | Neutral | 0.84 | Possibly damaging | 0.04 | Deleterious | | |
| p.Val25Leu | VUS | -0.06 | Neutral | 0.2415 | Benign | -3.55 | Benign | 0.12 | Neutral | 0.00 | Benign | 0.33 | Tolerated | 12.09 | Neutral |
| p.Val25Asp | VUS | -17.55 | Indeterminate | 0.2921 | Benign | -6.73 | Benign | 0.33 | Neutral | 0.96 | Probably damaging | 0.01 | Deleterious | | |
| p.Val25Glu | VUS | -0.18 | Neutral | 0.1475 | Benign | -3.45 | Benign | 0.16 | Neutral | 0.03 | Benign | 0.08 | Tolerated | 4.46 | Neutral |
| p.Val25Ala | VUS | -53.15 | Deleterious | 0.10 | Benign | -4.13 | Benign | 0.06 | Neutral | 0.01 | Benign | 0.38 | Tolerated | 0.34 | Neutral |
| p.Val25Gly | VUS | -19.94 | Indeterminate | 0.1571 | Benign | -7.74 | Pathogenic | 0.15 | Neutral | 0.91 | Possibly damaging | 0 | Deleterious | 5.28 | Neutral |
| p.Val25Tyr | VUS | -33.22 | Indeterminate | 0.5066 | Ambiguous | -6.86 | Benign | 0.27 | Neutral | 0.81 | Possibly damaging | 0.01 | Deleterious | | |
| p.Val25Cys | VUS | 0.00 | Neutral | 0.6914 | Pathogenic | -6.54 | Benign | 0.19 | Neutral | 0.93 | Possibly damaging | 0.04 | Deleterious | | |
| p.Val25Trp | VUS | -2.22 | Neutral | 0.6234 | Pathogenic | -7.79 | Pathogenic | 0.29 | Neutral | 0.99 | Probably damaging | 0 | Deleterious | | |
| p.Val25Phe | VUS | -3.13 | Neutral | 0.2408 | Benign | -7.96 | Pathogenic | 0.25 | Neutral | 0.68 | Possibly damaging | 0.03 | Deleterious | | |
| p.Glu26Asn | VUS | -1.41 | Neutral | 0.3554 | Ambiguous | -8.72 | Pathogenic | 0.27 | Neutral | 0.05 | Benign | 0.02 | Deleterious | | |
| p.Glu26Lys | VUS | -0.44 | Neutral | 0.3023 | Benign | -7.86 | Pathogenic | 0.19 | Neutral | 0.01 | Benign | 0.04 | Deleterious | 14.84 | Neutral |
| p.Glu26Thr | VUS | -9.37 | Indeterminate | 0.2484 | Benign | -6.86 | Benign | 0.29 | Neutral | 0.13 | Benign | 0.01 | Deleterious | | |
| p.Glu26Arg | VUS | -0.10 | Neutral | 0.2163 | Benign | -5.10 | Benign | 0.23 | Neutral | 0.00 | Benign | 0.02 | Deleterious | | |
| p.Glu26Ser | VUS | -27.38 | Indeterminate | 0.2275 | Benign | -5.96 | Benign | 0.31 | Neutral | 0.01 | Benign | 0.02 | Deleterious | | |
| p.Glu26Ile | VUS | -0.52 | Neutral | 0.5949 | Pathogenic | -10.16 | Pathogenic | 0.33 | Neutral | 0.11 | Benign | 0 | Deleterious | | |
| p.Glu26Met | VUS | -0.27 | Neutral | 0.6984 | Pathogenic | -9.89 | Pathogenic | 0.34 | Neutral | 0.02 | Benign | 0 | Deleterious | | |
| p.Glu26His | VUS | -4.15 | Neutral | 0.3936 | Ambiguous | -6.68 | Benign | 0.22 | Neutral | 0.00 | Benign | 0.21 | Tolerated | | |
| p.Glu26Gln | VUS | -2.58 | Neutral | 0.1534 | Benign | -5.86 | Benign | 0.11 | Neutral | 0.00 | Benign | 0.18 | Tolerated | 11.94 | Neutral |
| p.Glu26Pro | VUS | -0.39 | Neutral | 0.3451 | Ambiguous | -5.81 | Benign | 0.37 | Neutral | 0.21 | Benign | 0.02 | Deleterious | | |
| p.Glu26Leu | VUS | -0.53 | Neutral | 0.4372 | Ambiguous | -6.92 | Benign | 0.34 | Neutral | 0.02 | Benign | 0.01 | Deleterious | | |
| p.Glu26Asp | VUS | -3.00 | Neutral | 0.1188 | Benign | -3.77 | Benign | 0.11 | Neutral | 0.00 | Benign | 0.08 | Tolerated | 8.36 | Neutral |
| p.Glu26Ala | VUS | -1.96 | Neutral | 0.1106 | Benign | -4.01 | Benign | 0.12 | Neutral | 0.00 | Benign | 0.03 | Deleterious | 14.91 | Neutral |
| p.Glu26Gly | VUS | -4.39 | Neutral | 0.1345 | Benign | -5.02 | Benign | 0.21 | Neutral | 0.23 | Benign | 0.01 | Deleterious | 19.55 | Deleterious |
| p.Glu26Val | VUS | -0.20 | Neutral | 0.3096 | Benign | -6.90 | Benign | 0.28 | Neutral | 0.01 | Benign | 0.01 | Deleterious | 17.11 | Deleterious |
| p.Glu26Tyr | VUS | -5.42 | Neutral | 0.7028 | Pathogenic | -9.93 | Pathogenic | 0.32 | Neutral | 0.26 | Benign | 0 | Deleterious | | |
| p.Glu26Cys | VUS | -0.02 | Neutral | 0.8412 | Pathogenic | -8.04 | Pathogenic | 0.33 | Neutral | 1.00 | Probably damaging | 0 | Deleterious | | |
| p.Glu26Trp | VUS | -0.96 | Neutral | 0.8423 | Pathogenic | -8.77 | Pathogenic | 0.37 | Neutral | 0.89 | Possibly damaging | 0 | Deleterious | | |
| p.Glu26Phe | VUS | -3.59 | Neutral | 0.8086 | Pathogenic | -10.05 | Pathogenic | 0.33 | Neutral | 0.21 | Benign | 0 | Deleterious | | |
| p.Glu27Asn | VUS | -15.33 | Indeterminate | 0.8108 | Pathogenic | -8.82 | Pathogenic | 0.37 | Neutral | 0.97 | Probably damaging | 0.38 | Tolerated | | |
| p.Glu27Lys | VUS | -0.27 | Neutral | 0.6297 | Pathogenic | -7.32 | Benign | 0.18 | Neutral | 0.66 | Possibly damaging | 0.74 | Tolerated | 22.50 | Deleterious |
| p.Glu27Thr | VUS | -27.81 | Indeterminate | 0.582 | Pathogenic | -5.89 | Benign | 0.29 | Neutral | 0.25 | Benign | 0.47 | Tolerated | | |
| p.Glu27Arg | VUS | -6.08 | Indeterminate | 0.5915 | Pathogenic | -4.60 | Benign | 0.24 | Neutral | 0.90 | Possibly damaging | 0.4 | Tolerated | | |
| p.Glu27Ser | VUS | -4.04 | Neutral | 0.6781 | Pathogenic | -6.61 | Benign | 0.42 | Neutral | 0.91 | Possibly damaging | 0.56 | Tolerated | | |
| p.Glu27Ile | VUS | -2.89 | Neutral | 0.7775 | Pathogenic | -8.88 | Pathogenic | 0.44 | Neutral | 0.68 | Possibly damaging | 0.14 | Tolerated | | |
| p.Glu27Met | VUS | -2.28 | Neutral | 0.8549 | Pathogenic | -8.51 | Pathogenic | 0.35 | Neutral | 0.74 | Possibly damaging | 0.13 | Tolerated | | |
| p.Glu27His | VUS | -4.17 | Neutral | 0.7579 | Pathogenic | -6.22 | Benign | 0.40 | Neutral | 0.99 | Probably damaging | 0.1 | Tolerated | | |
| p.Glu27Gln | VUS | -3.17 | Neutral | 0.3021 | Benign | -4.07 | Benign | 0.13 | Neutral | 0.11 | Benign | 0.53 | Tolerated | 21.00 | Deleterious |
| p.Glu27Pro | VUS | -7.45 | Indeterminate | 0.9 | Pathogenic | -6.85 | Benign | 0.47 | Neutral | 0.95 | Possibly damaging | 0.21 | Tolerated | | |
| p.Glu27Leu | VUS | -2.14 | Neutral | 0.777 | Pathogenic | -4.75 | Benign | 0.31 | Neutral | 0.01 | Benign | 0.23 | Tolerated | | |
| p.Glu27Asp | VUS | -25.59 | Indeterminate | 0.3212 | Benign | -4.41 | Benign | 0.12 | Neutral | 0.01 | Benign | 0.51 | Tolerated | 21.80 | Deleterious |
| p.Glu27Ala | VUS | -3.09 | Neutral | 0.2891 | Benign | -3.86 | Benign | 0.21 | Neutral | 0.07 | Benign | 0.6 | Tolerated | 21.30 | Deleterious |
| p.Glu27Gly | VUS | -17.81 | Indeterminate | 0.6044 | Pathogenic | -6.56 | Benign | 0.24 | Neutral | 0.98 | Probably damaging | 0.29 | Tolerated | 23.10 | Deleterious |
| p.Glu27Val | VUS | -0.71 | Neutral | 0.5387 | Ambiguous | -5.68 | Benign | 0.43 | Neutral | 0.02 | Benign | 0.3 | Tolerated | 22.20 | Deleterious |
| p.Glu27Tyr | VUS | -53.15 | Deleterious | 0.9134 | Pathogenic | -10.09 | Pathogenic | 0.45 | Neutral | 0.98 | Probably damaging | 0.03 | Deleterious | | |
| p.Glu27Cys | VUS | -18.51 | Indeterminate | 0.9673 | Pathogenic | -8.40 | Pathogenic | 0.44 | Neutral | 1.00 | Probably damaging | 0.04 | Deleterious | | |
| p.Glu27Trp | VUS | -0.31 | Neutral | 0.9791 | Pathogenic | -9.54 | Pathogenic | 0.49 | Neutral | 1.00 | Probably damaging | 0.01 | Deleterious | | |
| p.Glu27Phe | VUS | -0.20 | Neutral | 0.9594 | Pathogenic | -10.44 | Pathogenic | 0.45 | Neutral | 0.90 | Possibly damaging | 0.04 | Deleterious | | |
| p.Val28Asn | VUS | -53.15 | Deleterious | 0.9413 | Pathogenic | -17.77 | Pathogenic | 0.75 | Disease | 1.00 | Probably damaging | 0 | Deleterious | | |
| p.Val28Lys | VUS | -53.15 | Deleterious | 0.9711 | Pathogenic | -19.40 | Pathogenic | 0.78 | Disease | 1.00 | Probably damaging | 0 | Deleterious | | |
| p.Val28Thr | VUS | -0.16 | Neutral | 0.5482 | Ambiguous | -7.25 | Benign | 0.66 | Disease | 1.00 | Probably damaging | 0.01 | Deleterious | | |
| p.Val28Arg | VUS | -53.15 | Deleterious | 0.9327 | Pathogenic | -15.40 | Pathogenic | 0.76 | Disease | 1.00 | Probably damaging | 0 | Deleterious | | |
| p.Val28Ser | VUS | -1.43 | Neutral | 0.8134 | Pathogenic | -10.22 | Pathogenic | 0.70 | Disease | 1.00 | Probably damaging | 0 | Deleterious | | |
| p.Val28Ile | VUS | -0.17 | Neutral | 0.2771 | Benign | -8.20 | Pathogenic | 0.44 | Neutral | 1.00 | Probably damaging | 0.05 | Deleterious | | |
| p.Val28Met | VUS | -0.29 | Neutral | 0.7568 | Pathogenic | -9.48 | Pathogenic | 0.62 | Disease | 1.00 | Probably damaging | 0.01 | Deleterious | 31.00 | Deleterious |
| p.Val28His | VUS | -53.15 | Deleterious | 0.9896 | Pathogenic | -16.15 | Pathogenic | 0.74 | Disease | 1.00 | Probably damaging | 0 | Deleterious | | |
| p.Val28Gln | VUS | -53.15 | Deleterious | 0.9319 | Pathogenic | -15.73 | Pathogenic | 0.76 | Disease | 1.00 | Probably damaging | 0 | Deleterious | | |
| p.Val28Pro | VUS | -53.15 | Deleterious | 0.9875 | Pathogenic | -12.89 | Pathogenic | 0.74 | Disease | 1.00 | Probably damaging | 0 | Deleterious | | |
| p.Val28Leu | VUS | 0.00 | Neutral | 0.7196 | Pathogenic | -7.49 | Benign | 0.43 | Neutral | 1.00 | Probably damaging | 0.2 | Tolerated | 25.95 | Deleterious |
| p.Val28Asp | VUS | -53.15 | Deleterious | 0.9632 | Pathogenic | -15.94 | Pathogenic | 0.81 | Disease | 1.00 | Probably damaging | 0 | Deleterious | | |
| p.Val28Glu | VUS | -53.15 | Deleterious | 0.9105 | Pathogenic | -14.49 | Pathogenic | 0.81 | Disease | 1.00 | Probably damaging | 0 | Deleterious | 32.00 | Deleterious |
| p.Val28Ala | VUS | -0.04 | Neutral | 0.4679 | Ambiguous | -5.32 | Benign | 0.56 | Disease | 1.00 | Probably damaging | 0.01 | Deleterious | 29.60 | Deleterious |
| p.Val28Gly | VUS | -20.39 | Indeterminate | 0.57 | Pathogenic | -11.97 | Pathogenic | 0.76 | Disease | 1.00 | Probably damaging | 0 | Deleterious | 32.00 | Deleterious |
| p.Val28Tyr | VUS | -53.15 | Deleterious | 0.9801 | Pathogenic | -16.38 | Pathogenic | 0.75 | Disease | 1.00 | Probably damaging | 0 | Deleterious | | |
| p.Val28Cys | VUS | -0.11 | Neutral | 0.9384 | Pathogenic | -5.74 | Benign | 0.68 | Disease | 1.00 | Probably damaging | 0 | Deleterious | | |
| p.Val28Trp | VUS | -53.15 | Deleterious | 0.9963 | Pathogenic | -15.93 | Pathogenic | 0.71 | Disease | 1.00 | Probably damaging | 0 | Deleterious | | |
| p.Val28Phe | VUS | -53.15 | Deleterious | 0.8948 | Pathogenic | -13.19 | Pathogenic | 0.77 | Disease | 1.00 | Probably damaging | 0.01 | Deleterious | | |
| p.Arg29Asn | VUS | -4.64 | Neutral | 0.7626 | Pathogenic | -10.66 | Pathogenic | 0.49 | Neutral | 0.98 | Probably damaging | 0.03 | Deleterious | | |
| p.Arg29Lys | VUS | -0.17 | Neutral | 0.191 | Benign | -7.10 | Benign | 0.20 | Neutral | 0.15 | Benign | 0.3 | Tolerated | | |
| p.Arg29Thr | VUS | -12.09 | Indeterminate | 0.2316 | Benign | -6.41 | Benign | 0.35 | Neutral | 0.55 | Possibly damaging | 0.26 | Tolerated | | |
| p.Arg29Ser | VUS | -0.79 | Neutral | 0.4961 | Ambiguous | -5.40 | Benign | 0.46 | Neutral | 0.99 | Probably damaging | 0.03 | Deleterious | | |
| p.Arg29Ile | VUS | -21.20 | Indeterminate | 0.335 | Benign | -10.02 | Pathogenic | 0.48 | Neutral | 1.00 | Probably damaging | 0.01 | Deleterious | | |
| p.Arg29Met | VUS | -1.63 | Neutral | 0.4175 | Ambiguous | -9.57 | Pathogenic | 0.47 | Neutral | 1.00 | Probably damaging | 0.01 | Deleterious | | |
| p.Arg29His | VUS | -3.49 | Neutral | 0.1851 | Benign | -8.02 | Pathogenic | 0.44 | Neutral | 0.98 | Probably damaging | 0.02 | Deleterious | | |
| p.Arg29Gln | VUS | -0.28 | Neutral | 0.1091 | Benign | -5.72 | Benign | 0.18 | Neutral | 0.79 | Possibly damaging | 0.06 | Tolerated | 22.70 | Deleterious |
| p.Arg29Pro | VUS | -53.15 | Deleterious | 0.929 | Pathogenic | -11.13 | Pathogenic | 0.59 | Disease | 1.00 | Probably damaging | 0.02 | Deleterious | 26.10 | Deleterious |
| p.Arg29Leu | VUS | -0.88 | Neutral | 0.1876 | Benign | -7.71 | Pathogenic | 0.28 | Neutral | 1.00 | Probably damaging | 0.03 | Deleterious | 25.30 | Deleterious |
| p.Arg29Asp | VUS | -18.67 | Indeterminate | 0.7182 | Pathogenic | -9.12 | Pathogenic | 0.54 | Disease | 0.99 | Probably damaging | 0.01 | Deleterious | | |
| p.Arg29Glu | VUS | -4.92 | Neutral | 0.2562 | Benign | -5.39 | Benign | 0.28 | Neutral | 0.31 | Benign | 0.04 | Deleterious | | |
| p.Arg29Ala | VUS | -0.01 | Neutral | 0.2363 | Benign | -2.97 | Benign | 0.45 | Neutral | 0.94 | Possibly damaging | 0.03 | Deleterious | | |
| p.Arg29Gly | VUS | -1.76 | Neutral | 0.2867 | Benign | -5.63 | Benign | 0.46 | Neutral | 0.81 | Possibly damaging | 0.02 | Deleterious | 25.40 | Deleterious |
| p.Arg29Val | VUS | -1.92 | Neutral | 0.2848 | Benign | -7.20 | Benign | 0.50 | Neutral | 1.00 | Probably damaging | 0.02 | Deleterious | | |
| p.Arg29Tyr | VUS | -0.27 | Neutral | 0.5312 | Ambiguous | -11.00 | Pathogenic | 0.45 | Neutral | 0.96 | Probably damaging | 0.01 | Deleterious | | |
| p.Arg29Cys | VUS | -0.99 | Neutral | 0.3189 | Benign | -7.40 | Benign | 0.47 | Neutral | 1.00 | Probably damaging | 0 | Deleterious | | |
| p.Arg29Trp | VUS | -0.91 | Neutral | 0.262 | Benign | -8.98 | Pathogenic | 0.46 | Neutral | 1.00 | Probably damaging | 0 | Deleterious | 32.00 | Deleterious |
| p.Arg29Phe | VUS | -3.19 | Neutral | 0.5847 | Pathogenic | -10.68 | Pathogenic | 0.48 | Neutral | 0.96 | Probably damaging | 0.01 | Deleterious | | |

| | | | | | | | | | | | | | | | |
|---|---|---|---|---|---|---|---|---|---|---|---|---|---|---|---|
| p.Ala30Asn | VUS | -53.15 | Deleterious | 0.2215 | Benign | -8.71 | Pathogenic | 0.35 | Neutral | 0.09 | Benign | 0.41 | Tolerated | | |
| p.Ala30Lys | VUS | -1.25 | Neutral | 0.2472 | Benign | -6.24 | Benign | 0.36 | Neutral | 0.60 | Possibly damaging | 0.5 | Tolerated | | |
| p.Ala30Thr | VUS | -23.96 | Indeterminate | 0.1066 | Benign | -6.59 | Benign | 0.24 | Neutral | 0.02 | Benign | 0.51 | Tolerated | 20.30 | Deleterious |
| p.Ala30Arg | VUS | -3.40 | Neutral | 0.1463 | Benign | -3.24 | Benign | 0.34 | Neutral | 0.84 | Possibly damaging | 0.47 | Tolerated | | |
| p.Ala30Ser | VUS | -11.06 | Indeterminate | 0.1102 | Benign | -5.07 | Benign | 0.26 | Neutral | 0.00 | Benign | 0.56 | Tolerated | 18.06 | Deleterious |
| p.Ala30Ile | VUS | -10.02 | Indeterminate | 0.2831 | Benign | -8.89 | Pathogenic | 0.47 | Neutral | 0.60 | Possibly damaging | 0.43 | Tolerated | | |
| p.Ala30Met | VUS | -5.52 | Neutral | 0.2343 | Benign | -8.27 | Pathogenic | 0.34 | Neutral | 0.38 | Benign | 0.24 | Tolerated | | |
| p.Ala30His | VUS | -6.94 | Indeterminate | 0.2277 | Benign | -7.13 | Benign | 0.44 | Neutral | 0.96 | Possibly damaging | 0.56 | Tolerated | | |
| p.Ala30Gln | VUS | -11.53 | Indeterminate | 0.1304 | Benign | -5.04 | Benign | 0.30 | Neutral | 0.09 | Benign | 0.42 | Tolerated | | |
| p.Ala30Pro | VUS | -53.15 | Deleterious | 0.7136 | Pathogenic | -8.71 | Pathogenic | 0.53 | Disease | 0.81 | Possibly damaging | 0.26 | Tolerated | 23.10 | Deleterious |
| p.Ala30Leu | VUS | -9.74 | Indeterminate | 0.1437 | Benign | -5.95 | Benign | 0.29 | Neutral | 0.04 | Benign | 0.71 | Tolerated | | |
| p.Ala30Asp | VUS | -33.22 | Indeterminate | 0.1445 | Benign | -6.67 | Benign | 0.33 | Neutral | 0.01 | Benign | 0.34 | Tolerated | | |
| p.Ala30Glu | VUS | -2.24 | Neutral | 0.0953 | Benign | -4.64 | Benign | 0.27 | Neutral | 0.01 | Benign | 0.49 | Tolerated | 14.41 | Neutral |
| p.Ala30Gly | VUS | -53.15 | Deleterious | 0.1178 | Benign | -5.15 | Benign | 0.36 | Neutral | 0.01 | Benign | 0.41 | Tolerated | 17.42 | Deleterious |
| p.Ala30Val | VUS | -7.22 | Indeterminate | 0.1334 | Benign | -6.66 | Benign | 0.35 | Neutral | 0.83 | Possibly damaging | 0.56 | Tolerated | 17.56 | Deleterious |
| p.Ala30Tyr | VUS | -25.61 | Indeterminate | 0.3408 | Ambiguous | -9.70 | Pathogenic | 0.45 | Neutral | 0.99 | Probably damaging | 1 | Tolerated | | |
| p.Ala30Cys | VUS | -21.54 | Indeterminate | 0.4009 | Ambiguous | -7.39 | Benign | 0.42 | Neutral | 0.99 | Probably damaging | 0.19 | Tolerated | | |
| p.Ala30Trp | VUS | -14.31 | Indeterminate | 0.4175 | Ambiguous | -8.02 | Pathogenic | 0.52 | Disease | 1.00 | Probably damaging | 0.19 | Tolerated | | |
| p.Ala30Phe | VUS | -9.80 | Indeterminate | 0.2629 | Benign | -8.95 | Pathogenic | 0.46 | Neutral | 0.96 | Possibly damaging | 0.7 | Tolerated | | |
| p.Leu31Asn | VUS | -15.80 | Indeterminate | 0.8943 | Pathogenic | -12.27 | Pathogenic | 0.74 | Disease | 1.00 | Probably damaging | 0 | Deleterious | | |
| p.Leu31Lys | VUS | -1.84 | Neutral | 0.6424 | Pathogenic | -10.83 | Pathogenic | 0.76 | Disease | 1.00 | Probably damaging | 0 | Deleterious | | |
| p.Leu31Thr | VUS | -7.23 | Indeterminate | 0.7298 | Pathogenic | -8.94 | Pathogenic | 0.72 | Disease | 1.00 | Probably damaging | 0 | Deleterious | | |
| p.Leu31Arg | VUS | -3.31 | Neutral | 0.3758 | Ambiguous | -6.01 | Benign | 0.84 | Disease | 1.00 | Probably damaging | 0 | Deleterious | 31.00 | Deleterious |
| p.Leu31Ser | VUS | -12.36 | Indeterminate | 0.8197 | Pathogenic | -6.81 | Benign | 0.82 | Disease | 1.00 | Probably damaging | 0 | Deleterious | | |
| p.Leu31Ile | VUS | -1.36 | Neutral | 0.3859 | Ambiguous | -8.78 | Pathogenic | 0.53 | Disease | 1.00 | Probably damaging | 0.04 | Deleterious | | |
| p.Leu31Met | VUS | -6.95 | Indeterminate | 0.3112 | Benign | -6.60 | Benign | 0.54 | Disease | 1.00 | Probably damaging | 0.02 | Deleterious | 28.90 | Deleterious |
| p.Leu31His | VUS | -5.62 | Neutral | 0.5671 | Pathogenic | -5.44 | Benign | 0.82 | Disease | 1.00 | Probably damaging | 0 | Deleterious | | |
| p.Leu31Gln | VUS | -18.98 | Indeterminate | 0.4554 | Ambiguous | -6.92 | Benign | 0.83 | Disease | 1.00 | Probably damaging | 0 | Deleterious | 30.00 | Deleterious |
| p.Leu31Pro | VUS | -53.15 | Deleterious | 0.9258 | Pathogenic | -12.46 | Pathogenic | 0.71 | Disease | 1.00 | Probably damaging | 0 | Deleterious | 31.00 | Deleterious |
| p.Leu31Asp | VUS | -53.15 | Deleterious | 0.9567 | Pathogenic | -11.70 | Pathogenic | 0.73 | Disease | 1.00 | Probably damaging | 0 | Deleterious | | |
| p.Leu31Glu | VUS | -9.76 | Indeterminate | 0.7905 | Pathogenic | -7.46 | Benign | 0.76 | Disease | 1.00 | Probably damaging | 0 | Deleterious | | |
| p.Leu31Ala | VUS | -9.65 | Indeterminate | 0.447 | Ambiguous | -2.53 | Benign | 0.70 | Disease | 1.00 | Probably damaging | 0.01 | Deleterious | | |
| p.Leu31Gly | VUS | -6.62 | Indeterminate | 0.7501 | Pathogenic | -8.20 | Pathogenic | 0.75 | Disease | 1.00 | Probably damaging | 0 | Deleterious | | |
| p.Leu31Val | VUS | -21.43 | Indeterminate | 0.3177 | Benign | -6.65 | Benign | 0.44 | Neutral | 1.00 | Probably damaging | 0.02 | Deleterious | 28.50 | Deleterious |
| p.Leu31Tyr | VUS | -21.21 | Indeterminate | 0.6131 | Pathogenic | -6.76 | Benign | 0.71 | Disease | 1.00 | Probably damaging | 0.01 | Deleterious | | |
| p.Leu31Cys | VUS | -33.20 | Indeterminate | 0.7024 | Pathogenic | -4.19 | Benign | 0.66 | Disease | 1.00 | Probably damaging | 0 | Deleterious | | |
| p.Leu31Trp | VUS | -18.55 | Indeterminate | 0.5192 | Ambiguous | -6.83 | Benign | 0.76 | Disease | 1.00 | Probably damaging | 0 | Deleterious | | |
| p.Leu31Phe | VUS | -5.13 | Neutral | 0.3289 | Benign | -6.42 | Benign | 0.54 | Disease | 1.00 | Probably damaging | 0.18 | Tolerated | | |
| p.Leu32Asn | VUS | -53.15 | Deleterious | 0.9879 | Pathogenic | -19.16 | Pathogenic | 0.72 | Disease | 1.00 | Probably damaging | 0 | Deleterious | | |
| p.Leu32Lys | VUS | -53.15 | Deleterious | 0.9561 | Pathogenic | -18.46 | Pathogenic | 0.72 | Disease | 1.00 | Probably damaging | 0 | Deleterious | | |
| p.Leu32Thr | VUS | -15.95 | Indeterminate | 0.8634 | Pathogenic | -14.45 | Pathogenic | 0.72 | Disease | 1.00 | Probably damaging | 0 | Deleterious | | |
| p.Leu32Arg | VUS | -53.15 | Deleterious | 0.8215 | Pathogenic | -13.76 | Pathogenic | 0.75 | Disease | 1.00 | Probably damaging | 0 | Deleterious | 32.00 | Deleterious |
| p.Leu32Ser | VUS | -32.53 | Indeterminate | 0.9828 | Pathogenic | -14.35 | Pathogenic | 0.82 | Disease | 1.00 | Probably damaging | 0 | Deleterious | | |
| p.Leu32Ile | VUS | -0.06 | Neutral | 0.256 | Benign | -10.71 | Pathogenic | 0.46 | Neutral | 1.00 | Probably damaging | 0 | Deleterious | | |
| p.Leu32Met | VUS | -0.11 | Neutral | 0.6061 | Pathogenic | -10.80 | Pathogenic | 0.47 | Neutral | 1.00 | Probably damaging | 0 | Deleterious | 29.00 | Deleterious |
| p.Leu32His | VUS | -53.15 | Deleterious | 0.9762 | Pathogenic | -15.47 | Pathogenic | 0.79 | Disease | 1.00 | Probably damaging | 0 | Deleterious | | |
| p.Leu32Gln | VUS | -10.66 | Indeterminate | 0.9367 | Pathogenic | -15.08 | Pathogenic | 0.82 | Disease | 1.00 | Probably damaging | 0 | Deleterious | 29.90 | Deleterious |
| p.Leu32Pro | Pathogenic | -53.15 | Deleterious | 0.9446 | Pathogenic | -13.13 | Pathogenic | 0.80 | Disease | 1.00 | Probably damaging | 0 | Deleterious | 31.00 | Deleterious |
| p.Leu32Asp | VUS | -53.15 | Deleterious | 0.9957 | Pathogenic | -18.43 | Pathogenic | 0.72 | Disease | 1.00 | Probably damaging | 0 | Deleterious | | |
| p.Leu32Glu | VUS | -53.15 | Deleterious | 0.9666 | Pathogenic | -17.68 | Pathogenic | 0.75 | Disease | 1.00 | Probably damaging | 0 | Deleterious | | |
| p.Leu32Ala | VUS | -18.61 | Indeterminate | 0.8518 | Pathogenic | -14.08 | Pathogenic | 0.70 | Disease | 1.00 | Probably damaging | 0 | Deleterious | | |
| p.Leu32Gly | VUS | -53.15 | Deleterious | 0.9602 | Pathogenic | -15.95 | Pathogenic | 0.71 | Disease | 1.00 | Probably damaging | 0 | Deleterious | | |
| p.Leu32Val | VUS | -0.05 | Neutral | 0.3013 | Benign | -11.51 | Pathogenic | 0.43 | Neutral | 1.00 | Probably damaging | 0 | Deleterious | 28.60 | Deleterious |
| p.Leu32Tyr | VUS | -53.15 | Deleterious | 0.9847 | Pathogenic | -15.88 | Pathogenic | 0.70 | Disease | 1.00 | Probably damaging | 0 | Deleterious | | |
| p.Leu32Cys | VUS | -3.18 | Neutral | 0.9346 | Pathogenic | -11.44 | Pathogenic | 0.65 | Disease | 1.00 | Probably damaging | 0 | Deleterious | | |
| p.Leu32Trp | VUS | -53.15 | Deleterious | 0.9562 | Pathogenic | -14.68 | Pathogenic | 0.74 | Disease | 1.00 | Probably damaging | 0 | Deleterious | | |
| p.Leu32Phe | VUS | -5.09 | Neutral | 0.8974 | Pathogenic | -9.81 | Pathogenic | 0.52 | Disease | 1.00 | Probably damaging | 0 | Deleterious | | |
| p.Glu33Asn | VUS | -0.12 | Neutral | 0.4922 | Ambiguous | -8.59 | Pathogenic | 0.26 | Neutral | 0.97 | Probably damaging | 0.06 | Tolerated | | |
| p.Glu33Lys | VUS | -0.41 | Neutral | 0.2662 | Benign | -7.44 | Benign | 0.17 | Neutral | 0.82 | Possibly damaging | 0.56 | Tolerated | 22.20 | Deleterious |
| p.Glu33Thr | VUS | -0.19 | Neutral | 0.3245 | Benign | -7.16 | Benign | 0.38 | Neutral | 0.99 | Probably damaging | 0.06 | Tolerated | | |
| p.Glu33Arg | VUS | -9.24 | Indeterminate | 0.1813 | Benign | -5.53 | Benign | 0.27 | Neutral | 0.98 | Probably damaging | 0.46 | Tolerated | | |
| p.Glu33Ser | VUS | -3.59 | Neutral | 0.2948 | Benign | -6.08 | Benign | 0.37 | Neutral | 0.99 | Probably damaging | 0.06 | Tolerated | | |
| p.Glu33Ile | VUS | -0.01 | Neutral | 0.6202 | Pathogenic | -10.00 | Pathogenic | 0.38 | Neutral | 0.99 | Probably damaging | 0.02 | Deleterious | | |
| p.Glu33Met | VUS | -0.01 | Neutral | 0.6565 | Pathogenic | -9.14 | Pathogenic | 0.44 | Neutral | 0.99 | Probably damaging | 0.02 | Deleterious | | |
| p.Glu33His | VUS | -0.18 | Neutral | 0.3876 | Ambiguous | -7.31 | Benign | 0.31 | Neutral | 0.98 | Probably damaging | 0.25 | Tolerated | | |
| p.Glu33Gln | VUS | -0.09 | Neutral | 0.1304 | Benign | -5.18 | Benign | 0.16 | Neutral | 0.20 | Benign | 0.38 | Tolerated | 15.26 | Deleterious |
| p.Glu33Pro | VUS | -0.04 | Neutral | 0.9028 | Pathogenic | -10.54 | Pathogenic | 0.43 | Neutral | 1.00 | Probably damaging | 0.04 | Deleterious | | |
| p.Glu33Leu | VUS | -0.82 | Neutral | 0.3512 | Ambiguous | -6.18 | Benign | 0.28 | Neutral | 0.95 | Possibly damaging | 0.05 | Deleterious | | |
| p.Glu33Asp | VUS | -0.15 | Neutral | 0.2215 | Benign | -4.30 | Benign | 0.17 | Neutral | 0.03 | Benign | 0.05 | Deleterious | 14.99 | Neutral |
| p.Glu33Ala | VUS | -11.39 | Indeterminate | 0.1146 | Benign | -3.90 | Benign | 0.21 | Neutral | 0.96 | Probably damaging | 0.07 | Tolerated | 19.25 | Deleterious |
| p.Glu33Gly | VUS | -0.32 | Neutral | 0.1955 | Benign | -6.08 | Benign | 0.21 | Neutral | 1.00 | Probably damaging | 0.04 | Deleterious | 23.50 | Deleterious |
| p.Glu33Val | VUS | -1.44 | Neutral | 0.3459 | Ambiguous | -7.67 | Pathogenic | 0.23 | Neutral | 0.95 | Possibly damaging | 0.03 | Deleterious | 23.10 | Deleterious |
| p.Glu33Tyr | VUS | -9.22 | Indeterminate | 0.7106 | Pathogenic | -10.60 | Pathogenic | 0.38 | Neutral | 1.00 | Probably damaging | 0.02 | Deleterious | | |
| p.Glu33Cys | VUS | -3.97 | Neutral | 0.8469 | Pathogenic | -8.41 | Pathogenic | 0.38 | Neutral | 1.00 | Probably damaging | 0.01 | Deleterious | | |
| p.Glu33Trp | VUS | -0.26 | Neutral | 0.8764 | Pathogenic | -10.04 | Pathogenic | 0.45 | Neutral | 1.00 | Probably damaging | 0.01 | Deleterious | | |
| p.Glu33Phe | VUS | -3.58 | Neutral | 0.7972 | Pathogenic | -10.77 | Pathogenic | 0.39 | Neutral | 1.00 | Probably damaging | 0.01 | Deleterious | | |
| p.Ala34Asn | VUS | -11.51 | Indeterminate | 0.275 | Benign | -8.01 | Pathogenic | 0.32 | Neutral | 0.99 | Probably damaging | 0.45 | Tolerated | | |
| p.Ala34Lys | VUS | -1.08 | Neutral | 0.3383 | Benign | -6.90 | Benign | 0.45 | Neutral | 1.00 | Probably damaging | 0.57 | Tolerated | | |
| p.Ala34Thr | VUS | -5.63 | Neutral | 0.1519 | Benign | -7.78 | Pathogenic | 0.31 | Neutral | 0.99 | Probably damaging | 0.57 | Tolerated | 20.50 | Deleterious |
| p.Ala34Arg | VUS | -0.01 | Neutral | 0.2107 | Benign | -3.95 | Benign | 0.32 | Neutral | 1.00 | Probably damaging | 0.51 | Tolerated | | |
| p.Ala34Ser | VUS | -2.77 | Neutral | 0.1039 | Benign | -5.61 | Benign | 0.21 | Neutral | 0.94 | Possibly damaging | 0.62 | Tolerated | 18.38 | Deleterious |
| p.Ala34Ile | VUS | -0.97 | Neutral | 0.4646 | Ambiguous | -11.26 | Pathogenic | 0.44 | Neutral | 1.00 | Probably damaging | 0.45 | Tolerated | | |
| p.Ala34Met | VUS | -0.41 | Neutral | 0.3905 | Ambiguous | -9.67 | Pathogenic | 0.42 | Neutral | 1.00 | Probably damaging | 0.25 | Tolerated | | |
| p.Ala34His | VUS | -0.11 | Neutral | 0.3442 | Ambiguous | -6.92 | Benign | 0.46 | Neutral | 1.00 | Probably damaging | 0.57 | Tolerated | | |
| p.Ala34Gln | VUS | -1.75 | Neutral | 0.1966 | Benign | -4.53 | Benign | 0.34 | Neutral | 1.00 | Probably damaging | 0.46 | Tolerated | | |
| p.Ala34Pro | VUS | -11.89 | Indeterminate | 0.8853 | Pathogenic | -10.72 | Pathogenic | 0.55 | Disease | 1.00 | Probably damaging | 0.29 | Tolerated | 23.10 | Deleterious |
| p.Ala34Leu | VUS | -0.40 | Neutral | 0.2923 | Benign | -7.01 | Benign | 0.44 | Neutral | 1.00 | Probably damaging | 0.75 | Tolerated | | |
| p.Ala34Asp | VUS | -1.54 | Neutral | 0.1828 | Benign | -5.86 | Benign | 0.33 | Neutral | 1.00 | Probably damaging | 0.38 | Tolerated | | |
| p.Ala34Glu | VUS | -2.19 | Neutral | 0.167 | Benign | -4.98 | Benign | 0.31 | Neutral | 1.00 | Probably damaging | 0.54 | Tolerated | 18.56 | Deleterious |
| p.Ala34Gly | VUS | -2.72 | Neutral | 0.1321 | Benign | -4.75 | Benign | 0.23 | Neutral | 1.00 | Probably damaging | 0.45 | Tolerated | 22.60 | Deleterious |
| p.Ala34Val | VUS | -12.28 | Indeterminate | 0.2212 | Benign | -8.47 | Pathogenic | 0.23 | Neutral | 1.00 | Probably damaging | 0.59 | Tolerated | 21.90 | Deleterious |
| p.Ala34Tyr | VUS | -0.23 | Neutral | 0.5224 | Ambiguous | -10.87 | Pathogenic | 0.44 | Neutral | 1.00 | Probably damaging | 1 | Tolerated | | |
| p.Ala34Cys | VUS | -0.28 | Neutral | 0.562 | Ambiguous | -8.03 | Pathogenic | 0.41 | Neutral | 1.00 | Probably damaging | 0.19 | Tolerated | | |
| p.Ala34Trp | VUS | -1.47 | Neutral | 0.6783 | Pathogenic | -10.10 | Pathogenic | 0.47 | Neutral | 1.00 | Probably damaging | 0.19 | Tolerated | | |
| p.Ala34Phe | VUS | -0.11 | Neutral | 0.483 | Ambiguous | -11.46 | Pathogenic | 0.42 | Neutral | 1.00 | Probably damaging | 0.71 | Tolerated | | |
| p.Gly35Asn | VUS | -7.34 | Indeterminate | 0.6621 | Pathogenic | -12.62 | Pathogenic | 0.49 | Neutral | 1.00 | Probably damaging | 0.49 | Tolerated | | |
| p.Gly35Lys | VUS | -13.25 | Indeterminate | 0.6573 | Pathogenic | -13.14 | Pathogenic | 0.65 | Disease | 1.00 | Probably damaging | 0.33 | Tolerated | | |
| p.Gly35Thr | VUS | -53.15 | Deleterious | 0.6714 | Pathogenic | -13.85 | Pathogenic | 0.62 | Disease | 1.00 | Probably damaging | 0.08 | Tolerated | | |
| p.Gly35Arg | VUS | -2.42 | Neutral | 0.4978 | Ambiguous | -10.24 | Pathogenic | 0.68 | Disease | 1.00 | Probably damaging | 0.09 | Tolerated | 27.55 | Deleterious |
| p.Gly35Ser | VUS | -13.90 | Indeterminate | 0.3827 | Ambiguous | -11.39 | Pathogenic | 0.62 | Disease | 1.00 | Probably damaging | 0.16 | Tolerated | | |
| p.Gly35Ile | VUS | -53.15 | Deleterious | 0.9411 | Pathogenic | -17.95 | Pathogenic | 0.65 | Disease | 1.00 | Probably damaging | 0.01 | Deleterious | | |
| p.Gly35Met | VUS | -21.31 | Indeterminate | 0.936 | Pathogenic | -15.80 | Pathogenic | 0.67 | Disease | 1.00 | Probably damaging | 0.01 | Deleterious | | |
| p.Gly35His | VUS | -2.71 | Neutral | 0.6966 | Pathogenic | -12.46 | Pathogenic | 0.63 | Disease | 1.00 | Probably damaging | 0.06 | Tolerated | | |
| p.Gly35Gln | VUS | -9.22 | Indeterminate | 0.6005 | Pathogenic | -12.67 | Pathogenic | 0.48 | Neutral | 1.00 | Probably damaging | 0.09 | Tolerated | | |
| p.Gly35Pro | VUS | -53.15 | Deleterious | 0.9693 | Pathogenic | -14.73 | Pathogenic | 0.61 | Disease | 1.00 | Probably damaging | 0.06 | Tolerated | | |
| p.Gly35Leu | VUS | -53.15 | Deleterious | 0.9122 | Pathogenic | -14.52 | Pathogenic | 0.67 | Disease | 1.00 | Probably damaging | 0.03 | Deleterious | | |
| p.Gly35Asp | VUS | -17.21 | Indeterminate | 0.4675 | Ambiguous | -11.44 | Pathogenic | 0.70 | Disease | 1.00 | Probably damaging | 0.28 | Tolerated | | |
| p.Gly35Glu | VUS | -10.41 | Indeterminate | 0.5927 | Pathogenic | -12.25 | Pathogenic | 0.71 | Disease | 1.00 | Probably damaging | 0.31 | Tolerated | 26.50 | Deleterious |
| p.Gly35Ala | VUS | -7.55 | Indeterminate | 0.483 | Ambiguous | -11.44 | Pathogenic | 0.62 | Disease | 1.00 | Probably damaging | 0.09 | Tolerated | 25.70 | Deleterious |
| p.Gly35Val | VUS | -53.15 | Deleterious | 0.8532 | Pathogenic | -14.01 | Pathogenic | 0.78 | Disease | 1.00 | Probably damaging | 0.02 | Deleterious | 31.00 | Deleterious |
| p.Gly35Tyr | VUS | -26.28 | Indeterminate | 0.8898 | Pathogenic | -15.99 | Pathogenic | 0.66 | Disease | 1.00 | Probably damaging | 0.03 | Deleterious | | |

| | | | | | | | | | | | | | | | |
|---|---|---|---|---|---|---|---|---|---|---|---|---|---|---|---|
| p.Gly35Cys | VUS | -12.62 | Indeterminate | 0.7264 | Pathogenic | -12.78 | Pathogenic | 0.75 | Disease | 1.00 | Probably damaging | 0.01 | Deleterious | | |
| p.Gly35Trp | VUS | -53.15 | Deleterious | 0.8878 | Pathogenic | -14.67 | Pathogenic | 0.70 | Disease | 1.00 | Probably damaging | 0.01 | Deleterious | 32.00 | Deleterious |
| p.Gly35Phe | VUS | -8.93 | Indeterminate | 0.9481 | Pathogenic | -16.86 | Pathogenic | 0.67 | Disease | 1.00 | Probably damaging | 0.01 | Deleterious | | |
| p.Ala36Asn | VUS | -21.29 | Indeterminate | 0.4997 | Ambiguous | -11.26 | Pathogenic | 0.44 | Neutral | 0.55 | Possibly damaging | 0.02 | Deleterious | | |
| p.Ala36Lys | VUS | -53.15 | Deleterious | 0.7005 | Pathogenic | -10.58 | Pathogenic | 0.49 | Neutral | 0.98 | Probably damaging | 0.03 | Deleterious | | |
| p.Ala36Thr | VUS | -15.57 | Indeterminate | 0.1621 | Benign | -5.22 | Benign | 0.20 | Neutral | 0.02 | Benign | 0.2 | Tolerated | 22.70 | Deleterious |
| p.Ala36Arg | VUS | -0.86 | Neutral | 0.4371 | Ambiguous | -7.93 | Pathogenic | 0.44 | Neutral | 0.98 | Probably damaging | 0.02 | Deleterious | | |
| p.Ala36Ser | VUS | -3.16 | Neutral | 0.1364 | Benign | -6.94 | Benign | 0.34 | Neutral | 0.08 | Benign | 0.04 | Deleterious | 23.70 | Deleterious |
| p.Ala36Ile | VUS | -0.14 | Neutral | 0.3407 | Ambiguous | -8.07 | Pathogenic | 0.45 | Neutral | 0.85 | Possibly damaging | 0.22 | Tolerated | | |
| p.Ala36Met | VUS | 0.00 | Neutral | 0.3645 | Ambiguous | -6.87 | Benign | 0.31 | Neutral | 0.99 | Probably damaging | 0.05 | Deleterious | | |
| p.Ala36His | VUS | -0.02 | Neutral | 0.4481 | Ambiguous | -7.60 | Pathogenic | 0.43 | Neutral | 1.00 | Probably damaging | 0.01 | Deleterious | | |
| p.Ala36Gln | VUS | -0.07 | Neutral | 0.3982 | Ambiguous | -5.70 | Benign | 0.44 | Neutral | 0.99 | Probably damaging | 0.02 | Deleterious | | |
| p.Ala36Pro | VUS | -53.15 | Deleterious | 0.7831 | Pathogenic | -11.46 | Pathogenic | 0.59 | Disease | 0.94 | Possibly damaging | 0.05 | Deleterious | 26.20 | Deleterious |
| p.Ala36Leu | VUS | -5.94 | Indeterminate | 0.2434 | Benign | -5.75 | Benign | 0.47 | Neutral | 0.85 | Possibly damaging | 0.29 | Tolerated | | |
| p.Ala36Asp | VUS | -3.54 | Neutral | 0.5068 | Ambiguous | -9.06 | Pathogenic | 0.52 | Disease | 0.91 | Possibly damaging | 0.01 | Deleterious | | |
| p.Ala36Glu | VUS | -6.72 | Indeterminate | 0.4466 | Ambiguous | -5.04 | Benign | 0.46 | Neutral | 0.97 | Probably damaging | 0.03 | Deleterious | 21.60 | Deleterious |
| p.Ala36Gly | VUS | -11.46 | Indeterminate | 0.2002 | Benign | -7.89 | Pathogenic | 0.32 | Neutral | 0.89 | Possibly damaging | 0.01 | Deleterious | 22.40 | Deleterious |
| p.Ala36Val | VUS | -0.06 | Neutral | 0.1435 | Benign | -3.71 | Benign | 0.18 | Neutral | 0.05 | Benign | 1 | Tolerated | 15.30 | Deleterious |
| p.Ala36Tyr | VUS | -4.78 | Neutral | 0.4751 | Ambiguous | -9.39 | Pathogenic | 0.43 | Neutral | 0.99 | Probably damaging | 0.01 | Deleterious | | |
| p.Ala36Cys | VUS | 0.00 | Neutral | 0.3531 | Ambiguous | -6.44 | Benign | 0.42 | Neutral | 1.00 | Probably damaging | 0.04 | Deleterious | | |
| p.Ala36Trp | VUS | 0.00 | Neutral | 0.6743 | Pathogenic | -8.05 | Pathogenic | 0.52 | Disease | 1.00 | Probably damaging | 0 | Deleterious | | |
| p.Ala36Phe | VUS | -2.39 | Neutral | 0.3369 | Benign | -9.00 | Pathogenic | 0.45 | Neutral | 0.98 | Probably damaging | 0.03 | Deleterious | | |
| p.Leu37Asn | VUS | -0.07 | Neutral | 0.1197 | Benign | -0.16 | Benign | 0.20 | Neutral | 0.08 | Benign | 0.53 | Tolerated | | |
| p.Leu37Lys | VUS | -8.20 | Indeterminate | 0.1038 | Benign | -5.18 | Benign | 0.27 | Neutral | 0.08 | Benign | 0.86 | Tolerated | | |
| p.Leu37Thr | VUS | -15.43 | Indeterminate | 0.1027 | Benign | -4.14 | Benign | 0.24 | Neutral | 0.02 | Benign | 0.54 | Tolerated | | |
| p.Leu37Arg | VUS | 0.00 | Neutral | 0.0572 | Benign | -3.63 | Benign | 0.12 | Neutral | 0.14 | Benign | 0.48 | Tolerated | 13.65 | Neutral |
| p.Leu37Ser | VUS | -3.05 | Neutral | 0.0723 | Benign | 0.97 | Benign | 0.12 | Neutral | 0.00 | Benign | 0.76 | Tolerated | | |
| p.Leu37Ile | VUS | -15.63 | Indeterminate | 0.1386 | Benign | -6.99 | Benign | 0.18 | Neutral | 0.15 | Benign | 0.18 | Tolerated | | |
| p.Leu37Met | VUS | -0.02 | Neutral | 0.1321 | Benign | -6.51 | Benign | 0.20 | Neutral | 0.78 | Possibly damaging | 0.1 | Tolerated | 13.49 | Neutral |
| p.Leu37His | VUS | -8.27 | Indeterminate | 0.0924 | Benign | -2.66 | Benign | 0.13 | Neutral | 0.82 | Possibly damaging | 0.13 | Tolerated | | |
| p.Leu37Gln | VUS | -9.51 | Indeterminate | 0.0564 | Benign | -3.34 | Benign | 0.11 | Neutral | 0.14 | Benign | 0.55 | Tolerated | 12.85 | Neutral |
| p.Leu37Pro | VUS | -27.67 | Indeterminate | 0.0683 | Benign | -0.36 | Benign | 0.12 | Neutral | 0.00 | Benign | 0.26 | Tolerated | 14.31 | Neutral |
| p.Leu37Asp | VUS | -2.34 | Neutral | 0.1044 | Benign | 3.18 | Benign | 0.21 | Neutral | 0.00 | Benign | 0.68 | Tolerated | | |
| p.Leu37Glu | VUS | -2.57 | Neutral | 0.0784 | Benign | -0.46 | Benign | 0.29 | Neutral | 0.02 | Benign | 1 | Tolerated | | |
| p.Leu37Ala | VUS | -0.14 | Neutral | 0.0689 | Benign | -1.70 | Benign | 0.20 | Neutral | 0.01 | Benign | 0.71 | Tolerated | | |
| p.Leu37Gly | VUS | -2.03 | Neutral | 0.2015 | Benign | -2.31 | Benign | 0.27 | Neutral | 0.02 | Benign | 0.36 | Tolerated | | |
| p.Leu37Val | VUS | -9.83 | Indeterminate | 0.076 | Benign | -4.47 | Benign | 0.13 | Neutral | 0.06 | Benign | 0.31 | Tolerated | 10.87 | Neutral |
| p.Leu37Tyr | VUS | -8.28 | Indeterminate | 0.2359 | Benign | -5.79 | Benign | 0.25 | Neutral | 0.82 | Possibly damaging | 0.06 | Tolerated | | |
| p.Leu37Cys | VUS | -0.85 | Neutral | 0.326 | Benign | -2.31 | Benign | 0.26 | Neutral | 0.82 | Possibly damaging | 0.05 | Deleterious | | |
| p.Leu37Trp | VUS | -53.15 | Deleterious | 0.1527 | Benign | -5.76 | Benign | 0.25 | Neutral | 0.95 | Possibly damaging | 0.02 | Deleterious | | |
| p.Leu37Phe | VUS | -0.24 | Neutral | 0.1341 | Benign | -5.89 | Benign | 0.10 | Neutral | 0.30 | Benign | 0.06 | Tolerated | | |
| p.Pro38Asn | VUS | -53.15 | Deleterious | 0.9812 | Pathogenic | -17.83 | Pathogenic | 0.47 | Neutral | 0.97 | Probably damaging | 0.03 | Deleterious | | |
| p.Pro38Lys | VUS | -53.15 | Deleterious | 0.9373 | Pathogenic | -15.62 | Pathogenic | 0.53 | Disease | 0.99 | Probably damaging | 0.06 | Tolerated | | |
| p.Pro38Thr | VUS | -6.12 | Indeterminate | 0.6142 | Pathogenic | -11.08 | Pathogenic | 0.40 | Neutral | 0.75 | Possibly damaging | 0.25 | Tolerated | 20.60 | Deleterious |
| p.Pro38Arg | VUS | -53.15 | Deleterious | 0.7822 | Pathogenic | -13.17 | Pathogenic | 0.58 | Disease | 0.99 | Probably damaging | 0.04 | Deleterious | 24.30 | Deleterious |
| p.Pro38Ser | VUS | -17.33 | Indeterminate | 0.7699 | Pathogenic | -11.06 | Pathogenic | 0.59 | Disease | 0.95 | Possibly damaging | 0.07 | Tolerated | 17.36 | Deleterious |
| p.Pro38Ile | VUS | -18.09 | Indeterminate | 0.7013 | Pathogenic | -12.16 | Pathogenic | 0.47 | Neutral | 0.99 | Probably damaging | 0.61 | Tolerated | | |
| p.Pro38Met | VUS | -27.70 | Indeterminate | 0.9268 | Pathogenic | -14.33 | Pathogenic | 0.47 | Neutral | 1.00 | Probably damaging | 0.1 | Tolerated | | |
| p.Pro38His | VUS | -53.15 | Deleterious | 0.9314 | Pathogenic | -14.31 | Pathogenic | 0.46 | Neutral | 1.00 | Probably damaging | 0.02 | Deleterious | 24.60 | Deleterious |
| p.Pro38Gln | VUS | -53.15 | Deleterious | 0.8702 | Pathogenic | -14.75 | Pathogenic | 0.44 | Neutral | 0.99 | Probably damaging | 0.04 | Deleterious | | |
| p.Pro38Leu | VUS | -33.22 | Indeterminate | 0.6615 | Pathogenic | -11.99 | Pathogenic | 0.42 | Neutral | 0.98 | Probably damaging | 0.39 | Tolerated | 24.50 | Deleterious |
| p.Pro38Asp | VUS | -53.15 | Deleterious | 0.9831 | Pathogenic | -16.69 | Pathogenic | 0.50 | Neutral | 0.96 | Probably damaging | 0.02 | Deleterious | | |
| p.Pro38Glu | VUS | -25.68 | Indeterminate | 0.9228 | Pathogenic | -14.43 | Pathogenic | 0.47 | Neutral | 0.78 | Possibly damaging | 0.06 | Tolerated | | |
| p.Pro38Ala | VUS | -53.15 | Deleterious | 0.297 | Benign | -6.83 | Benign | 0.19 | Neutral | 0.03 | Benign | 0.37 | Tolerated | 15.27 | Deleterious |
| p.Pro38Gly | VUS | -0.71 | Neutral | 0.8417 | Pathogenic | -11.38 | Pathogenic | 0.46 | Neutral | 0.95 | Possibly damaging | 0.02 | Deleterious | | |
| p.Pro38Val | VUS | -0.53 | Neutral | 0.3993 | Ambiguous | -7.82 | Pathogenic | 0.28 | Neutral | 0.74 | Possibly damaging | 1 | Tolerated | | |
| p.Pro38Tyr | VUS | -53.15 | Deleterious | 0.9766 | Pathogenic | -16.85 | Pathogenic | 0.48 | Neutral | 1.00 | Probably damaging | 0.02 | Deleterious | | |
| p.Pro38Cys | VUS | -0.04 | Neutral | 0.8647 | Pathogenic | -11.25 | Pathogenic | 0.47 | Neutral | 1.00 | Probably damaging | 0.07 | Tolerated | | |
| p.Pro38Trp | VUS | -53.15 | Deleterious | 0.9811 | Pathogenic | -16.36 | Pathogenic | 0.53 | Disease | 1.00 | Probably damaging | 0.01 | Deleterious | | |
| p.Pro38Phe | VUS | -53.15 | Deleterious | 0.9761 | Pathogenic | -16.36 | Pathogenic | 0.48 | Neutral | 1.00 | Probably damaging | 0.07 | Tolerated | | |
| p.Asn39Lys | VUS | -22.94 | Indeterminate | 0.9206 | Pathogenic | -10.77 | Pathogenic | 0.56 | Disease | 0.97 | Probably damaging | 0.02 | Deleterious | 22.45 | Deleterious |
| p.Asn39Thr | VUS | -0.01 | Neutral | 0.479 | Ambiguous | -7.19 | Benign | 0.47 | Neutral | 0.65 | Possibly damaging | 0.01 | Deleterious | 23.90 | Deleterious |
| p.Asn39Arg | VUS | -3.95 | Neutral | 0.8501 | Pathogenic | -8.02 | Pathogenic | 0.52 | Disease | 0.99 | Probably damaging | 0.01 | Deleterious | | |
| p.Asn39Ser | VUS | -4.05 | Neutral | 0.1805 | Benign | -5.65 | Benign | 0.64 | Disease | 0.95 | Possibly damaging | 0.03 | Deleterious | 22.20 | Deleterious |
| p.Asn39Ile | VUS | -7.95 | Indeterminate | 0.9261 | Pathogenic | -10.35 | Pathogenic | 0.68 | Disease | 0.98 | Probably damaging | 0 | Deleterious | 24.70 | Deleterious |
| p.Asn39Met | VUS | -25.20 | Indeterminate | 0.8921 | Pathogenic | -9.61 | Pathogenic | 0.61 | Disease | 1.00 | Probably damaging | 0 | Deleterious | | |
| p.Asn39His | VUS | -0.03 | Neutral | 0.4608 | Ambiguous | -7.72 | Pathogenic | 0.36 | Neutral | 0.99 | Probably damaging | 0.01 | Deleterious | 16.04 | Deleterious |
| p.Asn39Gln | VUS | -1.24 | Neutral | 0.7922 | Pathogenic | -8.55 | Pathogenic | 0.52 | Disease | 0.98 | Probably damaging | 0.01 | Deleterious | | |
| p.Asn39Pro | VUS | -53.15 | Deleterious | 0.9481 | Pathogenic | -11.48 | Pathogenic | 0.53 | Disease | 0.99 | Probably damaging | 0.01 | Deleterious | | |
| p.Asn39Leu | VUS | -53.15 | Deleterious | 0.7873 | Pathogenic | -8.94 | Pathogenic | 0.60 | Disease | 0.98 | Probably damaging | 0 | Deleterious | | |
| p.Asn39Asp | VUS | -0.66 | Neutral | 0.171 | Benign | -4.01 | Benign | 0.26 | Neutral | 0.05 | Benign | 0.62 | Tolerated | 12.26 | Neutral |
| p.Asn39Glu | VUS | -1.56 | Neutral | 0.7546 | Pathogenic | -6.00 | Benign | 0.51 | Disease | 0.72 | Possibly damaging | 0.02 | Deleterious | | |
| p.Asn39Ala | VUS | -6.60 | Indeterminate | 0.6305 | Pathogenic | -6.35 | Benign | 0.50 | Neutral | 0.74 | Possibly damaging | 0.01 | Deleterious | | |
| p.Asn39Gly | VUS | -0.34 | Neutral | 0.5207 | Ambiguous | -6.75 | Benign | 0.50 | Disease | 0.99 | Probably damaging | 0.04 | Deleterious | | |
| p.Asn39Val | VUS | -4.58 | Neutral | 0.8577 | Pathogenic | -7.98 | Pathogenic | 0.59 | Disease | 0.81 | Possibly damaging | 0 | Deleterious | | |
| p.Asn39Tyr | VUS | -0.72 | Neutral | 0.77 | Pathogenic | -12.28 | Pathogenic | 0.58 | Disease | 1.00 | Probably damaging | 0.01 | Deleterious | 22.00 | Deleterious |
| p.Asn39Cys | VUS | -53.15 | Deleterious | 0.7817 | Pathogenic | -7.10 | Benign | 0.59 | Disease | 1.00 | Probably damaging | 0 | Deleterious | | |
| p.Asn39Trp | VUS | -13.56 | Indeterminate | 0.9764 | Pathogenic | -12.04 | Pathogenic | 0.62 | Disease | 1.00 | Probably damaging | 0 | Deleterious | | |
| p.Asn39Phe | VUS | -53.15 | Deleterious | 0.9575 | Pathogenic | -12.35 | Pathogenic | 0.62 | Disease | 1.00 | Probably damaging | 0 | Deleterious | | |
| p.Ala40Asn | VUS | -6.02 | Indeterminate | 0.4103 | Ambiguous | -12.26 | Pathogenic | 0.37 | Neutral | 0.88 | Possibly damaging | 0.35 | Tolerated | | |
| p.Ala40Lys | VUS | -1.40 | Neutral | 0.3873 | Ambiguous | -9.78 | Pathogenic | 0.42 | Neutral | 0.94 | Possibly damaging | 0.65 | Tolerated | | |
| p.Ala40Thr | VUS | -53.15 | Deleterious | 0.1094 | Benign | -5.35 | Benign | 0.13 | Neutral | 0.98 | Probably damaging | 0.49 | Tolerated | 20.10 | Deleterious |
| p.Ala40Arg | VUS | -2.18 | Neutral | 0.2217 | Benign | -7.59 | Pathogenic | 0.40 | Neutral | 0.19 | Benign | 0.4 | Tolerated | | |
| p.Ala40Ser | VUS | -0.83 | Neutral | 0.1333 | Benign | -6.22 | Benign | 0.17 | Neutral | 0.54 | Possibly damaging | 0.52 | Tolerated | 22.60 | Deleterious |
| p.Ala40Ile | VUS | -6.06 | Indeterminate | 0.2938 | Benign | -9.12 | Pathogenic | 0.29 | Neutral | 1.00 | Probably damaging | 0.15 | Tolerated | | |
| p.Ala40Met | VUS | -0.01 | Neutral | 0.3271 | Benign | -9.25 | Pathogenic | 0.40 | Neutral | 1.00 | Probably damaging | 0.08 | Tolerated | | |
| p.Ala40His | VUS | -0.39 | Neutral | 0.3516 | Ambiguous | -10.41 | Pathogenic | 0.39 | Neutral | 1.00 | Probably damaging | 0.11 | Tolerated | | |
| p.Ala40Gln | VUS | -0.02 | Neutral | 0.1982 | Benign | -7.81 | Pathogenic | 0.38 | Neutral | 0.99 | Probably damaging | 0.41 | Tolerated | | |
| p.Ala40Pro | VUS | -12.20 | Indeterminate | 0.5607 | Ambiguous | -8.42 | Pathogenic | 0.36 | Neutral | 1.00 | Probably damaging | 0.2 | Tolerated | 24.40 | Deleterious |
| p.Ala40Leu | VUS | -53.15 | Deleterious | 0.1751 | Benign | -8.21 | Pathogenic | 0.47 | Neutral | 0.97 | Probably damaging | 0.36 | Tolerated | | |
| p.Ala40Asp | VUS | -17.56 | Indeterminate | 0.3014 | Benign | -9.74 | Pathogenic | 0.36 | Neutral | 0.98 | Probably damaging | 0.43 | Tolerated | | |
| p.Ala40Glu | VUS | -0.05 | Neutral | 0.1761 | Benign | -6.70 | Benign | 0.36 | Neutral | 0.90 | Possibly damaging | 0.8 | Tolerated | 16.96 | Deleterious |
| p.Ala40Gly | VUS | -0.94 | Neutral | 0.1478 | Benign | -5.36 | Benign | 0.15 | Neutral | 0.01 | Benign | 0.33 | Tolerated | 15.48 | Deleterious |
| p.Ala40Val | VUS | -6.48 | Indeterminate | 0.13 | Benign | -6.01 | Benign | 0.19 | Neutral | 0.95 | Possibly damaging | 0.26 | Tolerated | 21.60 | Deleterious |
| p.Ala40Tyr | VUS | -0.12 | Neutral | 0.4958 | Ambiguous | -11.90 | Pathogenic | 0.42 | Neutral | 1.00 | Probably damaging | 0.08 | Tolerated | | |
| p.Ala40Cys | VUS | -1.56 | Neutral | 0.5347 | Ambiguous | -8.14 | Pathogenic | 0.38 | Neutral | 1.00 | Probably damaging | 0.05 | Deleterious | | |
| p.Ala40Trp | VUS | -5.12 | Neutral | 0.6552 | Pathogenic | -10.32 | Pathogenic | 0.49 | Neutral | 1.00 | Probably damaging | 0.02 | Deleterious | | |
| p.Ala40Phe | VUS | -2.21 | Neutral | 0.3808 | Ambiguous | -10.62 | Pathogenic | 0.45 | Neutral | 1.00 | Probably damaging | 0.08 | Tolerated | | |
| p.Pro41Asn | VUS | -0.39 | Neutral | 0.5988 | Pathogenic | -10.56 | Pathogenic | 0.27 | Neutral | 0.67 | Possibly damaging | 0.07 | Tolerated | | |
| p.Pro41Lys | VUS | -4.11 | Neutral | 0.3597 | Ambiguous | -8.97 | Pathogenic | 0.24 | Neutral | 0.32 | Benign | 0.11 | Tolerated | | |
| p.Pro41Thr | VUS | -1.20 | Neutral | 0.1485 | Benign | -4.89 | Benign | 0.11 | Neutral | 0.08 | Benign | 0.35 | Tolerated | 11.57 | Neutral |
| p.Pro41Arg | VUS | -29.93 | Indeterminate | 0.1457 | Benign | -5.20 | Benign | 0.09 | Neutral | 0.06 | Benign | 0.08 | Tolerated | 4.17 | Neutral |
| p.Pro41Ser | VUS | -0.17 | Neutral | 0.1671 | Benign | -7.00 | Benign | 0.17 | Neutral | 0.84 | Possibly damaging | 0.13 | Tolerated | 12.33 | Neutral |
| p.Pro41Ile | VUS | -11.50 | Indeterminate | 0.4238 | Ambiguous | -4.31 | Benign | 0.28 | Neutral | 0.32 | Benign | 0.51 | Tolerated | | |
| p.Pro41Met | VUS | -0.01 | Neutral | 0.506 | Ambiguous | -5.75 | Benign | 0.26 | Neutral | 0.75 | Possibly damaging | 0.14 | Tolerated | | |
| p.Pro41His | VUS | -18.07 | Indeterminate | 0.3498 | Ambiguous | -5.42 | Benign | 0.17 | Neutral | 0.95 | Possibly damaging | 0.07 | Tolerated | | |
| p.Pro41Gln | VUS | -2.48 | Neutral | 0.1882 | Benign | -5.74 | Benign | 0.21 | Neutral | 0.85 | Possibly damaging | 0.09 | Tolerated | 12.54 | Neutral |
| p.Pro41Leu | VUS | -0.03 | Neutral | 0.2279 | Benign | -0.97 | Benign | 0.11 | Neutral | 0.01 | Benign | 0.51 | Tolerated | 8.79 | Neutral |
| p.Pro41Asp | VUS | -2.62 | Neutral | 0.584 | Pathogenic | -11.96 | Pathogenic | 0.28 | Neutral | 0.34 | Benign | 0.05 | Deleterious | | |
| p.Pro41Glu | VUS | -0.14 | Neutral | 0.3055 | Benign | -7.52 | Pathogenic | 0.24 | Neutral | 0.07 | Benign | 0.1 | Tolerated | | |
| p.Pro41Ala | VUS | -0.27 | Neutral | 0.07 | Benign | -4.16 | Benign | 0.15 | Neutral | 0.12 | Benign | 0.51 | Tolerated | 9.37 | Neutral |

| | | | | | | | | | | | | | | | |
|---|---|---|---|---|---|---|---|---|---|---|---|---|---|---|---|
| p.Pro41Gly | VUS | -1.82 | Neutral | 0.2599 | Benign | -7.59 | Pathogenic | 0.26 | Neutral | 0.84 | Possibly damaging | 0.05 | Deleterious | | |
| p.Pro41Val | VUS | -0.30 | Neutral | 0.2118 | Benign | -1.60 | Benign | 0.19 | Neutral | 0.00 | Benign | 1 | Tolerated | | |
| p.Pro41Tyr | VUS | -16.93 | Indeterminate | 0.6575 | Pathogenic | -7.76 | Pathogenic | 0.26 | Neutral | 0.94 | Possibly damaging | 0.11 | Tolerated | | |
| p.Pro41Cys | VUS | -3.07 | Neutral | 0.5137 | Ambiguous | -6.71 | Benign | 0.26 | Neutral | 1.00 | Probably damaging | 0.11 | Tolerated | | |
| p.Pro41Trp | VUS | -6.53 | Indeterminate | 0.6471 | Pathogenic | -10.69 | Pathogenic | 0.29 | Neutral | 1.00 | Probably damaging | 0.03 | Deleterious | | |
| p.Pro41Phe | VUS | -4.10 | Neutral | 0.6351 | Pathogenic | -6.87 | Benign | 0.26 | Neutral | 0.89 | Possibly damaging | 0.14 | Tolerated | | |
| p.Asn42Lys | VUS | -53.15 | Deleterious | 0.9801 | Pathogenic | -13.18 | Pathogenic | 0.71 | Disease | 1.00 | Probably damaging | 0 | Deleterious | 22.60 | Deleterious |
| p.Asn42Thr | VUS | -3.68 | Neutral | 0.6044 | Pathogenic | -9.85 | Pathogenic | 0.64 | Disease | 0.99 | Probably damaging | 0 | Deleterious | 22.70 | Deleterious |
| p.Asn42Arg | VUS | -53.15 | Deleterious | 0.944 | Pathogenic | -10.75 | Pathogenic | 0.67 | Disease | 1.00 | Probably damaging | 0 | Deleterious | | |
| p.Asn42Ser | VUS | -3.19 | Neutral | 0.2969 | Benign | -8.85 | Pathogenic | 0.61 | Disease | 1.00 | Probably damaging | 0 | Deleterious | 22.30 | Deleterious |
| p.Asn42Ile | VUS | -53.15 | Deleterious | 0.9567 | Pathogenic | -12.14 | Pathogenic | 0.82 | Disease | 1.00 | Probably damaging | 0 | Deleterious | 23.00 | Deleterious |
| p.Asn42Met | VUS | -53.15 | Deleterious | 0.9535 | Pathogenic | -13.26 | Pathogenic | 0.76 | Disease | 1.00 | Probably damaging | 0 | Deleterious | | |
| p.Asn42His | VUS | -53.15 | Deleterious | 0.7557 | Pathogenic | -10.08 | Pathogenic | 0.73 | Disease | 1.00 | Probably damaging | 0 | Deleterious | 23.60 | Deleterious |
| p.Asn42Gln | VUS | -53.15 | Deleterious | 0.9405 | Pathogenic | -12.35 | Pathogenic | 0.66 | Disease | 1.00 | Probably damaging | 0 | Deleterious | | |
| p.Asn42Pro | VUS | -53.15 | Deleterious | 0.966 | Pathogenic | -12.38 | Pathogenic | 0.71 | Disease | 1.00 | Probably damaging | 0 | Deleterious | | |
| p.Asn42Leu | VUS | -53.15 | Deleterious | 0.8813 | Pathogenic | -11.52 | Pathogenic | 0.75 | Disease | 1.00 | Probably damaging | 0 | Deleterious | | |
| p.Asn42Asp | VUS | -10.68 | Indeterminate | 0.5772 | Pathogenic | -7.97 | Pathogenic | 0.64 | Disease | 1.00 | Probably damaging | 0 | Deleterious | 23.80 | Deleterious |
| p.Asn42Glu | VUS | -53.15 | Deleterious | 0.9572 | Pathogenic | -14.75 | Pathogenic | 0.62 | Disease | 1.00 | Probably damaging | 0 | Deleterious | | |
| p.Asn42Ala | VUS | -3.46 | Neutral | 0.87 | Pathogenic | -10.39 | Pathogenic | 0.69 | Disease | 1.00 | Probably damaging | 0 | Deleterious | | |
| p.Asn42Gly | VUS | -0.24 | Neutral | 0.7349 | Pathogenic | -9.50 | Pathogenic | 0.65 | Disease | 1.00 | Probably damaging | 0 | Deleterious | | |
| p.Asn42Val | VUS | -53.15 | Deleterious | 0.9156 | Pathogenic | -12.01 | Pathogenic | 0.74 | Disease | 1.00 | Probably damaging | 0 | Deleterious | | |
| p.Asn42Tyr | VUS | -53.15 | Deleterious | 0.9089 | Pathogenic | -14.34 | Pathogenic | 0.82 | Disease | 1.00 | Probably damaging | 0 | Deleterious | 24.20 | Deleterious |
| p.Asn42Cys | VUS | -6.36 | Indeterminate | 0.776 | Pathogenic | -12.33 | Pathogenic | 0.73 | Disease | 1.00 | Probably damaging | 0 | Deleterious | | |
| p.Asn42Trp | VUS | -53.15 | Deleterious | 0.9933 | Pathogenic | -14.74 | Pathogenic | 0.76 | Disease | 1.00 | Probably damaging | 0 | Deleterious | | |
| p.Asn42Phe | VUS | -53.15 | Deleterious | 0.9846 | Pathogenic | -14.82 | Pathogenic | 0.75 | Disease | 1.00 | Probably damaging | 0 | Deleterious | | |
| p.Ser43Asn | VUS | -3.02 | Neutral | 0.2256 | Benign | -5.25 | Benign | 0.16 | Neutral | 0.02 | Benign | 0.53 | Tolerated | 15.04 | Deleterious |
| p.Ser43Lys | VUS | -0.48 | Neutral | 0.4613 | Ambiguous | -2.21 | Benign | 0.23 | Neutral | 0.17 | Benign | 0.87 | Tolerated | | |
| p.Ser43Thr | VUS | -0.40 | Neutral | 0.1071 | Benign | -4.90 | Benign | 0.13 | Neutral | 0.02 | Benign | 0.56 | Tolerated | 14.53 | Neutral |
| p.Ser43Arg | VUS | -0.31 | Neutral | 0.23 | Benign | 0.31 | Benign | 0.07 | Neutral | 0.00 | Benign | 0.56 | Tolerated | 5.19 | Neutral |
| p.Ser43Ile | VUS | -6.55 | Indeterminate | 0.2521 | Benign | -6.92 | Benign | 0.17 | Neutral | 0.77 | Possibly damaging | 0.17 | Tolerated | 16.76 | Deleterious |
| p.Ser43Met | VUS | -0.84 | Neutral | 0.4134 | Ambiguous | -7.16 | Benign | 0.22 | Neutral | 0.98 | Probably damaging | 0.1 | Tolerated | | |
| p.Ser43His | VUS | -0.33 | Neutral | 0.2069 | Benign | -4.76 | Benign | 0.21 | Neutral | 0.00 | Benign | 0.13 | Tolerated | | |
| p.Ser43Gln | VUS | -0.30 | Neutral | 0.279 | Benign | -4.04 | Benign | 0.22 | Neutral | 0.47 | Possibly damaging | 0.55 | Tolerated | | |
| p.Ser43Pro | VUS | -0.74 | Neutral | 0.1875 | Benign | -5.93 | Benign | 0.16 | Neutral | 0.78 | Possibly damaging | 0.26 | Tolerated | | |
| p.Ser43Leu | VUS | -1.74 | Neutral | 0.2431 | Benign | -6.35 | Benign | 0.16 | Neutral | 0.47 | Possibly damaging | 0.28 | Tolerated | | |
| p.Ser43Asp | VUS | -1.70 | Neutral | 0.3709 | Ambiguous | -6.10 | Benign | 0.21 | Neutral | 0.33 | Benign | 0.67 | Tolerated | | |
| p.Ser43Glu | VUS | -0.60 | Neutral | 0.3541 | Ambiguous | -4.04 | Benign | 0.20 | Neutral | 0.13 | Benign | 1 | Tolerated | | |
| p.Ser43Ala | VUS | -0.70 | Neutral | 0.0769 | Benign | -2.11 | Benign | 0.11 | Neutral | 0.01 | Benign | 0.71 | Tolerated | | |
| p.Ser43Gly | VUS | -4.02 | Neutral | 0.0829 | Benign | -4.55 | Benign | 0.13 | Neutral | 0.01 | Benign | 0.43 | Tolerated | 8.63 | Neutral |
| p.Ser43Val | VUS | -1.68 | Neutral | 0.2302 | Benign | -5.21 | Benign | 0.21 | Neutral | 0.62 | Possibly damaging | 0.26 | Tolerated | | |
| p.Ser43Tyr | VUS | -2.22 | Neutral | 0.2925 | Benign | -8.82 | Pathogenic | 0.22 | Neutral | 0.69 | Possibly damaging | 0.05 | Deleterious | | |
| p.Ser43Cys | VUS | -6.00 | Indeterminate | 0.1382 | Benign | -5.33 | Benign | 0.14 | Neutral | 0.91 | Possibly damaging | 0.05 | Deleterious | 10.53 | Neutral |
| p.Ser43Trp | VUS | -2.10 | Neutral | 0.3461 | Ambiguous | -7.76 | Pathogenic | 0.27 | Neutral | 0.99 | Probably damaging | 0.02 | Deleterious | | |
| p.Ser43Phe | VUS | -1.32 | Neutral | 0.3716 | Ambiguous | -9.45 | Pathogenic | 0.18 | Neutral | 0.82 | Possibly damaging | 0.05 | Deleterious | | |
| p.Tyr44Asn | VUS | -3.69 | Neutral | 0.5574 | Ambiguous | -12.78 | Pathogenic | 0.27 | Neutral | 0.99 | Probably damaging | 0.26 | Tolerated | 14.06 | Neutral |
| p.Tyr44Lys | VUS | -8.10 | Indeterminate | 0.7979 | Pathogenic | -13.39 | Pathogenic | 0.31 | Neutral | 0.97 | Probably damaging | 0.15 | Tolerated | | |
| p.Tyr44Thr | VUS | -1.08 | Neutral | 0.7134 | Pathogenic | -12.72 | Pathogenic | 0.35 | Neutral | 1.00 | Probably damaging | 0.2 | Tolerated | | |
| p.Tyr44Arg | VUS | -3.23 | Neutral | 0.5169 | Ambiguous | -10.93 | Pathogenic | 0.29 | Neutral | 1.00 | Probably damaging | 0.15 | Tolerated | | |
| p.Tyr44Ser | VUS | -3.02 | Neutral | 0.4284 | Ambiguous | -10.59 | Pathogenic | 0.23 | Neutral | 1.00 | Probably damaging | 0.32 | Tolerated | 13.61 | Neutral |
| p.Tyr44Ile | VUS | -3.05 | Neutral | 0.7981 | Pathogenic | -14.02 | Pathogenic | 0.30 | Neutral | 0.09 | Benign | 0.18 | Tolerated | | |
| p.Tyr44Met | VUS | -3.22 | Neutral | 0.816 | Pathogenic | -11.39 | Pathogenic | 0.30 | Neutral | 0.99 | Probably damaging | 0.1 | Tolerated | | |
| p.Tyr44His | VUS | -1.55 | Neutral | 0.3798 | Ambiguous | -9.04 | Pathogenic | 0.14 | Neutral | 0.09 | Benign | 0.24 | Tolerated | 9.59 | Neutral |
| p.Tyr44Gln | VUS | -18.21 | Indeterminate | 0.6722 | Pathogenic | -11.73 | Pathogenic | 0.23 | Neutral | 1.00 | Probably damaging | 0.14 | Tolerated | | |
| p.Tyr44Pro | VUS | -32.89 | Indeterminate | 0.9497 | Pathogenic | -13.45 | Pathogenic | 0.30 | Neutral | 1.00 | Probably damaging | 0.11 | Tolerated | | |
| p.Tyr44Leu | VUS | -2.69 | Neutral | 0.6422 | Pathogenic | -8.40 | Pathogenic | 0.27 | Neutral | 0.00 | Benign | 0.31 | Tolerated | | |
| p.Tyr44Asp | VUS | -6.48 | Indeterminate | 0.6152 | Pathogenic | -12.63 | Pathogenic | 0.25 | Neutral | 1.00 | Probably damaging | 0.32 | Tolerated | 14.26 | Neutral |
| p.Tyr44Glu | VUS | -13.61 | Indeterminate | 0.8125 | Pathogenic | -13.47 | Pathogenic | 0.31 | Neutral | 0.99 | Probably damaging | 0.16 | Tolerated | | |
| p.Tyr44Ala | VUS | -26.29 | Indeterminate | 0.5926 | Pathogenic | -11.13 | Pathogenic | 0.29 | Neutral | 0.98 | Probably damaging | 0.25 | Tolerated | | |
| p.Tyr44Gly | VUS | -11.44 | Indeterminate | 0.5878 | Pathogenic | -12.28 | Pathogenic | 0.30 | Neutral | 1.00 | Probably damaging | 0.22 | Tolerated | | |
| p.Tyr44Val | VUS | -0.88 | Neutral | 0.6303 | Pathogenic | -12.57 | Pathogenic | 0.32 | Neutral | 0.86 | Possibly damaging | 0.22 | Tolerated | | |
| p.Tyr44Cys | VUS | -2.08 | Neutral | 0.3951 | Ambiguous | -9.38 | Pathogenic | 0.20 | Neutral | 1.00 | Probably damaging | 0.08 | Tolerated | 17.56 | Deleterious |
| p.Tyr44Trp | VUS | -1.96 | Neutral | 0.3594 | Ambiguous | -11.11 | Pathogenic | 0.29 | Neutral | 0.96 | Probably damaging | 0.12 | Tolerated | | |
| p.Tyr44Phe | VUS | -2.38 | Neutral | 0.1446 | Benign | -3.70 | Benign | 0.18 | Neutral | 0.00 | Benign | 1 | Tolerated | 0.48 | Neutral |
| p.Gly45Asn | VUS | -1.75 | Neutral | 0.7484 | Pathogenic | -11.78 | Pathogenic | 0.77 | Disease | 1.00 | Probably damaging | 0.02 | Deleterious | | |
| p.Gly45Lys | VUS | -3.95 | Neutral | 0.877 | Pathogenic | -15.08 | Pathogenic | 0.79 | Disease | 1.00 | Probably damaging | 0.12 | Tolerated | | |
| p.Gly45Thr | VUS | -1.80 | Neutral | 0.8377 | Pathogenic | -13.43 | Pathogenic | 0.77 | Disease | 1.00 | Probably damaging | 0.01 | Deleterious | | |
| p.Gly45Arg | VUS | -0.50 | Neutral | 0.6263 | Pathogenic | -12.97 | Pathogenic | 0.83 | Disease | 1.00 | Probably damaging | 0.02 | Deleterious | 26.00 | Deleterious |
| p.Gly45Ser | VUS | -10.54 | Indeterminate | 0.4477 | Ambiguous | -10.50 | Pathogenic | 0.83 | Disease | 1.00 | Probably damaging | 0.03 | Deleterious | 26.00 | Deleterious |
| p.Gly45Ile | VUS | -13.67 | Indeterminate | 0.9659 | Pathogenic | -18.24 | Pathogenic | 0.79 | Disease | 1.00 | Probably damaging | 0 | Deleterious | | |
| p.Gly45Met | VUS | -2.56 | Neutral | 0.9583 | Pathogenic | -15.46 | Pathogenic | 0.79 | Disease | 1.00 | Probably damaging | 0 | Deleterious | | |
| p.Gly45His | VUS | -2.01 | Neutral | 0.7485 | Pathogenic | -14.87 | Pathogenic | 0.77 | Disease | 1.00 | Probably damaging | 0.01 | Deleterious | | |
| p.Gly45Gln | VUS | -0.28 | Neutral | 0.6642 | Pathogenic | -14.02 | Pathogenic | 0.78 | Disease | 1.00 | Probably damaging | 0.01 | Deleterious | | |
| p.Gly45Pro | VUS | -53.15 | Deleterious | 0.9945 | Pathogenic | -18.63 | Pathogenic | 0.76 | Disease | 1.00 | Probably damaging | 0.01 | Deleterious | | |
| p.Gly45Leu | VUS | -0.82 | Neutral | 0.8676 | Pathogenic | -14.32 | Pathogenic | 0.78 | Disease | 1.00 | Probably damaging | 0.01 | Deleterious | | |
| p.Gly45Asp | VUS | -0.26 | Neutral | 0.7037 | Pathogenic | -10.05 | Pathogenic | 0.86 | Disease | 1.00 | Probably damaging | 0.02 | Deleterious | 25.50 | Deleterious |
| p.Gly45Glu | VUS | -1.15 | Neutral | 0.8048 | Pathogenic | -12.40 | Pathogenic | 0.88 | Disease | 1.00 | Probably damaging | 0.01 | Deleterious | | |
| p.Gly45Ala | VUS | -2.19 | Neutral | 0.5934 | Pathogenic | -11.16 | Pathogenic | 0.83 | Disease | 1.00 | Probably damaging | 0.03 | Deleterious | 24.90 | Deleterious |
| p.Gly45Val | VUS | -21.77 | Indeterminate | 0.9261 | Pathogenic | -14.58 | Pathogenic | 0.87 | Disease | 1.00 | Probably damaging | 0.01 | Deleterious | 25.70 | Deleterious |
| p.Gly45Tyr | VUS | -3.37 | Neutral | 0.8664 | Pathogenic | -16.39 | Pathogenic | 0.80 | Disease | 1.00 | Probably damaging | 0 | Deleterious | | |
| p.Gly45Cys | VUS | -0.52 | Neutral | 0.7436 | Pathogenic | -13.81 | Pathogenic | 0.89 | Disease | 1.00 | Probably damaging | 0 | Deleterious | 26.90 | Deleterious |
| p.Gly45Trp | VUS | -3.99 | Neutral | 0.8636 | Pathogenic | -14.99 | Pathogenic | 0.85 | Disease | 1.00 | Probably damaging | 0 | Deleterious | | |
| p.Gly45Phe | VUS | -0.09 | Neutral | 0.9352 | Pathogenic | -16.79 | Pathogenic | 0.80 | Disease | 1.00 | Probably damaging | 0 | Deleterious | | |
| p.Arg46Asn | VUS | -0.71 | Neutral | 0.9806 | Pathogenic | -15.60 | Pathogenic | 0.67 | Disease | 1.00 | Probably damaging | 0.18 | Tolerated | | |
| p.Arg46Lys | VUS | -0.41 | Neutral | 0.731 | Pathogenic | -13.24 | Pathogenic | 0.48 | Neutral | 1.00 | Probably damaging | 0.58 | Tolerated | | |
| p.Arg46Thr | VUS | -33.07 | Indeterminate | 0.9723 | Pathogenic | -15.00 | Pathogenic | 0.80 | Disease | 1.00 | Probably damaging | 0.08 | Tolerated | | |
| p.Arg46Ser | VUS | -0.32 | Neutral | 0.9811 | Pathogenic | -13.34 | Pathogenic | 0.80 | Disease | 1.00 | Probably damaging | 0.06 | Tolerated | | |
| p.Arg46Ile | VUS | -53.15 | Deleterious | 0.9128 | Pathogenic | -17.10 | Pathogenic | 0.74 | Disease | 1.00 | Probably damaging | 0.03 | Deleterious | | |
| p.Arg46Met | VUS | -10.71 | Indeterminate | 0.9584 | Pathogenic | -16.82 | Pathogenic | 0.72 | Disease | 1.00 | Probably damaging | 0.02 | Deleterious | | |
| p.Arg46His | VUS | -0.53 | Neutral | 0.6345 | Pathogenic | -13.33 | Pathogenic | 0.66 | Disease | 1.00 | Probably damaging | 0.06 | Tolerated | | |
| p.Arg46Gln | VUS | -14.28 | Indeterminate | 0.5586 | Ambiguous | -13.16 | Pathogenic | 0.62 | Disease | 1.00 | Probably damaging | 0.16 | Tolerated | 26.20 | Deleterious |
| p.Arg46Pro | VUS | -53.15 | Deleterious | 0.9714 | Pathogenic | -16.65 | Pathogenic | 0.87 | Disease | 1.00 | Probably damaging | 0.05 | Deleterious | 25.50 | Deleterious |
| p.Arg46Leu | VUS | -5.97 | Indeterminate | 0.7987 | Pathogenic | -15.22 | Pathogenic | 0.81 | Disease | 1.00 | Probably damaging | 0.21 | Tolerated | 25.70 | Deleterious |
| p.Arg46Asp | VUS | -53.15 | Deleterious | 0.9842 | Pathogenic | -18.85 | Pathogenic | 0.77 | Disease | 1.00 | Probably damaging | 0.02 | Deleterious | | |
| p.Arg46Glu | VUS | -53.15 | Deleterious | 0.9098 | Pathogenic | -17.42 | Pathogenic | 0.68 | Disease | 1.00 | Probably damaging | 0.1 | Tolerated | | |
| p.Arg46Ala | VUS | -10.45 | Indeterminate | 0.9674 | Pathogenic | -13.76 | Pathogenic | 0.66 | Disease | 1.00 | Probably damaging | 0.07 | Tolerated | | |
| p.Arg46Gly | VUS | -6.54 | Indeterminate | 0.933 | Pathogenic | -11.73 | Pathogenic | 0.82 | Disease | 1.00 | Probably damaging | 0.05 | Deleterious | 23.50 | Deleterious |
| p.Arg46Val | VUS | -27.58 | Indeterminate | 0.9279 | Pathogenic | -16.09 | Pathogenic | 0.73 | Disease | 1.00 | Probably damaging | 0.04 | Deleterious | | |
| p.Arg46Tyr | VUS | -14.34 | Indeterminate | 0.883 | Pathogenic | -15.10 | Pathogenic | 0.70 | Disease | 1.00 | Probably damaging | 0.03 | Deleterious | | |
| p.Arg46Cys | VUS | -5.51 | Neutral | 0.8016 | Pathogenic | -11.14 | Pathogenic | 0.82 | Disease | 1.00 | Probably damaging | 0.01 | Deleterious | | |
| p.Arg46Trp | VUS | -0.33 | Neutral | 0.6392 | Pathogenic | -12.93 | Pathogenic | 0.80 | Disease | 1.00 | Probably damaging | 0.01 | Deleterious | 24.50 | Deleterious |
| p.Arg46Phe | VUS | -1.84 | Neutral | 0.9478 | Pathogenic | -15.17 | Pathogenic | 0.71 | Disease | 1.00 | Probably damaging | 0.02 | Deleterious | | |
| p.Arg47Asn | VUS | -2.20 | Neutral | 0.7485 | Pathogenic | -9.56 | Pathogenic | 0.35 | Neutral | 0.79 | Possibly damaging | 0.13 | Tolerated | | |
| p.Arg47Lys | VUS | -2.24 | Neutral | 0.3095 | Benign | -7.65 | Pathogenic | 0.14 | Neutral | 0.25 | Benign | 0.14 | Tolerated | 14.41 | Neutral |
| p.Arg47Thr | VUS | -1.27 | Neutral | 0.3093 | Benign | -5.10 | Benign | 0.15 | Neutral | 0.01 | Benign | 1 | Tolerated | 2.99 | Neutral |
| p.Arg47Ser | VUS | -8.25 | Indeterminate | 0.5326 | Ambiguous | -6.22 | Benign | 0.11 | Neutral | 0.05 | Benign | 0.52 | Tolerated | 9.65 | Neutral |
| p.Arg47Ile | VUS | -3.02 | Neutral | 0.6194 | Pathogenic | -11.61 | Pathogenic | 0.30 | Neutral | 0.98 | Probably damaging | 0.05 | Deleterious | | |
| p.Arg47Met | VUS | -1.94 | Neutral | 0.5667 | Pathogenic | -7.27 | Benign | 0.33 | Neutral | 0.96 | Probably damaging | 0.03 | Deleterious | 18.80 | Deleterious |
| p.Arg47His | VUS | -2.42 | Neutral | 0.2039 | Benign | -5.44 | Benign | 0.19 | Neutral | 0.97 | Probably damaging | 0.03 | Deleterious | | |
| p.Arg47Gln | VUS | -10.65 | Indeterminate | 0.2172 | Benign | -6.44 | Benign | 0.16 | Neutral | 0.87 | Possibly damaging | 0.08 | Tolerated | | |
| p.Arg47Pro | VUS | -11.61 | Indeterminate | 0.7548 | Pathogenic | -8.24 | Pathogenic | 0.29 | Neutral | 0.99 | Probably damaging | 0.09 | Tolerated | | |
| p.Arg47Leu | VUS | -0.37 | Neutral | 0.3066 | Benign | -5.62 | Benign | 0.24 | Neutral | 0.90 | Possibly damaging | 0.05 | Deleterious | | |

| | | | | | | | | | | | | | | | |
|---|---|---|---|---|---|---|---|---|---|---|---|---|---|---|---|
| p.Arg47Asp | VUS | -53.15 | Deleterious | 0.9177 | Pathogenic | -15.01 | Pathogenic | 0.37 | Neutral | 0.98 | Probably damaging | 0.08 | Tolerated | | |
| p.Arg47Glu | VUS | -3.92 | Neutral | 0.748 | Pathogenic | -13.09 | Pathogenic | 0.30 | Neutral | 0.53 | Possibly damaging | 0.06 | Tolerated | | |
| p.Arg47Ala | VUS | -2.77 | Neutral | 0.5277 | Ambiguous | -7.58 | Pathogenic | 0.30 | Neutral | 0.45 | Possibly damaging | 0.18 | Tolerated | | |
| p.Arg47Gly | VUS | -9.18 | Indeterminate | 0.4602 | Ambiguous | -10.59 | Pathogenic | 0.16 | Neutral | 0.86 | Possibly damaging | 0.07 | Tolerated | 16.41 | Deleterious |
| p.Arg47Val | VUS | -1.23 | Neutral | 0.5757 | Pathogenic | -9.79 | Pathogenic | 0.32 | Neutral | 0.94 | Possibly damaging | 0.08 | Tolerated | | |
| p.Arg47Tyr | VUS | -1.99 | Neutral | 0.5044 | Ambiguous | -5.05 | Benign | 0.30 | Neutral | 0.85 | Possibly damaging | 0.02 | Deleterious | | |
| p.Arg47Cys | VUS | -0.31 | Neutral | 0.2597 | Benign | -7.45 | Benign | 0.15 | Neutral | 1.00 | Probably damaging | 0.03 | Deleterious | | |
| p.Arg47Trp | VUS | -3.86 | Neutral | 0.2964 | Benign | -8.90 | Pathogenic | 0.26 | Neutral | 0.98 | Probably damaging | 0.01 | Deleterious | 22.40 | Deleterious |
| p.Arg47Phe | VUS | -0.20 | Neutral | 0.7065 | Pathogenic | -7.03 | Benign | 0.32 | Neutral | 0.65 | Possibly damaging | 0.02 | Deleterious | | |
| p.Pro48Asn | VUS | -53.15 | Deleterious | 0.9787 | Pathogenic | -17.26 | Pathogenic | 0.64 | Disease | 0.96 | Possibly damaging | 0.04 | Deleterious | | |
| p.Pro48Lys | VUS | -53.15 | Deleterious | 0.9943 | Pathogenic | -20.62 | Pathogenic | 0.70 | Disease | 0.99 | Probably damaging | 0.06 | Tolerated | | |
| p.Pro48Thr | Pathogenic | -19.30 | Indeterminate | 0.8084 | Pathogenic | -11.01 | Pathogenic | 0.61 | Disease | 0.69 | Possibly damaging | 0.11 | Tolerated | 16.24 | Deleterious |
| p.Pro48Arg | VUS | -53.15 | Deleterious | 0.9677 | Pathogenic | -16.85 | Pathogenic | 0.73 | Disease | 1.00 | Probably damaging | 0.04 | Deleterious | 25.60 | Deleterious |
| p.Pro48Ser | VUS | -4.30 | Neutral | 0.6914 | Pathogenic | -9.33 | Pathogenic | 0.57 | Disease | 0.93 | Possibly damaging | 0.29 | Tolerated | 16.81 | Deleterious |
| p.Pro48Ile | VUS | -53.15 | Deleterious | 0.9751 | Pathogenic | -16.41 | Pathogenic | 0.66 | Disease | 0.99 | Probably damaging | 0.04 | Deleterious | | |
| p.Pro48Met | VUS | -53.15 | Deleterious | 0.9761 | Pathogenic | -15.41 | Pathogenic | 0.65 | Disease | 1.00 | Probably damaging | 0.03 | Deleterious | | |
| p.Pro48His | VUS | -53.15 | Deleterious | 0.9692 | Pathogenic | -17.19 | Pathogenic | 0.64 | Disease | 1.00 | Probably damaging | 0.01 | Deleterious | | |
| p.Pro48Gln | VUS | -53.15 | Deleterious | 0.9466 | Pathogenic | -16.15 | Pathogenic | 0.63 | Disease | 0.99 | Probably damaging | 0.04 | Deleterious | 26.00 | Deleterious |
| p.Pro48Leu | VUS | -53.15 | Deleterious | 0.8968 | Pathogenic | -13.24 | Pathogenic | 0.75 | Disease | 0.98 | Probably damaging | 0.06 | Tolerated | 26.60 | Deleterious |
| p.Pro48Asp | VUS | -53.15 | Deleterious | 0.9925 | Pathogenic | -18.20 | Pathogenic | 0.68 | Disease | 0.95 | Possibly damaging | 0.05 | Deleterious | | |
| p.Pro48Glu | VUS | -53.15 | Deleterious | 0.9801 | Pathogenic | -16.52 | Pathogenic | 0.66 | Disease | 0.72 | Possibly damaging | 0.06 | Tolerated | | |
| p.Pro48Ala | VUS | -0.04 | Neutral | 0.314 | Benign | -5.17 | Benign | 0.33 | Neutral | 0.01 | Benign | 1 | Tolerated | 7.25 | Neutral |
| p.Pro48Gly | VUS | -14.99 | Indeterminate | 0.8399 | Pathogenic | -12.29 | Pathogenic | 0.49 | Neutral | 0.93 | Possibly damaging | 0.31 | Tolerated | | |
| p.Pro48Val | VUS | -17.05 | Indeterminate | 0.8808 | Pathogenic | -11.13 | Pathogenic | 0.63 | Disease | 0.68 | Possibly damaging | 0.11 | Tolerated | | |
| p.Pro48Tyr | VUS | -53.15 | Deleterious | 0.9952 | Pathogenic | -19.09 | Pathogenic | 0.65 | Disease | 1.00 | Probably damaging | 0.02 | Deleterious | | |
| p.Pro48Cys | VUS | -29.75 | Indeterminate | 0.9269 | Pathogenic | -12.01 | Pathogenic | 0.64 | Disease | 1.00 | Probably damaging | 0.06 | Tolerated | | |
| p.Pro48Trp | VUS | -53.15 | Deleterious | 0.9966 | Pathogenic | -19.53 | Pathogenic | 0.65 | Disease | 1.00 | Probably damaging | 0.01 | Deleterious | | |
| p.Pro48Phe | VUS | -53.15 | Deleterious | 0.9958 | Pathogenic | -17.68 | Pathogenic | 0.66 | Disease | 0.99 | Probably damaging | 0.02 | Deleterious | | |
| p.Ile49Asn | VUS | -53.15 | Deleterious | 0.9737 | Pathogenic | -15.57 | Pathogenic | 0.87 | Disease | 1.00 | Probably damaging | 0 | Deleterious | 26.40 | Deleterious |
| p.Ile49Lys | VUS | -53.15 | Deleterious | 0.9874 | Pathogenic | -19.12 | Pathogenic | 0.84 | Disease | 0.98 | Probably damaging | 0 | Deleterious | | |
| p.Ile49Thr | Likely pathogenic | -29.95 | Indeterminate | 0.9628 | Pathogenic | -12.04 | Pathogenic | 0.68 | Disease | 1.00 | Probably damaging | 0.01 | Deleterious | 25.70 | Deleterious |
| p.Ile49Arg | VUS | -53.15 | Deleterious | 0.9636 | Pathogenic | -15.14 | Pathogenic | 0.81 | Disease | 1.00 | Probably damaging | 0 | Deleterious | | |
| p.Ile49Ser | Likely pathogenic | -53.15 | Deleterious | 0.941 | Pathogenic | -14.12 | Pathogenic | 0.87 | Disease | 1.00 | Probably damaging | 0 | Deleterious | 26.50 | Deleterious |
| p.Ile49Met | VUS | -1.12 | Neutral | 0.4493 | Ambiguous | -9.30 | Pathogenic | 0.61 | Disease | 1.00 | Probably damaging | 0.04 | Deleterious | 19.76 | Deleterious |
| p.Ile49His | VUS | -53.15 | Deleterious | 0.9776 | Pathogenic | -14.20 | Pathogenic | 0.79 | Disease | 1.00 | Probably damaging | 0 | Deleterious | | |
| p.Ile49Gln | VUS | -53.15 | Deleterious | 0.9677 | Pathogenic | -13.76 | Pathogenic | 0.81 | Disease | 1.00 | Probably damaging | 0.01 | Deleterious | | |
| p.Ile49Pro | VUS | -53.15 | Deleterious | 0.9578 | Pathogenic | -13.00 | Pathogenic | 0.77 | Disease | 1.00 | Probably damaging | 0 | Deleterious | | |
| p.Ile49Leu | VUS | -0.37 | Neutral | 0.2537 | Benign | -1.42 | Benign | 0.30 | Neutral | 0.93 | Possibly damaging | 1 | Tolerated | 17.23 | Deleterious |
| p.Ile49Asp | VUS | -53.15 | Deleterious | 0.9963 | Pathogenic | -18.44 | Pathogenic | 0.79 | Disease | 1.00 | Probably damaging | 0 | Deleterious | | |
| p.Ile49Glu | VUS | -53.15 | Deleterious | 0.9845 | Pathogenic | -17.50 | Pathogenic | 0.79 | Disease | 1.00 | Probably damaging | 0 | Deleterious | | |
| p.Ile49Ala | VUS | -29.55 | Indeterminate | 0.9073 | Pathogenic | -13.41 | Pathogenic | 0.64 | Disease | 1.00 | Probably damaging | 0.01 | Deleterious | | |
| p.Ile49Gly | VUS | -53.15 | Deleterious | 0.9822 | Pathogenic | -16.64 | Pathogenic | 0.73 | Disease | 1.00 | Probably damaging | 0 | Deleterious | | |
| p.Ile49Val | VUS | -0.79 | Neutral | 0.3085 | Benign | -6.85 | Benign | 0.41 | Neutral | 0.95 | Possibly damaging | 0.06 | Tolerated | 19.03 | Deleterious |
| p.Ile49Tyr | VUS | -53.15 | Deleterious | 0.9512 | Pathogenic | -16.37 | Pathogenic | 0.69 | Disease | 0.99 | Probably damaging | 0 | Deleterious | | |
| p.Ile49Cys | VUS | -19.59 | Indeterminate | 0.9731 | Pathogenic | -14.94 | Pathogenic | 0.69 | Disease | 1.00 | Probably damaging | 0 | Deleterious | | |
| p.Ile49Trp | VUS | -53.15 | Deleterious | 0.9727 | Pathogenic | -16.39 | Pathogenic | 0.71 | Disease | 1.00 | Probably damaging | 0 | Deleterious | | |
| p.Ile49Phe | VUS | -27.97 | Indeterminate | 0.6763 | Pathogenic | -11.84 | Pathogenic | 0.75 | Disease | 1.00 | Probably damaging | 0.03 | Deleterious | 24.40 | Deleterious |
| p.Gln50Asn | VUS | -12.49 | Indeterminate | 0.8074 | Pathogenic | -10.43 | Pathogenic | 0.70 | Disease | 1.00 | Probably damaging | 0.03 | Deleterious | | |
| p.Gln50Lys | VUS | -53.15 | Deleterious | 0.9228 | Pathogenic | -17.58 | Pathogenic | 0.76 | Disease | 0.93 | Possibly damaging | 0.07 | Tolerated | 26.30 | Deleterious |
| p.Gln50Thr | VUS | -10.88 | Indeterminate | 0.6796 | Pathogenic | -13.74 | Pathogenic | 0.68 | Disease | 0.96 | Probably damaging | 0.02 | Deleterious | | |
| p.Gln50Arg | VUS | -53.15 | Deleterious | 0.84 | Pathogenic | -14.09 | Pathogenic | 0.78 | Disease | 0.96 | Possibly damaging | 0.05 | Deleterious | 28.50 | Deleterious |
| p.Gln50Ser | VUS | -5.39 | Neutral | 0.555 | Ambiguous | -12.73 | Pathogenic | 0.68 | Disease | 1.00 | Probably damaging | 0.02 | Deleterious | | |
| p.Gln50Ile | VUS | -53.15 | Deleterious | 0.9391 | Pathogenic | -17.14 | Pathogenic | 0.75 | Disease | 0.99 | Probably damaging | 0.01 | Deleterious | | |
| p.Gln50Met | VUS | -2.05 | Neutral | 0.8088 | Pathogenic | -14.73 | Pathogenic | 0.70 | Disease | 0.99 | Probably damaging | 0 | Deleterious | | |
| p.Gln50His | VUS | -53.15 | Deleterious | 0.7198 | Pathogenic | -3.60 | Benign | 0.75 | Disease | 0.99 | Probably damaging | 0.27 | Tolerated | 33.00 | Deleterious |
| p.Gln50Pro | VUS | -53.15 | Deleterious | 0.7863 | Pathogenic | -11.25 | Pathogenic | 0.86 | Disease | 1.00 | Probably damaging | 0.02 | Deleterious | 27.60 | Deleterious |
| p.Gln50Leu | VUS | -19.95 | Indeterminate | 0.7182 | Pathogenic | -13.31 | Pathogenic | 0.80 | Disease | 0.97 | Probably damaging | 0.01 | Deleterious | 29.80 | Deleterious |
| p.Gln50Asp | VUS | -53.15 | Deleterious | 0.8569 | Pathogenic | -12.67 | Pathogenic | 0.73 | Disease | 1.00 | Probably damaging | 0.03 | Deleterious | | |
| p.Gln50Glu | VUS | -1.37 | Neutral | 0.1941 | Benign | -13.92 | Pathogenic | 0.74 | Disease | 1.00 | Probably damaging | 0.04 | Deleterious | 25.50 | Deleterious |
| p.Gln50Ala | VUS | -10.39 | Indeterminate | 0.5825 | Pathogenic | -13.36 | Pathogenic | 0.72 | Disease | 1.00 | Probably damaging | 0.02 | Deleterious | | |
| p.Gln50Gly | VUS | -13.64 | Indeterminate | 0.7192 | Pathogenic | -12.93 | Pathogenic | 0.70 | Disease | 1.00 | Probably damaging | 0.03 | Deleterious | | |
| p.Gln50Val | VUS | -53.15 | Deleterious | 0.87 | Pathogenic | -15.58 | Pathogenic | 0.74 | Disease | 1.00 | Probably damaging | 0.01 | Deleterious | | |
| p.Gln50Tyr | VUS | -53.15 | Deleterious | 0.9728 | Pathogenic | -11.61 | Pathogenic | 0.76 | Disease | 1.00 | Probably damaging | 0.01 | Deleterious | | |
| p.Gln50Cys | VUS | -22.37 | Indeterminate | 0.9729 | Pathogenic | -15.46 | Pathogenic | 0.75 | Disease | 1.00 | Probably damaging | 0 | Deleterious | | |
| p.Gln50Trp | VUS | -53.15 | Deleterious | 0.9737 | Pathogenic | -15.75 | Pathogenic | 0.77 | Disease | 1.00 | Probably damaging | 0 | Deleterious | | |
| p.Gln50Phe | VUS | -53.15 | Deleterious | 0.9641 | Pathogenic | -15.27 | Pathogenic | 0.75 | Disease | 1.00 | Probably damaging | 0 | Deleterious | | |
| p.Val51Asn | VUS | -53.15 | Deleterious | 0.8296 | Pathogenic | -10.66 | Pathogenic | 0.78 | Disease | 1.00 | Probably damaging | 0.01 | Deleterious | | |
| p.Val51Lys | VUS | -53.15 | Deleterious | 0.9357 | Pathogenic | -12.52 | Pathogenic | 0.81 | Disease | 1.00 | Probably damaging | 0.01 | Deleterious | | |
| p.Val51Thr | VUS | -4.90 | Neutral | 0.6936 | Pathogenic | -7.68 | Pathogenic | 0.55 | Disease | 1.00 | Probably damaging | 0.03 | Deleterious | | |
| p.Val51Arg | VUS | -53.15 | Deleterious | 0.8385 | Pathogenic | -11.71 | Pathogenic | 0.81 | Disease | 1.00 | Probably damaging | 0.01 | Deleterious | | |
| p.Val51Ser | VUS | -0.22 | Neutral | 0.7805 | Pathogenic | -8.95 | Pathogenic | 0.73 | Disease | 1.00 | Probably damaging | 0.01 | Deleterious | | |
| p.Val51Ile | VUS | -5.58 | Neutral | 0.2785 | Benign | -7.63 | Pathogenic | 0.53 | Disease | 0.96 | Probably damaging | 0.41 | Tolerated | 28.50 | Deleterious |
| p.Val51Met | VUS | -53.15 | Deleterious | 0.7295 | Pathogenic | -9.08 | Pathogenic | 0.68 | Disease | 1.00 | Probably damaging | 0.02 | Deleterious | | |
| p.Val51His | VUS | -53.15 | Deleterious | 0.9476 | Pathogenic | -13.56 | Pathogenic | 0.80 | Disease | 1.00 | Probably damaging | 0 | Deleterious | | |
| p.Val51Gln | VUS | -53.15 | Deleterious | 0.8361 | Pathogenic | -11.73 | Pathogenic | 0.78 | Disease | 1.00 | Probably damaging | 0.01 | Deleterious | | |
| p.Val51Pro | VUS | -53.15 | Deleterious | 0.9692 | Pathogenic | -12.62 | Pathogenic | 0.80 | Disease | 1.00 | Probably damaging | 0.01 | Deleterious | | |
| p.Val51Leu | VUS | -1.84 | Neutral | 0.7283 | Pathogenic | -8.30 | Pathogenic | 0.75 | Disease | 0.96 | Probably damaging | 0.11 | Tolerated | 31.00 | Deleterious |
| p.Val51Asp | VUS | -53.15 | Deleterious | 0.9045 | Pathogenic | -15.14 | Pathogenic | 0.88 | Disease | 1.00 | Probably damaging | 0.01 | Deleterious | 26.90 | Deleterious |
| p.Val51Glu | VUS | -53.15 | Deleterious | 0.8206 | Pathogenic | -14.07 | Pathogenic | 0.86 | Disease | 1.00 | Probably damaging | 0.01 | Deleterious | | |
| p.Val51Ala | VUS | -0.01 | Neutral | 0.6612 | Pathogenic | -6.73 | Benign | 0.75 | Disease | 1.00 | Probably damaging | 0.04 | Deleterious | 26.30 | Deleterious |
| p.Val51Gly | VUS | -3.26 | Neutral | 0.5831 | Pathogenic | -8.94 | Pathogenic | 0.68 | Disease | 1.00 | Probably damaging | 0.01 | Deleterious | 26.40 | Deleterious |
| p.Val51Tyr | VUS | -53.15 | Deleterious | 0.909 | Pathogenic | -14.19 | Pathogenic | 0.77 | Disease | 1.00 | Probably damaging | 0.12 | Tolerated | | |
| p.Val51Cys | VUS | -0.37 | Neutral | 0.93 | Pathogenic | -11.25 | Pathogenic | 0.76 | Disease | 1.00 | Probably damaging | 0.16 | Tolerated | | |
| p.Val51Trp | VUS | -53.15 | Deleterious | 0.9773 | Pathogenic | -13.51 | Pathogenic | 0.76 | Disease | 1.00 | Probably damaging | 0 | Deleterious | | |
| p.Val51Phe | VUS | -53.15 | Deleterious | 0.7444 | Pathogenic | -12.38 | Pathogenic | 0.81 | Disease | 1.00 | Probably damaging | 0.03 | Deleterious | 32.00 | Deleterious |
| p.Met52Asn | VUS | -16.42 | Indeterminate | 0.9627 | Pathogenic | -13.68 | Pathogenic | 0.72 | Disease | 1.00 | Probably damaging | 0.14 | Tolerated | | |
| p.Met52Lys | VUS | -53.15 | Deleterious | 0.954 | Pathogenic | -14.14 | Pathogenic | 0.82 | Disease | 0.50 | Possibly damaging | 0.14 | Tolerated | 27.90 | Deleterious |
| p.Met52Thr | VUS | -1.17 | Neutral | 0.7826 | Pathogenic | -9.07 | Pathogenic | 0.77 | Disease | 0.82 | Possibly damaging | 0.5 | Tolerated | 26.40 | Deleterious |
| p.Met52Arg | VUS | -53.15 | Deleterious | 0.9272 | Pathogenic | -12.64 | Pathogenic | 0.82 | Disease | 0.68 | Possibly damaging | 0.1 | Tolerated | 28.50 | Deleterious |
| p.Met52Ser | VUS | -0.07 | Neutral | 0.8548 | Pathogenic | -11.63 | Pathogenic | 0.72 | Disease | 0.98 | Probably damaging | 0.72 | Tolerated | | |
| p.Met52Ile | VUS | -7.82 | Indeterminate | 0.9224 | Pathogenic | -11.19 | Pathogenic | 0.74 | Disease | 0.09 | Benign | 0.16 | Tolerated | 24.97 | Deleterious |
| p.Met52His | VUS | -4.19 | Neutral | 0.9567 | Pathogenic | -12.75 | Pathogenic | 0.74 | Disease | 1.00 | Probably damaging | 0.04 | Deleterious | | |
| p.Met52Gln | VUS | -1.31 | Neutral | 0.7907 | Pathogenic | -9.56 | Pathogenic | 0.70 | Disease | 0.99 | Probably damaging | 0.1 | Tolerated | | |
| p.Met52Pro | VUS | -13.05 | Indeterminate | 0.9812 | Pathogenic | -11.77 | Pathogenic | 0.73 | Disease | 0.99 | Probably damaging | 0.26 | Tolerated | | |
| p.Met52Leu | VUS | -0.01 | Neutral | 0.3933 | Ambiguous | -6.10 | Benign | 0.66 | Disease | 0.02 | Benign | 0.28 | Tolerated | 23.35 | Deleterious |
| p.Met52Asp | VUS | -28.20 | Indeterminate | 0.9925 | Pathogenic | -17.57 | Pathogenic | 0.71 | Disease | 0.99 | Probably damaging | 0.11 | Tolerated | | |
| p.Met52Glu | VUS | -11.11 | Indeterminate | 0.9593 | Pathogenic | -14.71 | Pathogenic | 0.71 | Disease | 0.98 | Probably damaging | 0.11 | Tolerated | | |
| p.Met52Ala | VUS | -0.01 | Neutral | 0.6987 | Pathogenic | -9.19 | Pathogenic | 0.72 | Disease | 0.54 | Possibly damaging | 1 | Tolerated | | |
| p.Met52Gly | VUS | 0.00 | Neutral | 0.9223 | Pathogenic | -13.23 | Pathogenic | 0.70 | Disease | 0.99 | Probably damaging | 0.5 | Tolerated | | |
| p.Met52Val | VUS | -0.01 | Neutral | 0.4159 | Ambiguous | -8.75 | Pathogenic | 0.46 | Neutral | 0.20 | Benign | 0.32 | Tolerated | 22.80 | Deleterious |
| p.Met52Tyr | VUS | -8.17 | Indeterminate | 0.9589 | Pathogenic | -16.79 | Pathogenic | 0.70 | Disease | 0.93 | Possibly damaging | 0.07 | Tolerated | | |
| p.Met52Cys | VUS | -0.01 | Neutral | 0.8433 | Pathogenic | -11.90 | Pathogenic | 0.70 | Disease | 1.00 | Probably damaging | 0.25 | Tolerated | | |
| p.Met52Trp | VUS | -9.17 | Indeterminate | 0.9696 | Pathogenic | -14.81 | Pathogenic | 0.70 | Disease | 1.00 | Probably damaging | 0.03 | Deleterious | | |
| p.Met52Phe | VUS | -4.81 | Neutral | 0.7519 | Pathogenic | -12.44 | Pathogenic | 0.66 | Disease | 0.14 | Benign | 0.09 | Tolerated | | |
| p.Met53Asn | VUS | -32.99 | Indeterminate | 0.9195 | Pathogenic | -15.29 | Pathogenic | 0.72 | Disease | 1.00 | Probably damaging | 0.3 | Tolerated | | |
| p.Met53Lys | VUS | -8.88 | Indeterminate | 0.8127 | Pathogenic | -11.47 | Pathogenic | 0.51 | Disease | 0.93 | Possibly damaging | 0.59 | Tolerated | 24.50 | Deleterious |
| p.Met53Thr | VUS | -33.22 | Indeterminate | 0.8028 | Pathogenic | -10.33 | Pathogenic | 0.76 | Disease | 1.00 | Probably damaging | 0.41 | Tolerated | 25.80 | Deleterious |
| p.Met53Arg | VUS | -3.54 | Neutral | 0.6785 | Pathogenic | -10.14 | Pathogenic | 0.58 | Disease | 0.99 | Probably damaging | 0.42 | Tolerated | 26.60 | Deleterious |
| p.Met53Ser | VUS | -6.07 | Indeterminate | 0.8555 | Pathogenic | -9.89 | Pathogenic | 0.72 | Disease | 1.00 | Probably damaging | 0.52 | Tolerated | | |
| p.Met53Ile | Pathogenic | -53.15 | Deleterious | 0.9254 | Pathogenic | -10.44 | Pathogenic | 0.64 | Disease | 0.94 | Possibly damaging | 0.59 | Tolerated | 28.80 | Deleterious |
| p.Met53His | VUS | -33.18 | Indeterminate | 0.889 | Pathogenic | -13.75 | Pathogenic | 0.73 | Disease | 1.00 | Probably damaging | 0.48 | Tolerated | | |

| | | | | | | | | | | | | | | | |
|---|---|---|---|---|---|---|---|---|---|---|---|---|---|---|---|
| p.Met53Gln | VUS | -1.49 | Neutral | 0.4944 | Ambiguous | -10.26 | Pathogenic | 0.49 | Neutral | 1.00 | Probably damaging | 0.31 | Tolerated | | |
| p.Met53Pro | VUS | -53.15 | Deleterious | 0.9676 | Pathogenic | -10.04 | Pathogenic | 0.72 | Disease | 1.00 | Probably damaging | 0.2 | Tolerated | | |
| p.Met53Leu | VUS | -6.97 | Indeterminate | 0.3533 | Ambiguous | -5.70 | Benign | 0.61 | Disease | 0.88 | Possibly damaging | 0.67 | Tolerated | 25.65 | Deleterious |
| p.Met53Asp | VUS | -53.15 | Deleterious | 0.9768 | Pathogenic | -16.24 | Pathogenic | 0.70 | Disease | 1.00 | Probably damaging | 0.2 | Tolerated | | |
| p.Met53Glu | VUS | -53.15 | Deleterious | 0.8544 | Pathogenic | -13.12 | Pathogenic | 0.70 | Disease | 1.00 | Probably damaging | 0.29 | Tolerated | | |
| p.Met53Ala | VUS | -4.27 | Neutral | 0.7197 | Pathogenic | -9.61 | Pathogenic | 0.69 | Disease | 1.00 | Probably damaging | 0.5 | Tolerated | | |
| p.Met53Gly | VUS | -53.15 | Deleterious | 0.8671 | Pathogenic | -12.13 | Pathogenic | 0.70 | Disease | 0.55 | Possibly damaging | 0.31 | Tolerated | | |
| p.Met53Val | Likely pathogenic | -53.15 | Deleterious | 0.4637 | Ambiguous | -8.22 | Pathogenic | 0.67 | Disease | 1.00 | Probably damaging | 0.51 | Tolerated | 25.00 | Deleterious |
| p.Met53Tyr | VUS | -53.15 | Deleterious | 0.8966 | Pathogenic | -14.30 | Pathogenic | 0.66 | Disease | 1.00 | Probably damaging | 1 | Tolerated | | |
| p.Met53Cys | VUS | -11.19 | Indeterminate | 0.9022 | Pathogenic | -12.07 | Pathogenic | 0.72 | Disease | 1.00 | Probably damaging | 0.16 | Tolerated | | |
| p.Met53Trp | VUS | -53.15 | Deleterious | 0.9197 | Pathogenic | -12.97 | Pathogenic | 0.67 | Disease | 1.00 | Probably damaging | 0.16 | Tolerated | | |
| p.Met53Phe | VUS | -53.15 | Deleterious | 0.647 | Pathogenic | -12.53 | Pathogenic | 0.64 | Disease | 0.93 | Possibly damaging | 0.61 | Tolerated | | |
| p.Met54Asn | VUS | -1.00 | Neutral | 0.7165 | Pathogenic | -10.22 | Pathogenic | 0.60 | Disease | 1.00 | Probably damaging | 0.1 | Tolerated | | |
| p.Met54Lys | VUS | -23.49 | Indeterminate | 0.5076 | Ambiguous | -13.23 | Pathogenic | 0.71 | Disease | 0.93 | Possibly damaging | 0.16 | Tolerated | 28.70 | Deleterious |
| p.Met54Thr | VUS | 0.00 | Neutral | 0.6814 | Pathogenic | -8.51 | Pathogenic | 0.58 | Disease | 0.98 | Probably damaging | 0.14 | Tolerated | 27.00 | Deleterious |
| p.Met54Arg | VUS | -24.87 | Indeterminate | 0.3292 | Benign | -12.14 | Pathogenic | 0.72 | Disease | 0.97 | Probably damaging | 0.12 | Tolerated | 29.00 | Deleterious |
| p.Met54Ser | VUS | -0.03 | Neutral | 0.6661 | Pathogenic | -9.69 | Pathogenic | 0.61 | Disease | 1.00 | Probably damaging | 0.13 | Tolerated | | |
| p.Met54Ile | VUS | -0.31 | Neutral | 0.8666 | Pathogenic | -7.90 | Pathogenic | 0.35 | Neutral | 0.84 | Possibly damaging | 0.36 | Tolerated | 28.60 | Deleterious |
| p.Met54His | VUS | -3.79 | Neutral | 0.584 | Pathogenic | -10.15 | Pathogenic | 0.62 | Disease | 1.00 | Probably damaging | 0.14 | Tolerated | | |
| p.Met54Gln | VUS | -5.09 | Neutral | 0.2563 | Benign | -10.50 | Pathogenic | 0.58 | Disease | 1.00 | Probably damaging | 0.11 | Tolerated | | |
| p.Met54Pro | VUS | -14.17 | Indeterminate | 0.8247 | Pathogenic | -10.81 | Pathogenic | 0.62 | Disease | 1.00 | Probably damaging | 0.08 | Tolerated | | |
| p.Met54Leu | VUS | -0.08 | Neutral | 0.324 | Benign | -5.24 | Benign | 0.30 | Neutral | 0.33 | Benign | 1 | Tolerated | 24.60 | Deleterious |
| p.Met54Asp | VUS | -12.93 | Indeterminate | 0.8666 | Pathogenic | -13.16 | Pathogenic | 0.59 | Disease | 1.00 | Probably damaging | 0.13 | Tolerated | | |
| p.Met54Glu | VUS | -14.17 | Indeterminate | 0.6117 | Pathogenic | -13.35 | Pathogenic | 0.59 | Disease | 1.00 | Probably damaging | 0.11 | Tolerated | | |
| p.Met54Ala | VUS | -0.10 | Neutral | 0.5595 | Ambiguous | -8.78 | Pathogenic | 0.59 | Disease | 0.94 | Possibly damaging | 0.19 | Tolerated | | |
| p.Met54Gly | VUS | -0.03 | Neutral | 0.6583 | Pathogenic | -9.91 | Pathogenic | 0.60 | Disease | 1.00 | Probably damaging | 0.1 | Tolerated | | |
| p.Met54Val | VUS | -0.07 | Neutral | 0.4549 | Ambiguous | -6.74 | Benign | 0.58 | Disease | 1.00 | Probably damaging | 0.27 | Tolerated | 26.20 | Deleterious |
| p.Met54Tyr | VUS | -0.82 | Neutral | 0.6609 | Pathogenic | -9.24 | Pathogenic | 0.59 | Disease | 1.00 | Probably damaging | 0.25 | Tolerated | | |
| p.Met54Cys | VUS | 0.00 | Neutral | 0.8017 | Pathogenic | -8.84 | Pathogenic | 0.60 | Disease | 1.00 | Probably damaging | 0.06 | Tolerated | | |
| p.Met54Trp | VUS | -0.70 | Neutral | 0.6948 | Pathogenic | -13.35 | Pathogenic | 0.55 | Disease | 1.00 | Probably damaging | 0.06 | Tolerated | | |
| p.Met54Phe | VUS | -0.04 | Neutral | 0.4123 | Ambiguous | -6.99 | Benign | 0.39 | Neutral | 0.93 | Possibly damaging | 0.35 | Tolerated | | |
| p.Gly55Asn | VUS | -53.15 | Deleterious | 0.9308 | Pathogenic | -12.60 | Pathogenic | 0.76 | Disease | 1.00 | Probably damaging | 0.21 | Tolerated | | |
| p.Gly55Lys | VUS | -53.15 | Deleterious | 0.9741 | Pathogenic | -14.48 | Pathogenic | 0.77 | Disease | 1.00 | Probably damaging | 0.01 | Deleterious | | |
| p.Gly55Thr | VUS | -53.15 | Deleterious | 0.9206 | Pathogenic | -13.28 | Pathogenic | 0.77 | Disease | 1.00 | Probably damaging | 0.01 | Deleterious | | |
| p.Gly55Arg | VUS | -53.15 | Deleterious | 0.9318 | Pathogenic | -10.66 | Pathogenic | 0.78 | Disease | 1.00 | Probably damaging | 0.01 | Deleterious | 30.00 | Deleterious |
| p.Gly55Ser | VUS | -31.93 | Indeterminate | 0.7309 | Pathogenic | -8.58 | Pathogenic | 0.75 | Disease | 1.00 | Probably damaging | 0.03 | Deleterious | 29.80 | Deleterious |
| p.Gly55Ile | VUS | -53.15 | Deleterious | 0.9786 | Pathogenic | -18.91 | Pathogenic | 0.84 | Disease | 1.00 | Probably damaging | 0 | Deleterious | | |
| p.Gly55Met | VUS | -53.15 | Deleterious | 0.987 | Pathogenic | -18.38 | Pathogenic | 0.83 | Disease | 1.00 | Probably damaging | 0 | Deleterious | | |
| p.Gly55His | VUS | -53.15 | Deleterious | 0.971 | Pathogenic | -14.25 | Pathogenic | 0.79 | Disease | 1.00 | Probably damaging | 0.01 | Deleterious | | |
| p.Gly55Gln | VUS | -53.15 | Deleterious | 0.9509 | Pathogenic | -14.01 | Pathogenic | 0.77 | Disease | 1.00 | Probably damaging | 0.01 | Deleterious | | |
| p.Gly55Pro | VUS | -53.15 | Deleterious | 0.9783 | Pathogenic | -12.38 | Pathogenic | 0.78 | Disease | 0.66 | Possibly damaging | 0.01 | Deleterious | | |
| p.Gly55Leu | VUS | -53.15 | Deleterious | 0.9734 | Pathogenic | -17.29 | Pathogenic | 0.81 | Disease | 1.00 | Probably damaging | 0 | Deleterious | | |
| p.Gly55Asp | VUS | -53.15 | Deleterious | 0.9053 | Pathogenic | -11.43 | Pathogenic | 0.81 | Disease | 1.00 | Probably damaging | 0.04 | Deleterious | 29.20 | Deleterious |
| p.Gly55Glu | VUS | -53.15 | Deleterious | 0.9242 | Pathogenic | -12.71 | Pathogenic | 0.83 | Disease | 1.00 | Probably damaging | 0.01 | Deleterious | | |
| p.Gly55Ala | VUS | -11.05 | Indeterminate | 0.6234 | Pathogenic | -7.81 | Pathogenic | 0.79 | Disease | 0.99 | Probably damaging | 0.02 | Deleterious | 27.40 | Deleterious |
| p.Gly55Val | VUS | -53.15 | Deleterious | 0.9375 | Pathogenic | -15.51 | Pathogenic | 0.89 | Disease | 1.00 | Probably damaging | 0 | Deleterious | 28.90 | Deleterious |
| p.Gly55Tyr | VUS | -53.15 | Deleterious | 0.9725 | Pathogenic | -16.46 | Pathogenic | 0.82 | Disease | 1.00 | Probably damaging | 0 | Deleterious | | |
| p.Gly55Cys | VUS | -53.15 | Deleterious | 0.9277 | Pathogenic | -12.03 | Pathogenic | 0.87 | Disease | 1.00 | Probably damaging | 0 | Deleterious | 32.00 | Deleterious |
| p.Gly55Trp | VUS | -53.15 | Deleterious | 0.9668 | Pathogenic | -16.50 | Pathogenic | 0.86 | Disease | 1.00 | Probably damaging | 0 | Deleterious | | |
| p.Gly55Phe | VUS | -53.15 | Deleterious | 0.9875 | Pathogenic | -18.26 | Pathogenic | 0.84 | Disease | 1.00 | Probably damaging | 0 | Deleterious | | |
| p.Ser56Asn | VUS | 0.00 | Neutral | 0.2869 | Benign | -3.98 | Benign | 0.16 | Neutral | 0.03 | Benign | 0.9 | Tolerated | 12.86 | Neutral |
| p.Ser56Lys | VUS | -0.10 | Neutral | 0.925 | Pathogenic | -9.82 | Pathogenic | 0.50 | Neutral | 0.99 | Probably damaging | 0.95 | Tolerated | | |
| p.Ser56Thr | VUS | -0.47 | Neutral | 0.3245 | Benign | -7.24 | Benign | 0.39 | Neutral | 0.68 | Possibly damaging | 0.62 | Tolerated | 17.51 | Deleterious |
| p.Ser56Arg | VUS | 0.00 | Neutral | 0.863 | Pathogenic | -8.36 | Pathogenic | 0.56 | Disease | 1.00 | Probably damaging | 0.78 | Tolerated | 24.70 | Deleterious |
| p.Ser56Ile | Pathogenic | -53.15 | Deleterious | 0.8369 | Pathogenic | -14.63 | Pathogenic | 0.57 | Disease | 1.00 | Probably damaging | 0.26 | Tolerated | 22.80 | Deleterious |
| p.Ser56Met | VUS | 0.00 | Neutral | 0.7197 | Pathogenic | -13.25 | Pathogenic | 0.54 | Disease | 1.00 | Probably damaging | 0.16 | Tolerated | | |
| p.Ser56His | VUS | 0.00 | Neutral | 0.4565 | Ambiguous | -5.87 | Benign | 0.33 | Neutral | 1.00 | Probably damaging | 0.37 | Tolerated | | |
| p.Ser56Gln | VUS | 0.00 | Neutral | 0.7042 | Pathogenic | -8.58 | Pathogenic | 0.48 | Neutral | 1.00 | Probably damaging | 0.62 | Tolerated | | |
| p.Ser56Pro | VUS | -53.15 | Deleterious | 0.8374 | Pathogenic | -7.37 | Benign | 0.64 | Disease | 1.00 | Probably damaging | 0.29 | Tolerated | | |
| p.Ser56Leu | VUS | -17.53 | Indeterminate | 0.6522 | Pathogenic | -10.78 | Pathogenic | 0.55 | Disease | 0.99 | Probably damaging | 0.45 | Tolerated | | |
| p.Ser56Asp | VUS | -0.20 | Neutral | 0.6454 | Pathogenic | -8.35 | Pathogenic | 0.48 | Neutral | 0.92 | Possibly damaging | 0.62 | Tolerated | | |
| p.Ser56Glu | VUS | -3.79 | Neutral | 0.7859 | Pathogenic | -9.46 | Pathogenic | 0.50 | Neutral | 0.97 | Probably damaging | 0.67 | Tolerated | | |
| p.Ser56Ala | VUS | -0.09 | Neutral | 0.2536 | Benign | -8.17 | Pathogenic | 0.43 | Neutral | 0.80 | Possibly damaging | 0.64 | Tolerated | | |
| p.Ser56Gly | VUS | -2.34 | Neutral | 0.2343 | Benign | -7.26 | Benign | 0.44 | Neutral | 0.39 | Benign | 0.52 | Tolerated | 25.70 | Deleterious |
| p.Ser56Val | VUS | -2.86 | Neutral | 0.7218 | Pathogenic | -11.82 | Pathogenic | 0.53 | Disease | 1.00 | Probably damaging | 0.36 | Tolerated | | |
| p.Ser56Tyr | VUS | -0.01 | Neutral | 0.5949 | Pathogenic | -8.44 | Pathogenic | 0.53 | Disease | 1.00 | Probably damaging | 0.47 | Tolerated | | |
| p.Ser56Cys | VUS | 0.00 | Neutral | 0.2089 | Benign | -8.00 | Pathogenic | 0.30 | Neutral | 1.00 | Probably damaging | 0.12 | Tolerated | 27.30 | Deleterious |
| p.Ser56Trp | VUS | -53.15 | Deleterious | 0.7018 | Pathogenic | -10.80 | Pathogenic | 0.61 | Disease | 1.00 | Probably damaging | 0.1 | Tolerated | | |
| p.Ser56Phe | VUS | -0.08 | Neutral | 0.7895 | Pathogenic | -10.42 | Pathogenic | 0.51 | Disease | 1.00 | Probably damaging | 0.42 | Tolerated | | |
| p.Ala57Asn | VUS | -0.57 | Neutral | 0.4468 | Ambiguous | -8.31 | Pathogenic | 0.37 | Neutral | 0.56 | Possibly damaging | 0.46 | Tolerated | | |
| p.Ala57Lys | VUS | -2.18 | Neutral | 0.4994 | Ambiguous | -10.63 | Pathogenic | 0.39 | Neutral | 0.92 | Possibly damaging | 0.87 | Tolerated | | |
| p.Ala57Thr | VUS | -3.45 | Neutral | 0.1004 | Benign | -4.76 | Benign | 0.10 | Neutral | 0.01 | Benign | 0.56 | Tolerated | 5.39 | Neutral |
| p.Ala57Arg | VUS | -0.24 | Neutral | 0.319 | Benign | -7.61 | Pathogenic | 0.39 | Neutral | 0.98 | Probably damaging | 0.49 | Tolerated | | |
| p.Ala57Ser | VUS | -4.10 | Neutral | 0.1098 | Benign | -5.17 | Benign | 0.20 | Neutral | 0.08 | Benign | 0.72 | Tolerated | 8.12 | Neutral |
| p.Ala57Ile | VUS | -1.13 | Neutral | 0.3926 | Ambiguous | -11.03 | Pathogenic | 0.29 | Neutral | 0.85 | Possibly damaging | 0.18 | Tolerated | | |
| p.Ala57Met | VUS | -3.27 | Neutral | 0.3899 | Ambiguous | -10.89 | Pathogenic | 0.37 | Neutral | 0.99 | Probably damaging | 0.1 | Tolerated | | |
| p.Ala57His | VUS | -0.01 | Neutral | 0.4664 | Ambiguous | -8.74 | Pathogenic | 0.38 | Neutral | 1.00 | Probably damaging | 0.12 | Tolerated | | |
| p.Ala57Gln | VUS | -1.67 | Neutral | 0.3253 | Benign | -9.50 | Pathogenic | 0.37 | Neutral | 0.99 | Probably damaging | 0.56 | Tolerated | | |
| p.Ala57Pro | VUS | -4.88 | Neutral | 0.1227 | Benign | -4.00 | Benign | 0.18 | Neutral | 0.02 | Benign | 0.3 | Tolerated | 7.88 | Neutral |
| p.Ala57Leu | VUS | -9.30 | Indeterminate | 0.267 | Benign | -8.91 | Pathogenic | 0.37 | Neutral | 0.71 | Possibly damaging | 0.29 | Tolerated | | |
| p.Ala57Asp | VUS | -0.12 | Neutral | 0.4412 | Ambiguous | -10.51 | Pathogenic | 0.35 | Neutral | 0.70 | Possibly damaging | 0.62 | Tolerated | 18.96 | Deleterious |
| p.Ala57Glu | VUS | -9.67 | Indeterminate | 0.2968 | Benign | -10.32 | Pathogenic | 0.32 | Neutral | 0.81 | Possibly damaging | 1 | Tolerated | | |
| p.Ala57Gly | VUS | -1.14 | Neutral | 0.1392 | Benign | -6.10 | Benign | 0.32 | Neutral | 0.57 | Possibly damaging | 0.36 | Tolerated | 14.86 | Neutral |
| p.Ala57Val | Benign | -0.16 | Neutral | 0.1532 | Benign | -7.22 | Benign | 0.15 | Neutral | 0.01 | Benign | 0.3 | Tolerated | 9.79 | Neutral |
| p.Ala57Tyr | VUS | -15.97 | Indeterminate | 0.6368 | Pathogenic | -9.29 | Pathogenic | 0.37 | Neutral | 0.99 | Probably damaging | 0.04 | Deleterious | | |
| p.Ala57Cys | VUS | -4.97 | Neutral | 0.471 | Ambiguous | -8.78 | Pathogenic | 0.26 | Neutral | 1.00 | Probably damaging | 0.05 | Deleterious | | |
| p.Ala57Trp | VUS | -8.41 | Indeterminate | 0.6804 | Pathogenic | -11.74 | Pathogenic | 0.42 | Neutral | 1.00 | Probably damaging | 0.02 | Deleterious | | |
| p.Ala57Phe | VUS | -15.14 | Indeterminate | 0.5515 | Ambiguous | -11.25 | Pathogenic | 0.38 | Neutral | 0.98 | Probably damaging | 0.05 | Deleterious | | |
| p.Arg58Asn | VUS | 0.00 | Neutral | 0.5027 | Ambiguous | -7.88 | Pathogenic | 0.46 | Neutral | 0.96 | Probably damaging | 0.35 | Tolerated | | |
| p.Arg58Lys | VUS | 0.00 | Neutral | 0.1612 | Benign | -5.06 | Benign | 0.12 | Neutral | 0.36 | Benign | 0.69 | Tolerated | | |
| p.Arg58Thr | VUS | -0.10 | Neutral | 0.1907 | Benign | -5.72 | Benign | 0.41 | Neutral | 0.96 | Probably damaging | 0.44 | Tolerated | | |
| p.Arg58Ser | VUS | 0.00 | Neutral | 0.311 | Benign | -4.61 | Benign | 0.17 | Neutral | 0.99 | Probably damaging | 0.51 | Tolerated | | |
| p.Arg58Ile | VUS | 0.00 | Neutral | 0.3846 | Ambiguous | -9.99 | Pathogenic | 0.50 | Neutral | 1.00 | Probably damaging | 0.13 | Tolerated | | |
| p.Arg58Met | VUS | 0.00 | Neutral | 0.4241 | Ambiguous | -8.79 | Pathogenic | 0.49 | Neutral | 0.99 | Probably damaging | 0.07 | Tolerated | | |
| p.Arg58His | VUS | 0.00 | Neutral | 0.1485 | Benign | -6.11 | Benign | 0.19 | Neutral | 0.01 | Benign | 0.13 | Tolerated | | |
| p.Arg58Gln | VUS | -0.05 | Neutral | 0.1117 | Benign | -4.30 | Benign | 0.17 | Neutral | 0.29 | Benign | 0.47 | Tolerated | 12.76 | Neutral |
| p.Arg58Pro | VUS | -0.22 | Neutral | 0.2054 | Benign | -3.63 | Benign | 0.18 | Neutral | 0.37 | Benign | 0.19 | Tolerated | 12.91 | Neutral |
| p.Arg58Leu | VUS | 0.00 | Neutral | 0.1855 | Benign | -6.61 | Benign | 0.47 | Neutral | 1.00 | Probably damaging | 0.21 | Tolerated | 14.45 | Neutral |
| p.Arg58Asp | VUS | 0.00 | Neutral | 0.3324 | Benign | -4.88 | Benign | 0.48 | Neutral | 0.99 | Probably damaging | 0.54 | Tolerated | | |
| p.Arg58Glu | VUS | 0.00 | Neutral | 0.203 | Benign | -3.13 | Benign | 0.25 | Neutral | 0.59 | Possibly damaging | 1 | Tolerated | | |
| p.Arg58Ala | VUS | -0.03 | Neutral | 0.1563 | Benign | -2.13 | Benign | 0.27 | Neutral | 0.90 | Possibly damaging | 0.72 | Tolerated | | |
| p.Arg58Gly | VUS | -0.04 | Neutral | 0.1835 | Benign | -4.40 | Benign | 0.20 | Neutral | 1.00 | Probably damaging | 0.27 | Tolerated | 17.53 | Deleterious |
| p.Arg58Val | VUS | 0.00 | Neutral | 0.303 | Benign | -6.25 | Benign | 0.52 | Disease | 0.99 | Probably damaging | 0.2 | Tolerated | | |
| p.Arg58Tyr | VUS | 0.00 | Neutral | 0.498 | Ambiguous | -9.91 | Pathogenic | 0.45 | Neutral | 0.69 | Possibly damaging | 0.03 | Deleterious | | |
| p.Arg58Cys | VUS | 0.00 | Neutral | 0.3604 | Ambiguous | -7.46 | Benign | 0.48 | Neutral | 1.00 | Probably damaging | 0.04 | Deleterious | | |
| p.Arg58Trp | VUS | 0.00 | Neutral | 0.282 | Benign | -9.55 | Pathogenic | 0.48 | Neutral | 1.00 | Probably damaging | 0.01 | Deleterious | | |
| p.Arg58Phe | VUS | -0.01 | Neutral | 0.5892 | Pathogenic | -9.88 | Pathogenic | 0.52 | Disease | 0.82 | Possibly damaging | 0.04 | Deleterious | | |
| p.Val59Asn | VUS | -53.15 | Deleterious | 0.9093 | Pathogenic | -15.23 | Pathogenic | 0.69 | Disease | 1.00 | Probably damaging | 0 | Deleterious | | |
| p.Val59Lys | VUS | -53.15 | Deleterious | 0.9586 | Pathogenic | -16.15 | Pathogenic | 0.75 | Disease | 0.86 | Possibly damaging | 0 | Deleterious | | |
| p.Val59Thr | VUS | -0.21 | Neutral | 0.4091 | Ambiguous | -8.59 | Pathogenic | 0.47 | Neutral | 1.00 | Probably damaging | 0.01 | Deleterious | | |
| p.Val59Arg | VUS | -53.15 | Deleterious | 0.9158 | Pathogenic | -14.30 | Pathogenic | 0.75 | Disease | 0.26 | Benign | 0 | Deleterious | | |

| | | | | | | | | | | | | | | | |
|---|---|---|---|---|---|---|---|---|---|---|---|---|---|---|---|
| p.Val59Ser | VUS | -4.70 | Neutral | 0.7444 | Pathogenic | -10.49 | Pathogenic | 0.68 | Disease | 1.00 | Probably damaging | 0 | Deleterious | | |
| p.Val59Ile | VUS | -0.16 | Neutral | 0.1519 | Benign | -7.73 | Pathogenic | 0.25 | Neutral | 0.32 | Benign | 0.93 | Tolerated | | |
| p.Val59Met | VUS | -1.55 | Neutral | 0.6448 | Pathogenic | -9.24 | Pathogenic | 0.42 | Neutral | 0.56 | Possibly damaging | 0.01 | Deleterious | 21.90 | Deleterious |
| p.Val59His | VUS | -53.15 | Deleterious | 0.979 | Pathogenic | -15.47 | Pathogenic | 0.68 | Disease | 1.00 | Probably damaging | 0 | Deleterious | | |
| p.Val59Gln | VUS | -53.15 | Deleterious | 0.905 | Pathogenic | -14.47 | Pathogenic | 0.72 | Disease | 0.97 | Probably damaging | 0 | Deleterious | | |
| p.Val59Pro | VUS | -53.15 | Deleterious | 0.8617 | Pathogenic | -12.37 | Pathogenic | 0.71 | Disease | 1.00 | Probably damaging | 0 | Deleterious | | |
| p.Val59Leu | VUS | -4.37 | Neutral | 0.6958 | Pathogenic | -9.84 | Pathogenic | 0.34 | Neutral | 0.26 | Benign | 0.14 | Tolerated | 16.58 | Deleterious |
| p.Val59Asp | VUS | -53.15 | Deleterious | 0.959 | Pathogenic | -14.03 | Pathogenic | 0.79 | Disease | 1.00 | Probably damaging | 0 | Deleterious | | |
| p.Val59Glu | VUS | -53.15 | Deleterious | 0.8776 | Pathogenic | -12.84 | Pathogenic | 0.82 | Disease | 1.00 | Probably damaging | 0 | Deleterious | 24.40 | Deleterious |
| p.Val59Ala | VUS | -5.43 | Neutral | 0.4051 | Ambiguous | -6.68 | Benign | 0.59 | Disease | 1.00 | Probably damaging | 0.01 | Deleterious | 23.80 | Deleterious |
| p.Val59Gly | Pathogenic | -33.22 | Indeterminate | 0.6413 | Pathogenic | -11.20 | Pathogenic | 0.60 | Disease | 1.00 | Probably damaging | 0 | Deleterious | 24.30 | Deleterious |
| p.Val59Tyr | VUS | -53.15 | Deleterious | 0.9861 | Pathogenic | -16.57 | Pathogenic | 0.69 | Disease | 1.00 | Probably damaging | 0.01 | Deleterious | | |
| p.Val59Cys | VUS | -0.94 | Neutral | 0.9254 | Pathogenic | -8.71 | Pathogenic | 0.64 | Disease | 1.00 | Probably damaging | 0 | Deleterious | | |
| p.Val59Trp | VUS | -53.15 | Deleterious | 0.9946 | Pathogenic | -17.04 | Pathogenic | 0.65 | Disease | 1.00 | Probably damaging | 0 | Deleterious | | |
| p.Val59Phe | VUS | -53.15 | Deleterious | 0.8967 | Pathogenic | -15.45 | Pathogenic | 0.74 | Disease | 0.99 | Probably damaging | 0.01 | Deleterious | | |
| p.Ala60Asn | VUS | -53.15 | Deleterious | 0.9602 | Pathogenic | -17.64 | Pathogenic | 0.72 | Disease | 0.98 | Probably damaging | 0.01 | Deleterious | | |
| p.Ala60Lys | VUS | -53.15 | Deleterious | 0.9921 | Pathogenic | -19.21 | Pathogenic | 0.76 | Disease | 0.99 | Probably damaging | 0.01 | Deleterious | | |
| p.Ala60Thr | VUS | -1.18 | Neutral | 0.3475 | Ambiguous | -7.22 | Benign | 0.62 | Disease | 0.99 | Probably damaging | 0.04 | Deleterious | 25.80 | Deleterious |
| p.Ala60Arg | VUS | -53.15 | Deleterious | 0.9613 | Pathogenic | -15.85 | Pathogenic | 0.73 | Disease | 1.00 | Probably damaging | 0.01 | Deleterious | | |
| p.Ala60Ser | VUS | -14.38 | Indeterminate | 0.3268 | Benign | -8.42 | Pathogenic | 0.62 | Disease | 0.79 | Possibly damaging | 0.03 | Deleterious | 25.10 | Deleterious |
| p.Ala60Ile | VUS | -22.52 | Indeterminate | 0.8372 | Pathogenic | -11.44 | Pathogenic | 0.74 | Disease | 1.00 | Probably damaging | 0.04 | Deleterious | | |
| p.Ala60Met | VUS | -2.94 | Neutral | 0.7943 | Pathogenic | -10.80 | Pathogenic | 0.68 | Disease | 1.00 | Probably damaging | 0.01 | Deleterious | | |
| p.Ala60His | VUS | -33.18 | Indeterminate | 0.9832 | Pathogenic | -18.02 | Pathogenic | 0.74 | Disease | 1.00 | Probably damaging | 0 | Deleterious | | |
| p.Ala60Gln | VUS | -53.15 | Deleterious | 0.9314 | Pathogenic | -16.75 | Pathogenic | 0.74 | Disease | 0.96 | Probably damaging | 0.01 | Deleterious | | |
| p.Ala60Pro | VUS | -18.86 | Indeterminate | 0.9469 | Pathogenic | -10.86 | Pathogenic | 0.80 | Disease | 1.00 | Probably damaging | 0.02 | Deleterious | 25.70 | Deleterious |
| p.Ala60Leu | VUS | -2.51 | Neutral | 0.6467 | Pathogenic | -8.43 | Pathogenic | 0.69 | Disease | 1.00 | Probably damaging | 0.03 | Deleterious | | |
| p.Ala60Asp | VUS | -33.22 | Indeterminate | 0.9837 | Pathogenic | -16.66 | Pathogenic | 0.84 | Disease | 1.00 | Probably damaging | 0.01 | Deleterious | | |
| p.Ala60Glu | Likely pathogenic | -33.22 | Indeterminate | 0.962 | Pathogenic | -15.15 | Pathogenic | 0.79 | Disease | 0.99 | Probably damaging | 0.01 | Deleterious | 25.40 | Deleterious |
| p.Ala60Gly | VUS | -1.98 | Neutral | 0.3107 | Benign | -9.00 | Pathogenic | 0.62 | Disease | 1.00 | Probably damaging | 0.03 | Deleterious | 26.20 | Deleterious |
| p.Ala60Val | VUS | -5.18 | Neutral | 0.4418 | Ambiguous | -7.31 | Benign | 0.49 | Neutral | 1.00 | Probably damaging | 0.37 | Tolerated | 25.80 | Deleterious |
| p.Ala60Tyr | VUS | -53.15 | Deleterious | 0.9901 | Pathogenic | -17.64 | Pathogenic | 0.75 | Disease | 1.00 | Probably damaging | 0 | Deleterious | | |
| p.Ala60Cys | VUS | -7.26 | Indeterminate | 0.6622 | Pathogenic | -8.74 | Pathogenic | 0.67 | Disease | 1.00 | Probably damaging | 0.01 | Deleterious | | |
| p.Ala60Trp | VUS | -53.15 | Deleterious | 0.9927 | Pathogenic | -15.47 | Pathogenic | 0.79 | Disease | 1.00 | Probably damaging | 0 | Deleterious | | |
| p.Ala60Phe | VUS | -53.15 | Deleterious | 0.9672 | Pathogenic | -13.94 | Pathogenic | 0.74 | Disease | 1.00 | Probably damaging | 0.01 | Deleterious | | |
| p.Glu61Asn | VUS | -1.02 | Neutral | 0.7007 | Pathogenic | -10.25 | Pathogenic | 0.41 | Neutral | 0.73 | Possibly damaging | 0.48 | Tolerated | | |
| p.Glu61Lys | VUS | -0.20 | Neutral | 0.3723 | Ambiguous | -9.29 | Pathogenic | 0.23 | Neutral | 0.12 | Benign | 0.89 | Tolerated | 11.10 | Neutral |
| p.Glu61Thr | VUS | -0.31 | Neutral | 0.5214 | Ambiguous | -8.09 | Pathogenic | 0.32 | Neutral | 0.79 | Possibly damaging | 0.52 | Tolerated | | |
| p.Glu61Arg | VUS | -4.80 | Neutral | 0.2715 | Benign | -6.99 | Benign | 0.23 | Neutral | 0.00 | Benign | 0.52 | Tolerated | | |
| p.Glu61Ser | VUS | -1.62 | Neutral | 0.3724 | Ambiguous | -6.91 | Benign | 0.29 | Neutral | 0.59 | Possibly damaging | 0.66 | Tolerated | | |
| p.Glu61Ile | VUS | -2.46 | Neutral | 0.8797 | Pathogenic | -12.01 | Pathogenic | 0.50 | Disease | 0.75 | Possibly damaging | 0.17 | Tolerated | | |
| p.Glu61Met | VUS | -2.35 | Neutral | 0.8462 | Pathogenic | -10.24 | Pathogenic | 0.54 | Disease | 0.62 | Possibly damaging | 0.1 | Tolerated | | |
| p.Glu61His | VUS | -1.51 | Neutral | 0.5801 | Pathogenic | -8.60 | Pathogenic | 0.39 | Neutral | 0.82 | Possibly damaging | 0.11 | Tolerated | | |
| p.Glu61Gln | VUS | -0.25 | Neutral | 0.1632 | Benign | -6.32 | Benign | 0.13 | Neutral | 0.02 | Benign | 0.53 | Tolerated | 7.74 | Neutral |
| p.Glu61Pro | VUS | -53.15 | Deleterious | 0.9409 | Pathogenic | -13.42 | Pathogenic | 0.52 | Disease | 0.86 | Possibly damaging | 0.25 | Tolerated | | |
| p.Glu61Leu | VUS | -0.27 | Neutral | 0.7588 | Pathogenic | -8.93 | Pathogenic | 0.38 | Neutral | 0.29 | Benign | 0.31 | Tolerated | | |
| p.Glu61Asp | VUS | -0.83 | Neutral | 0.4425 | Ambiguous | -7.21 | Benign | 0.33 | Neutral | 0.22 | Benign | 0.6 | Tolerated | 12.29 | Neutral |
| p.Glu61Ala | VUS | -0.62 | Neutral | 0.2695 | Benign | -6.58 | Benign | 0.19 | Neutral | 0.03 | Benign | 0.72 | Tolerated | 13.52 | Neutral |
| p.Glu61Gly | VUS | -0.43 | Neutral | 0.2971 | Benign | -8.01 | Pathogenic | 0.38 | Neutral | 0.02 | Benign | 0.34 | Tolerated | 18.67 | Deleterious |
| p.Glu61Val | VUS | -0.35 | Neutral | 0.6779 | Pathogenic | -9.71 | Pathogenic | 0.31 | Neutral | 0.18 | Benign | 0.26 | Tolerated | 17.51 | Deleterious |
| p.Glu61Tyr | VUS | -4.64 | Neutral | 0.871 | Pathogenic | -12.66 | Pathogenic | 0.50 | Neutral | 0.98 | Probably damaging | 0.04 | Deleterious | | |
| p.Glu61Cys | VUS | -0.54 | Neutral | 0.9517 | Pathogenic | -10.31 | Pathogenic | 0.53 | Disease | 1.00 | Probably damaging | 0.05 | Deleterious | | |
| p.Glu61Trp | VUS | -1.42 | Neutral | 0.9523 | Pathogenic | -11.63 | Pathogenic | 0.55 | Disease | 1.00 | Probably damaging | 0.02 | Deleterious | | |
| p.Glu61Phe | VUS | -2.00 | Neutral | 0.9411 | Pathogenic | -12.68 | Pathogenic | 0.51 | Disease | 0.95 | Possibly damaging | 0.05 | Deleterious | | |
| p.Leu62Asn | VUS | 0.00 | Neutral | 0.8298 | Pathogenic | -13.59 | Pathogenic | 0.68 | Disease | 1.00 | Probably damaging | 0.09 | Tolerated | | |
| p.Leu62Lys | VUS | -1.24 | Neutral | 0.7361 | Pathogenic | -12.18 | Pathogenic | 0.69 | Disease | 0.44 | Benign | 0.36 | Tolerated | | |
| p.Leu62Thr | VUS | -0.19 | Neutral | 0.5488 | Ambiguous | -7.20 | Benign | 0.64 | Disease | 1.00 | Probably damaging | 0.14 | Tolerated | | |
| p.Leu62Arg | VUS | -28.58 | Indeterminate | 0.5057 | Ambiguous | -10.44 | Pathogenic | 0.69 | Disease | 0.98 | Probably damaging | 0.31 | Tolerated | 24.20 | Deleterious |
| p.Leu62Ser | VUS | -0.05 | Neutral | 0.8354 | Pathogenic | -9.57 | Pathogenic | 0.70 | Disease | 1.00 | Probably damaging | 0.11 | Tolerated | | |
| p.Leu62Ile | VUS | -7.52 | Indeterminate | 0.1772 | Benign | -7.87 | Pathogenic | 0.28 | Neutral | 0.99 | Probably damaging | 0.37 | Tolerated | | |
| p.Leu62Met | VUS | 0.00 | Neutral | 0.2382 | Benign | -7.24 | Benign | 0.43 | Neutral | 1.00 | Probably damaging | 0.21 | Tolerated | 23.00 | Deleterious |
| p.Leu62His | VUS | 0.00 | Neutral | 0.6562 | Pathogenic | -8.78 | Pathogenic | 0.69 | Disease | 1.00 | Probably damaging | 0.08 | Tolerated | | |
| p.Leu62Gln | VUS | -29.45 | Indeterminate | 0.4796 | Ambiguous | -7.67 | Pathogenic | 0.69 | Disease | 0.99 | Probably damaging | 0.17 | Tolerated | 25.70 | Deleterious |
| p.Leu62Pro | VUS | -13.28 | Indeterminate | 0.9602 | Pathogenic | -12.35 | Pathogenic | 0.83 | Disease | 1.00 | Probably damaging | 0.09 | Tolerated | 26.00 | Deleterious |
| p.Leu62Asp | VUS | -0.04 | Neutral | 0.8998 | Pathogenic | -9.24 | Pathogenic | 0.68 | Disease | 1.00 | Probably damaging | 0.07 | Tolerated | | |
| p.Leu62Glu | VUS | -0.08 | Neutral | 0.5869 | Pathogenic | -4.51 | Benign | 0.68 | Disease | 1.00 | Probably damaging | 0.21 | Tolerated | | |
| p.Leu62Ala | VUS | -0.01 | Neutral | 0.5589 | Ambiguous | -5.18 | Benign | 0.46 | Neutral | 1.00 | Probably damaging | 0.19 | Tolerated | | |
| p.Leu62Gly | VUS | -4.09 | Neutral | 0.8549 | Pathogenic | -11.88 | Pathogenic | 0.63 | Disease | 1.00 | Probably damaging | 0.07 | Tolerated | | |
| p.Leu62Val | VUS | -0.18 | Neutral | 0.1734 | Benign | -4.38 | Benign | 0.26 | Neutral | 0.95 | Possibly damaging | 0.29 | Tolerated | 19.78 | Deleterious |
| p.Leu62Tyr | VUS | -8.39 | Indeterminate | 0.5464 | Ambiguous | -8.73 | Pathogenic | 0.61 | Disease | 0.98 | Probably damaging | 0.08 | Tolerated | | |
| p.Leu62Cys | VUS | -5.15 | Neutral | 0.7386 | Pathogenic | -8.67 | Pathogenic | 0.59 | Disease | 1.00 | Probably damaging | 0.03 | Deleterious | | |
| p.Leu62Trp | VUS | 0.00 | Neutral | 0.543 | Ambiguous | -10.97 | Pathogenic | 0.63 | Disease | 1.00 | Probably damaging | 0.03 | Deleterious | | |
| p.Leu62Phe | VUS | 0.00 | Neutral | 0.4535 | Ambiguous | -8.48 | Pathogenic | 0.47 | Neutral | 1.00 | Probably damaging | 0.19 | Tolerated | | |
| p.Leu63Asn | VUS | -53.15 | Deleterious | 0.9953 | Pathogenic | -17.75 | Pathogenic | 0.84 | Disease | 1.00 | Probably damaging | 0 | Deleterious | | |
| p.Leu63Lys | VUS | -53.15 | Deleterious | 0.9902 | Pathogenic | -18.24 | Pathogenic | 0.82 | Disease | 0.99 | Probably damaging | 0 | Deleterious | | |
| p.Leu63Thr | VUS | -53.15 | Deleterious | 0.9863 | Pathogenic | -13.92 | Pathogenic | 0.84 | Disease | 1.00 | Probably damaging | 0 | Deleterious | | |
| p.Leu63Arg | VUS | -53.15 | Deleterious | 0.9585 | Pathogenic | -13.96 | Pathogenic | 0.93 | Disease | 1.00 | Probably damaging | 0 | Deleterious | 27.60 | Deleterious |
| p.Leu63Ser | VUS | -53.15 | Deleterious | 0.9945 | Pathogenic | -12.87 | Pathogenic | 0.92 | Disease | 1.00 | Probably damaging | 0 | Deleterious | | |
| p.Leu63Ile | VUS | -5.21 | Neutral | 0.7278 | Pathogenic | -10.80 | Pathogenic | 0.70 | Disease | 1.00 | Probably damaging | 0 | Deleterious | | |
| p.Leu63Met | VUS | -0.30 | Neutral | 0.7228 | Pathogenic | -9.02 | Pathogenic | 0.76 | Disease | 1.00 | Probably damaging | 0 | Deleterious | 25.20 | Deleterious |
| p.Leu63His | VUS | -53.15 | Deleterious | 0.9796 | Pathogenic | -14.33 | Pathogenic | 0.91 | Disease | 1.00 | Probably damaging | 0 | Deleterious | | |
| p.Leu63Gln | VUS | -53.15 | Deleterious | 0.9753 | Pathogenic | -14.73 | Pathogenic | 0.93 | Disease | 1.00 | Probably damaging | 0 | Deleterious | 27.10 | Deleterious |
| p.Leu63Pro | VUS | -53.15 | Deleterious | 0.9779 | Pathogenic | -11.69 | Pathogenic | 0.93 | Disease | 1.00 | Probably damaging | 0 | Deleterious | 27.80 | Deleterious |
| p.Leu63Asp | VUS | -53.15 | Deleterious | 0.9988 | Pathogenic | -17.88 | Pathogenic | 0.84 | Disease | 1.00 | Probably damaging | 0 | Deleterious | | |
| p.Leu63Glu | VUS | -53.15 | Deleterious | 0.9929 | Pathogenic | -17.56 | Pathogenic | 0.85 | Disease | 1.00 | Probably damaging | 0 | Deleterious | | |
| p.Leu63Ala | VUS | -53.15 | Deleterious | 0.9699 | Pathogenic | -12.36 | Pathogenic | 0.83 | Disease | 1.00 | Probably damaging | 0 | Deleterious | | |
| p.Leu63Gly | VUS | -53.15 | Deleterious | 0.9899 | Pathogenic | -16.36 | Pathogenic | 0.86 | Disease | 1.00 | Probably damaging | 0 | Deleterious | | |
| p.Leu63Val | VUS | -53.15 | Deleterious | 0.7804 | Pathogenic | -10.79 | Pathogenic | 0.83 | Disease | 0.99 | Probably damaging | 0 | Deleterious | 25.00 | Deleterious |
| p.Leu63Tyr | VUS | -53.15 | Deleterious | 0.9691 | Pathogenic | -15.35 | Pathogenic | 0.82 | Disease | 0.99 | Probably damaging | 0 | Deleterious | | |
| p.Leu63Cys | VUS | -53.15 | Deleterious | 0.9661 | Pathogenic | -12.61 | Pathogenic | 0.79 | Disease | 1.00 | Probably damaging | 0 | Deleterious | | |
| p.Leu63Trp | VUS | -53.15 | Deleterious | 0.9618 | Pathogenic | -14.79 | Pathogenic | 0.87 | Disease | 1.00 | Probably damaging | 0 | Deleterious | | |
| p.Leu63Phe | VUS | -33.17 | Indeterminate | 0.768 | Pathogenic | -9.89 | Pathogenic | 0.81 | Disease | 1.00 | Probably damaging | 0 | Deleterious | | |
| p.Leu64Asn | VUS | -11.07 | Indeterminate | 0.9826 | Pathogenic | -17.64 | Pathogenic | 0.80 | Disease | 1.00 | Probably damaging | 0 | Deleterious | | |
| p.Leu64Lys | VUS | -2.27 | Neutral | 0.9752 | Pathogenic | -17.22 | Pathogenic | 0.80 | Disease | 0.99 | Probably damaging | 0 | Deleterious | | |
| p.Leu64Thr | VUS | -0.60 | Neutral | 0.8919 | Pathogenic | -12.34 | Pathogenic | 0.83 | Disease | 1.00 | Probably damaging | 0.01 | Deleterious | | |
| p.Leu64Arg | VUS | -2.37 | Neutral | 0.89 | Pathogenic | -13.68 | Pathogenic | 0.88 | Disease | 1.00 | Probably damaging | 0 | Deleterious | 27.70 | Deleterious |
| p.Leu64Ser | VUS | -0.65 | Neutral | 0.9742 | Pathogenic | -12.85 | Pathogenic | 0.90 | Disease | 1.00 | Probably damaging | 0 | Deleterious | | |
| p.Leu64Ile | VUS | -10.16 | Indeterminate | 0.2904 | Benign | -7.11 | Benign | 0.55 | Disease | 1.00 | Probably damaging | 0.04 | Deleterious | | |
| p.Leu64Met | VUS | -1.56 | Neutral | 0.5884 | Pathogenic | -8.33 | Pathogenic | 0.68 | Disease | 1.00 | Probably damaging | 0.02 | Deleterious | 25.30 | Deleterious |
| p.Leu64His | VUS | -2.25 | Neutral | 0.9626 | Pathogenic | -15.14 | Pathogenic | 0.86 | Disease | 1.00 | Probably damaging | 0 | Deleterious | | |
| p.Leu64Gln | VUS | -5.58 | Neutral | 0.944 | Pathogenic | -13.81 | Pathogenic | 0.87 | Disease | 1.00 | Probably damaging | 0 | Deleterious | 27.20 | Deleterious |
| p.Leu64Pro | VUS | -53.15 | Deleterious | 0.9709 | Pathogenic | -13.51 | Pathogenic | 0.87 | Disease | 1.00 | Probably damaging | 0 | Deleterious | 27.80 | Deleterious |
| p.Leu64Asp | VUS | -17.65 | Indeterminate | 0.997 | Pathogenic | -17.53 | Pathogenic | 0.81 | Disease | 1.00 | Probably damaging | 0 | Deleterious | | |
| p.Leu64Glu | VUS | -7.32 | Indeterminate | 0.9826 | Pathogenic | -16.64 | Pathogenic | 0.83 | Disease | 1.00 | Probably damaging | 0 | Deleterious | | |
| p.Leu64Ala | VUS | -1.79 | Neutral | 0.8713 | Pathogenic | -11.12 | Pathogenic | 0.81 | Disease | 1.00 | Probably damaging | 0.01 | Deleterious | | |
| p.Leu64Gly | VUS | -4.23 | Neutral | 0.9676 | Pathogenic | -13.74 | Pathogenic | 0.82 | Disease | 1.00 | Probably damaging | 0 | Deleterious | | |
| p.Leu64Val | VUS | -0.62 | Neutral | 0.3207 | Benign | -8.37 | Pathogenic | 0.69 | Disease | 1.00 | Probably damaging | 0.15 | Tolerated | 25.00 | Deleterious |
| p.Leu64Tyr | VUS | -1.41 | Neutral | 0.9719 | Pathogenic | -15.94 | Pathogenic | 0.79 | Disease | 1.00 | Probably damaging | 0 | Deleterious | | |
| p.Leu64Cys | VUS | -0.83 | Neutral | 0.9069 | Pathogenic | -10.92 | Pathogenic | 0.76 | Disease | 1.00 | Probably damaging | 0 | Deleterious | | |
| p.Leu64Trp | VUS | -1.00 | Neutral | 0.9526 | Pathogenic | -14.38 | Pathogenic | 0.81 | Disease | 1.00 | Probably damaging | 0 | Deleterious | | |
| p.Leu64Phe | VUS | -0.19 | Neutral | 0.8411 | Pathogenic | -11.58 | Pathogenic | 0.74 | Disease | 1.00 | Probably damaging | 0.02 | Deleterious | | |
| p.Leu65Asn | VUS | -12.04 | Indeterminate | 0.3576 | Ambiguous | -7.55 | Pathogenic | 0.33 | Neutral | 1.00 | Probably damaging | 0.54 | Tolerated | | |

| | | | | | | | | | | | | | | | |
|---|---|---|---|---|---|---|---|---|---|---|---|---|---|---|---|
| p.Leu65Lys | VUS | -0.29 | Neutral | 0.1691 | Benign | -5.40 | Benign | 0.32 | Neutral | 0.97 | Probably damaging | 1 | Tolerated | | |
| p.Leu65Thr | VUS | -3.35 | Neutral | 0.1612 | Benign | -5.99 | Benign | 0.28 | Neutral | 1.00 | Probably damaging | 0.75 | Tolerated | | |
| p.Leu65Arg | VUS | -0.66 | Neutral | 0.0718 | Benign | -2.33 | Benign | 0.24 | Neutral | 1.00 | Probably damaging | 0.6 | Tolerated | 17.13 | Deleterious |
| p.Leu65Ser | VUS | -2.83 | Neutral | 0.1969 | Benign | -4.68 | Benign | 0.21 | Neutral | 1.00 | Probably damaging | 0.86 | Tolerated | | |
| p.Leu65Ile | VUS | -1.69 | Neutral | 0.1806 | Benign | -8.58 | Pathogenic | 0.21 | Neutral | 0.17 | Benign | 0.21 | Tolerated | 17.98 | Deleterious |
| p.Leu65Met | VUS | -0.21 | Neutral | 0.1757 | Benign | -7.29 | Benign | 0.36 | Neutral | 1.00 | Probably damaging | 0.12 | Tolerated | | |
| p.Leu65His | VUS | -10.93 | Indeterminate | 0.1732 | Benign | -4.99 | Benign | 0.37 | Neutral | 1.00 | Probably damaging | 0.16 | Tolerated | 22.50 | Deleterious |
| p.Leu65Gln | VUS | -3.84 | Neutral | 0.0746 | Benign | -2.31 | Benign | 0.19 | Neutral | 1.00 | Probably damaging | 0.61 | Tolerated | | |
| p.Leu65Pro | Likely pathogenic | -53.15 | Deleterious | 0.7377 | Pathogenic | -9.74 | Pathogenic | 0.31 | Neutral | 1.00 | Probably damaging | 0.3 | Tolerated | 22.60 | Deleterious |
| p.Leu65Asp | VUS | -0.82 | Neutral | 0.4034 | Ambiguous | -5.28 | Benign | 0.33 | Neutral | 1.00 | Probably damaging | 0.62 | Tolerated | | |
| p.Leu65Glu | VUS | -0.42 | Neutral | 0.1215 | Benign | -3.10 | Benign | 0.26 | Neutral | 1.00 | Probably damaging | 0.98 | Tolerated | | |
| p.Leu65Ala | VUS | -4.46 | Neutral | 0.1044 | Benign | -2.87 | Benign | 0.27 | Neutral | 1.00 | Probably damaging | 0.78 | Tolerated | | |
| p.Leu65Gly | VUS | -8.22 | Indeterminate | 0.2405 | Benign | -5.06 | Benign | 0.43 | Neutral | 1.00 | Probably damaging | 0.4 | Tolerated | | |
| p.Leu65Val | VUS | -2.08 | Neutral | 0.1378 | Benign | -6.10 | Benign | 0.20 | Neutral | 0.62 | Possibly damaging | 0.31 | Tolerated | 16.61 | Deleterious |
| p.Leu65Tyr | VUS | -1.23 | Neutral | 0.3546 | Ambiguous | -8.87 | Pathogenic | 0.38 | Neutral | 0.98 | Probably damaging | 0.09 | Tolerated | | |
| p.Leu65Cys | VUS | -13.13 | Indeterminate | 0.4305 | Ambiguous | -6.24 | Benign | 0.30 | Neutral | 1.00 | Probably damaging | 0.07 | Tolerated | | |
| p.Leu65Trp | VUS | -7.73 | Indeterminate | 0.2329 | Benign | -7.17 | Benign | 0.43 | Neutral | 1.00 | Probably damaging | 0.03 | Deleterious | | |
| p.Leu65Phe | VUS | -1.85 | Neutral | 0.1992 | Benign | -9.10 | Pathogenic | 0.28 | Neutral | 0.99 | Probably damaging | 0.08 | Tolerated | 22.60 | Deleterious |
| p.His66Asn | VUS | -0.16 | Neutral | 0.3884 | Ambiguous | -7.58 | Pathogenic | 0.26 | Neutral | 1.00 | Probably damaging | 0.04 | Deleterious | 22.90 | Deleterious |
| p.His66Lys | VUS | -0.18 | Neutral | 0.5891 | Pathogenic | -6.91 | Benign | 0.30 | Neutral | 0.60 | Possibly damaging | 0.16 | Tolerated | | |
| p.His66Thr | VUS | -2.36 | Neutral | 0.7957 | Pathogenic | -8.87 | Pathogenic | 0.49 | Neutral | 0.99 | Probably damaging | 0.05 | Deleterious | | |
| p.His66Arg | VUS | -1.93 | Neutral | 0.1748 | Benign | -4.29 | Benign | 0.17 | Neutral | 0.01 | Benign | 0.41 | Tolerated | 22.90 | Deleterious |
| p.His66Ser | VUS | -0.93 | Neutral | 0.5326 | Ambiguous | -5.19 | Benign | 0.28 | Neutral | 1.00 | Probably damaging | 0.04 | Deleterious | | |
| p.His66Ile | VUS | -7.66 | Indeterminate | 0.8784 | Pathogenic | -12.43 | Pathogenic | 0.51 | Disease | 0.99 | Probably damaging | 0.03 | Deleterious | | |
| p.His66Met | VUS | -0.24 | Neutral | 0.8768 | Pathogenic | -8.28 | Pathogenic | 0.47 | Neutral | 1.00 | Probably damaging | 0.02 | Deleterious | | |
| p.His66Gln | VUS | -1.10 | Neutral | 0.4119 | Ambiguous | -6.28 | Benign | 0.24 | Neutral | 0.99 | Probably damaging | 0.29 | Tolerated | 22.85 | Deleterious |
| p.His66Pro | VUS | -53.15 | Deleterious | 0.7036 | Pathogenic | -10.87 | Pathogenic | 0.51 | Disease | 1.00 | Probably damaging | 0.03 | Deleterious | 25.00 | Deleterious |
| p.His66Leu | VUS | -1.55 | Neutral | 0.467 | Ambiguous | -6.67 | Benign | 0.28 | Neutral | 0.70 | Possibly damaging | 0.06 | Tolerated | 23.40 | Deleterious |
| p.His66Asp | VUS | -1.70 | Neutral | 0.7664 | Pathogenic | -8.82 | Pathogenic | 0.47 | Neutral | 1.00 | Probably damaging | 0.02 | Deleterious | 24.00 | Deleterious |
| p.His66Glu | VUS | -3.12 | Neutral | 0.6101 | Pathogenic | -6.34 | Benign | 0.43 | Neutral | 1.00 | Probably damaging | 0.05 | Deleterious | | |
| p.His66Ala | VUS | -2.15 | Neutral | 0.4953 | Ambiguous | -2.59 | Benign | 0.31 | Neutral | 1.00 | Probably damaging | 0.05 | Deleterious | | |
| p.His66Gly | VUS | -0.74 | Neutral | 0.5323 | Ambiguous | -5.55 | Benign | 0.46 | Neutral | 1.00 | Probably damaging | 0.04 | Deleterious | | |
| p.His66Val | VUS | -5.55 | Neutral | 0.735 | Pathogenic | -8.00 | Pathogenic | 0.46 | Neutral | 1.00 | Probably damaging | 0.04 | Deleterious | | |
| p.His66Tyr | VUS | -1.24 | Neutral | 0.1475 | Benign | -5.71 | Benign | 0.25 | Neutral | 0.95 | Possibly damaging | 0.54 | Tolerated | 18.56 | Deleterious |
| p.His66Cys | VUS | -0.01 | Neutral | 0.3504 | Ambiguous | -7.06 | Benign | 0.46 | Neutral | 1.00 | Probably damaging | 0.01 | Deleterious | | |
| p.His66Trp | VUS | -0.25 | Neutral | 0.4002 | Ambiguous | -6.01 | Benign | 0.50 | Neutral | 1.00 | Probably damaging | 0.02 | Deleterious | | |
| p.His66Phe | VUS | -0.70 | Neutral | 0.4537 | Ambiguous | -7.03 | Benign | 0.44 | Neutral | 0.99 | Probably damaging | 0.05 | Deleterious | | |
| p.Gly67Asn | VUS | -0.13 | Neutral | 0.7348 | Pathogenic | -10.55 | Pathogenic | 0.66 | Disease | 1.00 | Probably damaging | 0 | Deleterious | | |
| p.Gly67Lys | VUS | -0.88 | Neutral | 0.7226 | Pathogenic | -10.78 | Pathogenic | 0.69 | Disease | 1.00 | Probably damaging | 0 | Deleterious | | |
| p.Gly67Thr | VUS | -9.71 | Indeterminate | 0.7928 | Pathogenic | -13.00 | Pathogenic | 0.66 | Disease | 1.00 | Probably damaging | 0 | Deleterious | | |
| p.Gly67Arg | VUS | 0.00 | Neutral | 0.5034 | Ambiguous | -8.24 | Pathogenic | 0.76 | Disease | 1.00 | Probably damaging | 0 | Deleterious | 28.20 | Deleterious |
| p.Gly67Ser | VUS | -0.07 | Neutral | 0.4193 | Ambiguous | -9.72 | Pathogenic | 0.68 | Disease | 1.00 | Probably damaging | 0 | Deleterious | 27.90 | Deleterious |
| p.Gly67Ile | VUS | -53.15 | Deleterious | 0.9677 | Pathogenic | -17.56 | Pathogenic | 0.72 | Disease | 1.00 | Probably damaging | 0 | Deleterious | | |
| p.Gly67Met | VUS | -2.62 | Neutral | 0.9648 | Pathogenic | -14.65 | Pathogenic | 0.73 | Disease | 1.00 | Probably damaging | 0 | Deleterious | | |
| p.Gly67His | VUS | -0.01 | Neutral | 0.687 | Pathogenic | -9.44 | Pathogenic | 0.70 | Disease | 1.00 | Probably damaging | 0 | Deleterious | | |
| p.Gly67Gln | VUS | 0.00 | Neutral | 0.6087 | Pathogenic | -10.71 | Pathogenic | 0.67 | Disease | 1.00 | Probably damaging | 0 | Deleterious | | |
| p.Gly67Pro | VUS | -53.15 | Deleterious | 0.9938 | Pathogenic | -15.59 | Pathogenic | 0.66 | Disease | 1.00 | Probably damaging | 0 | Deleterious | | |
| p.Gly67Leu | VUS | -0.43 | Neutral | 0.9284 | Pathogenic | -12.85 | Pathogenic | 0.71 | Disease | 1.00 | Probably damaging | 0 | Deleterious | | |
| p.Gly67Asp | VUS | -0.22 | Neutral | 0.4936 | Ambiguous | -9.88 | Pathogenic | 0.74 | Disease | 1.00 | Probably damaging | 0 | Deleterious | 26.10 | Deleterious |
| p.Gly67Glu | VUS | -0.17 | Neutral | 0.7102 | Pathogenic | -11.05 | Pathogenic | 0.76 | Disease | 1.00 | Probably damaging | 0 | Deleterious | | |
| p.Gly67Ala | VUS | 0.00 | Neutral | 0.5522 | Ambiguous | -9.56 | Pathogenic | 0.49 | Neutral | 1.00 | Probably damaging | 0 | Deleterious | 25.00 | Deleterious |
| p.Gly67Val | VUS | -53.15 | Deleterious | 0.9212 | Pathogenic | -13.26 | Pathogenic | 0.81 | Disease | 1.00 | Probably damaging | 0 | Deleterious | 25.90 | Deleterious |
| p.Gly67Tyr | VUS | -0.17 | Neutral | 0.9176 | Pathogenic | -13.91 | Pathogenic | 0.73 | Disease | 1.00 | Probably damaging | 0 | Deleterious | | |
| p.Gly67Cys | VUS | -1.96 | Neutral | 0.7922 | Pathogenic | -11.77 | Pathogenic | 0.56 | Disease | 1.00 | Probably damaging | 0 | Deleterious | 29.30 | Deleterious |
| p.Gly67Trp | VUS | -5.92 | Indeterminate | 0.9304 | Pathogenic | -13.32 | Pathogenic | 0.75 | Disease | 1.00 | Probably damaging | 0 | Deleterious | | |
| p.Gly67Phe | VUS | -1.32 | Neutral | 0.9653 | Pathogenic | -14.69 | Pathogenic | 0.72 | Disease | 1.00 | Probably damaging | 0 | Deleterious | | |
| p.Ala68Asn | VUS | -53.15 | Deleterious | 0.9679 | Pathogenic | -18.24 | Pathogenic | 0.75 | Disease | 1.00 | Probably damaging | 0 | Deleterious | | |
| p.Ala68Lys | VUS | -53.15 | Deleterious | 0.9925 | Pathogenic | -22.22 | Pathogenic | 0.78 | Disease | 1.00 | Probably damaging | 0 | Deleterious | | |
| p.Ala68Thr | VUS | -2.88 | Neutral | 0.5914 | Pathogenic | -11.27 | Pathogenic | 0.73 | Disease | 1.00 | Probably damaging | 0 | Deleterious | 26.20 | Deleterious |
| p.Ala68Arg | VUS | -53.15 | Deleterious | 0.9558 | Pathogenic | -20.11 | Pathogenic | 0.76 | Disease | 1.00 | Probably damaging | 0 | Deleterious | | |
| p.Ala68Ser | VUS | -0.59 | Neutral | 0.293 | Benign | -10.10 | Pathogenic | 0.54 | Disease | 0.99 | Probably damaging | 0 | Deleterious | 25.50 | Deleterious |
| p.Ala68Ile | VUS | -53.15 | Deleterious | 0.9189 | Pathogenic | -17.35 | Pathogenic | 0.78 | Disease | 1.00 | Probably damaging | 0 | Deleterious | | |
| p.Ala68Met | VUS | -33.17 | Indeterminate | 0.9229 | Pathogenic | -16.44 | Pathogenic | 0.72 | Disease | 1.00 | Probably damaging | 0 | Deleterious | | |
| p.Ala68His | VUS | -24.88 | Indeterminate | 0.9743 | Pathogenic | -19.04 | Pathogenic | 0.79 | Disease | 1.00 | Probably damaging | 0 | Deleterious | | |
| p.Ala68Gln | VUS | -53.15 | Deleterious | 0.9384 | Pathogenic | -18.21 | Pathogenic | 0.77 | Disease | 1.00 | Probably damaging | 0 | Deleterious | | |
| p.Ala68Pro | VUS | -53.15 | Deleterious | 0.9539 | Pathogenic | -15.22 | Pathogenic | 0.82 | Disease | 1.00 | Probably damaging | 0 | Deleterious | 26.30 | Deleterious |
| p.Ala68Leu | Likely pathogenic | -53.15 | Deleterious | 0.8086 | Pathogenic | -16.58 | Pathogenic | 0.76 | Disease | 1.00 | Probably damaging | 0 | Deleterious | | |
| p.Ala68Asp | VUS | -32.95 | Indeterminate | 0.9549 | Pathogenic | -15.96 | Pathogenic | 0.84 | Disease | 1.00 | Probably damaging | 0 | Deleterious | | |
| p.Ala68Glu | VUS | -32.02 | Indeterminate | 0.9619 | Pathogenic | -16.60 | Pathogenic | 0.81 | Disease | 1.00 | Probably damaging | 0 | Deleterious | 25.70 | Deleterious |
| p.Ala68Gly | VUS | -1.38 | Neutral | 0.3176 | Benign | -8.95 | Pathogenic | 0.62 | Disease | 1.00 | Probably damaging | 0 | Deleterious | 26.00 | Deleterious |
| p.Ala68Val | VUS | -10.00 | Indeterminate | 0.676 | Pathogenic | -10.83 | Pathogenic | 0.70 | Disease | 1.00 | Probably damaging | 0 | Deleterious | 26.10 | Deleterious |
| p.Ala68Tyr | VUS | -53.15 | Deleterious | 0.9825 | Pathogenic | -19.51 | Pathogenic | 0.78 | Disease | 1.00 | Probably damaging | 0 | Deleterious | | |
| p.Ala68Cys | VUS | -0.84 | Neutral | 0.7551 | Pathogenic | -11.43 | Pathogenic | 0.70 | Disease | 1.00 | Probably damaging | 0 | Deleterious | | |
| p.Ala68Trp | VUS | -53.15 | Deleterious | 0.9882 | Pathogenic | -19.71 | Pathogenic | 0.78 | Disease | 1.00 | Probably damaging | 0 | Deleterious | | |
| p.Ala68Phe | VUS | -53.15 | Deleterious | 0.9666 | Pathogenic | -18.42 | Pathogenic | 0.79 | Disease | 1.00 | Probably damaging | 0 | Deleterious | | |
| p.Glu69Asn | VUS | -53.15 | Deleterious | 0.3183 | Benign | -3.89 | Benign | 0.21 | Neutral | 0.04 | Benign | 1 | Tolerated | | |
| p.Glu69Lys | VUS | -33.00 | Indeterminate | 0.472 | Ambiguous | -9.75 | Pathogenic | 0.25 | Neutral | 0.00 | Benign | 0.49 | Tolerated | 22.00 | Deleterious |
| p.Glu69Thr | VUS | -9.93 | Indeterminate | 0.4956 | Ambiguous | -8.39 | Pathogenic | 0.33 | Neutral | 0.11 | Benign | 0.22 | Tolerated | | |
| p.Glu69Arg | VUS | -1.75 | Neutral | 0.3922 | Ambiguous | -8.31 | Pathogenic | 0.24 | Neutral | 0.01 | Benign | 0.18 | Tolerated | | |
| p.Glu69Ser | VUS | -13.93 | Indeterminate | 0.3272 | Benign | -5.69 | Benign | 0.33 | Neutral | 0.04 | Benign | 0.51 | Tolerated | | |
| p.Glu69Ile | VUS | -3.57 | Neutral | 0.7085 | Pathogenic | -11.52 | Pathogenic | 0.37 | Neutral | 0.09 | Benign | 0.06 | Tolerated | | |
| p.Glu69Met | VUS | -30.66 | Indeterminate | 0.7565 | Pathogenic | -10.83 | Pathogenic | 0.38 | Neutral | 0.17 | Benign | 0.04 | Deleterious | | |
| p.Glu69His | VUS | -7.51 | Indeterminate | 0.5094 | Ambiguous | -6.31 | Benign | 0.30 | Neutral | 0.13 | Benign | 0.09 | Tolerated | | |
| p.Glu69Gln | VUS | -6.51 | Indeterminate | 0.2181 | Benign | -6.81 | Benign | 0.19 | Neutral | 0.01 | Benign | 0.25 | Tolerated | 20.60 | Deleterious |
| p.Glu69Pro | VUS | -53.15 | Deleterious | 0.6727 | Pathogenic | -10.70 | Pathogenic | 0.40 | Neutral | 0.06 | Benign | 0.13 | Tolerated | | |
| p.Glu69Leu | VUS | -33.22 | Indeterminate | 0.5672 | Pathogenic | -8.78 | Pathogenic | 0.35 | Neutral | 0.01 | Benign | 0.1 | Tolerated | | |
| p.Glu69Asp | VUS | -5.22 | Neutral | 0.0782 | Benign | 0.95 | Benign | 0.09 | Neutral | 0.00 | Benign | 0.65 | Tolerated | 5.42 | Neutral |
| p.Glu69Ala | VUS | -0.17 | Neutral | 0.2904 | Benign | -6.89 | Benign | 0.27 | Neutral | 0.01 | Benign | 0.28 | Tolerated | 23.60 | Deleterious |
| p.Glu69Gly | VUS | -13.03 | Indeterminate | 0.5256 | Ambiguous | -8.23 | Pathogenic | 0.26 | Neutral | 0.36 | Benign | 0.27 | Tolerated | 24.00 | Deleterious |
| p.Glu69Val | VUS | -25.39 | Indeterminate | 0.4395 | Ambiguous | -9.19 | Pathogenic | 0.32 | Neutral | 0.01 | Benign | 0.09 | Tolerated | 24.10 | Deleterious |
| p.Glu69Tyr | VUS | -8.93 | Indeterminate | 0.7874 | Pathogenic | -8.99 | Pathogenic | 0.36 | Neutral | 0.36 | Benign | 0.04 | Deleterious | | |
| p.Glu69Cys | VUS | -5.94 | Indeterminate | 0.8994 | Pathogenic | -7.55 | Pathogenic | 0.39 | Neutral | 1.00 | Probably damaging | 0.03 | Deleterious | | |
| p.Glu69Trp | VUS | -19.38 | Indeterminate | 0.9321 | Pathogenic | -10.11 | Pathogenic | 0.40 | Neutral | 0.86 | Possibly damaging | 0.01 | Deleterious | | |
| p.Glu69Phe | VUS | -11.30 | Indeterminate | 0.8778 | Pathogenic | -8.34 | Pathogenic | 0.38 | Neutral | 0.39 | Benign | 0.03 | Deleterious | | |
| p.Pro70Asn | VUS | -3.10 | Neutral | 0.9617 | Pathogenic | -19.19 | Pathogenic | 0.66 | Disease | 1.00 | Probably damaging | 0.44 | Tolerated | | |
| p.Pro70Lys | VUS | -33.21 | Indeterminate | 0.8971 | Pathogenic | -17.55 | Pathogenic | 0.71 | Disease | 1.00 | Probably damaging | 0.25 | Tolerated | | |
| p.Pro70Thr | VUS | -4.42 | Neutral | 0.4658 | Ambiguous | -10.15 | Pathogenic | 0.61 | Disease | 1.00 | Probably damaging | 0.97 | Tolerated | 24.30 | Deleterious |
| p.Pro70Arg | VUS | -0.46 | Neutral | 0.76 | Pathogenic | -14.70 | Pathogenic | 0.71 | Disease | 1.00 | Probably damaging | 0.16 | Tolerated | 27.30 | Deleterious |
| p.Pro70Ser | VUS | -32.63 | Indeterminate | 0.6826 | Pathogenic | -12.10 | Pathogenic | 0.31 | Neutral | 1.00 | Probably damaging | 1 | Tolerated | 24.50 | Deleterious |
| p.Pro70Ile | VUS | -0.38 | Neutral | 0.628 | Pathogenic | -9.42 | Pathogenic | 0.67 | Disease | 1.00 | Probably damaging | 0.18 | Tolerated | | |
| p.Pro70Met | VUS | 0.00 | Neutral | 0.8289 | Pathogenic | -11.40 | Pathogenic | 0.67 | Disease | 1.00 | Probably damaging | 0.08 | Tolerated | | |
| p.Pro70His | VUS | 0.00 | Neutral | 0.9162 | Pathogenic | -15.45 | Pathogenic | 0.66 | Disease | 1.00 | Probably damaging | 0.09 | Tolerated | 27.90 | Deleterious |
| p.Pro70Gln | VUS | 0.00 | Neutral | 0.805 | Pathogenic | -14.98 | Pathogenic | 0.63 | Disease | 1.00 | Probably damaging | 0.17 | Tolerated | | |
| p.Pro70Leu | VUS | -3.39 | Neutral | 0.6032 | Pathogenic | -9.00 | Pathogenic | 0.68 | Disease | 1.00 | Probably damaging | 0.19 | Tolerated | 28.80 | Deleterious |
| p.Pro70Asp | VUS | -53.15 | Deleterious | 0.9737 | Pathogenic | -19.20 | Pathogenic | 0.70 | Disease | 1.00 | Probably damaging | 0.29 | Tolerated | | |
| p.Pro70Glu | VUS | -53.15 | Deleterious | 0.8773 | Pathogenic | -17.44 | Pathogenic | 0.65 | Disease | 1.00 | Probably damaging | 0.22 | Tolerated | | |
| p.Pro70Ala | VUS | -2.01 | Neutral | 0.2232 | Benign | -7.48 | Benign | 0.33 | Neutral | 1.00 | Probably damaging | 0.82 | Tolerated | 23.80 | Deleterious |
| p.Pro70Gly | VUS | -2.54 | Neutral | 0.7794 | Pathogenic | -11.94 | Pathogenic | 0.61 | Disease | 1.00 | Probably damaging | 0.44 | Tolerated | | |
| p.Pro70Val | VUS | -0.02 | Neutral | 0.363 | Ambiguous | -5.39 | Benign | 0.64 | Disease | 1.00 | Probably damaging | 0.57 | Tolerated | | |
| p.Pro70Tyr | VUS | -1.90 | Neutral | 0.9689 | Pathogenic | -17.90 | Pathogenic | 0.66 | Disease | 1.00 | Probably damaging | 0.09 | Tolerated | | |
| p.Pro70Cys | VUS | 0.00 | Neutral | 0.8804 | Pathogenic | -13.04 | Pathogenic | 0.66 | Disease | 1.00 | Probably damaging | 0.11 | Tolerated | | |

| | | | | | | | | | | | | | | | |
|---|---|---|---|---|---|---|---|---|---|---|---|---|---|---|---|
| p.Pro70Trp | VUS | -53.15 | Deleterious | 0.9728 | Pathogenic | -16.09 | Pathogenic | 0.64 | Disease | 1.00 | Probably damaging | 0.03 | Deleterious | | |
| p.Pro70Phe | VUS | -0.11 | Neutral | 0.9674 | Pathogenic | -15.59 | Pathogenic | 0.68 | Disease | 1.00 | Probably damaging | 0.08 | Tolerated | | |
| p.Asn71Lys | VUS | -53.15 | Deleterious | 0.9616 | Pathogenic | -13.50 | Pathogenic | 0.76 | Disease | 1.00 | Probably damaging | 0.02 | Deleterious | 24.40 | Deleterious |
| p.Asn71Thr | VUS | -53.15 | Deleterious | 0.5918 | Pathogenic | -7.81 | Pathogenic | 0.69 | Disease | 0.98 | Probably damaging | 0.02 | Deleterious | 26.70 | Deleterious |
| p.Asn71Arg | VUS | -53.15 | Deleterious | 0.9005 | Pathogenic | -9.38 | Pathogenic | 0.69 | Disease | 1.00 | Probably damaging | 0.12 | Tolerated | | |
| p.Asn71Ser | Likely pathogenic | -53.15 | Deleterious | 0.2129 | Benign | -6.58 | Benign | 0.67 | Disease | 1.00 | Probably damaging | 0.04 | Deleterious | 25.30 | Deleterious |
| p.Asn71Ile | VUS | -53.15 | Deleterious | 0.9352 | Pathogenic | -11.05 | Pathogenic | 0.83 | Disease | 1.00 | Probably damaging | 0 | Deleterious | 28.10 | Deleterious |
| p.Asn71Met | VUS | -53.15 | Deleterious | 0.9288 | Pathogenic | -9.36 | Pathogenic | 0.77 | Disease | 1.00 | Probably damaging | 0 | Deleterious | | |
| p.Asn71His | VUS | -20.64 | Indeterminate | 0.5902 | Pathogenic | -8.69 | Pathogenic | 0.75 | Disease | 1.00 | Probably damaging | 0.02 | Deleterious | 25.90 | Deleterious |
| p.Asn71Gln | VUS | -13.47 | Indeterminate | 0.8669 | Pathogenic | -9.79 | Pathogenic | 0.67 | Disease | 1.00 | Probably damaging | 0.02 | Deleterious | | |
| p.Asn71Pro | VUS | -53.15 | Deleterious | 0.9656 | Pathogenic | -11.37 | Pathogenic | 0.73 | Disease | 1.00 | Probably damaging | 0.01 | Deleterious | | |
| p.Asn71Leu | VUS | -53.15 | Deleterious | 0.8076 | Pathogenic | -9.54 | Pathogenic | 0.74 | Disease | 1.00 | Probably damaging | 0 | Deleterious | | |
| p.Asn71Asp | VUS | -26.26 | Indeterminate | 0.223 | Benign | -2.53 | Benign | 0.43 | Neutral | 1.00 | Probably damaging | 0.16 | Tolerated | 24.00 | Deleterious |
| p.Asn71Glu | VUS | -12.42 | Indeterminate | 0.7821 | Pathogenic | -6.87 | Benign | 0.63 | Disease | 0.99 | Probably damaging | 0.02 | Deleterious | | |
| p.Asn71Ala | VUS | -20.51 | Indeterminate | 0.7043 | Pathogenic | -6.89 | Benign | 0.71 | Disease | 0.99 | Probably damaging | 0.02 | Deleterious | | |
| p.Asn71Gly | VUS | -1.93 | Neutral | 0.5961 | Pathogenic | -7.24 | Benign | 0.67 | Disease | 1.00 | Probably damaging | 0.05 | Deleterious | | |
| p.Asn71Val | VUS | -53.15 | Deleterious | 0.8789 | Pathogenic | -9.19 | Pathogenic | 0.75 | Disease | 0.98 | Probably damaging | 0 | Deleterious | | |
| p.Asn71Tyr | VUS | -53.15 | Deleterious | 0.7956 | Pathogenic | -12.45 | Pathogenic | 0.82 | Disease | 1.00 | Probably damaging | 0.01 | Deleterious | 26.80 | Deleterious |
| p.Asn71Cys | VUS | -16.50 | Indeterminate | 0.8283 | Pathogenic | -5.76 | Benign | 0.73 | Disease | 1.00 | Probably damaging | 0 | Deleterious | | |
| p.Asn71Trp | VUS | -53.15 | Deleterious | 0.9803 | Pathogenic | -11.43 | Pathogenic | 0.77 | Disease | 1.00 | Probably damaging | 0 | Deleterious | | |
| p.Asn71Phe | VUS | -53.15 | Deleterious | 0.9648 | Pathogenic | -11.10 | Pathogenic | 0.76 | Disease | 1.00 | Probably damaging | 0 | Deleterious | | |
| p.Cys72Asn | VUS | -5.60 | Neutral | 0.4565 | Ambiguous | -7.49 | Benign | 0.43 | Neutral | 0.79 | Possibly damaging | 0.27 | Tolerated | | |
| p.Cys72Lys | VUS | -6.71 | Indeterminate | 0.7865 | Pathogenic | -9.86 | Pathogenic | 0.43 | Neutral | 0.67 | Possibly damaging | 0.51 | Tolerated | | |
| p.Cys72Thr | VUS | -3.40 | Neutral | 0.256 | Benign | -3.64 | Benign | 0.25 | Neutral | 0.52 | Possibly damaging | 0.56 | Tolerated | | |
| p.Cys72Arg | VUS | -0.61 | Neutral | 0.3115 | Benign | -6.41 | Benign | 0.11 | Neutral | 0.01 | Benign | 0.3 | Tolerated | 14.36 | Neutral |
| p.Cys72Ser | VUS | -8.34 | Indeterminate | 0.2006 | Benign | -3.02 | Benign | 0.35 | Neutral | 0.98 | Probably damaging | 0.41 | Tolerated | 16.24 | Deleterious |
| p.Cys72Ile | VUS | -7.24 | Indeterminate | 0.3465 | Ambiguous | -6.65 | Benign | 0.42 | Neutral | 0.97 | Probably damaging | 0.49 | Tolerated | | |
| p.Cys72Met | VUS | -3.24 | Neutral | 0.5647 | Pathogenic | -6.74 | Benign | 0.43 | Neutral | 0.99 | Probably damaging | 0.16 | Tolerated | | |
| p.Cys72His | VUS | -10.13 | Indeterminate | 0.3639 | Ambiguous | -5.64 | Benign | 0.42 | Neutral | 0.98 | Probably damaging | 0.17 | Tolerated | | |
| p.Cys72Gln | VUS | -12.58 | Indeterminate | 0.4744 | Ambiguous | -5.57 | Benign | 0.41 | Neutral | 0.84 | Possibly damaging | 0.38 | Tolerated | | |
| p.Cys72Pro | VUS | -7.71 | Indeterminate | 0.7099 | Pathogenic | -5.40 | Benign | 0.43 | Neutral | 0.92 | Possibly damaging | 0.22 | Tolerated | | |
| p.Cys72Leu | VUS | -19.97 | Indeterminate | 0.2665 | Benign | -4.26 | Benign | 0.26 | Neutral | 0.80 | Possibly damaging | 0.55 | Tolerated | | |
| p.Cys72Asp | VUS | -1.91 | Neutral | 0.4226 | Ambiguous | -4.46 | Benign | 0.46 | Neutral | 0.78 | Possibly damaging | 0.29 | Tolerated | | |
| p.Cys72Glu | VUS | -3.08 | Neutral | 0.5005 | Ambiguous | -3.91 | Benign | 0.43 | Neutral | 0.84 | Possibly damaging | 0.48 | Tolerated | | |
| p.Cys72Ala | VUS | -4.81 | Neutral | 0.1581 | Benign | -0.64 | Benign | 0.27 | Neutral | 0.97 | Probably damaging | 0.79 | Tolerated | | |
| p.Cys72Gly | VUS | -2.97 | Neutral | 0.134 | Benign | -3.04 | Benign | 0.37 | Neutral | 0.99 | Probably damaging | 0.21 | Tolerated | 15.02 | Deleterious |
| p.Cys72Val | VUS | -5.23 | Neutral | 0.2264 | Benign | -3.23 | Benign | 0.21 | Neutral | 0.75 | Possibly damaging | 1 | Tolerated | | |
| p.Cys72Tyr | VUS | -0.68 | Neutral | 0.3384 | Benign | -7.29 | Benign | 0.19 | Neutral | 0.99 | Probably damaging | 0.23 | Tolerated | 18.34 | Deleterious |
| p.Cys72Trp | VUS | -11.02 | Indeterminate | 0.4589 | Ambiguous | -8.48 | Pathogenic | 0.40 | Neutral | 1.00 | Probably damaging | 0.05 | Deleterious | 24.00 | Deleterious |
| p.Cys72Phe | VUS | -8.51 | Indeterminate | 0.1717 | Benign | -6.38 | Benign | 0.37 | Neutral | 0.99 | Probably damaging | 0.2 | Tolerated | 19.45 | Deleterious |
| p.Ala73Asn | VUS | -7.83 | Indeterminate | 0.4928 | Ambiguous | -12.71 | Pathogenic | 0.38 | Neutral | 0.50 | Possibly damaging | 0.3 | Tolerated | | |
| p.Ala73Lys | VUS | -2.90 | Neutral | 0.6144 | Pathogenic | -15.55 | Pathogenic | 0.24 | Neutral | 0.82 | Possibly damaging | 0.65 | Tolerated | | |
| p.Ala73Thr | VUS | -0.56 | Neutral | 0.132 | Benign | -6.87 | Benign | 0.17 | Neutral | 0.87 | Possibly damaging | 0.35 | Tolerated | 23.20 | Deleterious |
| p.Ala73Arg | VUS | -4.04 | Neutral | 0.3606 | Ambiguous | -9.94 | Pathogenic | 0.24 | Neutral | 0.97 | Probably damaging | 0.31 | Tolerated | | |
| p.Ala73Ser | VUS | -3.48 | Neutral | 0.1403 | Benign | -5.74 | Benign | 0.30 | Neutral | 0.26 | Benign | 0.44 | Tolerated | 22.90 | Deleterious |
| p.Ala73Ile | VUS | -1.38 | Neutral | 0.3864 | Ambiguous | -10.33 | Pathogenic | 0.39 | Neutral | 0.99 | Probably damaging | 0.11 | Tolerated | | |
| p.Ala73Met | VUS | -2.96 | Neutral | 0.4562 | Ambiguous | -10.25 | Pathogenic | 0.40 | Neutral | 1.00 | Probably damaging | 0.07 | Tolerated | | |
| p.Ala73His | VUS | -0.20 | Neutral | 0.4991 | Ambiguous | -10.25 | Pathogenic | 0.39 | Neutral | 1.00 | Probably damaging | 0.09 | Tolerated | | |
| p.Ala73Gln | VUS | -5.57 | Neutral | 0.3172 | Benign | -9.39 | Pathogenic | 0.21 | Neutral | 0.94 | Possibly damaging | 0.45 | Tolerated | | |
| p.Ala73Pro | VUS | -2.47 | Neutral | 0.1182 | Benign | -4.70 | Benign | 0.16 | Neutral | 0.97 | Probably damaging | 0.23 | Tolerated | 20.90 | Deleterious |
| p.Ala73Leu | VUS | -1.54 | Neutral | 0.2858 | Benign | -7.24 | Benign | 0.38 | Neutral | 0.90 | Possibly damaging | 0.18 | Tolerated | | |
| p.Ala73Asp | VUS | -5.39 | Neutral | 0.3899 | Ambiguous | -12.17 | Pathogenic | 0.40 | Neutral | 0.48 | Possibly damaging | 0.58 | Tolerated | 21.90 | Deleterious |
| p.Ala73Glu | VUS | -1.95 | Neutral | 0.2439 | Benign | -10.59 | Pathogenic | 0.16 | Neutral | 0.01 | Benign | 1 | Tolerated | | |
| p.Ala73Gly | VUS | -2.99 | Neutral | 0.1805 | Benign | -6.60 | Benign | 0.19 | Neutral | 0.72 | Possibly damaging | 0.24 | Tolerated | 21.00 | Deleterious |
| p.Ala73Val | VUS | -2.21 | Neutral | 0.1819 | Benign | -6.67 | Benign | 0.20 | Neutral | 0.84 | Possibly damaging | 0.17 | Tolerated | 17.40 | Deleterious |
| p.Ala73Tyr | VUS | -4.06 | Neutral | 0.6755 | Pathogenic | -12.54 | Pathogenic | 0.42 | Neutral | 1.00 | Probably damaging | 0.03 | Deleterious | | |
| p.Ala73Cys | VUS | -2.99 | Neutral | 0.4259 | Ambiguous | -8.85 | Pathogenic | 0.36 | Neutral | 1.00 | Probably damaging | 0.03 | Deleterious | | |
| p.Ala73Trp | VUS | -7.09 | Indeterminate | 0.7274 | Pathogenic | -11.99 | Pathogenic | 0.48 | Neutral | 1.00 | Probably damaging | 0.01 | Deleterious | | |
| p.Ala73Phe | VUS | -3.00 | Neutral | 0.5232 | Ambiguous | -10.97 | Pathogenic | 0.39 | Neutral | 1.00 | Probably damaging | 0.03 | Deleterious | | |
| p.Asp74Asn | VUS | -53.15 | Deleterious | 0.8381 | Pathogenic | -9.62 | Pathogenic | 0.80 | Disease | 1.00 | Probably damaging | 0.17 | Tolerated | 31.00 | Deleterious |
| p.Asp74Lys | VUS | -53.15 | Deleterious | 0.9891 | Pathogenic | -14.48 | Pathogenic | 0.80 | Disease | 1.00 | Probably damaging | 0.01 | Deleterious | | |
| p.Asp74Thr | VUS | -53.15 | Deleterious | 0.9723 | Pathogenic | -10.83 | Pathogenic | 0.79 | Disease | 1.00 | Probably damaging | 0.01 | Deleterious | | |
| p.Asp74Arg | VUS | -53.15 | Deleterious | 0.9747 | Pathogenic | -13.09 | Pathogenic | 0.80 | Disease | 1.00 | Probably damaging | 0.01 | Deleterious | | |
| p.Asp74Ser | VUS | -12.50 | Indeterminate | 0.8258 | Pathogenic | -11.42 | Pathogenic | 0.74 | Disease | 1.00 | Probably damaging | 0.02 | Deleterious | | |
| p.Asp74Ile | VUS | -53.15 | Deleterious | 0.9679 | Pathogenic | -14.48 | Pathogenic | 0.80 | Disease | 1.00 | Probably damaging | 0 | Deleterious | | |
| p.Asp74Met | VUS | -53.15 | Deleterious | 0.9937 | Pathogenic | -15.23 | Pathogenic | 0.81 | Disease | 1.00 | Probably damaging | 0 | Deleterious | | |
| p.Asp74His | VUS | -5.47 | Neutral | 0.8861 | Pathogenic | -9.18 | Pathogenic | 0.90 | Disease | 1.00 | Probably damaging | 0.01 | Deleterious | 29.80 | Deleterious |
| p.Asp74Gln | VUS | -53.15 | Deleterious | 0.9741 | Pathogenic | -12.13 | Pathogenic | 0.79 | Disease | 1.00 | Probably damaging | 0.01 | Deleterious | | |
| p.Asp74Pro | VUS | -53.15 | Deleterious | 0.9834 | Pathogenic | -14.64 | Pathogenic | 0.75 | Disease | 0.98 | Probably damaging | 0.01 | Deleterious | | |
| p.Asp74Leu | VUS | -53.15 | Deleterious | 0.95 | Pathogenic | -13.73 | Pathogenic | 0.79 | Disease | 1.00 | Probably damaging | 0 | Deleterious | | |
| p.Asp74Glu | VUS | 0.00 | Neutral | 0.867 | Pathogenic | -7.60 | Pathogenic | 0.78 | Disease | 0.89 | Possibly damaging | 0.01 | Deleterious | 24.05 | Deleterious |
| p.Asp74Ala | VUS | -53.15 | Deleterious | 0.8943 | Pathogenic | -10.97 | Pathogenic | 0.85 | Disease | 1.00 | Probably damaging | 0.01 | Deleterious | 29.20 | Deleterious |
| p.Asp74Gly | VUS | -53.15 | Deleterious | 0.9501 | Pathogenic | -11.00 | Pathogenic | 0.88 | Disease | 1.00 | Probably damaging | 0.03 | Deleterious | 29.20 | Deleterious |
| p.Asp74Val | VUS | -53.15 | Deleterious | 0.9289 | Pathogenic | -12.81 | Pathogenic | 0.90 | Disease | 1.00 | Probably damaging | 0 | Deleterious | 29.00 | Deleterious |
| p.Asp74Tyr | VUS | -53.15 | Deleterious | 0.8956 | Pathogenic | -13.08 | Pathogenic | 0.90 | Disease | 1.00 | Probably damaging | 0 | Deleterious | 30.00 | Deleterious |
| p.Asp74Cys | VUS | -0.01 | Neutral | 0.929 | Pathogenic | -11.52 | Pathogenic | 0.79 | Disease | 1.00 | Probably damaging | 0 | Deleterious | | |
| p.Asp74Trp | VUS | -53.15 | Deleterious | 0.9936 | Pathogenic | -14.65 | Pathogenic | 0.81 | Disease | 1.00 | Probably damaging | 0 | Deleterious | | |
| p.Asp74Phe | VUS | -53.15 | Deleterious | 0.9831 | Pathogenic | -15.30 | Pathogenic | 0.82 | Disease | 1.00 | Probably damaging | 0 | Deleterious | | |
| p.Pro75Asn | VUS | -2.21 | Neutral | 0.6079 | Pathogenic | -11.44 | Pathogenic | 0.44 | Neutral | 1.00 | Probably damaging | 0.68 | Tolerated | | |
| p.Pro75Lys | VUS | -4.73 | Neutral | 0.5067 | Ambiguous | -10.14 | Pathogenic | 0.31 | Neutral | 1.00 | Probably damaging | 0.83 | Tolerated | | |
| p.Pro75Thr | VUS | -1.80 | Neutral | 0.2037 | Benign | -8.88 | Pathogenic | 0.40 | Neutral | 1.00 | Probably damaging | 0.53 | Tolerated | 25.20 | Deleterious |
| p.Pro75Arg | VUS | -0.73 | Neutral | 0.249 | Benign | -7.82 | Pathogenic | 0.23 | Neutral | 1.00 | Probably damaging | 0.64 | Tolerated | 23.60 | Deleterious |
| p.Pro75Ser | VUS | -1.67 | Neutral | 0.2505 | Benign | -7.87 | Pathogenic | 0.17 | Neutral | 1.00 | Probably damaging | 0.57 | Tolerated | 24.00 | Deleterious |
| p.Pro75Ile | VUS | -2.22 | Neutral | 0.5196 | Ambiguous | -12.88 | Pathogenic | 0.52 | Disease | 1.00 | Probably damaging | 0.14 | Tolerated | | |
| p.Pro75Met | VUS | -4.48 | Neutral | 0.5736 | Pathogenic | -12.25 | Pathogenic | 0.48 | Neutral | 1.00 | Probably damaging | 0.08 | Tolerated | | |
| p.Pro75His | VUS | -0.31 | Neutral | 0.3661 | Ambiguous | -10.10 | Pathogenic | 0.43 | Neutral | 1.00 | Probably damaging | 0.13 | Tolerated | 26.30 | Deleterious |
| p.Pro75Gln | VUS | -3.70 | Neutral | 0.3196 | Benign | -9.25 | Pathogenic | 0.40 | Neutral | 1.00 | Probably damaging | 0.48 | Tolerated | | |
| p.Pro75Leu | VUS | -1.79 | Neutral | 0.2612 | Benign | -9.65 | Pathogenic | 0.30 | Neutral | 1.00 | Probably damaging | 0.24 | Tolerated | 27.00 | Deleterious |
| p.Pro75Asp | VUS | -1.93 | Neutral | 0.5776 | Pathogenic | -10.89 | Pathogenic | 0.46 | Neutral | 1.00 | Probably damaging | 0.54 | Tolerated | | |
| p.Pro75Glu | VUS | -1.03 | Neutral | 0.3748 | Ambiguous | -10.34 | Pathogenic | 0.43 | Neutral | 1.00 | Probably damaging | 1 | Tolerated | | |
| p.Pro75Ala | VUS | -1.00 | Neutral | 0.1356 | Benign | -7.19 | Benign | 0.28 | Neutral | 1.00 | Probably damaging | 0.55 | Tolerated | 23.10 | Deleterious |
| p.Pro75Gly | VUS | -1.33 | Neutral | 0.4257 | Ambiguous | -8.31 | Pathogenic | 0.30 | Neutral | 1.00 | Probably damaging | 0.33 | Tolerated | | |
| p.Pro75Val | VUS | -1.31 | Neutral | 0.3043 | Benign | -10.14 | Pathogenic | 0.44 | Neutral | 1.00 | Probably damaging | 0.21 | Tolerated | | |
| p.Pro75Tyr | VUS | -1.28 | Neutral | 0.7005 | Pathogenic | -13.03 | Pathogenic | 0.47 | Neutral | 1.00 | Probably damaging | 0.07 | Tolerated | | |
| p.Pro75Cys | VUS | -1.58 | Neutral | 0.7905 | Pathogenic | -11.33 | Pathogenic | 0.49 | Neutral | 1.00 | Probably damaging | 0.05 | Deleterious | | |
| p.Pro75Trp | VUS | -2.95 | Neutral | 0.7735 | Pathogenic | -12.09 | Pathogenic | 0.52 | Disease | 1.00 | Probably damaging | 0.02 | Deleterious | | |
| p.Pro75Phe | VUS | -1.54 | Neutral | 0.7006 | Pathogenic | -13.16 | Pathogenic | 0.50 | Neutral | 1.00 | Probably damaging | 0.06 | Tolerated | | |
| p.Ala76Asn | VUS | -13.40 | Indeterminate | 0.31 | Benign | -4.57 | Benign | 0.29 | Neutral | 0.00 | Benign | 0.3 | Tolerated | | |
| p.Ala76Lys | VUS | -10.54 | Indeterminate | 0.4685 | Ambiguous | -3.81 | Benign | 0.29 | Neutral | 0.80 | Possibly damaging | 0.74 | Tolerated | | |
| p.Ala76Thr | VUS | -2.83 | Neutral | 0.1157 | Benign | -4.13 | Benign | 0.17 | Neutral | 0.04 | Benign | 0.66 | Tolerated | 15.16 | Deleterious |
| p.Ala76Arg | VUS | -12.40 | Indeterminate | 0.3049 | Benign | -2.13 | Benign | 0.24 | Neutral | 0.97 | Probably damaging | 0.31 | Tolerated | | |
| p.Ala76Ser | VUS | -7.14 | Indeterminate | 0.1173 | Benign | -4.44 | Benign | 0.17 | Neutral | 0.03 | Benign | 0.51 | Tolerated | 14.18 | Neutral |
| p.Ala76Ile | VUS | -20.17 | Indeterminate | 0.3135 | Benign | -5.74 | Benign | 0.30 | Neutral | 0.67 | Possibly damaging | 0.11 | Tolerated | | |
| p.Ala76Met | VUS | -1.83 | Neutral | 0.3062 | Benign | -7.02 | Benign | 0.30 | Neutral | 0.97 | Probably damaging | 0.06 | Tolerated | | |
| p.Ala76His | VUS | -4.71 | Neutral | 0.3524 | Ambiguous | -5.19 | Benign | 0.24 | Neutral | 0.99 | Probably damaging | 0.07 | Tolerated | | |
| p.Ala76Gln | VUS | -16.22 | Indeterminate | 0.256 | Benign | -5.01 | Benign | 0.28 | Neutral | 0.97 | Probably damaging | 0.35 | Tolerated | | |
| p.Ala76Pro | VUS | -3.41 | Neutral | 0.3909 | Ambiguous | -7.15 | Benign | 0.28 | Neutral | 0.94 | Possibly damaging | 0.17 | Tolerated | 21.00 | Deleterious |
| p.Ala76Leu | VUS | -9.05 | Indeterminate | 0.1986 | Benign | -5.09 | Benign | 0.31 | Neutral | 0.45 | Benign | 0.18 | Tolerated | | |
| p.Ala76Asp | VUS | -4.79 | Neutral | 0.2923 | Benign | -6.19 | Benign | 0.27 | Neutral | 0.00 | Benign | 0.38 | Tolerated | 14.35 | Neutral |
| p.Ala76Glu | VUS | -8.61 | Indeterminate | 0.2402 | Benign | -6.08 | Benign | 0.24 | Neutral | 0.43 | Benign | 0.82 | Tolerated | | |
| p.Ala76Gly | VUS | -2.53 | Neutral | 0.1588 | Benign | -3.44 | Benign | 0.27 | Neutral | 0.32 | Benign | 0.35 | Tolerated | 13.15 | Neutral |

| | | | | | | | | | | | | | | | |
|---|---|---|---|---|---|---|---|---|---|---|---|---|---|---|---|
| p.Ala76Val | VUS | -3.78 | Neutral | 0.1339 | Benign | -3.22 | Benign | 0.16 | Neutral | 0.00 | Benign | 0.17 | Tolerated | 13.15 | Neutral |
| p.Ala76Tyr | VUS | -3.09 | Neutral | 0.4651 | Ambiguous | -7.67 | Pathogenic | 0.35 | Neutral | 0.97 | Probably damaging | 0.03 | Deleterious | | |
| p.Ala76Cys | VUS | -14.50 | Indeterminate | 0.4836 | Ambiguous | -5.99 | Benign | 0.32 | Neutral | 0.99 | Probably damaging | 0.03 | Deleterious | | |
| p.Ala76Trp | VUS | -1.24 | Neutral | 0.6791 | Pathogenic | -8.05 | Pathogenic | 0.37 | Neutral | 1.00 | Probably damaging | 0.01 | Deleterious | | |
| p.Ala76Phe | VUS | -6.82 | Indeterminate | 0.3737 | Ambiguous | -8.80 | Pathogenic | 0.34 | Neutral | 0.93 | Possibly damaging | 0.03 | Deleterious | | |
| p.Thr77Asn | VUS | -4.52 | Neutral | 0.4401 | Ambiguous | -3.96 | Benign | 0.53 | Disease | 1.00 | Probably damaging | 0.22 | Tolerated | 25.80 | Deleterious |
| p.Thr77Lys | VUS | -4.40 | Neutral | 0.6986 | Pathogenic | -9.95 | Pathogenic | 0.60 | Disease | 1.00 | Probably damaging | 0.15 | Tolerated | | |
| p.Thr77Arg | VUS | -12.26 | Indeterminate | 0.5249 | Ambiguous | -9.09 | Pathogenic | 0.57 | Disease | 1.00 | Probably damaging | 0.1 | Tolerated | | |
| p.Thr77Ser | VUS | -6.48 | Indeterminate | 0.2652 | Benign | -5.23 | Benign | 0.28 | Neutral | 0.99 | Probably damaging | 1 | Tolerated | 23.40 | Deleterious |
| p.Thr77Ile | VUS | -2.46 | Neutral | 0.8141 | Pathogenic | -9.67 | Pathogenic | 0.57 | Disease | 1.00 | Probably damaging | 0.09 | Tolerated | 26.80 | Deleterious |
| p.Thr77Met | VUS | -6.23 | Indeterminate | 0.3921 | Ambiguous | -9.21 | Pathogenic | 0.57 | Disease | 1.00 | Probably damaging | 0.05 | Deleterious | | |
| p.Thr77His | VUS | -4.95 | Neutral | 0.5887 | Pathogenic | -6.65 | Benign | 0.57 | Disease | 1.00 | Probably damaging | 0.05 | Deleterious | | |
| p.Thr77Gln | VUS | -5.14 | Neutral | 0.6083 | Pathogenic | -8.48 | Pathogenic | 0.55 | Disease | 1.00 | Probably damaging | 0.11 | Tolerated | | |
| p.Thr77Pro | VUS | -33.22 | Indeterminate | 0.5 | Ambiguous | -12.09 | Pathogenic | 0.35 | Neutral | 1.00 | Probably damaging | 0.19 | Tolerated | 26.00 | Deleterious |
| p.Thr77Leu | VUS | -5.03 | Neutral | 0.3219 | Benign | -8.76 | Pathogenic | 0.56 | Disease | 0.99 | Probably damaging | 0.13 | Tolerated | | |
| p.Thr77Asp | VUS | -7.25 | Indeterminate | 0.7915 | Pathogenic | -5.72 | Benign | 0.57 | Disease | 1.00 | Probably damaging | 0.15 | Tolerated | | |
| p.Thr77Glu | VUS | -16.56 | Indeterminate | 0.7672 | Pathogenic | -7.61 | Pathogenic | 0.56 | Disease | 1.00 | Probably damaging | 0.12 | Tolerated | | |
| p.Thr77Ala | VUS | -6.61 | Indeterminate | 0.276 | Benign | -6.63 | Benign | 0.51 | Disease | 0.98 | Probably damaging | 0.56 | Tolerated | 23.30 | Deleterious |
| p.Thr77Gly | VUS | -1.41 | Neutral | 0.3456 | Ambiguous | -7.91 | Pathogenic | 0.55 | Disease | 1.00 | Probably damaging | 0.3 | Tolerated | | |
| p.Thr77Val | VUS | -8.27 | Indeterminate | 0.5823 | Pathogenic | -9.17 | Pathogenic | 0.48 | Neutral | 0.99 | Probably damaging | 0.16 | Tolerated | | |
| p.Thr77Tyr | VUS | -2.86 | Neutral | 0.7131 | Pathogenic | -9.02 | Pathogenic | 0.59 | Disease | 1.00 | Probably damaging | 0.06 | Tolerated | | |
| p.Thr77Cys | VUS | -2.08 | Neutral | 0.7422 | Pathogenic | -7.94 | Pathogenic | 0.58 | Disease | 1.00 | Probably damaging | 0.08 | Tolerated | | |
| p.Thr77Trp | VUS | -3.70 | Neutral | 0.8587 | Pathogenic | -11.81 | Pathogenic | 0.59 | Disease | 1.00 | Probably damaging | 0.02 | Deleterious | | |
| p.Thr77Phe | VUS | -3.56 | Neutral | 0.7102 | Pathogenic | -10.34 | Pathogenic | 0.58 | Disease | 1.00 | Probably damaging | 0.05 | Deleterious | | |
| p.Leu78Asn | VUS | 0.00 | Neutral | 0.3776 | Ambiguous | -8.32 | Pathogenic | 0.47 | Neutral | 1.00 | Probably damaging | 0.09 | Tolerated | | |
| p.Leu78Lys | VUS | -0.10 | Neutral | 0.2747 | Benign | -7.82 | Pathogenic | 0.47 | Neutral | 0.99 | Probably damaging | 0.1 | Tolerated | | |
| p.Leu78Thr | VUS | -0.22 | Neutral | 0.3528 | Ambiguous | -8.99 | Pathogenic | 0.45 | Neutral | 1.00 | Probably damaging | 0.14 | Tolerated | | |
| p.Leu78Arg | VUS | -16.95 | Indeterminate | 0.1211 | Benign | -4.66 | Benign | 0.50 | Neutral | 1.00 | Probably damaging | 0.11 | Tolerated | 27.00 | Deleterious |
| p.Leu78Ser | VUS | 0.00 | Neutral | 0.3317 | Benign | -5.84 | Benign | 0.42 | Neutral | 1.00 | Probably damaging | 0.13 | Tolerated | | |
| p.Leu78Ile | VUS | -0.32 | Neutral | 0.3375 | Benign | -13.34 | Pathogenic | 0.34 | Neutral | 1.00 | Probably damaging | 0.38 | Tolerated | 18.47 | Deleterious |
| p.Leu78Met | VUS | -11.97 | Indeterminate | 0.1891 | Benign | -7.19 | Benign | 0.36 | Neutral | 1.00 | Probably damaging | 0.23 | Tolerated | | |
| p.Leu78His | VUS | -1.17 | Neutral | 0.2993 | Benign | -6.42 | Benign | 0.44 | Neutral | 1.00 | Probably damaging | 0.15 | Tolerated | 27.00 | Deleterious |
| p.Leu78Gln | VUS | -5.66 | Neutral | 0.1461 | Benign | -6.01 | Benign | 0.45 | Neutral | 1.00 | Probably damaging | 0.1 | Tolerated | | |
| p.Leu78Pro | VUS | -10.97 | Indeterminate | 0.4528 | Ambiguous | -11.56 | Pathogenic | 0.52 | Disease | 1.00 | Probably damaging | 0.08 | Tolerated | 27.50 | Deleterious |
| p.Leu78Asp | VUS | 0.00 | Neutral | 0.606 | Pathogenic | -9.01 | Pathogenic | 0.49 | Neutral | 1.00 | Probably damaging | 0.06 | Tolerated | | |
| p.Leu78Glu | VUS | -5.41 | Neutral | 0.3396 | Benign | -8.87 | Pathogenic | 0.48 | Neutral | 1.00 | Probably damaging | 0.09 | Tolerated | | |
| p.Leu78Ala | VUS | -14.32 | Indeterminate | 0.2132 | Benign | -5.39 | Benign | 0.40 | Neutral | 1.00 | Probably damaging | 0.21 | Tolerated | | |
| p.Leu78Gly | VUS | 0.00 | Neutral | 0.1888 | Benign | -1.07 | Benign | 0.21 | Neutral | 1.00 | Probably damaging | 0.11 | Tolerated | | |
| p.Leu78Val | VUS | -0.08 | Neutral | 0.1929 | Benign | -9.48 | Pathogenic | 0.34 | Neutral | 0.99 | Probably damaging | 0.3 | Tolerated | 18.18 | Deleterious |
| p.Leu78Tyr | VUS | -3.31 | Neutral | 0.3986 | Ambiguous | -10.11 | Pathogenic | 0.40 | Neutral | 0.99 | Probably damaging | 0.52 | Tolerated | | |
| p.Leu78Cys | VUS | -2.33 | Neutral | 0.5088 | Ambiguous | -5.35 | Benign | 0.41 | Neutral | 1.00 | Probably damaging | 0.07 | Tolerated | | |
| p.Leu78Trp | VUS | -0.17 | Neutral | 0.3248 | Benign | -10.06 | Pathogenic | 0.42 | Neutral | 1.00 | Probably damaging | 0.13 | Tolerated | | |
| p.Leu78Phe | VUS | -0.03 | Neutral | 0.2207 | Benign | -9.28 | Pathogenic | 0.23 | Neutral | 1.00 | Probably damaging | 0.61 | Tolerated | 19.60 | Deleterious |
| p.Thr79Asn | VUS | -0.32 | Neutral | 0.5472 | Ambiguous | -10.60 | Pathogenic | 0.38 | Neutral | 0.93 | Possibly damaging | 0.04 | Deleterious | 25.30 | Deleterious |
| p.Thr79Lys | VUS | -2.10 | Neutral | 0.773 | Pathogenic | -13.86 | Pathogenic | 0.39 | Neutral | 0.85 | Possibly damaging | 0.02 | Deleterious | | |
| p.Thr79Arg | VUS | -12.64 | Indeterminate | 0.5924 | Pathogenic | -10.01 | Pathogenic | 0.41 | Neutral | 0.98 | Probably damaging | 0.02 | Deleterious | | |
| p.Thr79Ser | VUS | -0.66 | Neutral | 0.2456 | Benign | -3.94 | Benign | 0.12 | Neutral | 0.06 | Benign | 0.38 | Tolerated | 20.74 | Deleterious |
| p.Thr79Ile | VUS | -0.34 | Neutral | 0.8117 | Pathogenic | -6.63 | Benign | 0.17 | Neutral | 0.94 | Possibly damaging | 0.02 | Deleterious | 26.40 | Deleterious |
| p.Thr79Met | VUS | -1.24 | Neutral | 0.5079 | Ambiguous | -7.08 | Benign | 0.36 | Neutral | 1.00 | Probably damaging | 0.01 | Deleterious | | |
| p.Thr79His | VUS | -1.61 | Neutral | 0.6363 | Pathogenic | -11.02 | Pathogenic | 0.40 | Neutral | 1.00 | Probably damaging | 0.01 | Deleterious | | |
| p.Thr79Gln | VUS | -0.84 | Neutral | 0.5363 | Ambiguous | -8.94 | Pathogenic | 0.39 | Neutral | 0.95 | Possibly damaging | 0.02 | Deleterious | | |
| p.Thr79Pro | VUS | -53.15 | Deleterious | 0.6573 | Pathogenic | -11.22 | Pathogenic | 0.34 | Neutral | 1.00 | Probably damaging | 0.03 | Deleterious | 24.80 | Deleterious |
| p.Thr79Leu | VUS | -4.80 | Neutral | 0.48 | Ambiguous | -5.73 | Benign | 0.40 | Neutral | 0.85 | Possibly damaging | 0.02 | Deleterious | | |
| p.Thr79Asp | VUS | -3.25 | Neutral | 0.8238 | Pathogenic | -11.56 | Pathogenic | 0.41 | Neutral | 0.95 | Possibly damaging | 0.02 | Deleterious | | |
| p.Thr79Glu | VUS | -1.74 | Neutral | 0.7004 | Pathogenic | -10.32 | Pathogenic | 0.41 | Neutral | 0.81 | Possibly damaging | 0.02 | Deleterious | | |
| p.Thr79Ala | VUS | -1.95 | Neutral | 0.3031 | Benign | -4.35 | Benign | 0.09 | Neutral | 0.01 | Benign | 0.16 | Tolerated | 19.41 | Deleterious |
| p.Thr79Gly | VUS | -2.54 | Neutral | 0.4236 | Ambiguous | -8.45 | Pathogenic | 0.38 | Neutral | 1.00 | Probably damaging | 0.03 | Deleterious | | |
| p.Thr79Val | VUS | -1.05 | Neutral | 0.5979 | Pathogenic | -3.11 | Benign | 0.28 | Neutral | 0.67 | Possibly damaging | 0.03 | Deleterious | | |
| p.Thr79Tyr | VUS | -4.54 | Neutral | 0.8001 | Pathogenic | -9.48 | Pathogenic | 0.42 | Neutral | 1.00 | Probably damaging | 0.01 | Deleterious | | |
| p.Thr79Cys | VUS | -3.24 | Neutral | 0.7324 | Pathogenic | -6.03 | Benign | 0.23 | Neutral | 1.00 | Probably damaging | 0.01 | Deleterious | | |
| p.Thr79Trp | VUS | -5.36 | Neutral | 0.8715 | Pathogenic | -7.97 | Pathogenic | 0.42 | Neutral | 1.00 | Probably damaging | 0 | Deleterious | | |
| p.Thr79Phe | VUS | -1.91 | Neutral | 0.8125 | Pathogenic | -8.81 | Pathogenic | 0.23 | Neutral | 0.99 | Probably damaging | 0.09 | Tolerated | | |
| p.Arg80Asn | VUS | -17.84 | Indeterminate | 0.8009 | Pathogenic | -13.18 | Pathogenic | 0.46 | Neutral | 1.00 | Probably damaging | 0.47 | Tolerated | | |
| p.Arg80Lys | VUS | -4.55 | Neutral | 0.4406 | Ambiguous | -11.68 | Pathogenic | 0.32 | Neutral | 0.99 | Probably damaging | 0.9 | Tolerated | | |
| p.Arg80Thr | VUS | -10.86 | Indeterminate | 0.4117 | Ambiguous | -4.45 | Benign | 0.16 | Neutral | 1.00 | Probably damaging | 0.82 | Tolerated | | |
| p.Arg80Ser | VUS | -3.31 | Neutral | 0.6392 | Pathogenic | -6.99 | Benign | 0.13 | Neutral | 1.00 | Probably damaging | 0.93 | Tolerated | | |
| p.Arg80Ile | VUS | -53.15 | Deleterious | 0.5159 | Ambiguous | -12.71 | Pathogenic | 0.48 | Neutral | 1.00 | Probably damaging | 0.24 | Tolerated | | |
| p.Arg80Met | VUS | -12.61 | Indeterminate | 0.5624 | Ambiguous | -9.13 | Pathogenic | 0.47 | Neutral | 0.99 | Probably damaging | 0.13 | Tolerated | | |
| p.Arg80His | VUS | -7.91 | Indeterminate | 0.2263 | Benign | -7.35 | Benign | 0.15 | Neutral | 1.00 | Probably damaging | 0.18 | Tolerated | | |
| p.Arg80Gln | VUS | -17.50 | Indeterminate | 0.2243 | Benign | -7.28 | Benign | 0.14 | Neutral | 1.00 | Probably damaging | 0.54 | Tolerated | 22.70 | Deleterious |
| p.Arg80Pro | VUS | -53.15 | Deleterious | 0.8603 | Pathogenic | -9.74 | Pathogenic | 0.23 | Neutral | 1.00 | Probably damaging | 0.31 | Tolerated | 24.50 | Deleterious |
| p.Arg80Leu | VUS | -10.61 | Indeterminate | 0.2514 | Benign | -7.89 | Pathogenic | 0.14 | Neutral | 0.87 | Possibly damaging | 0.4 | Tolerated | 22.30 | Deleterious |
| p.Arg80Asp | VUS | -15.11 | Indeterminate | 0.9354 | Pathogenic | -15.84 | Pathogenic | 0.50 | Neutral | 1.00 | Probably damaging | 0.51 | Tolerated | | |
| p.Arg80Glu | VUS | -4.74 | Neutral | 0.6812 | Pathogenic | -13.85 | Pathogenic | 0.47 | Neutral | 1.00 | Probably damaging | 0.81 | Tolerated | | |
| p.Arg80Ala | VUS | -6.72 | Indeterminate | 0.5359 | Ambiguous | -7.56 | Pathogenic | 0.27 | Neutral | 1.00 | Probably damaging | 1 | Tolerated | | |
| p.Arg80Gly | VUS | -7.39 | Indeterminate | 0.5276 | Ambiguous | -9.90 | Pathogenic | 0.42 | Neutral | 1.00 | Probably damaging | 0.43 | Tolerated | 24.50 | Deleterious |
| p.Arg80Val | VUS | -20.05 | Indeterminate | 0.4465 | Ambiguous | -10.29 | Pathogenic | 0.49 | Neutral | 1.00 | Probably damaging | 0.38 | Tolerated | | |
| p.Arg80Tyr | VUS | -9.69 | Indeterminate | 0.4691 | Ambiguous | -6.99 | Benign | 0.42 | Neutral | 0.99 | Probably damaging | 0.16 | Tolerated | | |
| p.Arg80Cys | VUS | -2.00 | Neutral | 0.2644 | Benign | -8.00 | Pathogenic | 0.48 | Neutral | 1.00 | Probably damaging | 0.09 | Tolerated | | |
| p.Arg80Trp | VUS | -9.72 | Indeterminate | 0.2873 | Benign | -9.49 | Pathogenic | 0.46 | Neutral | 1.00 | Probably damaging | 0.04 | Deleterious | | |
| p.Arg80Phe | VUS | -14.98 | Indeterminate | 0.6496 | Pathogenic | -7.79 | Pathogenic | 0.34 | Neutral | 0.96 | Possibly damaging | 0.14 | Tolerated | | |
| p.Pro81Asn | VUS | -53.15 | Deleterious | 0.9894 | Pathogenic | -18.84 | Pathogenic | 0.82 | Disease | 1.00 | Probably damaging | 0.02 | Deleterious | | |
| p.Pro81Lys | VUS | -53.15 | Deleterious | 0.9967 | Pathogenic | -21.45 | Pathogenic | 0.85 | Disease | 1.00 | Probably damaging | 0.03 | Deleterious | | |
| p.Pro81Thr | VUS | -33.22 | Indeterminate | 0.8388 | Pathogenic | -13.29 | Pathogenic | 0.85 | Disease | 1.00 | Probably damaging | 0.07 | Tolerated | 27.80 | Deleterious |
| p.Pro81Arg | VUS | -53.15 | Deleterious | 0.9753 | Pathogenic | -17.08 | Pathogenic | 0.93 | Disease | 1.00 | Probably damaging | 0.02 | Deleterious | 29.10 | Deleterious |
| p.Pro81Ser | VUS | -53.15 | Deleterious | 0.8599 | Pathogenic | -12.55 | Pathogenic | 0.82 | Disease | 1.00 | Probably damaging | 0.07 | Tolerated | 29.00 | Deleterious |
| p.Pro81Ile | VUS | -53.15 | Deleterious | 0.9406 | Pathogenic | -18.39 | Pathogenic | 0.81 | Disease | 1.00 | Probably damaging | 0.07 | Tolerated | | |
| p.Pro81Met | VUS | -53.15 | Deleterious | 0.9665 | Pathogenic | -15.36 | Pathogenic | 0.83 | Disease | 1.00 | Probably damaging | 0.02 | Deleterious | | |
| p.Pro81His | VUS | -53.15 | Deleterious | 0.9778 | Pathogenic | -18.01 | Pathogenic | 0.87 | Disease | 1.00 | Probably damaging | 0.01 | Deleterious | 29.60 | Deleterious |
| p.Pro81Gln | VUS | -53.15 | Deleterious | 0.9612 | Pathogenic | -16.67 | Pathogenic | 0.82 | Disease | 1.00 | Probably damaging | 0.02 | Deleterious | | |
| p.Pro81Leu | VUS | -53.15 | Deleterious | 0.8411 | Pathogenic | -14.01 | Pathogenic | 0.90 | Disease | 1.00 | Probably damaging | 0.07 | Tolerated | 31.00 | Deleterious |
| p.Pro81Asp | VUS | -53.15 | Deleterious | 0.994 | Pathogenic | -19.16 | Pathogenic | 0.85 | Disease | 1.00 | Probably damaging | 0.02 | Deleterious | | |
| p.Pro81Glu | VUS | -53.15 | Deleterious | 0.9834 | Pathogenic | -17.89 | Pathogenic | 0.81 | Disease | 1.00 | Probably damaging | 0.03 | Deleterious | | |
| p.Pro81Ala | VUS | -20.08 | Indeterminate | 0.4524 | Ambiguous | -10.68 | Pathogenic | 0.71 | Disease | 1.00 | Probably damaging | 0.17 | Tolerated | 27.00 | Deleterious |
| p.Pro81Gly | VUS | -53.15 | Deleterious | 0.9254 | Pathogenic | -15.73 | Pathogenic | 0.77 | Disease | 1.00 | Probably damaging | 0.07 | Tolerated | | |
| p.Pro81Val | VUS | -33.22 | Indeterminate | 0.8183 | Pathogenic | -15.02 | Pathogenic | 0.62 | Disease | 1.00 | Probably damaging | 0.31 | Tolerated | | |
| p.Pro81Tyr | VUS | -53.15 | Deleterious | 0.9942 | Pathogenic | -20.20 | Pathogenic | 0.83 | Disease | 1.00 | Probably damaging | 0.01 | Deleterious | | |
| p.Pro81Cys | VUS | -19.66 | Indeterminate | 0.9474 | Pathogenic | -14.43 | Pathogenic | 0.82 | Disease | 1.00 | Probably damaging | 0.02 | Deleterious | | |
| p.Pro81Trp | VUS | -53.15 | Deleterious | 0.996 | Pathogenic | -20.71 | Pathogenic | 0.84 | Disease | 1.00 | Probably damaging | 0 | Deleterious | | |
| p.Pro81Phe | VUS | -53.15 | Deleterious | 0.994 | Pathogenic | -18.68 | Pathogenic | 0.82 | Disease | 1.00 | Probably damaging | 0.02 | Deleterious | | |
| p.Val82Asn | VUS | -53.15 | Deleterious | 0.9718 | Pathogenic | -16.39 | Pathogenic | 0.70 | Disease | 1.00 | Probably damaging | 0 | Deleterious | | |
| p.Val82Lys | VUS | -53.15 | Deleterious | 0.9855 | Pathogenic | -18.37 | Pathogenic | 0.77 | Disease | 0.99 | Probably damaging | 0.01 | Deleterious | | |
| p.Val82Thr | VUS | -8.29 | Indeterminate | 0.7058 | Pathogenic | -6.91 | Benign | 0.50 | Neutral | 1.00 | Probably damaging | 0.03 | Deleterious | | |
| p.Val82Arg | VUS | -53.15 | Deleterious | 0.9597 | Pathogenic | -14.36 | Pathogenic | 0.74 | Disease | 1.00 | Probably damaging | 0 | Deleterious | | |
| p.Val82Ser | VUS | -7.82 | Indeterminate | 0.7613 | Pathogenic | -8.13 | Pathogenic | 0.69 | Disease | 1.00 | Probably damaging | 0.01 | Deleterious | | |
| p.Val82Ile | VUS | -7.73 | Indeterminate | 0.1255 | Benign | -7.88 | Pathogenic | 0.25 | Neutral | 0.77 | Possibly damaging | 0.6 | Tolerated | | |
| p.Val82Met | VUS | -1.59 | Neutral | 0.5783 | Pathogenic | -8.20 | Pathogenic | 0.45 | Neutral | 1.00 | Probably damaging | 0.02 | Deleterious | 27.60 | Deleterious |
| p.Val82His | VUS | -33.22 | Indeterminate | 0.9819 | Pathogenic | -15.18 | Pathogenic | 0.68 | Disease | 1.00 | Probably damaging | 0 | Deleterious | | |
| p.Val82Gln | VUS | -13.11 | Indeterminate | 0.9388 | Pathogenic | -12.30 | Pathogenic | 0.71 | Disease | 1.00 | Probably damaging | 0 | Deleterious | | |
| p.Val82Pro | VUS | -30.46 | Indeterminate | 0.8379 | Pathogenic | -11.82 | Pathogenic | 0.68 | Disease | 1.00 | Probably damaging | 0.01 | Deleterious | | |
| p.Val82Leu | VUS | -5.30 | Neutral | 0.422 | Ambiguous | -6.55 | Benign | 0.22 | Neutral | 0.76 | Possibly damaging | 0.53 | Tolerated | 22.70 | Deleterious |

| | | | | | | | | | | | | | | | |
|---|---|---|---|---|---|---|---|---|---|---|---|---|---|---|---|
| p.Val82Asp | VUS | -53.15 | Deleterious | 0.9883 | Pathogenic | -16.64 | Pathogenic | 0.79 | Disease | 1.00 | Probably damaging | 0 | Deleterious | | |
| p.Val82Glu | VUS | -33.21 | Indeterminate | 0.9406 | Pathogenic | -13.40 | Pathogenic | 0.80 | Disease | 1.00 | Probably damaging | 0.01 | Deleterious | 27.70 | Deleterious |
| p.Val82Ala | VUS | -2.05 | Neutral | 0.3786 | Ambiguous | -1.31 | Benign | 0.32 | Neutral | 0.87 | Possibly damaging | 0.1 | Tolerated | 22.80 | Deleterious |
| p.Val82Gly | VUS | -33.22 | Indeterminate | 0.6991 | Pathogenic | -9.38 | Pathogenic | 0.55 | Disease | 1.00 | Probably damaging | 0 | Deleterious | 26.80 | Deleterious |
| p.Val82Tyr | VUS | -53.15 | Deleterious | 0.9744 | Pathogenic | -15.76 | Pathogenic | 0.67 | Disease | 1.00 | Probably damaging | 0.01 | Deleterious | | |
| p.Val82Cys | VUS | -13.90 | Indeterminate | 0.8291 | Pathogenic | -10.10 | Pathogenic | 0.64 | Disease | 1.00 | Probably damaging | 0.01 | Deleterious | | |
| p.Val82Trp | VUS | -53.15 | Deleterious | 0.9889 | Pathogenic | -14.31 | Pathogenic | 0.64 | Disease | 1.00 | Probably damaging | 0 | Deleterious | | |
| p.Val82Phe | VUS | -21.29 | Indeterminate | 0.7499 | Pathogenic | -13.04 | Pathogenic | 0.64 | Disease | 1.00 | Probably damaging | 0.02 | Deleterious | | |
| p.His83Asn | VUS | -53.15 | Deleterious | 0.7844 | Pathogenic | -16.37 | Pathogenic | 0.91 | Disease | 1.00 | Probably damaging | 0 | Deleterious | 29.40 | Deleterious |
| p.His83Lys | VUS | -53.15 | Deleterious | 0.9871 | Pathogenic | -18.73 | Pathogenic | 0.84 | Disease | 0.97 | Probably damaging | 0 | Deleterious | | |
| p.His83Thr | VUS | -53.15 | Deleterious | 0.96 | Pathogenic | -15.15 | Pathogenic | 0.84 | Disease | 1.00 | Probably damaging | 0 | Deleterious | | |
| p.His83Arg | Pathogenic | -53.15 | Deleterious | 0.9431 | Pathogenic | -14.58 | Pathogenic | 0.90 | Disease | 0.96 | Probably damaging | 0 | Deleterious | 27.10 | Deleterious |
| p.His83Ser | VUS | -53.15 | Deleterious | 0.9106 | Pathogenic | -15.71 | Pathogenic | 0.83 | Disease | 1.00 | Probably damaging | 0 | Deleterious | | |
| p.His83Ile | VUS | -53.15 | Deleterious | 0.9602 | Pathogenic | -14.15 | Pathogenic | 0.87 | Disease | 0.99 | Probably damaging | 0 | Deleterious | | |
| p.His83Met | VUS | -30.49 | Indeterminate | 0.9481 | Pathogenic | -12.73 | Pathogenic | 0.87 | Disease | 1.00 | Probably damaging | 0 | Deleterious | | |
| p.His83Gln | VUS | -53.15 | Deleterious | 0.8794 | Pathogenic | -10.99 | Pathogenic | 0.88 | Disease | 1.00 | Probably damaging | 0 | Deleterious | 25.70 | Deleterious |
| p.His83Pro | VUS | -53.15 | Deleterious | 0.8982 | Pathogenic | -12.25 | Pathogenic | 0.90 | Disease | 1.00 | Probably damaging | 0 | Deleterious | 28.50 | Deleterious |
| p.His83Leu | VUS | -53.15 | Deleterious | 0.8031 | Pathogenic | -12.52 | Pathogenic | 0.90 | Disease | 0.99 | Probably damaging | 0 | Deleterious | 29.40 | Deleterious |
| p.His83Asp | Likely pathogenic | -53.15 | Deleterious | 0.8932 | Pathogenic | -14.15 | Pathogenic | 0.90 | Disease | 1.00 | Probably damaging | 0 | Deleterious | 29.80 | Deleterious |
| p.His83Glu | VUS | -53.15 | Deleterious | 0.937 | Pathogenic | -13.40 | Pathogenic | 0.84 | Disease | 1.00 | Probably damaging | 0 | Deleterious | | |
| p.His83Ala | VUS | -33.22 | Indeterminate | 0.9429 | Pathogenic | -12.39 | Pathogenic | 0.82 | Disease | 1.00 | Probably damaging | 0 | Deleterious | | |
| p.His83Gly | VUS | -53.15 | Deleterious | 0.9562 | Pathogenic | -14.62 | Pathogenic | 0.83 | Disease | 1.00 | Probably damaging | 0 | Deleterious | | |
| p.His83Val | VUS | -53.15 | Deleterious | 0.9547 | Pathogenic | -13.93 | Pathogenic | 0.87 | Disease | 1.00 | Probably damaging | 0 | Deleterious | | |
| p.His83Tyr | Likely pathogenic | -53.15 | Deleterious | 0.7859 | Pathogenic | -14.36 | Pathogenic | 0.90 | Disease | 0.97 | Probably damaging | 0 | Deleterious | 29.40 | Deleterious |
| p.His83Cys | VUS | -53.15 | Deleterious | 0.7785 | Pathogenic | -14.26 | Pathogenic | 0.82 | Disease | 1.00 | Probably damaging | 0 | Deleterious | | |
| p.His83Trp | VUS | -53.15 | Deleterious | 0.8026 | Pathogenic | -16.77 | Pathogenic | 0.86 | Disease | 1.00 | Probably damaging | 0 | Deleterious | | |
| p.His83Phe | VUS | -53.15 | Deleterious | 0.8484 | Pathogenic | -14.90 | Pathogenic | 0.86 | Disease | 0.99 | Probably damaging | 0 | Deleterious | | |
| p.Asp84Asn | Likely pathogenic | -53.15 | Deleterious | 0.8357 | Pathogenic | -11.47 | Pathogenic | 0.71 | Disease | 1.00 | Probably damaging | 0.52 | Tolerated | 32.00 | Deleterious |
| p.Asp84Lys | VUS | -53.15 | Deleterious | 0.9685 | Pathogenic | -16.42 | Pathogenic | 0.78 | Disease | 1.00 | Probably damaging | 0.45 | Tolerated | | |
| p.Asp84Thr | VUS | -53.15 | Deleterious | 0.9381 | Pathogenic | -12.78 | Pathogenic | 0.77 | Disease | 1.00 | Probably damaging | 0.36 | Tolerated | | |
| p.Asp84Arg | VUS | -53.15 | Deleterious | 0.9309 | Pathogenic | -15.98 | Pathogenic | 0.77 | Disease | 1.00 | Probably damaging | 0.34 | Tolerated | | |
| p.Asp84Ser | VUS | -53.15 | Deleterious | 0.8767 | Pathogenic | -11.88 | Pathogenic | 0.69 | Disease | 1.00 | Probably damaging | 0.55 | Tolerated | | |
| p.Asp84Ile | VUS | -53.15 | Deleterious | 0.9329 | Pathogenic | -15.86 | Pathogenic | 0.78 | Disease | 1.00 | Probably damaging | 0.17 | Tolerated | | |
| p.Asp84Met | VUS | -53.15 | Deleterious | 0.9818 | Pathogenic | -15.99 | Pathogenic | 0.77 | Disease | 1.00 | Probably damaging | 0.1 | Tolerated | | |
| p.Asp84His | VUS | -53.15 | Deleterious | 0.8323 | Pathogenic | -12.46 | Pathogenic | 0.84 | Disease | 1.00 | Probably damaging | 0.24 | Tolerated | 32.00 | Deleterious |
| p.Asp84Gln | VUS | -53.15 | Deleterious | 0.9182 | Pathogenic | -14.08 | Pathogenic | 0.67 | Disease | 1.00 | Probably damaging | 0.36 | Tolerated | | |
| p.Asp84Pro | VUS | -53.15 | Deleterious | 0.9957 | Pathogenic | -18.92 | Pathogenic | 0.77 | Disease | 0.99 | Probably damaging | 0.26 | Tolerated | | |
| p.Asp84Leu | VUS | -53.15 | Deleterious | 0.8899 | Pathogenic | -15.56 | Pathogenic | 0.78 | Disease | 1.00 | Probably damaging | 0.29 | Tolerated | | |
| p.Asp84Glu | VUS | -11.57 | Indeterminate | 0.7934 | Pathogenic | -9.49 | Pathogenic | 0.69 | Disease | 0.98 | Probably damaging | 0.78 | Tolerated | 27.30 | Deleterious |
| p.Asp84Ala | VUS | -53.15 | Deleterious | 0.8398 | Pathogenic | -11.92 | Pathogenic | 0.83 | Disease | 1.00 | Probably damaging | 0.73 | Tolerated | 31.00 | Deleterious |
| p.Asp84Gly | VUS | -53.15 | Deleterious | 0.9245 | Pathogenic | -12.25 | Pathogenic | 0.82 | Disease | 1.00 | Probably damaging | 0.5 | Tolerated | 32.00 | Deleterious |
| p.Asp84Val | VUS | -53.15 | Deleterious | 0.8435 | Pathogenic | -13.70 | Pathogenic | 0.84 | Disease | 1.00 | Probably damaging | 0.26 | Tolerated | 31.00 | Deleterious |
| p.Asp84Tyr | VUS | -53.15 | Deleterious | 0.7537 | Pathogenic | -13.75 | Pathogenic | 0.84 | Disease | 1.00 | Probably damaging | 0.31 | Tolerated | 32.00 | Deleterious |
| p.Asp84Cys | VUS | -53.15 | Deleterious | 0.9561 | Pathogenic | -14.29 | Pathogenic | 0.78 | Disease | 1.00 | Probably damaging | 0.09 | Tolerated | | |
| p.Asp84Trp | VUS | -53.15 | Deleterious | 0.9764 | Pathogenic | -15.31 | Pathogenic | 0.78 | Disease | 1.00 | Probably damaging | 0.06 | Tolerated | | |
| p.Asp84Phe | VUS | -53.15 | Deleterious | 0.9455 | Pathogenic | -14.89 | Pathogenic | 0.79 | Disease | 1.00 | Probably damaging | 0.28 | Tolerated | | |
| p.Ala85Asn | VUS | -18.26 | Indeterminate | 0.9879 | Pathogenic | -15.33 | Pathogenic | 0.79 | Disease | 1.00 | Probably damaging | 0 | Deleterious | | |
| p.Ala85Lys | VUS | -33.22 | Indeterminate | 0.9973 | Pathogenic | -17.93 | Pathogenic | 0.83 | Disease | 1.00 | Probably damaging | 0 | Deleterious | | |
| p.Ala85Thr | VUS | -17.82 | Indeterminate | 0.6836 | Pathogenic | -7.99 | Pathogenic | 0.83 | Disease | 1.00 | Probably damaging | 0 | Deleterious | 32.00 | Deleterious |
| p.Ala85Arg | VUS | -33.22 | Indeterminate | 0.978 | Pathogenic | -14.78 | Pathogenic | 0.82 | Disease | 1.00 | Probably damaging | 0 | Deleterious | | |
| p.Ala85Ser | VUS | -3.75 | Neutral | 0.4337 | Ambiguous | -9.74 | Pathogenic | 0.70 | Disease | 0.99 | Probably damaging | 0 | Deleterious | 31.00 | Deleterious |
| p.Ala85Ile | VUS | -14.32 | Indeterminate | 0.9512 | Pathogenic | -15.31 | Pathogenic | 0.83 | Disease | 1.00 | Probably damaging | 0 | Deleterious | | |
| p.Ala85Met | VUS | -1.40 | Neutral | 0.9375 | Pathogenic | -12.33 | Pathogenic | 0.78 | Disease | 1.00 | Probably damaging | 0 | Deleterious | | |
| p.Ala85His | VUS | -27.36 | Indeterminate | 0.9894 | Pathogenic | -14.46 | Pathogenic | 0.83 | Disease | 1.00 | Probably damaging | 0 | Deleterious | | |
| p.Ala85Gln | VUS | -17.67 | Indeterminate | 0.9782 | Pathogenic | -10.38 | Pathogenic | 0.83 | Disease | 1.00 | Probably damaging | 0 | Deleterious | | |
| p.Ala85Pro | VUS | -13.79 | Indeterminate | 0.9715 | Pathogenic | -13.26 | Pathogenic | 0.86 | Disease | 1.00 | Probably damaging | 0 | Deleterious | 32.00 | Deleterious |
| p.Ala85Leu | VUS | -2.77 | Neutral | 0.8177 | Pathogenic | -11.90 | Pathogenic | 0.81 | Disease | 1.00 | Probably damaging | 0 | Deleterious | | |
| p.Ala85Asp | VUS | -33.22 | Indeterminate | 0.9893 | Pathogenic | -18.56 | Pathogenic | 0.87 | Disease | 1.00 | Probably damaging | 0 | Deleterious | 32.00 | Deleterious |
| p.Ala85Glu | VUS | -33.20 | Indeterminate | 0.986 | Pathogenic | -16.15 | Pathogenic | 0.87 | Disease | 1.00 | Probably damaging | 0 | Deleterious | | |
| p.Ala85Gly | VUS | -0.46 | Neutral | 0.5102 | Ambiguous | -10.76 | Pathogenic | 0.76 | Disease | 1.00 | Probably damaging | 0 | Deleterious | 31.00 | Deleterious |
| p.Ala85Val | VUS | -9.08 | Indeterminate | 0.7195 | Pathogenic | -9.69 | Pathogenic | 0.64 | Disease | 1.00 | Probably damaging | 0 | Deleterious | 27.20 | Deleterious |
| p.Ala85Tyr | VUS | -33.10 | Indeterminate | 0.9938 | Pathogenic | -16.68 | Pathogenic | 0.83 | Disease | 1.00 | Probably damaging | 0 | Deleterious | | |
| p.Ala85Cys | VUS | -1.78 | Neutral | 0.8193 | Pathogenic | -9.92 | Pathogenic | 0.76 | Disease | 1.00 | Probably damaging | 0 | Deleterious | | |
| p.Ala85Trp | VUS | -53.15 | Deleterious | 0.9971 | Pathogenic | -15.39 | Pathogenic | 0.85 | Disease | 1.00 | Probably damaging | 0 | Deleterious | | |
| p.Ala85Phe | VUS | -30.24 | Indeterminate | 0.9849 | Pathogenic | -15.98 | Pathogenic | 0.83 | Disease | 1.00 | Probably damaging | 0 | Deleterious | | |
| p.Ala86Asn | VUS | -53.15 | Deleterious | 0.9923 | Pathogenic | -21.15 | Pathogenic | 0.82 | Disease | 0.98 | Probably damaging | 0 | Deleterious | | |
| p.Ala86Lys | VUS | -53.15 | Deleterious | 0.9984 | Pathogenic | -22.11 | Pathogenic | 0.86 | Disease | 1.00 | Probably damaging | 0 | Deleterious | | |
| p.Ala86Thr | VUS | -0.06 | Neutral | 0.4991 | Ambiguous | -8.25 | Pathogenic | 0.87 | Disease | 0.96 | Probably damaging | 0.01 | Deleterious | 32.00 | Deleterious |
| p.Ala86Arg | VUS | -53.15 | Deleterious | 0.9881 | Pathogenic | -17.11 | Pathogenic | 0.82 | Disease | 1.00 | Probably damaging | 0 | Deleterious | | |
| p.Ala86Ser | VUS | -2.71 | Neutral | 0.281 | Benign | -7.75 | Pathogenic | 0.75 | Disease | 0.17 | Benign | 0.12 | Tolerated | 29.60 | Deleterious |
| p.Ala86Ile | VUS | -5.80 | Indeterminate | 0.8519 | Pathogenic | -14.37 | Pathogenic | 0.85 | Disease | 1.00 | Probably damaging | 0 | Deleterious | | |
| p.Ala86Met | VUS | -53.15 | Deleterious | 0.906 | Pathogenic | -15.48 | Pathogenic | 0.80 | Disease | 1.00 | Probably damaging | 0 | Deleterious | | |
| p.Ala86His | VUS | -53.15 | Deleterious | 0.9941 | Pathogenic | -21.22 | Pathogenic | 0.85 | Disease | 1.00 | Probably damaging | 0 | Deleterious | | |
| p.Ala86Gln | VUS | -53.15 | Deleterious | 0.9798 | Pathogenic | -20.63 | Pathogenic | 0.85 | Disease | 1.00 | Probably damaging | 0 | Deleterious | | |
| p.Ala86Pro | VUS | -53.15 | Deleterious | 0.9789 | Pathogenic | -15.35 | Pathogenic | 0.92 | Disease | 1.00 | Probably damaging | 0.01 | Deleterious | 32.00 | Deleterious |
| p.Ala86Leu | VUS | -53.15 | Deleterious | 0.7887 | Pathogenic | -13.86 | Pathogenic | 0.84 | Disease | 1.00 | Probably damaging | 0 | Deleterious | | |
| p.Ala86Asp | VUS | -53.15 | Deleterious | 0.9955 | Pathogenic | -18.42 | Pathogenic | 0.92 | Disease | 1.00 | Probably damaging | 0 | Deleterious | 32.00 | Deleterious |
| p.Ala86Glu | VUS | -53.15 | Deleterious | 0.9873 | Pathogenic | -18.10 | Pathogenic | 0.92 | Disease | 1.00 | Probably damaging | 0 | Deleterious | | |
| p.Ala86Gly | VUS | -0.02 | Neutral | 0.5397 | Ambiguous | -10.00 | Pathogenic | 0.72 | Disease | 0.99 | Probably damaging | 0.03 | Deleterious | 31.00 | Deleterious |
| p.Ala86Val | VUS | -0.10 | Neutral | 0.4564 | Ambiguous | -8.22 | Pathogenic | 0.84 | Disease | 1.00 | Probably damaging | 0.01 | Deleterious | 32.00 | Deleterious |
| p.Ala86Tyr | VUS | -53.15 | Deleterious | 0.9965 | Pathogenic | -20.61 | Pathogenic | 0.85 | Disease | 1.00 | Probably damaging | 0 | Deleterious | | |
| p.Ala86Cys | VUS | 0.00 | Neutral | 0.716 | Pathogenic | -9.01 | Pathogenic | 0.77 | Disease | 1.00 | Probably damaging | 0 | Deleterious | | |
| p.Ala86Trp | VUS | -53.15 | Deleterious | 0.998 | Pathogenic | -18.82 | Pathogenic | 0.86 | Disease | 1.00 | Probably damaging | 0 | Deleterious | | |
| p.Ala86Phe | VUS | -53.15 | Deleterious | 0.9859 | Pathogenic | -17.77 | Pathogenic | 0.87 | Disease | 1.00 | Probably damaging | 0 | Deleterious | | |
| p.Arg87Asn | VUS | -3.51 | Neutral | 0.972 | Pathogenic | -12.31 | Pathogenic | 0.66 | Disease | 1.00 | Probably damaging | 0.12 | Tolerated | | |
| p.Arg87Lys | VUS | 0.00 | Neutral | 0.583 | Pathogenic | -7.99 | Pathogenic | 0.58 | Disease | 0.99 | Probably damaging | 0.17 | Tolerated | | |
| p.Arg87Thr | VUS | -7.09 | Indeterminate | 0.9022 | Pathogenic | -9.79 | Pathogenic | 0.74 | Disease | 1.00 | Probably damaging | 0.88 | Tolerated | | |
| p.Arg87Ser | VUS | -53.15 | Deleterious | 0.9346 | Pathogenic | -6.62 | Benign | 0.73 | Disease | 1.00 | Probably damaging | 0.53 | Tolerated | | |
| p.Arg87Ile | VUS | -9.73 | Indeterminate | 0.8709 | Pathogenic | -11.77 | Pathogenic | 0.71 | Disease | 1.00 | Probably damaging | 0.04 | Deleterious | | |
| p.Arg87Met | VUS | -19.92 | Indeterminate | 0.9082 | Pathogenic | -11.21 | Pathogenic | 0.68 | Disease | 1.00 | Probably damaging | 0.02 | Deleterious | | |
| p.Arg87His | VUS | -33.21 | Indeterminate | 0.6262 | Pathogenic | -11.02 | Pathogenic | 0.64 | Disease | 1.00 | Probably damaging | 0.03 | Deleterious | | |
| p.Arg87Gln | VUS | -31.79 | Indeterminate | 0.4766 | Ambiguous | -9.61 | Pathogenic | 0.62 | Disease | 1.00 | Probably damaging | 0.08 | Tolerated | 32.00 | Deleterious |
| p.Arg87Pro | Pathogenic | -53.15 | Deleterious | 0.9811 | Pathogenic | -17.26 | Pathogenic | 0.81 | Disease | 1.00 | Probably damaging | 0.08 | Tolerated | 32.00 | Deleterious |
| p.Arg87Leu | VUS | -15.67 | Indeterminate | 0.7068 | Pathogenic | -12.08 | Pathogenic | 0.77 | Disease | 1.00 | Probably damaging | 0.05 | Deleterious | 32.00 | Deleterious |
| p.Arg87Asp | VUS | -53.15 | Deleterious | 0.9683 | Pathogenic | -16.23 | Pathogenic | 0.69 | Disease | 1.00 | Probably damaging | 0.07 | Tolerated | | |
| p.Arg87Glu | VUS | -53.15 | Deleterious | 0.8133 | Pathogenic | -13.22 | Pathogenic | 0.65 | Disease | 1.00 | Probably damaging | 0.06 | Tolerated | | |
| p.Arg87Ala | VUS | -33.22 | Indeterminate | 0.8485 | Pathogenic | -7.66 | Pathogenic | 0.60 | Disease | 1.00 | Probably damaging | 0.16 | Tolerated | | |
| p.Arg87Gly | VUS | -1.07 | Neutral | 0.8043 | Pathogenic | -8.75 | Pathogenic | 0.76 | Disease | 1.00 | Probably damaging | 0.07 | Tolerated | 27.80 | Deleterious |
| p.Arg87Val | VUS | -6.11 | Indeterminate | 0.8731 | Pathogenic | -11.39 | Pathogenic | 0.70 | Disease | 1.00 | Probably damaging | 0.07 | Tolerated | | |
| p.Arg87Tyr | VUS | -53.15 | Deleterious | 0.8984 | Pathogenic | -14.41 | Pathogenic | 0.66 | Disease | 0.98 | Probably damaging | 0.02 | Deleterious | | |
| p.Arg87Cys | VUS | -0.11 | Neutral | 0.7911 | Pathogenic | -7.86 | Pathogenic | 0.76 | Disease | 1.00 | Probably damaging | 0.03 | Deleterious | | |
| p.Arg87Trp | Likely pathogenic | -53.15 | Deleterious | 0.6316 | Pathogenic | -12.63 | Pathogenic | 0.58 | Disease | 0.40 | Benign | 0.01 | Deleterious | 31.00 | Deleterious |
| p.Arg87Phe | VUS | -6.79 | Indeterminate | 0.9377 | Pathogenic | -14.69 | Pathogenic | 0.68 | Disease | 0.98 | Probably damaging | 0.02 | Deleterious | | |
| p.Glu88Asn | VUS | -8.96 | Indeterminate | 0.7893 | Pathogenic | -10.27 | Pathogenic | 0.44 | Neutral | 1.00 | Probably damaging | 0.63 | Tolerated | | |
| p.Glu88Lys | VUS | -26.55 | Indeterminate | 0.7692 | Pathogenic | -11.55 | Pathogenic | 0.46 | Neutral | 0.98 | Probably damaging | 0.84 | Tolerated | 29.00 | Deleterious |
| p.Glu88Thr | VUS | -13.66 | Indeterminate | 0.8562 | Pathogenic | -7.38 | Benign | 0.26 | Neutral | 0.99 | Probably damaging | 0.93 | Tolerated | | |
| p.Glu88Arg | VUS | -23.65 | Indeterminate | 0.5865 | Pathogenic | -11.02 | Pathogenic | 0.46 | Neutral | 1.00 | Probably damaging | 0.49 | Tolerated | | |
| p.Glu88Ser | VUS | -8.02 | Indeterminate | 0.6826 | Pathogenic | -8.33 | Pathogenic | 0.29 | Neutral | 0.99 | Probably damaging | 1 | Tolerated | | |
| p.Glu88Ile | VUS | -5.97 | Indeterminate | 0.9304 | Pathogenic | -11.79 | Pathogenic | 0.48 | Neutral | 0.99 | Probably damaging | 0.23 | Tolerated | | |
| p.Glu88Met | VUS | -4.02 | Neutral | 0.8959 | Pathogenic | -10.44 | Pathogenic | 0.53 | Disease | 1.00 | Probably damaging | 0.12 | Tolerated | | |
| p.Glu88His | VUS | -13.18 | Indeterminate | 0.7291 | Pathogenic | -10.46 | Pathogenic | 0.43 | Neutral | 1.00 | Probably damaging | 0.16 | Tolerated | | |

| | | | | | | | | | | | | | | | |
|---|---|---|---|---|---|---|---|---|---|---|---|---|---|---|---|
| p.Glu88Gln | VUS | -9.17 | Indeterminate | 0.3834 | Ambiguous | -8.59 | Pathogenic | 0.36 | Neutral | 0.99 | Probably damaging | 0.55 | Tolerated | 26.90 | Deleterious |
| p.Glu88Pro | VUS | -53.15 | Deleterious | 0.9891 | Pathogenic | -14.03 | Pathogenic | 0.55 | Disease | 1.00 | Probably damaging | 0.33 | Tolerated | | |
| p.Glu88Leu | VUS | -11.28 | Indeterminate | 0.8529 | Pathogenic | -9.54 | Pathogenic | 0.49 | Neutral | 0.94 | Possibly damaging | 0.33 | Tolerated | | |
| p.Glu88Asp | VUS | -8.89 | Indeterminate | 0.7631 | Pathogenic | -8.59 | Pathogenic | 0.26 | Neutral | 0.98 | Probably damaging | 0.68 | Tolerated | 21.45 | Deleterious |
| p.Glu88Ala | VUS | -4.75 | Neutral | 0.5098 | Ambiguous | -5.30 | Benign | 0.18 | Neutral | 0.50 | Possibly damaging | 0.9 | Tolerated | 22.70 | Deleterious |
| p.Glu88Gly | VUS | -5.36 | Neutral | 0.5315 | Ambiguous | -8.43 | Pathogenic | 0.22 | Neutral | 1.00 | Probably damaging | 0.48 | Tolerated | 23.40 | Deleterious |
| p.Glu88Val | VUS | -3.02 | Neutral | 0.8221 | Pathogenic | -9.90 | Pathogenic | 0.44 | Neutral | 0.89 | Possibly damaging | 0.38 | Tolerated | 26.90 | Deleterious |
| p.Glu88Tyr | VUS | -7.59 | Indeterminate | 0.8342 | Pathogenic | -12.66 | Pathogenic | 0.50 | Neutral | 1.00 | Probably damaging | 0.11 | Tolerated | | |
| p.Glu88Cys | VUS | -4.88 | Neutral | 0.8882 | Pathogenic | -10.65 | Pathogenic | 0.53 | Disease | 1.00 | Probably damaging | 0.1 | Tolerated | | |
| p.Glu88Trp | VUS | -4.25 | Neutral | 0.9523 | Pathogenic | -14.19 | Pathogenic | 0.57 | Disease | 1.00 | Probably damaging | 0.03 | Deleterious | | |
| p.Glu88Phe | VUS | -10.21 | Indeterminate | 0.9337 | Pathogenic | -12.78 | Pathogenic | 0.50 | Neutral | 1.00 | Probably damaging | 0.09 | Tolerated | | |
| p.Gly89Asn | VUS | -16.97 | Indeterminate | 0.9434 | Pathogenic | -10.52 | Pathogenic | 0.81 | Disease | 1.00 | Probably damaging | 0 | Deleterious | | |
| p.Gly89Lys | VUS | -53.15 | Deleterious | 0.9852 | Pathogenic | -17.07 | Pathogenic | 0.84 | Disease | 1.00 | Probably damaging | 0 | Deleterious | | |
| p.Gly89Thr | VUS | -53.15 | Deleterious | 0.9692 | Pathogenic | -18.41 | Pathogenic | 0.82 | Disease | 1.00 | Probably damaging | 0 | Deleterious | | |
| p.Gly89Arg | VUS | -53.15 | Deleterious | 0.9629 | Pathogenic | -15.21 | Pathogenic | 0.87 | Disease | 1.00 | Probably damaging | 0 | Deleterious | 32.00 | Deleterious |
| p.Gly89Ser | VUS | -53.15 | Deleterious | 0.8246 | Pathogenic | -13.01 | Pathogenic | 0.84 | Disease | 1.00 | Probably damaging | 0 | Deleterious | 32.00 | Deleterious |
| p.Gly89Ile | VUS | -53.15 | Deleterious | 0.9909 | Pathogenic | -21.63 | Pathogenic | 0.86 | Disease | 1.00 | Probably damaging | 0 | Deleterious | | |
| p.Gly89Met | VUS | -53.15 | Deleterious | 0.9948 | Pathogenic | -19.70 | Pathogenic | 0.87 | Disease | 1.00 | Probably damaging | 0 | Deleterious | | |
| p.Gly89His | VUS | -53.15 | Deleterious | 0.9876 | Pathogenic | -16.30 | Pathogenic | 0.84 | Disease | 1.00 | Probably damaging | 0 | Deleterious | | |
| p.Gly89Gln | VUS | -53.15 | Deleterious | 0.9814 | Pathogenic | -17.42 | Pathogenic | 0.82 | Disease | 1.00 | Probably damaging | 0 | Deleterious | | |
| p.Gly89Pro | VUS | -53.15 | Deleterious | 0.998 | Pathogenic | -17.63 | Pathogenic | 0.83 | Disease | 1.00 | Probably damaging | 0 | Deleterious | | |
| p.Gly89Leu | VUS | -53.15 | Deleterious | 0.991 | Pathogenic | -18.75 | Pathogenic | 0.85 | Disease | 1.00 | Probably damaging | 0 | Deleterious | | |
| p.Gly89Asp | VUS | -53.15 | Deleterious | 0.9296 | Pathogenic | -10.89 | Pathogenic | 0.86 | Disease | 1.00 | Probably damaging | 0 | Deleterious | 32.00 | Deleterious |
| p.Gly89Glu | VUS | -53.15 | Deleterious | 0.9738 | Pathogenic | -15.05 | Pathogenic | 0.85 | Disease | 1.00 | Probably damaging | 0 | Deleterious | | |
| p.Gly89Ala | VUS | -23.61 | Indeterminate | 0.8544 | Pathogenic | -12.94 | Pathogenic | 0.87 | Disease | 1.00 | Probably damaging | 0 | Deleterious | 29.30 | Deleterious |
| p.Gly89Val | VUS | -53.15 | Deleterious | 0.981 | Pathogenic | -17.26 | Pathogenic | 0.91 | Disease | 1.00 | Probably damaging | 0 | Deleterious | 31.00 | Deleterious |
| p.Gly89Tyr | VUS | -53.15 | Deleterious | 0.9872 | Pathogenic | -18.70 | Pathogenic | 0.85 | Disease | 1.00 | Probably damaging | 0 | Deleterious | | |
| p.Gly89Cys | VUS | -53.15 | Deleterious | 0.9645 | Pathogenic | -14.56 | Pathogenic | 0.91 | Disease | 1.00 | Probably damaging | 0 | Deleterious | 32.00 | Deleterious |
| p.Gly89Trp | VUS | -53.15 | Deleterious | 0.9907 | Pathogenic | -17.79 | Pathogenic | 0.89 | Disease | 1.00 | Probably damaging | 0 | Deleterious | | |
| p.Gly89Phe | VUS | -53.15 | Deleterious | 0.9964 | Pathogenic | -19.66 | Pathogenic | 0.87 | Disease | 1.00 | Probably damaging | 0 | Deleterious | | |
| p.Phe90Asn | VUS | -0.08 | Neutral | 0.8549 | Pathogenic | -9.53 | Pathogenic | 0.83 | Disease | 1.00 | Probably damaging | 0.15 | Tolerated | | |
| p.Phe90Lys | VUS | -2.63 | Neutral | 0.9421 | Pathogenic | -5.70 | Benign | 0.79 | Disease | 1.00 | Probably damaging | 0.13 | Tolerated | | |
| p.Phe90Thr | VUS | -3.69 | Neutral | 0.9749 | Pathogenic | -11.43 | Pathogenic | 0.83 | Disease | 1.00 | Probably damaging | 0.19 | Tolerated | | |
| p.Phe90Arg | VUS | -0.31 | Neutral | 0.8195 | Pathogenic | -6.11 | Benign | 0.83 | Disease | 1.00 | Probably damaging | 0.14 | Tolerated | | |
| p.Phe90Ser | VUS | -0.47 | Neutral | 0.8915 | Pathogenic | -8.46 | Pathogenic | 0.87 | Disease | 1.00 | Probably damaging | 0.33 | Tolerated | 32.00 | Deleterious |
| p.Phe90Ile | VUS | -0.42 | Neutral | 0.9075 | Pathogenic | -9.55 | Pathogenic | 0.77 | Disease | 1.00 | Probably damaging | 0.15 | Tolerated | 32.00 | Deleterious |
| p.Phe90Met | VUS | -0.09 | Neutral | 0.9148 | Pathogenic | -6.68 | Benign | 0.76 | Disease | 1.00 | Probably damaging | 0.09 | Tolerated | | |
| p.Phe90His | VUS | -0.17 | Neutral | 0.6115 | Pathogenic | -6.31 | Benign | 0.85 | Disease | 1.00 | Probably damaging | 0.36 | Tolerated | | |
| p.Phe90Gln | VUS | -0.06 | Neutral | 0.8916 | Pathogenic | -7.91 | Pathogenic | 0.81 | Disease | 1.00 | Probably damaging | 0.12 | Tolerated | | |
| p.Phe90Pro | VUS | -8.66 | Indeterminate | 0.9992 | Pathogenic | -15.49 | Pathogenic | 0.83 | Disease | 1.00 | Probably damaging | 0.1 | Tolerated | | |
| p.Phe90Leu | VUS | -0.06 | Neutral | 0.9825 | Pathogenic | -4.19 | Benign | 0.80 | Disease | 1.00 | Probably damaging | 0.25 | Tolerated | 29.17 | Deleterious |
| p.Phe90Asp | VUS | -17.51 | Indeterminate | 0.9453 | Pathogenic | -13.21 | Pathogenic | 0.77 | Disease | 1.00 | Probably damaging | 0.11 | Tolerated | | |
| p.Phe90Glu | VUS | -1.91 | Neutral | 0.963 | Pathogenic | -12.39 | Pathogenic | 0.82 | Disease | 1.00 | Probably damaging | 0.12 | Tolerated | | |
| p.Phe90Ala | VUS | -0.40 | Neutral | 0.9492 | Pathogenic | -8.68 | Pathogenic | 0.81 | Disease | 1.00 | Probably damaging | 0.43 | Tolerated | | |
| p.Phe90Gly | VUS | -0.76 | Neutral | 0.9762 | Pathogenic | -11.21 | Pathogenic | 0.82 | Disease | 1.00 | Probably damaging | 0.19 | Tolerated | | |
| p.Phe90Val | VUS | -1.40 | Neutral | 0.8919 | Pathogenic | -9.16 | Pathogenic | 0.85 | Disease | 1.00 | Probably damaging | 0.2 | Tolerated | 31.00 | Deleterious |
| p.Phe90Tyr | VUS | -0.05 | Neutral | 0.1801 | Benign | -7.16 | Benign | 0.72 | Disease | 0.99 | Probably damaging | 0.36 | Tolerated | 32.00 | Deleterious |
| p.Phe90Cys | VUS | -0.46 | Neutral | 0.7307 | Pathogenic | -7.78 | Pathogenic | 0.88 | Disease | 1.00 | Probably damaging | 0.07 | Tolerated | 32.00 | Deleterious |
| p.Phe90Trp | VUS | -0.34 | Neutral | 0.6305 | Pathogenic | -9.88 | Pathogenic | 0.71 | Disease | 1.00 | Probably damaging | 0.07 | Tolerated | | |
| p.Leu91Asn | VUS | -0.02 | Neutral | 0.8419 | Pathogenic | -17.48 | Pathogenic | 0.70 | Disease | 1.00 | Probably damaging | 0 | Deleterious | | |
| p.Leu91Lys | VUS | -7.34 | Indeterminate | 0.5184 | Ambiguous | -13.81 | Pathogenic | 0.69 | Disease | 0.99 | Probably damaging | 0 | Deleterious | | |
| p.Leu91Thr | VUS | -8.95 | Indeterminate | 0.3834 | Ambiguous | -11.50 | Pathogenic | 0.66 | Disease | 1.00 | Probably damaging | 0.01 | Deleterious | | |
| p.Leu91Arg | VUS | -1.49 | Neutral | 0.2806 | Benign | -11.67 | Pathogenic | 0.83 | Disease | 1.00 | Probably damaging | 0 | Deleterious | 32.00 | Deleterious |
| p.Leu91Ser | VUS | -18.24 | Indeterminate | 0.6895 | Pathogenic | -13.58 | Pathogenic | 0.72 | Disease | 1.00 | Probably damaging | 0 | Deleterious | | |
| p.Leu91Ile | VUS | -19.53 | Indeterminate | 0.2114 | Benign | -10.91 | Pathogenic | 0.47 | Neutral | 0.57 | Possibly damaging | 0.24 | Tolerated | | |
| p.Leu91Met | VUS | -33.05 | Indeterminate | 0.3357 | Benign | -8.49 | Pathogenic | 0.46 | Neutral | 1.00 | Probably damaging | 0.02 | Deleterious | 23.90 | Deleterious |
| p.Leu91His | VUS | -16.61 | Indeterminate | 0.5598 | Ambiguous | -13.10 | Pathogenic | 0.72 | Disease | 1.00 | Probably damaging | 0 | Deleterious | | |
| p.Leu91Gln | VUS | -0.16 | Neutral | 0.4213 | Ambiguous | -11.72 | Pathogenic | 0.52 | Disease | 1.00 | Probably damaging | 0 | Deleterious | 31.00 | Deleterious |
| p.Leu91Pro | VUS | -7.49 | Indeterminate | 0.4079 | Ambiguous | -11.70 | Pathogenic | 0.84 | Disease | 1.00 | Probably damaging | 0 | Deleterious | 32.00 | Deleterious |
| p.Leu91Asp | VUS | -6.28 | Indeterminate | 0.9074 | Pathogenic | -15.92 | Pathogenic | 0.69 | Disease | 1.00 | Probably damaging | 0 | Deleterious | | |
| p.Leu91Glu | VUS | -4.76 | Neutral | 0.6245 | Pathogenic | -13.29 | Pathogenic | 0.68 | Disease | 1.00 | Probably damaging | 0 | Deleterious | | |
| p.Leu91Ala | VUS | -1.03 | Neutral | 0.4238 | Ambiguous | -11.83 | Pathogenic | 0.61 | Disease | 1.00 | Probably damaging | 0.01 | Deleterious | | |
| p.Leu91Gly | VUS | -4.74 | Neutral | 0.6454 | Pathogenic | -14.23 | Pathogenic | 0.68 | Disease | 1.00 | Probably damaging | 0 | Deleterious | | |
| p.Leu91Val | VUS | -2.65 | Neutral | 0.1689 | Benign | -8.72 | Pathogenic | 0.39 | Neutral | 0.89 | Possibly damaging | 0.18 | Tolerated | 23.40 | Deleterious |
| p.Leu91Tyr | VUS | -53.15 | Deleterious | 0.7318 | Pathogenic | -14.27 | Pathogenic | 0.60 | Disease | 1.00 | Probably damaging | 0 | Deleterious | | |
| p.Leu91Cys | VUS | -0.69 | Neutral | 0.749 | Pathogenic | -13.62 | Pathogenic | 0.61 | Disease | 1.00 | Probably damaging | 0 | Deleterious | | |
| p.Leu91Trp | VUS | -8.02 | Indeterminate | 0.5174 | Ambiguous | -13.64 | Pathogenic | 0.65 | Disease | 1.00 | Probably damaging | 0 | Deleterious | | |
| p.Leu91Phe | VUS | -21.09 | Indeterminate | 0.4145 | Ambiguous | -10.97 | Pathogenic | 0.61 | Disease | 1.00 | Probably damaging | 0.02 | Deleterious | | |
| p.Asp92Asn | VUS | -3.34 | Neutral | 0.4396 | Ambiguous | -10.26 | Pathogenic | 0.60 | Disease | 0.99 | Probably damaging | 0.35 | Tolerated | 32.00 | Deleterious |
| p.Asp92Lys | VUS | -7.05 | Indeterminate | 0.7779 | Pathogenic | -13.34 | Pathogenic | 0.66 | Disease | 1.00 | Probably damaging | 0.05 | Deleterious | | |
| p.Asp92Thr | VUS | -1.00 | Neutral | 0.6917 | Pathogenic | -11.07 | Pathogenic | 0.65 | Disease | 1.00 | Probably damaging | 0.04 | Deleterious | | |
| p.Asp92Arg | VUS | -1.01 | Neutral | 0.643 | Pathogenic | -11.51 | Pathogenic | 0.64 | Disease | 1.00 | Probably damaging | 0.02 | Deleterious | | |
| p.Asp92Ser | VUS | -3.99 | Neutral | 0.4136 | Ambiguous | -8.28 | Pathogenic | 0.58 | Disease | 1.00 | Probably damaging | 0.08 | Tolerated | | |
| p.Asp92Ile | VUS | -2.32 | Neutral | 0.9298 | Pathogenic | -16.37 | Pathogenic | 0.64 | Disease | 1.00 | Probably damaging | 0 | Deleterious | | |
| p.Asp92Met | VUS | -0.61 | Neutral | 0.9641 | Pathogenic | -14.36 | Pathogenic | 0.68 | Disease | 1.00 | Probably damaging | 0 | Deleterious | | |
| p.Asp92His | VUS | -4.28 | Neutral | 0.5663 | Pathogenic | -11.15 | Pathogenic | 0.66 | Disease | 1.00 | Probably damaging | 0.03 | Deleterious | 32.00 | Deleterious |
| p.Asp92Gln | VUS | -0.89 | Neutral | 0.5609 | Ambiguous | -10.85 | Pathogenic | 0.59 | Disease | 1.00 | Probably damaging | 0.03 | Deleterious | | |
| p.Asp92Pro | VUS | -1.59 | Neutral | 0.4936 | Ambiguous | -10.20 | Pathogenic | 0.66 | Disease | 0.99 | Probably damaging | 0.03 | Deleterious | | |
| p.Asp92Leu | VUS | -0.53 | Neutral | 0.8186 | Pathogenic | -12.76 | Pathogenic | 0.68 | Disease | 1.00 | Probably damaging | 0.01 | Deleterious | | |
| p.Asp92Glu | VUS | -3.93 | Neutral | 0.3017 | Benign | -6.36 | Benign | 0.19 | Neutral | 0.93 | Possibly damaging | 0.07 | Tolerated | 22.80 | Deleterious |
| p.Asp92Ala | VUS | -0.99 | Neutral | 0.3843 | Ambiguous | -7.59 | Pathogenic | 0.39 | Neutral | 1.00 | Probably damaging | 0.16 | Tolerated | 28.20 | Deleterious |
| p.Asp92Gly | VUS | -5.17 | Neutral | 0.3826 | Ambiguous | -6.77 | Benign | 0.40 | Neutral | 1.00 | Probably damaging | 0.1 | Tolerated | 31.00 | Deleterious |
| p.Asp92Val | VUS | -0.82 | Neutral | 0.7553 | Pathogenic | -12.77 | Pathogenic | 0.72 | Disease | 1.00 | Probably damaging | 0.01 | Deleterious | 30.00 | Deleterious |
| p.Asp92Tyr | VUS | -4.06 | Neutral | 0.7255 | Pathogenic | -15.33 | Pathogenic | 0.78 | Disease | 1.00 | Probably damaging | 0.02 | Deleterious | 32.00 | Deleterious |
| p.Asp92Cys | VUS | -0.57 | Neutral | 0.9254 | Pathogenic | -13.02 | Pathogenic | 0.63 | Disease | 1.00 | Probably damaging | 0.01 | Deleterious | | |
| p.Asp92Trp | VUS | -3.72 | Neutral | 0.9587 | Pathogenic | -13.87 | Pathogenic | 0.67 | Disease | 1.00 | Probably damaging | 0 | Deleterious | | |
| p.Asp92Phe | VUS | -4.82 | Neutral | 0.9489 | Pathogenic | -16.79 | Pathogenic | 0.68 | Disease | 1.00 | Probably damaging | 0.01 | Deleterious | | |
| p.Thr93Asn | VUS | -16.51 | Indeterminate | 0.697 | Pathogenic | -9.47 | Pathogenic | 0.81 | Disease | 1.00 | Probably damaging | 0.07 | Tolerated | | |
| p.Thr93Lys | VUS | -53.15 | Deleterious | 0.9499 | Pathogenic | -12.30 | Pathogenic | 0.86 | Disease | 1.00 | Probably damaging | 0.07 | Tolerated | 29.50 | Deleterious |
| p.Thr93Arg | VUS | -33.14 | Indeterminate | 0.914 | Pathogenic | -15.35 | Pathogenic | 0.87 | Disease | 1.00 | Probably damaging | 0.04 | Deleterious | 29.00 | Deleterious |
| p.Thr93Ser | VUS | -8.91 | Indeterminate | 0.4599 | Ambiguous | -8.45 | Pathogenic | 0.68 | Disease | 1.00 | Probably damaging | 0.29 | Tolerated | 26.90 | Deleterious |
| p.Thr93Ile | VUS | -0.81 | Neutral | 0.6269 | Pathogenic | -9.66 | Pathogenic | 0.86 | Disease | 1.00 | Probably damaging | 0.23 | Tolerated | | |
| p.Thr93Met | VUS | -13.29 | Indeterminate | 0.4101 | Ambiguous | -9.46 | Pathogenic | 0.80 | Disease | 1.00 | Probably damaging | 0.08 | Tolerated | 29.70 | Deleterious |
| p.Thr93His | VUS | -25.89 | Indeterminate | 0.8809 | Pathogenic | -11.97 | Pathogenic | 0.78 | Disease | 1.00 | Probably damaging | 0.02 | Deleterious | | |
| p.Thr93Gln | VUS | -20.83 | Indeterminate | 0.8851 | Pathogenic | -11.62 | Pathogenic | 0.75 | Disease | 1.00 | Probably damaging | 0.05 | Deleterious | | |
| p.Thr93Pro | VUS | -53.15 | Deleterious | 0.6064 | Pathogenic | -14.33 | Pathogenic | 0.72 | Disease | 1.00 | Probably damaging | 0.08 | Tolerated | 28.20 | Deleterious |
| p.Thr93Leu | VUS | -9.12 | Indeterminate | 0.4735 | Ambiguous | -11.97 | Pathogenic | 0.78 | Disease | 1.00 | Probably damaging | 0.39 | Tolerated | | |
| p.Thr93Asp | VUS | -53.15 | Deleterious | 0.933 | Pathogenic | -15.10 | Pathogenic | 0.79 | Disease | 1.00 | Probably damaging | 0.05 | Deleterious | | |
| p.Thr93Glu | VUS | -53.15 | Deleterious | 0.9352 | Pathogenic | -14.21 | Pathogenic | 0.79 | Disease | 1.00 | Probably damaging | 0.05 | Deleterious | | |
| p.Thr93Ala | VUS | -23.13 | Indeterminate | 0.2995 | Benign | -8.21 | Pathogenic | 0.68 | Disease | 1.00 | Probably damaging | 0.28 | Tolerated | 27.10 | Deleterious |
| p.Thr93Gly | VUS | -23.79 | Indeterminate | 0.6733 | Pathogenic | -10.98 | Pathogenic | 0.73 | Disease | 1.00 | Probably damaging | 0.06 | Tolerated | | |
| p.Thr93Val | VUS | -5.99 | Indeterminate | 0.3711 | Ambiguous | -7.09 | Benign | 0.51 | Disease | 1.00 | Probably damaging | 0.4 | Tolerated | | |
| p.Thr93Tyr | VUS | -33.22 | Indeterminate | 0.9494 | Pathogenic | -16.27 | Pathogenic | 0.81 | Disease | 1.00 | Probably damaging | 0.03 | Deleterious | | |
| p.Thr93Cys | VUS | -9.44 | Indeterminate | 0.6612 | Pathogenic | -10.25 | Pathogenic | 0.77 | Disease | 1.00 | Probably damaging | 0.14 | Tolerated | | |
| p.Thr93Trp | VUS | -24.67 | Indeterminate | 0.9837 | Pathogenic | -17.35 | Pathogenic | 0.79 | Disease | 1.00 | Probably damaging | 0.01 | Deleterious | | |
| p.Thr93Phe | VUS | -15.25 | Indeterminate | 0.9286 | Pathogenic | -14.73 | Pathogenic | 0.81 | Disease | 1.00 | Probably damaging | 0.06 | Tolerated | | |
| p.Leu94Asn | VUS | -32.69 | Indeterminate | 0.986 | Pathogenic | -19.97 | Pathogenic | 0.76 | Disease | 1.00 | Probably damaging | 0 | Deleterious | | |
| p.Leu94Lys | VUS | -53.15 | Deleterious | 0.9865 | Pathogenic | -23.40 | Pathogenic | 0.75 | Disease | 0.99 | Probably damaging | 0.01 | Deleterious | | |
| p.Leu94Thr | VUS | -32.22 | Indeterminate | 0.6064 | Pathogenic | -12.32 | Pathogenic | 0.70 | Disease | 1.00 | Probably damaging | 0.01 | Deleterious | | |
| p.Leu94Arg | VUS | -53.15 | Deleterious | 0.9491 | Pathogenic | -19.07 | Pathogenic | 0.71 | Disease | 1.00 | Probably damaging | 0 | Deleterious | 32.00 | Deleterious |
| p.Leu94Ser | VUS | -0.47 | Neutral | 0.9449 | Pathogenic | -15.68 | Pathogenic | 0.75 | Disease | 1.00 | Probably damaging | 0 | Deleterious | | |

| | | | | | | | | | | | | | | | |
|---|---|---|---|---|---|---|---|---|---|---|---|---|---|---|---|
| p.Leu94Ile | VUS | -0.01 | Neutral | 0.1938 | Benign | -6.65 | Benign | 0.31 | Neutral | 1.00 | Probably damaging | 0.09 | Tolerated | | |
| p.Leu94Met | VUS | -0.01 | Neutral | 0.2911 | Benign | -9.48 | Pathogenic | 0.47 | Neutral | 1.00 | Probably damaging | 0.03 | Deleterious | 24.10 | Deleterious |
| p.Leu94His | VUS | -53.15 | Deleterious | 0.9832 | Pathogenic | -17.38 | Pathogenic | 0.80 | Disease | 1.00 | Probably damaging | 0 | Deleterious | | |
| p.Leu94Gln | Likely pathogenic | -53.15 | Deleterious | 0.9434 | Pathogenic | -17.94 | Pathogenic | 0.86 | Disease | 1.00 | Probably damaging | 0.01 | Deleterious | 31.00 | Deleterious |
| p.Leu94Pro | Likely pathogenic | -53.15 | Deleterious | 0.9693 | Pathogenic | -14.99 | Pathogenic | 0.92 | Disease | 1.00 | Probably damaging | 0.01 | Deleterious | 31.00 | Deleterious |
| p.Leu94Asp | VUS | -53.15 | Deleterious | 0.9968 | Pathogenic | -19.02 | Pathogenic | 0.73 | Disease | 1.00 | Probably damaging | 0 | Deleterious | | |
| p.Leu94Glu | VUS | -53.15 | Deleterious | 0.9784 | Pathogenic | -18.62 | Pathogenic | 0.74 | Disease | 1.00 | Probably damaging | 0 | Deleterious | | |
| p.Leu94Ala | VUS | -33.22 | Indeterminate | 0.5738 | Pathogenic | -10.94 | Pathogenic | 0.64 | Disease | 1.00 | Probably damaging | 0.02 | Deleterious | | |
| p.Leu94Gly | VUS | -53.15 | Deleterious | 0.9556 | Pathogenic | -16.15 | Pathogenic | 0.70 | Disease | 1.00 | Probably damaging | 0 | Deleterious | | |
| p.Leu94Val | VUS | -3.56 | Neutral | 0.096 | Benign | -5.53 | Benign | 0.22 | Neutral | 1.00 | Probably damaging | 0.45 | Tolerated | 18.64 | Deleterious |
| p.Leu94Tyr | VUS | -53.15 | Deleterious | 0.9829 | Pathogenic | -15.31 | Pathogenic | 0.68 | Disease | 1.00 | Probably damaging | 0.01 | Deleterious | | |
| p.Leu94Cys | VUS | -8.77 | Indeterminate | 0.8797 | Pathogenic | -12.71 | Pathogenic | 0.63 | Disease | 1.00 | Probably damaging | 0 | Deleterious | | |
| p.Leu94Trp | VUS | -53.15 | Deleterious | 0.9769 | Pathogenic | -16.43 | Pathogenic | 0.72 | Disease | 1.00 | Probably damaging | 0 | Deleterious | | |
| p.Leu94Phe | VUS | -9.94 | Indeterminate | 0.8065 | Pathogenic | -10.88 | Pathogenic | 0.52 | Disease | 1.00 | Probably damaging | 0.03 | Deleterious | | |
| p.Val95Asn | VUS | -4.64 | Neutral | 0.5539 | Ambiguous | -12.79 | Pathogenic | 0.37 | Neutral | 0.90 | Possibly damaging | 0.38 | Tolerated | | |
| p.Val95Lys | VUS | -0.83 | Neutral | 0.3308 | Benign | -8.07 | Pathogenic | 0.19 | Neutral | 0.33 | Benign | 1 | Tolerated | | |
| p.Val95Thr | VUS | -4.38 | Neutral | 0.2304 | Benign | -7.25 | Benign | 0.28 | Neutral | 0.67 | Possibly damaging | 0.45 | Tolerated | | |
| p.Val95Arg | VUS | -3.32 | Neutral | 0.1836 | Benign | -6.75 | Benign | 0.17 | Neutral | 0.52 | Possibly damaging | 0.65 | Tolerated | | |
| p.Val95Ser | VUS | -4.99 | Neutral | 0.3177 | Benign | -9.04 | Pathogenic | 0.33 | Neutral | 0.83 | Possibly damaging | 0.52 | Tolerated | | |
| p.Val95Ile | VUS | -2.11 | Neutral | 0.1425 | Benign | -6.02 | Benign | 0.20 | Neutral | 0.01 | Benign | 0.18 | Tolerated | | |
| p.Val95Met | VUS | -5.88 | Indeterminate | 0.2892 | Benign | -7.98 | Pathogenic | 0.12 | Neutral | 0.02 | Benign | 0.1 | Tolerated | 15.05 | Deleterious |
| p.Val95His | VUS | -0.48 | Neutral | 0.5271 | Ambiguous | -10.30 | Pathogenic | 0.23 | Neutral | 0.95 | Possibly damaging | 0.15 | Tolerated | | |
| p.Val95Gln | VUS | -11.31 | Indeterminate | 0.2673 | Benign | -7.77 | Pathogenic | 0.20 | Neutral | 0.42 | Benign | 0.56 | Tolerated | | |
| p.Val95Pro | VUS | -53.15 | Deleterious | 0.7501 | Pathogenic | -13.48 | Pathogenic | 0.38 | Neutral | 0.82 | Possibly damaging | 0.25 | Tolerated | | |
| p.Val95Leu | VUS | -1.93 | Neutral | 0.1973 | Benign | -7.70 | Pathogenic | 0.10 | Neutral | 0.00 | Benign | 0.28 | Tolerated | 13.66 | Neutral |
| p.Val95Asp | VUS | -2.24 | Neutral | 0.5199 | Ambiguous | -13.25 | Pathogenic | 0.37 | Neutral | 0.99 | Probably damaging | 0.53 | Tolerated | | |
| p.Val95Glu | VUS | -3.95 | Neutral | 0.2418 | Benign | -10.87 | Pathogenic | 0.41 | Neutral | 0.91 | Possibly damaging | 0.97 | Tolerated | 12.13 | Neutral |
| p.Val95Ala | VUS | -0.53 | Neutral | 0.1371 | Benign | -5.31 | Benign | 0.09 | Neutral | 0.02 | Benign | 0.6 | Tolerated | 10.96 | Neutral |
| p.Val95Gly | VUS | -2.19 | Neutral | 0.2822 | Benign | -9.77 | Pathogenic | 0.27 | Neutral | 0.93 | Possibly damaging | 0.29 | Tolerated | 14.08 | Neutral |
| p.Val95Tyr | VUS | -5.69 | Neutral | 0.6414 | Pathogenic | -12.32 | Pathogenic | 0.37 | Neutral | 0.68 | Possibly damaging | 0.08 | Tolerated | | |
| p.Val95Cys | VUS | -1.01 | Neutral | 0.7598 | Pathogenic | -7.33 | Benign | 0.37 | Neutral | 0.96 | Probably damaging | 0.05 | Deleterious | | |
| p.Val95Trp | VUS | -8.18 | Indeterminate | 0.8085 | Pathogenic | -11.12 | Pathogenic | 0.40 | Neutral | 0.99 | Probably damaging | 0.03 | Deleterious | | |
| p.Val95Phe | VUS | -6.73 | Indeterminate | 0.2799 | Benign | -12.40 | Pathogenic | 0.35 | Neutral | 0.52 | Possibly damaging | 0.07 | Tolerated | | |
| p.Val96Asn | VUS | -5.06 | Neutral | 0.7289 | Pathogenic | -13.54 | Pathogenic | 0.51 | Disease | 0.77 | Possibly damaging | 0.02 | Deleterious | | |
| p.Val96Lys | VUS | -0.44 | Neutral | 0.7191 | Pathogenic | -14.08 | Pathogenic | 0.51 | Disease | 0.05 | Benign | 0.04 | Deleterious | | |
| p.Val96Thr | VUS | -0.49 | Neutral | 0.2976 | Benign | -6.39 | Benign | 0.23 | Neutral | 0.56 | Possibly damaging | 0.23 | Tolerated | | |
| p.Val96Arg | VUS | -0.44 | Neutral | 0.4778 | Ambiguous | -10.88 | Pathogenic | 0.50 | Disease | 0.19 | Benign | 0.12 | Tolerated | | |
| p.Val96Ser | VUS | -0.05 | Neutral | 0.4376 | Ambiguous | -8.35 | Pathogenic | 0.41 | Neutral | 0.61 | Possibly damaging | 0.04 | Deleterious | | |
| p.Val96Ile | VUS | -5.81 | Indeterminate | 0.1363 | Benign | -7.68 | Pathogenic | 0.10 | Neutral | 0.00 | Benign | 0.48 | Tolerated | | |
| p.Val96Met | VUS | -0.70 | Neutral | 0.3646 | Ambiguous | -8.53 | Pathogenic | 0.11 | Neutral | 0.01 | Benign | 0.06 | Tolerated | 20.90 | Deleterious |
| p.Val96His | VUS | -4.73 | Neutral | 0.7703 | Pathogenic | -12.65 | Pathogenic | 0.46 | Neutral | 0.54 | Possibly damaging | 0.01 | Deleterious | | |
| p.Val96Gln | VUS | -2.82 | Neutral | 0.538 | Ambiguous | -11.35 | Pathogenic | 0.45 | Neutral | 0.07 | Benign | 0.03 | Deleterious | | |
| p.Val96Pro | VUS | -1.23 | Neutral | 0.7634 | Pathogenic | -10.28 | Pathogenic | 0.48 | Neutral | 0.66 | Possibly damaging | 0.05 | Deleterious | | |
| p.Val96Leu | VUS | -12.83 | Indeterminate | 0.25 | Benign | -5.17 | Benign | 0.07 | Neutral | 0.00 | Benign | 0.36 | Tolerated | 15.88 | Deleterious |
| p.Val96Asp | VUS | -10.72 | Indeterminate | 0.7853 | Pathogenic | -13.82 | Pathogenic | 0.59 | Disease | 0.90 | Possibly damaging | 0.02 | Deleterious | | |
| p.Val96Glu | VUS | -1.00 | Neutral | 0.5405 | Ambiguous | -12.30 | Pathogenic | 0.29 | Neutral | 0.01 | Benign | 0.04 | Deleterious | 14.61 | Neutral |
| p.Val96Ala | VUS | -53.15 | Deleterious | 0.2136 | Benign | -4.20 | Benign | 0.08 | Neutral | 0.00 | Benign | 0.32 | Tolerated | 6.24 | Neutral |
| p.Val96Gly | VUS | -5.35 | Neutral | 0.4053 | Ambiguous | -10.47 | Pathogenic | 0.37 | Neutral | 0.80 | Possibly damaging | 0.01 | Deleterious | 18.21 | Deleterious |
| p.Val96Tyr | VUS | -7.43 | Indeterminate | 0.7911 | Pathogenic | -13.57 | Pathogenic | 0.40 | Neutral | 0.83 | Possibly damaging | 0.02 | Deleterious | | |
| p.Val96Cys | VUS | -8.48 | Indeterminate | 0.766 | Pathogenic | -6.53 | Benign | 0.38 | Neutral | 0.83 | Possibly damaging | 0.05 | Deleterious | | |
| p.Val96Trp | VUS | -17.98 | Indeterminate | 0.8816 | Pathogenic | -11.55 | Pathogenic | 0.40 | Neutral | 0.98 | Probably damaging | 0.01 | Deleterious | | |
| p.Val96Phe | VUS | -3.30 | Neutral | 0.3632 | Ambiguous | -10.93 | Pathogenic | 0.38 | Neutral | 0.45 | Benign | 0.04 | Deleterious | | |
| p.Leu97Asn | VUS | -53.15 | Deleterious | 0.9929 | Pathogenic | -18.02 | Pathogenic | 0.90 | Disease | 1.00 | Probably damaging | 0 | Deleterious | | |
| p.Leu97Lys | VUS | -53.15 | Deleterious | 0.9869 | Pathogenic | -20.26 | Pathogenic | 0.87 | Disease | 0.99 | Probably damaging | 0 | Deleterious | | |
| p.Leu97Thr | VUS | -53.15 | Deleterious | 0.974 | Pathogenic | -14.28 | Pathogenic | 0.89 | Disease | 1.00 | Probably damaging | 0 | Deleterious | | |
| p.Leu97Arg | VUS | -53.15 | Deleterious | 0.9325 | Pathogenic | -14.97 | Pathogenic | 0.98 | Disease | 1.00 | Probably damaging | 0 | Deleterious | 29.90 | Deleterious |
| p.Leu97Ser | VUS | -53.15 | Deleterious | 0.9912 | Pathogenic | -12.95 | Pathogenic | 0.97 | Disease | 1.00 | Probably damaging | 0 | Deleterious | | |
| p.Leu97Ile | VUS | -0.26 | Neutral | 0.5969 | Pathogenic | -12.02 | Pathogenic | 0.76 | Disease | 1.00 | Probably damaging | 0 | Deleterious | | |
| p.Leu97Met | VUS | -0.03 | Neutral | 0.6247 | Pathogenic | -8.97 | Pathogenic | 0.82 | Disease | 1.00 | Probably damaging | 0 | Deleterious | 28.20 | Deleterious |
| p.Leu97His | VUS | -53.15 | Deleterious | 0.9765 | Pathogenic | -13.98 | Pathogenic | 0.96 | Disease | 1.00 | Probably damaging | 0 | Deleterious | | |
| p.Leu97Gln | VUS | -53.15 | Deleterious | 0.9516 | Pathogenic | -13.04 | Pathogenic | 0.98 | Disease | 1.00 | Probably damaging | 0 | Deleterious | 29.20 | Deleterious |
| p.Leu97Pro | VUS | -53.15 | Deleterious | 0.9583 | Pathogenic | -11.16 | Pathogenic | 0.99 | Disease | 1.00 | Probably damaging | 0 | Deleterious | 29.70 | Deleterious |
| p.Leu97Asp | VUS | -53.15 | Deleterious | 0.9983 | Pathogenic | -18.54 | Pathogenic | 0.88 | Disease | 1.00 | Probably damaging | 0 | Deleterious | | |
| p.Leu97Glu | VUS | -53.15 | Deleterious | 0.9855 | Pathogenic | -17.43 | Pathogenic | 0.90 | Disease | 1.00 | Probably damaging | 0 | Deleterious | | |
| p.Leu97Ala | VUS | -53.15 | Deleterious | 0.9489 | Pathogenic | -12.12 | Pathogenic | 0.87 | Disease | 1.00 | Probably damaging | 0 | Deleterious | | |
| p.Leu97Gly | VUS | -53.15 | Deleterious | 0.98 | Pathogenic | -16.62 | Pathogenic | 0.90 | Disease | 1.00 | Probably damaging | 0 | Deleterious | | |
| p.Leu97Val | VUS | -11.89 | Indeterminate | 0.6824 | Pathogenic | -12.00 | Pathogenic | 0.86 | Disease | 1.00 | Probably damaging | 0 | Deleterious | 27.70 | Deleterious |
| p.Leu97Tyr | VUS | -0.83 | Neutral | 0.9699 | Pathogenic | -14.71 | Pathogenic | 0.86 | Disease | 1.00 | Probably damaging | 0 | Deleterious | | |
| p.Leu97Cys | VUS | -0.82 | Neutral | 0.9414 | Pathogenic | -13.95 | Pathogenic | 0.82 | Disease | 1.00 | Probably damaging | 0 | Deleterious | | |
| p.Leu97Trp | VUS | -53.15 | Deleterious | 0.9348 | Pathogenic | -13.49 | Pathogenic | 0.89 | Disease | 1.00 | Probably damaging | 0 | Deleterious | | |
| p.Leu97Phe | VUS | -0.45 | Neutral | 0.7408 | Pathogenic | -11.26 | Pathogenic | 0.86 | Disease | 1.00 | Probably damaging | 0 | Deleterious | | |
| p.His98Asn | VUS | -0.67 | Neutral | 0.6006 | Pathogenic | -11.17 | Pathogenic | 0.57 | Disease | 1.00 | Probably damaging | 0.05 | Deleterious | 24.30 | Deleterious |
| p.His98Lys | VUS | -1.47 | Neutral | 0.8132 | Pathogenic | -11.46 | Pathogenic | 0.54 | Disease | 0.65 | Possibly damaging | 0.08 | Tolerated | | |
| p.His98Thr | VUS | -2.77 | Neutral | 0.8741 | Pathogenic | -13.12 | Pathogenic | 0.58 | Disease | 1.00 | Probably damaging | 0.16 | Tolerated | | |
| p.His98Arg | VUS | -0.45 | Neutral | 0.4032 | Ambiguous | -6.80 | Benign | 0.32 | Neutral | 0.02 | Benign | 0.07 | Tolerated | 21.30 | Deleterious |
| p.His98Ser | VUS | -3.55 | Neutral | 0.6155 | Pathogenic | -10.39 | Pathogenic | 0.55 | Disease | 1.00 | Probably damaging | 0.07 | Tolerated | | |
| p.His98Ile | VUS | -0.50 | Neutral | 0.8886 | Pathogenic | -13.55 | Pathogenic | 0.61 | Disease | 0.98 | Probably damaging | 0.45 | Tolerated | | |
| p.His98Met | VUS | -0.87 | Neutral | 0.9337 | Pathogenic | -13.32 | Pathogenic | 0.56 | Disease | 1.00 | Probably damaging | 0.23 | Tolerated | | |
| p.His98Gln | VUS | -4.71 | Neutral | 0.5458 | Ambiguous | -7.97 | Pathogenic | 0.54 | Disease | 0.80 | Possibly damaging | 0.07 | Tolerated | 21.20 | Deleterious |
| p.His98Pro | VUS | -53.15 | Deleterious | 0.9124 | Pathogenic | -16.70 | Pathogenic | 0.60 | Disease | 1.00 | Probably damaging | 0.07 | Tolerated | 23.90 | Deleterious |
| p.His98Leu | VUS | -2.42 | Neutral | 0.4531 | Ambiguous | -10.14 | Pathogenic | 0.33 | Neutral | 0.87 | Possibly damaging | 1 | Tolerated | 21.10 | Deleterious |
| p.His98Asp | VUS | -3.97 | Neutral | 0.6312 | Pathogenic | -11.83 | Pathogenic | 0.62 | Disease | 1.00 | Probably damaging | 0.03 | Deleterious | 24.60 | Deleterious |
| p.His98Glu | VUS | -0.45 | Neutral | 0.714 | Pathogenic | -12.40 | Pathogenic | 0.55 | Disease | 0.99 | Probably damaging | 0.06 | Tolerated | | |
| p.His98Ala | VUS | -0.20 | Neutral | 0.6583 | Pathogenic | -10.81 | Pathogenic | 0.56 | Disease | 1.00 | Probably damaging | 0.23 | Tolerated | | |
| p.His98Gly | VUS | -2.35 | Neutral | 0.7349 | Pathogenic | -11.64 | Pathogenic | 0.56 | Disease | 1.00 | Probably damaging | 0.04 | Deleterious | | |
| p.His98Val | VUS | -0.16 | Neutral | 0.7307 | Pathogenic | -10.20 | Pathogenic | 0.29 | Neutral | 1.00 | Probably damaging | 0.49 | Tolerated | | |
| p.His98Tyr | VUS | -0.56 | Neutral | 0.2623 | Benign | -8.49 | Pathogenic | 0.54 | Disease | 0.98 | Probably damaging | 0.08 | Tolerated | 23.20 | Deleterious |
| p.His98Cys | VUS | -0.94 | Neutral | 0.6208 | Pathogenic | -10.68 | Pathogenic | 0.54 | Disease | 1.00 | Probably damaging | 0.05 | Deleterious | | |
| p.His98Trp | VUS | -0.29 | Neutral | 0.6278 | Pathogenic | -10.62 | Pathogenic | 0.59 | Disease | 1.00 | Probably damaging | 0.03 | Deleterious | | |
| p.His98Phe | VUS | -0.37 | Neutral | 0.7347 | Pathogenic | -10.97 | Pathogenic | 0.38 | Neutral | 1.00 | Probably damaging | 0.21 | Tolerated | | |
| p.Arg99Asn | VUS | -8.76 | Indeterminate | 0.5313 | Ambiguous | -11.41 | Pathogenic | 0.50 | Neutral | 0.96 | Probably damaging | 0.48 | Tolerated | | |
| p.Arg99Lys | VUS | -6.84 | Indeterminate | 0.2293 | Benign | -7.88 | Pathogenic | 0.40 | Neutral | 0.02 | Benign | 0.79 | Tolerated | | |
| p.Arg99Thr | VUS | -18.03 | Indeterminate | 0.3207 | Benign | -12.37 | Pathogenic | 0.53 | Disease | 0.96 | Probably damaging | 0.25 | Tolerated | | |
| p.Arg99Ser | VUS | -10.56 | Indeterminate | 0.4364 | Ambiguous | -10.45 | Pathogenic | 0.30 | Neutral | 0.99 | Probably damaging | 0.3 | Tolerated | | |
| p.Arg99Ile | VUS | -7.17 | Indeterminate | 0.5564 | Ambiguous | -15.76 | Pathogenic | 0.54 | Disease | 1.00 | Probably damaging | 0.08 | Tolerated | | |
| p.Arg99Met | VUS | -15.32 | Indeterminate | 0.5306 | Ambiguous | -14.31 | Pathogenic | 0.51 | Disease | 0.99 | Probably damaging | 0.05 | Deleterious | | |
| p.Arg99His | VUS | -1.53 | Neutral | 0.1437 | Benign | -8.54 | Pathogenic | 0.47 | Neutral | 0.96 | Probably damaging | 0.07 | Tolerated | | |
| p.Arg99Gln | VUS | -2.51 | Neutral | 0.1211 | Benign | -8.63 | Pathogenic | 0.31 | Neutral | 0.49 | Possibly damaging | 0.68 | Tolerated | 14.30 | Neutral |
| p.Arg99Pro | Likely pathogenic | -53.15 | Deleterious | 0.9146 | Pathogenic | -15.88 | Pathogenic | 0.65 | Disease | 1.00 | Probably damaging | 0.13 | Tolerated | 22.40 | Deleterious |
| p.Arg99Leu | VUS | -26.04 | Indeterminate | 0.2472 | Benign | -11.47 | Pathogenic | 0.35 | Neutral | 1.00 | Probably damaging | 0.14 | Tolerated | 17.52 | Deleterious |
| p.Arg99Asp | VUS | -30.54 | Indeterminate | 0.4469 | Ambiguous | -12.39 | Pathogenic | 0.39 | Neutral | 0.99 | Probably damaging | 0.25 | Tolerated | | |
| p.Arg99Glu | VUS | -10.51 | Indeterminate | 0.2565 | Benign | -11.97 | Pathogenic | 0.36 | Neutral | 0.59 | Possibly damaging | 1 | Tolerated | | |
| p.Arg99Ala | VUS | -33.08 | Indeterminate | 0.2617 | Benign | -10.29 | Pathogenic | 0.39 | Neutral | 0.90 | Possibly damaging | 0.31 | Tolerated | | |
| p.Arg99Gly | VUS | -1.22 | Neutral | 0.1947 | Benign | -9.37 | Pathogenic | 0.32 | Neutral | 1.00 | Probably damaging | 0.26 | Tolerated | 22.90 | Deleterious |
| p.Arg99Val | VUS | -33.22 | Indeterminate | 0.4828 | Ambiguous | -13.31 | Pathogenic | 0.57 | Disease | 1.00 | Probably damaging | 0.12 | Tolerated | | |
| p.Arg99Tyr | VUS | -3.07 | Neutral | 0.5805 | Pathogenic | -14.21 | Pathogenic | 0.49 | Neutral | 0.93 | Possibly damaging | 0.03 | Deleterious | | |
| p.Arg99Cys | VUS | -21.73 | Indeterminate | 0.3804 | Ambiguous | -10.67 | Pathogenic | 0.55 | Disease | 1.00 | Probably damaging | 0.02 | Deleterious | | |
| p.Arg99Trp | VUS | -21.35 | Indeterminate | 0.3409 | Ambiguous | -11.97 | Pathogenic | 0.52 | Disease | 1.00 | Probably damaging | 0.01 | Deleterious | 26.30 | Deleterious |
| p.Arg99Phe | VUS | -4.69 | Neutral | 0.6897 | Pathogenic | -15.15 | Pathogenic | 0.55 | Disease | 0.93 | Possibly damaging | 0.02 | Deleterious | | |
| p.Ala100Asn | VUS | -1.08 | Neutral | 0.4496 | Ambiguous | -6.83 | Benign | 0.33 | Neutral | 1.00 | Probably damaging | 0.32 | Tolerated | | |
| p.Ala100Lys | VUS | 0.00 | Neutral | 0.6223 | Pathogenic | -8.59 | Pathogenic | 0.53 | Disease | 1.00 | Probably damaging | 0.29 | Tolerated | | |

| | | | | | | | | | | | | | | | |
|---|---|---|---|---|---|---|---|---|---|---|---|---|---|---|---|
| p.Ala100Thr | VUS | -1.44 | Neutral | 0.2533 | Benign | -10.20 | Pathogenic | 0.41 | Neutral | 1.00 | Probably damaging | 0.39 | Tolerated | 18.57 | Deleterious |
| p.Ala100Arg | VUS | 0.00 | Neutral | 0.3656 | Ambiguous | -4.54 | Benign | 0.52 | Disease | 1.00 | Probably damaging | 0.35 | Tolerated | | |
| p.Ala100Ser | Benign | -0.01 | Neutral | 0.1502 | Benign | -5.63 | Benign | 0.25 | Neutral | 0.97 | Probably damaging | 0.4 | Tolerated | 16.80 | Deleterious |
| p.Ala100Ile | VUS | -0.08 | Neutral | 0.6148 | Pathogenic | -13.16 | Pathogenic | 0.37 | Neutral | 1.00 | Probably damaging | 0.39 | Tolerated | | |
| p.Ala100Met | VUS | -0.05 | Neutral | 0.4862 | Ambiguous | -10.76 | Pathogenic | 0.47 | Neutral | 1.00 | Probably damaging | 0.22 | Tolerated | | |
| p.Ala100His | VUS | 0.00 | Neutral | 0.2773 | Benign | -2.80 | Benign | 0.47 | Neutral | 1.00 | Probably damaging | 0.54 | Tolerated | | |
| p.Ala100Gln | VUS | -0.06 | Neutral | 0.3302 | Benign | -8.15 | Pathogenic | 0.48 | Neutral | 1.00 | Probably damaging | 0.29 | Tolerated | | |
| p.Ala100Pro | VUS | -53.15 | Deleterious | 0.9671 | Pathogenic | -12.76 | Pathogenic | 0.56 | Disease | 1.00 | Probably damaging | 0.2 | Tolerated | 22.60 | Deleterious |
| p.Ala100Leu | VUS | -0.01 | Neutral | 0.4268 | Ambiguous | -8.48 | Pathogenic | 0.34 | Neutral | 1.00 | Probably damaging | 0.65 | Tolerated | | |
| p.Ala100Asp | VUS | -1.07 | Neutral | 0.6518 | Pathogenic | -11.39 | Pathogenic | 0.60 | Disease | 1.00 | Probably damaging | 0.21 | Tolerated | 22.80 | Deleterious |
| p.Ala100Glu | VUS | -3.66 | Neutral | 0.4117 | Ambiguous | -10.24 | Pathogenic | 0.32 | Neutral | 1.00 | Probably damaging | 0.28 | Tolerated | | |
| p.Ala100Gly | VUS | -53.15 | Deleterious | 0.1742 | Benign | -5.19 | Benign | 0.27 | Neutral | 1.00 | Probably damaging | 0.34 | Tolerated | 19.20 | Deleterious |
| p.Ala100Val | VUS | 0.00 | Neutral | 0.2647 | Benign | -8.74 | Pathogenic | 0.26 | Neutral | 1.00 | Probably damaging | 0.5 | Tolerated | 22.40 | Deleterious |
| p.Ala100Tyr | VUS | -1.42 | Neutral | 0.5139 | Ambiguous | -4.04 | Benign | 0.51 | Disease | 1.00 | Probably damaging | 1 | Tolerated | | |
| p.Ala100Cys | VUS | 0.00 | Neutral | 0.4116 | Ambiguous | -6.21 | Benign | 0.46 | Neutral | 1.00 | Probably damaging | 0.18 | Tolerated | | |
| p.Ala100Trp | VUS | 0.00 | Neutral | 0.6812 | Pathogenic | -4.53 | Benign | 0.55 | Disease | 1.00 | Probably damaging | 0.18 | Tolerated | | |
| p.Ala100Phe | VUS | 0.00 | Neutral | 0.4423 | Ambiguous | -4.04 | Benign | 0.36 | Neutral | 1.00 | Probably damaging | 0.7 | Tolerated | | |
| p.Gly101Asn | VUS | -6.43 | Indeterminate | 0.8106 | Pathogenic | -12.58 | Pathogenic | 0.56 | Disease | 1.00 | Probably damaging | 0.09 | Tolerated | | |
| p.Gly101Lys | VUS | -13.62 | Indeterminate | 0.7814 | Pathogenic | -12.51 | Pathogenic | 0.42 | Neutral | 1.00 | Probably damaging | 0.09 | Tolerated | | |
| p.Gly101Thr | VUS | -26.84 | Indeterminate | 0.8392 | Pathogenic | -13.88 | Pathogenic | 0.47 | Neutral | 1.00 | Probably damaging | 0.04 | Deleterious | | |
| p.Gly101Arg | VUS | -10.33 | Indeterminate | 0.6226 | Pathogenic | -9.29 | Pathogenic | 0.53 | Disease | 1.00 | Probably damaging | 0.04 | Deleterious | 26.75 | Deleterious |
| p.Gly101Ser | VUS | -7.05 | Indeterminate | 0.5163 | Ambiguous | -10.72 | Pathogenic | 0.54 | Disease | 1.00 | Probably damaging | 0.07 | Tolerated | | |
| p.Gly101Ile | VUS | -53.15 | Deleterious | 0.9774 | Pathogenic | -19.70 | Pathogenic | 0.65 | Disease | 1.00 | Probably damaging | 0.01 | Deleterious | | |
| p.Gly101Met | VUS | -14.55 | Indeterminate | 0.9705 | Pathogenic | -15.37 | Pathogenic | 0.66 | Disease | 1.00 | Probably damaging | 0.01 | Deleterious | | |
| p.Gly101His | VUS | -12.96 | Indeterminate | 0.8225 | Pathogenic | -11.05 | Pathogenic | 0.40 | Neutral | 1.00 | Probably damaging | 0.03 | Deleterious | | |
| p.Gly101Gln | VUS | -7.18 | Indeterminate | 0.7227 | Pathogenic | -12.08 | Pathogenic | 0.43 | Neutral | 1.00 | Probably damaging | 0.25 | Tolerated | | |
| p.Gly101Pro | VUS | -53.15 | Deleterious | 0.9934 | Pathogenic | -17.02 | Pathogenic | 0.62 | Disease | 1.00 | Probably damaging | 0.04 | Deleterious | | |
| p.Gly101Leu | VUS | -53.15 | Deleterious | 0.9599 | Pathogenic | -14.19 | Pathogenic | 0.65 | Disease | 1.00 | Probably damaging | 0.02 | Deleterious | | |
| p.Gly101Asp | VUS | -3.90 | Neutral | 0.6576 | Pathogenic | -11.78 | Pathogenic | 0.61 | Disease | 1.00 | Probably damaging | 0.17 | Tolerated | | |
| p.Gly101Glu | VUS | -15.02 | Indeterminate | 0.759 | Pathogenic | -13.11 | Pathogenic | 0.63 | Disease | 1.00 | Probably damaging | 0.35 | Tolerated | 26.80 | Deleterious |
| p.Gly101Ala | VUS | -11.65 | Indeterminate | 0.6391 | Pathogenic | -10.21 | Pathogenic | 0.51 | Disease | 1.00 | Probably damaging | 0.09 | Tolerated | 25.60 | Deleterious |
| p.Gly101Val | VUS | -53.15 | Deleterious | 0.9342 | Pathogenic | -15.06 | Pathogenic | 0.67 | Disease | 1.00 | Probably damaging | 0.02 | Deleterious | 26.60 | Deleterious |
| p.Gly101Tyr | VUS | -33.17 | Indeterminate | 0.9532 | Pathogenic | -15.85 | Pathogenic | 0.65 | Disease | 1.00 | Probably damaging | 0.01 | Deleterious | | |
| p.Gly101Cys | VUS | -5.64 | Neutral | 0.8484 | Pathogenic | -12.72 | Pathogenic | 0.65 | Disease | 1.00 | Probably damaging | 0.01 | Deleterious | | |
| p.Gly101Trp | Pathogenic | -53.15 | Deleterious | 0.9495 | Pathogenic | -14.73 | Pathogenic | 0.71 | Disease | 1.00 | Probably damaging | 0 | Deleterious | 28.90 | Deleterious |
| p.Gly101Phe | VUS | -32.01 | Indeterminate | 0.9773 | Pathogenic | -15.88 | Pathogenic | 0.65 | Disease | 1.00 | Probably damaging | 0.01 | Deleterious | | |
| p.Ala102Asn | VUS | -53.15 | Deleterious | 0.9525 | Pathogenic | -17.84 | Pathogenic | 0.80 | Disease | 0.91 | Possibly damaging | 0 | Deleterious | | |
| p.Ala102Lys | VUS | -53.15 | Deleterious | 0.9904 | Pathogenic | -22.38 | Pathogenic | 0.83 | Disease | 1.00 | Probably damaging | 0 | Deleterious | | |
| p.Ala102Thr | VUS | -53.15 | Deleterious | 0.6023 | Pathogenic | -11.13 | Pathogenic | 0.79 | Disease | 0.92 | Possibly damaging | 0.01 | Deleterious | 28.40 | Deleterious |
| p.Ala102Arg | VUS | -53.15 | Deleterious | 0.9445 | Pathogenic | -18.16 | Pathogenic | 0.79 | Disease | 1.00 | Probably damaging | 0 | Deleterious | | |
| p.Ala102Ser | VUS | -53.15 | Deleterious | 0.2741 | Benign | -9.44 | Pathogenic | 0.74 | Disease | 0.39 | Benign | 0.02 | Deleterious | 27.00 | Deleterious |
| p.Ala102Ile | VUS | -53.15 | Deleterious | 0.9049 | Pathogenic | -16.43 | Pathogenic | 0.83 | Disease | 1.00 | Probably damaging | 0 | Deleterious | | |
| p.Ala102Met | VUS | -53.15 | Deleterious | 0.8938 | Pathogenic | -15.76 | Pathogenic | 0.78 | Disease | 1.00 | Probably damaging | 0 | Deleterious | | |
| p.Ala102His | VUS | -53.15 | Deleterious | 0.9635 | Pathogenic | -18.36 | Pathogenic | 0.81 | Disease | 1.00 | Probably damaging | 0 | Deleterious | | |
| p.Ala102Gln | VUS | -53.15 | Deleterious | 0.9233 | Pathogenic | -17.47 | Pathogenic | 0.84 | Disease | 1.00 | Probably damaging | 0 | Deleterious | | |
| p.Ala102Pro | VUS | -53.15 | Deleterious | 0.9353 | Pathogenic | -13.22 | Pathogenic | 0.86 | Disease | 1.00 | Probably damaging | 0.01 | Deleterious | 28.30 | Deleterious |
| p.Ala102Leu | VUS | -53.15 | Deleterious | 0.7181 | Pathogenic | -14.15 | Pathogenic | 0.82 | Disease | 0.99 | Probably damaging | 0 | Deleterious | | |
| p.Ala102Asp | VUS | -53.15 | Deleterious | 0.9578 | Pathogenic | -15.69 | Pathogenic | 0.88 | Disease | 0.99 | Probably damaging | 0 | Deleterious | | |
| p.Ala102Glu | VUS | -53.15 | Deleterious | 0.9565 | Pathogenic | -16.01 | Pathogenic | 0.86 | Disease | 1.00 | Probably damaging | 0 | Deleterious | 27.70 | Deleterious |
| p.Ala102Gly | VUS | -0.03 | Neutral | 0.2432 | Benign | -8.21 | Pathogenic | 0.66 | Disease | 0.04 | Benign | 0.06 | Tolerated | 28.80 | Deleterious |
| p.Ala102Val | VUS | -18.01 | Indeterminate | 0.639 | Pathogenic | -9.19 | Pathogenic | 0.75 | Disease | 1.00 | Probably damaging | 0.01 | Deleterious | 28.70 | Deleterious |
| p.Ala102Tyr | VUS | -53.15 | Deleterious | 0.9663 | Pathogenic | -19.07 | Pathogenic | 0.83 | Disease | 1.00 | Probably damaging | 0 | Deleterious | | |
| p.Ala102Cys | VUS | -12.41 | Indeterminate | 0.6812 | Pathogenic | -10.37 | Pathogenic | 0.75 | Disease | 1.00 | Probably damaging | 0 | Deleterious | | |
| p.Ala102Trp | VUS | -53.15 | Deleterious | 0.981 | Pathogenic | -17.78 | Pathogenic | 0.83 | Disease | 1.00 | Probably damaging | 0 | Deleterious | | |
| p.Ala102Phe | VUS | -53.15 | Deleterious | 0.9308 | Pathogenic | -18.43 | Pathogenic | 0.84 | Disease | 1.00 | Probably damaging | 0 | Deleterious | | |
| p.Arg103Asn | VUS | -0.01 | Neutral | 0.4251 | Ambiguous | -11.54 | Pathogenic | 0.38 | Neutral | 1.00 | Probably damaging | 0.89 | Tolerated | | |
| p.Arg103Lys | VUS | -0.48 | Neutral | 0.2025 | Benign | -7.78 | Pathogenic | 0.36 | Neutral | 0.78 | Possibly damaging | 0.26 | Tolerated | | |
| p.Arg103Thr | VUS | -6.16 | Indeterminate | 0.1995 | Benign | -10.46 | Pathogenic | 0.50 | Neutral | 1.00 | Probably damaging | 0.17 | Tolerated | | |
| p.Arg103Ser | VUS | -0.01 | Neutral | 0.2587 | Benign | -9.19 | Pathogenic | 0.30 | Neutral | 1.00 | Probably damaging | 0.41 | Tolerated | | |
| p.Arg103Ile | VUS | -3.14 | Neutral | 0.3389 | Benign | -12.16 | Pathogenic | 0.53 | Disease | 1.00 | Probably damaging | 0.02 | Deleterious | | |
| p.Arg103Met | VUS | -3.90 | Neutral | 0.3543 | Ambiguous | -11.90 | Pathogenic | 0.54 | Disease | 1.00 | Probably damaging | 0.02 | Deleterious | | |
| p.Arg103His | VUS | -0.99 | Neutral | 0.1323 | Benign | -7.95 | Pathogenic | 0.32 | Neutral | 0.99 | Probably damaging | 0.16 | Tolerated | | |
| p.Arg103Gln | VUS | -0.97 | Neutral | 0.1029 | Benign | -8.15 | Pathogenic | 0.28 | Neutral | 0.94 | Possibly damaging | 0.17 | Tolerated | 19.34 | Deleterious |
| p.Arg103Pro | VUS | -0.63 | Neutral | 0.3065 | Benign | -10.10 | Pathogenic | 0.58 | Disease | 1.00 | Probably damaging | 0.13 | Tolerated | 22.90 | Deleterious |
| p.Arg103Leu | VUS | -0.01 | Neutral | 0.1593 | Benign | -9.92 | Pathogenic | 0.55 | Disease | 1.00 | Probably damaging | 0.04 | Deleterious | 20.30 | Deleterious |
| p.Arg103Asp | VUS | 0.00 | Neutral | 0.3581 | Ambiguous | -10.15 | Pathogenic | 0.29 | Neutral | 1.00 | Probably damaging | 1 | Tolerated | | |
| p.Arg103Glu | VUS | -2.72 | Neutral | 0.2591 | Benign | -10.02 | Pathogenic | 0.37 | Neutral | 0.90 | Possibly damaging | 0.28 | Tolerated | | |
| p.Arg103Ala | VUS | 0.00 | Neutral | 0.1801 | Benign | -8.30 | Pathogenic | 0.45 | Neutral | 0.98 | Probably damaging | 0.17 | Tolerated | | |
| p.Arg103Gly | VUS | -0.01 | Neutral | 0.1779 | Benign | -10.95 | Pathogenic | 0.32 | Neutral | 1.00 | Probably damaging | 0.52 | Tolerated | 22.60 | Deleterious |
| p.Arg103Val | VUS | -0.16 | Neutral | 0.2474 | Benign | -9.95 | Pathogenic | 0.55 | Disease | 1.00 | Probably damaging | 0.03 | Deleterious | | |
| p.Arg103Tyr | VUS | -0.63 | Neutral | 0.4737 | Ambiguous | -11.61 | Pathogenic | 0.50 | Neutral | 0.99 | Probably damaging | 0.08 | Tolerated | | |
| p.Arg103Cys | VUS | -0.01 | Neutral | 0.2144 | Benign | -9.27 | Pathogenic | 0.56 | Disease | 1.00 | Probably damaging | 0.04 | Deleterious | | |
| p.Arg103Trp | VUS | -0.24 | Neutral | 0.2676 | Benign | -11.13 | Pathogenic | 0.54 | Disease | 1.00 | Probably damaging | 0.02 | Deleterious | 26.40 | Deleterious |
| p.Arg103Phe | VUS | 0.00 | Neutral | 0.5257 | Ambiguous | -12.21 | Pathogenic | 0.53 | Disease | 0.99 | Probably damaging | 0.03 | Deleterious | | |
| p.Leu104Asn | VUS | 0.00 | Neutral | 0.8502 | Pathogenic | -17.13 | Pathogenic | 0.62 | Disease | 1.00 | Probably damaging | 0 | Deleterious | | |
| p.Leu104Lys | VUS | -11.35 | Indeterminate | 0.7882 | Pathogenic | -17.24 | Pathogenic | 0.64 | Disease | 0.98 | Probably damaging | 0.01 | Deleterious | | |
| p.Leu104Thr | VUS | 0.00 | Neutral | 0.3934 | Ambiguous | -9.66 | Pathogenic | 0.61 | Disease | 1.00 | Probably damaging | 0.02 | Deleterious | | |
| p.Leu104Arg | VUS | -53.15 | Deleterious | 0.5814 | Pathogenic | -13.14 | Pathogenic | 0.73 | Disease | 1.00 | Probably damaging | 0.01 | Deleterious | 28.00 | Deleterious |
| p.Leu104Ser | VUS | 0.00 | Neutral | 0.7933 | Pathogenic | -12.46 | Pathogenic | 0.65 | Disease | 1.00 | Probably damaging | 0.01 | Deleterious | | |
| p.Leu104Ile | VUS | 0.00 | Neutral | 0.2168 | Benign | -7.73 | Pathogenic | 0.45 | Neutral | 1.00 | Probably damaging | 0.41 | Tolerated | | |
| p.Leu104Met | VUS | 0.00 | Neutral | 0.2598 | Benign | -5.42 | Benign | 0.46 | Neutral | 1.00 | Probably damaging | 0.04 | Deleterious | 25.70 | Deleterious |
| p.Leu104His | VUS | 0.00 | Neutral | 0.7805 | Pathogenic | -14.47 | Pathogenic | 0.62 | Disease | 1.00 | Probably damaging | 0 | Deleterious | | |
| p.Leu104Gln | VUS | 0.00 | Neutral | 0.6621 | Pathogenic | -13.35 | Pathogenic | 0.69 | Disease | 1.00 | Probably damaging | 0.01 | Deleterious | 27.40 | Deleterious |
| p.Leu104Pro | VUS | -0.01 | Neutral | 0.2401 | Benign | -9.68 | Pathogenic | 0.27 | Neutral | 1.00 | Probably damaging | 0.01 | Deleterious | 24.50 | Deleterious |
| p.Leu104Asp | VUS | 0.00 | Neutral | 0.9278 | Pathogenic | -16.57 | Pathogenic | 0.64 | Disease | 1.00 | Probably damaging | 0 | Deleterious | | |
| p.Leu104Glu | VUS | 0.00 | Neutral | 0.7798 | Pathogenic | -15.90 | Pathogenic | 0.64 | Disease | 1.00 | Probably damaging | 0.01 | Deleterious | | |
| p.Leu104Ala | VUS | 0.00 | Neutral | 0.5071 | Ambiguous | -10.47 | Pathogenic | 0.37 | Neutral | 1.00 | Probably damaging | 0.02 | Deleterious | | |
| p.Leu104Gly | VUS | -53.15 | Deleterious | 0.7488 | Pathogenic | -12.87 | Pathogenic | 0.61 | Disease | 1.00 | Probably damaging | 0.01 | Deleterious | | |
| p.Leu104Val | VUS | 0.00 | Neutral | 0.1816 | Benign | -7.26 | Benign | 0.16 | Neutral | 0.99 | Probably damaging | 0.84 | Tolerated | 23.50 | Deleterious |
| p.Leu104Tyr | VUS | 0.00 | Neutral | 0.8168 | Pathogenic | -14.67 | Pathogenic | 0.55 | Disease | 0.99 | Probably damaging | 0.01 | Deleterious | | |
| p.Leu104Cys | VUS | 0.00 | Neutral | 0.7713 | Pathogenic | -11.82 | Pathogenic | 0.56 | Disease | 1.00 | Probably damaging | 0.01 | Deleterious | | |
| p.Leu104Trp | VUS | 0.00 | Neutral | 0.5661 | Pathogenic | -12.84 | Pathogenic | 0.57 | Disease | 1.00 | Probably damaging | 0 | Deleterious | | |
| p.Leu104Phe | VUS | 0.00 | Neutral | 0.439 | Ambiguous | -10.35 | Pathogenic | 0.50 | Neutral | 1.00 | Probably damaging | 0.03 | Deleterious | | |
| p.Asp105Asn | VUS | -2.86 | Neutral | 0.2727 | Benign | -8.41 | Pathogenic | 0.24 | Neutral | 0.98 | Probably damaging | 1 | Tolerated | 19.47 | Deleterious |
| p.Asp105Lys | VUS | -15.87 | Indeterminate | 0.8635 | Pathogenic | -11.29 | Pathogenic | 0.50 | Neutral | 1.00 | Probably damaging | 0.15 | Tolerated | | |
| p.Asp105Thr | VUS | -12.21 | Indeterminate | 0.5806 | Pathogenic | -7.97 | Pathogenic | 0.47 | Neutral | 1.00 | Probably damaging | 0.12 | Tolerated | | |
| p.Asp105Arg | VUS | -32.93 | Indeterminate | 0.7381 | Pathogenic | -7.21 | Benign | 0.48 | Neutral | 1.00 | Probably damaging | 0.08 | Tolerated | | |
| p.Asp105Ser | VUS | -53.15 | Deleterious | 0.4241 | Ambiguous | -7.28 | Benign | 0.43 | Neutral | 1.00 | Probably damaging | 0.26 | Tolerated | | |
| p.Asp105Ile | VUS | -53.15 | Deleterious | 0.861 | Pathogenic | -13.45 | Pathogenic | 0.50 | Neutral | 1.00 | Probably damaging | 0.02 | Deleterious | | |
| p.Asp105Met | VUS | -15.69 | Indeterminate | 0.9269 | Pathogenic | -11.01 | Pathogenic | 0.52 | Disease | 1.00 | Probably damaging | 0.01 | Deleterious | | |
| p.Asp105His | VUS | -11.71 | Indeterminate | 0.5656 | Pathogenic | -6.84 | Benign | 0.30 | Neutral | 1.00 | Probably damaging | 0.1 | Tolerated | 23.10 | Deleterious |
| p.Asp105Gln | VUS | -18.85 | Indeterminate | 0.6735 | Pathogenic | -7.92 | Pathogenic | 0.29 | Neutral | 1.00 | Probably damaging | 0.1 | Tolerated | | |
| p.Asp105Pro | VUS | -53.15 | Deleterious | 0.9092 | Pathogenic | -11.63 | Pathogenic | 0.51 | Disease | 0.97 | Probably damaging | 0.08 | Tolerated | | |
| p.Asp105Leu | VUS | -18.17 | Indeterminate | 0.8148 | Pathogenic | -8.91 | Pathogenic | 0.50 | Disease | 1.00 | Probably damaging | 0.18 | Tolerated | | |
| p.Asp105Glu | VUS | -17.38 | Indeterminate | 0.4885 | Ambiguous | -6.27 | Benign | 0.27 | Neutral | 0.92 | Possibly damaging | 0.2 | Tolerated | 13.08 | Neutral |
| p.Asp105Ala | VUS | -6.08 | Indeterminate | 0.4992 | Ambiguous | -6.15 | Benign | 0.42 | Neutral | 1.00 | Probably damaging | 0.11 | Tolerated | 24.10 | Deleterious |
| p.Asp105Gly | VUS | -53.15 | Deleterious | 0.4131 | Ambiguous | -4.39 | Benign | 0.38 | Neutral | 1.00 | Probably damaging | 0.33 | Tolerated | 24.60 | Deleterious |
| p.Asp105Val | VUS | -29.16 | Indeterminate | 0.6699 | Pathogenic | -9.73 | Pathogenic | 0.53 | Disease | 1.00 | Probably damaging | 0.03 | Deleterious | 25.90 | Deleterious |
| p.Asp105Tyr | VUS | 0.00 | Neutral | 0.6581 | Pathogenic | -11.05 | Pathogenic | 0.57 | Disease | 1.00 | Probably damaging | 0.05 | Deleterious | 23.40 | Deleterious |
| p.Asp105Cys | VUS | -0.23 | Neutral | 0.888 | Pathogenic | -9.07 | Pathogenic | 0.49 | Neutral | 1.00 | Probably damaging | 0.02 | Deleterious | | |
| p.Asp105Trp | VUS | -6.73 | Indeterminate | 0.9483 | Pathogenic | -10.19 | Pathogenic | 0.42 | Neutral | 1.00 | Probably damaging | 0.01 | Deleterious | | |

| | | | | | | | | | | | | | | | |
|---|---|---|---|---|---|---|---|---|---|---|---|---|---|---|---|
| p.Asp105Phe | VUS | -16.15 | Indeterminate | 0.9163 | Pathogenic | -12.06 | Pathogenic | 0.51 | Disease | 1.00 | Probably damaging | 0.03 | Deleterious | | |
| p.Val106Asn | VUS | -8.17 | Indeterminate | 0.5709 | Pathogenic | -13.95 | Pathogenic | 0.51 | Disease | 1.00 | Probably damaging | 0.03 | Deleterious | | |
| p.Val106Lys | VUS | -16.70 | Indeterminate | 0.5522 | Ambiguous | -15.08 | Pathogenic | 0.55 | Disease | 1.00 | Probably damaging | 0.05 | Deleterious | | |
| p.Val106Thr | VUS | -1.68 | Neutral | 0.2756 | Benign | -7.63 | Pathogenic | 0.30 | Neutral | 1.00 | Probably damaging | 0.16 | Tolerated | | |
| p.Val106Arg | VUS | -24.14 | Indeterminate | 0.3403 | Ambiguous | -10.26 | Pathogenic | 0.51 | Disease | 1.00 | Probably damaging | 0.04 | Deleterious | | |
| p.Val106Ser | VUS | -0.72 | Neutral | 0.3524 | Ambiguous | -8.63 | Pathogenic | 0.48 | Neutral | 1.00 | Probably damaging | 0.14 | Tolerated | | |
| p.Val106Ile | VUS | -1.09 | Neutral | 0.1505 | Benign | -7.97 | Pathogenic | 0.11 | Neutral | 0.97 | Probably damaging | 0.55 | Tolerated | | |
| p.Val106Met | VUS | -0.81 | Neutral | 0.3682 | Ambiguous | -7.68 | Pathogenic | 0.36 | Neutral | 1.00 | Probably damaging | 0.07 | Tolerated | 25.60 | Deleterious |
| p.Val106His | VUS | -0.71 | Neutral | 0.6751 | Pathogenic | -10.67 | Pathogenic | 0.51 | Disease | 1.00 | Probably damaging | 0.03 | Deleterious | | |
| p.Val106Gln | VUS | -1.67 | Neutral | 0.3212 | Benign | -8.12 | Pathogenic | 0.30 | Neutral | 1.00 | Probably damaging | 0.04 | Deleterious | | |
| p.Val106Pro | VUS | -21.83 | Indeterminate | 0.8011 | Pathogenic | -9.45 | Pathogenic | 0.51 | Disease | 1.00 | Probably damaging | 0.05 | Deleterious | | |
| p.Val106Leu | VUS | -9.78 | Indeterminate | 0.3179 | Benign | -5.92 | Benign | 0.09 | Neutral | 0.98 | Probably damaging | 0.25 | Tolerated | 21.55 | Deleterious |
| p.Val106Asp | VUS | -1.29 | Neutral | 0.4428 | Ambiguous | -10.12 | Pathogenic | 0.59 | Disease | 1.00 | Probably damaging | 0.02 | Deleterious | | |
| p.Val106Glu | VUS | -29.44 | Indeterminate | 0.2415 | Benign | -8.47 | Pathogenic | 0.60 | Disease | 1.00 | Probably damaging | 0.04 | Deleterious | 26.30 | Deleterious |
| p.Val106Ala | VUS | -5.62 | Neutral | 0.2098 | Benign | -4.32 | Benign | 0.13 | Neutral | 1.00 | Probably damaging | 0.4 | Tolerated | 23.40 | Deleterious |
| p.Val106Gly | VUS | -9.59 | Indeterminate | 0.2586 | Benign | -8.95 | Pathogenic | 0.43 | Neutral | 1.00 | Probably damaging | 0.03 | Deleterious | 26.10 | Deleterious |
| p.Val106Tyr | VUS | -1.56 | Neutral | 0.7948 | Pathogenic | -14.22 | Pathogenic | 0.50 | Neutral | 1.00 | Probably damaging | 0.06 | Tolerated | | |
| p.Val106Cys | VUS | -1.64 | Neutral | 0.6917 | Pathogenic | -9.60 | Pathogenic | 0.44 | Neutral | 1.00 | Probably damaging | 0.05 | Deleterious | | |
| p.Val106Trp | VUS | -13.39 | Indeterminate | 0.8982 | Pathogenic | -12.41 | Pathogenic | 0.47 | Neutral | 1.00 | Probably damaging | 0.01 | Deleterious | | |
| p.Val106Phe | VUS | -1.15 | Neutral | 0.3839 | Ambiguous | -11.84 | Pathogenic | 0.49 | Neutral | 1.00 | Probably damaging | 0.17 | Tolerated | | |
| p.Arg107Asn | VUS | -2.70 | Neutral | 0.6572 | Pathogenic | -14.08 | Pathogenic | 0.44 | Neutral | 0.27 | Benign | 0.5 | Tolerated | | |
| p.Arg107Lys | VUS | -0.88 | Neutral | 0.2096 | Benign | -7.96 | Pathogenic | 0.20 | Neutral | 0.02 | Benign | 0.92 | Tolerated | | |
| p.Arg107Thr | VUS | -0.06 | Neutral | 0.2177 | Benign | -9.41 | Pathogenic | 0.43 | Neutral | 0.43 | Benign | 0.6 | Tolerated | | |
| p.Arg107Ser | VUS | -3.51 | Neutral | 0.4114 | Ambiguous | -10.34 | Pathogenic | 0.38 | Neutral | 0.86 | Possibly damaging | 0.73 | Tolerated | 23.80 | Deleterious |
| p.Arg107Ile | VUS | -1.12 | Neutral | 0.3829 | Ambiguous | -13.41 | Pathogenic | 0.49 | Neutral | 0.93 | Possibly damaging | 0.22 | Tolerated | | |
| p.Arg107Met | VUS | -0.08 | Neutral | 0.4767 | Ambiguous | -11.83 | Pathogenic | 0.49 | Neutral | 0.72 | Possibly damaging | 0.12 | Tolerated | | |
| p.Arg107His | VUS | -1.66 | Neutral | 0.2041 | Benign | -8.80 | Pathogenic | 0.19 | Neutral | 0.00 | Benign | 0.16 | Tolerated | 18.27 | Deleterious |
| p.Arg107Gln | VUS | 0.00 | Neutral | 0.1418 | Benign | -8.82 | Pathogenic | 0.19 | Neutral | 0.02 | Benign | 0.6 | Tolerated | | |
| p.Arg107Pro | VUS | -0.08 | Neutral | 0.1271 | Benign | -6.34 | Benign | 0.22 | Neutral | 0.98 | Probably damaging | 0.29 | Tolerated | 16.64 | Deleterious |
| p.Arg107Leu | VUS | -2.08 | Neutral | 0.2011 | Benign | -8.63 | Pathogenic | 0.26 | Neutral | 0.88 | Possibly damaging | 0.35 | Tolerated | 22.00 | Deleterious |
| p.Arg107Asp | VUS | -1.62 | Neutral | 0.6577 | Pathogenic | -14.97 | Pathogenic | 0.47 | Neutral | 0.76 | Possibly damaging | 0.61 | Tolerated | | |
| p.Arg107Glu | VUS | -5.67 | Neutral | 0.3408 | Ambiguous | -10.44 | Pathogenic | 0.29 | Neutral | 0.04 | Benign | 1 | Tolerated | | |
| p.Arg107Ala | VUS | -0.01 | Neutral | 0.1877 | Benign | -7.89 | Pathogenic | 0.24 | Neutral | 0.24 | Benign | 0.77 | Tolerated | | |
| p.Arg107Gly | VUS | -1.60 | Neutral | 0.1981 | Benign | -9.62 | Pathogenic | 0.43 | Neutral | 0.86 | Possibly damaging | 0.39 | Tolerated | 24.10 | Deleterious |
| p.Arg107Val | VUS | -0.10 | Neutral | 0.3024 | Benign | -10.32 | Pathogenic | 0.51 | Disease | 0.73 | Possibly damaging | 0.35 | Tolerated | | |
| p.Arg107Tyr | VUS | -1.06 | Neutral | 0.5547 | Ambiguous | -10.26 | Pathogenic | 0.45 | Neutral | 0.04 | Benign | 0.12 | Tolerated | | |
| p.Arg107Cys | VUS | -8.51 | Indeterminate | 0.2239 | Benign | -9.83 | Pathogenic | 0.50 | Neutral | 0.22 | Benign | 0.08 | Tolerated | 28.50 | Deleterious |
| p.Arg107Trp | VUS | -0.08 | Neutral | 0.3155 | Benign | -12.74 | Pathogenic | 0.49 | Neutral | 0.86 | Possibly damaging | 0.03 | Deleterious | | |
| p.Arg107Phe | VUS | -1.19 | Neutral | 0.6012 | Pathogenic | -10.67 | Pathogenic | 0.51 | Disease | 0.07 | Benign | 0.11 | Tolerated | | |
| p.Asp108Asn | VUS | -0.78 | Neutral | 0.8442 | Pathogenic | -9.82 | Pathogenic | 0.70 | Disease | 1.00 | Probably damaging | 0.43 | Tolerated | 31.00 | Deleterious |
| p.Asp108Lys | VUS | -53.15 | Deleterious | 0.9973 | Pathogenic | -18.69 | Pathogenic | 0.78 | Disease | 1.00 | Probably damaging | 0.02 | Deleterious | | |
| p.Asp108Thr | VUS | -31.02 | Indeterminate | 0.9831 | Pathogenic | -13.75 | Pathogenic | 0.77 | Disease | 1.00 | Probably damaging | 0.02 | Deleterious | | |
| p.Asp108Arg | VUS | -53.15 | Deleterious | 0.9944 | Pathogenic | -19.00 | Pathogenic | 0.78 | Disease | 1.00 | Probably damaging | 0.01 | Deleterious | | |
| p.Asp108Ser | VUS | -53.15 | Deleterious | 0.9524 | Pathogenic | -14.48 | Pathogenic | 0.74 | Disease | 1.00 | Probably damaging | 0.22 | Tolerated | | |
| p.Asp108Ile | VUS | -53.15 | Deleterious | 0.9979 | Pathogenic | -19.67 | Pathogenic | 0.82 | Disease | 1.00 | Probably damaging | 0 | Deleterious | | |
| p.Asp108Met | VUS | -53.15 | Deleterious | 0.9983 | Pathogenic | -18.14 | Pathogenic | 0.81 | Disease | 1.00 | Probably damaging | 0 | Deleterious | | |
| p.Asp108His | VUS | -53.15 | Deleterious | 0.9917 | Pathogenic | -16.58 | Pathogenic | 0.87 | Disease | 1.00 | Probably damaging | 0.02 | Deleterious | 29.20 | Deleterious |
| p.Asp108Gln | VUS | -53.15 | Deleterious | 0.9959 | Pathogenic | -18.53 | Pathogenic | 0.74 | Disease | 1.00 | Probably damaging | 0.02 | Deleterious | | |
| p.Asp108Pro | VUS | -53.15 | Deleterious | 0.9971 | Pathogenic | -20.06 | Pathogenic | 0.79 | Disease | 0.97 | Probably damaging | 0.01 | Deleterious | | |
| p.Asp108Leu | VUS | -53.15 | Deleterious | 0.9897 | Pathogenic | -19.60 | Pathogenic | 0.81 | Disease | 1.00 | Probably damaging | 0 | Deleterious | | |
| p.Asp108Glu | VUS | -11.70 | Indeterminate | 0.9837 | Pathogenic | -10.69 | Pathogenic | 0.73 | Disease | 0.82 | Possibly damaging | 0.03 | Deleterious | 22.30 | Deleterious |
| p.Asp108Ala | VUS | -53.15 | Deleterious | 0.9849 | Pathogenic | -14.98 | Pathogenic | 0.83 | Disease | 1.00 | Probably damaging | 0.02 | Deleterious | 29.00 | Deleterious |
| p.Asp108Gly | VUS | -53.15 | Deleterious | 0.9533 | Pathogenic | -13.96 | Pathogenic | 0.87 | Disease | 1.00 | Probably damaging | 0.06 | Tolerated | 29.60 | Deleterious |
| p.Asp108Val | VUS | -53.15 | Deleterious | 0.989 | Pathogenic | -17.56 | Pathogenic | 0.90 | Disease | 1.00 | Probably damaging | 0 | Deleterious | 29.30 | Deleterious |
| p.Asp108Tyr | VUS | -53.15 | Deleterious | 0.9811 | Pathogenic | -16.66 | Pathogenic | 0.73 | Disease | 1.00 | Probably damaging | 0.01 | Deleterious | 31.00 | Deleterious |
| p.Asp108Cys | VUS | -53.15 | Deleterious | 0.9907 | Pathogenic | -12.83 | Pathogenic | 0.82 | Disease | 1.00 | Probably damaging | 0 | Deleterious | | |
| p.Asp108Trp | VUS | -53.15 | Deleterious | 0.9992 | Pathogenic | -21.27 | Pathogenic | 0.81 | Disease | 1.00 | Probably damaging | 0 | Deleterious | | |
| p.Asp108Phe | VUS | -53.15 | Deleterious | 0.9984 | Pathogenic | -21.33 | Pathogenic | 0.83 | Disease | 1.00 | Probably damaging | 0 | Deleterious | | |
| p.Ala109Asn | VUS | 0.00 | Neutral | 0.4117 | Ambiguous | -13.00 | Pathogenic | 0.22 | Neutral | 0.91 | Possibly damaging | 0.87 | Tolerated | | |
| p.Ala109Lys | VUS | -0.84 | Neutral | 0.5698 | Pathogenic | -12.24 | Pathogenic | 0.25 | Neutral | 0.99 | Probably damaging | 0.74 | Tolerated | | |
| p.Ala109Thr | VUS | -0.16 | Neutral | 0.1665 | Benign | -9.23 | Pathogenic | 0.20 | Neutral | 0.07 | Benign | 0.49 | Tolerated | 20.30 | Deleterious |
| p.Ala109Arg | VUS | -0.44 | Neutral | 0.345 | Ambiguous | -8.44 | Pathogenic | 0.28 | Neutral | 1.00 | Probably damaging | 0.38 | Tolerated | | |
| p.Ala109Ser | VUS | 0.00 | Neutral | 0.1441 | Benign | -6.48 | Benign | 0.23 | Neutral | 0.25 | Benign | 0.91 | Tolerated | 18.06 | Deleterious |
| p.Ala109Ile | VUS | 0.00 | Neutral | 0.4344 | Ambiguous | -13.28 | Pathogenic | 0.36 | Neutral | 0.99 | Probably damaging | 0.13 | Tolerated | | |
| p.Ala109Met | VUS | 0.00 | Neutral | 0.491 | Ambiguous | -12.83 | Pathogenic | 0.28 | Neutral | 1.00 | Probably damaging | 0.08 | Tolerated | | |
| p.Ala109His | VUS | 0.00 | Neutral | 0.5129 | Ambiguous | -12.47 | Pathogenic | 0.25 | Neutral | 1.00 | Probably damaging | 0.16 | Tolerated | | |
| p.Ala109Gln | VUS | -0.04 | Neutral | 0.3616 | Ambiguous | -10.25 | Pathogenic | 0.21 | Neutral | 1.00 | Probably damaging | 0.42 | Tolerated | | |
| p.Ala109Pro | VUS | -53.15 | Deleterious | 0.376 | Ambiguous | -8.64 | Pathogenic | 0.35 | Neutral | 1.00 | Probably damaging | 0.29 | Tolerated | 22.90 | Deleterious |
| p.Ala109Leu | VUS | -0.83 | Neutral | 0.3332 | Benign | -11.69 | Pathogenic | 0.27 | Neutral | 0.98 | Probably damaging | 0.22 | Tolerated | | |
| p.Ala109Asp | VUS | -0.01 | Neutral | 0.4589 | Ambiguous | -13.00 | Pathogenic | 0.37 | Neutral | 0.98 | Probably damaging | 0.73 | Tolerated | 18.50 | Deleterious |
| p.Ala109Glu | VUS | -1.19 | Neutral | 0.3553 | Ambiguous | -11.40 | Pathogenic | 0.31 | Neutral | 0.99 | Probably damaging | 0.69 | Tolerated | | |
| p.Ala109Gly | VUS | 0.00 | Neutral | 0.142 | Benign | -5.89 | Benign | 0.22 | Neutral | 0.96 | Probably damaging | 0.67 | Tolerated | 16.99 | Deleterious |
| p.Ala109Val | VUS | -0.01 | Neutral | 0.2152 | Benign | -9.47 | Pathogenic | 0.19 | Neutral | 0.92 | Possibly damaging | 0.22 | Tolerated | 18.10 | Deleterious |
| p.Ala109Tyr | VUS | -0.01 | Neutral | 0.6138 | Pathogenic | -15.63 | Pathogenic | 0.39 | Neutral | 1.00 | Probably damaging | 0.08 | Tolerated | | |
| p.Ala109Cys | VUS | 0.00 | Neutral | 0.4761 | Ambiguous | -9.32 | Pathogenic | 0.37 | Neutral | 1.00 | Probably damaging | 0.08 | Tolerated | | |
| p.Ala109Trp | VUS | -0.02 | Neutral | 0.7885 | Pathogenic | -14.26 | Pathogenic | 0.43 | Neutral | 1.00 | Probably damaging | 0.02 | Deleterious | | |
| p.Ala109Phe | VUS | -0.40 | Neutral | 0.5707 | Pathogenic | -15.34 | Pathogenic | 0.40 | Neutral | 1.00 | Probably damaging | 0.06 | Tolerated | | |
| p.Trp110Asn | VUS | -0.41 | Neutral | 0.7983 | Pathogenic | -8.67 | Pathogenic | 0.31 | Neutral | 1.00 | Probably damaging | 0.14 | Tolerated | | |
| p.Trp110Lys | VUS | -1.92 | Neutral | 0.8356 | Pathogenic | -7.22 | Benign | 0.46 | Neutral | 1.00 | Probably damaging | 0.21 | Tolerated | | |
| p.Trp110Thr | VUS | -0.13 | Neutral | 0.699 | Pathogenic | -6.84 | Benign | 0.43 | Neutral | 1.00 | Probably damaging | 0.14 | Tolerated | | |
| p.Trp110Arg | VUS | -0.12 | Neutral | 0.618 | Pathogenic | -5.19 | Benign | 0.23 | Neutral | 1.00 | Probably damaging | 0.07 | Tolerated | 18.20 | Deleterious |
| p.Trp110Ser | VUS | -1.65 | Neutral | 0.4616 | Ambiguous | -5.68 | Benign | 0.24 | Neutral | 1.00 | Probably damaging | 0.16 | Tolerated | 22.30 | Deleterious |
| p.Trp110Ile | VUS | -0.14 | Neutral | 0.8704 | Pathogenic | -8.29 | Pathogenic | 0.47 | Neutral | 1.00 | Probably damaging | 0.03 | Deleterious | | |
| p.Trp110Met | VUS | -1.18 | Neutral | 0.8504 | Pathogenic | -5.64 | Benign | 0.48 | Neutral | 1.00 | Probably damaging | 0.02 | Deleterious | | |
| p.Trp110His | VUS | -4.10 | Neutral | 0.5532 | Ambiguous | -6.71 | Benign | 0.45 | Neutral | 1.00 | Probably damaging | 0.06 | Tolerated | | |
| p.Trp110Gln | VUS | -1.38 | Neutral | 0.7159 | Pathogenic | -6.04 | Benign | 0.31 | Neutral | 1.00 | Probably damaging | 0.25 | Tolerated | | |
| p.Trp110Pro | VUS | -6.95 | Indeterminate | 0.9657 | Pathogenic | -9.39 | Pathogenic | 0.42 | Neutral | 1.00 | Probably damaging | 0.12 | Tolerated | | |
| p.Trp110Leu | VUS | -0.06 | Neutral | 0.6424 | Pathogenic | -4.43 | Benign | 0.31 | Neutral | 1.00 | Probably damaging | 0.05 | Deleterious | 21.60 | Deleterious |
| p.Trp110Asp | VUS | -0.01 | Neutral | 0.7311 | Pathogenic | -7.29 | Benign | 0.29 | Neutral | 1.00 | Probably damaging | 0.54 | Tolerated | | |
| p.Trp110Glu | VUS | -1.76 | Neutral | 0.7096 | Pathogenic | -7.33 | Benign | 0.32 | Neutral | 1.00 | Probably damaging | 1 | Tolerated | | |
| p.Trp110Ala | VUS | -6.88 | Indeterminate | 0.5241 | Ambiguous | -4.22 | Benign | 0.29 | Neutral | 1.00 | Probably damaging | 0.21 | Tolerated | | |
| p.Trp110Gly | VUS | -0.16 | Neutral | 0.4853 | Ambiguous | -5.73 | Benign | 0.25 | Neutral | 0.38 | Benign | 0.11 | Tolerated | 18.78 | Deleterious |
| p.Trp110Val | VUS | -0.02 | Neutral | 0.7336 | Pathogenic | -6.25 | Benign | 0.45 | Neutral | 1.00 | Probably damaging | 0.06 | Tolerated | | |
| p.Trp110Tyr | VUS | -0.11 | Neutral | 0.368 | Ambiguous | -6.32 | Benign | 0.42 | Neutral | 1.00 | Probably damaging | 0.03 | Deleterious | | |
| p.Trp110Cys | VUS | -0.29 | Neutral | 0.8142 | Pathogenic | -5.17 | Benign | 0.45 | Neutral | 1.00 | Probably damaging | 0.01 | Deleterious | 25.85 | Deleterious |
| p.Trp110Phe | VUS | -0.03 | Neutral | 0.2728 | Benign | -2.04 | Benign | 0.33 | Neutral | 1.00 | Probably damaging | 0.02 | Deleterious | | |
| p.Gly111Asn | VUS | -0.20 | Neutral | 0.46 | Ambiguous | -8.63 | Pathogenic | 0.34 | Neutral | 1.00 | Probably damaging | 0 | Deleterious | | |
| p.Gly111Lys | VUS | -4.17 | Neutral | 0.8154 | Pathogenic | -13.08 | Pathogenic | 0.46 | Neutral | 1.00 | Probably damaging | 0 | Deleterious | | |
| p.Gly111Thr | VUS | -33.22 | Indeterminate | 0.774 | Pathogenic | -13.32 | Pathogenic | 0.62 | Disease | 1.00 | Probably damaging | 0 | Deleterious | | |
| p.Gly111Arg | VUS | -1.14 | Neutral | 0.6573 | Pathogenic | -10.53 | Pathogenic | 0.63 | Disease | 1.00 | Probably damaging | 0 | Deleterious | 32.00 | Deleterious |
| p.Gly111Ser | VUS | -0.30 | Neutral | 0.3456 | Ambiguous | -8.72 | Pathogenic | 0.43 | Neutral | 1.00 | Probably damaging | 0 | Deleterious | 31.00 | Deleterious |
| p.Gly111Ile | VUS | -53.15 | Deleterious | 0.9449 | Pathogenic | -17.92 | Pathogenic | 0.66 | Disease | 1.00 | Probably damaging | 0 | Deleterious | | |
| p.Gly111Met | VUS | -0.76 | Neutral | 0.9062 | Pathogenic | -14.61 | Pathogenic | 0.66 | Disease | 1.00 | Probably damaging | 0 | Deleterious | | |
| p.Gly111His | VUS | -6.80 | Indeterminate | 0.6661 | Pathogenic | -10.62 | Pathogenic | 0.63 | Disease | 1.00 | Probably damaging | 0 | Deleterious | | |
| p.Gly111Gln | VUS | -0.03 | Neutral | 0.5589 | Ambiguous | -10.39 | Pathogenic | 0.60 | Disease | 1.00 | Probably damaging | 0 | Deleterious | | |
| p.Gly111Pro | VUS | -53.15 | Deleterious | 0.994 | Pathogenic | -17.33 | Pathogenic | 0.61 | Disease | 1.00 | Probably damaging | 0 | Deleterious | | |
| p.Gly111Leu | VUS | -0.79 | Neutral | 0.8297 | Pathogenic | -13.47 | Pathogenic | 0.65 | Disease | 1.00 | Probably damaging | 0 | Deleterious | | |
| p.Gly111Asp | VUS | -0.28 | Neutral | 0.3488 | Ambiguous | -5.39 | Benign | 0.39 | Neutral | 1.00 | Probably damaging | 0 | Deleterious | 23.30 | Deleterious |
| p.Gly111Glu | VUS | -1.42 | Neutral | 0.4928 | Ambiguous | -8.31 | Pathogenic | 0.65 | Disease | 1.00 | Probably damaging | 0 | Deleterious | | |
| p.Gly111Ala | VUS | -4.33 | Neutral | 0.5207 | Ambiguous | -9.36 | Pathogenic | 0.36 | Neutral | 1.00 | Probably damaging | 0 | Deleterious | 23.00 | Deleterious |
| p.Gly111Val | VUS | -53.15 | Deleterious | 0.8854 | Pathogenic | -13.29 | Pathogenic | 0.71 | Disease | 1.00 | Probably damaging | 0 | Deleterious | 25.40 | Deleterious |

| | | | | | | | | | | | | | | | |
|---|---|---|---|---|---|---|---|---|---|---|---|---|---|---|---|
| p.Gly111Tyr | VUS | -1.17 | Neutral | 0.8014 | Pathogenic | -14.27 | Pathogenic | 0.66 | Disease | 1.00 | Probably damaging | 0 | Deleterious | | |
| p.Gly111Cys | VUS | -0.80 | Neutral | 0.6902 | Pathogenic | -11.37 | Pathogenic | 0.66 | Disease | 1.00 | Probably damaging | 0 | Deleterious | 32.00 | Deleterious |
| p.Gly111Trp | VUS | -0.54 | Neutral | 0.8024 | Pathogenic | -14.45 | Pathogenic | 0.69 | Disease | 1.00 | Probably damaging | 0 | Deleterious | | |
| p.Gly111Phe | VUS | -0.47 | Neutral | 0.9281 | Pathogenic | -15.88 | Pathogenic | 0.66 | Disease | 1.00 | Probably damaging | 0 | Deleterious | | |
| p.Arg112Asn | VUS | -0.21 | Neutral | 0.8545 | Pathogenic | -11.02 | Pathogenic | 0.34 | Neutral | 1.00 | Probably damaging | 0.9 | Tolerated | | |
| p.Arg112Lys | VUS | -2.10 | Neutral | 0.4874 | Ambiguous | -6.79 | Benign | 0.50 | Neutral | 0.99 | Probably damaging | 0.92 | Tolerated | | |
| p.Arg112Thr | VUS | -0.24 | Neutral | 0.8492 | Pathogenic | -12.48 | Pathogenic | 0.67 | Disease | 1.00 | Probably damaging | 0.46 | Tolerated | | |
| p.Arg112Ser | VUS | -0.35 | Neutral | 0.8846 | Pathogenic | -11.11 | Pathogenic | 0.35 | Neutral | 1.00 | Probably damaging | 0.66 | Tolerated | 27.30 | Deleterious |
| p.Arg112Ile | VUS | -1.31 | Neutral | 0.782 | Pathogenic | -15.13 | Pathogenic | 0.68 | Disease | 1.00 | Probably damaging | 0.14 | Tolerated | | |
| p.Arg112Met | VUS | -0.01 | Neutral | 0.8234 | Pathogenic | -12.83 | Pathogenic | 0.67 | Disease | 1.00 | Probably damaging | 0.08 | Tolerated | | |
| p.Arg112His | VUS | -1.24 | Neutral | 0.2973 | Benign | -8.43 | Pathogenic | 0.32 | Neutral | 1.00 | Probably damaging | 0.12 | Tolerated | 27.60 | Deleterious |
| p.Arg112Gln | VUS | -2.70 | Neutral | 0.2994 | Benign | -9.10 | Pathogenic | 0.56 | Disease | 1.00 | Probably damaging | 0.47 | Tolerated | | |
| p.Arg112Pro | VUS | -53.15 | Deleterious | 0.9671 | Pathogenic | -14.73 | Pathogenic | 0.76 | Disease | 1.00 | Probably damaging | 0.22 | Tolerated | 32.00 | Deleterious |
| p.Arg112Leu | VUS | -14.52 | Indeterminate | 0.5683 | Pathogenic | -10.93 | Pathogenic | 0.70 | Disease | 1.00 | Probably damaging | 0.24 | Tolerated | 31.00 | Deleterious |
| p.Arg112Asp | VUS | -7.02 | Indeterminate | 0.915 | Pathogenic | -13.50 | Pathogenic | 0.69 | Disease | 1.00 | Probably damaging | 0.7 | Tolerated | | |
| p.Arg112Glu | VUS | -15.47 | Indeterminate | 0.7571 | Pathogenic | -12.45 | Pathogenic | 0.61 | Disease | 1.00 | Probably damaging | 0.79 | Tolerated | | |
| p.Arg112Ala | VUS | -4.46 | Neutral | 0.8545 | Pathogenic | -11.59 | Pathogenic | 0.35 | Neutral | 1.00 | Probably damaging | 0.71 | Tolerated | | |
| p.Arg112Gly | Pathogenic | -29.42 | Indeterminate | 0.7369 | Pathogenic | -11.75 | Pathogenic | 0.69 | Disease | 1.00 | Probably damaging | 0.35 | Tolerated | 32.00 | Deleterious |
| p.Arg112Val | VUS | -5.85 | Indeterminate | 0.7993 | Pathogenic | -13.18 | Pathogenic | 0.67 | Disease | 1.00 | Probably damaging | 0.22 | Tolerated | | |
| p.Arg112Tyr | VUS | -0.95 | Neutral | 0.7175 | Pathogenic | -13.47 | Pathogenic | 0.61 | Disease | 0.99 | Probably damaging | 0.04 | Deleterious | | |
| p.Arg112Cys | VUS | -16.03 | Indeterminate | 0.5496 | Ambiguous | -10.16 | Pathogenic | 0.42 | Neutral | 1.00 | Probably damaging | 0.05 | Deleterious | 33.00 | Deleterious |
| p.Arg112Trp | VUS | -1.48 | Neutral | 0.4479 | Ambiguous | -12.78 | Pathogenic | 0.69 | Disease | 1.00 | Probably damaging | 0.02 | Deleterious | | |
| p.Arg112Phe | VUS | -4.69 | Neutral | 0.8447 | Pathogenic | -14.05 | Pathogenic | 0.67 | Disease | 0.99 | Probably damaging | 0.04 | Deleterious | | |
| p.Leu113Asn | VUS | -7.95 | Indeterminate | 0.4174 | Ambiguous | -14.49 | Pathogenic | 0.66 | Disease | 1.00 | Probably damaging | 0.03 | Deleterious | | |
| p.Leu113Lys | VUS | -0.38 | Neutral | 0.1897 | Benign | -11.50 | Pathogenic | 0.68 | Disease | 0.98 | Probably damaging | 0.03 | Deleterious | | |
| p.Leu113Thr | VUS | -3.67 | Neutral | 0.211 | Benign | -5.96 | Benign | 0.45 | Neutral | 1.00 | Probably damaging | 0.24 | Tolerated | | |
| p.Leu113Arg | VUS | -5.57 | Neutral | 0.1019 | Benign | -5.81 | Benign | 0.45 | Neutral | 1.00 | Probably damaging | 0.02 | Deleterious | 23.30 | Deleterious |
| p.Leu113Ser | VUS | -0.49 | Neutral | 0.3002 | Benign | -8.80 | Pathogenic | 0.69 | Disease | 1.00 | Probably damaging | 0.22 | Tolerated | | |
| p.Leu113Ile | VUS | -1.77 | Neutral | 0.2739 | Benign | -9.90 | Pathogenic | 0.49 | Neutral | 1.00 | Probably damaging | 0.13 | Tolerated | | |
| p.Leu113Met | VUS | -5.88 | Indeterminate | 0.1896 | Benign | -6.39 | Benign | 0.49 | Neutral | 1.00 | Probably damaging | 0.22 | Tolerated | 26.00 | Deleterious |
| p.Leu113His | VUS | -14.07 | Indeterminate | 0.2142 | Benign | -10.56 | Pathogenic | 0.75 | Disease | 1.00 | Probably damaging | 0.01 | Deleterious | | |
| p.Leu113Gln | VUS | -1.19 | Neutral | 0.1148 | Benign | -8.15 | Pathogenic | 0.47 | Neutral | 1.00 | Probably damaging | 0.03 | Deleterious | 26.40 | Deleterious |
| p.Leu113Pro | VUS | -53.15 | Deleterious | 0.4469 | Ambiguous | -11.83 | Pathogenic | 0.80 | Disease | 1.00 | Probably damaging | 0.03 | Deleterious | 28.20 | Deleterious |
| p.Leu113Asp | VUS | -1.75 | Neutral | 0.6505 | Pathogenic | -13.75 | Pathogenic | 0.70 | Disease | 1.00 | Probably damaging | 0.02 | Deleterious | | |
| p.Leu113Glu | VUS | -0.57 | Neutral | 0.2218 | Benign | -11.71 | Pathogenic | 0.68 | Disease | 1.00 | Probably damaging | 0.02 | Deleterious | | |
| p.Leu113Ala | VUS | -22.19 | Indeterminate | 0.2436 | Benign | -9.41 | Pathogenic | 0.65 | Disease | 1.00 | Probably damaging | 0.08 | Tolerated | | |
| p.Leu113Gly | VUS | -6.02 | Indeterminate | 0.4156 | Ambiguous | -11.03 | Pathogenic | 0.69 | Disease | 1.00 | Probably damaging | 0.02 | Deleterious | | |
| p.Leu113Val | VUS | -0.13 | Neutral | 0.1729 | Benign | -8.54 | Pathogenic | 0.52 | Disease | 0.99 | Probably damaging | 0.11 | Tolerated | 25.20 | Deleterious |
| p.Leu113Tyr | VUS | -2.67 | Neutral | 0.2983 | Benign | -11.35 | Pathogenic | 0.62 | Disease | 0.99 | Probably damaging | 0.02 | Deleterious | | |
| p.Leu113Cys | VUS | -0.85 | Neutral | 0.4819 | Ambiguous | -9.89 | Pathogenic | 0.64 | Disease | 1.00 | Probably damaging | 0.02 | Deleterious | | |
| p.Leu113Trp | VUS | -3.29 | Neutral | 0.2657 | Benign | -9.90 | Pathogenic | 0.69 | Disease | 1.00 | Probably damaging | 0.01 | Deleterious | | |
| p.Leu113Phe | VUS | -13.98 | Indeterminate | 0.1755 | Benign | -9.75 | Pathogenic | 0.55 | Disease | 1.00 | Probably damaging | 0.07 | Tolerated | | |
| p.Pro114Asn | VUS | -53.15 | Deleterious | 0.995 | Pathogenic | -18.60 | Pathogenic | 0.89 | Disease | 1.00 | Probably damaging | 0 | Deleterious | | |
| p.Pro114Lys | VUS | -53.15 | Deleterious | 0.9972 | Pathogenic | -20.74 | Pathogenic | 0.92 | Disease | 1.00 | Probably damaging | 0 | Deleterious | | |
| p.Pro114Thr | Likely pathogenic | -53.15 | Deleterious | 0.8814 | Pathogenic | -14.09 | Pathogenic | 0.93 | Disease | 1.00 | Probably damaging | 0 | Deleterious | 28.90 | Deleterious |
| p.Pro114Arg | VUS | -53.15 | Deleterious | 0.973 | Pathogenic | -15.01 | Pathogenic | 0.94 | Disease | 1.00 | Probably damaging | 0 | Deleterious | 31.00 | Deleterious |
| p.Pro114Ser | VUS | -9.07 | Indeterminate | 0.8619 | Pathogenic | -9.87 | Pathogenic | 0.91 | Disease | 1.00 | Probably damaging | 0 | Deleterious | 29.90 | Deleterious |
| p.Pro114Ile | VUS | -53.15 | Deleterious | 0.9784 | Pathogenic | -19.22 | Pathogenic | 0.88 | Disease | 1.00 | Probably damaging | 0 | Deleterious | | |
| p.Pro114Met | VUS | -53.15 | Deleterious | 0.9889 | Pathogenic | -17.99 | Pathogenic | 0.91 | Disease | 1.00 | Probably damaging | 0 | Deleterious | | |
| p.Pro114His | Likely pathogenic | -53.15 | Deleterious | 0.9886 | Pathogenic | -15.83 | Pathogenic | 0.91 | Disease | 1.00 | Probably damaging | 0 | Deleterious | 31.00 | Deleterious |
| p.Pro114Gln | VUS | -53.15 | Deleterious | 0.9769 | Pathogenic | -17.49 | Pathogenic | 0.90 | Disease | 1.00 | Probably damaging | 0 | Deleterious | | |
| p.Pro114Leu | VUS | -53.15 | Deleterious | 0.8805 | Pathogenic | -15.28 | Pathogenic | 0.93 | Disease | 1.00 | Probably damaging | 0 | Deleterious | 32.00 | Deleterious |
| p.Pro114Asp | VUS | -53.15 | Deleterious | 0.9963 | Pathogenic | -18.40 | Pathogenic | 0.91 | Disease | 1.00 | Probably damaging | 0 | Deleterious | | |
| p.Pro114Glu | VUS | -53.15 | Deleterious | 0.9878 | Pathogenic | -17.85 | Pathogenic | 0.87 | Disease | 1.00 | Probably damaging | 0 | Deleterious | | |
| p.Pro114Ala | VUS | -8.63 | Indeterminate | 0.4367 | Ambiguous | -5.69 | Benign | 0.81 | Disease | 1.00 | Probably damaging | 0 | Deleterious | 27.90 | Deleterious |
| p.Pro114Gly | VUS | -53.15 | Deleterious | 0.9382 | Pathogenic | -11.20 | Pathogenic | 0.86 | Disease | 1.00 | Probably damaging | 0 | Deleterious | | |
| p.Pro114Val | VUS | -32.81 | Indeterminate | 0.9005 | Pathogenic | -13.48 | Pathogenic | 0.85 | Disease | 1.00 | Probably damaging | 0 | Deleterious | | |
| p.Pro114Tyr | VUS | -53.15 | Deleterious | 0.9981 | Pathogenic | -18.72 | Pathogenic | 0.91 | Disease | 1.00 | Probably damaging | 0 | Deleterious | | |
| p.Pro114Cys | VUS | -7.47 | Indeterminate | 0.9625 | Pathogenic | -10.61 | Pathogenic | 0.90 | Disease | 1.00 | Probably damaging | 0 | Deleterious | | |
| p.Pro114Trp | VUS | -53.15 | Deleterious | 0.9989 | Pathogenic | -17.70 | Pathogenic | 0.90 | Disease | 1.00 | Probably damaging | 0 | Deleterious | | |
| p.Pro114Phe | VUS | -53.15 | Deleterious | 0.9984 | Pathogenic | -16.89 | Pathogenic | 0.90 | Disease | 1.00 | Probably damaging | 0 | Deleterious | | |
| p.Val115Asn | VUS | -2.41 | Neutral | 0.5598 | Ambiguous | -12.03 | Pathogenic | 0.69 | Disease | 1.00 | Probably damaging | 0.01 | Deleterious | | |
| p.Val115Lys | VUS | -3.03 | Neutral | 0.4431 | Ambiguous | -9.06 | Pathogenic | 0.75 | Disease | 1.00 | Probably damaging | 0.01 | Deleterious | | |
| p.Val115Thr | VUS | -1.57 | Neutral | 0.2098 | Benign | -6.92 | Benign | 0.53 | Disease | 1.00 | Probably damaging | 0.03 | Deleterious | | |
| p.Val115Arg | VUS | -21.93 | Indeterminate | 0.2736 | Benign | -6.10 | Benign | 0.75 | Disease | 1.00 | Probably damaging | 0.01 | Deleterious | | |
| p.Val115Ser | VUS | -4.66 | Neutral | 0.2381 | Benign | -6.40 | Benign | 0.62 | Disease | 1.00 | Probably damaging | 0.01 | Deleterious | | |
| p.Val115Ile | VUS | -2.54 | Neutral | 0.1129 | Benign | -5.62 | Benign | 0.11 | Neutral | 0.45 | Benign | 0.72 | Tolerated | | |
| p.Val115Met | VUS | -1.44 | Neutral | 0.2528 | Benign | -7.13 | Benign | 0.44 | Neutral | 1.00 | Probably damaging | 0.05 | Deleterious | 26.30 | Deleterious |
| p.Val115His | VUS | -7.13 | Indeterminate | 0.6119 | Pathogenic | -8.48 | Pathogenic | 0.67 | Disease | 1.00 | Probably damaging | 0 | Deleterious | | |
| p.Val115Gln | VUS | -19.06 | Indeterminate | 0.3317 | Benign | -5.81 | Benign | 0.68 | Disease | 1.00 | Probably damaging | 0.01 | Deleterious | | |
| p.Val115Pro | VUS | -33.13 | Indeterminate | 0.5724 | Pathogenic | -9.79 | Pathogenic | 0.66 | Disease | 1.00 | Probably damaging | 0.01 | Deleterious | | |
| p.Val115Leu | VUS | -2.89 | Neutral | 0.2356 | Benign | -5.26 | Benign | 0.11 | Neutral | 0.01 | Benign | 0.97 | Tolerated | 22.20 | Deleterious |
| p.Val115Asp | VUS | -9.37 | Indeterminate | 0.5529 | Ambiguous | -10.51 | Pathogenic | 0.82 | Disease | 1.00 | Probably damaging | 0.01 | Deleterious | | |
| p.Val115Glu | VUS | -9.20 | Indeterminate | 0.3085 | Benign | -4.96 | Benign | 0.78 | Disease | 1.00 | Probably damaging | 0.01 | Deleterious | 29.60 | Deleterious |
| p.Val115Ala | VUS | -22.86 | Indeterminate | 0.1017 | Benign | -2.70 | Benign | 0.24 | Neutral | 1.00 | Probably damaging | 0.03 | Deleterious | 27.50 | Deleterious |
| p.Val115Gly | VUS | -3.24 | Neutral | 0.1657 | Benign | -5.53 | Benign | 0.59 | Disease | 1.00 | Probably damaging | 0.01 | Deleterious | 29.20 | Deleterious |
| p.Val115Tyr | VUS | -2.24 | Neutral | 0.6985 | Pathogenic | -11.15 | Pathogenic | 0.63 | Disease | 1.00 | Probably damaging | 0.25 | Tolerated | | |
| p.Val115Cys | VUS | -2.73 | Neutral | 0.6361 | Pathogenic | -6.77 | Benign | 0.59 | Disease | 1.00 | Probably damaging | 0.01 | Deleterious | | |
| p.Val115Trp | VUS | -3.56 | Neutral | 0.7798 | Pathogenic | -8.45 | Pathogenic | 0.59 | Disease | 1.00 | Probably damaging | 0.01 | Deleterious | | |
| p.Val115Phe | VUS | -3.04 | Neutral | 0.342 | Ambiguous | -10.76 | Pathogenic | 0.65 | Disease | 0.99 | Probably damaging | 0.05 | Deleterious | | |
| p.Asp116Asn | VUS | -3.49 | Neutral | 0.4 | Ambiguous | -10.46 | Pathogenic | 0.65 | Disease | 0.98 | Probably damaging | 0.59 | Tolerated | 32.00 | Deleterious |
| p.Asp116Lys | VUS | -0.50 | Neutral | 0.8451 | Pathogenic | -11.20 | Pathogenic | 0.74 | Disease | 1.00 | Probably damaging | 0.18 | Tolerated | | |
| p.Asp116Thr | VUS | -5.85 | Indeterminate | 0.6412 | Pathogenic | -9.90 | Pathogenic | 0.70 | Disease | 0.96 | Probably damaging | 0.13 | Tolerated | | |
| p.Asp116Arg | VUS | -4.81 | Neutral | 0.7232 | Pathogenic | -8.52 | Pathogenic | 0.73 | Disease | 1.00 | Probably damaging | 0.09 | Tolerated | | |
| p.Asp116Ser | VUS | -1.21 | Neutral | 0.4038 | Ambiguous | -9.67 | Pathogenic | 0.64 | Disease | 1.00 | Probably damaging | 0.29 | Tolerated | | |
| p.Asp116Ile | VUS | -3.94 | Neutral | 0.8019 | Pathogenic | -13.22 | Pathogenic | 0.69 | Disease | 0.45 | Benign | 0.02 | Deleterious | | |
| p.Asp116Met | VUS | -0.33 | Neutral | 0.8692 | Pathogenic | -10.86 | Pathogenic | 0.70 | Disease | 1.00 | Probably damaging | 0.01 | Deleterious | | |
| p.Asp116His | VUS | -1.16 | Neutral | 0.3546 | Ambiguous | -8.67 | Pathogenic | 0.30 | Neutral | 1.00 | Probably damaging | 0.49 | Tolerated | 25.20 | Deleterious |
| p.Asp116Gln | VUS | -1.92 | Neutral | 0.5174 | Ambiguous | -5.49 | Benign | 0.64 | Disease | 1.00 | Probably damaging | 0.14 | Tolerated | | |
| p.Asp116Pro | VUS | -53.15 | Deleterious | 0.9689 | Pathogenic | -13.02 | Pathogenic | 0.70 | Disease | 0.92 | Possibly damaging | 0.11 | Tolerated | | |
| p.Asp116Leu | VUS | -2.83 | Neutral | 0.6897 | Pathogenic | -10.07 | Pathogenic | 0.71 | Disease | 0.93 | Possibly damaging | 0.04 | Deleterious | | |
| p.Asp116Glu | VUS | -1.73 | Neutral | 0.319 | Benign | -4.80 | Benign | 0.28 | Neutral | 0.79 | Possibly damaging | 0.32 | Tolerated | 24.95 | Deleterious |
| p.Asp116Ala | VUS | -2.29 | Neutral | 0.4161 | Ambiguous | -7.45 | Benign | 0.75 | Disease | 0.97 | Probably damaging | 0.14 | Tolerated | 32.00 | Deleterious |
| p.Asp116Gly | VUS | -7.40 | Indeterminate | 0.5678 | Pathogenic | -8.44 | Pathogenic | 0.77 | Disease | 1.00 | Probably damaging | 0.37 | Tolerated | 32.00 | Deleterious |
| p.Asp116Val | VUS | -2.63 | Neutral | 0.5947 | Pathogenic | -9.75 | Pathogenic | 0.77 | Disease | 0.63 | Possibly damaging | 0.03 | Deleterious | 32.00 | Deleterious |
| p.Asp116Tyr | VUS | -0.83 | Neutral | 0.5072 | Ambiguous | -12.24 | Pathogenic | 0.78 | Disease | 1.00 | Probably damaging | 0.06 | Tolerated | 32.00 | Deleterious |
| p.Asp116Cys | VUS | -0.88 | Neutral | 0.8589 | Pathogenic | -10.57 | Pathogenic | 0.69 | Disease | 1.00 | Probably damaging | 0.03 | Deleterious | | |
| p.Asp116Trp | VUS | -8.45 | Indeterminate | 0.9209 | Pathogenic | -12.10 | Pathogenic | 0.70 | Disease | 1.00 | Probably damaging | 0.01 | Deleterious | | |
| p.Asp116Phe | VUS | -3.01 | Neutral | 0.7835 | Pathogenic | -12.89 | Pathogenic | 0.69 | Disease | 0.99 | Probably damaging | 0.03 | Deleterious | | |
| p.Leu117Asn | VUS | -1.87 | Neutral | 0.9031 | Pathogenic | -12.87 | Pathogenic | 0.70 | Disease | 1.00 | Probably damaging | 0 | Deleterious | | |
| p.Leu117Lys | VUS | -1.02 | Neutral | 0.7514 | Pathogenic | -9.51 | Pathogenic | 0.74 | Disease | 0.99 | Probably damaging | 0 | Deleterious | | |
| p.Leu117Thr | VUS | -0.87 | Neutral | 0.6291 | Pathogenic | -8.55 | Pathogenic | 0.72 | Disease | 1.00 | Probably damaging | 0 | Deleterious | | |
| p.Leu117Arg | VUS | -1.66 | Neutral | 0.4913 | Ambiguous | -7.85 | Pathogenic | 0.87 | Disease | 1.00 | Probably damaging | 0 | Deleterious | 32.00 | Deleterious |
| p.Leu117Ser | VUS | -1.67 | Neutral | 0.8948 | Pathogenic | -9.99 | Pathogenic | 0.77 | Disease | 1.00 | Probably damaging | 0 | Deleterious | | |
| p.Leu117Ile | VUS | -1.14 | Neutral | 0.1638 | Benign | -8.89 | Pathogenic | 0.27 | Neutral | 1.00 | Probably damaging | 0.14 | Tolerated | | |
| p.Leu117Met | VUS | -2.16 | Neutral | 0.3229 | Benign | -7.59 | Pathogenic | 0.47 | Neutral | 1.00 | Probably damaging | 0.16 | Tolerated | 26.10 | Deleterious |
| p.Leu117His | VUS | -0.50 | Neutral | 0.6986 | Pathogenic | -9.81 | Pathogenic | 0.75 | Disease | 1.00 | Probably damaging | 0 | Deleterious | | |
| p.Leu117Gln | VUS | -3.86 | Neutral | 0.6392 | Pathogenic | -9.14 | Pathogenic | 0.80 | Disease | 1.00 | Probably damaging | 0 | Deleterious | 31.00 | Deleterious |
| p.Leu117Pro | VUS | -33.22 | Indeterminate | 0.9393 | Pathogenic | -12.56 | Pathogenic | 0.88 | Disease | 1.00 | Probably damaging | 0 | Deleterious | 32.00 | Deleterious |
| p.Leu117Asp | VUS | -0.74 | Neutral | 0.9488 | Pathogenic | -12.70 | Pathogenic | 0.71 | Disease | 1.00 | Probably damaging | 0 | Deleterious | | |
| p.Leu117Glu | VUS | -1.45 | Neutral | 0.7894 | Pathogenic | -10.29 | Pathogenic | 0.73 | Disease | 1.00 | Probably damaging | 0 | Deleterious | | |

| | | | | | | | | | | | | | | | |
|---|---|---|---|---|---|---|---|---|---|---|---|---|---|---|---|
| p.Leu117Ala | VUS | -3.34 | Neutral | 0.6184 | Pathogenic | -7.30 | Benign | 0.64 | Disease | 1.00 | Probably damaging | 0.01 | Deleterious | | |
| p.Leu117Gly | VUS | -1.70 | Neutral | 0.8729 | Pathogenic | -10.95 | Pathogenic | 0.71 | Disease | 1.00 | Probably damaging | 0 | Deleterious | | |
| p.Leu117Val | VUS | -1.09 | Neutral | 0.1608 | Benign | -5.80 | Benign | 0.18 | Neutral | 1.00 | Probably damaging | 0.02 | Deleterious | 23.80 | Deleterious |
| p.Leu117Tyr | VUS | -0.42 | Neutral | 0.7173 | Pathogenic | -10.15 | Pathogenic | 0.63 | Disease | 1.00 | Probably damaging | 0 | Deleterious | | |
| p.Leu117Cys | VUS | -2.13 | Neutral | 0.8485 | Pathogenic | -9.32 | Pathogenic | 0.65 | Disease | 1.00 | Probably damaging | 0 | Deleterious | | |
| p.Leu117Trp | VUS | -1.72 | Neutral | 0.6017 | Pathogenic | -6.36 | Benign | 0.71 | Disease | 1.00 | Probably damaging | 0 | Deleterious | | |
| p.Leu117Phe | VUS | -1.86 | Neutral | 0.4215 | Ambiguous | -9.89 | Pathogenic | 0.51 | Disease | 1.00 | Probably damaging | 0.02 | Deleterious | | |
| p.Ala118Asn | VUS | -32.02 | Indeterminate | 0.9907 | Pathogenic | -18.64 | Pathogenic | 0.85 | Disease | 1.00 | Probably damaging | 0.01 | Deleterious | | |
| p.Ala118Lys | VUS | -33.22 | Indeterminate | 0.9948 | Pathogenic | -19.83 | Pathogenic | 0.89 | Disease | 1.00 | Probably damaging | 0.01 | Deleterious | | |
| p.Ala118Thr | VUS | -4.89 | Neutral | 0.8464 | Pathogenic | -12.28 | Pathogenic | 0.90 | Disease | 1.00 | Probably damaging | 0.02 | Deleterious | 32.00 | Deleterious |
| p.Ala118Arg | VUS | -27.39 | Indeterminate | 0.9534 | Pathogenic | -15.22 | Pathogenic | 0.87 | Disease | 1.00 | Probably damaging | 0.01 | Deleterious | | |
| p.Ala118Ser | VUS | -6.37 | Indeterminate | 0.418 | Ambiguous | -10.49 | Pathogenic | 0.80 | Disease | 0.97 | Probably damaging | 0.04 | Deleterious | 29.80 | Deleterious |
| p.Ala118Ile | VUS | -12.28 | Indeterminate | 0.9817 | Pathogenic | -17.46 | Pathogenic | 0.89 | Disease | 1.00 | Probably damaging | 0.01 | Deleterious | | |
| p.Ala118Met | VUS | -26.26 | Indeterminate | 0.9763 | Pathogenic | -17.51 | Pathogenic | 0.83 | Disease | 1.00 | Probably damaging | 0 | Deleterious | | |
| p.Ala118His | VUS | -33.22 | Indeterminate | 0.9896 | Pathogenic | -18.95 | Pathogenic | 0.87 | Disease | 1.00 | Probably damaging | 0 | Deleterious | | |
| p.Ala118Gln | VUS | -29.56 | Indeterminate | 0.966 | Pathogenic | -18.83 | Pathogenic | 0.89 | Disease | 1.00 | Probably damaging | 0.01 | Deleterious | | |
| p.Ala118Pro | VUS | -33.04 | Indeterminate | 0.92 | Pathogenic | -13.05 | Pathogenic | 0.90 | Disease | 1.00 | Probably damaging | 0.02 | Deleterious | 31.00 | Deleterious |
| p.Ala118Leu | VUS | -10.18 | Indeterminate | 0.8613 | Pathogenic | -16.67 | Pathogenic | 0.85 | Disease | 1.00 | Probably damaging | 0.01 | Deleterious | | |
| p.Ala118Asp | VUS | -28.32 | Indeterminate | 0.98 | Pathogenic | -16.04 | Pathogenic | 0.94 | Disease | 1.00 | Probably damaging | 0.01 | Deleterious | 32.00 | Deleterious |
| p.Ala118Glu | VUS | -31.29 | Indeterminate | 0.9708 | Pathogenic | -16.79 | Pathogenic | 0.91 | Disease | 1.00 | Probably damaging | 0.01 | Deleterious | | |
| p.Ala118Gly | VUS | -2.90 | Neutral | 0.3954 | Ambiguous | -10.74 | Pathogenic | 0.82 | Disease | 1.00 | Probably damaging | 0.05 | Deleterious | 32.00 | Deleterious |
| p.Ala118Val | VUS | -6.66 | Indeterminate | 0.8495 | Pathogenic | -12.26 | Pathogenic | 0.85 | Disease | 1.00 | Probably damaging | 0.02 | Deleterious | 32.00 | Deleterious |
| p.Ala118Tyr | VUS | -32.85 | Indeterminate | 0.9941 | Pathogenic | -19.07 | Pathogenic | 0.89 | Disease | 1.00 | Probably damaging | 0 | Deleterious | | |
| p.Ala118Cys | VUS | -2.10 | Neutral | 0.8877 | Pathogenic | -12.96 | Pathogenic | 0.80 | Disease | 1.00 | Probably damaging | 0.16 | Tolerated | | |
| p.Ala118Trp | VUS | -33.22 | Indeterminate | 0.9968 | Pathogenic | -19.92 | Pathogenic | 0.90 | Disease | 1.00 | Probably damaging | 0 | Deleterious | | |
| p.Ala118Phe | VUS | -31.19 | Indeterminate | 0.9903 | Pathogenic | -19.66 | Pathogenic | 0.89 | Disease | 1.00 | Probably damaging | 0 | Deleterious | | |
| p.Glu119Asn | VUS | -1.46 | Neutral | 0.5101 | Ambiguous | -10.85 | Pathogenic | 0.39 | Neutral | 1.00 | Probably damaging | 0.29 | Tolerated | | |
| p.Glu119Lys | VUS | -0.30 | Neutral | 0.2273 | Benign | -9.38 | Pathogenic | 0.25 | Neutral | 1.00 | Probably damaging | 0.55 | Tolerated | 19.51 | Deleterious |
| p.Glu119Thr | VUS | -0.06 | Neutral | 0.294 | Benign | -7.87 | Pathogenic | 0.42 | Neutral | 1.00 | Probably damaging | 0.57 | Tolerated | | |
| p.Glu119Arg | VUS | -10.02 | Indeterminate | 0.1837 | Benign | -7.03 | Benign | 0.25 | Neutral | 1.00 | Probably damaging | 0.32 | Tolerated | | |
| p.Glu119Ser | VUS | 0.00 | Neutral | 0.3272 | Benign | -9.06 | Pathogenic | 0.43 | Neutral | 1.00 | Probably damaging | 0.45 | Tolerated | | |
| p.Glu119Ile | VUS | -0.05 | Neutral | 0.5279 | Ambiguous | -9.42 | Pathogenic | 0.31 | Neutral | 1.00 | Probably damaging | 0.51 | Tolerated | | |
| p.Glu119Met | VUS | -4.89 | Neutral | 0.5649 | Pathogenic | -9.59 | Pathogenic | 0.49 | Neutral | 1.00 | Probably damaging | 0.15 | Tolerated | | |
| p.Glu119His | VUS | -30.83 | Indeterminate | 0.453 | Ambiguous | -9.34 | Pathogenic | 0.42 | Neutral | 1.00 | Probably damaging | 0.11 | Tolerated | | |
| p.Glu119Gln | VUS | -23.79 | Indeterminate | 0.1483 | Benign | -6.81 | Benign | 0.17 | Neutral | 0.99 | Probably damaging | 0.37 | Tolerated | 17.96 | Deleterious |
| p.Glu119Pro | VUS | -33.21 | Indeterminate | 0.945 | Pathogenic | -13.45 | Pathogenic | 0.47 | Neutral | 1.00 | Probably damaging | 0.23 | Tolerated | | |
| p.Glu119Leu | VUS | -1.13 | Neutral | 0.3708 | Ambiguous | -8.04 | Pathogenic | 0.29 | Neutral | 1.00 | Probably damaging | 0.49 | Tolerated | | |
| p.Glu119Asp | VUS | -9.98 | Indeterminate | 0.2843 | Benign | -7.45 | Benign | 0.33 | Neutral | 0.99 | Probably damaging | 0.37 | Tolerated | 15.22 | Deleterious |
| p.Glu119Ala | VUS | -0.01 | Neutral | 0.1304 | Benign | -6.79 | Benign | 0.16 | Neutral | 1.00 | Probably damaging | 0.84 | Tolerated | 18.66 | Deleterious |
| p.Glu119Gly | VUS | -0.65 | Neutral | 0.1783 | Benign | -8.90 | Pathogenic | 0.25 | Neutral | 1.00 | Probably damaging | 0.22 | Tolerated | 22.20 | Deleterious |
| p.Glu119Val | VUS | -5.91 | Indeterminate | 0.2797 | Benign | -6.98 | Benign | 0.22 | Neutral | 1.00 | Probably damaging | 1 | Tolerated | 21.70 | Deleterious |
| p.Glu119Tyr | VUS | 0.00 | Neutral | 0.6978 | Pathogenic | -12.23 | Pathogenic | 0.42 | Neutral | 1.00 | Probably damaging | 0.1 | Tolerated | | |
| p.Glu119Cys | VUS | 0.00 | Neutral | 0.8272 | Pathogenic | -10.09 | Pathogenic | 0.44 | Neutral | 1.00 | Probably damaging | 0.11 | Tolerated | | |
| p.Glu119Trp | VUS | -31.32 | Indeterminate | 0.8444 | Pathogenic | -11.54 | Pathogenic | 0.49 | Neutral | 1.00 | Probably damaging | 0.03 | Deleterious | | |
| p.Glu119Phe | VUS | -33.21 | Indeterminate | 0.733 | Pathogenic | -12.37 | Pathogenic | 0.43 | Neutral | 1.00 | Probably damaging | 0.12 | Tolerated | | |
| p.Glu120Asn | VUS | -1.82 | Neutral | 0.6455 | Pathogenic | -8.71 | Pathogenic | 0.46 | Neutral | 1.00 | Probably damaging | 0.2 | Tolerated | | |
| p.Glu120Lys | VUS | -1.90 | Neutral | 0.5939 | Pathogenic | -7.47 | Benign | 0.53 | Disease | 0.79 | Possibly damaging | 0.54 | Tolerated | 24.20 | Deleterious |
| p.Glu120Thr | VUS | -1.40 | Neutral | 0.4767 | Ambiguous | -6.27 | Benign | 0.50 | Neutral | 1.00 | Probably damaging | 0.24 | Tolerated | | |
| p.Glu120Arg | VUS | -5.70 | Neutral | 0.4604 | Ambiguous | -6.28 | Benign | 0.46 | Neutral | 0.91 | Possibly damaging | 0.3 | Tolerated | | |
| p.Glu120Ser | VUS | -1.67 | Neutral | 0.4157 | Ambiguous | -5.36 | Benign | 0.46 | Neutral | 0.99 | Probably damaging | 0.3 | Tolerated | | |
| p.Glu120Ile | VUS | -5.03 | Neutral | 0.6566 | Pathogenic | -7.99 | Pathogenic | 0.49 | Neutral | 0.99 | Probably damaging | 0.19 | Tolerated | | |
| p.Glu120Met | VUS | -12.94 | Indeterminate | 0.7051 | Pathogenic | -6.86 | Benign | 0.55 | Disease | 0.99 | Probably damaging | 0.04 | Deleterious | | |
| p.Glu120His | VUS | -2.96 | Neutral | 0.6129 | Pathogenic | -6.68 | Benign | 0.46 | Neutral | 0.98 | Probably damaging | 0.06 | Tolerated | | |
| p.Glu120Gln | VUS | -4.15 | Neutral | 0.2187 | Benign | -3.82 | Benign | 0.28 | Neutral | 0.17 | Benign | 0.31 | Tolerated | 21.90 | Deleterious |
| p.Glu120Pro | VUS | -4.57 | Neutral | 0.9068 | Pathogenic | -10.42 | Pathogenic | 0.56 | Disease | 1.00 | Probably damaging | 0.12 | Tolerated | | |
| p.Glu120Leu | VUS | -1.27 | Neutral | 0.6466 | Pathogenic | -6.07 | Benign | 0.50 | Neutral | 0.94 | Possibly damaging | 0.13 | Tolerated | | |
| p.Glu120Asp | VUS | -2.28 | Neutral | 0.2992 | Benign | -5.91 | Benign | 0.26 | Neutral | 0.75 | Possibly damaging | 0.45 | Tolerated | 21.85 | Deleterious |
| p.Glu120Ala | VUS | -0.79 | Neutral | 0.2356 | Benign | -4.11 | Benign | 0.30 | Neutral | 0.96 | Possibly damaging | 0.32 | Tolerated | 22.90 | Deleterious |
| p.Glu120Gly | VUS | -1.95 | Neutral | 0.3373 | Benign | -6.30 | Benign | 0.46 | Neutral | 1.00 | Probably damaging | 0.16 | Tolerated | 25.20 | Deleterious |
| p.Glu120Val | VUS | -1.61 | Neutral | 0.482 | Ambiguous | -6.19 | Benign | 0.50 | Disease | 0.95 | Possibly damaging | 0.12 | Tolerated | 26.50 | Deleterious |
| p.Glu120Tyr | VUS | -11.75 | Indeterminate | 0.7664 | Pathogenic | -8.90 | Pathogenic | 0.49 | Neutral | 1.00 | Probably damaging | 0.02 | Deleterious | | |
| p.Glu120Cys | VUS | -1.14 | Neutral | 0.9161 | Pathogenic | -8.29 | Pathogenic | 0.54 | Disease | 1.00 | Probably damaging | 0.02 | Deleterious | | |
| p.Glu120Trp | VUS | -2.61 | Neutral | 0.9213 | Pathogenic | -10.28 | Pathogenic | 0.43 | Neutral | 1.00 | Probably damaging | 0.01 | Deleterious | | |
| p.Glu120Phe | VUS | -0.12 | Neutral | 0.8696 | Pathogenic | -9.00 | Pathogenic | 0.52 | Disease | 1.00 | Probably damaging | 0.02 | Deleterious | | |
| p.Leu121Asn | VUS | -4.57 | Neutral | 0.2029 | Benign | 0.79 | Benign | 0.18 | Neutral | 0.05 | Benign | 0.38 | Tolerated | | |
| p.Leu121Lys | VUS | -1.77 | Neutral | 0.147 | Benign | -2.38 | Benign | 0.20 | Neutral | 0.00 | Benign | 0.79 | Tolerated | | |
| p.Leu121Thr | VUS | -1.63 | Neutral | 0.1938 | Benign | -4.38 | Benign | 0.24 | Neutral | 0.01 | Benign | 0.47 | Tolerated | | |
| p.Leu121Arg | VUS | -12.00 | Indeterminate | 0.0651 | Benign | 1.68 | Benign | 0.11 | Neutral | 0.00 | Benign | 0.85 | Tolerated | 11.52 | Neutral |
| p.Leu121Ser | VUS | -2.80 | Neutral | 0.2532 | Benign | -0.34 | Benign | 0.09 | Neutral | 0.02 | Benign | 0.48 | Tolerated | | |
| p.Leu121Ile | VUS | -1.47 | Neutral | 0.19 | Benign | -8.56 | Pathogenic | 0.18 | Neutral | 0.00 | Benign | 0.4 | Tolerated | | |
| p.Leu121Met | VUS | -53.15 | Deleterious | 0.1454 | Benign | -5.63 | Benign | 0.22 | Neutral | 0.00 | Benign | 0.23 | Tolerated | 21.30 | Deleterious |
| p.Leu121His | VUS | -0.28 | Neutral | 0.1246 | Benign | 0.35 | Benign | 0.15 | Neutral | 0.18 | Benign | 0.64 | Tolerated | | |
| p.Leu121Gln | VUS | -1.06 | Neutral | 0.0787 | Benign | -1.63 | Benign | 0.11 | Neutral | 0.00 | Benign | 0.55 | Tolerated | 13.36 | Neutral |
| p.Leu121Pro | VUS | -0.56 | Neutral | 0.7069 | Pathogenic | -7.54 | Pathogenic | 0.23 | Neutral | 0.07 | Benign | 0.26 | Tolerated | 14.80 | Neutral |
| p.Leu121Asp | VUS | 0.00 | Neutral | 0.4718 | Ambiguous | -5.15 | Benign | 0.29 | Neutral | 0.05 | Benign | 0.27 | Tolerated | | |
| p.Leu121Glu | VUS | -3.77 | Neutral | 0.1833 | Benign | -2.75 | Benign | 0.18 | Neutral | 0.01 | Benign | 0.76 | Tolerated | | |
| p.Leu121Ala | VUS | -2.68 | Neutral | 0.1338 | Benign | -0.09 | Benign | 0.22 | Neutral | 0.01 | Benign | 0.59 | Tolerated | | |
| p.Leu121Gly | VUS | -3.55 | Neutral | 0.168 | Benign | 0.18 | Benign | 0.15 | Neutral | 0.02 | Benign | 0.38 | Tolerated | | |
| p.Leu121Val | VUS | -0.04 | Neutral | 0.1629 | Benign | -5.47 | Benign | 0.14 | Neutral | 0.00 | Benign | 0.51 | Tolerated | 17.90 | Deleterious |
| p.Leu121Tyr | VUS | -0.11 | Neutral | 0.2306 | Benign | -1.88 | Benign | 0.24 | Neutral | 0.00 | Benign | 1 | Tolerated | | |
| p.Leu121Cys | VUS | -0.25 | Neutral | 0.2808 | Benign | -0.78 | Benign | 0.25 | Neutral | 0.56 | Possibly damaging | 0.17 | Tolerated | | |
| p.Leu121Trp | VUS | -3.11 | Neutral | 0.1596 | Benign | -5.17 | Benign | 0.26 | Neutral | 0.26 | Benign | 0.18 | Tolerated | | |
| p.Leu121Phe | VUS | -0.56 | Neutral | 0.1576 | Benign | -2.40 | Benign | 0.12 | Neutral | 0.00 | Benign | 0.65 | Tolerated | | |
| p.Gly122Asn | VUS | -0.06 | Neutral | 0.6676 | Pathogenic | -9.46 | Pathogenic | 0.41 | Neutral | 1.00 | Probably damaging | 0.27 | Tolerated | | |
| p.Gly122Lys | VUS | -0.45 | Neutral | 0.8542 | Pathogenic | -8.95 | Pathogenic | 0.64 | Disease | 1.00 | Probably damaging | 0.05 | Deleterious | | |
| p.Gly122Thr | VUS | -0.11 | Neutral | 0.7899 | Pathogenic | -9.54 | Pathogenic | 0.43 | Neutral | 1.00 | Probably damaging | 0.04 | Deleterious | | |
| p.Gly122Arg | VUS | -2.36 | Neutral | 0.6425 | Pathogenic | -6.20 | Benign | 0.66 | Disease | 1.00 | Probably damaging | 0.03 | Deleterious | 25.80 | Deleterious |
| p.Gly122Ser | VUS | -0.02 | Neutral | 0.4055 | Ambiguous | -8.12 | Pathogenic | 0.55 | Disease | 1.00 | Probably damaging | 0.1 | Tolerated | 25.80 | Deleterious |
| p.Gly122Ile | VUS | -3.72 | Neutral | 0.9444 | Pathogenic | -12.62 | Pathogenic | 0.68 | Disease | 1.00 | Probably damaging | 0.01 | Deleterious | | |
| p.Gly122Met | VUS | -0.24 | Neutral | 0.9514 | Pathogenic | -6.51 | Benign | 0.68 | Disease | 1.00 | Probably damaging | 0 | Deleterious | | |
| p.Gly122His | VUS | -0.06 | Neutral | 0.7213 | Pathogenic | -7.17 | Benign | 0.62 | Disease | 1.00 | Probably damaging | 0.03 | Deleterious | | |
| p.Gly122Gln | VUS | -0.10 | Neutral | 0.6852 | Pathogenic | -6.15 | Benign | 0.59 | Disease | 1.00 | Probably damaging | 0.03 | Deleterious | | |
| p.Gly122Pro | VUS | -1.75 | Neutral | 0.972 | Pathogenic | -10.91 | Pathogenic | 0.43 | Neutral | 1.00 | Probably damaging | 0.19 | Tolerated | | |
| p.Gly122Leu | VUS | -0.96 | Neutral | 0.8733 | Pathogenic | -4.19 | Benign | 0.66 | Disease | 1.00 | Probably damaging | 0.01 | Deleterious | | |
| p.Gly122Asp | VUS | -0.29 | Neutral | 0.4509 | Ambiguous | -8.31 | Pathogenic | 0.66 | Disease | 1.00 | Probably damaging | 0.14 | Tolerated | 25.50 | Deleterious |
| p.Gly122Glu | VUS | -0.41 | Neutral | 0.6464 | Pathogenic | -7.32 | Benign | 0.67 | Disease | 1.00 | Probably damaging | 0.05 | Deleterious | | |
| p.Gly122Ala | VUS | -0.34 | Neutral | 0.5326 | Ambiguous | -6.70 | Benign | 0.48 | Neutral | 1.00 | Probably damaging | 0.08 | Tolerated | 24.70 | Deleterious |
| p.Gly122Val | VUS | -0.08 | Neutral | 0.8833 | Pathogenic | -9.66 | Pathogenic | 0.74 | Disease | 1.00 | Probably damaging | 0.01 | Deleterious | 25.30 | Deleterious |
| p.Gly122Tyr | VUS | -0.60 | Neutral | 0.8741 | Pathogenic | -10.48 | Pathogenic | 0.66 | Disease | 1.00 | Probably damaging | 0.02 | Deleterious | | |
| p.Gly122Cys | VUS | -4.58 | Neutral | 0.6653 | Pathogenic | -7.96 | Pathogenic | 0.68 | Disease | 1.00 | Probably damaging | 0.01 | Deleterious | 26.40 | Deleterious |
| p.Gly122Trp | VUS | -5.31 | Neutral | 0.8381 | Pathogenic | -9.89 | Pathogenic | 0.72 | Disease | 1.00 | Probably damaging | 0 | Deleterious | | |
| p.Gly122Phe | VUS | -1.39 | Neutral | 0.9486 | Pathogenic | -9.80 | Pathogenic | 0.69 | Disease | 1.00 | Probably damaging | 0.01 | Deleterious | | |
| p.His123Asn | VUS | -6.00 | Indeterminate | 0.4955 | Ambiguous | -9.45 | Pathogenic | 0.64 | Disease | 1.00 | Probably damaging | 0.12 | Tolerated | 25.50 | Deleterious |
| p.His123Lys | VUS | -0.43 | Neutral | 0.7284 | Pathogenic | -11.00 | Pathogenic | 0.65 | Disease | 1.00 | Probably damaging | 0.1 | Tolerated | | |
| p.His123Thr | VUS | -26.63 | Indeterminate | 0.9262 | Pathogenic | -14.31 | Pathogenic | 0.67 | Disease | 1.00 | Probably damaging | 0.37 | Tolerated | | |
| p.His123Arg | VUS | -3.23 | Neutral | 0.3271 | Benign | -7.70 | Pathogenic | 0.65 | Disease | 0.99 | Probably damaging | 0.07 | Tolerated | 24.20 | Deleterious |
| p.His123Ser | VUS | -4.18 | Neutral | 0.7649 | Pathogenic | -11.43 | Pathogenic | 0.68 | Disease | 1.00 | Probably damaging | 0.17 | Tolerated | | |
| p.His123Ile | VUS | -5.85 | Indeterminate | 0.9083 | Pathogenic | -18.43 | Pathogenic | 0.73 | Disease | 1.00 | Probably damaging | 0.03 | Deleterious | | |
| p.His123Met | VUS | -13.15 | Indeterminate | 0.8867 | Pathogenic | -12.84 | Pathogenic | 0.72 | Disease | 1.00 | Probably damaging | 0.02 | Deleterious | | |
| p.His123Gln | Benign | -1.24 | Neutral | 0.56 | Ambiguous | -6.66 | Benign | 0.44 | Neutral | 1.00 | Probably damaging | 0.05 | Deleterious | 21.45 | Deleterious |
| p.His123Pro | VUS | -53.15 | Deleterious | 0.8604 | Pathogenic | -14.98 | Pathogenic | 0.82 | Disease | 1.00 | Probably damaging | 0.05 | Deleterious | 25.00 | Deleterious |

| | | | | | | | | | | | | | | | |
|---|---|---|---|---|---|---|---|---|---|---|---|---|---|---|---|
| p.His123Leu | VUS | -8.40 | Indeterminate | 0.5414 | Ambiguous | -11.64 | Pathogenic | 0.77 | Disease | 1.00 | Probably damaging | 0.04 | Deleterious | 25.20 | Deleterious |
| p.His123Asp | VUS | -19.23 | Indeterminate | 0.8124 | Pathogenic | -12.97 | Pathogenic | 0.75 | Disease | 1.00 | Probably damaging | 0.1 | Tolerated | 25.60 | Deleterious |
| p.His123Glu | VUS | -16.55 | Indeterminate | 0.8202 | Pathogenic | -11.12 | Pathogenic | 0.64 | Disease | 1.00 | Probably damaging | 0.06 | Tolerated | | |
| p.His123Ala | VUS | -5.96 | Indeterminate | 0.8083 | Pathogenic | -11.49 | Pathogenic | 0.69 | Disease | 1.00 | Probably damaging | 0.09 | Tolerated | | |
| p.His123Gly | VUS | -53.15 | Deleterious | 0.8408 | Pathogenic | -12.93 | Pathogenic | 0.71 | Disease | 1.00 | Probably damaging | 0.09 | Tolerated | | |
| p.His123Val | VUS | -27.77 | Indeterminate | 0.8496 | Pathogenic | -15.85 | Pathogenic | 0.72 | Disease | 1.00 | Probably damaging | 0.05 | Deleterious | | |
| p.His123Tyr | VUS | -0.91 | Neutral | 0.2345 | Benign | -9.44 | Pathogenic | 0.69 | Disease | 1.00 | Probably damaging | 0.03 | Deleterious | 25.50 | Deleterious |
| p.His123Cys | VUS | -1.67 | Neutral | 0.4191 | Ambiguous | -11.97 | Pathogenic | 0.46 | Neutral | 1.00 | Probably damaging | 0.02 | Deleterious | | |
| p.His123Trp | VUS | -0.86 | Neutral | 0.5355 | Ambiguous | -10.89 | Pathogenic | 0.71 | Disease | 1.00 | Probably damaging | 0.01 | Deleterious | | |
| p.His123Phe | VUS | -24.07 | Indeterminate | 0.5384 | Ambiguous | -10.33 | Pathogenic | 0.71 | Disease | 1.00 | Probably damaging | 0.02 | Deleterious | | |
| p.Arg124Asn | VUS | -0.04 | Neutral | 0.3597 | Ambiguous | -8.79 | Pathogenic | 0.39 | Neutral | 0.82 | Possibly damaging | 0.5 | Tolerated | | |
| p.Arg124Lys | VUS | -0.74 | Neutral | 0.1647 | Benign | -6.83 | Benign | 0.23 | Neutral | 0.18 | Benign | 1 | Tolerated | | |
| p.Arg124Thr | VUS | -8.50 | Indeterminate | 0.1369 | Benign | -7.10 | Benign | 0.21 | Neutral | 0.82 | Possibly damaging | 0.59 | Tolerated | | |
| p.Arg124Ser | VUS | -0.05 | Neutral | 0.211 | Benign | -7.20 | Benign | 0.18 | Neutral | 0.97 | Probably damaging | 0.72 | Tolerated | 6.28 | Neutral |
| p.Arg124Ile | VUS | -32.97 | Indeterminate | 0.2337 | Benign | -10.01 | Pathogenic | 0.38 | Neutral | 0.99 | Probably damaging | 0.24 | Tolerated | | |
| p.Arg124Met | VUS | -0.21 | Neutral | 0.3405 | Ambiguous | -9.58 | Pathogenic | 0.37 | Neutral | 0.94 | Possibly damaging | 0.14 | Tolerated | | |
| p.Arg124His | VUS | -0.31 | Neutral | 0.1059 | Benign | -6.32 | Benign | 0.10 | Neutral | 0.01 | Benign | 0.23 | Tolerated | 11.48 | Neutral |
| p.Arg124Gln | VUS | -0.20 | Neutral | 0.1057 | Benign | -6.73 | Benign | 0.12 | Neutral | 0.87 | Possibly damaging | 0.61 | Tolerated | | |
| p.Arg124Pro | VUS | -0.31 | Neutral | 0.1531 | Benign | -8.06 | Pathogenic | 0.42 | Neutral | 1.00 | Probably damaging | 0.29 | Tolerated | 16.31 | Deleterious |
| p.Arg124Leu | VUS | -0.46 | Neutral | 0.1291 | Benign | -7.80 | Pathogenic | 0.16 | Neutral | 0.96 | Possibly damaging | 0.41 | Tolerated | 10.34 | Neutral |
| p.Arg124Asp | VUS | -0.04 | Neutral | 0.2316 | Benign | -6.81 | Benign | 0.29 | Neutral | 0.98 | Probably damaging | 0.58 | Tolerated | | |
| p.Arg124Glu | VUS | -0.26 | Neutral | 0.1569 | Benign | -6.75 | Benign | 0.36 | Neutral | 0.58 | Possibly damaging | 0.94 | Tolerated | | |
| p.Arg124Ala | VUS | -0.23 | Neutral | 0.118 | Benign | -6.06 | Benign | 0.27 | Neutral | 0.48 | Possibly damaging | 0.76 | Tolerated | | |
| p.Arg124Gly | VUS | -11.05 | Indeterminate | 0.0959 | Benign | -6.10 | Benign | 0.11 | Neutral | 0.05 | Benign | 0.4 | Tolerated | 6.52 | Neutral |
| p.Arg124Val | VUS | -1.45 | Neutral | 0.1826 | Benign | -7.60 | Pathogenic | 0.27 | Neutral | 0.89 | Possibly damaging | 0.35 | Tolerated | | |
| p.Arg124Tyr | VUS | -15.60 | Indeterminate | 0.3596 | Ambiguous | -10.13 | Pathogenic | 0.36 | Neutral | 0.31 | Benign | 0.23 | Tolerated | | |
| p.Arg124Cys | VUS | -1.37 | Neutral | 0.2354 | Benign | -9.11 | Pathogenic | 0.17 | Neutral | 0.63 | Possibly damaging | 0.08 | Tolerated | 12.16 | Neutral |
| p.Arg124Trp | VUS | -0.21 | Neutral | 0.1848 | Benign | -8.64 | Pathogenic | 0.19 | Neutral | 0.98 | Probably damaging | 0.05 | Deleterious | | |
| p.Arg124Phe | VUS | -0.13 | Neutral | 0.3872 | Ambiguous | -10.36 | Pathogenic | 0.40 | Neutral | 0.47 | Possibly damaging | 0.18 | Tolerated | | |
| p.Asp125Asn | VUS | -0.69 | Neutral | 0.238 | Benign | -6.08 | Benign | 0.41 | Neutral | 0.11 | Benign | 0.27 | Tolerated | 15.76 | Deleterious |
| p.Asp125Lys | VUS | -0.03 | Neutral | 0.4591 | Ambiguous | -8.56 | Pathogenic | 0.47 | Neutral | 0.19 | Benign | 0.47 | Tolerated | | |
| p.Asp125Thr | VUS | 0.00 | Neutral | 0.3727 | Ambiguous | -6.64 | Benign | 0.45 | Neutral | 0.06 | Benign | 0.31 | Tolerated | | |
| p.Asp125Arg | VUS | -0.24 | Neutral | 0.3515 | Ambiguous | -6.43 | Benign | 0.45 | Neutral | 0.62 | Possibly damaging | 0.35 | Tolerated | | |
| p.Asp125Ser | VUS | -0.29 | Neutral | 0.2393 | Benign | -5.78 | Benign | 0.39 | Neutral | 0.27 | Benign | 0.49 | Tolerated | | |
| p.Asp125Ile | VUS | 0.00 | Neutral | 0.6176 | Pathogenic | -10.50 | Pathogenic | 0.50 | Neutral | 0.77 | Possibly damaging | 0.1 | Tolerated | | |
| p.Asp125Met | VUS | -0.01 | Neutral | 0.8052 | Pathogenic | -9.40 | Pathogenic | 0.52 | Disease | 0.94 | Possibly damaging | 0.06 | Tolerated | | |
| p.Asp125His | VUS | -0.08 | Neutral | 0.3071 | Benign | -4.75 | Benign | 0.27 | Neutral | 0.00 | Benign | 0.08 | Tolerated | 11.51 | Neutral |
| p.Asp125Gln | VUS | 0.00 | Neutral | 0.2999 | Benign | -5.99 | Benign | 0.26 | Neutral | 0.24 | Benign | 0.3 | Tolerated | | |
| p.Asp125Pro | VUS | -0.63 | Neutral | 0.4534 | Ambiguous | -6.85 | Benign | 0.30 | Neutral | 0.06 | Benign | 0.23 | Tolerated | | |
| p.Asp125Leu | VUS | -0.52 | Neutral | 0.5063 | Ambiguous | -7.72 | Pathogenic | 0.30 | Neutral | 0.62 | Possibly damaging | 0.22 | Tolerated | | |
| p.Asp125Glu | VUS | -0.31 | Neutral | 0.1526 | Benign | -4.16 | Benign | 0.08 | Neutral | 0.00 | Benign | 0.53 | Tolerated | 3.76 | Neutral |
| p.Asp125Ala | VUS | -3.17 | Neutral | 0.1713 | Benign | -3.78 | Benign | 0.17 | Neutral | 0.01 | Benign | 0.47 | Tolerated | 14.59 | Neutral |
| p.Asp125Gly | VUS | -1.02 | Neutral | 0.2285 | Benign | -5.41 | Benign | 0.41 | Neutral | 0.34 | Benign | 0.22 | Tolerated | 18.70 | Deleterious |
| p.Asp125Val | VUS | 0.00 | Neutral | 0.354 | Ambiguous | -7.27 | Benign | 0.32 | Neutral | 0.06 | Benign | 0.16 | Tolerated | 18.83 | Deleterious |
| p.Asp125Tyr | VUS | -53.15 | Deleterious | 0.4002 | Ambiguous | -10.54 | Pathogenic | 0.54 | Disease | 0.62 | Possibly damaging | 0.04 | Deleterious | 19.73 | Deleterious |
| p.Asp125Cys | VUS | -0.23 | Neutral | 0.8053 | Pathogenic | -8.69 | Pathogenic | 0.51 | Disease | 1.00 | Probably damaging | 0.03 | Deleterious | | |
| p.Asp125Trp | VUS | -0.08 | Neutral | 0.8612 | Pathogenic | -10.59 | Pathogenic | 0.51 | Disease | 0.98 | Probably damaging | 0.01 | Deleterious | | |
| p.Asp125Phe | VUS | -0.75 | Neutral | 0.7897 | Pathogenic | -11.42 | Pathogenic | 0.52 | Disease | 0.62 | Possibly damaging | 0.04 | Deleterious | | |
| p.Val126Asn | VUS | -0.47 | Neutral | 0.9138 | Pathogenic | -15.68 | Pathogenic | 0.71 | Disease | 1.00 | Probably damaging | 0 | Deleterious | | |
| p.Val126Lys | VUS | -53.15 | Deleterious | 0.9309 | Pathogenic | -16.52 | Pathogenic | 0.78 | Disease | 1.00 | Probably damaging | 0 | Deleterious | | |
| p.Val126Thr | VUS | 0.00 | Neutral | 0.4014 | Ambiguous | -8.18 | Pathogenic | 0.60 | Disease | 1.00 | Probably damaging | 0.01 | Deleterious | | |
| p.Val126Arg | VUS | -53.15 | Deleterious | 0.8469 | Pathogenic | -14.45 | Pathogenic | 0.78 | Disease | 1.00 | Probably damaging | 0 | Deleterious | | |
| p.Val126Ser | VUS | 0.00 | Neutral | 0.7069 | Pathogenic | -11.39 | Pathogenic | 0.71 | Disease | 1.00 | Probably damaging | 0 | Deleterious | | |
| p.Val126Ile | VUS | 0.00 | Neutral | 0.1435 | Benign | -4.94 | Benign | 0.24 | Neutral | 0.02 | Benign | 0.18 | Tolerated | 14.41 | Neutral |
| p.Val126Met | VUS | 0.00 | Neutral | 0.6606 | Pathogenic | -9.28 | Pathogenic | 0.58 | Disease | 1.00 | Probably damaging | 0.01 | Deleterious | | |
| p.Val126His | VUS | -33.22 | Indeterminate | 0.9725 | Pathogenic | -14.49 | Pathogenic | 0.72 | Disease | 1.00 | Probably damaging | 0 | Deleterious | | |
| p.Val126Gln | VUS | -1.39 | Neutral | 0.9026 | Pathogenic | -15.09 | Pathogenic | 0.75 | Disease | 1.00 | Probably damaging | 0 | Deleterious | | |
| p.Val126Pro | VUS | 0.00 | Neutral | 0.8918 | Pathogenic | -11.16 | Pathogenic | 0.73 | Disease | 1.00 | Probably damaging | 0 | Deleterious | | |
| p.Val126Leu | VUS | 0.00 | Neutral | 0.6571 | Pathogenic | -6.58 | Benign | 0.55 | Disease | 0.38 | Benign | 0.03 | Deleterious | 18.01 | Deleterious |
| p.Val126Asp | Pathogenic | -53.15 | Deleterious | 0.9359 | Pathogenic | -15.06 | Pathogenic | 0.85 | Disease | 1.00 | Probably damaging | 0 | Deleterious | 24.20 | Deleterious |
| p.Val126Glu | VUS | -0.02 | Neutral | 0.876 | Pathogenic | -13.79 | Pathogenic | 0.89 | Disease | 1.00 | Probably damaging | 0 | Deleterious | | |
| p.Val126Ala | VUS | -0.01 | Neutral | 0.3085 | Benign | -6.81 | Benign | 0.60 | Disease | 1.00 | Probably damaging | 0.01 | Deleterious | 23.60 | Deleterious |
| p.Val126Gly | VUS | -0.41 | Neutral | 0.627 | Pathogenic | -12.00 | Pathogenic | 0.69 | Disease | 1.00 | Probably damaging | 0 | Deleterious | 23.80 | Deleterious |
| p.Val126Tyr | VUS | -53.15 | Deleterious | 0.982 | Pathogenic | -16.16 | Pathogenic | 0.71 | Disease | 1.00 | Probably damaging | 0 | Deleterious | | |
| p.Val126Cys | VUS | 0.00 | Neutral | 0.9051 | Pathogenic | -8.45 | Pathogenic | 0.67 | Disease | 1.00 | Probably damaging | 0 | Deleterious | | |
| p.Val126Trp | VUS | -53.15 | Deleterious | 0.9938 | Pathogenic | -15.53 | Pathogenic | 0.68 | Disease | 1.00 | Probably damaging | 0 | Deleterious | | |
| p.Val126Phe | VUS | -0.42 | Neutral | 0.8624 | Pathogenic | -13.84 | Pathogenic | 0.78 | Disease | 0.99 | Probably damaging | 0 | Deleterious | 22.60 | Deleterious |
| p.Ala127Asn | VUS | -1.01 | Neutral | 0.6811 | Pathogenic | -13.00 | Pathogenic | 0.62 | Disease | 0.97 | Probably damaging | 0.01 | Deleterious | | |
| p.Ala127Lys | VUS | -6.63 | Indeterminate | 0.7578 | Pathogenic | -13.09 | Pathogenic | 0.67 | Disease | 1.00 | Probably damaging | 0.02 | Deleterious | | |
| p.Ala127Thr | VUS | -24.84 | Indeterminate | 0.1629 | Benign | -7.48 | Benign | 0.39 | Neutral | 0.97 | Probably damaging | 0.08 | Tolerated | 22.80 | Deleterious |
| p.Ala127Arg | VUS | -19.93 | Indeterminate | 0.501 | Ambiguous | -10.02 | Pathogenic | 0.51 | Disease | 1.00 | Probably damaging | 0.01 | Deleterious | | |
| p.Ala127Ser | Likely benign | -2.10 | Neutral | 0.1514 | Benign | -6.46 | Benign | 0.33 | Neutral | 0.77 | Possibly damaging | 0.02 | Deleterious | 19.18 | Deleterious |
| p.Ala127Ile | VUS | -19.64 | Indeterminate | 0.384 | Ambiguous | -9.36 | Pathogenic | 0.51 | Disease | 0.96 | Possibly damaging | 0.41 | Tolerated | | |
| p.Ala127Met | VUS | -0.87 | Neutral | 0.4705 | Ambiguous | -9.42 | Pathogenic | 0.53 | Disease | 1.00 | Probably damaging | 0.03 | Deleterious | | |
| p.Ala127His | VUS | -26.04 | Indeterminate | 0.7222 | Pathogenic | -11.09 | Pathogenic | 0.61 | Disease | 1.00 | Probably damaging | 0 | Deleterious | | |
| p.Ala127Gln | VUS | -2.66 | Neutral | 0.4734 | Ambiguous | -9.09 | Pathogenic | 0.62 | Disease | 1.00 | Probably damaging | 0.01 | Deleterious | | |
| p.Ala127Pro | VUS | -53.15 | Deleterious | 0.729 | Pathogenic | -8.93 | Pathogenic | 0.72 | Disease | 1.00 | Probably damaging | 0.02 | Deleterious | 23.10 | Deleterious |
| p.Ala127Leu | VUS | -3.30 | Neutral | 0.2703 | Benign | -6.02 | Benign | 0.36 | Neutral | 0.96 | Possibly damaging | 0.12 | Tolerated | | |
| p.Ala127Asp | VUS | -13.68 | Indeterminate | 0.6679 | Pathogenic | -12.66 | Pathogenic | 0.72 | Disease | 1.00 | Probably damaging | 0.01 | Deleterious | | |
| p.Ala127Glu | VUS | -0.78 | Neutral | 0.4097 | Ambiguous | -9.59 | Pathogenic | 0.51 | Disease | 1.00 | Probably damaging | 0.02 | Deleterious | 19.09 | Deleterious |
| p.Ala127Gly | VUS | -7.66 | Indeterminate | 0.1776 | Benign | -7.74 | Pathogenic | 0.34 | Neutral | 0.99 | Probably damaging | 0 | Deleterious | 12.32 | Neutral |
| p.Ala127Val | VUS | -0.26 | Neutral | 0.1436 | Benign | -4.38 | Benign | 0.16 | Neutral | 0.14 | Benign | 1 | Tolerated | 6.92 | Neutral |
| p.Ala127Tyr | VUS | -6.56 | Indeterminate | 0.7619 | Pathogenic | -12.16 | Pathogenic | 0.60 | Disease | 1.00 | Probably damaging | 0.01 | Deleterious | | |
| p.Ala127Cys | VUS | -0.09 | Neutral | 0.5407 | Ambiguous | -8.02 | Pathogenic | 0.49 | Neutral | 1.00 | Probably damaging | 0.02 | Deleterious | | |
| p.Ala127Trp | VUS | -5.00 | Neutral | 0.8559 | Pathogenic | -11.20 | Pathogenic | 0.59 | Disease | 1.00 | Probably damaging | 0 | Deleterious | | |
| p.Ala127Phe | VUS | -3.71 | Neutral | 0.6267 | Pathogenic | -10.86 | Pathogenic | 0.54 | Disease | 1.00 | Probably damaging | 0.02 | Deleterious | | |
| p.Arg128Asn | VUS | 0.00 | Neutral | 0.3288 | Benign | -6.64 | Benign | 0.23 | Neutral | 0.79 | Possibly damaging | 0.24 | Tolerated | | |
| p.Arg128Lys | VUS | -2.71 | Neutral | 0.2077 | Benign | -6.72 | Benign | 0.12 | Neutral | 0.08 | Benign | 0.44 | Tolerated | | |
| p.Arg128Thr | VUS | -0.32 | Neutral | 0.1483 | Benign | -6.10 | Benign | 0.29 | Neutral | 0.79 | Possibly damaging | 0.27 | Tolerated | | |
| p.Arg128Ser | VUS | -19.13 | Indeterminate | 0.1872 | Benign | -4.93 | Benign | 0.14 | Neutral | 0.76 | Possibly damaging | 0.53 | Tolerated | | |
| p.Arg128Ile | VUS | -8.33 | Indeterminate | 0.2361 | Benign | -8.69 | Pathogenic | 0.36 | Neutral | 0.90 | Possibly damaging | 0.08 | Tolerated | | |
| p.Arg128Met | VUS | -1.08 | Neutral | 0.3178 | Benign | -8.05 | Pathogenic | 0.37 | Neutral | 0.68 | Possibly damaging | 0.05 | Deleterious | | |
| p.Arg128His | VUS | -3.37 | Neutral | 0.1143 | Benign | -5.66 | Benign | 0.14 | Neutral | 0.81 | Possibly damaging | 0.07 | Tolerated | | |
| p.Arg128Gln | VUS | -0.11 | Neutral | 0.118 | Benign | -5.62 | Benign | 0.12 | Neutral | 0.13 | Benign | 0.28 | Tolerated | 17.20 | Deleterious |
| p.Arg128Pro | VUS | -53.15 | Deleterious | 0.6461 | Pathogenic | -10.17 | Pathogenic | 0.40 | Neutral | 1.00 | Probably damaging | 0.15 | Tolerated | 19.02 | Deleterious |
| p.Arg128Leu | VUS | -1.25 | Neutral | 0.1061 | Benign | -5.79 | Benign | 0.14 | Neutral | 0.04 | Benign | 0.14 | Tolerated | 18.39 | Deleterious |
| p.Arg128Asp | VUS | -0.12 | Neutral | 0.2566 | Benign | -5.30 | Benign | 0.21 | Neutral | 0.94 | Possibly damaging | 0.31 | Tolerated | | |
| p.Arg128Glu | VUS | -0.71 | Neutral | 0.2037 | Benign | -5.06 | Benign | 0.23 | Neutral | 0.19 | Benign | 1 | Tolerated | | |
| p.Arg128Ala | VUS | -0.48 | Neutral | 0.1075 | Benign | -3.42 | Benign | 0.23 | Neutral | 0.25 | Benign | 0.5 | Tolerated | | |
| p.Arg128Gly | VUS | -6.37 | Indeterminate | 0.0979 | Benign | -4.91 | Benign | 0.12 | Neutral | 0.01 | Benign | 0.31 | Tolerated | 10.62 | Neutral |
| p.Arg128Val | VUS | -0.07 | Neutral | 0.1998 | Benign | -6.73 | Benign | 0.38 | Neutral | 0.15 | Benign | 0.14 | Tolerated | | |
| p.Arg128Tyr | VUS | -0.76 | Neutral | 0.3589 | Ambiguous | -7.14 | Benign | 0.36 | Neutral | 0.69 | Possibly damaging | 0.03 | Deleterious | | |
| p.Arg128Cys | VUS | 0.00 | Neutral | 0.1985 | Benign | -7.01 | Benign | 0.33 | Neutral | 1.00 | Probably damaging | 0.04 | Deleterious | | |
| p.Arg128Trp | VUS | -0.10 | Neutral | 0.1984 | Benign | -7.82 | Pathogenic | 0.17 | Neutral | 0.99 | Probably damaging | 0.01 | Deleterious | 21.80 | Deleterious |
| p.Arg128Phe | VUS | -6.69 | Indeterminate | 0.3814 | Ambiguous | -6.18 | Benign | 0.40 | Neutral | 0.26 | Benign | 0.03 | Deleterious | | |
| p.Tyr129Asn | VUS | -33.22 | Indeterminate | 0.9556 | Pathogenic | -15.00 | Pathogenic | 0.66 | Disease | 1.00 | Probably damaging | 0.09 | Tolerated | 32.00 | Deleterious |
| p.Tyr129Lys | VUS | -3.97 | Neutral | 0.9891 | Pathogenic | -15.47 | Pathogenic | 0.61 | Disease | 1.00 | Probably damaging | 0.09 | Tolerated | | |
| p.Tyr129Thr | VUS | -27.35 | Indeterminate | 0.9689 | Pathogenic | -12.26 | Pathogenic | 0.59 | Disease | 1.00 | Probably damaging | 0.27 | Tolerated | | |
| p.Tyr129Arg | VUS | -0.71 | Neutral | 0.9606 | Pathogenic | -11.05 | Pathogenic | 0.57 | Disease | 1.00 | Probably damaging | 0.1 | Tolerated | | |
| p.Tyr129Ser | VUS | -2.79 | Neutral | 0.9417 | Pathogenic | -10.96 | Pathogenic | 0.62 | Disease | 1.00 | Probably damaging | 0.14 | Tolerated | 30.00 | Deleterious |
| p.Tyr129Ile | VUS | -4.50 | Neutral | 0.9257 | Pathogenic | -11.98 | Pathogenic | 0.56 | Disease | 0.98 | Probably damaging | 0.15 | Tolerated | | |

| | | | | | | | | | | | | | | | |
|---|---|---|---|---|---|---|---|---|---|---|---|---|---|---|---|
| p.Tyr129Met | VUS | -0.78 | Neutral | 0.9612 | Pathogenic | -12.23 | Pathogenic | 0.53 | Disease | 1.00 | Probably damaging | 0.08 | Tolerated | | |
| p.Tyr129His | VUS | -1.08 | Neutral | 0.8201 | Pathogenic | -7.47 | Benign | 0.53 | Disease | 1.00 | Probably damaging | 0.16 | Tolerated | 28.00 | Deleterious |
| p.Tyr129Gln | VUS | -0.73 | Neutral | 0.9784 | Pathogenic | -11.98 | Pathogenic | 0.54 | Disease | 1.00 | Probably damaging | 0.09 | Tolerated | | |
| p.Tyr129Pro | VUS | -53.15 | Deleterious | 0.9962 | Pathogenic | -14.76 | Pathogenic | 0.59 | Disease | 1.00 | Probably damaging | 0.07 | Tolerated | | |
| p.Tyr129Leu | VUS | -0.36 | Neutral | 0.8573 | Pathogenic | -9.20 | Pathogenic | 0.52 | Disease | 0.94 | Possibly damaging | 0.28 | Tolerated | | |
| p.Tyr129Asp | VUS | -16.50 | Indeterminate | 0.9656 | Pathogenic | -13.41 | Pathogenic | 0.64 | Disease | 1.00 | Probably damaging | 0.06 | Tolerated | 32.00 | Deleterious |
| p.Tyr129Glu | VUS | -0.57 | Neutral | 0.9774 | Pathogenic | -12.49 | Pathogenic | 0.60 | Disease | 1.00 | Probably damaging | 0.08 | Tolerated | | |
| p.Tyr129Ala | VUS | -0.56 | Neutral | 0.9607 | Pathogenic | -10.40 | Pathogenic | 0.39 | Neutral | 1.00 | Probably damaging | 0.17 | Tolerated | | |
| p.Tyr129Gly | VUS | -3.93 | Neutral | 0.9617 | Pathogenic | -14.19 | Pathogenic | 0.56 | Disease | 1.00 | Probably damaging | 0.1 | Tolerated | | |
| p.Tyr129Val | VUS | -0.90 | Neutral | 0.8873 | Pathogenic | -9.78 | Pathogenic | 0.57 | Disease | 1.00 | Probably damaging | 0.18 | Tolerated | | |
| p.Tyr129Cys | VUS | -0.95 | Neutral | 0.8807 | Pathogenic | -9.02 | Pathogenic | 0.41 | Neutral | 1.00 | Probably damaging | 0.15 | Tolerated | 27.90 | Deleterious |
| p.Tyr129Trp | VUS | -0.17 | Neutral | 0.7312 | Pathogenic | -7.42 | Benign | 0.52 | Disease | 1.00 | Probably damaging | 0.14 | Tolerated | | |
| p.Tyr129Phe | VUS | -1.69 | Neutral | 0.2033 | Benign | -2.91 | Benign | 0.24 | Neutral | 0.18 | Benign | 0.96 | Tolerated | 22.70 | Deleterious |
| p.Leu130Asn | VUS | -29.02 | Indeterminate | 0.9902 | Pathogenic | -16.37 | Pathogenic | 0.76 | Disease | 1.00 | Probably damaging | 0.01 | Deleterious | | |
| p.Leu130Lys | VUS | -53.15 | Deleterious | 0.9732 | Pathogenic | -16.08 | Pathogenic | 0.78 | Disease | 0.56 | Possibly damaging | 0.02 | Deleterious | | |
| p.Leu130Thr | VUS | -2.31 | Neutral | 0.9455 | Pathogenic | -11.34 | Pathogenic | 0.79 | Disease | 1.00 | Probably damaging | 0.19 | Tolerated | | |
| p.Leu130Arg | VUS | -53.15 | Deleterious | 0.8515 | Pathogenic | -12.89 | Pathogenic | 0.90 | Disease | 0.99 | Probably damaging | 0.01 | Deleterious | 26.80 | Deleterious |
| p.Leu130Ser | VUS | -3.13 | Neutral | 0.9847 | Pathogenic | -12.78 | Pathogenic | 0.87 | Disease | 1.00 | Probably damaging | 0.04 | Deleterious | | |
| p.Leu130Ile | VUS | -0.30 | Neutral | 0.4013 | Ambiguous | -6.51 | Benign | 0.60 | Disease | 1.00 | Probably damaging | 0.11 | Tolerated | | |
| p.Leu130Met | VUS | -0.88 | Neutral | 0.6895 | Pathogenic | -8.81 | Pathogenic | 0.66 | Disease | 1.00 | Probably damaging | 0.05 | Deleterious | 24.90 | Deleterious |
| p.Leu130His | VUS | -26.23 | Indeterminate | 0.9657 | Pathogenic | -14.64 | Pathogenic | 0.87 | Disease | 1.00 | Probably damaging | 0.01 | Deleterious | | |
| p.Leu130Gln | VUS | -21.89 | Indeterminate | 0.9262 | Pathogenic | -13.79 | Pathogenic | 0.89 | Disease | 0.99 | Probably damaging | 0.01 | Deleterious | 27.00 | Deleterious |
| p.Leu130Pro | VUS | -53.15 | Deleterious | 0.9537 | Pathogenic | -13.21 | Pathogenic | 0.92 | Disease | 1.00 | Probably damaging | 0.02 | Deleterious | 27.00 | Deleterious |
| p.Leu130Asp | VUS | -53.15 | Deleterious | 0.9973 | Pathogenic | -15.79 | Pathogenic | 0.74 | Disease | 1.00 | Probably damaging | 0.01 | Deleterious | | |
| p.Leu130Glu | VUS | -33.22 | Indeterminate | 0.9695 | Pathogenic | -14.38 | Pathogenic | 0.79 | Disease | 1.00 | Probably damaging | 0.01 | Deleterious | | |
| p.Leu130Ala | VUS | -3.07 | Neutral | 0.8823 | Pathogenic | -9.53 | Pathogenic | 0.76 | Disease | 1.00 | Probably damaging | 0.08 | Tolerated | | |
| p.Leu130Gly | VUS | -2.06 | Neutral | 0.9579 | Pathogenic | -14.41 | Pathogenic | 0.79 | Disease | 1.00 | Probably damaging | 0.01 | Deleterious | | |
| p.Leu130Val | VUS | -0.32 | Neutral | 0.3976 | Ambiguous | -6.54 | Benign | 0.67 | Disease | 0.97 | Probably damaging | 0.13 | Tolerated | 25.80 | Deleterious |
| p.Leu130Tyr | VUS | -3.10 | Neutral | 0.9672 | Pathogenic | -14.10 | Pathogenic | 0.74 | Disease | 0.99 | Probably damaging | 0.01 | Deleterious | | |
| p.Leu130Cys | VUS | -1.46 | Neutral | 0.8681 | Pathogenic | -11.99 | Pathogenic | 0.72 | Disease | 1.00 | Probably damaging | 0.16 | Tolerated | | |
| p.Leu130Trp | VUS | -8.91 | Indeterminate | 0.9236 | Pathogenic | -12.53 | Pathogenic | 0.62 | Disease | 1.00 | Probably damaging | 0.01 | Deleterious | | |
| p.Leu130Phe | VUS | -0.48 | Neutral | 0.7227 | Pathogenic | -8.71 | Pathogenic | 0.67 | Disease | 1.00 | Probably damaging | 0.04 | Deleterious | | |
| p.Arg131Asn | VUS | -0.68 | Neutral | 0.451 | Ambiguous | -8.74 | Pathogenic | 0.52 | Disease | 0.99 | Probably damaging | 0.27 | Tolerated | | |
| p.Arg131Lys | VUS | -0.01 | Neutral | 0.1996 | Benign | -7.54 | Pathogenic | 0.48 | Neutral | 0.87 | Possibly damaging | 0.7 | Tolerated | | |
| p.Arg131Thr | VUS | -8.63 | Indeterminate | 0.2073 | Benign | -7.55 | Pathogenic | 0.54 | Disease | 1.00 | Probably damaging | 0.38 | Tolerated | | |
| p.Arg131Ser | VUS | -1.15 | Neutral | 0.3002 | Benign | -6.80 | Benign | 0.53 | Disease | 1.00 | Probably damaging | 0.34 | Tolerated | 25.70 | Deleterious |
| p.Arg131Ile | VUS | -0.04 | Neutral | 0.2953 | Benign | -9.77 | Pathogenic | 0.59 | Disease | 1.00 | Probably damaging | 0.34 | Tolerated | | |
| p.Arg131Met | VUS | -0.35 | Neutral | 0.3648 | Ambiguous | -9.38 | Pathogenic | 0.58 | Disease | 1.00 | Probably damaging | 0.35 | Tolerated | | |
| p.Arg131His | VUS | -0.64 | Neutral | 0.1147 | Benign | -6.76 | Benign | 0.33 | Neutral | 0.23 | Benign | 0.41 | Tolerated | 24.10 | Deleterious |
| p.Arg131Gln | VUS | -22.33 | Indeterminate | 0.1263 | Benign | -6.16 | Benign | 0.34 | Neutral | 1.00 | Probably damaging | 0.37 | Tolerated | | |
| p.Arg131Pro | VUS | -19.41 | Indeterminate | 0.7075 | Pathogenic | -10.28 | Pathogenic | 0.64 | Disease | 1.00 | Probably damaging | 0.2 | Tolerated | 23.80 | Deleterious |
| p.Arg131Leu | VUS | -0.02 | Neutral | 0.1195 | Benign | -6.72 | Benign | 0.37 | Neutral | 1.00 | Probably damaging | 0.54 | Tolerated | 22.80 | Deleterious |
| p.Arg131Asp | VUS | 0.00 | Neutral | 0.3993 | Ambiguous | -8.15 | Pathogenic | 0.59 | Disease | 1.00 | Probably damaging | 0.17 | Tolerated | | |
| p.Arg131Glu | VUS | -0.04 | Neutral | 0.2321 | Benign | -6.63 | Benign | 0.53 | Disease | 0.98 | Probably damaging | 0.32 | Tolerated | | |
| p.Arg131Ala | VUS | -0.74 | Neutral | 0.1688 | Benign | -5.48 | Benign | 0.53 | Disease | 1.00 | Probably damaging | 0.52 | Tolerated | | |
| p.Arg131Gly | VUS | -1.08 | Neutral | 0.1641 | Benign | -6.87 | Benign | 0.38 | Neutral | 1.00 | Probably damaging | 0.26 | Tolerated | 25.70 | Deleterious |
| p.Arg131Val | VUS | -0.61 | Neutral | 0.2515 | Benign | -8.11 | Pathogenic | 0.58 | Disease | 1.00 | Probably damaging | 0.56 | Tolerated | | |
| p.Arg131Tyr | VUS | -0.93 | Neutral | 0.3346 | Benign | -9.54 | Pathogenic | 0.53 | Disease | 0.93 | Possibly damaging | 0.58 | Tolerated | | |
| p.Arg131Cys | VUS | -53.15 | Deleterious | 0.2663 | Benign | -8.50 | Pathogenic | 0.41 | Neutral | 1.00 | Probably damaging | 0.12 | Tolerated | 28.30 | Deleterious |
| p.Arg131Trp | VUS | -0.79 | Neutral | 0.1928 | Benign | -8.69 | Pathogenic | 0.58 | Disease | 1.00 | Probably damaging | 0.11 | Tolerated | | |
| p.Arg131Phe | VUS | 0.00 | Neutral | 0.3727 | Ambiguous | -10.29 | Pathogenic | 0.57 | Disease | 0.97 | Probably damaging | 0.41 | Tolerated | | |
| p.Ala132Asn | VUS | -33.21 | Indeterminate | 0.2461 | Benign | -6.61 | Benign | 0.35 | Neutral | 0.00 | Benign | 0.08 | Tolerated | | |
| p.Ala132Lys | VUS | -3.78 | Neutral | 0.3425 | Ambiguous | -6.43 | Benign | 0.46 | Neutral | 0.85 | Possibly damaging | 0.41 | Tolerated | | |
| p.Ala132Thr | VUS | -20.37 | Indeterminate | 0.1104 | Benign | -5.18 | Benign | 0.31 | Neutral | 0.05 | Benign | 0.35 | Tolerated | 12.63 | Neutral |
| p.Ala132Arg | VUS | -1.61 | Neutral | 0.2268 | Benign | -4.54 | Benign | 0.38 | Neutral | 0.95 | Possibly damaging | 0.07 | Tolerated | | |
| p.Ala132Ser | VUS | -33.21 | Indeterminate | 0.1128 | Benign | -4.01 | Benign | 0.29 | Neutral | 0.04 | Benign | 0.56 | Tolerated | 10.21 | Neutral |
| p.Ala132Ile | VUS | -12.89 | Indeterminate | 0.3068 | Benign | -7.48 | Benign | 0.47 | Neutral | 0.74 | Possibly damaging | 0.03 | Deleterious | | |
| p.Ala132Met | VUS | -8.62 | Indeterminate | 0.2506 | Benign | -7.04 | Benign | 0.43 | Neutral | 0.98 | Probably damaging | 0.02 | Deleterious | | |
| p.Ala132His | VUS | -19.91 | Indeterminate | 0.2825 | Benign | -6.65 | Benign | 0.46 | Neutral | 0.99 | Probably damaging | 0.02 | Deleterious | | |
| p.Ala132Gln | VUS | -1.62 | Neutral | 0.1939 | Benign | -4.86 | Benign | 0.36 | Neutral | 0.98 | Probably damaging | 0.07 | Tolerated | | |
| p.Ala132Pro | VUS | -3.53 | Neutral | 0.5877 | Pathogenic | -6.77 | Benign | 0.52 | Disease | 0.95 | Possibly damaging | 0.06 | Tolerated | 18.35 | Deleterious |
| p.Ala132Leu | VUS | -9.42 | Indeterminate | 0.1643 | Benign | -5.30 | Benign | 0.45 | Neutral | 0.74 | Possibly damaging | 0.05 | Deleterious | | |
| p.Ala132Asp | VUS | -29.25 | Indeterminate | 0.1618 | Benign | -4.14 | Benign | 0.36 | Neutral | 0.01 | Benign | 0.08 | Tolerated | | |
| p.Ala132Glu | VUS | -9.67 | Indeterminate | 0.1341 | Benign | -2.89 | Benign | 0.31 | Neutral | 0.67 | Possibly damaging | 0.1 | Tolerated | 10.63 | Neutral |
| p.Ala132Gly | VUS | -53.15 | Deleterious | 0.1244 | Benign | -3.15 | Benign | 0.29 | Neutral | 0.63 | Possibly damaging | 0.09 | Tolerated | 14.01 | Neutral |
| p.Ala132Val | VUS | -4.37 | Neutral | 0.1471 | Benign | -5.64 | Benign | 0.30 | Neutral | 0.11 | Benign | 0.06 | Tolerated | 10.93 | Neutral |
| p.Ala132Tyr | VUS | -8.74 | Indeterminate | 0.3593 | Ambiguous | -8.46 | Pathogenic | 0.46 | Neutral | 0.99 | Probably damaging | 0.01 | Deleterious | | |
| p.Ala132Cys | VUS | -14.40 | Indeterminate | 0.4755 | Ambiguous | -6.02 | Benign | 0.44 | Neutral | 0.99 | Probably damaging | 0.02 | Deleterious | | |
| p.Ala132Trp | VUS | -4.92 | Neutral | 0.4766 | Ambiguous | -7.45 | Benign | 0.51 | Disease | 1.00 | Probably damaging | 0.01 | Deleterious | | |
| p.Ala132Phe | VUS | -4.29 | Neutral | 0.29 | Benign | -8.13 | Pathogenic | 0.46 | Neutral | 0.98 | Probably damaging | 0.01 | Deleterious | | |
| p.Ala133Asn | VUS | -0.02 | Neutral | 0.4307 | Ambiguous | -7.78 | Pathogenic | 0.40 | Neutral | 0.00 | Benign | 0.33 | Tolerated | | |
| p.Ala133Lys | VUS | -0.09 | Neutral | 0.5727 | Pathogenic | -5.53 | Benign | 0.51 | Disease | 0.65 | Possibly damaging | 0.36 | Tolerated | | |
| p.Ala133Thr | VUS | -0.13 | Neutral | 0.125 | Benign | -5.14 | Benign | 0.43 | Neutral | 0.00 | Benign | 0.42 | Tolerated | 18.39 | Deleterious |
| p.Ala133Arg | VUS | -2.27 | Neutral | 0.3825 | Ambiguous | -3.20 | Benign | 0.50 | Disease | 0.87 | Possibly damaging | 0.39 | Tolerated | | |
| p.Ala133Ser | VUS | -0.11 | Neutral | 0.1281 | Benign | -5.87 | Benign | 0.31 | Neutral | 0.01 | Benign | 0.43 | Tolerated | 16.65 | Deleterious |
| p.Ala133Ile | VUS | -0.71 | Neutral | 0.3424 | Ambiguous | -5.89 | Benign | 0.39 | Neutral | 0.48 | Possibly damaging | 0.4 | Tolerated | | |
| p.Ala133Met | VUS | -0.01 | Neutral | 0.3164 | Benign | -4.42 | Benign | 0.49 | Neutral | 0.93 | Possibly damaging | 0.22 | Tolerated | | |
| p.Ala133His | VUS | -0.26 | Neutral | 0.449 | Ambiguous | -5.71 | Benign | 0.50 | Neutral | 0.98 | Probably damaging | 0.55 | Tolerated | | |
| p.Ala133Gln | VUS | -0.08 | Neutral | 0.3327 | Benign | -3.97 | Benign | 0.51 | Disease | 0.87 | Possibly damaging | 0.33 | Tolerated | | |
| p.Ala133Pro | VUS | -0.37 | Neutral | 0.5773 | Pathogenic | -6.87 | Benign | 0.38 | Neutral | 0.73 | Possibly damaging | 0.22 | Tolerated | 22.30 | Deleterious |
| p.Ala133Leu | VUS | -0.18 | Neutral | 0.2337 | Benign | -3.65 | Benign | 0.49 | Neutral | 0.48 | Possibly damaging | 0.66 | Tolerated | | |
| p.Ala133Asp | VUS | -7.43 | Indeterminate | 0.4543 | Ambiguous | -7.93 | Pathogenic | 0.57 | Disease | 0.16 | Benign | 0.24 | Tolerated | 17.88 | Deleterious |
| p.Ala133Glu | VUS | -2.21 | Neutral | 0.2918 | Benign | -5.22 | Benign | 0.50 | Neutral | 0.41 | Benign | 0.32 | Tolerated | | |
| p.Ala133Gly | VUS | -1.09 | Neutral | 0.178 | Benign | -5.15 | Benign | 0.45 | Neutral | 0.00 | Benign | 0.35 | Tolerated | 16.79 | Deleterious |
| p.Ala133Val | VUS | -2.44 | Neutral | 0.1407 | Benign | -3.13 | Benign | 0.31 | Neutral | 0.01 | Benign | 0.51 | Tolerated | 13.93 | Neutral |
| p.Ala133Tyr | VUS | -1.01 | Neutral | 0.5866 | Pathogenic | -8.24 | Pathogenic | 0.53 | Disease | 0.98 | Probably damaging | 1 | Tolerated | | |
| p.Ala133Cys | VUS | -0.27 | Neutral | 0.4304 | Ambiguous | -6.36 | Benign | 0.50 | Neutral | 0.98 | Probably damaging | 0.18 | Tolerated | | |
| p.Ala133Trp | VUS | -0.43 | Neutral | 0.7089 | Pathogenic | -6.23 | Benign | 0.55 | Disease | 0.99 | Probably damaging | 0.18 | Tolerated | | |
| p.Ala133Phe | VUS | -0.96 | Neutral | 0.4143 | Ambiguous | -7.72 | Pathogenic | 0.53 | Disease | 0.93 | Possibly damaging | 0.7 | Tolerated | | |
| p.Ala134Asn | VUS | -0.10 | Neutral | 0.2328 | Benign | -7.73 | Pathogenic | 0.37 | Neutral | 0.03 | Benign | 0.2 | Tolerated | | |
| p.Ala134Lys | VUS | -0.93 | Neutral | 0.3067 | Benign | -7.53 | Pathogenic | 0.38 | Neutral | 0.29 | Benign | 0.31 | Tolerated | | |
| p.Ala134Thr | VUS | -2.44 | Neutral | 0.0865 | Benign | -5.69 | Benign | 0.23 | Neutral | 0.00 | Benign | 1 | Tolerated | 0.94 | Neutral |
| p.Ala134Arg | VUS | -2.27 | Neutral | 0.1773 | Benign | -5.05 | Benign | 0.37 | Neutral | 0.45 | Possibly damaging | 0.19 | Tolerated | | |
| p.Ala134Ser | VUS | -1.36 | Neutral | 0.0891 | Benign | -5.10 | Benign | 0.19 | Neutral | 0.00 | Benign | 0.47 | Tolerated | 0.19 | Neutral |
| p.Ala134Ile | VUS | -0.33 | Neutral | 0.2793 | Benign | -7.47 | Benign | 0.39 | Neutral | 0.29 | Benign | 0.55 | Tolerated | | |
| p.Ala134Met | VUS | -0.23 | Neutral | 0.2727 | Benign | -5.48 | Benign | 0.33 | Neutral | 0.86 | Possibly damaging | 0.11 | Tolerated | | |
| p.Ala134His | VUS | -0.11 | Neutral | 0.2608 | Benign | -5.82 | Benign | 0.37 | Neutral | 0.86 | Possibly damaging | 0.08 | Tolerated | | |
| p.Ala134Gln | VUS | -0.26 | Neutral | 0.1889 | Benign | -3.90 | Benign | 0.29 | Neutral | 0.59 | Possibly damaging | 0.38 | Tolerated | | |
| p.Ala134Pro | VUS | -0.55 | Neutral | 0.1837 | Benign | -5.78 | Benign | 0.39 | Neutral | 0.25 | Benign | 0.18 | Tolerated | 9.21 | Neutral |
| p.Ala134Leu | VUS | -0.05 | Neutral | 0.1652 | Benign | -4.97 | Benign | 0.31 | Neutral | 0.15 | Benign | 0.35 | Tolerated | | |
| p.Ala134Asp | VUS | -0.09 | Neutral | 0.1632 | Benign | -6.35 | Benign | 0.37 | Neutral | 0.10 | Benign | 0.19 | Tolerated | | |
| p.Ala134Glu | VUS | -0.09 | Neutral | 0.1294 | Benign | -4.64 | Benign | 0.25 | Neutral | 0.00 | Benign | 0.3 | Tolerated | 10.08 | Neutral |
| p.Ala134Gly | VUS | -0.40 | Neutral | 0.0788 | Benign | -4.12 | Benign | 0.28 | Neutral | 0.00 | Benign | 0.14 | Tolerated | 10.93 | Neutral |
| p.Ala134Val | VUS | -0.10 | Neutral | 0.1345 | Benign | -5.35 | Benign | 0.27 | Neutral | 0.02 | Benign | 0.81 | Tolerated | 10.60 | Neutral |
| p.Ala134Tyr | VUS | -0.75 | Neutral | 0.3656 | Ambiguous | -8.41 | Pathogenic | 0.39 | Neutral | 0.95 | Possibly damaging | 0.08 | Tolerated | | |
| p.Ala134Cys | VUS | -1.54 | Neutral | 0.3103 | Benign | -6.67 | Benign | 0.37 | Neutral | 0.95 | Possibly damaging | 0.1 | Tolerated | | |
| p.Ala134Trp | VUS | -0.83 | Neutral | 0.4264 | Ambiguous | -7.45 | Benign | 0.40 | Neutral | 0.99 | Probably damaging | 0.02 | Deleterious | | |
| p.Ala134Phe | VUS | -0.06 | Neutral | 0.2995 | Benign | -8.52 | Pathogenic | 0.39 | Neutral | 0.86 | Possibly damaging | 0.09 | Tolerated | | |
| p.Gly135Asn | VUS | -0.01 | Neutral | 0.2191 | Benign | -6.23 | Benign | 0.26 | Neutral | 0.98 | Probably damaging | 0.07 | Tolerated | | |
| p.Gly135Lys | VUS | -3.29 | Neutral | 0.3025 | Benign | -6.29 | Benign | 0.46 | Neutral | 0.96 | Probably damaging | 0.11 | Tolerated | | |
| p.Gly135Thr | VUS | -0.86 | Neutral | 0.1142 | Benign | -4.91 | Benign | 0.42 | Neutral | 0.98 | Probably damaging | 0.16 | Tolerated | | |

| | | | | | | | | | | | | | | | |
|---|---|---|---|---|---|---|---|---|---|---|---|---|---|---|---|
| p.Gly135Arg | VUS | 0.00 | Neutral | 0.1684 | Benign | -4.16 | Benign | 0.29 | Neutral | 0.97 | Probably damaging | 0.08 | Tolerated | 14.69 | Neutral |
| p.Gly135Ser | VUS | -1.39 | Neutral | 0.1081 | Benign | -4.09 | Benign | 0.18 | Neutral | 0.93 | Possibly damaging | 0.44 | Tolerated | | |
| p.Gly135Ile | VUS | -2.16 | Neutral | 0.2628 | Benign | -7.30 | Benign | 0.45 | Neutral | 1.00 | Probably damaging | 0.06 | Tolerated | | |
| p.Gly135Met | VUS | -0.29 | Neutral | 0.4175 | Ambiguous | -6.81 | Benign | 0.46 | Neutral | 1.00 | Probably damaging | 0.05 | Deleterious | | |
| p.Gly135His | VUS | -0.47 | Neutral | 0.2218 | Benign | -6.08 | Benign | 0.45 | Neutral | 1.00 | Probably damaging | 0.03 | Deleterious | | |
| p.Gly135Gln | VUS | -0.74 | Neutral | 0.2094 | Benign | -4.77 | Benign | 0.42 | Neutral | 0.96 | Probably damaging | 0.07 | Tolerated | | |
| p.Gly135Pro | VUS | -0.58 | Neutral | 0.3056 | Benign | -4.07 | Benign | 0.41 | Neutral | 0.99 | Probably damaging | 0.22 | Tolerated | | |
| p.Gly135Leu | VUS | -1.45 | Neutral | 0.2535 | Benign | -5.17 | Benign | 0.43 | Neutral | 0.98 | Probably damaging | 0.23 | Tolerated | | |
| p.Gly135Asp | VUS | -1.16 | Neutral | 0.1262 | Benign | -4.55 | Benign | 0.30 | Neutral | 0.88 | Possibly damaging | 0.08 | Tolerated | | |
| p.Gly135Glu | VUS | -31.01 | Indeterminate | 0.1313 | Benign | -3.64 | Benign | 0.20 | Neutral | 0.07 | Benign | 0.1 | Tolerated | 11.99 | Neutral |
| p.Gly135Ala | VUS | -0.30 | Neutral | 0.0882 | Benign | -3.00 | Benign | 0.31 | Neutral | 0.91 | Possibly damaging | 1 | Tolerated | 14.70 | Neutral |
| p.Gly135Val | VUS | -1.45 | Neutral | 0.1561 | Benign | -5.17 | Benign | 0.40 | Neutral | 0.99 | Probably damaging | 0.16 | Tolerated | 19.92 | Deleterious |
| p.Gly135Tyr | VUS | -0.60 | Neutral | 0.3453 | Ambiguous | -7.46 | Benign | 0.45 | Neutral | 1.00 | Probably damaging | 0.04 | Deleterious | | |
| p.Gly135Cys | VUS | -0.05 | Neutral | 0.2046 | Benign | -6.28 | Benign | 0.39 | Neutral | 1.00 | Probably damaging | 0.09 | Tolerated | | |
| p.Gly135Trp | VUS | -0.45 | Neutral | 0.2645 | Benign | -6.78 | Benign | 0.43 | Neutral | 1.00 | Probably damaging | 0.02 | Deleterious | 21.20 | Deleterious |
| p.Gly135Phe | VUS | -0.98 | Neutral | 0.4177 | Ambiguous | -7.49 | Benign | 0.45 | Neutral | 1.00 | Probably damaging | 0.05 | Deleterious | | |
| p.Gly136Asn | VUS | -7.38 | Indeterminate | 0.1736 | Benign | -5.90 | Benign | 0.24 | Neutral | 0.00 | Benign | 0.21 | Tolerated | | |
| p.Gly136Lys | VUS | -0.08 | Neutral | 0.2592 | Benign | -5.97 | Benign | 0.32 | Neutral | 0.36 | Benign | 0.23 | Tolerated | | |
| p.Gly136Thr | VUS | -0.05 | Neutral | 0.1004 | Benign | -4.43 | Benign | 0.26 | Neutral | 0.52 | Possibly damaging | 0.41 | Tolerated | | |
| p.Gly136Arg | VUS | -0.04 | Neutral | 0.1522 | Benign | -4.07 | Benign | 0.19 | Neutral | 0.03 | Benign | 0.19 | Tolerated | 12.18 | Neutral |
| p.Gly136Ser | VUS | -5.18 | Neutral | 0.0975 | Benign | -3.77 | Benign | 0.16 | Neutral | 0.15 | Benign | 0.65 | Tolerated | 12.00 | Neutral |
| p.Gly136Ile | VUS | -0.50 | Neutral | 0.1965 | Benign | -6.12 | Benign | 0.35 | Neutral | 0.79 | Possibly damaging | 0.24 | Tolerated | | |
| p.Gly136Met | VUS | -0.11 | Neutral | 0.3032 | Benign | -6.66 | Benign | 0.33 | Neutral | 0.99 | Probably damaging | 0.1 | Tolerated | | |
| p.Gly136His | VUS | -8.50 | Indeterminate | 0.1781 | Benign | -5.62 | Benign | 0.33 | Neutral | 0.02 | Benign | 0.16 | Tolerated | | |
| p.Gly136Gln | VUS | -0.68 | Neutral | 0.1665 | Benign | -4.91 | Benign | 0.29 | Neutral | 0.89 | Possibly damaging | 0.18 | Tolerated | | |
| p.Gly136Pro | VUS | -17.41 | Indeterminate | 0.2606 | Benign | -3.68 | Benign | 0.34 | Neutral | 0.89 | Possibly damaging | 0.27 | Tolerated | | |
| p.Gly136Leu | VUS | -6.43 | Indeterminate | 0.2113 | Benign | -4.75 | Benign | 0.33 | Neutral | 0.52 | Possibly damaging | 0.29 | Tolerated | | |
| p.Gly136Asp | VUS | -22.04 | Indeterminate | 0.0963 | Benign | -4.26 | Benign | 0.15 | Neutral | 0.00 | Benign | 0.19 | Tolerated | 0.01 | Neutral |
| p.Gly136Glu | VUS | -0.28 | Neutral | 0.0992 | Benign | -3.98 | Benign | 0.19 | Neutral | 0.36 | Benign | 0.22 | Tolerated | | |
| p.Gly136Ala | VUS | -0.61 | Neutral | 0.09 | Benign | -3.42 | Benign | 0.12 | Neutral | 0.00 | Benign | 1 | Tolerated | 0.04 | Neutral |
| p.Gly136Val | VUS | -0.13 | Neutral | 0.1269 | Benign | -4.79 | Benign | 0.26 | Neutral | 0.29 | Benign | 0.35 | Tolerated | 7.01 | Neutral |
| p.Gly136Tyr | VUS | -2.12 | Neutral | 0.2855 | Benign | -7.52 | Pathogenic | 0.37 | Neutral | 0.89 | Possibly damaging | 0.27 | Tolerated | | |
| p.Gly136Cys | VUS | -9.10 | Indeterminate | 0.1768 | Benign | -6.13 | Benign | 0.32 | Neutral | 0.01 | Benign | 0.13 | Tolerated | 13.10 | Neutral |
| p.Gly136Trp | VUS | -3.05 | Neutral | 0.2066 | Benign | -6.85 | Benign | 0.34 | Neutral | 0.99 | Probably damaging | 0.06 | Tolerated | | |
| p.Gly136Phe | VUS | -3.24 | Neutral | 0.3221 | Benign | -7.08 | Benign | 0.34 | Neutral | 0.96 | Probably damaging | 0.21 | Tolerated | | |
| p.Thr137Asn | VUS | -0.55 | Neutral | 0.0965 | Benign | -4.57 | Benign | 0.23 | Neutral | 0.65 | Possibly damaging | 0.55 | Tolerated | 14.08 | Neutral |
| p.Thr137Lys | VUS | -0.52 | Neutral | 0.1405 | Benign | -4.41 | Benign | 0.35 | Neutral | 0.55 | Possibly damaging | 0.23 | Tolerated | | |
| p.Thr137Arg | VUS | -1.34 | Neutral | 0.0794 | Benign | -2.14 | Benign | 0.29 | Neutral | 0.02 | Benign | 0.15 | Tolerated | | |
| p.Thr137Ser | VUS | -0.38 | Neutral | 0.0779 | Benign | -2.45 | Benign | 0.16 | Neutral | 0.01 | Benign | 0.45 | Tolerated | 6.14 | Neutral |
| p.Thr137Ile | VUS | -3.13 | Neutral | 0.1468 | Benign | -5.11 | Benign | 0.35 | Neutral | 0.48 | Possibly damaging | 0.06 | Tolerated | 15.18 | Deleterious |
| p.Thr137Met | VUS | -0.62 | Neutral | 0.1281 | Benign | -5.43 | Benign | 0.37 | Neutral | 0.95 | Possibly damaging | 0.03 | Deleterious | | |
| p.Thr137His | VUS | -1.76 | Neutral | 0.1225 | Benign | -3.97 | Benign | 0.41 | Neutral | 0.99 | Probably damaging | 0.06 | Tolerated | | |
| p.Thr137Gln | VUS | -3.36 | Neutral | 0.1096 | Benign | -3.44 | Benign | 0.36 | Neutral | 0.90 | Possibly damaging | 0.36 | Tolerated | | |
| p.Thr137Pro | VUS | -0.52 | Neutral | 0.056 | Benign | -2.24 | Benign | 0.19 | Neutral | 0.01 | Benign | 0.1 | Tolerated | 6.04 | Neutral |
| p.Thr137Leu | VUS | -0.74 | Neutral | 0.0931 | Benign | -3.41 | Benign | 0.40 | Neutral | 0.34 | Benign | 0.08 | Tolerated | | |
| p.Thr137Asp | VUS | -1.83 | Neutral | 0.1409 | Benign | -2.83 | Benign | 0.40 | Neutral | 0.71 | Possibly damaging | 0.24 | Tolerated | | |
| p.Thr137Glu | VUS | -1.17 | Neutral | 0.1202 | Benign | -2.54 | Benign | 0.40 | Neutral | 0.71 | Possibly damaging | 0.2 | Tolerated | | |
| p.Thr137Ala | VUS | -3.04 | Neutral | 0.0728 | Benign | -2.00 | Benign | 0.13 | Neutral | 0.00 | Benign | 0.21 | Tolerated | 2.64 | Neutral |
| p.Thr137Gly | VUS | -0.57 | Neutral | 0.0918 | Benign | -1.59 | Benign | 0.43 | Neutral | 0.34 | Benign | 0.16 | Tolerated | | |
| p.Thr137Val | VUS | -0.45 | Neutral | 0.1196 | Benign | -3.47 | Benign | 0.39 | Neutral | 0.03 | Benign | 0.09 | Tolerated | | |
| p.Thr137Tyr | VUS | -0.56 | Neutral | 0.1962 | Benign | -6.09 | Benign | 0.42 | Neutral | 0.98 | Probably damaging | 0.04 | Deleterious | | |
| p.Thr137Cys | VUS | -6.22 | Indeterminate | 0.2876 | Benign | -4.48 | Benign | 0.41 | Neutral | 0.99 | Probably damaging | 0.03 | Deleterious | | |
| p.Thr137Trp | VUS | -0.80 | Neutral | 0.2606 | Benign | -4.81 | Benign | 0.47 | Neutral | 1.00 | Probably damaging | 0.01 | Deleterious | | |
| p.Thr137Phe | VUS | -0.80 | Neutral | 0.1667 | Benign | -5.57 | Benign | 0.43 | Neutral | 0.95 | Possibly damaging | 0.03 | Deleterious | | |
| p.Arg138Asn | VUS | -1.57 | Neutral | 0.2106 | Benign | -4.35 | Benign | 0.27 | Neutral | 0.00 | Benign | 0.34 | Tolerated | | |
| p.Arg138Lys | VUS | -0.03 | Neutral | 0.142 | Benign | -5.07 | Benign | 0.17 | Neutral | 0.00 | Benign | 1 | Tolerated | 0.10 | Neutral |
| p.Arg138Thr | VUS | -0.25 | Neutral | 0.0777 | Benign | -3.90 | Benign | 0.13 | Neutral | 0.00 | Benign | 0.38 | Tolerated | 0.07 | Neutral |
| p.Arg138Ser | VUS | 0.00 | Neutral | 0.1182 | Benign | -3.18 | Benign | 0.19 | Neutral | 0.00 | Benign | 0.43 | Tolerated | 7.95 | Neutral |
| p.Arg138Ile | VUS | -0.01 | Neutral | 0.1574 | Benign | -5.73 | Benign | 0.25 | Neutral | 0.01 | Benign | 0.16 | Tolerated | 6.18 | Neutral |
| p.Arg138Met | VUS | 0.00 | Neutral | 0.2112 | Benign | -5.98 | Benign | 0.26 | Neutral | 0.11 | Benign | 0.09 | Tolerated | | |
| p.Arg138His | VUS | -1.18 | Neutral | 0.0894 | Benign | -4.65 | Benign | 0.17 | Neutral | 0.07 | Benign | 0.17 | Tolerated | | |
| p.Arg138Gln | VUS | -0.05 | Neutral | 0.0888 | Benign | -4.23 | Benign | 0.12 | Neutral | 0.00 | Benign | 0.47 | Tolerated | | |
| p.Arg138Pro | VUS | -4.52 | Neutral | 0.081 | Benign | -2.99 | Benign | 0.24 | Neutral | 0.03 | Benign | 0.2 | Tolerated | | |
| p.Arg138Leu | VUS | 0.00 | Neutral | 0.1027 | Benign | -4.21 | Benign | 0.22 | Neutral | 0.00 | Benign | 0.27 | Tolerated | | |
| p.Arg138Asp | VUS | -0.07 | Neutral | 0.1354 | Benign | -3.15 | Benign | 0.17 | Neutral | 0.00 | Benign | 0.32 | Tolerated | | |
| p.Arg138Glu | VUS | -0.95 | Neutral | 0.1114 | Benign | -3.13 | Benign | 0.15 | Neutral | 0.00 | Benign | 0.57 | Tolerated | | |
| p.Arg138Ala | VUS | -1.36 | Neutral | 0.0753 | Benign | -2.56 | Benign | 0.26 | Neutral | 0.00 | Benign | 0.46 | Tolerated | | |
| p.Arg138Gly | VUS | -0.38 | Neutral | 0.0609 | Benign | -2.55 | Benign | 0.10 | Neutral | 0.00 | Benign | 0.26 | Tolerated | 9.06 | Neutral |
| p.Arg138Val | VUS | -0.15 | Neutral | 0.1281 | Benign | -4.09 | Benign | 0.29 | Neutral | 0.00 | Benign | 0.23 | Tolerated | | |
| p.Arg138Tyr | VUS | 0.00 | Neutral | 0.225 | Benign | -6.28 | Benign | 0.28 | Neutral | 0.18 | Benign | 0.14 | Tolerated | | |
| p.Arg138Cys | VUS | -0.01 | Neutral | 0.1701 | Benign | -5.08 | Benign | 0.23 | Neutral | 0.43 | Benign | 0.05 | Deleterious | | |
| p.Arg138Trp | VUS | 0.00 | Neutral | 0.1311 | Benign | -6.12 | Benign | 0.21 | Neutral | 0.43 | Benign | 0.04 | Deleterious | | |
| p.Arg138Phe | VUS | -0.11 | Neutral | 0.2553 | Benign | -6.12 | Benign | 0.29 | Neutral | 0.03 | Benign | 0.1 | Tolerated | | |
| p.Gly139Asn | VUS | -2.55 | Neutral | 0.1977 | Benign | -5.50 | Benign | 0.40 | Neutral | 0.30 | Benign | 0 | Deleterious | | |
| p.Gly139Lys | VUS | -4.57 | Neutral | 0.3065 | Benign | -5.76 | Benign | 0.43 | Neutral | 0.02 | Benign | 0 | Deleterious | | |
| p.Gly139Thr | VUS | -7.65 | Indeterminate | 0.1081 | Benign | -4.38 | Benign | 0.31 | Neutral | 0.18 | Benign | 0 | Deleterious | | |
| p.Gly139Arg | VUS | -1.61 | Neutral | 0.17 | Benign | -3.75 | Benign | 0.26 | Neutral | 0.00 | Benign | 0 | Deleterious | 10.44 | Neutral |
| p.Gly139Ser | VUS | -4.27 | Neutral | 0.1007 | Benign | -3.68 | Benign | 0.18 | Neutral | 0.00 | Benign | 0 | Deleterious | 4.19 | Neutral |
| p.Gly139Ile | VUS | -1.29 | Neutral | 0.1859 | Benign | -5.82 | Benign | 0.46 | Neutral | 0.18 | Benign | 0 | Deleterious | | |
| p.Gly139Met | VUS | -5.11 | Neutral | 0.3266 | Benign | -6.63 | Benign | 0.46 | Neutral | 0.90 | Possibly damaging | 0 | Deleterious | | |
| p.Gly139His | VUS | -1.68 | Neutral | 0.191 | Benign | -5.23 | Benign | 0.32 | Neutral | 0.00 | Benign | 0 | Deleterious | | |
| p.Gly139Gln | VUS | -1.02 | Neutral | 0.185 | Benign | -4.97 | Benign | 0.42 | Neutral | 0.04 | Benign | 0 | Deleterious | | |
| p.Gly139Pro | VUS | -6.78 | Indeterminate | 0.2134 | Benign | -3.36 | Benign | 0.42 | Neutral | 0.75 | Possibly damaging | 0 | Deleterious | | |
| p.Gly139Leu | VUS | -3.93 | Neutral | 0.2184 | Benign | -4.61 | Benign | 0.44 | Neutral | 0.18 | Benign | 0 | Deleterious | | |
| p.Gly139Asp | VUS | -7.08 | Indeterminate | 0.1074 | Benign | -4.34 | Benign | 0.27 | Neutral | 0.01 | Benign | 0 | Deleterious | 15.62 | Deleterious |
| p.Gly139Glu | VUS | -3.30 | Neutral | 0.1158 | Benign | -4.40 | Benign | 0.38 | Neutral | 0.15 | Benign | 0 | Deleterious | | |
| p.Gly139Ala | VUS | -7.05 | Indeterminate | 0.0943 | Benign | -3.44 | Benign | 0.19 | Neutral | 0.03 | Benign | 0 | Deleterious | 14.30 | Neutral |
| p.Gly139Val | VUS | -0.26 | Neutral | 0.1219 | Benign | -4.71 | Benign | 0.29 | Neutral | 0.01 | Benign | 0 | Deleterious | 15.89 | Deleterious |
| p.Gly139Tyr | VUS | -11.79 | Indeterminate | 0.2451 | Benign | -6.64 | Benign | 0.47 | Neutral | 0.60 | Possibly damaging | 0 | Deleterious | | |
| p.Gly139Cys | VUS | -0.55 | Neutral | 0.1649 | Benign | -5.70 | Benign | 0.42 | Neutral | 0.00 | Benign | 0 | Deleterious | 11.46 | Neutral |
| p.Gly139Trp | VUS | -2.00 | Neutral | 0.1747 | Benign | -6.41 | Benign | 0.44 | Neutral | 0.97 | Probably damaging | 0 | Deleterious | | |
| p.Gly139Phe | VUS | -4.58 | Neutral | 0.3282 | Benign | -6.45 | Benign | 0.45 | Neutral | 0.75 | Possibly damaging | 0 | Deleterious | | |
| p.Ser140Asn | VUS | -0.75 | Neutral | 0.1255 | Benign | -4.78 | Benign | 0.20 | Neutral | 0.07 | Benign | 0.86 | Tolerated | 10.99 | Neutral |
| p.Ser140Lys | VUS | -0.33 | Neutral | 0.3318 | Benign | -4.95 | Benign | 0.28 | Neutral | 0.05 | Benign | 0.48 | Tolerated | | |
| p.Ser140Thr | VUS | -0.31 | Neutral | 0.0958 | Benign | -4.00 | Benign | 0.19 | Neutral | 0.26 | Benign | 0.74 | Tolerated | 14.43 | Neutral |
| p.Ser140Arg | VUS | -3.36 | Neutral | 0.1902 | Benign | -3.15 | Benign | 0.21 | Neutral | 0.00 | Benign | 0.34 | Tolerated | 3.22 | Neutral |
| p.Ser140Ile | VUS | -0.10 | Neutral | 0.2008 | Benign | -5.33 | Benign | 0.30 | Neutral | 0.55 | Possibly damaging | 0.14 | Tolerated | 16.39 | Deleterious |
| p.Ser140Met | VUS | -0.98 | Neutral | 0.2592 | Benign | -6.10 | Benign | 0.34 | Neutral | 0.83 | Possibly damaging | 0.08 | Tolerated | | |
| p.Ser140His | VUS | -1.04 | Neutral | 0.1677 | Benign | -4.67 | Benign | 0.28 | Neutral | 0.00 | Benign | 0.33 | Tolerated | | |
| p.Ser140Gln | VUS | -0.44 | Neutral | 0.2021 | Benign | -4.32 | Benign | 0.28 | Neutral | 0.01 | Benign | 0.3 | Tolerated | | |
| p.Ser140Pro | VUS | -6.82 | Indeterminate | 0.0917 | Benign | -3.10 | Benign | 0.20 | Neutral | 0.59 | Possibly damaging | 0.26 | Tolerated | | |
| p.Ser140Leu | VUS | -0.36 | Neutral | 0.1651 | Benign | -4.23 | Benign | 0.21 | Neutral | 0.19 | Benign | 0.23 | Tolerated | | |
| p.Ser140Asp | VUS | -0.17 | Neutral | 0.1572 | Benign | -3.67 | Benign | 0.25 | Neutral | 0.00 | Benign | 0.73 | Tolerated | | |
| p.Ser140Glu | VUS | -0.32 | Neutral | 0.1861 | Benign | -3.54 | Benign | 0.28 | Neutral | 0.05 | Benign | 0.41 | Tolerated | | |
| p.Ser140Ala | VUS | -5.04 | Neutral | 0.0814 | Benign | -3.00 | Benign | 0.21 | Neutral | 0.02 | Benign | 0.55 | Tolerated | | |
| p.Ser140Gly | VUS | -7.24 | Indeterminate | 0.0608 | Benign | -2.61 | Benign | 0.11 | Neutral | 0.00 | Benign | 0.62 | Tolerated | 0.04 | Neutral |
| p.Ser140Val | VUS | -0.10 | Neutral | 0.1746 | Benign | -4.28 | Benign | 0.30 | Neutral | 0.32 | Benign | 0.22 | Tolerated | | |
| p.Ser140Tyr | VUS | -0.45 | Neutral | 0.1917 | Benign | -6.16 | Benign | 0.34 | Neutral | 0.45 | Benign | 0.25 | Tolerated | | |
| p.Ser140Cys | VUS | -0.72 | Neutral | 0.1042 | Benign | -5.25 | Benign | 0.26 | Neutral | 0.93 | Possibly damaging | 0.1 | Tolerated | 14.40 | Neutral |
| p.Ser140Trp | VUS | -1.21 | Neutral | 0.15 | Benign | -5.53 | Benign | 0.36 | Neutral | 0.95 | Possibly damaging | 0.06 | Tolerated | | |
| p.Ser140Phe | VUS | -1.17 | Neutral | 0.2242 | Benign | -6.05 | Benign | 0.31 | Neutral | 0.62 | Possibly damaging | 0.16 | Tolerated | | |

| | | | | | | | | | | | | | | | |
|---|---|---|---|---|---|---|---|---|---|---|---|---|---|---|---|
| p.Asn141Lys | VUS | -0.39 | Neutral | 0.2591 | Benign | -0.77 | Benign | 0.11 | Neutral | 0.01 | Benign | 0.72 | Tolerated | 11.59 | Neutral |
| p.Asn141Thr | VUS | -2.77 | Neutral | 0.0964 | Benign | -1.83 | Benign | 0.11 | Neutral | 0.01 | Benign | 0.25 | Tolerated | 1.26 | Neutral |
| p.Asn141Arg | VUS | -0.97 | Neutral | 0.1387 | Benign | -0.94 | Benign | 0.15 | Neutral | 0.00 | Benign | 1 | Tolerated | | |
| p.Asn141Ser | VUS | -0.22 | Neutral | 0.06 | Benign | -0.87 | Benign | 0.05 | Neutral | 0.00 | Benign | 0.31 | Tolerated | 0.01 | Neutral |
| p.Asn141Ile | VUS | -0.37 | Neutral | 0.1859 | Benign | -3.06 | Benign | 0.22 | Neutral | 0.01 | Benign | 0.08 | Tolerated | 9.60 | Neutral |
| p.Asn141Met | VUS | -0.27 | Neutral | 0.3507 | Ambiguous | -3.84 | Benign | 0.25 | Neutral | 0.10 | Benign | 0.05 | Deleterious | | |
| p.Asn141His | VUS | -2.15 | Neutral | 0.0972 | Benign | -2.50 | Benign | 0.15 | Neutral | 0.14 | Benign | 0.15 | Tolerated | 11.25 | Neutral |
| p.Asn141Gln | VUS | -0.80 | Neutral | 0.2108 | Benign | -2.04 | Benign | 0.13 | Neutral | 0.03 | Benign | 0.31 | Tolerated | | |
| p.Asn141Pro | VUS | -1.00 | Neutral | 0.1131 | Benign | -0.78 | Benign | 0.20 | Neutral | 0.03 | Benign | 0.14 | Tolerated | | |
| p.Asn141Leu | VUS | -0.14 | Neutral | 0.1576 | Benign | -1.46 | Benign | 0.21 | Neutral | 0.00 | Benign | 0.15 | Tolerated | | |
| p.Asn141Asp | VUS | -0.21 | Neutral | 0.10 | Benign | -1.68 | Benign | 0.07 | Neutral | 0.00 | Benign | 0.33 | Tolerated | 5.52 | Neutral |
| p.Asn141Glu | VUS | -1.28 | Neutral | 0.2015 | Benign | -1.38 | Benign | 0.13 | Neutral | 0.01 | Benign | 0.32 | Tolerated | | |
| p.Asn141Ala | VUS | -0.22 | Neutral | 0.1119 | Benign | -0.72 | Benign | 0.19 | Neutral | 0.00 | Benign | 0.25 | Tolerated | | |
| p.Asn141Gly | VUS | -0.33 | Neutral | 0.128 | Benign | -0.38 | Benign | 0.10 | Neutral | 0.00 | Benign | 0.28 | Tolerated | | |
| p.Asn141Val | VUS | -0.04 | Neutral | 0.1653 | Benign | -1.97 | Benign | 0.23 | Neutral | 0.01 | Benign | 0.11 | Tolerated | | |
| p.Asn141Tyr | VUS | -0.90 | Neutral | 0.14 | Benign | -4.40 | Benign | 0.19 | Neutral | 0.14 | Benign | 0.1 | Tolerated | 12.95 | Neutral |
| p.Asn141Cys | VUS | -2.57 | Neutral | 0.1935 | Benign | -2.75 | Benign | 0.24 | Neutral | 0.18 | Benign | 0.03 | Deleterious | | |
| p.Asn141Trp | VUS | -0.06 | Neutral | 0.4218 | Ambiguous | -3.02 | Benign | 0.28 | Neutral | 0.71 | Possibly damaging | 0.03 | Deleterious | | |
| p.Asn141Phe | VUS | -0.47 | Neutral | 0.3924 | Ambiguous | -3.45 | Benign | 0.25 | Neutral | 0.10 | Benign | 0.06 | Tolerated | | |
| p.His142Asn | VUS | 0.00 | Neutral | 0.1197 | Benign | -3.33 | Benign | 0.25 | Neutral | 0.02 | Benign | 1 | Tolerated | 11.88 | Neutral |
| p.His142Lys | VUS | 0.00 | Neutral | 0.2146 | Benign | -3.01 | Benign | 0.28 | Neutral | 0.01 | Benign | 0.17 | Tolerated | | |
| p.His142Thr | VUS | 0.00 | Neutral | 0.1444 | Benign | -2.18 | Benign | 0.30 | Neutral | 0.01 | Benign | 0.12 | Tolerated | | |
| p.His142Arg | VUS | 0.00 | Neutral | 0.0718 | Benign | -1.68 | Benign | 0.23 | Neutral | 0.00 | Benign | 0.11 | Tolerated | 0.02 | Neutral |
| p.His142Ser | VUS | 0.00 | Neutral | 0.1269 | Benign | -1.78 | Benign | 0.25 | Neutral | 0.00 | Benign | 0.22 | Tolerated | | |
| p.His142Ile | VUS | 0.00 | Neutral | 0.2557 | Benign | -3.13 | Benign | 0.31 | Neutral | 0.09 | Benign | 0.03 | Deleterious | | |
| p.His142Met | VUS | 0.00 | Neutral | 0.3804 | Ambiguous | -3.62 | Benign | 0.32 | Neutral | 0.52 | Possibly damaging | 0.02 | Deleterious | | |
| p.His142Gln | VUS | -0.01 | Neutral | 0.1289 | Benign | -2.77 | Benign | 0.19 | Neutral | 0.00 | Benign | 0.1 | Tolerated | 0.01 | Neutral |
| p.His142Pro | VUS | 0.00 | Neutral | 0.0702 | Benign | -1.34 | Benign | 0.18 | Neutral | 0.00 | Benign | 0.07 | Tolerated | 0.05 | Neutral |
| p.His142Leu | VUS | 0.00 | Neutral | 0.1061 | Benign | -1.95 | Benign | 0.31 | Neutral | 0.02 | Benign | 0.06 | Tolerated | 4.22 | Neutral |
| p.His142Asp | VUS | 0.00 | Neutral | 0.1004 | Benign | -2.16 | Benign | 0.22 | Neutral | 0.02 | Benign | 0.35 | Tolerated | 12.83 | Neutral |
| p.His142Glu | VUS | 0.00 | Neutral | 0.1352 | Benign | -1.77 | Benign | 0.27 | Neutral | 0.01 | Benign | 0.15 | Tolerated | | |
| p.His142Ala | VUS | 0.00 | Neutral | 0.1025 | Benign | -1.08 | Benign | 0.33 | Neutral | 0.01 | Benign | 0.11 | Tolerated | | |
| p.His142Gly | VUS | 0.00 | Neutral | 0.1126 | Benign | -0.62 | Benign | 0.26 | Neutral | 0.01 | Benign | 0.25 | Tolerated | | |
| p.His142Val | VUS | 0.00 | Neutral | 0.1537 | Benign | -2.18 | Benign | 0.32 | Neutral | 0.05 | Benign | 0.05 | Deleterious | | |
| p.His142Tyr | VUS | 0.00 | Neutral | 0.1167 | Benign | -4.44 | Benign | 0.29 | Neutral | 0.07 | Benign | 0.09 | Tolerated | 15.56 | Deleterious |
| p.His142Cys | VUS | 0.00 | Neutral | 0.2166 | Benign | -3.85 | Benign | 0.26 | Neutral | 0.81 | Possibly damaging | 0.03 | Deleterious | | |
| p.His142Trp | VUS | 0.00 | Neutral | 0.2488 | Benign | -4.24 | Benign | 0.32 | Neutral | 0.81 | Possibly damaging | 0.02 | Deleterious | | |
| p.His142Phe | VUS | -0.01 | Neutral | 0.2291 | Benign | -3.98 | Benign | 0.32 | Neutral | 0.27 | Benign | 0.05 | Deleterious | | |
| p.Ala143Asn | VUS | -1.04 | Neutral | 0.1998 | Benign | -4.62 | Benign | 0.35 | Neutral | 0.69 | Possibly damaging | 0.54 | Tolerated | | |
| p.Ala143Lys | VUS | -0.58 | Neutral | 0.2701 | Benign | -4.70 | Benign | 0.27 | Neutral | 0.69 | Possibly damaging | 0.67 | Tolerated | | |
| p.Ala143Thr | VUS | -0.48 | Neutral | 0.0847 | Benign | -4.30 | Benign | 0.28 | Neutral | 0.00 | Benign | 0.56 | Tolerated | 8.17 | Neutral |
| p.Ala143Arg | VUS | -0.80 | Neutral | 0.1332 | Benign | -3.14 | Benign | 0.35 | Neutral | 0.69 | Possibly damaging | 0.52 | Tolerated | | |
| p.Ala143Ser | VUS | -0.86 | Neutral | 0.0842 | Benign | -3.57 | Benign | 0.26 | Neutral | 0.19 | Benign | 0.72 | Tolerated | 12.41 | Neutral |
| p.Ala143Ile | VUS | -0.71 | Neutral | 0.2309 | Benign | -5.25 | Benign | 0.27 | Neutral | 0.24 | Benign | 0.33 | Tolerated | | |
| p.Ala143Met | VUS | -0.89 | Neutral | 0.2533 | Benign | -5.97 | Benign | 0.39 | Neutral | 0.94 | Possibly damaging | 0.18 | Tolerated | | |
| p.Ala143His | VUS | -1.17 | Neutral | 0.1867 | Benign | -4.21 | Benign | 0.37 | Neutral | 0.98 | Probably damaging | 0.42 | Tolerated | | |
| p.Ala143Gln | VUS | -0.91 | Neutral | 0.1507 | Benign | -4.16 | Benign | 0.33 | Neutral | 0.82 | Possibly damaging | 0.44 | Tolerated | | |
| p.Ala143Pro | VUS | -1.82 | Neutral | 0.0784 | Benign | -3.37 | Benign | 0.27 | Neutral | 0.77 | Possibly damaging | 0.34 | Tolerated | 16.83 | Deleterious |
| p.Ala143Leu | VUS | -0.93 | Neutral | 0.1214 | Benign | -4.00 | Benign | 0.37 | Neutral | 0.24 | Benign | 0.53 | Tolerated | | |
| p.Ala143Asp | VUS | -0.87 | Neutral | 0.118 | Benign | -3.80 | Benign | 0.38 | Neutral | 0.62 | Possibly damaging | 0.54 | Tolerated | 15.10 | Deleterious |
| p.Ala143Glu | VUS | -0.69 | Neutral | 0.117 | Benign | -3.79 | Benign | 0.33 | Neutral | 0.69 | Possibly damaging | 0.65 | Tolerated | | |
| p.Ala143Gly | VUS | -0.83 | Neutral | 0.0776 | Benign | -3.01 | Benign | 0.30 | Neutral | 0.32 | Benign | 0.61 | Tolerated | 14.00 | Neutral |
| p.Ala143Val | VUS | -0.68 | Neutral | 0.1113 | Benign | -4.29 | Benign | 0.28 | Neutral | 0.00 | Benign | 0.53 | Tolerated | 10.91 | Neutral |
| p.Ala143Tyr | VUS | -1.11 | Neutral | 0.2593 | Benign | -5.92 | Benign | 0.39 | Neutral | 0.93 | Possibly damaging | 0.53 | Tolerated | | |
| p.Ala143Cys | VUS | -1.23 | Neutral | 0.2989 | Benign | -5.09 | Benign | 0.39 | Neutral | 0.98 | Probably damaging | 0.16 | Tolerated | | |
| p.Ala143Trp | VUS | -1.68 | Neutral | 0.3151 | Benign | -5.77 | Benign | 0.44 | Neutral | 0.98 | Probably damaging | 0.11 | Tolerated | | |
| p.Ala143Phe | VUS | -0.74 | Neutral | 0.2193 | Benign | -5.72 | Benign | 0.40 | Neutral | 0.82 | Possibly damaging | 0.38 | Tolerated | | |
| p.Arg144Asn | VUS | -0.11 | Neutral | 0.2474 | Benign | -4.32 | Benign | 0.31 | Neutral | 0.01 | Benign | 0.09 | Tolerated | | |
| p.Arg144Lys | VUS | -0.74 | Neutral | 0.1626 | Benign | -4.96 | Benign | 0.20 | Neutral | 0.01 | Benign | 1 | Tolerated | | |
| p.Arg144Thr | VUS | -0.34 | Neutral | 0.1087 | Benign | -3.81 | Benign | 0.26 | Neutral | 0.01 | Benign | 0.1 | Tolerated | | |
| p.Arg144Ser | VUS | -0.41 | Neutral | 0.1479 | Benign | -3.10 | Benign | 0.17 | Neutral | 0.00 | Benign | 0.11 | Tolerated | 0.04 | Neutral |
| p.Arg144Ile | VUS | -0.67 | Neutral | 0.1729 | Benign | -4.85 | Benign | 0.23 | Neutral | 0.03 | Benign | 0.04 | Deleterious | | |
| p.Arg144Met | VUS | -0.35 | Neutral | 0.218 | Benign | -5.62 | Benign | 0.31 | Neutral | 0.39 | Benign | 0.03 | Deleterious | | |
| p.Arg144His | VUS | -0.56 | Neutral | 0.0933 | Benign | -4.43 | Benign | 0.11 | Neutral | 0.00 | Benign | 0.06 | Tolerated | 4.38 | Neutral |
| p.Arg144Gln | VUS | -0.43 | Neutral | 0.0911 | Benign | -4.18 | Benign | 0.17 | Neutral | 0.03 | Benign | 0.17 | Tolerated | | |
| p.Arg144Pro | VUS | -1.27 | Neutral | 0.0937 | Benign | -3.26 | Benign | 0.24 | Neutral | 0.00 | Benign | 0.07 | Tolerated | 6.21 | Neutral |
| p.Arg144Leu | VUS | -0.17 | Neutral | 0.1097 | Benign | -4.07 | Benign | 0.19 | Neutral | 0.01 | Benign | 0.08 | Tolerated | 2.32 | Neutral |
| p.Arg144Asp | VUS | -0.45 | Neutral | 0.1536 | Benign | -2.90 | Benign | 0.29 | Neutral | 0.01 | Benign | 0.08 | Tolerated | | |
| p.Arg144Glu | VUS | -0.46 | Neutral | 0.127 | Benign | -3.09 | Benign | 0.26 | Neutral | 0.01 | Benign | 0.66 | Tolerated | | |
| p.Arg144Ala | VUS | -0.38 | Neutral | 0.1021 | Benign | -2.55 | Benign | 0.27 | Neutral | 0.00 | Benign | 0.12 | Tolerated | | |
| p.Arg144Gly | VUS | -0.84 | Neutral | 0.0753 | Benign | -2.52 | Benign | 0.14 | Neutral | 0.00 | Benign | 0.07 | Tolerated | 0.14 | Neutral |
| p.Arg144Val | VUS | -0.16 | Neutral | 0.1463 | Benign | -3.69 | Benign | 0.25 | Neutral | 0.01 | Benign | 0.05 | Deleterious | | |
| p.Arg144Tyr | VUS | -0.32 | Neutral | 0.2413 | Benign | -5.87 | Benign | 0.27 | Neutral | 0.02 | Benign | 0.03 | Deleterious | | |
| p.Arg144Cys | Likely benign | -0.03 | Neutral | 0.1551 | Benign | -5.12 | Benign | 0.26 | Neutral | 0.00 | Benign | 0.01 | Deleterious | 4.63 | Neutral |
| p.Arg144Trp | VUS | -0.28 | Neutral | 0.1153 | Benign | -5.87 | Benign | 0.27 | Neutral | 0.71 | Possibly damaging | 0.01 | Deleterious | | |
| p.Arg144Phe | VUS | -0.11 | Neutral | 0.2614 | Benign | -5.77 | Benign | 0.34 | Neutral | 0.03 | Benign | 0.02 | Deleterious | | |
| p.Ile145Asn | VUS | -0.68 | Neutral | 0.1287 | Benign | -2.73 | Benign | 0.13 | Neutral | 0.03 | Benign | 0.13 | Tolerated | | |
| p.Ile145Lys | VUS | -0.74 | Neutral | 0.2074 | Benign | -2.37 | Benign | 0.25 | Neutral | 0.02 | Benign | 0.16 | Tolerated | 9.74 | Neutral |
| p.Ile145Thr | VUS | -1.03 | Neutral | 0.1185 | Benign | -1.66 | Benign | 0.09 | Neutral | 0.00 | Benign | 0.2 | Tolerated | 5.59 | Neutral |
| p.Ile145Arg | VUS | -1.68 | Neutral | 0.1187 | Benign | -0.84 | Benign | 0.19 | Neutral | 0.05 | Benign | 0.12 | Tolerated | 8.58 | Neutral |
| p.Ile145Ser | VUS | -0.64 | Neutral | 0.1266 | Benign | -1.11 | Benign | 0.12 | Neutral | 0.00 | Benign | 0.29 | Tolerated | | |
| p.Ile145Met | VUS | -3.12 | Neutral | 0.0964 | Benign | -3.42 | Benign | 0.13 | Neutral | 0.29 | Benign | 0.13 | Tolerated | 0.23 | Neutral |
| p.Ile145His | VUS | -0.55 | Neutral | 0.2337 | Benign | -2.10 | Benign | 0.20 | Neutral | 0.62 | Possibly damaging | 0.06 | Tolerated | | |
| p.Ile145Gln | VUS | -1.06 | Neutral | 0.1739 | Benign | -1.49 | Benign | 0.19 | Neutral | 0.13 | Benign | 0.1 | Tolerated | | |
| p.Ile145Pro | VUS | -0.38 | Neutral | 0.1772 | Benign | -0.58 | Benign | 0.21 | Neutral | 0.00 | Benign | 0.14 | Tolerated | | |
| p.Ile145Leu | VUS | -0.77 | Neutral | 0.1002 | Benign | -1.63 | Benign | 0.22 | Neutral | 0.01 | Benign | 0.61 | Tolerated | 0.07 | Neutral |
| p.Ile145Asp | VUS | -1.09 | Neutral | 0.2116 | Benign | -0.72 | Benign | 0.30 | Neutral | 0.03 | Benign | 0.13 | Tolerated | | |
| p.Ile145Glu | VUS | -0.60 | Neutral | 0.1666 | Benign | -0.54 | Benign | 0.30 | Neutral | 0.03 | Benign | 0.13 | Tolerated | | |
| p.Ile145Ala | VUS | -3.86 | Neutral | 0.1271 | Benign | -0.29 | Benign | 0.18 | Neutral | 0.00 | Benign | 0.47 | Tolerated | | |
| p.Ile145Gly | VUS | -0.33 | Neutral | 0.2128 | Benign | -0.19 | Benign | 0.17 | Neutral | 0.00 | Benign | 0.3 | Tolerated | | |
| p.Ile145Val | VUS | -2.24 | Neutral | 0.0996 | Benign | -1.76 | Benign | 0.12 | Neutral | 0.00 | Benign | 0.97 | Tolerated | 0.00 | Neutral |
| p.Ile145Tyr | VUS | -0.82 | Neutral | 0.2778 | Benign | -4.33 | Benign | 0.30 | Neutral | 0.32 | Benign | 0.12 | Tolerated | | |
| p.Ile145Cys | VUS | -0.23 | Neutral | 0.4874 | Ambiguous | -2.99 | Benign | 0.29 | Neutral | 0.35 | Benign | 0.07 | Tolerated | | |
| p.Ile145Trp | VUS | -1.01 | Neutral | 0.3666 | Ambiguous | -3.77 | Benign | 0.32 | Neutral | 0.86 | Possibly damaging | 0.03 | Deleterious | | |
| p.Ile145Phe | VUS | -0.95 | Neutral | 0.1128 | Benign | -3.99 | Benign | 0.25 | Neutral | 0.13 | Benign | 0.16 | Tolerated | | |
| p.Asp146Asn | VUS | -3.08 | Neutral | 0.148 | Benign | -4.76 | Benign | 0.13 | Neutral | 0.00 | Benign | 0 | Deleterious | 6.95 | Neutral |
| p.Asp146Lys | VUS | -0.48 | Neutral | 0.2876 | Benign | -4.30 | Benign | 0.35 | Neutral | 0.03 | Benign | 0 | Deleterious | | |
| p.Asp146Thr | VUS | -2.33 | Neutral | 0.161 | Benign | -3.26 | Benign | 0.23 | Neutral | 0.03 | Benign | 0 | Deleterious | | |
| p.Asp146Arg | VUS | -0.62 | Neutral | 0.1628 | Benign | -2.67 | Benign | 0.23 | Neutral | 0.00 | Benign | 0 | Deleterious | | |
| p.Asp146Ser | VUS | -0.81 | Neutral | 0.1182 | Benign | -2.77 | Benign | 0.25 | Neutral | 0.00 | Benign | 0 | Deleterious | | |
| p.Asp146Ile | VUS | -0.60 | Neutral | 0.2896 | Benign | -4.82 | Benign | 0.34 | Neutral | 0.03 | Benign | 0 | Deleterious | | |
| p.Asp146Met | VUS | -1.10 | Neutral | 0.5411 | Ambiguous | -5.12 | Benign | 0.38 | Neutral | 0.27 | Benign | 0 | Deleterious | | |
| p.Asp146His | VUS | -2.20 | Neutral | 0.1334 | Benign | -4.22 | Benign | 0.28 | Neutral | 0.12 | Benign | 0 | Deleterious | 15.17 | Deleterious |
| p.Asp146Gln | VUS | -2.14 | Neutral | 0.1883 | Benign | -3.57 | Benign | 0.35 | Neutral | 0.08 | Benign | 0 | Deleterious | | |
| p.Asp146Pro | VUS | -1.91 | Neutral | 0.2096 | Benign | -2.18 | Benign | 0.34 | Neutral | 0.27 | Benign | 0 | Deleterious | | |
| p.Asp146Leu | VUS | -1.92 | Neutral | 0.2342 | Benign | -3.28 | Benign | 0.37 | Neutral | 0.03 | Benign | 0 | Deleterious | | |
| p.Asp146Glu | VUS | -0.72 | Neutral | 0.1395 | Benign | -2.93 | Benign | 0.09 | Neutral | 0.00 | Benign | 0 | Deleterious | 0.64 | Neutral |
| p.Asp146Ala | VUS | -1.04 | Neutral | 0.0971 | Benign | -2.12 | Benign | 0.13 | Neutral | 0.00 | Benign | 0 | Deleterious | 9.20 | Neutral |
| p.Asp146Gly | VUS | -0.79 | Neutral | 0.1138 | Benign | -2.52 | Benign | 0.16 | Neutral | 0.00 | Benign | 0 | Deleterious | 11.57 | Neutral |
| p.Asp146Val | VUS | -0.77 | Neutral | 0.1656 | Benign | -3.61 | Benign | 0.30 | Neutral | 0.00 | Benign | 0 | Deleterious | 14.02 | Neutral |
| p.Asp146Tyr | VUS | -3.39 | Neutral | 0.1614 | Benign | -5.88 | Benign | 0.28 | Neutral | 0.00 | Benign | 0 | Deleterious | 11.95 | Neutral |

| | | | | | | | | | | | | | | | |
|---|---|---|---|---|---|---|---|---|---|---|---|---|---|---|---|
| p.Asp146Cys | VUS | -3.78 | Neutral | 0.3974 | Ambiguous | -4.71 | Benign | 0.37 | Neutral | 0.58 | Possibly damaging | 0 | Deleterious | | |
| p.Asp146Trp | VUS | -0.56 | Neutral | 0.5798 | Pathogenic | -5.33 | Benign | 0.37 | Neutral | 0.58 | Possibly damaging | 0 | Deleterious | | |
| p.Asp146Phe | VUS | -1.81 | Neutral | 0.3732 | Ambiguous | -5.37 | Benign | 0.35 | Neutral | 0.08 | Benign | 0 | Deleterious | | |
| p.Ala147Asn | VUS | -0.93 | Neutral | 0.2019 | Benign | -5.67 | Benign | 0.38 | Neutral | 0.45 | Benign | 0.17 | Tolerated | | |
| p.Ala147Lys | VUS | -1.03 | Neutral | 0.2553 | Benign | -5.44 | Benign | 0.39 | Neutral | 0.19 | Benign | 0.18 | Tolerated | | |
| p.Ala147Thr | VUS | -0.92 | Neutral | 0.0804 | Benign | -4.49 | Benign | 0.16 | Neutral | 0.08 | Benign | 0.91 | Tolerated | 15.28 | Deleterious |
| p.Ala147Arg | VUS | -0.26 | Neutral | 0.1538 | Benign | -4.27 | Benign | 0.29 | Neutral | 0.45 | Benign | 0.12 | Tolerated | | |
| p.Ala147Ser | VUS | -1.41 | Neutral | 0.0998 | Benign | -4.21 | Benign | 0.15 | Neutral | 0.00 | Benign | 0.7 | Tolerated | 7.95 | Neutral |
| p.Ala147Ile | VUS | -0.68 | Neutral | 0.1893 | Benign | -5.47 | Benign | 0.38 | Neutral | 0.10 | Benign | 0.3 | Tolerated | | |
| p.Ala147Met | VUS | -1.38 | Neutral | 0.2114 | Benign | -6.10 | Benign | 0.36 | Neutral | 0.62 | Possibly damaging | 0.1 | Tolerated | | |
| p.Ala147His | VUS | -0.23 | Neutral | 0.2085 | Benign | -5.71 | Benign | 0.29 | Neutral | 0.95 | Possibly damaging | 0.06 | Tolerated | | |
| p.Ala147Gln | VUS | -5.59 | Neutral | 0.1828 | Benign | -5.29 | Benign | 0.38 | Neutral | 0.45 | Benign | 0.13 | Tolerated | | |
| p.Ala147Pro | VUS | -1.86 | Neutral | 0.0928 | Benign | -3.80 | Benign | 0.21 | Neutral | 0.00 | Benign | 0.24 | Tolerated | 12.40 | Neutral |
| p.Ala147Leu | VUS | -1.49 | Neutral | 0.1278 | Benign | -4.42 | Benign | 0.37 | Neutral | 0.05 | Benign | 0.27 | Tolerated | | |
| p.Ala147Asp | VUS | -2.43 | Neutral | 0.1068 | Benign | -4.04 | Benign | 0.25 | Neutral | 0.15 | Benign | 0.13 | Tolerated | 14.09 | Neutral |
| p.Ala147Glu | VUS | -0.24 | Neutral | 0.1148 | Benign | -3.75 | Benign | 0.35 | Neutral | 0.19 | Benign | 0.16 | Tolerated | | |
| p.Ala147Gly | VUS | -0.97 | Neutral | 0.09 | Benign | -3.42 | Benign | 0.18 | Neutral | 0.00 | Benign | 0.34 | Tolerated | 9.12 | Neutral |
| p.Ala147Val | VUS | -1.68 | Neutral | 0.1016 | Benign | -4.38 | Benign | 0.17 | Neutral | 0.00 | Benign | 0.64 | Tolerated | 9.43 | Neutral |
| p.Ala147Tyr | VUS | -0.40 | Neutral | 0.2901 | Benign | -6.98 | Benign | 0.31 | Neutral | 0.83 | Possibly damaging | 0.08 | Tolerated | | |
| p.Ala147Cys | VUS | -0.81 | Neutral | 0.3345 | Benign | -6.15 | Benign | 0.27 | Neutral | 0.95 | Possibly damaging | 0.23 | Tolerated | | |
| p.Ala147Trp | VUS | -0.55 | Neutral | 0.3676 | Ambiguous | -6.70 | Benign | 0.40 | Neutral | 0.95 | Possibly damaging | 0.03 | Deleterious | | |
| p.Ala147Phe | VUS | -0.22 | Neutral | 0.2218 | Benign | -6.35 | Benign | 0.39 | Neutral | 0.62 | Possibly damaging | 0.09 | Tolerated | | |
| p.Ala148Asn | VUS | -6.43 | Indeterminate | 0.2079 | Benign | -5.63 | Benign | 0.46 | Neutral | 0.77 | Possibly damaging | 0 | Deleterious | | |
| p.Ala148Lys | VUS | -26.41 | Indeterminate | 0.2504 | Benign | -5.49 | Benign | 0.48 | Neutral | 0.33 | Benign | 0 | Deleterious | | |
| p.Ala148Thr | Likely benign | -1.32 | Neutral | 0.0801 | Benign | -4.63 | Benign | 0.12 | Neutral | 0.80 | Possibly damaging | 0 | Deleterious | 15.70 | Deleterious |
| p.Ala148Arg | VUS | -8.95 | Indeterminate | 0.1519 | Benign | -4.24 | Benign | 0.36 | Neutral | 0.63 | Possibly damaging | 0 | Deleterious | | |
| p.Ala148Ser | VUS | -2.30 | Neutral | 0.0873 | Benign | -4.09 | Benign | 0.17 | Neutral | 0.07 | Benign | 0 | Deleterious | 14.77 | Neutral |
| p.Ala148Ile | VUS | -3.57 | Neutral | 0.2128 | Benign | -5.17 | Benign | 0.46 | Neutral | 0.63 | Possibly damaging | 0 | Deleterious | | |
| p.Ala148Met | VUS | -12.46 | Indeterminate | 0.219 | Benign | -5.53 | Benign | 0.46 | Neutral | 0.96 | Probably damaging | 0 | Deleterious | | |
| p.Ala148His | VUS | -6.12 | Indeterminate | 0.2355 | Benign | -5.65 | Benign | 0.45 | Neutral | 0.96 | Probably damaging | 0 | Deleterious | | |
| p.Ala148Gln | VUS | -9.43 | Indeterminate | 0.1675 | Benign | -4.87 | Benign | 0.31 | Neutral | 0.03 | Benign | 0 | Deleterious | | |
| p.Ala148Pro | VUS | -4.20 | Neutral | 0.0883 | Benign | -3.89 | Benign | 0.48 | Neutral | 0.01 | Benign | 0 | Deleterious | 15.93 | Deleterious |
| p.Ala148Leu | VUS | -4.52 | Neutral | 0.1365 | Benign | -4.13 | Benign | 0.47 | Neutral | 0.33 | Benign | 0 | Deleterious | | |
| p.Ala148Asp | VUS | -3.31 | Neutral | 0.1202 | Benign | -3.97 | Benign | 0.48 | Neutral | 0.33 | Benign | 0 | Deleterious | | |
| p.Ala148Glu | VUS | -7.75 | Indeterminate | 0.1156 | Benign | -3.57 | Benign | 0.26 | Neutral | 0.01 | Benign | 0 | Deleterious | 13.03 | Neutral |
| p.Ala148Gly | VUS | -1.38 | Neutral | 0.0837 | Benign | -3.48 | Benign | 0.22 | Neutral | 0.01 | Benign | 0 | Deleterious | 10.10 | Neutral |
| p.Ala148Val | VUS | -10.84 | Indeterminate | 0.1065 | Benign | -4.20 | Benign | 0.16 | Neutral | 0.03 | Benign | 0 | Deleterious | 9.76 | Neutral |
| p.Ala148Tyr | VUS | -0.81 | Neutral | 0.3042 | Benign | -6.69 | Benign | 0.47 | Neutral | 0.87 | Possibly damaging | 0 | Deleterious | | |
| p.Ala148Cys | VUS | -0.04 | Neutral | 0.3346 | Benign | -5.91 | Benign | 0.41 | Neutral | 0.99 | Probably damaging | 0 | Deleterious | | |
| p.Ala148Trp | VUS | -23.29 | Indeterminate | 0.3527 | Ambiguous | -6.33 | Benign | 0.51 | Disease | 0.99 | Probably damaging | 0 | Deleterious | | |
| p.Ala148Phe | VUS | -24.04 | Indeterminate | 0.2336 | Benign | -6.05 | Benign | 0.49 | Neutral | 0.77 | Possibly damaging | 0 | Deleterious | | |
| p.Glu149Asn | VUS | -0.21 | Neutral | 0.2363 | Benign | -4.72 | Benign | 0.38 | Neutral | 0.12 | Benign | 0.08 | Tolerated | | |
| p.Glu149Lys | VUS | -1.24 | Neutral | 0.1483 | Benign | -5.09 | Benign | 0.35 | Neutral | 0.04 | Benign | 0.12 | Tolerated | 16.13 | Deleterious |
| p.Glu149Thr | VUS | -3.47 | Neutral | 0.1487 | Benign | -3.51 | Benign | 0.39 | Neutral | 0.05 | Benign | 0.06 | Tolerated | | |
| p.Glu149Arg | VUS | -0.32 | Neutral | 0.1196 | Benign | -3.11 | Benign | 0.37 | Neutral | 0.01 | Benign | 0.04 | Deleterious | | |
| p.Glu149Ser | VUS | -5.62 | Neutral | 0.1372 | Benign | -3.06 | Benign | 0.39 | Neutral | 0.03 | Benign | 0.08 | Tolerated | | |
| p.Glu149Ile | VUS | -3.18 | Neutral | 0.3233 | Benign | -5.12 | Benign | 0.46 | Neutral | 0.00 | Benign | 0.02 | Deleterious | | |
| p.Glu149Met | VUS | -0.48 | Neutral | 0.4196 | Ambiguous | -5.39 | Benign | 0.34 | Neutral | 0.01 | Benign | 0.01 | Deleterious | | |
| p.Glu149His | VUS | -1.17 | Neutral | 0.215 | Benign | -4.58 | Benign | 0.25 | Neutral | 0.74 | Possibly damaging | 0.03 | Deleterious | | |
| p.Glu149Gln | VUS | -0.22 | Neutral | 0.1027 | Benign | -4.20 | Benign | 0.15 | Neutral | 0.17 | Benign | 0.13 | Tolerated | 10.92 | Neutral |
| p.Glu149Pro | VUS | -0.16 | Neutral | 0.1886 | Benign | -2.71 | Benign | 0.43 | Neutral | 0.00 | Benign | 0.06 | Tolerated | | |
| p.Glu149Leu | VUS | -0.12 | Neutral | 0.258 | Benign | -3.72 | Benign | 0.45 | Neutral | 0.01 | Benign | 0.03 | Deleterious | | |
| p.Glu149Asp | VUS | -1.68 | Neutral | 0.1 | Benign | -3.21 | Benign | 0.25 | Neutral | 0.04 | Benign | 0.76 | Tolerated | 6.26 | Neutral |
| p.Glu149Ala | VUS | -2.07 | Neutral | 0.0853 | Benign | -2.16 | Benign | 0.23 | Neutral | 0.00 | Benign | 0.11 | Tolerated | 11.61 | Neutral |
| p.Glu149Gly | VUS | -1.58 | Neutral | 0.0873 | Benign | -2.74 | Benign | 0.19 | Neutral | 0.00 | Benign | 0.06 | Tolerated | 12.78 | Neutral |
| p.Glu149Val | VUS | -1.34 | Neutral | 0.1769 | Benign | -3.95 | Benign | 0.42 | Neutral | 0.00 | Benign | 0.03 | Deleterious | 16.59 | Deleterious |
| p.Glu149Tyr | VUS | -1.10 | Neutral | 0.3748 | Ambiguous | -5.81 | Benign | 0.43 | Neutral | 0.74 | Possibly damaging | 0.01 | Deleterious | | |
| p.Glu149Cys | VUS | -1.40 | Neutral | 0.5637 | Ambiguous | -4.96 | Benign | 0.33 | Neutral | 0.91 | Possibly damaging | 0 | Deleterious | | |
| p.Glu149Trp | VUS | -1.72 | Neutral | 0.5825 | Pathogenic | -6.09 | Benign | 0.46 | Neutral | 0.91 | Possibly damaging | 0 | Deleterious | | |
| p.Glu149Phe | VUS | -4.67 | Neutral | 0.4663 | Ambiguous | -5.34 | Benign | 0.46 | Neutral | 0.31 | Benign | 0.01 | Deleterious | | |
| p.Gly150Asn | VUS | -0.22 | Neutral | 0.1638 | Benign | -5.06 | Benign | 0.39 | Neutral | 0.02 | Benign | 0.41 | Tolerated | | |
| p.Gly150Lys | VUS | -2.08 | Neutral | 0.2131 | Benign | -5.12 | Benign | 0.41 | Neutral | 0.51 | Possibly damaging | 1 | Tolerated | | |
| p.Gly150Thr | VUS | -1.15 | Neutral | 0.0936 | Benign | -4.15 | Benign | 0.27 | Neutral | 0.51 | Possibly damaging | 0.27 | Tolerated | | |
| p.Gly150Arg | VUS | -0.34 | Neutral | 0.1683 | Benign | -3.78 | Benign | 0.41 | Neutral | 0.03 | Benign | 0.47 | Tolerated | 14.19 | Neutral |
| p.Gly150Ser | VUS | -1.31 | Neutral | 0.0985 | Benign | -3.67 | Benign | 0.14 | Neutral | 0.05 | Benign | 0.42 | Tolerated | 10.05 | Neutral |
| p.Gly150Ile | VUS | -0.16 | Neutral | 0.1346 | Benign | -4.58 | Benign | 0.44 | Neutral | 0.94 | Possibly damaging | 0.09 | Tolerated | | |
| p.Gly150Met | VUS | -0.78 | Neutral | 0.231 | Benign | -5.39 | Benign | 0.45 | Neutral | 0.99 | Probably damaging | 0.06 | Tolerated | | |
| p.Gly150His | VUS | -1.35 | Neutral | 0.1691 | Benign | -5.13 | Benign | 0.32 | Neutral | 0.98 | Probably damaging | 0.14 | Tolerated | | |
| p.Gly150Gln | VUS | -1.46 | Neutral | 0.1539 | Benign | -4.40 | Benign | 0.38 | Neutral | 0.88 | Possibly damaging | 0.32 | Tolerated | | |
| p.Gly150Pro | VUS | -0.78 | Neutral | 0.2298 | Benign | -3.31 | Benign | 0.39 | Neutral | 0.94 | Possibly damaging | 0.19 | Tolerated | | |
| p.Gly150Leu | VUS | -0.48 | Neutral | 0.1639 | Benign | -3.85 | Benign | 0.30 | Neutral | 0.88 | Possibly damaging | 0.17 | Tolerated | | |
| p.Gly150Asp | VUS | -1.76 | Neutral | 0.0881 | Benign | -3.69 | Benign | 0.29 | Neutral | 0.01 | Benign | 0.39 | Tolerated | 10.58 | Neutral |
| p.Gly150Glu | VUS | -0.18 | Neutral | 0.084 | Benign | -3.41 | Benign | 0.37 | Neutral | 0.51 | Possibly damaging | 0.42 | Tolerated | | |
| p.Gly150Ala | VUS | -0.14 | Neutral | 0.0866 | Benign | -3.32 | Benign | 0.13 | Neutral | 0.06 | Benign | 0.47 | Tolerated | 8.81 | Neutral |
| p.Gly150Val | VUS | -0.76 | Neutral | 0.0977 | Benign | -3.95 | Benign | 0.29 | Neutral | 0.85 | Possibly damaging | 0.14 | Tolerated | 17.64 | Deleterious |
| p.Gly150Tyr | VUS | -0.13 | Neutral | 0.1895 | Benign | -6.43 | Benign | 0.44 | Neutral | 0.98 | Probably damaging | 0.09 | Tolerated | | |
| p.Gly150Cys | VUS | -0.02 | Neutral | 0.1781 | Benign | -5.86 | Benign | 0.43 | Neutral | 0.99 | Probably damaging | 0.05 | Deleterious | 15.99 | Deleterious |
| p.Gly150Trp | VUS | -0.49 | Neutral | 0.1743 | Benign | -6.25 | Benign | 0.44 | Neutral | 0.99 | Probably damaging | 0.03 | Deleterious | | |
| p.Gly150Phe | VUS | -0.87 | Neutral | 0.2684 | Benign | -5.52 | Benign | 0.44 | Neutral | 0.98 | Probably damaging | 0.06 | Tolerated | | |
| p.Pro151Asn | VUS | -0.72 | Neutral | 0.1899 | Benign | -4.86 | Benign | 0.44 | Neutral | 0.77 | Possibly damaging | 0.41 | Tolerated | | |
| p.Pro151Lys | VUS | -0.10 | Neutral | 0.1679 | Benign | -5.17 | Benign | 0.48 | Neutral | 0.49 | Possibly damaging | 0.45 | Tolerated | | |
| p.Pro151Thr | VUS | -1.96 | Neutral | 0.0763 | Benign | -3.99 | Benign | 0.20 | Neutral | 0.00 | Benign | 0.46 | Tolerated | 9.37 | Neutral |
| p.Pro151Arg | VUS | -0.63 | Neutral | 0.0798 | Benign | -3.63 | Benign | 0.29 | Neutral | 0.72 | Possibly damaging | 0.42 | Tolerated | 14.93 | Neutral |
| p.Pro151Ser | VUS | -15.51 | Indeterminate | 0.0798 | Benign | -3.16 | Benign | 0.24 | Neutral | 0.01 | Benign | 0.54 | Tolerated | 9.05 | Neutral |
| p.Pro151Ile | VUS | -0.49 | Neutral | 0.1737 | Benign | -4.99 | Benign | 0.51 | Disease | 0.63 | Possibly damaging | 0.37 | Tolerated | | |
| p.Pro151Met | VUS | -0.07 | Neutral | 0.2229 | Benign | -5.77 | Benign | 0.49 | Neutral | 0.96 | Probably damaging | 0.2 | Tolerated | | |
| p.Pro151His | VUS | -0.88 | Neutral | 0.0925 | Benign | -4.94 | Benign | 0.28 | Neutral | 0.01 | Benign | 0.42 | Tolerated | 14.78 | Neutral |
| p.Pro151Gln | VUS | -0.19 | Neutral | 0.0882 | Benign | -4.84 | Benign | 0.41 | Neutral | 0.77 | Possibly damaging | 0.33 | Tolerated | | |
| p.Pro151Leu | VUS | -3.56 | Neutral | 0.1004 | Benign | -4.02 | Benign | 0.26 | Neutral | 0.27 | Benign | 0.64 | Tolerated | 15.41 | Deleterious |
| p.Pro151Asp | VUS | -2.62 | Neutral | 0.1412 | Benign | -3.26 | Benign | 0.48 | Neutral | 0.49 | Possibly damaging | 0.36 | Tolerated | | |
| p.Pro151Glu | VUS | -0.84 | Neutral | 0.1103 | Benign | -3.74 | Benign | 0.44 | Neutral | 0.49 | Possibly damaging | 0.4 | Tolerated | | |
| p.Pro151Ala | VUS | -2.27 | Neutral | 0.063 | Benign | -3.15 | Benign | 0.32 | Neutral | 0.13 | Benign | 0.67 | Tolerated | 11.33 | Neutral |
| p.Pro151Gly | VUS | -8.60 | Indeterminate | 0.1164 | Benign | -2.83 | Benign | 0.44 | Neutral | 0.33 | Benign | 0.47 | Tolerated | | |
| p.Pro151Val | VUS | -0.18 | Neutral | 0.1199 | Benign | -4.25 | Benign | 0.31 | Neutral | 0.03 | Benign | 0.48 | Tolerated | | |
| p.Pro151Tyr | VUS | -0.04 | Neutral | 0.204 | Benign | -5.61 | Benign | 0.49 | Neutral | 0.77 | Possibly damaging | 1 | Tolerated | | |
| p.Pro151Cys | VUS | -0.48 | Neutral | 0.2543 | Benign | -5.01 | Benign | 0.49 | Neutral | 0.99 | Probably damaging | 0.16 | Tolerated | | |
| p.Pro151Trp | VUS | -0.23 | Neutral | 0.2269 | Benign | -5.83 | Benign | 0.51 | Disease | 0.99 | Probably damaging | 0.21 | Tolerated | | |
| p.Pro151Phe | VUS | -0.37 | Neutral | 0.1909 | Benign | -5.15 | Benign | 0.51 | Disease | 0.87 | Possibly damaging | 0.84 | Tolerated | | |
| p.Ser152Asn | VUS | -3.22 | Neutral | 0.1709 | Benign | -4.88 | Benign | 0.21 | Neutral | 0.19 | Benign | 0.46 | Tolerated | | |
| p.Ser152Lys | VUS | -2.25 | Neutral | 0.3202 | Benign | -4.52 | Benign | 0.31 | Neutral | 0.09 | Benign | 0.28 | Tolerated | | |
| p.Ser152Thr | VUS | -2.54 | Neutral | 0.093 | Benign | -3.89 | Benign | 0.11 | Neutral | 0.00 | Benign | 0.38 | Tolerated | 3.25 | Neutral |
| p.Ser152Arg | VUS | -1.68 | Neutral | 0.204 | Benign | -3.04 | Benign | 0.29 | Neutral | 0.19 | Benign | 0.18 | Tolerated | | |
| p.Ser152Ile | VUS | -0.47 | Neutral | 0.1923 | Benign | -4.47 | Benign | 0.33 | Neutral | 0.19 | Benign | 0.08 | Tolerated | | |
| p.Ser152Met | VUS | -3.57 | Neutral | 0.2632 | Benign | -4.60 | Benign | 0.36 | Neutral | 0.83 | Possibly damaging | 0.06 | Tolerated | | |
| p.Ser152His | VUS | -0.63 | Neutral | 0.1793 | Benign | -4.84 | Benign | 0.34 | Neutral | 0.83 | Possibly damaging | 0.11 | Tolerated | | |
| p.Ser152Gln | VUS | -0.32 | Neutral | 0.219 | Benign | -3.80 | Benign | 0.30 | Neutral | 0.32 | Benign | 0.19 | Tolerated | | |
| p.Ser152Pro | VUS | -2.64 | Neutral | 0.0892 | Benign | -3.07 | Benign | 0.15 | Neutral | 0.00 | Benign | 0.29 | Tolerated | 0.77 | Neutral |
| p.Ser152Leu | VUS | -3.01 | Neutral | 0.1385 | Benign | -3.57 | Benign | 0.23 | Neutral | 0.04 | Benign | 0.15 | Tolerated | 21.40 | Deleterious |
| p.Ser152Asp | VUS | -0.38 | Neutral | 0.1717 | Benign | -3.02 | Benign | 0.30 | Neutral | 0.19 | Benign | 0.43 | Tolerated | | |
| p.Ser152Glu | VUS | -1.44 | Neutral | 0.1777 | Benign | -2.63 | Benign | 0.29 | Neutral | 0.09 | Benign | 0.28 | Tolerated | | |
| p.Ser152Ala | VUS | -3.53 | Neutral | 0.0696 | Benign | -2.69 | Benign | 0.10 | Neutral | 0.00 | Benign | 1 | Tolerated | 0.00 | Neutral |

| | | | | | | | | | | | | | | | |
|---|---|---|---|---|---|---|---|---|---|---|---|---|---|---|---|
| p.Ser152Gly | VUS | -4.94 | Neutral | 0.0752 | Benign | -2.64 | Benign | 0.15 | Neutral | 0.05 | Benign | 0.79 | Tolerated | | |
| p.Ser152Val | VUS | -0.74 | Neutral | 0.1548 | Benign | -3.64 | Benign | 0.25 | Neutral | 0.05 | Benign | 0.19 | Tolerated | | |
| p.Ser152Tyr | VUS | -6.00 | Indeterminate | 0.1686 | Benign | -5.83 | Benign | 0.35 | Neutral | 0.83 | Possibly damaging | 0.09 | Tolerated | | |
| p.Ser152Cys | VUS | -0.14 | Neutral | 0.1286 | Benign | -5.27 | Benign | 0.32 | Neutral | 0.62 | Possibly damaging | 0.11 | Tolerated | | |
| p.Ser152Trp | VUS | -0.77 | Neutral | 0.1637 | Benign | -5.34 | Benign | 0.40 | Neutral | 0.95 | Possibly damaging | 0.03 | Deleterious | | |
| p.Ser152Phe | VUS | -0.82 | Neutral | 0.2066 | Benign | -5.34 | Benign | 0.35 | Neutral | 0.62 | Possibly damaging | 0.07 | Tolerated | | |
| p.Asp153Asn | VUS | -3.03 | Neutral | 0.1258 | Benign | -4.57 | Benign | 0.22 | Neutral | 0.08 | Benign | 1 | Tolerated | 25.20 | Deleterious |
| p.Asp153Lys | VUS | -1.73 | Neutral | 0.2406 | Benign | -4.74 | Benign | 0.29 | Neutral | 0.05 | Benign | 0.13 | Tolerated | | |
| p.Asp153Thr | VUS | -5.10 | Neutral | 0.1456 | Benign | -3.57 | Benign | 0.29 | Neutral | 0.10 | Benign | 0.1 | Tolerated | | |
| p.Asp153Arg | VUS | -4.54 | Neutral | 0.1458 | Benign | -3.09 | Benign | 0.28 | Neutral | 0.29 | Benign | 0.06 | Tolerated | | |
| p.Asp153Ser | VUS | 0.00 | Neutral | 0.0995 | Benign | -2.91 | Benign | 0.28 | Neutral | 0.05 | Benign | 0.2 | Tolerated | | |
| p.Asp153Ile | VUS | -0.70 | Neutral | 0.2505 | Benign | -4.40 | Benign | 0.30 | Neutral | 0.29 | Benign | 0.01 | Deleterious | | |
| p.Asp153Met | VUS | -1.49 | Neutral | 0.4459 | Ambiguous | -5.20 | Benign | 0.31 | Neutral | 0.90 | Possibly damaging | 0.01 | Deleterious | | |
| p.Asp153His | VUS | -2.40 | Neutral | 0.1339 | Benign | -4.29 | Benign | 0.19 | Neutral | 0.88 | Possibly damaging | 0.07 | Tolerated | 24.40 | Deleterious |
| p.Asp153Gln | VUS | -2.53 | Neutral | 0.1533 | Benign | -4.09 | Benign | 0.27 | Neutral | 0.29 | Benign | 0.09 | Tolerated | | |
| p.Asp153Pro | VUS | 0.00 | Neutral | 0.2242 | Benign | -2.92 | Benign | 0.31 | Neutral | 0.45 | Benign | 0.06 | Tolerated | | |
| p.Asp153Leu | VUS | -1.93 | Neutral | 0.217 | Benign | -3.61 | Benign | 0.32 | Neutral | 0.17 | Benign | 0.03 | Deleterious | | |
| p.Asp153Glu | VUS | -0.72 | Neutral | 0.11 | Benign | -3.39 | Benign | 0.07 | Neutral | 0.00 | Benign | 0.17 | Tolerated | 0.30 | Neutral |
| p.Asp153Ala | VUS | -0.06 | Neutral | 0.0941 | Benign | -2.90 | Benign | 0.19 | Neutral | 0.02 | Benign | 0.1 | Tolerated | 3.18 | Neutral |
| p.Asp153Gly | VUS | 0.00 | Neutral | 0.0907 | Benign | -2.74 | Benign | 0.13 | Neutral | 0.00 | Benign | 0.24 | Tolerated | 1.87 | Neutral |
| p.Asp153Val | VUS | -3.27 | Neutral | 0.1424 | Benign | -3.92 | Benign | 0.27 | Neutral | 0.13 | Benign | 0.02 | Deleterious | 6.46 | Neutral |
| p.Asp153Tyr | VUS | -3.98 | Neutral | 0.1422 | Benign | -4.27 | Benign | 0.28 | Neutral | 0.65 | Possibly damaging | 0.04 | Deleterious | 49.00 | Deleterious |
| p.Asp153Cys | VUS | -2.76 | Neutral | 0.3314 | Benign | -4.38 | Benign | 0.31 | Neutral | 0.00 | Benign | 0.02 | Deleterious | | |
| p.Asp153Trp | VUS | -2.73 | Neutral | 0.4708 | Ambiguous | -5.07 | Benign | 0.36 | Neutral | 0.90 | Possibly damaging | 0.01 | Deleterious | | |
| p.Asp153Phe | VUS | -0.54 | Neutral | 0.3314 | Benign | -4.49 | Benign | 0.31 | Neutral | 0.45 | Benign | 0.02 | Deleterious | | |
| p.Ile154Asn | VUS | -4.12 | Neutral | 0.1052 | Benign | -3.50 | Benign | 0.10 | Neutral | 0.00 | Benign | 0.1 | Tolerated | 4.16 | Neutral |
| p.Ile154Lys | VUS | -0.82 | Neutral | 0.243 | Benign | -3.21 | Benign | 0.19 | Neutral | 0.00 | Benign | 0.13 | Tolerated | | |
| p.Ile154Thr | VUS | -4.01 | Neutral | 0.20 | Benign | -2.18 | Benign | 0.05 | Neutral | 0.00 | Benign | 0.79 | Tolerated | 6.37 | Neutral |
| p.Ile154Arg | VUS | -1.80 | Neutral | 0.1514 | Benign | -1.74 | Benign | 0.18 | Neutral | 0.00 | Benign | 0.09 | Tolerated | | |
| p.Ile154Ser | VUS | -1.22 | Neutral | 0.12 | Benign | -1.48 | Benign | 0.08 | Neutral | 0.00 | Benign | 0.29 | Tolerated | 7.22 | Neutral |
| p.Ile154Met | VUS | -2.85 | Neutral | 0.1331 | Benign | -3.99 | Benign | 0.08 | Neutral | 0.03 | Benign | 0.12 | Tolerated | 6.97 | Neutral |
| p.Ile154His | VUS | -7.49 | Indeterminate | 0.1902 | Benign | -3.01 | Benign | 0.21 | Neutral | 0.20 | Benign | 0.05 | Deleterious | | |
| p.Ile154Gln | VUS | -2.71 | Neutral | 0.1991 | Benign | -2.54 | Benign | 0.23 | Neutral | 0.01 | Benign | 0.09 | Tolerated | | |
| p.Ile154Pro | VUS | -11.15 | Indeterminate | 0.2623 | Benign | -1.45 | Benign | 0.23 | Neutral | 0.03 | Benign | 0.11 | Tolerated | | |
| p.Ile154Leu | VUS | -1.30 | Neutral | 0.0907 | Benign | -2.13 | Benign | 0.13 | Neutral | 0.00 | Benign | 0.48 | Tolerated | 0.60 | Neutral |
| p.Ile154Asp | VUS | -1.80 | Neutral | 0.1529 | Benign | -1.86 | Benign | 0.17 | Neutral | 0.00 | Benign | 0.08 | Tolerated | | |
| p.Ile154Glu | VUS | -2.16 | Neutral | 0.1869 | Benign | -1.53 | Benign | 0.26 | Neutral | 0.00 | Benign | 0.11 | Tolerated | | |
| p.Ile154Ala | VUS | -11.26 | Indeterminate | 0.1848 | Benign | -1.19 | Benign | 0.18 | Neutral | 0.00 | Benign | 0.37 | Tolerated | | |
| p.Ile154Gly | VUS | -0.86 | Neutral | 0.2514 | Benign | -0.95 | Benign | 0.14 | Neutral | 0.00 | Benign | 0.07 | Tolerated | | |
| p.Ile154Val | VUS | -5.37 | Neutral | 0.12 | Benign | -2.12 | Benign | 0.05 | Neutral | 0.00 | Benign | 0.94 | Tolerated | 0.70 | Neutral |
| p.Ile154Tyr | VUS | -6.33 | Indeterminate | 0.2394 | Benign | -4.89 | Benign | 0.18 | Neutral | 0.01 | Benign | 0.08 | Tolerated | | |
| p.Ile154Cys | VUS | -7.96 | Indeterminate | 0.457 | Ambiguous | -3.47 | Benign | 0.18 | Neutral | 0.20 | Benign | 0.07 | Tolerated | | |
| p.Ile154Trp | VUS | -5.04 | Neutral | 0.4093 | Ambiguous | -4.07 | Benign | 0.20 | Neutral | 0.43 | Benign | 0.02 | Deleterious | | |
| p.Ile154Phe | VUS | -2.57 | Neutral | 0.102 | Benign | -3.93 | Benign | 0.09 | Neutral | 0.00 | Benign | 0.11 | Tolerated | 0.84 | Neutral |
| p.Pro155Asn | VUS | -1.38 | Neutral | 0.245 | Benign | -4.51 | Benign | 0.24 | Neutral | 0.77 | Possibly damaging | 0 | Deleterious | | |
| p.Pro155Lys | VUS | -4.81 | Neutral | 0.2015 | Benign | -3.82 | Benign | 0.21 | Neutral | 0.49 | Possibly damaging | 0 | Deleterious | | |
| p.Pro155Thr | VUS | 0.00 | Neutral | 0.075 | Benign | -3.48 | Benign | 0.14 | Neutral | 0.27 | Benign | 0 | Deleterious | 5.66 | Neutral |
| p.Pro155Arg | VUS | -0.34 | Neutral | 0.0812 | Benign | -3.02 | Benign | 0.16 | Neutral | 0.72 | Possibly damaging | 0 | Deleterious | 5.27 | Neutral |
| p.Pro155Ser | VUS | -28.22 | Indeterminate | 0.0838 | Benign | -2.51 | Benign | 0.08 | Neutral | 0.01 | Benign | 0 | Deleterious | 6.39 | Neutral |
| p.Pro155Ile | VUS | -7.35 | Indeterminate | 0.2085 | Benign | -3.53 | Benign | 0.26 | Neutral | 0.33 | Benign | 0 | Deleterious | | |
| p.Pro155Met | VUS | 0.00 | Neutral | 0.2982 | Benign | -4.40 | Benign | 0.23 | Neutral | 0.77 | Possibly damaging | 0 | Deleterious | | |
| p.Pro155His | VUS | -10.71 | Indeterminate | 0.123 | Benign | -4.21 | Benign | 0.19 | Neutral | 0.98 | Probably damaging | 0 | Deleterious | 5.43 | Neutral |
| p.Pro155Gln | VUS | 0.00 | Neutral | 0.0963 | Benign | -3.33 | Benign | 0.15 | Neutral | 0.87 | Possibly damaging | 0 | Deleterious | | |
| p.Pro155Leu | VUS | 0.00 | Neutral | 0.125 | Benign | -3.06 | Benign | 0.18 | Neutral | 0.00 | Benign | 0 | Deleterious | 6.17 | Neutral |
| p.Pro155Asp | VUS | -32.02 | Indeterminate | 0.1623 | Benign | -2.61 | Benign | 0.26 | Neutral | 0.49 | Possibly damaging | 0 | Deleterious | | |
| p.Pro155Glu | VUS | 0.00 | Neutral | 0.1133 | Benign | -2.35 | Benign | 0.24 | Neutral | 0.49 | Possibly damaging | 0 | Deleterious | | |
| p.Pro155Ala | VUS | 0.00 | Neutral | 0.0586 | Benign | -2.99 | Benign | 0.14 | Neutral | 0.00 | Benign | 0 | Deleterious | 5.51 | Neutral |
| p.Pro155Gly | VUS | -0.04 | Neutral | 0.1214 | Benign | -2.95 | Benign | 0.18 | Neutral | 0.33 | Benign | 0 | Deleterious | | |
| p.Pro155Val | VUS | -21.58 | Indeterminate | 0.138 | Benign | -3.09 | Benign | 0.22 | Neutral | 0.33 | Benign | 0 | Deleterious | | |
| p.Pro155Tyr | VUS | -0.02 | Neutral | 0.2893 | Benign | -5.04 | Benign | 0.25 | Neutral | 0.95 | Possibly damaging | 0 | Deleterious | | |
| p.Pro155Cys | VUS | 0.00 | Neutral | 0.3012 | Benign | -4.75 | Benign | 0.21 | Neutral | 0.96 | Probably damaging | 0 | Deleterious | | |
| p.Pro155Trp | VUS | -13.71 | Indeterminate | 0.3032 | Benign | -4.95 | Benign | 0.27 | Neutral | 0.99 | Probably damaging | 0 | Deleterious | | |
| p.Pro155Phe | VUS | 0.00 | Neutral | 0.3021 | Benign | -4.46 | Benign | 0.25 | Neutral | 0.77 | Possibly damaging | 0 | Deleterious | | |
| p.Asp156Asn | VUS | -3.19 | Neutral | 0.13 | Benign | -4.37 | Benign | 0.05 | Neutral | 0.00 | Benign | 0 | Deleterious | 10.02 | Neutral |
| p.Asp156Lys | VUS | -7.54 | Indeterminate | 0.2314 | Benign | -4.42 | Benign | 0.19 | Neutral | 0.21 | Benign | 0 | Deleterious | | |
| p.Asp156Thr | VUS | -1.63 | Neutral | 0.1663 | Benign | -3.45 | Benign | 0.20 | Neutral | 0.34 | Benign | 0 | Deleterious | | |
| p.Asp156Arg | VUS | -4.36 | Neutral | 0.1677 | Benign | -2.84 | Benign | 0.18 | Neutral | 0.65 | Possibly damaging | 0 | Deleterious | | |
| p.Asp156Ser | VUS | -0.77 | Neutral | 0.0915 | Benign | -2.61 | Benign | 0.19 | Neutral | 0.10 | Benign | 0 | Deleterious | | |
| p.Asp156Ile | VUS | -2.46 | Neutral | 0.3162 | Benign | -4.16 | Benign | 0.23 | Neutral | 0.65 | Possibly damaging | 0 | Deleterious | | |
| p.Asp156Met | VUS | -1.90 | Neutral | 0.5632 | Ambiguous | -4.77 | Benign | 0.24 | Neutral | 0.98 | Probably damaging | 0 | Deleterious | | |
| p.Asp156His | VUS | -4.44 | Neutral | 0.1454 | Benign | -4.09 | Benign | 0.10 | Neutral | 0.88 | Possibly damaging | 0 | Deleterious | 9.67 | Neutral |
| p.Asp156Gln | VUS | -6.15 | Indeterminate | 0.1475 | Benign | -3.65 | Benign | 0.17 | Neutral | 0.48 | Possibly damaging | 0 | Deleterious | | |
| p.Asp156Pro | VUS | -0.96 | Neutral | 0.2106 | Benign | -2.74 | Benign | 0.22 | Neutral | 0.79 | Possibly damaging | 0 | Deleterious | | |
| p.Asp156Leu | VUS | -3.31 | Neutral | 0.2758 | Benign | -2.83 | Benign | 0.25 | Neutral | 0.21 | Benign | 0 | Deleterious | | |
| p.Asp156Glu | VUS | -1.87 | Neutral | 0.11 | Benign | -3.22 | Benign | 0.06 | Neutral | 0.00 | Benign | 0 | Deleterious | 4.93 | Neutral |
| p.Asp156Ala | VUS | -9.22 | Indeterminate | 0.10 | Benign | -2.29 | Benign | 0.06 | Neutral | 0.00 | Benign | 0 | Deleterious | 7.78 | Neutral |
| p.Asp156Gly | VUS | -3.71 | Neutral | 0.12 | Benign | -2.67 | Benign | 0.07 | Neutral | 0.08 | Benign | 0 | Deleterious | 8.04 | Neutral |
| p.Asp156Val | VUS | -3.79 | Neutral | 0.188 | Benign | -3.23 | Benign | 0.17 | Neutral | 0.16 | Benign | 0 | Deleterious | 8.63 | Neutral |
| p.Asp156Tyr | VUS | -6.31 | Indeterminate | 0.1729 | Benign | -4.03 | Benign | 0.18 | Neutral | 0.96 | Possibly damaging | 0 | Deleterious | 11.12 | Neutral |
| p.Asp156Cys | VUS | -6.69 | Indeterminate | 0.3819 | Ambiguous | -3.93 | Benign | 0.23 | Neutral | 0.93 | Possibly damaging | 0 | Deleterious | | |
| p.Asp156Trp | VUS | -1.51 | Neutral | 0.6033 | Pathogenic | -4.70 | Benign | 0.26 | Neutral | 0.98 | Probably damaging | 0 | Deleterious | | |
| p.Asp156Phe | VUS | -1.31 | Neutral | 0.4633 | Ambiguous | -3.52 | Benign | 0.22 | Neutral | 0.79 | Possibly damaging | 0 | Deleterious | | |
