## Appendix 1-table 8 for "Functional characterization of all *CDKN2A* missense variants and comparison to in silico models of pathogenicity"

**Appendix 1-table 8. Assessment of in silico variant effect prediction models.**

| In silico model | Accuracy (%) | Sensitivity | Specificity | Positive predictive value | Negative predictive value |
|---|---|---|---|---|---|
| CADD | 45.1 | 0.97 | 0.35 | 0.23 | 0.98 |
| Polyphen-2 | 39.5 | 0.98 | 0.27 | 0.22 | 0.98 |
| SIFT | 60.9 | 0.79 | 0.57 | 0.28 | 0.93 |
| VEST | 71.9 | 0.91 | 0.68 | 0.38 | 0.97 |
| AlphaMissense | 71.6 | 0.94 | 0.67 | 0.38 | 0.98 |
| ESM1b | 59.2 | 0.95 | 0.51 | 0.30 | 0.98 |
| PrimateAI-3D | 85.4 | 0.25 | 0.98 | 0.68 | 0.87 |
