## Appendix 1-table 9 for "Functional characterization of all *CDKN2A* missense variants and comparison to in silico models of pathogenicity"

**Appendix 1-table 9. Missense somatic mutations in *CDKN2A* reported in COSMIC, TCGA, JHU, MSK-IMPACT.**

| CDKN2A amino acid residue | Variant | Number of patietns with mutation reported | | | | Characteritic |
|---|---|---|---|---|---|---|
| | | COSMIC | TCGA | JHU | MSK-IMPACT | |
| 2 | p.Glu2Lys | 3 | - | - | 1 | Neutral |
| 4 | p.Ala4Ser | 1 | - | - | - | Deleterious |
| 5 | p.Ala5Thr | 2 | - | - | - | Neutral |
| 6 | p.Gly6Glu | 2 | - | - | - | Neutral |
| 7 | p.Ser7Arg | 1 | - | - | - | Neutral |
| 8 | p.Ser8Ile | 1 | 1 | - | - | Neutral |
| 8 | p.Ser8Arg | 1 | 1 | - | - | Neutral |
| 9 | p.Met9Ile | 2 | - | - | - | Neutral |
| 10 | p.Glu10Val | 1 | - | - | 1 | Neutral |
| 12 | p.Ser12Leu | 5 | 1 | - | 3 | Indeterminate |
| 13 | p.Ala13Thr | 1 | 1 | - | - | Neutral |
| 14 | p.Asp14Glu | 3 | - | - | 1 | Neutral |
| 14 | p.Asp14Gly | 2 | - | - | - | Neutral |
| 14 | p.Asp14Asn | 2 | - | - | 1 | Neutral |
| 14 | p.Asp14Val | 1 | - | - | - | Indeterminate |
| 16 | p.Leu16Pro | 4 | - | - | - | Deleterious |
| 16 | p.Leu16Gln | 1 | 1 | - | - | Deleterious |
| 16 | p.Leu16Arg | 1 | 1 | - | - | Deleterious |
| 17 | p.Ala17Gly | 1 | - | - | - | Neutral |
| 18 | p.Thr18Ala | 1 | - | - | - | Indeterminate |
| 18 | p.Thr18Met | 1 | - | - | - | Indeterminate |
| 19 | p.Ala19Pro | 1 | - | - | - | Deleterious |
| 20 | p.Ala20Glu | 3 | 1 | - | - | Deleterious |
| 20 | p.Ala20Pro | 2 | 1 | - | - | Deleterious |
| 20 | p.Ala20Arg | 1 | - | - | - | Deleterious |
| 20 | p.Ala20Ser | 2 | - | - | - | Indeterminate |
| 20 | p.Ala20Thr | 4 | - | - | - | Indeterminate |
| 21 | p.Ala21Asp | 3 | - | - | 1 | Deleterious |
| 21 | p.Ala21Pro | 1 | - | - | - | Deleterious |
| 22 | p.Arg22Pro | 4 | 1 | - | - | Deleterious |
| 23 | p.Gly23Cys | 1 | 1 | - | - | Deleterious |
| 23 | p.Gly23Asp | 1 | 1 | - | - | Deleterious |
| 23 | p.Gly23Arg | 2 | 1 | 1 | - | Deleterious |
| 23 | p.Gly23Ser | 3 | - | - | - | Deleterious |
| 23 | p.Gly23Val | 5 | 1 | - | - | Deleterious |
| 24 | p.Arg24Gly | 1 | 1 | - | - | Neutral |
| 24 | p.Arg24Pro | 8 | - | - | 1 | Deleterious |
| 24 | p.Arg24Gln | 1 | - | - | - | Indeterminate |
| 25 | p.Val25Gly | 2 | - | - | - | Indeterminate |
| 26 | p.Glu26Asp | 1 | - | - | - | Neutral |
| 26 | p.Glu26Lys | 1 | - | - | - | Neutral |
| 28 | p.Val28Gly | 2 | - | - | - | Indeterminate |
| 29 | p.Arg29Pro | 3 | - | - | - | Deleterious |
| 29 | p.Arg29Trp | 1 | - | - | - | Neutral |
| 30 | p.Ala30Pro | 1 | - | - | - | Deleterious |
| 30 | p.Ala30Val | 4 | - | - | - | Indeterminate |
| 31 | p.Leu31Pro | 1 | - | - | - | Deleterious |
| 31 | p.Leu31Arg | 1 | 1 | - | - | Neutral |
| 31 | p.Leu31Val | 1 | 1 | - | - | Indeterminate |
| 32 | p.Leu32Pro | 3 | 1 | - | 2 | Deleterious |
| 32 | p.Leu32Gln | 2 | - | - | 2 | Indeterminate |
| 33 | p.Glu33Asp | 1 | - | - | - | Neutral |
| 34 | p.Ala34Asp | 1 | - | - | - | Neutral |
| 35 | p.Gly35Ala | 1 | - | - | - | Indeterminate |
| 35 | p.Gly35Glu | 1 | - | - | - | Indeterminate |
| 35 | p.Gly35Arg | 2 | - | - | - | Neutral |
| 35 | p.Gly35Val | 3 | - | - | - | Deleterious |
| 35 | p.Gly35Trp | 2 | - | - | 1 | Deleterious |
| 36 | p.Ala36Gly | 3 | - | - | - | Indeterminate |

| | | | | | | |
|---|---|---|---|---|---|---|
| 36 | p.Ala36Ser | - | 1 | - | - | Neutral |
| 36 | p.Ala36Thr | 1 | - | - | - | Indeterminate |
| 38 | p.Pro38His | 1 | - | - | 1 | Deleterious |
| 38 | p.Pro38Leu | 2 | - | - | - | Indeterminate |
| 38 | p.Pro38Ser | 1 | - | - | - | Indeterminate |
| 38 | p.Pro38Thr | - | - | 1 | - | Indeterminate |
| 39 | p.Asn39Ile | 1 | - | - | 1 | Indeterminate |
| 39 | p.Asn39Lys | 1 | - | - | - | Indeterminate |
| 40 | p.Ala40Pro | 1 | - | - | - | Indeterminate |
| 42 | p.Asn42Asp | 1 | - | - | - | Indeterminate |
| 42 | p.Asn42His | 4 | 1 | - | 1 | Deleterious |
| 42 | p.Asn42Ile | 2 | 1 | - | - | Deleterious |
| 42 | p.Asn42Tyr | 2 | - | - | 1 | Deleterious |
| 43 | p.Ser43Ile | - | 1 | - | - | Indeterminate |
| 43 | p.Ser43Arg | 1 | - | - | - | Neutral |
| 44 | p.Tyr44Cys | 1 | 1 | - | - | Neutral |
| 45 | p.Gly45Asp | 3 | - | - | - | Neutral |
| 45 | p.Gly45Ser | 1 | - | - | - | Indeterminate |
| 47 | p.Arg47Met | 1 | - | - | 1 | Neutral |
| 48 | p.Pro48Leu | 28 | 6 | - | 4 | Deleterious |
| 48 | p.Pro48Gln | 1 | 2 | - | - | Deleterious |
| 48 | p.Pro48Arg | 3 | 1 | - | - | Deleterious |
| 48 | p.Pro48Ser | 2 | 1 | - | - | Neutral |
| 49 | p.Ile49Asn | 2 | - | - | 1 | Deleterious |
| 49 | p.Ile49Ser | 2 | - | - | - | Deleterious |
| 49 | p.Ile49Thr | 3 | - | - | - | Indeterminate |
| 50 | p.Gln50His | 3 | 1 | - | 1 | Deleterious |
| 50 | p.Gln50Leu | 2 | - | - | - | Indeterminate |
| 50 | p.Gln50Arg | 3 | 1 | - | 2 | Deleterious |
| 51 | p.Val51Ala | 3 | - | - | - | Neutral |
| 51 | p.Val51Asp | 4 | - | - | - | Deleterious |
| 51 | p.Val51Phe | 2 | - | 1 | - | Deleterious |
| 51 | p.Val51Ile | 3 | - | - | - | Neutral |
| 52 | p.Met52Ile | 1 | - | 1 | - | Indeterminate |
| 52 | p.Met52Lys | 3 | - | - | - | Deleterious |
| 52 | p.Met52Leu | 1 | - | - | - | Neutral |
| 52 | p.Met52Arg | 3 | 1 | - | - | Deleterious |
| 53 | p.Met53Ile | 9 | 1 | 3 | 1 | Deleterious |
| 53 | p.Met53Val | - | - | 2 | - | Deleterious |
| 54 | p.Met54Leu | - | - | 1 | - | Neutral |
| 55 | p.Gly55Cys | 1 | - | 1 | - | Deleterious |
| 55 | p.Gly55Asp | 2 | - | - | - | Deleterious |
| 55 | p.Gly55Arg | 3 | - | - | - | Deleterious |
| 55 | p.Gly55Val | 3 | - | 1 | 1 | Deleterious |
| 56 | p.Ser56Asn | 1 | - | 1 | - | Neutral |
| 56 | p.Ser56Arg | 1 | 1 | - | - | Neutral |
| 57 | p.Ala57Phe | 1 | - | - | - | Indeterminate |
| 57 | p.Ala57Pro | 1 | - | - | - | Neutral |
| 57 | p.Ala57Ser | 2 | - | - | - | Neutral |
| 57 | p.Ala57Thr | 2 | - | - | - | Neutral |
| 57 | p.Ala57Val | 6 | - | 2 | - | Neutral |
| 58 | p.Arg58Gln | 2 | - | - | - | Neutral |
| 58 | p.Val58Leu | - | - | 1 | - | Neutral |
| 59 | p.Val59Met | 1 | - | - | - | Neutral |
| 60 | p.Ala60Glu | 1 | - | - | - | Indeterminate |
| 60 | p.Ala60Ser | 1 | - | - | - | Indeterminate |
| 60 | p.Ala60Thr | - | - | 1 | - | Neutral |
| 60 | p.Ala60Val | 5 | 1 | - | 1 | Neutral |
| 61 | p.Glu61Lys | 1 | 1 | - | - | Neutral |
| 61 | p.Glu61Val | - | - | 1 | - | Neutral |
| 62 | p.Leu62Met | - | 1 | 1 | - | Neutral |
| 62 | p.Leu62Pro | 3 | - | - | 2 | Indeterminate |
| 63 | p.Leu63Gln | 5 | - | 1 | 3 | Deleterious |

| | | | | | | |
|---|---|---|---|---|---|---|
| 63 | p.Leu63Arg | 1 | - | 1 | - | Deleterious |
| 63 | p.Leu63Val | 1 | - | - | - | Deleterious |
| 64 | p.Leu64Pro | 2 | 1 | 2 | - | Deleterious |
| 65 | p.Leu65Pro | 1 | - | - | - | Deleterious |
| 65 | p.Leu65Val | - | - | 1 | - | Neutral |
| 66 | p.His66Leu | 1 | - | - | - | Neutral |
| 66 | p.His66Arg | 5 | - | 3 | - | Neutral |
| 66 | p.His66Tyr | 1 | - | 1 | - | Neutral |
| 67 | p.Gly67Cys | 2 | - | - | 1 | Neutral |
| 67 | p.Gly67Asp | 2 | - | - | - | Neutral |
| 67 | p.Gly67Ser | 5 | - | 1 | - | Neutral |
| 67 | p.Gly67Val | - | 1 | - | - | Deleterious |
| 68 | p.Ala68Glu | 2 | - | - | 1 | Indeterminate |
| 68 | p.Ala68Gly | 1 | - | - | - | Neutral |
| 68 | p.Ala68Pro | 2 | 1 | 1 | - | Deleterious |
| 68 | p.Ala68Thr | 8 | - | - | 1 | Neutral |
| 68 | p.Ala68Val | 2 | - | 3 | - | Indeterminate |
| 69 | p.Glu69Asp | 3 | - | - | - | Neutral |
| 69 | p.Glu69Gly | - | - | 4 | - | Indeterminate |
| 69 | p.Glu69Lys | 1 | - | - | - | Indeterminate |
| 69 | p.Glu69Val | 1 | - | - | - | Indeterminate |
| 70 | p.Pro70Leu | 4 | - | 2 | 2 | Neutral |
| 70 | p.Pro70Arg | 1 | - | - | - | Neutral |
| 70 | p.Pro70Ser | 1 | - | - | - | Indeterminate |
| 71 | p.Asn71Asp | 1 | - | - | - | Indeterminate |
| 71 | p.Asn71His | - | - | 1 | - | Indeterminate |
| 71 | p.Asn71Ile | 2 | - | - | - | Deleterious |
| 71 | p.Asn71Lys | - | 1 | - | - | Deleterious |
| 71 | p.Asn71Ser | - | - | 1 | - | Deleterious |
| 71 | p.Asn71Tyr | 1 | - | - | - | Deleterious |
| 72 | p.Cys72Gly | 1 | - | - | - | Neutral |
| 72 | p.Cys72ser | - | - | 1 | - | Indeterminate |
| 72 | p.Cys72Tyr | - | - | 1 | - | Neutral |
| 73 | p.Ala73Asp | 2 | 1 | - | - | Neutral |
| 73 | p.Ala73Ser | 1 | - | - | - | Neutral |
| 74 | p.Asp74Ala | 5 | - | - | - | Deleterious |
| 74 | p.Asp74Gly | 1 | - | 3 | - | Deleterious |
| 74 | p.Asp74Asn | 10 | 1 | 2 | 2 | Deleterious |
| 74 | p.Asp74Val | 3 | 1 | 1 | - | Deleterious |
| 74 | p.Asp74Tyr | 9 | 1 | 2 | 2 | Deleterious |
| 75 | p.Pro75Ser | 1 | - | 1 | - | Neutral |
| 75 | p.Pro75Thr | 1 | - | - | - | Neutral |
| 76 | p.Ala76Gly | 1 | - | - | 1 | Neutral |
| 76 | p.Ala76Ser | 1 | - | - | - | Indeterminate |
| 76 | p.Ala76Thr | 7 | 1 | - | - | Neutral |
| 76 | p.Ala76Val | 6 | - | - | - | Neutral |
| 77 | p.Thr77Ile | 1 | 1 | 1 | - | Neutral |
| 77 | p.Thr77Ser | 1 | - | - | - | Indeterminate |
| 78 | p.Leu78Phe | - | - | 2 | - | Neutral |
| 78 | p.Leu78His | 1 | - | - | - | Neutral |
| 79 | p.Thr79Ala | 1 | - | - | - | Neutral |
| 79 | p.Thr79Ile | 5 | - | - | - | Neutral |
| 79 | p.Thr79Asn | - | - | 1 | - | Neutral |
| 79 | p.Thr79Pro | 1 | - | - | 1 | Deleterious |
| 80 | p.Arg80Leu | - | - | 1 | - | Indeterminate |
| 80 | p.Arg80Pro | - | - | 1 | - | Deleterious |
| 80 | p.Arg80Gln | 4 | 1 | - | - | Indeterminate |
| 81 | p.Pro81Ala | 1 | - | - | - | Indeterminate |
| 81 | p.Pro81His | 3 | 1 | 1 | - | Deleterious |
| 81 | p.Pro81Leu | 28 | 5 | 6 | 3 | Deleterious |
| 81 | p.Pro81Arg | 3 | - | - | 1 | Deleterious |
| 81 | p.Pro81Ser | 2 | - | 1 | - | Deleterious |
| 82 | p.Val82Glu | 2 | 1 | - | - | Indeterminate |

| | | | | | | |
|---|---|---|---|---|---|---|
| 82 | p.Val82Gly | 1 | - | 1 | - | Indeterminate |
| 82 | p.Val82Leu | 2 | - | - | 1 | Neutral |
| 82 | p.Val82Met | 8 | 2 | 1 | - | Neutral |
| 83 | p.His83Asp | 8 | 3 | 2 | 2 | Deleterious |
| 83 | p.His83Leu | 2 | 1 | 2 | - | Deleterious |
| 83 | p.His83Asn | 4 | - | 1 | 1 | Deleterious |
| 83 | p.His83Pro | 4 | - | - | - | Deleterious |
| 83 | p.His83Gln | 2 | 1 | - | 1 | Deleterious |
| 83 | p.His83Arg | 13 | 2 | 2 | 1 | Deleterious |
| 83 | p.His83Tyr | 139 | 18 | 17 | 32 | Deleterious |
| 84 | p.Asp84Ala | 2 | - | 1 | 1 | Deleterious |
| 84 | p.Asp84Gly | 15 | 3 | 2 | 2 | Deleterious |
| 84 | p.Asp84His | 3 | - | 1 | - | Deleterious |
| 84 | p.Asp84Asn | 40 | 10 | 3 | 4 | Deleterious |
| 84 | p.Asp84Val | 5 | 1 | - | 2 | Deleterious |
| 84 | p.Asp84Tyr | 24 | 4 | 8 | 5 | Deleterious |
| 85 | p.Ala85Pro | 5 | 1 | 1 | 2 | Indeterminate |
| 85 | p.Ala85Ser | 1 | - | 1 | - | Neutral |
| 85 | p.Ala85Thr | 3 | 2 | - | - | Indeterminate |
| 86 | p.Ala86Asp | 4 | 1 | - | - | Deleterious |
| 86 | p.Ala86Pro | - | 1 | - | - | Deleterious |
| 87 | p.Arg87Leu | 1 | - | 1 | - | Indeterminate |
| 87 | p.Arg87Pro | 3 | - | 1 | 2 | Deleterious |
| 87 | p.Arg87Trp | 3 | - | 2 | - | Deleterious |
| 88 | p.Glu88Ala | 1 | - | - | - | Neutral |
| 88 | p.Glu88Asp | 2 | - | - | - | Indeterminate |
| 88 | p.Glu88Lys | 15 | 5 | 1 | 3 | Indeterminate |
| 88 | p.Glu88Val | 1 | - | - | - | Neutral |
| 89 | p.Gly89Cys | 2 | - | - | 2 | Deleterious |
| 89 | p.Gly89Asp | - | - | 1 | - | Deleterious |
| 89 | p.Gly89Phe | - | - | 1 | - | Deleterious |
| 89 | p.Gly89Ser | 4 | - | 1 | 1 | Deleterious |
| 89 | p.Gly89Val | 2 | 1 | - | - | Deleterious |
| 90 | p.Phe90Leu | 10 | 1 | - | 1 | Neutral |
| 91 | p.Leu91Gln | 1 | - | - | - | Neutral |
| 92 | p.Asp92Tyr | 1 | - | - | 1 | Neutral |
| 93 | p.Thr93Ala | 1 | - | - | - | Indeterminate |
| 93 | p.Thr93Lys | 6 | - | - | 1 | Deleterious |
| 93 | p.Thr93Met | 6 | - | - | 1 | Indeterminate |
| 93 | p.Thr93Arg | 1 | - | - | - | Indeterminate |
| 94 | p.Leu94Pro | 2 | 1 | - | - | Deleterious |
| 94 | p.Leu94Gln | - | 1 | - | - | Deleterious |
| 95 | p.Val95Ala | 1 | - | - | - | Neutral |
| 95 | p.Val95Leu | 2 | - | - | - | Neutral |
| 95 | p.Val95Met | 1 | - | - | - | Indeterminate |
| 97 | p.Leu97Pro | 3 | - | - | - | Deleterious |
| 97 | p.Leu97Arg | 4 | - | - | - | Deleterious |
| 98 | p.His98Pro | 4 | 2 | 1 | 1 | Deleterious |
| 98 | p.His98Tyr | 2 | - | - | 1 | Neutral |
| 99 | p.Arg99Gln | 4 | - | - | 1 | Neutral |
| 99 | p.Arg99Trp | 1 | - | 1 | - | Indeterminate |
| 100 | p.Ala100Pro | 1 | - | - | - | Deleterious |
| 100 | p.Ala100Ser | 3 | - | 2 | - | Neutral |
| 100 | p.Ala100Thr | 2 | - | - | - | Neutral |
| 100 | p.Ala100Val | 2 | - | - | - | Neutral |
| 101 | p.Gly101Val | 4 | 2 | 4 | - | Deleterious |
| 101 | p.Gly101Trp | 8 | 1 | 4 | 1 | Deleterious |
| 102 | p.Ala102Glu | 7 | 1 | 2 | - | Deleterious |
| 102 | p.Ala102Thr | 3 | - | 1 | - | Deleterious |
| 102 | p.Ala102Val | 10 | 1 | 4 | 2 | Indeterminate |
| 103 | p.Arg103Gln | 1 | - | 1 | - | Neutral |
| 103 | p.Arg103Trp | 2 | 1 | - | 1 | Neutral |
| 104 | p.Leu104Gln | 1 | 1 | - | - | Neutral |

| | | | | | | |
|---|---|---|---|---|---|---|
| 104 | p.Leu104Arg | 2 | - | 1 | 1 | Deleterious |
| 104 | p.Leu104Val | 1 | - | - | - | Neutral |
| 105 | p.Asp105Asn | 1 | - | - | - | Neutral |
| 106 | p.Val106Met | 4 | - | - | - | Neutral |
| 107 | p.Arg107Cys | 2 | - | 2 | - | Indeterminate |
| 107 | p.Arg107His | 2 | 1 | - | - | Neutral |
| 108 | p.Asp108Ala | 1 | - | - | - | Deleterious |
| 108 | p.Asp108Gly | 6 | 1 | 1 | - | Deleterious |
| 108 | p.Asp108His | 15 | 1 | - | 3 | Deleterious |
| 108 | p.Asp108Asn | 20 | 5 | 1 | 5 | Neutral |
| 108 | p.Asp108Val | 3 | 1 | - | - | Deleterious |
| 108 | p.Asp108Tyr | 39 | 11 | 5 | 5 | Deleterious |
| 109 | p.Ala109Pro | - | - | 1 | - | Deleterious |
| 109 | p.Ala109Thr | 2 | - | - | - | Neutral |
| 109 | p.Ala109Val | 1 | - | - | - | Neutral |
| 110 | p.Trp110Cys | 1 | - | - | - | Neutral |
| 110 | p.Trp110Arg | 1 | - | - | - | Neutral |
| 111 | p.Gly111Asp | 2 | 1 | - | - | Neutral |
| 111 | p.Gly111Ser | 1 | - | - | - | Neutral |
| 111 | p.Gly111Val | 1 | - | 1 | - | Deleterious |
| 112 | p.Arg112Cys | 3 | - | 1 | - | Indeterminate |
| 112 | p.Arg112His | 2 | 2 | 1 | - | Neutral |
| 112 | p.Arg112Pro | 2 | - | - | 1 | Deleterious |
| 112 | p.Arg112ser | 2 | - | - | - | Neutral |
| 114 | p.Pro114Phe | 1 | - | 1 | 1 | Deleterious |
| 114 | p.Pro114His | 2 | - | - | - | Deleterious |
| 114 | p.Pro114Leu | 78 | 13 | 11 | 11 | Deleterious |
| 114 | p.Pro114Arg | 2 | - | - | - | Deleterious |
| 114 | p.Pro114Ser | 4 | - | - | 1 | Indeterminate |
| 114 | p.Pro114Thr | 4 | 3 | - | - | Deleterious |
| 115 | p.Val115Glu | 1 | - | - | - | Indeterminate |
| 115 | p.Val115Leu | 4 | - | - | - | Neutral |
| 116 | p.Asp116Asn | - | - | 1 | - | Neutral |
| 116 | p.Asp116Tyr | 3 | - | - | 1 | Neutral |
| 118 | p.Ala118Pro | 2 | 1 | - | - | Indeterminate |
| 118 | p.Ala118Thr | 1 | 1 | - | - | Neutral |
| 118 | p.Ala118Val | 1 | - | - | - | Indeterminate |
| 119 | p.Glu119Asp | 2 | 2 | - | - | Indeterminate |
| 119 | p.Glu119Lys | 2 | 1 | 2 | - | Neutral |
| 119 | p.Glu119Gln | 2 | 1 | - | - | Indeterminate |
| 120 | p.Glu120Ala | 1 | - | - | - | Neutral |
| 120 | p.Glu120Lys | 4 | - | - | 1 | Neutral |
| 122 | p.Gly122Cys | 1 | - | - | 1 | Neutral |
| 122 | p.Gly122Asp | 2 | - | - | - | Neutral |
| 122 | p.Gly122Ser | 2 | - | - | - | Neutral |
| 122 | p.Gly122Val | - | - | 1 | - | Neutral |
| 123 | p.His123Asn | 1 | 1 | - | 1 | Indeterminate |
| 123 | p.His123Gln | 2 | - | - | - | Neutral |
| 124 | p.Arg124His | 6 | - | - | 2 | Neutral |
| 124 | p.Arg124Cys | 1 | - | 1 | - | Neutral |
| 125 | p.Asp125His | - | - | 3 | - | Neutral |
| 125 | p.Asp125Asn | 3 | 1 | - | - | Neutral |
| 126 | p.Val126Ala | 1 | - | - | 1 | Neutral |
| 126 | p.Val126Asp | 3 | 1 | 2 | - | Deleterious |
| 126 | p.Val126Phe | 3 | - | - | - | Neutral |
| 126 | p.Val126Gly | - | 1 | - | - | Neutral |
| 126 | p.Val126Ile | 3 | - | - | - | Neutral |
| 127 | p.Ala127Pro | 1 | - | - | - | Deleterious |
| 127 | p.Ala127Ser | 2 | - | - | 1 | Neutral |
| 128 | p.Arg128Cys | - | - | - | 1 | Neutral |
| 128 | p.Arg128Leu | - | - | 1 | - | Neutral |
| 128 | p.Arg128Pro | - | - | 1 | - | Deleterious |
| 128 | p.Arg128Gln | 2 | 1 | - | - | Neutral |

| | | | | | | |
|---|---|---|---|---|---|---|
| 128 | p.Arg128Trp | 5 | - | - | - | Neutral |
| 129 | p.Tyr129Cys | 3 | - | - | 2 | Neutral |
| 129 | p.Tyr129Phe | 1 | - | - | - | Neutral |
| 129 | p.Tyr129His | 1 | - | - | - | Neutral |
| 129 | p.Tyr129Asn | 1 | 1 | - | - | Indeterminate |
| 130 | p.Leu130Met | - | - | 1 | - | Neutral |
| 130 | p.Leu130Pro | 4 | 1 | 1 | - | Deleterious |
| 130 | p.Leu130Gln | 8 | - | - | 2 | Indeterminate |
| 130 | p.Leu130Arg | 5 | - | 2 | - | Deleterious |
| 131 | p.Arg131Cys | 4 | 1 | - | 1 | Deleterious |
| 131 | p.Arg131His | 5 | 1 | - | - | Neutral |
| 131 | p.Arg131Leu | 1 | - | - | - | Neutral |
| 131 | p.Arg131Pro | 2 | - | - | 1 | Indeterminate |
| 132 | p.Ala132Pro | 2 | - | - | - | Neutral |
| 132 | p.Ala132Val | - | 1 | - | - | Neutral |
| 133 | p.Ala133ser | - | - | 1 | - | Neutral |
| 133 | p.Ala133Thr | - | 1 | - | - | Neutral |
| 134 | p.Ala134Val | 1 | - | - | - | Neutral |
| 135 | p.Gly135Ala | 1 | - | - | - | Neutral |
| 135 | p.Gly135Glu | 3 | - | - | 1 | Indeterminate |
| 136 | p.Gly136Asp | 2 | - | - | - | Indeterminate |
| 136 | p.Gly136Arg | - | - | 1 | - | Neutral |
| 137 | p.Thr137Ala | 1 | - | - | - | Neutral |
| 138 | p.Arg138Thr | 2 | - | - | 1 | Neutral |
| 139 | p.Gly139Ser | - | - | 2 | - | Neutral |
| 140 | p.Ser140Cys | 1 | - | - | - | Neutral |
| 140 | p.Ser140Asn | 1 | - | - | - | Neutral |
| 142 | p.His142Asn | 1 | - | - | - | Neutral |
| 142 | p.His142Gln | 1 | - | - | - | Neutral |
| 142 | p.His142Tyr | 1 | - | - | - | Neutral |
| 144 | p.Arg144His | 2 | - | - | 2 | Neutral |
| 146 | p.Asp146Ala | 1 | - | - | - | Neutral |
| 146 | p.Asp146Glu | 1 | - | - | - | Neutral |
| 146 | p.Asp146Gly | - | 1 | - | - | Neutral |
| 146 | p.Asp146Asn | 1 | - | - | - | Neutral |
| 148 | p.Ala148Gly | 1 | - | - | - | Neutral |
| 148 | p.Ala148Thr | 10 | - | - | - | Neutral |
| 148 | p.Ala148Val | 1 | 1 | - | - | Indeterminate |
| 149 | p.Glu149Lys | 1 | - | - | - | Neutral |
| 150 | p.Gly150Ser | 1 | 1 | - | - | Neutral |
| 150 | p.Gly150Val | 1 | - | - | - | Neutral |
| 153 | p.Asp153Asn | 9 | - | 1 | 4 | Neutral |
| 154 | p.Ile154Thr | 1 | - | - | - | Neutral |
| 155 | p.Pro155Ser | 1 | - | - | - | Indeterminate |

COSMIC - Catalogue Of Somatic Mutations In Cancer; TCGA - The Cancer Genome Atlas; JHU - Johns Hopkins University ; MSKCC-IMPACT - Memorial Sloan Kettering-Integrated Mutation Profiling of Actionable Cancer Targets Clinical Sequencing Cohort.
