## Appendix 1-table 10 for "Functional characterization of all *CDKN2A* missense variants and comparison to in silico models of pathogenicity"

**Appendix 1-table 10.** ***CDKN2A*** **missense and synonymous variants reproted in gnomAD.**

| Residue | Variant | Transcript Consequence | Missense or synonymous | Benchmark | Functionally reported VUS | P value | Fucntional characterization | Allele Count | Allele Number | Allele Frequency | ClinVar Clinical Significance | ClinVar Variation ID |
|---|---|---|---|---|---|---|---|---|---|---|---|---|
| 1 | p.Met1Arg | c.2T>G | Missense | | | 0.86 | Neutral | 2 | 1440078 | 1.38881E-06 | Uncertain significance | 660169 |
| 1 | p.Met1Lys | c.2T>A | Missense | | | 0.99 | Neutral | 11 | 1591956 | 6.90974E-06 | Uncertain significance | 487018 |
| 1 | p.Met1Val | c.1A>G | Missense | | | 0.06 | Neutral | 1 | 606364 | 1.64917E-06 | Uncertain significance | 820481 |
| 2 | p.Glu2Gln | c.4G>C | Missense | | | 1.00 | Neutral | 2 | 1592086 | 1.25621E-06 | Uncertain significance | 406722 |
| 2 | p.Glu2Glu | c.6G>A | Synonymous | | | 0.48 | Neutral | 3 | 1439810 | 2.08361E-06 | Likely benign | 483322 |
| 2 | p.Glu2Gly | c.5A>G | Missense | | | 1.00 | Neutral | 1 | 1439958 | 6.94E-07 | | |
| 3 | p.Pro3Ala | c.7C>G | Missense | | | 0.08 | Neutral | 2 | 1439510 | 1.38936E-06 | | |
| 3 | p.Pro3Leu | c.8C>T | Missense | | | 0.00 | Deleterious | 2 | 1446148 | 1.38298E-06 | Uncertain significance | 1448238 |
| 3 | p.Pro3Pro | c.9G>A | Synonymous | | | 0.48 | Neutral | 12 | 1598072 | 7.50905E-06 | Likely benign | 389468 |
| 3 | p.Pro3Ser | c.7C>T | Missense | | | 0.00 | Deleterious | 3 | 1591586 | 1.88491E-06 | Uncertain significance | 532281 |
| 3 | p.Pro3Thr | c.7C>A | Missense | | | 0.00 | Deleterious | 4 | 1439508 | 2.77873E-06 | | |
| 4 | p.Ala4Ala | c.12G>A | Synonymous | | | 0.48 | Neutral | 1 | 152000 | 6.57895E-06 | Likely benign | 1101964 |
| 4 | p.Ala4Ala | c.12G>T | Synonymous | | | 0.48 | Neutral | 1 | 1448522 | 6.90E-07 | | |
| 4 | p.Ala4Gly | c.11C>G | Missense | | | 0.01 | Deleterious | 5 | 1447740 | 3.45366E-06 | Uncertain significance | 628437 |
| 4 | p.Ala4Val | c.11C>T | Missense | | | 0.93 | Neutral | 1 | 1447738 | 6.91E-07 | Uncertain significance | 1058585 |
| 5 | p.Ala5Ala | c.15G>C | Synonymous | | | 0.48 | Neutral | 2 | 1449148 | 1.38012E-06 | | |
| 5 | p.Ala5Ala | c.15G>A | Synonymous | | | 0.48 | Neutral | 2 | 1449148 | 1.38012E-06 | Likely benign | 2129711 |
| 5 | p.Ala5Ala | c.15G>T | Synonymous | | | 0.48 | Neutral | 3 | 1449148 | 2.07018E-06 | Likely benign | 229890 |
| 5 | p.Ala5Glu | c.14C>A | Missense | | | 0.01 | Deleterious | 1 | 151816 | 6.58692E-06 | Uncertain significance | 489864 |
| 5 | p.Ala5Thr | c.13G>A | Missense | | | 0.59 | Neutral | 6 | 1448480 | 4.14227E-06 | Uncertain significance | 233536 |
| 6 | p.Gly6Glu | c.17G>A | Missense | | | 0.61 | Neutral | 2 | 1449782 | 1.37952E-06 | Uncertain significance | 483323 |
| 6 | p.Gly6Trp | c.16G>T | Missense | | | 0.11 | Neutral | 1 | 616342 | 1.62248E-06 | Uncertain significance | 1525531 |
| 6 | p.Gly6Val | c.17G>T | Missense | | | 0.97 | Neutral | 1 | 1449780 | 6.90E-07 | Uncertain significance | 220777 |
| 7 | p.Ser7Asn | c.20G>A | Missense | | | 0.95 | Neutral | 1 | 617530 | 1.61935E-06 | | |
| 7 | p.Ser7Cys | c.19A>T | Missense | | | 0.11 | Neutral | 2 | 1450466 | 1.37887E-06 | Uncertain significance | 489865 |
| 7 | p.Ser7Gly | c.19A>G | Missense | | | 0.99 | Neutral | 1 | 151766 | 6.58909E-06 | | |
| 8 | p.Ser8Arg | c.22A>C | Missense | | | 0.97 | Neutral | 1 | 1451436 | 6.89E-07 | Uncertain significance | 968555 |
| 8 | p.Ser8Gly | c.22A>G | Missense | | | 0.75 | Neutral | 1 | 1451436 | 6.89E-07 | Uncertain significance | 480814 |
| 9 | p.Met9Arg | c.26T>G | Missense | | | 0.95 | Neutral | 7 | 1604276 | 4.36334E-06 | Uncertain significance | 821690 |
| 9 | p.Met9Ile | c.27G>A | Missense | | | 0.82 | Neutral | 2 | 619538 | 3.22821E-06 | Uncertain significance | 656960 |
| 9 | p.Met9Leu | c.25A>T | Missense | | | 0.99 | Neutral | 9 | 1604328 | 5.60983E-06 | Uncertain significance | 483321 |
| 9 | p.Met9Lys | c.26T>A | Missense | | | 0.85 | Neutral | 48 | 1604394 | 2.99178E-05 | flicting interpretations of pathogen | 236985 |
| 9 | p.Met9Thr | c.26T>C | Missense | | Functionally neutral | 0.83 | Neutral | 2 | 1452556 | 1.37688E-06 | Uncertain significance | 483319 |
| 9 | p.Met9Val | c.25A>G | Missense | | | 0.73 | Neutral | 1 | 151942 | 6.58146E-06 | Uncertain significance | 483345 |
| 10 | p.Glu10Asp | c.30G>C | Missense | | | 0.23 | Neutral | 1 | 619520 | 1.61415E-06 | Uncertain significance | 822896 |
| 10 | p.Glu10Asp | c.30G>T | Missense | | | 0.23 | Neutral | 2 | 619520 | 3.22831E-06 | | |
| 11 | p.Pro11Leu | c.32C>T | Missense | | Functionally neutral | 0.29 | Neutral | 2 | 1454026 | 1.37549E-06 | flicting interpretations of pathogen | 584734 |
| 11 | p.Pro11Pro | c.33T>G | Synonymous | | | 0.48 | Neutral | 1 | 1454498 | 6.88E-07 | | |
| 11 | p.Pro11Pro | c.33T>C | Synonymous | | | 0.48 | Neutral | 1 | 1454498 | 6.88E-07 | | |
| 11 | p.Pro11Ser | c.31C>T | Missense | | | 0.21 | Neutral | 1 | 619456 | 1.61432E-06 | | |
| 12 | p.Ser12Ser | c.36G>T | Synonymous | | | 0.48 | Neutral | 1 | 1455028 | 6.87E-07 | Likely benign | 1559185 |
| 12 | p.Ser12Ser | c.36G>A | Synonymous | | | 0.48 | Neutral | 2 | 1455028 | 1.37454E-06 | Likely benign | 1172273 |
| 13 | p.Ala13Ala | c.39T>G | Synonymous | | | 0.48 | Neutral | 3 | 1607248 | 1.86654E-06 | Likely benign | 185104 |
| 13 | p.Ala13Asp | c.38C>A | Missense | | | 1.00 | Neutral | 2 | 622070 | 3.21507E-06 | Uncertain significance | 483348 |
| 13 | p.Ala13Gly | c.38C>G | Missense | | | 1.00 | Neutral | 1 | 622070 | 1.60754E-06 | | |
| 13 | p.Ala13Pro | c.37G>C | Missense | | | 1.00 | Neutral | 1 | 1454934 | 6.87E-07 | | |
| 13 | p.Ala13Thr | c.37G>A | Missense | | | 0.98 | Neutral | 3 | 1454934 | 2.06195E-06 | | |
| 14 | p.Asp14Glu | c.42C>A | Missense | | | 0.25 | Neutral | 2 | 622370 | 3.21352E-06 | Uncertain significance | 863866 |
| 15 | p.Trp15Arg | c.43T>C | Missense | | | 0.32 | Neutral | 3 | 622444 | 4.81971E-06 | | |
| 15 | p.Trp15Leu | c.44G>T | Missense | | | 0.70 | Neutral | 1 | 833110 | 1.20032E-06 | | |
| 16 | p.Leu16Arg | c.47T>G | Missense | Pathogenic | | 0.00 | Deleterious | 1 | 1455866 | 6.87E-07 | Pathogenic/Likely pathogenic | 219815 |
| 16 | p.Leu16Pro | c.47T>C | Missense | Pathogenic | | 0.00 | Deleterious | 3 | 1455866 | 2.06063E-06 | Pathogenic/Likely pathogenic | 649266 |
| 17 | p.Ala17Ala | c.51C>T | Synonymous | | | 0.48 | Neutral | 2 | 1454700 | 1.37485E-06 | Likely benign | 631355 |
| 17 | p.Ala17Ala | c.51C>G | Synonymous | | | 0.48 | Neutral | 3 | 1606690 | 1.86719E-06 | Likely benign | 532299 |
| 17 | p.Ala17Ala | c.51C>A | Synonymous | | | 0.48 | Neutral | 37 | 1606690 | 2.30287E-05 | Benign/Likely benign | 215440 |
| 17 | p.Ala17Pro | c.49G>C | Missense | | | 0.32 | Neutral | 1 | 1456126 | 6.87E-07 | | |
| 17 | p.Ala17Val | c.50C>T | Missense | | | 0.49 | Neutral | 1 | 619864 | 1.61326E-06 | Uncertain significance | 825441 |
| 18 | p.Thr18Ala | c.52A>G | Missense | | | 0.00 | Deleterious | 2 | 620722 | 3.22205E-06 | flicting interpretations of pathogen | 920669 |
| 18 | p.Thr18Lys | c.53C>A | Missense | | | 0.00 | Deleterious | 1 | 152026 | 6.57782E-06 | Uncertain significance | 491580 |
| 18 | p.Thr18Pro | c.52A>C | Missense | | Functionally deleterious | 0.00 | Deleterious | 2 | 620722 | 3.22205E-06 | Uncertain significance | 232702 |
| 18 | p.Thr18Thr | c.54G>C | Synonymous | | | 0.48 | Neutral | 1 | 621138 | 1.60995E-06 | Likely benign | 2560570 |
| 18 | p.Thr18Thr | c.54G>T | Synonymous | | | 0.48 | Neutral | 1 | 621138 | 1.60995E-06 | Likely benign | 514155 |
| 18 | p.Thr18Thr | c.54G>A | Synonymous | | | 0.48 | Neutral | 7 | 773146 | 9.05392E-06 | Likely benign | 1376569 |
| 19 | p.Ala19Ala | c.57C>G | Synonymous | | | 0.48 | Neutral | 1 | 1455558 | 6.87E-07 | Likely benign | 1077933 |
| 19 | p.Ala19Ala | c.57C>A | Synonymous | | | 0.48 | Neutral | 2 | 1455558 | 1.37404E-06 | | |
| 19 | p.Ala19Ala | c.57C>T | Synonymous | | | 0.48 | Neutral | 9 | 1607400 | 5.5991E-06 | Likely benign | 414094 |
| 19 | p.Ala19Thr | c.55G>A | Missense | | | 1.00 | Neutral | 1 | 621320 | 1.60948E-06 | | |
| 20 | p.Ala20Gly | c.59C>G | Missense | | Functionally deleterious | 0.00 | Deleterious | 2 | 1455702 | 1.37391E-06 | Uncertain significance | 220344 |
| 20 | p.Ala20Ser | c.58G>T | Missense | | | 0.00 | Deleterious | 2 | 1455404 | 1.37419E-06 | Uncertain significance | 664488 |
| 21 | p.Ala21Ala | c.63C>G | Synonymous | | | 0.48 | Neutral | 1 | 623564 | 1.60368E-06 | | |
| 21 | p.Ala21Asp | c.62C>A | Missense | | | 0.00 | Deleterious | 1 | 623328 | 1.60429E-06 | Uncertain significance | 1392805 |
| 22 | p.Arg22Arg | c.66G>C | Synonymous | | | 0.48 | Neutral | 1 | 152034 | 6.57748E-06 | | |
| 22 | p.Arg22Arg | c.64C>A | Synonymous | | | 0.48 | Neutral | 2 | 1457054 | 1.37263E-06 | Likely benign | 630709 |
| 22 | p.Arg22Arg | c.66G>A | Synonymous | | | 0.48 | Neutral | 2 | 624216 | 3.20402E-06 | Likely benign | 230052 |
| 22 | p.Arg22Gln | c.65G>A | Missense | | | 1.00 | Neutral | 2 | 1456978 | 1.3727E-06 | Uncertain significance | 491581 |
| 22 | p.Arg22Gly | c.64C>G | Missense | | | 0.00 | Deleterious | 2 | 1457056 | 1.37263E-06 | Uncertain significance | 863230 |
| 23 | p.Gly23Asp | c.68G>A | Missense | Pathogenic | | 0.00 | Deleterious | 2 | 833110 | 2.40064E-06 | Pathogenic/Likely pathogenic | 420108 |
| 24 | p.Arg24Arg | c.72G>T | Synonymous | | | 0.57 | Neutral | 3 | 1458286 | 2.05721E-06 | Likely benign | 463511 |
| 24 | p.Arg24Arg | c.72G>A | Synonymous | | | 0.57 | Neutral | 4 | 1458286 | 2.74295E-06 | | |
| 24 | p.Arg24Gln | c.71G>A | Missense | | | 0.00 | Deleterious | 6 | 1457942 | 4.11539E-06 | Uncertain significance | 406714 |
| 24 | p.Arg24Gly | c.70C>G | Missense | | | 0.13 | Neutral | 1 | 624724 | 1.60071E-06 | Uncertain significance | 1481554 |
| 24 | p.Arg24Pro | c.71G>C | Missense | Pathogenic | | 0.00 | Deleterious | 13 | 1609864 | 8.07522E-06 | Pathogenic/Likely pathogenic | 9415 |
| 25 | p.Val25Ile | c.73G>A | Missense | | | 0.71 | Neutral | 1 | 833110 | 1.20032E-06 | Uncertain significance | 1376284 |
| 25 | p.Val25Val | c.75A>T | Synonymous | | | 0.48 | Neutral | 1 | 1458796 | 6.85E-07 | | |
| 25 | p.Val25Val | c.75A>G | Synonymous | | | 0.48 | Neutral | 11 | 1458796 | 7.54046E-06 | Likely benign | 827146 |
| 26 | p.Glu26Gln | c.76G>C | Missense | | | 0.17 | Neutral | 2 | 625858 | 3.19561E-06 | Uncertain significance | 919572 |
| 26 | p.Glu26Glu | c.78G>A | Synonymous | | | 0.48 | Neutral | 1 | 833108 | 1.20032E-06 | Likely benign | 1761123 |
| 26 | p.Glu26Val | c.77A>T | Missense | | | 0.87 | Neutral | 1 | 1459254 | 6.85E-07 | | |
| 27 | p.Glu27Ala | c.80A>C | Missense | | | 0.12 | Neutral | 1 | 626382 | 1.59647E-06 | Uncertain significance | 628014 |
| 27 | p.Glu27Asp | c.81G>C | Missense | | | 0.00 | Deleterious | 1 | 626370 | 1.5965E-06 | Uncertain significance | 483354 |
| 27 | p.Glu27Asp | c.81G>T | Missense | | | 0.00 | Deleterious | 2 | 151976 | 1.316E-05 | Uncertain significance | 949317 |
| 28 | p.Val28Met | c.82G>A | Missense | | | 0.82 | Neutral | 1 | 626354 | 1.59654E-06 | Uncertain significance | 827567 |
| 28 | p.Val28Val | c.84G>A | Synonymous | | | 0.48 | Neutral | 2 | 1460032 | 1.36983E-06 | flicting interpretations of pathogen | 917571 |
| 29 | p.Arg29Arg | c.87G>C | Synonymous | | | 0.48 | Neutral | 1 | 1460268 | 6.85E-07 | Likely benign | 927182 |
| 29 | p.Arg29Arg | c.87G>A | Synonymous | | | 0.48 | Neutral | 20 | 1612530 | 1.24029E-05 | Likely benign | 230671 |
| 29 | p.Arg29Pro | c.86G>C | Missense | | | 0.00 | Deleterious | 1 | 626996 | 1.59491E-06 | Uncertain significance | 1041690 |
| 30 | p.Ala30Thr | c.88G>A | Missense | | | 0.00 | Deleterious | 2 | 833108 | 2.40065E-06 | | |
| 30 | p.Ala30Val | c.89C>T | Missense | | | 0.01 | Deleterious | 3 | 1460210 | 2.0545E-06 | Uncertain significance | 246070 |
| 31 | p.Leu31Leu | c.93G>A | Synonymous | | | 0.48 | Neutral | 2 | 1460414 | 1.36947E-06 | | |
| 32 | p.Leu32Pro | c.95T>C | Missense | Pathogenic | | 0.00 | Deleterious | 6 | 833108 | 7.20195E-06 | Pathogenic/Likely pathogenic | 236992 |
| 32 | p.Leu32Val | c.94C>G | Missense | | | 0.97 | Neutral | 3 | 1460394 | 2.05424E-06 | Uncertain significance | 532278 |
| 33 | p.Glu33Glu | c.99G>A | Synonymous | | | 0.48 | Neutral | 1 | 627582 | 1.59342E-06 | | |
| 34 | p.Ala34Ala | c.102G>C | Synonymous | | | 0.57 | Neutral | 33 | 779890 | 4.23137E-05 | | |
| 34 | p.Ala34Glu | c.101C>A | Missense | | | 0.22 | Neutral | 1 | 1460594 | 6.85E-07 | | |
| 34 | p.Ala34Ser | c.100G>T | Missense | | | 0.15 | Neutral | 1 | 1460706 | 6.85E-07 | | |
| 34 | p.Ala34Thr | c.100G>A | Missense | | | 0.02 | Neutral | 10 | 1460706 | 6.846E-06 | Uncertain significance | 489862 |
| 34 | p.Ala34Val | c.101C>T | Missense | | | 0.00 | Deleterious | 2 | 1460594 | 1.36931E-06 | Uncertain significance | 483329 |
| 35 | p.Gly35Ala | c.104G>C | Missense | | | 0.01 | Deleterious | 64 | 1612992 | 3.96778E-05 | flicting interpretations of pathogen | 463481 |
| 35 | p.Gly35Arg | c.103G>A | Missense | | Functionally neutral | 0.19 | Neutral | 4 | 1460806 | 2.73821E-06 | Uncertain significance | 483347 |
| 35 | p.Gly35Glu | c.104G>A | Missense | | Functionally neutral | 0.00 | Deleterious | 2 | 1460806 | 1.36911E-06 | Uncertain significance | 188292 |
| 35 | p.Gly35Val | c.104G>T | Missense | | Functionally deleterious | 0.00 | Deleterious | 1 | 1460806 | 6.85E-07 | Uncertain significance | 406709 |
| 36 | p.Ala36Ala | c.108G>C | Synonymous | | | 0.48 | Neutral | 5 | 1458686 | 3.42774E-06 | Likely benign | 822139 |
| 36 | p.Ala36Pro | c.106G>C | Missense | | | 0.00 | Deleterious | 1 | 1460710 | 6.85E-07 | | |
| 36 | p.Ala36Ser | c.106G>T | Missense | | | 0.11 | Neutral | 3 | 1460710 | 2.0538E-06 | Uncertain significance | 532277 |
| 36 | p.Ala36Thr | c.106G>A | Missense | | | 0.00 | Deleterious | 7 | 1612882 | 4.34006E-06 | Uncertain significance | 489863 |
| 36 | p.Ala36Val | c.107C>T | Missense | | | 0.96 | Neutral | 7 | 617654 | 1.13332E-05 | Uncertain significance | 483324 |
| 37 | p.Leu37Arg | c.110T>G | Missense | | | 1.00 | Neutral | 4 | 833108 | 4.8013E-06 | Uncertain significance | 1331763 |
| 37 | p.Leu37Leu | c.109C>T | Synonymous | | | 0.48 | Neutral | 1 | 1458504 | 6.86E-07 | Likely benign | 2563917 |
| 37 | p.Leu37Leu | c.111G>A | Synonymous | | | 0.48 | Neutral | 1 | 625942 | 1.59759E-06 | Likely benign | 532301 |
| 38 | p.Pro38Arg | c.113C>G | Missense | | | 0.00 | Deleterious | 1 | 1459894 | 6.85E-07 | Likely pathogenic | 1730766 |
| 38 | p.Pro38His | c.113C>A | Missense | | | 0.00 | Deleterious | 1 | 1459894 | 6.85E-07 | | |
| 38 | p.Pro38Ser | c.112C>T | Missense | | | 0.00 | Deleterious | 1 | 626538 | 1.59607E-06 | Uncertain significance | 822217 |
| 39 | p.Asn39Lys | c.117C>A | Missense | | | 0.00 | Deleterious | 1 | 626904 | 1.59514E-06 | Uncertain significance | 220249 |
| 40 | p.Ala40Gly | c.119C>G | Missense | | | 0.52 | Neutral | 1 | 626832 | 1.59532E-06 | | |
| 40 | p.Ala40Ser | c.118G>T | Missense | | | 0.56 | Neutral | 7 | 1459870 | 4.79495E-06 | Uncertain significance | 491563 |
| 41 | p.Pro41Ala | c.121C>G | Missense | | | 0.83 | Neutral | 2 | 1612966 | 1.23995E-06 | Uncertain significance | 406713 |
| 41 | p.Pro41Gln | c.122C>A | Missense | | Functionally neutral | 0.18 | Neutral | 6 | 1612890 | 3.72003E-06 | flicting interpretations of pathogen | 141111 |
| 41 | p.Pro41Leu | c.122C>T | Missense | | | 0.98 | Neutral | 3 | 1460848 | 2.0536E-06 | Uncertain significance | 1755177 |
| 41 | p.Pro41Pro | c.123G>A | Synonymous | | | 0.48 | Neutral | 1 | 627930 | 1.59253E-06 | flicting interpretations of pathogen | 923492 |
| 42 | p.Asn42Asn | c.126T>C | Synonymous | | | 0.48 | Neutral | 3 | 628174 | 4.77575E-06 | Likely benign | 1101776 |
| 42 | p.Asn42Ser | c.125A>G | Missense | | | 0.11 | Neutral | 3 | 1461208 | 2.0531E-06 | Uncertain significance | 406711 |
| 43 | p.Ser43Gly | c.127A>G | Missense | | | 0.06 | Neutral | 4 | 780364 | 5.12581E-06 | Uncertain significance | 630389 |
| 44 | p.Tyr44Cys | c.131A>G | Missense | | | 0.24 | Neutral | 1 | 833108 | 1.20032E-06 | Uncertain significance | 630012 |
| 44 | p.Tyr44Tyr | c.132C>T | Synonymous | | | 0.48 | Neutral | 4 | 1461440 | 2.73703E-06 | Likely benign | 483351 |
| 45 | p.Gly45Arg | c.133G>C | Missense | | | 0.71 | Neutral | 1 | 1461522 | 6.84E-07 | Uncertain significance | 463483 |
| 45 | p.Gly45Ser | c.133G>A | Missense | | | 0.00 | Deleterious | 1 | 1461522 | 6.84E-07 | Uncertain significance | 463482 |
| 46 | p.Arg46Gln | c.137G>A | Missense | | | 0.00 | Deleterious | 1 | 628584 | 1.59088E-06 | Uncertain significance | 631208 |
| 47 | p.Arg47Arg | c.139A>C | Synonymous | | | 0.48 | Neutral | 1 | 628580 | 1.59089E-06 | Likely benign | 1160771 |
| 48 | p.Pro48Pro | c.144G>A | Synonymous | | | 0.48 | Neutral | 3 | 780826 | 3.84209E-06 | Likely benign | 232233 |
| 48 | p.Pro48Thr | c.142C>A | Missense | Pathogenic | | 0.00 | Deleterious | 1 | 628604 | 1.59083E-06 | Pathogenic/Likely pathogenic | 188286 |

| | | | | | | | | | | | | |
|---|---|---|---|---|---|---|---|---|---|---|---|---|
| 49 | p.Ile49Ile | c.147C>A | Synonymous | | | 0.48 | Neutral | 77 | 1613900 | 4.77105E-05 | Likely benign | 184680 |
| 49 | p.Ile49Ser | c.146T>G | Missense | Likely pathogenic | | 0.00 | Deleterious | 2 | 1461744 | 1.36823E-06 | Pathogenic/Likely pathogenic | 430217 |
| 49 | p.Ile49Thr | c.146T>C | Missense | Likely pathogenic | | 0.00 | Deleterious | 178 | 1613956 | 0.000110288 | flicting interpretations of pathogen | 127523 |
| 50 | p.Gln50Arg | c.149A>G | Missense | | Functionally deleterious | 0.00 | Deleterious | 1 | 628592 | 1.59086E-06 | Pathogenic/Likely pathogenic | 232304 |
| 50 | p.Gln50Leu | c.149A>T | Missense | | | 0.00 | Deleterious | 1 | 628592 | 1.59086E-06 | Uncertain significance | 133878 |
| 51 | p.Val51Ile | c.151G>A | Missense | | | 0.02 | Neutral | 7 | 1597372 | 4.3822E-06 | Uncertain significance | 463485 |
| 51 | p.Val51Leu | c.151G>C | Missense | | | 0.28 | Neutral | 5 | 1445136 | 3.45988E-06 | | |
| 53 | p.Met53Ile | c.159G>C | Missense | Pathogenic | | 0.00 | Deleterious | 29 | 1597670 | 1.81514E-05 | Pathogenic | 9414 |
| 54 | p.Met54Ile | c.162G>A | Missense | | | 0.81 | Neutral | 1 | 612304 | 1.63318E-06 | Uncertain significance | 819708 |
| 54 | p.Met54Leu | c.160A>T | Missense | | | 0.94 | Neutral | 2 | 1445492 | 1.38361E-06 | Uncertain significance | 1483043 |
| 54 | p.Met54Leu | c.160A>C | Missense | | | 0.94 | Neutral | 11 | 1597750 | 6.88468E-06 | flicting interpretations of pathogen | 463486 |
| 55 | p.Gly55Ala | c.164G>C | Missense | | | 0.00 | Deleterious | 1 | 1445362 | 6.92E-07 | Uncertain significance | 2563916 |
| 55 | p.Gly55Ser | c.163G>A | Missense | | | 0.00 | Deleterious | 1 | 612334 | 1.6331E-06 | | |
| 55 | p.Gly55Val | c.164G>T | Missense | | | 0.00 | Deleterious | 1 | 152332 | 6.56461E-06 | Uncertain significance | 2136747 |
| 56 | p.Ser56Ile | c.167G>T | Missense | Pathogenic | | 0.00 | Deleterious | 2 | 1445436 | 1.38367E-06 | Pathogenic/Likely pathogenic | 9425 |
| 57 | p.Ala57Asp | c.170C>A | Missense | | | 0.92 | Neutral | 11 | 1597598 | 6.88534E-06 | Uncertain significance | 483325 |
| 57 | p.Ala57Gly | c.170C>G | Missense | | | 0.45 | Neutral | 22 | 1597598 | 1.37707E-05 | Uncertain significance | 187272 |
| 57 | p.Ala57Val | c.170C>T | Missense | Benign | | 0.89 | Neutral | 163 | 1597598 | 0.000102028 | flicting interpretations of pathogen | 220562 |
| 58 | p.Arg58Gly | c.172C>G | Missense | | | 0.98 | Neutral | 8 | 1445042 | 5.53617E-06 | Uncertain significance | 532285 |
| 59 | p.Val59Gly | c.176T>G | Missense | Pathogenic | | 0.00 | Deleterious | 3 | 1444950 | 2.0762E-06 | Pathogenic/Likely pathogenic | 9423 |
| 59 | p.Val59Met | c.175G>A | Missense | | | 0.34 | Neutral | 1 | 612098 | 1.63373E-06 | Uncertain significance | 1007891 |
| 60 | p.Ala60Thr | c.178G>A | Missense | | | 0.44 | Neutral | 2 | 764162 | 2.61725E-06 | Uncertain significance | 406705 |
| 61 | p.Glu61Asp | c.183G>C | Missense | | | 0.56 | Neutral | 1 | 1443704 | 6.93E-07 | Uncertain significance | 236982 |
| 61 | p.Glu61Lys | c.181G>A | Missense | | | 0.87 | Neutral | 1 | 611732 | 1.6347E-06 | | |
| 62 | p.Leu62Val | c.184C>G | Missense | | | 0.98 | Neutral | 1 | 152214 | 6.5697E-06 | Uncertain significance | 969307 |
| 63 | p.Leu63Pro | c.188T>C | Missense | | | 0.00 | Deleterious | 2 | 1443526 | 1.3855E-06 | flicting interpretations of pathogen | 1367946 |
| 66 | p.His66Arg | c.197A>G | Missense | | Functionally neutral | 0.26 | Neutral | 303 | 1594708 | 0.000190003 | flicting interpretations of pathogen | 246117 |
| 66 | p.His66Gln | c.198C>G | Missense | | | 0.47 | Neutral | 9 | 1442770 | 6.238E-06 | Uncertain significance | 573530 |
| 67 | p.Gly67Arg | c.199G>C | Missense | | Functionally neutral | 1.00 | Neutral | 4 | 1442608 | 2.77276E-06 | flicting interpretations of pathogen | 216272 |
| 67 | p.Gly67Asp | c.200G>A | Missense | | | 0.86 | Neutral | 3 | 761738 | 3.93836E-06 | Uncertain significance | 216273 |
| 67 | p.Gly67Ser | c.199G>A | Missense | | | 0.96 | Neutral | 2 | 1442606 | 1.38638E-06 | Uncertain significance | 925139 |
| 68 | p.Ala68Gly | c.203C>G | Missense | | | 0.38 | Neutral | 13 | 1441598 | 9.01777E-06 | Uncertain significance | 628976 |
| 68 | p.Ala68Val | c.203C>T | Missense | | | 0.00 | Deleterious | 1 | 1441598 | 6.94E-07 | flicting interpretations of pathogen | 406701 |
| 69 | p.Glu69Gly | c.206A>G | Missense | | | 0.00 | Deleterious | 67 | 1593674 | 4.20412E-05 | flicting interpretations of pathogen | 186615 |
| 70 | p.Pro70Arg | c.209C>G | Missense | | | 0.72 | Neutral | 1 | 1442188 | 6.93E-07 | Uncertain significance | 185936 |
| 70 | p.Pro70Thr | c.208C>A | Missense | | | 0.05 | Neutral | 4 | 1441902 | 2.77411E-06 | Uncertain significance | 1374577 |
| 71 | p.Asn71Ser | c.212A>G | Missense | Likely pathogenic | | 0.00 | Deleterious | 6 | 1593656 | 3.76493E-06 | Likely pathogenic | 418121 |
| 71 | p.Asn71Thr | c.212A>C | Missense | | | 0.00 | Deleterious | 4 | 1441448 | 2.77499E-06 | Uncertain significance | 1786361 |
| 73 | p.Ala73Ala | c.219C>A | Synonymous | | | 0.57 | Neutral | 2 | 152226 | 1.31384E-05 | Likely benign | 653470 |
| 73 | p.Ala73Thr | c.217G>A | Missense | | | 0.68 | Neutral | 1 | 606262 | 1.64945E-06 | Uncertain significance | 1986070 |
| 74 | p.Asp74Asn | c.220G>A | Missense | | | 0.00 | Deleterious | 1 | 605542 | 1.65141E-06 | Uncertain significance | 483340 |
| 75 | p.Pro75Leu | c.224C>T | Missense | | | 0.29 | Neutral | 9 | 1596408 | 5.63766E-06 | Uncertain significance | 483326 |
| 75 | p.Pro75Ser | c.223C>T | Missense | | | 0.31 | Neutral | 1 | 610794 | 1.63721E-06 | Uncertain significance | 644311 |
| 76 | p.Ala76Ser | c.226G>T | Missense | | | 0.01 | Deleterious | 1 | 152148 | 6.57255E-06 | Uncertain significance | 2059365 |
| 76 | p.Ala76Thr | c.226G>A | Missense | | | 0.14 | Neutral | 18 | 1595206 | 1.12838E-05 | Uncertain significance | 489866 |
| 77 | p.Thr77Ala | c.229A>G | Missense | | | 0.01 | Deleterious | 1 | 611264 | 1.63595E-06 | Uncertain significance | 567670 |
| 77 | p.Thr77Asn | c.230C>A | Missense | | | 0.04 | Neutral | 2 | 1445234 | 1.38386E-06 | | |
| 78 | p.Leu78Phe | c.232C>T | Missense | | | 0.98 | Neutral | 2 | 1445644 | 1.38347E-06 | Uncertain significance | 630762 |
| 79 | p.Thr79Asn | c.236C>A | Missense | | | 0.80 | Neutral | 3 | 1603784 | 1.87058E-06 | Uncertain significance | 491570 |
| 79 | p.Thr79Ile | c.236C>T | Missense | | | 0.79 | Neutral | 5 | 1451540 | 3.44462E-06 | Uncertain significance | 630548 |
| 80 | p.Arg80Gln | c.239G>A | Missense | | | 0.00 | Deleterious | 6 | 1451726 | 4.13301E-06 | Uncertain significance | 376385 |
| 80 | p.Arg80Leu | c.239G>T | Missense | | | 0.00 | Deleterious | 2 | 1451724 | 1.37767E-06 | | |
| 80 | p.Arg80Pro | c.239G>C | Missense | | | 0.00 | Deleterious | 1 | 152196 | 6.57047E-06 | | |
| 81 | p.Pro81Leu | c.242C>T | Missense | | Functionally deleterious | 0.00 | Deleterious | 1 | 618752 | 1.61616E-06 | Likely pathogenic | 833629 |
| 81 | p.Pro81Ser | c.241C>T | Missense | | | 0.00 | Deleterious | 1 | 152230 | 6.56901E-06 | Uncertain significance | 664812 |
| 82 | p.Val82Leu | c.244G>T | Missense | | | 0.03 | Neutral | 2 | 1451914 | 1.37749E-06 | Uncertain significance | 406718 |
| 82 | p.Val82Met | c.244G>A | Missense | | | 0.33 | Neutral | 2 | 1451914 | 1.37749E-06 | Uncertain significance | 1063226 |
| 83 | p.His83Arg | c.248A>G | Missense | Pathogenic | | 0.00 | Deleterious | 1 | 152338 | 6.56435E-06 | flicting interpretations of pathogen | 376379 |
| 83 | p.His83Asn | c.247C>A | Missense | | | 0.00 | Deleterious | 1 | 618816 | 1.61599E-06 | | |
| 83 | p.His83Gln | c.249C>G | Missense | | | 0.00 | Deleterious | 1 | 1452118 | 6.89E-07 | Pathogenic | 376381 |
| 83 | p.His83Gln | c.249C>A | Missense | | | 0.00 | Deleterious | 2 | 1452116 | 1.3773E-06 | flicting interpretations of pathogen | 429110 |
| 84 | p.Asp84Ala | c.251A>C | Missense | | Functionally deleterious | 0.00 | Deleterious | 7 | 1452176 | 4.82035E-06 | flicting interpretations of pathogen | 142882 |
| 84 | p.Asp84Tyr | c.250G>T | Missense | | Functionally deleterious | 0.00 | Deleterious | 1 | 619038 | 1.61541E-06 | flicting interpretations of pathogen | 376306 |
| 85 | p.Ala85Ser | c.253G>T | Missense | | | 0.07 | Neutral | 5 | 771240 | 6.48307E-06 | Uncertain significance | 532289 |
| 85 | p.Ala85Thr | c.253G>A | Missense | | | 0.00 | Deleterious | 3 | 619004 | 4.8465E-06 | Uncertain significance | 236983 |
| 86 | p.Ala86Ser | c.256G>T | Missense | | | 0.15 | Neutral | 1 | 619124 | 1.61519E-06 | Uncertain significance | 1793230 |
| 86 | p.Ala86Thr | c.256G>A | Missense | | | 0.96 | Neutral | 3 | 771478 | 3.88864E-06 | | |
| 87 | p.Arg87Trp | c.259C>T | Missense | Likely pathogenic | | 0.00 | Deleterious | 6 | 1604616 | 3.73921E-06 | Likely pathogenic | 406707 |
| 88 | p.Glu88Gln | c.262G>C | Missense | | | 0.00 | Deleterious | 1 | 619464 | 1.6143E-06 | Uncertain significance | 619828 |
| 89 | p.Gly89Ser | c.265G>A | Missense | | | 0.00 | Deleterious | 4 | 619634 | 6.45542E-06 | Uncertain significance | 9408 |
| 91 | p.Leu91Arg | c.272T>G | Missense | | | 0.35 | Neutral | 1 | 1453016 | 6.88E-07 | | |
| 92 | p.Asp92Glu | c.276C>A | Missense | | | 0.07 | Neutral | 2 | 619946 | 3.22609E-06 | Uncertain significance | 645261 |
| 93 | p.Thr93Met | c.278C>T | Missense | | | 0.00 | Deleterious | 1 | 619998 | 1.61291E-06 | Uncertain significance | 1796019 |
| 96 | p.Val96Gly | c.287T>G | Missense | | | 0.02 | Neutral | 1 | 1453218 | 6.88E-07 | | |
| 96 | p.Val96Leu | c.286G>T | Missense | | | 0.00 | Deleterious | 1 | 1453266 | 6.88E-07 | Uncertain significance | 2106366 |
| 96 | p.Val96Met | c.286G>A | Missense | | | 0.62 | Neutral | 1 | 1453266 | 6.88E-07 | Uncertain significance | 821875 |
| 98 | p.His98Asn | c.292C>A | Missense | | | 0.63 | Neutral | 1 | 620158 | 1.61249E-06 | | |
| 98 | p.His98Tyr | c.292C>T | Missense | | | 0.68 | Neutral | 1 | 620158 | 1.61249E-06 | Uncertain significance | 419560 |
| 99 | p.Arg99Gln | c.296G>A | Missense | | | 0.18 | Neutral | 4 | 1453046 | 2.75284E-06 | flicting interpretations of pathogen | 628568 |
| 99 | p.Arg99Gly | c.295C>G | Missense | | Functionally neutral | 0.43 | Neutral | 7 | 1453094 | 4.81731E-06 | flicting interpretations of pathogen | 372062 |
| 99 | p.Arg99Trp | c.295C>T | Missense | | | 0.00 | Deleterious | 3 | 1453094 | 2.06456E-06 | Uncertain significance | 483350 |
| 100 | p.Ala100Ser | c.298G>T | Missense | Benign | | 0.99 | Neutral | 98 | 1605346 | 6.1046E-05 | flicting interpretations of pathogen | 127524 |
| 100 | p.Ala100Thr | c.298G>A | Missense | | | 0.37 | Neutral | 1 | 1453014 | 6.88E-07 | Uncertain significance | 629701 |
| 100 | p.Ala100Val | c.299C>T | Missense | | | 1.00 | Neutral | 2 | 772064 | 2.59046E-06 | Uncertain significance | 924911 |
| 101 | p.Gly101Arg | c.301G>A | Missense | | Functionally neutral | 0.00 | Deleterious | 4 | 1605208 | 2.49189E-06 | Uncertain significance | 463494 |
| 101 | p.Gly101Arg | c.301G>C | Missense | | Functionally neutral | 0.00 | Deleterious | 7 | 1452982 | 4.81768E-06 | Uncertain significance | 216274 |
| 101 | p.Gly101Trp | c.301G>T | Missense | Pathogenic | | 0.00 | Deleterious | 12 | 1605210 | 7.47566E-06 | Pathogenic | 9412 |
| 102 | p.Ala102Thr | c.304G>A | Missense | | | 0.00 | Deleterious | 2 | 772270 | 2.58977E-06 | Uncertain significance | 463495 |
| 102 | p.Ala102Val | c.305C>T | Missense | | | 0.00 | Deleterious | 5 | 1452980 | 3.4412E-06 | Uncertain significance | 234071 |
| 103 | p.Arg103Gln | c.308G>A | Missense | | | 0.51 | Neutral | 1 | 833110 | 1.20032E-06 | Uncertain significance | 1141816 |
| 103 | p.Arg103Trp | c.307C>T | Missense | | | 0.85 | Neutral | 12 | 1605324 | 7.47513E-06 | Uncertain significance | 423630 |
| 105 | p.Asp105Asn | c.313G>A | Missense | | | 0.14 | Neutral | 2 | 1453262 | 1.37621E-06 | Uncertain significance | 491573 |
| 105 | p.Asp105Glu | c.315C>A | Missense | | | 0.00 | Deleterious | 20 | 1605504 | 1.24571E-05 | flicting interpretations of pathogen | 406702 |
| 106 | p.Val106Ala | c.317T>C | Missense | | | 0.02 | Neutral | 2 | 833110 | 2.40064E-06 | Uncertain significance | 463497 |
| 106 | p.Val106Gly | c.317T>G | Missense | | | 0.00 | Deleterious | 1 | 833110 | 1.20032E-06 | | |
| 106 | p.Val106Leu | c.316G>C | Missense | | | 0.00 | Deleterious | 1 | 1453226 | 6.88E-07 | Uncertain significance | 1318653 |
| 106 | p.Val106Leu | c.316G>T | Missense | | | 0.00 | Deleterious | 1 | 152232 | 6.56892E-06 | Uncertain significance | 1728421 |
| 106 | p.Val106Met | c.316G>A | Missense | | | 0.57 | Neutral | 1 | 1453226 | 6.88E-07 | Uncertain significance | 1392243 |
| 107 | p.Arg107Cys | c.319C>T | Missense | | | 0.00 | Deleterious | 3 | 1453408 | 2.06411E-06 | flicting interpretations of pathogen | 483352 |
| 107 | p.Arg107Gly | c.319C>G | Missense | | | 0.33 | Neutral | 2 | 1453408 | 1.37608E-06 | | |
| 107 | p.Arg107His | c.320G>A | Missense | | | 0.32 | Neutral | 25 | 1605852 | 1.55681E-05 | flicting interpretations of pathogen | 182413 |
| 107 | p.Arg107Leu | c.320G>T | Missense | | | 0.24 | Neutral | 1 | 1453632 | 6.88E-07 | | |
| 107 | p.Arg107Ser | c.319C>A | Missense | | | 0.09 | Neutral | 1 | 1453408 | 6.88E-07 | Uncertain significance | 463498 |
| 108 | p.Asp108Val | c.323A>T | Missense | | | 0.00 | Deleterious | 1 | 620726 | 1.61102E-06 | | |
| 109 | p.Ala109Ala | c.327C>T | Synonymous | | | 0.57 | Neutral | 1 | 1454144 | 6.88E-07 | Likely benign | 1123298 |
| 109 | p.Ala109Pro | c.325G>C | Missense | | Functionally deleterious | 0.00 | Deleterious | 19 | 1606256 | 1.18287E-05 | Uncertain significance | 127525 |
| 110 | p.Trp110Arg | c.328T>C | Missense | | | 0.92 | Neutral | 1 | 152190 | 6.57073E-06 | Uncertain significance | 2109926 |
| 110 | p.Trp110Gly | c.328T>G | Missense | | | 0.89 | Neutral | 1 | 621072 | 1.61012E-06 | | |
| 111 | p.Gly111Ser | c.331G>A | Missense | | Functionally neutral | 0.81 | Neutral | 6 | 1454378 | 4.12547E-06 | Uncertain significance | 463499 |
| 111 | p.Gly111Val | c.332G>T | Missense | | | 0.00 | Deleterious | 1 | 621180 | 1.60984E-06 | Uncertain significance | 1487529 |
| 112 | p.Arg112Gly | c.334C>G | Missense | Pathogenic | | 0.00 | Deleterious | 2 | 1454592 | 1.37496E-06 | Pathogenic/Likely pathogenic | 233484 |
| 112 | p.Arg112His | c.335G>A | Missense | | | 0.42 | Neutral | 1 | 1454582 | 6.87E-07 | Uncertain significance | 142887 |
| 112 | p.Arg112Leu | c.335G>T | Missense | | | 0.00 | Deleterious | 7 | 1606934 | 4.35612E-06 | Uncertain significance | 823661 |
| 113 | p.Leu113Val | c.337C>G | Missense | | | 0.91 | Neutral | 1 | 152250 | 6.56814E-06 | | |
| 114 | p.Pro114Ser | c.340C>T | Missense | | | 0.00 | Deleterious | 1 | 152346 | 6.56401E-06 | Uncertain significance | 1405931 |
| 115 | p.Val115Glu | c.344T>A | Missense | | | 0.00 | Deleterious | 10 | 1607670 | 6.22018E-06 | Uncertain significance | 246437 |
| 115 | p.Val115Leu | c.343G>T | Missense | | | 0.13 | Neutral | 32 | 1607650 | 1.99048E-05 | Uncertain significance | 419130 |
| 115 | p.Val115Met | c.343G>A | Missense | | | 0.37 | Neutral | 3 | 1455412 | 2.06127E-06 | Uncertain significance | 1469323 |
| 116 | p.Asp116Glu | c.348C>G | Missense | | | 0.30 | Neutral | 1 | 622926 | 1.60533E-06 | | |
| 116 | p.Asp116Tyr | c.346G>T | Missense | | | 0.56 | Neutral | 1 | 622770 | 1.60573E-06 | Uncertain significance | 2563909 |
| 117 | p.Leu117Gln | c.350T>A | Missense | | | 0.07 | Neutral | 1 | 833110 | 1.20032E-06 | | |
| 118 | p.Ala118Gly | c.353C>G | Missense | | | 0.13 | Neutral | 1 | 623472 | 1.60392E-06 | | |
| 118 | p.Ala118Thr | c.352G>A | Missense | | | 0.03 | Neutral | 7 | 1608806 | 4.35105E-06 | Uncertain significance | 491574 |
| 118 | p.Ala118Val | c.353C>T | Missense | | | 0.01 | Deleterious | 1 | 623472 | 1.60392E-06 | flicting interpretations of pathogen | 406719 |
| 119 | p.Glu119Glu | c.357G>A | Synonymous | | | 0.48 | Neutral | 1 | 623920 | 1.60277E-06 | | |
| 119 | p.Glu119Lys | c.355G>A | Missense | | | 0.81 | Neutral | 2 | 1456610 | 1.37305E-06 | Uncertain significance | 570565 |
| 120 | p.Glu120Asp | c.360G>T | Missense | | | 0.21 | Neutral | 5 | 1457182 | 3.43128E-06 | Uncertain significance | 823988 |
| 120 | p.Glu120Gln | c.358G>C | Missense | | | 0.06 | Neutral | 1 | 623948 | 1.6027E-06 | | |
| 120 | p.Glu120Glu | c.360G>A | Synonymous | | | 0.57 | Neutral | 2 | 1457184 | 1.37251E-06 | Likely benign | 463502 |
| 120 | p.Glu120Val | c.359A>T | Missense | | | 0.33 | Neutral | 1 | 624016 | 1.60252E-06 | | |
| 121 | p.Leu121Arg | c.362T>G | Missense | | | 0.00 | Deleterious | 1 | 152228 | 6.56909E-06 | | |
| 121 | p.Leu121Leu | c.361C>T | Synonymous | | | 0.48 | Neutral | 146 | 1609492 | 9.07119E-05 | Benign/Likely benign | 630387 |
| 121 | p.Leu121Met | c.361C>A | Missense | | | 0.00 | Deleterious | 1 | 1457230 | 6.86E-07 | | |
| 121 | p.Leu121Val | c.361C>G | Missense | | | 0.97 | Neutral | 29 | 1609610 | 1.80168E-05 | flicting interpretations of pathogen | 219905 |
| 122 | p.Gly122Asp | c.365G>A | Missense | | | 0.95 | Neutral | 1 | 1457356 | 6.86E-07 | | |
| 122 | p.Gly122Gly | c.366C>G | Synonymous | | | 0.62 | Neutral | 1 | 624330 | 1.60172E-06 | | |
| 122 | p.Gly122Gly | c.366C>T | Synonymous | | | 0.62 | Neutral | 1 | 624330 | 1.60172E-06 | Likely benign | 1133481 |
| 122 | p.Gly122Val | c.365G>T | Missense | | | 0.49 | Neutral | 38 | 1609606 | 2.36083E-05 | flicting interpretations of pathogen | 182419 |
| 123 | p.His123Gln | c.369T>A | Missense | Benign | | 0.10 | Neutral | 359 | 1610194 | 0.000222955 | flicting interpretations of pathogen | 127526 |
| 123 | p.His123His | c.369T>C | Synonymous | | | 0.62 | Neutral | 15 | 1610076 | 9.31633E-06 | Likely benign | 220829 |
| 124 | p.Arg124Cys | c.370C>T | Missense | | Functionally neutral | 0.39 | Neutral | 73 | 1610132 | 4.53379E-05 | flicting interpretations of pathogen | 231362 |
| 124 | p.Arg124Gly | c.370C>G | Missense | | | 0.00 | Deleterious | 9 | 1610014 | 5.59001E-06 | Uncertain significance | 582036 |
| 124 | p.Arg124His | c.371G>A | Missense | | | 0.80 | Neutral | 1 | 624836 | 1.60042E-06 | | |

| | | | | | | | | | | | | |
|---|---|---|---|---|---|---|---|---|---|---|---|---|
| 125 | p.Asp125Asn | c.373G>A | Missense | | | 0.62 | Neutral | 2 | 1610302 | 1.242E-06 | Uncertain significance | 858988 |
| 125 | p.Asp125Glu | c.375T>A | Missense | | | 0.80 | Neutral | 2 | 777462 | 2.57247E-06 | Uncertain significance | 1329079 |
| 125 | p.Asp125His | c.373G>C | Missense | | Functionally neutral | 0.95 | Neutral | 569 | 1610420 | 0.000353324 | flicting interpretations of pathoger | 41577 |
| 126 | p.Val126Asp | c.377T>A | Missense | Pathogenic | | 0.00 | Deleterious | 3 | 1458484 | 2.05693E-06 | Pathogenic | 9420 |
| 126 | p.Val126Leu | c.376G>C | Missense | | | 1.00 | Neutral | 1 | 1458368 | 6.86E-07 | Uncertain significance | 489868 |
| 126 | p.Val126Phe | c.376G>T | Missense | | | 0.75 | Neutral | 1 | 1458368 | 6.86E-07 | | |
| 126 | p.Val126Val | c.378C>T | Synonymous | | | 0.48 | Neutral | 1 | 1458494 | 6.86E-07 | Likely benign | 1692141 |
| 126 | p.Val126Val | c.378C>G | Synonymous | | | 0.48 | Neutral | 1 | 1458494 | 6.86E-07 | | |
| 127 | p.Ala127Pro | c.379G>C | Missense | | Functionally deleterious | 0.00 | Deleterious | 1 | 1458444 | 6.86E-07 | flicting interpretations of pathoger | 571866 |
| 127 | p.Ala127Ser | c.379G>T | Missense | Likely benign | | 0.23 | Neutral | 1532 | 1610798 | 0.000951081 | Benign/Likely benign | 41578 |
| 127 | p.Ala127Val | c.380C>T | Missense | | | 0.83 | Neutral | 1 | 152238 | 6.56866E-06 | Uncertain significance | 2030143 |
| 128 | p.Arg128Gln | c.383G>A | Missense | | | 0.93 | Neutral | 19 | 1610916 | 1.17945E-05 | Uncertain significance | 581636 |
| 128 | p.Arg128Pro | c.383G>C | Missense | | | 0.00 | Deleterious | 1 | 1458694 | 6.86E-07 | Uncertain significance | 483344 |
| 130 | p.Leu130Leu | c.388C>T | Synonymous | | | 0.48 | Neutral | 2 | 778188 | 2.57007E-06 | flicting interpretations of pathoger | 491575 |
| 131 | p.Arg131Cys | c.391C>T | Missense | | | 0.00 | Deleterious | 1 | 1459134 | 6.85E-07 | Uncertain significance | 1350948 |
| 131 | p.Arg131His | c.392G>A | Missense | | | 0.64 | Neutral | 2 | 1459184 | 1.37063E-06 | Uncertain significance | 620615 |
| 132 | p.Ala132Ala | c.396G>C | Synonymous | | | 0.48 | Neutral | 1 | 152250 | 6.56814E-06 | Likely benign | 382703 |
| 132 | p.Ala132Ala | c.396G>T | Synonymous | | | 0.48 | Neutral | 1 | 626296 | 1.59669E-06 | | |
| 132 | p.Ala132Val | c.395C>T | Missense | | | 0.05 | Neutral | 5 | 1459290 | 3.42632E-06 | Uncertain significance | 630452 |
| 133 | p.Ala133Ala | c.399T>C | Synonymous | | | 0.48 | Neutral | 2 | 1459450 | 1.37038E-06 | | |
| 134 | p.Ala134Ala | c.402G>A | Synonymous | | | 0.57 | Neutral | 1 | 1459422 | 6.85E-07 | flicting interpretations of pathoger | 583214 |
| 134 | p.Ala134Ala | c.402G>T | Synonymous | | | 0.57 | Neutral | 9 | 1611668 | 5.58428E-06 | Likely benign | 236989 |
| 134 | p.Ala134Pro | c.400G>C | Missense | | | 0.68 | Neutral | 5 | 1611668 | 3.10238E-06 | Uncertain significance | 495536 |
| 134 | p.Ala134Val | c.401C>T | Missense | | | 0.93 | Neutral | 3 | 626164 | 4.79108E-06 | Uncertain significance | 650117 |
| 135 | p.Gly135Glu | c.404G>A | Missense | | | 0.00 | Deleterious | 5 | 1611926 | 3.10188E-06 | Uncertain significance | 463504 |
| 135 | p.Gly135Gly | c.405G>A | Synonymous | | | 0.48 | Neutral | 306 | 1611984 | 0.000189828 | Benign/Likely benign | 383182 |
| 135 | p.Gly135Val | c.404G>T | Missense | | | 0.37 | Neutral | 1 | 1459674 | 6.85E-07 | Uncertain significance | 843913 |
| 136 | p.Gly136Ala | c.407G>C | Missense | | | 0.66 | Neutral | 2 | 152212 | 1.31396E-05 | Uncertain significance | 483335 |
| 136 | p.Gly136Asp | c.407G>A | Missense | | | 0.00 | Deleterious | 5 | 626354 | 7.98271E-06 | Uncertain significance | 925925 |
| 136 | p.Gly136Ser | c.406G>A | Missense | | | 0.03 | Neutral | 5 | 1612104 | 3.10154E-06 | Uncertain significance | 483342 |
| 137 | p.Thr137Pro | c.409A>C | Missense | | | 0.70 | Neutral | 1 | 1459840 | 6.85E-07 | | |
| 137 | p.Thr137Ser | c.410C>G | Missense | | | 0.77 | Neutral | 1 | 626710 | 1.59563E-06 | | |
| 138 | p.Arg138Gly | c.412A>G | Missense | | | 0.77 | Neutral | 15 | 1612210 | 9.304E-06 | Uncertain significance | 233990 |
| 139 | p.Gly139Arg | c.415G>C | Missense | | Functionally neutral | 0.33 | Neutral | 27 | 1460010 | 1.8493E-05 | flicting interpretations of pathoger | 216276 |
| 139 | p.Gly139Asp | c.416G>A | Missense | | | 0.01 | Deleterious | 9 | 1612280 | 5.58216E-06 | Uncertain significance | 491576 |
| 139 | p.Gly139Ser | c.415G>A | Missense | | | 0.05 | Neutral | 4 | 1460010 | 2.73971E-06 | Uncertain significance | 141419 |
| 139 | p.Gly139Val | c.416G>T | Missense | | | 0.83 | Neutral | 1 | 1459922 | 6.85E-07 | Uncertain significance | 2172999 |
| 140 | p.Ser140Ser | c.420T>C | Synonymous | | | 0.48 | Neutral | 2 | 1460096 | 1.36977E-06 | Likely benign | 1738745 |
| 141 | p.Asn141Asn | c.423C>T | Synonymous | | | 0.48 | Neutral | 1 | 1460072 | 6.85E-07 | Likely benign | 924813 |
| 141 | p.Asn141Asp | c.421A>G | Missense | | | 0.87 | Neutral | 1 | 152230 | 6.56901E-06 | Uncertain significance | 631362 |
| 142 | p.His142Arg | c.425A>G | Missense | | | 1.00 | Neutral | 25 | 1612402 | 1.55048E-05 | Uncertain significance | 184564 |
| 142 | p.His142Gln | c.426T>G | Missense | | | 1.00 | Neutral | 1 | 833110 | 1.20032E-06 | Uncertain significance | 246046 |
| 142 | p.His142Tyr | c.424C>T | Missense | | | 1.00 | Neutral | 2 | 626994 | 3.18982E-06 | Uncertain significance | 824721 |
| 143 | p.Ala143Ala | c.429C>A | Synonymous | | | 0.48 | Neutral | 1 | 1460104 | 6.85E-07 | Likely benign | 414090 |
| 143 | p.Ala143Gly | c.428C>G | Missense | | | 0.56 | Neutral | 5 | 626918 | 7.97552E-06 | Uncertain significance | 630386 |
| 143 | p.Ala143Thr | c.427G>A | Missense | | Functionally neutral | 0.72 | Neutral | 16 | 1460132 | 1.09579E-05 | Uncertain significance | 245681 |
| 144 | p.Arg144Arg | c.432C>T | Synonymous | | | 0.48 | Neutral | 1 | 627030 | 1.59482E-06 | | |
| 144 | p.Arg144Cys | c.430C>T | Missense | Likely benign | | 0.98 | Neutral | 763 | 1612398 | 0.000473208 | Benign/Likely benign | 41579 |
| 144 | p.Arg144His | c.431G>A | Missense | | | 0.68 | Neutral | 1 | 152236 | 6.56875E-06 | Uncertain significance | 406723 |
| 144 | p.Arg144Leu | c.431G>T | Missense | | | 0.89 | Neutral | 10 | 1612274 | 6.20242E-06 | Uncertain significance | 463505 |
| 144 | p.Arg144Ser | c.430C>A | Missense | | | 0.75 | Neutral | 1 | 152216 | 6.56961E-06 | | |
| 145 | p.Ile145Met | c.435A>G | Missense | | | 0.12 | Neutral | 1 | 627006 | 1.59488E-06 | | |
| 145 | p.Ile145Thr | c.434T>C | Missense | | | 0.49 | Neutral | 7 | 1460132 | 4.79409E-06 | flicting interpretations of pathoger | 182420 |
| 145 | p.Ile145Val | c.433A>G | Missense | | | 0.21 | Neutral | 1 | 1460158 | 6.85E-07 | Uncertain significance | 645545 |
| 146 | p.Asp146Asn | c.436G>A | Missense | | | 0.12 | Neutral | 1 | 1460116 | 6.85E-07 | Uncertain significance | 578446 |
| 146 | p.Asp146Glu | c.438T>A | Missense | | | 0.61 | Neutral | 1 | 833110 | 1.20032E-06 | | |
| 146 | p.Asp146His | c.436G>C | Missense | | | 0.22 | Neutral | 1 | 1460116 | 6.85E-07 | Uncertain significance | 824834 |
| 146 | p.Asp146Val | c.437A>T | Missense | | | 0.59 | Neutral | 1 | 152216 | 6.56961E-06 | Uncertain significance | 928297 |
| 147 | p.Ala147Ala | c.441C>A | Synonymous | | | 0.48 | Neutral | 4 | 1459910 | 2.73989E-06 | Likely benign | 391388 |
| 147 | p.Ala147Val | c.440C>T | Missense | | | 0.31 | Neutral | 3 | 779052 | 3.85083E-06 | Uncertain significance | 406716 |
| 148 | p.Ala148Ala | c.444G>A | Synonymous | | | 0.48 | Neutral | 5 | 1459850 | 3.42501E-06 | Likely benign | 923208 |
| 148 | p.Ala148Gly | c.443C>G | Missense | | | 0.38 | Neutral | 1 | 1459882 | 6.85E-07 | | |
| 148 | p.Ala148Thr | c.442G>A | Missense | Likely benign | | 0.40 | Neutral | 39384 | 1612214 | 0.024428519 | Benign | 41580 |
| 148 | p.Ala148Val | c.443C>T | Missense | | | 0.00 | Deleterious | 5 | 1612130 | 3.10149E-06 | Uncertain significance | 231700 |
| 150 | p.Gly150Asp | c.449G>A | Missense | | | 0.30 | Neutral | 3 | 778834 | 3.85191E-06 | Uncertain significance | 483331 |
| 150 | p.Gly150Gly | c.450T>C | Synonymous | | | 0.48 | Neutral | 5 | 1459644 | 3.42549E-06 | Likely benign | 630385 |
| 150 | p.Gly150Ser | c.448G>A | Missense | | | 0.40 | Neutral | 1 | 626658 | 1.59577E-06 | | |
| 151 | p.Pro151His | c.452C>A | Missense | | | 0.54 | Neutral | 1 | 1459568 | 6.85E-07 | | |
| 151 | p.Pro151Leu | c.452C>T | Missense | | | 0.08 | Neutral | 4 | 1611804 | 2.48169E-06 | Uncertain significance | 406724 |
| 154 | p.Ile154Asn | c.461T>A | Missense | | | 0.06 | Neutral | 6 | 628748 | 9.54277E-06 | Uncertain significance | 491579 |
| 156 | p.Asp156Ala | c.467A>C | Missense | | | 0.00 | Deleterious | 1 | 833022 | 1.20045E-06 | | |
| 156 | p.Asp156Asn | c.466G>A | Missense | | | 0.11 | Neutral | 1 | 1461724 | 6.84E-07 | Uncertain significance | 629192 |
| 156 | p.Asp156Asp | c.468T>C | Synonymous | | | 0.48 | Neutral | 4 | 780878 | 5.12244E-06 | Likely benign | 215633 |
| 156 | p.Asp156Tyr | c.466G>T | Missense | | | 0.01 | Deleterious | 2 | 1461722 | 1.36825E-06 | Uncertain significance | 1171898 |
