## Appendix 1-table 11 for "Functional characterization of all *CDKN2A* missense variants and comparison to in silico models of pathogenicity"

**Appendix 1-table 11.** ***CDKN2A*** **missense VUSs reproted in ClinVar.**

| Residue | Variant | Fucntional characterization | Accession |
|---|---|---|---|
| 1 | p.Met1Arg | Neutral | VCV000660169 |
| 1 | p.Met1Leu | Neutral | VCV001475625 |
| 1 | p.Met1Lys | Neutral | VCV000487018 |
| 1 | p.Met1Thr | Neutral | VCV000483341 |
| 1 | p.Met1Val | Neutral | VCV000820481 |
| 2 | p.Glu2Asp | Neutral | VCV001756617 |
| 2 | p.Glu2Gln | Neutral | VCV000406722 |
| 3 | p.Pro3Ser | Indeterminate | VCV000532281 |
| 4 | p.Ala4Gly | Indeterminate | VCV000628437 |
| 4 | p.Ala4Thr | Indeterminate | VCV001500852 |
| 4 | p.Ala4Val | Neutral | VCV001058585 |
| 5 | p.Ala5Glu | Indeterminate | VCV000489864 |
| 5 | p.Ala5Thr | Neutral | VCV000233536 |
| 5 | p.Ala5Val | Neutral | VCV000580530 |
| 6 | p.Gly6Ala | Neutral | VCV000572416 |
| 6 | p.Gly6Arg | Deleterious | VCV000133879 |
| 6 | p.Gly6Trp | Neutral | VCV001525531 |
| 6 | p.Gly6Val | Neutral | VCV000220777 |
| 7 | p.Ser7Cys | Neutral | VCV000489865 |
| 8 | p.Ser8Arg | Neutral | VCV000968555 |
| 8 | p.Ser8Gly | Neutral | VCV000480814 |
| 8 | p.Ser8Thr | Neutral | VCV000532290 |
| 9 | p.Met9Arg | Neutral | VCV000821690 |
| 9 | p.Met9Ile | Neutral | VCV000656960 |
| 9 | p.Met9Leu | Neutral | VCV000483321 |
| 9 | p.Met9Thr | Neutral | VCV000483319 |
| 9 | p.Met9Val | Neutral | VCV000483345 |
| 10 | p.Glu10Ala | Indeterminate | VCV001798627 |
| 10 | p.Glu10Asp | Neutral | VCV000822896 |
| 10 | p.Glu10Lys | Indeterminate | VCV001797426 |
| 11 | p.Pro11Ala | Neutral | VCV000651162 |
| 11 | p.Pro11Arg | Neutral | VCV001332591 |
| 11 | p.Pro11Thr | Indeterminate | VCV001405946 |
| 12 | p.Ser12Ala | Neutral | VCV001507985 |
| 12 | p.Ser12Leu | Indeterminate | VCV000236988 |
| 12 | p.Ser12Trp | Neutral | VCV001022671 |
| 13 | p.Ala13Asp | Neutral | VCV000483348 |
| 13 | p.Ala13Ser | Neutral | VCV000569262 |
| 13 | p.Ala13Val | Neutral | VCV000187508 |
| 14 | p.Asp14Asn | Neutral | VCV000628442 |
| 14 | p.Asp14Glu | Neutral | VCV001466187 |
| 14 | p.Asp14Gly | Neutral | VCV001059920 |
| 15 | p.Trp15Cys | Neutral | VCV000182422 |
| 16 | p.Leu16Gln | Deleterious | VCV000463509 |
| 17 | p.Ala17Asp | Indeterminate | VCV000923872 |

| | | | |
|---|---|---|---|
| 17 | p.Ala17Gly | Neutral | VCV001803801 |
| 17 | p.Ala17Val | Neutral | VCV000825441 |
| 18 | .Thr18Arg | Indeterminate | VCV000630845 |
| 18 | p.Thr18Lys | Indeterminate | VCV000491580 |
| 18 | p.Thr18Met | Indeterminate | VCV000485513 |
| 18 | p.Thr18Pro | Deleterious | VCV000232702 |
| 19 | p.Ala19Asp | Neutral | VCV002775057 |
| 19 | p.Ala19Ser | Neutral | VCV001041797 |
| 20 | p.Ala20Glu | Deleterious | VCV001750950 |
| 20 | p.Ala20Gly | Deleterious | VCV000220344 |
| 20 | p.Ala20Ser | Indeterminate | VCV000664488 |
| 20 | p.Ala20Thr | Indeterminate | VCV001062691 |
| 20 | p.Ala20Val | Indeterminate | VCV001001191 |
| 21 | p.Ala21Asp | Deleterious | VCV001392805 |
| 21 | p.Ala21Ser | Neutral | VCV000928884 |
| 21 | p.Ala21Thr | Indeterminate | VCV000418120 |
| 21 | p.Ala21Val | Neutral | VCV000631322 |
| 22 | p.Arg22Gln | Neutral | VCV000491581 |
| 22 | p.Arg22Gly | Deleterious | VCV000863230 |
| 22 | p.Arg22Leu | Neutral | VCV000463510 |
| 22 | p.Arg22Trp | Indeterminate | VCV000833936 |
| 24 | p.Arg24Gly | Neutral | VCV001481554 |
| 24 | p.Arg24Leu | Neutral | VCV000532272 |
| 25 | p.Val25Ile | Neutral | VCV001376284 |
| 25 | p.Val25Leu | Neutral | VCV001400825 |
| 26 | p.Glu26Ala | Neutral | VCV002748212 |
| 26 | p.Glu26Gln | Neutral | VCV000919572 |
| 27 | p.Glu27Ala | Neutral | VCV000628014 |
| 27 | p.Glu27Asp | Indeterminate | VCV000949317 |
| 27 | p.Glu27Gly | Indeterminate | VCV000483333 |
| 27 | p.Glu27Lys | Neutral | VCV001761576 |
| 28 | p.Val28Leu | Neutral | VCV000231005 |
| 28 | p.Val28Met | Neutral | VCV000827567 |
| 29 | p.Arg29Gln | Neutral | VCV002127104 |
| 29 | p.Arg29Leu | Neutral | VCV002023352 |
| 29 | p.Arg29Pro | Deleterious | VCV001041690 |
| 29 | p.Arg29Trp | Neutral | VCV000483334 |
| 30 | p.Ala30Gly | Deleterious | VCV002775056 |
| 30 | p.Ala30Val | Indeterminate | VCV000246070 |
| 32 | p.Leu32Val | Neutral | VCV000532278 |
| 33 | p.Glu33Gly | Neutral | VCV002005989 |
| 33 | p.Glu33Lys | Neutral | VCV002809872 |
| 33 | p.Glu33Val | Neutral | VCV002863240 |
| 34 | p.Ala34Gly | Neutral | VCV002449061 |
| 34 | p.Ala34Thr | Neutral | VCV000489862 |
| 34 | p.Ala34Val | Indeterminate | VCV000483329 |
| 35 | p.Gly35Arg | Neutral | VCV000483347 |
| 35 | p.Gly35Glu | Indeterminate | VCV000188292 |

| | | | |
|---|---|---|---|
| 35 | p.Gly35Val | Deleterious | VCV000406709 |
| 36 | p.Ala36Ser | Neutral | VCV000532277 |
| 36 | p.Ala36Thr | Indeterminate | VCV000489863 |
| 36 | p.Ala36Val | Neutral | VCV000483324 |
| 37 | p.Leu37Arg | Neutral | VCV001331763 |
| 38 | p.Pro38Ser | Indeterminate | VCV000822217 |
| 39 | p.Asn39Ile | Indeterminate | VCV002680551 |
| 39 | p.Asn39Lys | Indeterminate | VCV000629162 |
| 39 | p.Asn39Ser | Neutral | VCV002136748 |
| 40 | p.Ala40Pro | Indeterminate | VCV001063406 |
| 40 | p.Ala40Ser | Neutral | VCV000491563 |
| 40 | p.Ala40Val | Indeterminate | VCV003230252 |
| 41 | p.Pro41Ala | Neutral | VCV000406713 |
| 41 | p.Pro41Arg | Indeterminate | VCV001755139 |
| 41 | p.Pro41Leu | Neutral | VCV001755177 |
| 41 | p.Pro41Ser | Neutral | VCV002804807 |
| 42 | p.Asn42Asp | Indeterminate | VCV000656316 |
| 42 | p.Asn42Lys | Deleterious | VCV000664863 |
| 42 | p.Asn42Ser | Neutral | VCV000406711 |
| 43 | p.Ser43Asn | Neutral | VCV000954477 |
| 43 | p.Ser43Gly | Neutral | VCV000630389 |
| 43 | p.Ser43Thr | Neutral | VCV000569634 |
| 44 | p.Tyr44Cys | Neutral | VCV000630012 |
| 45 | p.Gly45Ala | Neutral | VCV001054175 |
| 45 | p.Gly45Arg | Neutral | VCV000463483 |
| 45 | p.Gly45Cys | Neutral | VCV001052395 |
| 45 | p.Gly45Ser | Indeterminate | VCV000463482 |
| 46 | p.Arg46Gln | Indeterminate | VCV000631208 |
| 46 | p.Arg46Pro | Deleterious | VCV000532274 |
| 46 | p.Arg46Trp | Neutral | VCV000573855 |
| 47 | p.Arg47Lys | Neutral | VCV002976242 |
| 48 | p.Pro48Gln | Deleterious | VCV001042953 |
| 49 | p.Ile49Met | Neutral | VCV000483330 |
| 50 | p.Gln50His | Deleterious | VCV001045852 |
| 50 | p.Gln50Leu | Indeterminate | VCV000133878 |
| 50 | p.Gln50Lys | Deleterious | VCV000584732 |
| 51 | p.Val51Ile | Neutral | VCV000463485 |
| 51 | p.Val51Phe | Deleterious | VCV002735266 |
| 52 | p.Met52Ile | Indeterminate | VCV002132254 |
| 52 | p.Met52Thr | Neutral | VCV000418881 |
| 53 | p.Met53Thr | Indeterminate | VCV001480737 |
| 54 | p.Met54Arg | Indeterminate | VCV000819690 |
| 54 | p.Met54Ile | Neutral | VCV000819708 |
| 54 | p.Met54Leu | Neutral | VCV001483043 |
| 54 | p.Met54Thr | Neutral | VCV000819689 |
| 55 | p.Gly55Ala | Indeterminate | VCV002563916 |
| 55 | p.Gly55Arg | Deleterious | VCV000579094 |
| 55 | p.Gly55Val | Deleterious | VCV002136747 |

| | | | |
|---|---|---|---|
| 56 | p.Ser56Arg | Neutral | VCV000463488 |
| 56 | p.Ser56Asn | Neutral | VCV000463487 |
| 56 | p.Ser56Gly | Neutral | VCV002563910 |
| 57 | p.Ala57Asp | Neutral | VCV000483325 |
| 57 | p.Ala57Gly | Neutral | VCV000187272 |
| 58 | p.Arg58Gln | Neutral | VCV002083590 |
| 58 | p.Arg58Gly | Neutral | VCV000532285 |
| 58 | p.Arg58Leu | Neutral | VCV002416022 |
| 59 | p.Val59Ala | Neutral | VCV001434858 |
| 59 | p.Val59Glu | Deleterious | VCV000491569 |
| 59 | p.Val59Leu | Neutral | VCV002122385 |
| 59 | p.Val59Met | Neutral | VCV001007891 |
| 60 | p.Ala60Pro | Indeterminate | VCV000532284 |
| 60 | p.Ala60Thr | Neutral | VCV000406705 |
| 60 | p.Ala60Val | Neutral | VCV000570547 |
| 61 | p.Glu61Asp | Neutral | VCV000236982 |
| 62 | p.Leu62Val | Neutral | VCV000969307 |
| 64 | p.Leu64Val | Neutral | VCV002775055 |
| 65 | p.Leu65Phe | Neutral | VCV000966551 |
| 66 | p.His66Asp | Neutral | VCV000969054 |
| 66 | p.His66Gln | Neutral | VCV000573530 |
| 66 | p.His66Pro | Deleterious | VCV000419802 |
| 66 | p.His66Tyr | Neutral | VCV001348846 |
| 67 | p.Gly67Asp | Neutral | VCV000216273 |
| 67 | p.Gly67Cys | Neutral | VCV002563913 |
| 67 | p.Gly67Ser | Neutral | VCV000925139 |
| 68 | p.Ala68Gly | Neutral | VCV000628976 |
| 68 | p.Ala68Ser | Neutral | VCV001355040 |
| 68 | p.Ala68Thr | Neutral | VCV001784739 |
| 69 | p.Glu69Ala | Neutral | VCV002891892 |
| 69 | p.Glu69Asp | Neutral | VCV001052397 |
| 69 | p.Glu69Lys | Indeterminate | VCV000966903 |
| 70 | p.Pro70Arg | Neutral | VCV000185936 |
| 70 | p.Pro70Leu | Neutral | VCV000463490 |
| 70 | p.Pro70Ser | Indeterminate | VCV001486509 |
| 70 | p.Pro70Thr | Neutral | VCV001374577 |
| 71 | p.Asn71Thr | Deleterious | VCV001786361 |
| 71 | p.Asn71Tyr | Deleterious | VCV000485514 |
| 72 | p.Cys72Gly | Neutral | VCV001436072 |
| 72 | p.Cys72Phe | Indeterminate | VCV001320714 |
| 72 | p.Cys72Ser | Indeterminate | VCV000532279 |
| 73 | p.Ala73Thr | Neutral | VCV001986070 |
| 74 | p.Asp74Ala | Deleterious | VCV000802463 |
| 74 | p.Asp74Asn | Deleterious | VCV000483340 |
| 74 | p.Asp74Glu | Neutral | VCV000660824 |
| 74 | p.Asp74His | Neutral | VCV000802464 |
| 74 | p.Asp74Val | Deleterious | VCV000820911 |
| 75 | p.Pro75Ala | Neutral | VCV001788244 |

| | | | |
|---|---|---|---|
| 75 | p.Pro75Leu | Neutral | VCV000483326 |
| 75 | p.Pro75Ser | Neutral | VCV000644311 |
| 76 | p.Ala76Ser | Indeterminate | VCV002059365 |
| 76 | p.Ala76Thr | Neutral | VCV000489866 |
| 77 | p.Thr77Ala | Indeterminate | VCV000567670 |
| 77 | p.Thr77Ser | Indeterminate | VCV000582983 |
| 78 | p.Leu78Arg | Indeterminate | VCV001493577 |
| 78 | p.Leu78Phe | Neutral | VCV000630762 |
| 78 | p.Leu78Pro | Indeterminate | VCV000483349 |
| 78 | p.Leu78Val | Neutral | VCV002814776 |
| 79 | p.Thr79Asn | Neutral | VCV000491570 |
| 79 | p.Thr79Ile | Neutral | VCV000630548 |
| 79 | p.Thr79Pro | Deleterious | VCV000463491 |
| 79 | p.Thr79Ser | Neutral | VCV000967063 |
| 80 | p.Arg80Gln | Indeterminate | VCV000376385 |
| 80 | p.Arg80Gly | Indeterminate | VCV000483353 |
| 80 | p.Arg80Pro | Deleterious | VCV002688757 |
| 81 | p.Pro81Ser | Deleterious | VCV000664812 |
| 81 | p.Pro81Thr | Indeterminate | VCV000629992 |
| 82 | p.Val82Glu | Indeterminate | VCV000833905 |
| 82 | p.Val82Leu | Neutral | VCV000628722 |
| 82 | p.Val82Met | Neutral | VCV001063226 |
| 84 | p.Asp84Glu | Indeterminate | VCV002868822 |
| 84 | p.Asp84His | Deleterious | VCV002200354 |
| 85 | p.Ala85Phe | Indeterminate | VCV000423274 |
| 85 | p.Ala85Pro | Indeterminate | VCV001376090 |
| 85 | p.Ala85Ser | Neutral | VCV000532289 |
| 85 | p.Ala85Thr | Indeterminate | VCV000236983 |
| 85 | p.Ala85Val | Indeterminate | VCV001438157 |
| 86 | p.Ala86Gly | Neutral | VCV001793349 |
| 86 | p.Ala86Ser | Neutral | VCV001793230 |
| 87 | p.Arg87Gln | Indeterminate | VCV003230255 |
| 88 | p.Glu88Gln | Indeterminate | VCV000619828 |
| 88 | p.Glu88Lys | Indeterminate | VCV000821566 |
| 88 | p.Glu88Val | Neutral | VCV001794163 |
| 89 | p.Gly89Ala | Indeterminate | VCV000565793 |
| 89 | p.Gly89Cys | Deleterious | VCV002087960 |
| 89 | p.Gly89Ser | Deleterious | VCV000009408 |
| 89 | p.Gly89Val | Deleterious | VCV001802421 |
| 90 | p.Phe90Cys | Neutral | VCV000532291 |
| 90 | p.Phe90Leu | Neutral | VCV002095801 |
| 91 | p.Leu91Gln | Neutral | VCV000619829 |
| 91 | p.Leu91Met | Indeterminate | VCV002762000 |
| 92 | p.Asp92Glu | Neutral | VCV001795780 |
| 92 | p.Asp92His | Neutral | VCV002030149 |
| 93 | p.Thr93Ala | Indeterminate | VCV001795894 |
| 93 | p.Thr93Arg | Indeterminate | VCV000232366 |
| 93 | p.Thr93Lys | Deleterious | VCV001796014 |

| | | | |
|---|---|---|---|
| 93 | p.Thr93Met | Indeterminate | VCV001796019 |
| 93 | p.Thr93Pro | Deleterious | VCV001795891 |
| 94 | p.Leu94Val | Neutral | VCV001796309 |
| 95 | p.Val95Glu | Neutral | VCV001796819 |
| 95 | p.Val95Gly | Neutral | VCV002001636 |
| 96 | p.Val96Leu | Indeterminate | VCV002106366 |
| 96 | p.Val96Met | Neutral | VCV000821875 |
| 97 | p.Leu97Pro | Deleterious | VCV000962431 |
| 98 | p.His98Arg | Neutral | VCV002151380 |
| 98 | p.His98Gln | Neutral | VCV000822346 |
| 98 | p.His98Tyr | Neutral | VCV000419560 |
| 99 | p.Arg99Leu | Indeterminate | VCV002867606 |
| 99 | p.Arg99Trp | Indeterminate | VCV000483350 |
| 100 | p.Ala100Thr | Neutral | VCV000629701 |
| 100 | p.Ala100Val | Neutral | VCV000924911 |
| 101 | p.Gly101Ala | Indeterminate | VCV001129005 |
| 101 | p.Gly101Arg | Indeterminate | VCV000463494 |
| 101 | p.Gly101Glu | Indeterminate | VCV000419180 |
| 102 | p.Ala102Leu | Deleterious | VCV001799155 |
| 102 | p.Ala102Thr | Deleterious | VCV000463495 |
| 102 | p.Ala102Val | Indeterminate | VCV000234071 |
| 103 | p.Arg103Gln | Neutral | VCV001141816 |
| 103 | p.Arg103Pro | Neutral | VCV000406703 |
| 103 | p.Arg103Trp | Neutral | VCV000423630 |
| 104 | p.Leu104Pro | Neutral | VCV001014833 |
| 105 | p.Asp105Ala | Indeterminate | VCV001728158 |
| 105 | p.Asp105Asn | Neutral | VCV000491573 |
| 105 | p.Asp105Glu | Indeterminate | VCV001728273 |
| 105 | p.Asp105Tyr | Neutral | VCV000584518 |
| 106 | p.Val106Ala | Neutral | VCV000463497 |
| 106 | p.Val106Glu | Indeterminate | VCV000823098 |
| 106 | p.Val106Leu | Indeterminate | VCV001728421 |
| 106 | p.Val106Met | Neutral | VCV001392243 |
| 107 | p.Arg107Leu | Neutral | VCV001005179 |
| 107 | p.Arg107Pro | Neutral | VCV000630455 |
| 107 | p.Arg107Ser | Neutral | VCV000463498 |
| 108 | p.Asp108Glu | Indeterminate | VCV001498736 |
| 108 | p.Asp108Val | Deleterious | VCV002767690 |
| 109 | p.Ala109Asp | Neutral | VCV000420995 |
| 109 | p.Ala109Pro | Deleterious | VCV000127525 |
| 109 | p.Ala109Ser | Neutral | VCV000628079 |
| 110 | p.Trp110Arg | Neutral | VCV002109926 |
| 111 | p.Gly111Arg | Neutral | VCV000230231 |
| 111 | p.Gly111Asp | Neutral | VCV001416042 |
| 111 | p.Gly111Cys | Neutral | VCV002625891 |
| 111 | p.Gly111Ser | Neutral | VCV000463499 |
| 111 | p.Gly111Val | Deleterious | VCV001487529 |
| 112 | p.Arg112Cys | Indeterminate | VCV002775054 |

| | | | |
|---|---|---|---|
| 112 | p.Arg112His | Neutral | VCV000142887 |
| 112 | p.Arg112Leu | Indeterminate | VCV000823661 |
| 112 | p.Arg112Pro | Deleterious | VCV001730600 |
| 112 | p.Arg112Ser | Neutral | VCV001995012 |
| 113 | p.Leu113Pro | Deleterious | VCV000833598 |
| 114 | p.Pro114Ser | Indeterminate | VCV001405931 |
| 114 | p.Pro114Thr | Deleterious | VCV000376382 |
| 115 | p.Val115Glu | Indeterminate | VCV000246437 |
| 115 | p.Val115Gly | Neutral | VCV000236987 |
| 115 | p.Val115Leu | Neutral | VCV002013862 |
| 115 | p.Val115Met | Neutral | VCV001469323 |
| 116 | p.Asp116Glu | Neutral | VCV000489867 |
| 116 | p.Asp116Tyr | Neutral | VCV002563909 |
| 117 | p.Leu117Pro | Indeterminate | VCV000833600 |
| 118 | p.Ala118Thr | Neutral | VCV000491574 |
| 119 | p.Glu119Lys | Neutral | VCV000570565 |
| 120 | p.Glu120Asp | Neutral | VCV000823988 |
| 121 | p.Leu121Arg | Indeterminate | VCV002723614 |
| 121 | p.Leu121Pro | Neutral | VCV000566923 |
| 122 | p.Gly122Arg | Neutral | VCV000009422 |
| 122 | p.Gly122Ser | Neutral | VCV001468600 |
| 123 | p.His123Tyr | Neutral | VCV002836865 |
| 124 | p.Arg124Gly | Indeterminate | VCV000582036 |
| 124 | p.Arg124Pro | Neutral | VCV000824122 |
| 124 | p.Arg124Ser | Neutral | VCV000824106 |
| 125 | p.Asp125Asn | Neutral | VCV000858988 |
| 125 | p.Asp125Glu | Neutral | VCV002497223 |
| 125 | p.Asp125Gly | Neutral | VCV000834371 |
| 126 | p.Val126Leu | Neutral | VCV000489868 |
| 127 | p.Ala127Val | Neutral | VCV002030143 |
| 128 | p.Arg128Gln | Neutral | VCV000581636 |
| 128 | p.Arg128Gly | Indeterminate | VCV001735329 |
| 128 | p.Arg128Pro | Deleterious | VCV000483344 |
| 128 | p.Arg128Trp | Neutral | VCV000575361 |
| 129 | p.Tyr129Asp | Indeterminate | VCV000480813 |
| 129 | p.Tyr129His | Neutral | VCV001735601 |
| 130 | p.Leu130Gln | Indeterminate | VCV002744302 |
| 130 | p.Leu130Met | Neutral | VCV000406704 |
| 131 | p.Arg131Cys | Deleterious | VCV001350948 |
| 131 | p.Arg131His | Neutral | VCV000620615 |
| 131 | p.Arg131Leu | Neutral | VCV002625894 |
| 131 | p.Arg131Ser | Neutral | VCV001366069 |
| 132 | p.Ala132Thr | Indeterminate | VCV001362170 |
| 132 | p.Ala132Val | Neutral | VCV000630452 |
| 133 | p.Ala133Ser | Neutral | VCV000933583 |
| 133 | p.Ala133Thr | Neutral | VCV000630684 |
| 134 | p.Ala134Gly | Neutral | VCV001737089 |
| 134 | p.Ala134Pro | Neutral | VCV000495536 |

| | | | |
|---|---|---|---|
| 134 | p.Ala134Ser | Neutral | VCV000532288 |
| 134 | p.Ala134Val | Neutral | VCV000650117 |
| 135 | p.Gly135Ala | Neutral | VCV001061502 |
| 135 | p.Gly135Arg | Neutral | VCV000928883 |
| 135 | p.Gly135Glu | Indeterminate | VCV000463504 |
| 135 | p.Gly135Trp | Neutral | VCV000937971 |
| 135 | p.Gly135Val | Neutral | VCV000843913 |
| 136 | p.Gly136Ala | Neutral | VCV000483335 |
| 136 | p.Gly136Asp | Indeterminate | VCV000925925 |
| 136 | p.Gly136Cys | Indeterminate | VCV002563912 |
| 136 | p.Gly136Ser | Neutral | VCV000483342 |
| 137 | p.Thr137Ile | Neutral | VCV000824601 |
| 138 | p.Arg138Gly | Neutral | VCV000233990 |
| 138 | p.Arg138Lys | Neutral | VCV001353011 |
| 138 | p.Arg138Thr | Neutral | VCV001738148 |
| 139 | p.Gly139Asp | Indeterminate | VCV000491576 |
| 139 | p.Gly139Cys | Neutral | VCV001316508 |
| 139 | p.Gly139Ser | Neutral | VCV000141419 |
| 139 | p.Gly139Val | Neutral | VCV002172999 |
| 140 | p.Ser140Arg | Neutral | VCV000824656 |
| 140 | p.Ser140Asn | Neutral | VCV000485512 |
| 141 | p.Asn141Asp | Neutral | VCV000631362 |
| 141 | p.Asn141Lys | Neutral | VCV000953252 |
| 142 | p.His142Arg | Neutral | VCV000184564 |
| 142 | p.His142Asp | Neutral | VCV000840158 |
| 142 | p.His142Gln | Neutral | VCV000246046 |
| 142 | p.His142Tyr | Neutral | VCV000824721 |
| 143 | p.Ala143Gly | Neutral | VCV000630386 |
| 143 | p.Ala143Val | Neutral | VCV001739412 |
| 144 | p.Arg144Gly | Neutral | VCV000532270 |
| 144 | p.Arg144His | Neutral | VCV000406723 |
| 144 | p.Arg144Leu | Neutral | VCV000463505 |
| 144 | p.Arg144Pro | Neutral | VCV000576389 |
| 145 | p.Ile145Arg | Neutral | VCV002563906 |
| 145 | p.Ile145Lys | Neutral | VCV000658624 |
| 145 | p.Ile145Val | Neutral | VCV000645545 |
| 146 | p.Asp146Asn | Neutral | VCV000578446 |
| 146 | p.Asp146Glu | Neutral | VCV002012076 |
| 146 | p.Asp146Gly | Neutral | VCV002680552 |
| 146 | p.Asp146His | Neutral | VCV000824834 |
| 146 | p.Asp146Tyr | Neutral | VCV002820920 |
| 146 | p.Asp146Val | Neutral | VCV000928297 |
| 147 | p.Ala147Val | Neutral | VCV000406716 |
| 148 | p.Ala148Pro | Neutral | VCV000483343 |
| 148 | p.Ala148Ser | Neutral | VCV000406706 |
| 148 | p.Ala148Val | Indeterminate | VCV000231700 |
| 150 | p.Gly150Arg | Neutral | VCV002775052 |
| 150 | p.Gly150Asp | Neutral | VCV000483331 |

| | | | |
|---|---|---|---|
| 150 | p.Gly150Cys | Neutral | VCV002094524 |
| 151 | p.Pro151Leu | Neutral | VCV000406724 |
| 151 | p.Pro151Ser | Indeterminate | VCV001741192 |
| 152 | p.Ser152Leu | Neutral | VCV000630384 |
| 153 | p.Asp153Asn | Neutral | VCV000532292 |
| 153 | p.Asp153Gly | Neutral | VCV000939445 |
| 153 | p.Asp153His | Neutral | VCV001741633 |
| 153 | p.Asp153Val | Neutral | VCV000406721 |
| 154 | p.Ile154Asn | Neutral | VCV000491579 |
| 155 | p.Pro155Ala | Neutral | VCV000406720 |
| 155 | p.Pro155Arg | Neutral | VCV000859084 |
| 156 | p.Asp156Asn | Neutral | VCV000629192 |
| 156 | p.Asp156Tyr | Indeterminate | VCV001171898 |
| 156 | p.Asp156Val | Neutral | VCV001742377 |
