## Appendix 1-table 12 for "Functional characterization of all *CDKN2A* missense variants and comparison to in silico models of pathogenicity"

**Appendix 1-table 12. Codon optimized *CDKN2A* sequence.**

| Sequence |
| --- |
| ATGGAACCCGCCGCTGGCTCATCAATGGAACCCTCTGCAGATTGGCTCGCT |
| ACCGCAGCCGCGCGAGGACGAGTGGAAGAAGTCCGAGCGCTCCTTGAAG |
| CCGGTGCATTGCCAAATGCGCCTAACTCCTATGGACGCCGCCCAATACAAG |
| TTATGATGATGGGATCTGCAAGGGTAGCTGAACTTCTCCTCTTGCATGGAG |
| CAGAACCTAATTGTGCTGATCCAGCGACCCTGACTAGACCTGTACATGATG |
| CCGCGCGTGAAGGGTTTCTCGATACCCTTGTCGTCCTTCATCGAGCTGGTGC |
| CCGCCTCGATGTCCGGGACGCATGGGGACGGCTCCCAGTCGATCTCGCAG |
| AAGAACTTGGGCACAGGGACGTAGCCAGATATTTGCGTGCAGCCGCCGGT |
| GGTACGAGGGGATCAAATCACGCTAGAATCGACGCTGCCGAGGGCCCAA |
| [illegible] |
