## Appendix 1-table 13 for "Functional characterization of all *CDKN2A* missense variants and comparison to in silico models of pathogenicity"

**Appendix 1-table 13. Sequences of primers used in study.**

| Oligos | Sequence |
|---|---|
| **Primers used to generate variant *CDKN2A* expression plasmids** | |
| p.Ser7Gly_FWD | CGCCGCTGGCGGCTCAATGGAAC |
| p.Ser7Gly_REV | GGTTCCATTTAGTCTGGTATG |
| p.Glu26Cys_FWD | AGGACGAGTGTGCGAAGTCCGAGCGCTCCTTGAAGCCGG |
| p.Glu26Cys_REV | CGCGCGGCTGCGGTAGCG |
| p.Glu26Pro_FWD | AGGACGAGTGCCCGAAGTCCGAGCGCTCCTTGAAGCCG |
| p.Glu26Pro_REV | CGCGCGGCTGCGGTAGCG |
| p.Glu27Ala_FWD | ACGAGTGGAAGCCGTCCGAGCGCTCCTTGAAGC |
| p.Glu27Ala_REV | CCTCGCGCGGCTGCGGTA |
| p.Glu27Thr_FWD | ACGAGTGGAAACCGTCCGAGCGCTCCTTGAAGCCG |
| p.Glu27Thr_REV | CCTCGCGCGGCTGCGGTA |
| p.Glu27His_FWD | ACGAGTGGAACACGTCCGAGCGCTCCTTGAAGC |
| p.Glu27His_REV | CCTCGCGCGGCTGCGGTA |
| p.Glu27Pro_FWD | ACGAGTGGAACCCGTCCGAGCGCTCCTTGAAGCCGGTG |
| p.Glu27Pro_REV | CCTCGCGCGGCTGCGGTA |
| p.Glu27Ile_FWD | ACGAGTGGAAATCGTCCGAGCGCTCCTTGAAGCCG |
| p.Glu27Ile_REV | CCTCGCGCGGCTGCGGTA |
| p.Glu69Ala_FWD | GCATGGAGCAGCCCCTAATTGTG |
| p.Glu69Ala_REV | AAGAGGAGAAGTTCAGCTAC |
| p.Val95His_FWD | CGATACCCTTCACGTCCTTCATCGAG |
| p.Val95His_REV | AGAAACCCTTCACGCGCG |
| p.Ala100Gly_FWD | CCGCCTCGATGTCCGGGACGCATG |
| p.Ala100Gly_REV | GCACCGCCTCGATGAAGGACGAC |
| p.Arg103Lys_FWD | AGCTGGTGCCAAGCTCGATGTCC |
| p.Arg103Lys_REV | CGATGAAGGACGACAAGG |
| p.Arg103Leu_FWD | AGCTGGTGCCCTGCTCGATGTCC |
| p.Arg103Leu_REV | CGATGAAGGACGACAAGG |
| p.Arg103Gln_FWD | AGCTGGTGCCCAGCTCGATGTCC |
| p.Arg103Gln_REV | CGATGAAGGACGACAAGG |
| p.Ala109Asp_FWD | TGTCCGGGACGACTGGGGACGGCTC |
| p.Ala109Asp_REV | TCGAGGCGGGCACCAGCT |
| p.Trp110Pro_FWD | CCGGGACGCACCCGGACGGCTCCCAG |
| p.Trp110Pro_REV | ACATCGAGGCGGGCACCA |
| p.Val115Cys_FWD | ACGGCTCCCATGCGATCTCGCAGAAGAAC |
| p.Val115Cys_REV | CCCCATGCGTCCCGGACA |
| p.Gly136Cys_FWD | AGCCGCCGGTTGCACGAGGGGAT |
| p.Gly136Cys_REV | GCACGCAAATATCTGGCTACG |
| p.Pro151Gly_FWD | TGCCGAGGGCGGCAGCGACATAC |
| p.Pro151Gly_REV | GCGTCGATTCTAGCGTGA |
| p.Ile154Gly_FWD | CCCAAGCGACGGCCCAGACTAAGAATTCTCGACC |
| p.Ile154Gly_REV | CCCTCGGCAGCGTCGATT |
| p.L32P_FWD | CCGAGCGCTCCCCGAAGCCGGTG |
| p.L32P_REV | ACTTCTTCCACTCGTCCTCG |
| p.G101W_FWD | TCATCGAGCTTGGGCCCGCCTCG |
| p.G101W_REV | AGGACGACAAGGGTATCGAGAAAC |
| p.V126D_FWD | GCACAGGGACGACGCCAGATATTTG |
| p.V126D_REV | CCAAGTTCTTCTGCGAGATC |
| p.L32L_FWD | CCGAGCGCTCCTGGAAGCCGGTG |
| p.L32L_REV | ACTTCTTCCACTCGTCCTCGCGC |
| p.G101G_FWD | TCATCGAGCTGGCGCCCGCCTCG |
| p.G101G_REV | AGGACGACAAGGGTATCGAGAAACC |
| p.V126V_FWD | GCACAGGGACGTGGCCAGATATT |
| p.V126V_REV | CCAAGTTCTTCTGCGAGATCG |
| **Primers used to generate CellTag plasmids** | |
| Barcode 1_FWD | CTGCGGCAGCGGCGCCACCAAC |
| Barcode 1_REV | TACCACTTATCGTCATCGTCTTTGTAATCCATGGTGGC |
| Barcode 2_FWD | CGATAAGGATCATAGCGGCAGCGGCG |
| Barcode 2_REV | TCATCGTCTTTGTAATCC |
| Barcode 3_FWD | GAATGGCAGCGGCGCCACCAAC |
| Barcode 3_REV | TGAATCTTATCGTCATCGTCTTTGTAATCCATGGTGGC |
| Barcode 4_FWD | GTACGGCAGCGGCGCCACCAAC |
| Barcode 4_REV | TGCTCCTTATCGTCATCGTCTTTGTAATCCATGGTGGC |
| Barcode 5_FWD | GTACGGCAGCGGCGCCACCAAC |
| Barcode 5_REV | ATATCCTTATCGTCATCGTCTTTGTAATCCATGGTGGC |
| Barcode 6_FWD | GTTCGGCAGCGGCGCCACCAAC |
| Barcode 6_REV | TCAACCTTATCGTCATCGTCTTTGTAATCCATGGTGGC |
| Barcode 7_FWD | ACATGGCAGCGGCGCCACCAAC |
| Barcode 7_REV | TGATACTTATCGTCATCGTCTTTGTAATCCATGGTGGC |
| Barcode 8_FWD | AGCTGGCAGCGGCGCCACCAAC |
| Barcode 8_REV | TCGCACTTATCGTCATCGTCTTTGTAATCCATGGTGGC |
| Barcode 9_FWD | GACGATAAGGATTGTACCGGCAGCGG |
| Barcode 9_REV | ATCGTCTTTGTAATCCATGGTG |
| Barcode 10_FWD | GGTCGGCAGCGGCGCCACCAAC |
| Barcode 10_REV | CCTGGCTTATCGTCATCGTCTTTGTAATCCATGGTGGC |
| Barcode 11_FWD | ACGATAAGATCACGATCGGCAGCGGC |
| Barcode 11_REV | CATCGTCTTTGTAATCCATG |
| Barcode 12_FWD | CGATAAGGTGAGGAGCGGCAGCGGCG |
| Barcode 12_REV | TCATCGTCTTTGTAATCCATGG |
| Barcode 13_FWD | TATAGGCAGCGGCGCCACCAAC |
| Barcode 13_REV | CAACCCTTATCGTCATCGTCTTTGTAATCCATGGTGGC |
| Barcode 14_FWD | GATAAGGTTCGTATCGGCAGCGG |
| Barcode 14_REV | GTCATCGTCTTTGTAATCC |
| Barcode 15_FWD | GTACGGCAGCGGCGCCACCAAC |
| Barcode 15_REV | AGAACCTTATCGTCATCGTCTTTGTAATCCATGGTGGC |
| Barcode 16_FWD | AGCCGGCAGCGGCGCCACCAAC |
| Barcode 16_REV | ACCGCCTTATCGTCATCGTCTTTGTAATCCATGGTGGC |
| Barcode 17_FWD | TGATGGCAGCGGCGCCACCAAC |
| Barcode 17_REV | ACGTACTTATCGTCATCGTCTTTGTAATCCATGGTGGC |
| Barcode 18_FWD | CTCAGGCAGCGGCGCCACCAAC |
| Barcode 18_REV | CACTGCTTATCGTCATCGTCTTTGTAATCCATGGTGGC |
| Barcode 19_FWD | CTGTGGCAGCGGCGCCACCAAC |
| Barcode 19_REV | TTCTGCTTATCGTCATCGTCTTTGTAATCCATGGTGGC |
| Barcode 20_FWD | CAATGGCAGCGGCGCCACCAAC |
| Barcode 20_REV | TATCCCTTATCGTCATCGTCTTTGTAATCCATGGTGGC |
| **1st stage primers used to generate sequence libraries** | |
| CellTag_FWD | GTAAAACGACGGCCAGGTGTCGTGACGCGGGATCCG |
| CellTag_REV | CAGGAAACAGCTATGACTGTAGCCGTTGCTCTTTCAA |
| p.AA1-53_FWD | GTAAAACGACGGCCAGCGCTACCGGGTACAAGTCC |
| p.AA1-53_REV | CAGGAAACAGCTATGACTCAGCTACCCTTGCAGATCC |
| p.AA54-110_FWD | GTAAAACGACGGCCAGCCGCCCAATACAAGTTATG |
| p.AA54-110_REV | CAGGAAACAGCTATGACCCCAAGTTCTTCTGCGAGA |
| p.AA111-156_FWD | GTAAAACGACGGCCAGCGATACCCTTGTCGTCCTTC |
| p.AA111-156_REV | CAGGAAACAGCTATGACACTGCCATTTGTCTCGAGGT |
| **2nd stage primers used to generate sequence libraries** | |
| FWD1 | AATGATACGGCGACCACCGAGATCTACACTCTTTCCCTACACGACGCTCTTCCGATCTATGTAAAACGACGGCCAG |
| FWD2 | AATGATACGGCGACCACCGAGATCTACACTCTTTCCCTACACGACGCTCTTCCGATCTGATGTAAAACGACGGCCAG |
| FWD3 | AATGATACGGCGACCACCGAGATCTACACTCTTTCCCTACACGACGCTCTTCCGATCTCGATGTAAAACGACGGCCAG |
| FWD4 | AATGATACGGCGACCACCGAGATCTACACTCTTTCCCTACACGACGCTCTTCCGATCTTCGATGTAAAACGACGGCCAG |
| FWD5 | AATGATACGGCGACCACCGAGATCTACACTCTTTCCCTACACGACGCTCTTCCGATCTATCGATGTAAAACGACGGCCAG |
| FWD6 | AATGATACGGCGACCACCGAGATCTACACTCTTTCCCTACACGACGCTCTTCCGATCTGATCGATGTAAAACGACGGCCAG |
| FWD7 | AATGATACGGCGACCACCGAGATCTACACTCTTTCCCTACACGACGCTCTTCCGATCTCGATCGATGTAAAACGACGGCCAG |
| FWD8 | AATGATACGGCGACCACCGAGATCTACACTCTTTCCCTACACGACGCTCTTCCGATCTTCGATCGATGTAAAACGACGGCCAG |
| REV1 | CAAGCAGAAGACGGCATACGAGATGCTGGATTGTGACTGGAGTTCAGACGTGTGCTCTTCCGATCTATCAGGAAACAGCTATGAC |
| REV2 | CAAGCAGAAGACGGCATACGAGATTAACTCGGGTGACTGGAGTTCAGACGTGTGCTCTTCCGATCTGATCAGGAAACAGCTATGAC |
| REV3 | CAAGCAGAAGACGGCATACGAGATTAACAGTTGTGACTGGAGTTCAGACGTGTGCTCTTCCGATCTCGATCAGGAAACAGCTATGAC |
| REV4 | CAAGCAGAAGACGGCATACGAGATATACTCAAGTGACTGGAGTTCAGACGTGTGCTCTTCCGATCTTCGATCAGGAAACAGCTATGAC |
| REV5 | CAAGCAGAAGACGGCATACGAGATGCTGAGAAGTGACTGGAGTTCAGACGTGTGCTCTTCCGATCTATCGATCAGGAAACAGCTATGAC |
| REV6 | CAAGCAGAAGACGGCATACGAGATATTGGAGGGTGACTGGAGTTCAGACGTGTGCTCTTCCGATCTGATCGATCAGGAAACAGCTATGAC |
| REV7 | CAAGCAGAAGACGGCATACGAGATTAGTCTAAGTGACTGGAGTTCAGACGTGTGCTCTTCCGATCTCGATCGATCAGGAAACAGCTATGAC |
| REV8 | CAAGCAGAAGACGGCATACGAGATCGGTGACCGTGACTGGAGTTCAGACGTGTGCTCTTCCGATCTTCGATCGATCAGGAAACAGCTATGAC |
| REV9 | CAAGCAGAAGACGGCATACGAGATTACAGAGGGTGACTGGAGTTCAGACGTGTGCTCTTCCGATCTATCAGGAAACAGCTATGAC |
| REV10 | CAAGCAGAAGACGGCATACGAGATATTGTCAAGTGACTGGAGTTCAGACGTGTGCTCTTCCGATCTGATCAGGAAACAGCTATGAC |
| REV11 | CAAGCAGAAGACGGCATACGAGATTATGTCTTGTGACTGGAGTTCAGACGTGTGCTCTTCCGATCTCGATCAGGAAACAGCTATGAC |
| REV12 | CAAGCAGAAGACGGCATACGAGATATTGGATTGTGACTGGAGTTCAGACGTGTGCTCTTCCGATCTTCGATCAGGAAACAGCTATGAC |
| REV13 | CAAGCAGAAGACGGCATACGAGATATACTCGGGTGACTGGAGTTCAGACGTGTGCTCTTCCGATCTATCGATCAGGAAACAGCTATGAC |
| REV14 | CAAGCAGAAGACGGCATACGAGATTATGAGAAGTGACTGGAGTTCAGACGTGTGCTCTTCCGATCTGATCGATCAGGAAACAGCTATGAC |
| REV15 | CAAGCAGAAGACGGCATACGAGATGCACAGTTGTGACTGGAGTTCAGACGTGTGCTCTTCCGATCTCGATCGATCAGGAAACAGCTATGAC |
| REV16 | CAAGCAGAAGACGGCATACGAGATCGTGGATTGTGACTGGAGTTCAGACGTGTGCTCTTCCGATCTTCGATCGATCAGGAAACAGCTATGAC |

FWD - forward primer; REV - reverse primer.
