## Supplementary figures and images for "Functional characterization of all *CDKN2A* missense variants and comparison to in silico models of pathogenicity"

### Figure 1 -figure supplement 1

#### Figure 1-figure supplement 1

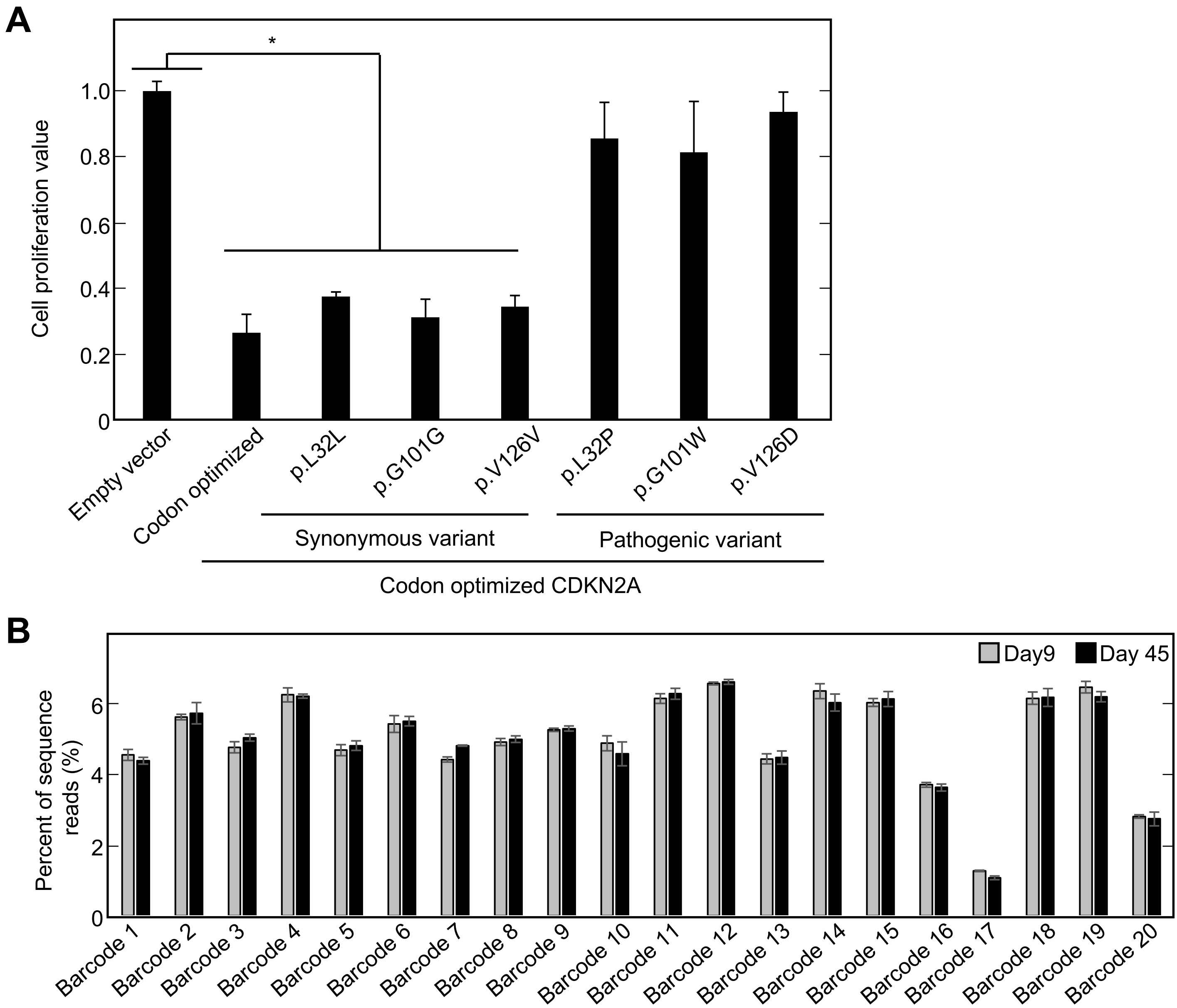

### Figure 1 -figure supplement 2

#### Figure 1-figure supplement 2

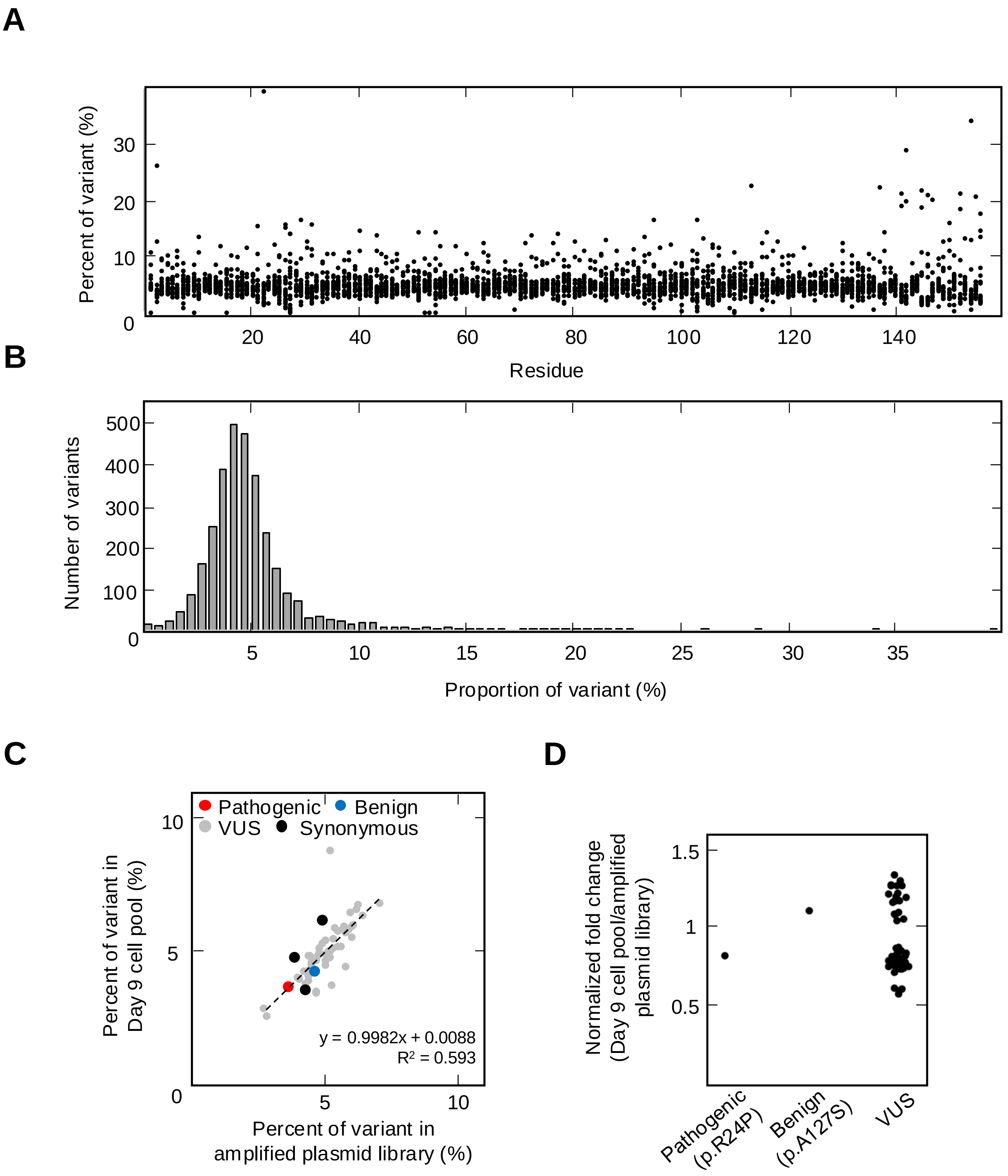

### Figure 2 -figure supplement 1

#### Figure 2-figure supplement 1

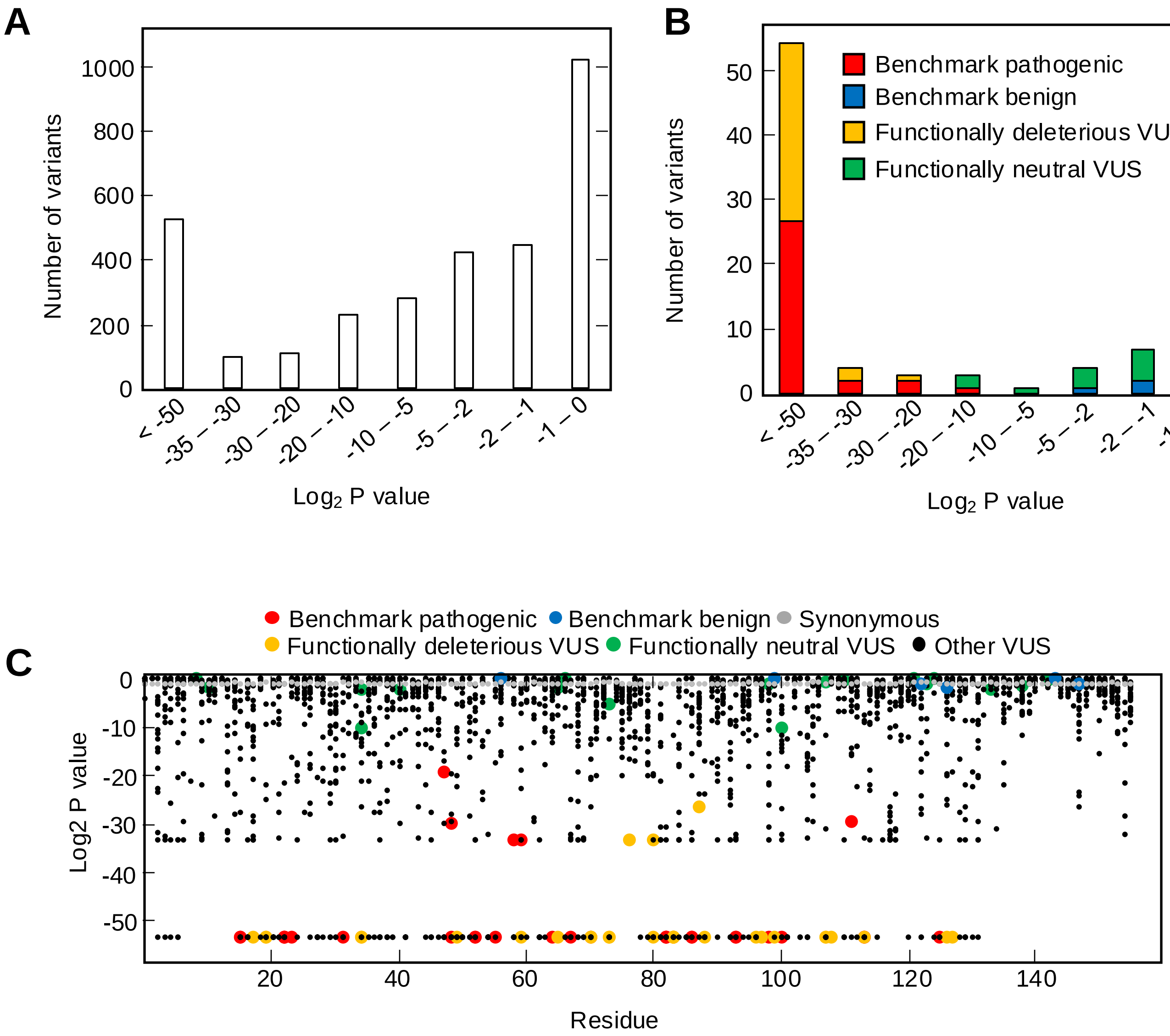

### Figure 2 -figure supplement 2

#### Figure 2-figure supplement 2

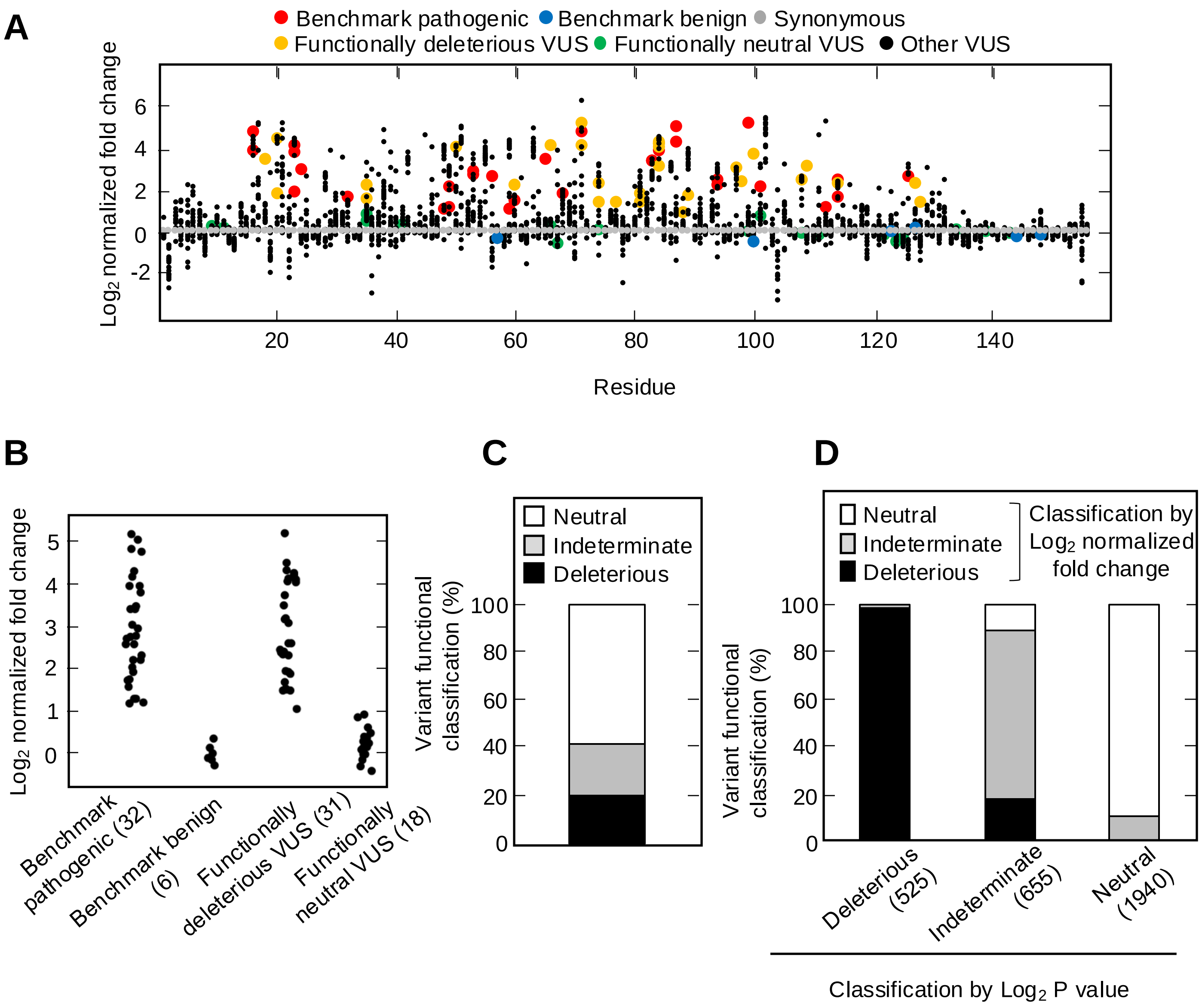

### Figure 2 -figure supplement 3

#### Figure 2-figure supplement 3

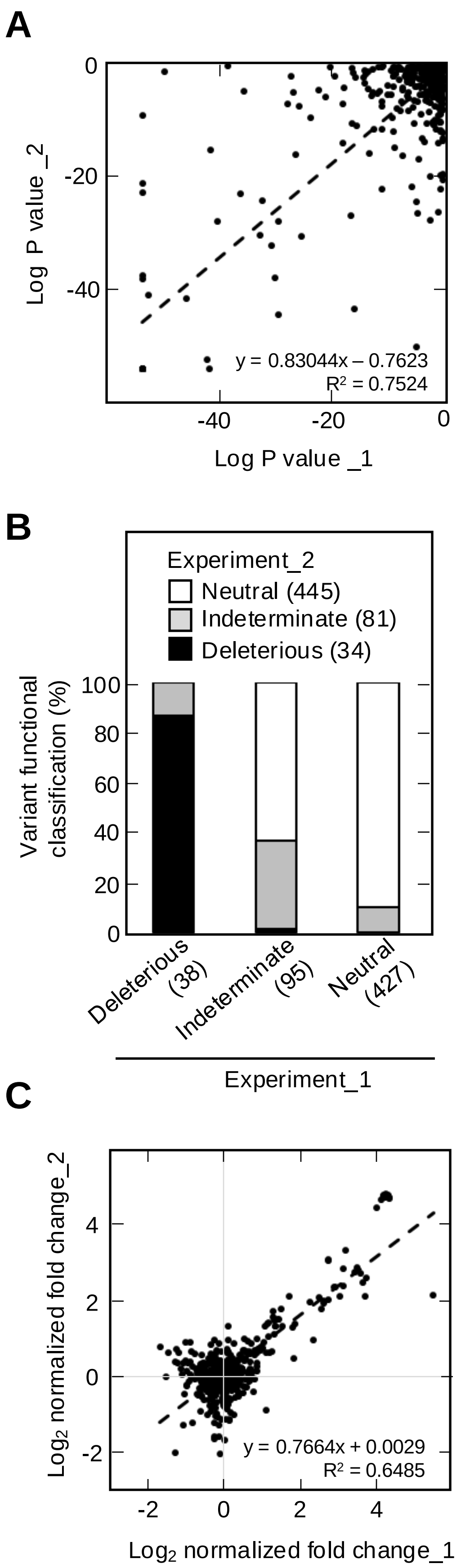

### Figure 2 -figure supplement 4

#### Figure 2-figure supplement 4

**A**

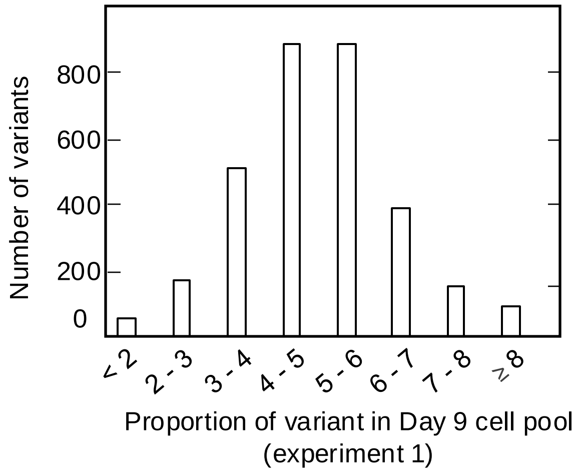


**B**

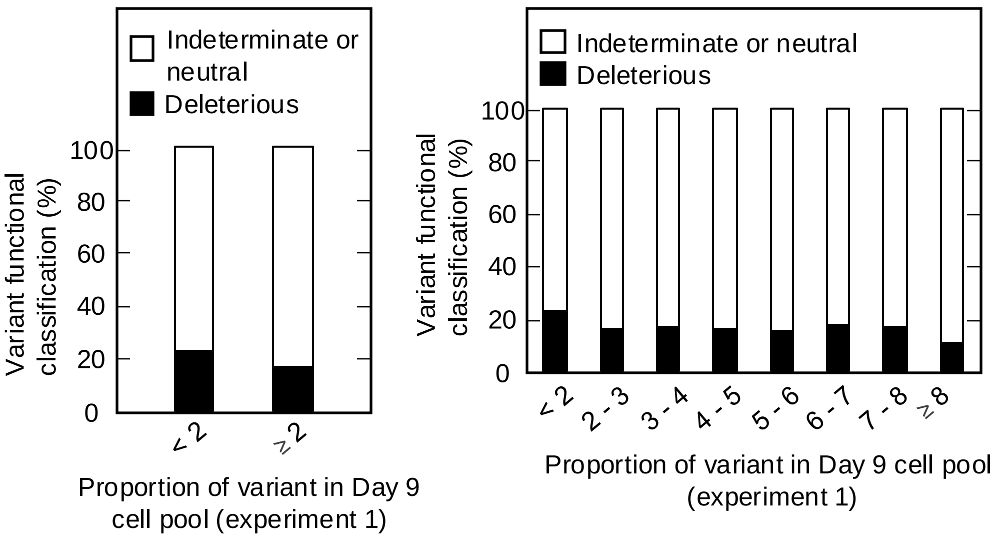

### Figure 2 -figure supplement 5

#### Figure 2-figure supplement 5

A

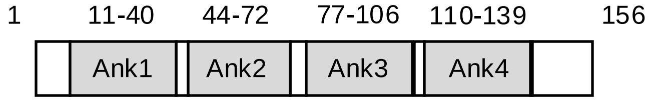


B

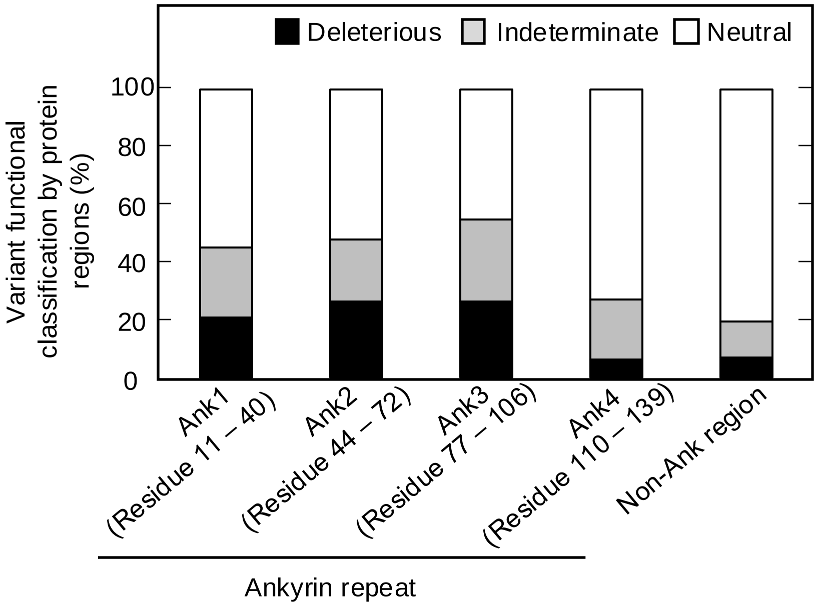


C

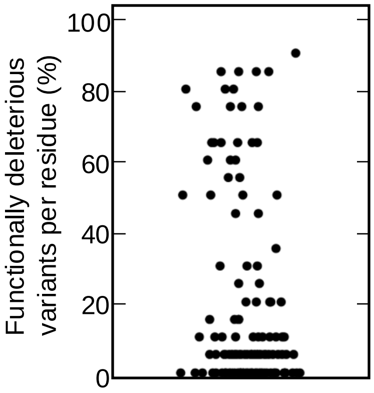

### Figure 3 -figure supplement 1

#### Figure 3-figure supplement 1

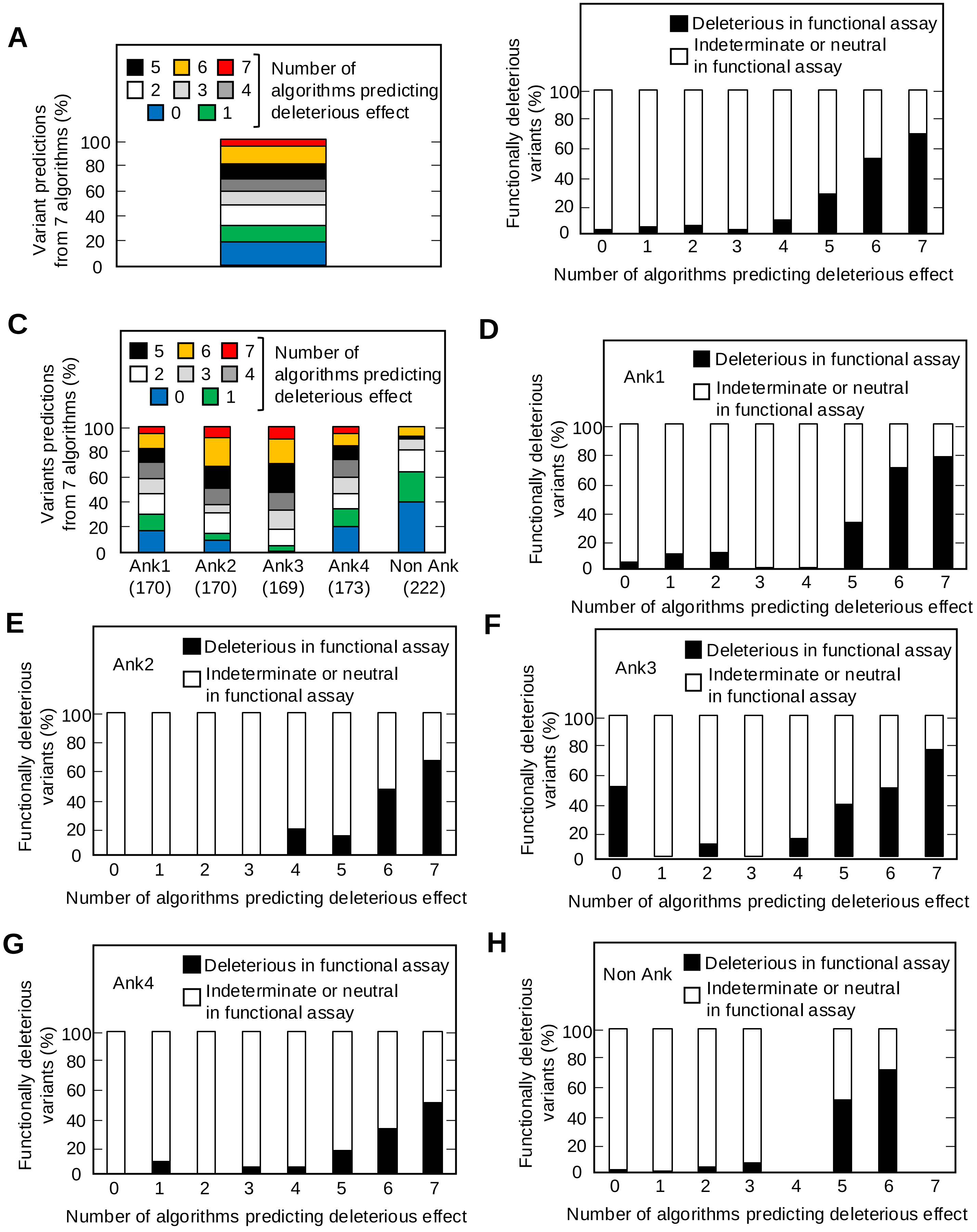

### Figure 3 -figure supplement 2

#### Figure 3-figure supplement 2

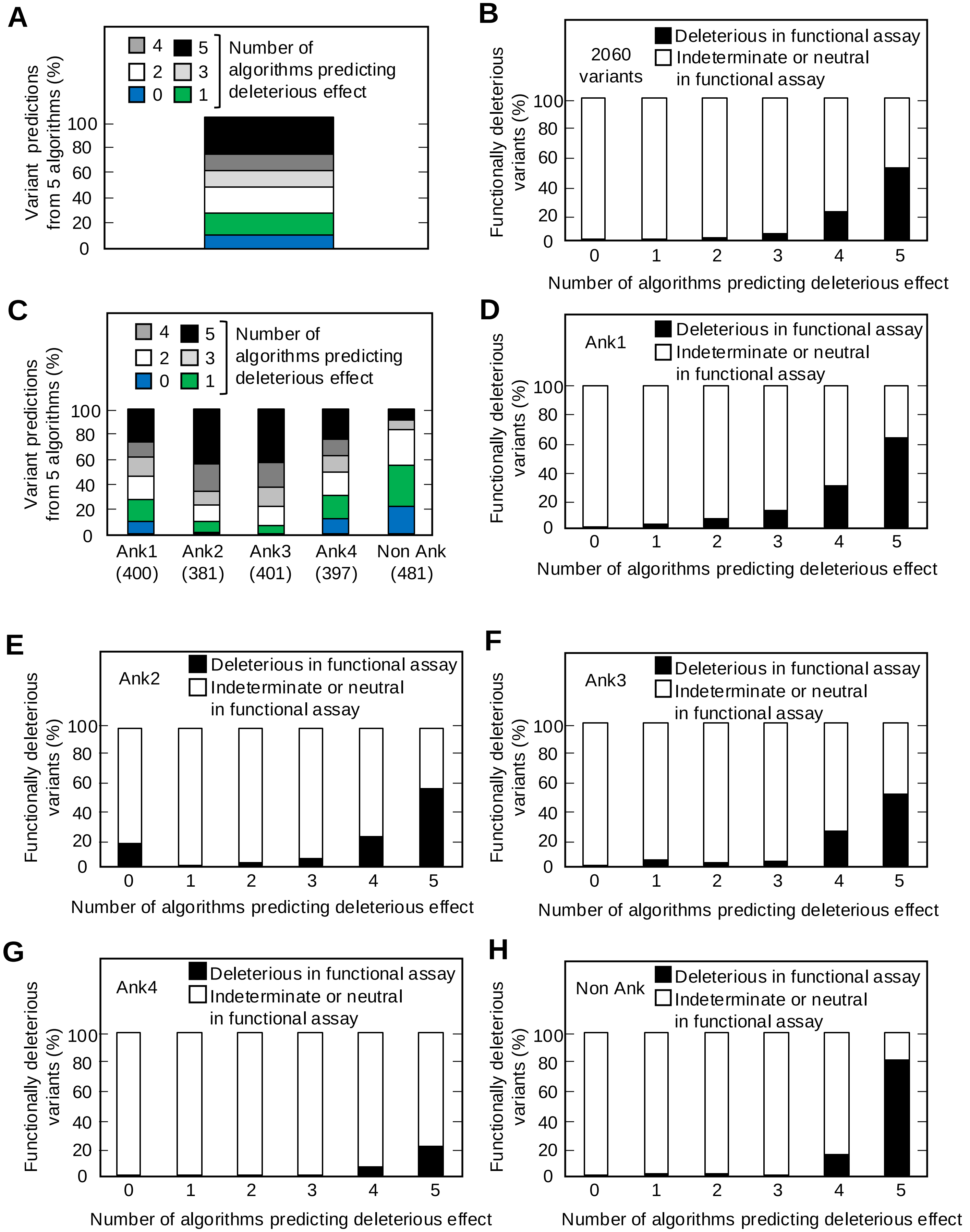

### Figure 4 -figure supplement 1

#### Figure 4-figure supplement 1

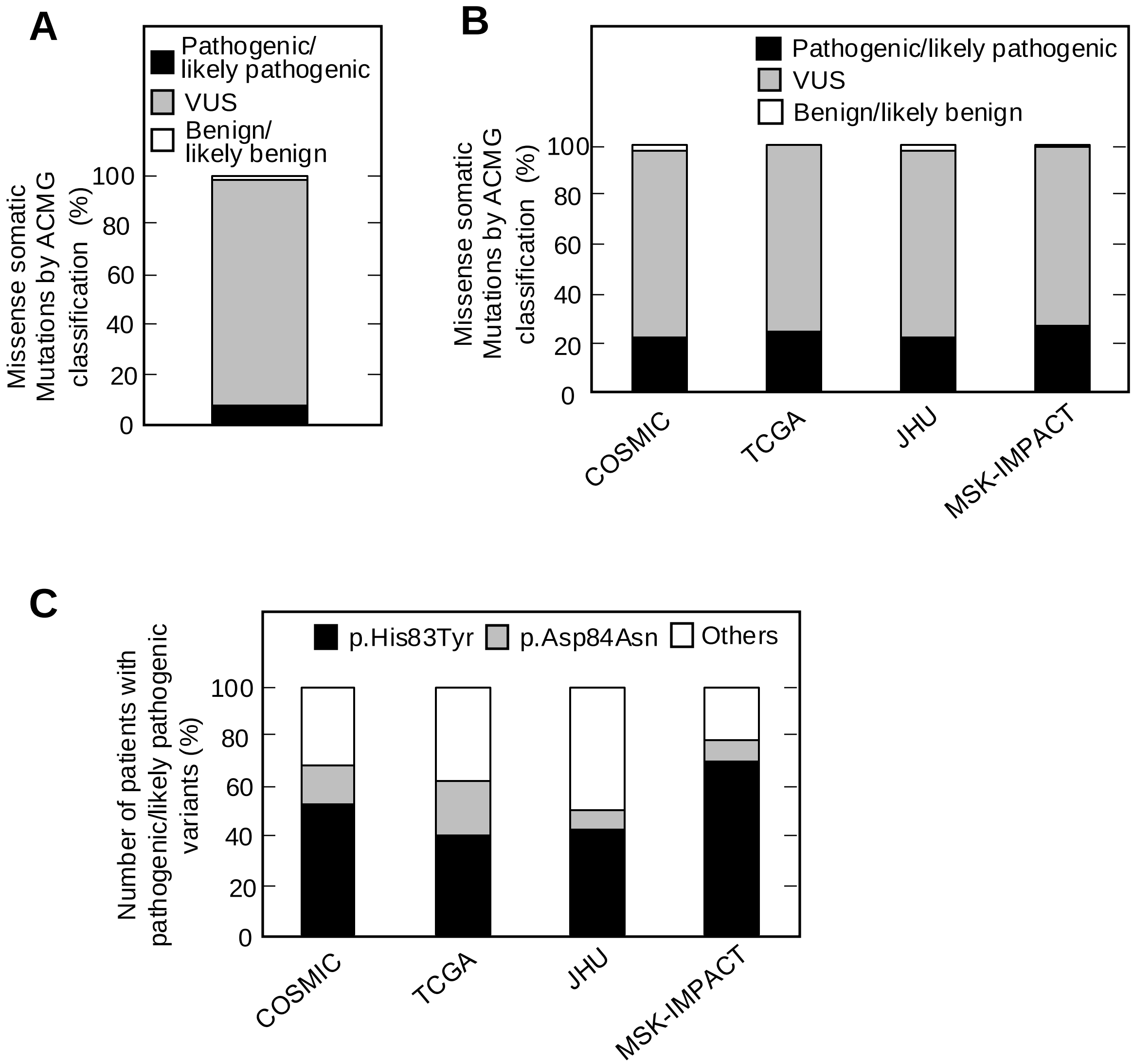

### Figure 4 -figure supplement 2

#### Figure 4-figure supplement 2

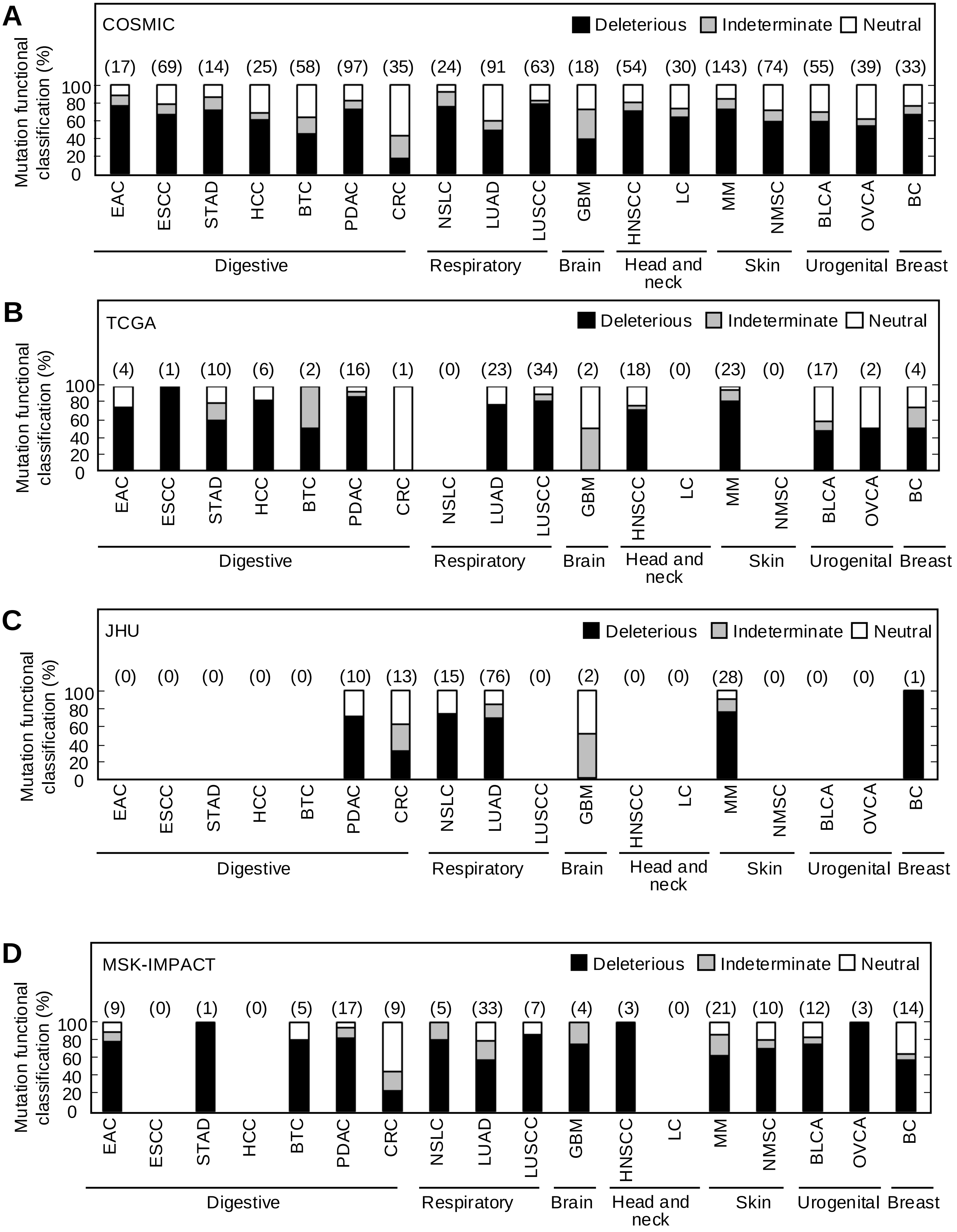
